# A structural solution to functional HGT: Gene chimerism bypasses mitochondrial expression barriers in parasitic plants

**DOI:** 10.1101/2025.11.14.688545

**Authors:** ME Roulet, LF Ceriotti, L Gatica-Soria, WD Tulle, MV Sanchez-Puerta

## Abstract

Horizontal Gene Transfer (HGT) in plant mitochondria is frequent, yet acquired genes are rarely functional due to expression barriers. The holoparasitic plant *Lophophytum mirabile* (Balanophoraceae) is an exceptional case, having functionally replaced numerous native mitochondrial genes with host-derived xenologs. This system provides a unique opportunity to investigate the mechanisms of functional HGT assimilation. Here, we assembled mitochondrial genomes of the sister species *L. pyramidale* and their mimosoid hosts and analyzed expression data from both holoparasites. We show that this extensive functional integration occurred without the co-transfer of nuclear regulatory factors; *Lophophytum* relies entirely on its pre-existing, native machinery. Our results demonstrate that the primary mechanism enabling *Lophophytum* to overcome the transcription barrier is structural: most functional xenologs are chimeric and retain native 5′ regions that likely place foreign coding sequences under the control of a recognizable native promoter. This structural solution is complemented by post-transcriptional flexibility, as the RNA editing machinery efficiently processes novel host-specific sites. However, functional replacement appears biased towards genes with inherently low editing requirements and no introns, highlighting a strong selective filter. Taken together, our results show that functional integration is driven by a combination of structural integration and the flexibility of the native regulatory system.

## INTRODUCTION

The acquisition of novel traits is a fundamental driver of evolution, achieved through mechanisms such as duplication and divergence of pre-existing genes, *de novo* from noncoding DNA, or the incorporation of foreign genes (xenologs) via horizontal gene transfer (HGT) [1–8]. HGT, the movement of genetic material between non-mating organisms, is a key evolutionary force extensively studied in prokaryotes, but increasingly recognized in eukaryotes. Among eukaryotes, plant mitochondria are a notable hotspot for frequent HGT, accumulating DNA fragments from diverse plant mitochondria [9–22]. However, this foreign DNA is only rarely transcribed into functional RNAs, often rendering these xenologs non-functional. This pattern suggests a high tolerance within the plant mitochondrial genome (mtDNA) for integrating non-functional foreign DNA, with most HGT events contributing to genetic diversity rather than functional novelty [12].

The limited evolutionary impact of most mitochondrial HGT events stands in contrast to the exceptional cases where horizontally acquired or intracellularly transferred genes (e.g., from nucleus or plastid) have successfully integrated and become functional in the recipient mtDNA [23–25]. Acquiring a functional role, however, is severely constrained by several expression barriers, (Figure 1), as the native, nuclear-encoded machinery often fails to process foreign transcripts. These barriers include poor recognition of foreign promoters, inefficient splicing of novel introns, or incomplete RNA editing, modifications essential for producing functional mitochondrial proteins in plants [26–28]. The scarcity of documented functional foreign genes in plant mitochondria precludes a thorough understanding of the specific strength and nature of these regulatory barriers, and how they can be effectively overcome by the recipient cell.

**Figure 1.**
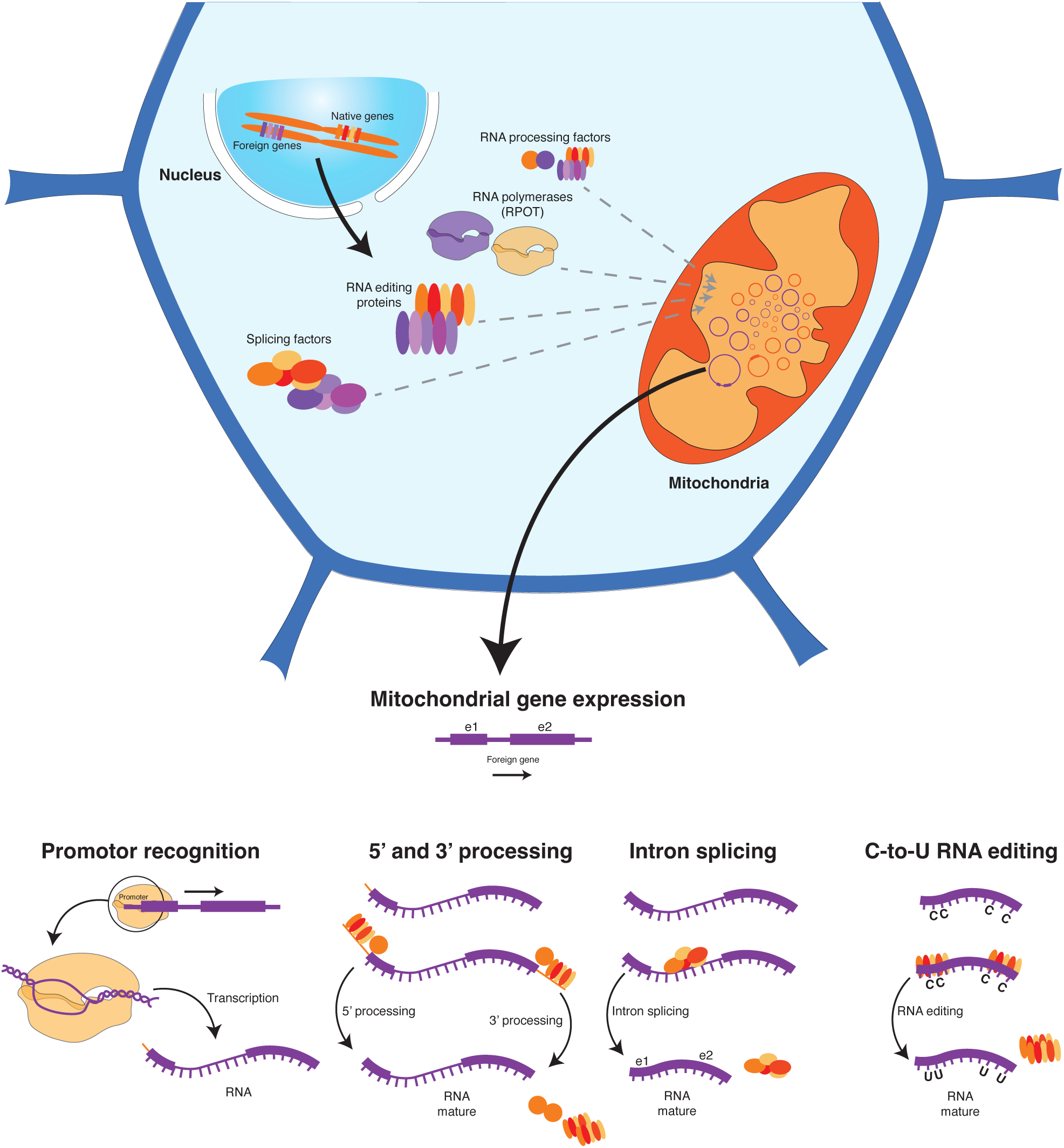
Functional barriers to the expression of foreign mitochondrial genes in *Lophophytum*. Schematic representation of the main nuclear-encoded factors involved in mitochondrial gene expression.

While most foreign genes are non-functional, recurrent HGT events inherently increase the likelihood of successful functional replacements over evolutionary time. Recent work from our group has identified a truly exceptional system: the mtDNA of the holoparasite *Lophophytum mirabile* (Balanophoraceae, Santalales). This is the first mtDNA, plant or otherwise, in which a substantial number of native protein-coding genes (roughly two-thirds) have been functionally replaced by intact foreign homologs, all acquired via HGT from its legume hosts [15,29]. Most of these foreign genes that have replaced the native copies are highly transcribed, undergo efficient RNA editing, and evolve under purifying selection, providing robust evidence for their functional role [15,29]. The underlying mechanisms enabling *Lophophytum* to successfully express these foreign genes—acquired from donors that diverged ∼125 million years ago [30]—remain unknown.

This study exploits the distinctive characteristics of the *Lophophytum* system to analyze the molecular pathways through which foreign mitochondrial genes achieved successful functional incorporation. Specifically, we address how the expression barriers were overcome, focusing on three major regulatory checkpoints: (1) transcriptional initiation: Are the foreign mitochondrial genes transcribed using foreign promoter regions, or have they been placed under the control of native *Lophophytum* regulatory sequences? (2) post-transcriptional processing: Are *Lophophytum* spp. capable of efficient RNA editing of novel C-to-U sites that are specific to the host lineage? Are foreign introns accurately spliced? And (3) nuclear-encoded machinery: Did *Lophophytum* co-acquire foreign nuclear factors (e.g., PPR proteins) responsible for the transcription and post-transcriptional processes necessary for xenolog expression? To address these questions, we combined genomic and deep transcriptomic data from two *Lophophytum* species and their mimosoid host plants. Together, these combined analyses provide a comprehensive framework for understanding how horizontally acquired genes can achieve functional assimilation within a highly diverged mitochondrial background.

## MATERIALS AND METHODS

### Assembly of organellar genomes of mimosoid hosts

DNA sequencing data from the mimosoid *Anadenanthera colubrina* (a host species of *L. mirabile*; tribe Mimoseae - Fabaceae [31]) were obtained from recent sequencing efforts from our group (NCBI BioProject PRJNA1197971 [32]). Additionally, the DNA-seq data from the mimosoid *Vachellia collinsii* were downloaded from public databases (SRX4061545; tribe Mimoseae - Fabaceae [33]). To assemble the plastid genome (ptDNA) of both species, we used NOVOplasty v.2.6.2 [34] with different references as seed: *Acacia ligulata* (NC_026134.2) and *Vachellia tortilis* subsp. *raddiana* (KY100266.1) ptDNAs for *Anadenanthera* and *Vachellia* assemblies, respectively, followed by visualization and manual curation using Consed v.29.0 [35]. The ptDNA of *Anadenanthera* was deposited in GenBank (PX438801).

To obtain the mitochondrial genomes (mtDNAs) of both mimosoid species, we performed *de novo* assemblies of the DNAseq data using SPAdes v.3.15.2 [36] with coverage cutoff set to automatic (assembly details in Table S1). Putative mitochondrial contigs were identified and visualized in Bandage v.0.8.1 [37] through BLASTn analysis [38] using a custom database of Mimoseae mitochondrial DNA downloaded from NCBI (Table S2A). The putative mitochondrial contigs were extended individually using SSAKE v.3.8.5 [39] (Table S1). Then, mitochondrial contigs were manually edited, joined, and circularized based on consistent paired-end reads visualized in Consed v.29 [35].

The total read depth for each organellar genome was calculated using Bowtie2 v.2.2.2 [40] with the following presets: –end-to-end –very-fast, SAMtools v.1.10 [41] and BEDtools v.2.26.0 [42]. Plots were generated using R software v.4.2.2. No gaps or low-coverage regions were detected. Mitochondrial genomes of *Anadenanthera* and *Vachellia* were deposited in the GenBank (PV798382 and PV826156). The characterization of the mimosoid organellar genomes is available in Note S1.

### Phylogenetic analyses of mitochondrial genes in *Lophophytum* spp

To infer the origin of the mitochondrial genes in *Lophophytum pyramidale* and re-assess those in *L. mirabile*, phylogenetic analyses were performed under Maximum Likelihood (ML). Nucleotide sequences of diverse angiosperms (Table S2B), were aligned with muscle in AliView v.1.27 [43]. ML trees were obtained with RAxML v.8.2.11 [44] using the GTR + gamma model including 1,000 rapid bootstrap (BS) pseudoreplicates. The trees were visualized with FigTree v.1.4.4. Genes were considered foreign when *Lophophytum sp.* was affiliated with Fabaceae with BS >70%. Host contamination has been ruled out through careful dissection of the inflorescences used for DNA extraction, consistent read depth between native and foreign mitochondrial regions, and the analyses of other individuals of these holoparasitic species [32,45].

To evaluate the presence of chimeric genes (partly native and partly foreign) in both *Lophophytum* spp., gene alignments were visually examined and analyzed with the program Geneconv v.1.81a [46]. For these alignments, we incorporated gene information from various Santalales and mimosoid species, including those mimosoid species that we assembled (Table S2C). For those conversion tracts considered significant in Geneconv (*p* <0.05), phylogenetic analyses were performed on subregions of the gene alignments, and BLASTn searches were conducted on 100 bp flanking sequences, when possible. Subregions were classified as foreign if sequences from *Lophophytum* spp. were in a well supported clade (BS > 70%) with Fabaceae species.

### Mitochondrial genome expression analysis of *Lophophytum spp*

Sequence data from total RNA reads, depleted in rRNA (NCBI SRA: SRR31014705; [32]) were aligned over each mitochondrial circular chromosome of *L. pyramidale* individual #1 using the Bowtie2 program [40] with the default settings: –end-to-end, –sensitive, -no-discordant, -no-mixed, and -R=10. Additionally, the -fr and -nofw options were used to adjust the strand-specific alignment and the alignment was done on each strand separately. The mapping was visualized in Geneious v.2023.1.1, and the results were plotted with R v.4.2.2. We also aligned all RNA reads using Bowtie2 [40] to all protein genes in the *L. pyramidale* mtDNA, with these CDS extended by adding the flanking 300 bp at their 5’ and 3’ end to obtain the appropriate read depth at the ends of the genes and the results were plotted with R v.4.2.2.

To assess the transcription levels of foreign and native mitochondrial protein genes, we first calculated the background transcription of the whole mtDNA of *L. pyramidale*. For this, we analyzed RNA reads derived from noncoding chromosomes of *L. pyramidale* (without known mitochondrial genes, pseudogenes, and MTPTs) to estimate the background transcription level (i.e., the average RNA level for those regions that probably have no function but are transcribed nonetheless). Then, we determined the read depth threshold for significance at the *p* ≤0.05 level following Wu et al. [47]. This method calculates the probability of obtaining an RNA read depth, averaged from 500 bp genomic windows, equal to or exceeding the 5% tail of the read depth distribution across the mtDNA. The transcription of *L. mirabile* mtDNA has been previously analyzed [29].

### Identification of RNA editing sites in mitochondrial genes

We identified the RNA editing sites and the editing efficiency in the mitochondrial protein-coding genes of *L. pyramidale* (SRR31014705) and of a close relative with native functional genes, *Ombrophytum subterraneum* (SRR31014706; Balanophoraceae; [48]). For this, we aligned RNAseq data over each of the CDS plus 300 bp flanking sequences and analyzed the alignments with the bam-readcount program [49] and customs scripts. The RNA editing sites and the editing efficiency for *L. mirabile* were previously reported [29]. For comparison purposes, predicted editing sites were identified using PREPACT3 v.3.12.0 [50] with the BLASTX editing mode against a curated set of angiosperm mitochondrial genomes.

The editing efficiency at non-synonymous RNA editing sites was compared between the native and foreign regions of mitochondrial genes in *L. pyramidale* and *L. mirabile*. In both species, data distributions deviated significantly from normality (Shapiro–Wilk test), supporting the use of the non-parametric Wilcoxon rank-sum test. To evaluate whether the lower editing efficiency in foreign regions is due to the suboptimal editing of foreign editing sites or to intrinsic properties of the genes impacted by HGT, we performed a comparative analysis using the dataset of editing sites across 17 diverse angiosperms from Edera et al. [51]. We analyzed the editing efficiency across angiosperms of genes homologous to those that are native vs those foreign ones in *L. mirabile*.

A potential barrier for foreign gene expression is the presence of host-specific non-synonymous C-to-U editing sites. Of the RNA editing sites identified in *L. mirabile* and in *L. pyramidale*, we inferred the host-specific non-synonymous C-to-U RNA editing sites in foreign regions of *Lophophytum* mitochondrial genes by comparing the editing profiles across four taxa: *L. pyramidale*, *L. mirabile*, *Ombrophytum*, and the mimosoid *Acacia ligulata* (representing the host lineage). We assumed that *Lophophytum* and *Ombrophytum* share ancestral editing patterns in native genes. A host-specific editing site acquired by *Lophophytum* was defined as a position where (i) a cytosine (C) is edited in a foreign region of *Lophophytum*, and (ii) the homologous site in *Ombrophytum* contains a thymine (T) or an unedited C.

### Phylogenetic origin of nuclear genes involved in mitochondrial RNA maturation

To describe and assess the phylogenetic origin of the nuclear-encoded machinery required for mitochondrial gene expression in *Lophophytum* spp., we performed a targeted survey of nuclear genes involved in transcription and post-transcriptional regulation within plant mitochondria, such as transcription initiation, transcript stabilization, RNA splicing, and RNA editing. We assembled a curated reference set of nuclear-encoded proteins known to function in mitochondrial RNA maturation, encompassing several functional classes: mitochondrial transcription termination factors (mTERFs), CRM-domain proteins, RNA-binding proteins, RNA helicases, RNases, and RNA polymerases (Table S4, [52]). For site-specific RNA editing factors, we followed Guo et al. [53].

We searched for homologs of *Arabidopsis* genes in the transcriptomes of *Lophophytum mirabile, L. pyramidale,* and *Ombrophytum subterraneum*. For this, we used the transcriptome assembly of *L. mirabile* [29] and assembled the RNAseq data of *L. pyramidale* and *O. subterraneum* using Trinity v2.15.0 (parameters --SS_lib_type RF and --min_contig_length 100). We extracted >100 bp ORFs with TransDecoder v.5.7.0 (https://github.com/TransDecoder/TransDecoder). *Arabidopsis* accession numbers were also used to identify orthogroups across the species of interest using OrthoFinder clustering results (Table S4; Ceriotti et al. unpublished). Multiple protein sequence alignments were performed using MAFFT [54], and poorly aligned regions or divergent sequences were trimmed with TrimAl (parameters: -resoverlap 0.5 -seqoverlap 50 and -gt 0.3) to improve phylogenetic resolution. Maximum Likelihood phylogenetic trees were inferred with IQ-TREE, using a substitution model specific to plants (Q.plant). Bootstrap support values were calculated based on 1,000 ultrafast replicates and those >95% were considered statistically robust [55] to infer a HGT event from mimosoid hosts.

## RESULTS

### More chimeric than fully foreign mitochondrial genes in *Lophophytum* spp

Based on phylogenetic and gene conversion analyses using new comparative data from two *Lophophytum* species and mimosoid host mtDNAs, we reclassified several protein-coding genes previously identified as fully foreign in the holoparasite *L. mirabile* [16,29,56]. Specifically, 12 genes were reclassified as chimeric and four (including *nad4L*, *rpl16*, *rps4*, and *rps19*) as native. Conversely, the native gene *cob* was reclassified as chimeric. Overall, the protein-coding gene complement of *L. mirabile* (44 genes) is roughly equally distributed across the three categories: 14 chimeric (32%), 15 foreign (34%), and 15 native (34%) genes (Figure 2 and Table S5).

**Figure 2.**
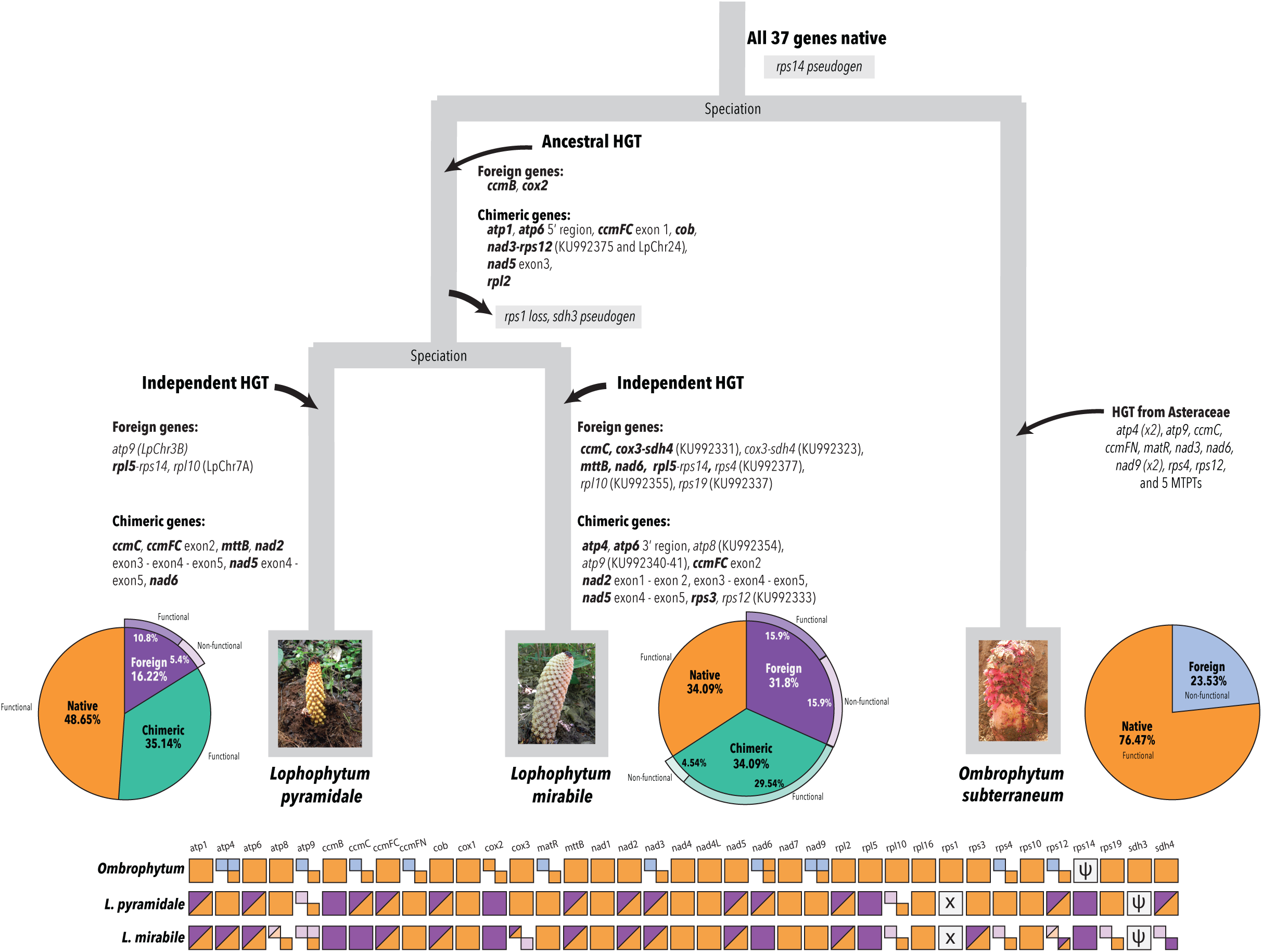
Ancestral and independent HGT events from diverse hosts (Fabaceae and Asteraceae) in the mitochondrial genomes (mtDNAs) of *Lophophytum* spp. and *Ombrophytum*. The phylogenetic timing of HGT events resulted in foreign or chimeric genes. Functional genes in *L. mirabile* and *L. pyramidale* are shown in boldface. The evolutionary origins of intact protein-coding genes are depicted, with native genes in orange, foreign genes from Fabaceae in purple, and those from Asteraceae in blue. Non-functional genes are represented in pale colors. The percentages of native (orange), foreign (purple or blue), and chimeric (green) protein-coding genes in *Lophophytum* spp. and *Ombrophytum* are indicated. Chimeric genes are represented by boxes with diagonal lines. The photographs show inflorescences of each holoparasite (courtesy of H. A. Sato).

Our analyses also confirmed the presence of six foreign genes (16% [32]) and revealed 13 chimeric protein-coding genes (35% of all protein-coding genes) in the mtDNA of the sister species *L. pyramidale*. The remaining 18 of the 37 protein-coding genes were classified as native, most grouping with other Balanophoraceae in phylogenetic trees (Tables 1, S6, and Figure S3).

The presence of HGT-impacted introns in both holoparasites is low: 6 (25%) and 4 (16%) of the 24 typical mimosoid introns were found to be only partially foreign in *L. mirabile* and *L. pyramidale*, respectively. No fully foreign introns were detected in either species, except for the *cox1* intron, which was ancestrally acquired in the Balanophoraceae [16].

### Mitochondrial genes have been acquired in ancestral and independent HGT events

By combining phylogenetic trees and analyses of foreign DNA integration sites (Figure S3), we inferred the evolutionary timing of the HGT acquisitions in *Lophophytum* spp. (Figure 2). Foreign DNA was incorporated through both ancestral events, predating the divergence of *L. pyramidale* and *L. mirabile*, and more recent independent HGT events. Ancestral HGT led to two foreign and seven chimeric protein-coding genes or gene clusters. The long-term maintenance of these HGT-impacted genes is consistent with their inferred functionality in both *Lophophytum* species (as detailed below). After speciation, independent HGT events from mimosoid donors introduced additional foreign and chimeric genes into both species, including instances of convergent HGT where the same genes were transferred independently to both *L. mirabile* and *L. pyramidale* (Figure 2). Overall, the impact of HGT on coding regions is greater in *L. mirabile*, with 29 chimeric or foreign protein-coding genes, compared to 19 in *L. pyramidale*.

### Single-copy foreign and chimeric mitochondrial genes in *Lophophytum pyramidale* mtDNA are highly transcribed

Building upon previous work on *L. mirabile* mitochondria [29], we evaluated the RNA read depth and editing efficiency of all mitochondrial genes in *L. pyramidale* to assess their functionality (Table S6). Of the 237 million high-quality paired-end RNA reads, approximately 43 and 30 million aligned to each strand of the *L. pyramidale* mitochondrial genome, respectively. The entire *L. pyramidale* mtDNA, including all 81 chromosomes, is fully transcribed on both strands (Figure S4 and Table S7). All protein-coding genes exhibit distinct peaks in read depth; however, the lack of apparent distinction between exonic and intronic read depth suggests reduced splicing efficiency, which is not restricted to genes impacted by HGT [48]. We estimated the background transcription level using non-coding regions (732.2; standard deviation = 630.5) and determined that significant transcription following the method described by Wu et al. 2015 [47] requires a mean read depth exceeding 10,930 reads (*p* ≤ 0.05).

All native genes (except for *rps4*), all chimeric genes (except for *ccmFC*), and all non-duplicated foreign genes (except for *rsp14*) in *L. pyramidale* mtDNA show significant transcription levels (Table S6). The three genes that fall below the significance threshold, show an RNA read depth well above the background transcription level and no evidence of nuclear compensation, except for *rps14* (see Note S2 and Figure S5). Thus, 34 native, chimeric or foreign mitochondrial genes are putatively functional in *L. pyramidale*.

In stark contrast, the two foreign genes that co-exist as duplicates with native homologs exhibit non-significant and substantially lower transcription levels, while their native counterparts show high and significant transcription (Figure S6 and Table S6; [32]). This suggests that in the presence of a functional native copy, the foreign duplicate remains non-functional, which aligns with previous observations in *L. mirabile* mtDNA [29].

### The putatively functional mitochondrial genes in *L. pyramidale* are accurately and efficiently RNA-edited regardless of their phylogenetic origin

A total of 568 C-to-U editing sites were identified in the 34 putatively-functional protein genes in *L. pyramidale*, of which 464 (81.7%) are non-synonymous editing sites (Tables 1, S8 and S9). As reported for *L. mirabile* [29], in three *L. pyramidale* genes (*atp6*, *ccmFc,* and *cox2*), RNA editing produce premature stop codons, while three others (*nad1*, *nad3*, and *rps10*) acquire a start codon through RNA editing (Table S9). Like in diverse photosynthetic angiosperms [51,57], most editing events in *L. pyramidale* occur at first and second codon positions, with lower frequency at third codon positions (Table S9). We also observed similar proportions in *L. mirabile* and *Ombrophytum*, with editing sites concentrated in the first and second codon positions (Tables 2, S10 and S11). Notably, 90% of all the predicted editing sites in foreign regions of protein-coding genes were actually edited in *L. pyramidale,* similar to the percentage observed in native regions or native genes (91%) (Table S6).

Moreover, editing efficiency per codon position in these holoparasitic species closely matches, or exceeds, the average values reported in photosynthetic species (Table 2). In particular, the putatively functional foreign and chimeric genes in *L. pyramidale* undergo efficient C-to-U editing (>70% average efficiency for most genes, Table S6).

**Table 1.**
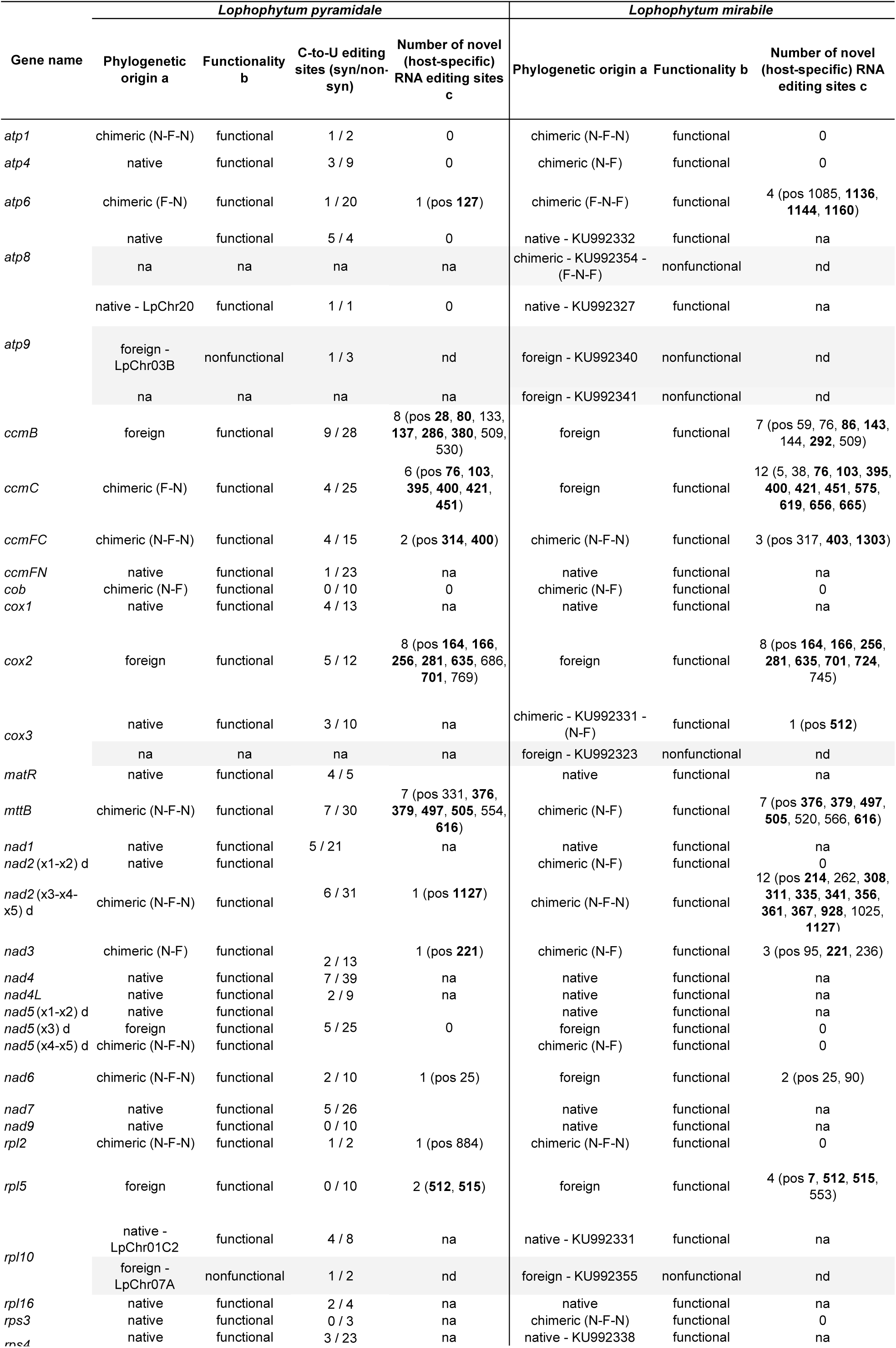
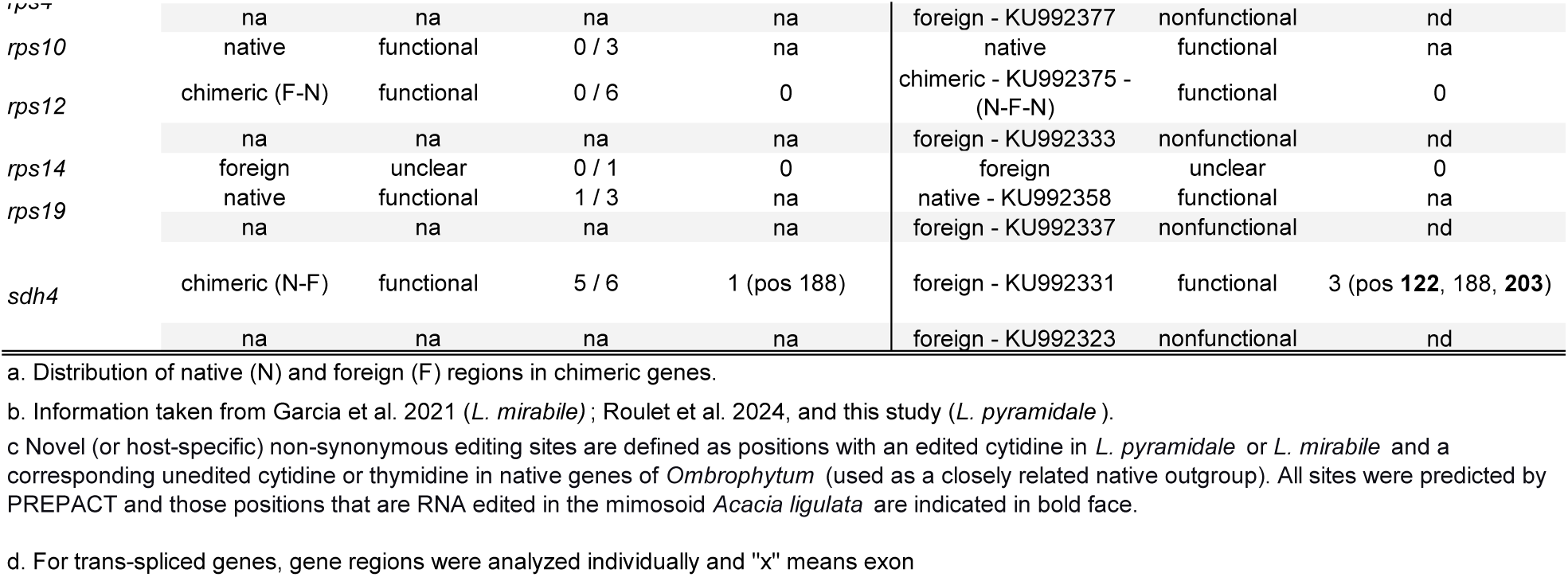
Main features of mitochondrial protein-coding genes in *Lophophytum* spp.

**Table 2.** RNA editing in 24 mitochondrial protein genes in *Lophophytum* spp., *Ombrophytum* and selected photosynthetic angiosperms.

| Species | Number of editing sites a |  |  |  | Average editing efficiency (%) |  |  |
| --- | --- | --- | --- | --- | --- | --- | --- |
|  | 1st Codon Position | 2nd Codon Position | 3rd Codon Position | TOTAL | 1st Codon Position | 2nd Codon Position | 3rd Codon Position |
| *L. pyramidale* | 143 | 233 | 70 | 446 | 70.9 | 78.97 | 38.74 |
| *L. mirabile* b | 152 | 223 | 87 | 462 | 81.05 | 86.15 | 42.23 |
| *Ombrophytum* | 128 | 211 | 65 | 404 | 84.98 | 87.44 | 42.96 |
| Photosynthetic angiosperms c | 159 | 275 | 56 | 489 | 78.5 | 83.2 | 40.9 |

a. C-to-U editing sites in 24 genes: *atp1*, *atp4*, *atp6*, *atp8*, *atp9*, *ccmB*, *ccmC*, *ccmFc*, *ccmFn*, *cob*, *cox1*, *cox2*, *cox3*, *matR*, *nad1*, *nad2*, *nad3*, *nad4*, *nad4L*, *nad5*, *nad6*, *nad7*, *nad9*, and *rps12*.
b. Data taken from Garcia et al. (2021).
c. Average values taken from Edera et al. (2018) including *Allium cepa*, *Arabidopsis thaliana*, *Cucumis sativus*, *Daucus carota*, *Glycine max*, *Gossypium hirsutum*,

Taken together, these results indicate that the mitochondrial gene complement of *L. pyramidale* retains broad functional capacity. Based on a combination of significant transcription, accurate and efficient non-synonymous C-to-U editing, and the absence of nuclear homologs, a total of 34 mitochondrial protein-coding genes in *L. pyramidale* were classified as functional, including 4 foreign and 12 chimeric genes (Tables 1 and S6).

### Editing efficiency differs between native and foreign regions of functional genes

RNA editing analyses of functional genes revealed that the editing efficiency was significantly higher in native than in foreign mitochondrial gene regions of both *Lophophytum* spp. In *L. pyramidale*, the mean efficiency was 75.6% in native regions versus 68.4% in foreign ones, while in *L. mirabile*, means were 84.1% and 75.9%, respectively (Figure S7). To evaluate whether the lower editing efficiency in foreign regions is due to the suboptimal editing of foreign editing sites or to intrinsic properties of the genes impacted by HGT, we performed a comparative analysis. We analyzed the editing efficiency of genes homologous to those that are native vs those foreign ones in *L. mirabile* and in diverse free-living angiosperms. Even in these non-parasitic species, where all genes are native, editing efficiency was lower in the group of genes homologous to the foreign genes in *L. mirabile* (Wilcoxon test: W = 10,586,941, *p* < 2.2 × 10⁻^16^) than in the group of genes homologous to native genes in *L. mirabile* (Figure 3). This finding strongly indicates that the reduced editing efficiency is likely gene-intrinsic, rather than a general feature imposed by their foreign origin or the *Lophophytum* editing machinery.

**Figure 3.**
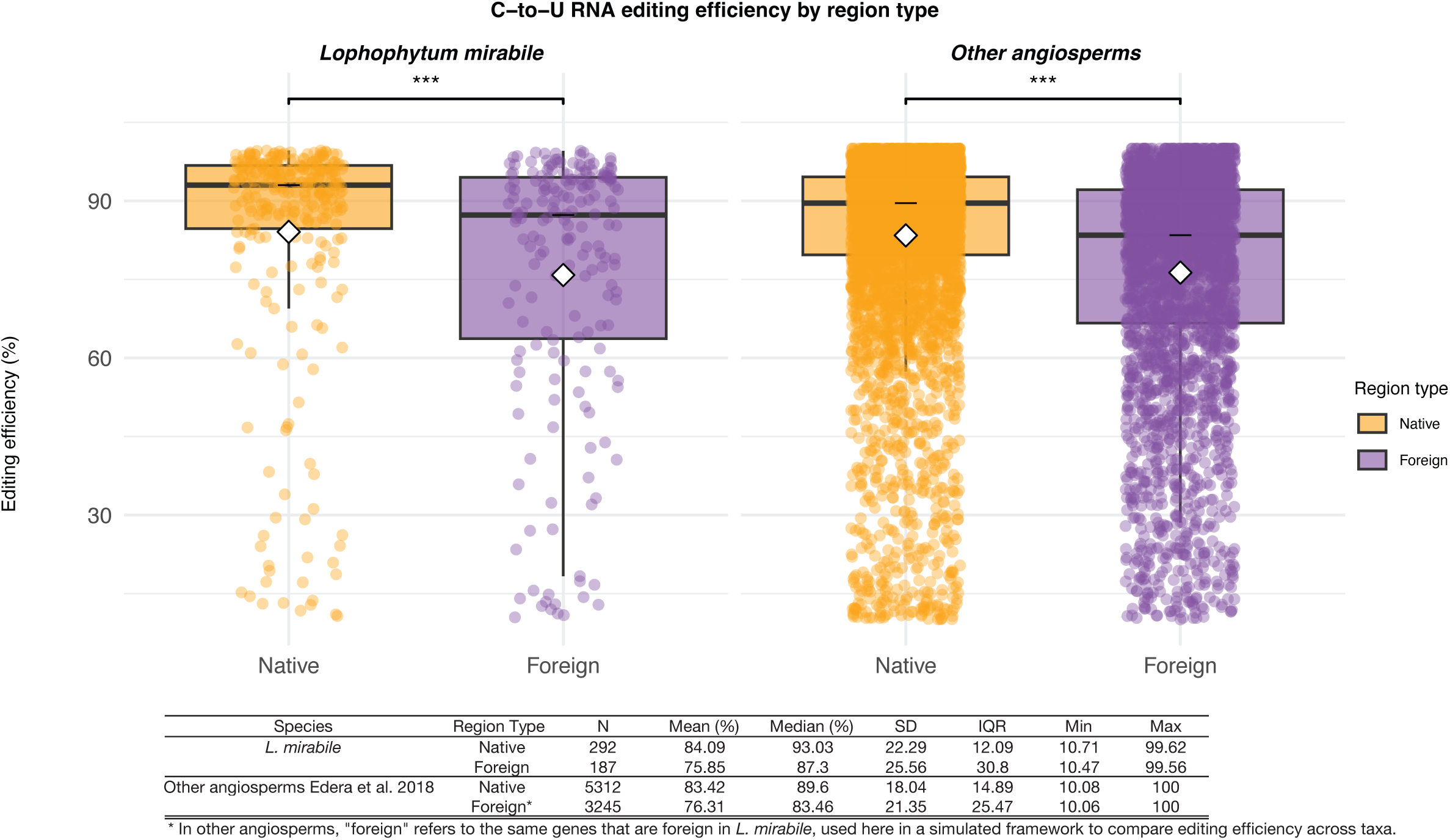
Boxplots showing C-to-U RNA editing efficiency (%) at non-synonymous sites in native vs. foreign regions for *L. mirabile* compared to other angiosperms. For *L. mirabile*, categories correspond to gene origin (native vs. foreign). Editing efficiency is significantly higher in native regions (Wilcoxon rank-sum test, W = 34,286, *p* = 2.3 × 10−6). Shapiro–Wilk tests indicated non-normal distributions for both native (W = 0.642, *p* < 2.2 × 10−16) and foreign (W = 0.797, *p* = 7.5 × 10−15) sites, supporting the use of the non-parametric test. For other angiosperms, “native” and “foreign” labels were assigned by replicating the *L. mirabile* gene classification, although all genes are native in these species (dataset from Edera et al., 2018). Editing efficiency was again higher in the “native” group (Wilcoxon rank-sum test, W = 10,586,941, *p* < 2.2 × 10−16). Normality testing (Shapiro–Wilk) on a random subset of 5,000 sites confirmed non-normal distributions (native: W = 0.742; foreign: W = 0.855; both *p* < 2.2 × 10−16). Boxplots show medians (black line) and means (white diamond). *** *p* < 0.001.

### Novel RNA editing sites in horizontally acquired regions

To determine whether *Lophophytum* spp. can recognize and edit efficiently novel RNA editing sites acquired by HGT, we searched for host-specific C-to-U RNA editing sites in foreign and chimeric mitochondrial genes. Novel (or host-specific) sites were defined as positions with an edited cytidine in *Lophophytum* and a corresponding unedited cytidine or thymidine in *Ombrophytum* (used as a closely related native outgroup). Of the non-synonymous editing sites identified within foreign coding regions in *L. pyramidale* (127 sites), 31% (39 sites) were classified as novel (Table 1). Similarly, 31% (66 of 210 sites) were novel in *L. mirabile*. The majority of these novel sites are RNA edited in *Acacia* [29] and all of them are predicted by PREPACT, indicating that their edition leads to conserved amino acids. These novel sites show high average editing efficiencies of 72.5% in *L. pyramidale* and 75.6% in *L. mirabile* (Tables S9 and S10). This result demonstrates the successful recognition and processing of editing sites specific to the mimosoid donor lineage by the *Lophophytum* machinery.

### Chimeric genes and co-transcriptional units in *Lophophytum* mitochondria

To address whether transcription of foreign genes is facilitated by co-acquired foreign RNA polymerases (RPOTs) capable of recognizing foreign promoters, we analyzed the *Lophophytum* spp. transcriptomes. The mitochondrial (RpoTm) and dual-targeted (RpoTpm) RNA polymerases were identified and phylogenetic analyses confirmed their native origin (Figure S8), excluding the co-acquisition of foreign nuclear transcription factors.

Given that a majority of functional HGT-impacted genes are chimeric, we examined the arrangement of native and foreign regions. Strikingly, nearly all functional chimeric genes (all but three) exhibit a native 5’-region followed by a foreign downstream region (Tables 1 and S6). This 5’-native configuration strongly suggests the retention of a native promoter upstream, which is efficiently recognized by the native RPOTs. Furthermore, several foreign or chimeric genes are likely co-transcribed as part of gene clusters ([29]; Figure S9), in which the upstream gene is either native (e.g. *cox3*-native-*sdh4*-foreign) or chimeric (e.g. *nad3*-*rps12*-*cob*). Thus, promoter retention via chimeric gene formation is a major mechanism for overcoming the transcription barrier. Finally, two ancestral foreign genes (*cox2* and *ccmB*) retain native upstream regions, consistent with homologous recombination-mediated replacement at the native genomic region [29].

Unexpectedly, four functional foreign or chimeric genes or co-transcriptional units in *L. mirabile* and in *L. pyramidale* (*atp6*, *ccmC*, *nad6*, and *rpl5*-*rps14*) are embedded within extended mimosoid-derived tracts [32,56], suggesting they may use foreign promoters while still achieving significant transcription levels. Conversely, nine foreign or chimeric nonfunctional genes in *L. mirabile* (*atp8*, *atp9-two copies*, *cox3-sdh4*, *rpl10*, *rps4*, *rps12*, and *rps19*) and two in *L. pyramidale* (*atp9* and *rpl10*) are embedded in large foreign tracts [32,56], suggesting that while foreign promoters are present, their efficiency is highly variable, there is non-significant transcription in these cases.

### Nuclear genes involved in mitochondrial RNA maturation in *Lophophytum* spp. are native

Mitochondrial RNA maturation, encompassing intron splicing, transcript stabilization, and RNA editing, is largely mediated by nuclear-encoded factors, such as PPR proteins. We examined the possibility of horizontal acquisition of these factors to aid in the expression of foreign mitochondrial genes in *Lophophytum* spp. Homologs to *Arabidopsis* nuclear genes were identified in the transcriptomes of *Lophophytum* spp. and *Ombrophytum* (Table S4). Phylogenetic analyses confirm their native origin, with one exception in which native and foreign homologs co-exist (Figure S10). This finding indicates that the complex RNA maturation machinery required for the successful expression of HGT-acquired mitochondrial genes is almost entirely native in *Lophophytum* spp., suggesting high flexibility of the existing cellular components.

## DISCUSSION

The massive influx of horizontally acquired DNA in plant mitochondria is well-documented, yet it rarely translates into functional novelty [14,16,20]. This discrepancy establishes a central paradox: despite high tolerance for foreign DNA integration, severe expression barriers typically render these xenologs non-functional. Holoparasitic *Lophophytum* shatters this paradigm, representing the most extraordinary known case of functional mitochondrial HGT, with 6 foreign and 13 chimeric functional genes in *L. mirabile* and 4 foreign and 12 chimeric functional genes in *L. pyramidale* (Table 1). Here, we provide a comprehensive molecular framework explaining this functional assimilation, revealing that *Lophophytum* overcomes these barriers not by co-opting foreign nuclear machinery, but through a novel combination of structural gene chimerism and high adaptability of its native regulatory systems.

### Functional barriers to foreign gene expression

An outstanding question is how *Lophophytum* achieved this functional integration despite the well-known barriers to heterologous gene expression: promoter recognition, intron splicing, and RNA editing (Figure 1; [52]), in addition to the potential cytonuclear incompatibilities [58]. Our findings confirm these barriers were overcome, as evidenced by high transcription, efficient RNA editing, accurate splicing, and, in some cases, protein translation [29,48,59]. Furthermore, this functional integration scales to the protein-complex and physiological levels: mito-nuclear coevolution within oxidative phosphorylation complexes is not altered by HGT [60] and overall oxygen consumption rates are similar to those of free-living plants [59]. Given this robust functional assimilation, a functionally coherent possibility is that *Lophophytum* spp. may have co-acquired some nuclear-encoded regulatory factors from their hosts via HGT. However, this hypothesis is definitively refuted by our data. We found that all identified nuclear factors involved in mitochondrial expression (including RPOs, PPR proteins, and CRM-domain proteins) are of native origin. Furthermore, even though RNAs are known to move across haustorial connections in other parasitic plants [61–64], we found no evidence of host-derived mRNAs in the *Lophophytum* inflorescence transcriptome. This unequivocally demonstrates that the functional assimilation of dozens of foreign genes relies entirely on the pre-existing, native nuclear machinery of the parasite.

### Chimerism and promoter retention as drivers of functional integration

A primary barrier to functional HGT is transcription initiation [27]. Plant mitochondrial promoters are not universally conserved across angiosperms, and multiple promoters per gene with non-canonical motifs have been described [26,27]. One of the most striking findings of this study is that *Lophophytum* seems to bypass this barrier primarily through structural integration: the vast majority of functional foreign genes are, in fact, chimeric or part of a chimeric transcriptional unit. The distinction between fully foreign and chimeric genes, enabled by our new comparative data, is critical for two reasons. First, it provides direct evidence of the integration mechanism: the mosaic nature of chimeric genes strongly suggests they are produced by homologous recombination between foreign and native DNA, in contrast to fully foreign genes which may integrate at different loci via less frequent non-homologous repair pathways [65]. Second, this chimeric structure could be the key to their functional success. In nearly all functional cases, these genes retain native 5′ gene regions (Tables 1 and S6).

This 5′-native configuration strongly implies the retention of endogenous, recognizable promoters upstream of the foreign coding sequence, ensuring transcriptional compatibility with the native transcription machinery. Although direct mapping of mitochondrial transcription start sites was beyond the scope of this study, the consistent retention of the native 5’ region in functional genes makes the preservation of endogenous promoter sequences the most parsimonious explanation for their successful transcription. Furthermore, this mosaicism can simultaneously preserve other essential native signals, such as those for intron splicing or RNA editing sites. This prevalence of chimerism indicates that partial, recombination-mediated replacements are more readily tolerated than full replacements, providing immediate transcriptional activity and buffering potential cytonuclear incompatibilities. This conclusion is supported by other documented cases of functional chimerism in plant mitochondria [9–12,66] and highlights the need to re-evaluate inferences of complete gene replacement in other systems as more comparative genomic data become available.

### Genes with foreign promoters show contrasting transcriptional outcomes

While chimerism may be a successful bypass for promoter recognition, our data show it is not the only one. A few functional genes (i.e. *atp6*, *ccmC*, *nad6*, and *rpl5*-*rps14*) in both species are embedded in extended foreign tracts, suggesting they are transcribed from putatively foreign promoters. This suggests that the native transcription machinery of *Lophophytum* may exhibit sufficient flexibility to recognize at least some non-native promoter motifs [67], perhaps aided by the low substitution rates of *Lophophytum* mtDNA [60] and of other angiosperm mtDNAs [68]. This contrasts sharply with the 11 non-functional, duplicated foreign or chimeric genes, which are also embedded in large foreign tracts. In these cases, the low affinity of the native machinery for these specific foreign promoters may result in non-significant transcription [29], rendering them non-functional and destined to become pseudogenes. This contrasting transcription pattern of genes carrying foreign promoters raises the possibility that other molecular mechanisms regulate the expression of mitochondrial genes, such as transcription factors, methylation or non-coding RNAs [69–73].

Therefore, the differential expression observed in *Lophophytum*—where some foreign-promoter genes are active while others are silenced—may be determined by additional layers of transcriptional or post-transcriptional control that differ in their specificity.

### Compatibility of the native RNA editing machinery facilitates integration of xenologs

Besides transcription, accurate RNA editing is essential for functionality, as the disruption of specific sites can compromise vital protein functions [28,74,75]. Our results reveal a two-part solution to overcome this barrier. First, the lower editing efficiency seen in foreign regions is gene-intrinsic, not a failure of the *Lophophytum* machinery (Figure 3). This suggests that foreign genes with intrinsically lower RNA editing requirements are more likely to be functionally retained. In addition, chimerism enables the exclusion of unrecognized RNA editing sites of vital functional importance. Second, the Lophophytum editing machinery successfully recognizes and edits numerous novel, host-specific C-to-U sites (∼30% of non-synonymous sites). This demonstrates a remarkable flexibility, likely mediated by the broad specificity of native PPR proteins and accessory cofactors like MORF proteins [28,76,77], allowing the parasite to process the editing requirements of phylogenetically distant donor genes with likely distinct RNA editing site content [51]. This outcome supports findings showing that exogenous editing sites can be edited when introduced into phylogenetically distant mitochondria [78,79] or the nuclear context of an existing mitochondrial genome is altered [80]. Given that individual editing factors often target multiple sites [81], *Lophophytum*’s pre-existing machinery appears sufficiently plastic to edit new targets. Together, these findings indicate that functional integration is a highly selective process: xenologs whose editing requirements are either intrinsically low or can be edited by the native machinery are more likely to achieve functional integration, while sequences with incompatible editing patterns perish.

### The splicing likely represents a functional filter against foreign introns

Finally, splicing appears to act as a strong functional barrier that filters out foreign introns. Unlike the remarkable flexibility observed in the RNA editing machinery, the native splicing system appears highly recalcitrant to foreign sequences. Despite the 24 introns typically present in the mitochondrial genes of their mimosoid donors, we found no fully foreign introns in *Lophophytum* spp., and only short foreign patches in less than 25% of the introns of *Lophophytum* spp. The ancestral acquisition of an intronless *cox2* gene provides a notable example of this functional filter in action. While the mimosoid donors possess both intron-containing and intronless *cox2* copies, only the intronless version—which bypasses this splicing barrier entirely—was successfully integrated and functionally retained in the *Lophophytum* lineage. This finding is likely explained by the fact that most plant mitochondrial introns are Group II introns, for which very few cases of foreign or chimeric transfer have been reported [11,82,83]. The tight cytonuclear co-adaptation required for efficient splicing of these Group II introns, which are assisted by numerous nuclear-encoded factors [84,85], likely prevents their proper processing by the recipient’s endogenous machinery. This is consistent with our broader finding that no foreign nuclear maturation factors were co-transferred.

This stands in striking contrast to the Group I intron in the *cox1* gene, which has been horizontally transferred hundreds of times among angiosperms, likely aided by its intron-encoded homing endonuclease [25,86]. The critical difference could be that the *cox1* intron may require only a generalized environment rather than highly specialized, co-evolved protein factors for its excision [87]. This suggests that the success of functional HGT is biased towards foreign genes or gene pieces that lack specific splicing requirements.

## CONCLUSIONS

In conclusion, the case of *Lophophytum* provides a unique look into the functional assimilation of foreign mitochondrial genes, helping to explain how massive HGT can, in rare cases, lead to functional novelty. We show that this large-scale integration does not depend on the co-transfer of nuclear regulatory factors, as the *Lophophytum* expression machinery remains entirely native. Instead, functional success appears to be primarily achieved by the formation of chimeric genes. By retaining native 5′ regions, these chimeric structures likely keep promoter recognition by the native RNA polymerases. This structural solution is complemented by a highly selective process. The retention of foreign genes is not random, but favors xenologs that lack foreign introns that the native machinery cannot process, and possess intrinsically low RNA editing demands or RNA editing requirements that can be fulfilled. Taken together, our findings show that the successful integration of foreign genes is dictated by both structural compatibility and the inherent plasticity of the native host machinery. Future studies will be critical to resolve the role of additional regulatory layers, such as epigenetic modifications, additional transcription factors, or non-coding RNAs, in silencing or activating mitochondrial genes.

## Acknowledgements

This work used the SARTOI Cluster from IBAM (CONICET-UNCuyo).

## Author contributions

MER and MVS-P conceived and designed the research. MER, LFC, LG-S, and WDT performed data analysis. MER and MVS-P drafted and wrote the manuscript. MER, LFC, LG-S, and MVS-P revised the manuscript.

