## Supplementary Text for "A structural solution to functional HGT: Gene chimerism bypasses mitochondrial expression barriers in parasitic plants"

### ELECTRONIC SUPPLEMENTARY MATERIAL

The following Supporting Information is available for this article:

**Note S1: Mimosoid organellar genome characterization**

**Note S2: The native homologs of the foreign or chimeric mitochondrial genes in *Lophophytum* were not functionally transferred to the nuclear genome, except for *rps14***

**Figure S1. Total read depth and map of organellar genomes of *Anadenanthera colubrina*.** A. The mitochondrial genome (mtDNA) is 677,949 bp long. The average total read depth was 194x. Shown are full-length genes, pseudogenes (indicated by 'Ψ'), repeats > 1 kb (labeled 'R'), and sequences derived from the chloroplast (cp) longer than 100 bp. B. The chloroplast genome (ptDNA) is 163,911 bp long. The average total read depth was 1115.4x. Genes drawn inside and outside each circle are transcribed clockwise and counterclockwise, respectively. Below, circular genome is shown linearized for clarity and the curves depict the DNA read depth along the organellar genome. C. PCR primer pairs and conditions used for the validation of the assembly of the MTPTs in the *Anadenanthera* mtDNA. D. Agarose gel images showing the PCR products.

**Figure S2. Total read depth and map of organellar genomes of *Vachellia collinsii*.** A. The mitochondrial genome (mtDNA) is 674,577 bp long. The average total read depth was 2,071x. Shown are full-length genes, pseudogenes (indicated by 'Ψ'), repeats > 1 kb (labeled 'R'), and sequences derived from the chloroplast (cp) longer than 100 bp. B. The chloroplast genome (ptDNA) is 164,605 bp long. The average total read depth was 11,508x. Genes drawn inside and outside each circle are transcribed clockwise and counterclockwise, respectively. Below, circular genome is shown linearized for clarity and the curves depict the DNA read depth along the organellar genome.

**Figure S3. Maximum Likelihood (ML) phylogenetic analyses of mitochondrial protein coding genes in *Lophophytum* spp.** The phylogenetic trees for foreign, chimeric, and native genes in *Lophophytum* spp. are shown. ML bootstrap support values ≥50% are shown. The chromosome number is indicated for each *L. pyramidale* and *L. mirabile* gene. Pseudogenes are depicted by 'Ψ'. Scale bars correspond to substitutions per site. For chimeric genes the ML trees were performed based on native (in orange) or foreign (in purple) subregions of chimeric mitochondrial genes.

**Figure S4. RNA read depth of the 81 chromosomes in the *Lophophytum pyramidale* mtDNA.** The curves represent the number of RNA reads aligned to each nucleotide position. Reads corresponding to transcription from left to right and right to left are shown as green and orange shaded curves on the positive and negative y-axes, respectively. For each chromosome, an additional magnified plot Y-axis (*ylim* = 10,000) to enhance the visualization of regions with lower read coverage is included. Grey boxes indicate the locations of mitochondrial genes, plastid-derived sequences (cp-like), and repeats ("R" followed by the length of the repeat). The RNA reads cover 99.77% of the mitochondrial genome considering both strands. The coverage is not 100% likely due to a small difference in the mitochondrial genome of the individual from which the RNA was extracted and the individual used for mtDNA assembly. For each protein-coding gene, the phylogenetic origin is indicated: chimeric ("C"), foreign (F) or native (N). Noticeably, a few non-coding regions show elevated coverage, such as regions in *LpChr1A* (5–6 kb) and *LpChr04C* (3.5–5.5 kb) with over 10,000 aligned reads. BLASTn searches indicate that these regions are of foreign origin, associated with Fabaceae species, and do not contain any annotated coding sequences.

**Figure S5. Maximum Likelihood (ML) phylogenetic analyses of mitochondrial and nuclear *rps14* genes in *Lophophytum* spp. and *Ombrophytum*.** To determine the phylogenetic origin of the nuclear gene encoding the organellar ribosomal protein RPS14, homologous sequences were retrieved from photosynthetic species available in the PLAZA v5.0 database (https://bioinformatics.psb.ugent.be/plaza.dev/instances/dicots_05/). Codon-based alignments were generated using MAFFT v7.407 (Katoh & Standley, 2013) and PAL2NAL (Suyama et al., 2006). Poorly aligned or highly divergent regions were filtered out with BMGE v1.12 (Criscuolo & Gribaldo, 2010). The phylogenetic tree was reconstructed under a maximum likelihood framework with RAxML v8.2.11 (Stamatakis, 2014), employing the GTR+GAMMA+I substitution model and 1,000 bootstrap replicates. The resulting tree was visualized using FigTree (http://tree.bio.ed.ac.uk/software/figtree/).

**Figure S6. RNA read depth of protein-coding genes (CDSs) in the *Lophophytum pyramidale* mtDNA.** The curves show the number of RNA reads aligned to each nucleotide position. Each gene (represented as a grey rectangle) is extended by 300 nucleotides upstream and downstream to include flanking regions.

**Figure S7. Boxplots showing C-to-U RNA editing efficiency (%) at non-synonymous sites in native vs. foreign regions for (A) *Lophophytum pyramidale* and (B) *L. mirabile*.** Black line: median; white diamond: mean; *** $p < 0.001$.

**Figure S8. Maximum Likelihood (ML) phylogenetic analyses of RNA polymerases in *Lophophytum* spp. and *Ombrophytum*.** ML bootstrap support values ≥50% are shown. Scale bars correspond to substitutions per site. The plastid-targeted RpoTp was not found in the transcriptomes of *Lophophytum* spp. and *Ombrophytum*.

**Figure S9. Evolutionary model proposed for the mitochondrial gene clusters in Balanophoraceae mitochondria.** Red lines indicate losses of clusters or genes within a cluster.

**Figure S10. Maximum-likelihood (ML) phylogenetic analyses of the detected orthologs in *Lophophytum* spp. and *Ombrophytum*.** ML bootstrap support values ≥50% are shown next to nodes. Scale bars correspond to substitutions per site. The orthogroup identifiers corresponding to each gene (as listed in Table S4) are indicated in the upper-right corner of each tree. Color code: *Arabidopsis thaliana* (red), Fabaceae species (blue), Santalales (brown), *Lophophytum mirabile* (orange), *L. pyramidale* (yellow), and *Ombrophytum subterraneum* (pink).

**Table S1.** Assembly details of mimosoid species.

**Table S2.** Accession numbers and references of the mitochondrial sequences used in different analyses (A); phylogenetic analyses (B); Geneconv analyses (C).

**Table S3.** General features of the mitochondrial genome of mimosoids species.

**TableS4.** Proteins involved in RNA maturation in plant mitochondria.

**Table S5**. Phylogenetic origin of protein-coding genes in the *Lophophytum mirabile* mtDNA.

**Table S6**. Features of mitochondrial genes in the *Lophophytum pyramidale* mtDNA.

**Table S7.** RNA read depth of each mitochondrial chromosome of *Lophophytum pyramidale.*
**Table S8.** Total number of C-to-U non-synonymous RNA editing sites in *Lophophytum* spp., *Ombrophytum*, and the mimosoid *Acacia ligulata.*
**Table S9.** C-to-U RNA editing sites in putatively functional protein genes in *Lophophytum pyramidale* mtDNA.
**Table S10.** C-to-U RNA editing sites in mitochondrial genes in *Lophophytum mirabile* mtDNA.
**Table S11.** C-to-U RNA editing sites in native mitochondrial genes in *Ombrophytum subterraneum* mtDNA.

**Note S1: Mimosoid organellar genome characterization**

Organellar genomes were annotated using Geneious v.11.1.5. The mitochondrial or plastid origin of the tRNAs was determined by BLASTn searches. Plastid-derived mitochondrial sequences (MTPTs) > 200 bp were detected by BLASTn searches against a custom angiosperm chloroplast database, including those assembled in this study. Some MTPTs found in the *Anadenanthera* mtDNA were larger than the library size (> 250 pb). To corroborate the assembly, we designed primers to validate the length of the MTPT by PCR (Figure S1C-D). For those MTPTs smaller than the library size, the presence of paired-end reads having one mate mapping to the flanking mitochondrial region of the MTPT and the other mate in the opposite flanking region of the MTPT support the assembly of the plastid-derived regions in the *Anadenanthera* mtDNA. The repetitive content of both mitochondrial genomes was analyzed using the *get_repeats.sh* script developed by Gandini *et al.* (2019) and the maps were generated using OGDraw software [1].

Mitochondrial genomes (mtDNAs) of *Anandenanthera* and *Vachellia* were assembled into one circular chromosome (674-677 kb in length; Table S3) with an average read depth of 194x and 2017x, respectively (Figure S1A-S2A). Both mtDNAs show a continuous read depth except for those regions with spikes corresponding to plastid-derived mitochondrial sequences (MTPTs). The GC content of both mtDNAs is 45% and they contain 36 unique protein, 3 rRNA, and 20-25 tRNA-encoding genes (Table S3). Besides, both genomes contain 17 cis-splicing and 6 trans-splicing group II introns and the repetitive content occupies 12-16% of the genome and 11% of the total mtDNA corresponds to coding genes (Table S3). The plastid genome of both legumes was assembled into a single circular chromosome of 163,911 bp in length for *Anadenanthera* and 164,605 bp for *Vachellia*. Both cpDNAs show a continuous read depth along the genome (Figure S1B-S2B).

**Note S2: The native homologs of the foreign or chimeric mitochondrial genes in *Lophophytum* were not functionally transferred to the nuclear genome, except for *rps14***

To test whether genes with lower transcription levels could be compensated by nuclear-encoded homologs (*ccmFC*, *rps4*, and *rps14*), we searched for functional nuclear transfers of the mitochondrial genes. Homology searches of these genes in the transcriptome of *L. pyramidale* found no nuclear-encoded homologs, except for *rps14* (Figure S5). We conducted BLASTn and tBLASTn searches against the transcriptomes of *Lophophytum* spp. and *Ombrophytum subterraneum*, using the three ribosomal *rps14* genes of *Arabidopsis thaliana* as queries: plastid-encoded (ATCG00330), three lineage specific nuclear copies (AT4G33865, AT3G44010, and AT3G43980), and a nuclear-encoded mitochondrial-targeted gene (AT2G34520). Based on previous results [2], we retained transcripts with > 40% identity and > 60% query coverage to minimize the risk of missing divergent homologs. Open reading frames (ORFs) were predicted using Geneious v.2023.1.1, and homology was confirmed through reciprocal BLASTn searches against the *Arabidopsis* genome. In both *Lophophytum* species and in *Ombrophytum*, we found multiple transcripts of the mitochondrial *rps14* genes: one transcript derived from the mitochondrial-encoded *rps14* and three from nuclear-encoded copies of *rps14*. One of the latter is affiliated to mitochondrial-targeted nuclear-encoded *rps14* genes, such as that of *Arabidopsis* [3]. These results suggest that these copies originated from an ancestral intracellular transfer of the *rps14* from the mtDNA to the nuclear genome in the ancestor of *Lophophytum* and *Ombrophytum*. It is unclear whether the mitochondrial-encoded, the nuclear-encoded mitochondrial-targeted RPS14 or both are functional in *Lophophytum* spp. In contrast, *Ombrophytum* does not carry a functional mitochondrial-encoded *rps14* gene and the nuclear-encoded copy is likely functional.
