## Supplementary Figures for "A structural solution to functional HGT: Gene chimerism bypasses mitochondrial expression barriers in parasitic plants"

**Figure S1. Total read depth and map of organellar genomes of *Anadenanthera colubrina*. A.** The mitochondrial genome (mtDNA) is 677,949 bp long. The average total read depth was 194x. Shown are full-length genes, pseudogenes (indicated by 'Ψ'), repeats > 1 kb (labeled 'R'), and sequences derived from the chloroplast (cp) longer than 100 bp. **B.** The chloroplast genome (ptDNA) is 163,911 bp long. The average total read depth was 1115.4x. Genes drawn inside and outside each circle are transcribed clockwise and counterclockwise, respectively. Below, circular genome is shown linearized for clarity and the curves depict the DNA read depth along the organellar genome. **C.** PCR primer pairs and conditions used for the validation of the assembly of the MTPTs in the *Anadenanthera* mtDNA. **D.** Agarose gel images showing the PCR products.

**A**

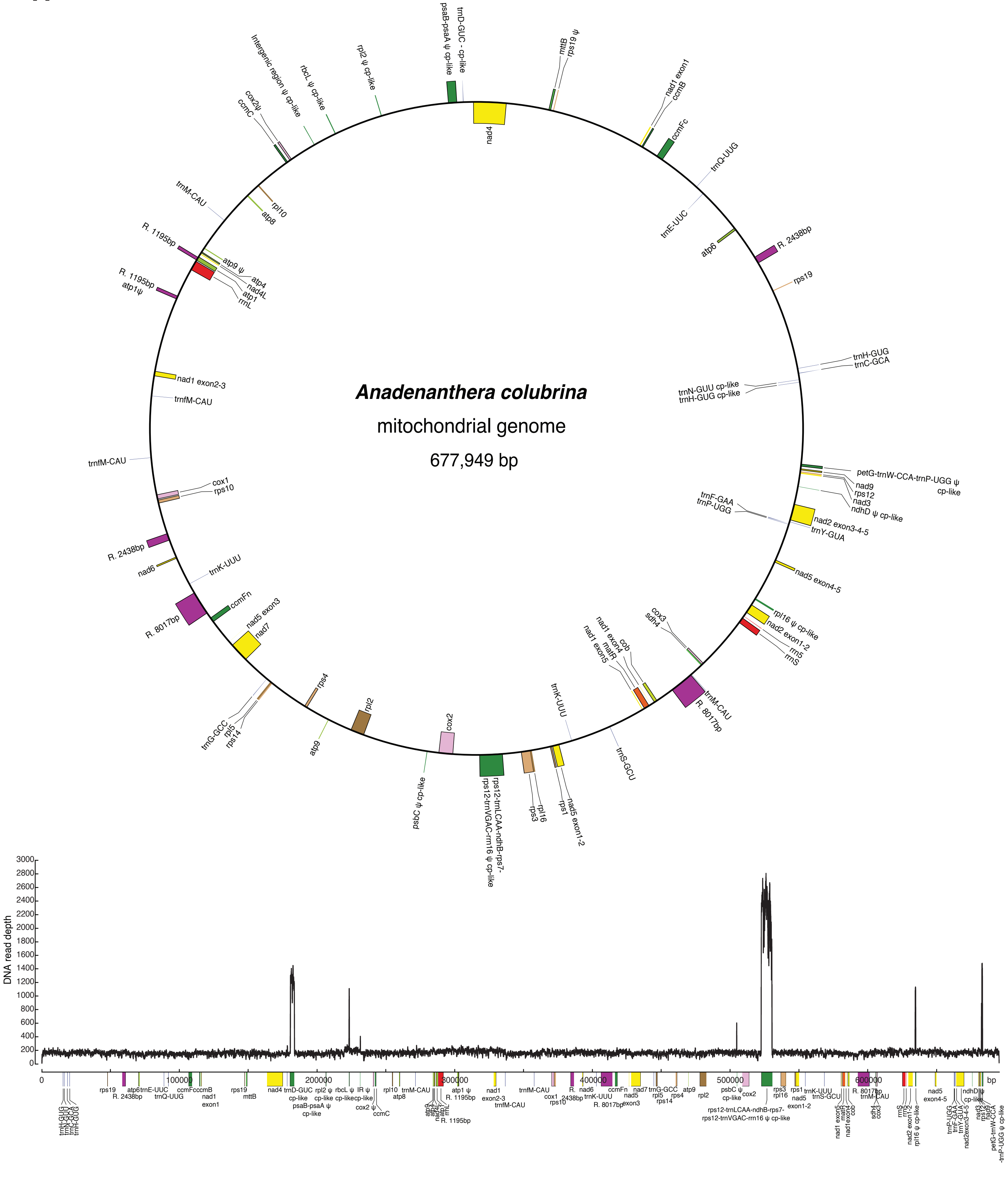

**B**

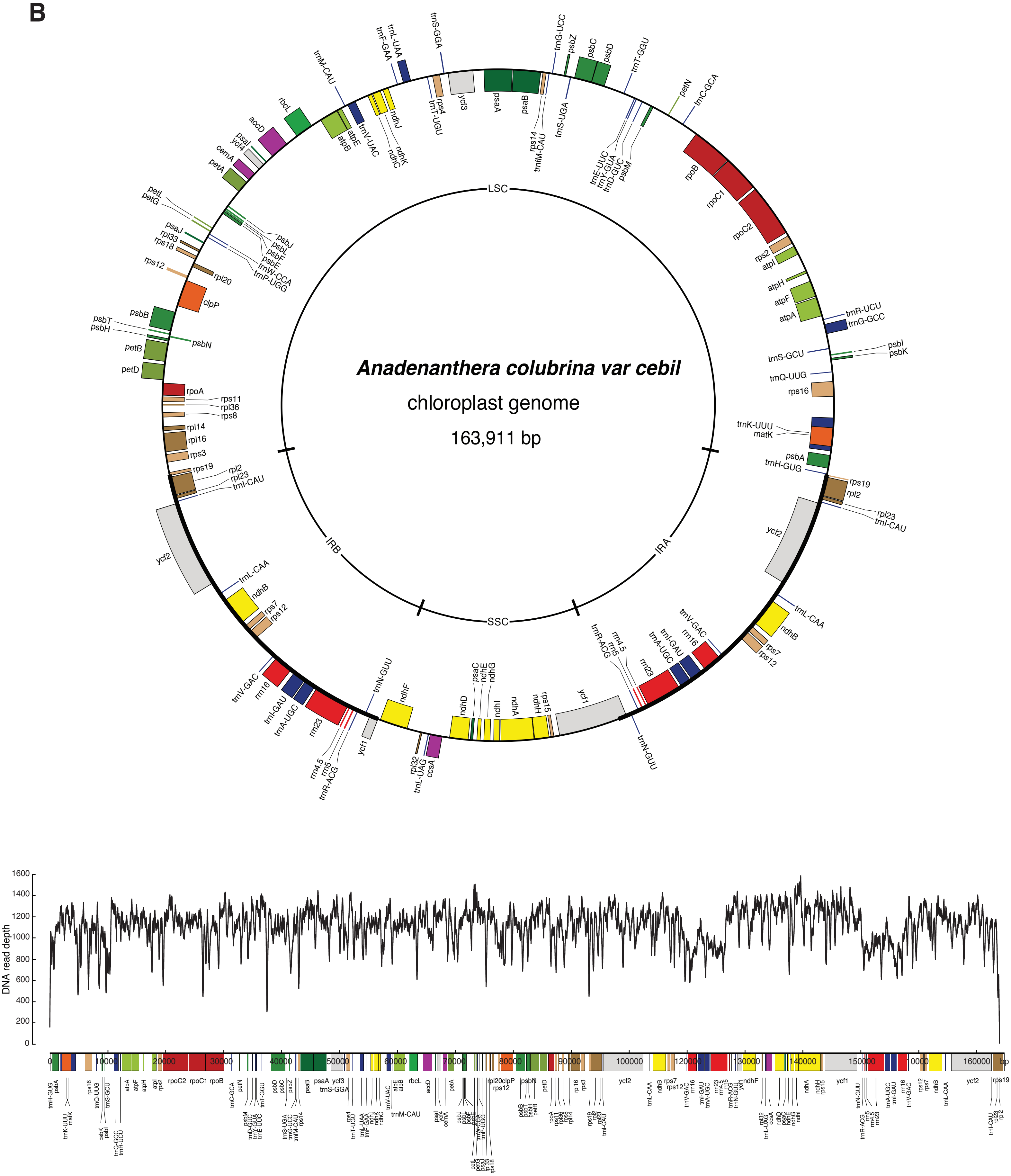

Anadenanthera colubrina var cebil
chloroplast genome
163,911 bp
LSC
SSC
IRA
IRB
DNA read depth
0
200
400
600
800
1000
1200
1400
1600
bp

**C.** PCR primer pairs and conditions used for the validation of the assembly of the MTPTs in the *A. colubrina* mtDNA

| Oligo name | Sequence | Length | Start | End | Tm | %GC | Product length |
|---|---|---|---|---|---|---|---|
| MTPT1-5'_F | GCCCATCACATTCTAGTGTGC | 21 | 509357 | 509377 | 59.33 | 52.38 | 3971 |
| MTPT1-5'_R | AAGCACTTGCCATTCGTTGG | 20 | 513308 | 513327 | 59.69 | 50 | 3971 |
| MTPT1-3'_F | TATGGCATCGCCACTCCCTT | 20 | 513212 | 513231 | 61.34 | 55 | 3938 |
| MTPT1-3'_R | ATTGGGTGGGTCGATTCAAGC | 21 | 517129 | 517149 | 60.96 | 52.38 | 3938 |
| MTPT2_F | CGGGGAAGCCTCGCTTATTC | 20 | 665251 | 665270 | 60.88 | 60 | 998 |
| MTPT2_R | GATGCCCAGGGTGTCCATTG | 20 | 666229 | 666248 | 61.04 | 60 | 998 |
| MTPT3_F | TACCGGGTGCATTCATTGGT | 20 | 175812 | 175831 | 59.67 | 50 | 3223 |
| MTPT3_R | AAGGCTCTGACTCTTGCACC | 20 | 179015 | 179034 | 59.96 | 55 | 3223 |
| MTPT4_F | CCAGCCACGAGGGTCAAATA | 20 | 200089 | 200108 | 59.75 | 55 | 1515 |
| MTPT4_R | GACACCCAAGCATCAACTCG | 20 | 201584 | 201603 | 59.2 | 55 | 1515 |

**MTPT1**

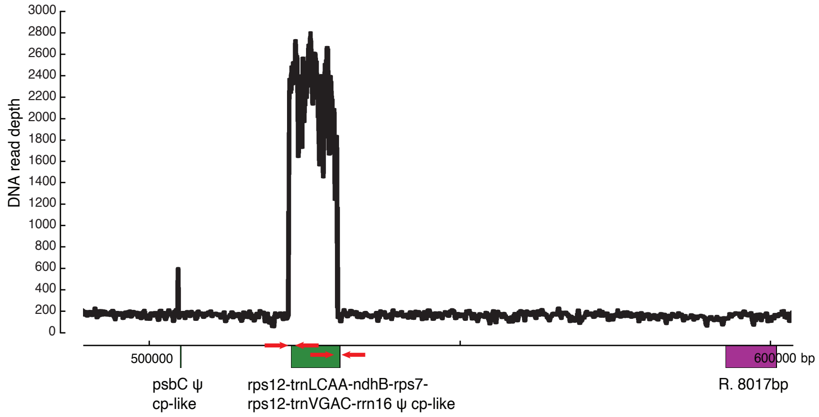


**MTPT2**

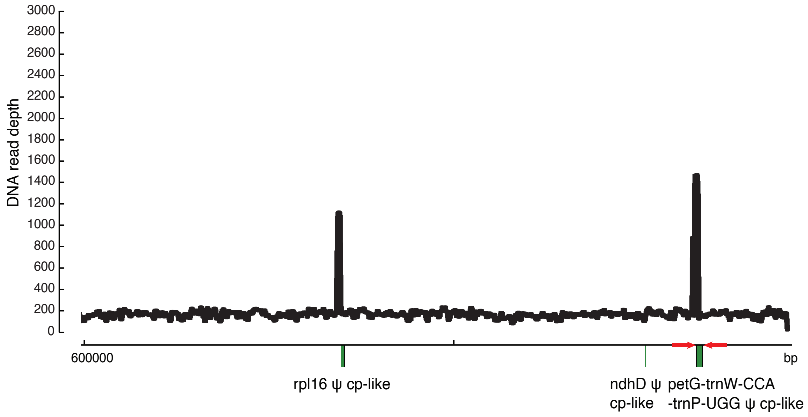


**MTPT3 - MTPT4**

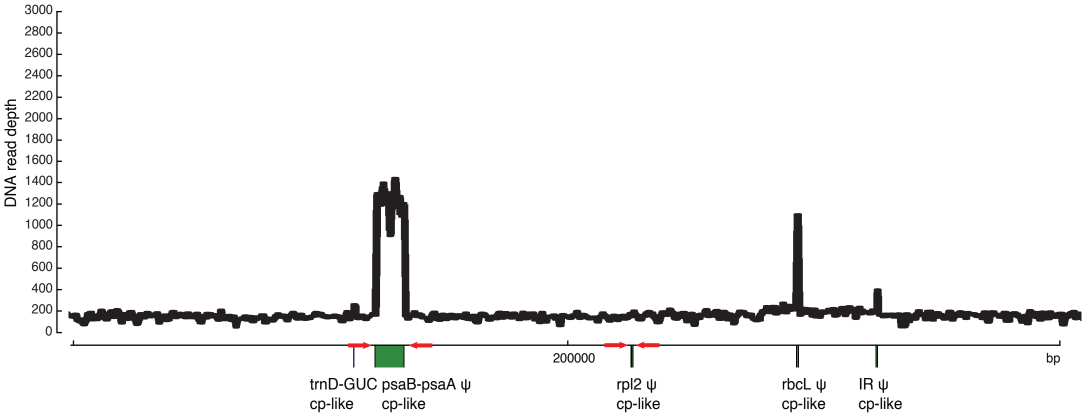

**D**

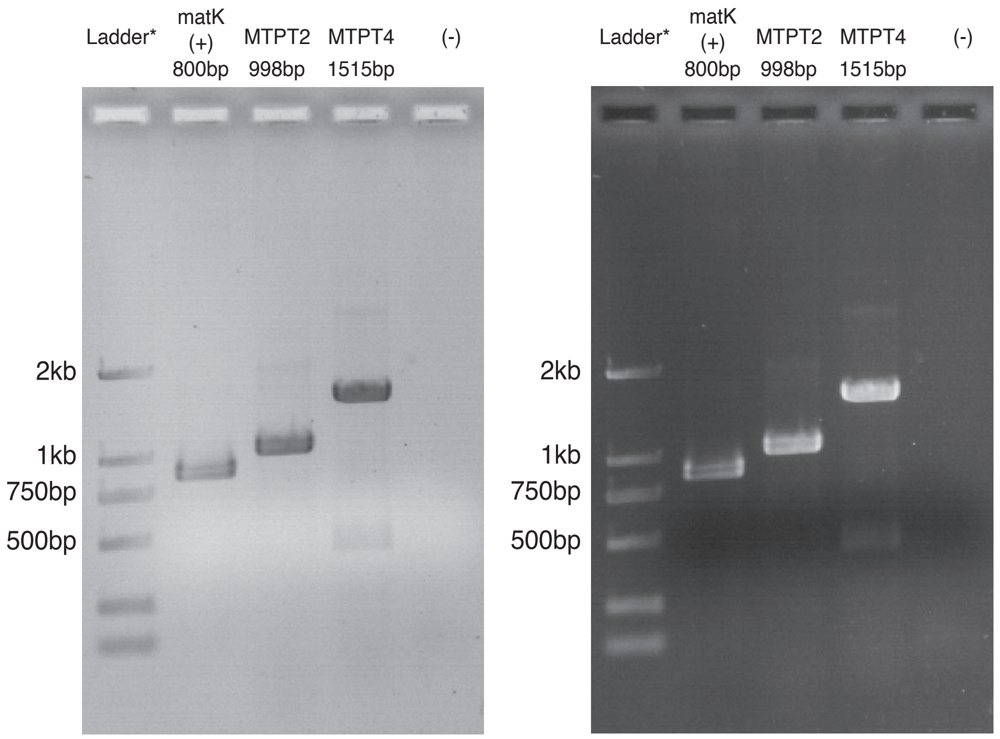


MTPT2 -> petG-trnW-CCA-trnP-UGG. Length MTPT: 889 bp

MTPT4 -> rpl2. Length MTPT: 256 bp

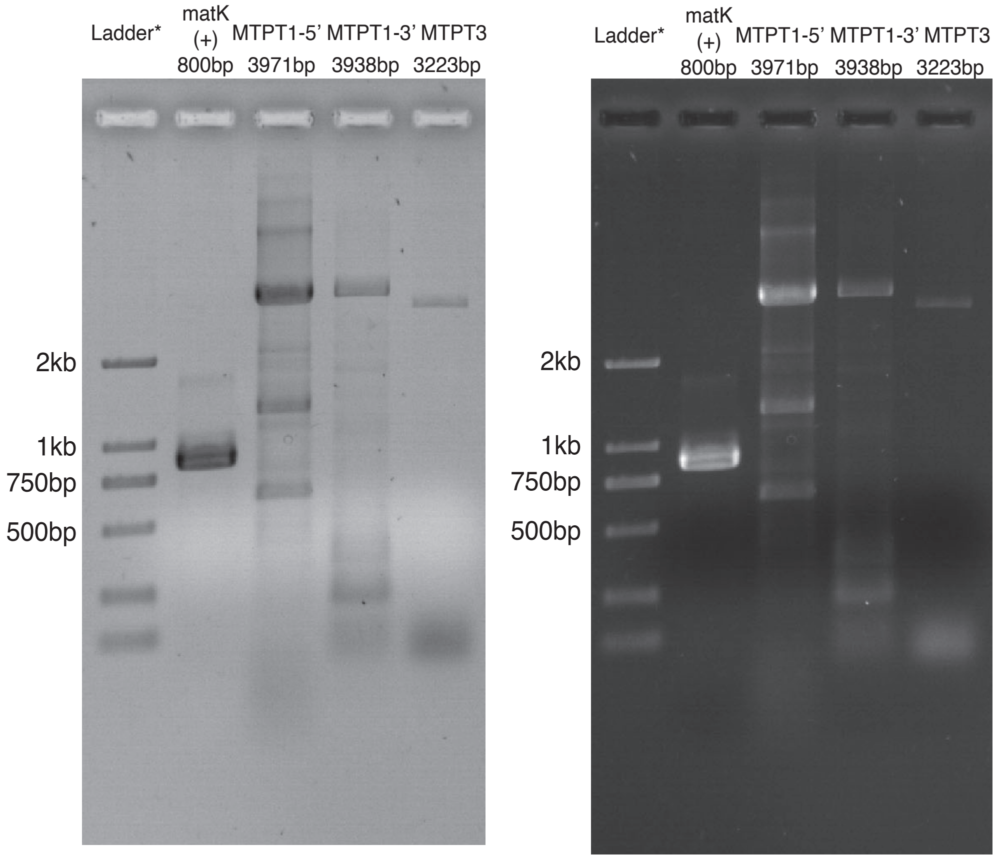


MTPT1 - 5’ region -> rps12-trnLCAA-ndhB-rps7-rps12-trnVGAC-rrn16. Length MTPT: 7692 bp

MTPT1 - 3’ region -> rps12-trnLCAA-ndhB-rps7-rps12-trnVGAC-rrn16. Length MTPT: 7692 bp

MTPT3 -> psaB-psaA. Length MTPT: 2952 bp

*Trans2K® DNA Marker

**Figure S2. Total read depth and map of organellar genomes of *Vachellia collinsii*. A.** The mitochondrial genome (mtDNA) is 674,577 bp long. The average total read depth was 2,071x. Shown are full-length genes, pseudogenes (indicated by 'Ψ'), repeats > 1 kb (labeled 'R'), and sequences derived from the chloroplast (cp) longer than 100 bp. **B.** The chloroplast genome (ptDNA) is 164,605 bp long. The average total read depth was 11,508x. Genes drawn inside and outside each circle are transcribed clockwise and counterclockwise, respectively. Below, circular genome is shown linearized for clarity and the curves depict the DNA read depth along the organellar genome.

# A

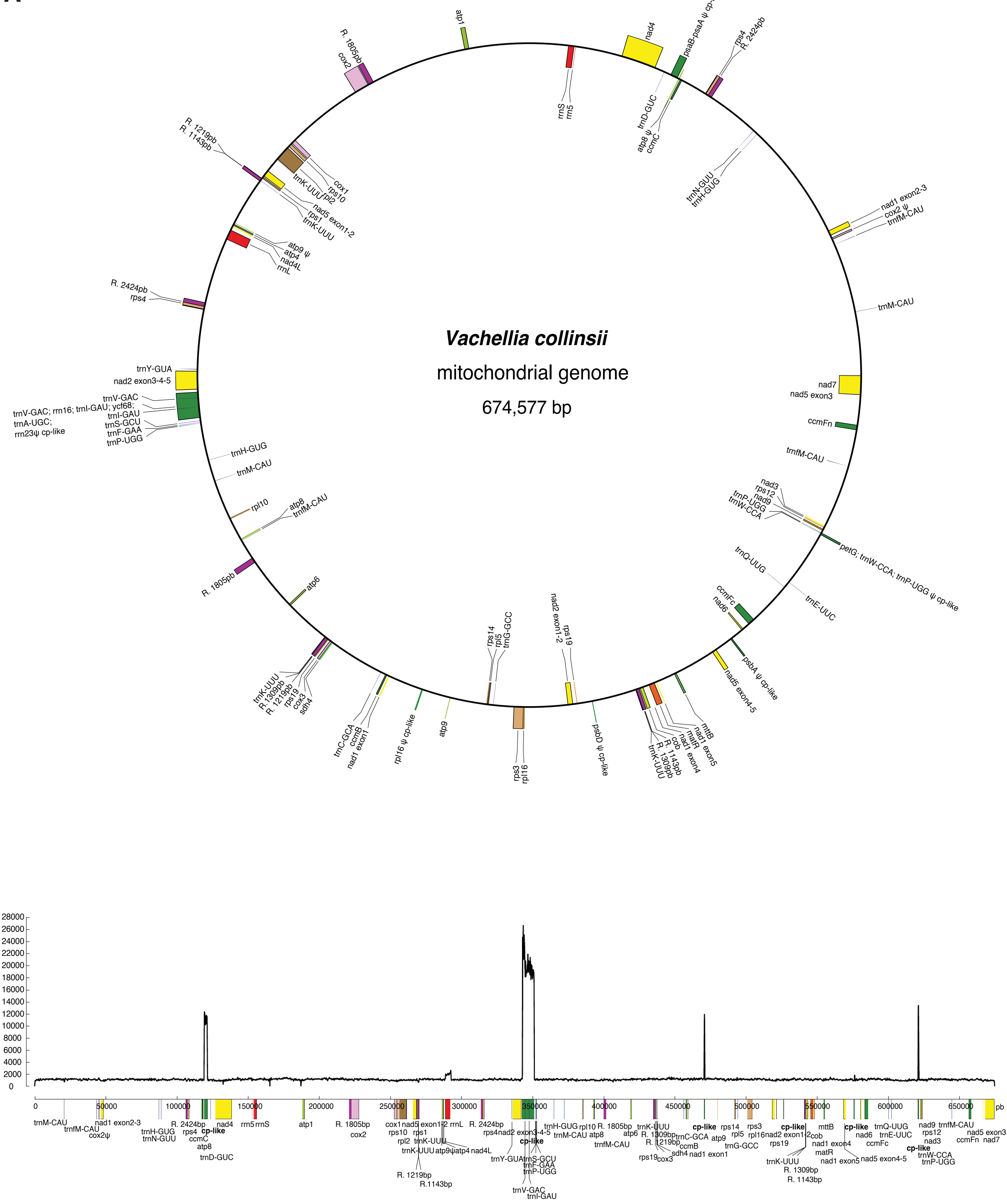

B

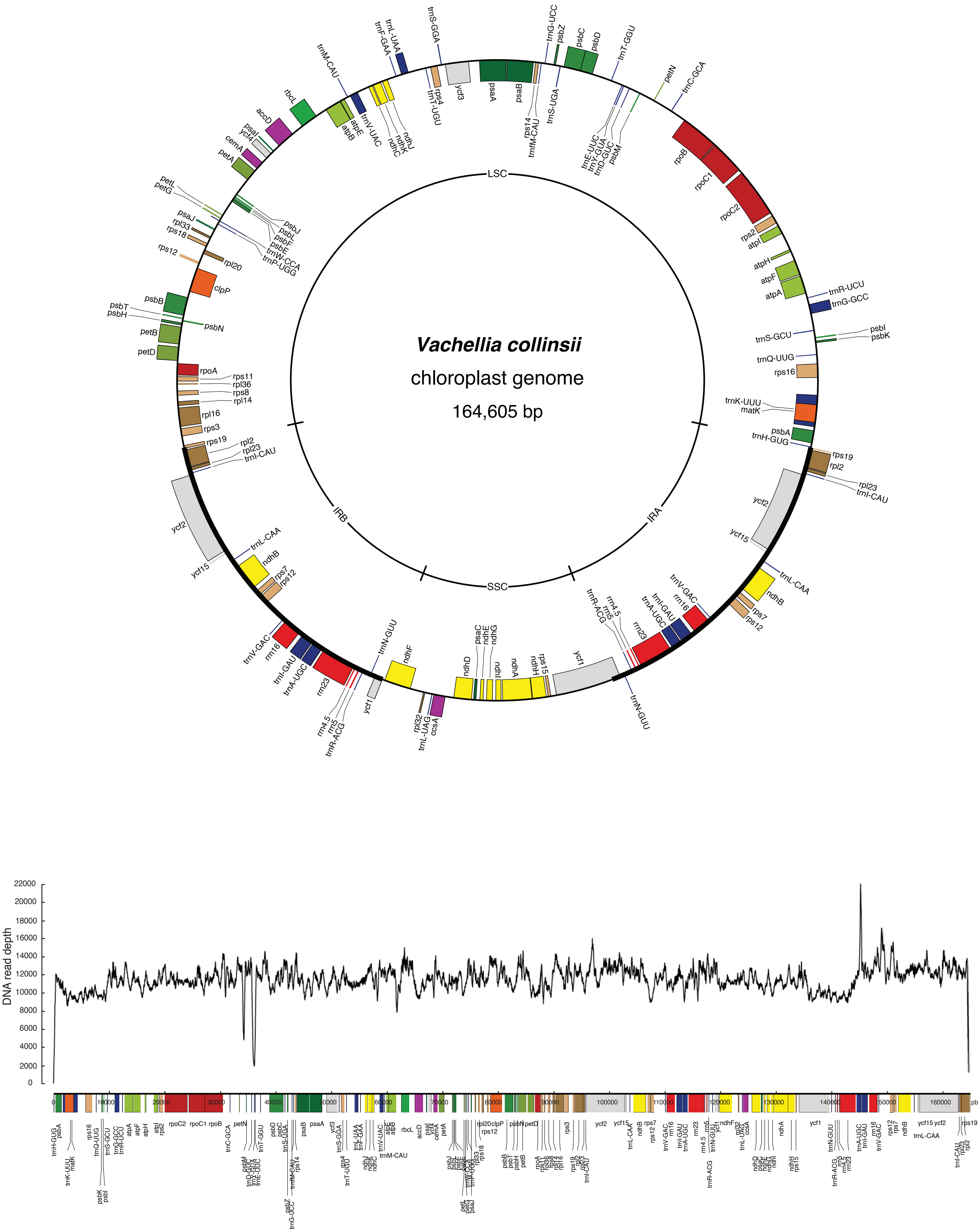

Vachellia collinsii
chloroplast genome
164,605 bp
LSC
IRA
SSC
IRB
trnH-GUG
psbA
matK
trnK-UUU
rps16
trnQ-UUG
trnS-GCU
psbI
psbK
trnG-GCC
trnR-UCU
atpA
atpF
atpH
atpI
rps2
rpoC2
rpoC1
rpoB
trnC-GCA
petN
psbM
trnD-GUC
trnY-GUA
trnE-UUC
trnT-GGU
psbD
psbC
psbZ
trnG-UCC
trnS-UGA
trnfM-CAU
rps14
psaB
psaA
ycf3
trnS-GGA
rps4
trnT-UGU
trnL-UAA
trnF-GAA
ndhJ
ndhK
ndhC
trnV-UAC
trnM-CAU
atpE
atpB
rbcL
accD
psaI
ycf4
cemA
petA
petL
petG
psbJ
psbL
psbF
psbE
trnW-CCA
trnP-UGG
psaJ
rpl33
rps18
rps12
rpl20
clpP
psbB
psbT
psbH
psbN
petB
petD
rpoA
rps11
rpl36
rps8
rpl14
rpl16
rps3
rps19
rpl2
rpl23
trnI-CAU
ycf2
ycf15
trnL-CAA
ndhB
rps7
trnV-GAC
rrn16
trnI-GAU
trnA-UGC
rrn23
rrn4.5
rrn5
trnR-ACG
trnN-GUU
ycf1
ndhF
rpl32
trnL-UAG
ccsA
ndhD
psaC
ndhE
ndhG
ndhI
ndhA
ndhH
rps15
DNA read depth
22000
20000
18000
16000
14000
12000
10000
8000
6000
4000
2000
0
10000
20000
30000
40000
50000
60000
70000
80000
90000
100000
110000
120000
130000
140000
150000
160000

**Figure S3. Maximum Likelihood (ML) phylogenetic analyses of mitochondrial genes in *Lophophytum* spp.**
The phylogenetic trees for foreign, chimeric, and native genes in *Lophophytum* spp. are shown. ML bootstrap support values ≥50% are shown. The chromosome number is indicated for each *L. pyramidale* and *L.mirabile* gene. Pseudogenes are depicted by 'Ψ'. Scale bars correspond to substitutions per site. For chimeric genes the ML trees were performed based on native (in orange) or foreign (in purple) subregions of chimeric mitochondrial genes.

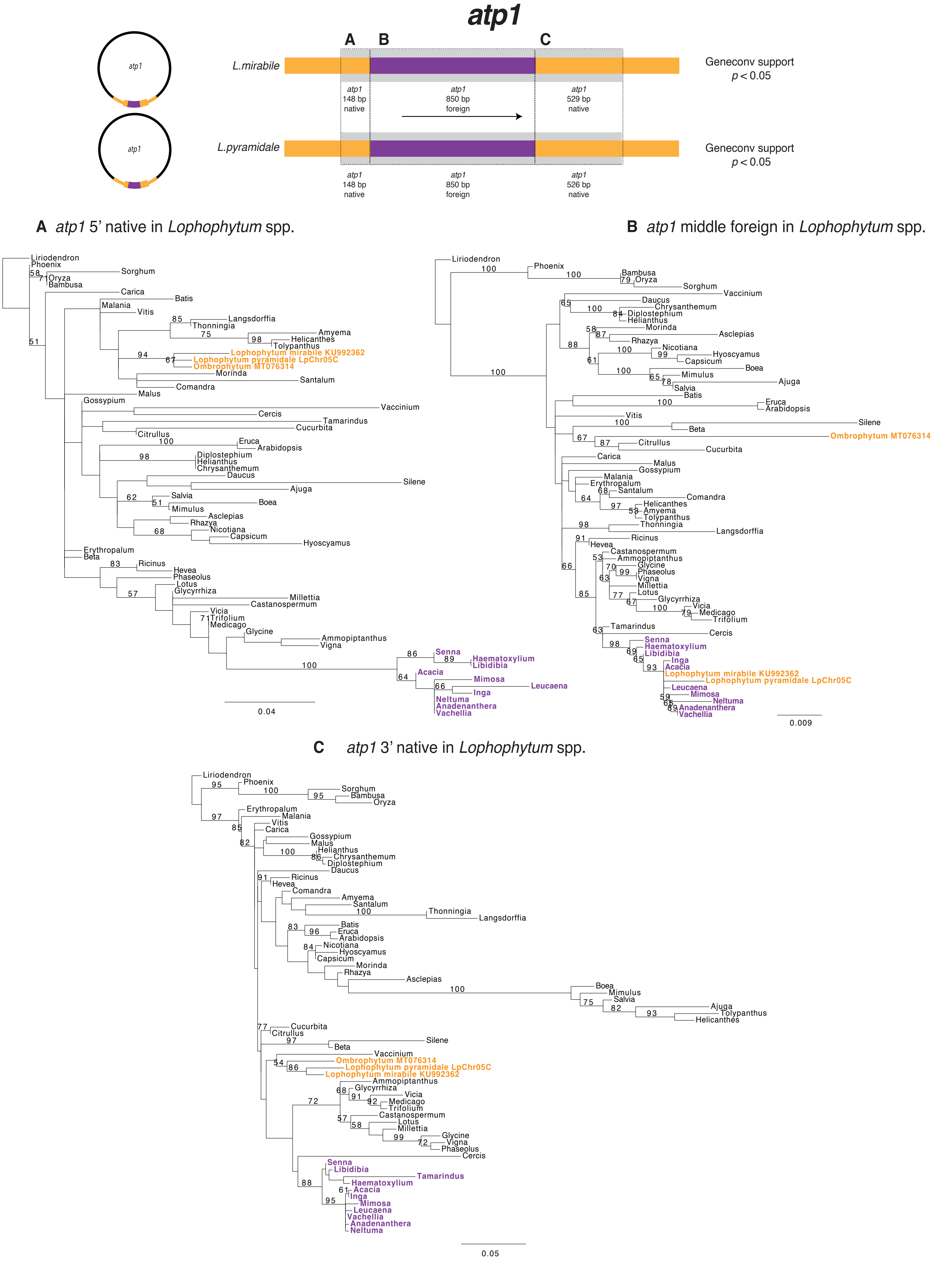

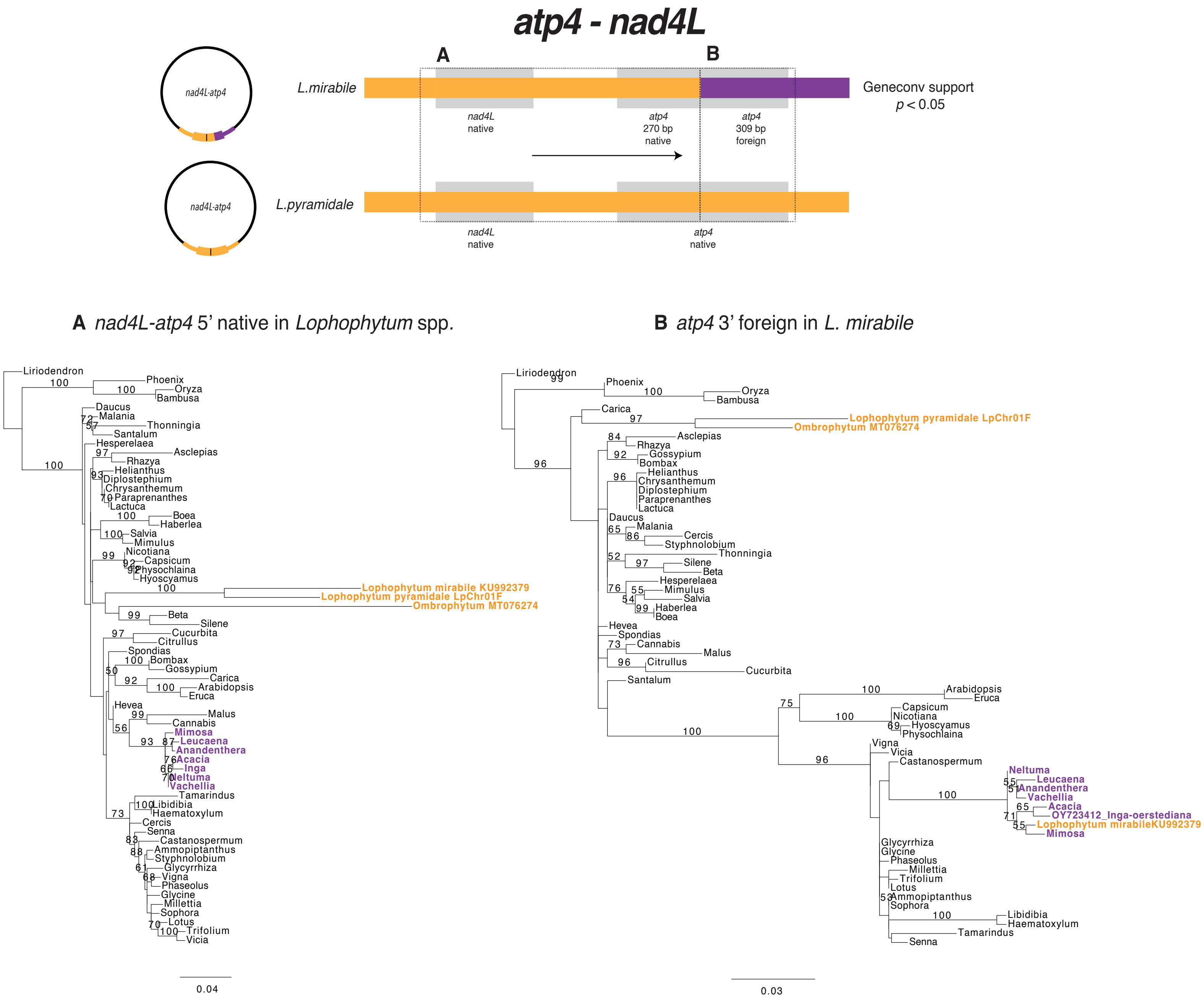

atp4 - nad4L
A
B
nad4L-atp4
L.mirabile
L.pyramidale
Geneconv support
p < 0.05
nad4L
native
atp4
270 bp
native
atp4
309 bp
foreign
nad4L
native
atp4
native
A nad4L-atp4 5' native in Lophophytum spp.
B atp4 3' foreign in L. mirabile
Liriodendron
Phoenix
Oryza
Bambusa
Daucus
Malania
Thonningia
Santalum
Hesperelaea
Asclepias
Rhazya
Helianthus
Diplostephium
Chrysanthemum
Paraprenanthes
Lactuca
Boea
Haberlea
Salvia
Mimulus
Nicotiana
Capsicum
Physochlaina
Hyoscyamus
Lophophytum mirabile KU992379
Lophophytum pyramidale LpChr01F
Ombrophytum MT076274
Beta
Silene
Cucurbita
Citrullus
Spondias
Bombax
Gossypium
Carica
Arabidopsis
Eruca
Hevea
Malus
Cannabis
Mimosa
Leucaena
Anandenthera
Acacia
Inga
Neltuma
Vachellia
Tamarindus
Libidibia
Haematoxylum
Cercis
Senna
Castanospermum
Ammopiptanthus
Styphnolobium
Glycyrrhiza
Vigna
Phaseolus
Glycine
Millettia
Sophora
Lotus
Trifolium
Vicia
0.04
Liriodendron
Phoenix
Oryza
Bambusa
Carica
Lophophytum pyramidale LpChr01F
Ombrophytum MT076274
Asclepias
Rhazya
Gossypium
Bombax
Helianthus
Chrysanthemum
Diplostephium
Paraprenanthes
Lactuca
Daucus
Malania
Cercis
Styphnolobium
Thonningia
Silene
Beta
Hesperelaea
Mimulus
Salvia
Haberlea
Boea
Hevea
Spondias
Cannabis
Malus
Citrullus
Cucurbita
Santalum
Arabidopsis
Eruca
Capsicum
Nicotiana
Hyoscyamus
Physochlaina
Vigna
Vicia
Castanospermum
Neltuma
Leucaena
Anandenthera
Vachellia
Acacia
OY723412_Inga-oerstediana
Lophophytum mirabileKU992379
Mimosa
Glycyrrhiza
Glycine
Phaseolus
Millettia
Trifolium
Lotus
Ammopiptanthus
Sophora
Libidibia
Haematoxylum
Tamarindus
Senna
0.03

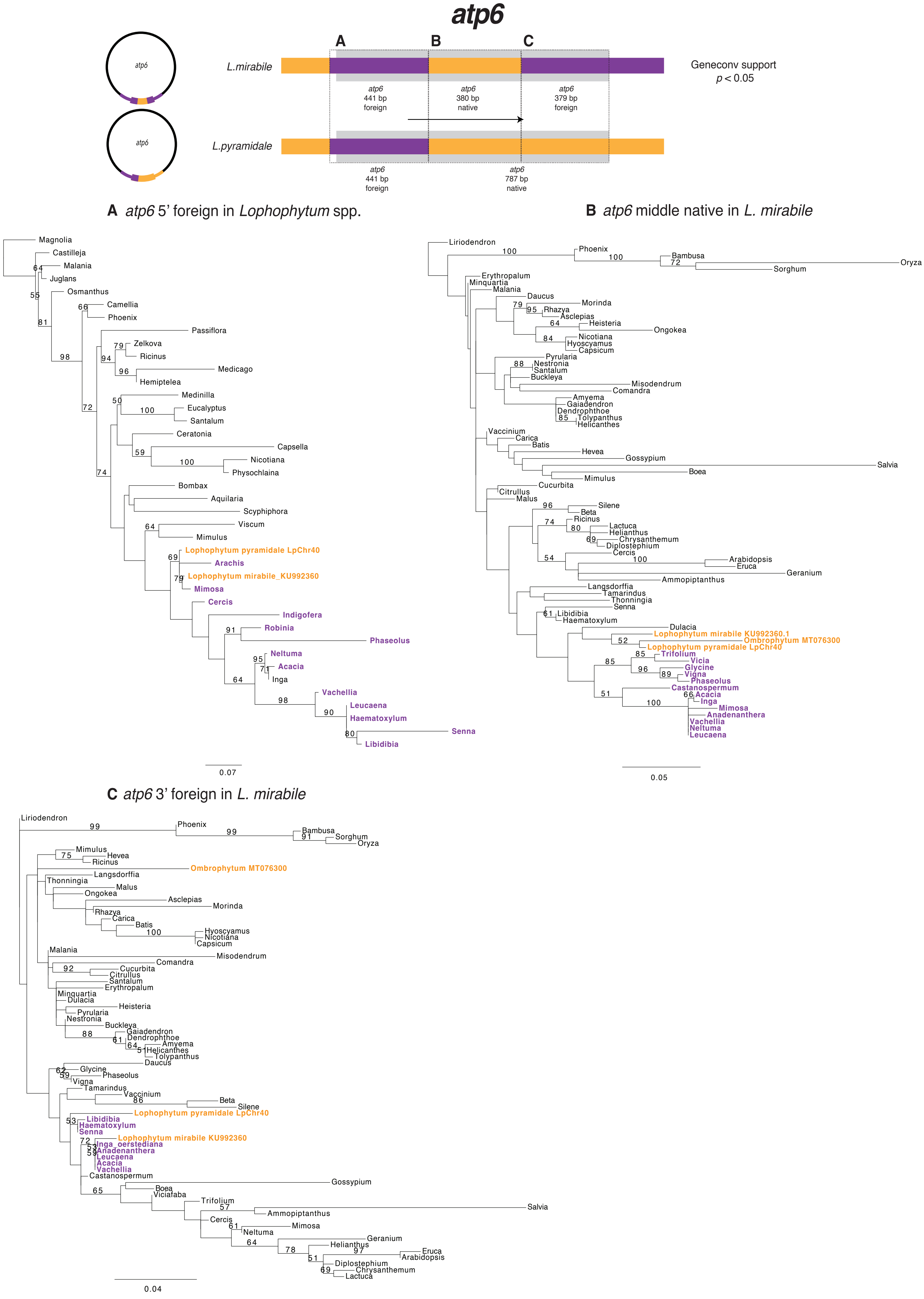

atp6
A
B
C
L.mirabile
L.pyramidale
Geneconv support
p < 0.05
atp6 441 bp foreign
atp6 380 bp native
atp6 379 bp foreign
atp6 441 bp foreign
atp6 787 bp native
A atp6 5' foreign in Lophophytum spp.
Magnolia
Castilleja
Malania
Juglans
Osmanthus
Camellia
Phoenix
Passiflora
Zelkova
Ricinus
Medicago
Hemiptelea
Medinilla
Eucalyptus
Santalum
Ceratonia
Capsella
Nicotiana
Physochlaina
Bombax
Aquilaria
Scyphiphora
Viscum
Mimulus
Lophophytum pyramidale LpChr40
Arachis
Lophophytum mirabile_KU992360
Mimosa
Cercis
Indigofera
Robinia
Phaseolus
Neltuma
Acacia
Inga
Vachellia
Leucaena
Haematoxylum
Senna
Libidibia
0.07
B atp6 middle native in L. mirabile
Liriodendron
Phoenix
Bambusa
Oryza
Sorghum
Erythropalum
Minquartia
Malania
Daucus
Morinda
Rhazya
Asclepias
Heisteria
Ongokea
Nicotiana
Hyoscyamus
Capsicum
Pyrularia
Nestronia
Santalum
Buckleya
Misodendrum
Comandra
Amyema
Gaiadendron
Dendrophthoe
Tolypanthus
Helicanthes
Vaccinium
Carica
Batis
Hevea
Gossypium
Salvia
Boea
Mimulus
Cucurbita
Citrullus
Malus
Silene
Beta
Ricinus
Lactuca
Helianthus
Chrysanthemum
Diplostephium
Cercis
Arabidopsis
Eruca
Geranium
Ammopiptanthus
Langsdorffia
Tamarindus
Thonningia
Senna
Libidibia
Haematoxylum
Dulacia
Lophophytum mirabile KU992360.1
Ombrophytum MT076300
Lophophytum pyramidale LpChr40
Trifolium
Vicia
Glycine
Vigna
Phaseolus
Castanospermum
Acacia
Inga
Mimosa
Anadenanthera
Vachellia
Neltuma
Leucaena
0.05
C atp6 3' foreign in L. mirabile
Liriodendron
Phoenix
Bambusa
Sorghum
Oryza
Mimulus
Hevea
Ricinus
Ombrophytum MT076300
Langsdorffia
Thonningia
Malus
Ongokea
Asclepias
Morinda
Rhazya
Carica
Batis
Hyoscyamus
Nicotiana
Capsicum
Malania
Misodendrum
Comandra
Cucurbita
Citrullus
Santalum
Erythropalum
Minquartia
Dulacia
Heisteria
Pyrularia
Nestronia
Buckleya
Gaiadendron
Dendrophthoe
Amyema
Helicanthes
Tolypanthus
Daucus
Glycine
Phaseolus
Vigna
Tamarindus
Vaccinium
Beta
Silene
Lophophytum pyramidale LpChr40
Libidibia
Haematoxylum
Senna
Lophophytum mirabile KU992360
Inga_oerstediana
Anadenanthera
Leucaena
Acacia
Vachellia
Castanospermum
Gossypium
Boea
Viciafaba
Trifolium
Salvia
Ammopiptanthus
Cercis
Mimosa
Neltuma
Geranium
Helianthus
Eruca
Arabidopsis
Diplostephium
Chrysanthemum
Lactuca
0.04

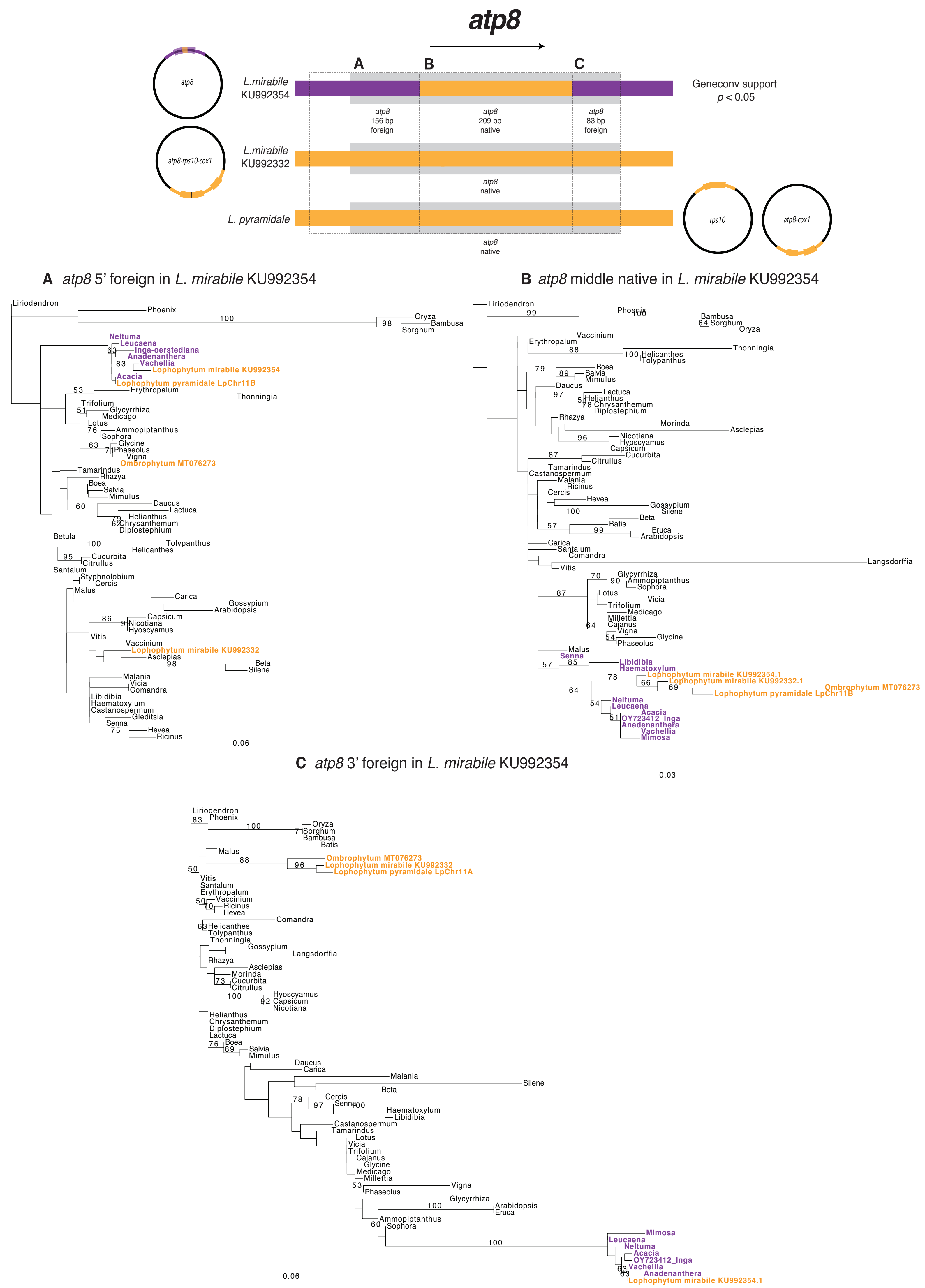
atp8
A
B
C
atp8
L.mirabile KU992354
atp8 156 bp foreign
atp8 209 bp native
atp8 83 bp foreign
Geneconv support $p < 0.05$
atp8-rps10-cox1
L.mirabile KU992332
atp8 native
L. pyramidale
atp8 native
rps10
atp8-cox1
A atp8 5' foreign in L. mirabile KU992354
Liriodendron
Phoenix
Oryza
Bambusa
Sorghum
Neltuma
Leucaena
Inga-oerstediana
Anadenanthera
Vachellia
Lophophytum mirabile KU992354
Acacia
Lophophytum pyramidale LpChr11B
Erythropalum
Thonningia
Trifolium
Glycyrrhiza
Medicago
Lotus
Ammopiptanthus
Sophora
Glycine
Phaseolus
Vigna
Ombrophytum MT076273
Tamarindus
Rhazya
Boea
Salvia
Mimulus
Daucus
Lactuca
Helianthus
Chrysanthemum
Diplostephium
Betula
Tolypanthus
Helicanthes
Cucurbita
Citrullus
Santalum
Styphnolobium
Cercis
Malus
Carica
Gossypium
Arabidopsis
Capsicum
Nicotiana
Hyoscyamus
Vitis
Vaccinium
Lophophytum mirabile KU992332
Asclepias
Beta
Silene
Malania
Vicia
Comandra
Libidibia
Haematoxylum
Castanospermum
Gleditsia
Senna
Hevea
Ricinus
0.06
B atp8 middle native in L. mirabile KU992354
Liriodendron
Phoenix
Bambusa
Sorghum
Oryza
Vaccinium
Erythropalum
Thonningia
Helicanthes
Tolypanthus
Boea
Salvia
Mimulus
Daucus
Lactuca
Helianthus
Chrysanthemum
Diplostephium
Rhazya
Morinda
Asclepias
Nicotiana
Hyoscyamus
Capsicum
Cucurbita
Citrullus
Tamarindus
Castanospermum
Malania
Ricinus
Cercis
Hevea
Gossypium
Silene
Beta
Batis
Eruca
Arabidopsis
Carica
Santalum
Comandra
Langsdorffia
Vitis
Glycyrrhiza
Ammopiptanthus
Sophora
Lotus
Vicia
Trifolium
Medicago
Millettia
Cajanus
Vigna
Glycine
Phaseolus
Malus
Senna
Libidibia
Haematoxylum
Lophophytum mirabile KU992354.1
Lophophytum mirabile KU992332.1
Ombrophytum MT076273
Lophophytum pyramidale LpChr11B
Neltuma
Leucaena
Acacia
OY723412_Inga
Anadenanthera
Vachellia
Mimosa
0.03
C atp8 3' foreign in L. mirabile KU992354
Liriodendron
Phoenix
Oryza
Sorghum
Bambusa
Batis
Malus
Ombrophytum MT076273
Lophophytum mirabile KU992332
Lophophytum pyramidale LpChr11A
Vitis
Santalum
Erythropalum
Vaccinium
Ricinus
Hevea
Comandra
Helicanthes
Tolypanthus
Thonningia
Gossypium
Langsdorffia
Rhazya
Asclepias
Morinda
Cucurbita
Citrullus
Hyoscyamus
Capsicum
Nicotiana
Helianthus
Chrysanthemum
Diplostephium
Lactuca
Boea
Salvia
Mimulus
Daucus
Carica
Malania
Silene
Beta
Cercis
Senna
Haematoxylum
Libidibia
Castanospermum
Tamarindus
Lotus
Vicia
Trifolium
Cajanus
Glycine
Medicago
Millettia
Vigna
Phaseolus
Glycyrrhiza
Arabidopsis
Eruca
Ammopiptanthus
Sophora
Mimosa
Leucaena
Neltuma
Acacia
OY723412_Inga
Vachellia
Anadenanthera
Lophophytum mirabile KU992354.1
0.06

### *atp9* ; *nad5x1x2*

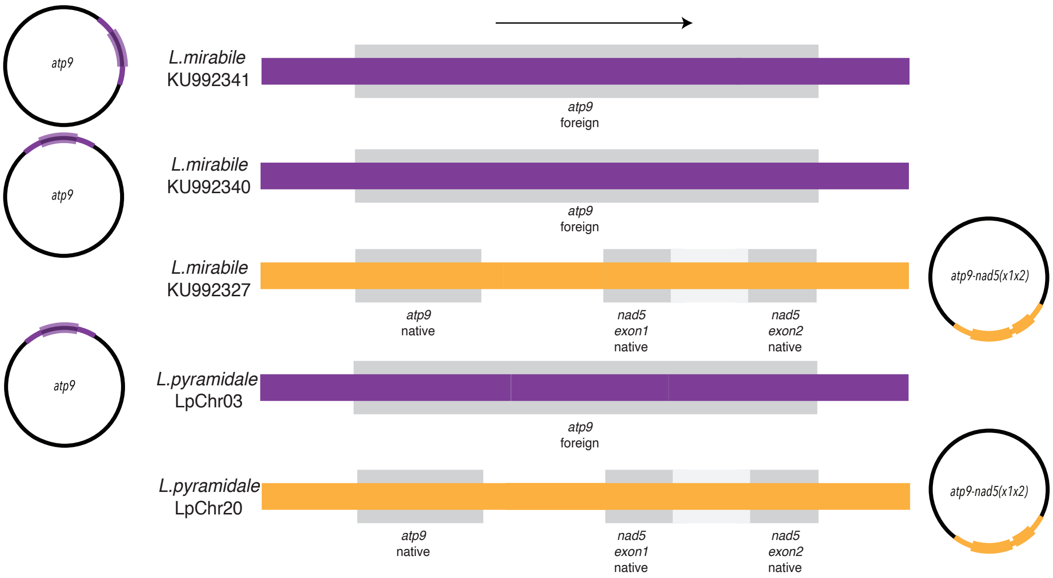


#### *atp9*

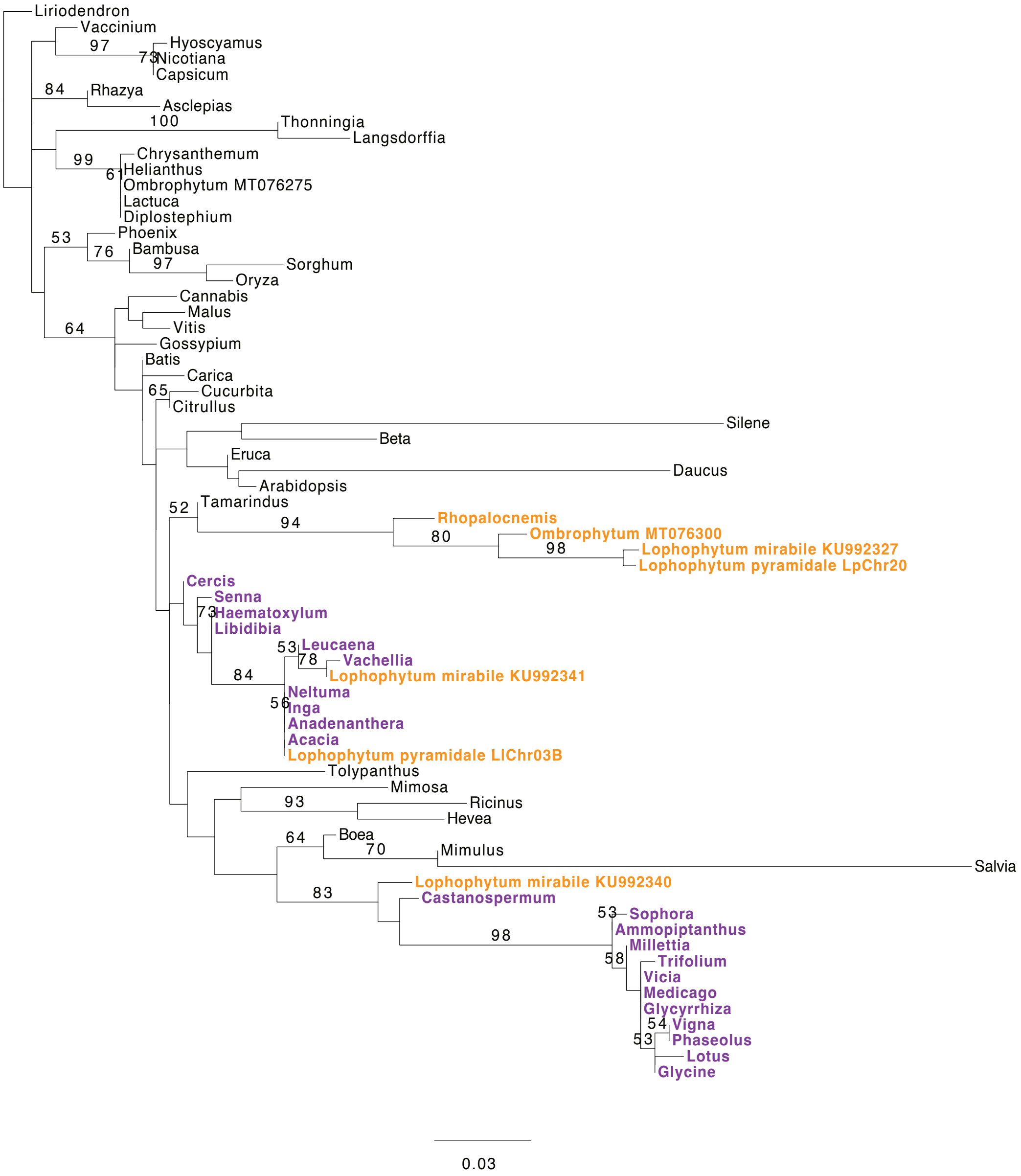

### cox1

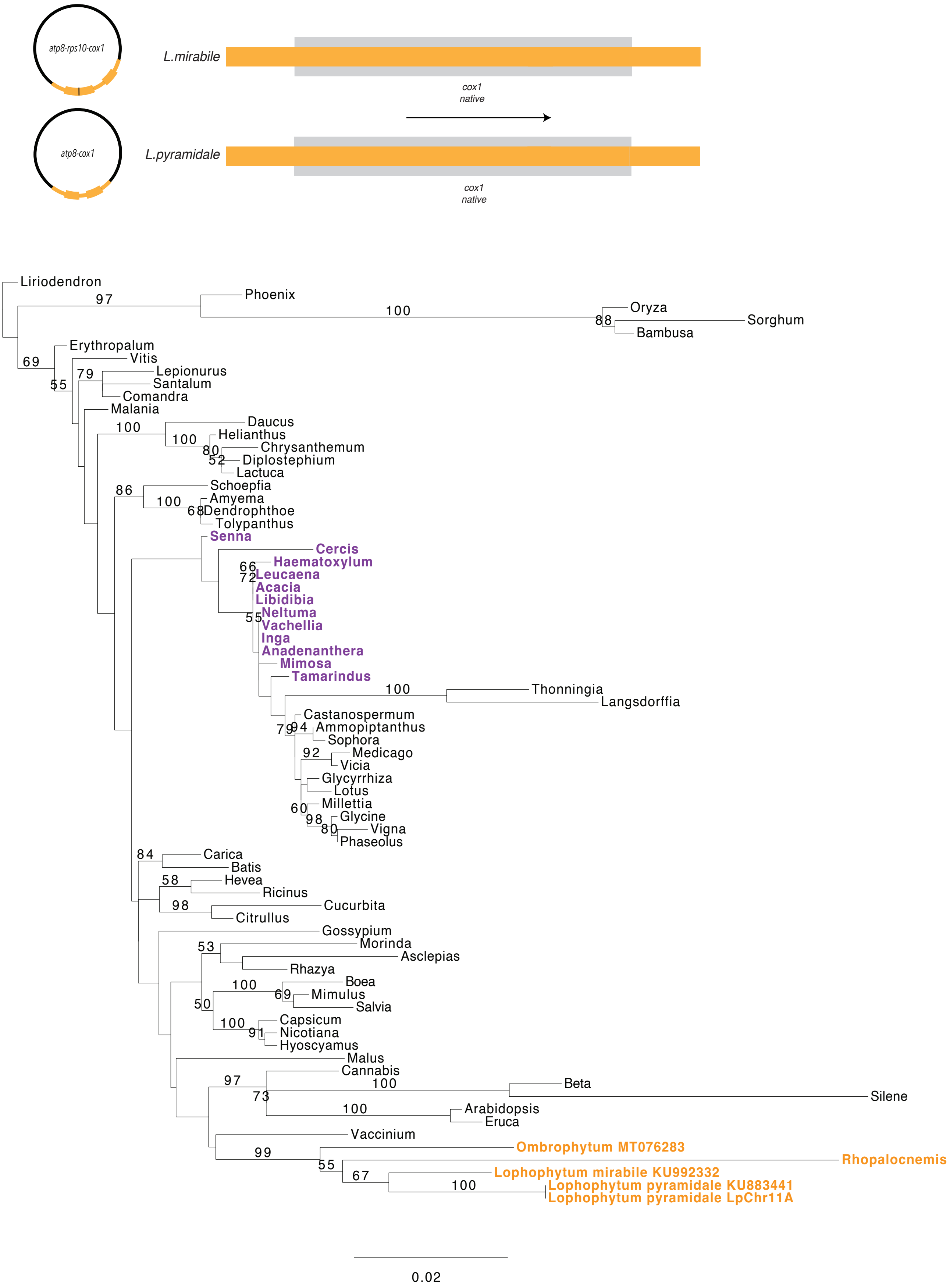
atp8-rps10-cox1
L.mirabile
cox1
native
atp8-cox1
L.pyramidale
cox1
native
Liriodendron
Phoenix
Oryza
Sorghum
Bambusa
Erythropalum
Vitis
Lepionurus
Santalum
Comandra
Malania
Daucus
Helianthus
Chrysanthemum
Diplostephium
Lactuca
Schoepfia
Amyema
Dendrophthoe
Tolypanthus
Senna
Cercis
Haematoxylum
Leucaena
Acacia
Libidibia
Neltuma
Vachellia
Inga
Anadenanthera
Mimosa
Tamarindus
Thonningia
Langsdorffia
Castanospermum
Ammopiptanthus
Sophora
Medicago
Vicia
Glycyrrhiza
Lotus
Millettia
Glycine
Vigna
Phaseolus
Carica
Batis
Hevea
Ricinus
Cucurbita
Citrullus
Gossypium
Morinda
Asclepias
Rhazya
Boea
Mimulus
Salvia
Capsicum
Nicotiana
Hyoscyamus
Malus
Cannabis
Beta
Silene
Arabidopsis
Eruca
Vaccinium
Ombrophytum MT076283
Rhopalocnemis
Lophophytum mirabile KU992332
Lophophytum pyramidale KU883441
Lophophytum pyramidale LpChr11A
0.02

### *cox2 ; ccmB*

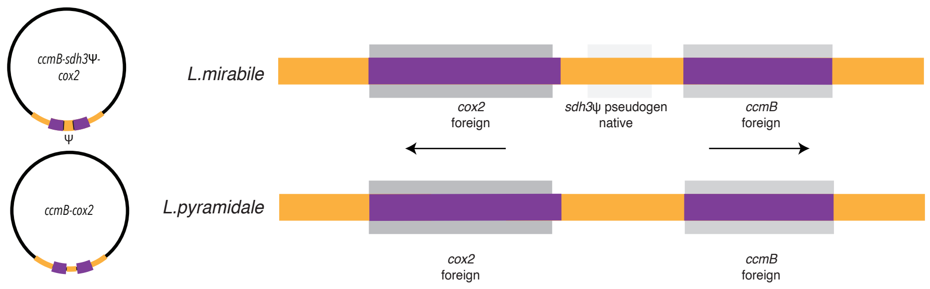


#### *cox2*

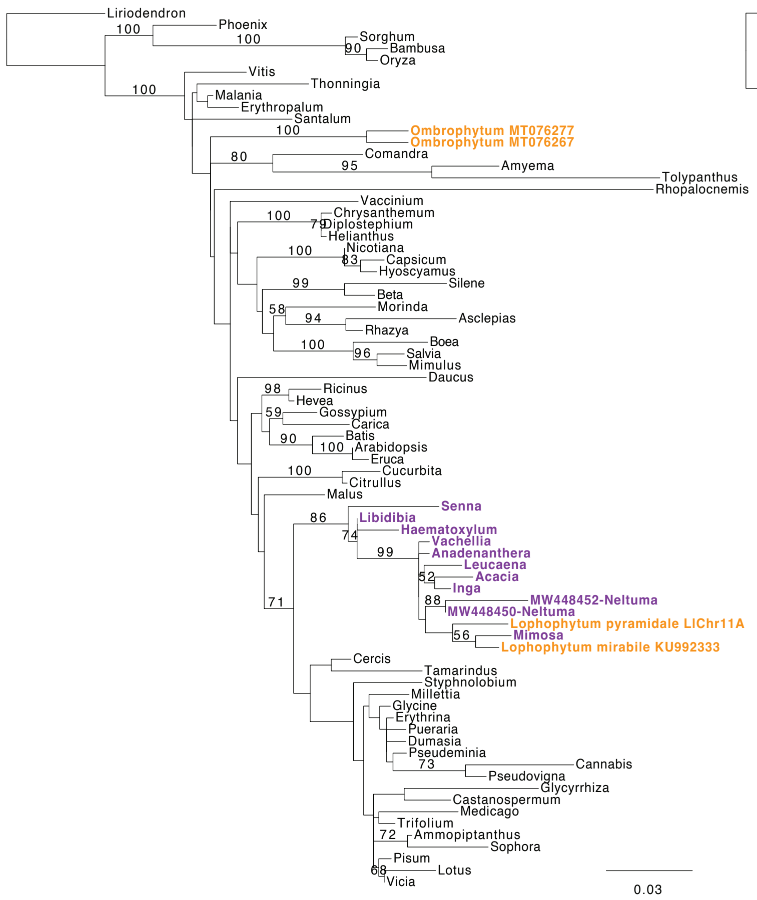


#### *ccmB*

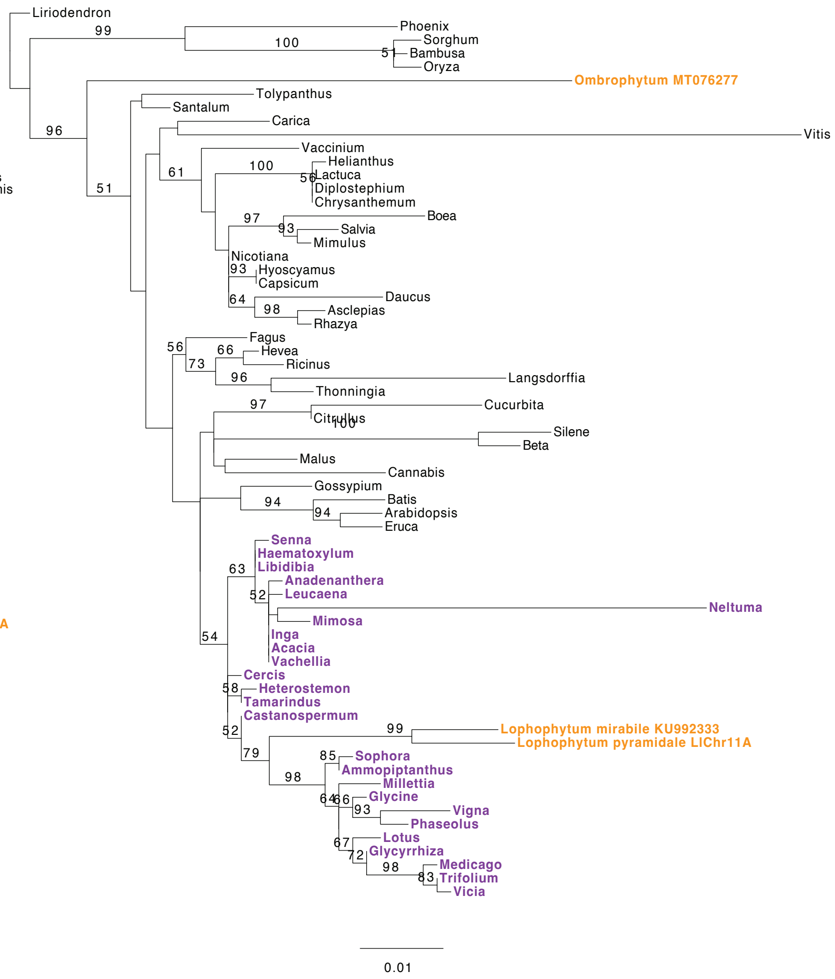

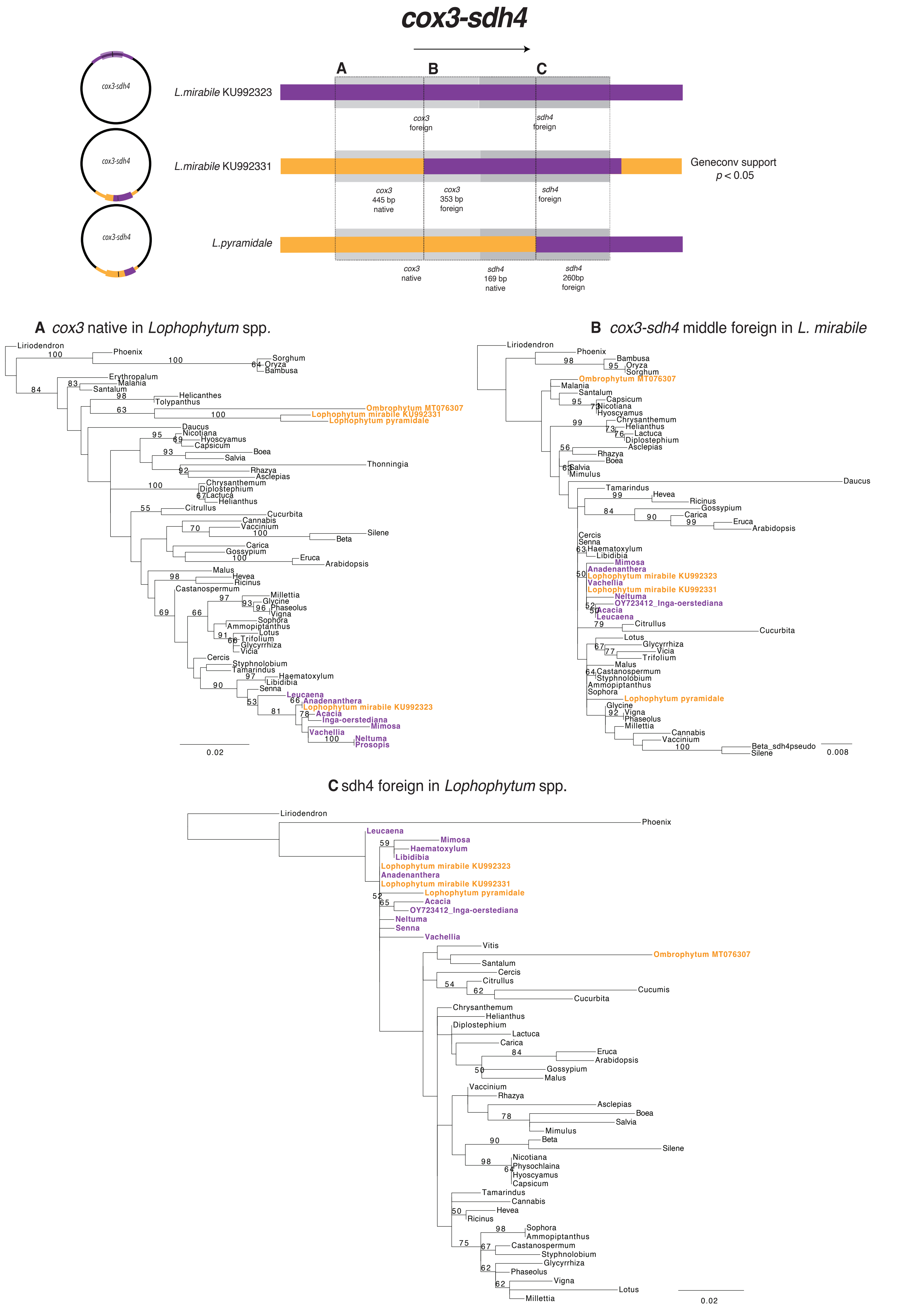
cox3-sdh4
cox3-sdh4
cox3-sdh4
cox3-sdh4
A
B
C
L.mirabile KU992323
cox3 foreign
sdh4 foreign
L.mirabile KU992331
Geneconv support
p < 0.05
cox3 445 bp native
cox3 353 bp foreign
sdh4 foreign
L.pyramidale
cox3 native
sdh4 169 bp native
sdh4 260bp foreign
A cox3 native in Lophophytum spp.
Liriodendron
Phoenix
Sorghum
Oryza
Bambusa
Erythropalum
Malania
Santalum
Helicanthes
Tolypanthus
Ombrophytum MT076307
Lophophytum mirabile KU992331
Lophophytum pyramidale
Daucus
Nicotiana
Hyoscyamus
Capsicum
Boea
Salvia
Thonningia
Rhazya
Asclepias
Chrysanthemum
Diplostephium
Lactuca
Helianthus
Citrullus
Cucurbita
Cannabis
Vaccinium
Silene
Beta
Carica
Gossypium
Eruca
Arabidopsis
Malus
Hevea
Ricinus
Castanospermum
Millettia
Glycine
Phaseolus
Vigna
Sophora
Ammopiptanthus
Lotus
Trifolium
Glycyrrhiza
Vicia
Cercis
Styphnolobium
Tamarindus
Haematoxylum
Libidibia
Senna
Leucaena
Anadenanthera
Lophophytum mirabile KU992323
Acacia
Inga-oerstediana
Mimosa
Vachellia
Neltuma
Prosopis
0.02
B cox3-sdh4 middle foreign in L. mirabile
Liriodendron
Phoenix
Bambusa
Oryza
Sorghum
Ombrophytum MT076307
Malania
Santalum
Capsicum
Nicotiana
Hyoscyamus
Chrysanthemum
Helianthus
Lactuca
Diplostephium
Asclepias
Rhazya
Boea
Salvia
Mimulus
Daucus
Tamarindus
Hevea
Ricinus
Gossypium
Carica
Eruca
Arabidopsis
Cercis
Senna
Haematoxylum
Libidibia
Mimosa
Anadenanthera
Lophophytum mirabile KU992323
Vachellia
Lophophytum mirabile KU992331
Neltuma
OY723412_Inga-oerstediana
Acacia
Leucaena
Citrullus
Cucurbita
Lotus
Glycyrrhiza
Vicia
Trifolium
Malus
Castanospermum
Styphnolobium
Ammopiptanthus
Sophora
Lophophytum pyramidale
Glycine
Vigna
Phaseolus
Millettia
Cannabis
Vaccinium
Beta_sdh4pseudo
Silene
0.008
C sdh4 foreign in Lophophytum spp.
Liriodendron
Phoenix
Leucaena
Mimosa
Haematoxylum
Libidibia
Lophophytum mirabile KU992323
Anadenanthera
Lophophytum mirabile KU992331
Lophophytum pyramidale
Acacia
OY723412_Inga-oerstediana
Neltuma
Senna
Vachellia
Vitis
Ombrophytum MT076307
Santalum
Cercis
Citrullus
Cucumis
Cucurbita
Chrysanthemum
Helianthus
Diplostephium
Lactuca
Carica
Eruca
Arabidopsis
Gossypium
Malus
Vaccinium
Rhazya
Asclepias
Boea
Salvia
Mimulus
Beta
Silene
Nicotiana
Physochlaina
Hyoscyamus
Capsicum
Tamarindus
Cannabis
Hevea
Ricinus
Sophora
Ammopiptanthus
Castanospermum
Styphnolobium
Glycyrrhiza
Phaseolus
Vigna
Lotus
Millettia
0.02

### *ccmC*

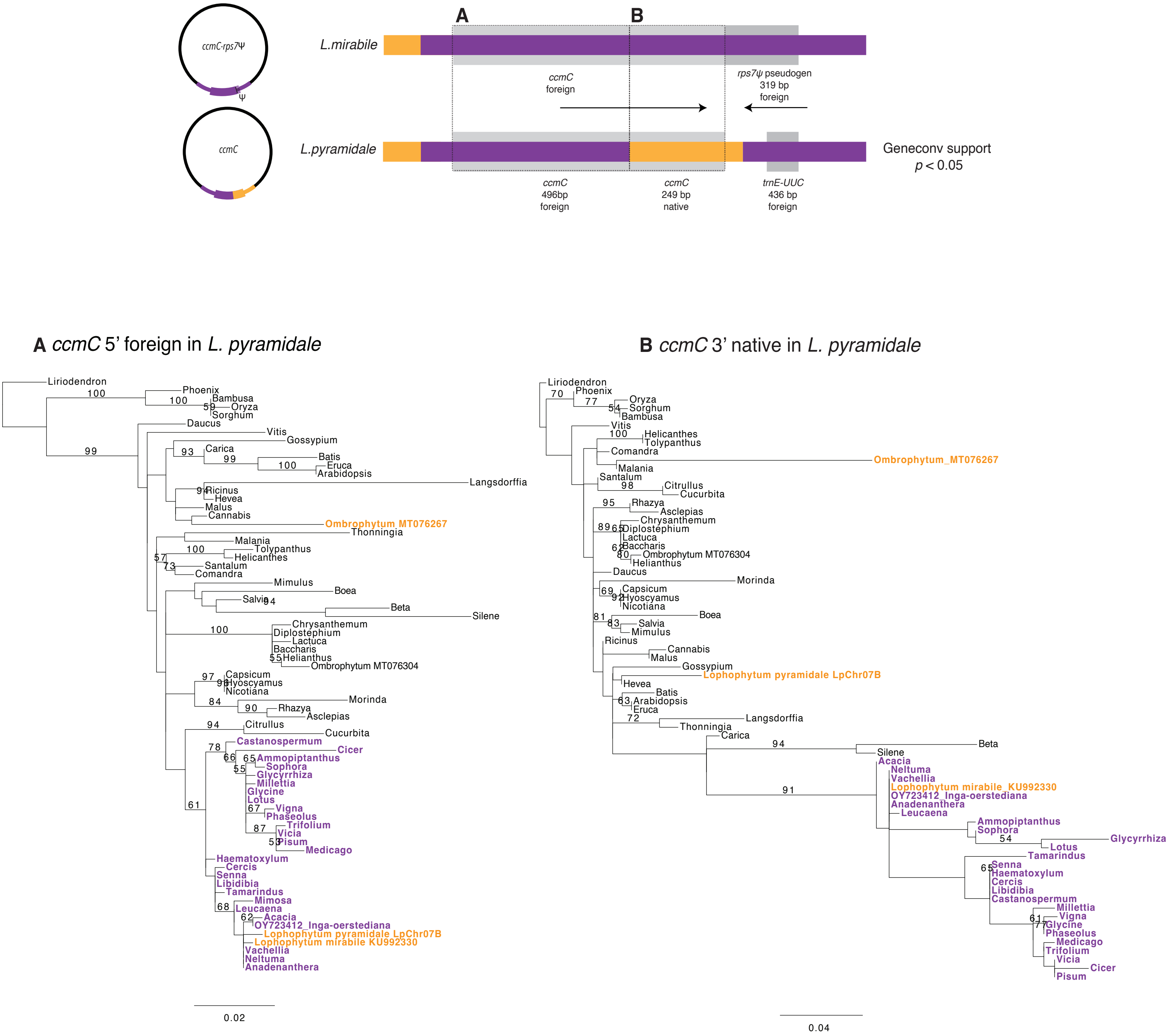

A
B
ccmC-rps7Ψ
Ψ
ccmC
L.mirabile
L.pyramidale
ccmC
foreign
rps7ψ pseudogen
319 bp
foreign
Geneconv support
p < 0.05
ccmC
496bp
foreign
ccmC
249 bp
native
trnE-UUC
436 bp
foreign
A ccmC 5' foreign in L. pyramidale
Liriodendron
Phoenix
Bambusa
Oryza
Sorghum
Daucus
Vitis
Gossypium
Carica
Batis
Eruca
Arabidopsis
Langsdorffia
Ricinus
Hevea
Malus
Cannabis
Ombrophytum MT076267
Thonningia
Malania
Tolypanthus
Helicanthes
Santalum
Comandra
Mimulus
Boea
Salvia
Beta
Silene
Chrysanthemum
Diplostephium
Lactuca
Baccharis
Helianthus
Ombrophytum MT076304
Capsicum
Hyoscyamus
Nicotiana
Morinda
Rhazya
Asclepias
Citrullus
Cucurbita
Castanospermum
Cicer
Ammopiptanthus
Sophora
Glycyrrhiza
Millettia
Glycine
Lotus
Vigna
Phaseolus
Trifolium
Vicia
Pisum
Medicago
Haematoxylum
Cercis
Senna
Libidibia
Tamarindus
Mimosa
Leucaena
Acacia
OY723412_Inga-oerstediana
Lophophytum pyramidale LpChr07B
Lophophytum mirabile KU992330
Vachellia
Neltuma
Anadenanthera
0.02
B ccmC 3' native in L. pyramidale
Liriodendron
Phoenix
Oryza
Sorghum
Bambusa
Vitis
Helicanthes
Tolypanthus
Comandra
Ombrophytum_MT076267
Malania
Santalum
Citrullus
Cucurbita
Rhazya
Asclepias
Chrysanthemum
Diplostephium
Lactuca
Baccharis
Ombrophytum MT076304
Helianthus
Daucus
Morinda
Capsicum
Hyoscyamus
Nicotiana
Boea
Salvia
Mimulus
Ricinus
Cannabis
Malus
Gossypium
Lophophytum pyramidale LpChr07B
Hevea
Batis
Arabidopsis
Eruca
Langsdorffia
Thonningia
Carica
Beta
Silene
Acacia
Neltuma
Vachellia
Lophophytum mirabile_KU992330
OY723412_Inga-oerstediana
Anadenanthera
Leucaena
Ammopiptanthus
Sophora
Glycyrrhiza
Lotus
Tamarindus
Senna
Haematoxylum
Cercis
Libidibia
Castanospermum
Millettia
Vigna
Glycine
Phaseolus
Medicago
Trifolium
Vicia
Cicer
Pisum
0.04

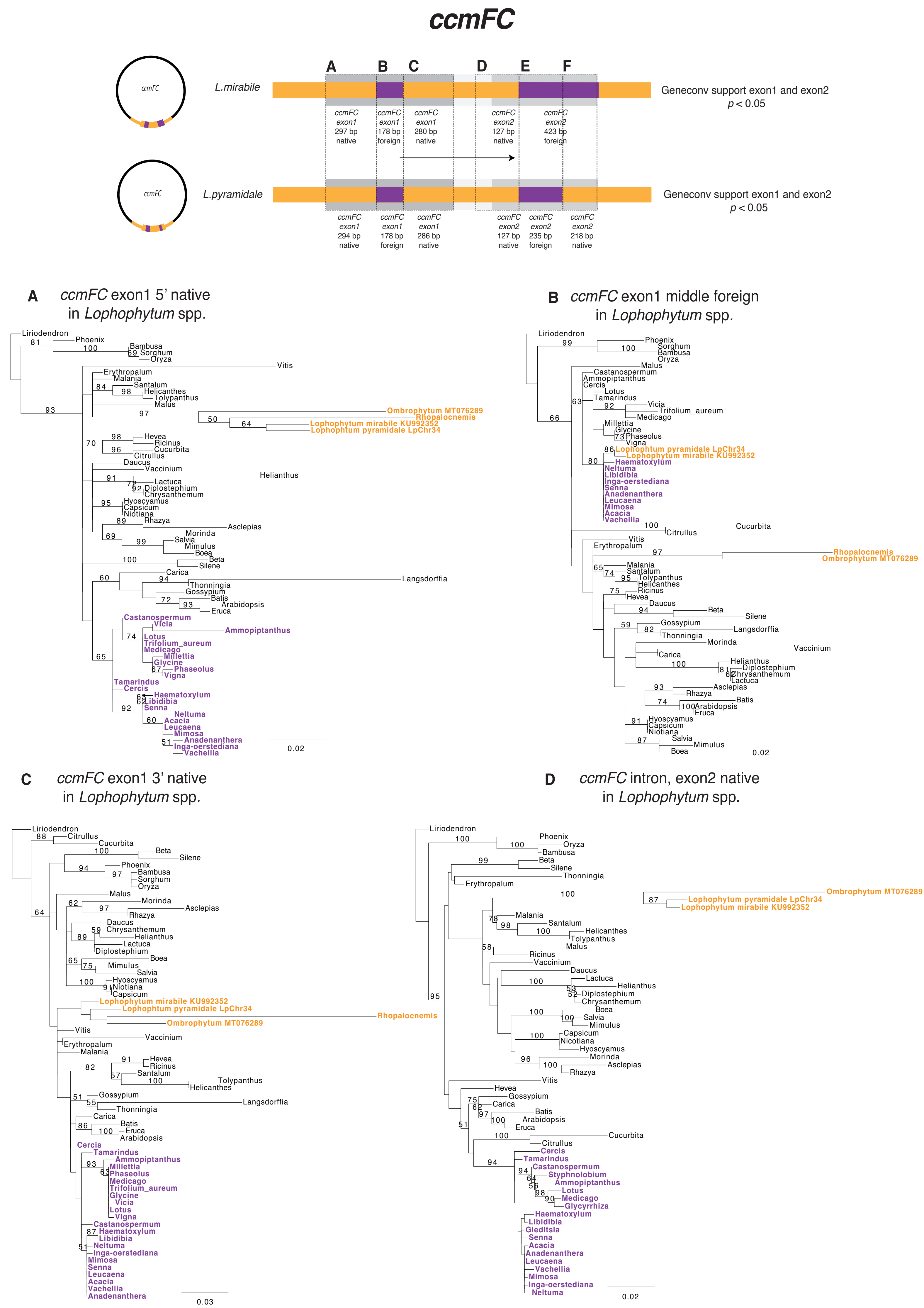
ccmFC
A
B
C
D
E
F
L.mirabile
ccmFC exon1 297 bp native
ccmFC exon1 178 bp foreign
ccmFC exon1 280 bp native
ccmFC exon2 127 bp native
ccmFC exon2 423 bp foreign
Geneconv support exon1 and exon2
p < 0.05
L.pyramidale
ccmFC exon1 294 bp native
ccmFC exon1 178 bp foreign
ccmFC exon1 286 bp native
ccmFC exon2 127 bp native
ccmFC exon2 235 bp foreign
ccmFC exon2 218 bp native
Geneconv support exon1 and exon2
p < 0.05
A ccmFC exon1 5' native in Lophophytum spp.
Liriodendron
Phoenix
Bambusa
Sorghum
Oryza
Vitis
Erythropalum
Malania
Santalum
Helicanthes
Tolypanthus
Malus
Ombrophytum MT076289
Rhopalocnemis
Lophophytum mirabile KU992352
Lophophtum pyramidale LpChr34
Hevea
Ricinus
Cucurbita
Citrullus
Daucus
Vaccinium
Helianthus
Lactuca
Diplostephium
Chrysanthemum
Hyoscyamus
Capsicum
Niotiana
Rhazya
Asclepias
Morinda
Salvia
Mimulus
Boea
Beta
Silene
Carica
Langsdorffia
Thonningia
Gossypium
Batis
Arabidopsis
Eruca
Castanospermum
Vicia
Ammopiptanthus
Lotus
Trifolium_aureum
Medicago
Millettia
Glycine
Phaseolus
Vigna
Tamarindus
Cercis
Haematoxylum
Libidibia
Senna
Neltuma
Acacia
Leucaena
Mimosa
Anadenanthera
Inga-oerstediana
Vachellia
0.02
B ccmFC exon1 middle foreign in Lophophytum spp.
Liriodendron
Phoenix
Sorghum
Bambusa
Oryza
Malus
Castanospermum
Ammopiptanthus
Cercis
Lotus
Tamarindus
Vicia
Trifolium_aureum
Medicago
Millettia
Glycine
Phaseolus
Vigna
Lophophtum pyramidale LpChr34
Lophophytum mirabile KU992352
Haematoxylum
Neltuma
Libidibia
Inga-oerstediana
Senna
Anadenanthera
Leucaena
Mimosa
Acacia
Vachellia
Citrullus
Cucurbita
Vitis
Erythropalum
Rhopalocnemis
Ombrophytum MT076289
Malania
Santalum
Tolypanthus
Helicanthes
Ricinus
Hevea
Daucus
Beta
Silene
Gossypium
Langsdorffia
Thonningia
Morinda
Vaccinium
Carica
Helianthus
Diplostephium
Chrysanthemum
Lactuca
Asclepias
Rhazya
Batis
Arabidopsis
Eruca
Hyoscyamus
Capscicum
Niotiana
Salvia
Mimulus
Boea
0.02
C ccmFC exon1 3' native in Lophophytum spp.
Liriodendron
Citrullus
Cucurbita
Beta
Silene
Phoenix
Bambusa
Sorghum
Oryza
Malus
Morinda
Asclepias
Rhazya
Daucus
Chrysanthemum
Helianthus
Lactuca
Diplostephium
Boea
Mimulus
Salvia
Hyoscyamus
Niotiana
Capsicum
Lophophytum mirabile KU992352
Lophophtum pyramidale LpChr34
Ombrophytum MT076289
Rhopalocnemis
Vitis
Vaccinium
Erythropalum
Malania
Hevea
Ricinus
Santalum
Tolypanthus
Helicanthes
Gossypium
Langsdorffia
Thonningia
Carica
Batis
Eruca
Arabidopsis
Cercis
Tamarindus
Ammopiptanthus
Millettia
Phaseolus
Medicago
Trifolium_aureum
Glycine
Vicia
Lotus
Vigna
Castanospermum
Haematoxylum
Libidibia
Neltuma
Inga-oerstediana
Mimosa
Senna
Leucaena
Acacia
Vachellia
Anadenanthera
0.03
D ccmFC intron, exon2 native in Lophophytum spp.
Liriodendron
Phoenix
Oryza
Bambusa
Beta
Silene
Thonningia
Erythropalum
Ombrophytum MT076289
Lophophytum pyramidale LpChr34
Lophophytum mirabile KU992352
Malania
Santalum
Helicanthes
Tolypanthus
Malus
Ricinus
Vaccinium
Daucus
Lactuca
Helianthus
Diplostephium
Chrysanthemum
Boea
Salvia
Mimulus
Capscicum
Nicotiana
Hyoscyamus
Morinda
Asclepias
Rhazya
Vitis
Hevea
Gossypium
Carica
Batis
Arabidopsis
Eruca
Cucurbita
Citrullus
Cercis
Tamarindus
Castanospermum
Styphnolobium
Ammopiptanthus
Lotus
Medicago
Glycyrrhiza
Haematoxylum
Libidibia
Gleditsia
Senna
Acacia
Anadenanthera
Leucaena
Vachellia
Mimosa
Inga-oerstediana
Neltuma
0.02

**E** *ccmFC* exon2 foreign in *Lophophytum* spp.

**F** *ccmFC* exon2 3' foreign in *L. mirabile*

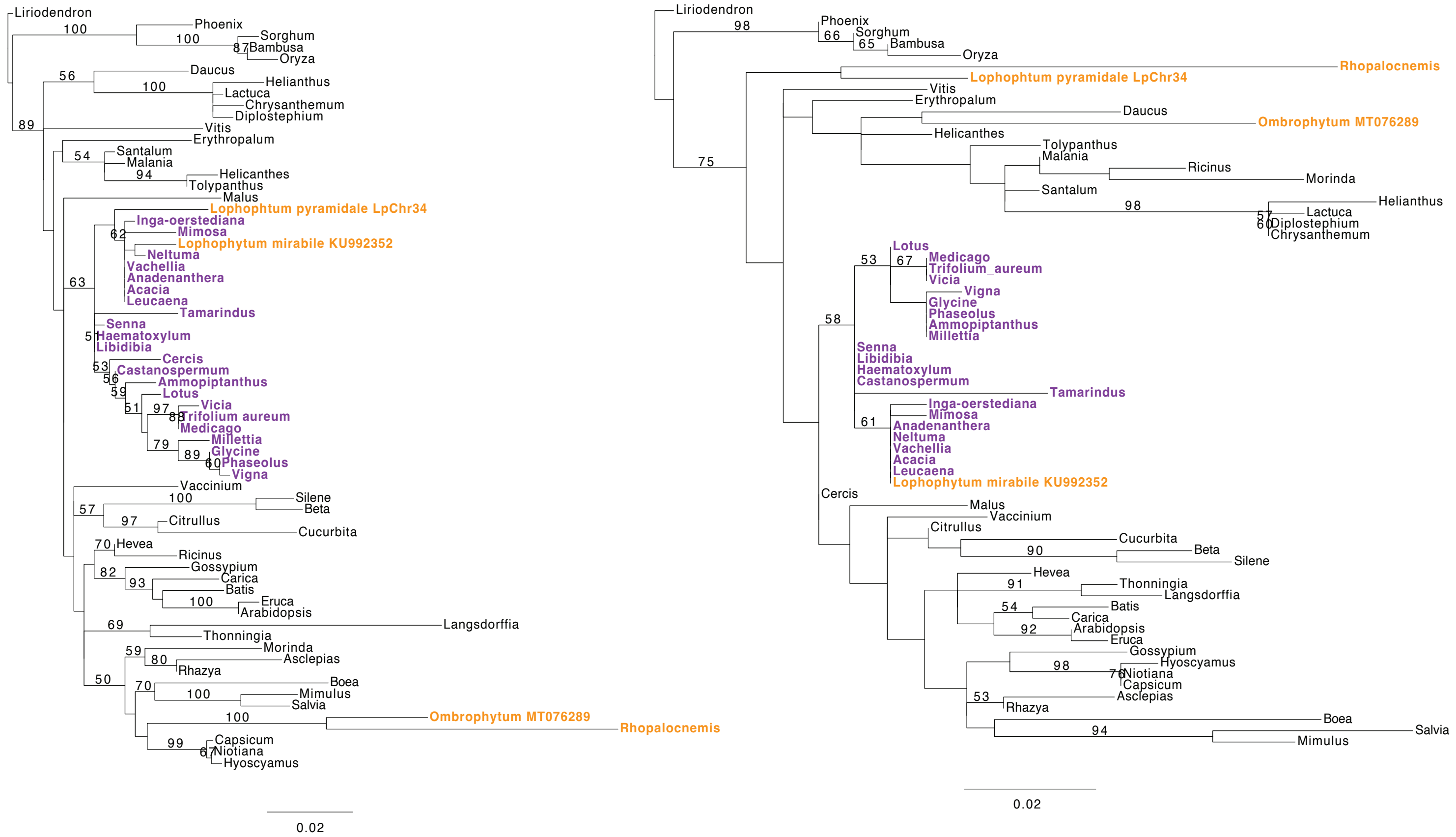

### ccmFN

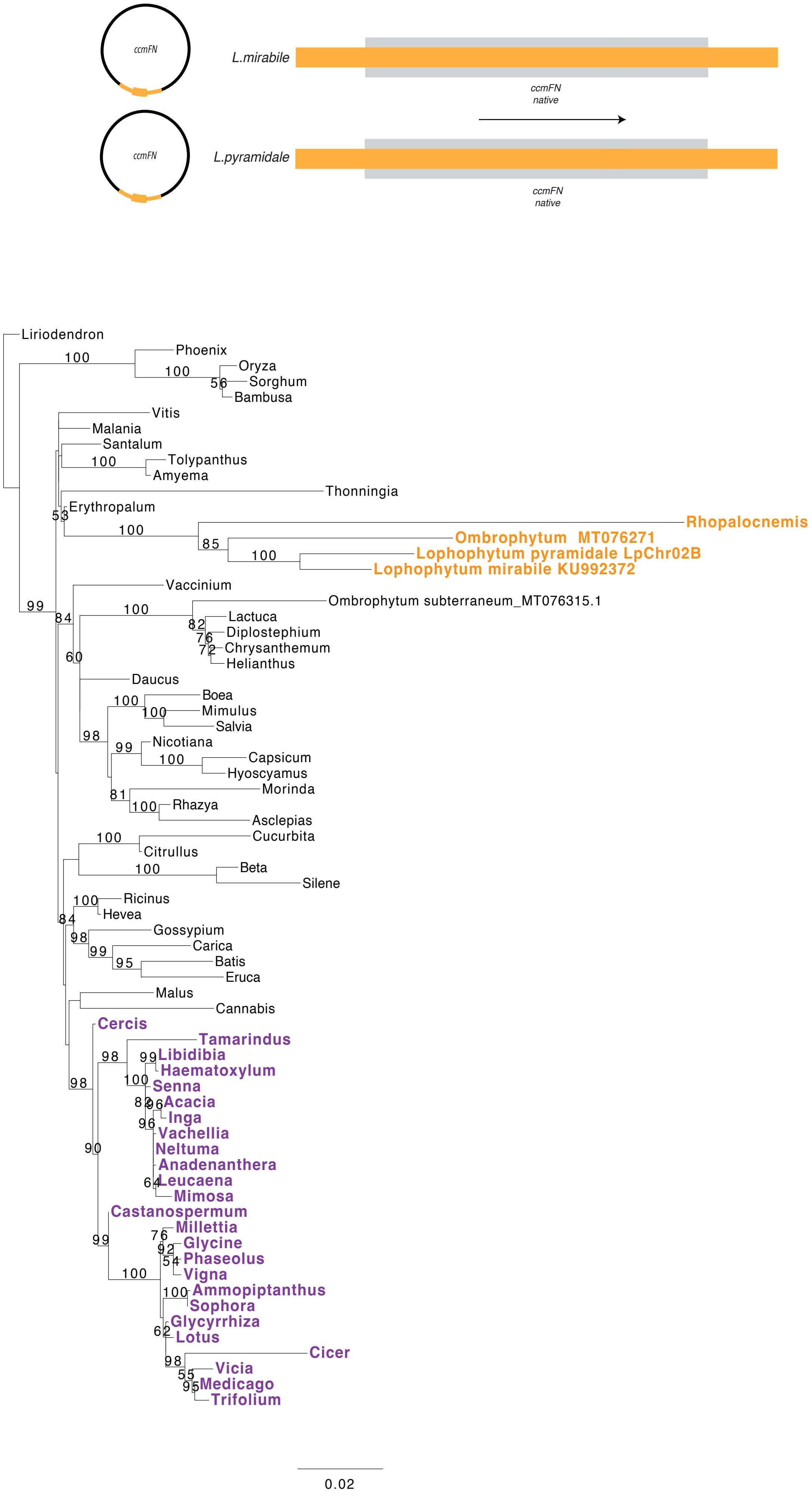

ccmFN
L.mirabile
ccmFN native
ccmFN
L.pyramidale
ccmFN native
Liriodendron
Phoenix
Oryza
Sorghum
Bambusa
Vitis
Malania
Santalum
Tolypanthus
Amyema
Thonningia
Erythropalum
Rhopalocnemis
Ombrophytum MT076271
Lophophytum pyramidale LpChr02B
Lophophytum mirabile KU992372
Vaccinium
Ombrophytum subterraneum_MT076315.1
Lactuca
Diplostephium
Chrysanthemum
Helianthus
Daucus
Boea
Mimulus
Salvia
Nicotiana
Capsicum
Hyoscyamus
Morinda
Rhazya
Asclepias
Cucurbita
Citrullus
Beta
Silene
Ricinus
Hevea
Gossypium
Carica
Batis
Eruca
Malus
Cannabis
Cercis
Tamarindus
Libidibia
Haematoxylum
Senna
Acacia
Inga
Vachellia
Neltuma
Anadenanthera
Leucaena
Mimosa
Castanospermum
Millettia
Glycine
Phaseolus
Vigna
Ammopiptanthus
Sophora
Glycyrrhiza
Lotus
Cicer
Vicia
Medicago
Trifolium
0.02

### matR

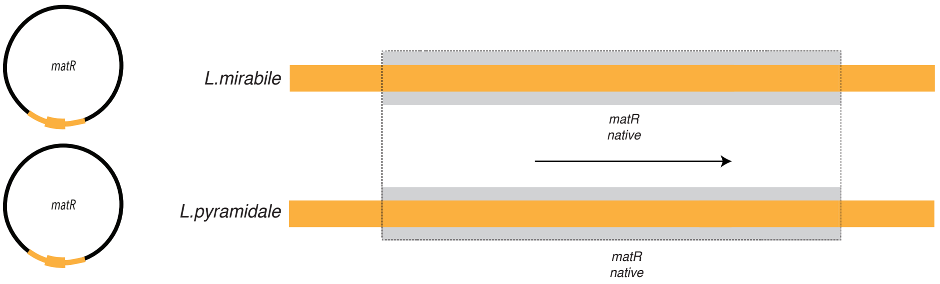
matR
matR
L.mirabile
matR
native
L.pyramidale
matR
native

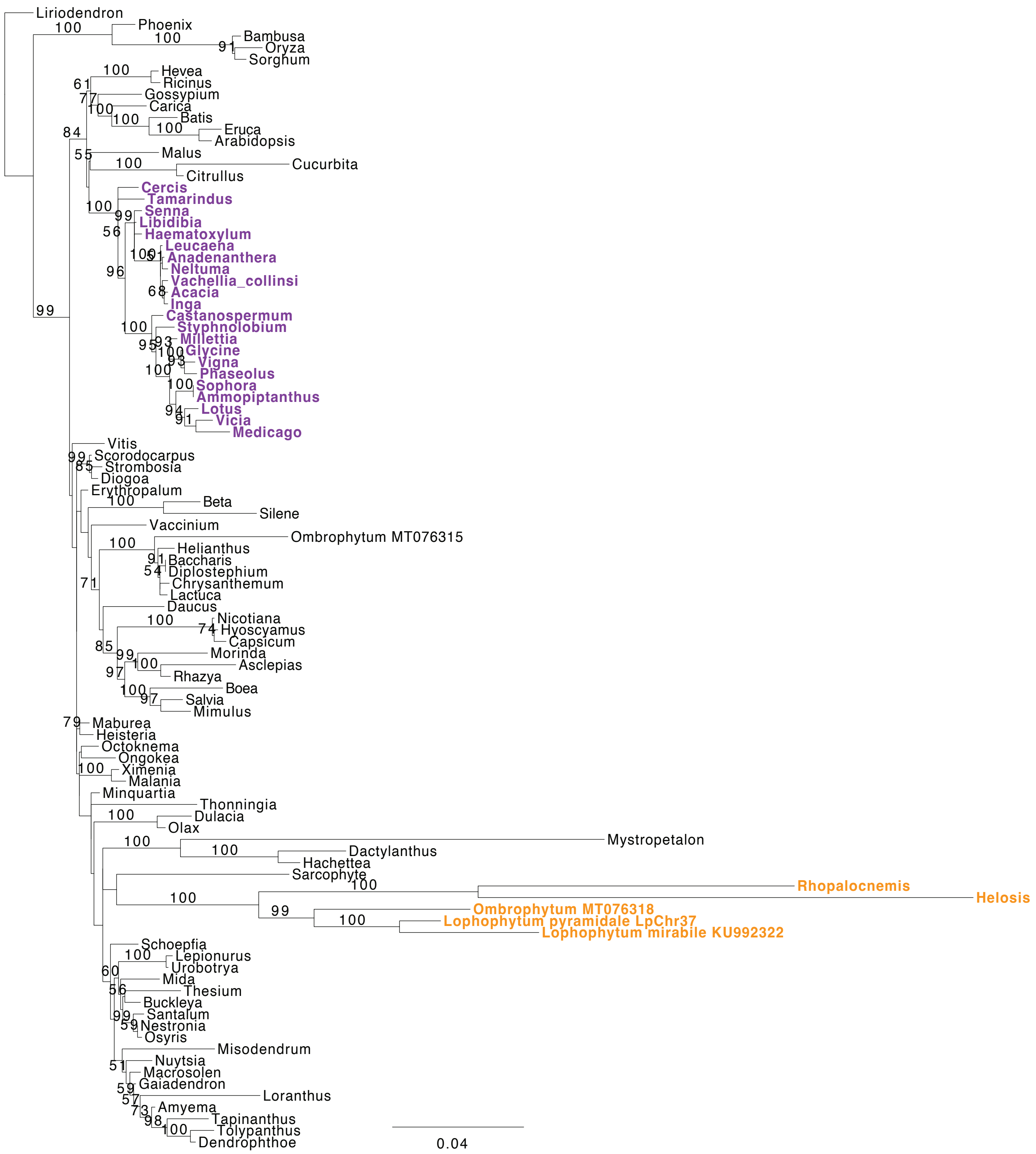
Liriodendron
Phoenix
Bambusa
Oryza
Sorghum
Hevea
Ricinus
Gossypium
Carica
Batis
Eruca
Arabidopsis
Malus
Cucurbita
Citrullus
Cercis
Tamarindus
Senna
Libidibia
Haematoxylum
Leucaena
Anadenanthera
Neltuma
Vachellia_collinsi
Acacia
Inga
Castanospermum
Styphnolobium
Millettia
Glycine
Vigna
Phaseolus
Sophora
Ammopiptanthus
Lotus
Vicia
Medicago
Vitis
Scorodocarpus
Strombosia
Diogoa
Erythropalum
Beta
Silene
Vaccinium
Ombrophytum MT076315
Helianthus
Baccharis
Diplostephium
Chrysanthemum
Lactuca
Daucus
Nicotiana
Hyoscyamus
Capsicum
Morinda
Asclepias
Rhazya
Boea
Salvia
Mimulus
Maburea
Heisteria
Octoknema
Ongokea
Ximenia
Malania
Minquartia
Thonningia
Dulacia
Olax
Mystropetalon
Dactylanthus
Hachettea
Sarcophyte
Rhopalocnemis
Helosis
Ombrophytum MT076318
Lophophytum pyramidale LpChr37
Lophophytum mirabile KU992322
Schoepfia
Lepionurus
Urobotrya
Mida
Thesium
Buckleya
Santalum
Nestronia
Osyris
Misodendrum
Nuytsia
Macrosolen
Gaiadendron
Loranthus
Amyema
Tapinanthus
Tolypanthus
Dendrophthoe
0.04

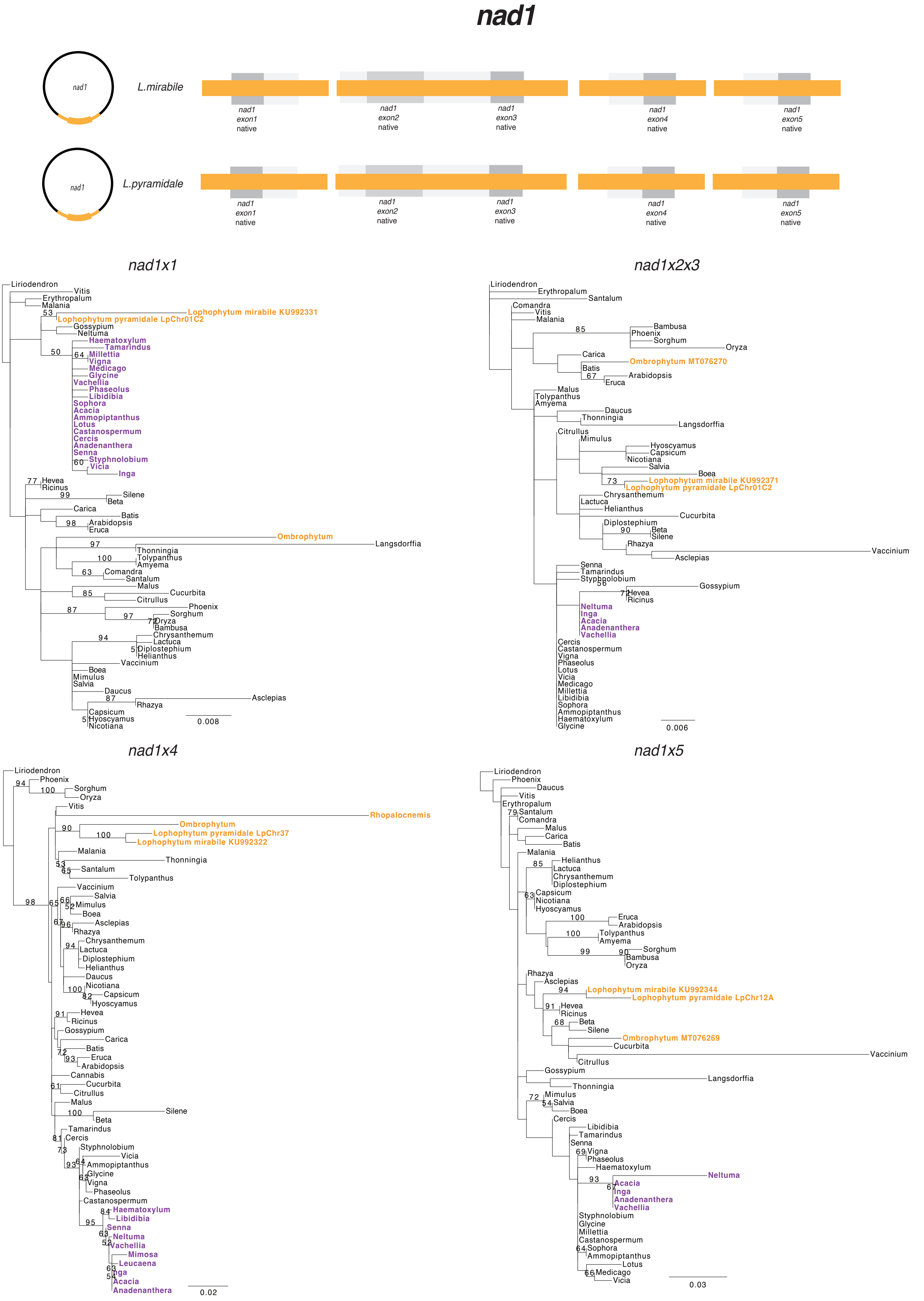
nad1
nad1
L.mirabile
nad1 exon1 native
nad1 exon2 native
nad1 exon3 native
nad1 exon4 native
nad1 exon5 native
nad1
L.pyramidale
nad1 exon1 native
nad1 exon2 native
nad1 exon3 native
nad1 exon4 native
nad1 exon5 native
nad1x1
Liriodendron
Vitis
Erythropalum
Malania
53
Lophophytum mirabile KU992331
Lophophytum pyramidale LpChr01C2
Gossypium
Neltuma
50
Haematoxylum
Tamarindus
64
Milletia
Vigna
Medicago
Glycine
Vachellia
Phaseolus
Libidibia
Sophora
Acacia
Ammopiptanthus
Lotus
Castanospermum
Cercis
Anadenanthera
Senna
Styphnolobium
60
Vicia
Inga
77
Hevea
Ricinus
99
Silene
Beta
Carica
Batis
98
Arabidopsis
Eruca
Ombrophytum
97
Langsdorffia
Thonningia
100
Tolypanthus
Amyema
63
Comandra
Santalum
Malus
85
Cucurbita
Citrullus
87
Phoenix
97
Sorghum
72
Oryza
Bambusa
94
Chrysanthemum
Lactuca
51
Diplostephium
Helianthus
Vaccinium
Boea
Mimulus
Salvia
Daucus
87
Asclepias
Rhazya
Capsicum
51
Hyoscyamus
Nicotiana
0.008
nad1x2x3
Liriodendron
Erythropalum
Santalum
Comandra
Vitis
Malania
85
Bambusa
Phoenix
Sorghum
Oryza
Carica
Ombrophytum MT076270
Batis
67
Arabidopsis
Eruca
Malus
Tolypanthus
Amyema
Daucus
Thonningia
Langsdorffia
Citrullus
Mimulus
Hyoscyamus
Capsicum
Nicotiana
Salvia
Boea
73
Lophophytum mirabile KU992371
Lophophytum pyramidale LpChr01C2
Chrysanthemum
Lactuca
Helianthus
Cucurbita
Diplostephium
90
Beta
Silene
Rhazya
Vaccinium
Asclepias
Senna
Tamarindus
Styphnolobium
56
Gossypium
72
Hevea
Ricinus
Neltuma
Inga
Acacia
Anadenanthera
Vachellia
Cercis
Castanospermum
Vigna
Phaseolus
Lotus
Vicia
Medicago
Millettia
Libidibia
Sophora
Ammopiptanthus
Haematoxylum
Glycine
0.006
nad1x4
Liriodendron
94
Phoenix
100
Sorghum
Oryza
Vitis
Rhopalocnemis
90
Ombrophytum
100
Lophophytum pyramidale LpChr37
Lophophytum mirabile KU992322
Malania
Thonningia
53
65
Santalum
Tolypanthus
Vaccinium
98
65
66
Salvia
52
Mimulus
Boea
67
96
Asclepias
Rhazya
94
Chrysanthemum
Lactuca
Diplostephium
Helianthus
Daucus
100
Nicotiana
82
Capsicum
Hyoscyamus
91
Hevea
Ricinus
Gossypium
Carica
72
Batis
93
Eruca
Arabidopsis
Cannabis
61
Cucurbita
Citrullus
Malus
100
Silene
Beta
Tamarindus
81
Cercis
73
Styphnolobium
Vicia
93
64
Ammopiptanthus
Glycine
63
Vigna
Phaseolus
Castanospermum
84
Haematoxylum
Libidibia
95
Senna
63
Neltuma
52
Vachellia
Mimosa
Leucaena
63
Inga
54
Acacia
Anadenanthera
0.02
nad1x5
Liriodendron
Phoenix
Daucus
Vitis
Erythropalum
79
Santalum
Comandra
Malus
Carica
Batis
Malania
85
Helianthus
Lactuca
Chrysanthemum
Diplostephium
63
Capsicum
Nicotiana
Hyoscyamus
100
Eruca
Arabidopsis
100
Tolypanthus
Amyema
99
90
Sorghum
Bambusa
Oryza
Rhazya
Asclepias
94
Lophophytum mirabile KU992344
Lophophytum pyramidale LpChr12A
91
Hevea
Ricinus
68
Beta
Silene
Ombrophytum MT076269
Cucurbita
Vaccinium
Citrullus
Gossypium
Langsdorffia
Thonningia
72
Mimulus
54
Salvia
Boea
Cercis
Libidibia
Tamarindus
Senna
69
Vigna
Phaseolus
Haematoxylum
Neltuma
93
67
Acacia
Inga
Anadenanthera
Vachellia
Styphnolobium
Glycine
Millettia
Castanospermum
64
Sophora
Ammopiptanthus
Lotus
66
Medicago
Vicia
0.03

### *nad2x1x2*

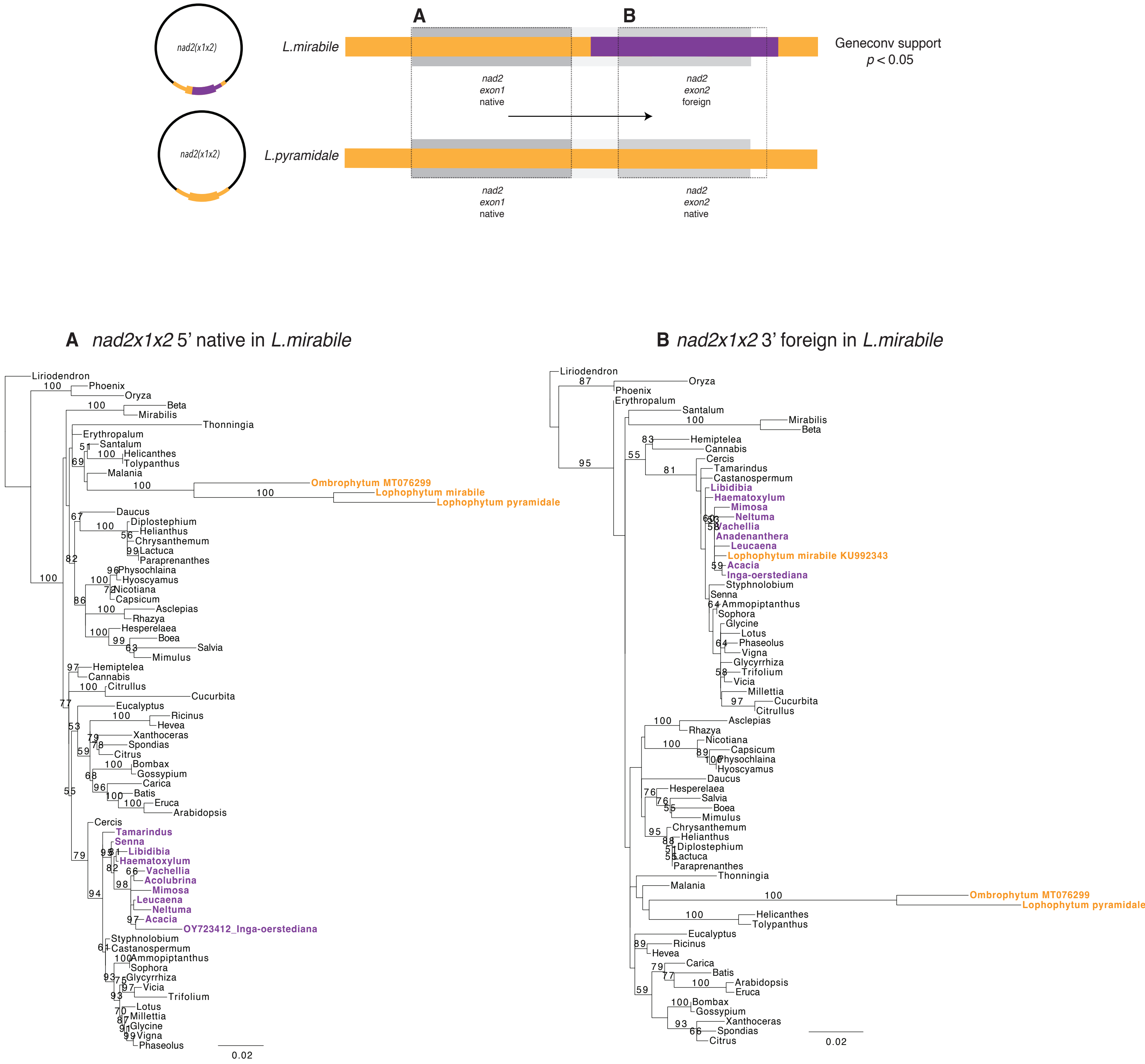
nad2(x1x2)
L.mirabile
A
B
Geneconv support
p < 0.05
nad2
exon1
native
nad2
exon2
foreign
nad2(x1x2)
L.pyramidale
nad2
exon1
native
nad2
exon2
native
A nad2x1x2 5' native in L.mirabile
B nad2x1x2 3' foreign in L.mirabile
Liriodendron
Phoenix
Oryza
Beta
Mirabilis
Thonningia
Erythropalum
Santalum
Helicanthes
Tolypanthus
Malania
Ombrophytum MT076299
Lophophytum mirabile
Lophophytum pyramidale
Daucus
Diplostephium
Helianthus
Chrysanthemum
Lactuca
Paraprenanthes
Physochlaina
Hyoscyamus
Nicotiana
Capsicum
Asclepias
Rhazya
Hesperelaea
Boea
Salvia
Mimulus
Hemiptelea
Cannabis
Citrullus
Cucurbita
Eucalyptus
Ricinus
Hevea
Xanthoceras
Spondias
Citrus
Bombax
Gossypium
Carica
Batis
Eruca
Arabidopsis
Cercis
Tamarindus
Senna
Libidibia
Haematoxylum
Vachellia
Acolubrina
Mimosa
Leucaena
Neltuma
Acacia
OY723412_Inga-oerstediana
Styphnolobium
Castanospermum
Ammopiptanthus
Sophora
Glycyrrhiza
Vicia
Trifolium
Lotus
Millettia
Glycine
Vigna
Phaseolus
0.02
Liriodendron
Oryza
Phoenix
Erythropalum
Santalum
Mirabilis
Beta
Hemiptelea
Cannabis
Cercis
Tamarindus
Castanospermum
Libidibia
Haematoxylum
Mimosa
Neltuma
Vachellia
Anadenanthera
Leucaena
Lophophytum mirabile KU992343
Acacia
Inga-oerstediana
Styphnolobium
Senna
Ammopiptanthus
Sophora
Glycine
Lotus
Phaseolus
Vigna
Glycyrrhiza
Trifolium
Vicia
Millettia
Cucurbita
Citrullus
Asclepias
Rhazya
Nicotiana
Capsicum
Physochlaina
Hyoscyamus
Daucus
Hesperelaea
Salvia
Boea
Mimulus
Chrysanthemum
Helianthus
Diplostephium
Lactuca
Paraprenanthes
Thonningia
Malania
Ombrophytum MT076299
Lophophytum pyramidale
Helicanthes
Tolypanthus
Eucalyptus
Ricinus
Hevea
Carica
Batis
Arabidopsis
Eruca
Bombax
Gossypium
Xanthoceras
Spondias
Citrus
0.02

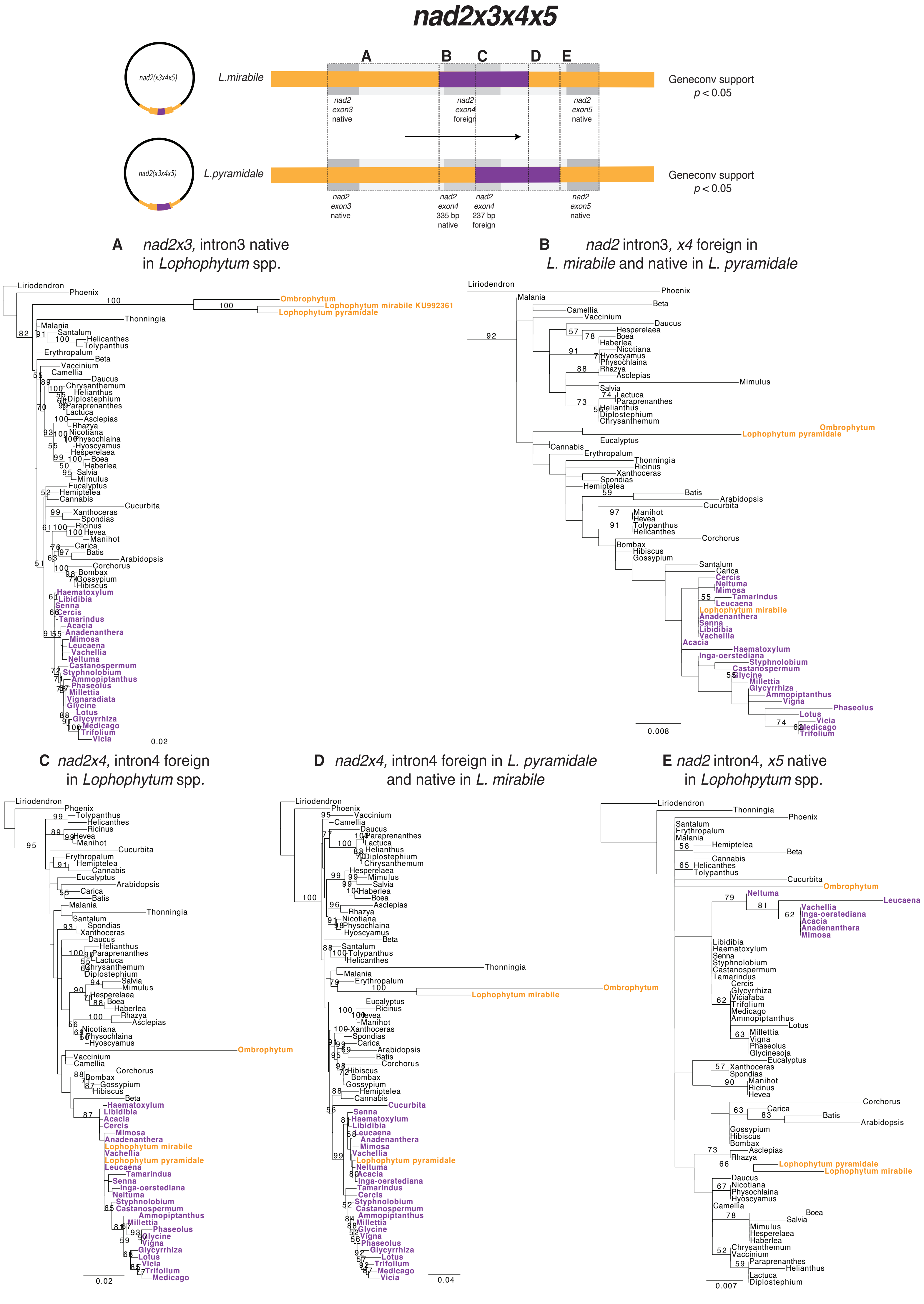
nad2x3x4x5
nad2(x3x4x5)
L.mirabile
A
B
C
D
E
nad2 exon3 native
nad2 exon4 foreign
nad2 exon5 native
Geneconv support
p < 0.05
nad2(x3x4x5)
L.pyramidale
nad2 exon3 native
nad2 exon4 335 bp native
nad2 exon4 237 bp foreign
nad2 exon5 native
Geneconv support
p < 0.05
A nad2x3, intron3 native in Lophophytum spp.
B nad2 intron3, x4 foreign in L. mirabile and native in L. pyramidale
C nad2x4, intron4 foreign in Lophophytum spp.
D nad2x4, intron4 foreign in L. pyramidale and native in L. mirabile
E nad2 intron4, x5 native in Lophohpytum spp.
0.02
0.008
0.02
0.04
0.007

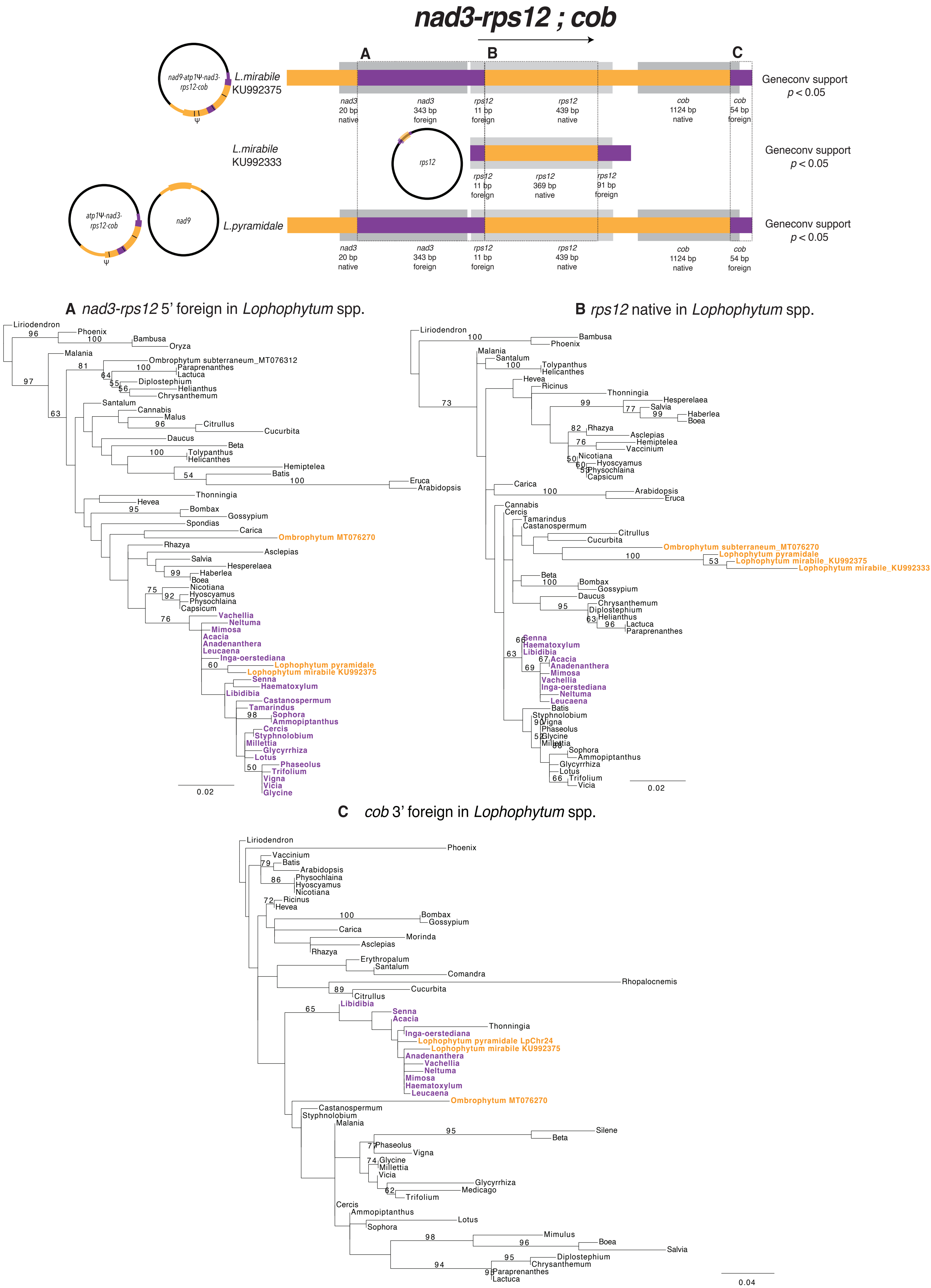
nad3-rps12 ; cob
A
B
C
nad9-atp1Ψ-nad3-rps12-cob
Ψ
L.mirabile KU992375
nad3 20 bp native
nad3 343 bp foreign
rps12 11 bp foreign
rps12 439 bp native
cob 1124 bp native
cob 54 bp foreign
Geneconv support $p < 0.05$
L.mirabile KU992333
rps12
rps12 11 bp foreign
rps12 369 bp native
rps12 91 bp foreign
Geneconv support $p < 0.05$
atp1Ψ-nad3-rps12-cob
Ψ
nad9
L.pyramidale
nad3 20 bp native
nad3 343 bp foreign
rps12 11 bp foreign
rps12 439 bp native
cob 1124 bp native
cob 54 bp foreign
Geneconv support $p < 0.05$
A nad3-rps12 5' foreign in Lophophytum spp.
Liriodendron
96
Phoenix
100
Bambusa
Oryza
Malania
81
Ombrophytum subterraneum_MT076312
100
Paraprenanthes
Lactuca
64
55
Diplostephium
56
Helianthus
Chrysanthemum
97
63
Santalum
Cannabis
Malus
96
Citrullus
Cucurbita
Daucus
Beta
100
Tolypanthus
Helicanthes
Hemiptelea
54
Batis
100
Eruca
Arabidopsis
Thonningia
Hevea
95
Bombax
Gossypium
Spondias
Carica
Ombrophytum MT076270
Rhazya
Asclepias
Salvia
Hesperelaea
99
Haberlea
Boea
75
Nicotiana
92
Hyoscyamus
Physochlaina
Capsicum
76
Vachellia
Neltuma
Mimosa
Acacia
Anadenanthera
Leucaena
Inga-oerstediana
60
Lophophytum pyramidale
Lophophytum mirabile KU992375
Senna
Haematoxylum
Libidibia
Castanospermum
Tamarindus
98
Sophora
Ammopiptanthus
Cercis
Styphnolobium
Millettia
Glycyrrhiza
Lotus
50
Phaseolus
Trifolium
Vigna
Vicia
Glycine
0.02
B rps12 native in Lophophytum spp.
Liriodendron
100
Bambusa
Phoenix
Malania
Santalum
100
Tolypanthus
Helicanthes
73
Hevea
Ricinus
Thonningia
99
Hesperelaea
77
Salvia
99
Haberlea
Boea
82
Rhazya
Asclepias
76
Hemiptelea
Vaccinium
50
Nicotiana
60
Hyoscyamus
58
Physochlaina
Capsicum
Carica
100
Arabidopsis
Eruca
Cannabis
Cercis
Tamarindus
Castanospermum
Citrullus
Cucurbita
Ombrophytum subterraneum_MT076270
100
Lophophytum pyramidale
53
Lophophytum mirabile_KU992375
Lophophytum mirabile_KU992333
Beta
100
Bombax
Gossypium
Daucus
95
Chrysanthemum
Diplostephium
63
Helianthus
96
Lactuca
Paraprenanthes
66
Senna
Haematoxylum
63
Libidibia
67
Acacia
69
Anadenanthera
Mimosa
Vachellia
Inga-oerstediana
Neltuma
Leucaena
Batis
Styphnolobium
90
Vigna
Phaseolus
52
Glycine
Millettia
68
Sophora
Ammopiptanthus
Glycyrrhiza
Lotus
66
Trifolium
Vicia
0.02
C cob 3' foreign in Lophophytum spp.
Liriodendron
Phoenix
Vaccinium
79
Batis
Arabidopsis
86
Physochlaina
Hyoscyamus
Nicotiana
72
Ricinus
Hevea
100
Bombax
Gossypium
Carica
Morinda
Asclepias
Rhazya
Erythropalum
Santalum
Comandra
Rhopalocnemis
89
Cucurbita
Citrullus
65
Libidibia
Senna
Acacia
Thonningia
Inga-oerstediana
Lophophytum pyramidale LpChr24
Lophophytum mirabile KU992375
Anadenanthera
Vachellia
Neltuma
Mimosa
Haematoxylum
Leucaena
Ombrophytum MT076270
Castanospermum
Styphnolobium
Malania
95
Silene
Beta
77
Phaseolus
Vigna
74
Glycine
Millettia
Vicia
Glycyrrhiza
62
Medicago
Trifolium
Cercis
Ammopiptanthus
Lotus
Sophora
98
Mimulus
96
Boea
Salvia
95
Diplostephium
94
Chrysanthemum
98
Paraprenanthes
Lactuca
0.04

### nad4

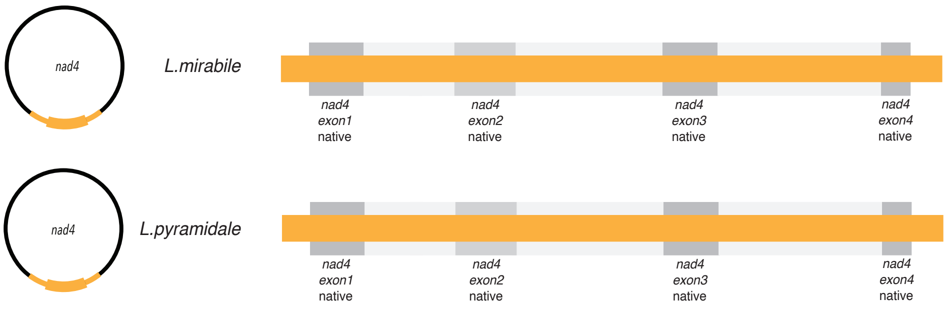
nad4
L.mirabile
nad4 exon1 native
nad4 exon2 native
nad4 exon3 native
nad4 exon4 native
nad4
L.pyramidale
nad4 exon1 native
nad4 exon2 native
nad4 exon3 native
nad4 exon4 native

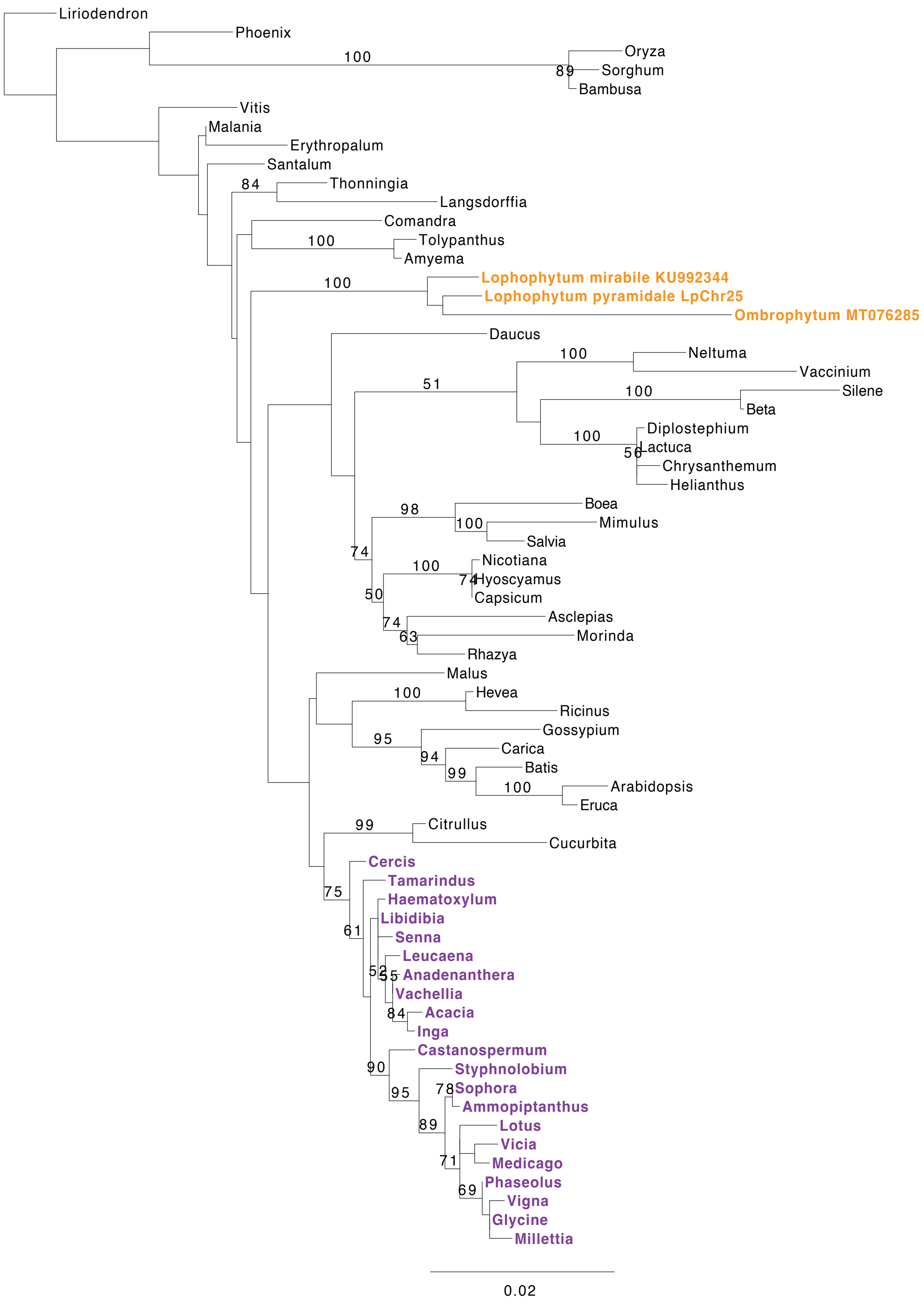
Liriodendron
Phoenix
Oryza
Sorghum
Bambusa
Vitis
Malania
Erythropalum
Santalum
Thonningia
Langsdorffia
Comandra
Tolypanthus
Amyema
Lophophytum mirabile KU992344
Lophophytum pyramidale LpChr25
Ombrophytum MT076285
Daucus
Neltuma
Vaccinium
Silene
Beta
Diplostephium
Lactuca
Chrysanthemum
Helianthus
Boea
Mimulus
Salvia
Nicotiana
Hyoscyamus
Capsicum
Asclepias
Morinda
Rhazya
Malus
Hevea
Ricinus
Gossypium
Carica
Batis
Arabidopsis
Eruca
Citrullus
Cucurbita
Cercis
Tamarindus
Haematoxylum
Libidibia
Senna
Leucaena
Anadenanthera
Vachellia
Acacia
Inga
Castanospermum
Styphnolobium
Sophora
Ammopiptanthus
Lotus
Vicia
Medicago
Phaseolus
Vigna
Glycine
Millettia
0.02

### *nad5x1x2*

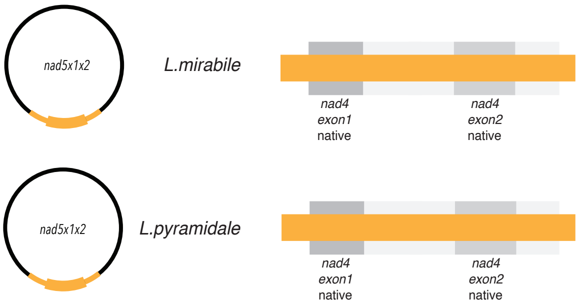
nad5x1x2
L.mirabile
nad4
exon1
native
nad4
exon2
native
nad5x1x2
L.pyramidale
nad4
exon1
native
nad4
exon2
native

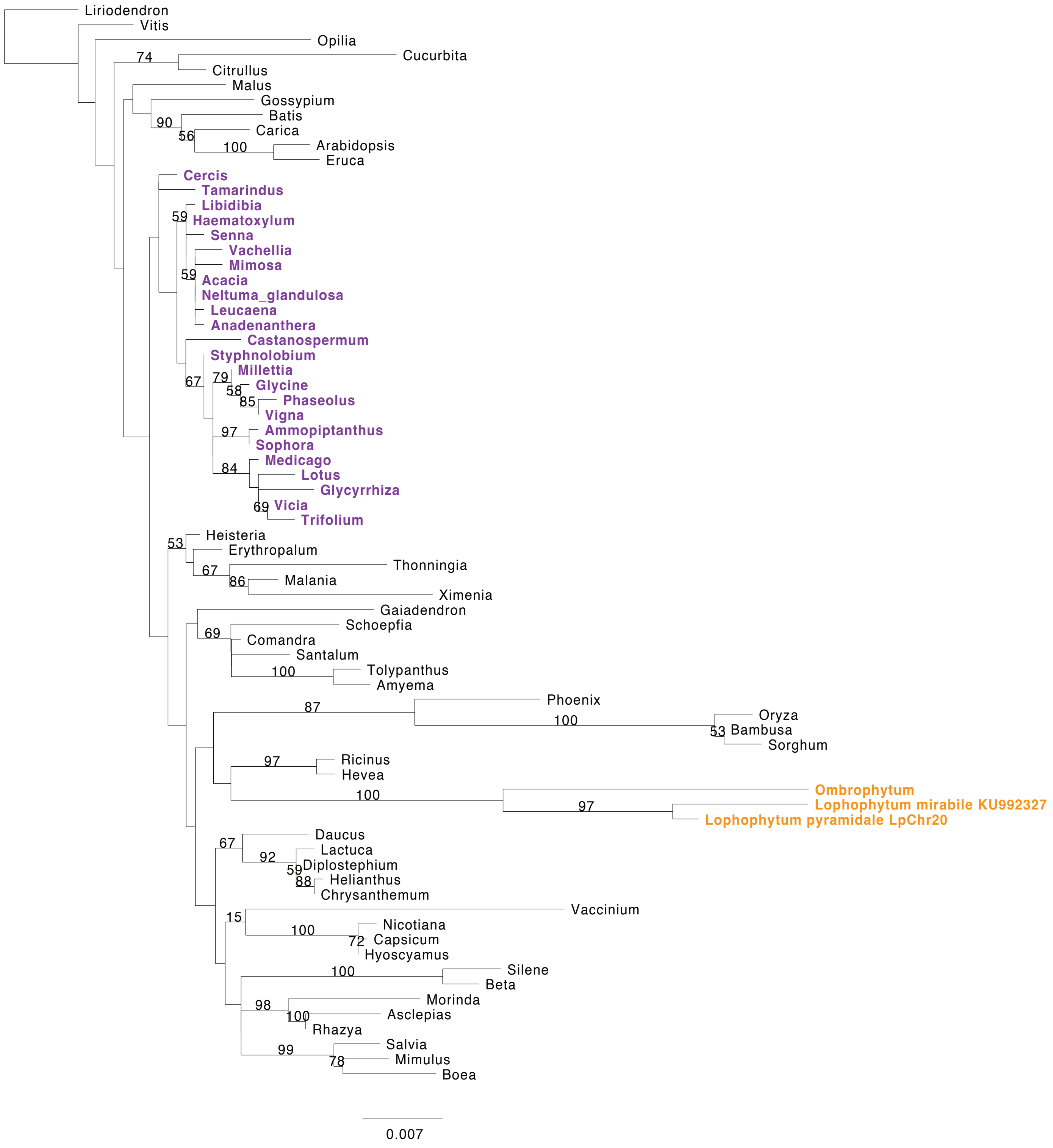
Liriodendron
Vitis
Opilia
Cucurbita
Citrullus
Malus
Gossypium
Batis
Carica
Arabidopsis
Eruca
Cercis
Tamarindus
Libidibia
Haematoxylum
Senna
Vachellia
Mimosa
Acacia
Neltuma_glandulosa
Leucaena
Anadenanthera
Castanospermum
Styphnolobium
Millettia
Glycine
Phaseolus
Vigna
Ammopiptanthus
Sophora
Medicago
Lotus
Glycyrrhiza
Vicia
Trifolium
Heisteria
Erythropalum
Thonningia
Malania
Ximenia
Gaiadendron
Schoepfia
Comandra
Santalum
Tolypanthus
Amyema
Phoenix
Oryza
Bambusa
Sorghum
Ricinus
Hevea
Ombrophytum
Lophophytum mirabile KU992327
Lophophytum pyramidale LpChr20
Daucus
Lactuca
Diplostephium
Helianthus
Chrysanthemum
Vaccinium
Nicotiana
Capsicum
Hyoscyamus
Silene
Beta
Morinda
Asclepias
Rhazya
Salvia
Mimulus
Boea
0.007

### nad5x3

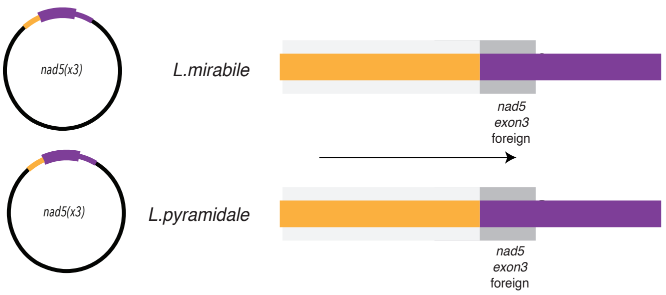
nad5(x3)
L.mirabile
nad5
exon3
foreign
nad5(x3)
L.pyramidale
nad5
exon3
foreign

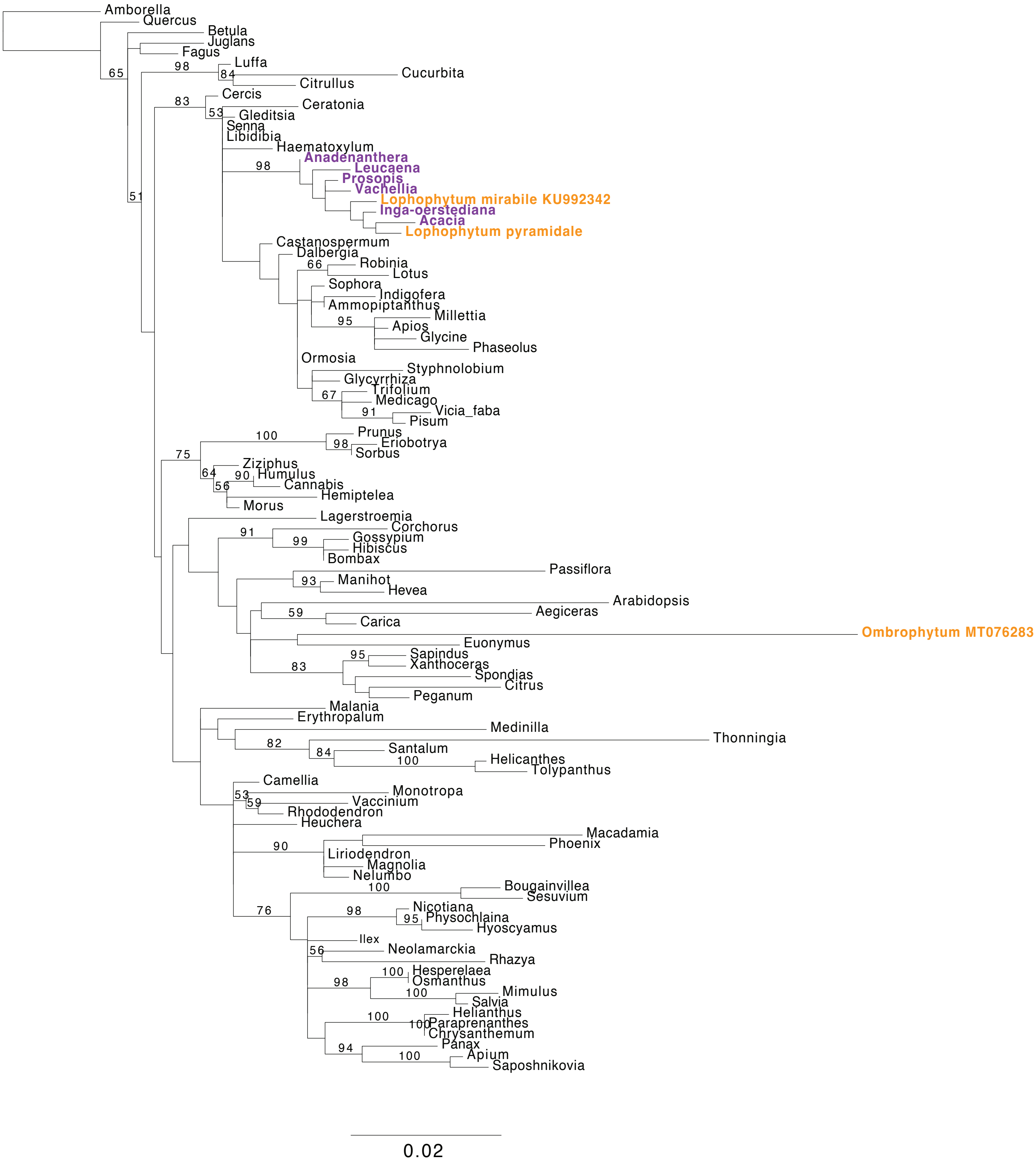
Amborella
Quercus
Betula
Juglans
Fagus
Luffa
Cucurbita
Citrullus
Cercis
Ceratonia
Gleditsia
Senna
Libidibia
Haematoxylum
Anadenanthera
Leucaena
Prosopis
Vachellia
Lophophytum mirabile KU992342
Inga-oerstediana
Acacia
Lophophytum pyramidale
Castanospermum
Dalbergia
Robinia
Lotus
Sophora
Indigofera
Ammopiptanthus
Millettia
Apios
Glycine
Phaseolus
Ormosia
Styphnolobium
Glycyrrhiza
Trifolium
Medicago
Vicia_faba
Pisum
Prunus
Eriobotrya
Sorbus
Ziziphus
Humulus
Cannabis
Hemiptelea
Morus
Lagerstroemia
Corchorus
Gossypium
Hibiscus
Bombax
Passiflora
Manihot
Hevea
Arabidopsis
Aegiceras
Carica
Ombrophytum MT076283
Euonymus
Sapindus
Xanthoceras
Spondias
Citrus
Peganum
Malania
Erythropalum
Medinilla
Thonningia
Santalum
Helicanthes
Tolypanthus
Camellia
Monotropa
Vaccinium
Rhododendron
Heuchera
Macadamia
Phoenix
Liriodendron
Magnolia
Nelumbo
Bougainvillea
Sesuvium
Nicotiana
Physochlaina
Hyoscyamus
Ilex
Neolamarckia
Rhazya
Hesperelaea
Osmanthus
Mimulus
Salvia
Helianthus
Paraprenanthes
Chrysanthemum
Panax
Apium
Saposhnikovia
0.02

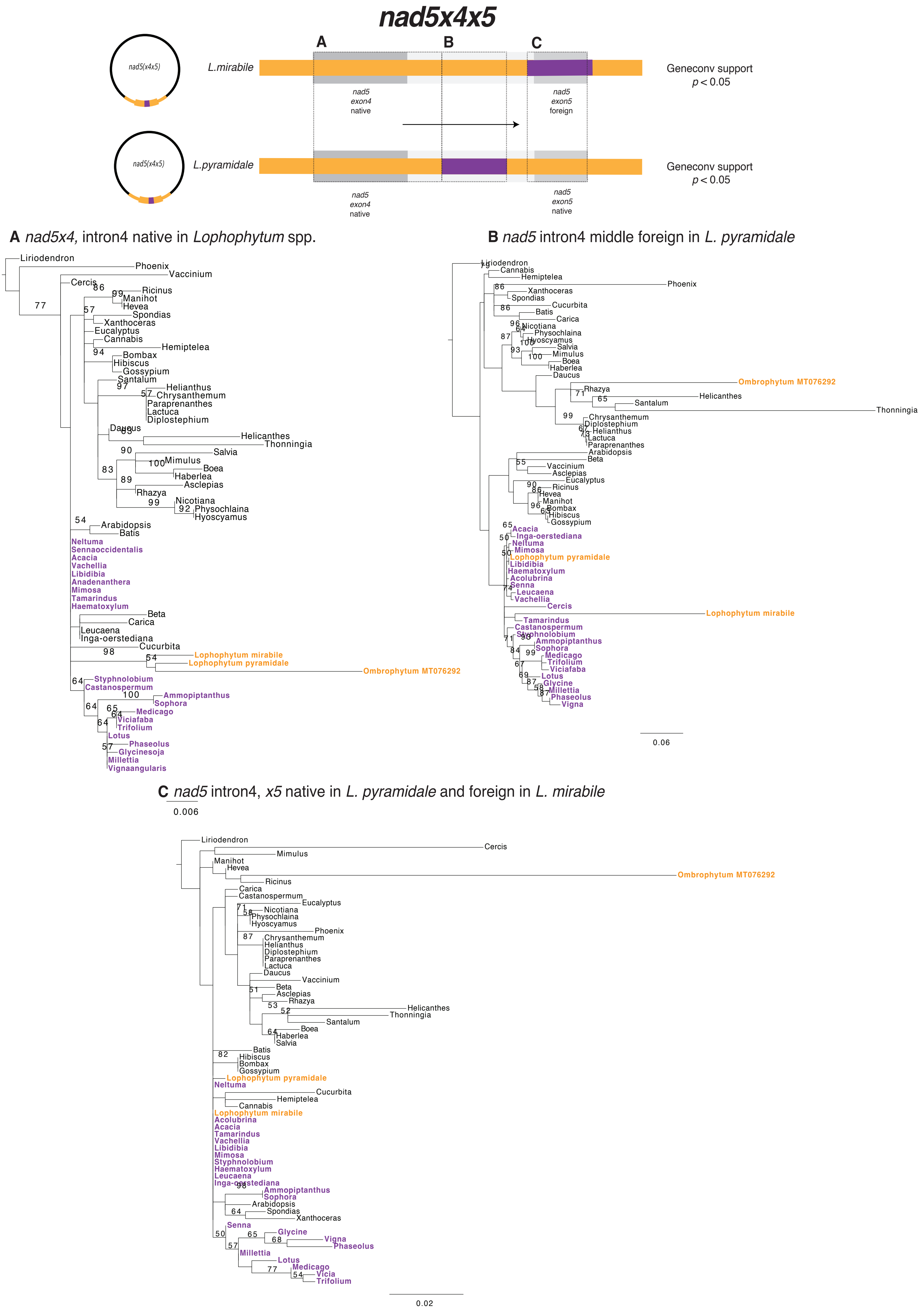
nad5x4x5
nad5(x4x5)
L.mirabile
L.pyramidale
A
B
C
Geneconv support
p < 0.05
nad5
exon4
native
exon5
foreign
A nad5x4, intron4 native in Lophophytum spp.
B nad5 intron4 middle foreign in L. pyramidale
C nad5 intron4, x5 native in L. pyramidale and foreign in L. mirabile
Liriodendron
Phoenix
Vaccinium
Cercis
Ricinus
Manihot
Hevea
Spondias
Xanthoceras
Eucalyptus
Cannabis
Hemiptelea
Bombax
Hibiscus
Gossypium
Santalum
Helianthus
Chrysanthemum
Paraprenanthes
Lactuca
Diplostephium
Daucus
Helicanthes
Thonningia
Salvia
Mimulus
Boea
Haberlea
Asclepias
Rhazya
Nicotiana
Physochlaina
Hyoscyamus
Arabidopsis
Batis
Neltuma
Sennaoccidentalis
Acacia
Vachellia
Libidibia
Anadenanthera
Mimosa
Tamarindus
Haematoxylum
Beta
Carica
Leucaena
Inga-oerstediana
Cucurbita
Lophophytum mirabile
Lophophytum pyramidale
Ombrophytum MT076292
Styphnolobium
Castanospermum
Ammopiptanthus
Sophora
Medicago
Viciafaba
Trifolium
Lotus
Phaseolus
Glycinesoja
Millettia
Vignaangularis
Acolubrina
Senna
Glycine
Vigna
Vicia
0.06
0.006
0.02

### *nad6*

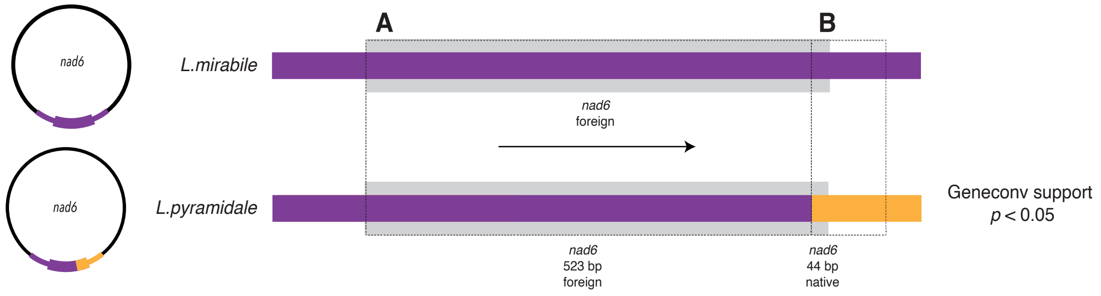


**A** *nad6* foreign in *Lophophytum* spp.

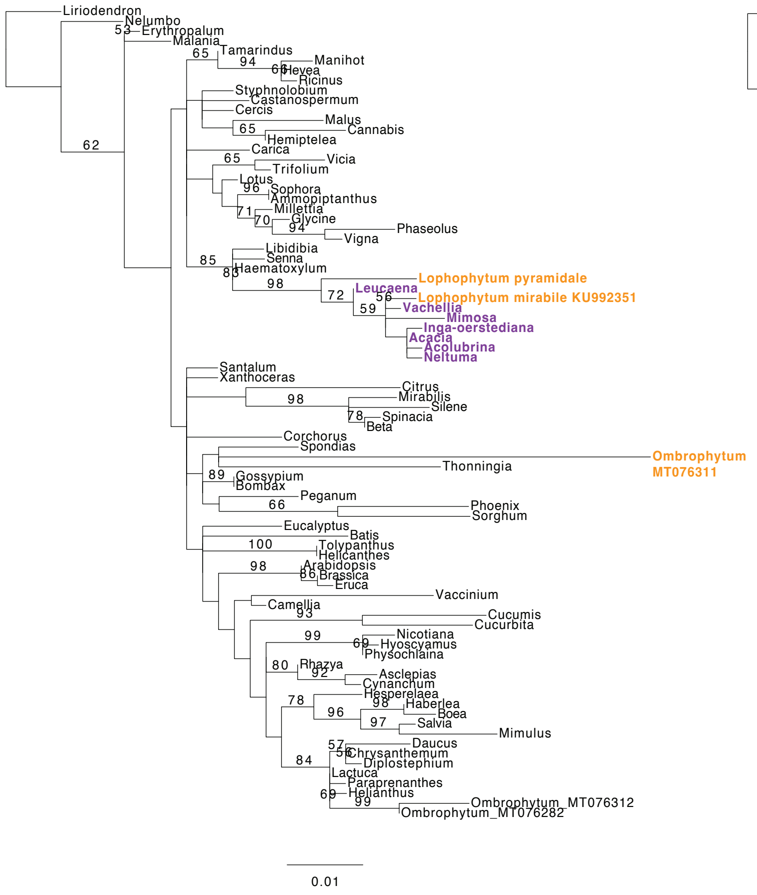


**B** *nad6* 3’ native in *L. pyramidale*

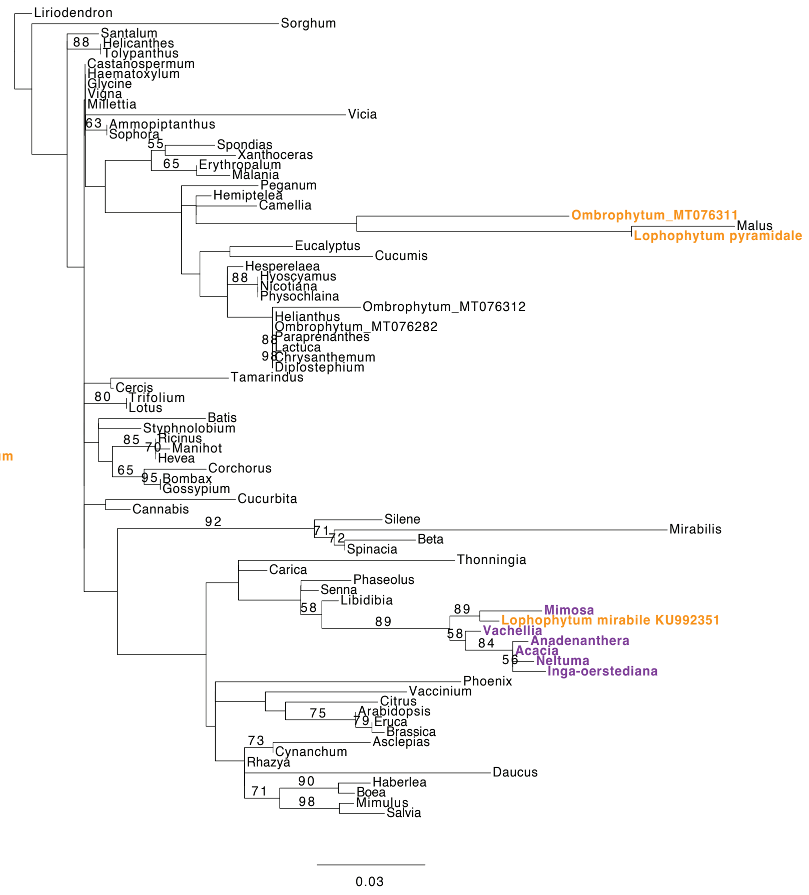

### nad7

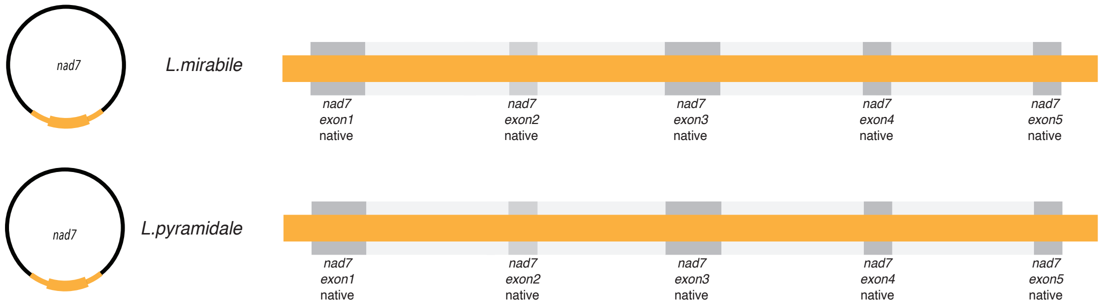
nad7
L.mirabile
nad7 exon1 native
nad7 exon2 native
nad7 exon3 native
nad7 exon4 native
nad7 exon5 native
nad7
L.pyramidale
nad7 exon1 native
nad7 exon2 native
nad7 exon3 native
nad7 exon4 native
nad7 exon5 native

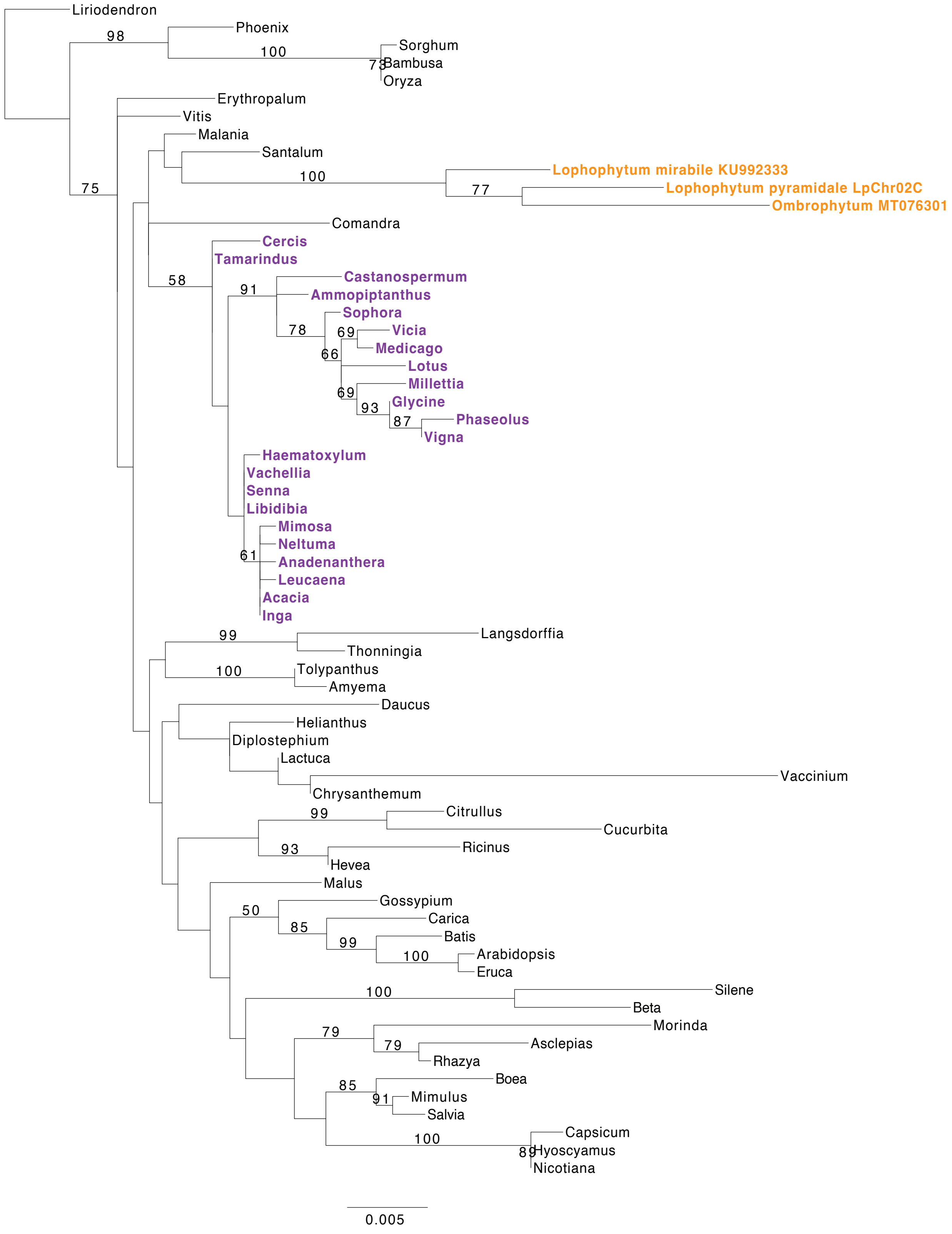
Liriodendron
98
Phoenix
100
Sorghum
73
Bambusa
Oryza
Erythropalum
Vitis
Malania
Santalum
100
Lophophytum mirabile KU992333
77
Lophophytum pyramidale LpChr02C
Ombrophytum MT076301
75
Comandra
Cercis
Tamarindus
58
Castanospermum
91
Ammopiptanthus
Sophora
78
69
Vicia
66
Medicago
Lotus
69
Millettia
93
Glycine
87
Phaseolus
Vigna
Haematoxylum
Vachellia
Senna
Libidibia
Mimosa
Neltuma
61
Anadenanthera
Leucaena
Acacia
Inga
99
Langsdorffia
Thonningia
100
Tolypanthus
Amyema
Daucus
Helianthus
Diplostephium
Lactuca
Vaccinium
Chrysanthemum
99
Citrullus
Cucurbita
93
Ricinus
Hevea
Malus
Gossypium
50
Carica
85
Batis
99
100
Arabidopsis
Eruca
100
Silene
Beta
Morinda
79
79
Asclepias
Rhazya
Boea
85
91
Mimulus
Salvia
Capsicum
100
89
Hyoscyamus
Nicotiana
0.005

### nad9

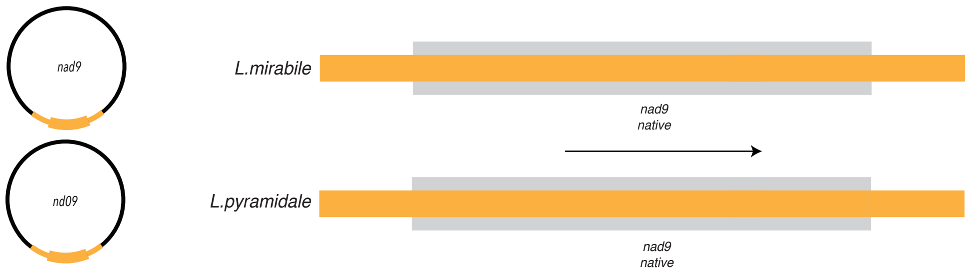

nad9
nd09
L.mirabile
nad9
native
L.pyramidale
nad9
native


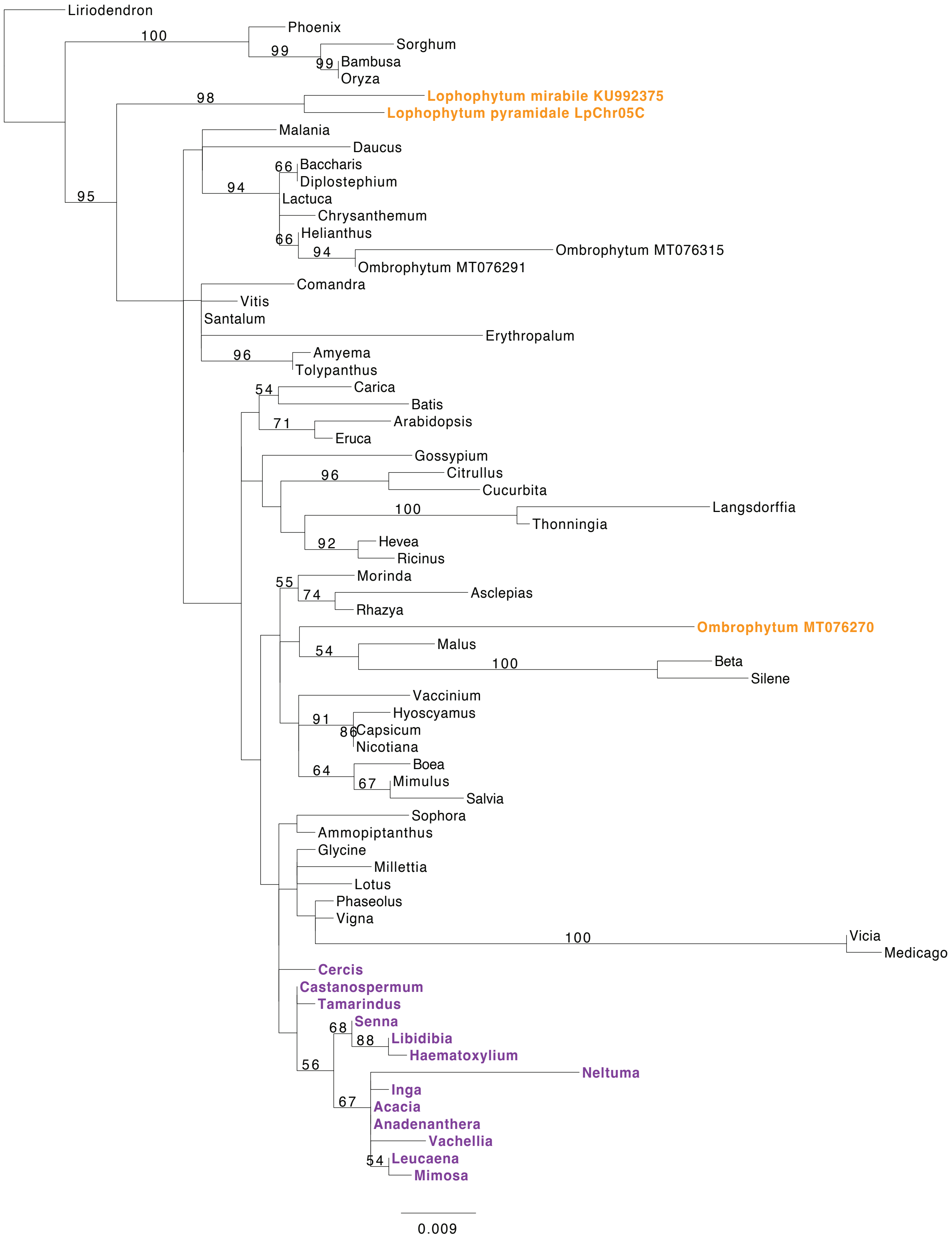

Liriodendron
Phoenix
Sorghum
Bambusa
Oryza
Lophophytum mirabile KU992375
Lophophytum pyramidale LpChr05C
Malania
Daucus
Baccharis
Diplostephium
Lactuca
Chrysanthemum
Helianthus
Ombrophytum MT076315
Ombrophytum MT076291
Comandra
Vitis
Santalum
Erythropalum
Amyema
Tolypanthus
Carica
Batis
Arabidopsis
Eruca
Gossypium
Citrullus
Cucurbita
Langsdorffia
Thonningia
Hevea
Ricinus
Morinda
Asclepias
Rhazya
Ombrophytum MT076270
Malus
Beta
Silene
Vaccinium
Hyoscyamus
Capsicum
Nicotiana
Boea
Mimulus
Salvia
Sophora
Ammopiptanthus
Glycine
Millettia
Lotus
Phaseolus
Vigna
Vicia
Medicago
Cercis
Castanospermum
Tamarindus
Senna
Libidibia
Haematoxylium
Neltuma
Inga
Acacia
Anadenanthera
Vachellia
Leucaena
Mimosa
0.009

### *rpl2-mttB*

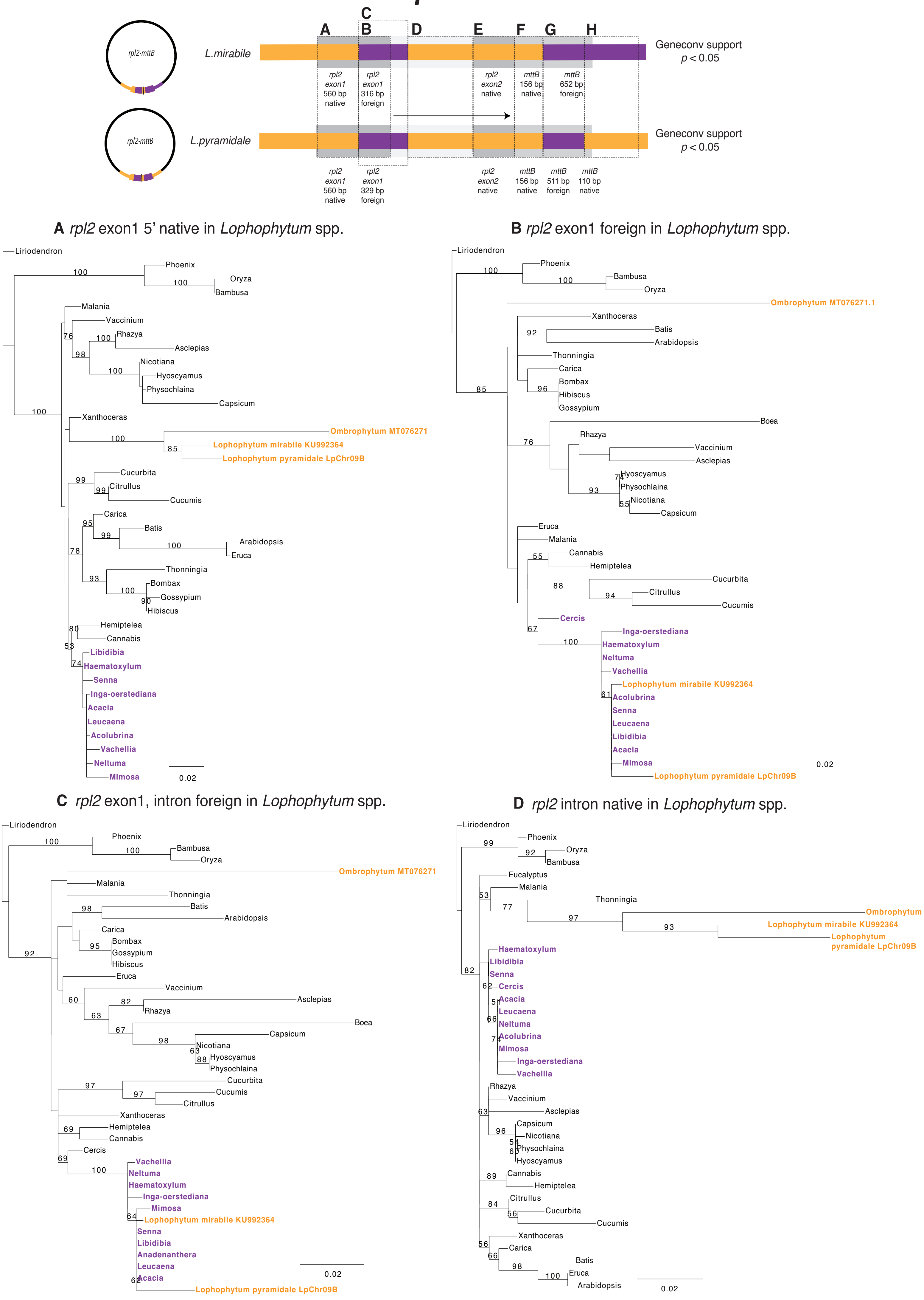

rpl2-mttB
L.mirabile
A
B
C
D
E
F
G
H
Geneconv support
$p < 0.05$
rpl2 exon1 560 bp native
rpl2 exon1 316 bp foreign
rpl2 exon2 native
mttB 156 bp native
mttB 652 bp foreign
rpl2-mttB
L.pyramidale
Geneconv support
$p < 0.05$
rpl2 exon1 560 bp native
rpl2 exon1 329 bp foreign
rpl2 exon2 native
mttB 156 bp native
mttB 511 bp foreign
mttB 110 bp native
A rpl2 exon1 5' native in Lophophytum spp.
B rpl2 exon1 foreign in Lophophytum spp.
C rpl2 exon1, intron foreign in Lophophytum spp.
D rpl2 intron native in Lophophytum spp.
Liriodendron
Phoenix
Oryza
Bambusa
Malania
Vaccinium
Rhazya
Asclepias
Nicotiana
Hyoscyamus
Physochlaina
Capsicum
Xanthoceras
Ombrophytum MT076271
Lophophytum mirabile KU992364
Lophophytum pyramidale LpChr09B
Cucurbita
Citrullus
Cucumis
Carica
Batis
Arabidopsis
Eruca
Thonningia
Bombax
Gossypium
Hibiscus
Hemiptelea
Cannabis
Libidibia
Haematoxylum
Senna
Inga-oerstediana
Acacia
Leucaena
Acolubrina
Vachellia
Neltuma
Mimosa
0.02
Ombrophytum MT076271.1
Boea
Cercis
Anadenanthera
Eucalyptus
Ombrophytum

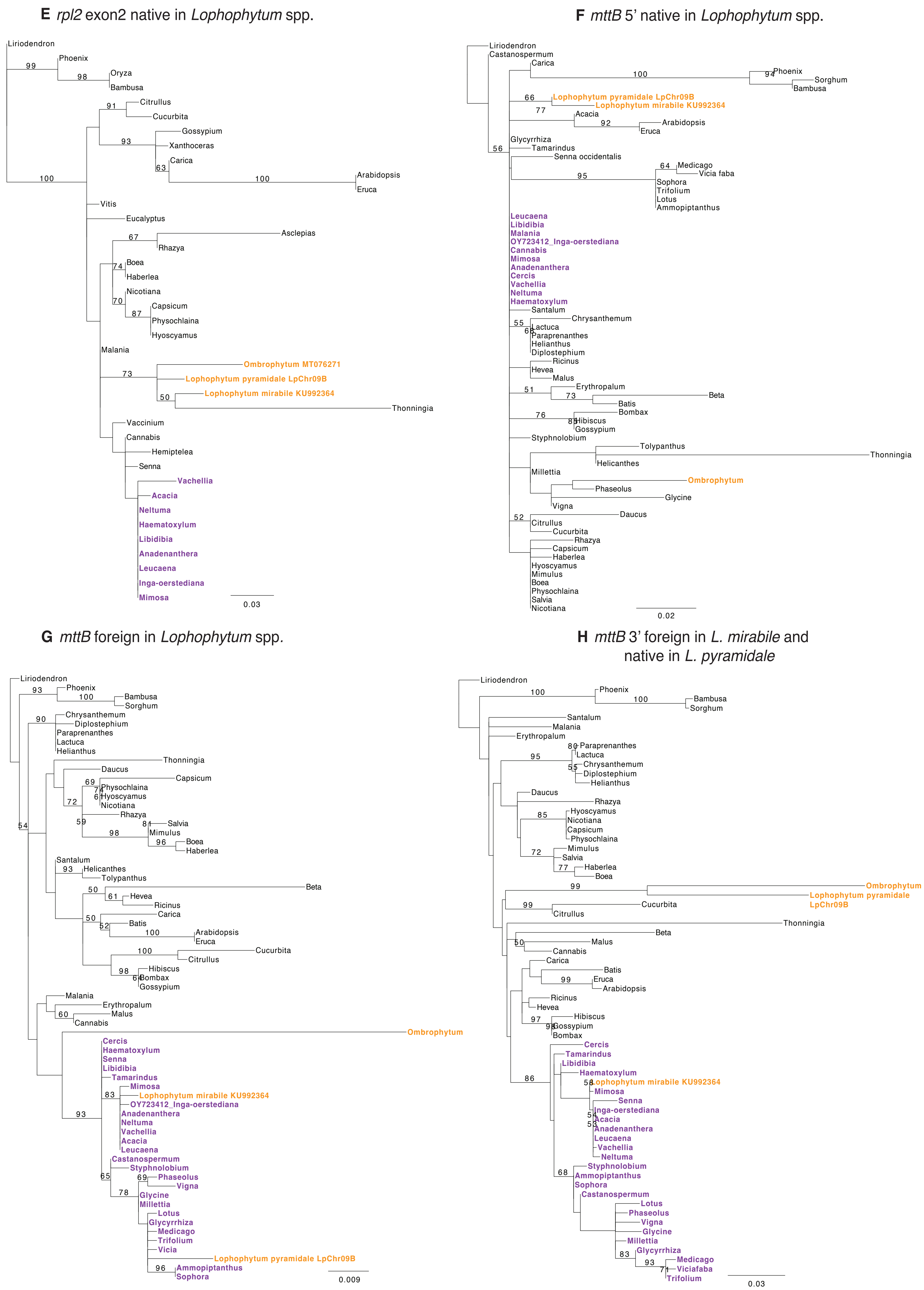

E rpl2 exon2 native in Lophophytum spp.
Liriodendron
Phoenix
Oryza
Bambusa
Citrullus
Cucurbita
Gossypium
Xanthoceras
Carica
Arabidopsis
Eruca
Vitis
Eucalyptus
Asclepias
Rhazya
Boea
Haberlea
Nicotiana
Capsicum
Physochlaina
Hyoscyamus
Malania
Ombrophytum MT076271
Lophophytum pyramidale LpChr09B
Lophophytum mirabile KU992364
Thonningia
Vaccinium
Cannabis
Hemiptelea
Senna
Vachellia
Acacia
Neltuma
Haematoxylum
Libidibia
Anadenanthera
Leucaena
Inga-oerstediana
Mimosa
0.03
F mttB 5' native in Lophophytum spp.
Liriodendron
Castanospermum
Carica
Phoenix
Sorghum
Bambusa
Lophophytum pyramidale LpChr09B
Lophophytum mirabile KU992364
Acacia
Arabidopsis
Eruca
Glycyrrhiza
Tamarindus
Senna occidentalis
Medicago
Vicia faba
Sophora
Trifolium
Lotus
Ammopiptanthus
Leucaena
Libidibia
Malania
OY723412_Inga-oerstediana
Cannabis
Mimosa
Anadenanthera
Cercis
Vachellia
Neltuma
Haematoxylum
Santalum
Chrysanthemum
Lactuca
Paraprenanthes
Helianthus
Diplostephium
Ricinus
Hevea
Malus
Erythropalum
Beta
Batis
Bombax
Hibiscus
Gossypium
Styphnolobium
Tolypanthus
Thonningia
Helicanthes
Millettia
Ombrophytum
Phaseolus
Glycine
Vigna
Daucus
Citrullus
Cucurbita
Rhazya
Capsicum
Haberlea
Hyoscyamus
Mimulus
Boea
Physochlaina
Salvia
Nicotiana
0.02
G mttB foreign in Lophophytum spp.
Liriodendron
Phoenix
Bambusa
Sorghum
Chrysanthemum
Diplostephium
Paraprenanthes
Lactuca
Helianthus
Thonningia
Daucus
Capsicum
Physochlaina
Hyoscyamus
Nicotiana
Rhazya
Salvia
Mimulus
Boea
Haberlea
Santalum
Helicanthes
Tolypanthus
Beta
Hevea
Ricinus
Carica
Batis
Arabidopsis
Eruca
Cucurbita
Citrullus
Hibiscus
Bombax
Gossypium
Malania
Erythropalum
Malus
Cannabis
Ombrophytum
Cercis
Haematoxylum
Senna
Libidibia
Tamarindus
Mimosa
Lophophytum mirabile KU992364
OY723412_Inga-oerstediana
Anadenanthera
Neltuma
Vachellia
Acacia
Leucaena
Castanospermum
Styphnolobium
Phaseolus
Vigna
Glycine
Millettia
Lotus
Glycyrrhiza
Medicago
Trifolium
Vicia
Lophophytum pyramidale LpChr09B
Ammopiptanthus
Sophora
0.009
H mttB 3' foreign in L. mirabile and native in L. pyramidale
Liriodendron
Phoenix
Bambusa
Sorghum
Santalum
Malania
Erythropalum
Paraprenanthes
Lactuca
Chrysanthemum
Diplostephium
Helianthus
Daucus
Rhazya
Hyoscyamus
Nicotiana
Capsicum
Physochlaina
Mimulus
Salvia
Haberlea
Boea
Ombrophytum
Lophophytum pyramidale LpChr09B
Cucurbita
Citrullus
Thonningia
Beta
Malus
Cannabis
Carica
Batis
Eruca
Arabidopsis
Ricinus
Hevea
Hibiscus
Gossypium
Bombax
Cercis
Tamarindus
Libidibia
Haematoxylum
Lophophytum mirabile KU992364
Mimosa
Senna
Inga-oerstediana
Acacia
Anadenanthera
Leucaena
Vachellia
Neltuma
Styphnolobium
Ammopiptanthus
Sophora
Castanospermum
Lotus
Phaseolus
Vigna
Glycine
Millettia
Glycyrrhiza
Medicago
Viciafaba
Trifolium
0.03

### *rpl5-rps14*

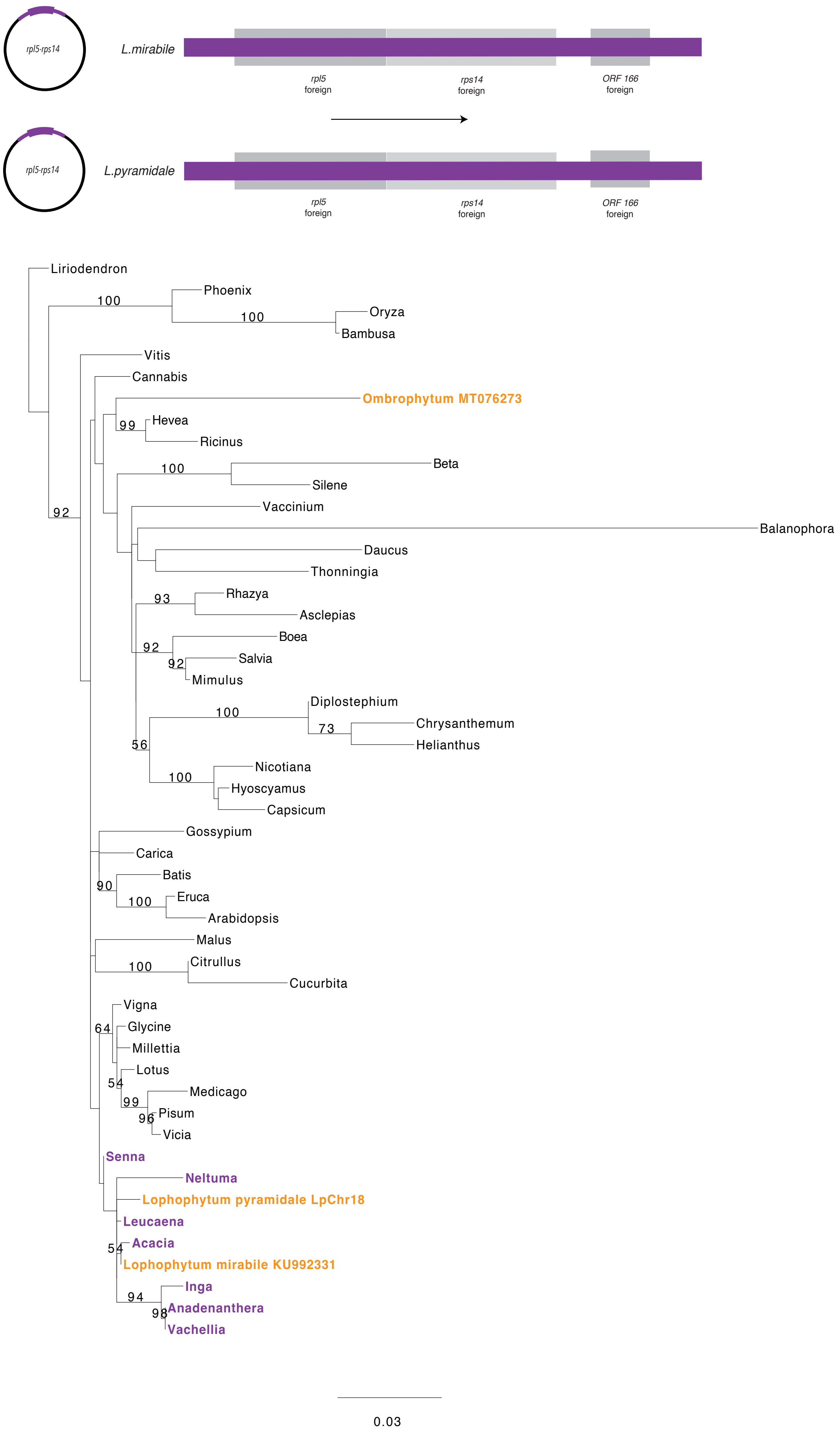
rpl5-rps14
L.mirabile
rpl5
foreign
rps14
foreign
ORF 166
foreign
rpl5-rps14
L.pyramidale
rpl5
foreign
rps14
foreign
ORF 166
foreign
Liriodendron
100
Phoenix
100
Oryza
Bambusa
Vitis
Cannabis
Ombrophytum MT076273
99
Hevea
Ricinus
100
Beta
Silene
92
Vaccinium
Balanophora
Daucus
Thonningia
93
Rhazya
Asclepias
Boea
92
Salvia
92
Mimulus
100
Diplostephium
73
Chrysanthemum
Helianthus
56
100
Nicotiana
Hyoscyamus
Capsicum
Gossypium
Carica
Batis
90
100
Eruca
Arabidopsis
Malus
100
Citrullus
Cucurbita
Vigna
64
Glycine
Millettia
Lotus
54
Medicago
99
96
Pisum
Vicia
Senna
Neltuma
Lophophytum pyramidale LpChr18
Leucaena
Acacia
54
Lophophytum mirabile KU992331
94
Inga
98
Anadenanthera
Vachellia
0.03

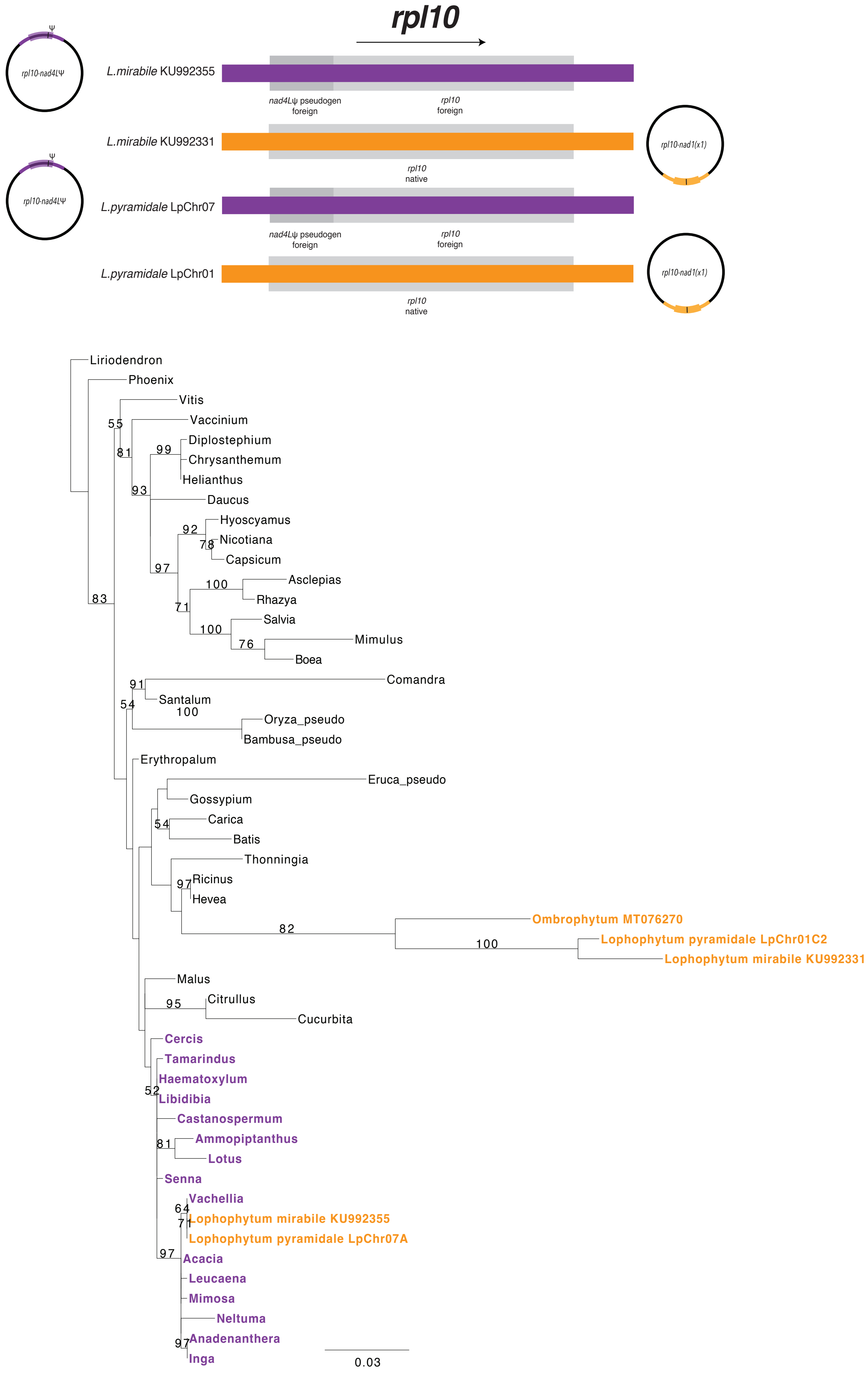

rpl10
rpl10-nad4LΨ
L.mirabile KU992355
nad4Lψ pseudogen
foreign
rpl10
foreign
L.mirabile KU992331
rpl10-nad1(x1)
rpl10
native
rpl10-nad4LΨ
L.pyramidale LpChr07
nad4Lψ pseudogen
foreign
rpl10
foreign
L.pyramidale LpChr01
rpl10-nad1(x1)
rpl10
native
Liriodendron
Phoenix
Vitis
Vaccinium
Diplostephium
Chrysanthemum
Helianthus
Daucus
Hyoscyamus
Nicotiana
Capsicum
Asclepias
Rhazya
Salvia
Mimulus
Boea
Comandra
Santalum
Oryza_pseudo
Bambusa_pseudo
Erythropalum
Eruca_pseudo
Gossypium
Carica
Batis
Thonningia
Ricinus
Hevea
Ombrophytum MT076270
Lophophytum pyramidale LpChr01C2
Lophophytum mirabile KU992331
Malus
Citrullus
Cucurbita
Cercis
Tamarindus
Haematoxylum
Libidibia
Castanospermum
Ammopiptanthus
Lotus
Senna
Vachellia
Lophophytum mirabile KU992355
Lophophytum pyramidale LpChr07A
Acacia
Leucaena
Mimosa
Neltuma
Anadenanthera
Inga
0.03

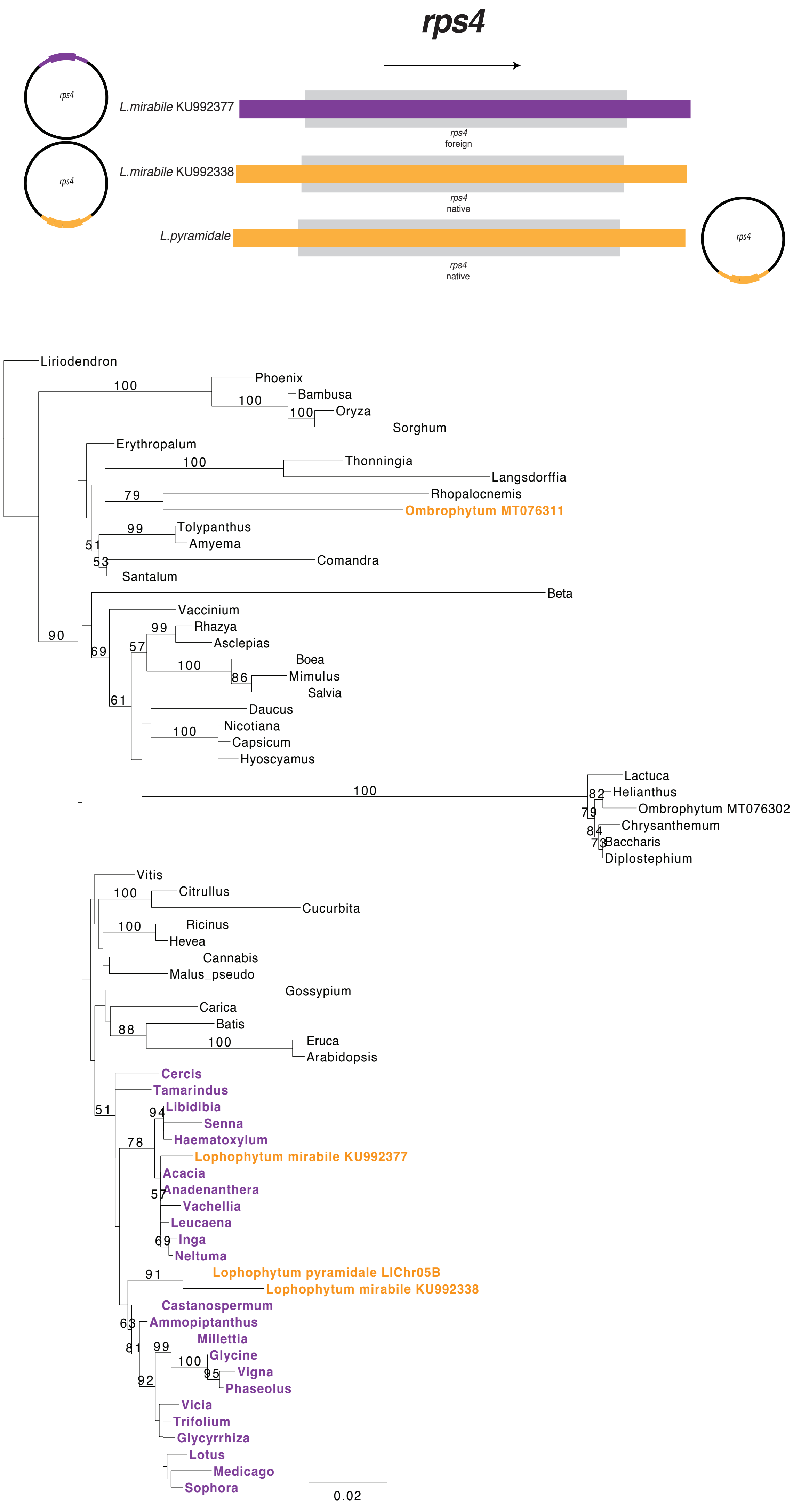
rps4
rps4
L.mirabile KU992377
rps4
foreign
rps4
L.mirabile KU992338
rps4
native
L.pyramidale
rps4
native
rps4
Liriodendron
100
Phoenix
100
Bambusa
100
Oryza
Sorghum
Erythropalum
100
Thonningia
Langsdorffia
79
Rhopalocnemis
Ombrophytum MT076311
99
Tolypanthus
51
Amyema
53
Comandra
Santalum
Beta
Vaccinium
99
Rhazya
90
69
57
Asclepias
100
Boea
86
Mimulus
Salvia
61
Daucus
100
Nicotiana
Capsicum
Hyoscyamus
100
Lactuca
82
Helianthus
79
Ombrophytum MT076302
84
Chrysanthemum
73
Baccharis
Diplostephium
Vitis
100
Citrullus
Cucurbita
100
Ricinus
Hevea
Cannabis
Malus_pseudo
Gossypium
Carica
88
Batis
100
Eruca
Arabidopsis
Cercis
Tamarindus
51
94
Libidibia
Senna
Haematoxylum
78
Lophophytum mirabile KU992377
Acacia
57
Anadenanthera
Vachellia
Leucaena
69
Inga
Neltuma
91
Lophophytum pyramidale LlChr05B
Lophophytum mirabile KU992338
Castanospermum
63
Ammopiptanthus
81
99
Millettia
100
Glycine
92
95
Vigna
Phaseolus
Vicia
Trifolium
Glycyrrhiza
Lotus
Medicago
Sophora
0.02

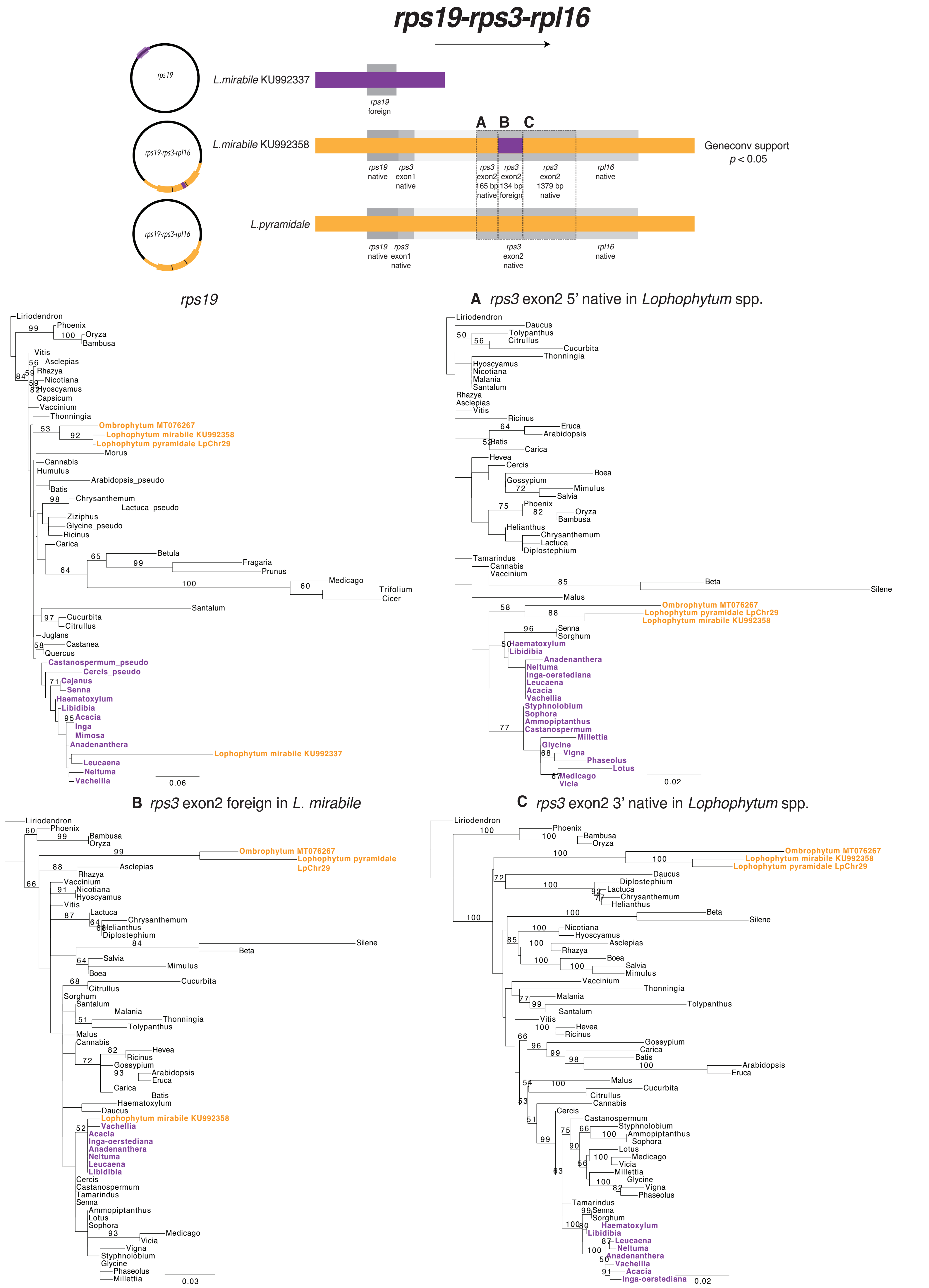
rps19-rps3-rpl16
rps19
L.mirabile KU992337
rps19 foreign
rps19-rps3-rpl16
L.mirabile KU992358
A B C
rps19 native
rps3 exon1 native
rps3 exon2 165 bp native
rps3 exon2 134 bp foreign
rps3 exon2 1379 bp native
rpl16 native
Geneconv support
p < 0.05
rps19-rps3-rpl16
L.pyramidale
rps19 native
rps3 exon1 native
rps3 exon2 native
rpl16 native
rps19
Liriodendron
Phoenix
Oryza
Bambusa
Vitis
Asclepias
Rhazya
Nicotiana
Hyoscyamus
Capsicum
Vaccinium
Thonningia
Ombrophytum MT076267
Lophophytum mirabile KU992358
Lophophytum pyramidale LpChr29
Morus
Cannabis
Humulus
Arabidopsis_pseudo
Batis
Chrysanthemum
Lactuca_pseudo
Ziziphus
Glycine_pseudo
Ricinus
Carica
Betula
Fragaria
Prunus
Medicago
Trifolium
Cicer
Santalum
Cucurbita
Citrullus
Juglans
Castanea
Quercus
Castanospermum_pseudo
Cercis_pseudo
Cajanus
Senna
Haematoxylum
Libidibia
Acacia
Inga
Mimosa
Anadenanthera
Lophophytum mirabile KU992337
Leucaena
Neltuma
Vachellia
0.06
A rps3 exon2 5' native in Lophophytum spp.
Liriodendron
Daucus
Tolypanthus
Citrullus
Cucurbita
Thonningia
Hyoscyamus
Nicotiana
Malania
Santalum
Rhazya
Asclepias
Vitis
Ricinus
Eruca
Arabidopsis
Batis
Carica
Hevea
Cercis
Boea
Gossypium
Mimulus
Salvia
Phoenix
Oryza
Bambusa
Helianthus
Chrysanthemum
Lactuca
Diplostephium
Tamarindus
Cannabis
Vaccinium
Beta
Silene
Malus
Ombrophytum MT076267
Lophophytum pyramidale LpChr29
Lophophytum mirabile KU992358
Senna
Sorghum
Haematoxylum
Libidibia
Anadenanthera
Neltuma
Inga-oerstediana
Leucaena
Acacia
Vachellia
Styphnolobium
Sophora
Ammopiptanthus
Castanospermum
Millettia
Glycine
Vigna
Phaseolus
Lotus
Medicago
Vicia
0.02
B rps3 exon2 foreign in L. mirabile
Liriodendron
Phoenix
Bambusa
Oryza
Ombrophytum MT076267
Lophophytum pyramidale LpChr29
Asclepias
Rhazya
Vaccinium
Nicotiana
Hyoscyamus
Vitis
Lactuca
Chrysanthemum
Helianthus
Diplostephium
Silene
Beta
Salvia
Mimulus
Boea
Cucurbita
Citrullus
Sorghum
Santalum
Malania
Thonningia
Tolypanthus
Malus
Cannabis
Hevea
Ricinus
Gossypium
Arabidopsis
Eruca
Carica
Batis
Haematoxylum
Daucus
Lophophytum mirabile KU992358
Vachellia
Acacia
Inga-oerstediana
Anadenanthera
Neltuma
Leucaena
Libidibia
Cercis
Castanospermum
Tamarindus
Senna
Ammopiptanthus
Lotus
Sophora
Medicago
Vicia
Vigna
Styphnolobium
Glycine
Phaseolus
Millettia
0.03
C rps3 exon2 3' native in Lophophytum spp.
Liriodendron
Phoenix
Bambusa
Oryza
Ombrophytum MT076267
Lophophytum mirabile KU992358
Lophophytum pyramidale LpChr29
Daucus
Diplostephium
Lactuca
Chrysanthemum
Helianthus
Beta
Silene
Nicotiana
Hyoscyamus
Asclepias
Rhazya
Boea
Salvia
Mimulus
Vaccinium
Thonningia
Malania
Tolypanthus
Santalum
Vitis
Hevea
Ricinus
Gossypium
Carica
Batis
Arabidopsis
Eruca
Malus
Cucurbita
Citrullus
Cannabis
Cercis
Castanospermum
Styphnolobium
Ammopiptanthus
Sophora
Lotus
Medicago
Vicia
Millettia
Glycine
Vigna
Phaseolus
Tamarindus
Senna
Sorghum
Haematoxylum
Libidibia
Leucaena
Neltuma
Anadenanthera
Vachellia
Acacia
Inga-oerstediana
0.02

### *rpl16*

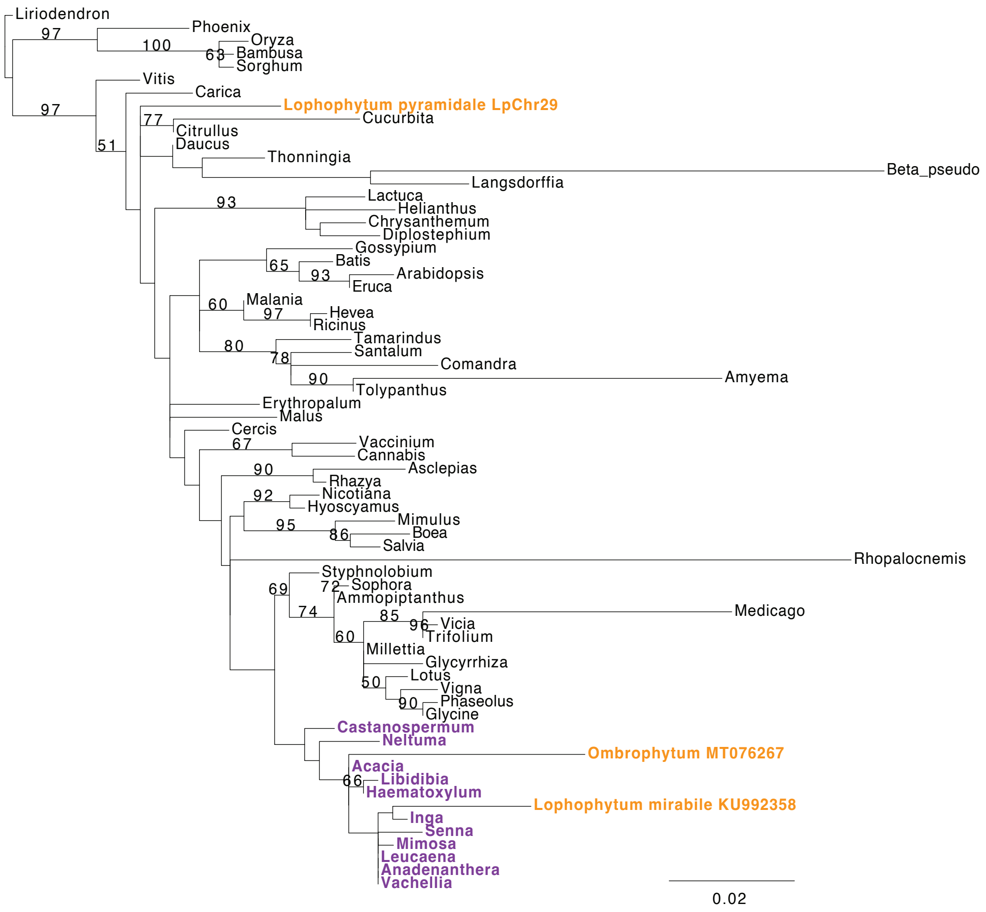
Liriodendron
Phoenix
Oryza
Bambusa
Sorghum
Vitis
Carica
Lophophytum pyramidale LpChr29
Cucurbita
Citrullus
Daucus
Thonningia
Beta_pseudo
Langsdorffia
Lactuca
Helianthus
Chrysanthemum
Diplostephium
Gossypium
Batis
Arabidopsis
Eruca
Malania
Hevea
Ricinus
Tamarindus
Santalum
Comandra
Amyema
Tolypanthus
Erythropalum
Malus
Cercis
Vaccinium
Cannabis
Asclepias
Rhazya
Nicotiana
Hyoscyamus
Mimulus
Boea
Salvia
Rhopalocnemis
Styphnolobium
Sophora
Ammopiptanthus
Medicago
Vicia
Trifolium
Millettia
Glycyrrhiza
Lotus
Vigna
Phaseolus
Glycine
Castanospermum
Neltuma
Ombrophytum MT076267
Acacia
Libidibia
Haematoxylum
Lophophytum mirabile KU992358
Inga
Senna
Mimosa
Leucaena
Anadenanthera
Vachellia
0.02

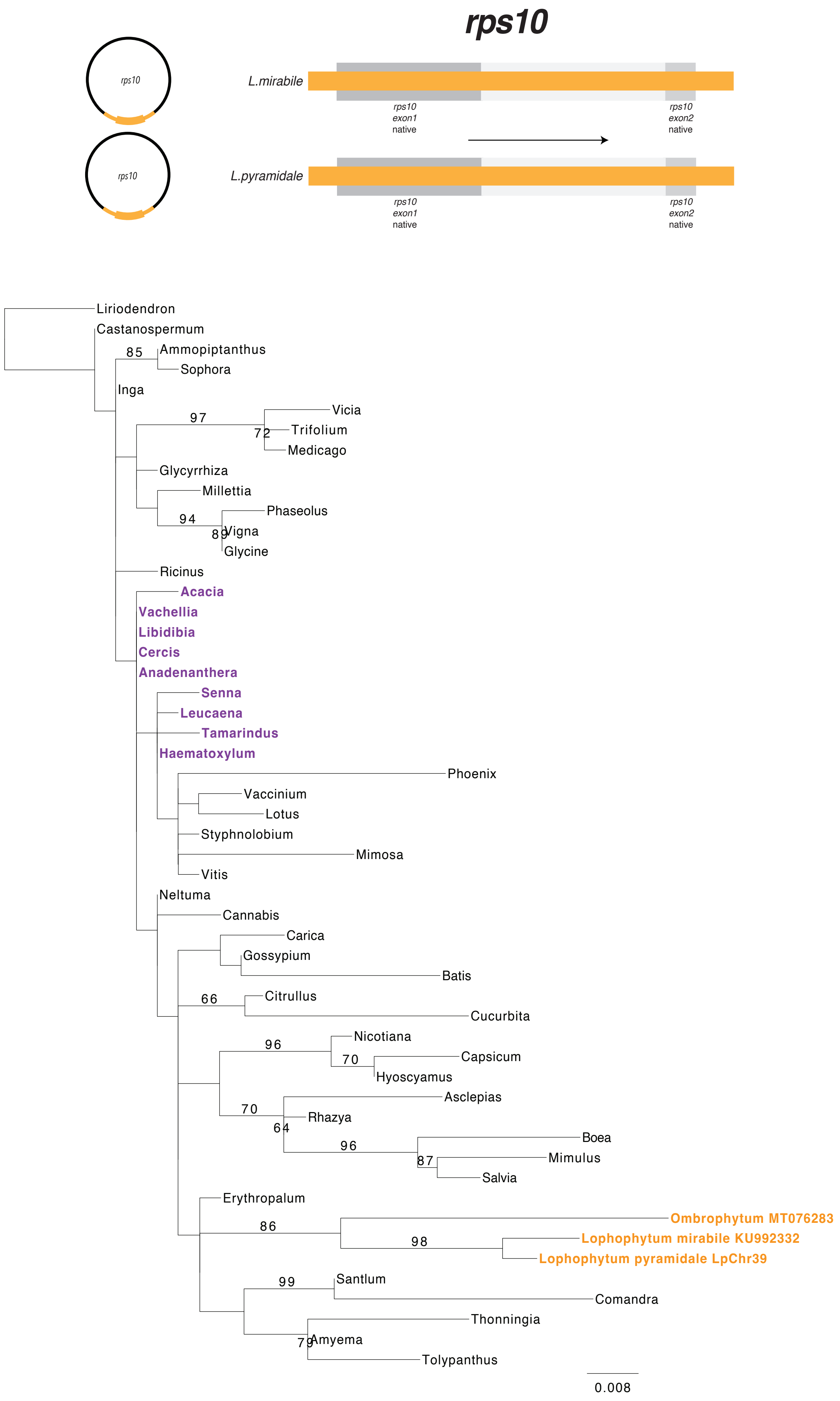

rps10
rps10
rps10
L.mirabile
rps10
exon1
native
rps10
exon2
native
L.pyramidale
rps10
exon1
native
rps10
exon2
native
Liriodendron
Castanospermum
85
Ammopiptanthus
Sophora
Inga
97
Vicia
72
Trifolium
Medicago
Glycyrrhiza
Millettia
94
Phaseolus
89
Vigna
Glycine
Ricinus
Acacia
Vachellia
Libidibia
Cercis
Anadenanthera
Senna
Leucaena
Tamarindus
Haematoxylum
Phoenix
Vaccinium
Lotus
Styphnolobium
Mimosa
Vitis
Neltuma
Cannabis
Carica
Gossypium
Batis
66
Citrullus
Cucurbita
96
Nicotiana
70
Capsicum
Hyoscyamus
Asclepias
70
Rhazya
64
96
Boea
87
Mimulus
Salvia
Erythropalum
86
Ombrophytum MT076283
98
Lophophytum mirabile KU992332
Lophophytum pyramidale LpChr39
99
Santlum
Comandra
Thonningia
79
Amyema
Tolypanthus
0.008

### *rrnS L. pyramidale genes*

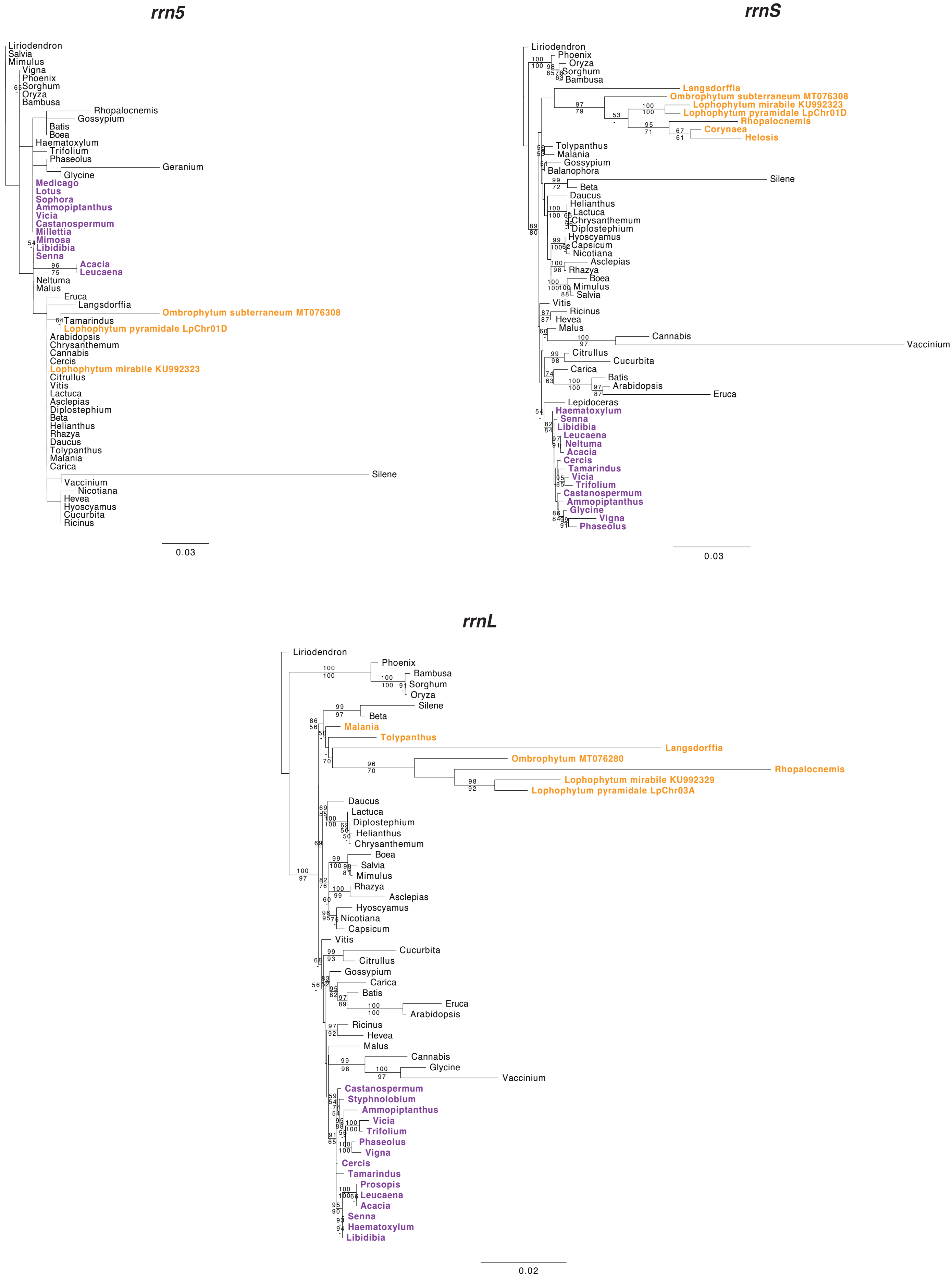

### *trnS L. pyramidale genes*

#### *trnE-UUC* (LpChr07B 19125-20468)

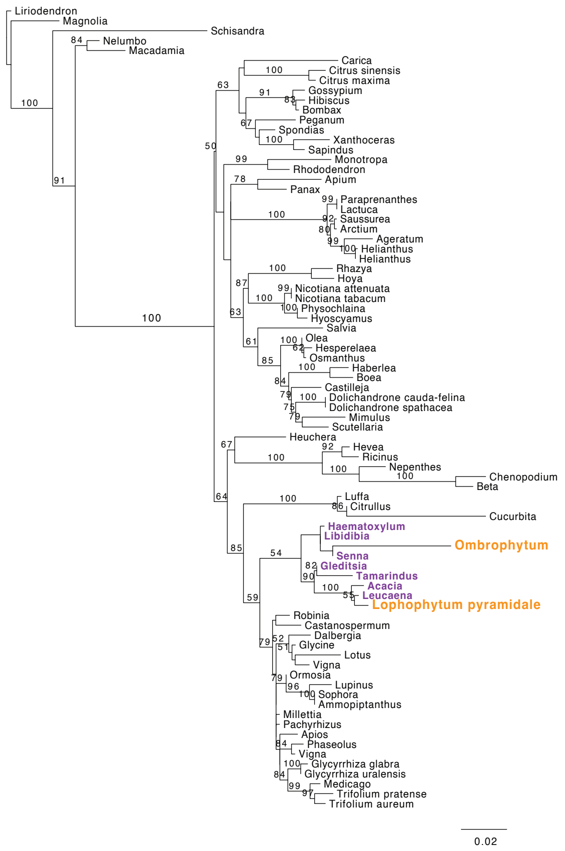


#### *trnK-UUU* (LpChr01C2 66900-67300)

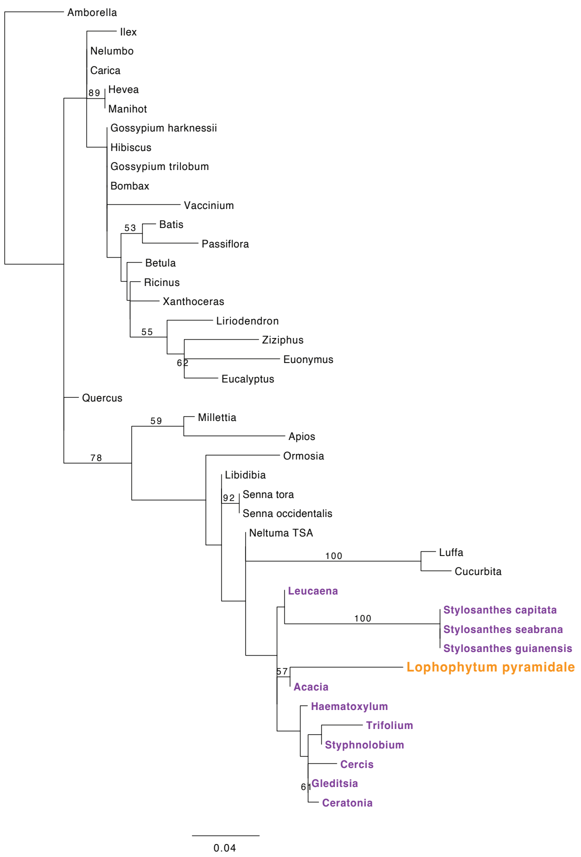


#### *trnI-CAU* (LpChr36 4700-5198)

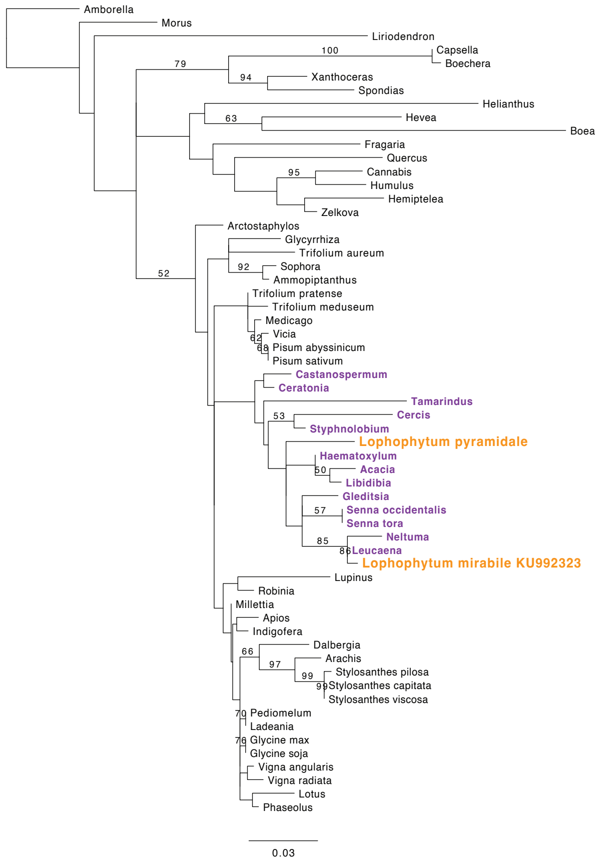


#### *trnM-CAU* cp-like (LpChr43 3900-4299)

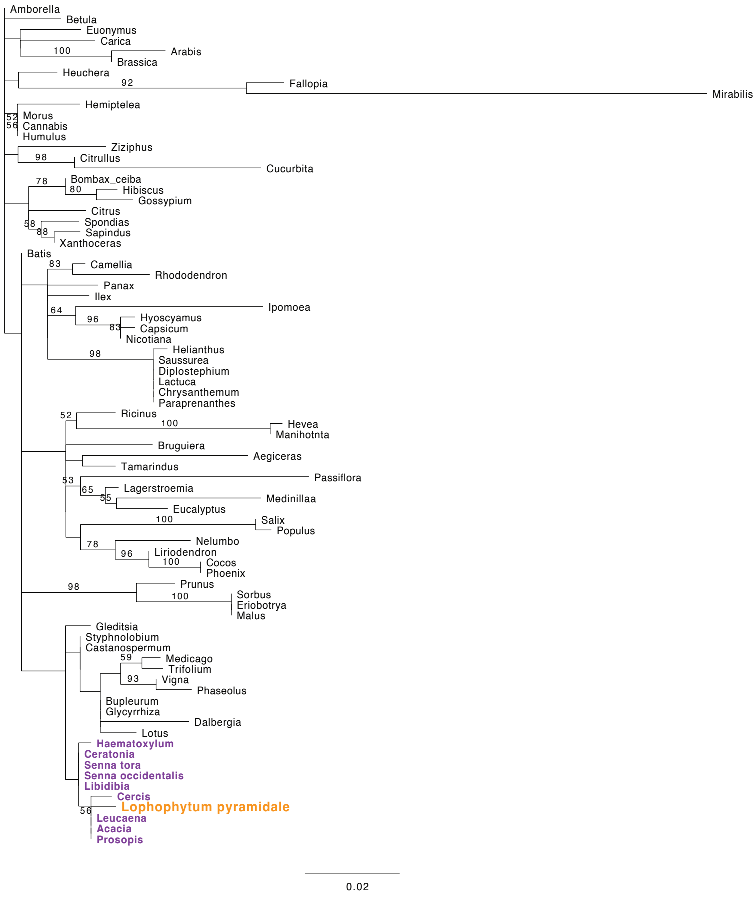

#### *trnfM-CAU* (LpChr03A 13411-13711)

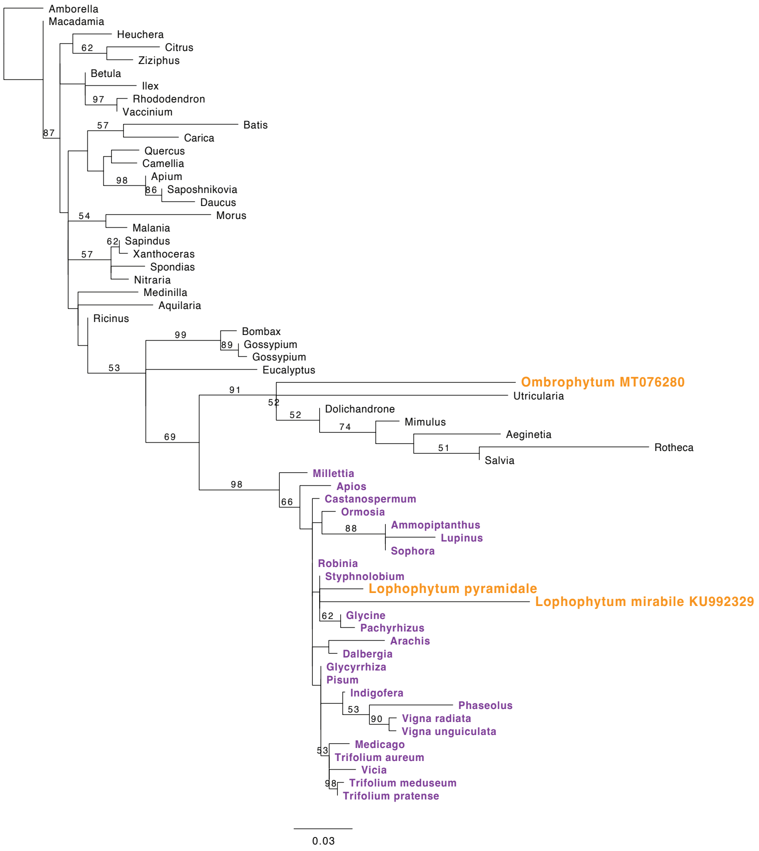


#### *trnW-CCA* cp-like (LpChr04A 6955-7094)
##### (within a foreign MTPT)

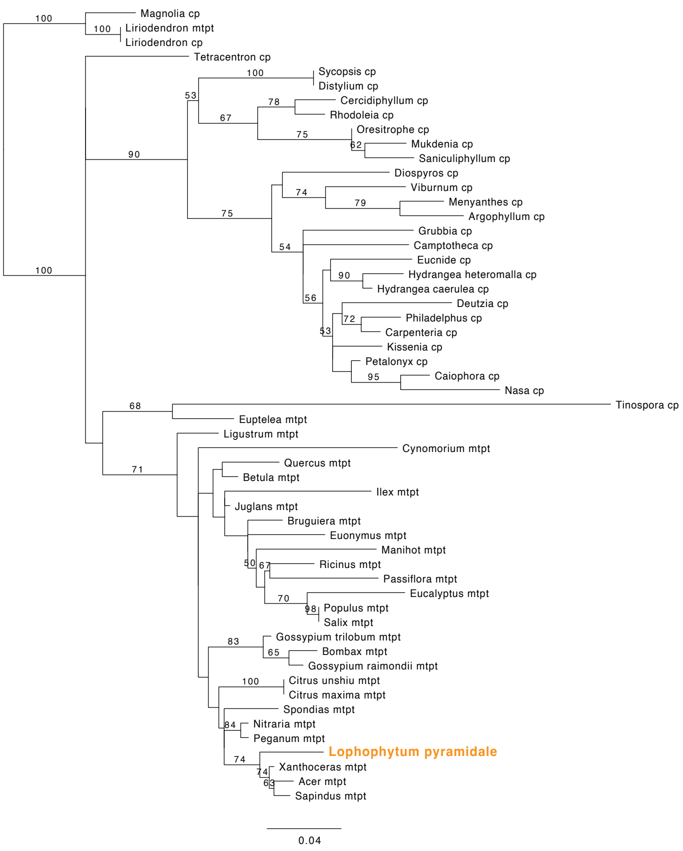

**Figure S4. RNA read depth of the 81 chromosomes in the *Lophophytum pyramidale* mtDNA.** The curves represent the number of RNA reads aligned to each nucleotide position. Reads corresponding to transcription from left to right and right to left are shown as green and orange shaded curves on the positive and negative y-axes, respectively. For each chromosome, an additional magnified plot Y-axis (ylim = 10,000) to enhance the visualization of regions with lower read coverage is included. Grey boxes indicate the locations of mitochondrial genes, plastid-derived sequences (cp-like), and repeats ("R" followed by the length of the repeat). The RNA reads cover 99.77% of the mitochondrial genome considering both strands. The coverage is not 100% likely due to a small difference in the mitochondrial genome of the individual from which the RNA was extracted and the individual used for mtDNA assembly. For each protein-coding gene, the phylogenetic origin is indicated: chimeric ("C"), foreign (F) or native (N).Noticeably, a few non-coding regions show elevated coverage, such as regions in LpChr1A (5–6 kb) and LpChr04C (3.5–5.5 kb) with over 10,000 aligned reads. BLASTn searches indicate that these regions are of foreign origin, associated with Fabaceae species, and do not contain any annotated coding sequences.

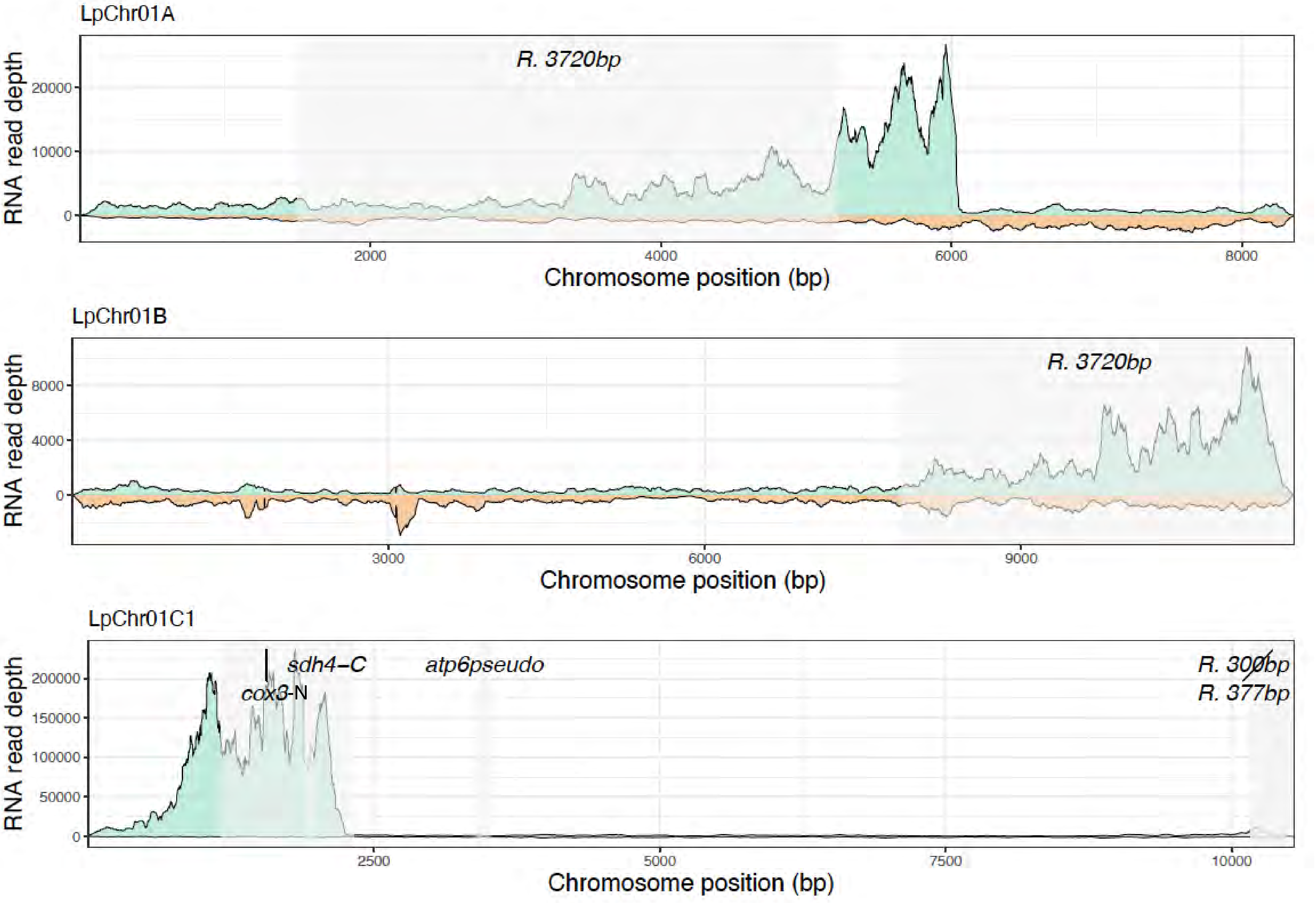

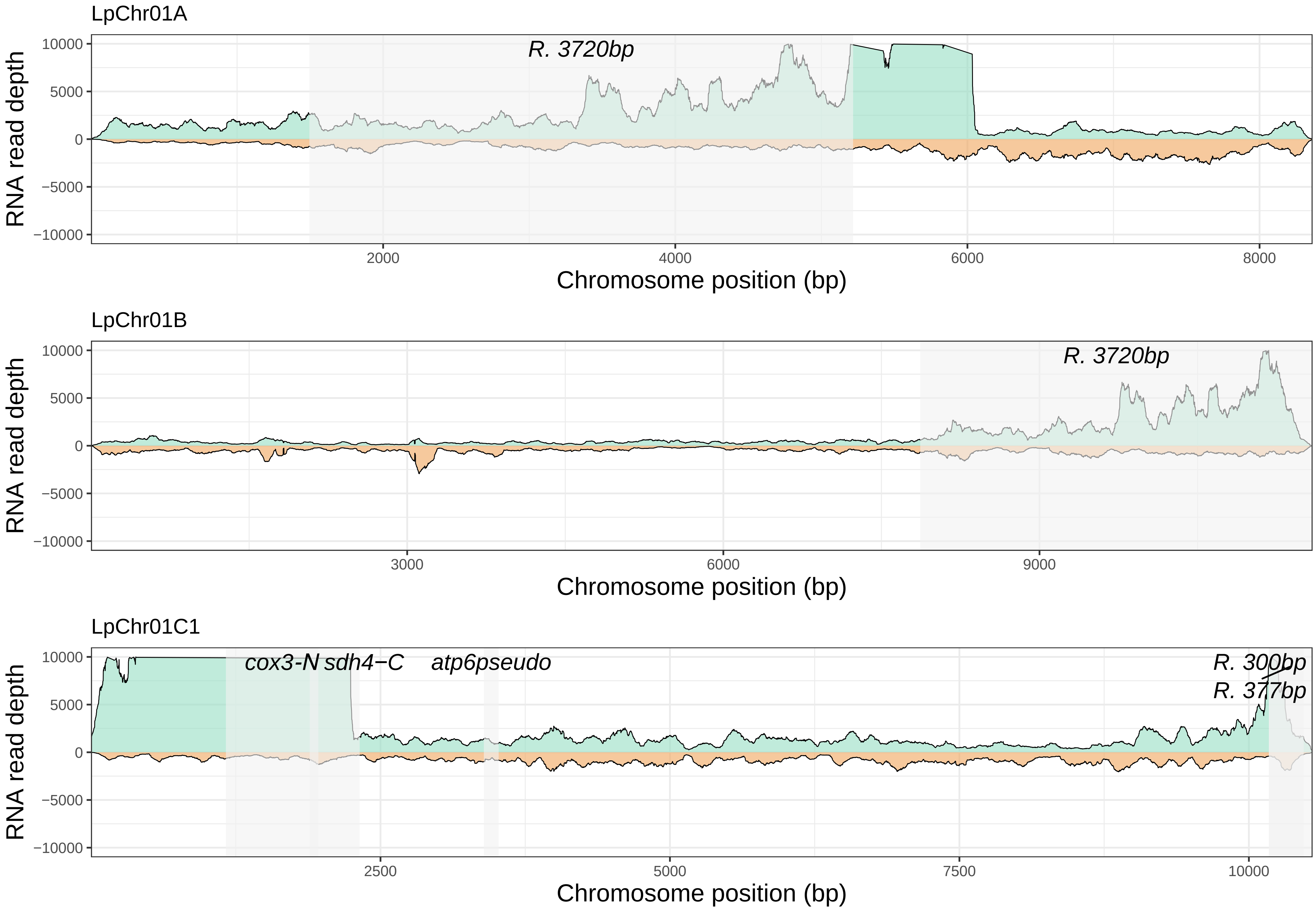
LpChr01A
R. 3720bp
RNA read depth
10000
5000
0
−5000
−10000
2000
4000
6000
8000
Chromosome position (bp)
LpChr01B
R. 3720bp
RNA read depth
10000
5000
0
−5000
−10000
3000
6000
9000
Chromosome position (bp)
LpChr01C1
cox3-N
sdh4−C
atp6pseudo
R. 300bp
R. 377bp
RNA read depth
10000
5000
0
−5000
−10000
2500
5000
7500
10000
Chromosome position (bp)

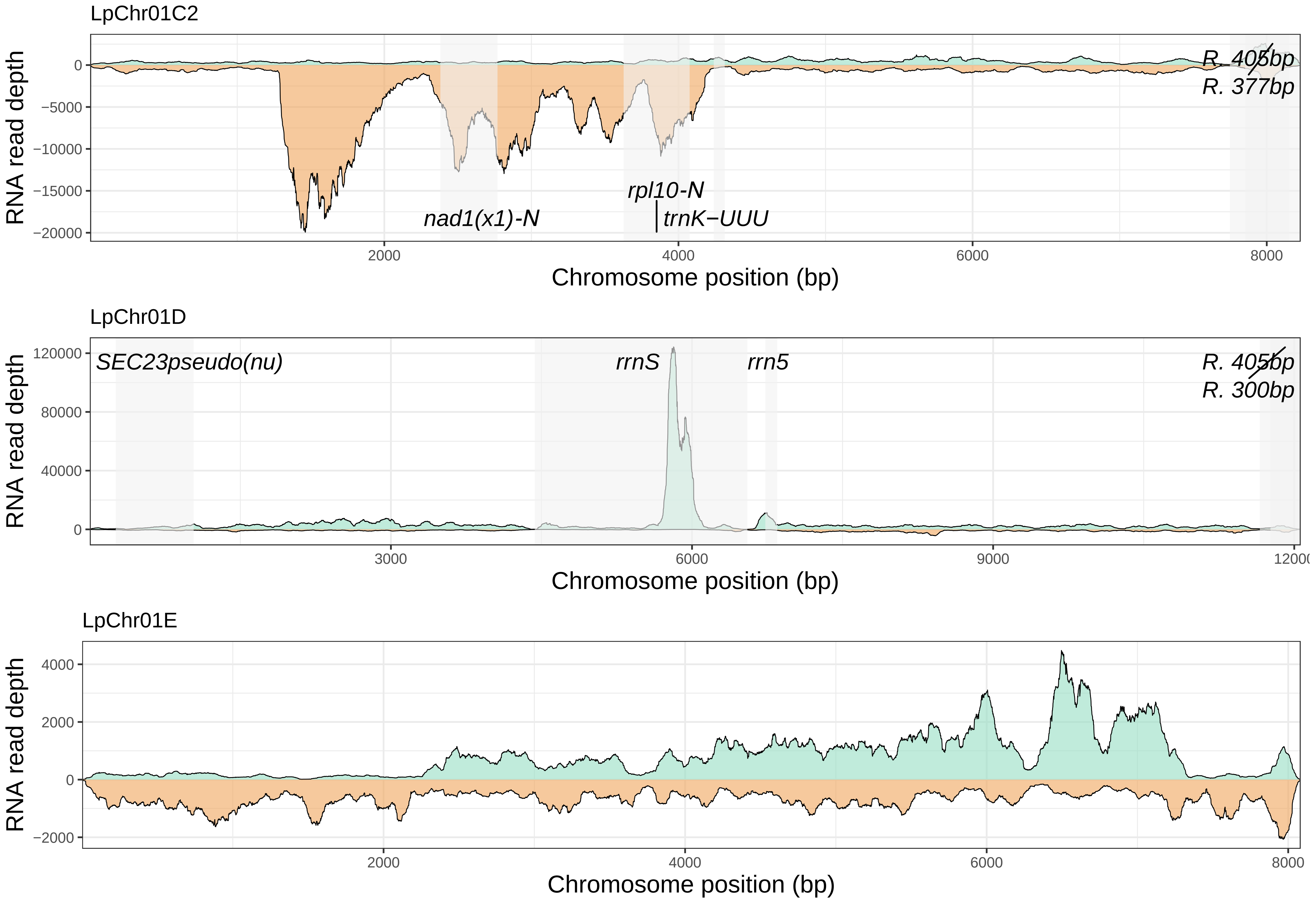

LpChr01C2
RNA read depth
0
−5000
−10000
−15000
−20000
nad1(x1)-N
rpl10-N
trnK−UUU
R. 405bp
R. 377bp
2000
4000
6000
8000
Chromosome position (bp)
LpChr01D
RNA read depth
120000
80000
40000
0
SEC23pseudo(nu)
rrnS
rrn5
R. 405bp
R. 300bp
3000
6000
9000
12000
Chromosome position (bp)
LpChr01E
RNA read depth
4000
2000
0
−2000
2000
4000
6000
8000
Chromosome position (bp)

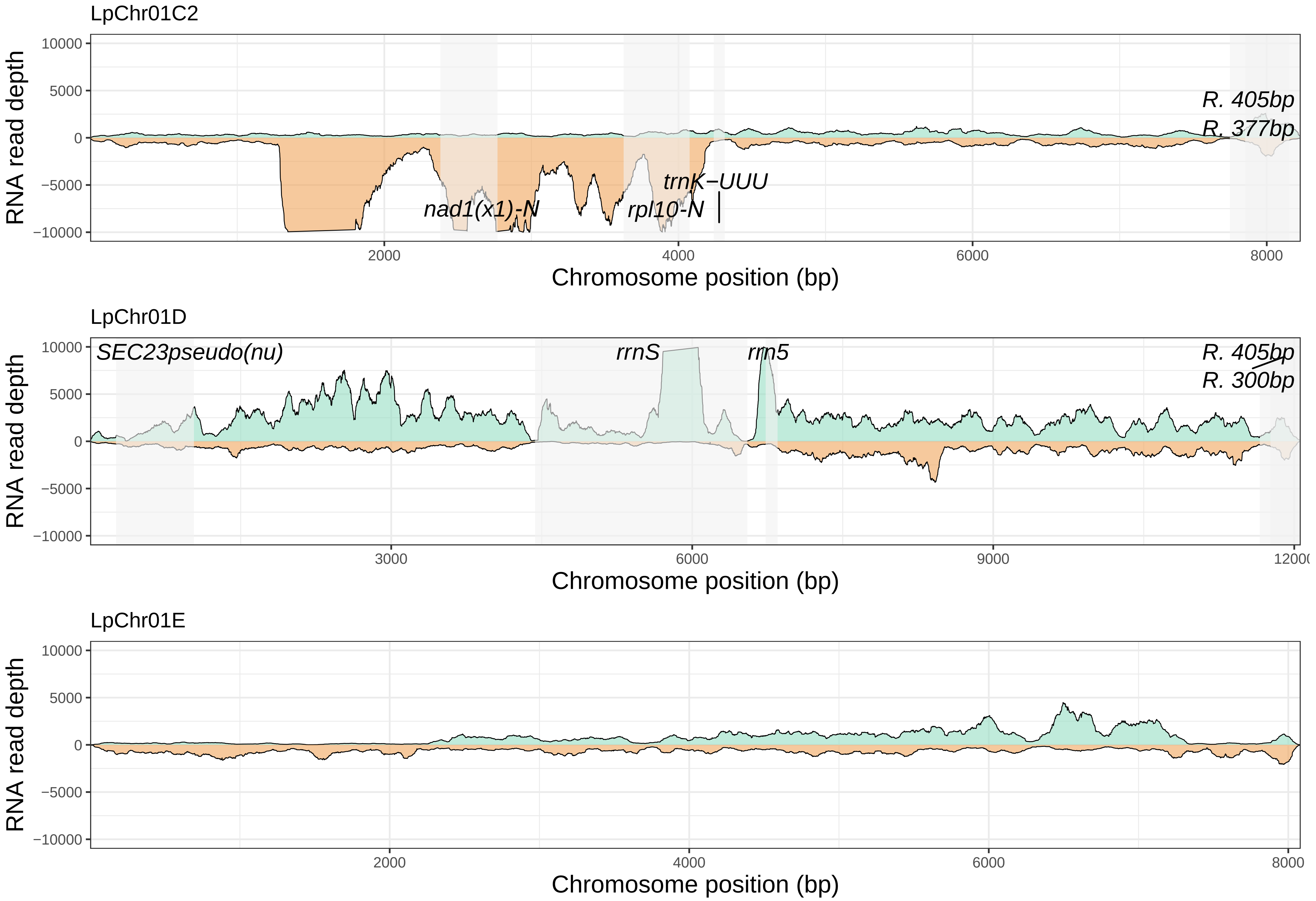

LpChr01C2
RNA read depth
10000
5000
0
−5000
−10000
R. 405bp
R. 377bp
trnK−UUU
nad1(x1)-N
rpl10-N
2000
4000
6000
8000
Chromosome position (bp)
LpChr01D
SEC23pseudo(nu)
rrnS
rrn5
R. 405bp
R. 300bp
3000
6000
9000
Chromosome position (bp)
LpChr01E
2000
4000
6000
8000
Chromosome position (bp)

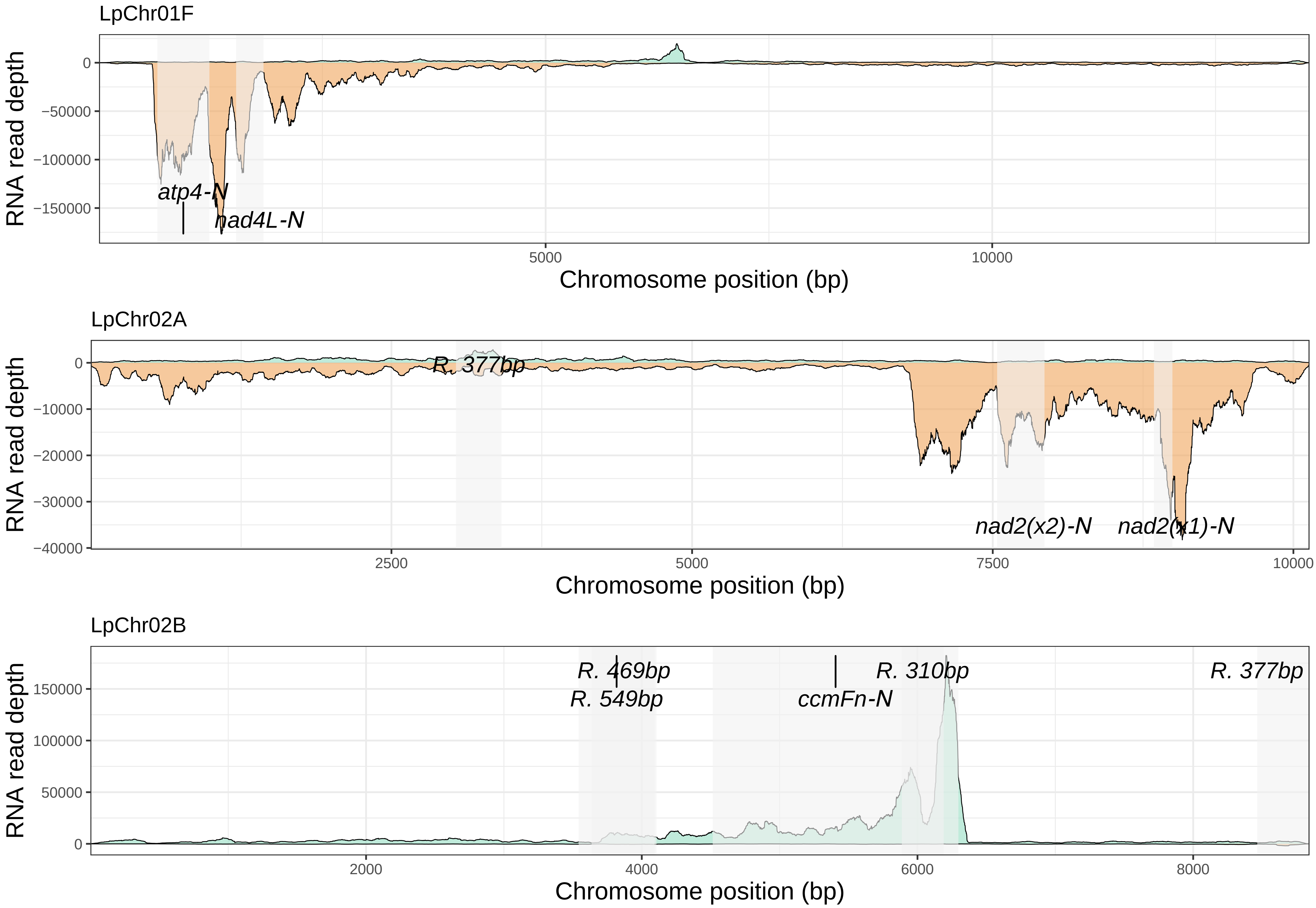
LpChr01F
RNA read depth
0
−50000
−100000
−150000
atp4-N
nad4L-N
5000
10000
Chromosome position (bp)
LpChr02A
RNA read depth
0
−10000
−20000
−30000
−40000
R. 377bp
nad2(x2)-N
nad2(x1)-N
2500
5000
7500
10000
Chromosome position (bp)
LpChr02B
RNA read depth
150000
100000
50000
0
R. 469bp
R. 549bp
ccmFn-N
R. 310bp
R. 377bp
2000
4000
6000
8000
Chromosome position (bp)

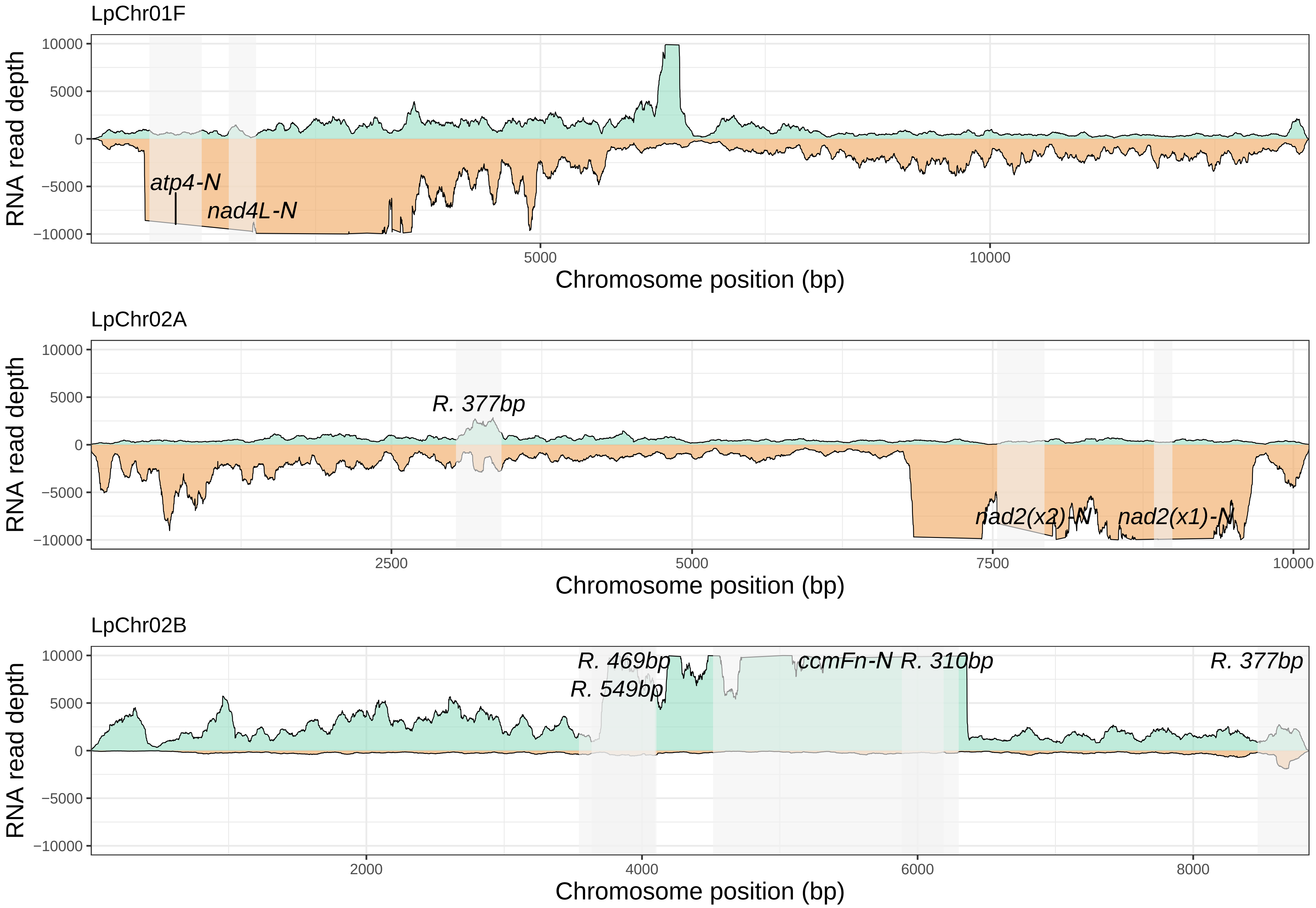

LpChr01F
RNA read depth
10000
5000
0
−5000
−10000
atp4-N
nad4L-N
5000
10000
Chromosome position (bp)
LpChr02A
RNA read depth
R. 377bp
nad2(x2)-N
nad2(x1)-N
2500
5000
7500
10000
Chromosome position (bp)
LpChr02B
RNA read depth
R. 469bp
R. 549bp
ccmFn-N
R. 310bp
R. 377bp
2000
4000
6000
8000
Chromosome position (bp)

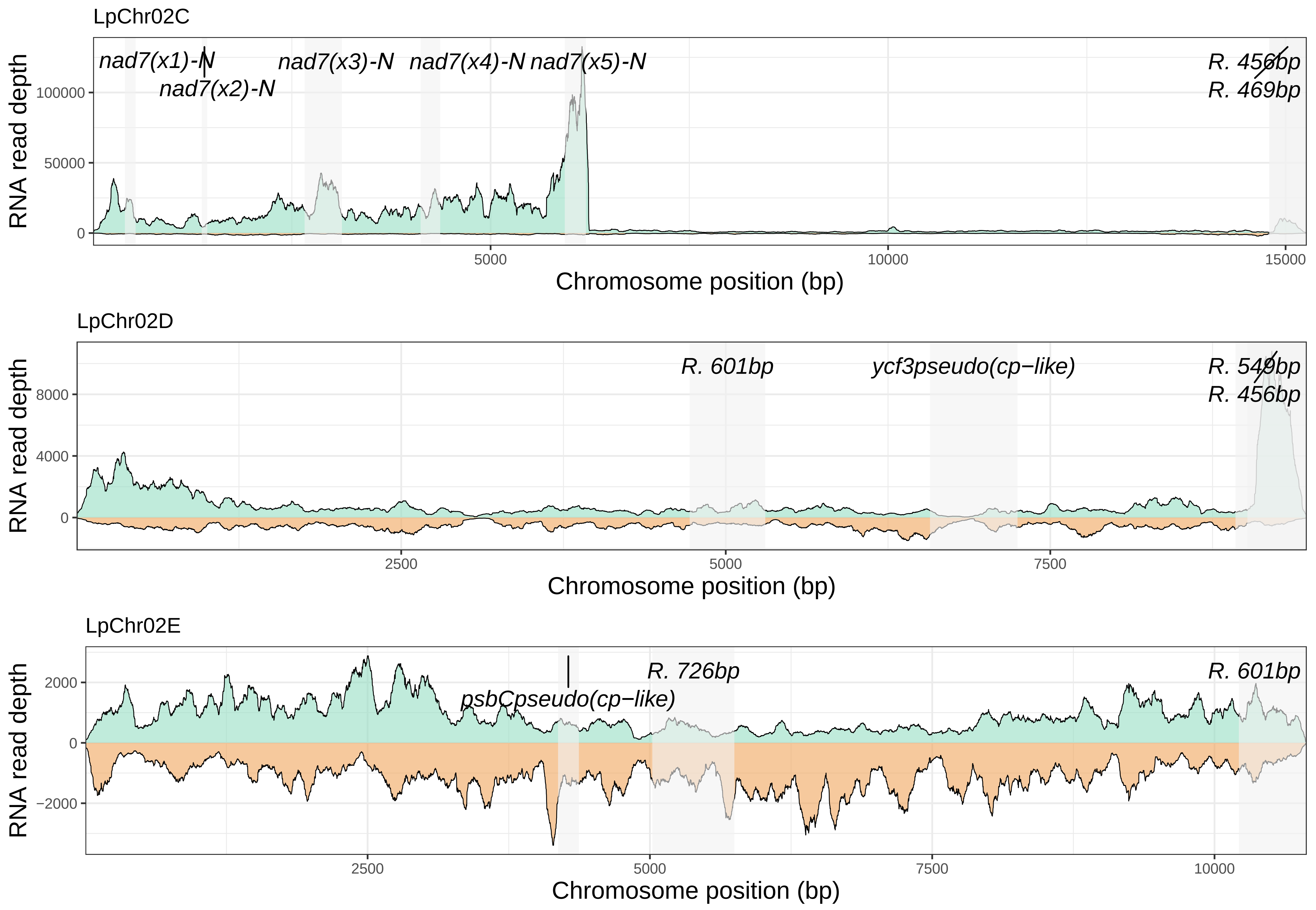
LpChr02C
nad7(x1)-N
nad7(x2)-N
nad7(x3)-N
nad7(x4)-N
nad7(x5)-N
R. 456bp
R. 469bp
RNA read depth
100000
50000
0
5000
10000
15000
Chromosome position (bp)
LpChr02D
R. 601bp
ycf3pseudo(cp−like)
R. 549bp
R. 456bp
RNA read depth
8000
4000
0
2500
5000
7500
Chromosome position (bp)
LpChr02E
R. 726bp
psbCpseudo(cp−like)
R. 601bp
RNA read depth
2000
0
−2000
2500
5000
7500
10000
Chromosome position (bp)

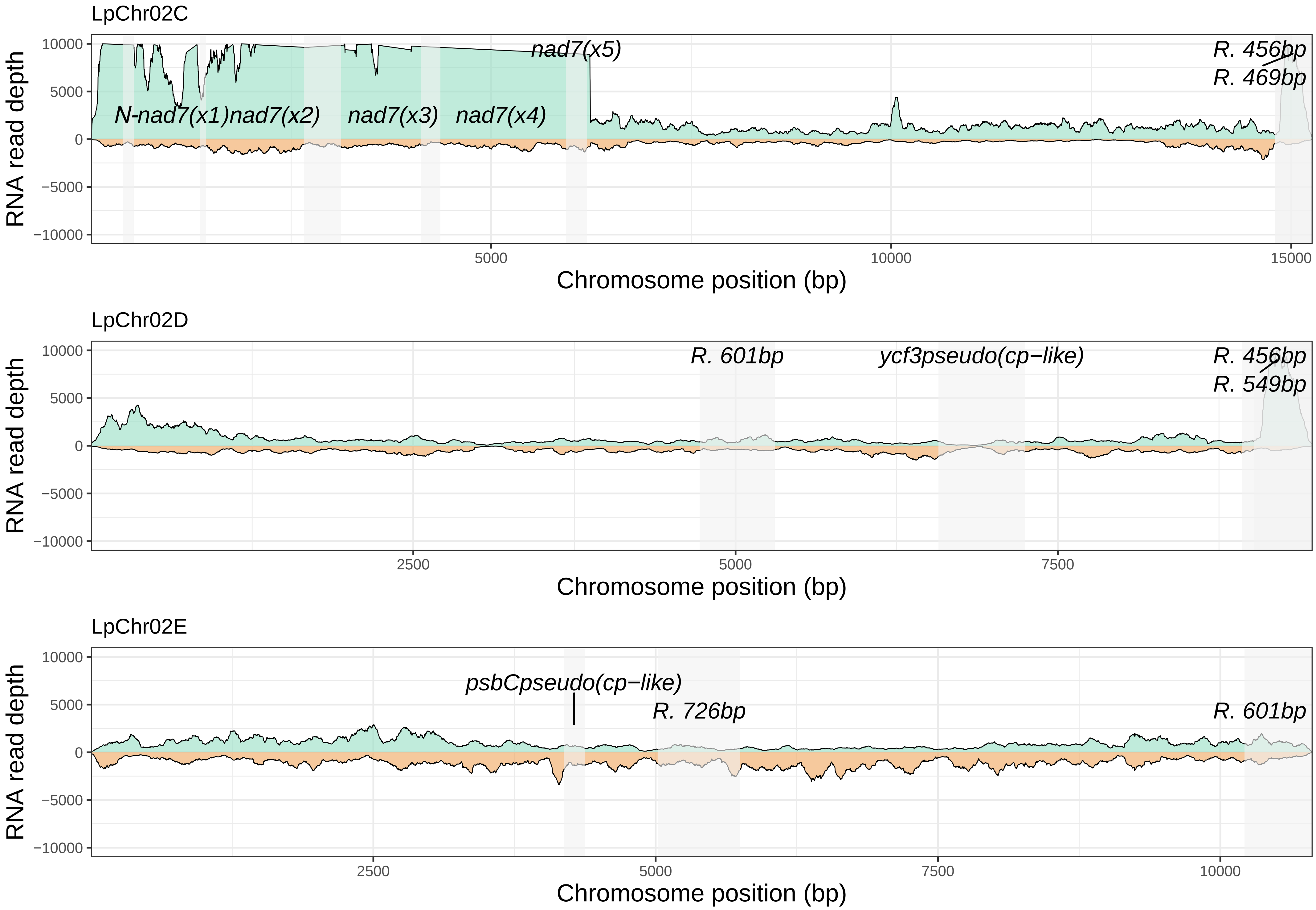

LpChr02C
nad7(x5)
R. 456bp
R. 469bp
N-nad7(x1)
nad7(x2)
nad7(x3)
nad7(x4)
RNA read depth
Chromosome position (bp)
LpChr02D
R. 601bp
ycf3pseudo(cp−like)
R. 456bp
R. 549bp
RNA read depth
Chromosome position (bp)
LpChr02E
psbCpseudo(cp−like)
R. 726bp
R. 601bp
RNA read depth
Chromosome position (bp)

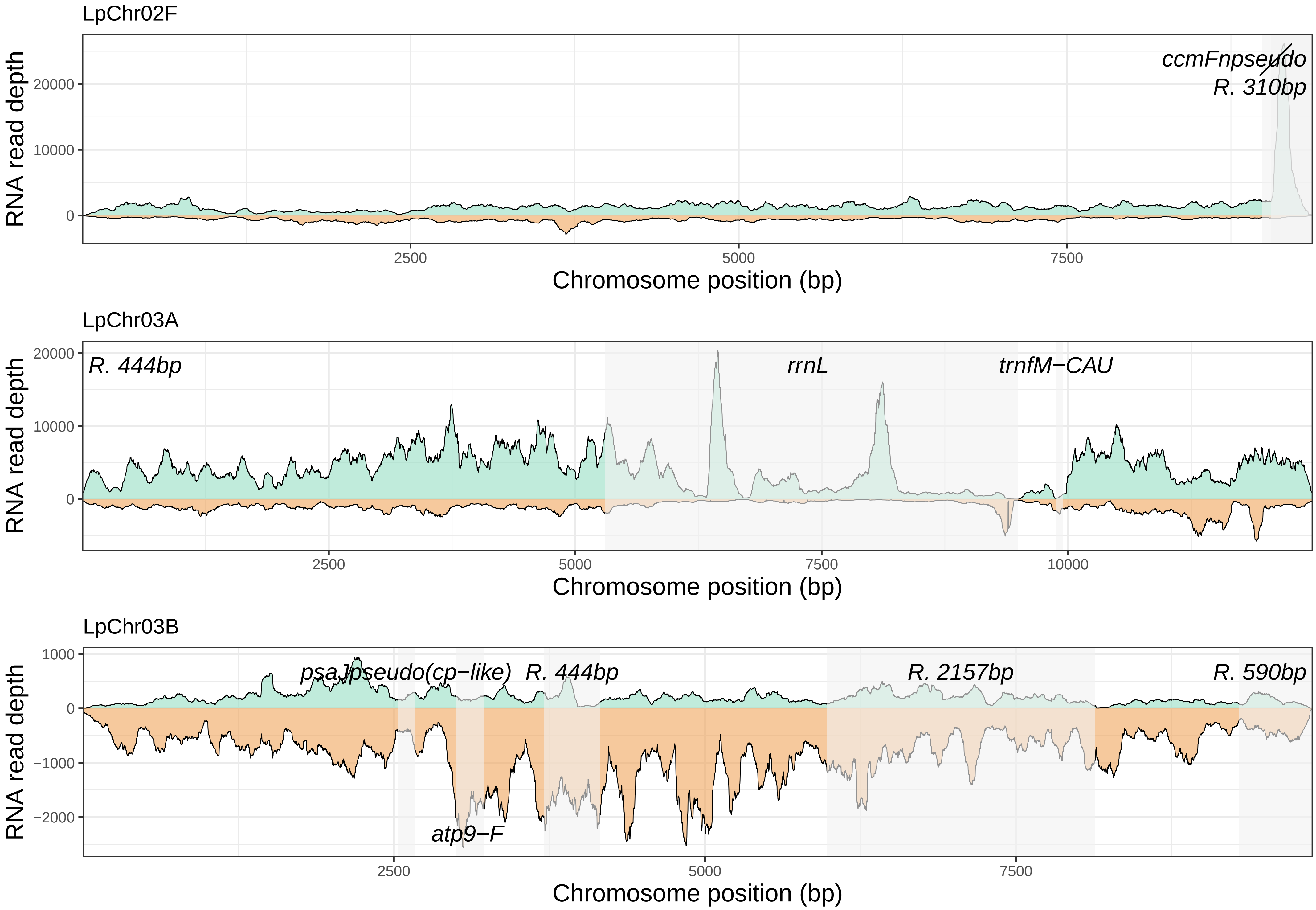

LpChr02F
RNA read depth
20000
10000
0
ccmFnpseudo
R. 310bp
2500
5000
7500
Chromosome position (bp)
LpChr03A
RNA read depth
20000
10000
0
R. 444bp
rrnL
trnfM−CAU
2500
5000
7500
10000
Chromosome position (bp)
LpChr03B
RNA read depth
1000
0
−1000
−2000
psaJpseudo(cp−like)
R. 444bp
R. 2157bp
R. 590bp
atp9−F
2500
5000
7500
Chromosome position (bp)

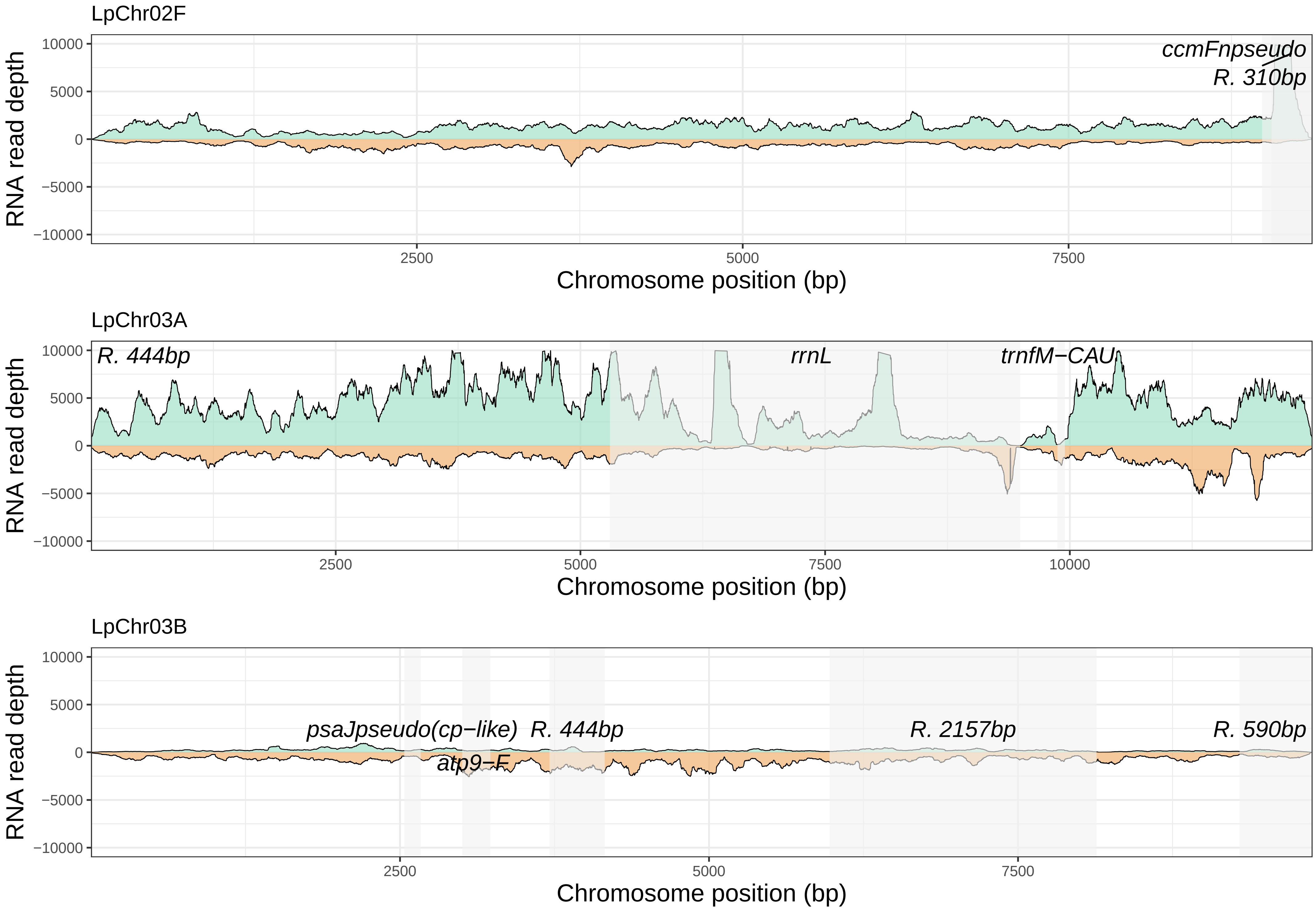

LpChr02F
ccmFnpseudo
R. 310bp
RNA read depth
10000
5000
0
−5000
−10000
2500
5000
7500
Chromosome position (bp)
LpChr03A
R. 444bp
rrnL
trnfM−CAU
RNA read depth
10000
5000
0
−5000
−10000
2500
5000
7500
10000
Chromosome position (bp)
LpChr03B
psaJpseudo(cp−like)
R. 444bp
atp9−F
R. 2157bp
R. 590bp
RNA read depth
10000
5000
0
−5000
−10000
2500
5000
7500
Chromosome position (bp)

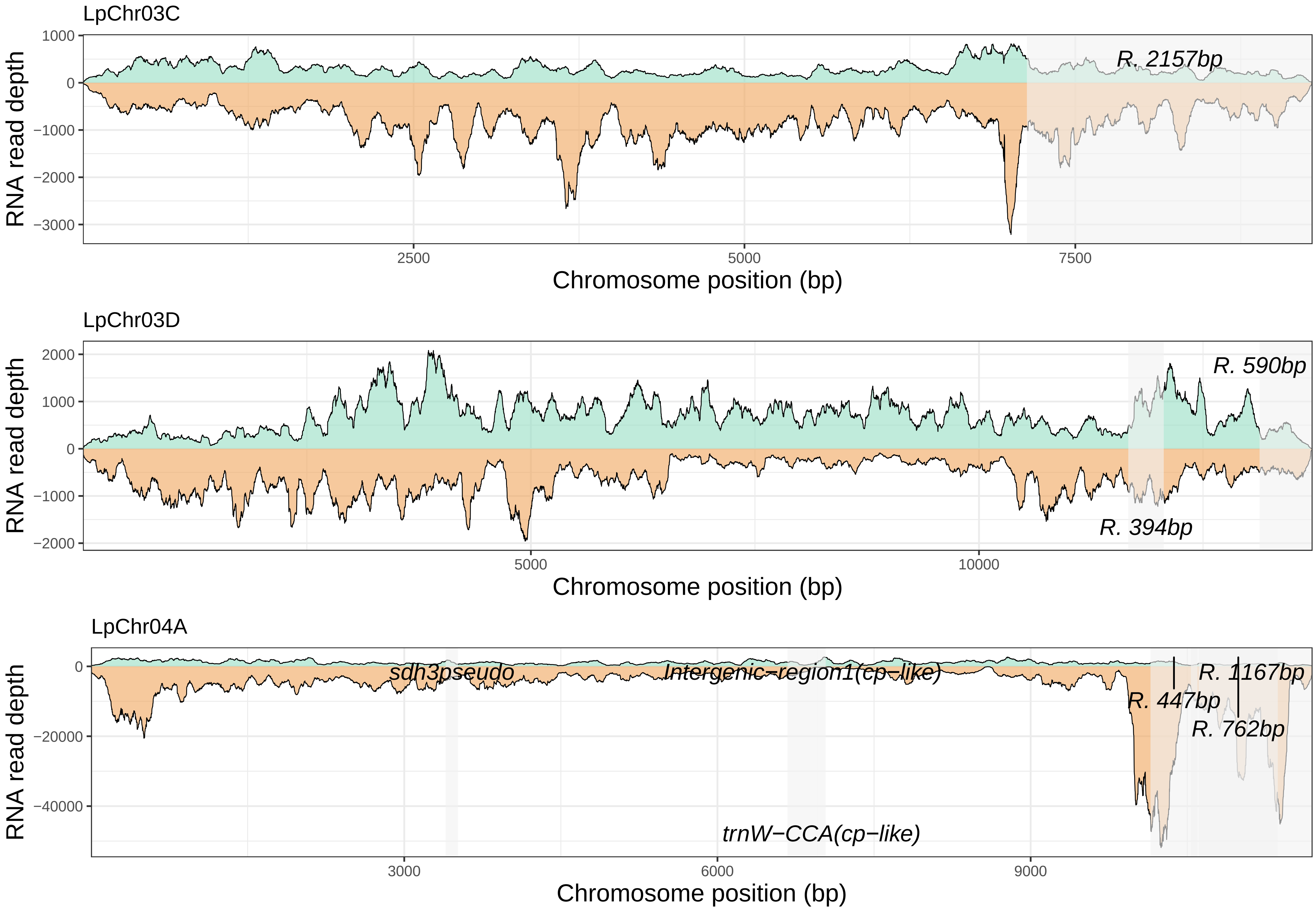
LpChr03C
R. 2157bp
RNA read depth
1000
0
−1000
−2000
−3000
2500
5000
7500
Chromosome position (bp)
LpChr03D
R. 590bp
R. 394bp
RNA read depth
2000
1000
0
−1000
−2000
5000
10000
Chromosome position (bp)
LpChr04A
sdh3pseudo
Intergenic−region1(cp−like)
R. 1167bp
R. 447bp
R. 762bp
trnW−CCA(cp−like)
RNA read depth
0
−20000
−40000
3000
6000
9000
Chromosome position (bp)

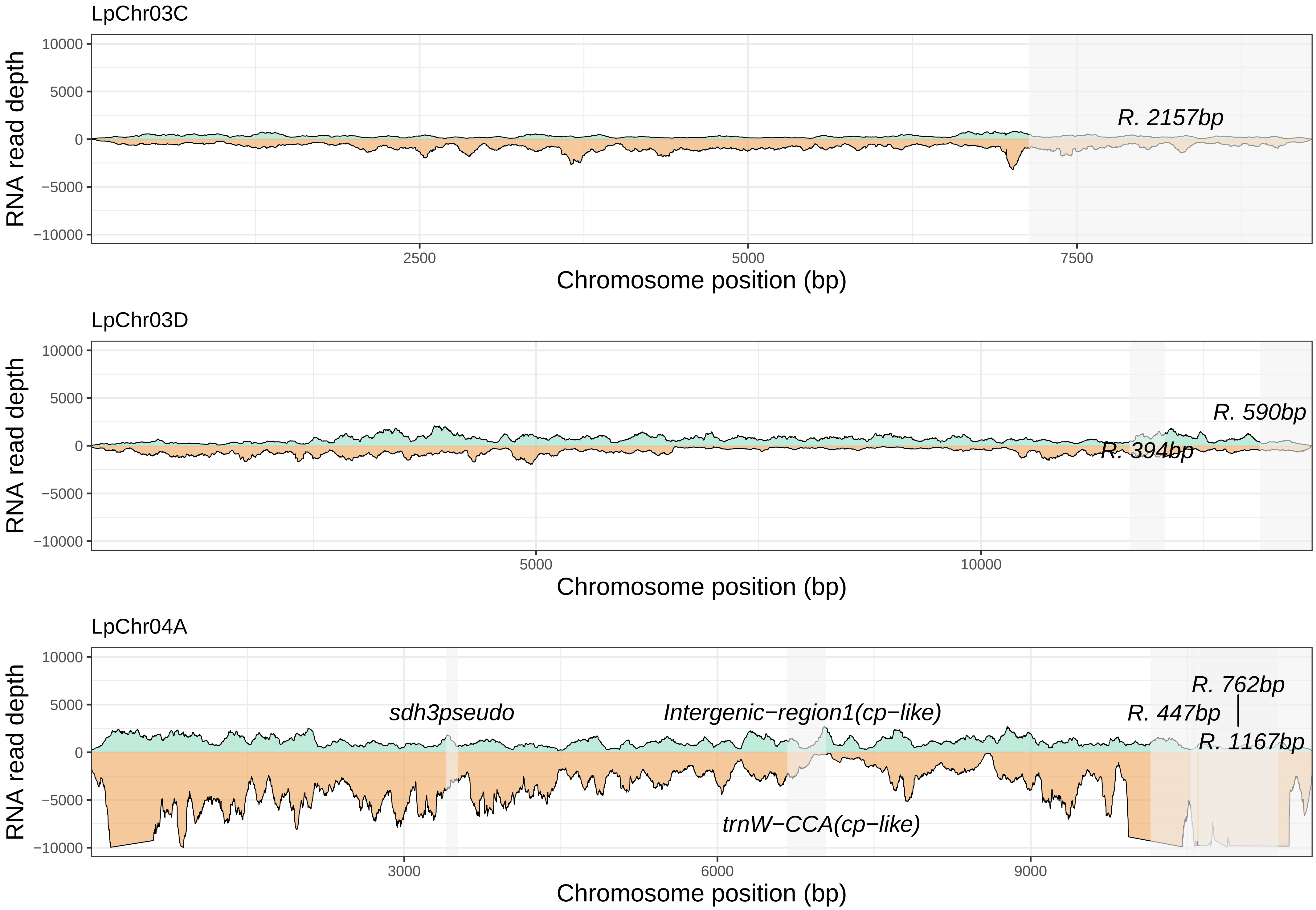
LpChr03C
RNA read depth
10000
5000
0
−5000
−10000
2500
5000
7500
Chromosome position (bp)
R. 2157bp
LpChr03D
RNA read depth
10000
5000
0
−5000
−10000
5000
10000
Chromosome position (bp)
R. 590bp
R. 394bp
LpChr04A
RNA read depth
10000
5000
0
−5000
−10000
3000
6000
9000
Chromosome position (bp)
sdh3pseudo
Intergenic−region1(cp−like)
trnW−CCA(cp−like)
R. 762bp
R. 447bp
R. 1167bp

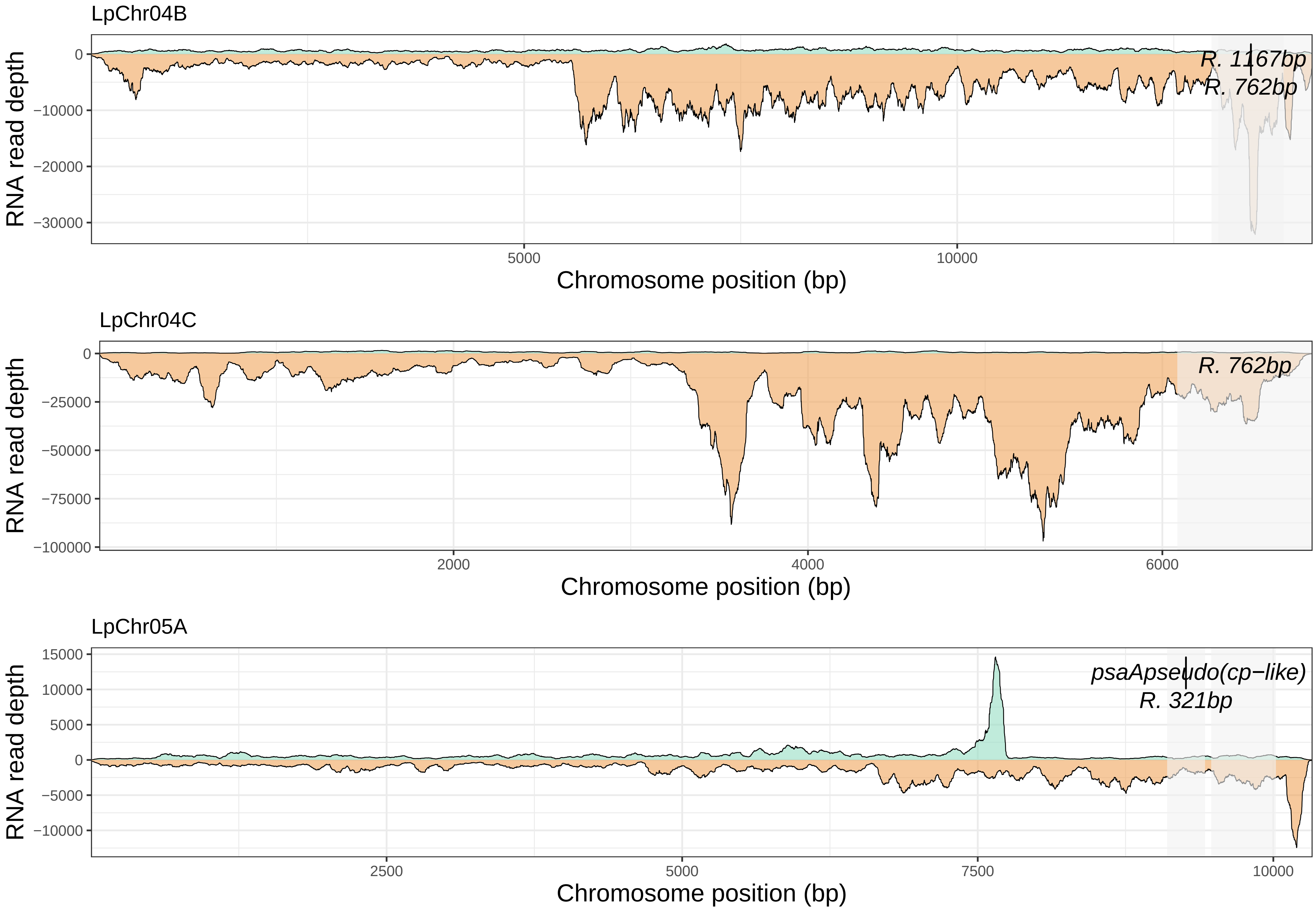
LpChr04B
RNA read depth
0
−10000
−20000
−30000
R. 1167bp
R. 762bp
5000
10000
Chromosome position (bp)
LpChr04C
RNA read depth
0
−25000
−50000
−75000
−100000
R. 762bp
2000
4000
6000
Chromosome position (bp)
LpChr05A
RNA read depth
15000
10000
5000
0
−5000
−10000
psaApseudo(cp−like)
R. 321bp
2500
5000
7500
10000
Chromosome position (bp)

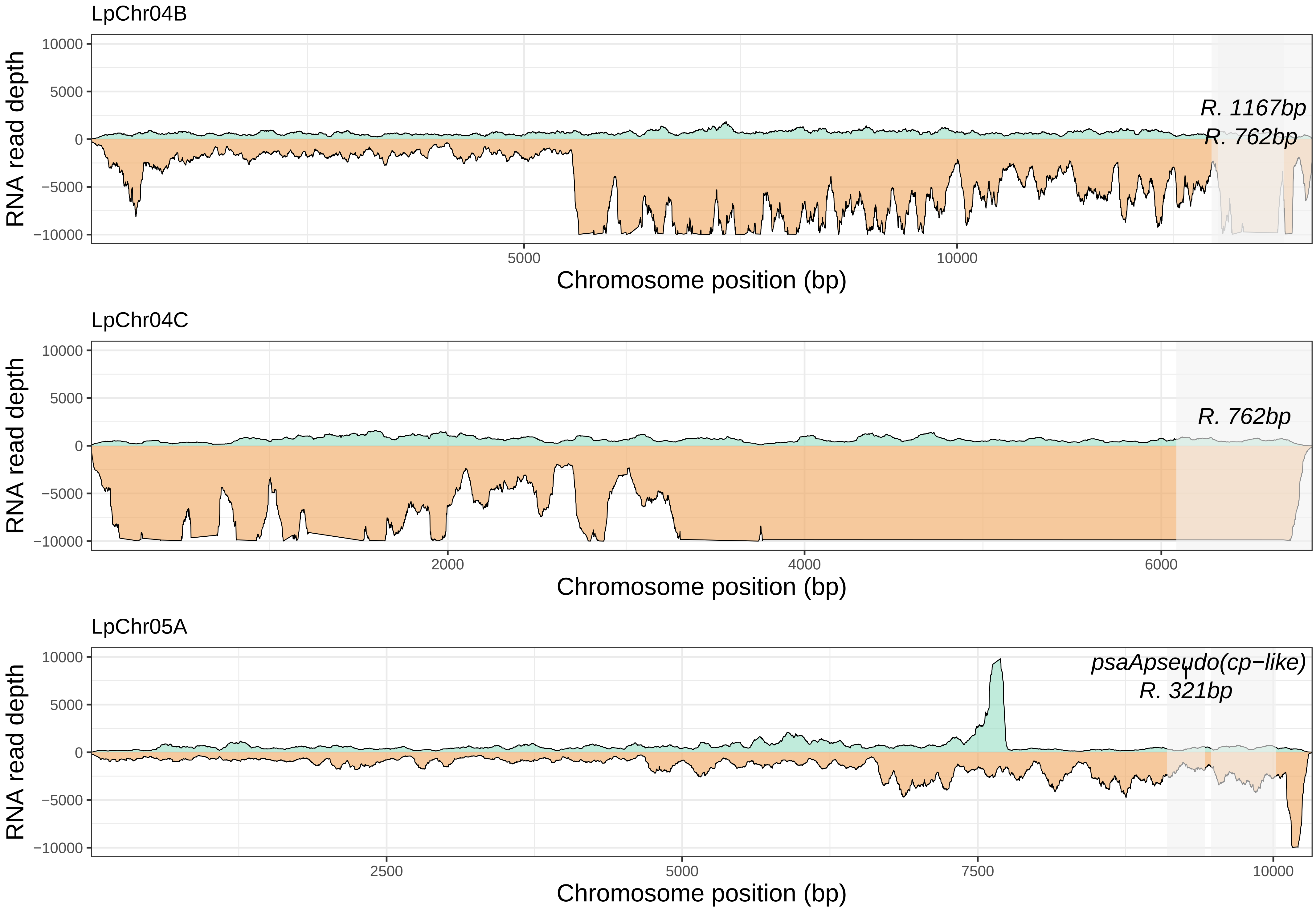

LpChr04B
RNA read depth
10000
5000
0
−5000
−10000
5000
10000
Chromosome position (bp)
R. 1167bp
R. 762bp
LpChr04C
RNA read depth
10000
5000
0
−5000
−10000
2000
4000
6000
Chromosome position (bp)
R. 762bp
LpChr05A
RNA read depth
10000
5000
0
−5000
−10000
2500
5000
7500
10000
Chromosome position (bp)
psaApseudo(cp−like)
R. 321bp

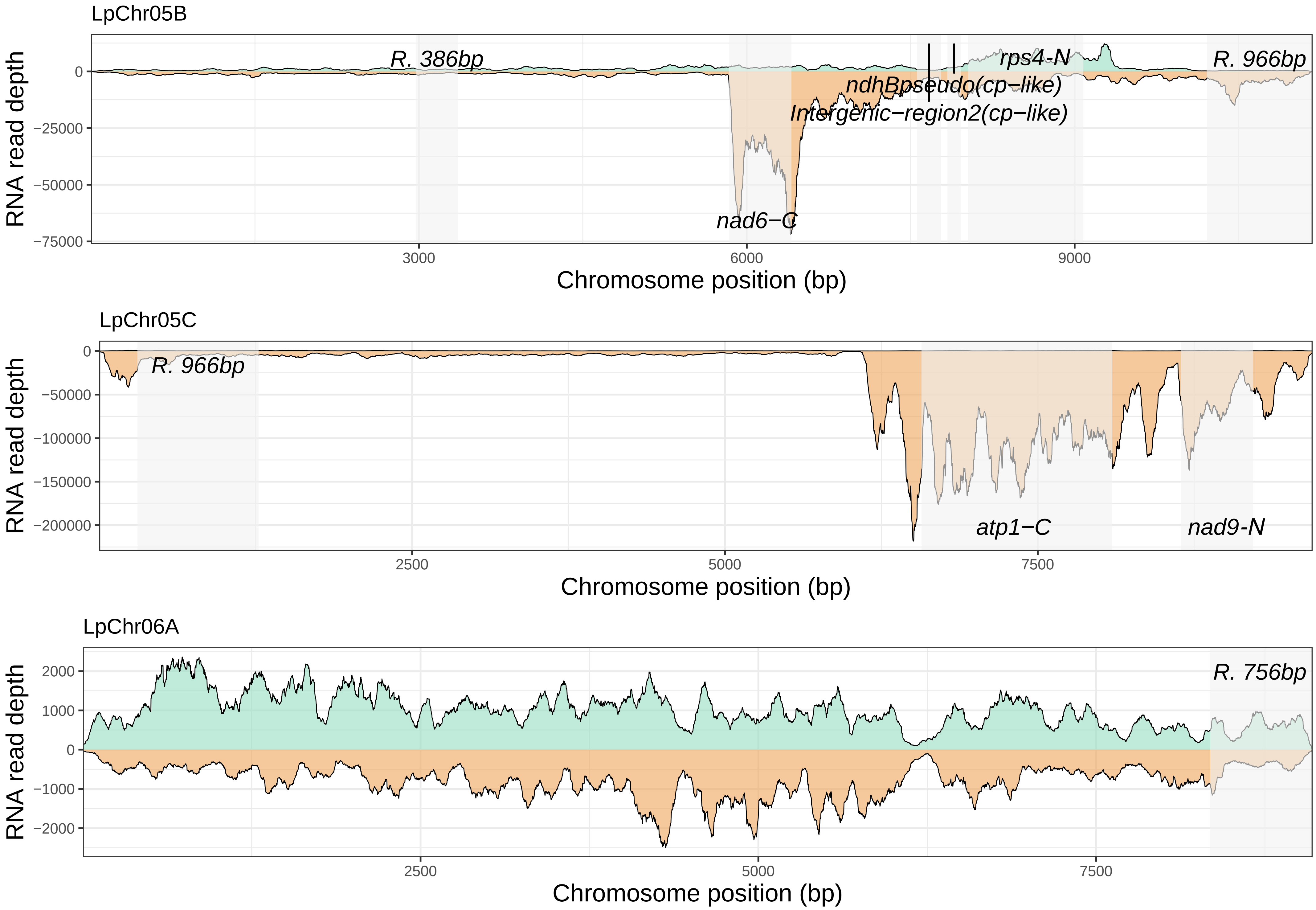

LpChr05B
R. 386bp
rps4-N
R. 966bp
ndhBpseudo(cp−like)
Intergenic−region2(cp−like)
nad6−C
RNA read depth
0
−25000
−50000
−75000
3000
6000
9000
Chromosome position (bp)
LpChr05C
R. 966bp
atp1−C
nad9-N
RNA read depth
0
−50000
−100000
−150000
−200000
2500
5000
7500
Chromosome position (bp)
LpChr06A
R. 756bp
RNA read depth
2000
1000
0
−1000
−2000
2500
5000
7500
Chromosome position (bp)

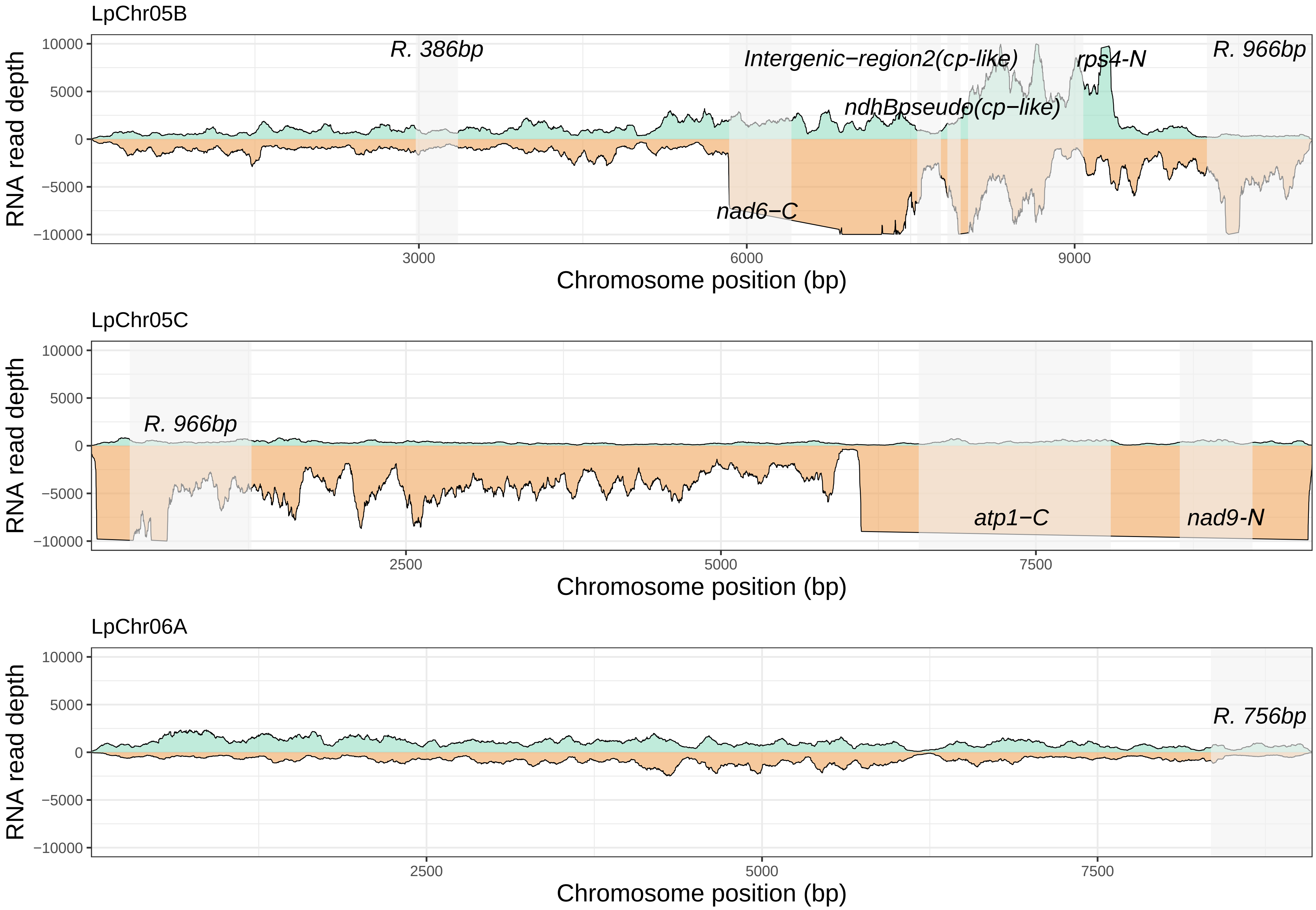
LpChr05B
R. 386bp
Intergenic−region2(cp-like)
rps4-N
R. 966bp
ndhBpseudo(cp−like)
nad6−C
RNA read depth
10000
5000
0
−5000
−10000
3000
6000
9000
Chromosome position (bp)
LpChr05C
R. 966bp
atp1−C
nad9-N
RNA read depth
10000
5000
0
−5000
−10000
2500
5000
7500
Chromosome position (bp)
LpChr06A
R. 756bp
RNA read depth
10000
5000
0
−5000
−10000
2500
5000
7500
Chromosome position (bp)

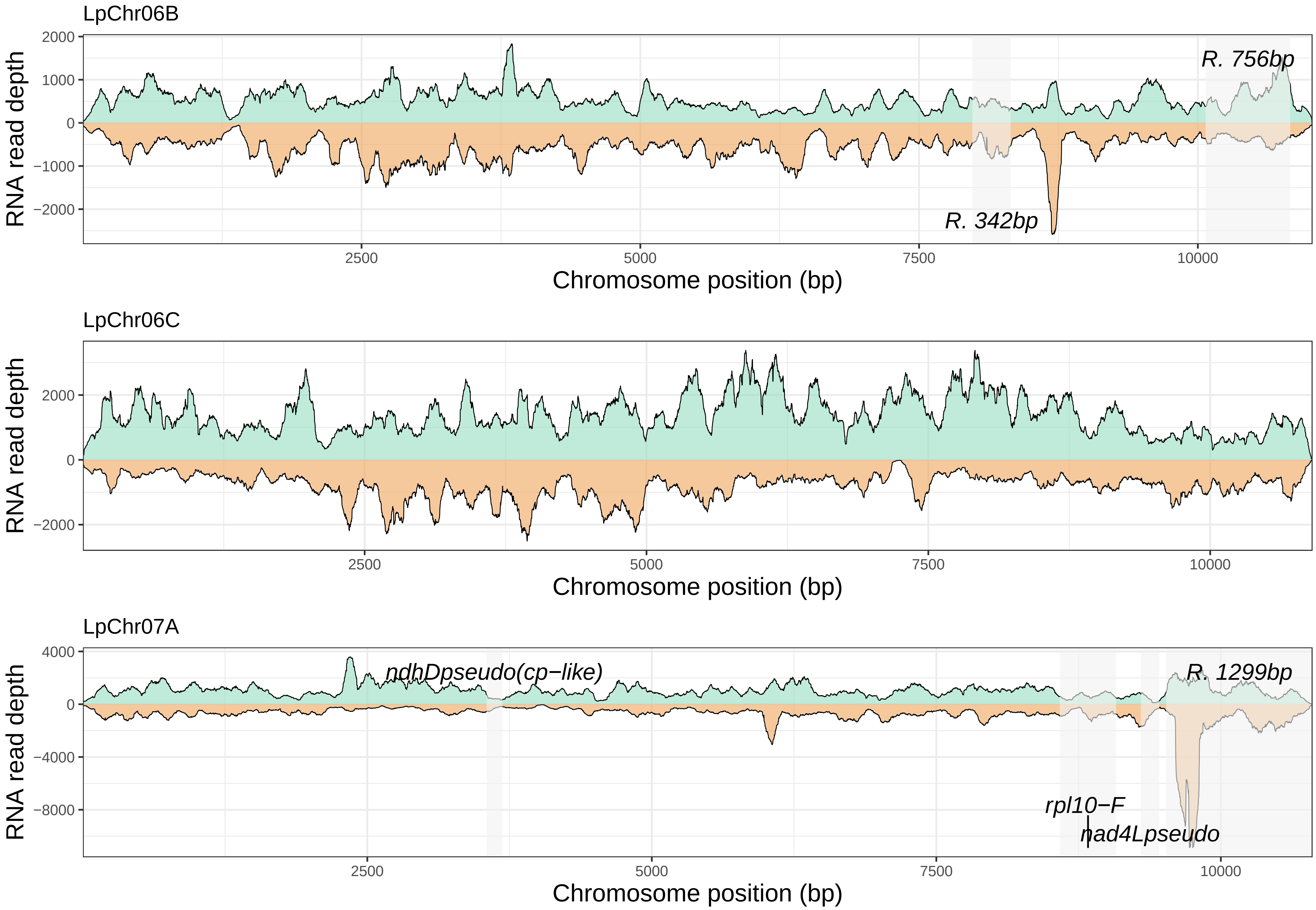

LpChr06B
RNA read depth
2000
1000
0
−1000
−2000
2500
5000
7500
10000
Chromosome position (bp)
R. 756bp
R. 342bp
LpChr06C
RNA read depth
2000
0
−2000
2500
5000
7500
10000
Chromosome position (bp)
LpChr07A
RNA read depth
4000
0
−4000
−8000
2500
5000
7500
10000
Chromosome position (bp)
ndhDpseudo(cp−like)
R. 1299bp
rpl10−F
nad4Lpseudo

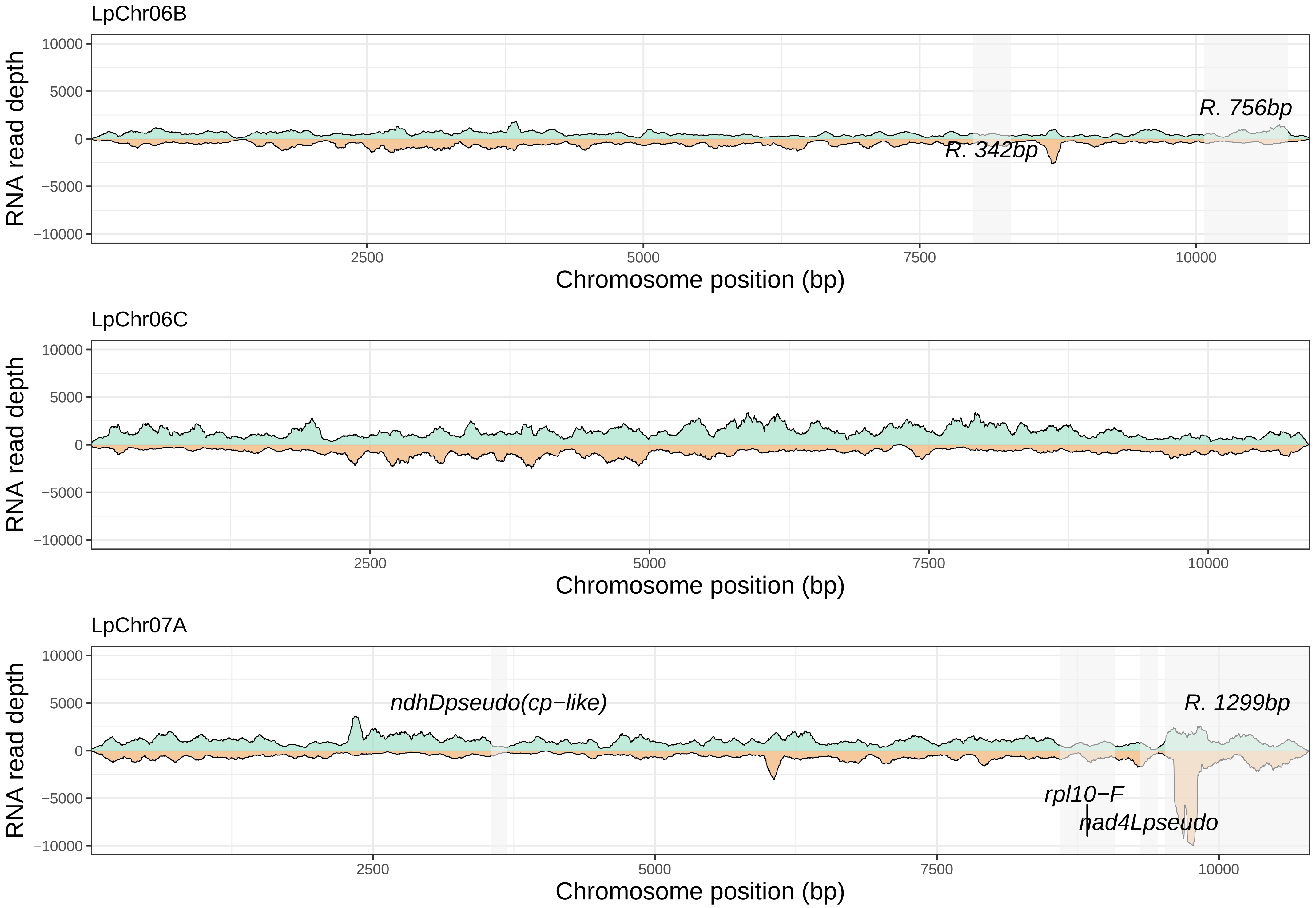
LpChr06B
RNA read depth
10000
5000
0
−5000
−10000
2500
5000
7500
10000
Chromosome position (bp)
R. 756bp
R. 342bp
LpChr06C
RNA read depth
Chromosome position (bp)
LpChr07A
RNA read depth
ndhDpseudo(cp−like)
R. 1299bp
rpl10−F
nad4Lpseudo
Chromosome position (bp)

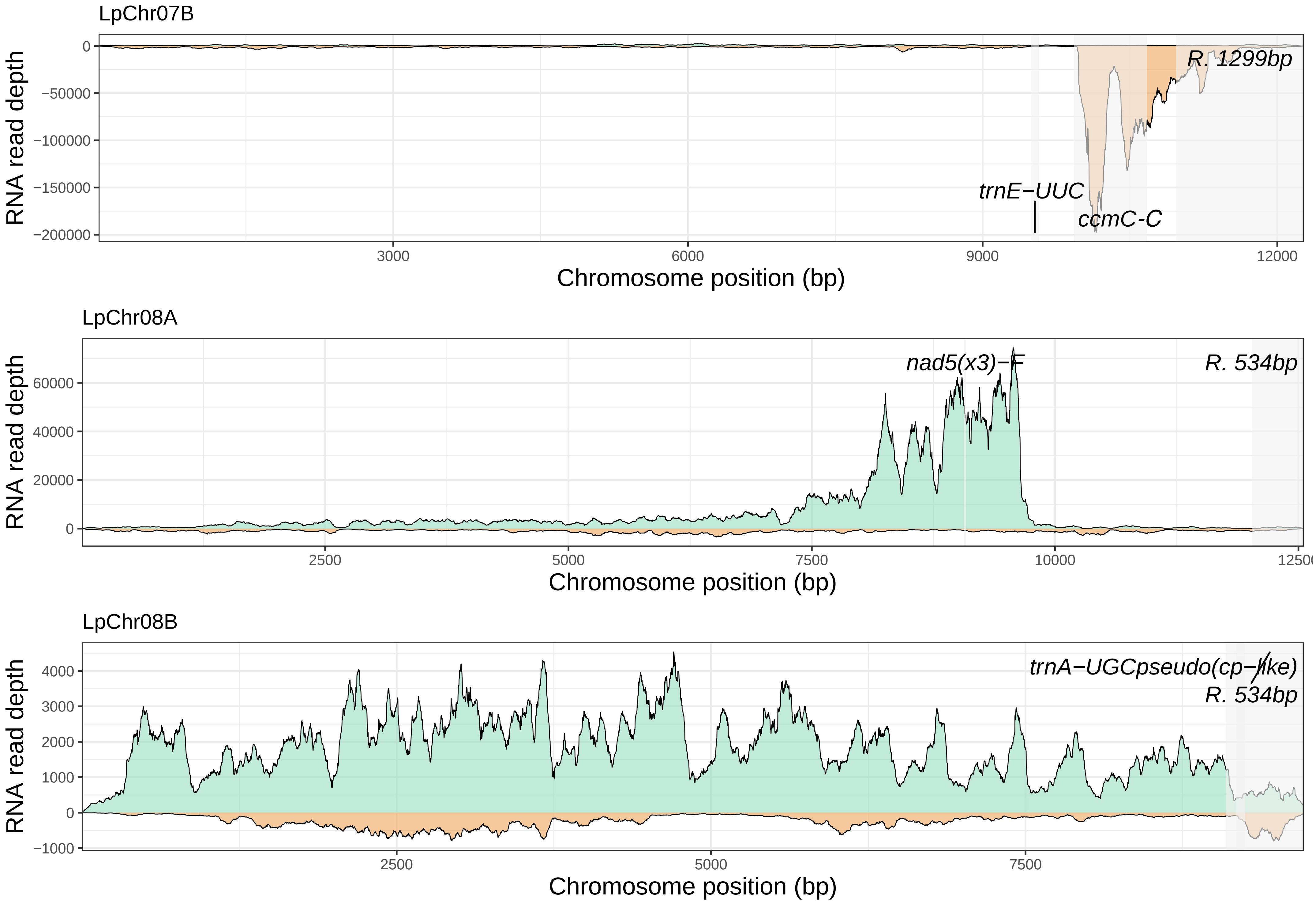

LpChr07B
RNA read depth
0
−50000
−100000
−150000
−200000
R. 1299bp
trnE−UUC
ccmC-C
3000
6000
9000
12000
Chromosome position (bp)
LpChr08A
RNA read depth
60000
40000
20000
0
nad5(x3)−F
R. 534bp
2500
5000
7500
10000
Chromosome position (bp)
LpChr08B
RNA read depth
4000
3000
2000
1000
0
−1000
trnA−UGCpseudo(cp−like)
R. 534bp
2500
5000
7500
Chromosome position (bp)

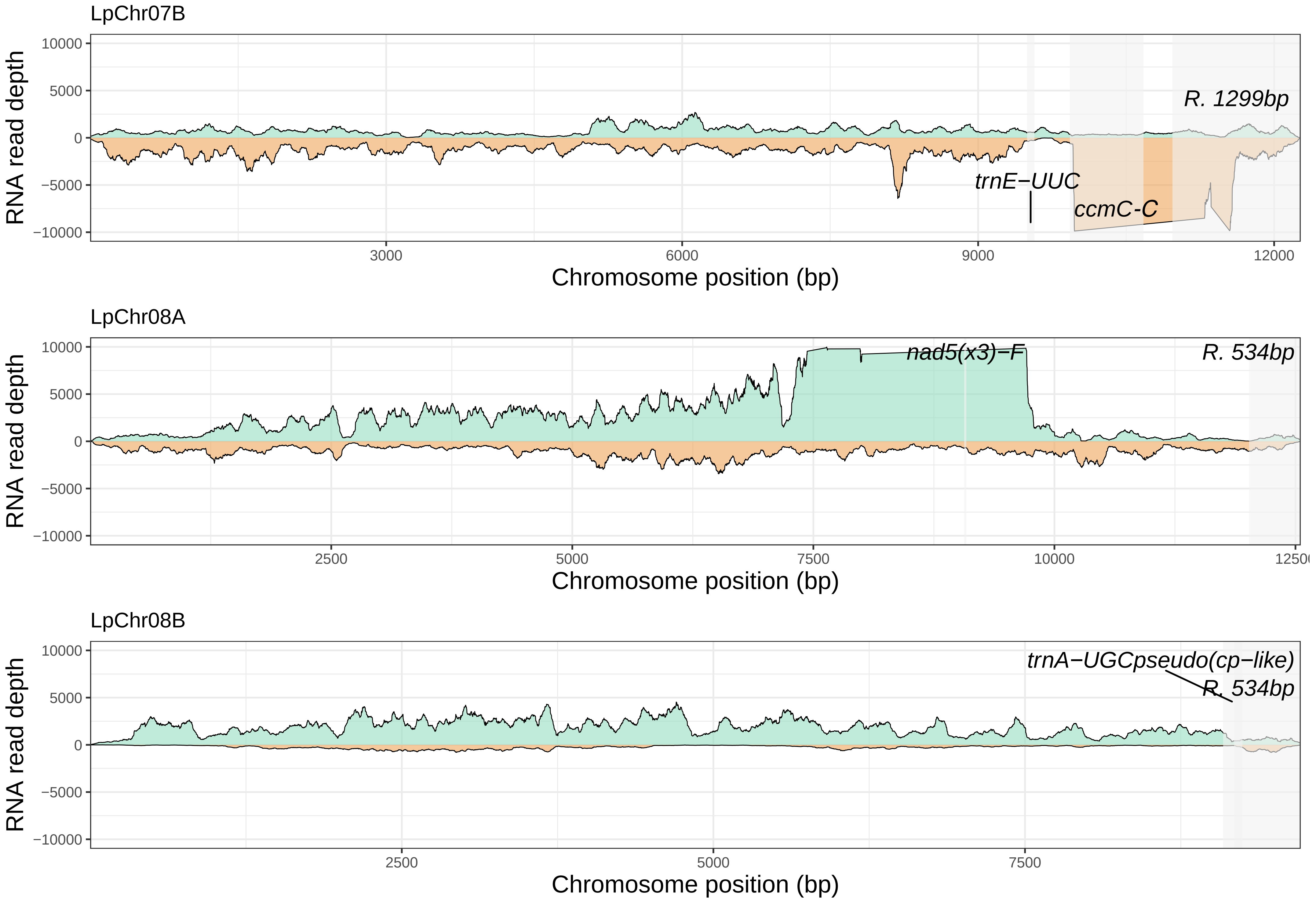
LpChr07B
RNA read depth
10000
5000
0
−5000
−10000
3000
6000
9000
12000
Chromosome position (bp)
R. 1299bp
trnE−UUC
ccmC-C
LpChr08A
RNA read depth
2500
5000
7500
10000
Chromosome position (bp)
nad5(x3)−F
R. 534bp
LpChr08B
RNA read depth
2500
5000
7500
Chromosome position (bp)
trnA−UGCpseudo(cp−like)
R. 534bp

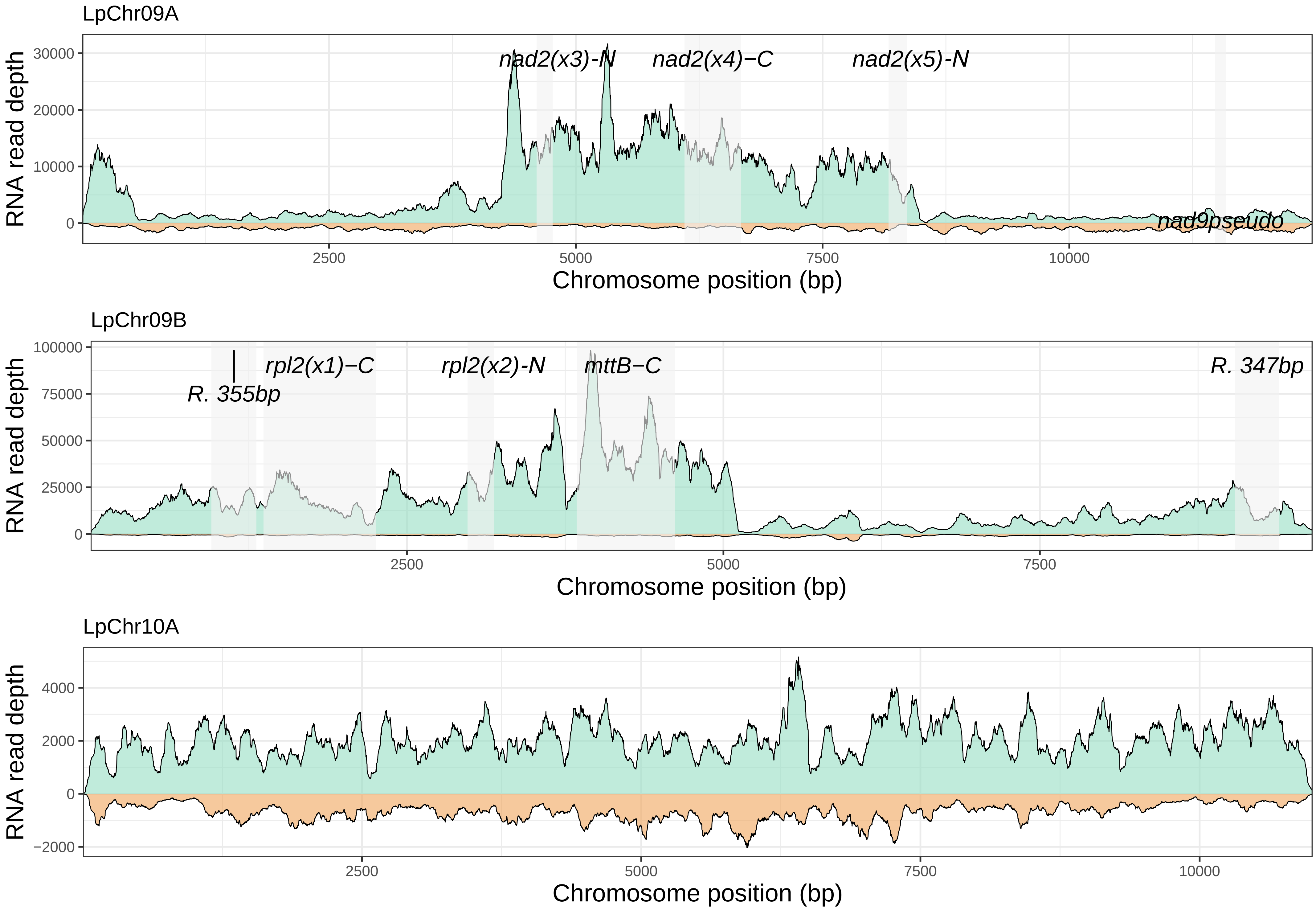

LpChr09A
RNA read depth
30000
20000
10000
0
nad2(x3)-N
nad2(x4)−C
nad2(x5)-N
nad9pseudo
2500
5000
7500
10000
Chromosome position (bp)
LpChr09B
RNA read depth
100000
75000
50000
25000
0
R. 355bp
rpl2(x1)−C
rpl2(x2)-N
mttB−C
R. 347bp
2500
5000
7500
Chromosome position (bp)
LpChr10A
RNA read depth
4000
2000
0
−2000
2500
5000
7500
10000
Chromosome position (bp)

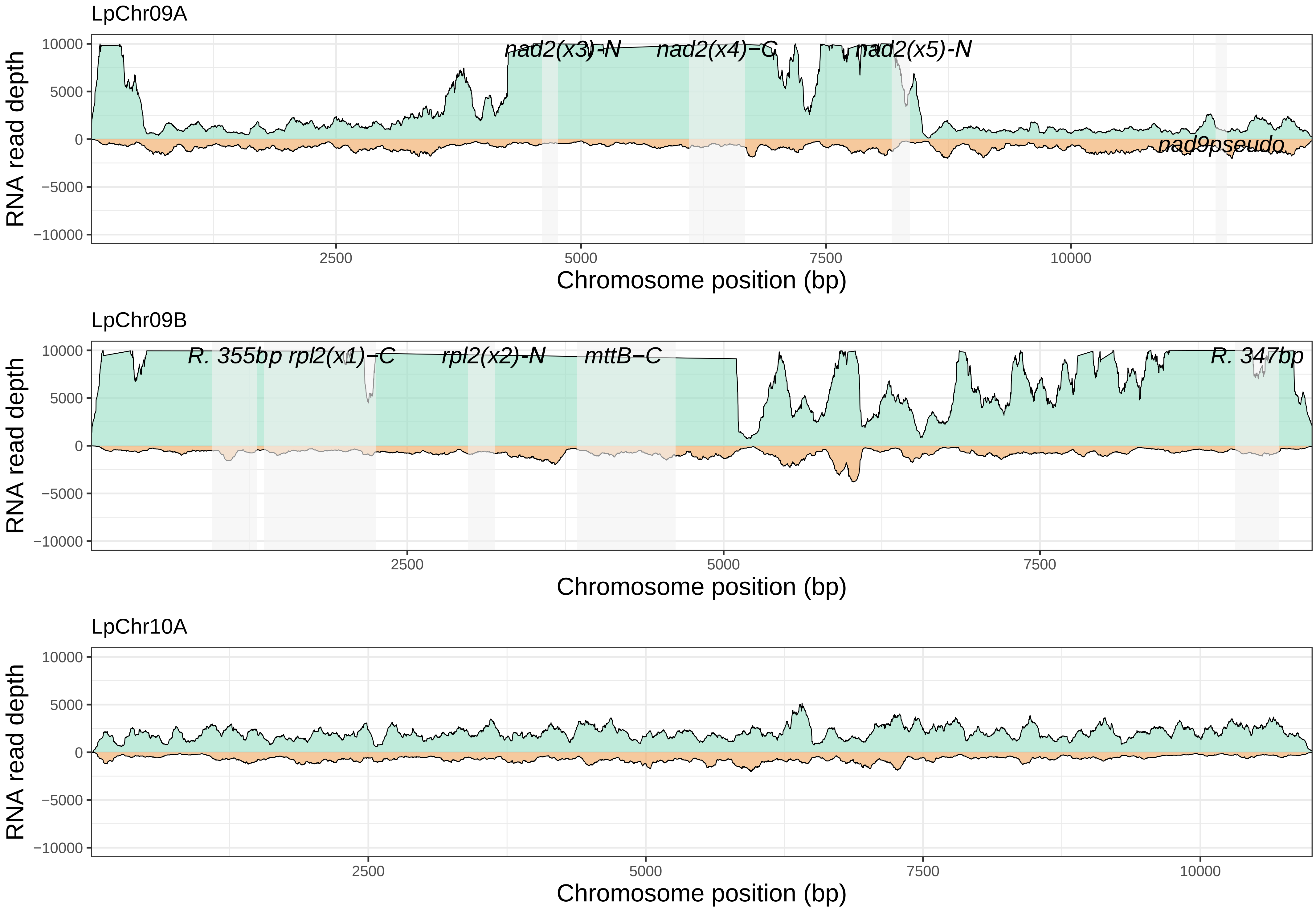
LpChr09A
nad2(x3)-N
nad2(x4)-C
nad2(x5)-N
nad9pseudo
RNA read depth
10000
5000
0
−5000
−10000
2500
5000
7500
10000
Chromosome position (bp)
LpChr09B
R. 355bp
rpl2(x1)−C
rpl2(x2)-N
mttB−C
R. 347bp
RNA read depth
2500
5000
7500
Chromosome position (bp)
LpChr10A
RNA read depth
2500
5000
7500
10000
Chromosome position (bp)

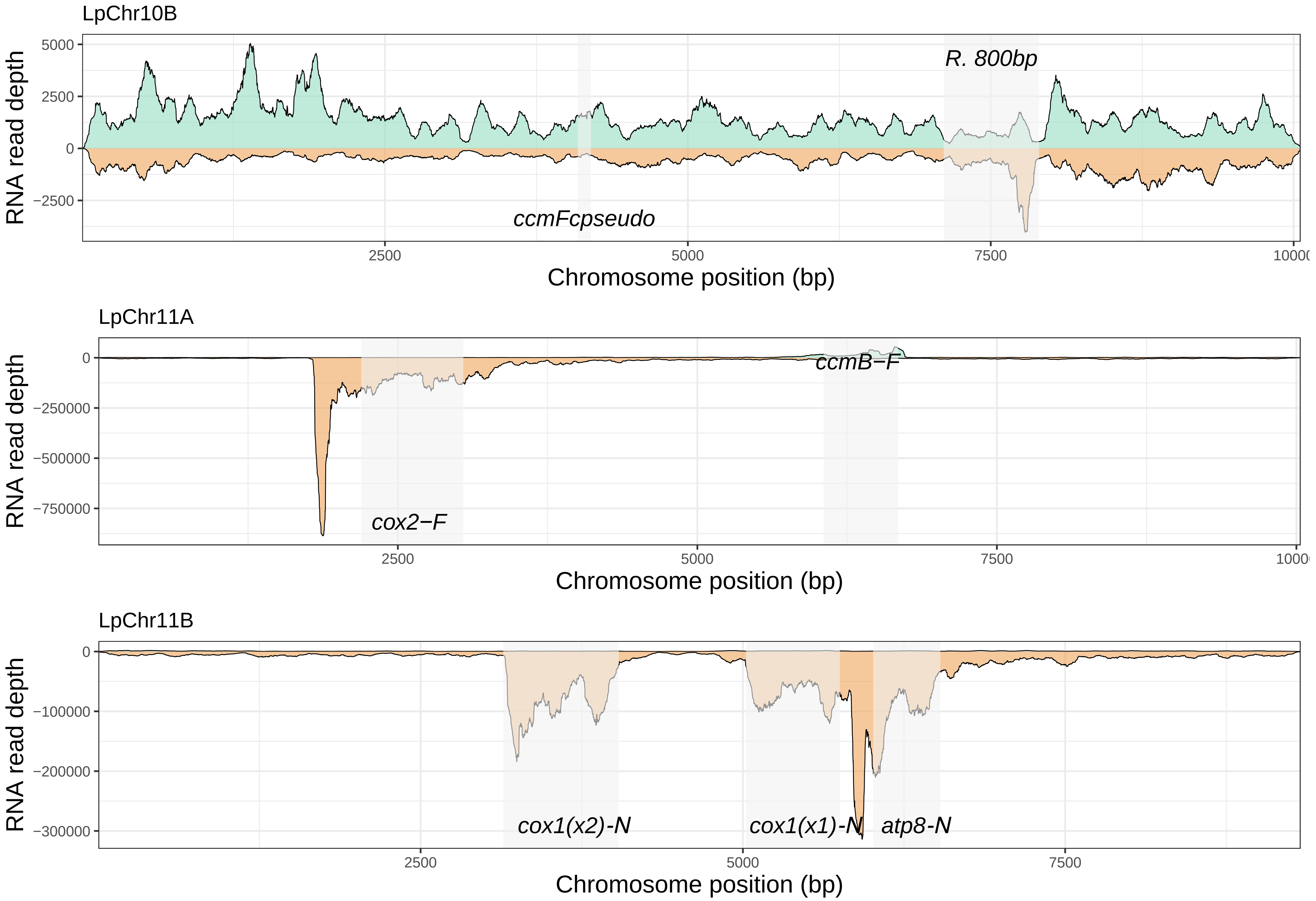

LpChr10B
RNA read depth
5000
2500
0
−2500
R. 800bp
ccmFcpseudo
2500
5000
7500
10000
Chromosome position (bp)
LpChr11A
RNA read depth
0
−250000
−500000
−750000
ccmB−F
cox2−F
2500
5000
7500
10000
Chromosome position (bp)
LpChr11B
RNA read depth
0
−100000
−200000
−300000
cox1(x2)-N
cox1(x1)-N
atp8-N
2500
5000
7500
Chromosome position (bp)

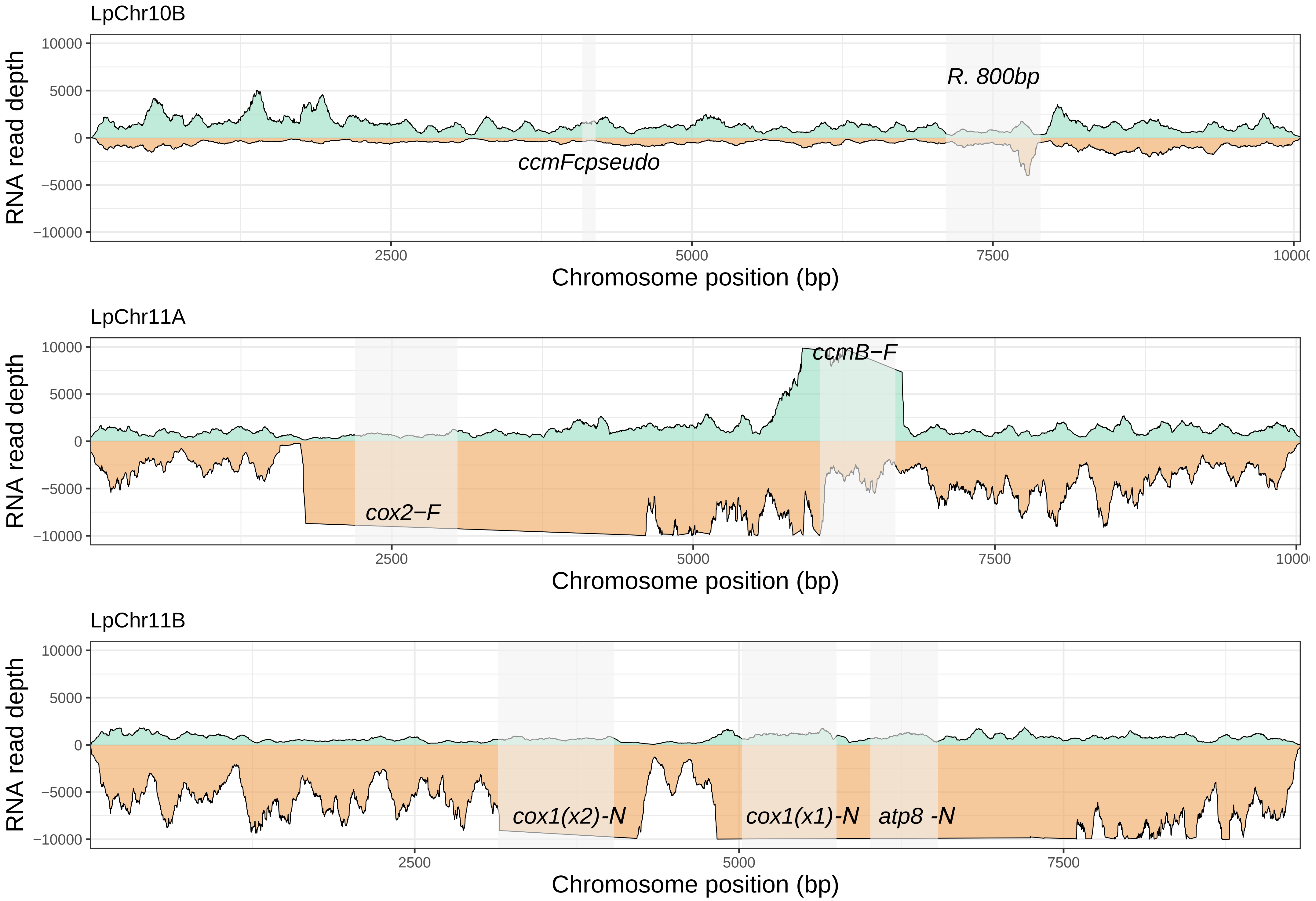

LpChr10B
RNA read depth
10000
5000
0
−5000
−10000
R. 800bp
ccmFcpseudo
2500
5000
7500
10000
Chromosome position (bp)
LpChr11A
RNA read depth
ccmB−F
cox2−F
Chromosome position (bp)
LpChr11B
RNA read depth
cox1(x2)-N
cox1(x1)-N
atp8 -N
Chromosome position (bp)

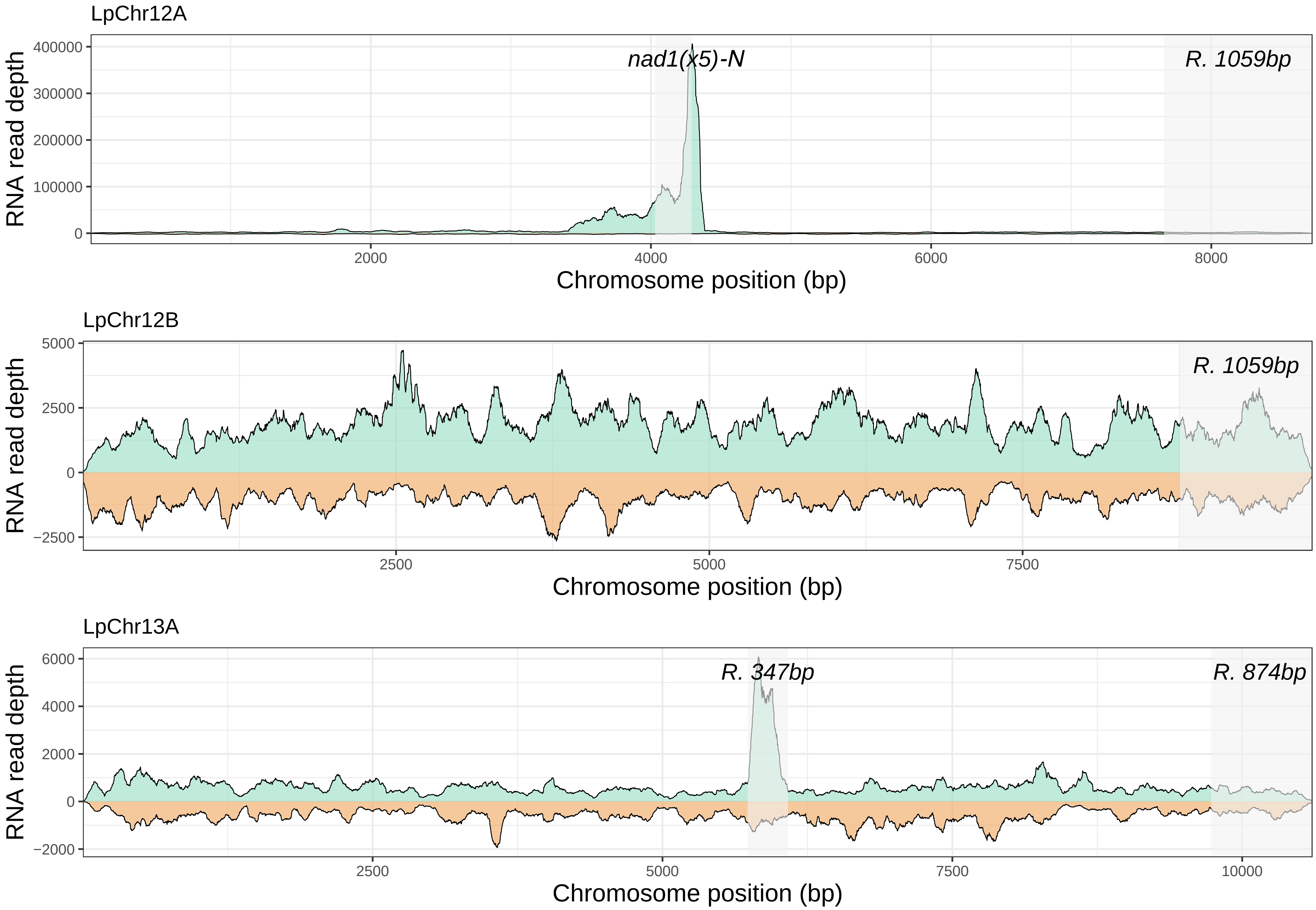

LpChr12A
nad1(x5)-N
R. 1059bp
RNA read depth
400000
300000
200000
100000
0
2000
4000
6000
8000
Chromosome position (bp)
LpChr12B
R. 1059bp
RNA read depth
5000
2500
0
−2500
2500
5000
7500
Chromosome position (bp)
LpChr13A
R. 347bp
R. 874bp
RNA read depth
6000
4000
2000
0
−2000
2500
5000
7500
10000
Chromosome position (bp)

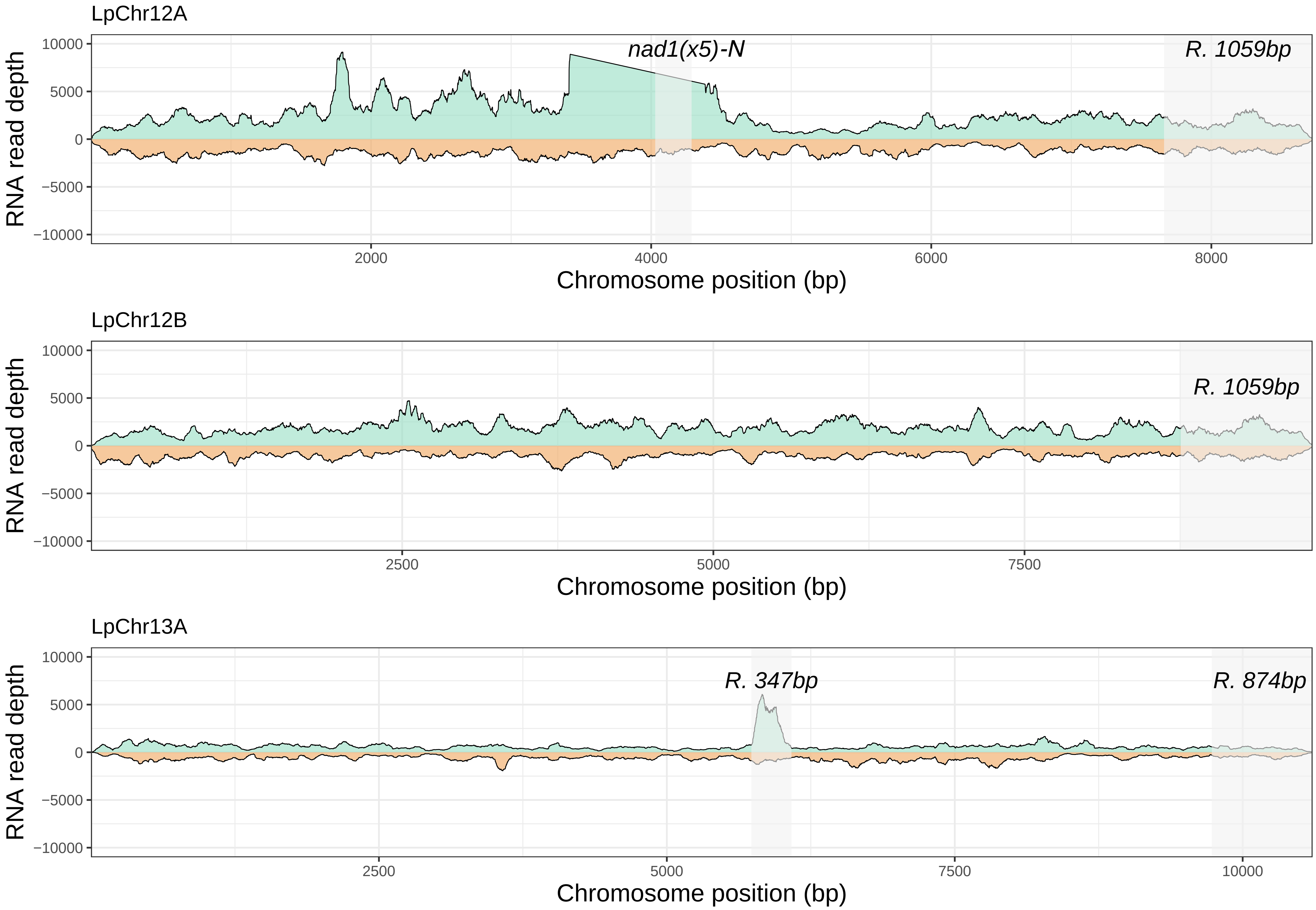
LpChr12A
nad1(x5)-N
R. 1059bp
RNA read depth
Chromosome position (bp)
LpChr12B
R. 1059bp
RNA read depth
Chromosome position (bp)
LpChr13A
R. 347bp
R. 874bp
RNA read depth
Chromosome position (bp)

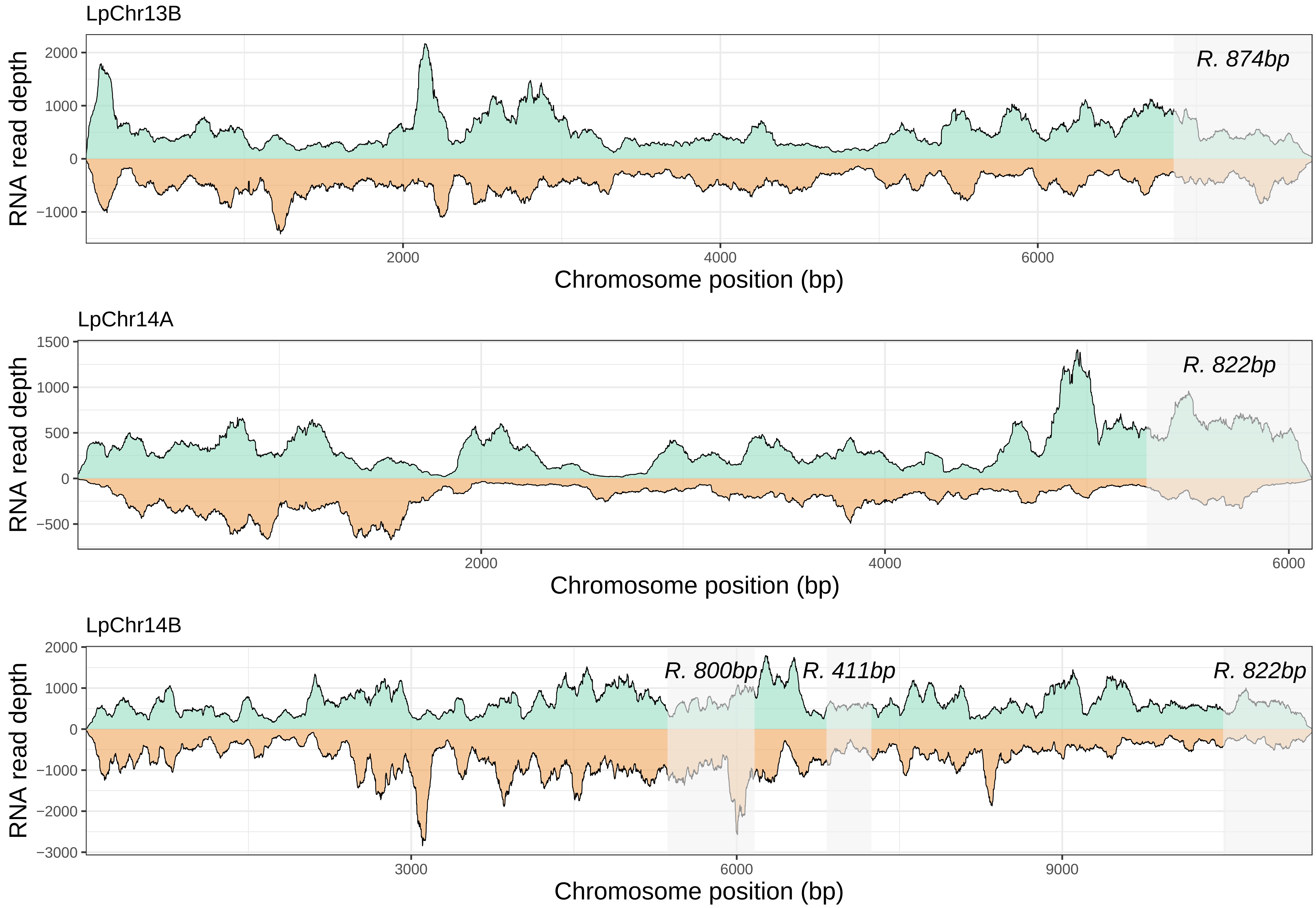

LpChr13B
RNA read depth
2000
1000
0
−1000
R. 874bp
2000
4000
6000
Chromosome position (bp)
LpChr14A
RNA read depth
1500
1000
500
0
−500
R. 822bp
2000
4000
6000
Chromosome position (bp)
LpChr14B
RNA read depth
2000
1000
0
−1000
−2000
−3000
R. 800bp
R. 411bp
R. 822bp
3000
6000
9000
Chromosome position (bp)

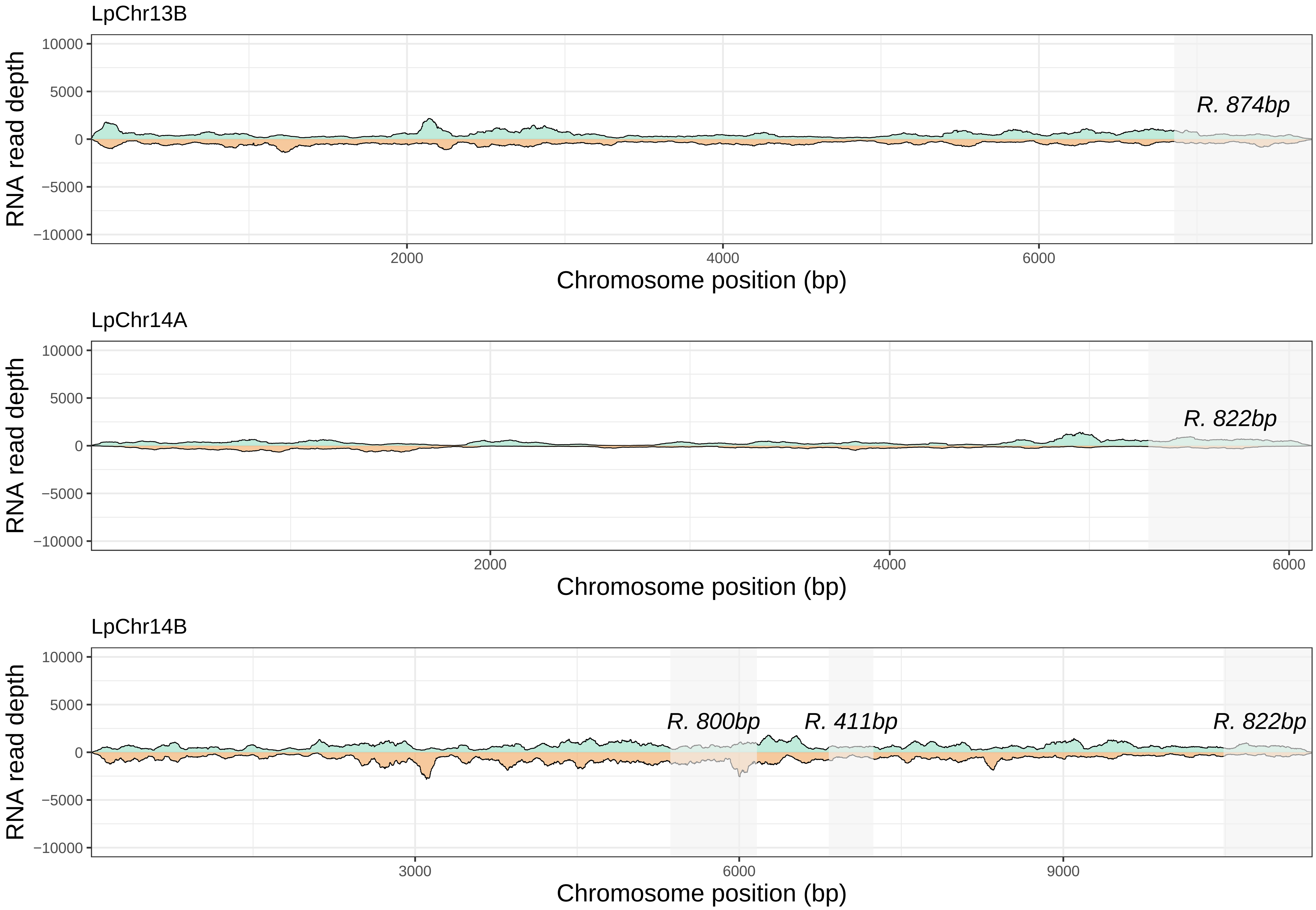
LpChr13B
RNA read depth
10000
5000
0
−5000
−10000
R. 874bp
2000
4000
6000
Chromosome position (bp)
LpChr14A
RNA read depth
10000
5000
0
−5000
−10000
R. 822bp
2000
4000
6000
Chromosome position (bp)
LpChr14B
RNA read depth
10000
5000
0
−5000
−10000
R. 800bp
R. 411bp
R. 822bp
3000
6000
9000
Chromosome position (bp)

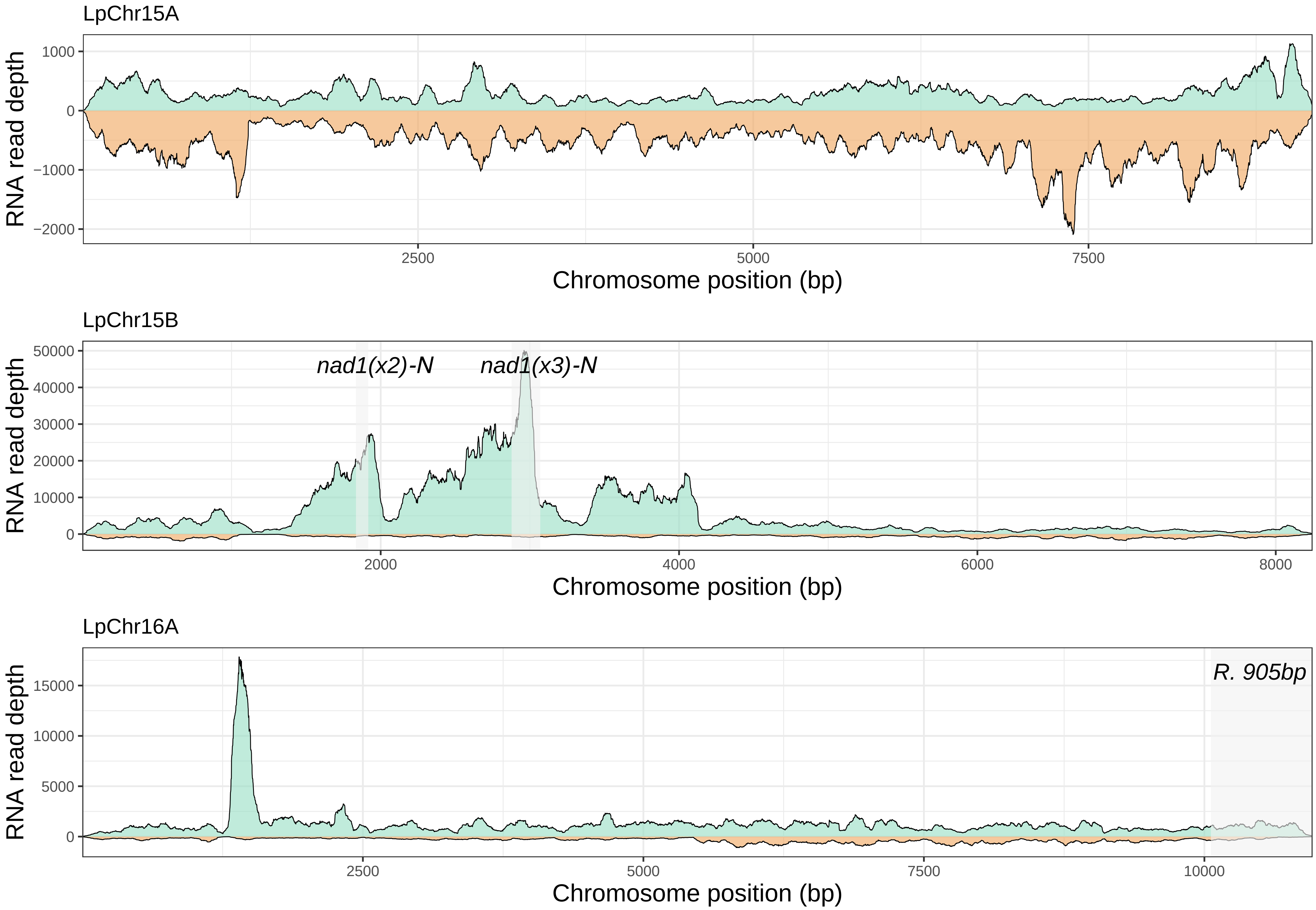

LpChr15A
RNA read depth
1000
0
−1000
−2000
2500
5000
7500
Chromosome position (bp)
LpChr15B
RNA read depth
50000
40000
30000
20000
10000
0
nad1(x2)-N
nad1(x3)-N
2000
4000
6000
8000
Chromosome position (bp)
LpChr16A
RNA read depth
15000
10000
5000
0
R. 905bp
2500
5000
7500
10000
Chromosome position (bp)

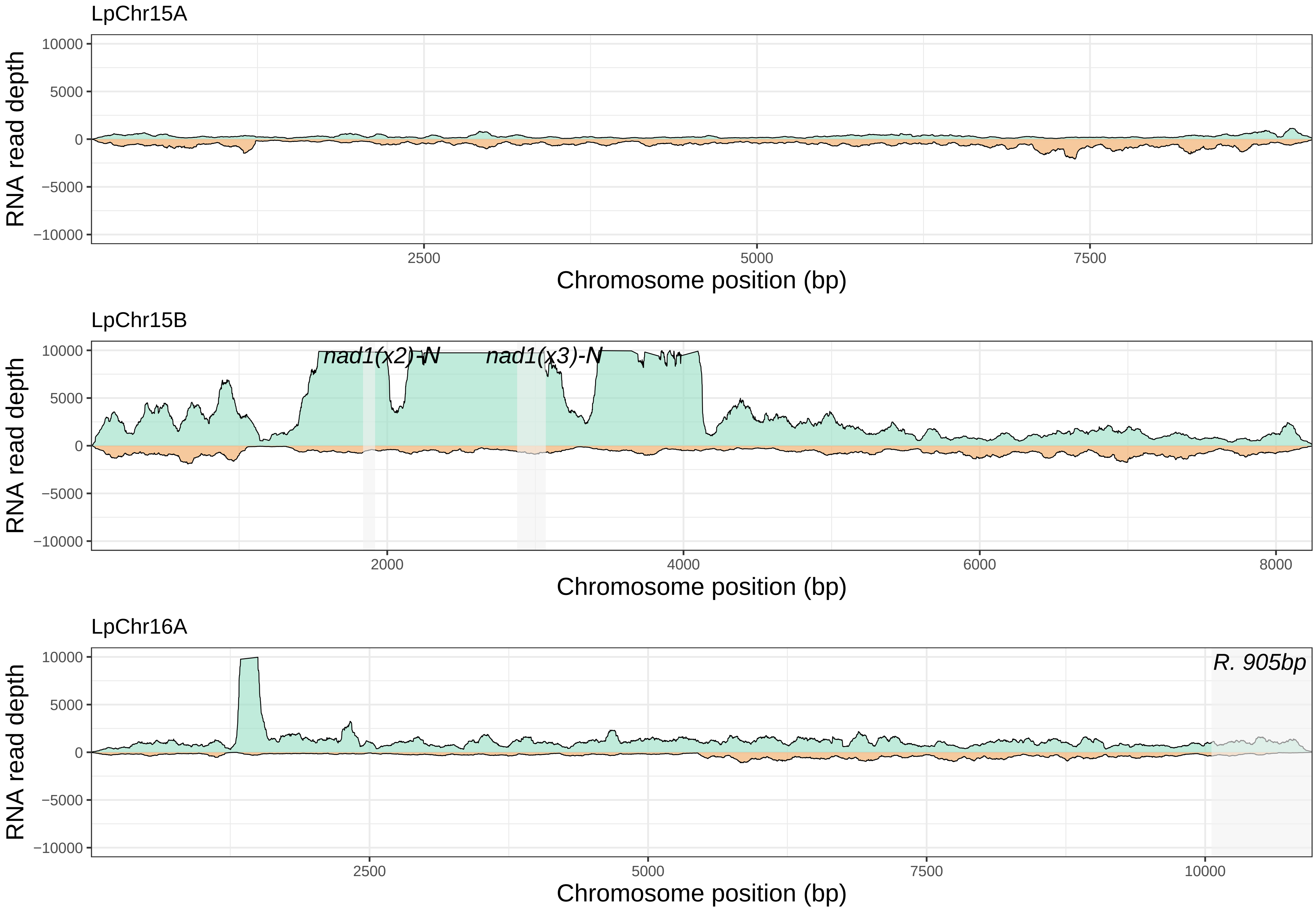
LpChr15A
RNA read depth
10000
5000
0
−5000
−10000
2500
5000
7500
Chromosome position (bp)
LpChr15B
nad1(x2)-N
nad1(x3)-N
2000
4000
6000
8000
LpChr16A
R. 905bp
10000

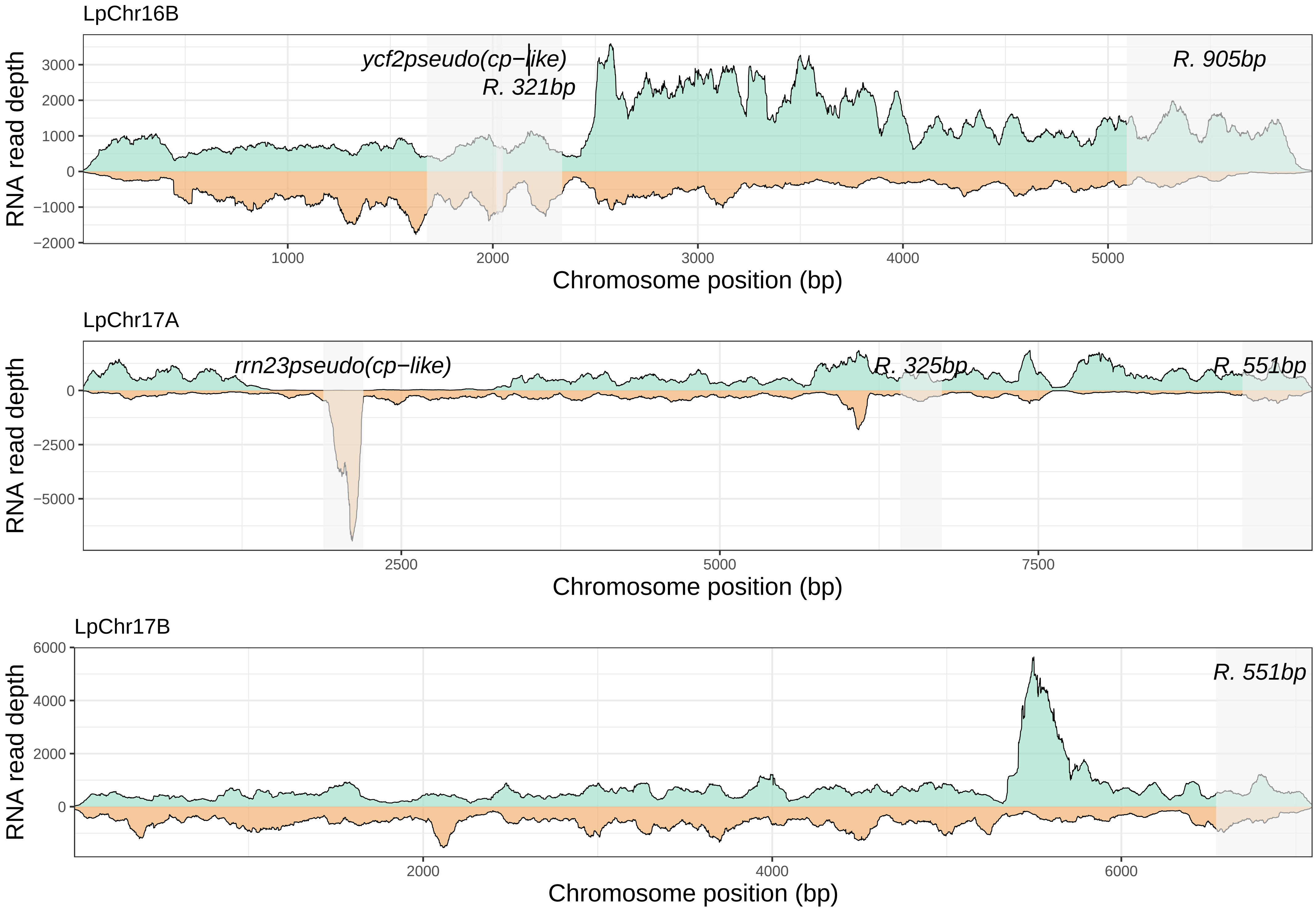

LpChr16B
ycf2pseudo(cp−like)
R. 321bp
R. 905bp
RNA read depth
3000
2000
1000
0
−1000
−2000
1000
2000
3000
4000
5000
Chromosome position (bp)
LpChr17A
rrn23pseudo(cp−like)
R. 325bp
R. 551bp
RNA read depth
0
−2500
−5000
2500
5000
7500
Chromosome position (bp)
LpChr17B
R. 551bp
RNA read depth
6000
4000
2000
0
2000
4000
6000
Chromosome position (bp)

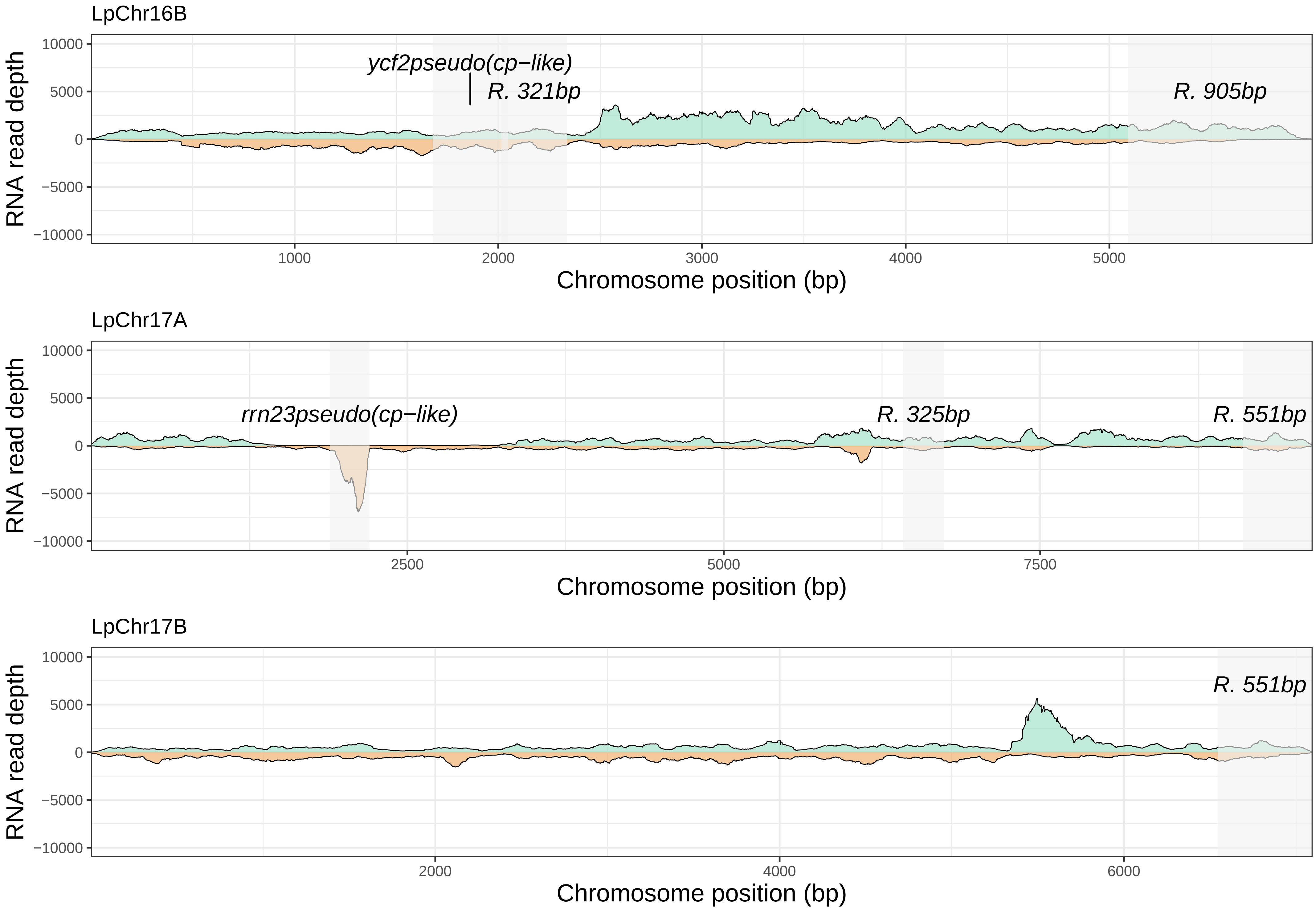
LpChr16B
ycf2pseudo(cp−like)
R. 321bp
R. 905bp
RNA read depth
10000
5000
0
−5000
−10000
1000
2000
3000
4000
5000
Chromosome position (bp)
LpChr17A
rrn23pseudo(cp−like)
R. 325bp
R. 551bp
RNA read depth
2500
5000
7500
Chromosome position (bp)
LpChr17B
R. 551bp
RNA read depth
2000
4000
6000
Chromosome position (bp)

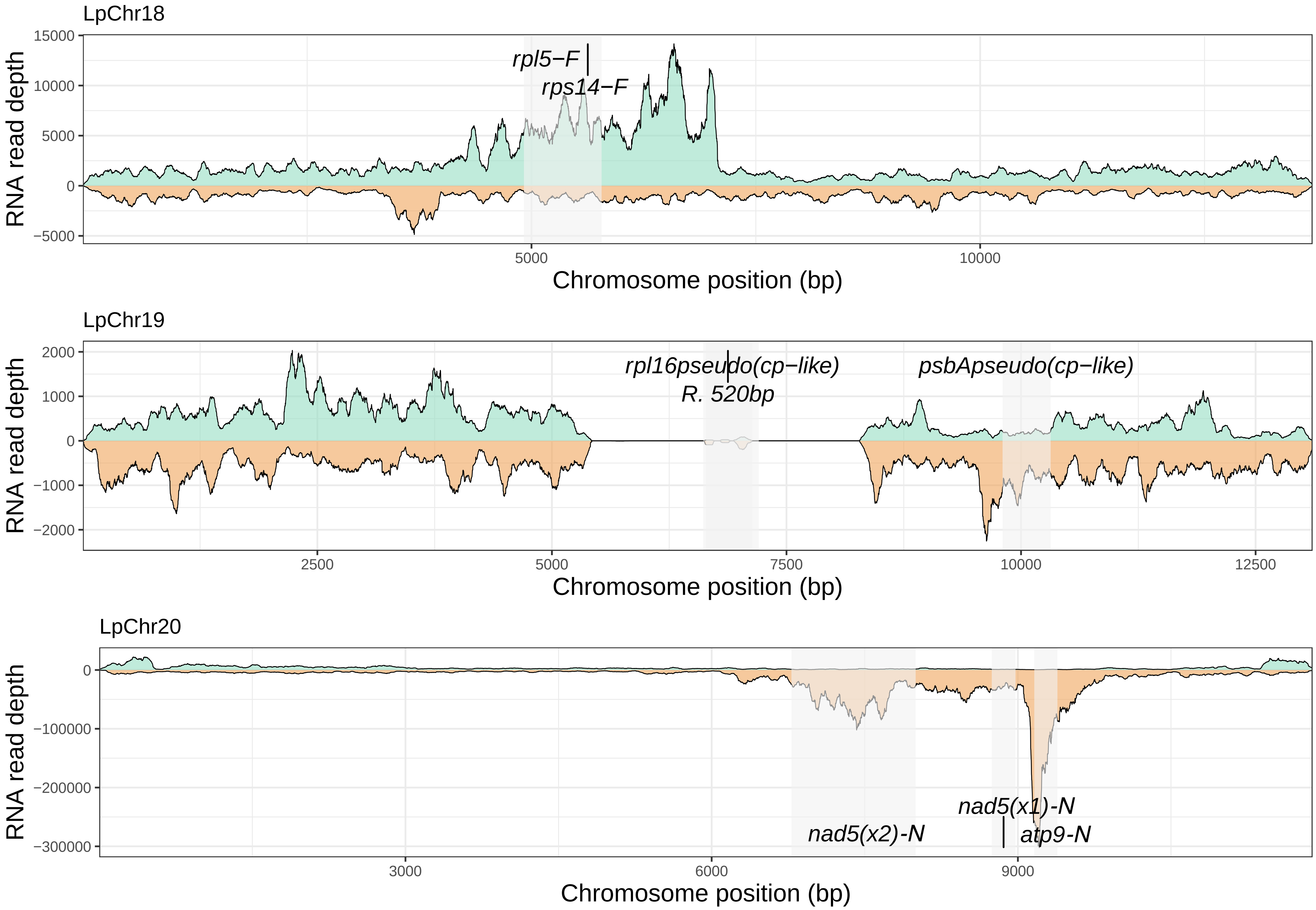

LpChr18
RNA read depth
15000
10000
5000
0
−5000
rpl5−F
rps14−F
5000
10000
Chromosome position (bp)
LpChr19
RNA read depth
2000
1000
0
−1000
−2000
rpl16pseudo(cp−like)
R. 520bp
psbApseudo(cp−like)
2500
5000
7500
10000
12500
Chromosome position (bp)
LpChr20
RNA read depth
0
−100000
−200000
−300000
nad5(x2)-N
nad5(x1)-N
atp9-N
3000
6000
9000
Chromosome position (bp)

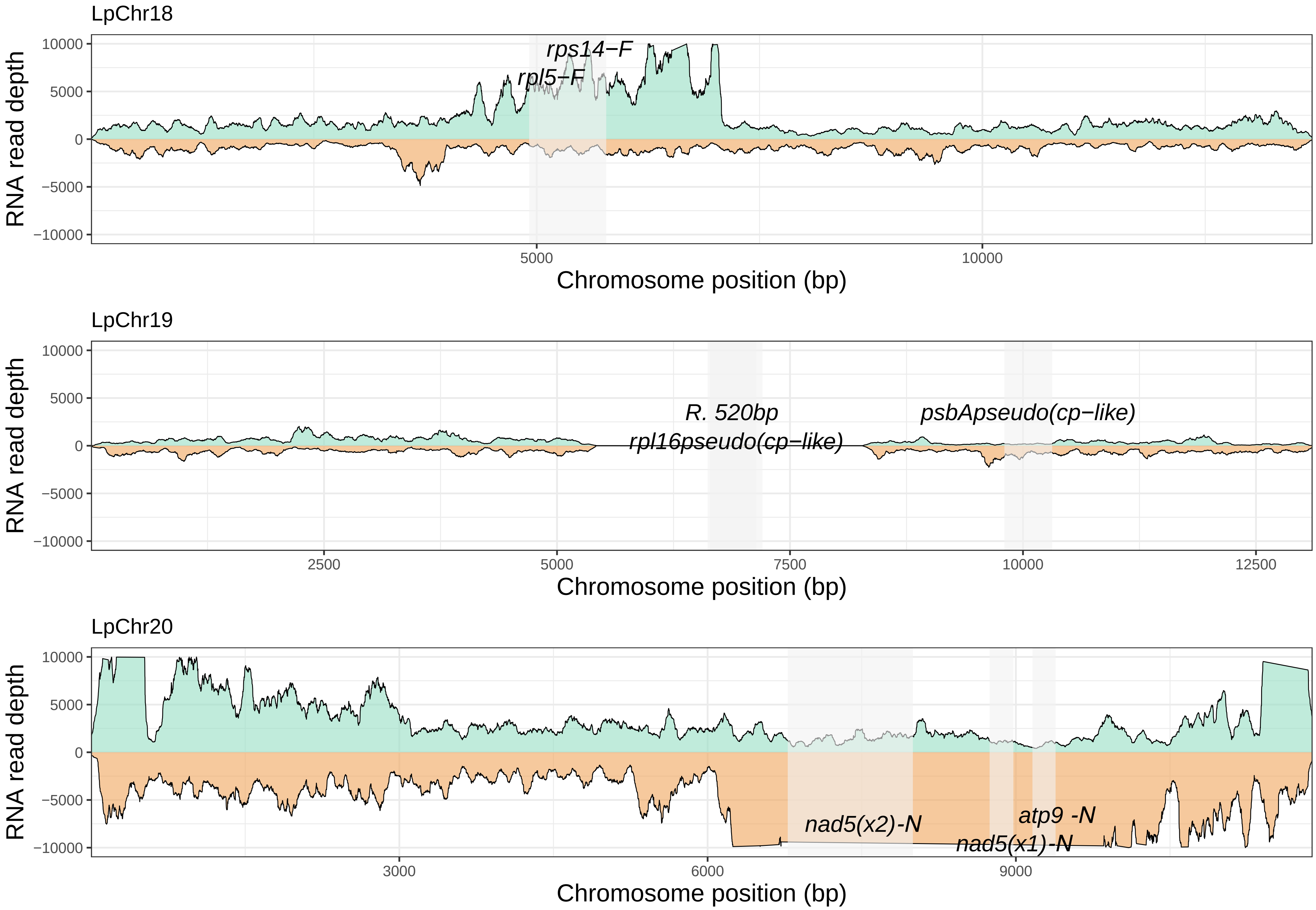
LpChr18
RNA read depth
10000
5000
0
−5000
−10000
rps14−F
rpl5−F
5000
10000
Chromosome position (bp)
LpChr19
RNA read depth
10000
5000
0
−5000
−10000
R. 520bp
rpl16pseudo(cp−like)
psbApseudo(cp−like)
2500
5000
7500
10000
12500
Chromosome position (bp)
LpChr20
RNA read depth
10000
5000
0
−5000
−10000
nad5(x2)-N
atp9 -N
nad5(x1)-N
3000
6000
9000
Chromosome position (bp)

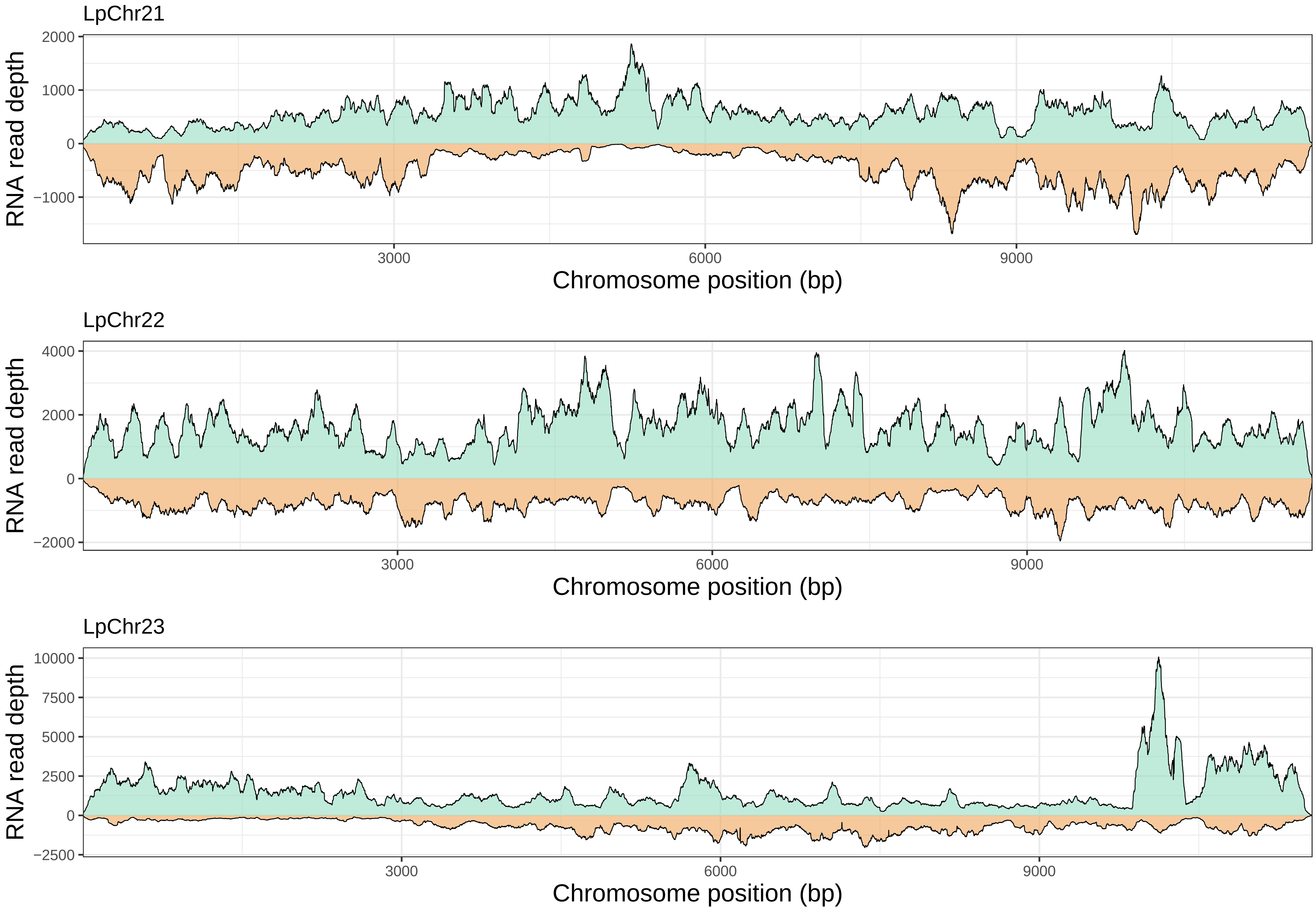

LpChr21
2000
1000
0
−1000
RNA read depth
3000
6000
9000
Chromosome position (bp)
LpChr22
4000
2000
0
−2000
RNA read depth
3000
6000
9000
Chromosome position (bp)
LpChr23
10000
7500
5000
2500
0
−2500
RNA read depth
3000
6000
9000
Chromosome position (bp)

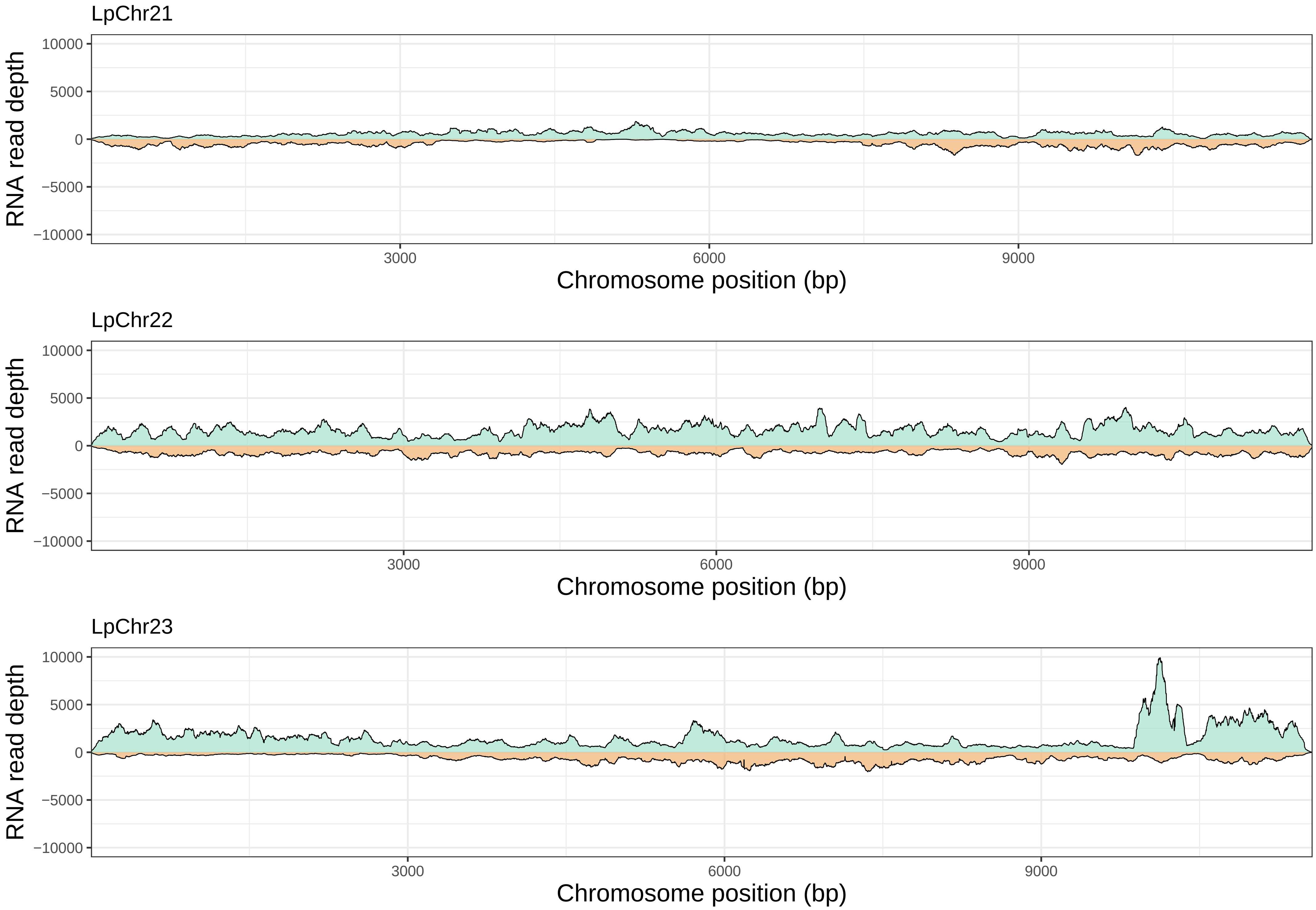
LpChr21
RNA read depth
10000
5000
0
−5000
−10000
3000
6000
9000
Chromosome position (bp)
LpChr22
RNA read depth
10000
5000
0
−5000
−10000
3000
6000
9000
Chromosome position (bp)
LpChr23
RNA read depth
10000
5000
0
−5000
−10000
3000
6000
9000
Chromosome position (bp)

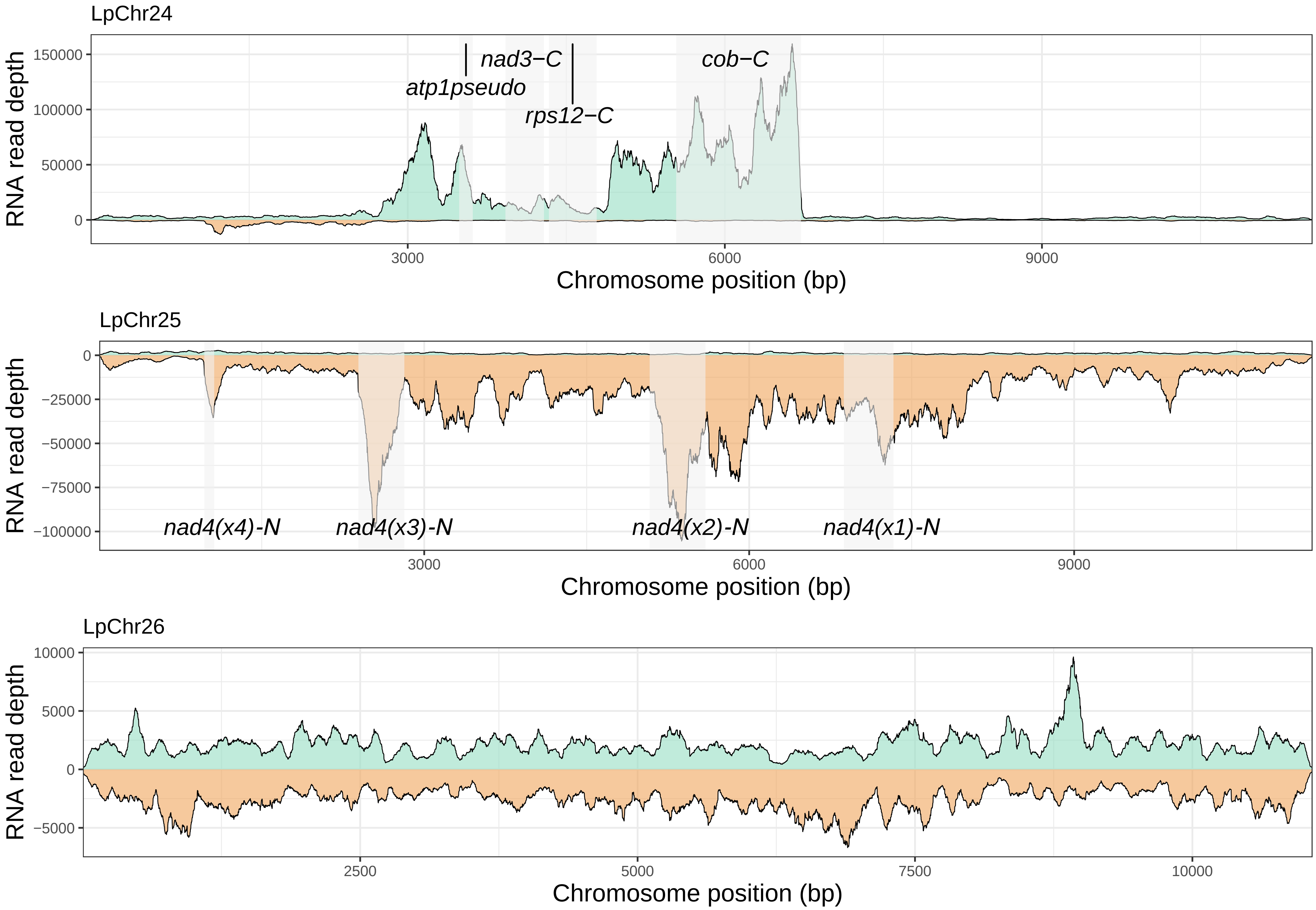

LpChr24
RNA read depth
150000
100000
50000
0
nad3−C
atp1pseudo
rps12−C
cob−C
3000
6000
9000
Chromosome position (bp)
LpChr25
RNA read depth
0
−25000
−50000
−75000
−100000
nad4(x4)-N
nad4(x3)-N
nad4(x2)-N
nad4(x1)-N
3000
6000
9000
Chromosome position (bp)
LpChr26
RNA read depth
10000
5000
0
−5000
2500
5000
7500
10000
Chromosome position (bp)

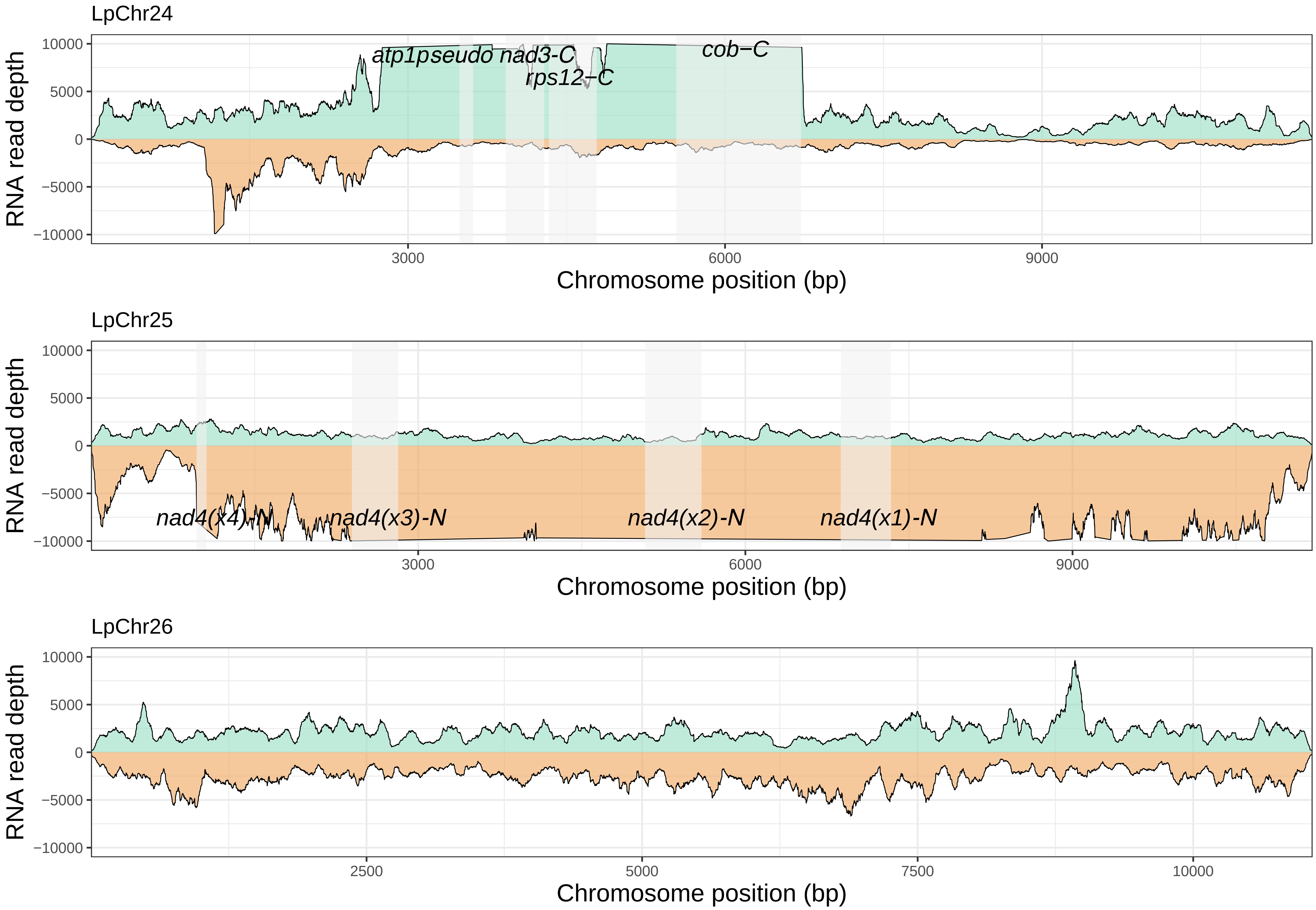

LpChr24
atp1pseudo
nad3-C
rps12–C
cob–C
RNA read depth
10000
5000
0
−5000
−10000
3000
6000
9000
Chromosome position (bp)
LpChr25
nad4(x4)-N
nad4(x3)-N
nad4(x2)-N
nad4(x1)-N
RNA read depth
10000
5000
0
−5000
−10000
3000
6000
9000
Chromosome position (bp)
LpChr26
RNA read depth
10000
5000
0
−5000
−10000
2500
5000
7500
10000
Chromosome position (bp)

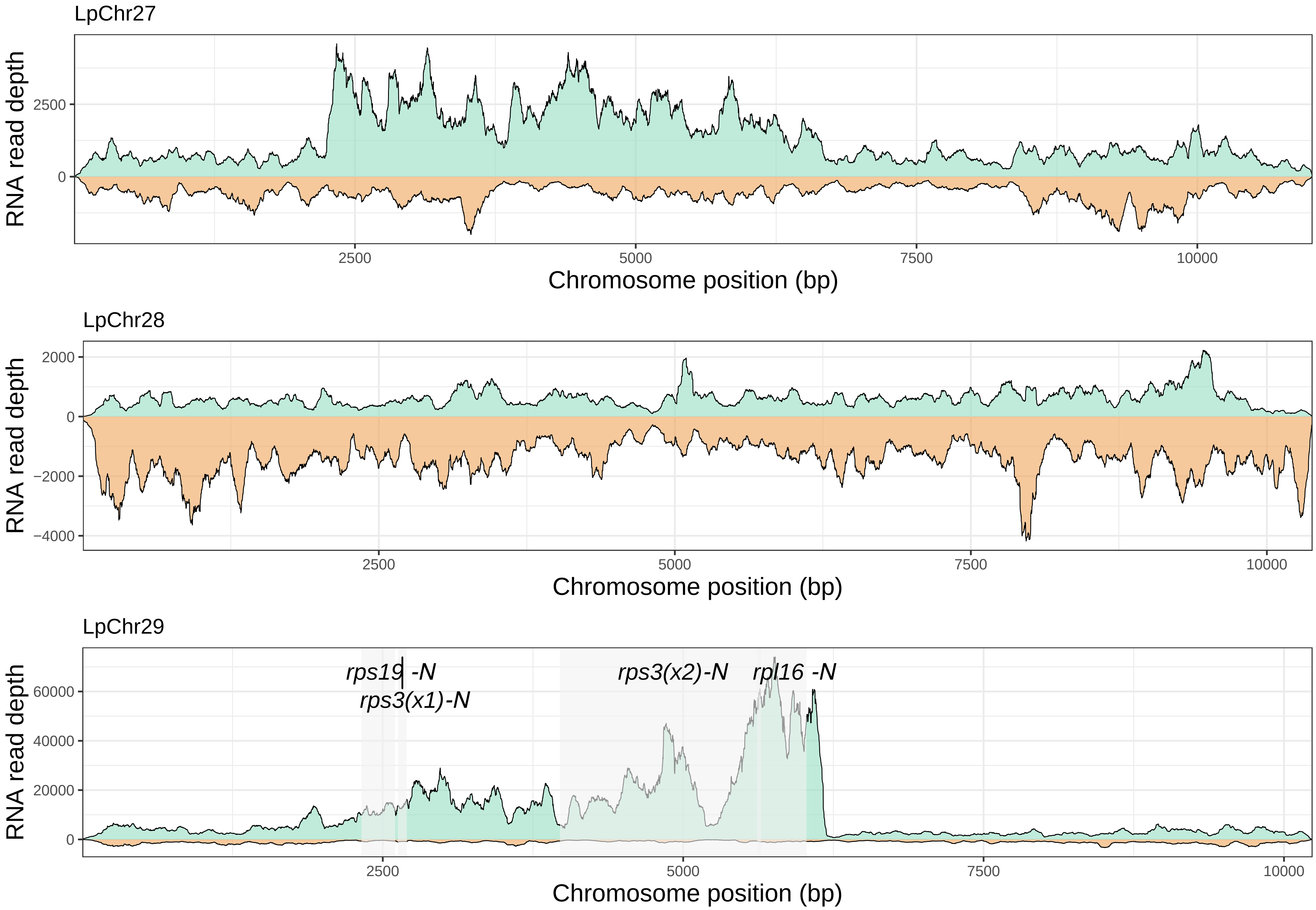
LpChr27
RNA read depth
2500
0
2500
5000
7500
10000
Chromosome position (bp)
LpChr28
RNA read depth
2000
0
−2000
−4000
2500
5000
7500
10000
Chromosome position (bp)
LpChr29
RNA read depth
60000
40000
20000
0
rps19 -N
rps3(x1)-N
rps3(x2)-N
rpl16 -N
2500
5000
7500
10000
Chromosome position (bp)

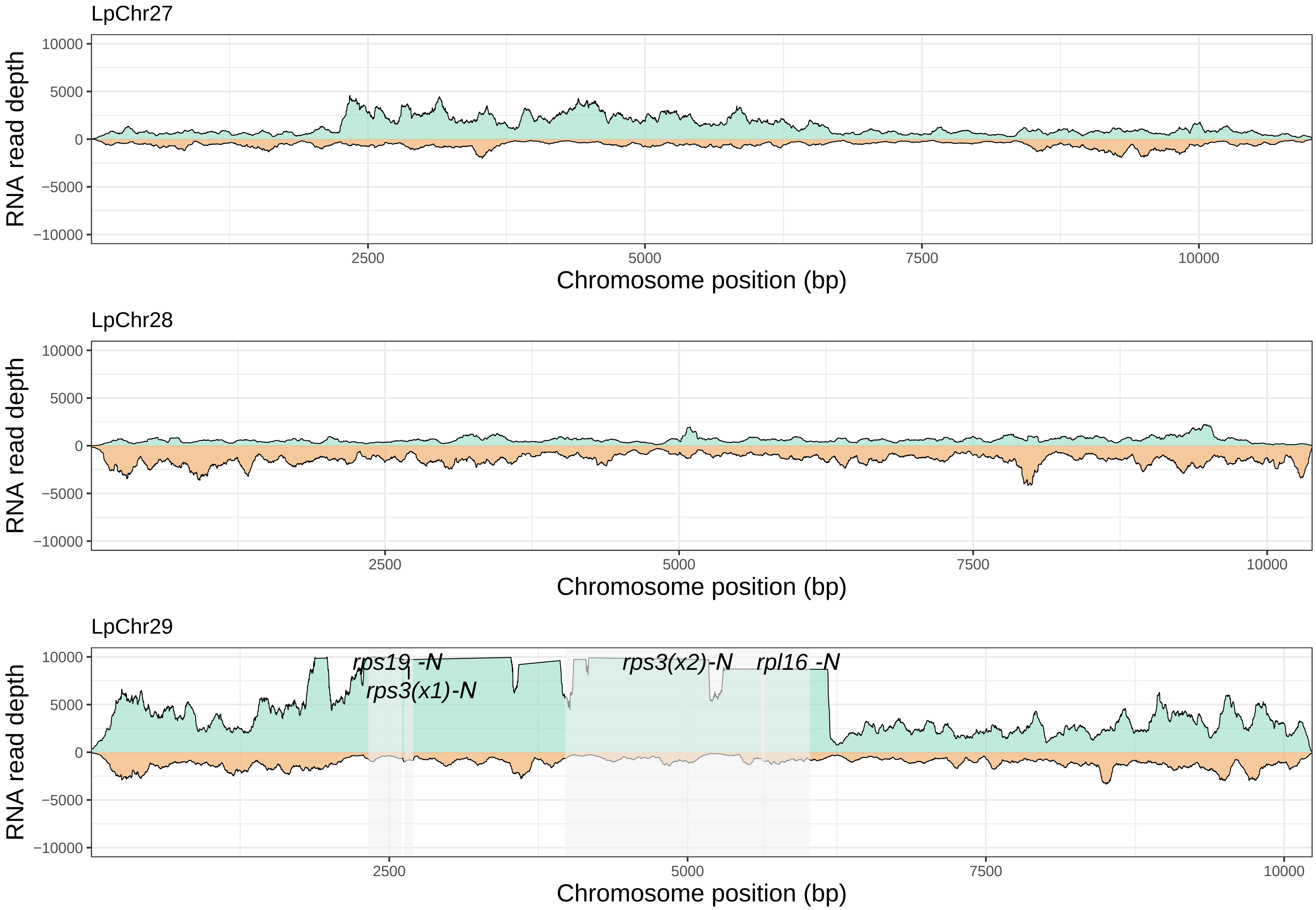
LpChr27
RNA read depth
10000
5000
0
−5000
−10000
2500
5000
7500
10000
Chromosome position (bp)
LpChr28
RNA read depth
10000
5000
0
−5000
−10000
2500
5000
7500
10000
Chromosome position (bp)
LpChr29
rps19 -N
rps3(x1)-N
rps3(x2)-N
rpl16 -N
RNA read depth
10000
5000
0
−5000
−10000
2500
5000
7500
10000
Chromosome position (bp)

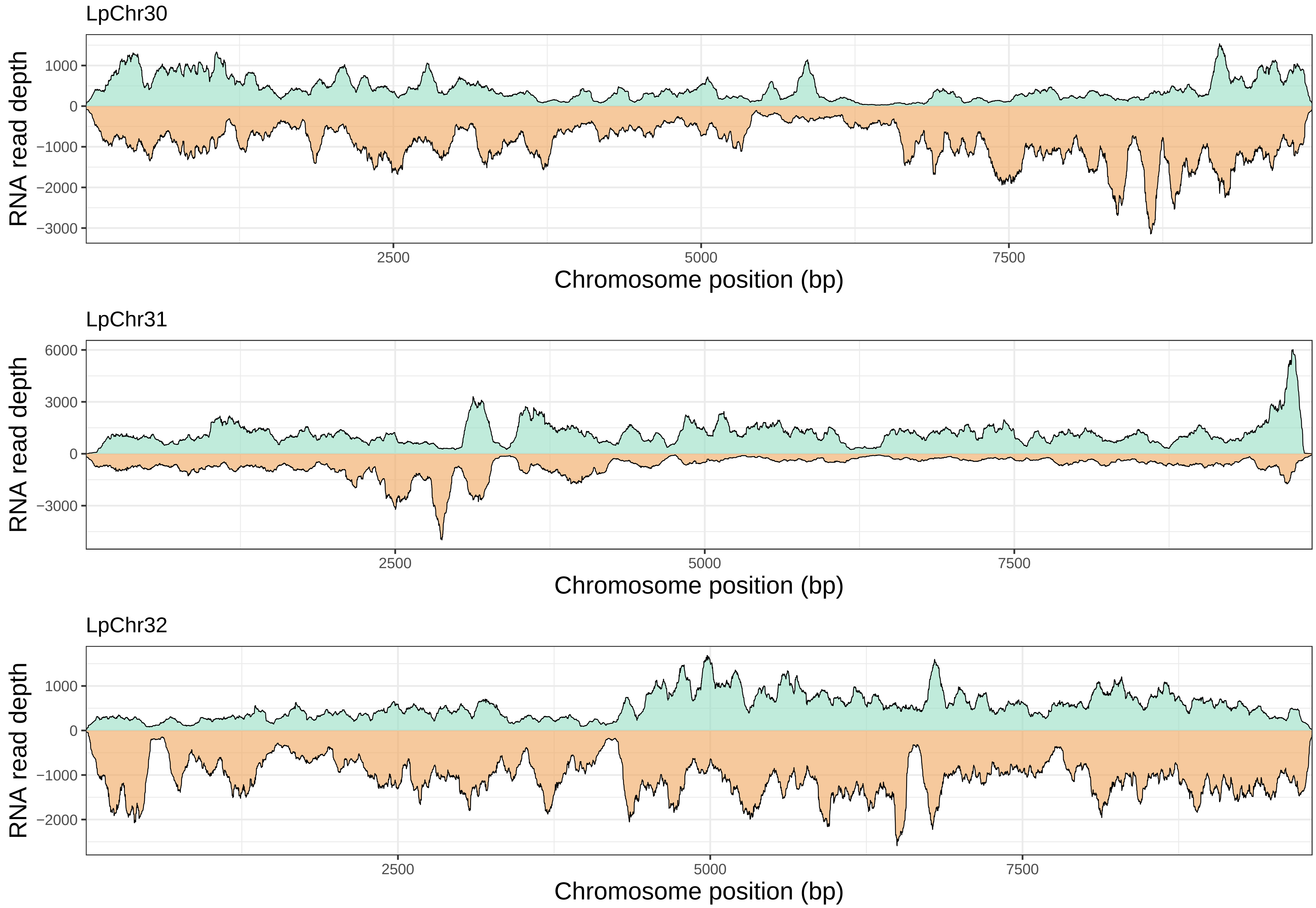

LpChr30
RNA read depth
1000
0
−1000
−2000
−3000
2500
5000
7500
Chromosome position (bp)
LpChr31
RNA read depth
6000
3000
0
−3000
2500
5000
7500
Chromosome position (bp)
LpChr32
RNA read depth
1000
0
−1000
−2000
2500
5000
7500
Chromosome position (bp)

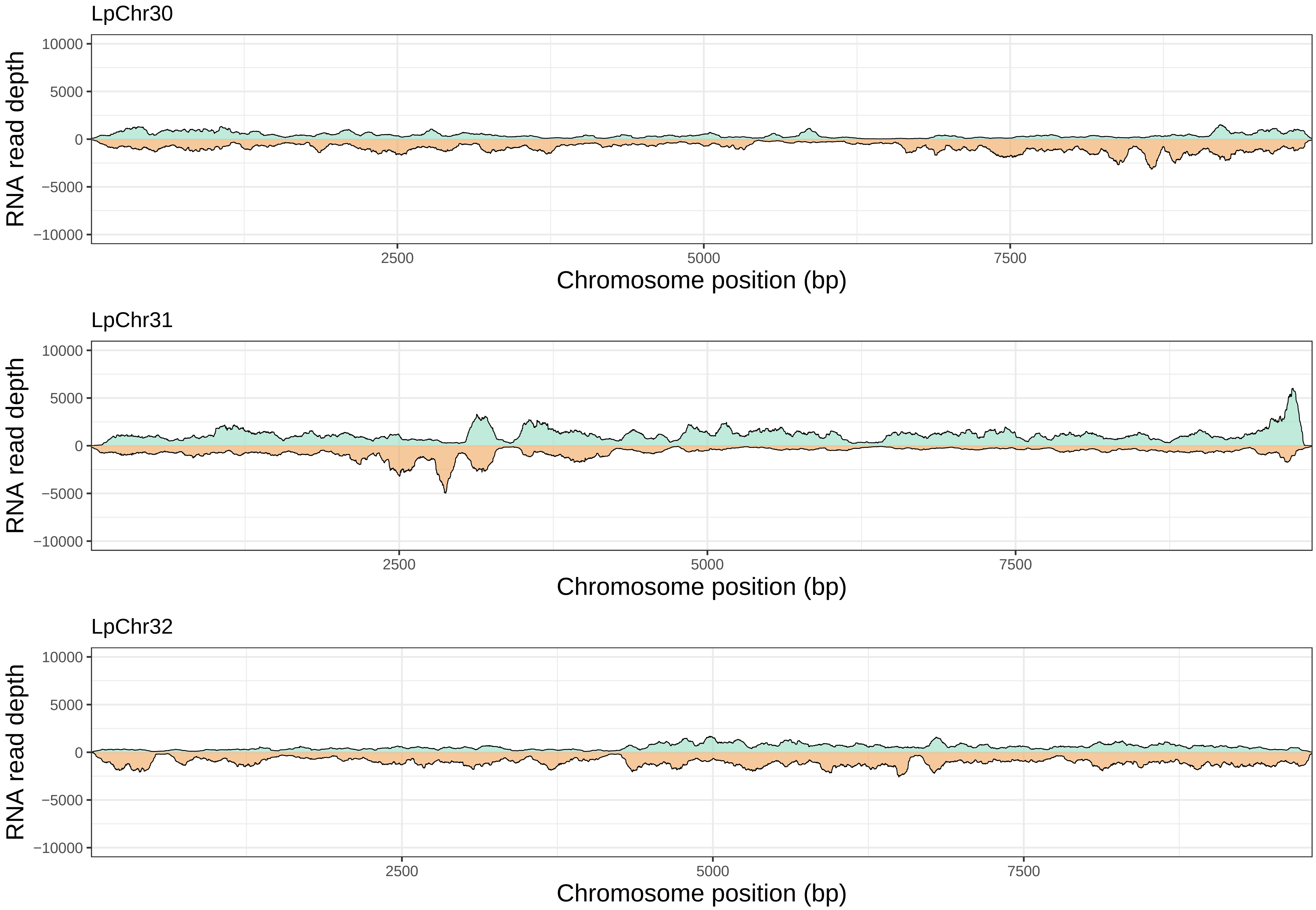

LpChr30
RNA read depth
10000
5000
0
−5000
−10000
2500
5000
7500
Chromosome position (bp)
LpChr31
RNA read depth
10000
5000
0
−5000
−10000
2500
5000
7500
Chromosome position (bp)
LpChr32
RNA read depth
10000
5000
0
−5000
−10000
2500
5000
7500
Chromosome position (bp)

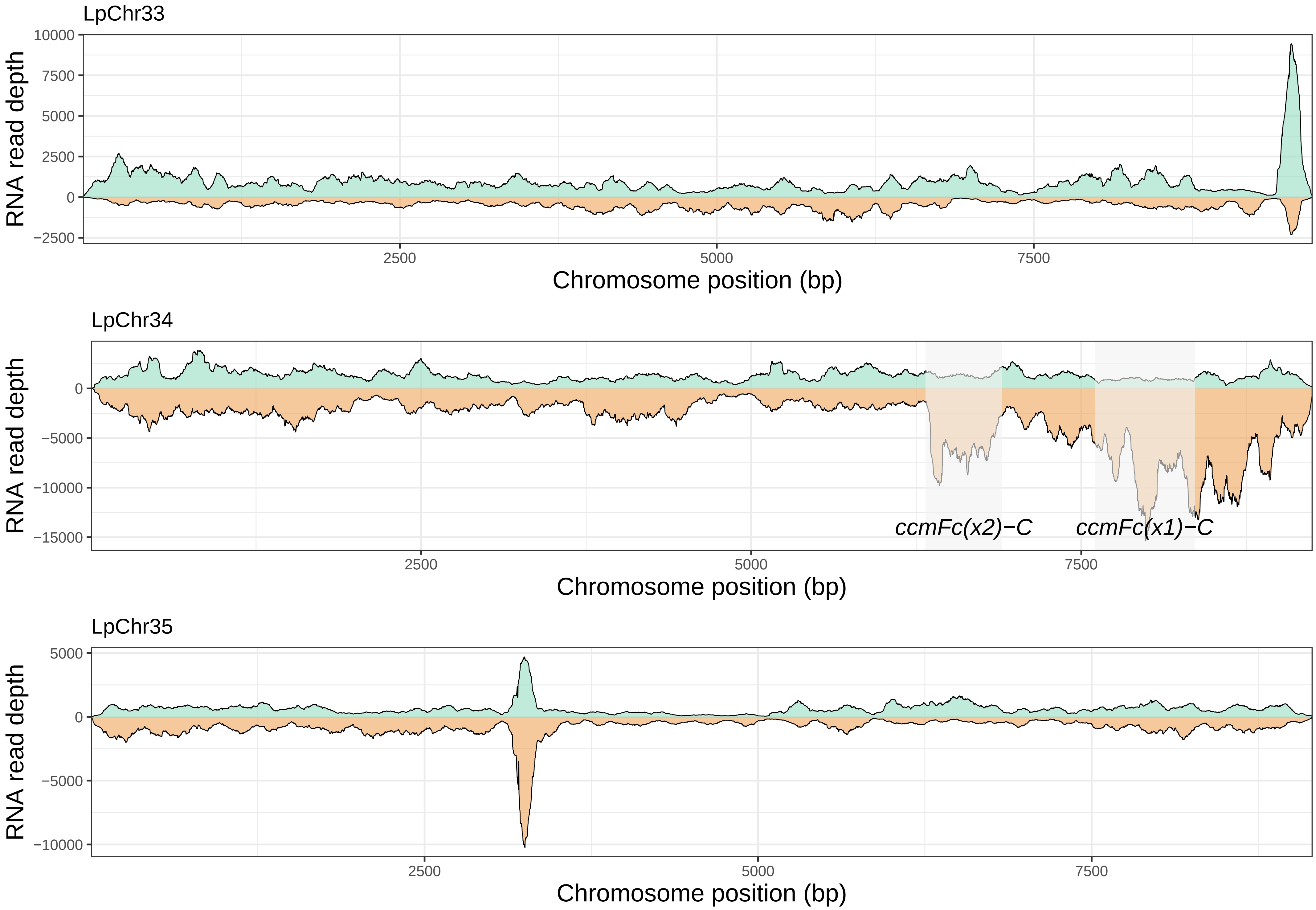

LpChr33
10000
7500
5000
2500
0
−2500
RNA read depth
2500
5000
7500
Chromosome position (bp)
LpChr34
0
−5000
−10000
−15000
RNA read depth
ccmFc(x2)−C
ccmFc(x1)−C
2500
5000
7500
Chromosome position (bp)
LpChr35
5000
0
−5000
−10000
RNA read depth
2500
5000
7500
Chromosome position (bp)

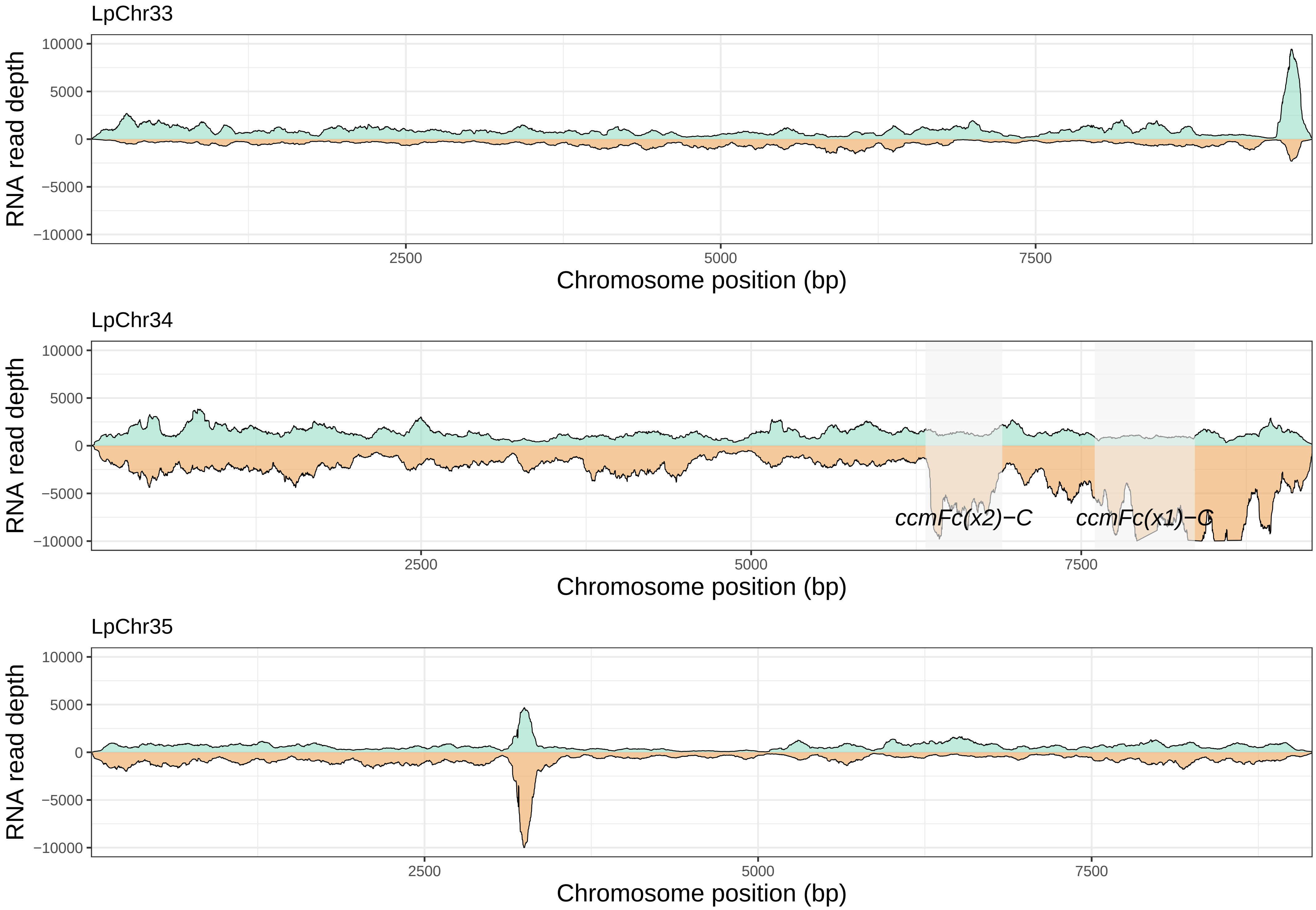
LpChr33
RNA read depth
10000
5000
0
−5000
−10000
2500
5000
7500
Chromosome position (bp)
LpChr34
ccmFc(x2)−C
ccmFc(x1)−C
LpChr35

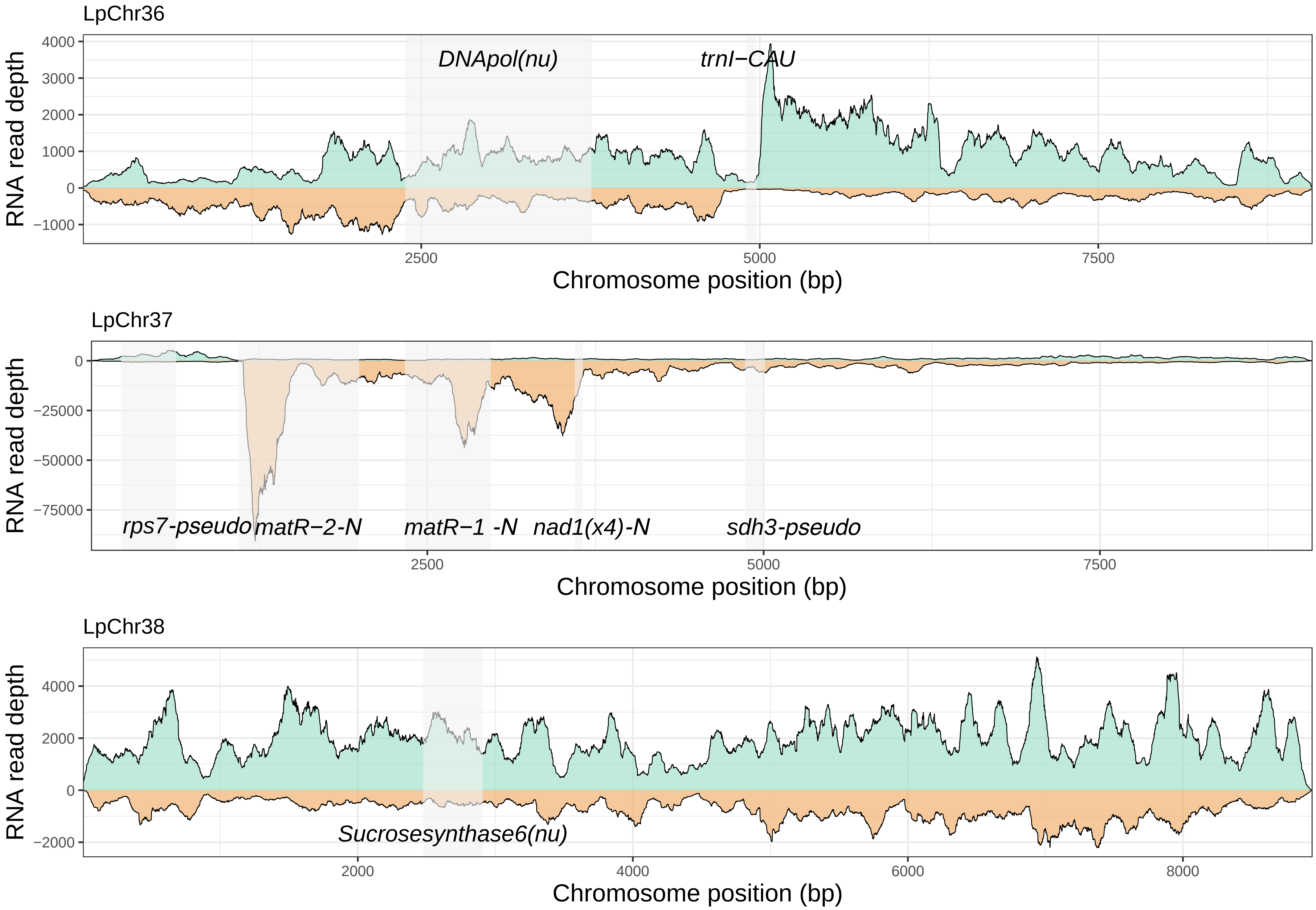

LpChr36
DNApol(nu)
trnI−CAU
RNA read depth
4000
3000
2000
1000
0
−1000
2500
5000
7500
Chromosome position (bp)
LpChr37
rps7-pseudo
matR−2-N
matR−1 -N
nad1(x4)-N
sdh3-pseudo
RNA read depth
0
−25000
−50000
−75000
2500
5000
7500
Chromosome position (bp)
LpChr38
Sucrosesynthase6(nu)
RNA read depth
4000
2000
0
−2000
2000
4000
6000
8000
Chromosome position (bp)

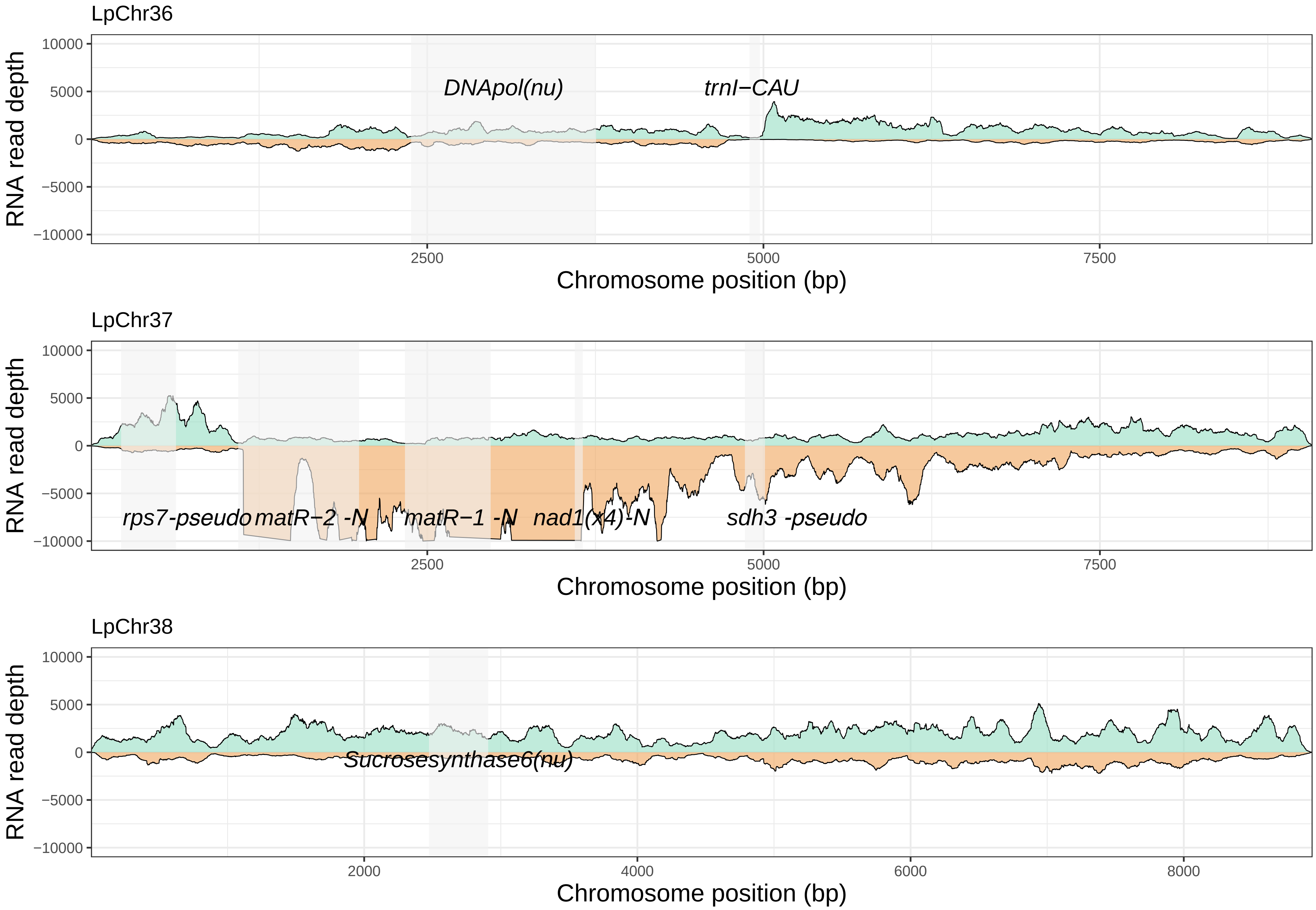

LpChr36
RNA read depth
10000
5000
0
−5000
−10000
DNApol(nu)
trnI−CAU
2500
5000
7500
Chromosome position (bp)
LpChr37
RNA read depth
10000
5000
0
−5000
−10000
rps7-pseudo
matR−2 -N
matR−1 -N
nad1(x4)-N
sdh3 -pseudo
2500
5000
7500
Chromosome position (bp)
LpChr38
RNA read depth
10000
5000
0
−5000
−10000
Sucrosesynthase6(nu)
2000
4000
6000
8000
Chromosome position (bp)

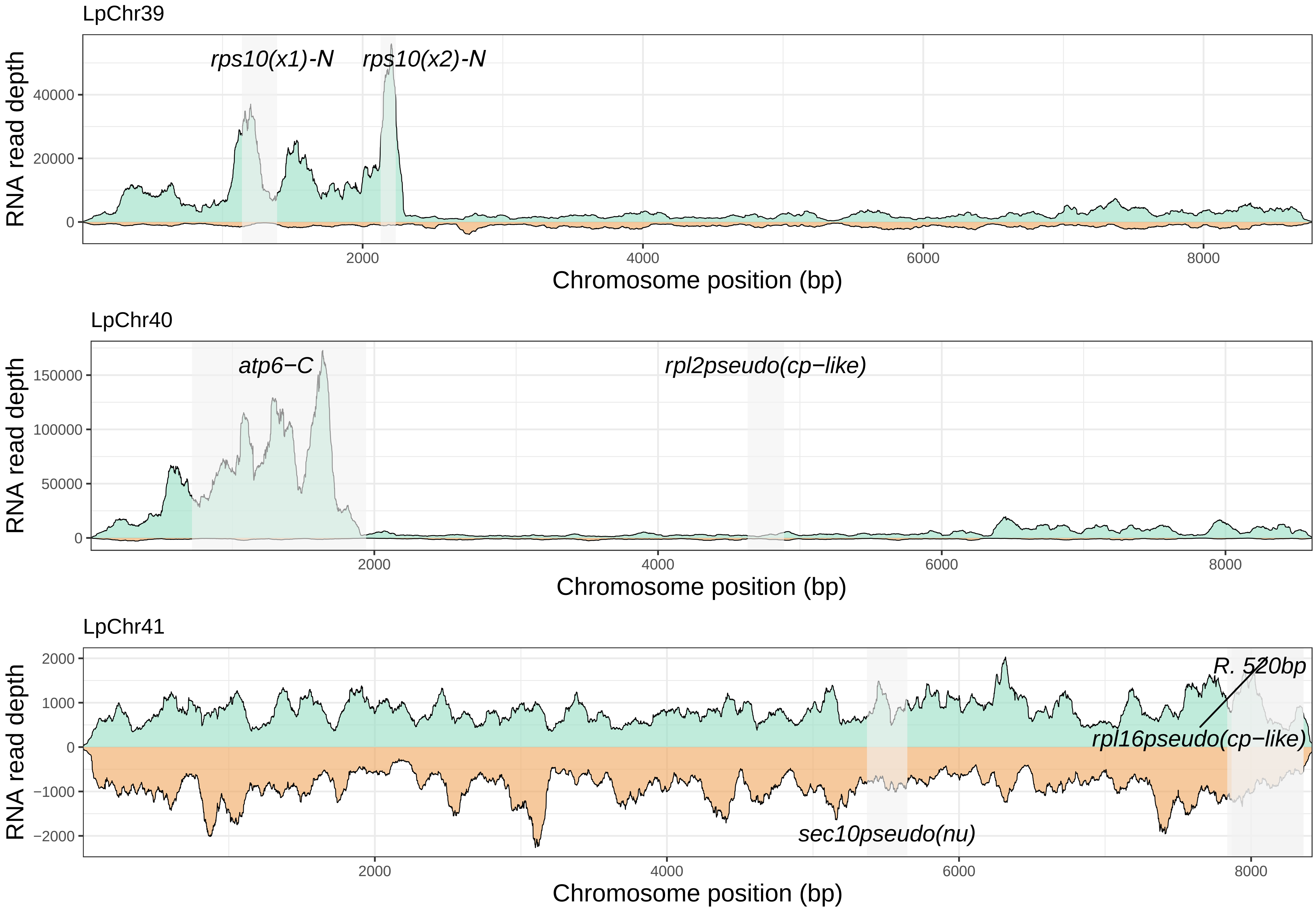

LpChr39
rps10(x1)-N
rps10(x2)-N
RNA read depth
40000
20000
0
2000
4000
6000
8000
Chromosome position (bp)
LpChr40
atp6−C
rpl2pseudo(cp−like)
RNA read depth
150000
100000
50000
0
2000
4000
6000
8000
Chromosome position (bp)
LpChr41
R. 520bp
rpl16pseudo(cp−like)
sec10pseudo(nu)
RNA read depth
2000
1000
0
−1000
−2000
2000
4000
6000
8000
Chromosome position (bp)

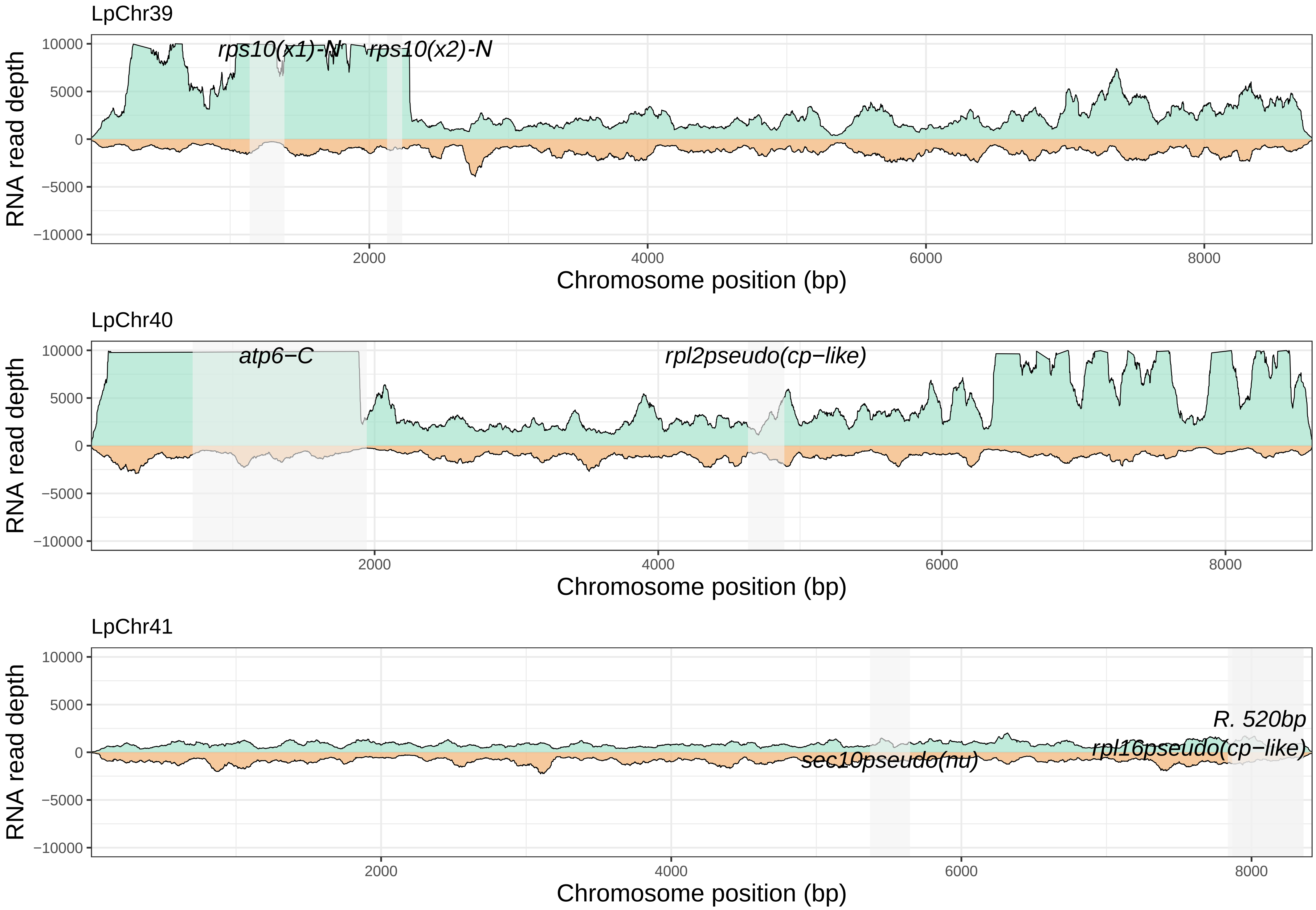
LpChr39
RNA read depth
10000
5000
0
−5000
−10000
rps10(x1)-N
rps10(x2)-N
2000
4000
6000
8000
Chromosome position (bp)
LpChr40
RNA read depth
10000
5000
0
−5000
−10000
atp6−C
rpl2pseudo(cp−like)
2000
4000
6000
8000
Chromosome position (bp)
LpChr41
RNA read depth
10000
5000
0
−5000
−10000
R. 520bp
rpl16pseudo(cp−like)
sec10pseudo(nu)
2000
4000
6000
8000
Chromosome position (bp)

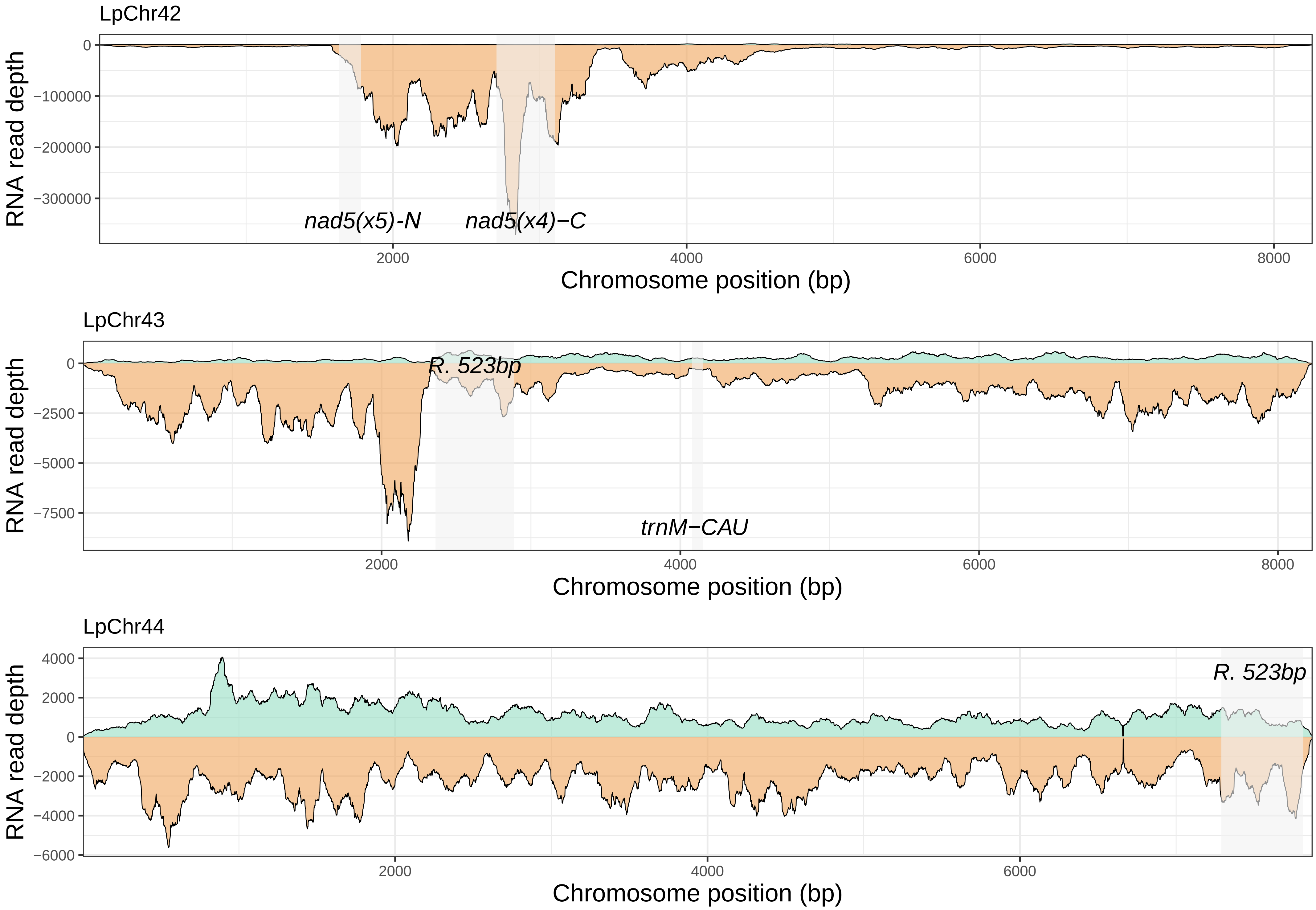
LpChr42
RNA read depth
0
−100000
−200000
−300000
nad5(x5)-N
nad5(x4)−C
2000
4000
6000
8000
Chromosome position (bp)
LpChr43
RNA read depth
0
−2500
−5000
−7500
R. 523bp
trnM−CAU
2000
4000
6000
8000
Chromosome position (bp)
LpChr44
RNA read depth
4000
2000
0
−2000
−4000
−6000
R. 523bp
2000
4000
6000
Chromosome position (bp)

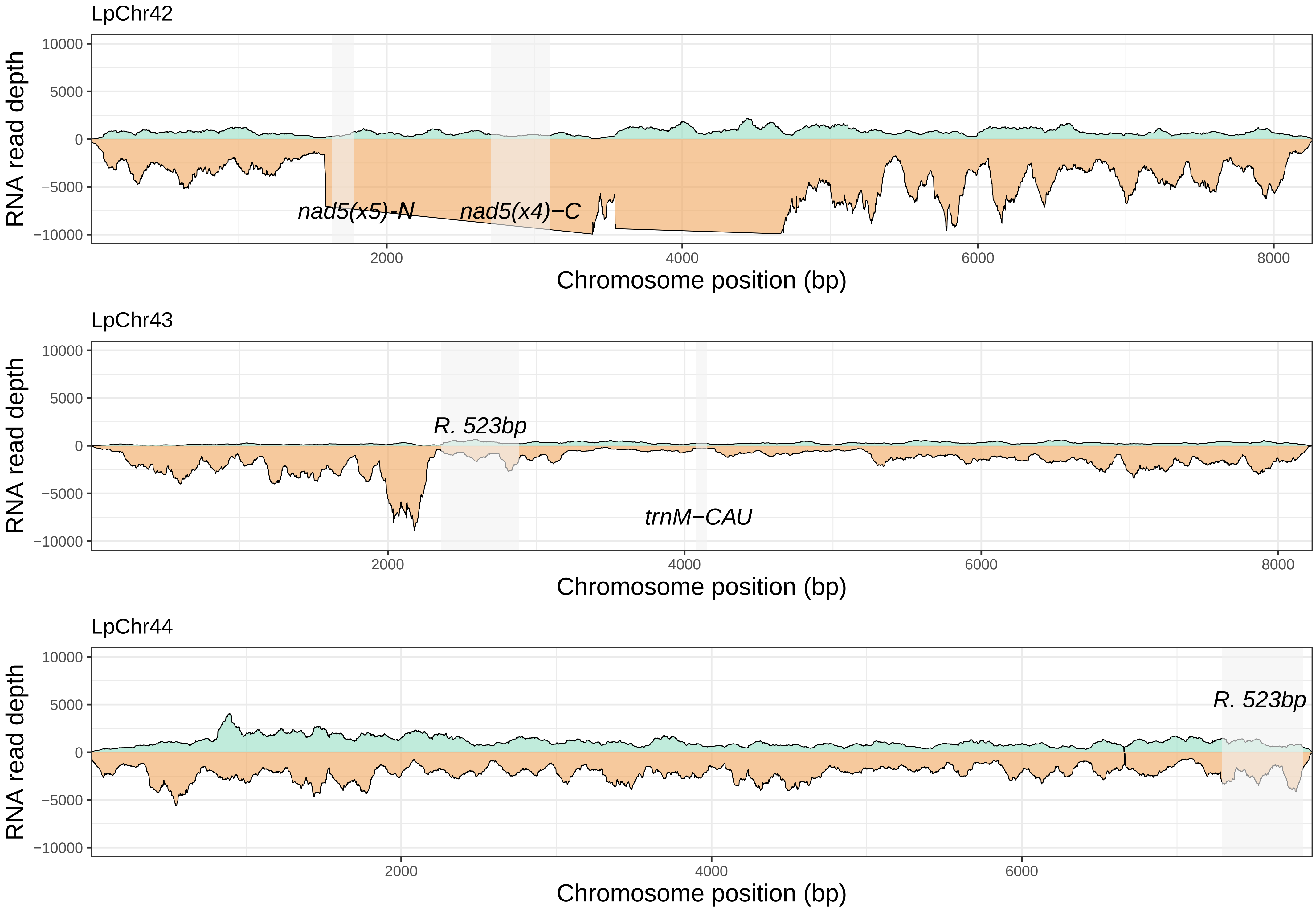

LpChr42
RNA read depth
10000
5000
0
−5000
−10000
nad5(x5)-N
nad5(x4)−C
2000
4000
6000
8000
Chromosome position (bp)
LpChr43
RNA read depth
10000
5000
0
−5000
−10000
R. 523bp
trnM−CAU
2000
4000
6000
8000
Chromosome position (bp)
LpChr44
RNA read depth
10000
5000
0
−5000
−10000
R. 523bp
2000
4000
6000
Chromosome position (bp)

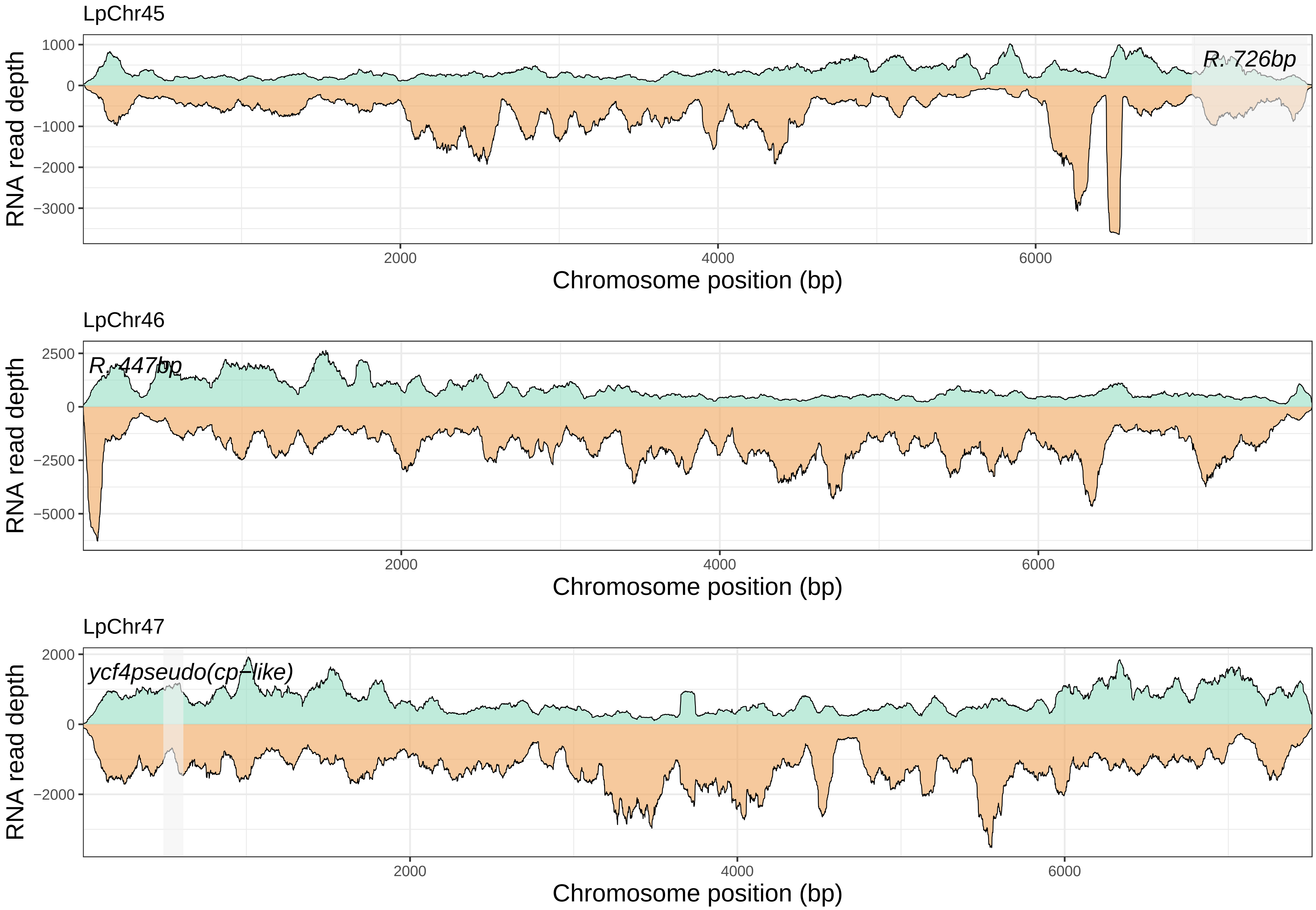

LpChr45
R. 726bp
1000
0
−1000
−2000
−3000
2000
4000
6000
RNA read depth
Chromosome position (bp)
LpChr46
R. 447bp
2500
0
−2500
−5000
2000
4000
6000
RNA read depth
Chromosome position (bp)
LpChr47
ycf4pseudo(cp−like)
2000
0
−2000
2000
4000
6000
RNA read depth
Chromosome position (bp)

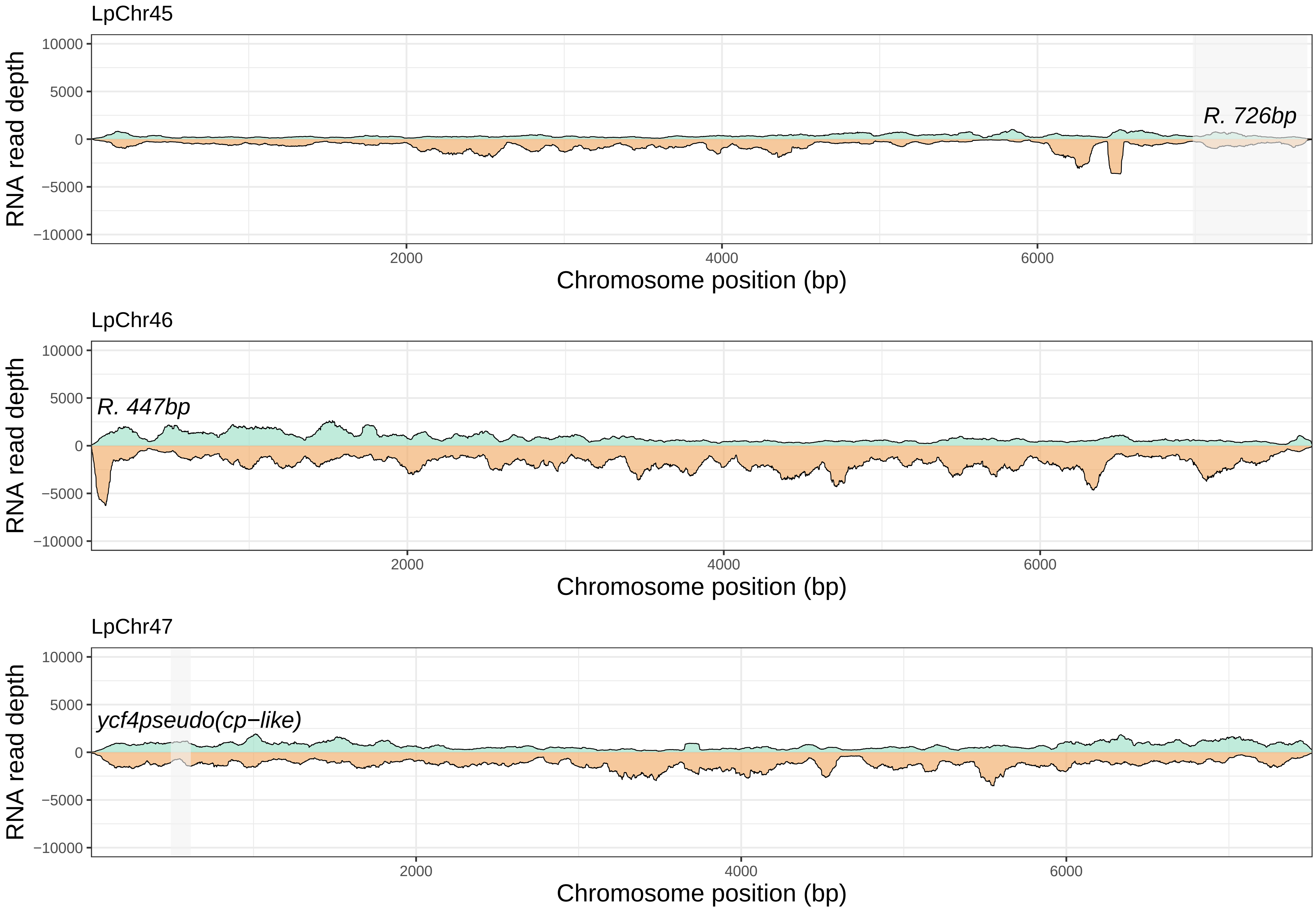

LpChr45
R. 726bp
RNA read depth
10000
5000
0
−5000
−10000
2000
4000
6000
Chromosome position (bp)
LpChr46
R. 447bp
RNA read depth
10000
5000
0
−5000
−10000
2000
4000
6000
Chromosome position (bp)
LpChr47
ycf4pseudo(cp−like)
RNA read depth
10000
5000
0
−5000
−10000
2000
4000
6000
Chromosome position (bp)

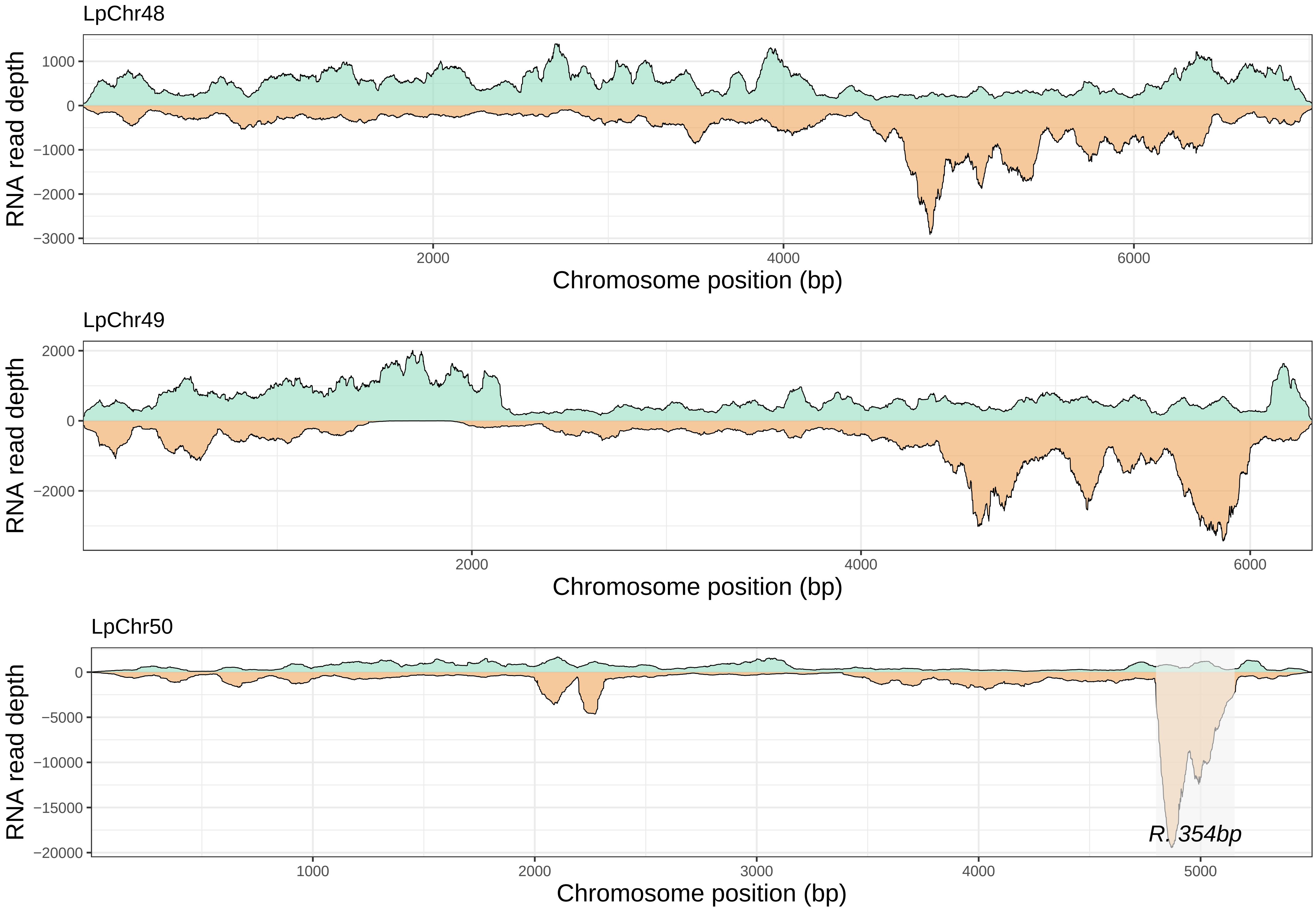

LpChr48
RNA read depth
1000
0
−1000
−2000
−3000
2000
4000
6000
Chromosome position (bp)
LpChr49
RNA read depth
2000
0
−2000
2000
4000
6000
Chromosome position (bp)
LpChr50
RNA read depth
0
−5000
−10000
−15000
−20000
1000
2000
3000
4000
5000
R. 354bp
Chromosome position (bp)

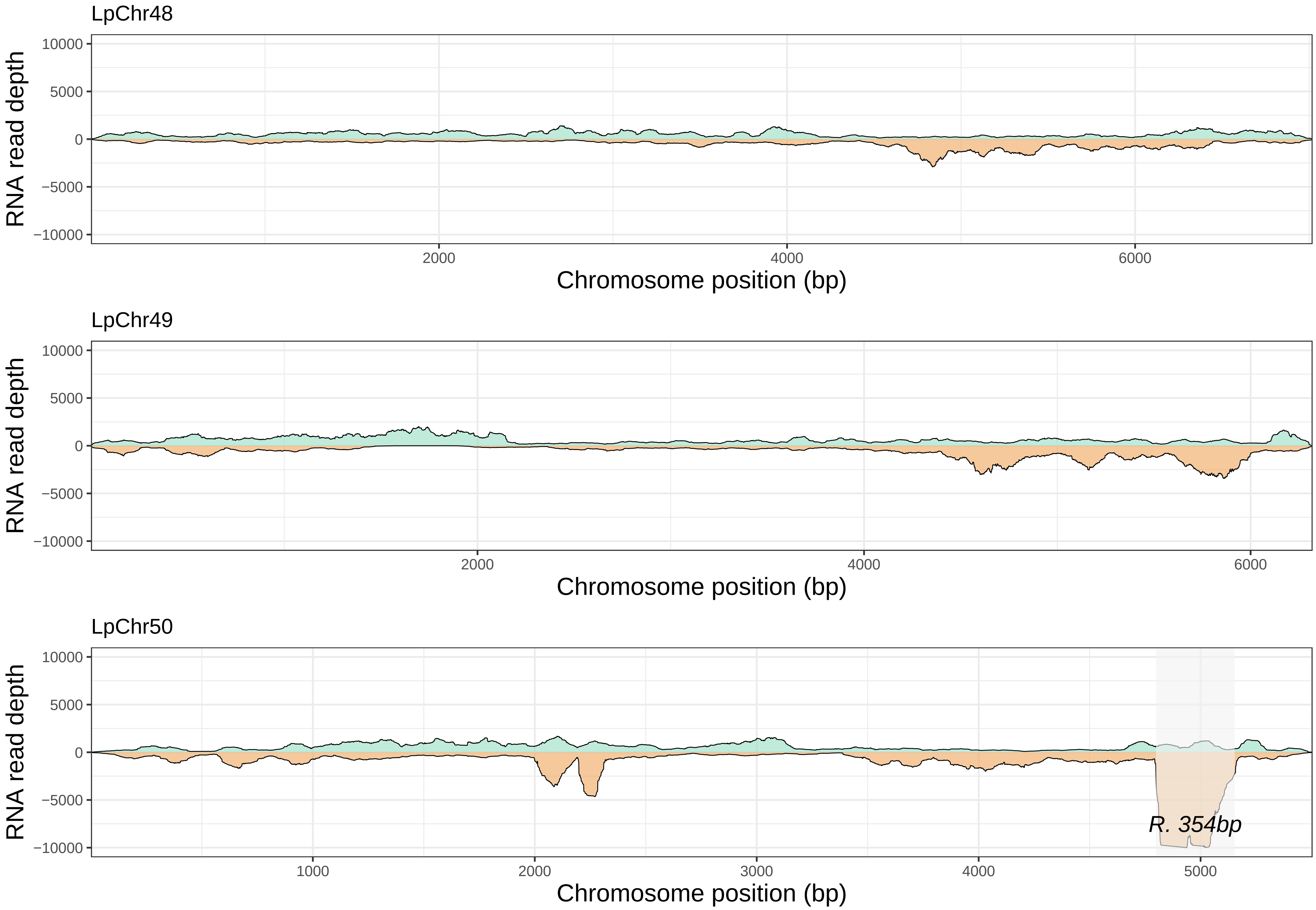

LpChr48
RNA read depth
10000
5000
0
−5000
−10000
2000
4000
6000
Chromosome position (bp)
LpChr49
RNA read depth
10000
5000
0
−5000
−10000
2000
4000
6000
Chromosome position (bp)
LpChr50
RNA read depth
10000
5000
0
−5000
−10000
1000
2000
3000
4000
5000
R. 354bp
Chromosome position (bp)

**Figure S5. Maximum Likelihood (ML) phylogenetic analyses of mitochondrial and nuclear rps14 genes in *Lophophytum* spp. and *Ombrophytum*.** To determine the phylogenetic origin of the nuclear gene encoding the organellar ribosomal protein RPS14, homologous sequences were retrieved from photosynthetic species available in the PLAZA v5.0 database (https://bioinformatics.psb.ugent.be/plaza.dev/instances/dicots_05/). Codon-based alignments were generated using MAFFT v7.407 (Katoh & Standley, 2013) and PAL2NAL (Suyama et al., 2006). Poorly aligned or highly divergent regions were filtered out with BMGE v1.12 (Criscuolo & Gribaldo, 2010). The phylogenetic tree was reconstructed under a maximum likelihood framework with RAxML v8.2.11 (Stamatakis, 2014), employing the GTR+GAMMA+I substitution model and 1,000 bootstrap replicates. The resulting tree was visualized using FigTree (http://tree.bio.ed.ac.uk/software/figtree/).

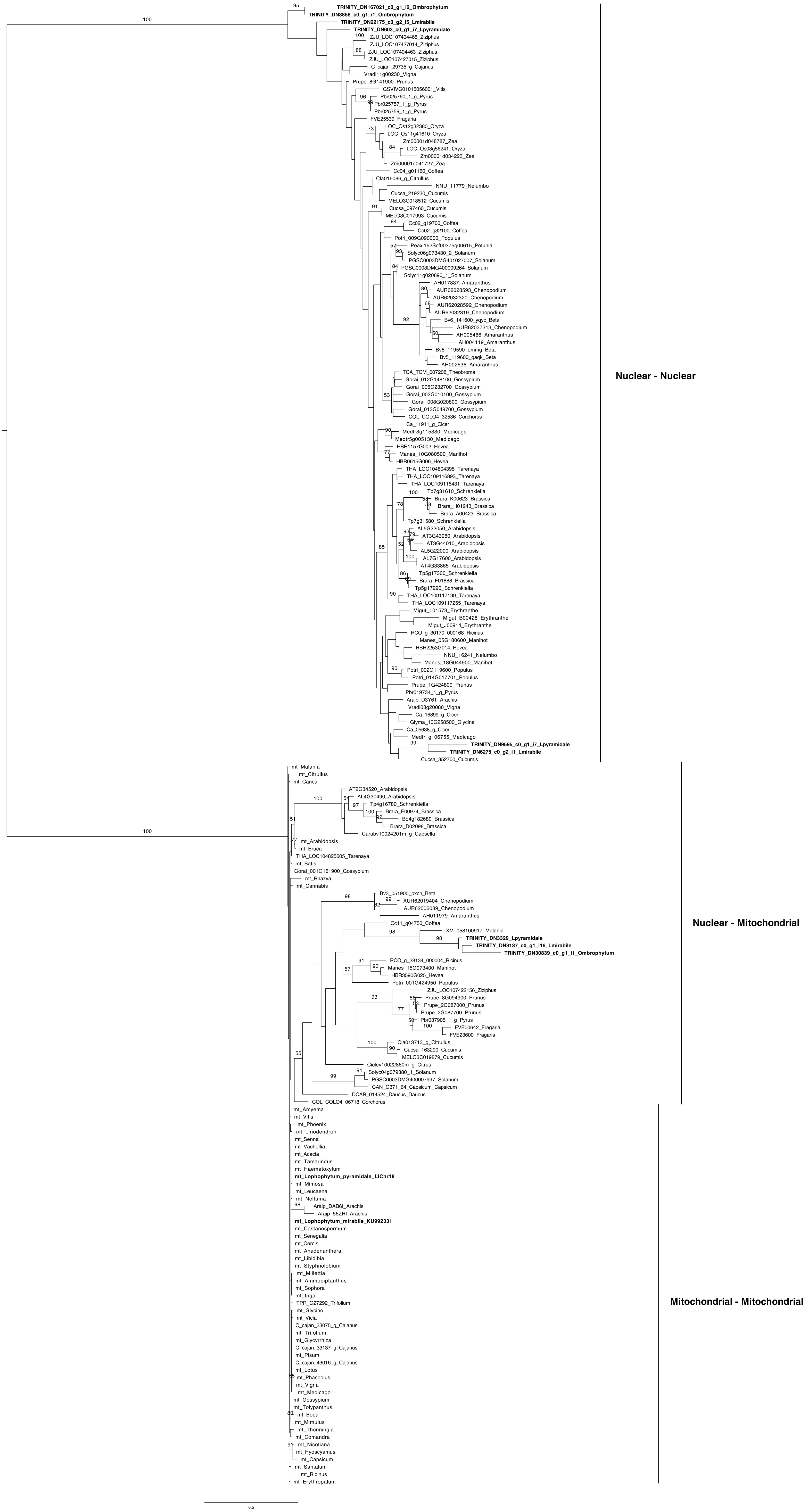

**Figure S6. RNA read depth of protein-coding genes (CDSs) in the *Lophophytum pyramidale* mtDNA.** The curves show the number of RNA reads aligned to each nucleotide position. Each gene (represented as a grey rectangle) is extended by 300 nucleotides upstream and downstream to include flanking regions.

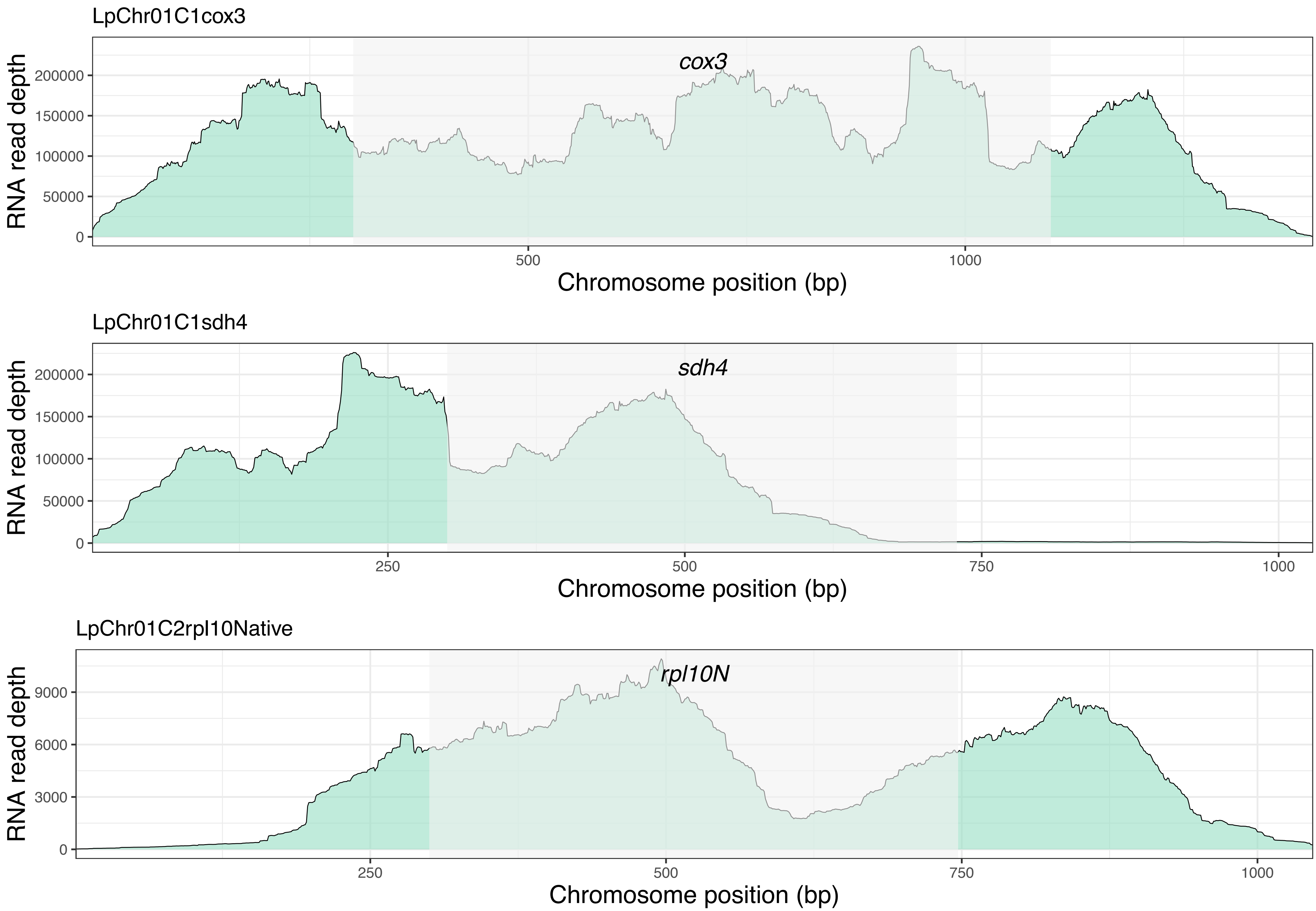

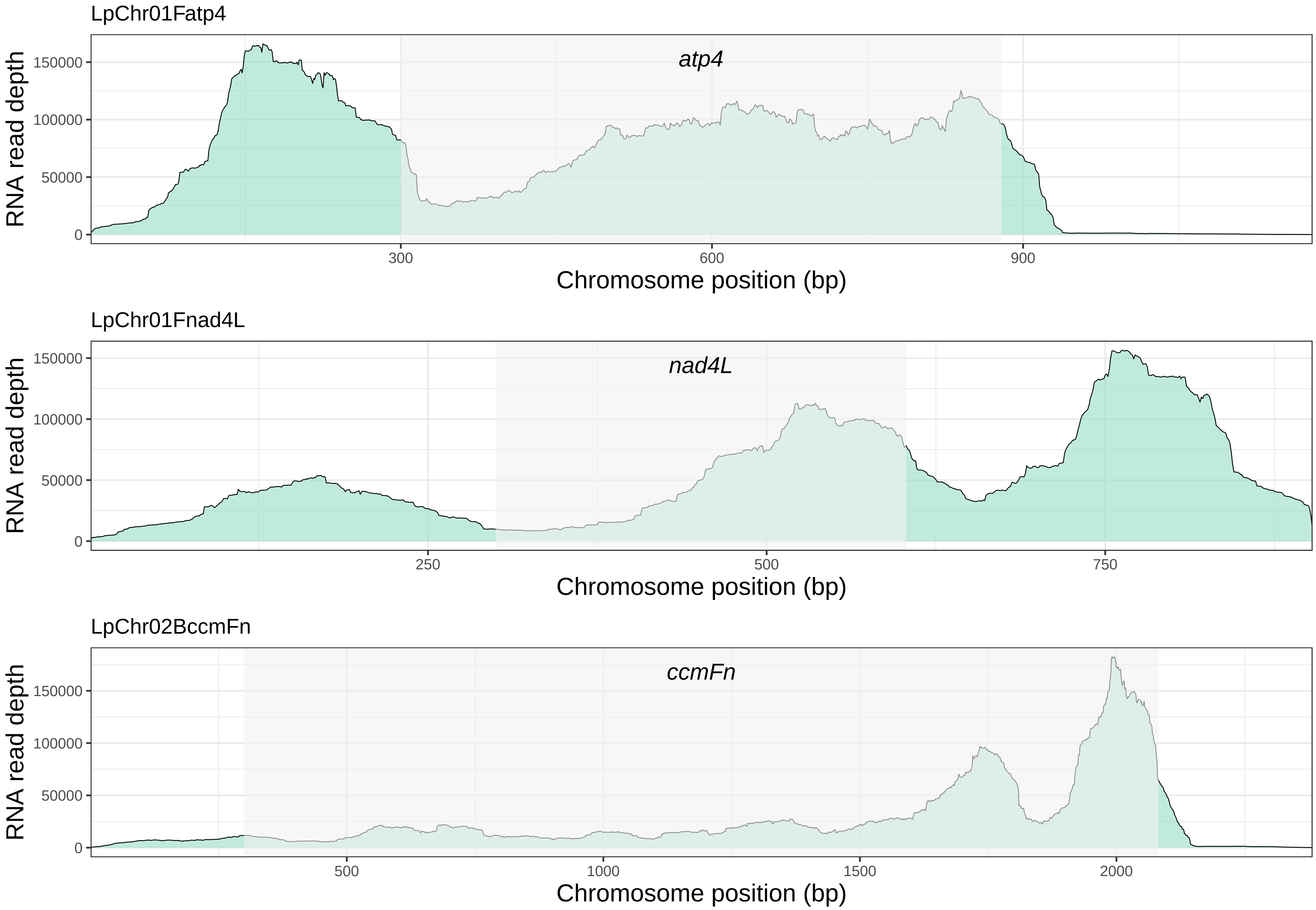

LpChr01Fatp4
atp4
RNA read depth
0
50000
100000
150000
300
600
900
Chromosome position (bp)
LpChr01Fnad4L
nad4L
RNA read depth
0
50000
100000
150000
250
500
750
Chromosome position (bp)
LpChr02BccmFn
ccmFn
RNA read depth
0
50000
100000
150000
500
1000
1500
2000
Chromosome position (bp)

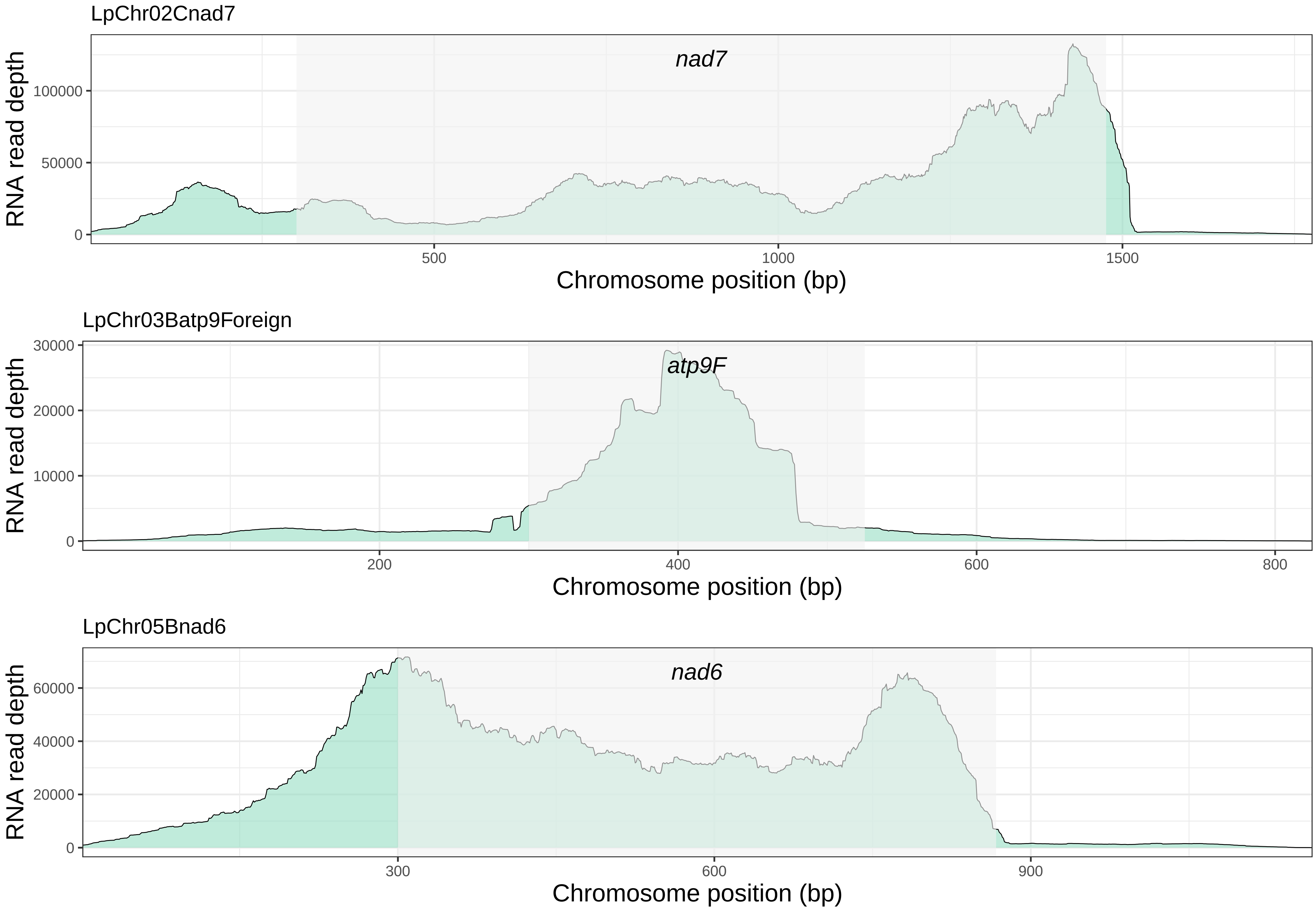
LpChr02Cnad7
nad7
RNA read depth
100000
50000
0
500
1000
1500
Chromosome position (bp)
LpChr03Batp9Foreign
atp9F
RNA read depth
30000
20000
10000
0
200
400
600
800
Chromosome position (bp)
LpChr05Bnad6
nad6
RNA read depth
60000
40000
20000
0
300
600
900
Chromosome position (bp)

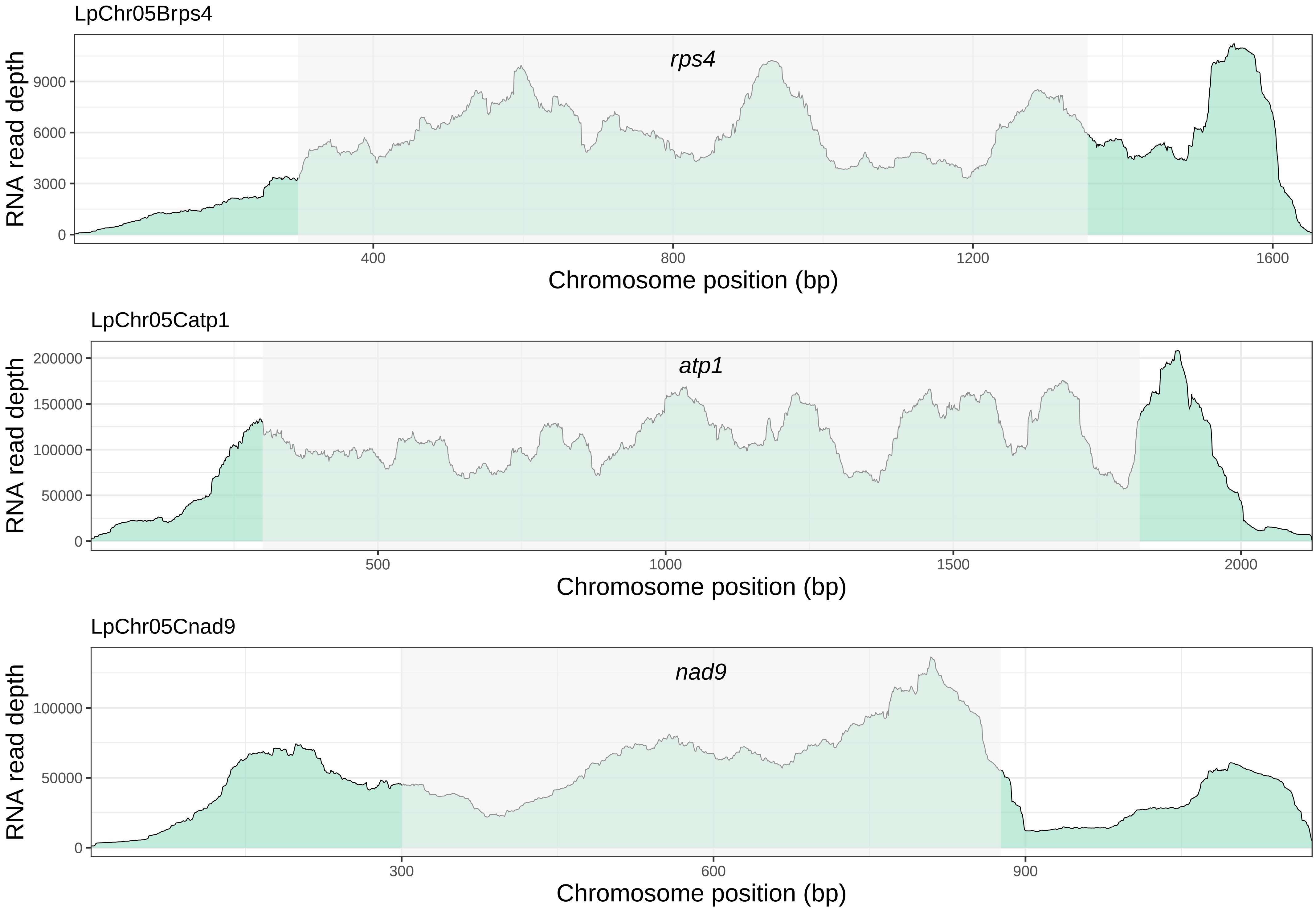

LpChr05Brps4
rps4
RNA read depth
0
3000
6000
9000
400
800
1200
1600
Chromosome position (bp)
LpChr05Catp1
atp1
RNA read depth
0
50000
100000
150000
200000
500
1000
1500
2000
Chromosome position (bp)
LpChr05Cnad9
nad9
RNA read depth
0
50000
100000
300
600
900
Chromosome position (bp)

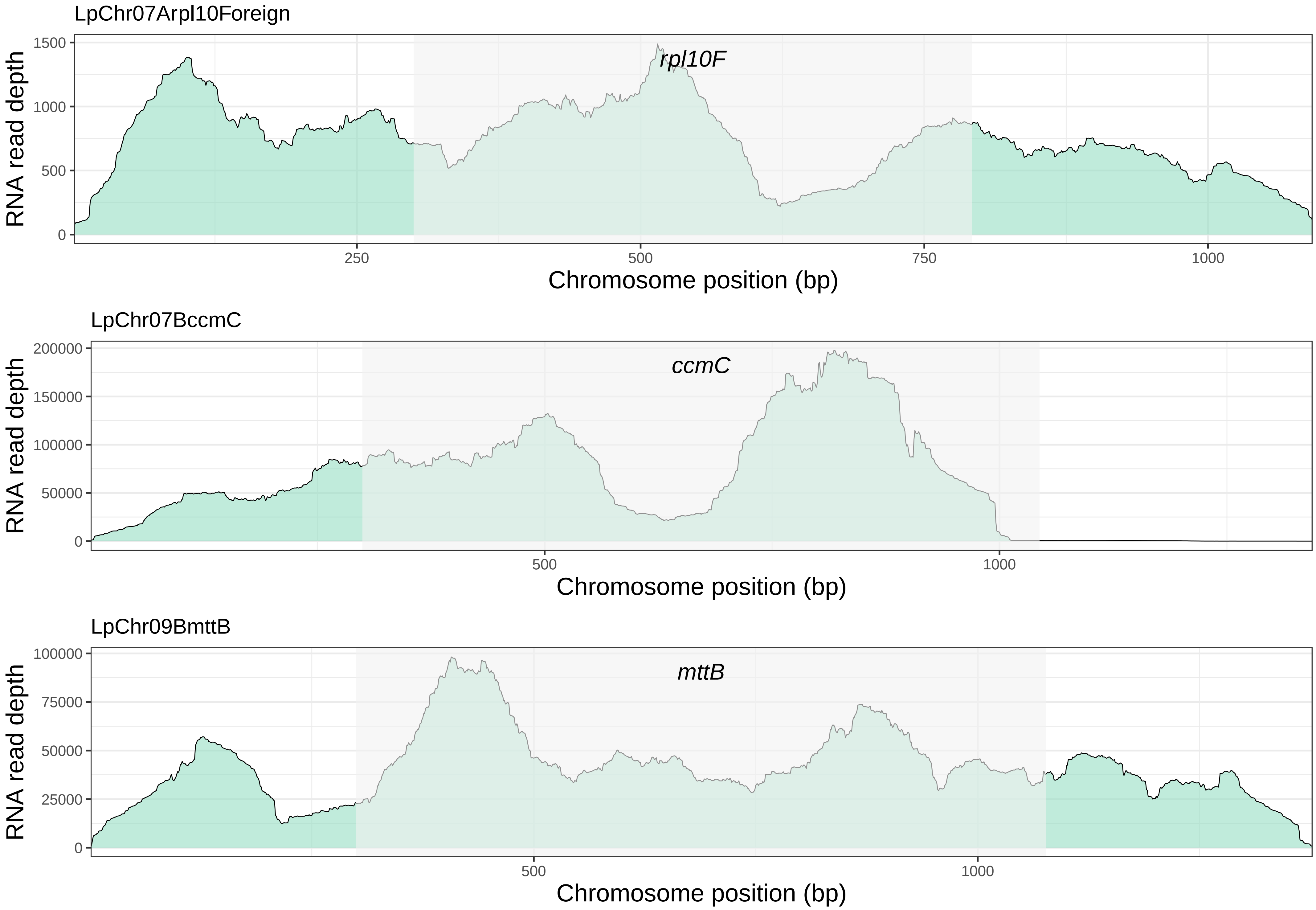

LpChr07Arpl10Foreign
rpl10F
RNA read depth
0
500
1000
1500
250
500
750
1000
Chromosome position (bp)
LpChr07BccmC
ccmC
RNA read depth
0
50000
100000
150000
200000
500
1000
Chromosome position (bp)
LpChr09BmttB
mttB
RNA read depth
0
25000
50000
75000
100000
500
1000
Chromosome position (bp)

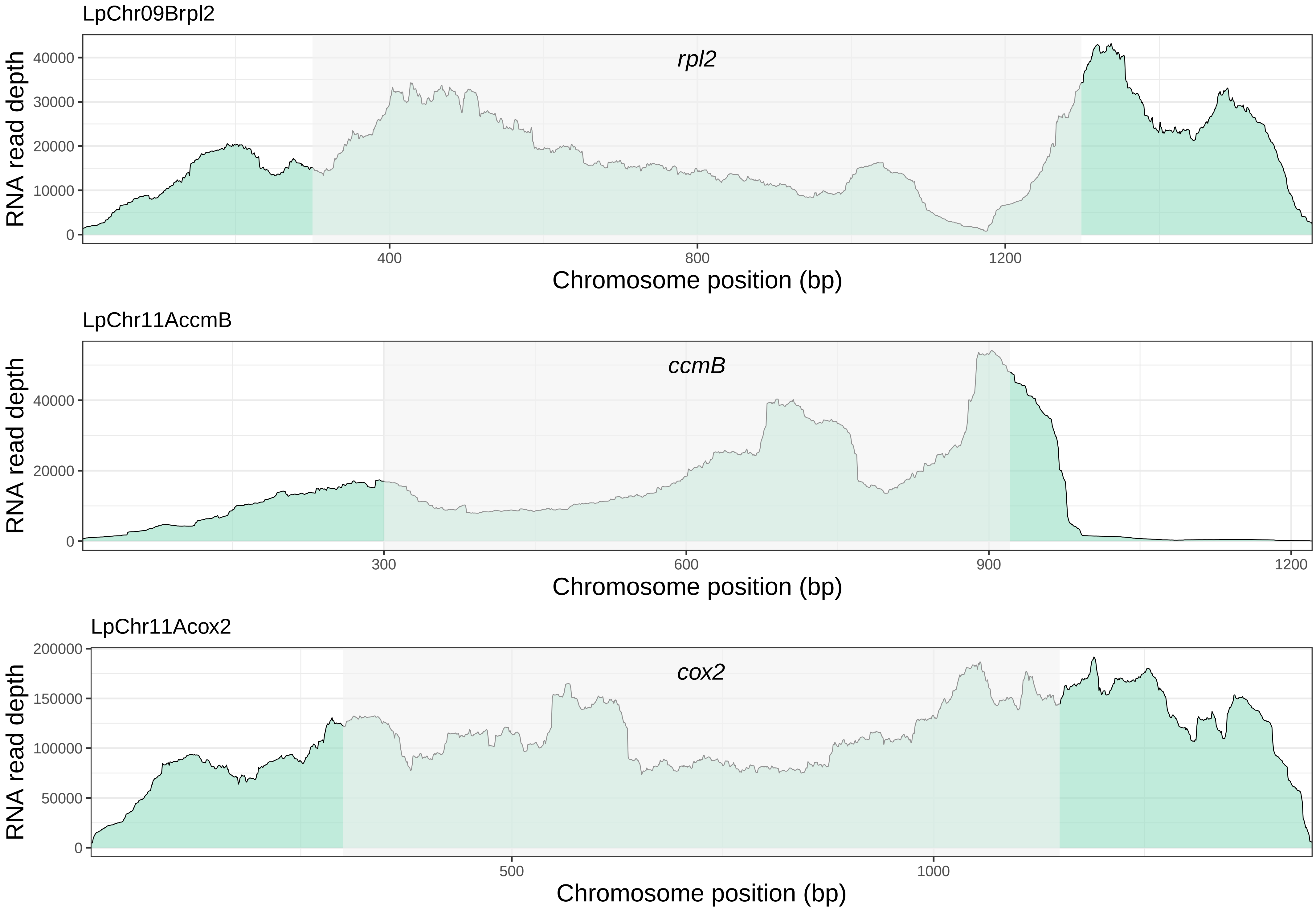

LpChr09Brpl2
rpl2
RNA read depth
40000
30000
20000
10000
0
400
800
1200
Chromosome position (bp)
LpChr11AccmB
ccmB
RNA read depth
40000
20000
0
300
600
900
1200
Chromosome position (bp)
LpChr11Acox2
cox2
RNA read depth
200000
150000
100000
50000
0
500
1000
Chromosome position (bp)

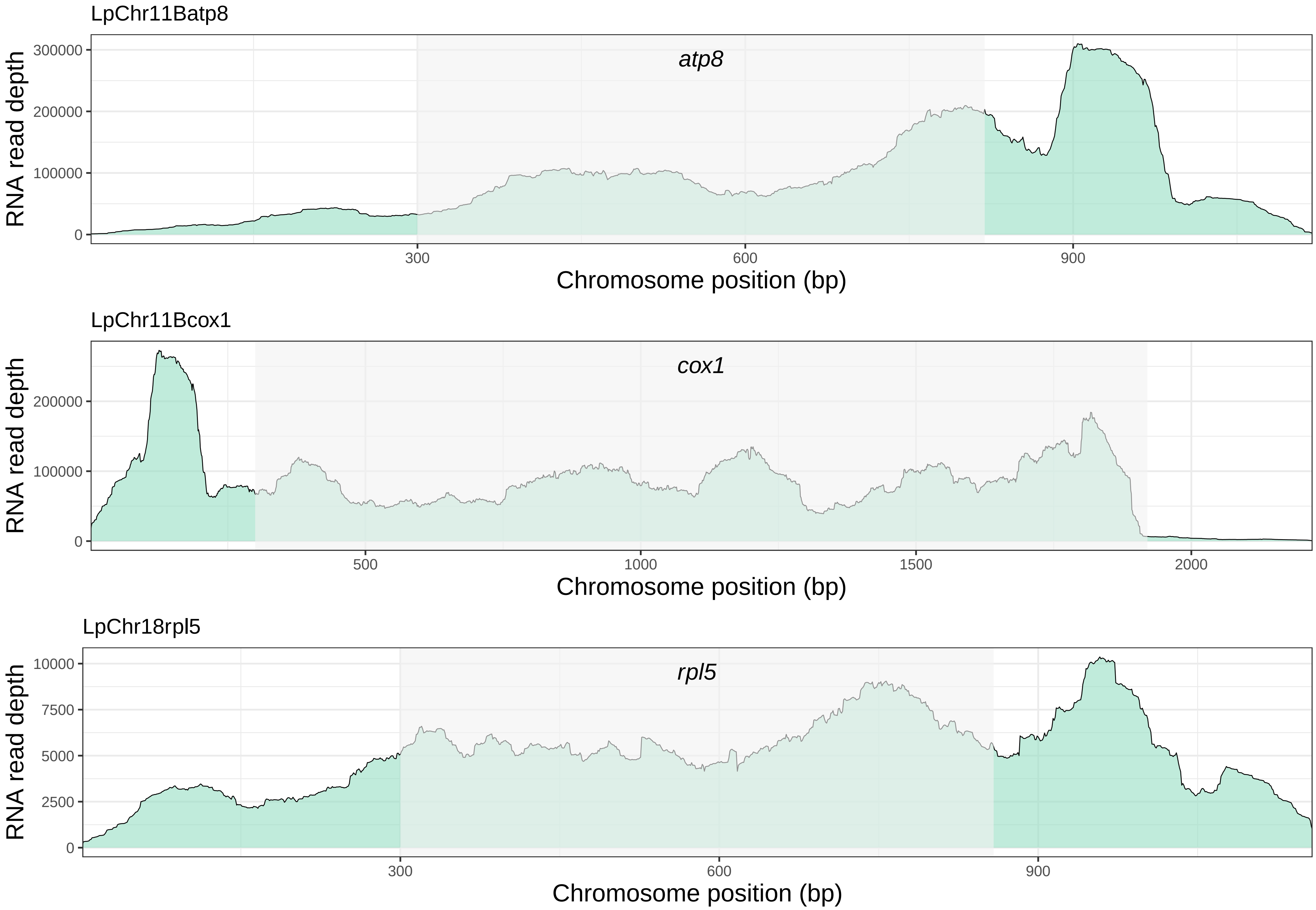
LpChr11Batp8
atp8
RNA read depth
300000
200000
100000
0
300
600
900
Chromosome position (bp)
LpChr11Bcox1
cox1
RNA read depth
200000
100000
0
500
1000
1500
2000
Chromosome position (bp)
LpChr18rpl5
rpl5
RNA read depth
10000
7500
5000
2500
0
300
600
900
Chromosome position (bp)

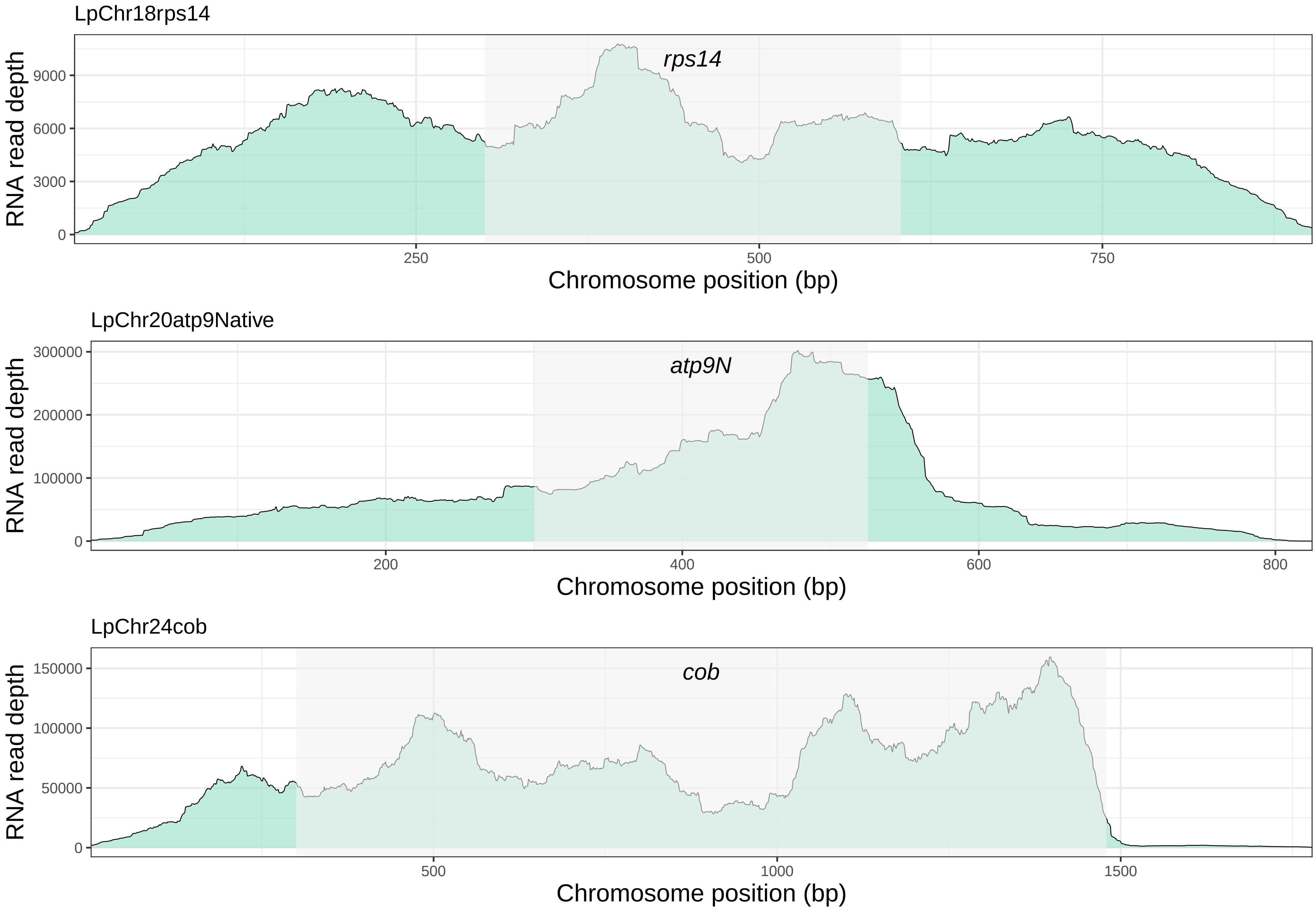
LpChr18rps14
rps14
RNA read depth
0
3000
6000
9000
250
500
750
Chromosome position (bp)
LpChr20atp9Native
atp9N
RNA read depth
0
100000
200000
300000
200
400
600
800
Chromosome position (bp)
LpChr24cob
cob
RNA read depth
0
50000
100000
150000
500
1000
1500
Chromosome position (bp)

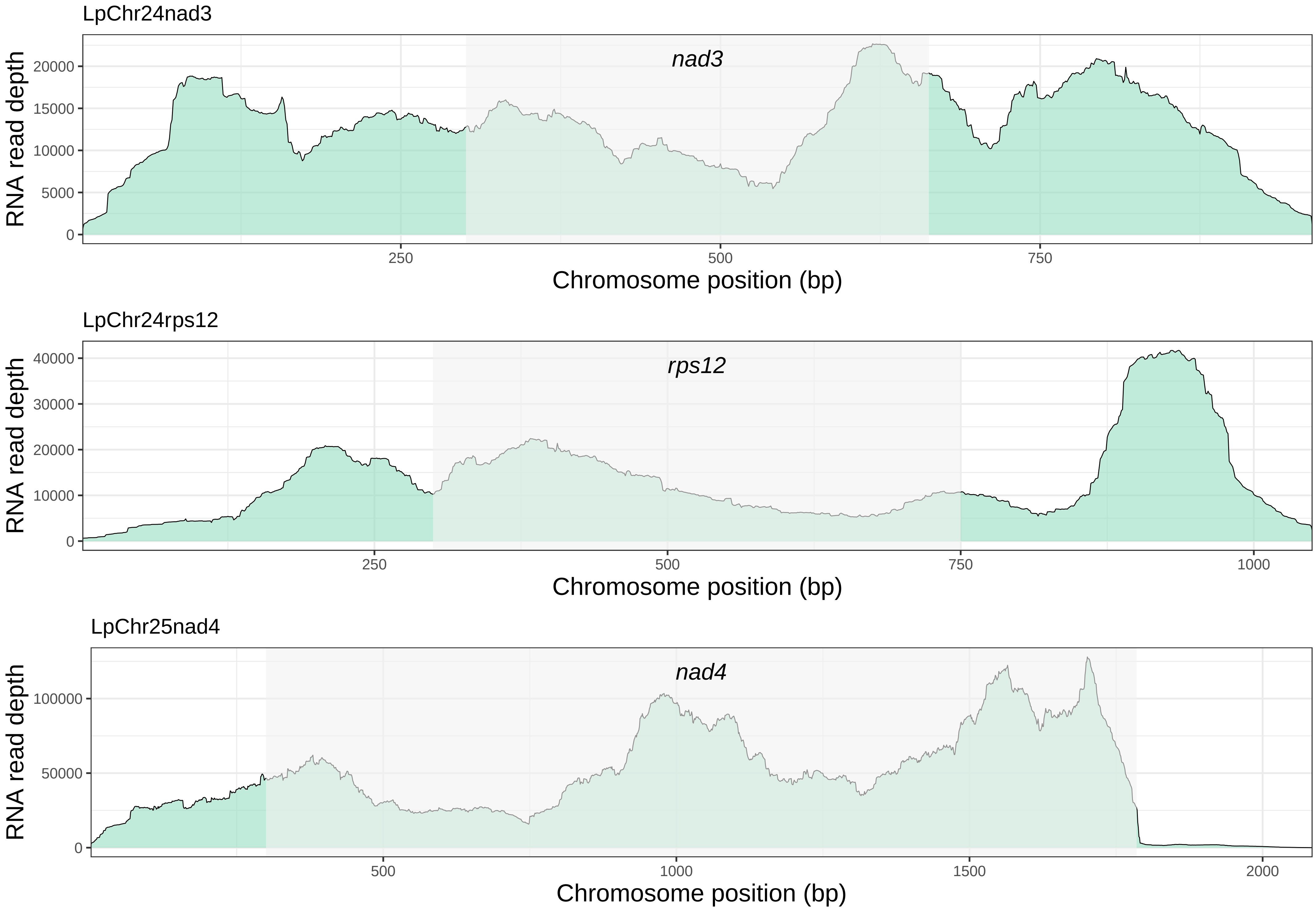
LpChr24nad3
nad3
RNA read depth
0
5000
10000
15000
20000
250
500
750
Chromosome position (bp)
LpChr24rps12
rps12
RNA read depth
0
10000
20000
30000
40000
250
500
750
1000
Chromosome position (bp)
LpChr25nad4
nad4
RNA read depth
0
50000
100000
500
1000
1500
2000
Chromosome position (bp)

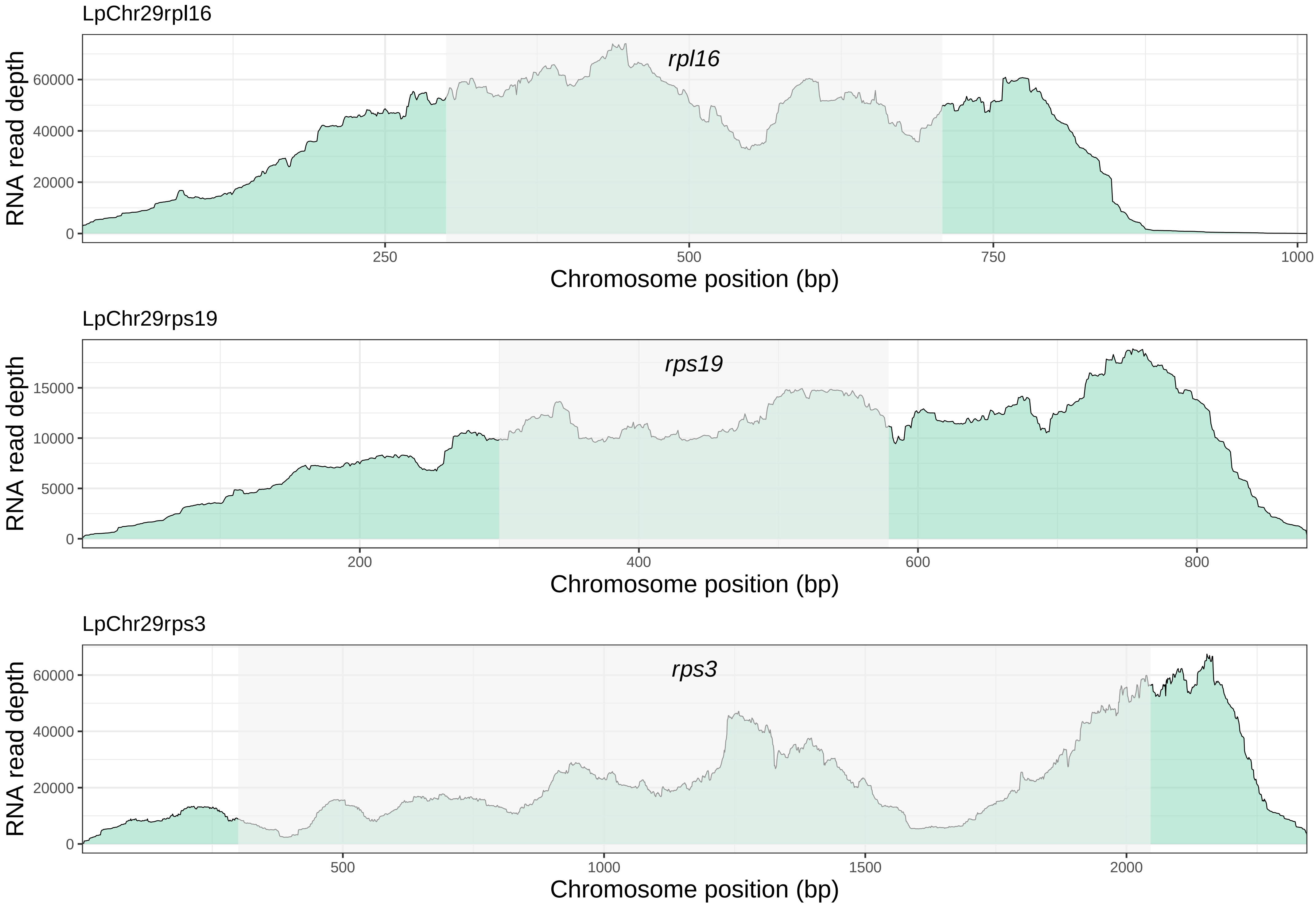

LpChr29rpl16
rpl16
RNA read depth
0
20000
40000
60000
250
500
750
1000
Chromosome position (bp)
LpChr29rps19
rps19
RNA read depth
0
5000
10000
15000
200
400
600
800
Chromosome position (bp)
LpChr29rps3
rps3
RNA read depth
0
20000
40000
60000
500
1000
1500
2000
Chromosome position (bp)

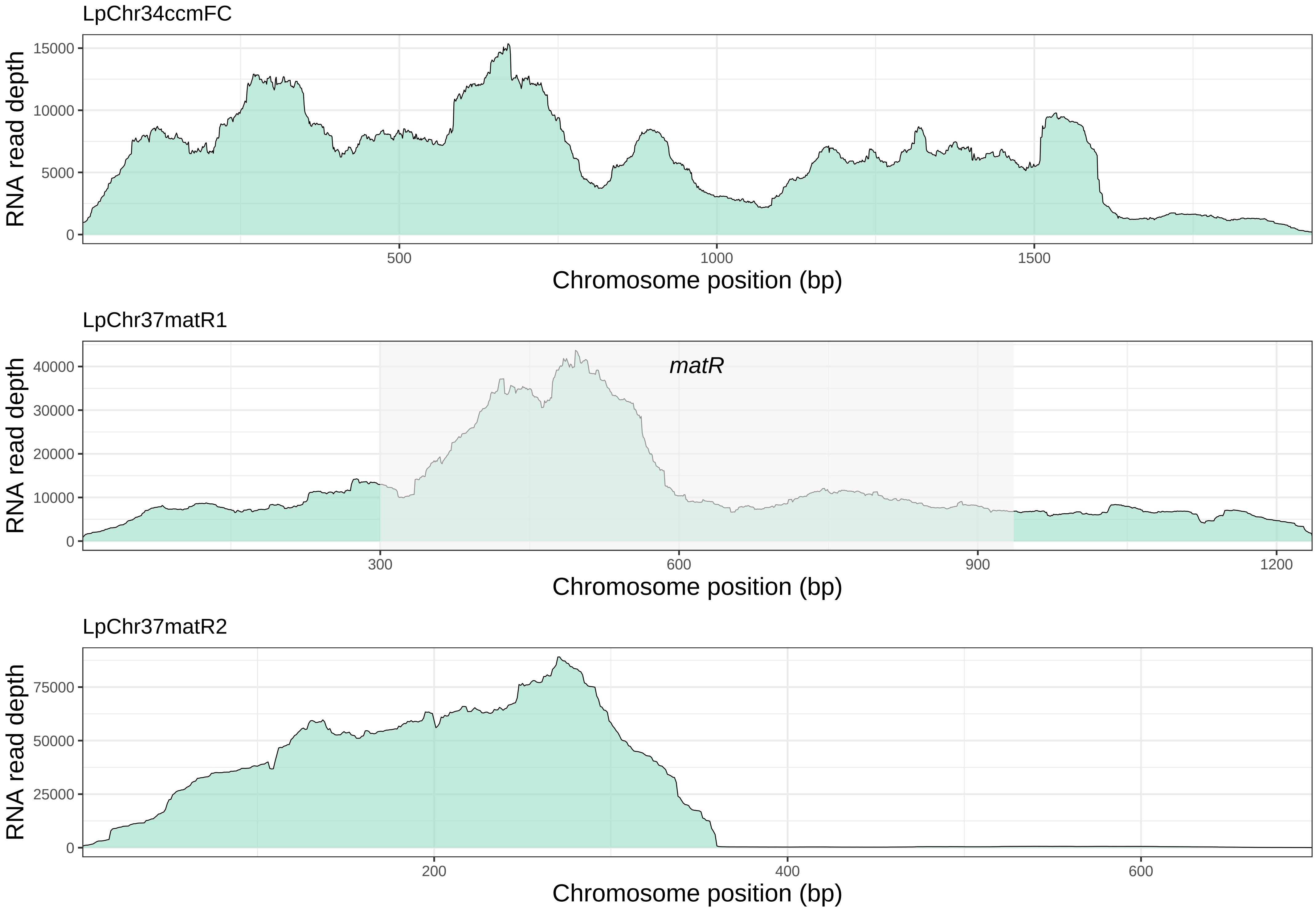
LpChr34ccmFC
RNA read depth
15000
10000
5000
0
500
1000
1500
Chromosome position (bp)
LpChr37matR1
matR
RNA read depth
40000
30000
20000
10000
0
300
600
900
1200
Chromosome position (bp)
LpChr37matR2
RNA read depth
75000
50000
25000
0
200
400
600
Chromosome position (bp)

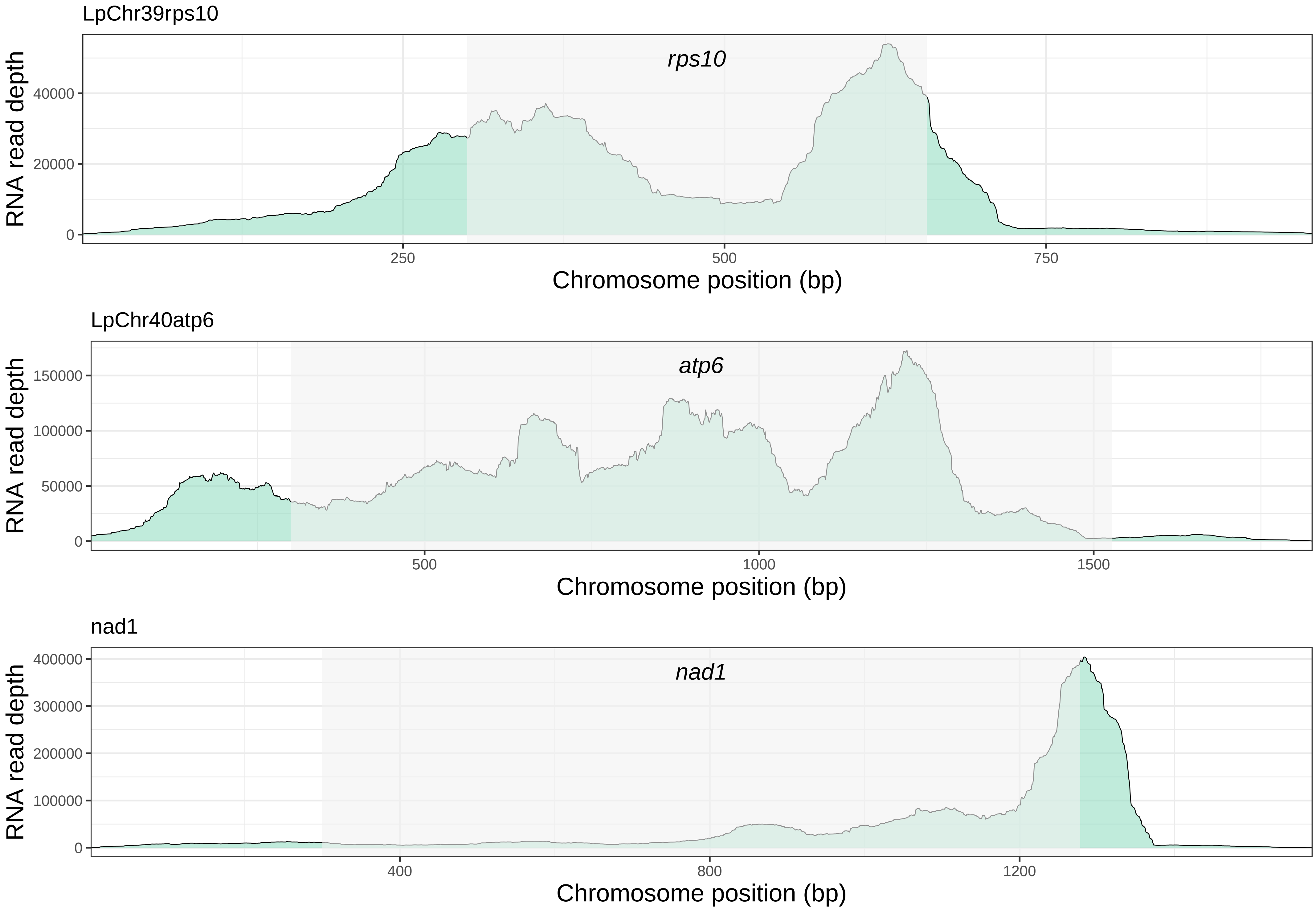

LpChr39rps10
rps10
RNA read depth
40000
20000
0
250
500
750
Chromosome position (bp)
LpChr40atp6
atp6
RNA read depth
150000
100000
50000
0
500
1000
1500
Chromosome position (bp)
nad1
nad1
RNA read depth
400000
300000
200000
100000
0
400
800
1200
Chromosome position (bp)

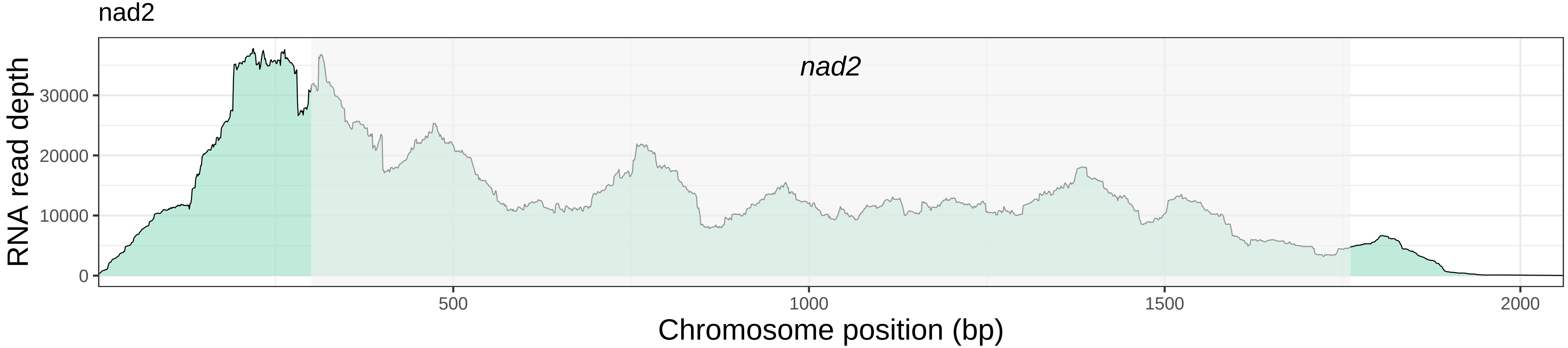
nad2
nad2
RNA read depth
30000
20000
10000
0
500
1000
1500
2000
Chromosome position (bp)

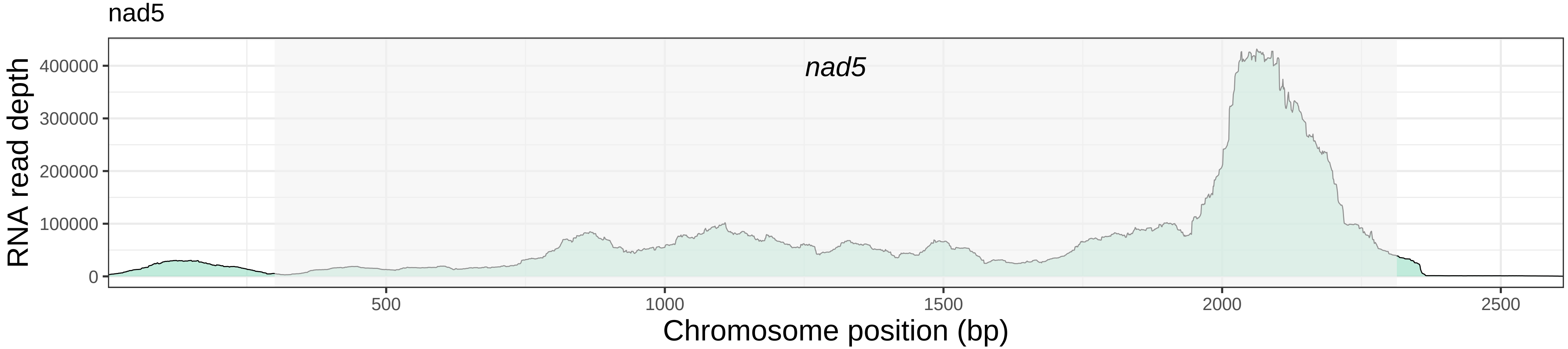
nad5
nad5
RNA read depth
400000
300000
200000
100000
0
500
1000
1500
2000
2500
Chromosome position (bp)

**Figure S7. Boxplots showing C-to-U RNA editing efficiency (%) at non-synonymous sites in native vs. foreign regions for (A) *Lophophytum pyramidale* and (B) *L. mirabile*.** Black line: median; white diamond: mean; *** $p < 0.001$.

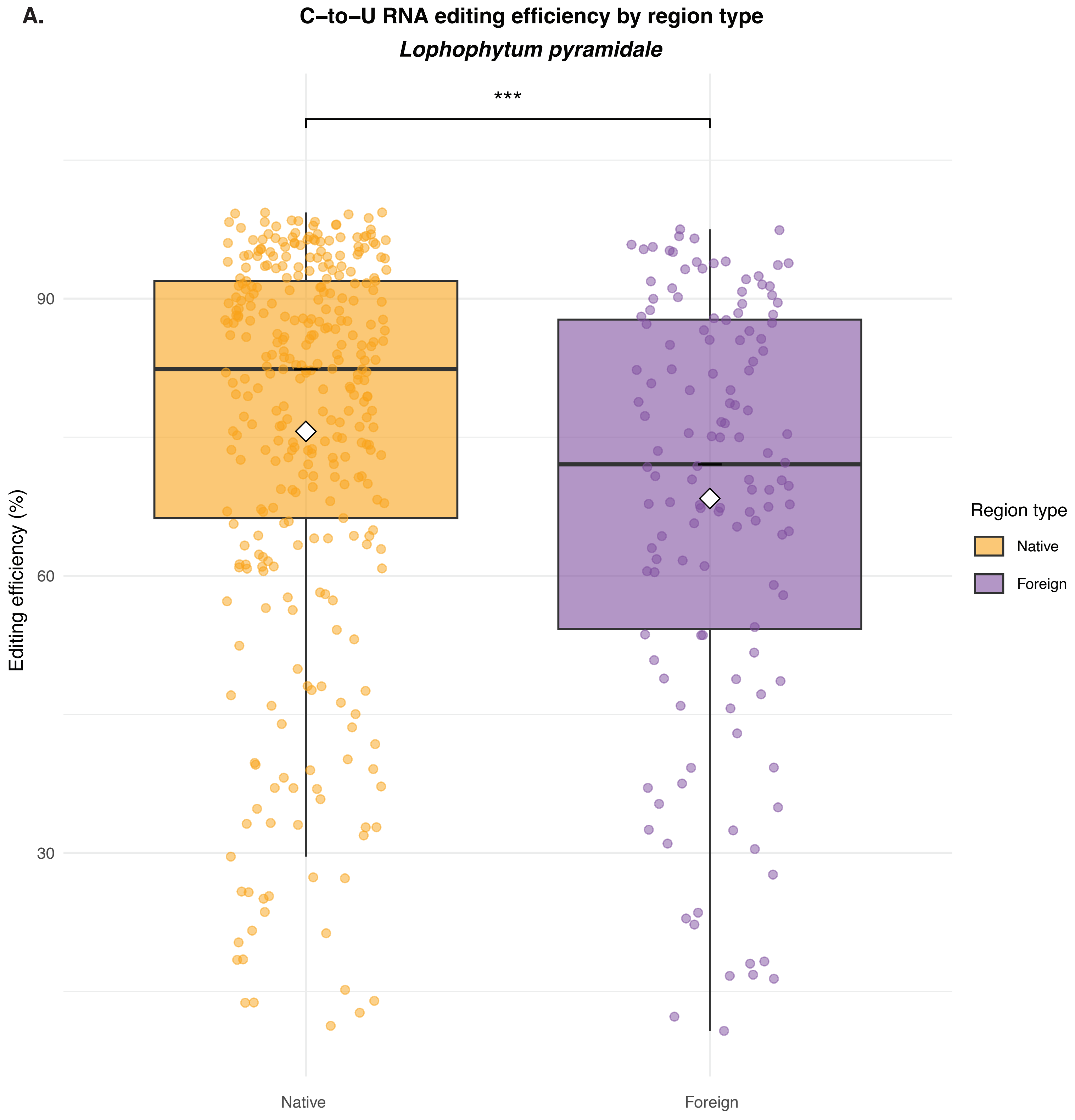


| Species | Region Type | N | Mean (%) | Median (%) | SD | IQR | Min | Max |
|---|---|---|---|---|---|---|---|---|
| *L.pyramidale* | Native | 333 | 75.63 | 82.36 | 21.4 | 25.68 | 11.27 | 99.33 |
| | Foreign | 132 | 68.36 | 72.06 | 22.95 | 33.49 | 10.72 | 97.5 |

Editing efficiency is significantly higher in native regions (Wilcoxon rank sum test, W = 26684, $p = 0.0003167$). Data normality was assessed using the Shapiro–Wilk test. Editing efficiencies in both native and foreign sites deviated significantly from normality (Native: W = 0.859, $p < 2.2 \times 10^{-16}$; Foreign: W = 0.912, $p = 3.112 \times 10^{-7}$), supporting the use of the non-parametric Wilcoxon rank sum test.

**B.**

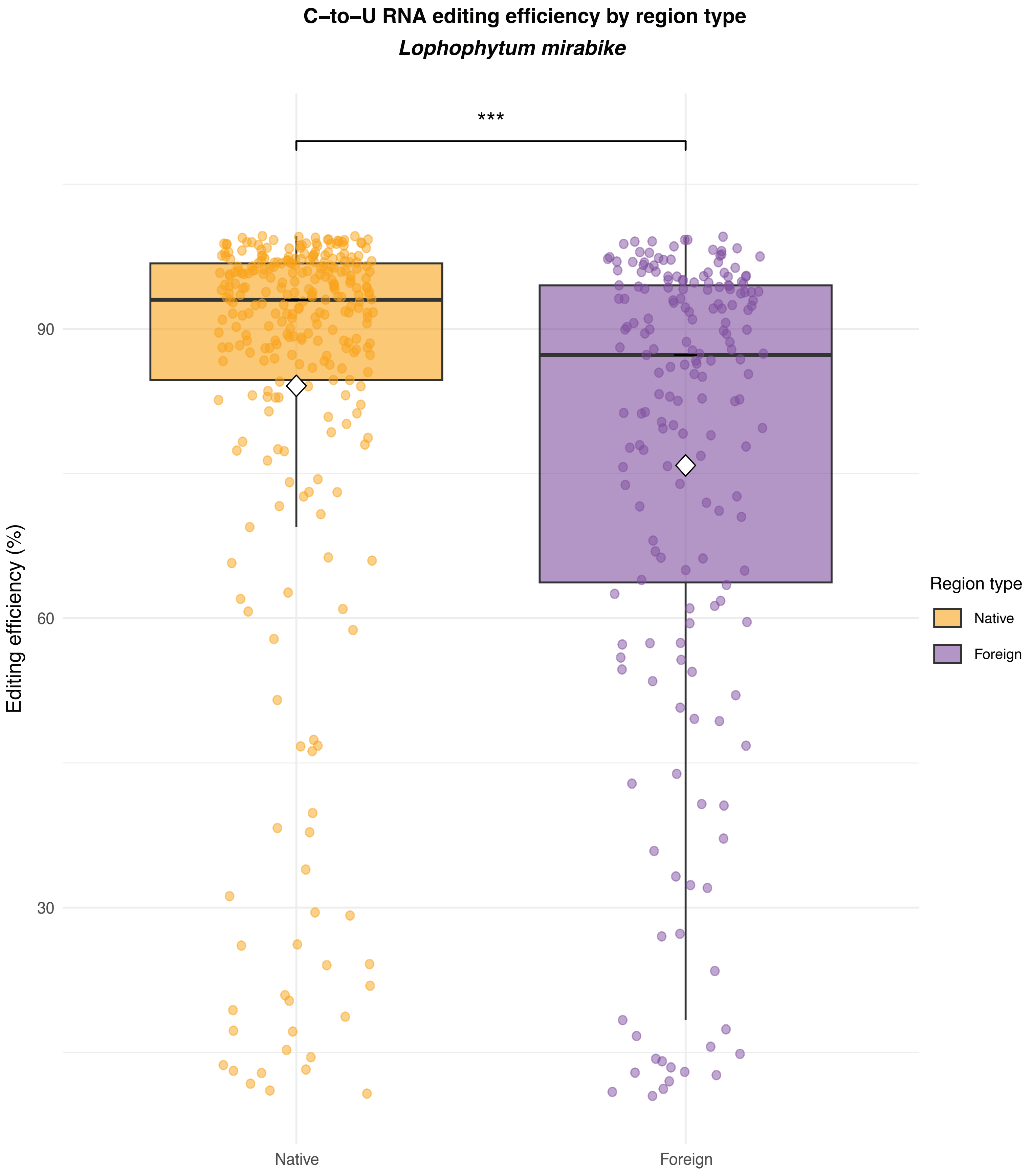


| Species | Region Type | N | Mean (%) | Median (%) | SD | IQR | Min | Max |
|---|---|---|---|---|---|---|---|---|
| *L. mirabile* | Native | 292 | 84.09 | 93.03 | 22.29 | 12.09 | 10.71 | 99.62 |
| | Foreign | 187 | 75.85 | 87.3 | 25.56 | 30.8 | 10.47 | 99.56 |

Editing efficiency is significantly higher in native regions (Wilcoxon rank sum test, W = 34286, $p = 2.3 \times 10^{-6}$). Data normality was assessed using the Shapiro–Wilk test. Editing efficiencies in both native and foreign sites deviated significantly from normality (Native: W = 0.642, $p < 2.2 \times 10^{-16}$; Foreign: W = 0.797, $p = 7.5 \times 10^{-15}$), supporting the use of the non-parametric Wilcoxon rank sum test.

**Figure S8. Maximum Likelihood (ML) phylogenetic analyses of RNA polymerases in *Lophophytum* spp. and *Ombrophytum*.** ML bootstrap support values ≥50% are shown. Scale bars correspond to substitutions per site.The plastid-targeted RpoTp was not found in the transcriptomes of *Lophophytum* spp. and *Ombrophytum*.

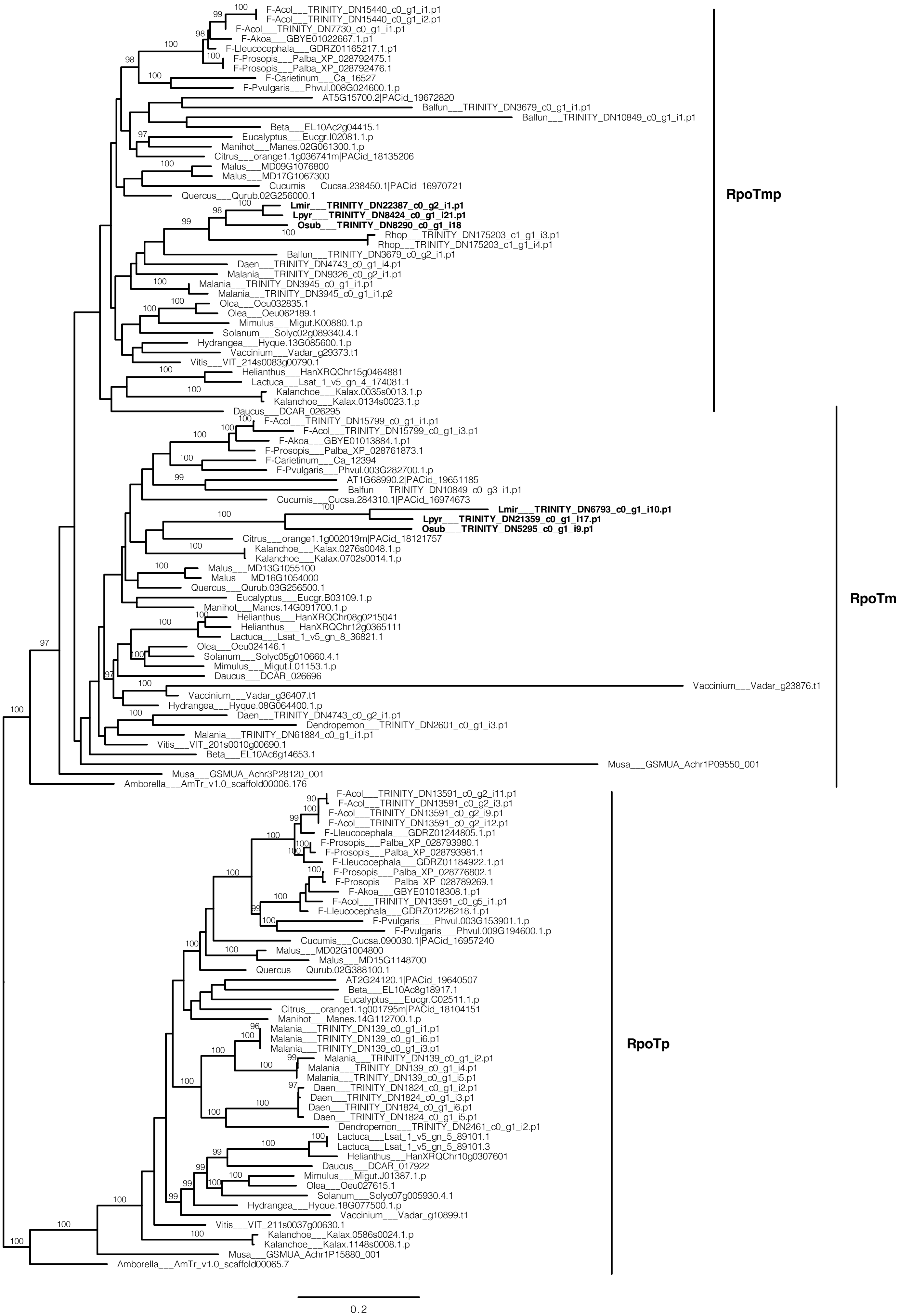

**Figure S9. Evolutionary model proposed for the mitochondrial gene clusters in Balanophoraceae mitochondria.** Red lines indicate losses of clusters or genes within a cluster.

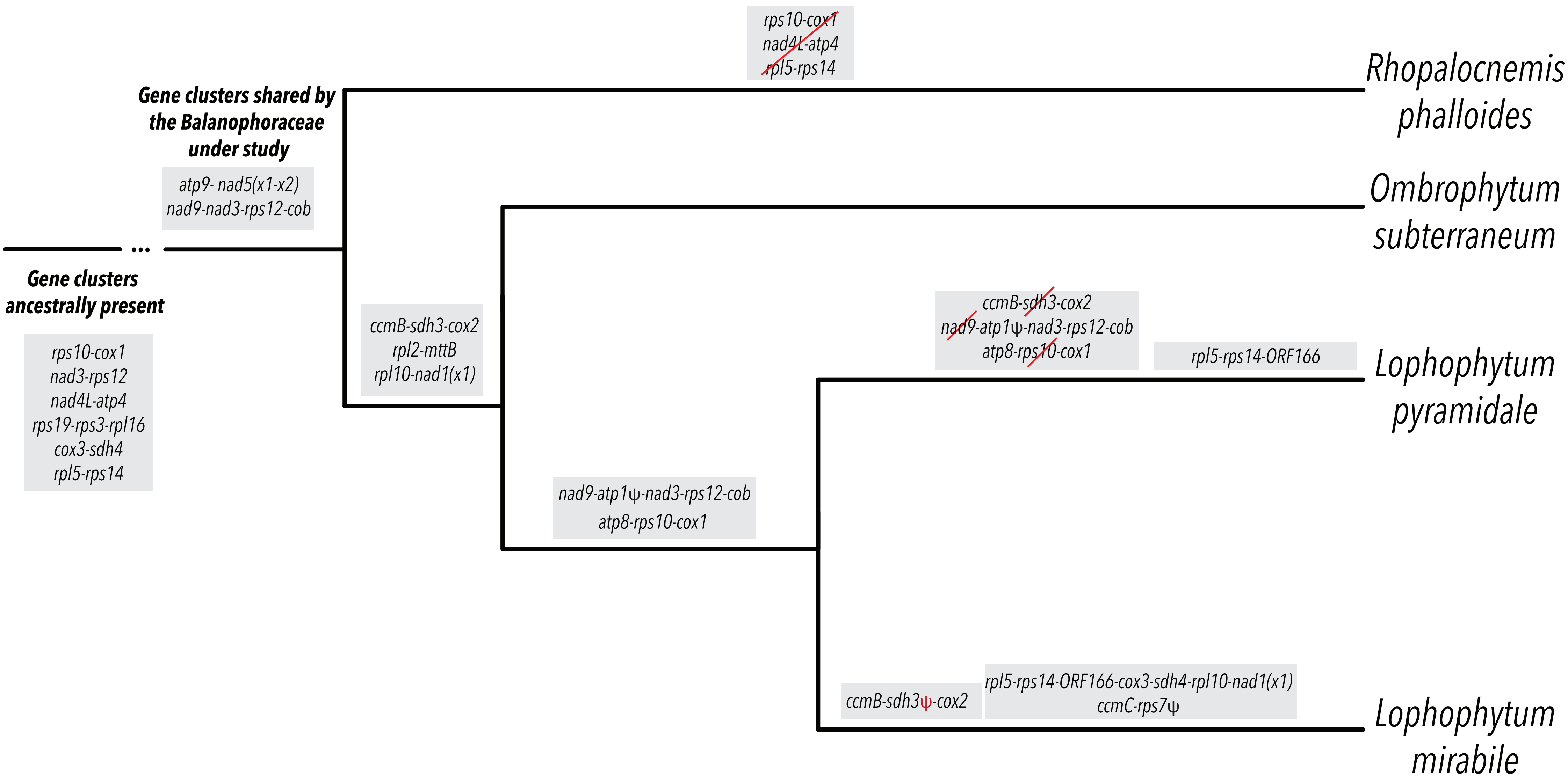

**Figure S10.** Maximum-likelihood (ML) phylogenetic analyses of the detected orthologs in *Lophophytum* spp. and *Ombrophytum*. ML bootstrap support values ≥50% are shown next to nodes. Scale bars correspond to substitutions per site. The orthogroup identifiers corresponding to each gene (as listed in Table S4) are indicated in the upper-right corner of each tree. Color code: *Arabidopsis thaliana* (red), Fabaceae species (blue), Santalales (brown), *Lophophytum mirabile* (orange), *L. pyramidale* (yellow), and *Ombrophytum subterraneum* (pink).

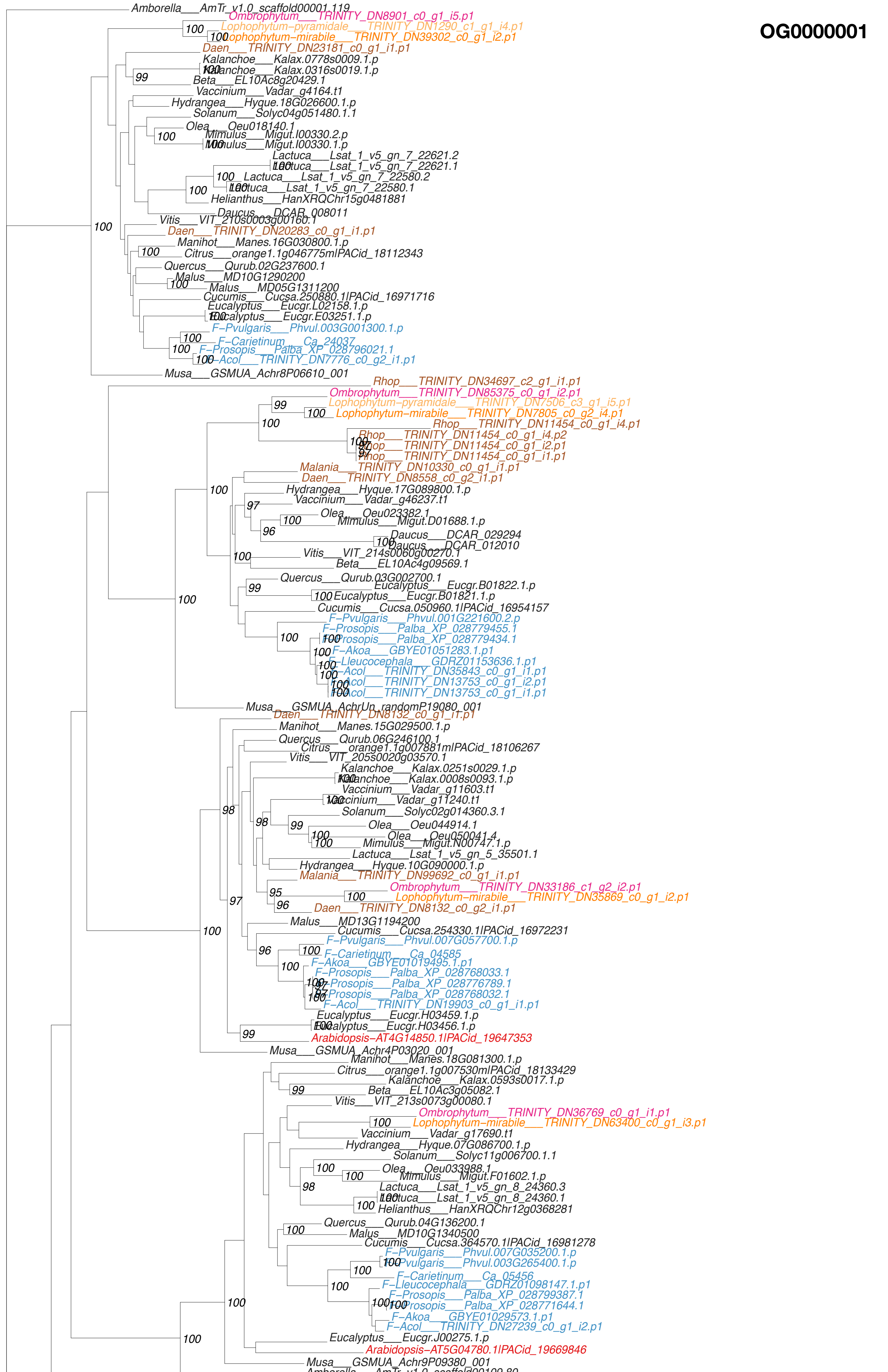

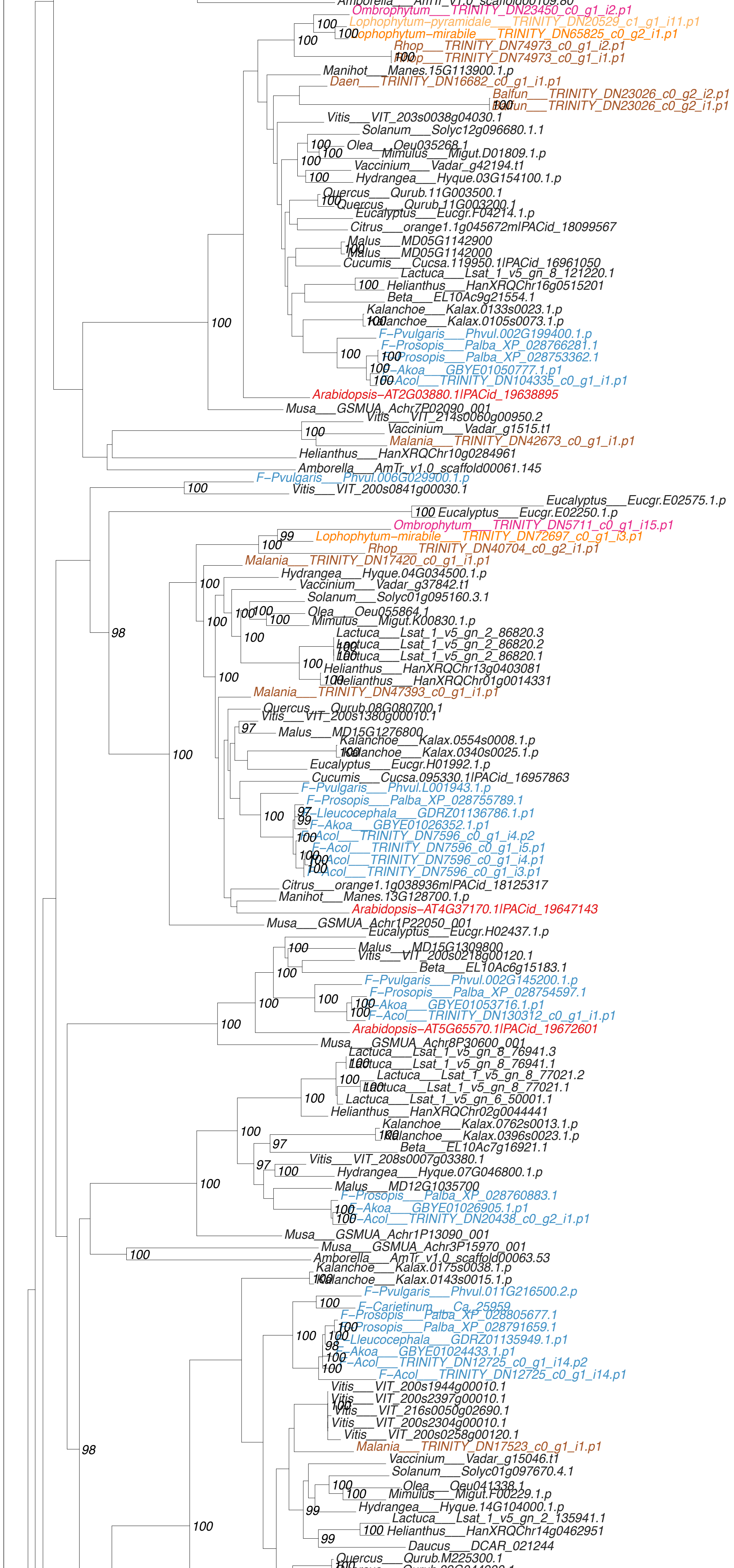

Ombrophytum___TRINITY_DN23450_c0_g1_i2.p1
Lophophytum-pyramidale___TRINITY_DN20529_c1_g1_i11.p1
Lophophytum-mirabile___TRINITY_DN65825_c0_g2_i1.p1
Rhop___TRINITY_DN74973_c0_g1_i2.p1
Rhop___TRINITY_DN74973_c0_g1_i1.p1
Manihot___Manes.15G113900.1.p
Daen___TRINITY_DN16682_c0_g1_i1.p1
Balfun___TRINITY_DN23026_c0_g2_i2.p1
Balfun___TRINITY_DN23026_c0_g2_i1.p1
Vitis___VIT_203s0038g04030.1
Solanum___Solyc12g096680.1.1
Olea___Oeu035268.1
Mimulus___Migut.D01809.1.p
Vaccinium___Vadar_g42194.t1
Hydrangea___Hyque.03G154100.1.p
Quercus___Qurub.11G003500.1
Quercus___Qurub.11G003200.1
Eucalyptus___Eucgr.F04214.1.p
Citrus___orange1.1g045672mIPACid_18099567
Malus___MD05G1142900
Malus___MD05G1142000
Cucumis___Cucsa.119950.1IPACid_16961050
Lactuca___Lsat_1_v5_gn_8_121220.1
Helianthus___HanXRQChr16g0515201
Beta___EL10Ac9g21554.1
Kalanchoe___Kalax.0133s0023.1.p
Kalanchoe___Kalax.0105s0073.1.p
F-Pvulgaris___Phvul.002G199400.1.p
F-Prosopis___Palba_XP_028766281.1
F-Prosopis___Palba_XP_028753362.1
F-Akoa___GBYE01050777.1.p1
F-Acol___TRINITY_DN104335_c0_g1_i1.p1
Arabidopsis-AT2G03880.1IPACid_19638895
Musa___GSMUA_Achr7P02090_001
Vitis___VIT_214s0060g00950.2
Vaccinium___Vadar_g1515.t1
Malania___TRINITY_DN42673_c0_g1_i1.p1
Helianthus___HanXRQChr10g0284961
Amborella___AmTr_v1.0_scaffold00061.145
F-Pvulgaris___Phvul.006G029900.1.p
Vitis___VIT_200s0841g00030.1
Eucalyptus___Eucgr.E02575.1.p
Eucalyptus___Eucgr.E02250.1.p
Ombrophytum___TRINITY_DN5711_c0_g1_i15.p1
Lophophytum-mirabile___TRINITY_DN72697_c0_g1_i3.p1
Rhop___TRINITY_DN40704_c0_g2_i1.p1
Malania___TRINITY_DN17420_c0_g1_i1.p1
Hydrangea___Hyque.04G034500.1.p
Vaccinium___Vadar_g37842.t1
Solanum___Solyc01g095160.3.1
Olea___Oeu055864.1
Mimulus___Migut.K00830.1.p
Lactuca___Lsat_1_v5_gn_2_86820.3
Lactuca___Lsat_1_v5_gn_2_86820.2
Lactuca___Lsat_1_v5_gn_2_86820.1
Helianthus___HanXRQChr13g0403081
Helianthus___HanXRQChr01g0014331
Malania___TRINITY_DN47393_c0_g1_i1.p1
Quercus___Qurub.08G080700.1
Vitis___VIT_200s1380g00010.1
Malus___MD15G1276800
Kalanchoe___Kalax.0554s0008.1.p
Kalanchoe___Kalax.0340s0025.1.p
Eucalyptus___Eucgr.H01992.1.p
Cucumis___Cucsa.095330.1IPACid_16957863
F-Pvulgaris___Phvul.L001943.1.p
F-Prosopis___Palba_XP_028755789.1
F-Lleucocephala___GDRZ01136786.1.p1
F-Akoa___GBYE01026352.1.p1
F-Acol___TRINITY_DN7596_c0_g1_i4.p2
F-Acol___TRINITY_DN7596_c0_g1_i5.p1
F-Acol___TRINITY_DN7596_c0_g1_i4.p1
F-Acol___TRINITY_DN7596_c0_g1_i3.p1
Citrus___orange1.1g038936mIPACid_18125317
Manihot___Manes.13G128700.1.p
Arabidopsis-AT4G37170.1IPACid_19647143
Musa___GSMUA_Achr1P22050_001
Eucalyptus___Eucgr.H02437.1.p
Malus___MD15G1309800
Vitis___VIT_200s0218g00120.1
Beta___EL10Ac6g15183.1
F-Pvulgaris___Phvul.002G145200.1.p
F-Prosopis___Palba_XP_028754597.1
F-Akoa___GBYE01053716.1.p1
F-Acol___TRINITY_DN130312_c0_g1_i1.p1
Arabidopsis-AT5G65570.1IPACid_19672601
Musa___GSMUA_Achr8P30600_001
Lactuca___Lsat_1_v5_gn_8_76941.3
Lactuca___Lsat_1_v5_gn_8_76941.1
Lactuca___Lsat_1_v5_gn_8_77021.2
Lactuca___Lsat_1_v5_gn_8_77021.1
Lactuca___Lsat_1_v5_gn_6_50001.1
Helianthus___HanXRQChr02g0044441
Kalanchoe___Kalax.0762s0013.1.p
Kalanchoe___Kalax.0396s0023.1.p
Beta___EL10Ac7g16921.1
Vitis___VIT_208s0007g03380.1
Hydrangea___Hyque.07G046800.1.p
Malus___MD12G1035700
F-Prosopis___Palba_XP_028760883.1
F-Akoa___GBYE01026905.1.p1
F-Acol___TRINITY_DN20438_c0_g2_i1.p1
Musa___GSMUA_Achr1P13090_001
Musa___GSMUA_Achr3P15970_001
Amborella___AmTr_v1.0_scaffold00063.53
Kalanchoe___Kalax.0175s0038.1.p
Kalanchoe___Kalax.0143s0015.1.p
F-Pvulgaris___Phvul.011G216500.2.p
F-Carietinum___Ca_25959
F-Prosopis___Palba_XP_028805677.1
F-Prosopis___Palba_XP_028791659.1
F-Lleucocephala___GDRZ01135949.1.p1
F-Akoa___GBYE01024433.1.p1
F-Acol___TRINITY_DN12725_c0_g1_i14.p2
F-Acol___TRINITY_DN12725_c0_g1_i14.p1
Vitis___VIT_200s1944g00010.1
Vitis___VIT_200s2397g00010.1
Vitis___VIT_216s0050g02690.1
Vitis___VIT_200s2304g00010.1
Vitis___VIT_200s0258g00120.1
Malania___TRINITY_DN17523_c0_g1_i1.p1
Vaccinium___Vadar_g15046.t1
Solanum___Solyc01g097670.4.1
Olea___Oeu041338.1
Mimulus___Migut.F00229.1.p
Hydrangea___Hyque.14G104000.1.p
Lactuca___Lsat_1_v5_gn_2_135941.1
Helianthus___HanXRQChr14g0462951
Daucus___DCAR_021244
Quercus___Qurub.M225300.1

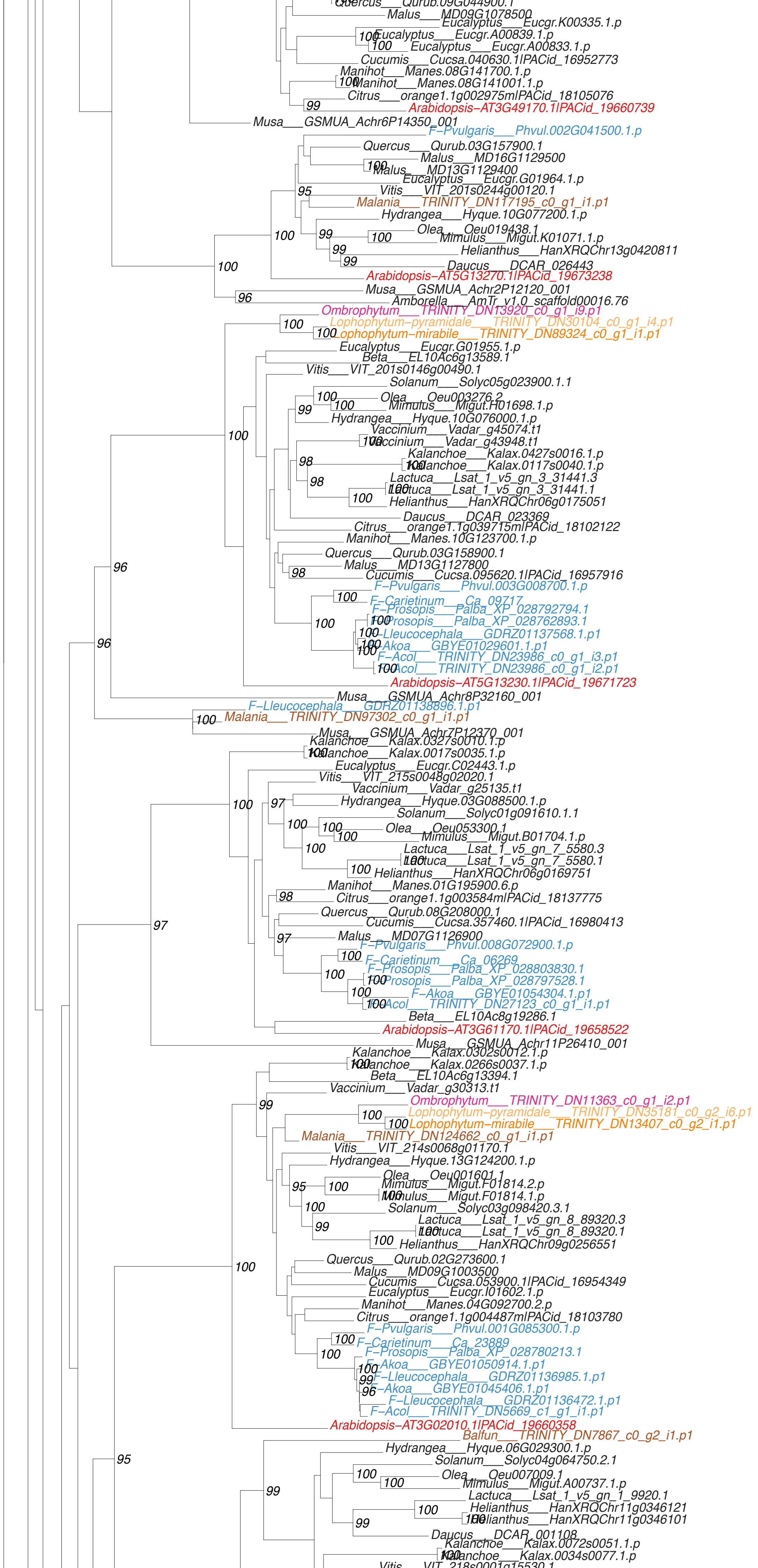
Quercus___Qurub.09G044900.1
Malus___MD09G1078500
Eucalyptus___Eucgr.K00335.1.p
100 Eucalyptus___Eucgr.A00839.1.p
100 Eucalyptus___Eucgr.A00833.1.p
Cucumis___Cucsa.040630.1lPACid_16952773
Manihot___Manes.08G141700.1.p
100 Manihot___Manes.08G141001.1.p
Citrus___orange1.1g002975mlPACid_18105076
99 Arabidopsis−AT3G49170.1lPACid_19660739
Musa___GSMUA_Achr6P14350_001
F−Pvulgaris___Phvul.002G041500.1.p
Quercus___Qurub.03G157900.1
Malus___MD16G1129500
100 Malus___MD13G1129400
Eucalyptus___Eucgr.G01964.1.p
95 Vitis___VIT_201s0244g00120.1
Malania___TRINITY_DN117195_c0_g1_i1.p1
100 Hydrangea___Hyque.10G077200.1.p
Olea___Oeu019438.1
99 100 Mimulus___Migut.K01071.1.p
99 Helianthus___HanXRQChr13g0420811
100 99 Daucus___DCAR_026443
Arabidopsis−AT5G13270.1lPACid_19673238
Musa___GSMUA_Achr2P12120_001
96 Amborella___AmTr_v1.0_scaffold00016.76
Ombrophytum___TRINITY_DN13920_c0_g1_i9.p1
100 Lophophytum−pyramidale___TRINITY_DN30104_c0_g1_i4.p1
100 Lophophytum−mirabile___TRINITY_DN89324_c0_g1_i1.p1
Eucalyptus___Eucgr.G01955.1.p
Beta___EL10Ac6g13589.1
Vitis___VIT_201s0146g00490.1
Solanum___Solyc05g023900.1.1
100 Olea___Oeu003276.2
100 Mimulus___Migut.H01698.1.p
99 Hydrangea___Hyque.10G076000.1.p
Vaccinium___Vadar_g45074.t1
100 Vaccinium___Vadar_g43948.t1
Kalanchoe___Kalax.0427s0016.1.p
98 100 Kalanchoe___Kalax.0117s0040.1.p
Lactuca___Lsat_1_v5_gn_3_31441.3
98 100 Lactuca___Lsat_1_v5_gn_3_31441.1
100 Helianthus___HanXRQChr06g0175051
Daucus___DCAR_023369
Citrus___orange1.1g039715mlPACid_18102122
Manihot___Manes.10G123700.1.p
Quercus___Qurub.03G158900.1
Malus___MD13G1127800
98 Cucumis___Cucsa.095620.1lPACid_16957916
F−Pvulgaris___Phvul.003G008700.1.p
100 F−Carietinum___Ca_09717
F−Prosopis___Palba_XP_028792794.1
100 F−Prosopis___Palba_XP_028762893.1
100 F−Lleucocephala___GDRZ01137568.1.p1
100 F−Akoa___GBYE01029601.1.p1
100 F−Acol___TRINITY_DN23986_c0_g1_i3.p1
100 F−Acol___TRINITY_DN23986_c0_g1_i2.p1
Arabidopsis−AT5G13230.1lPACid_19671723
96
Musa___GSMUA_Achr8P32160_001
F−Lleucocephala___GDRZ01138896.1.p1
100 Malania___TRINITY_DN97302_c0_g1_i1.p1
96
Musa___GSMUA_Achr7P12370_001
Kalanchoe___Kalax.0327s0010.1.p
100 Kalanchoe___Kalax.0017s0035.1.p
Eucalyptus___Eucgr.C02443.1.p
Vitis___VIT_215s0048g02020.1
Vaccinium___Vadar_g25135.t1
97 Hydrangea___Hyque.03G088500.1.p
Solanum___Solyc01g091610.1.1
100 100 Olea___Oeu053300.1
100 Mimulus___Migut.B01704.1.p
100 Lactuca___Lsat_1_v5_gn_7_5580.3
100 Lactuca___Lsat_1_v5_gn_7_5580.1
100 Helianthus___HanXRQChr06g0169751
Manihot___Manes.01G195900.6.p
98 Citrus___orange1.1g003584mlPACid_18137775
Quercus___Qurub.08G208000.1
Cucumis___Cucsa.357460.1lPACid_16980413
97 Malus___MD07G1126900
F−Pvulgaris___Phvul.008G072900.1.p
100 F−Carietinum___Ca_06269
F−Prosopis___Palba_XP_028803830.1
100 F−Prosopis___Palba_XP_028797528.1
100 F−Akoa___GBYE01054304.1.p1
100 F−Acol___TRINITY_DN27123_c0_g1_i1.p1
Beta___EL10Ac8g19286.1
Arabidopsis−AT3G61170.1lPACid_19658522
97
Musa___GSMUA_Achr11P26410_001
Kalanchoe___Kalax.0302s0012.1.p
100 Kalanchoe___Kalax.0266s0037.1.p
Beta___EL10Ac6g13394.1
Vaccinium___Vadar_g30313.t1
99 Ombrophytum___TRINITY_DN11363_c0_g1_i2.p1
100 Lophophytum−pyramidale___TRINITY_DN35181_c0_g2_i6.p1
100 Lophophytum−mirabile___TRINITY_DN13407_c0_g2_i1.p1
Malania___TRINITY_DN124662_c0_g1_i1.p1
Vitis___VIT_214s0068g01170.1
Hydrangea___Hyque.13G124200.1.p
Olea___Oeu001601.1
95 100 Mimulus___Migut.F01814.2.p
100 Mimulus___Migut.F01814.1.p
100 Solanum___Solyc03g098420.3.1
Lactuca___Lsat_1_v5_gn_8_89320.3
99 100 Lactuca___Lsat_1_v5_gn_8_89320.1
100 Helianthus___HanXRQChr09g0256551
Quercus___Qurub.02G273600.1
Malus___MD09G1003500
Cucumis___Cucsa.053900.1lPACid_16954349
Eucalyptus___Eucgr.I01602.1.p
Manihot___Manes.04G092700.2.p
Citrus___orange1.1g004487mlPACid_18103780
F−Pvulgaris___Phvul.001G085300.1.p
100 F−Carietinum___Ca_23889
100 F−Prosopis___Palba_XP_028780213.1
F−Akoa___GBYE01050914.1.p1
100 F−Lleucocephala___GDRZ01136985.1.p1
99 F−Akoa___GBYE01045406.1.p1
96 F−Lleucocephala___GDRZ01136472.1.p1
F−Acol___TRINITY_DN5669_c1_g1_i1.p1
100 Arabidopsis−AT3G02010.1lPACid_19660358
Balfun___TRINITY_DN7867_c0_g2_i1.p1
95
Hydrangea___Hyque.06G029300.1.p
Solanum___Solyc04g064750.2.1
100 Olea___Oeu007009.1
100 Mimulus___Migut.A00737.1.p
99 Lactuca___Lsat_1_v5_gn_1_9920.1
100 Helianthus___HanXRQChr11g0346121
99 100 Helianthus___HanXRQChr11g0346101
Daucus___DCAR_001108
Kalanchoe___Kalax.0072s0051.1.p
100 Kalanchoe___Kalax.0034s0077.1.p

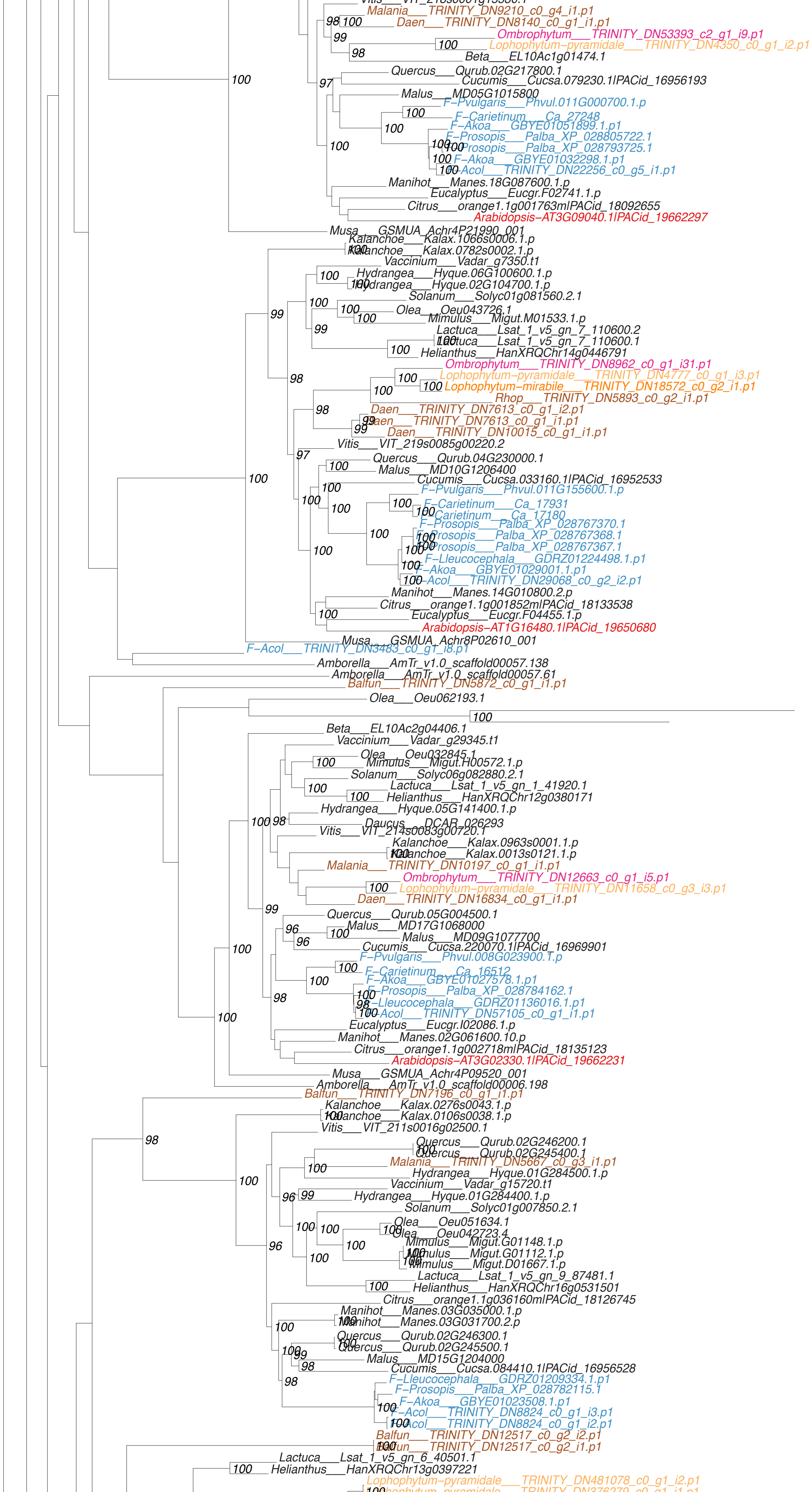

Malania___TRINITY_DN9210_c0_g4_i1.p1
Daen___TRINITY_DN8140_c0_g1_i1.p1
Ombrophytum___TRINITY_DN53393_c2_g1_i9.p1
Lophophytum-pyramidale___TRINITY_DN4350_c0_g1_i2.p1
Beta___EL10Ac1g01474.1
Quercus___Qurub.02G217800.1
Cucumis___Cucsa.079230.1lPACid_16956193
Malus___MD05G1015800
F-Pvulgaris___Phvul.011G000700.1.p
F-Carietinum___Ca_27248
F-Akoa___GBYE01051899.1.p1
F-Prosopis___Palba_XP_028805722.1
F-Prosopis___Palba_XP_028793725.1
F-Akoa___GBYE01032298.1.p1
F-Acol___TRINITY_DN22256_c0_g5_i1.p1
Manihot___Manes.18G087600.1.p
Eucalyptus___Eucgr.F02741.1.p
Citrus___orange1.1g001763mlPACid_18092655
Arabidopsis-AT3G09040.1lPACid_19662297
Musa___GSMUA_Achr4P21990_001
Kalanchoe___Kalax.1066s0006.1.p
Kalanchoe___Kalax.0782s0002.1.p
Vaccinium___Vadar_g7350.t1
Hydrangea___Hyque.06G100600.1.p
Hydrangea___Hyque.02G104700.1.p
Solanum___Solyc01g081560.2.1
Olea___Oeu043726.1
Mimulus___Migut.M01533.1.p
Lactuca___Lsat_1_v5_gn_7_110600.2
Lactuca___Lsat_1_v5_gn_7_110600.1
Helianthus___HanXRQChr14g0446791
Ombrophytum___TRINITY_DN8962_c0_g1_i31.p1
Lophophytum-pyramidale___TRINITY_DN4777_c0_g1_i3.p1
Lophophytum-mirabile___TRINITY_DN18572_c0_g2_i1.p1
Rhop___TRINITY_DN5893_c0_g2_i1.p1
Daen___TRINITY_DN7613_c0_g1_i2.p1
Daen___TRINITY_DN7613_c0_g1_i1.p1
Daen___TRINITY_DN10015_c0_g1_i1.p1
Vitis___VIT_219s0085g00220.2
Quercus___Qurub.04G230000.1
Malus___MD10G1206400
Cucumis___Cucsa.033160.1lPACid_16952533
F-Pvulgaris___Phvul.011G155600.1.p
F-Carietinum___Ca_17931
F-Carietinum___Ca_17180
F-Prosopis___Palba_XP_028767370.1
F-Prosopis___Palba_XP_028767368.1
F-Prosopis___Palba_XP_028767367.1
F-Lleucocephala___GDRZ01224498.1.p1
F-Akoa___GBYE01029001.1.p1
F-Acol___TRINITY_DN29068_c0_g2_i2.p1
Manihot___Manes.14G010800.2.p
Citrus___orange1.1g001852mlPACid_18133538
Eucalyptus___Eucgr.F04455.1.p
Arabidopsis-AT1G16480.1lPACid_19650680
Musa___GSMUA_Achr8P02610_001
F-Acol___TRINITY_DN3483_c0_g1_i8.p1
Amborella___AmTr_v1.0_scaffold00057.138
Amborella___AmTr_v1.0_scaffold00057.61
Balfun___TRINITY_DN5872_c0_g1_i1.p1
Olea___Oeu062193.1
Beta___EL10Ac2g04406.1
Vaccinium___Vadar_g29345.t1
Olea___Oeu032845.1
Mimulus___Migut.H00572.1.p
Solanum___Solyc06g082880.2.1
Lactuca___Lsat_1_v5_gn_1_41920.1
Helianthus___HanXRQChr12g0380171
Hydrangea___Hyque.05G141400.1.p
Daucus___DCAR_026293
Vitis___VIT_214s0083g00720.1
Kalanchoe___Kalax.0963s0001.1.p
Kalanchoe___Kalax.0013s0121.1.p
Malania___TRINITY_DN10197_c0_g1_i1.p1
Ombrophytum___TRINITY_DN12663_c0_g1_i5.p1
Lophophytum-pyramidale___TRINITY_DN11658_c0_g3_i3.p1
Daen___TRINITY_DN16834_c0_g1_i1.p1
Quercus___Qurub.05G004500.1
Malus___MD17G1068000
Malus___MD09G1077700
Cucumis___Cucsa.220070.1lPACid_16969901
F-Pvulgaris___Phvul.008G023900.1.p
F-Carietinum___Ca_16512
F-Akoa___GBYE01027578.1.p1
F-Prosopis___Palba_XP_028784162.1
F-Lleucocephala___GDRZ01136016.1.p1
F-Acol___TRINITY_DN57105_c0_g1_i1.p1
Eucalyptus___Eucgr.I02086.1.p
Manihot___Manes.02G061600.10.p
Citrus___orange1.1g002718mlPACid_18135123
Arabidopsis-AT3G02330.1lPACid_19662231
Musa___GSMUA_Achr4P09520_001
Amborella___AmTr_v1.0_scaffold00006.198
Balfun___TRINITY_DN7196_c0_g1_i1.p1
Kalanchoe___Kalax.0276s0043.1.p
Kalanchoe___Kalax.0106s0038.1.p
Vitis___VIT_211s0016g02500.1
Quercus___Qurub.02G246200.1
Quercus___Qurub.02G245400.1
Malania___TRINITY_DN5667_c0_g3_i1.p1
Hydrangea___Hyque.01G284500.1.p
Vaccinium___Vadar_g15720.t1
Hydrangea___Hyque.01G284400.1.p
Solanum___Solyc01g007850.2.1
Olea___Oeu051634.1
Olea___Oeu042723.4
Mimulus___Migut.G01148.1.p
Mimulus___Migut.G01112.1.p
Mimulus___Migut.D01667.1.p
Lactuca___Lsat_1_v5_gn_9_87481.1
Helianthus___HanXRQChr16g0531501
Citrus___orange1.1g036160mlPACid_18126745
Manihot___Manes.03G035000.1.p
Manihot___Manes.03G031700.2.p
Quercus___Qurub.02G246300.1
Quercus___Qurub.02G245500.1
Malus___MD15G1204000
Cucumis___Cucsa.084410.1lPACid_16956528
F-Lleucocephala___GDRZ01209334.1.p1
F-Prosopis___Palba_XP_028782115.1
F-Akoa___GBYE01023508.1.p1
F-Acol___TRINITY_DN8824_c0_g1_i3.p1
F-Acol___TRINITY_DN8824_c0_g1_i2.p1
Balfun___TRINITY_DN12517_c0_g2_i2.p1
Balfun___TRINITY_DN12517_c0_g2_i1.p1
Lactuca___Lsat_1_v5_gn_6_40501.1
Helianthus___HanXRQChr13g0397221
Lophophytum-pyramidale___TRINITY_DN481078_c0_g1_i2.p1

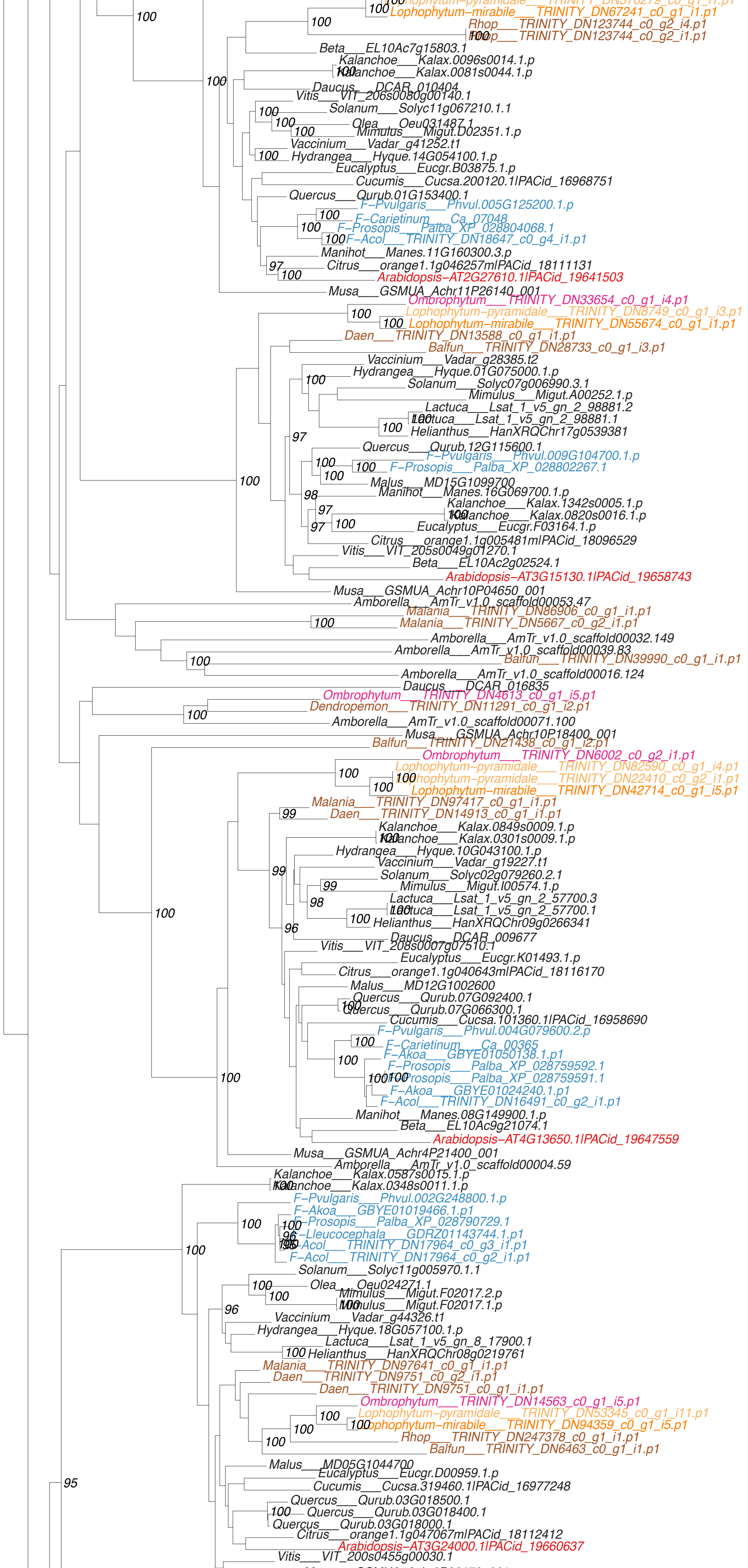
Lophophytum-mirabile___TRINITY_DN67241_c0_g1_i1.p1
Rhop___TRINITY_DN123744_c0_g2_i4.p1
Rhop___TRINITY_DN123744_c0_g2_i1.p1
Beta___EL10Ac7g15803.1
Kalanchoe___Kalax.0096s0014.1.p
Kalanchoe___Kalax.0081s0044.1.p
Daucus___DCAR_010404
Vitis___VIT_206s0080g00140.1
Solanum___Solyc11g067210.1.1
Olea___Oeu031487.1
Mimulus___Migut.D02351.1.p
Vaccinium___Vadar_g41252.t1
Hydrangea___Hyque.14G054100.1.p
Eucalyptus___Eucgr.B03875.1.p
Cucumis___Cucsa.200120.1IPACid_16968751
Quercus___Qurub.01G153400.1
F-Pvulgaris___Phvul.005G125200.1.p
F-Carietinum___Ca_07048
F-Prosopis___Palba_XP_028804068.1
F-Acol___TRINITY_DN18647_c0_g4_i1.p1
Manihot___Manes.11G160300.3.p
Citrus___orange1.1g046257mIPACid_18111131
Arabidopsis-AT2G27610.1IPACid_19641503
Musa___GSMUA_Achr11P26140_001
Ombrophytum___TRINITY_DN33654_c0_g1_i4.p1
Lophophytum-pyramidale___TRINITY_DN8749_c0_g1_i3.p1
Lophophytum-mirabile___TRINITY_DN55674_c0_g1_i1.p1
Daen___TRINITY_DN13588_c0_g1_i1.p1
Balfun___TRINITY_DN28733_c0_g1_i3.p1
Vaccinium___Vadar_g28385.t2
Hydrangea___Hyque.01G075000.1.p
Solanum___Solyc07g006990.3.1
Mimulus___Migut.A00252.1.p
Lactuca___Lsat_1_v5_gn_2_98881.2
Lactuca___Lsat_1_v5_gn_2_98881.1
Helianthus___HanXRQChr17g0539381
Quercus___Qurub.12G115600.1
F-Pvulgaris___Phvul.009G104700.1.p
F-Prosopis___Palba_XP_028802267.1
Malus___MD15G1099700
Manihot___Manes.16G069700.1.p
Kalanchoe___Kalax.1342s0005.1.p
Kalanchoe___Kalax.0820s0016.1.p
Eucalyptus___Eucgr.F03164.1.p
Citrus___orange1.1g005481mIPACid_18096529
Vitis___VIT_205s0049g01270.1
Beta___EL10Ac2g02524.1
Arabidopsis-AT3G15130.1IPACid_19658743
Musa___GSMUA_Achr10P04650_001
Amborella___AmTr_v1.0_scaffold00053.47
Malania___TRINITY_DN86906_c0_g1_i1.p1
Malania___TRINITY_DN5667_c0_g2_i1.p1
Amborella___AmTr_v1.0_scaffold00032.149
Amborella___AmTr_v1.0_scaffold00039.83
Balfun___TRINITY_DN39990_c0_g1_i1.p1
Amborella___AmTr_v1.0_scaffold00016.124
Daucus___DCAR_016835
Ombrophytum___TRINITY_DN4613_c0_g1_i5.p1
Dendropemon___TRINITY_DN11291_c0_g1_i2.p1
Amborella___AmTr_v1.0_scaffold00071.100
Musa___GSMUA_Achr10P18400_001
Balfun___TRINITY_DN21438_c0_g1_i2.p1
Ombrophytum___TRINITY_DN6002_c0_g2_i1.p1
Lophophytum-pyramidale___TRINITY_DN82590_c0_g1_i4.p1
Lophophytum-pyramidale___TRINITY_DN22410_c0_g2_i1.p1
Lophophytum-mirabile___TRINITY_DN42714_c0_g1_i5.p1
Malania___TRINITY_DN97417_c0_g1_i1.p1
Daen___TRINITY_DN14913_c0_g1_i1.p1
Kalanchoe___Kalax.0849s0009.1.p
Kalanchoe___Kalax.0301s0009.1.p
Hydrangea___Hyque.10G043100.1.p
Vaccinium___Vadar_g19227.t1
Solanum___Solyc02g079260.2.1
Mimulus___Migut.I00574.1.p
Lactuca___Lsat_1_v5_gn_2_57700.3
Lactuca___Lsat_1_v5_gn_2_57700.1
Helianthus___HanXRQChr09g0266341
Daucus___DCAR_009677
Vitis___VIT_208s0007g07510.1
Eucalyptus___Eucgr.K01493.1.p
Citrus___orange1.1g040643mIPACid_18116170
Malus___MD12G1002600
Quercus___Qurub.07G092400.1
Quercus___Qurub.07G066300.1
Cucumis___Cucsa.101360.1IPACid_16958690
F-Pvulgaris___Phvul.004G079600.2.p
F-Carietinum___Ca_00365
F-Akoa___GBYE01050138.1.p1
F-Prosopis___Palba_XP_028759592.1
F-Prosopis___Palba_XP_028759591.1
F-Akoa___GBYE01024240.1.p1
F-Acol___TRINITY_DN16491_c0_g2_i1.p1
Manihot___Manes.08G149900.1.p
Beta___EL10Ac9g21074.1
Arabidopsis-AT4G13650.1IPACid_19647559
Musa___GSMUA_Achr4P21400_001
Amborella___AmTr_v1.0_scaffold00004.59
Kalanchoe___Kalax.0587s0015.1.p
Kalanchoe___Kalax.0348s0011.1.p
F-Pvulgaris___Phvul.002G248800.1.p
F-Akoa___GBYE01019466.1.p1
F-Prosopis___Palba_XP_028790729.1
F-Lleucocephala___GDRZ01143744.1.p1
F-Acol___TRINITY_DN17964_c0_g3_i1.p1
F-Acol___TRINITY_DN17964_c0_g2_i1.p1
Solanum___Solyc11g005970.1.1
Olea___Oeu024271.1
Mimulus___Migut.F02017.2.p
Mimulus___Migut.F02017.1.p
Vaccinium___Vadar_g44326.t1
Hydrangea___Hyque.18G057100.1.p
Lactuca___Lsat_1_v5_gn_8_17900.1
Helianthus___HanXRQChr08g0219761
Malania___TRINITY_DN97641_c0_g1_i1.p1
Daen___TRINITY_DN9751_c0_g2_i1.p1
Daen___TRINITY_DN9751_c0_g1_i1.p1
Ombrophytum___TRINITY_DN14563_c0_g1_i5.p1
Lophophytum-pyramidale___TRINITY_DN53345_c0_g1_i11.p1
Lophophytum-mirabile___TRINITY_DN94359_c0_g1_i5.p1
Rhop___TRINITY_DN247378_c0_g1_i1.p1
Balfun___TRINITY_DN6463_c0_g1_i1.p1
Malus___MD05G1044700
Eucalyptus___Eucgr.D00959.1.p
Cucumis___Cucsa.319460.1IPACid_16977248
Quercus___Qurub.03G018500.1
Quercus___Qurub.03G018400.1
Quercus___Qurub.03G018000.1
Citrus___orange1.1g047067mIPACid_18112412
Arabidopsis-AT3G24000.1IPACid_19660637
Vitis___VIT_200s0455g00030.1

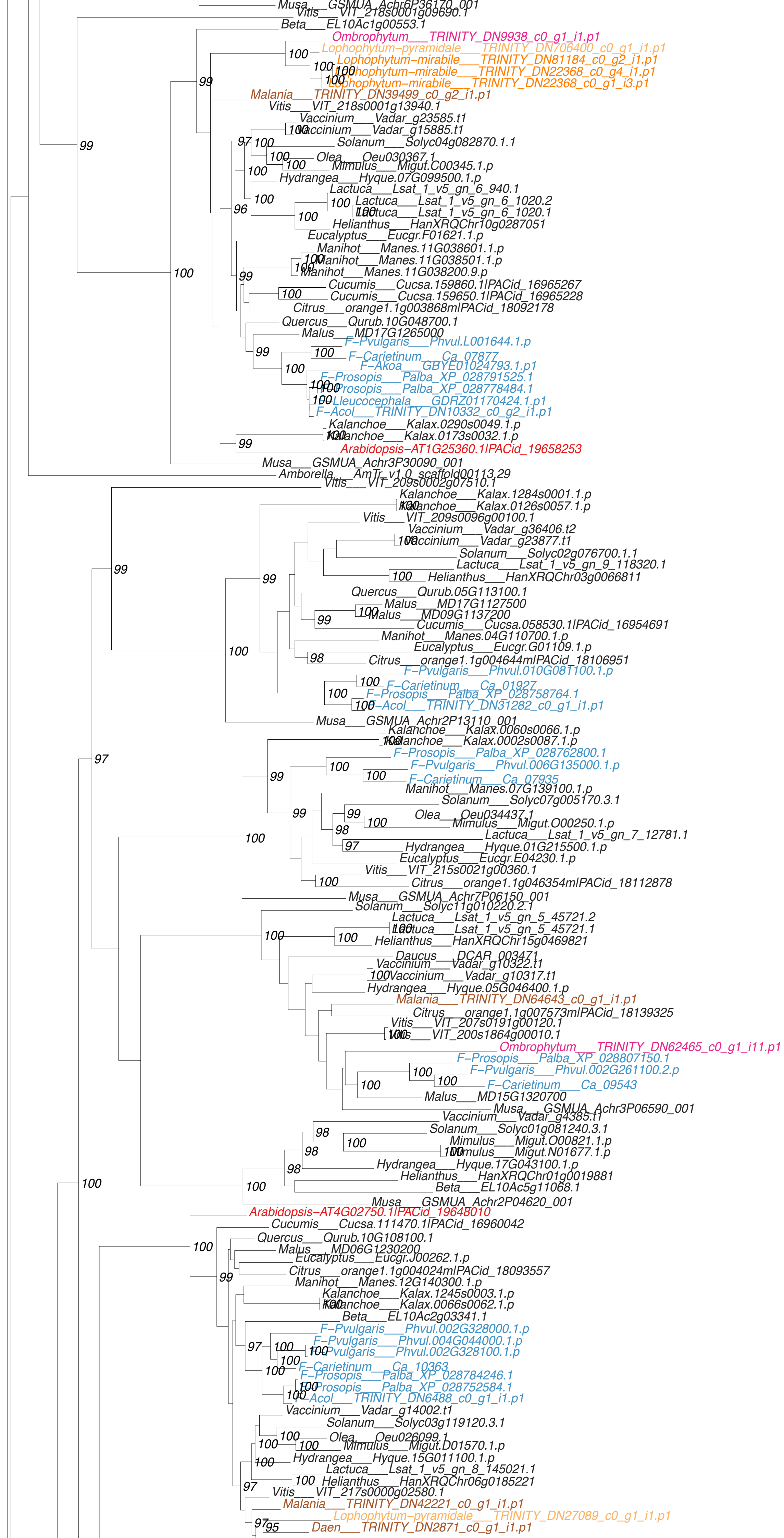
Musa___GSMUA_Achr6P36170_001
Vitis___VIT_218s0001g09690.1
Beta___EL10Ac1g00553.1
Ombrophytum___TRINITY_DN9938_c0_g1_i1.p1
Lophophytum–pyramidale___TRINITY_DN706400_c0_g1_i1.p1
Lophophytum–mirabile___TRINITY_DN81184_c0_g2_i1.p1
Lophophytum–mirabile___TRINITY_DN22368_c0_g4_i1.p1
Lophophytum–mirabile___TRINITY_DN22368_c0_g1_i3.p1
Malania___TRINITY_DN39499_c0_g2_i1.p1
Vitis___VIT_218s0001g13940.1
Vaccinium___Vadar_g23585.t1
Vaccinium___Vadar_g15885.t1
Solanum___Solyc04g082870.1.1
Olea___Oeu030367.1
Mimulus___Migut.C00345.1.p
Hydrangea___Hyque.07G099500.1.p
Lactuca___Lsat_1_v5_gn_6_940.1
Lactuca___Lsat_1_v5_gn_6_1020.2
Lactuca___Lsat_1_v5_gn_6_1020.1
Helianthus___HanXRQChr10g0287051
Eucalyptus___Eucgr.F01621.1.p
Manihot___Manes.11G038601.1.p
Manihot___Manes.11G038501.1.p
Manihot___Manes.11G038200.9.p
Cucumis___Cucsa.159860.1lPACid_16965267
Cucumis___Cucsa.159650.1lPACid_16965228
Citrus___orange1.1g003868mlPACid_18092178
Quercus___Qurub.10G048700.1
Malus___MD17G1265000
F–Pvulgaris___Phvul.L001644.1.p
F–Carietinum___Ca_07877
F–Akoa___GBYE01024793.1.p1
F–Prosopis___Palba_XP_028791525.1
F–Prosopis___Palba_XP_028778484.1
F–Lleucocephala___GDRZ01170424.1.p1
F–Acol___TRINITY_DN10332_c0_g2_i1.p1
Kalanchoe___Kalax.0290s0049.1.p
Kalanchoe___Kalax.0173s0032.1.p
Arabidopsis–AT1G25360.1lPACid_19658253
Musa___GSMUA_Achr3P30090_001
Amborella___AmTr_v1.0_scaffold00113.29
Vitis___VIT_209s0002g07510.1
Kalanchoe___Kalax.1284s0001.1.p
Kalanchoe___Kalax.0126s0057.1.p
Vitis___VIT_209s0096g00100.1
Vaccinium___Vadar_g36406.t2
Vaccinium___Vadar_g23877.t1
Solanum___Solyc02g076700.1.1
Lactuca___Lsat_1_v5_gn_9_118320.1
Helianthus___HanXRQChr03g0066811
Quercus___Qurub.05G113100.1
Malus___MD17G1127500
Malus___MD09G1137200
Cucumis___Cucsa.058530.1lPACid_16954691
Manihot___Manes.04G110700.1.p
Eucalyptus___Eucgr.G01109.1.p
Citrus___orange1.1g004644mlPACid_18106951
F–Pvulgaris___Phvul.010G08T100.1.p
F–Carietinum___Ca_01927
F–Prosopis___Palba_XP_028758764.1
F–Acol___TRINITY_DN31282_c0_g1_i1.p1
Musa___GSMUA_Achr2P13110_001
Kalanchoe___Kalax.0060s0066.1.p
Kalanchoe___Kalax.0002s0087.1.p
F–Prosopis___Palba_XP_028762800.1
F–Pvulgaris___Phvul.006G135000.1.p
F–Carietinum___Ca_07935
Manihot___Manes.07G139100.1.p
Solanum___Solyc07g005170.3.1
Olea___Oeu034437.1
Mimulus___Migut.O00250.1.p
Lactuca___Lsat_1_v5_gn_7_12781.1
Hydrangea___Hyque.01G215500.1.p
Eucalyptus___Eucgr.E04230.1.p
Vitis___VIT_215s0021g00360.1
Citrus___orange1.1g046354mlPACid_18112878
Musa___GSMUA_Achr7P06150_001
Solanum___Solyc11g010220.2.1
Lactuca___Lsat_1_v5_gn_5_45721.2
Lactuca___Lsat_1_v5_gn_5_45721.1
Helianthus___HanXRQChr15g0469821
Daucus___DCAR_003471
Vaccinium___Vadar_g10322.t1
Vaccinium___Vadar_g10317.t1
Hydrangea___Hyque.05G046400.1.p
Malania___TRINITY_DN64643_c0_g1_i1.p1
Citrus___orange1.1g007573mlPACid_18139325
Vitis___VIT_207s0191g00120.1
Vitis___VIT_200s1864g00010.1
Ombrophytum___TRINITY_DN62465_c0_g1_i11.p1
F–Prosopis___Palba_XP_028807150.1
F–Pvulgaris___Phvul.002G261100.2.p
F–Carietinum___Ca_09543
Malus___MD15G1320700
Musa___GSMUA_Achr3P06590_001
Vaccinium___Vadar_g4385.t1
Solanum___Solyc01g081240.3.1
Mimulus___Migut.O00821.1.p
Mimulus___Migut.N01677.1.p
Hydrangea___Hyque.17G043100.1.p
Helianthus___HanXRQChr01g0019881
Beta___EL10Ac5g11068.1
Musa___GSMUA_Achr2P04620_001
Arabidopsis–AT4G02750.1lPACid_19648010
Cucumis___Cucsa.111470.1lPACid_16960042
Quercus___Qurub.10G108100.1
Malus___MD06G1230200
Eucalyptus___Eucgr.J00262.1.p
Citrus___orange1.1g004024mlPACid_18093557
Manihot___Manes.12G140300.1.p
Kalanchoe___Kalax.1245s0003.1.p
Kalanchoe___Kalax.0066s0062.1.p
Beta___EL10Ac2g03341.1
F–Pvulgaris___Phvul.002G328000.1.p
F–Pvulgaris___Phvul.004G044000.1.p
F–Pvulgaris___Phvul.002G328100.1.p
F–Carietinum___Ca_10363
F–Prosopis___Palba_XP_028784246.1
F–Prosopis___Palba_XP_028752584.1
F–Acol___TRINITY_DN6488_c0_g1_i1.p1
Vaccinium___Vadar_g14002.t1
Solanum___Solyc03g119120.3.1
Olea___Oeu026099.1
Mimulus___Migut.D01570.1.p
Hydrangea___Hyque.15G011100.1.p
Lactuca___Lsat_1_v5_gn_8_145021.1
Helianthus___HanXRQChr06g0185221
Vitis___VIT_217s0000g02580.1
Malania___TRINITY_DN42221_c0_g1_i1.p1
Lophophytum–pyramidale___TRINITY_DN27089_c0_g1_i1.p1
Daen___TRINITY_DN2871_c0_g1_i1.p1

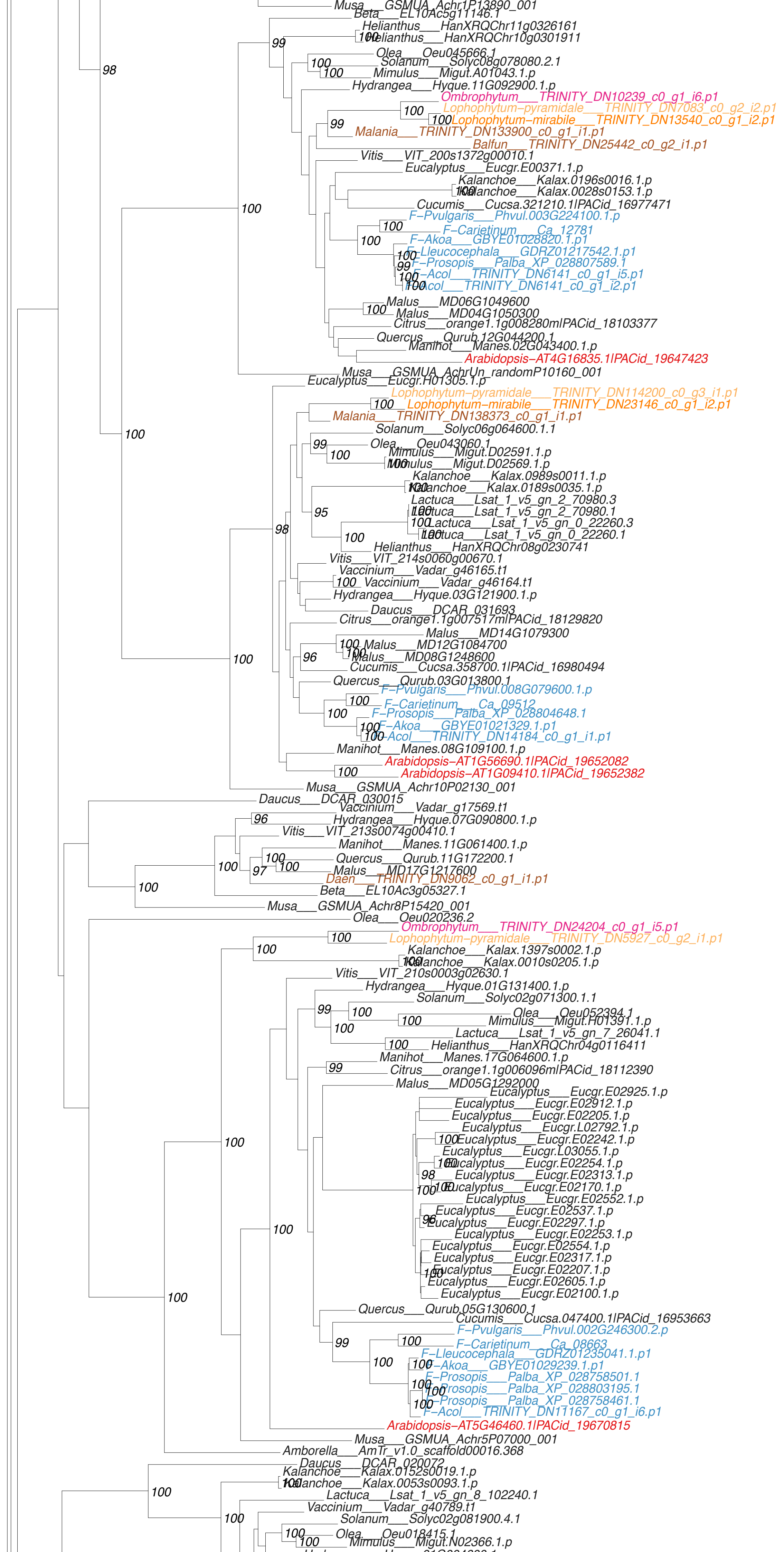

Musa___GSMUA_Achr1P13890_001
Beta___EL10Ac5g11146.1
Helianthus___HanXRQChr11g0326161
Helianthus___HanXRQChr10g0301911
Olea___Oeu045666.1
Solanum___Solyc08g078080.2.1
Mimulus___Migut.A01043.1.p
Hydrangea___Hyque.11G092900.1.p
Ombrophytum___TRINITY_DN10239_c0_g1_i6.p1
Lophophytum−pyramidale___TRINITY_DN7083_c0_g2_i2.p1
Lophophytum−mirabile___TRINITY_DN13540_c0_g1_i2.p1
Malania___TRINITY_DN133900_c0_g1_i1.p1
Balfun___TRINITY_DN25442_c0_g2_i1.p1
Vitis___VIT_200s1372g00010.1
Eucalyptus___Eucgr.E00371.1.p
Kalanchoe___Kalax.0196s0016.1.p
Kalanchoe___Kalax.0028s0153.1.p
Cucumis___Cucsa.321210.1|PACid_16977471
F−Pvulgaris___Phvul.003G224100.1.p
F−Carietinum___Ca_12781
F−Akoa___GBYE01028820.1.p1
F−Lleucocephala___GDRZ01217542.1.p1
F−Prosopis___Palba_XP_028807589.1
F−Acol___TRINITY_DN6141_c0_g1_i5.p1
F−Acol___TRINITY_DN6141_c0_g1_i2.p1
Malus___MD06G1049600
Malus___MD04G1050300
Citrus___orange1.1g008280m|PACid_18103377
Quercus___Qurub.12G044200.1
Manihot___Manes.02G043400.1.p
Arabidopsis−AT4G16835.1|PACid_19647423
Musa___GSMUA_AchrUn_randomP10160_001
Eucalyptus___Eucgr.H01305.1.p
Lophophytum−pyramidale___TRINITY_DN114200_c0_g3_i1.p1
Lophophytum−mirabile___TRINITY_DN23146_c0_g1_i2.p1
Malania___TRINITY_DN138373_c0_g1_i1.p1
Solanum___Solyc06g064600.1.1
Olea___Oeu043060.1
Mimulus___Migut.D02591.1.p
Mimulus___Migut.D02569.1.p
Kalanchoe___Kalax.0989s0011.1.p
Kalanchoe___Kalax.0189s0035.1.p
Lactuca___Lsat_1_v5_gn_2_70980.3
Lactuca___Lsat_1_v5_gn_2_70980.1
Lactuca___Lsat_1_v5_gn_0_22260.3
Lactuca___Lsat_1_v5_gn_0_22260.1
Helianthus___HanXRQChr08g0230741
Vitis___VIT_214s0060g00670.1
Vaccinium___Vadar_g46165.t1
Vaccinium___Vadar_g46164.t1
Hydrangea___Hyque.03G121900.1.p
Daucus___DCAR_031693
Citrus___orange1.1g007517m|PACid_18129820
Malus___MD14G1079300
Malus___MD12G1084700
Malus___MD08G1248600
Cucumis___Cucsa.358700.1|PACid_16980494
Quercus___Qurub.03G013800.1
F−Pvulgaris___Phvul.008G079600.1.p
F−Carietinum___Ca_09512
F−Prosopis___Palba_XP_028804648.1
F−Akoa___GBYE01021329.1.p1
F−Acol___TRINITY_DN14184_c0_g1_i1.p1
Manihot___Manes.08G109100.1.p
Arabidopsis−AT1G56690.1|PACid_19652082
Arabidopsis−AT1G09410.1|PACid_19652382
Musa___GSMUA_Achr10P02130_001
Daucus___DCAR_030015
Vaccinium___Vadar_g17569.t1
Hydrangea___Hyque.07G090800.1.p
Vitis___VIT_213s0074g00410.1
Manihot___Manes.11G061400.1.p
Quercus___Qurub.11G172200.1
Malus___MD17G1217600
Daen___TRINITY_DN9062_c0_g1_i1.p1
Beta___EL10Ac3g05327.1
Musa___GSMUA_Achr8P15420_001
Olea___Oeu020236.2
Ombrophytum___TRINITY_DN24204_c0_g1_i5.p1
Lophophytum−pyramidale___TRINITY_DN5927_c0_g2_i1.p1
Kalanchoe___Kalax.1397s0002.1.p
Kalanchoe___Kalax.0010s0205.1.p
Vitis___VIT_210s0003g02630.1
Hydrangea___Hyque.01G131400.1.p
Solanum___Solyc02g071300.1.1
Olea___Oeu052394.1
Mimulus___Migut.H01391.1.p
Lactuca___Lsat_1_v5_gn_7_26041.1
Helianthus___HanXRQChr04g0116411
Manihot___Manes.17G064600.1.p
Citrus___orange1.1g006096m|PACid_18112390
Malus___MD05G1292000
Eucalyptus___Eucgr.E02925.1.p
Eucalyptus___Eucgr.E02912.1.p
Eucalyptus___Eucgr.E02205.1.p
Eucalyptus___Eucgr.L02792.1.p
Eucalyptus___Eucgr.E02242.1.p
Eucalyptus___Eucgr.L03055.1.p
Eucalyptus___Eucgr.E02254.1.p
Eucalyptus___Eucgr.E02313.1.p
Eucalyptus___Eucgr.E02170.1.p
Eucalyptus___Eucgr.E02552.1.p
Eucalyptus___Eucgr.E02537.1.p
Eucalyptus___Eucgr.E02297.1.p
Eucalyptus___Eucgr.E02253.1.p
Eucalyptus___Eucgr.E02554.1.p
Eucalyptus___Eucgr.E02317.1.p
Eucalyptus___Eucgr.E02207.1.p
Eucalyptus___Eucgr.E02605.1.p
Eucalyptus___Eucgr.E02100.1.p
Quercus___Qurub.05G130600.1
Cucumis___Cucsa.047400.1|PACid_16953663
F−Pvulgaris___Phvul.002G246300.2.p
F−Carietinum___Ca_08663
F−Lleucocephala___GDRZ01235041.1.p1
F−Akoa___GBYE01029239.1.p1
F−Prosopis___Palba_XP_028758501.1
F−Prosopis___Palba_XP_028803195.1
F−Prosopis___Palba_XP_028758461.1
F−Acol___TRINITY_DN11167_c0_g1_i6.p1
Arabidopsis−AT5G46460.1|PACid_19670815
Musa___GSMUA_Achr5P07000_001
Amborella___AmTr_v1.0_scaffold00016.368
Daucus___DCAR_020072
Kalanchoe___Kalax.0152s0019.1.p
Kalanchoe___Kalax.0053s0093.1.p
Lactuca___Lsat_1_v5_gn_8_102240.1
Vaccinium___Vadar_g40789.t1
Solanum___Solyc02g081900.4.1
Olea___Oeu018415.1
Mimulus___Migut.N02366.1.p

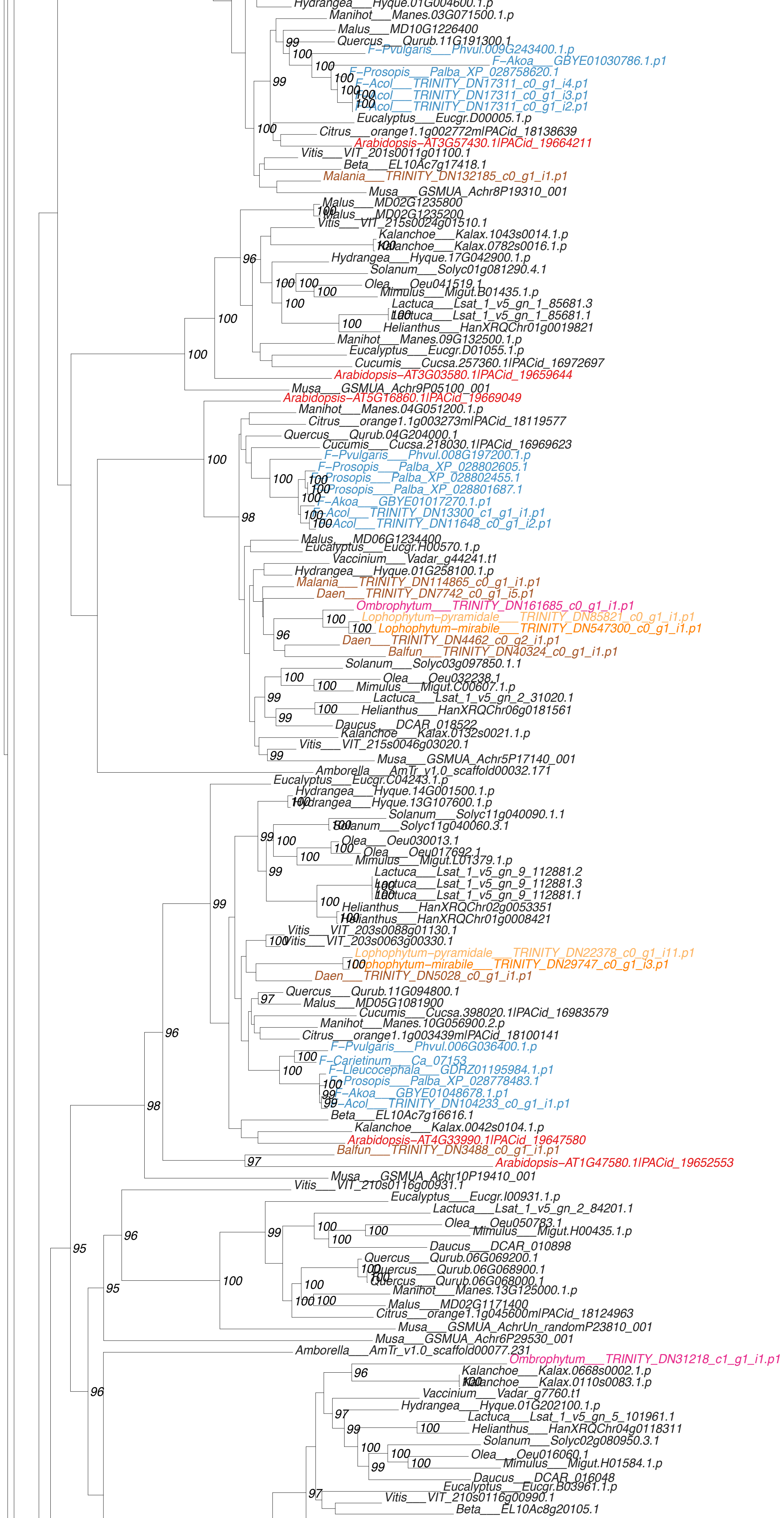

Hydrangea___Hyque.01G004600.1.p
Manihot___Manes.03G071500.1.p
Malus___MD10G1226400
Quercus___Qurub.11G191300.1
F–Pvulgaris___Phvul.009G243400.1.p
F–Akoa___GBYE01030786.1.p1
F–Prosopis___Palba_XP_028758620.1
F–Acol___TRINITY_DN17311_c0_g1_i4.p1
F–Acol___TRINITY_DN17311_c0_g1_i3.p1
F–Acol___TRINITY_DN17311_c0_g1_i2.p1
Eucalyptus___Eucgr.D00005.1.p
Citrus___orange1.1g002772mlPACid_18138639
Arabidopsis–AT3G57430.1lPACid_19664211
Vitis___VIT_201s0011g01100.1
Beta___EL10Ac7g17418.1
Malania___TRINITY_DN132185_c0_g1_i1.p1
Musa___GSMUA_Achr8P19310_001
Malus___MD02G1235800
Malus___MD02G1235200
Vitis___VIT_215s0024g01510.1
Kalanchoe___Kalax.1043s0014.1.p
Kalanchoe___Kalax.0782s0016.1.p
Hydrangea___Hyque.17G042900.1.p
Solanum___Solyc01g081290.4.1
Olea___Oeu041519.1
Mimulus___Migut.B01435.1.p
Lactuca___Lsat_1_v5_gn_1_85681.3
Lactuca___Lsat_1_v5_gn_1_85681.1
Helianthus___HanXRQChr01g0019821
Manihot___Manes.09G132500.1.p
Eucalyptus___Eucgr.D01055.1.p
Cucumis___Cucsa.257360.1lPACid_16972697
Arabidopsis–AT3G03580.1lPACid_19659644
Musa___GSMUA_Achr9P05100_001
Arabidopsis–AT5G16860.1lPACid_19669049
Manihot___Manes.04G051200.1.p
Citrus___orange1.1g003273mlPACid_18119577
Quercus___Qurub.04G204000.1
Cucumis___Cucsa.218030.1lPACid_16969623
F–Pvulgaris___Phvul.008G197200.1.p
F–Prosopis___Palba_XP_028802605.1
F–Prosopis___Palba_XP_028802455.1
F–Prosopis___Palba_XP_028801687.1
F–Akoa___GBYE01017270.1.p1
F–Acol___TRINITY_DN13300_c1_g1_i1.p1
F–Acol___TRINITY_DN11648_c0_g1_i2.p1
Malus___MD06G1234400
Eucalyptus___Eucgr.H00570.1.p
Vaccinium___Vadar_g44241.t1
Hydrangea___Hyque.01G258100.1.p
Malania___TRINITY_DN114865_c0_g1_i1.p1
Daen___TRINITY_DN7742_c0_g1_i5.p1
Ombrophytum___TRINITY_DN161685_c0_g1_i1.p1
Lophophytum–pyramidale___TRINITY_DN85821_c0_g1_i1.p1
Lophophytum–mirabile___TRINITY_DN547300_c0_g1_i1.p1
Daen___TRINITY_DN4462_c0_g2_i1.p1
Balfun___TRINITY_DN40324_c0_g1_i1.p1
Solanum___Solyc03g097850.1.1
Olea___Oeu032238.1
Mimulus___Migut.C00607.1.p
Lactuca___Lsat_1_v5_gn_2_31020.1
Helianthus___HanXRQChr06g0181561
Daucus___DCAR_018522
Kalanchoe___Kalax.0132s0021.1.p
Vitis___VIT_215s0046g03020.1
Musa___GSMUA_Achr5P17140_001
Amborella___AmTr_v1.0_scaffold00032.171
Eucalyptus___Eucgr.C04243.1.p
Hydrangea___Hyque.14G001500.1.p
Hydrangea___Hyque.13G107600.1.p
Solanum___Solyc11g040090.1.1
Solanum___Solyc11g040060.3.1
Olea___Oeu030013.1
Olea___Oeu017692.1
Mimulus___Migut.L01379.1.p
Lactuca___Lsat_1_v5_gn_9_112881.2
Lactuca___Lsat_1_v5_gn_9_112881.3
Lactuca___Lsat_1_v5_gn_9_112881.1
Helianthus___HanXRQChr02g0053351
Helianthus___HanXRQChr01g0008421
Vitis___VIT_203s0088g01130.1
Vitis___VIT_203s0063g00330.1
Lophophytum–pyramidale___TRINITY_DN22378_c0_g1_i11.p1
Lophophytum–mirabile___TRINITY_DN29747_c0_g1_i3.p1
Daen___TRINITY_DN5028_c0_g1_i1.p1
Quercus___Qurub.11G094800.1
Malus___MD05G1081900
Cucumis___Cucsa.398020.1lPACid_16983579
Manihot___Manes.10G056900.2.p
Citrus___orange1.1g003439mlPACid_18100141
F–Pvulgaris___Phvul.006G036400.1.p
F–Carietinum___Ca_07153
F–Lleucocephala___GDRZ01195984.1.p1
F–Prosopis___Palba_XP_028778483.1
F–Akoa___GBYE01048678.1.p1
F–Acol___TRINITY_DN104233_c0_g1_i1.p1
Beta___EL10Ac7g16616.1
Kalanchoe___Kalax.0042s0104.1.p
Arabidopsis–AT4G33990.1lPACid_19647580
Balfun___TRINITY_DN3488_c0_g1_i1.p1
Arabidopsis–AT1G47580.1lPACid_19652553
Musa___GSMUA_Achr10P19410_001
Vitis___VIT_210s0116g00931.1
Eucalyptus___Eucgr.I00931.1.p
Lactuca___Lsat_1_v5_gn_2_84201.1
Olea___Oeu050783.1
Mimulus___Migut.H00435.1.p
Daucus___DCAR_010898
Quercus___Qurub.06G069200.1
Quercus___Qurub.06G068900.1
Quercus___Qurub.06G068000.1
Manihot___Manes.13G125000.1.p
Malus___MD02G1171400
Citrus___orange1.1g045600mlPACid_18124963
Musa___GSMUA_AchrUn_randomP23810_001
Musa___GSMUA_Achr6P29530_001
Amborella___AmTr_v1.0_scaffold00077.231
Ombrophytum___TRINITY_DN31218_c1_g1_i1.p1
Kalanchoe___Kalax.0668s0002.1.p
Kalanchoe___Kalax.0110s0083.1.p
Vaccinium___Vadar_g7760.t1
Hydrangea___Hyque.01G202100.1.p
Lactuca___Lsat_1_v5_gn_5_101961.1
Helianthus___HanXRQChr04g0118311
Solanum___Solyc02g080950.3.1
Olea___Oeu016060.1
Mimulus___Migut.H01584.1.p
Daucus___DCAR_016048
Eucalyptus___Eucgr.B03961.1.p
Vitis___VIT_210s0116g00990.1
Beta___EL10Ac8g20105.1

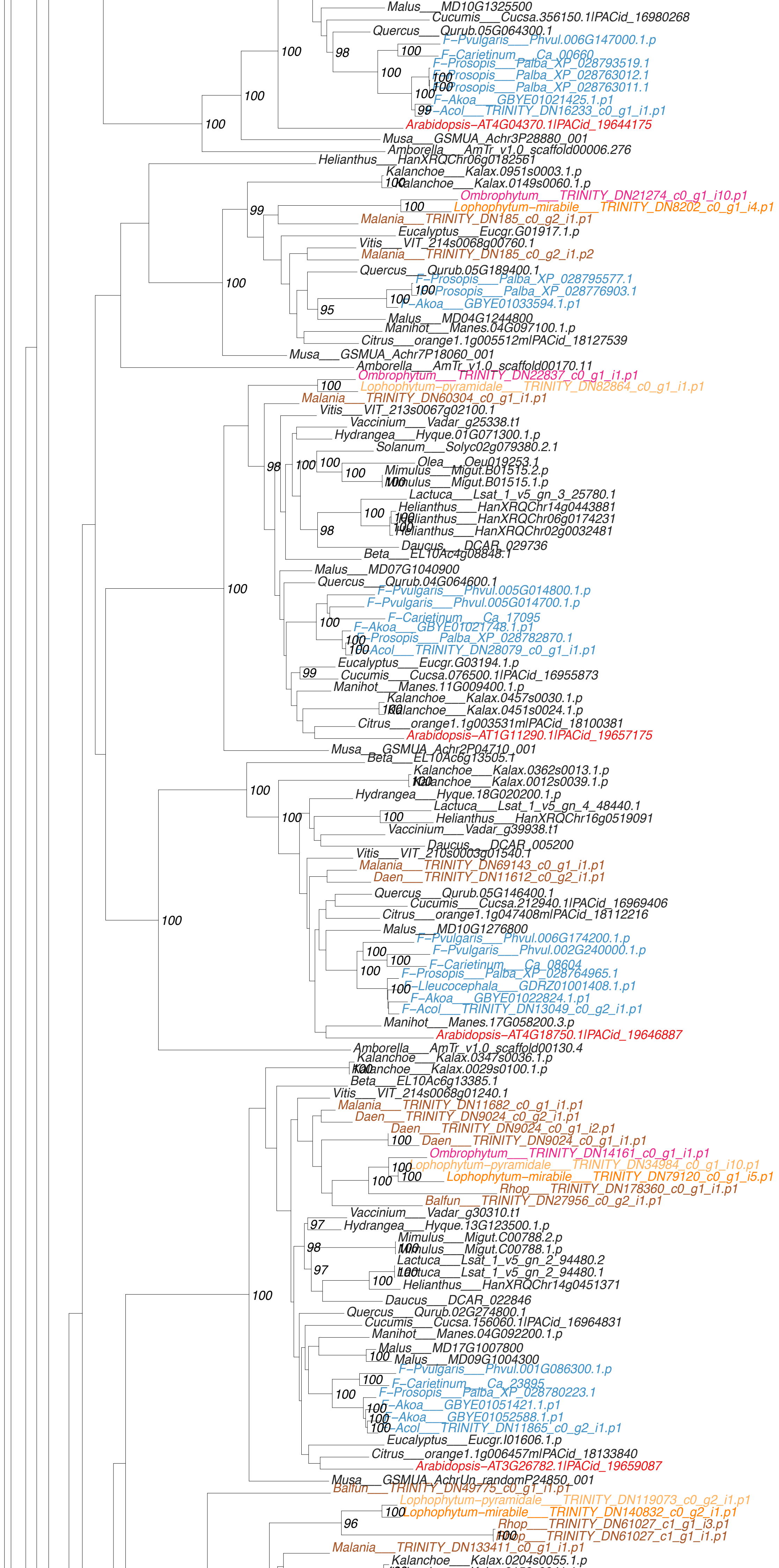
Malus___MD10G1325500
Cucumis___Cucsa.356150.1IPACid_16980268
Quercus___Qurub.05G064300.1
F–Pvulgaris___Phvul.006G147000.1.p
F–Carietinum___Ca_00660
F–Prosopis___Palba_XP_028793519.1
F–Prosopis___Palba_XP_028763012.1
F–Prosopis___Palba_XP_028763011.1
F–Akoa___GBYE01021425.1.p1
F–Acol___TRINITY_DN16233_c0_g1_i1.p1
Arabidopsis–AT4G04370.1IPACid_19644175
Musa___GSMUA_Achr3P28880_001
Amborella___AmTr_v1.0_scaffold00006.276
Helianthus___HanXRQChr06g0182561
Kalanchoe___Kalax.0951s0003.1.p
Kalanchoe___Kalax.0149s0060.1.p
Ombrophytum___TRINITY_DN21274_c0_g1_i10.p1
Lophophytum–mirabile___TRINITY_DN8202_c0_g1_i4.p1
Malania___TRINITY_DN185_c0_g2_i1.p1
Eucalyptus___Eucgr.G01917.1.p
Vitis___VIT_214s0068g00760.1
Malania___TRINITY_DN185_c0_g2_i1.p2
Quercus___Qurub.05G189400.1
F–Prosopis___Palba_XP_028795577.1
F–Prosopis___Palba_XP_028776903.1
F–Akoa___GBYE01033594.1.p1
Malus___MD04G1244800
Manihot___Manes.04G097100.1.p
Citrus___orange1.1g005512mIPACid_18127539
Musa___GSMUA_Achr7P18060_001
Amborella___AmTr_v1.0_scaffold00170.11
Ombrophytum___TRINITY_DN22837_c0_g1_i1.p1
Lophophytum–pyramidale___TRINITY_DN82864_c0_g1_i1.p1
Malania___TRINITY_DN60304_c0_g1_i1.p1
Vitis___VIT_213s0067g02100.1
Vaccinium___Vadar_g25338.t1
Hydrangea___Hyque.01G071300.1.p
Solanum___Solyc02g079380.2.1
Olea___Oeu019253.1
Mimulus___Migut.B01515.2.p
Mimulus___Migut.B01515.1.p
Lactuca___Lsat_1_v5_gn_3_25780.1
Helianthus___HanXRQChr14g0443881
Helianthus___HanXRQChr06g0174231
Helianthus___HanXRQChr02g0032481
Daucus___DCAR_029736
Beta___EL10Ac4g08848.1
Malus___MD07G1040900
Quercus___Qurub.04G064600.1
F–Pvulgaris___Phvul.005G014800.1.p
F–Pvulgaris___Phvul.005G014700.1.p
F–Carietinum___Ca_17095
F–Akoa___GBYE01021748.1.p1
F–Prosopis___Palba_XP_028782870.1
F–Acol___TRINITY_DN28079_c0_g1_i1.p1
Eucalyptus___Eucgr.G03194.1.p
Cucumis___Cucsa.076500.1IPACid_16955873
Manihot___Manes.11G009400.1.p
Kalanchoe___Kalax.0457s0030.1.p
Kalanchoe___Kalax.0451s0024.1.p
Citrus___orange1.1g003531mIPACid_18100381
Arabidopsis–AT1G11290.1IPACid_19657175
Musa___GSMUA_Achr2P04710_001
Beta___EL10Ac6g13505.1
Kalanchoe___Kalax.0362s0013.1.p
Kalanchoe___Kalax.0012s0039.1.p
Hydrangea___Hyque.18G020200.1.p
Lactuca___Lsat_1_v5_gn_4_48440.1
Helianthus___HanXRQChr16g0519091
Vaccinium___Vadar_g39938.t1
Daucus___DCAR_005200
Vitis___VIT_210s0003g01540.1
Malania___TRINITY_DN69143_c0_g1_i1.p1
Daen___TRINITY_DN11612_c0_g2_i1.p1
Quercus___Qurub.05G146400.1
Cucumis___Cucsa.212940.1IPACid_16969406
Citrus___orange1.1g047408mIPACid_18112216
Malus___MD10G1276800
F–Pvulgaris___Phvul.006G174200.1.p
F–Pvulgaris___Phvul.002G240000.1.p
F–Carietinum___Ca_08604
F–Prosopis___Palba_XP_028764965.1
F–Lleucocephala___GDRZ01001408.1.p1
F–Akoa___GBYE01022824.1.p1
F–Acol___TRINITY_DN13049_c0_g2_i1.p1
Manihot___Manes.17G058200.3.p
Arabidopsis–AT4G18750.1IPACid_19646887
Amborella___AmTr_v1.0_scaffold00130.4
Kalanchoe___Kalax.0347s0036.1.p
Kalanchoe___Kalax.0029s0100.1.p
Beta___EL10Ac6g13385.1
Vitis___VIT_214s0068g01240.1
Malania___TRINITY_DN11682_c0_g1_i1.p1
Daen___TRINITY_DN9024_c0_g2_i1.p1
Daen___TRINITY_DN9024_c0_g1_i2.p1
Daen___TRINITY_DN9024_c0_g1_i1.p1
Ombrophytum___TRINITY_DN14161_c0_g1_i1.p1
Lophophytum–pyramidale___TRINITY_DN34984_c0_g1_i10.p1
Lophophytum–mirabile___TRINITY_DN79120_c0_g1_i5.p1
Rhop___TRINITY_DN178360_c0_g1_i1.p1
Balfun___TRINITY_DN27956_c0_g2_i1.p1
Vaccinium___Vadar_g30310.t1
Hydrangea___Hyque.13G123500.1.p
Mimulus___Migut.C00788.2.p
Mimulus___Migut.C00788.1.p
Lactuca___Lsat_1_v5_gn_2_94480.2
Lactuca___Lsat_1_v5_gn_2_94480.1
Helianthus___HanXRQChr14g0451371
Daucus___DCAR_022846
Quercus___Qurub.02G274800.1
Cucumis___Cucsa.156060.1IPACid_16964831
Manihot___Manes.04G092200.1.p
Malus___MD17G1007800
Malus___MD09G1004300
F–Pvulgaris___Phvul.001G086300.1.p
F–Carietinum___Ca_23895
F–Prosopis___Palba_XP_028780223.1
F–Akoa___GBYE01051421.1.p1
F–Akoa___GBYE01052588.1.p1
F–Acol___TRINITY_DN11865_c0_g2_i1.p1
Eucalyptus___Eucgr.I01606.1.p
Citrus___orange1.1g006457mIPACid_18133840
Arabidopsis–AT3G26782.1IPACid_19659087
Musa___GSMUA_AchrUn_randomP24850_001
Balfun___TRINITY_DN49775_c0_g1_i1.p1
Lophophytum–pyramidale___TRINITY_DN119073_c0_g2_i1.p1
Lophophytum–mirabile___TRINITY_DN140832_c0_g2_i1.p1
Rhop___TRINITY_DN61027_c1_g1_i3.p1
Rhop___TRINITY_DN61027_c1_g1_i1.p1
Malania___TRINITY_DN133411_c0_g1_i1.p1
Kalanchoe___Kalax.0204s0055.1.p

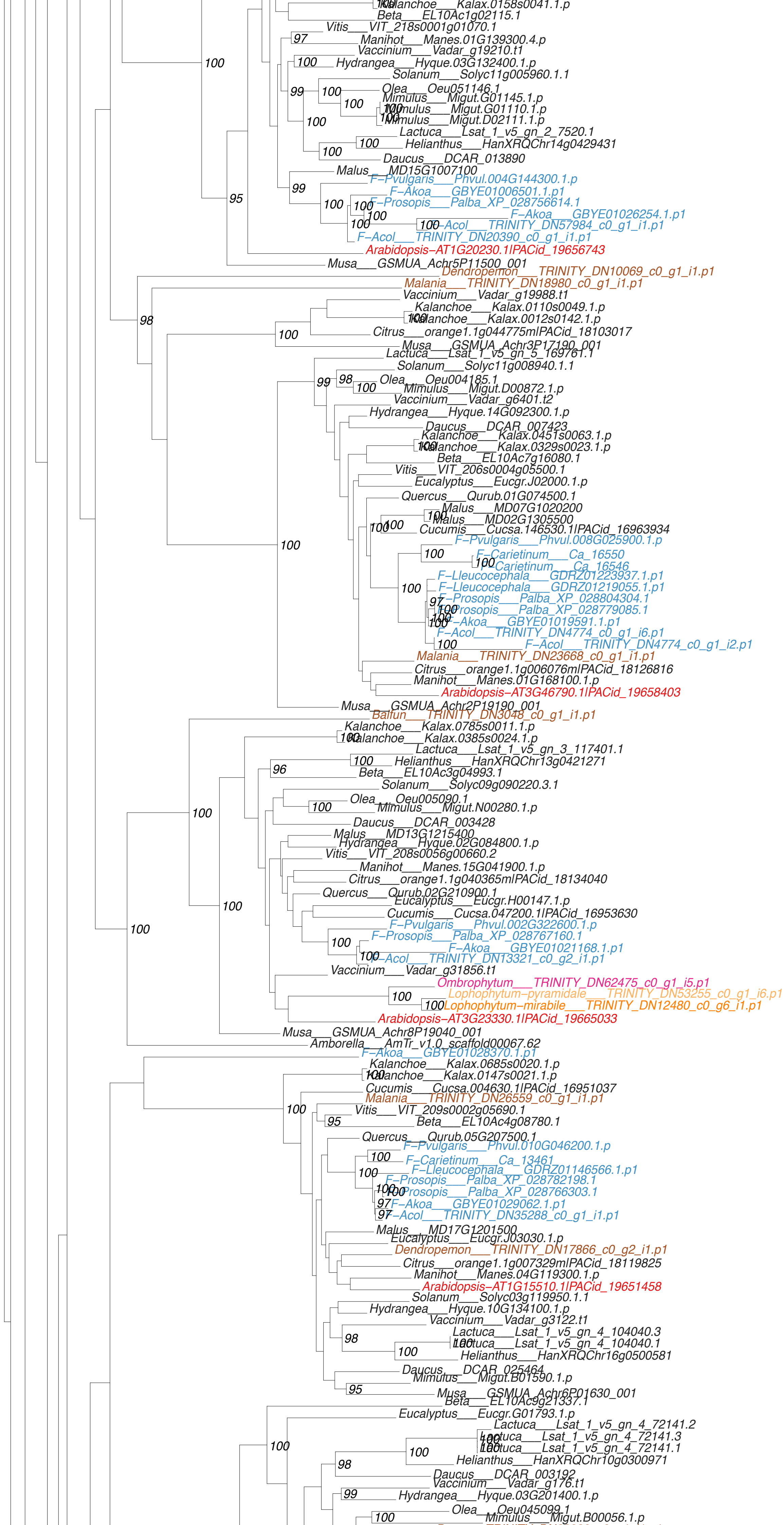
Kalanchoe___Kalax.0158s0041.1.p
Beta___EL10Ac1g02115.1
Vitis___VIT_218s0001g01070.1
Manihot___Manes.01G139300.4.p
Vaccinium___Vadar_g19210.t1
Hydrangea___Hyque.03G132400.1.p
Solanum___Solyc11g005960.1.1
Olea___Oeu051146.1
Mimulus___Migut.G01145.1.p
Mimulus___Migut.G01110.1.p
Mimulus___Migut.D02111.1.p
Lactuca___Lsat_1_v5_gn_2_7520.1
Helianthus___HanXRQChr14g0429431
Daucus___DCAR_013890
Malus___MD15G1007100
F–Pvulgaris___Phvul.004G144300.1.p
F–Akoa___GBYE01006501.1.p1
F–Prosopis___Palba_XP_028756614.1
F–Akoa___GBYE01026254.1.p1
F–Acol___TRINITY_DN57984_c0_g1_i1.p1
F–Acol___TRINITY_DN20390_c0_g1_i1.p1
Arabidopsis–AT1G20230.1|PACid_19656743
Musa___GSMUA_Achr5P11500_001
Dendropemon___TRINITY_DN10069_c0_g1_i1.p1
Malania___TRINITY_DN18980_c0_g1_i1.p1
Vaccinium___Vadar_g19988.t1
Kalanchoe___Kalax.0110s0049.1.p
Kalanchoe___Kalax.0012s0142.1.p
Citrus___orange1.1g044775m|PACid_18103017
Musa___GSMUA_Achr3P17190_001
Lactuca___Lsat_1_v5_gn_5_169761.1
Solanum___Solyc11g008940.1.1
Olea___Oeu004185.1
Mimulus___Migut.D00872.1.p
Vaccinium___Vadar_g6401.t2
Hydrangea___Hyque.14G092300.1.p
Daucus___DCAR_007423
Kalanchoe___Kalax.0451s0063.1.p
Kalanchoe___Kalax.0329s0023.1.p
Beta___EL10Ac7g16080.1
Vitis___VIT_206s0004g05500.1
Eucalyptus___Eucgr.J02000.1.p
Quercus___Qurub.01G074500.1
Malus___MD07G1020200
Malus___MD02G1305500
Cucumis___Cucsa.146530.1|PACid_16963934
F–Pvulgaris___Phvul.008G025900.1.p
F–Carietinum___Ca_16550
F–Carietinum___Ca_16546
F–Lleucocephala___GDRZ01223937.1.p1
F–Lleucocephala___GDRZ01219055.1.p1
F–Prosopis___Palba_XP_028804304.1
F–Prosopis___Palba_XP_028779085.1
F–Akoa___GBYE01019591.1.p1
F–Acol___TRINITY_DN4774_c0_g1_i6.p1
F–Acol___TRINITY_DN4774_c0_g1_i2.p1
Malania___TRINITY_DN23668_c0_g1_i1.p1
Citrus___orange1.1g006076m|PACid_18126816
Manihot___Manes.01G168100.1.p
Arabidopsis–AT3G46790.1|PACid_19658403
Musa___GSMUA_Achr2P19190_001
Balfun___TRINITY_DN3048_c0_g1_i1.p1
Kalanchoe___Kalax.0785s0011.1.p
Kalanchoe___Kalax.0385s0024.1.p
Lactuca___Lsat_1_v5_gn_3_117401.1
Helianthus___HanXRQChr13g0421271
Beta___EL10Ac3g04993.1
Solanum___Solyc09g090220.3.1
Olea___Oeu005090.1
Mimulus___Migut.N00280.1.p
Daucus___DCAR_003428
Malus___MD13G1215400
Hydrangea___Hyque.02G084800.1.p
Vitis___VIT_208s0056g00660.2
Manihot___Manes.15G041900.1.p
Citrus___orange1.1g040365m|PACid_18134040
Quercus___Qurub.02G210900.1
Eucalyptus___Eucgr.H00147.1.p
Cucumis___Cucsa.047200.1|PACid_16953630
F–Pvulgaris___Phvul.002G322600.1.p
F–Prosopis___Palba_XP_028767160.1
F–Akoa___GBYE01021168.1.p1
F–Acol___TRINITY_DN13321_c0_g2_i1.p1
Vaccinium___Vadar_g31856.t1
Ombrophytum___TRINITY_DN62475_c0_g1_i5.p1
Lophophytum–pyramidale___TRINITY_DN53255_c0_g1_i6.p1
Lophophytum–mirabile___TRINITY_DN12480_c0_g6_i1.p1
Arabidopsis–AT3G23330.1|PACid_19665033
Musa___GSMUA_Achr8P19040_001
Amborella___AmTr_v1.0_scaffold00067.62
F–Akoa___GBYE01028370.1.p1
Kalanchoe___Kalax.0685s0020.1.p
Kalanchoe___Kalax.0147s0021.1.p
Cucumis___Cucsa.004630.1|PACid_16951037
Malania___TRINITY_DN26559_c0_g1_i1.p1
Vitis___VIT_209s0002g05690.1
Beta___EL10Ac4g08780.1
Quercus___Qurub.05G207500.1
F–Pvulgaris___Phvul.010G046200.1.p
F–Carietinum___Ca_13461
F–Lleucocephala___GDRZ01146566.1.p1
F–Prosopis___Palba_XP_028782198.1
F–Prosopis___Palba_XP_028766303.1
F–Akoa___GBYE01029062.1.p1
F–Acol___TRINITY_DN35288_c0_g1_i1.p1
Malus___MD17G1201500
Eucalyptus___Eucgr.J03030.1.p
Dendropemon___TRINITY_DN17866_c0_g2_i1.p1
Citrus___orange1.1g007329m|PACid_18119825
Manihot___Manes.04G119300.1.p
Arabidopsis–AT1G15510.1|PACid_19651458
Solanum___Solyc03g119950.1.1
Hydrangea___Hyque.10G134100.1.p
Vaccinium___Vadar_g3122.t1
Lactuca___Lsat_1_v5_gn_4_104040.3
Lactuca___Lsat_1_v5_gn_4_104040.1
Helianthus___HanXRQChr16g0500581
Daucus___DCAR_025464
Mimulus___Migut.B01590.1.p
Musa___GSMUA_Achr6P01630_001
Beta___EL10Ac9g21337.1
Eucalyptus___Eucgr.G01793.1.p
Lactuca___Lsat_1_v5_gn_4_72141.2
Lactuca___Lsat_1_v5_gn_4_72141.3
Lactuca___Lsat_1_v5_gn_4_72141.1
Helianthus___HanXRQChr10g0300971
Daucus___DCAR_003192
Vaccinium___Vadar_g176.t1
Hydrangea___Hyque.03G201400.1.p
Olea___Oeu045099.1
Mimulus___Migut.B00056.1.p

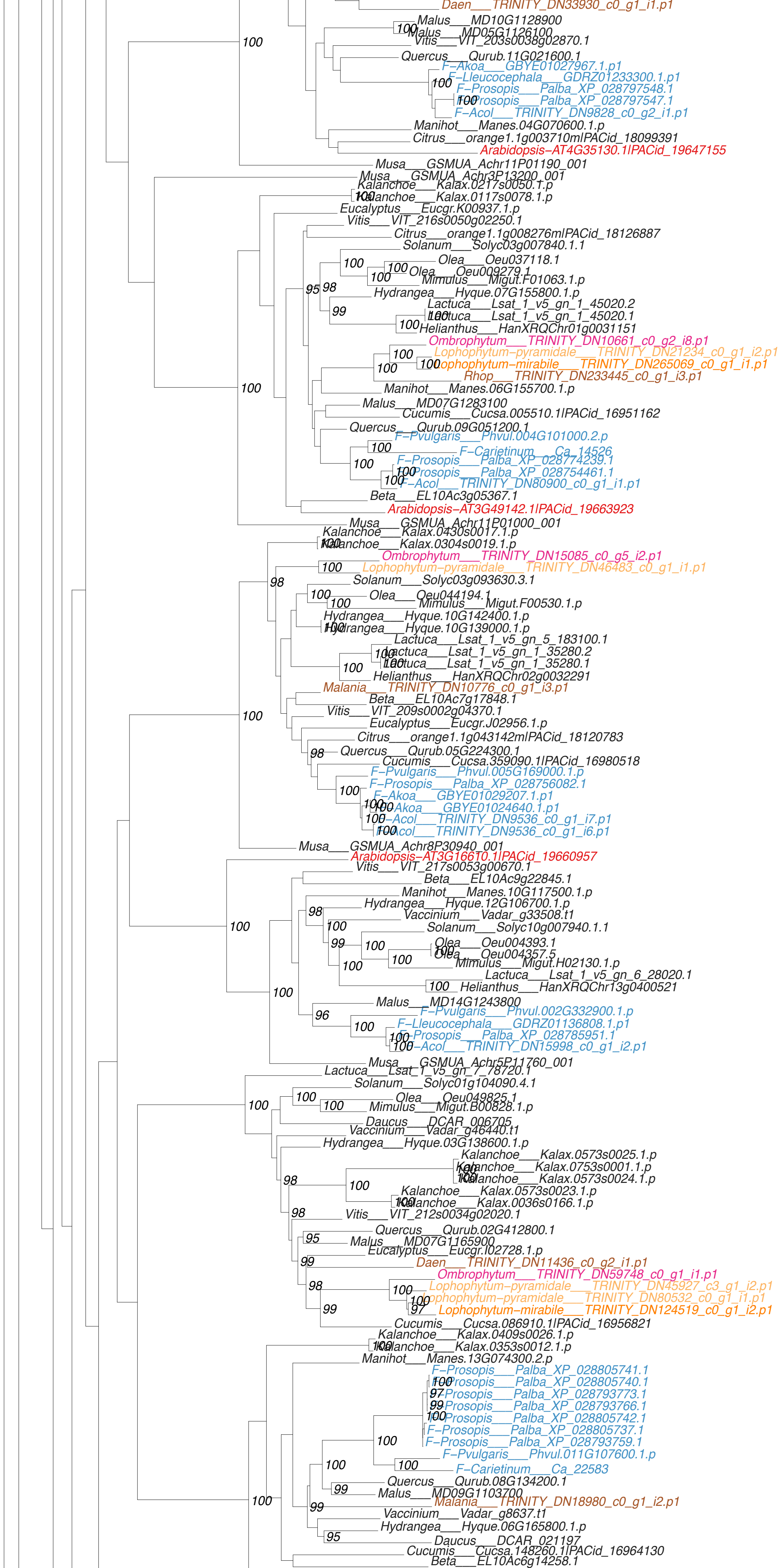
Daen___TRINITY_DN33930_c0_g1_i1.p1
Malus___MD10G1128900
Malus___MD05G1126100
Vitis___VIT_203s0038g02870.1
Quercus___Qurub.11G021600.1
F–Akoa___GBYE01027967.1.p1
F–Lleucocephala___GDRZ01233300.1.p1
F–Prosopis___Palba_XP_028797548.1
F–Prosopis___Palba_XP_028797547.1
F–Acol___TRINITY_DN9828_c0_g2_i1.p1
Manihot___Manes.04G070600.1.p
Citrus___orange1.1g003710mIPACid_18099391
Arabidopsis–AT4G35130.1IPACid_19647155
Musa___GSMUA_Achr11P01190_001
Musa___GSMUA_Achr3P13200_001
Kalanchoe___Kalax.0217s0050.1.p
Kalanchoe___Kalax.0117s0078.1.p
Eucalyptus___Eucgr.K00937.1.p
Vitis___VIT_216s0050g02250.1
Citrus___orange1.1g008276mIPACid_18126887
Solanum___Solyc03g007840.1.1
Olea___Oeu037118.1
Olea___Oeu009279.1
Mimulus___Migut.F01063.1.p
Hydrangea___Hyque.07G155800.1.p
Lactuca___Lsat_1_v5_gn_1_45020.2
Lactuca___Lsat_1_v5_gn_1_45020.1
Helianthus___HanXRQChr01g0031151
Ombrophytum___TRINITY_DN10661_c0_g2_i8.p1
Lophophytum–pyramidale___TRINITY_DN21234_c0_g1_i2.p1
Lophophytum–mirabile___TRINITY_DN265069_c0_g1_i1.p1
Rhop___TRINITY_DN233445_c0_g1_i3.p1
Manihot___Manes.06G155700.1.p
Malus___MD07G1283100
Cucumis___Cucsa.005510.1IPACid_16951162
Quercus___Qurub.09G051200.1
F–Pvulgaris___Phvul.004G101000.2.p
F–Carietinum___Ca_14526
F–Prosopis___Palba_XP_028774239.1
F–Prosopis___Palba_XP_028754461.1
F–Acol___TRINITY_DN80900_c0_g1_i1.p1
Beta___EL10Ac3g05367.1
Arabidopsis–AT3G49142.1IPACid_19663923
Musa___GSMUA_Achr11P01000_001
Kalanchoe___Kalax.0430s0017.1.p
Kalanchoe___Kalax.0304s0019.1.p
Ombrophytum___TRINITY_DN15085_c0_g5_i2.p1
Lophophytum–pyramidale___TRINITY_DN46483_c0_g1_i1.p1
Solanum___Solyc03g093630.3.1
Olea___Oeu044194.1
Mimulus___Migut.F00530.1.p
Hydrangea___Hyque.10G142400.1.p
Hydrangea___Hyque.10G139000.1.p
Lactuca___Lsat_1_v5_gn_5_183100.1
Lactuca___Lsat_1_v5_gn_1_35280.2
Lactuca___Lsat_1_v5_gn_1_35280.1
Helianthus___HanXRQChr02g0032291
Malania___TRINITY_DN10776_c0_g1_i3.p1
Beta___EL10Ac7g17848.1
Vitis___VIT_209s0002g04370.1
Eucalyptus___Eucgr.J02956.1.p
Citrus___orange1.1g043142mIPACid_18120783
Quercus___Qurub.05G224300.1
Cucumis___Cucsa.359090.1IPACid_16980518
F–Pvulgaris___Phvul.005G169000.1.p
F–Prosopis___Palba_XP_028756082.1
F–Akoa___GBYE01029207.1.p1
F–Akoa___GBYE01024640.1.p1
F–Acol___TRINITY_DN9536_c0_g1_i7.p1
F–Acol___TRINITY_DN9536_c0_g1_i6.p1
Musa___GSMUA_Achr8P30940_001
Arabidopsis–AT3G16610.1IPACid_19660957
Vitis___VIT_217s0053g00670.1
Beta___EL10Ac9g22845.1
Manihot___Manes.10G117500.1.p
Hydrangea___Hyque.12G106700.1.p
Vaccinium___Vadar_g33508.t1
Solanum___Solyc10g007940.1.1
Olea___Oeu004393.1
Olea___Oeu004357.5
Mimulus___Migut.H02130.1.p
Lactuca___Lsat_1_v5_gn_6_28020.1
Helianthus___HanXRQChr13g0400521
Malus___MD14G1243800
F–Pvulgaris___Phvul.002G332900.1.p
F–Lleucocephala___GDRZ01136808.1.p1
F–Prosopis___Palba_XP_028785951.1
F–Acol___TRINITY_DN15998_c0_g1_i2.p1
Musa___GSMUA_Achr5P11760_001
Lactuca___Lsat_1_v5_gn_7_78720.1
Solanum___Solyc01g104090.4.1
Olea___Oeu049825.1
Mimulus___Migut.B00828.1.p
Daucus___DCAR_006705
Vaccinium___Vadar_g46440.t1
Hydrangea___Hyque.03G138600.1.p
Kalanchoe___Kalax.0573s0025.1.p
Kalanchoe___Kalax.0753s0001.1.p
Kalanchoe___Kalax.0573s0024.1.p
Kalanchoe___Kalax.0573s0023.1.p
Kalanchoe___Kalax.0036s0166.1.p
Vitis___VIT_212s0034g02020.1
Quercus___Qurub.02G412800.1
Malus___MD07G1165900
Eucalyptus___Eucgr.I02728.1.p
Daen___TRINITY_DN11436_c0_g2_i1.p1
Ombrophytum___TRINITY_DN59748_c0_g1_i1.p1
Lophophytum–pyramidale___TRINITY_DN45927_c3_g1_i2.p1
Lophophytum–pyramidale___TRINITY_DN80532_c0_g1_i1.p1
Lophophytum–mirabile___TRINITY_DN124519_c0_g1_i2.p1
Cucumis___Cucsa.086910.1IPACid_16956821
Kalanchoe___Kalax.0409s0026.1.p
Kalanchoe___Kalax.0353s0012.1.p
Manihot___Manes.13G074300.2.p
F–Prosopis___Palba_XP_028805741.1
F–Prosopis___Palba_XP_028805740.1
F–Prosopis___Palba_XP_028793773.1
F–Prosopis___Palba_XP_028793766.1
F–Prosopis___Palba_XP_028805742.1
F–Prosopis___Palba_XP_028805737.1
F–Prosopis___Palba_XP_028793759.1
F–Pvulgaris___Phvul.011G107600.1.p
F–Carietinum___Ca_22583
Quercus___Qurub.08G134200.1
Malus___MD09G1103700
Malania___TRINITY_DN18980_c0_g1_i2.p1
Vaccinium___Vadar_g8637.t1
Hydrangea___Hyque.06G165800.1.p
Daucus___DCAR_021197
Cucumis___Cucsa.148260.1IPACid_16964130
Beta___EL10Ac6g14258.1

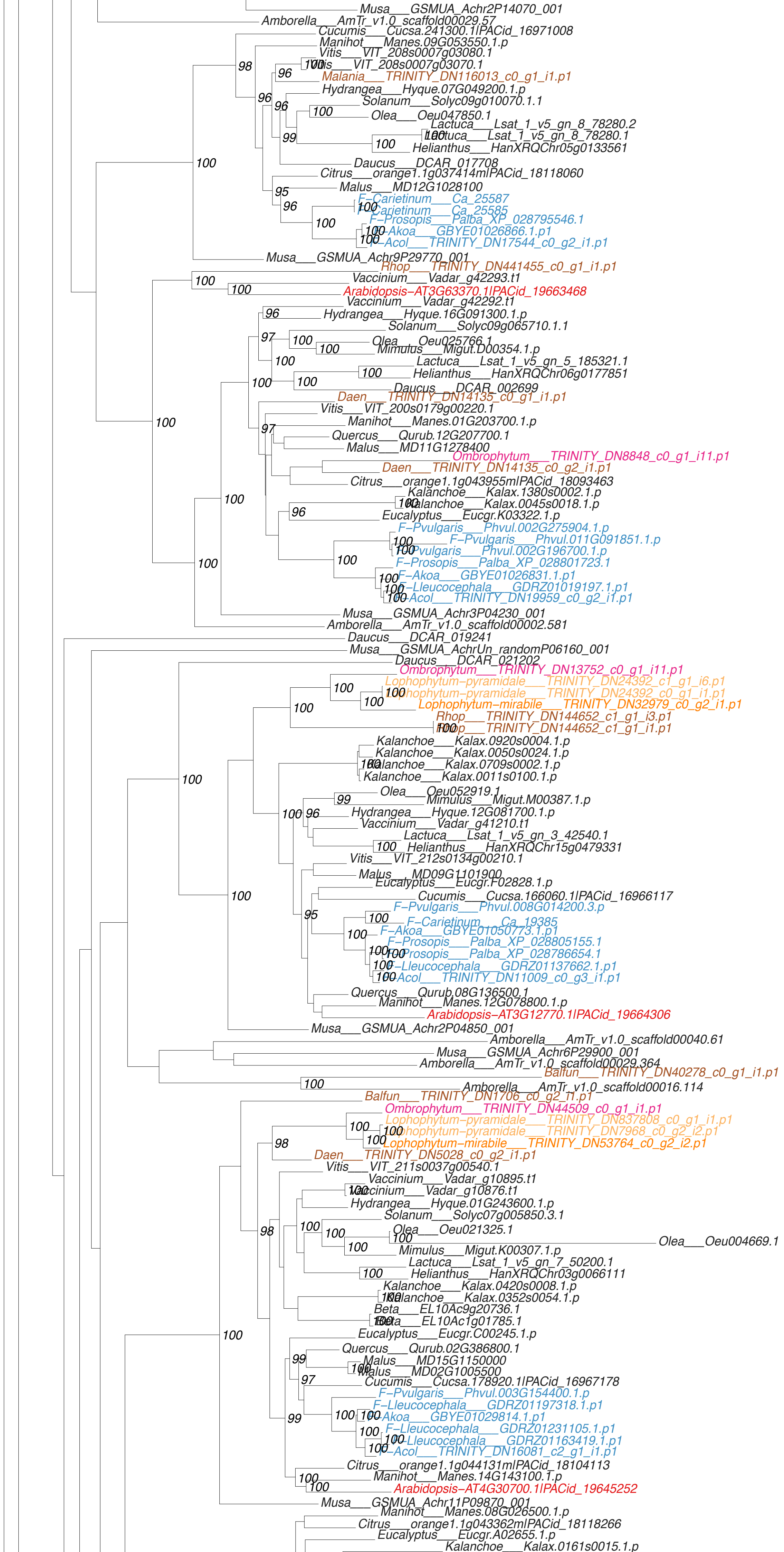

Musa___GSMUA_Achr2P14070_001
Amborella___AmTr_v1.0_scaffold00029.57
Cucumis___Cucsa.241300.1IPACid_16971008
Manihot___Manes.09G053550.1.p
Vitis___VIT_208s0007g03080.1
Vitis___VIT_208s0007g03070.1
Malania___TRINITY_DN116013_c0_g1_i1.p1
Hydrangea___Hyque.07G049200.1.p
Solanum___Solyc09g010070.1.1
Olea___Oeu047850.1
Lactuca___Lsat_1_v5_gn_8_78280.2
Lactuca___Lsat_1_v5_gn_8_78280.1
Helianthus___HanXRQChr05g0133561
Daucus___DCAR_017708
Citrus___orange1.1g037414mIPACid_18118060
Malus___MD12G1028100
F–Carietinum___Ca_25587
F–Carietinum___Ca_25585
F–Prosopis___Palba_XP_028795546.1
F–Akoa___GBYE01026866.1.p1
F–Acol___TRINITY_DN17544_c0_g2_i1.p1
Musa___GSMUA_Achr9P29770_001
Rhop___TRINITY_DN441455_c0_g1_i1.p1
Vaccinium___Vadar_g42293.t1
Arabidopsis–AT3G63370.1IPACid_19663468
Vaccinium___Vadar_g42292.t1
Hydrangea___Hyque.16G091300.1.p
Solanum___Solyc09g065710.1.1
Olea___Oeu025766.1
Mimulus___Migut.D00354.1.p
Lactuca___Lsat_1_v5_gn_5_185321.1
Helianthus___HanXRQChr06g0177851
Daucus___DCAR_002699
Daen___TRINITY_DN14135_c0_g1_i1.p1
Vitis___VIT_200s0179g00220.1
Manihot___Manes.01G203700.1.p
Quercus___Qurub.12G207700.1
Malus___MD11G1278400
Ombrophytum___TRINITY_DN8848_c0_g1_i11.p1
Daen___TRINITY_DN14135_c0_g2_i1.p1
Citrus___orange1.1g043955mIPACid_18093463
Kalanchoe___Kalax.1380s0002.1.p
Kalanchoe___Kalax.0045s0018.1.p
Eucalyptus___Eucgr.K03322.1.p
F–Pvulgaris___Phvul.002G275904.1.p
F–Pvulgaris___Phvul.011G091851.1.p
F–Pvulgaris___Phvul.002G196700.1.p
F–Prosopis___Palba_XP_028801723.1
F–Akoa___GBYE01026831.1.p1
F–Lleucocephala___GDRZ01019197.1.p1
F–Acol___TRINITY_DN19959_c0_g2_i1.p1
Musa___GSMUA_Achr3P04230_001
Amborella___AmTr_v1.0_scaffold00002.581
Daucus___DCAR_019241
Musa___GSMUA_AchrUn_randomP06160_001
Daucus___DCAR_021202
Ombrophytum___TRINITY_DN13752_c0_g1_i11.p1
Lophophytum–pyramidale___TRINITY_DN24392_c1_g1_i6.p1
Lophophytum–pyramidale___TRINITY_DN24392_c0_g1_i1.p1
Lophophytum–mirabile___TRINITY_DN32979_c0_g2_i1.p1
Rhop___TRINITY_DN144652_c1_g1_i3.p1
Rhop___TRINITY_DN144652_c1_g1_i1.p1
Kalanchoe___Kalax.0920s0004.1.p
Kalanchoe___Kalax.0050s0024.1.p
Kalanchoe___Kalax.0709s0002.1.p
Kalanchoe___Kalax.0011s0100.1.p
Olea___Oeu052919.1
Mimulus___Migut.M00387.1.p
Hydrangea___Hyque.12G081700.1.p
Vaccinium___Vadar_g41210.t1
Lactuca___Lsat_1_v5_gn_3_42540.1
Helianthus___HanXRQChr15g0479331
Vitis___VIT_212s0134g00210.1
Malus___MD09G1101900
Eucalyptus___Eucgr.F02828.1.p
Cucumis___Cucsa.166060.1IPACid_16966117
F–Pvulgaris___Phvul.008G014200.3.p
F–Carietinum___Ca_19385
F–Akoa___GBYE01050773.1.p1
F–Prosopis___Palba_XP_028805155.1
F–Prosopis___Palba_XP_028786654.1
F–Lleucocephala___GDRZ01137662.1.p1
F–Acol___TRINITY_DN11009_c0_g3_i1.p1
Quercus___Qurub.08G136500.1
Manihot___Manes.12G078800.1.p
Arabidopsis–AT3G12770.1IPACid_19664306
Musa___GSMUA_Achr2P04850_001
Amborella___AmTr_v1.0_scaffold00040.61
Musa___GSMUA_Achr6P29900_001
Amborella___AmTr_v1.0_scaffold00029.364
Balfun___TRINITY_DN40278_c0_g1_i1.p1
Amborella___AmTr_v1.0_scaffold00016.114
Balfun___TRINITY_DN1706_c0_g2_i1.p1
Ombrophytum___TRINITY_DN44509_c0_g1_i1.p1
Lophophytum–pyramidale___TRINITY_DN837808_c0_g1_i1.p1
Lophophytum–pyramidale___TRINITY_DN7968_c0_g2_i2.p1
Lophophytum–mirabile___TRINITY_DN53764_c0_g2_i2.p1
Daen___TRINITY_DN5028_c0_g2_i1.p1
Vitis___VIT_211s0037g00540.1
Vaccinium___Vadar_g10895.t1
Vaccinium___Vadar_g10876.t1
Hydrangea___Hyque.01G243600.1.p
Solanum___Solyc07g005850.3.1
Olea___Oeu021325.1
Olea___Oeu004669.1
Mimulus___Migut.K00307.1.p
Lactuca___Lsat_1_v5_gn_7_50200.1
Helianthus___HanXRQChr03g0066111
Kalanchoe___Kalax.0420s0008.1.p
Kalanchoe___Kalax.0352s0054.1.p
Beta___EL10Ac9g20736.1
Beta___EL10Ac1g01785.1
Eucalyptus___Eucgr.C00245.1.p
Quercus___Qurub.02G386800.1
Malus___MD15G1150000
Malus___MD02G1005500
Cucumis___Cucsa.178920.1IPACid_16967178
F–Pvulgaris___Phvul.003G154400.1.p
F–Lleucocephala___GDRZ01197318.1.p1
F–Akoa___GBYE01029814.1.p1
F–Lleucocephala___GDRZ01231105.1.p1
F–Lleucocephala___GDRZ01163419.1.p1
F–Acol___TRINITY_DN16081_c2_g1_i1.p1
Citrus___orange1.1g044131mIPACid_18104113
Manihot___Manes.14G143100.1.p
Arabidopsis–AT4G30700.1IPACid_19645252
Musa___GSMUA_Achr11P09870_001
Manihot___Manes.08G026500.1.p
Citrus___orange1.1g043362mIPACid_18118266
Eucalyptus___Eucgr.A02655.1.p
Kalanchoe___Kalax.0161s0015.1.p

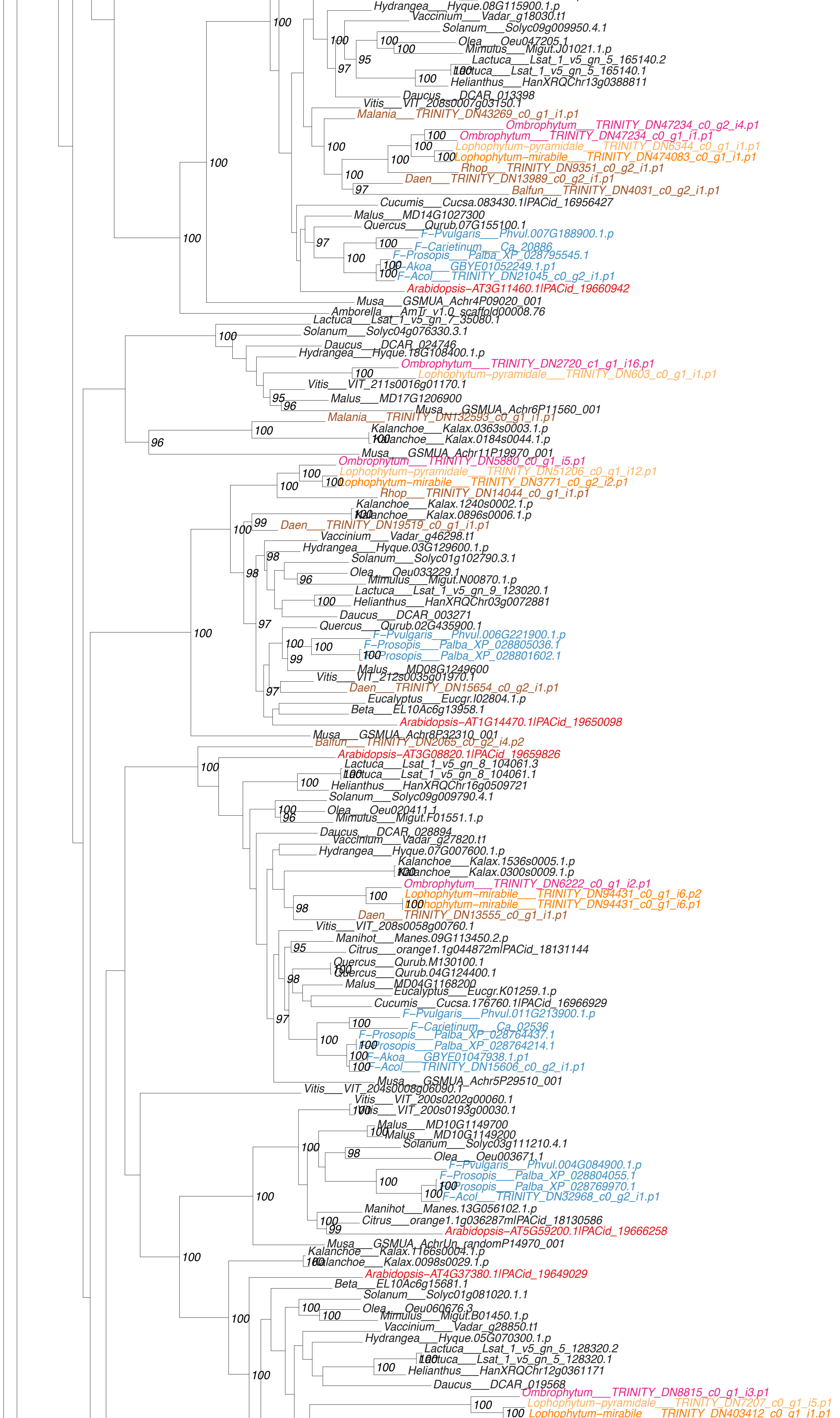
Hydrangea___Hyque.08G115900.1.p
Vaccinium___Vadar_g18030.t1
Solanum___Solyc09g009950.4.1
Olea___Oeu047205.1
Mimulus___Migut.J01021.1.p
Lactuca___Lsat_1_v5_gn_5_165140.2
Lactuca___Lsat_1_v5_gn_5_165140.1
Helianthus___HanXRQChr13g0388811
Daucus___DCAR_013398
Vitis___VIT_208s0007g03150.1
Malania___TRINITY_DN43269_c0_g1_i1.p1
Ombrophytum___TRINITY_DN47234_c0_g2_i4.p1
Ombrophytum___TRINITY_DN47234_c0_g1_i1.p1
Lophophytum−pyramidale___TRINITY_DN6344_c0_g1_i1.p1
Lophophytum−mirabile___TRINITY_DN474083_c0_g1_i1.p1
Rhop___TRINITY_DN9351_c0_g2_i1.p1
Daen___TRINITY_DN13989_c0_g2_i1.p1
Balfun___TRINITY_DN4031_c0_g2_i1.p1
Cucumis___Cucsa.083430.1IPACid_16956427
Malus___MD14G1027300
Quercus___Qurub.07G155100.1
F−Pvulgaris___Phvul.007G188900.1.p
F−Carietinum___Ca_20886
F−Prosopis___Palba_XP_028795545.1
F−Akoa___GBYE01052249.1.p1
F−Acol___TRINITY_DN21045_c0_g2_i1.p1
Arabidopsis−AT3G11460.1IPACid_19660942
Musa___GSMUA_Achr4P09020_001
Amborella___AmTr_v1.0_scaffold00008.76
Lactuca___Lsat_1_v5_gn_7_35080.1
Solanum___Solyc04g076330.3.1
Daucus___DCAR_024746
Hydrangea___Hyque.18G108400.1.p
Ombrophytum___TRINITY_DN2720_c1_g1_i16.p1
Lophophytum−pyramidale___TRINITY_DN603_c0_g1_i1.p1
Vitis___VIT_211s0016g01170.1
Malus___MD17G1206900
Musa___GSMUA_Achr6P11560_001
Malania___TRINITY_DN132593_c0_g1_i1.p1
Kalanchoe___Kalax.0363s0003.1.p
Kalanchoe___Kalax.0184s0044.1.p
Musa___GSMUA_Achr11P19970_001
Ombrophytum___TRINITY_DN5880_c0_g1_i5.p1
Lophophytum−pyramidale___TRINITY_DN51206_c0_g1_i12.p1
Lophophytum−mirabile___TRINITY_DN3771_c0_g2_i2.p1
Rhop___TRINITY_DN14044_c0_g1_i1.p1
Kalanchoe___Kalax.1240s0002.1.p
Kalanchoe___Kalax.0896s0006.1.p
Daen___TRINITY_DN19519_c0_g1_i1.p1
Vaccinium___Vadar_g46298.t1
Hydrangea___Hyque.03G129600.1.p
Solanum___Solyc01g102790.3.1
Olea___Oeu033229.1
Mimulus___Migut.N00870.1.p
Lactuca___Lsat_1_v5_gn_9_123020.1
Helianthus___HanXRQChr03g0072881
Daucus___DCAR_003271
Quercus___Qurub.02G435900.1
F−Pvulgaris___Phvul.006G221900.1.p
F−Prosopis___Palba_XP_028805036.1
F−Prosopis___Palba_XP_028801602.1
Malus___MD08G1249600
Vitis___VIT_212s0035g01970.1
Daen___TRINITY_DN15654_c0_g2_i1.p1
Eucalyptus___Eucgr.I02804.1.p
Beta___EL10Ac6g13958.1
Arabidopsis−AT1G14470.1IPACid_19650098
Musa___GSMUA_Achr8P32310_001
Balfun___TRINITY_DN2065_c0_g2_i4.p2
Arabidopsis−AT3G08820.1IPACid_19659826
Lactuca___Lsat_1_v5_gn_8_104061.3
Lactuca___Lsat_1_v5_gn_8_104061.1
Helianthus___HanXRQChr16g0509721
Solanum___Solyc09g009790.4.1
Olea___Oeu020411.1
Mimulus___Migut.F01551.1.p
Daucus___DCAR_028894
Vaccinium___Vadar_g27820.t1
Hydrangea___Hyque.07G007600.1.p
Kalanchoe___Kalax.1536s0005.1.p
Kalanchoe___Kalax.0300s0009.1.p
Ombrophytum___TRINITY_DN6222_c0_g1_i2.p1
Lophophytum−mirabile___TRINITY_DN94431_c0_g1_i6.p2
Lophophytum−mirabile___TRINITY_DN94431_c0_g1_i6.p1
Daen___TRINITY_DN13555_c0_g1_i1.p1
Vitis___VIT_208s0058g00760.1
Manihot___Manes.09G113450.2.p
Citrus___orange1.1g044872mIPACid_18131144
Quercus___Qurub.M130100.1
Quercus___Qurub.04G124400.1
Malus___MD04G1168200
Eucalyptus___Eucgr.K01259.1.p
Cucumis___Cucsa.176760.1IPACid_16966929
F−Pvulgaris___Phvul.011G213900.1.p
F−Carietinum___Ca_02536
F−Prosopis___Palba_XP_028764437.1
F−Prosopis___Palba_XP_028764214.1
F−Akoa___GBYE01047938.1.p1
F−Acol___TRINITY_DN15606_c0_g2_i1.p1
Musa___GSMUA_Achr5P29510_001
Vitis___VIT_204s0008g06090.1
Vitis___VIT_200s0202g00060.1
Vitis___VIT_200s0193g00030.1
Malus___MD10G1149700
Malus___MD10G1149200
Solanum___Solyc03g111210.4.1
Olea___Oeu003671.1
F−Pvulgaris___Phvul.004G084900.1.p
F−Prosopis___Palba_XP_028804055.1
F−Prosopis___Palba_XP_028769970.1
F−Acol___TRINITY_DN32968_c0_g2_i1.p1
Manihot___Manes.13G056102.1.p
Citrus___orange1.1g036287mIPACid_18130586
Arabidopsis−AT5G59200.1IPACid_19666258
Musa___GSMUA_AchrUn_randomP14970_001
Kalanchoe___Kalax.1166s0004.1.p
Kalanchoe___Kalax.0098s0029.1.p
Arabidopsis−AT4G37380.1IPACid_19649029
Beta___EL10Ac6g15681.1
Solanum___Solyc01g081020.1.1
Olea___Oeu060676.3
Mimulus___Migut.B01450.1.p
Vaccinium___Vadar_g28850.t1
Hydrangea___Hyque.05G070300.1.p
Lactuca___Lsat_1_v5_gn_5_128320.2
Lactuca___Lsat_1_v5_gn_5_128320.1
Helianthus___HanXRQChr12g0361171
Daucus___DCAR_019568
Ombrophytum___TRINITY_DN8815_c0_g1_i3.p1
Lophophytum−pyramidale___TRINITY_DN7207_c0_g1_i5.p1
Lophophytum−mirabile___TRINITY_DN403412_c0_g1_i1.p1

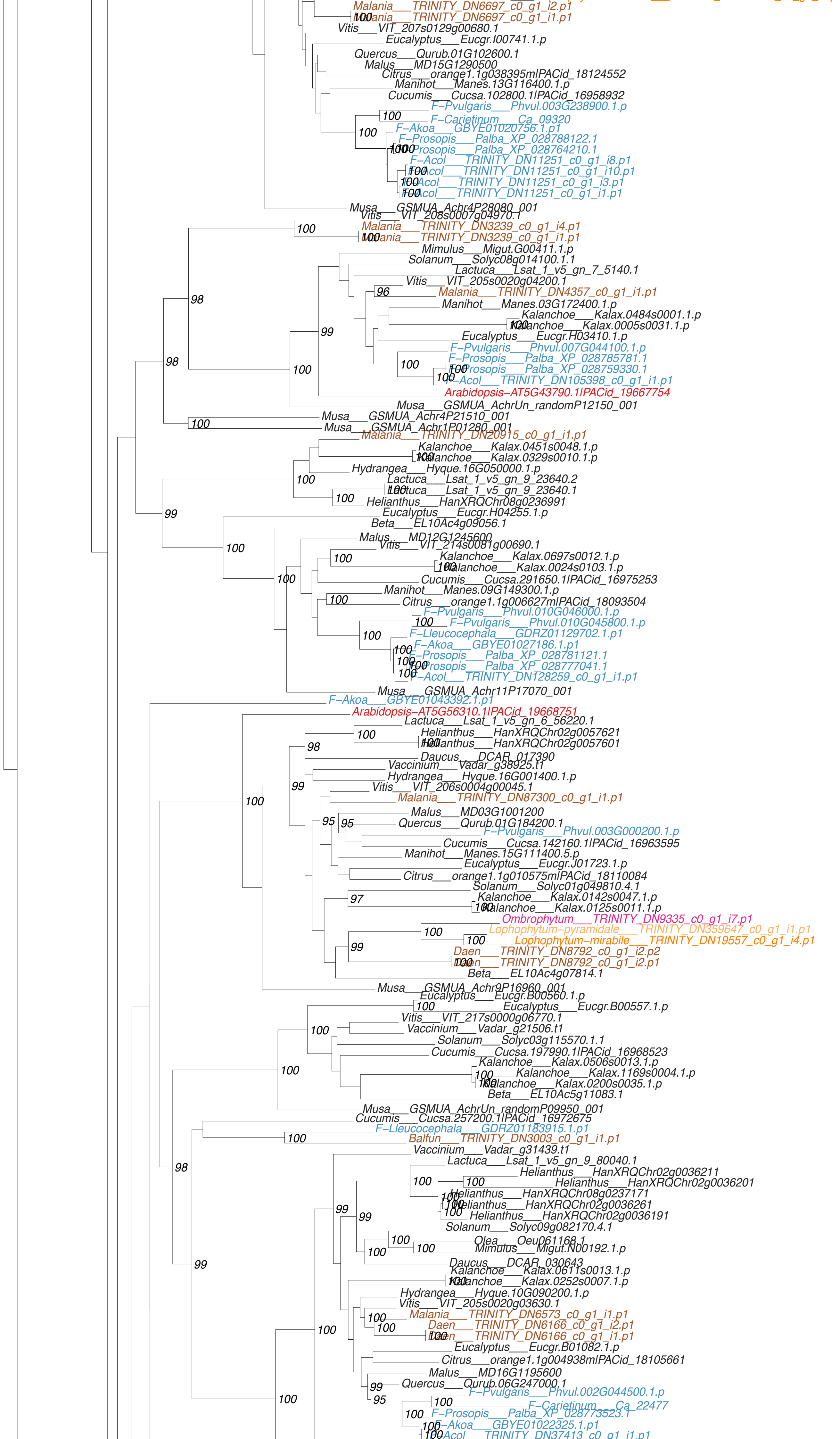

Malania___TRINITY_DN6697_c0_g1_i2.p1
Malania___TRINITY_DN6697_c0_g1_i1.p1
Vitis___VIT_207s0129g00680.1
Eucalyptus___Eucgr.I00741.1.p
Quercus___Qurub.01G102600.1
Malus___MD15G1290500
Citrus___orange1.1g038395mIPACid_18124552
Manihot___Manes.13G116400.1.p
Cucumis___Cucsa.102800.1IPACid_16958932
F-Pvulgaris___Phvul.003G238900.1.p
F-Carietinum___Ca_09320
F-Akoa___GBYE01020756.1.p1
F-Prosopis___Palba_XP_028788122.1
F-Prosopis___Palba_XP_028764210.1
F-Acol___TRINITY_DN11251_c0_g1_i8.p1
F-Acol___TRINITY_DN11251_c0_g1_i10.p1
F-Acol___TRINITY_DN11251_c0_g1_i3.p1
F-Acol___TRINITY_DN11251_c0_g1_i1.p1
Musa___GSMUA_Achr4P28080_001
Vitis___VIT_208s0007g04970.1
Malania___TRINITY_DN3239_c0_g1_i4.p1
Malania___TRINITY_DN3239_c0_g1_i1.p1
Mimulus___Migut.G00411.1.p
Solanum___Solyc08g014100.1.1
Lactuca___Lsat_1_v5_gn_7_5140.1
Vitis___VIT_205s0020g04200.1
Malania___TRINITY_DN4357_c0_g1_i1.p1
Manihot___Manes.03G172400.1.p
Kalanchoe___Kalax.0484s0001.1.p
Kalanchoe___Kalax.0005s0031.1.p
Eucalyptus___Eucgr.H03410.1.p
F-Pvulgaris___Phvul.007G044100.1.p
F-Prosopis___Palba_XP_028785781.1
F-Prosopis___Palba_XP_028759330.1
F-Acol___TRINITY_DN105398_c0_g1_i1.p1
Arabidopsis-AT5G43790.1IPACid_19667754
Musa___GSMUA_AchrUn_randomP12150_001
Musa___GSMUA_Achr4P21510_001
Musa___GSMUA_Achr1P01280_001
Malania___TRINITY_DN20915_c0_g1_i1.p1
Kalanchoe___Kalax.0451s0048.1.p
Kalanchoe___Kalax.0329s0010.1.p
Hydrangea___Hyque.16G050000.1.p
Lactuca___Lsat_1_v5_gn_9_23640.2
Lactuca___Lsat_1_v5_gn_9_23640.1
Helianthus___HanXRQChr08g0236991
Eucalyptus___Eucgr.H04255.1.p
Beta___EL10Ac4g09056.1
Malus___MD12G1245600
Vitis___VIT_214s0081g00690.1
Kalanchoe___Kalax.0697s0012.1.p
Kalanchoe___Kalax.0024s0103.1.p
Cucumis___Cucsa.291650.1IPACid_16975253
Manihot___Manes.09G149300.1.p
Citrus___orange1.1g006627mIPACid_18093504
F-Pvulgaris___Phvul.010G046000.1.p
F-Pvulgaris___Phvul.010G045800.1.p
F-Lleucocephala___GDRZ01129702.1.p1
F-Akoa___GBYE01027186.1.p1
F-Prosopis___Palba_XP_028781121.1
F-Prosopis___Palba_XP_028777041.1
F-Acol___TRINITY_DN128259_c0_g1_i1.p1
Musa___GSMUA_Achr11P17070_001
F-Akoa___GBYE01043392.1.p1
Arabidopsis-AT5G56310.1IPACid_19668751
Lactuca___Lsat_1_v5_gn_6_56220.1
Helianthus___HanXRQChr02g0057621
Helianthus___HanXRQChr02g0057601
Daucus___DCAR_017390
Vaccinium___Vadar_g38925.t1
Hydrangea___Hyque.16G001400.1.p
Vitis___VIT_206s0004g00045.1
Malania___TRINITY_DN87300_c0_g1_i1.p1
Malus___MD03G1001200
Quercus___Qurub.01G184200.1
F-Pvulgaris___Phvul.003G000200.1.p
Cucumis___Cucsa.142160.1IPACid_16963595
Manihot___Manes.15G111400.5.p
Eucalyptus___Eucgr.J01723.1.p
Citrus___orange1.1g010575mIPACid_18110084
Solanum___Solyc01g049810.4.1
Kalanchoe___Kalax.0142s0047.1.p
Kalanchoe___Kalax.0125s0011.1.p
Ombrophytum___TRINITY_DN9335_c0_g1_i7.p1
Lophophytum-pyramidale___TRINITY_DN359647_c0_g1_i1.p1
Lophophytum-mirabile___TRINITY_DN19557_c0_g1_i4.p1
Daen___TRINITY_DN8792_c0_g1_i2.p2
Daen___TRINITY_DN8792_c0_g1_i2.p1
Beta___EL10Ac4g07814.1
Musa___GSMUA_Achr9P16960_001
Eucalyptus___Eucgr.B00560.1.p
Eucalyptus___Eucgr.B00557.1.p
Vitis___VIT_217s0000g06770.1
Vaccinium___Vadar_g21506.t1
Solanum___Solyc03g115570.1.1
Cucumis___Cucsa.197990.1IPACid_16968523
Kalanchoe___Kalax.0506s0013.1.p
Kalanchoe___Kalax.1169s0004.1.p
Kalanchoe___Kalax.0200s0035.1.p
Beta___EL10Ac5g11083.1
Musa___GSMUA_AchrUn_randomP09950_001
Cucumis___Cucsa.257200.1IPACid_16972675
F-Lleucocephala___GDRZ01183915.1.p1
Balfun___TRINITY_DN3003_c0_g1_i1.p1
Vaccinium___Vadar_g31439.t1
Lactuca___Lsat_1_v5_gn_9_80040.1
Helianthus___HanXRQChr02g0036211
Helianthus___HanXRQChr02g0036201
Helianthus___HanXRQChr08g0237171
Helianthus___HanXRQChr02g0036261
Helianthus___HanXRQChr02g0036191
Solanum___Solyc09g082170.4.1
Olea___Oeu061168.1
Mimulus___Migut.N00192.1.p
Daucus___DCAR_030643
Kalanchoe___Kalax.0611s0013.1.p
Kalanchoe___Kalax.0252s0007.1.p
Hydrangea___Hyque.10G090200.1.p
Vitis___VIT_205s0020g03630.1
Malania___TRINITY_DN6573_c0_g1_i1.p1
Daen___TRINITY_DN6166_c0_g1_i2.p1
Daen___TRINITY_DN6166_c0_g1_i1.p1
Eucalyptus___Eucgr.B01082.1.p
Citrus___orange1.1g004938mIPACid_18105661
Malus___MD16G1195600
Quercus___Qurub.06G247000.1
F-Pvulgaris___Phvul.002G044500.1.p
F-Carietinum___Ca_22477
F-Prosopis___Palba_XP_028773523.1
F-Akoa___GBYE01022325.1.p1
F-Acol___TRINITY_DN37413_c0_g1_i1.p1

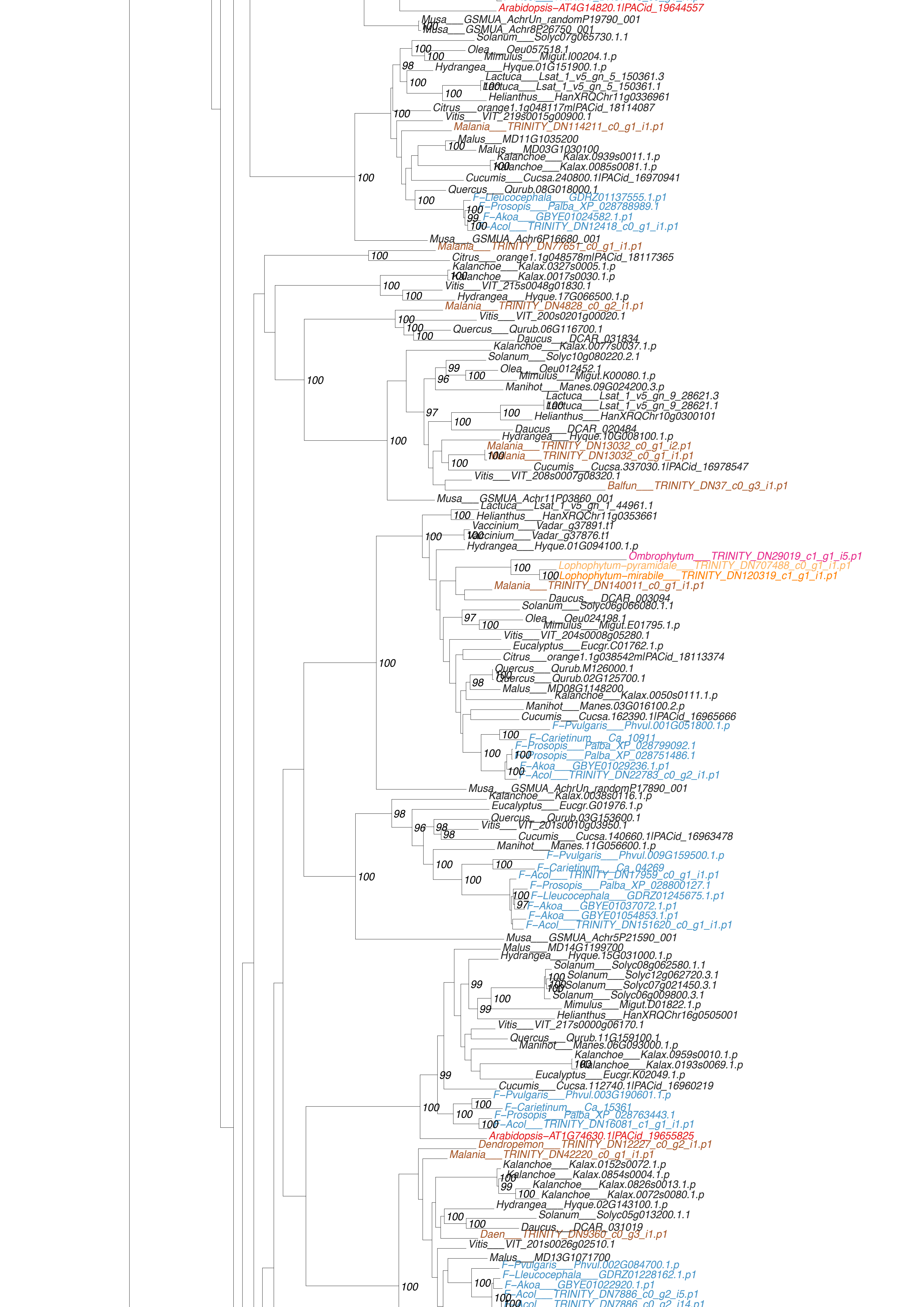
Arabidopsis–AT4G14820.1lPACid_19644557
Musa___GSMUA_AchrUn_randomP19790_001
Musa___GSMUA_Achr8P26750_001
Solanum___Solyc07g065730.1.1
Olea___Oeu057518.1
Mimulus___Migut.I00204.1.p
Hydrangea___Hyque.01G151900.1.p
Lactuca___Lsat_1_v5_gn_5_150361.3
Lactuca___Lsat_1_v5_gn_5_150361.1
Helianthus___HanXRQChr11g0336961
Citrus___orange1.1g048117mlPACid_18114087
Vitis___VIT_219s0015g00900.1
Malania___TRINITY_DN114211_c0_g1_i1.p1
Malus___MD11G1035200
Malus___MD03G1030100
Kalanchoe___Kalax.0939s0011.1.p
Kalanchoe___Kalax.0085s0081.1.p
Cucumis___Cucsa.240800.1lPACid_16970941
Quercus___Qurub.08G018000.1
F–Lleucocephala___GDRZ01137555.1.p1
F–Prosopis___Palba_XP_028788989.1
F–Akoa___GBYE01024582.1.p1
F–Acol___TRINITY_DN12418_c0_g1_i1.p1
Musa___GSMUA_Achr6P16680_001
Malania___TRINITY_DN7765l_c0_g1_i1.p1
Citrus___orange1.1g048578mlPACid_18117365
Kalanchoe___Kalax.0327s0005.1.p
Kalanchoe___Kalax.0017s0030.1.p
Vitis___VIT_215s0048g01830.1
Hydrangea___Hyque.17G066500.1.p
Malania___TRINITY_DN4828_c0_g2_i1.p1
Vitis___VIT_200s0201g00020.1
Quercus___Qurub.06G116700.1
Daucus___DCAR_031834
Kalanchoe___Kalax.0077s0037.1.p
Solanum___Solyc10g080220.2.1
Olea___Oeu012452.1
Mimulus___Migut.K00080.1.p
Manihot___Manes.09G024200.3.p
Lactuca___Lsat_1_v5_gn_9_28621.3
Lactuca___Lsat_1_v5_gn_9_28621.1
Helianthus___HanXRQChr10g0300101
Daucus___DCAR_020484
Hydrangea___Hyque.10G008100.1.p
Malania___TRINITY_DN13032_c0_g1_i2.p1
Malania___TRINITY_DN13032_c0_g1_i1.p1
Cucumis___Cucsa.337030.1lPACid_16978547
Vitis___VIT_208s0007g08320.1
Balfun___TRINITY_DN37_c0_g3_i1.p1
Musa___GSMUA_Achr11P03860_001
Lactuca___Lsat_1_v5_gn_1_44961.1
Helianthus___HanXRQChr11g0353661
Vaccinium___Vadar_g37891.t1
Vaccinium___Vadar_g37876.t1
Hydrangea___Hyque.01G094100.1.p
Ombrophytum___TRINITY_DN29019_c1_g1_i5.p1
Lophophytum–pyramidale___TRINITY_DN707488_c0_g1_i1.p1
Lophophytum–mirabile___TRINITY_DN120319_c1_g1_i1.p1
Malania___TRINITY_DN140011_c0_g1_i1.p1
Daucus___DCAR_003094
Solanum___Solyc06g066080.1.1
Olea___Oeu024198.1
Mimulus___Migut.E01795.1.p
Vitis___VIT_204s0008g05280.1
Eucalyptus___Eucgr.C01762.1.p
Citrus___orange1.1g038542mlPACid_18113374
Quercus___Qurub.M126000.1
Quercus___Qurub.02G125700.1
Malus___MD08G1148200
Kalanchoe___Kalax.0050s0111.1.p
Manihot___Manes.03G016100.2.p
Cucumis___Cucsa.162390.1lPACid_16965666
F–Pvulgaris___Phvul.001G051800.1.p
F–Carietinum___Ca_10911
F–Prosopis___Palba_XP_028799092.1
F–Prosopis___Palba_XP_028751486.1
F–Akoa___GBYE01029236.1.p1
F–Acol___TRINITY_DN22783_c0_g2_i1.p1
Musa___GSMUA_AchrUn_randomP17890_001
Kalanchoe___Kalax.0038s0116.1.p
Eucalyptus___Eucgr.G01976.1.p
Quercus___Qurub.03G153600.1
Vitis___VIT_201s0010g03950.1
Cucumis___Cucsa.140660.1lPACid_16963478
Manihot___Manes.11G056600.1.p
F–Pvulgaris___Phvul.009G159500.1.p
F–Carietinum___Ca_04269
F–Acol___TRINITY_DN17959_c0_g1_i1.p1
F–Prosopis___Palba_XP_028800127.1
F–Lleucocephala___GDRZ01245675.1.p1
F–Akoa___GBYE01037072.1.p1
F–Akoa___GBYE01054853.1.p1
F–Acol___TRINITY_DN151620_c0_g1_i1.p1
Musa___GSMUA_Achr5P21590_001
Malus___MD14G1199700
Hydrangea___Hyque.15G031000.1.p
Solanum___Solyc08g062580.1.1
Solanum___Solyc12g062720.3.1
Solanum___Solyc07g021450.3.1
Solanum___Solyc06g009800.3.1
Mimulus___Migut.D01822.1.p
Helianthus___HanXRQChr16g0505001
Vitis___VIT_217s0000g06170.1
Quercus___Qurub.11G159100.1
Manihot___Manes.06G093000.1.p
Kalanchoe___Kalax.0959s0010.1.p
Kalanchoe___Kalax.0193s0069.1.p
Eucalyptus___Eucgr.K02049.1.p
Cucumis___Cucsa.112740.1lPACid_16960219
F–Pvulgaris___Phvul.003G190601.1.p
F–Carietinum___Ca_15361
F–Prosopis___Palba_XP_028763443.1
F–Acol___TRINITY_DN16081_c1_g1_i1.p1
Arabidopsis–AT1G74630.1lPACid_19655825
Dendropemon___TRINITY_DN12227_c0_g2_i1.p1
Malania___TRINITY_DN42220_c0_g1_i1.p1
Kalanchoe___Kalax.0152s0072.1.p
Kalanchoe___Kalax.0854s0004.1.p
Kalanchoe___Kalax.0826s0013.1.p
Kalanchoe___Kalax.0072s0080.1.p
Hydrangea___Hyque.02G143100.1.p
Solanum___Solyc05g013200.1.1
Daucus___DCAR_031019
Daen___TRINITY_DN9360_c0_g3_i1.p1
Vitis___VIT_201s0026g02510.1
Malus___MD13G1071700
F–Pvulgaris___Phvul.002G084700.1.p
F–Lleucocephala___GDRZ01228162.1.p1
F–Akoa___GBYE01022920.1.p1
F–Acol___TRINITY_DN7886_c0_g2_i5.p1
F–Acol___TRINITY_DN7886_c0_g2_i14.p1

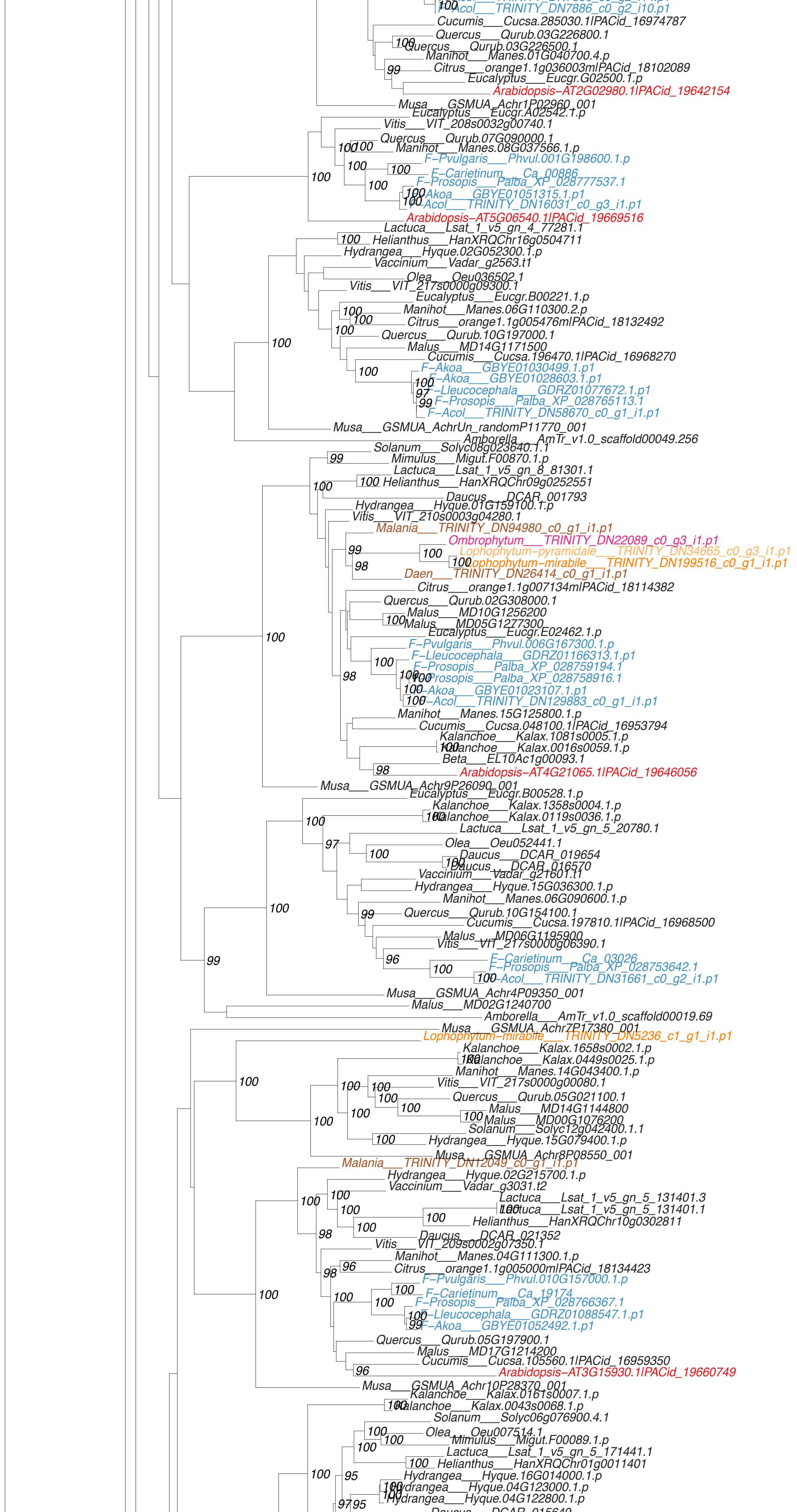
F–Acol___TRINITY_DN7886_c0_g2_i10.p1
Cucumis___Cucsa.285030.1IPACid_16974787
Quercus___Qurub.03G226800.1
Quercus___Qurub.03G226500.1
Manihot___Manes.01G040700.4.p
Citrus___orange1.1g036003mIPACid_18102089
Eucalyptus___Eucgr.G02500.1.p
Arabidopsis–AT2G02980.1IPACid_19642154
Musa___GSMUA_Achr1P02960_001
Eucalyptus___Eucgr.A02542.1.p
Vitis___VIT_208s0032g00740.1
Quercus___Qurub.07G090000.1
Manihot___Manes.08G037566.1.p
F–Pvulgaris___Phvul.001G198600.1.p
F–Carietinum___Ca_00886
F–Prosopis___Palba_XP_028777537.1
F–Akoa___GBYE01051315.1.p1
F–Acol___TRINITY_DN16031_c0_g3_i1.p1
Arabidopsis–AT5G06540.1IPACid_19669516
Lactuca___Lsat_1_v5_gn_4_77281.1
Helianthus___HanXRQChr16g0504711
Hydrangea___Hyque.02G052300.1.p
Vaccinium___Vadar_g2563.t1
Olea___Oeu036502.1
Vitis___VIT_217s0000g09300.1
Eucalyptus___Eucgr.B00221.1.p
Manihot___Manes.06G110300.2.p
Citrus___orange1.1g005476mIPACid_18132492
Quercus___Qurub.10G197000.1
Malus___MD14G1171500
Cucumis___Cucsa.196470.1IPACid_16968270
F–Akoa___GBYE01030499.1.p1
F–Akoa___GBYE01028603.1.p1
F–Lleucocephala___GDRZ01077672.1.p1
F–Prosopis___Palba_XP_028765113.1
F–Acol___TRINITY_DN58670_c0_g1_i1.p1
Musa___GSMUA_AchrUn_randomP11770_001
Amborella___AmTr_v1.0_scaffold00049.256
Solanum___Solyc08g023640.1.1
Mimulus___Migut.F00870.1.p
Lactuca___Lsat_1_v5_gn_8_81301.1
Helianthus___HanXRQChr09g0252551
Daucus___DCAR_001793
Hydrangea___Hyque.01G159100.1.p
Vitis___VIT_210s0003g04280.1
Malania___TRINITY_DN94980_c0_g1_i1.p1
Ombrophytum___TRINITY_DN22089_c0_g3_i1.p1
Lophophytum–pyramidale___TRINITY_DN34665_c0_g3_i1.p1
Lophophytum–mirabile___TRINITY_DN199516_c0_g1_i1.p1
Daen___TRINITY_DN26414_c0_g1_i1.p1
Citrus___orange1.1g007134mIPACid_18114382
Quercus___Qurub.02G308000.1
Malus___MD10G1256200
Malus___MD05G1277300
Eucalyptus___Eucgr.E02462.1.p
F–Pvulgaris___Phvul.006G167300.1.p
F–Lleucocephala___GDRZ01166313.1.p1
F–Prosopis___Palba_XP_028759194.1
F–Prosopis___Palba_XP_028758916.1
F–Akoa___GBYE01023107.1.p1
F–Acol___TRINITY_DN129883_c0_g1_i1.p1
Manihot___Manes.15G125800.1.p
Cucumis___Cucsa.048100.1IPACid_16953794
Kalanchoe___Kalax.1081s0005.1.p
Kalanchoe___Kalax.0016s0059.1.p
Beta___EL10Ac1g00093.1
Arabidopsis–AT4G21065.1IPACid_19646056
Musa___GSMUA_Achr9P26090_001
Eucalyptus___Eucgr.B00528.1.p
Kalanchoe___Kalax.1358s0004.1.p
Kalanchoe___Kalax.0119s0036.1.p
Lactuca___Lsat_1_v5_gn_5_20780.1
Olea___Oeu052441.1
Daucus___DCAR_019654
Daucus___DCAR_016570
Vaccinium___Vadar_g21601.t1
Hydrangea___Hyque.15G036300.1.p
Manihot___Manes.06G090600.1.p
Quercus___Qurub.10G154100.1
Cucumis___Cucsa.197810.1IPACid_16968500
Malus___MD06G1195900
Vitis___VIT_217s0000g06390.1
F–Carietinum___Ca_03026
F–Prosopis___Palba_XP_028753642.1
F–Acol___TRINITY_DN31661_c0_g2_i1.p1
Musa___GSMUA_Achr4P09350_001
Malus___MD02G1240700
Amborella___AmTr_v1.0_scaffold00019.69
Musa___GSMUA_Achr7P17380_001
Lophophytum–mirabile___TRINITY_DN5236_c1_g1_i1.p1
Kalanchoe___Kalax.1658s0002.1.p
Kalanchoe___Kalax.0449s0025.1.p
Manihot___Manes.14G043400.1.p
Vitis___VIT_217s0000g00080.1
Quercus___Qurub.05G021100.1
Malus___MD14G1144800
Malus___MD00G1076200
Solanum___Solyc12g042400.1.1
Hydrangea___Hyque.15G079400.1.p
Musa___GSMUA_Achr8P08550_001
Malania___TRINITY_DN12049_c0_g1_i1.p1
Hydrangea___Hyque.02G215700.1.p
Vaccinium___Vadar_g3031.t2
Lactuca___Lsat_1_v5_gn_5_131401.3
Lactuca___Lsat_1_v5_gn_5_131401.1
Helianthus___HanXRQChr10g0302811
Daucus___DCAR_021352
Vitis___VIT_209s0002g07350.1
Manihot___Manes.04G111300.1.p
Citrus___orange1.1g005000mIPACid_18134423
F–Pvulgaris___Phvul.010G157000.1.p
F–Carietinum___Ca_19174
F–Prosopis___Palba_XP_028766367.1
F–Lleucocephala___GDRZ01088547.1.p1
F–Akoa___GBYE01052492.1.p1
Quercus___Qurub.05G197900.1
Malus___MD17G1214200
Cucumis___Cucsa.105560.1IPACid_16959350
Arabidopsis–AT3G15930.1IPACid_19660749
Musa___GSMUA_Achr10P28370_001
Kalanchoe___Kalax.0161s0007.1.p
Kalanchoe___Kalax.0043s0068.1.p
Solanum___Solyc06g076900.4.1
Olea___Oeu007514.1
Mimulus___Migut.F00089.1.p
Lactuca___Lsat_1_v5_gn_5_171441.1
Helianthus___HanXRQChr01g0011401
Hydrangea___Hyque.16G014000.1.p
Hydrangea___Hyque.04G123000.1.p
Hydrangea___Hyque.04G122800.1.p

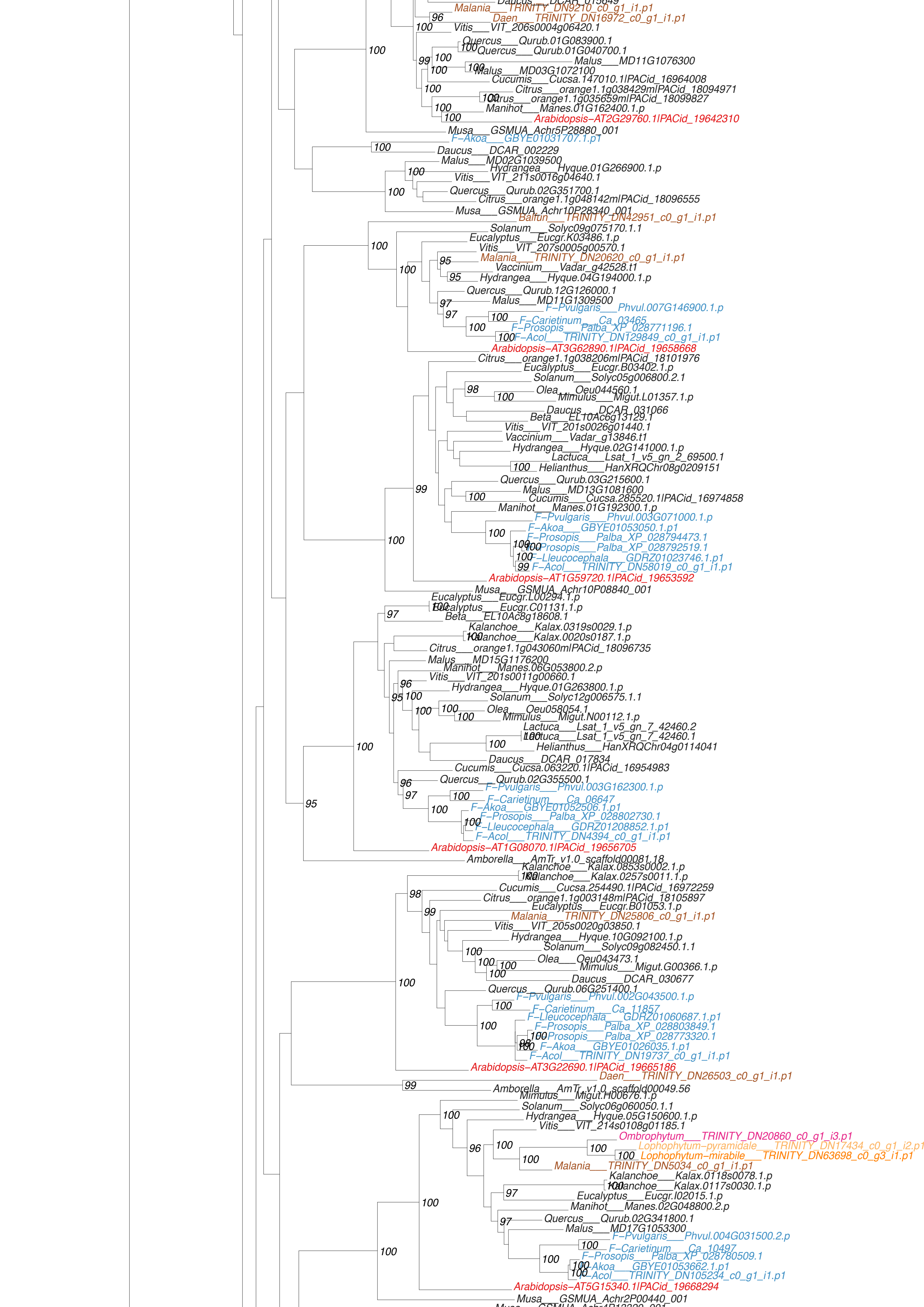

Malania___TRINITY_DN9210_c0_g1_i1.p1
Daen___TRINITY_DN16972_c0_g1_i1.p1
Vitis___VIT_206s0004g06420.1
Quercus___Qurub.01G083900.1
Quercus___Qurub.01G040700.1
Malus___MD11G1076300
Malus___MD03G1072100
Cucumis___Cucsa.147010.1lPACid_16964008
Citrus___orange1.1g038429mlPACid_18094971
Citrus___orange1.1g035659mlPACid_18099827
Manihot___Manes.01G162400.1.p
Arabidopsis−AT2G29760.1lPACid_19642310
Musa___GSMUA_Achr5P28880_001
F−Akoa___GBYE01031707.1.p1
Daucus___DCAR_002229
Malus___MD02G1039500
Hydrangea___Hyque.01G266900.1.p
Vitis___VIT_211s0016g04640.1
Quercus___Qurub.02G351700.1
Citrus___orange1.1g048142mlPACid_18096555
Musa___GSMUA_Achr10P28340_001
Balfun___TRINITY_DN42951_c0_g1_i1.p1
Solanum___Solyc09g075170.1.1
Eucalyptus___Eucgr.K03486.1.p
Vitis___VIT_207s0005g00570.1
Malania___TRINITY_DN20620_c0_g1_i1.p1
Vaccinium___Vadar_g42528.t1
Hydrangea___Hyque.04G194000.1.p
Quercus___Qurub.12G126000.1
Malus___MD11G1309500
F−Pvulgaris___Phvul.007G146900.1.p
F−Carietinum___Ca_03465
F−Prosopis___Palba_XP_028771196.1
F−Acol___TRINITY_DN129849_c0_g1_i1.p1
Arabidopsis−AT3G62890.1lPACid_19658668
Citrus___orange1.1g038206mlPACid_18101976
Eucalyptus___Eucgr.B03402.1.p
Solanum___Solyc05g006800.2.1
Olea___Oeu044560.1
Mimulus___Migut.L01357.1.p
Daucus___DCAR_031066
Beta___EL10Ac6g13129.1
Vitis___VIT_201s0026g01440.1
Vaccinium___Vadar_g13846.t1
Hydrangea___Hyque.02G141000.1.p
Lactuca___Lsat_1_v5_gn_2_69500.1
Helianthus___HanXRQChr08g0209151
Quercus___Qurub.03G215600.1
Malus___MD13G1081600
Cucumis___Cucsa.285520.1lPACid_16974858
Manihot___Manes.01G192300.1.p
F−Pvulgaris___Phvul.003G071000.1.p
F−Akoa___GBYE01053050.1.p1
F−Prosopis___Palba_XP_028794473.1
F−Prosopis___Palba_XP_028792519.1
F−Lleucocephala___GDRZ01023746.1.p1
F−Acol___TRINITY_DN58019_c0_g1_i1.p1
Arabidopsis−AT1G59720.1lPACid_19653592
Musa___GSMUA_Achr10P08840_001
Eucalyptus___Eucgr.L00294.1.p
Eucalyptus___Eucgr.C01131.1.p
Beta___EL10Ac8g18608.1
Kalanchoe___Kalax.0319s0029.1.p
Kalanchoe___Kalax.0020s0187.1.p
Citrus___orange1.1g043060mlPACid_18096735
Malus___MD15G1176200
Manihot___Manes.06G053800.2.p
Vitis___VIT_201s0011g00660.1
Hydrangea___Hyque.01G263800.1.p
Solanum___Solyc12g006575.1.1
Olea___Oeu058054.1
Mimulus___Migut.N00112.1.p
Lactuca___Lsat_1_v5_gn_7_42460.2
Lactuca___Lsat_1_v5_gn_7_42460.1
Helianthus___HanXRQChr04g0114041
Daucus___DCAR_017834
Cucumis___Cucsa.063220.1lPACid_16954983
Quercus___Qurub.02G355500.1
F−Pvulgaris___Phvul.003G162300.1.p
F−Carietinum___Ca_06647
F−Akoa___GBYE01052506.1.p1
F−Prosopis___Palba_XP_028802730.1
F−Lleucocephala___GDRZ01208852.1.p1
F−Acol___TRINITY_DN4394_c0_g1_i1.p1
Arabidopsis−AT1G08070.1lPACid_19656705
Amborella___AmTr_v1.0_scaffold00081.18
Kalanchoe___Kalax.0853s0002.1.p
Kalanchoe___Kalax.0257s0011.1.p
Cucumis___Cucsa.254490.1lPACid_16972259
Citrus___orange1.1g003148mlPACid_18105897
Eucalyptus___Eucgr.B01053.1.p
Malania___TRINITY_DN25806_c0_g1_i1.p1
Vitis___VIT_205s0020g03850.1
Hydrangea___Hyque.10G092100.1.p
Solanum___Solyc09g082450.1.1
Olea___Oeu043473.1
Mimulus___Migut.G00366.1.p
Daucus___DCAR_030677
Quercus___Qurub.06G251400.1
F−Pvulgaris___Phvul.002G043500.1.p
F−Carietinum___Ca_11857
F−Lleucocephala___GDRZ01060687.1.p1
F−Prosopis___Palba_XP_028803849.1
F−Prosopis___Palba_XP_028773320.1
F−Akoa___GBYE01026035.1.p1
F−Acol___TRINITY_DN19737_c0_g1_i1.p1
Arabidopsis−AT3G22690.1lPACid_19665186
Daen___TRINITY_DN26503_c0_g1_i1.p1
Amborella___AmTr_v1.0_scaffold00049.56
Mimulus___Migut.H00676.1.p
Solanum___Solyc06g060050.1.1
Hydrangea___Hyque.05G150600.1.p
Vitis___VIT_214s0108g01185.1
Ombrophytum___TRINITY_DN20860_c0_g1_i3.p1
Lophophytum−pyramidale___TRINITY_DN17434_c0_g1_i2.p1
Lophophytum−mirabile___TRINITY_DN63698_c0_g3_i1.p1
Malania___TRINITY_DN5034_c0_g1_i1.p1
Kalanchoe___Kalax.0118s0078.1.p
Kalanchoe___Kalax.0117s0030.1.p
Eucalyptus___Eucgr.I02015.1.p
Manihot___Manes.02G048800.2.p
Quercus___Qurub.02G341800.1
Malus___MD17G1053300
F−Pvulgaris___Phvul.004G031500.2.p
F−Carietinum___Ca_10497
F−Prosopis___Palba_XP_028780509.1
F−Akoa___GBYE01053662.1.p1
F−Acol___TRINITY_DN105234_c0_g1_i1.p1
Arabidopsis−AT5G15340.1lPACid_19668294
Musa___GSMUA_Achr2P00440_001

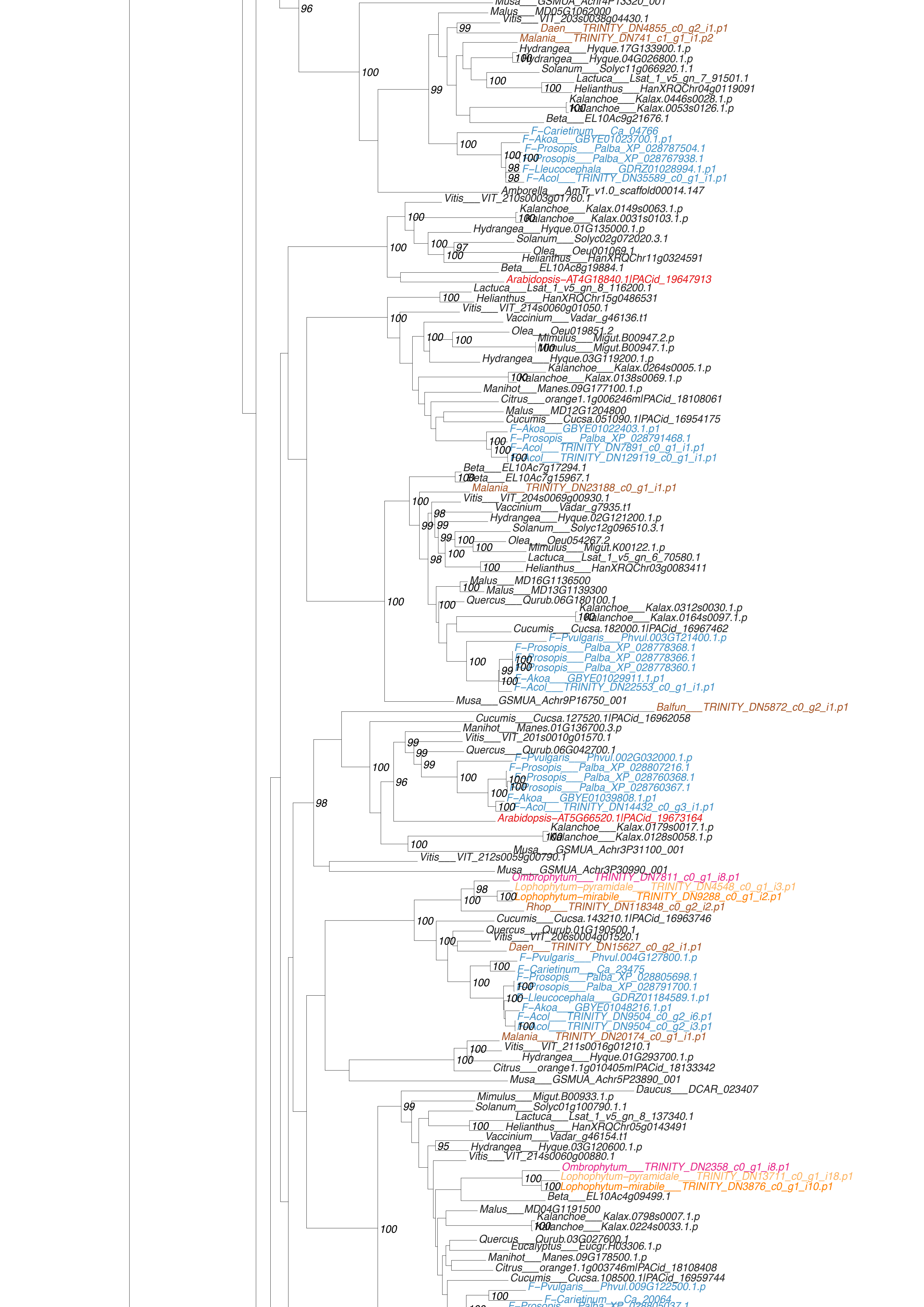
Malus___MD05G1062000
Vitis___VIT_203s0038g04430.1
Daen___TRINITY_DN4855_c0_g2_i1.p1
Malania___TRINITY_DN741_c1_g1_i1.p2
Hydrangea___Hyque.17G133900.1.p
Hydrangea___Hyque.04G026800.1.p
Solanum___Solyc11g066920.1.1
Lactuca___Lsat_1_v5_gn_7_91501.1
Helianthus___HanXRQChr04g0119091
Kalanchoe___Kalax.0446s0028.1.p
Kalanchoe___Kalax.0053s0126.1.p
Beta___EL10Ac9g21676.1
F−Carietinum___Ca_04766
F−Akoa___GBYE01023700.1.p1
F−Prosopis___Palba_XP_028787504.1
F−Prosopis___Palba_XP_028767938.1
F−Lleucocephala___GDRZ01028994.1.p1
F−Acol___TRINITY_DN35589_c0_g1_i1.p1
Amborella___AmTr_v1.0_scaffold00014.147
Vitis___VIT_210s0003g01760.1
Kalanchoe___Kalax.0149s0063.1.p
Kalanchoe___Kalax.0031s0103.1.p
Hydrangea___Hyque.01G135000.1.p
Solanum___Solyc02g072020.3.1
Olea___Oeu001069.1
Helianthus___HanXRQChr11g0324591
Beta___EL10Ac8g19884.1
Arabidopsis−AT4G18840.1lPACid_19647913
Lactuca___Lsat_1_v5_gn_8_116200.1
Helianthus___HanXRQChr15g0486531
Vitis___VIT_214s0060g01050.1
Vaccinium___Vadar_g46136.t1
Olea___Oeu019851.2
Mimulus___Migut.B00947.2.p
Mimulus___Migut.B00947.1.p
Hydrangea___Hyque.03G119200.1.p
Kalanchoe___Kalax.0264s0005.1.p
Kalanchoe___Kalax.0138s0069.1.p
Manihot___Manes.09G177100.1.p
Citrus___orange1.1g006246mlPACid_18108061
Malus___MD12G1204800
Cucumis___Cucsa.051090.1lPACid_16954175
F−Akoa___GBYE01022403.1.p1
F−Prosopis___Palba_XP_028791468.1
F−Acol___TRINITY_DN7891_c0_g1_i1.p1
F−Acol___TRINITY_DN129119_c0_g1_i1.p1
Beta___EL10Ac7g17294.1
Beta___EL10Ac7g15967.1
Malania___TRINITY_DN23188_c0_g1_i1.p1
Vitis___VIT_204s0069g00930.1
Vaccinium___Vadar_g7935.t1
Hydrangea___Hyque.02G121200.1.p
Solanum___Solyc12g096510.3.1
Olea___Oeu054267.2
Mimulus___Migut.K00122.1.p
Lactuca___Lsat_1_v5_gn_6_70580.1
Helianthus___HanXRQChr03g0083411
Malus___MD16G1136500
Malus___MD13G1139300
Quercus___Qurub.06G180100.1
Kalanchoe___Kalax.0312s0030.1.p
Kalanchoe___Kalax.0164s0097.1.p
Cucumis___Cucsa.182000.1lPACid_16967462
F−Pvulgaris___Phvul.003G121400.1.p
F−Prosopis___Palba_XP_028778368.1
F−Prosopis___Palba_XP_028778366.1
F−Prosopis___Palba_XP_028778360.1
F−Akoa___GBYE01029911.1.p1
F−Acol___TRINITY_DN22553_c0_g1_i1.p1
Musa___GSMUA_Achr9P16750_001
Balfun___TRINITY_DN5872_c0_g2_i1.p1
Cucumis___Cucsa.127520.1lPACid_16962058
Manihot___Manes.01G136700.3.p
Vitis___VIT_201s0010g01570.1
Quercus___Qurub.06G042700.1
F−Pvulgaris___Phvul.002G032000.1.p
F−Prosopis___Palba_XP_028807216.1
F−Prosopis___Palba_XP_028760368.1
F−Prosopis___Palba_XP_028760367.1
F−Akoa___GBYE01039808.1.p1
F−Acol___TRINITY_DN14432_c0_g3_i1.p1
Arabidopsis−AT5G66520.1lPACid_19673164
Kalanchoe___Kalax.0179s0017.1.p
Kalanchoe___Kalax.0128s0058.1.p
Musa___GSMUA_Achr3P31100_001
Vitis___VIT_212s0059g00790.1
Musa___GSMUA_Achr3P30990_001
Ombrophytum___TRINITY_DN7811_c0_g1_i8.p1
Lophophytum−pyramidale___TRINITY_DN4548_c0_g1_i3.p1
Lophophytum−mirabile___TRINITY_DN9288_c0_g1_i2.p1
Rhop___TRINITY_DN118348_c0_g2_i2.p1
Cucumis___Cucsa.143210.1lPACid_16963746
Quercus___Qurub.01G190500.1
Vitis___VIT_206s0004g01520.1
Daen___TRINITY_DN15627_c0_g2_i1.p1
F−Pvulgaris___Phvul.004G127800.1.p
F−Carietinum___Ca_23475
F−Prosopis___Palba_XP_028805698.1
F−Prosopis___Palba_XP_028791700.1
F−Lleucocephala___GDRZ01184589.1.p1
F−Akoa___GBYE01048216.1.p1
F−Acol___TRINITY_DN9504_c0_g2_i6.p1
F−Acol___TRINITY_DN9504_c0_g2_i3.p1
Malania___TRINITY_DN20174_c0_g1_i1.p1
Vitis___VIT_211s0016g01210.1
Hydrangea___Hyque.01G293700.1.p
Citrus___orange1.1g010405mlPACid_18133342
Musa___GSMUA_Achr5P23890_001
Daucus___DCAR_023407
Mimulus___Migut.B00933.1.p
Solanum___Solyc01g100790.1.1
Lactuca___Lsat_1_v5_gn_8_137340.1
Helianthus___HanXRQChr05g0143491
Vaccinium___Vadar_g46154.t1
Hydrangea___Hyque.03G120600.1.p
Vitis___VIT_214s0060g00880.1
Ombrophytum___TRINITY_DN2358_c0_g1_i8.p1
Lophophytum−pyramidale___TRINITY_DN13711_c0_g1_i18.p1
Lophophytum−mirabile___TRINITY_DN3876_c0_g1_i10.p1
Beta___EL10Ac4g09499.1
Malus___MD04G1191500
Kalanchoe___Kalax.0798s0007.1.p
Kalanchoe___Kalax.0224s0033.1.p
Quercus___Qurub.03G027600.1
Eucalyptus___Eucgr.H03306.1.p
Manihot___Manes.09G178500.1.p
Citrus___orange1.1g003746mlPACid_18108408
Cucumis___Cucsa.108500.1lPACid_16959744
F−Pvulgaris___Phvul.009G122500.1.p
F−Carietinum___Ca_20064

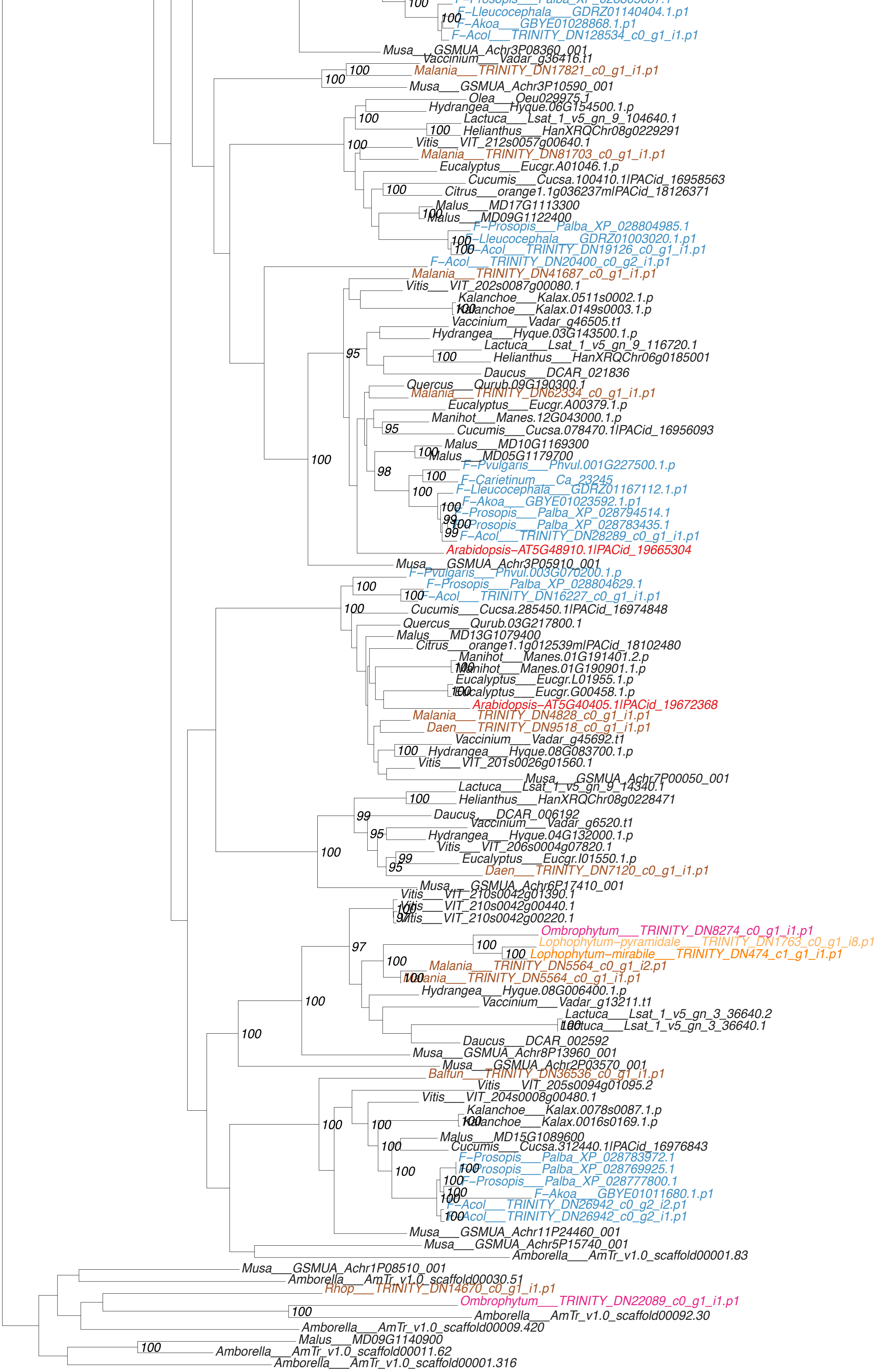

F-Lleucocephala___GDRZ01140404.1.p1
F-Akoa___GBYE01028868.1.p1
F-Acol___TRINITY_DN128534_c0_g1_i1.p1
Musa___GSMUA_Achr3P08360_001
Vaccinium___Vadar_g36416.t1
Malania___TRINITY_DN17821_c0_g1_i1.p1
Musa___GSMUA_Achr3P10590_001
Olea___Oeu029975.1
Hydrangea___Hyque.06G154500.1.p
Lactuca___Lsat_1_v5_gn_9_104640.1
Helianthus___HanXRQChr08g0229291
Vitis___VIT_212s0057g00640.1
Malania___TRINITY_DN81703_c0_g1_i1.p1
Eucalyptus___Eucgr.A01046.1.p
Cucumis___Cucsa.100410.1lPACid_16958563
Citrus___orange1.1g036237mlPACid_18126371
Malus___MD17G1113300
Malus___MD09G1122400
F-Prosopis___Palba_XP_028804985.1
F-Lleucocephala___GDRZ01003020.1.p1
F-Acol___TRINITY_DN19126_c0_g1_i1.p1
F-Acol___TRINITY_DN20400_c0_g2_i1.p1
Malania___TRINITY_DN41687_c0_g1_i1.p1
Vitis___VIT_202s0087g00080.1
Kalanchoe___Kalax.0511s0002.1.p
Kalanchoe___Kalax.0149s0003.1.p
Vaccinium___Vadar_g46505.t1
Hydrangea___Hyque.03G143500.1.p
Lactuca___Lsat_1_v5_gn_9_116720.1
Helianthus___HanXRQChr06g0185001
Daucus___DCAR_021836
Quercus___Qurub.09G190300.1
Malania___TRINITY_DN62334_c0_g1_i1.p1
Eucalyptus___Eucgr.A00379.1.p
Manihot___Manes.12G043000.1.p
Cucumis___Cucsa.078470.1lPACid_16956093
Malus___MD10G1169300
Malus___MD05G1179700
F-Pvulgaris___Phvul.001G227500.1.p
F-Carietinum___Ca_23245
F-Lleucocephala___GDRZ01167112.1.p1
F-Akoa___GBYE01023592.1.p1
F-Prosopis___Palba_XP_028794514.1
F-Prosopis___Palba_XP_028783435.1
F-Acol___TRINITY_DN28289_c0_g1_i1.p1
Arabidopsis-AT5G48910.1lPACid_19665304
Musa___GSMUA_Achr3P05910_001
F-Pvulgaris___Phvul.003G070200.1.p
F-Prosopis___Palba_XP_028804629.1
F-Acol___TRINITY_DN16227_c0_g1_i1.p1
Cucumis___Cucsa.285450.1lPACid_16974848
Quercus___Qurub.03G217800.1
Malus___MD13G1079400
Citrus___orange1.1g012539mlPACid_18102480
Manihot___Manes.01G191401.2.p
Manihot___Manes.01G190901.1.p
Eucalyptus___Eucgr.L01955.1.p
Eucalyptus___Eucgr.G00458.1.p
Arabidopsis-AT5G40405.1lPACid_19672368
Malania___TRINITY_DN4828_c0_g1_i1.p1
Daen___TRINITY_DN9518_c0_g1_i1.p1
Vaccinium___Vadar_g45692.t1
Hydrangea___Hyque.08G083700.1.p
Vitis___VIT_201s0026g01560.1
Musa___GSMUA_Achr7P00050_001
Lactuca___Lsat_1_v5_gn_9_14340.1
Helianthus___HanXRQChr08g0228471
Daucus___DCAR_006192
Vaccinium___Vadar_g6520.t1
Hydrangea___Hyque.04G132000.1.p
Vitis___VIT_206s0004g07820.1
Eucalyptus___Eucgr.I01550.1.p
Daen___TRINITY_DN7120_c0_g1_i1.p1
Musa___GSMUA_Achr6P17410_001
Vitis___VIT_210s0042g01390.1
Vitis___VIT_210s0042g00440.1
Vitis___VIT_210s0042g00220.1
Ombrophytum___TRINITY_DN8274_c0_g1_i1.p1
Lophophytum-pyramidale___TRINITY_DN1763_c0_g1_i8.p1
Lophophytum-mirabile___TRINITY_DN474_c1_g1_i1.p1
Malania___TRINITY_DN5564_c0_g1_i2.p1
Malania___TRINITY_DN5564_c0_g1_i1.p1
Hydrangea___Hyque.08G006400.1.p
Vaccinium___Vadar_g13211.t1
Lactuca___Lsat_1_v5_gn_3_36640.2
Lactuca___Lsat_1_v5_gn_3_36640.1
Daucus___DCAR_002592
Musa___GSMUA_Achr8P13960_001
Musa___GSMUA_Achr2P03570_001
Balfun___TRINITY_DN36536_c0_g1_i1.p1
Vitis___VIT_205s0094g01095.2
Vitis___VIT_204s0008g00480.1
Kalanchoe___Kalax.0078s0087.1.p
Kalanchoe___Kalax.0016s0169.1.p
Malus___MD15G1089600
Cucumis___Cucsa.312440.1lPACid_16976843
F-Prosopis___Palba_XP_028783972.1
F-Prosopis___Palba_XP_028769925.1
F-Prosopis___Palba_XP_028777800.1
F-Akoa___GBYE01011680.1.p1
F-Acol___TRINITY_DN26942_c0_g2_i2.p1
F-Acol___TRINITY_DN26942_c0_g2_i1.p1
Musa___GSMUA_Achr11P24460_001
Musa___GSMUA_Achr5P15740_001
Amborella___AmTr_v1.0_scaffold00001.83
Musa___GSMUA_Achr1P08510_001
Amborella___AmTr_v1.0_scaffold00030.51
Rhop___TRINITY_DN14670_c0_g1_i1.p1
Ombrophytum___TRINITY_DN22089_c0_g1_i1.p1
Amborella___AmTr_v1.0_scaffold00092.30
Amborella___AmTr_v1.0_scaffold00009.420
Malus___MD09G1140900
Amborella___AmTr_v1.0_scaffold00011.62
Amborella___AmTr_v1.0_scaffold00001.316

# OG0000022

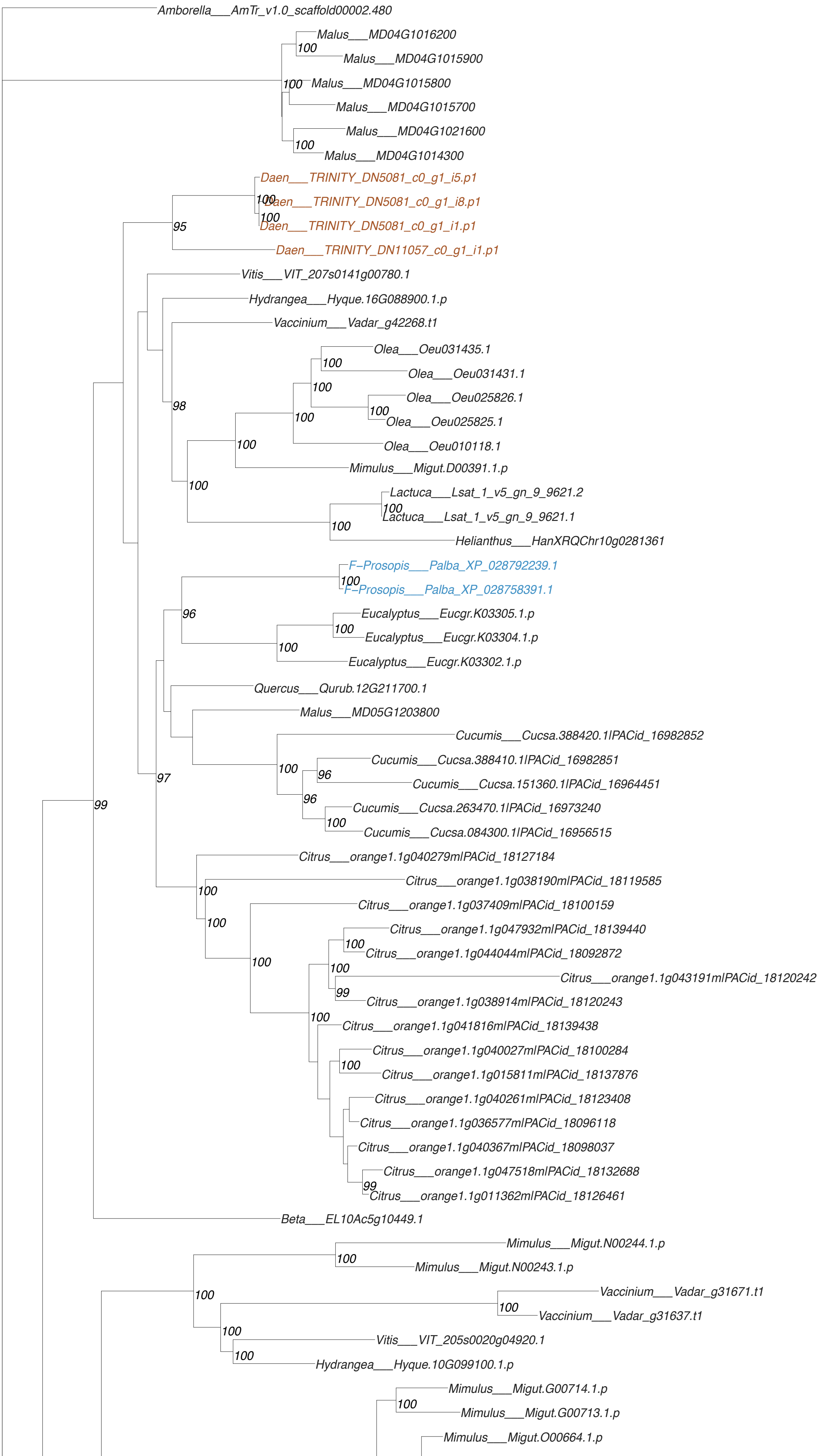

Amborella___AmTr_v1.0_scaffold00002.480
Malus___MD04G1016200
Malus___MD04G1015900
Malus___MD04G1015800
Malus___MD04G1015700
Malus___MD04G1021600
Malus___MD04G1014300
Daen___TRINITY_DN5081_c0_g1_i5.p1
Daen___TRINITY_DN5081_c0_g1_i8.p1
Daen___TRINITY_DN5081_c0_g1_i1.p1
Daen___TRINITY_DN11057_c0_g1_i1.p1
Vitis___VIT_207s0141g00780.1
Hydrangea___Hyque.16G088900.1.p
Vaccinium___Vadar_g42268.t1
Olea___Oeu031435.1
Olea___Oeu031431.1
Olea___Oeu025826.1
Olea___Oeu025825.1
Olea___Oeu010118.1
Mimulus___Migut.D00391.1.p
Lactuca___Lsat_1_v5_gn_9_9621.2
Lactuca___Lsat_1_v5_gn_9_9621.1
Helianthus___HanXRQChr10g0281361
F−Prosopis___Palba_XP_028792239.1
F−Prosopis___Palba_XP_028758391.1
Eucalyptus___Eucgr.K03305.1.p
Eucalyptus___Eucgr.K03304.1.p
Eucalyptus___Eucgr.K03302.1.p
Quercus___Qurub.12G211700.1
Malus___MD05G1203800
Cucumis___Cucsa.388420.1|PACid_16982852
Cucumis___Cucsa.388410.1|PACid_16982851
Cucumis___Cucsa.151360.1|PACid_16964451
Cucumis___Cucsa.263470.1|PACid_16973240
Cucumis___Cucsa.084300.1|PACid_16956515
Citrus___orange1.1g040279m|PACid_18127184
Citrus___orange1.1g038190m|PACid_18119585
Citrus___orange1.1g037409m|PACid_18100159
Citrus___orange1.1g047932m|PACid_18139440
Citrus___orange1.1g044044m|PACid_18092872
Citrus___orange1.1g043191m|PACid_18120242
Citrus___orange1.1g038914m|PACid_18120243
Citrus___orange1.1g041816m|PACid_18139438
Citrus___orange1.1g040027m|PACid_18100284
Citrus___orange1.1g015811m|PACid_18137876
Citrus___orange1.1g040261m|PACid_18123408
Citrus___orange1.1g036577m|PACid_18096118
Citrus___orange1.1g040367m|PACid_18098037
Citrus___orange1.1g047518m|PACid_18132688
Citrus___orange1.1g011362m|PACid_18126461
Beta___EL10Ac5g10449.1
Mimulus___Migut.N00244.1.p
Mimulus___Migut.N00243.1.p
Vaccinium___Vadar_g31671.t1
Vaccinium___Vadar_g31637.t1
Vitis___VIT_205s0020g04920.1
Hydrangea___Hyque.10G099100.1.p
Mimulus___Migut.G00714.1.p
Mimulus___Migut.G00713.1.p
Mimulus___Migut.O00664.1.p

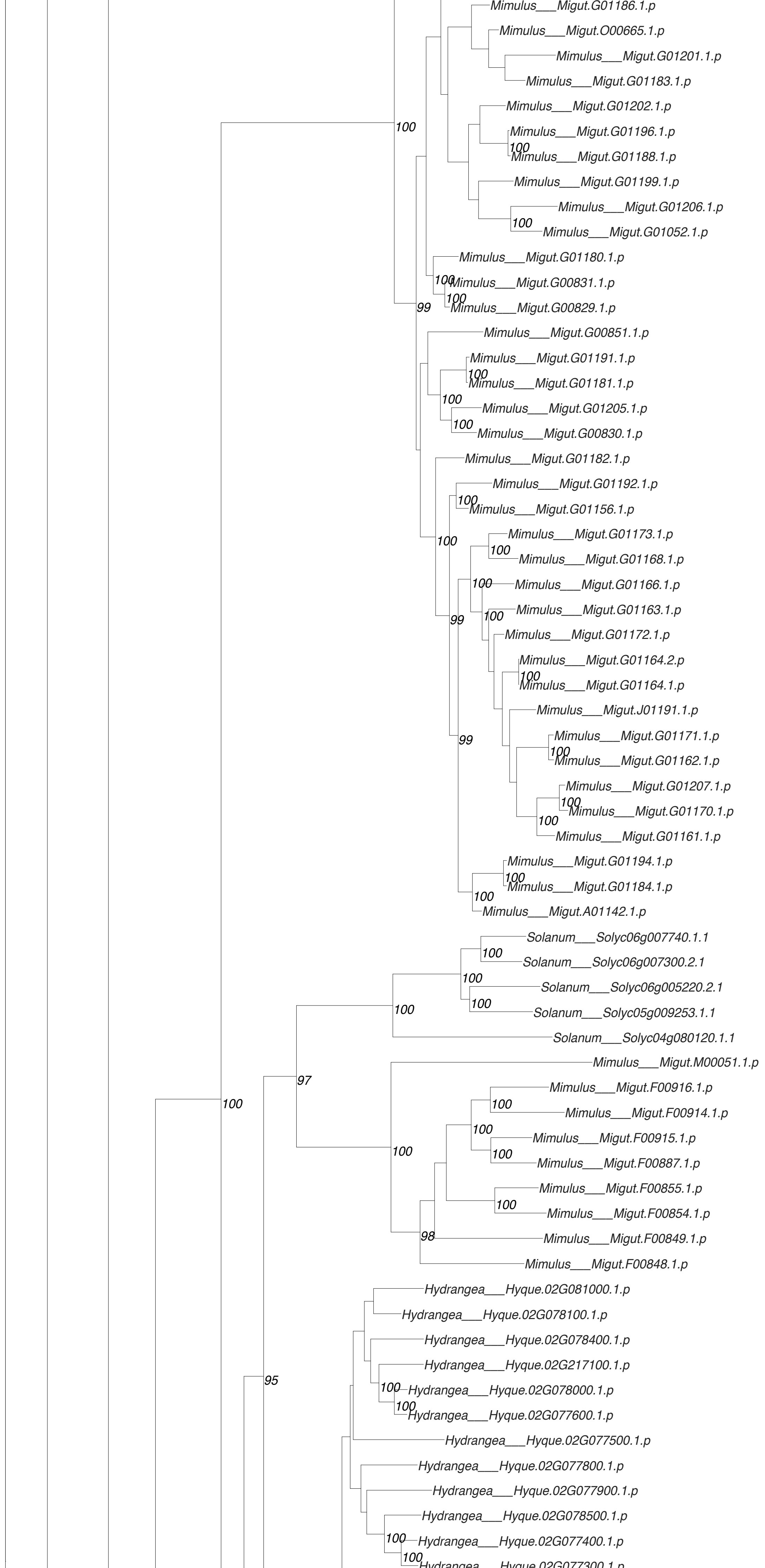

Mimulus___Migut.G01186.1.p
Mimulus___Migut.O00665.1.p
Mimulus___Migut.G01201.1.p
Mimulus___Migut.G01183.1.p
Mimulus___Migut.G01202.1.p
Mimulus___Migut.G01196.1.p
Mimulus___Migut.G01188.1.p
Mimulus___Migut.G01199.1.p
Mimulus___Migut.G01206.1.p
Mimulus___Migut.G01052.1.p
Mimulus___Migut.G01180.1.p
Mimulus___Migut.G00831.1.p
Mimulus___Migut.G00829.1.p
Mimulus___Migut.G00851.1.p
Mimulus___Migut.G01191.1.p
Mimulus___Migut.G01181.1.p
Mimulus___Migut.G01205.1.p
Mimulus___Migut.G00830.1.p
Mimulus___Migut.G01182.1.p
Mimulus___Migut.G01192.1.p
Mimulus___Migut.G01156.1.p
Mimulus___Migut.G01173.1.p
Mimulus___Migut.G01168.1.p
Mimulus___Migut.G01166.1.p
Mimulus___Migut.G01163.1.p
Mimulus___Migut.G01172.1.p
Mimulus___Migut.G01164.2.p
Mimulus___Migut.G01164.1.p
Mimulus___Migut.J01191.1.p
Mimulus___Migut.G01171.1.p
Mimulus___Migut.G01162.1.p
Mimulus___Migut.G01207.1.p
Mimulus___Migut.G01170.1.p
Mimulus___Migut.G01161.1.p
Mimulus___Migut.G01194.1.p
Mimulus___Migut.G01184.1.p
Mimulus___Migut.A01142.1.p
Solanum___Solyc06g007740.1.1
Solanum___Solyc06g007300.2.1
Solanum___Solyc06g005220.2.1
Solanum___Solyc05g009253.1.1
Solanum___Solyc04g080120.1.1
Mimulus___Migut.M00051.1.p
Mimulus___Migut.F00916.1.p
Mimulus___Migut.F00914.1.p
Mimulus___Migut.F00915.1.p
Mimulus___Migut.F00887.1.p
Mimulus___Migut.F00855.1.p
Mimulus___Migut.F00854.1.p
Mimulus___Migut.F00849.1.p
Mimulus___Migut.F00848.1.p
Hydrangea___Hyque.02G081000.1.p
Hydrangea___Hyque.02G078100.1.p
Hydrangea___Hyque.02G078400.1.p
Hydrangea___Hyque.02G217100.1.p
Hydrangea___Hyque.02G078000.1.p
Hydrangea___Hyque.02G077600.1.p
Hydrangea___Hyque.02G077500.1.p
Hydrangea___Hyque.02G077800.1.p
Hydrangea___Hyque.02G077900.1.p
Hydrangea___Hyque.02G078500.1.p
Hydrangea___Hyque.02G077400.1.p

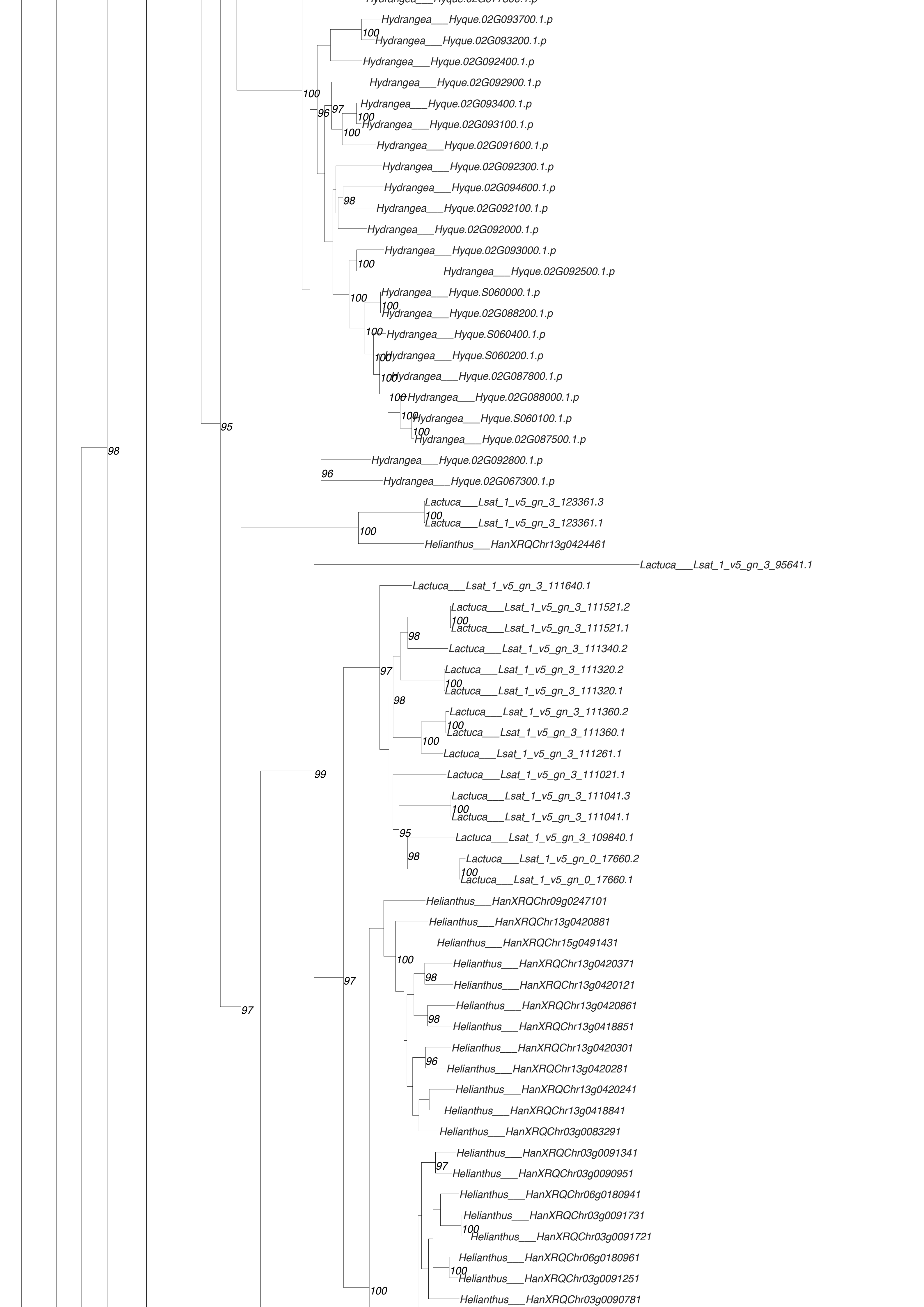
Hydrangea___Hyque.02G093700.1.p
Hydrangea___Hyque.02G093200.1.p
Hydrangea___Hyque.02G092400.1.p
Hydrangea___Hyque.02G092900.1.p
Hydrangea___Hyque.02G093400.1.p
Hydrangea___Hyque.02G093100.1.p
Hydrangea___Hyque.02G091600.1.p
Hydrangea___Hyque.02G092300.1.p
Hydrangea___Hyque.02G094600.1.p
Hydrangea___Hyque.02G092100.1.p
Hydrangea___Hyque.02G092000.1.p
Hydrangea___Hyque.02G093000.1.p
Hydrangea___Hyque.02G092500.1.p
Hydrangea___Hyque.S060000.1.p
Hydrangea___Hyque.02G088200.1.p
Hydrangea___Hyque.S060400.1.p
Hydrangea___Hyque.S060200.1.p
Hydrangea___Hyque.02G087800.1.p
Hydrangea___Hyque.02G088000.1.p
Hydrangea___Hyque.S060100.1.p
Hydrangea___Hyque.02G087500.1.p
Hydrangea___Hyque.02G092800.1.p
Hydrangea___Hyque.02G067300.1.p
Lactuca___Lsat_1_v5_gn_3_123361.3
Lactuca___Lsat_1_v5_gn_3_123361.1
Helianthus___HanXRQChr13g0424461
Lactuca___Lsat_1_v5_gn_3_95641.1
Lactuca___Lsat_1_v5_gn_3_111640.1
Lactuca___Lsat_1_v5_gn_3_111521.2
Lactuca___Lsat_1_v5_gn_3_111521.1
Lactuca___Lsat_1_v5_gn_3_111340.2
Lactuca___Lsat_1_v5_gn_3_111320.2
Lactuca___Lsat_1_v5_gn_3_111320.1
Lactuca___Lsat_1_v5_gn_3_111360.2
Lactuca___Lsat_1_v5_gn_3_111360.1
Lactuca___Lsat_1_v5_gn_3_111261.1
Lactuca___Lsat_1_v5_gn_3_111021.1
Lactuca___Lsat_1_v5_gn_3_111041.3
Lactuca___Lsat_1_v5_gn_3_111041.1
Lactuca___Lsat_1_v5_gn_3_109840.1
Lactuca___Lsat_1_v5_gn_0_17660.2
Lactuca___Lsat_1_v5_gn_0_17660.1
Helianthus___HanXRQChr09g0247101
Helianthus___HanXRQChr13g0420881
Helianthus___HanXRQChr15g0491431
Helianthus___HanXRQChr13g0420371
Helianthus___HanXRQChr13g0420121
Helianthus___HanXRQChr13g0420861
Helianthus___HanXRQChr13g0418851
Helianthus___HanXRQChr13g0420301
Helianthus___HanXRQChr13g0420281
Helianthus___HanXRQChr13g0420241
Helianthus___HanXRQChr13g0418841
Helianthus___HanXRQChr03g0083291
Helianthus___HanXRQChr03g0091341
Helianthus___HanXRQChr03g0090951
Helianthus___HanXRQChr06g0180941
Helianthus___HanXRQChr03g0091731
Helianthus___HanXRQChr03g0091721
Helianthus___HanXRQChr06g0180961
Helianthus___HanXRQChr03g0091251
Helianthus___HanXRQChr03g0090781

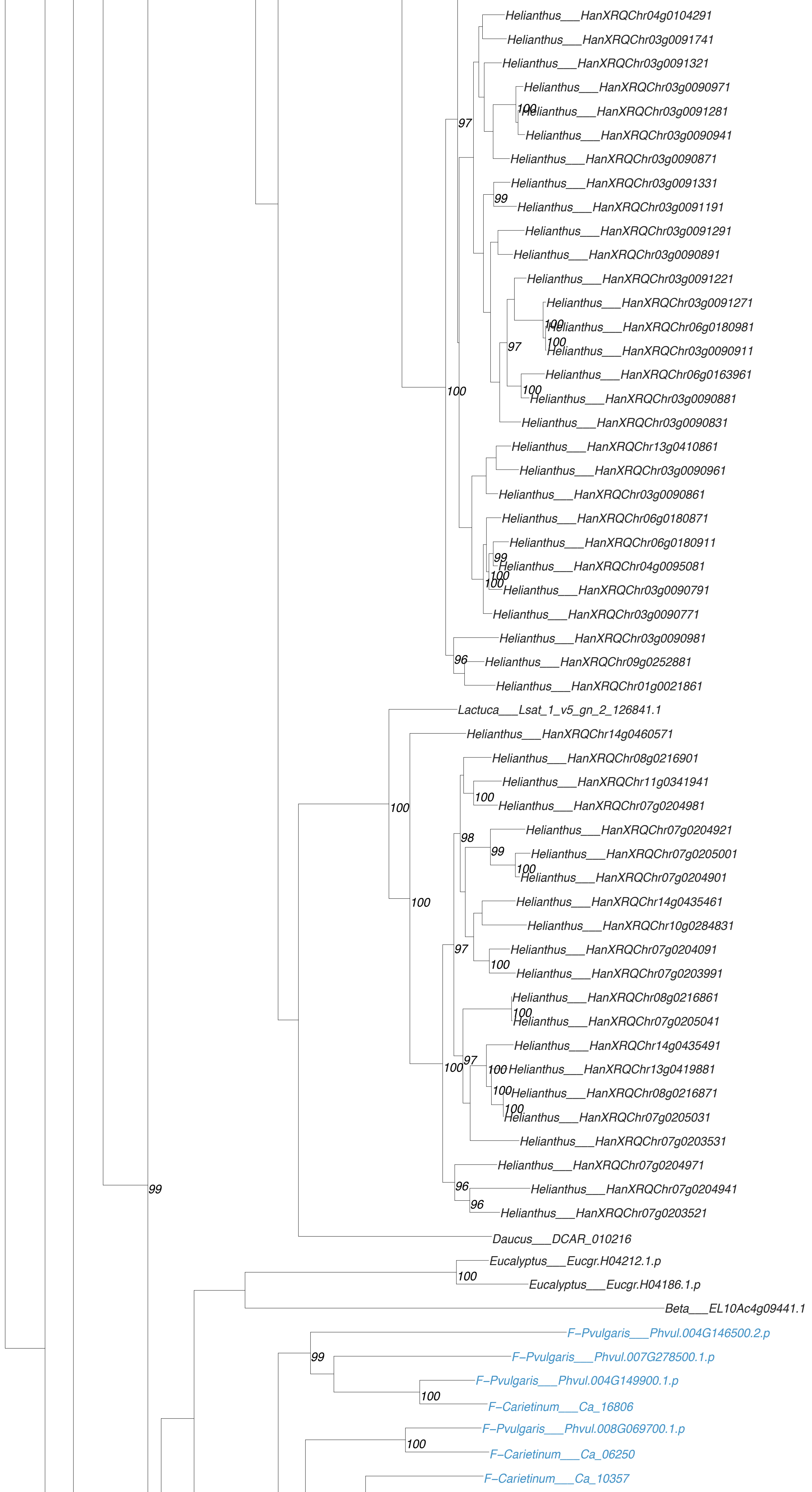
Helianthus___HanXRQChr04g0104291
Helianthus___HanXRQChr03g0091741
Helianthus___HanXRQChr03g0091321
Helianthus___HanXRQChr03g0090971
Helianthus___HanXRQChr03g0091281
Helianthus___HanXRQChr03g0090941
Helianthus___HanXRQChr03g0090871
Helianthus___HanXRQChr03g0091331
Helianthus___HanXRQChr03g0091191
Helianthus___HanXRQChr03g0091291
Helianthus___HanXRQChr03g0090891
Helianthus___HanXRQChr03g0091221
Helianthus___HanXRQChr03g0091271
Helianthus___HanXRQChr06g0180981
Helianthus___HanXRQChr03g0090911
Helianthus___HanXRQChr06g0163961
Helianthus___HanXRQChr03g0090881
Helianthus___HanXRQChr03g0090831
Helianthus___HanXRQChr13g0410861
Helianthus___HanXRQChr03g0090961
Helianthus___HanXRQChr03g0090861
Helianthus___HanXRQChr06g0180871
Helianthus___HanXRQChr06g0180911
Helianthus___HanXRQChr04g0095081
Helianthus___HanXRQChr03g0090791
Helianthus___HanXRQChr03g0090771
Helianthus___HanXRQChr03g0090981
Helianthus___HanXRQChr09g0252881
Helianthus___HanXRQChr01g0021861
Lactuca___Lsat_1_v5_gn_2_126841.1
Helianthus___HanXRQChr14g0460571
Helianthus___HanXRQChr08g0216901
Helianthus___HanXRQChr11g0341941
Helianthus___HanXRQChr07g0204981
Helianthus___HanXRQChr07g0204921
Helianthus___HanXRQChr07g0205001
Helianthus___HanXRQChr07g0204901
Helianthus___HanXRQChr14g0435461
Helianthus___HanXRQChr10g0284831
Helianthus___HanXRQChr07g0204091
Helianthus___HanXRQChr07g0203991
Helianthus___HanXRQChr08g0216861
Helianthus___HanXRQChr07g0205041
Helianthus___HanXRQChr14g0435491
Helianthus___HanXRQChr13g0419881
Helianthus___HanXRQChr08g0216871
Helianthus___HanXRQChr07g0205031
Helianthus___HanXRQChr07g0203531
Helianthus___HanXRQChr07g0204971
Helianthus___HanXRQChr07g0204941
Helianthus___HanXRQChr07g0203521
Daucus___DCAR_010216
Eucalyptus___Eucgr.H04212.1.p
Eucalyptus___Eucgr.H04186.1.p
Beta___EL10Ac4g09441.1
F−Pvulgaris___Phvul.004G146500.2.p
F−Pvulgaris___Phvul.007G278500.1.p
F−Pvulgaris___Phvul.004G149900.1.p
F−Carietinum___Ca_16806
F−Pvulgaris___Phvul.008G069700.1.p
F−Carietinum___Ca_06250
F−Carietinum___Ca_10357

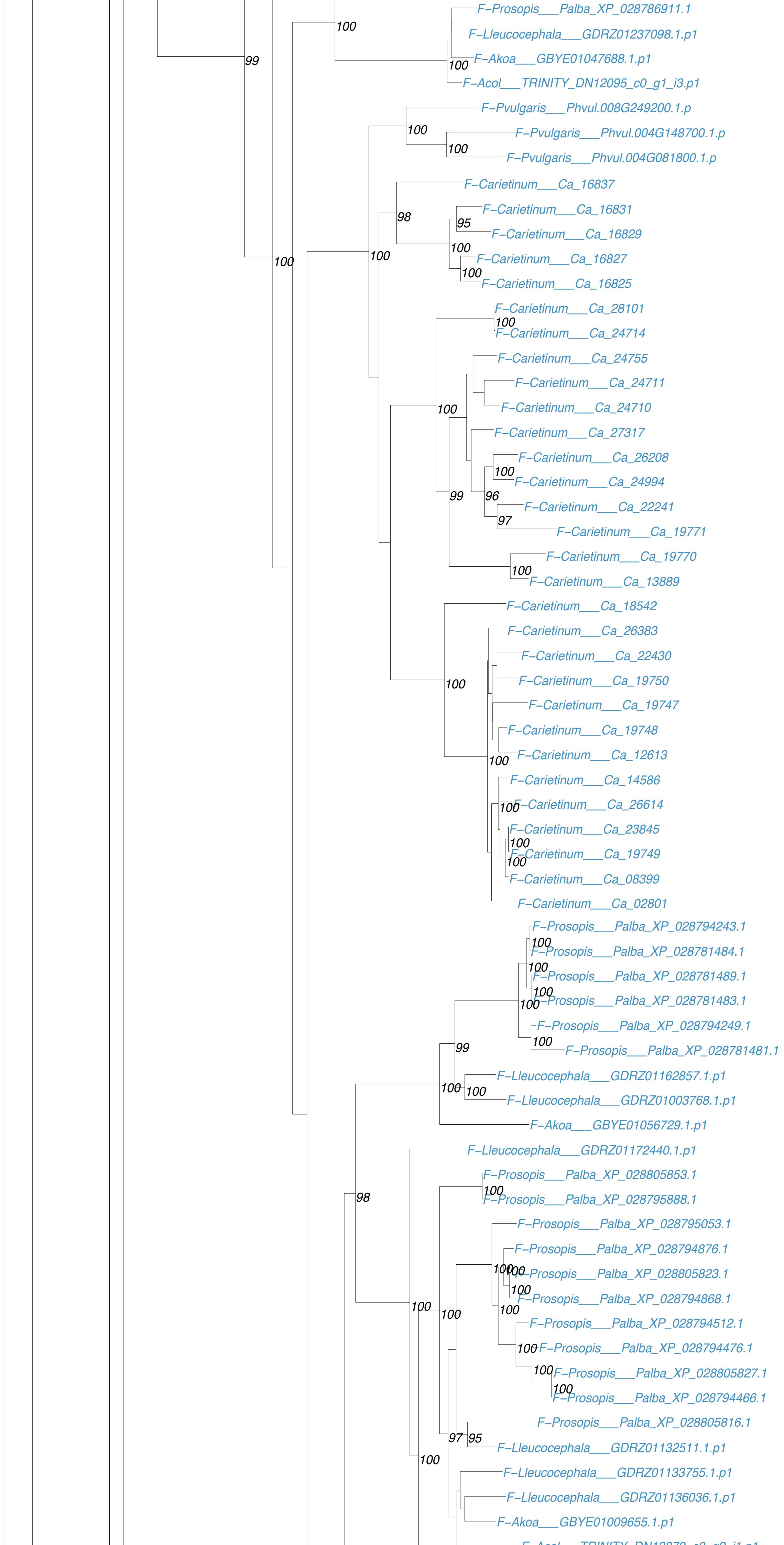
F–Prosopis___Palba_XP_028786911.1
F–Lleucocephala___GDRZ01237098.1.p1
F–Akoa___GBYE01047688.1.p1
F–Acol___TRINITY_DN12095_c0_g1_i3.p1
F–Pvulgaris___Phvul.008G249200.1.p
F–Pvulgaris___Phvul.004G148700.1.p
F–Pvulgaris___Phvul.004G081800.1.p
F–Carietinum___Ca_16837
F–Carietinum___Ca_16831
F–Carietinum___Ca_16829
F–Carietinum___Ca_16827
F–Carietinum___Ca_16825
F–Carietinum___Ca_28101
F–Carietinum___Ca_24714
F–Carietinum___Ca_24755
F–Carietinum___Ca_24711
F–Carietinum___Ca_24710
F–Carietinum___Ca_27317
F–Carietinum___Ca_26208
F–Carietinum___Ca_24994
F–Carietinum___Ca_22241
F–Carietinum___Ca_19771
F–Carietinum___Ca_19770
F–Carietinum___Ca_13889
F–Carietinum___Ca_18542
F–Carietinum___Ca_26383
F–Carietinum___Ca_22430
F–Carietinum___Ca_19750
F–Carietinum___Ca_19747
F–Carietinum___Ca_19748
F–Carietinum___Ca_12613
F–Carietinum___Ca_14586
F–Carietinum___Ca_26614
F–Carietinum___Ca_23845
F–Carietinum___Ca_19749
F–Carietinum___Ca_08399
F–Carietinum___Ca_02801
F–Prosopis___Palba_XP_028794243.1
F–Prosopis___Palba_XP_028781484.1
F–Prosopis___Palba_XP_028781489.1
F–Prosopis___Palba_XP_028781483.1
F–Prosopis___Palba_XP_028794249.1
F–Prosopis___Palba_XP_028781481.1
F–Lleucocephala___GDRZ01162857.1.p1
F–Lleucocephala___GDRZ01003768.1.p1
F–Akoa___GBYE01056729.1.p1
F–Lleucocephala___GDRZ01172440.1.p1
F–Prosopis___Palba_XP_028805853.1
F–Prosopis___Palba_XP_028795888.1
F–Prosopis___Palba_XP_028795053.1
F–Prosopis___Palba_XP_028794876.1
F–Prosopis___Palba_XP_028805823.1
F–Prosopis___Palba_XP_028794868.1
F–Prosopis___Palba_XP_028794512.1
F–Prosopis___Palba_XP_028794476.1
F–Prosopis___Palba_XP_028805827.1
F–Prosopis___Palba_XP_028794466.1
F–Prosopis___Palba_XP_028805816.1
F–Lleucocephala___GDRZ01132511.1.p1
F–Lleucocephala___GDRZ01133755.1.p1
F–Lleucocephala___GDRZ01136036.1.p1
F–Akoa___GBYE01009655.1.p1

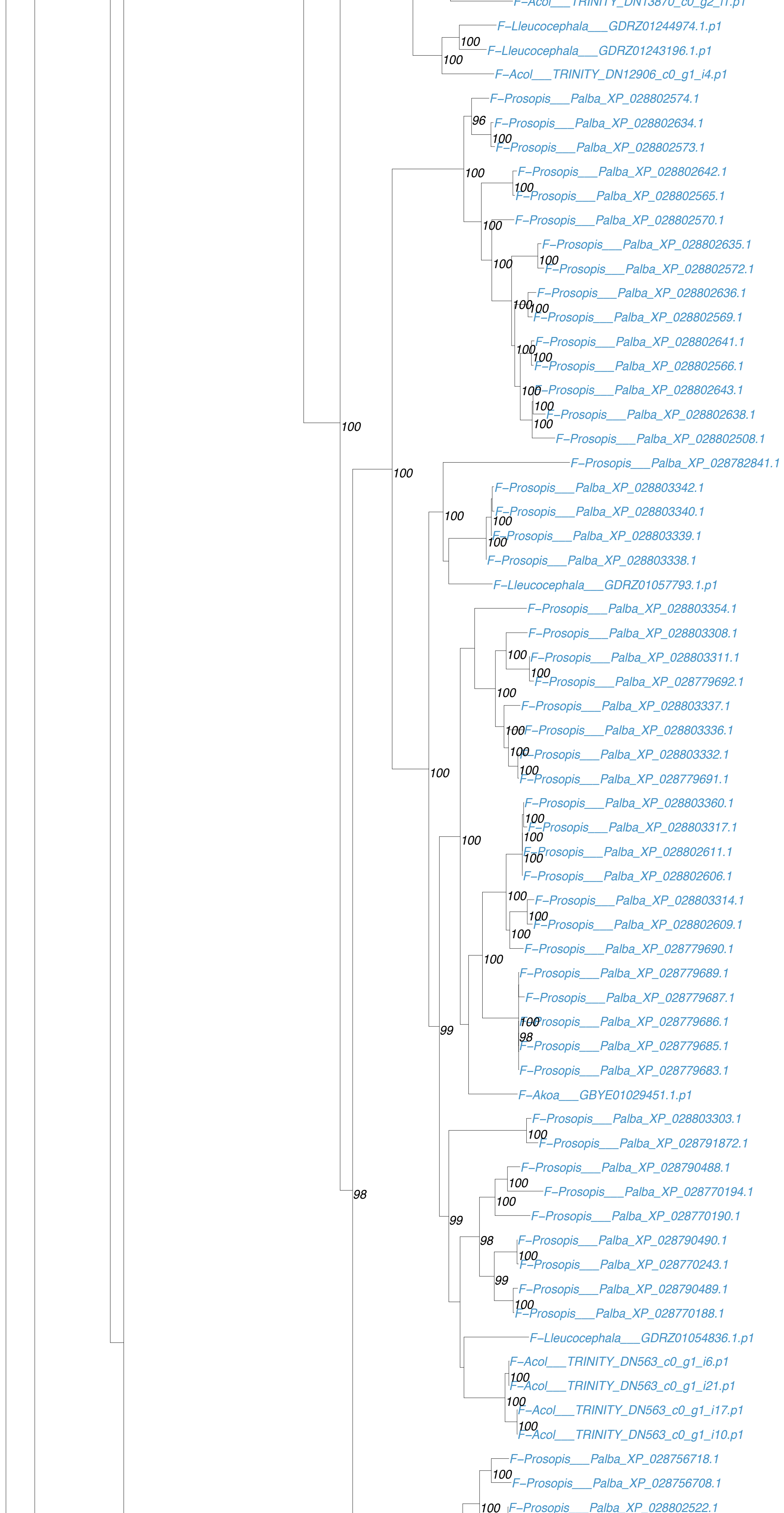

F–Lleucocephala___GDRZ01244974.1.p1
F–Lleucocephala___GDRZ01243196.1.p1
F–Acol___TRINITY_DN12906_c0_g1_i4.p1
F–Prosopis___Palba_XP_028802574.1
F–Prosopis___Palba_XP_028802634.1
F–Prosopis___Palba_XP_028802573.1
F–Prosopis___Palba_XP_028802642.1
F–Prosopis___Palba_XP_028802565.1
F–Prosopis___Palba_XP_028802570.1
F–Prosopis___Palba_XP_028802635.1
F–Prosopis___Palba_XP_028802572.1
F–Prosopis___Palba_XP_028802636.1
F–Prosopis___Palba_XP_028802569.1
F–Prosopis___Palba_XP_028802641.1
F–Prosopis___Palba_XP_028802566.1
F–Prosopis___Palba_XP_028802643.1
F–Prosopis___Palba_XP_028802638.1
F–Prosopis___Palba_XP_028802508.1
F–Prosopis___Palba_XP_028782841.1
F–Prosopis___Palba_XP_028803342.1
F–Prosopis___Palba_XP_028803340.1
F–Prosopis___Palba_XP_028803339.1
F–Prosopis___Palba_XP_028803338.1
F–Lleucocephala___GDRZ01057793.1.p1
F–Prosopis___Palba_XP_028803354.1
F–Prosopis___Palba_XP_028803308.1
F–Prosopis___Palba_XP_028803311.1
F–Prosopis___Palba_XP_028779692.1
F–Prosopis___Palba_XP_028803337.1
F–Prosopis___Palba_XP_028803336.1
F–Prosopis___Palba_XP_028803332.1
F–Prosopis___Palba_XP_028779691.1
F–Prosopis___Palba_XP_028803360.1
F–Prosopis___Palba_XP_028803317.1
F–Prosopis___Palba_XP_028802611.1
F–Prosopis___Palba_XP_028802606.1
F–Prosopis___Palba_XP_028803314.1
F–Prosopis___Palba_XP_028802609.1
F–Prosopis___Palba_XP_028779690.1
F–Prosopis___Palba_XP_028779689.1
F–Prosopis___Palba_XP_028779687.1
F–Prosopis___Palba_XP_028779686.1
F–Prosopis___Palba_XP_028779685.1
F–Prosopis___Palba_XP_028779683.1
F–Akoa___GBYE01029451.1.p1
F–Prosopis___Palba_XP_028803303.1
F–Prosopis___Palba_XP_028791872.1
F–Prosopis___Palba_XP_028790488.1
F–Prosopis___Palba_XP_028770194.1
F–Prosopis___Palba_XP_028770190.1
F–Prosopis___Palba_XP_028790490.1
F–Prosopis___Palba_XP_028770243.1
F–Prosopis___Palba_XP_028790489.1
F–Prosopis___Palba_XP_028770188.1
F–Lleucocephala___GDRZ01054836.1.p1
F–Acol___TRINITY_DN563_c0_g1_i6.p1
F–Acol___TRINITY_DN563_c0_g1_i21.p1
F–Acol___TRINITY_DN563_c0_g1_i17.p1
F–Acol___TRINITY_DN563_c0_g1_i10.p1
F–Prosopis___Palba_XP_028756718.1
F–Prosopis___Palba_XP_028756708.1
F–Prosopis___Palba_XP_028802522.1

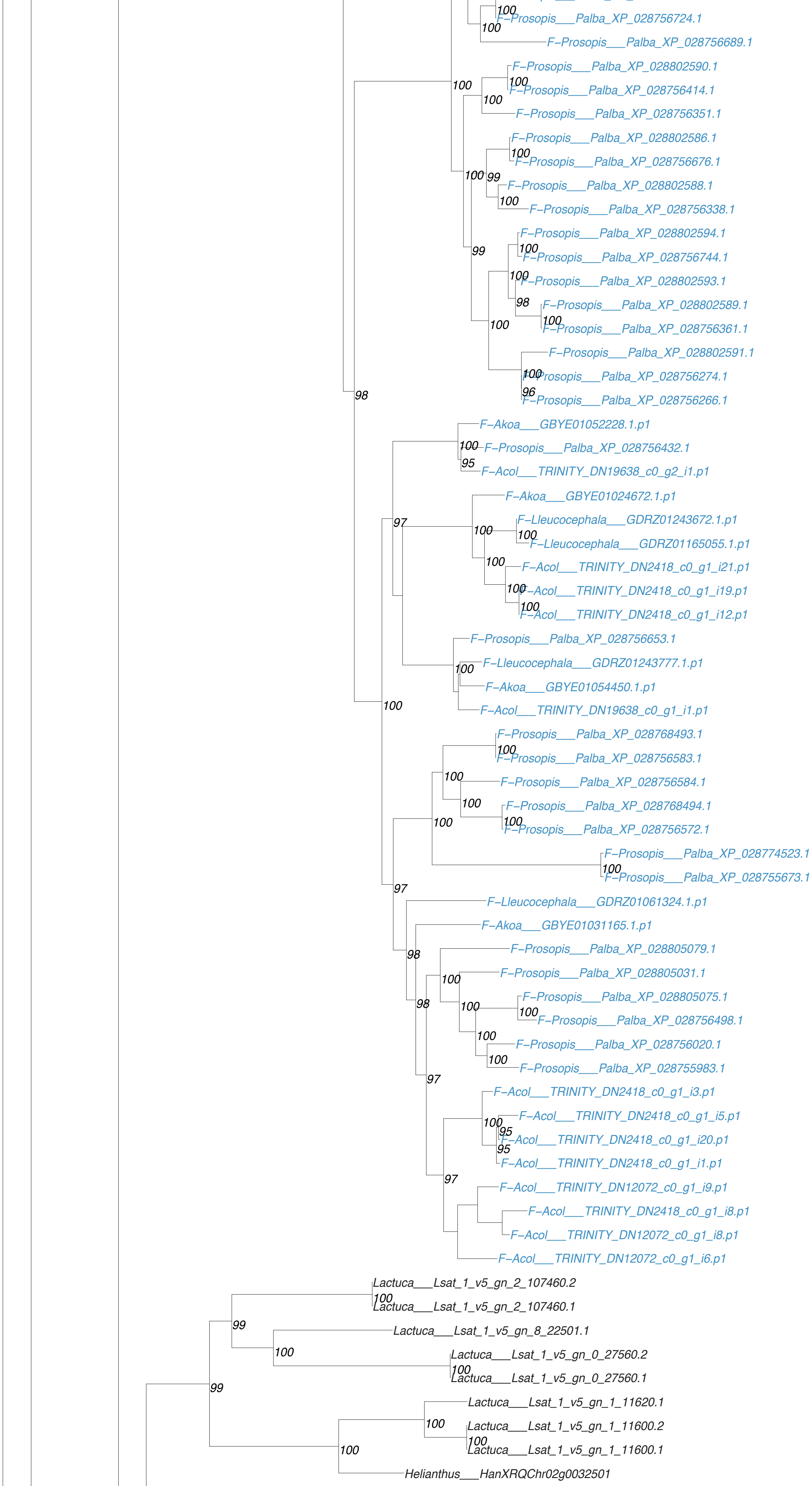
F–Prosopis___Palba_XP_028756724.1
F–Prosopis___Palba_XP_028756689.1
F–Prosopis___Palba_XP_028802590.1
F–Prosopis___Palba_XP_028756414.1
F–Prosopis___Palba_XP_028756351.1
F–Prosopis___Palba_XP_028802586.1
F–Prosopis___Palba_XP_028756676.1
F–Prosopis___Palba_XP_028802588.1
F–Prosopis___Palba_XP_028756338.1
F–Prosopis___Palba_XP_028802594.1
F–Prosopis___Palba_XP_028756744.1
F–Prosopis___Palba_XP_028802593.1
F–Prosopis___Palba_XP_028802589.1
F–Prosopis___Palba_XP_028756361.1
F–Prosopis___Palba_XP_028802591.1
F–Prosopis___Palba_XP_028756274.1
F–Prosopis___Palba_XP_028756266.1
F–Akoa___GBYE01052228.1.p1
F–Prosopis___Palba_XP_028756432.1
F–Acol___TRINITY_DN19638_c0_g2_i1.p1
F–Akoa___GBYE01024672.1.p1
F–Lleucocephala___GDRZ01243672.1.p1
F–Lleucocephala___GDRZ01165055.1.p1
F–Acol___TRINITY_DN2418_c0_g1_i21.p1
F–Acol___TRINITY_DN2418_c0_g1_i19.p1
F–Acol___TRINITY_DN2418_c0_g1_i12.p1
F–Prosopis___Palba_XP_028756653.1
F–Lleucocephala___GDRZ01243777.1.p1
F–Akoa___GBYE01054450.1.p1
F–Acol___TRINITY_DN19638_c0_g1_i1.p1
F–Prosopis___Palba_XP_028768493.1
F–Prosopis___Palba_XP_028756583.1
F–Prosopis___Palba_XP_028756584.1
F–Prosopis___Palba_XP_028768494.1
F–Prosopis___Palba_XP_028756572.1
F–Prosopis___Palba_XP_028774523.1
F–Prosopis___Palba_XP_028755673.1
F–Lleucocephala___GDRZ01061324.1.p1
F–Akoa___GBYE01031165.1.p1
F–Prosopis___Palba_XP_028805079.1
F–Prosopis___Palba_XP_028805031.1
F–Prosopis___Palba_XP_028805075.1
F–Prosopis___Palba_XP_028756498.1
F–Prosopis___Palba_XP_028756020.1
F–Prosopis___Palba_XP_028755983.1
F–Acol___TRINITY_DN2418_c0_g1_i3.p1
F–Acol___TRINITY_DN2418_c0_g1_i5.p1
F–Acol___TRINITY_DN2418_c0_g1_i20.p1
F–Acol___TRINITY_DN2418_c0_g1_i1.p1
F–Acol___TRINITY_DN12072_c0_g1_i9.p1
F–Acol___TRINITY_DN2418_c0_g1_i8.p1
F–Acol___TRINITY_DN12072_c0_g1_i8.p1
F–Acol___TRINITY_DN12072_c0_g1_i6.p1
Lactuca___Lsat_1_v5_gn_2_107460.2
Lactuca___Lsat_1_v5_gn_2_107460.1
Lactuca___Lsat_1_v5_gn_8_22501.1
Lactuca___Lsat_1_v5_gn_0_27560.2
Lactuca___Lsat_1_v5_gn_0_27560.1
Lactuca___Lsat_1_v5_gn_1_11620.1
Lactuca___Lsat_1_v5_gn_1_11600.2
Lactuca___Lsat_1_v5_gn_1_11600.1
Helianthus___HanXRQChr02g0032501

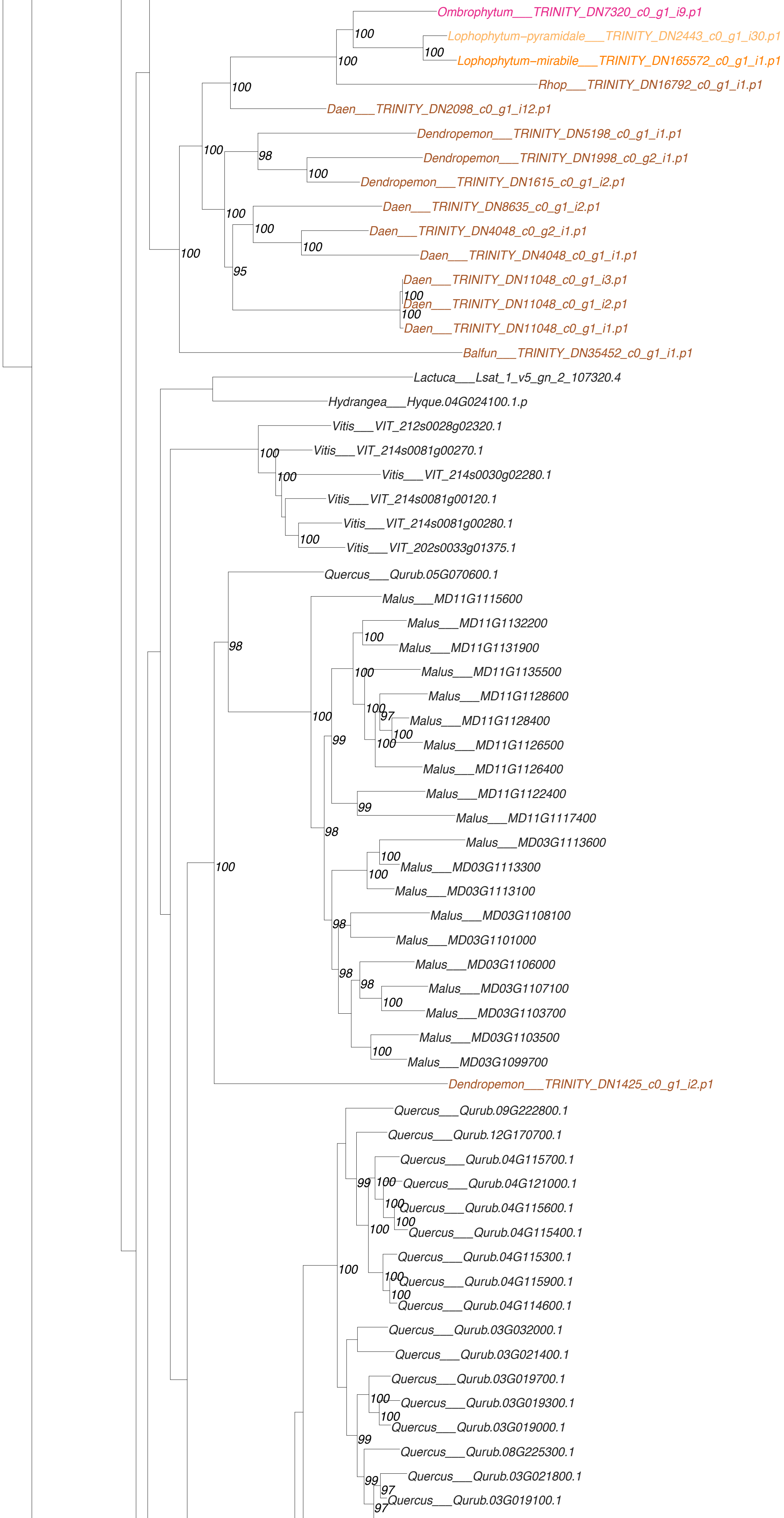
Ombrophytum___TRINITY_DN7320_c0_g1_i9.p1
Lophophytum−pyramidale___TRINITY_DN2443_c0_g1_i30.p1
Lophophytum−mirabile___TRINITY_DN165572_c0_g1_i1.p1
Rhop___TRINITY_DN16792_c0_g1_i1.p1
Daen___TRINITY_DN2098_c0_g1_i12.p1
Dendropemon___TRINITY_DN5198_c0_g1_i1.p1
Dendropemon___TRINITY_DN1998_c0_g2_i1.p1
Dendropemon___TRINITY_DN1615_c0_g1_i2.p1
Daen___TRINITY_DN8635_c0_g1_i2.p1
Daen___TRINITY_DN4048_c0_g2_i1.p1
Daen___TRINITY_DN4048_c0_g1_i1.p1
Daen___TRINITY_DN11048_c0_g1_i3.p1
Daen___TRINITY_DN11048_c0_g1_i2.p1
Daen___TRINITY_DN11048_c0_g1_i1.p1
Balfun___TRINITY_DN35452_c0_g1_i1.p1
Lactuca___Lsat_1_v5_gn_2_107320.4
Hydrangea___Hyque.04G024100.1.p
Vitis___VIT_212s0028g02320.1
Vitis___VIT_214s0081g00270.1
Vitis___VIT_214s0030g02280.1
Vitis___VIT_214s0081g00120.1
Vitis___VIT_214s0081g00280.1
Vitis___VIT_202s0033g01375.1
Quercus___Qurub.05G070600.1
Malus___MD11G1115600
Malus___MD11G1132200
Malus___MD11G1131900
Malus___MD11G1135500
Malus___MD11G1128600
Malus___MD11G1128400
Malus___MD11G1126500
Malus___MD11G1126400
Malus___MD11G1122400
Malus___MD11G1117400
Malus___MD03G1113600
Malus___MD03G1113300
Malus___MD03G1113100
Malus___MD03G1108100
Malus___MD03G1101000
Malus___MD03G1106000
Malus___MD03G1107100
Malus___MD03G1103700
Malus___MD03G1103500
Malus___MD03G1099700
Dendropemon___TRINITY_DN1425_c0_g1_i2.p1
Quercus___Qurub.09G222800.1
Quercus___Qurub.12G170700.1
Quercus___Qurub.04G115700.1
Quercus___Qurub.04G121000.1
Quercus___Qurub.04G115600.1
Quercus___Qurub.04G115400.1
Quercus___Qurub.04G115300.1
Quercus___Qurub.04G115900.1
Quercus___Qurub.04G114600.1
Quercus___Qurub.03G032000.1
Quercus___Qurub.03G021400.1
Quercus___Qurub.03G019700.1
Quercus___Qurub.03G019300.1
Quercus___Qurub.03G019000.1
Quercus___Qurub.08G225300.1
Quercus___Qurub.03G021800.1
Quercus___Qurub.03G019100.1

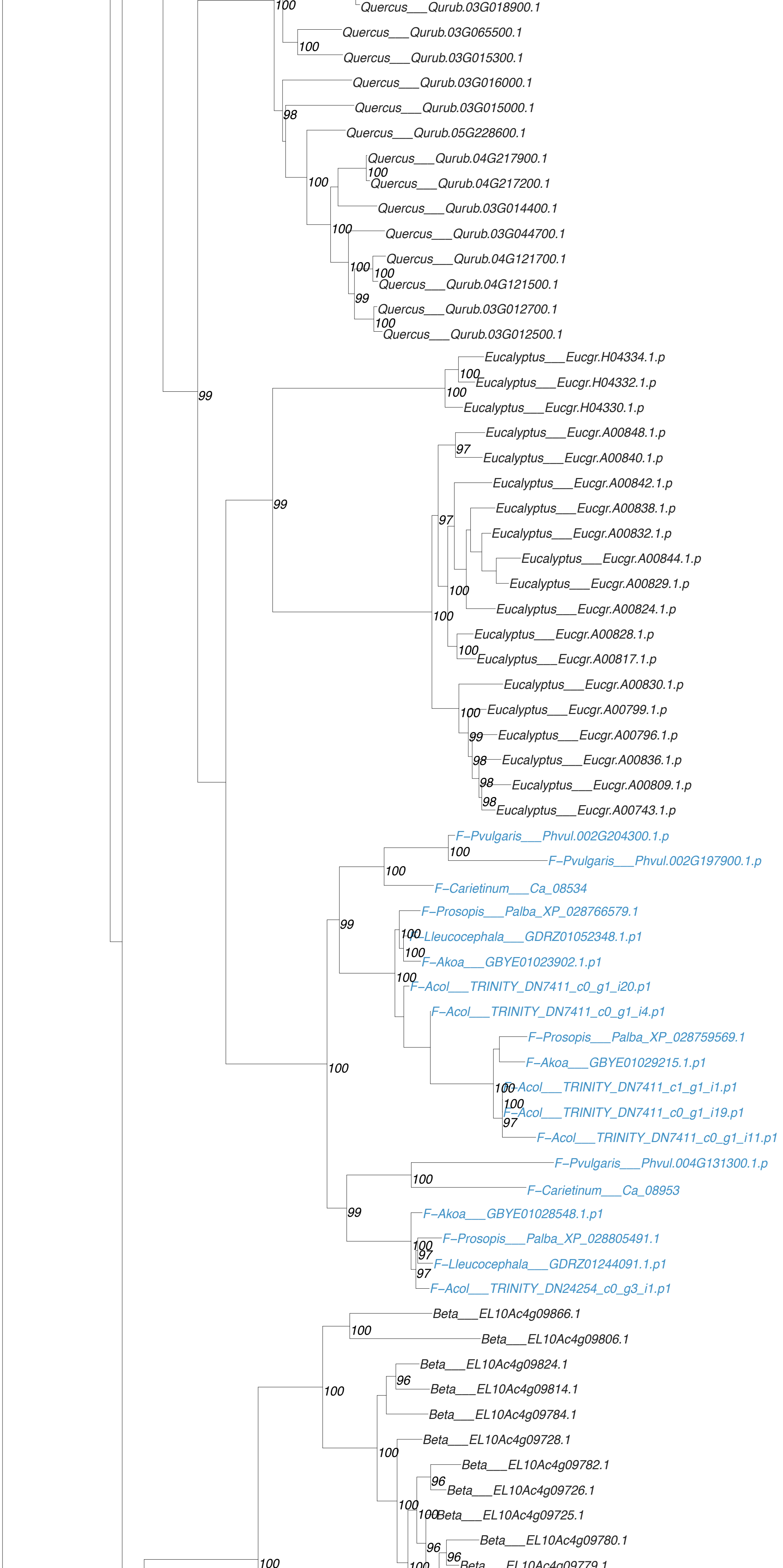

Quercus___Qurub.03G018900.1
Quercus___Qurub.03G065500.1
Quercus___Qurub.03G015300.1
Quercus___Qurub.03G016000.1
Quercus___Qurub.03G015000.1
Quercus___Qurub.05G228600.1
Quercus___Qurub.04G217900.1
Quercus___Qurub.04G217200.1
Quercus___Qurub.03G014400.1
Quercus___Qurub.03G044700.1
Quercus___Qurub.04G121700.1
Quercus___Qurub.04G121500.1
Quercus___Qurub.03G012700.1
Quercus___Qurub.03G012500.1
Eucalyptus___Eucgr.H04334.1.p
Eucalyptus___Eucgr.H04332.1.p
Eucalyptus___Eucgr.H04330.1.p
Eucalyptus___Eucgr.A00848.1.p
Eucalyptus___Eucgr.A00840.1.p
Eucalyptus___Eucgr.A00842.1.p
Eucalyptus___Eucgr.A00838.1.p
Eucalyptus___Eucgr.A00832.1.p
Eucalyptus___Eucgr.A00844.1.p
Eucalyptus___Eucgr.A00829.1.p
Eucalyptus___Eucgr.A00824.1.p
Eucalyptus___Eucgr.A00828.1.p
Eucalyptus___Eucgr.A00817.1.p
Eucalyptus___Eucgr.A00830.1.p
Eucalyptus___Eucgr.A00799.1.p
Eucalyptus___Eucgr.A00796.1.p
Eucalyptus___Eucgr.A00836.1.p
Eucalyptus___Eucgr.A00809.1.p
Eucalyptus___Eucgr.A00743.1.p
F−Pvulgaris___Phvul.002G204300.1.p
F−Pvulgaris___Phvul.002G197900.1.p
F−Carietinum___Ca_08534
F−Prosopis___Palba_XP_028766579.1
F−Lleucocephala___GDRZ01052348.1.p1
F−Akoa___GBYE01023902.1.p1
F−Acol___TRINITY_DN7411_c0_g1_i20.p1
F−Acol___TRINITY_DN7411_c0_g1_i4.p1
F−Prosopis___Palba_XP_028759569.1
F−Akoa___GBYE01029215.1.p1
F−Acol___TRINITY_DN7411_c1_g1_i1.p1
F−Acol___TRINITY_DN7411_c0_g1_i19.p1
F−Acol___TRINITY_DN7411_c0_g1_i11.p1
F−Pvulgaris___Phvul.004G131300.1.p
F−Carietinum___Ca_08953
F−Akoa___GBYE01028548.1.p1
F−Prosopis___Palba_XP_028805491.1
F−Lleucocephala___GDRZ01244091.1.p1
F−Acol___TRINITY_DN24254_c0_g3_i1.p1
Beta___EL10Ac4g09866.1
Beta___EL10Ac4g09806.1
Beta___EL10Ac4g09824.1
Beta___EL10Ac4g09814.1
Beta___EL10Ac4g09784.1
Beta___EL10Ac4g09728.1
Beta___EL10Ac4g09782.1
Beta___EL10Ac4g09726.1
Beta___EL10Ac4g09725.1
Beta___EL10Ac4g09780.1

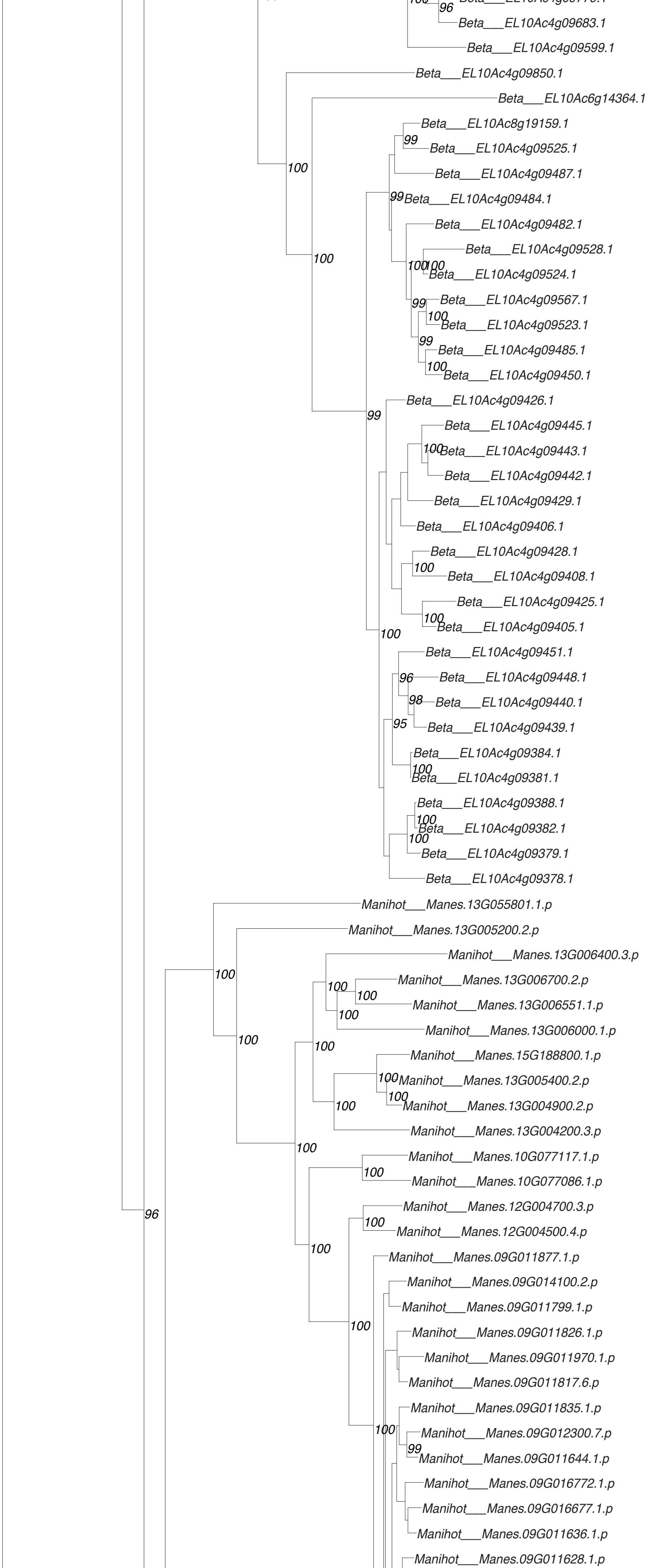
Beta___EL10Ac4g09683.1
Beta___EL10Ac4g09599.1
Beta___EL10Ac4g09850.1
Beta___EL10Ac6g14364.1
Beta___EL10Ac8g19159.1
Beta___EL10Ac4g09525.1
Beta___EL10Ac4g09487.1
Beta___EL10Ac4g09484.1
Beta___EL10Ac4g09482.1
Beta___EL10Ac4g09528.1
Beta___EL10Ac4g09524.1
Beta___EL10Ac4g09567.1
Beta___EL10Ac4g09523.1
Beta___EL10Ac4g09485.1
Beta___EL10Ac4g09450.1
Beta___EL10Ac4g09426.1
Beta___EL10Ac4g09445.1
Beta___EL10Ac4g09443.1
Beta___EL10Ac4g09442.1
Beta___EL10Ac4g09429.1
Beta___EL10Ac4g09406.1
Beta___EL10Ac4g09428.1
Beta___EL10Ac4g09408.1
Beta___EL10Ac4g09425.1
Beta___EL10Ac4g09405.1
Beta___EL10Ac4g09451.1
Beta___EL10Ac4g09448.1
Beta___EL10Ac4g09440.1
Beta___EL10Ac4g09439.1
Beta___EL10Ac4g09384.1
Beta___EL10Ac4g09381.1
Beta___EL10Ac4g09388.1
Beta___EL10Ac4g09382.1
Beta___EL10Ac4g09379.1
Beta___EL10Ac4g09378.1
Manihot___Manes.13G055801.1.p
Manihot___Manes.13G005200.2.p
Manihot___Manes.13G006400.3.p
Manihot___Manes.13G006700.2.p
Manihot___Manes.13G006551.1.p
Manihot___Manes.13G006000.1.p
Manihot___Manes.15G188800.1.p
Manihot___Manes.13G005400.2.p
Manihot___Manes.13G004900.2.p
Manihot___Manes.13G004200.3.p
Manihot___Manes.10G077117.1.p
Manihot___Manes.10G077086.1.p
Manihot___Manes.12G004700.3.p
Manihot___Manes.12G004500.4.p
Manihot___Manes.09G011877.1.p
Manihot___Manes.09G014100.2.p
Manihot___Manes.09G011799.1.p
Manihot___Manes.09G011826.1.p
Manihot___Manes.09G011970.1.p
Manihot___Manes.09G011817.6.p
Manihot___Manes.09G011835.1.p
Manihot___Manes.09G012300.7.p
Manihot___Manes.09G011644.1.p
Manihot___Manes.09G016772.1.p
Manihot___Manes.09G016677.1.p
Manihot___Manes.09G011636.1.p
Manihot___Manes.09G011628.1.p

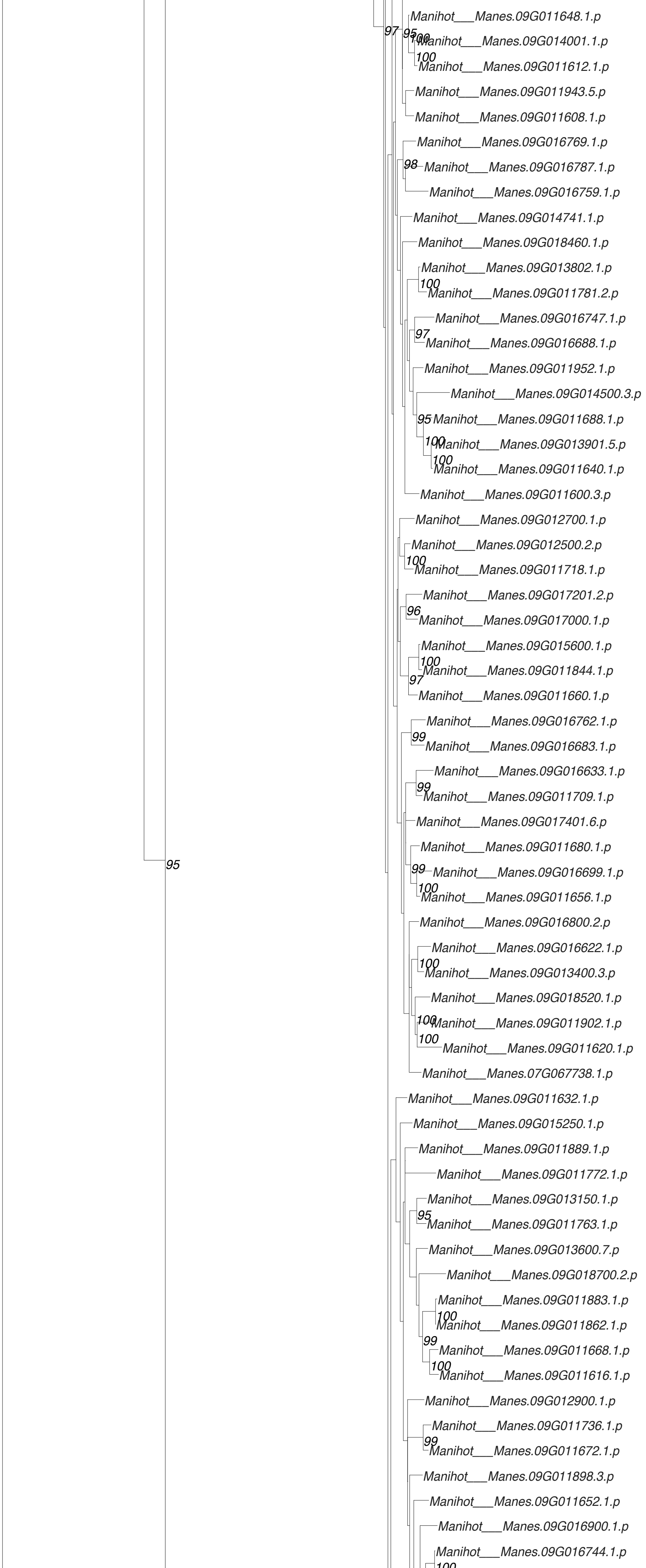
Manihot___Manes.09G011648.1.p
97
95
100
Manihot___Manes.09G014001.1.p
100
Manihot___Manes.09G011612.1.p
Manihot___Manes.09G011943.5.p
Manihot___Manes.09G011608.1.p
Manihot___Manes.09G016769.1.p
98
Manihot___Manes.09G016787.1.p
Manihot___Manes.09G016759.1.p
Manihot___Manes.09G014741.1.p
Manihot___Manes.09G018460.1.p
Manihot___Manes.09G013802.1.p
100
Manihot___Manes.09G011781.2.p
Manihot___Manes.09G016747.1.p
97
Manihot___Manes.09G016688.1.p
Manihot___Manes.09G011952.1.p
Manihot___Manes.09G014500.3.p
95
Manihot___Manes.09G011688.1.p
100
Manihot___Manes.09G013901.5.p
100
Manihot___Manes.09G011640.1.p
Manihot___Manes.09G011600.3.p
Manihot___Manes.09G012700.1.p
Manihot___Manes.09G012500.2.p
100
Manihot___Manes.09G011718.1.p
Manihot___Manes.09G017201.2.p
96
Manihot___Manes.09G017000.1.p
Manihot___Manes.09G015600.1.p
100
Manihot___Manes.09G011844.1.p
97
Manihot___Manes.09G011660.1.p
Manihot___Manes.09G016762.1.p
99
Manihot___Manes.09G016683.1.p
Manihot___Manes.09G016633.1.p
99
Manihot___Manes.09G011709.1.p
Manihot___Manes.09G017401.6.p
Manihot___Manes.09G011680.1.p
95
99
Manihot___Manes.09G016699.1.p
100
Manihot___Manes.09G011656.1.p
Manihot___Manes.09G016800.2.p
Manihot___Manes.09G016622.1.p
100
Manihot___Manes.09G013400.3.p
Manihot___Manes.09G018520.1.p
100
Manihot___Manes.09G011902.1.p
100
Manihot___Manes.09G011620.1.p
Manihot___Manes.07G067738.1.p
Manihot___Manes.09G011632.1.p
Manihot___Manes.09G015250.1.p
Manihot___Manes.09G011889.1.p
Manihot___Manes.09G011772.1.p
Manihot___Manes.09G013150.1.p
95
Manihot___Manes.09G011763.1.p
Manihot___Manes.09G013600.7.p
Manihot___Manes.09G018700.2.p
Manihot___Manes.09G011883.1.p
100
Manihot___Manes.09G011862.1.p
99
Manihot___Manes.09G011668.1.p
100
Manihot___Manes.09G011616.1.p
Manihot___Manes.09G012900.1.p
Manihot___Manes.09G011736.1.p
99
Manihot___Manes.09G011672.1.p
Manihot___Manes.09G011898.3.p
Manihot___Manes.09G011652.1.p
Manihot___Manes.09G016900.1.p
Manihot___Manes.09G016744.1.p
100

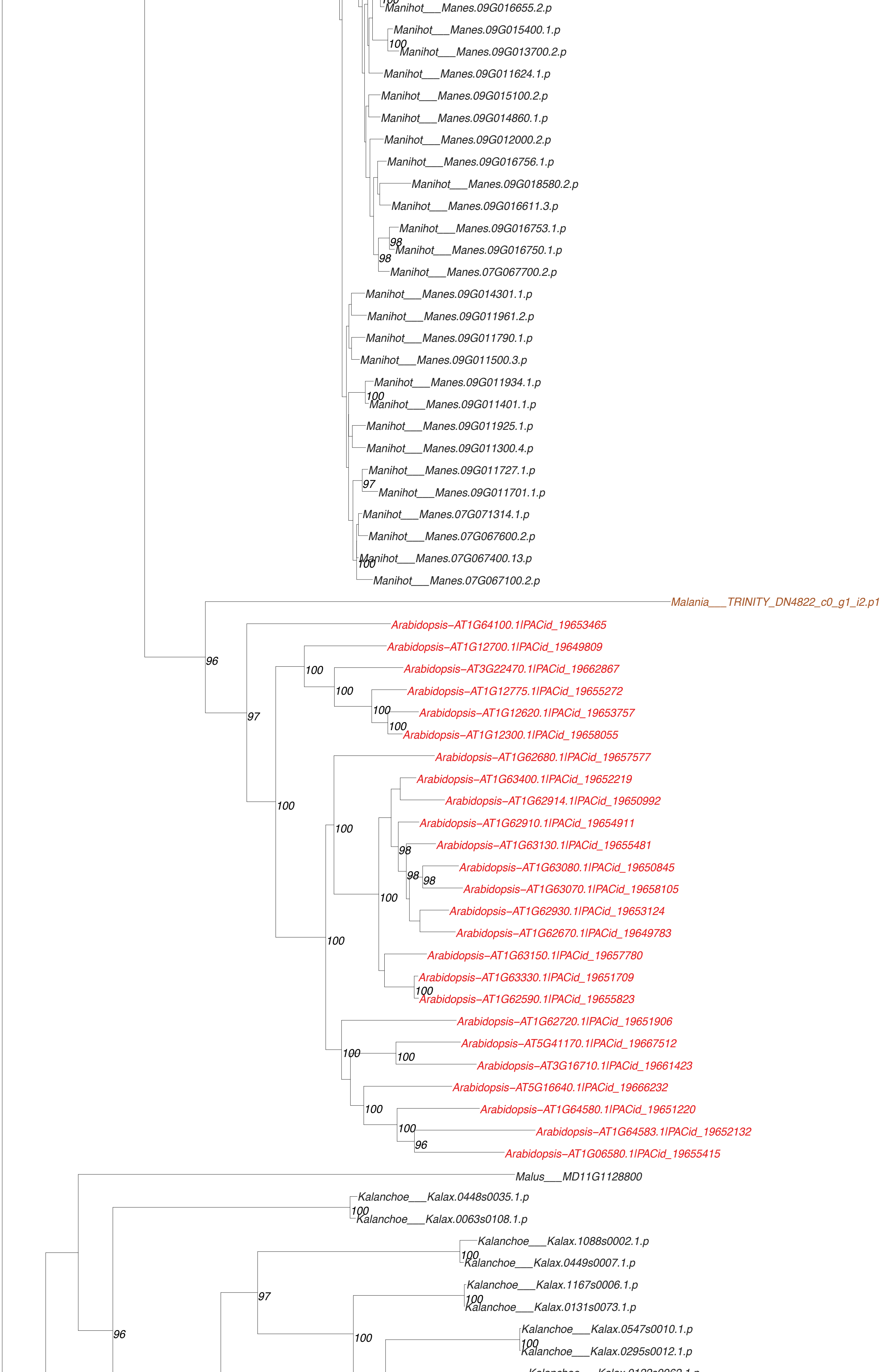

Manihot___Manes.09G016655.2.p
Manihot___Manes.09G015400.1.p
Manihot___Manes.09G013700.2.p
Manihot___Manes.09G011624.1.p
Manihot___Manes.09G015100.2.p
Manihot___Manes.09G014860.1.p
Manihot___Manes.09G012000.2.p
Manihot___Manes.09G016756.1.p
Manihot___Manes.09G018580.2.p
Manihot___Manes.09G016611.3.p
Manihot___Manes.09G016753.1.p
Manihot___Manes.09G016750.1.p
Manihot___Manes.07G067700.2.p
Manihot___Manes.09G014301.1.p
Manihot___Manes.09G011961.2.p
Manihot___Manes.09G011790.1.p
Manihot___Manes.09G011500.3.p
Manihot___Manes.09G011934.1.p
Manihot___Manes.09G011401.1.p
Manihot___Manes.09G011925.1.p
Manihot___Manes.09G011300.4.p
Manihot___Manes.09G011727.1.p
Manihot___Manes.09G011701.1.p
Manihot___Manes.07G071314.1.p
Manihot___Manes.07G067600.2.p
Manihot___Manes.07G067400.13.p
Manihot___Manes.07G067100.2.p
Malania___TRINITY_DN4822_c0_g1_i2.p1
Arabidopsis–AT1G64100.1|PACid_19653465
Arabidopsis–AT1G12700.1|PACid_19649809
Arabidopsis–AT3G22470.1|PACid_19662867
Arabidopsis–AT1G12775.1|PACid_19655272
Arabidopsis–AT1G12620.1|PACid_19653757
Arabidopsis–AT1G12300.1|PACid_19658055
Arabidopsis–AT1G62680.1|PACid_19657577
Arabidopsis–AT1G63400.1|PACid_19652219
Arabidopsis–AT1G62914.1|PACid_19650992
Arabidopsis–AT1G62910.1|PACid_19654911
Arabidopsis–AT1G63130.1|PACid_19655481
Arabidopsis–AT1G63080.1|PACid_19650845
Arabidopsis–AT1G63070.1|PACid_19658105
Arabidopsis–AT1G62930.1|PACid_19653124
Arabidopsis–AT1G62670.1|PACid_19649783
Arabidopsis–AT1G63150.1|PACid_19657780
Arabidopsis–AT1G63330.1|PACid_19651709
Arabidopsis–AT1G62590.1|PACid_19655823
Arabidopsis–AT1G62720.1|PACid_19651906
Arabidopsis–AT5G41170.1|PACid_19667512
Arabidopsis–AT3G16710.1|PACid_19661423
Arabidopsis–AT5G16640.1|PACid_19666232
Arabidopsis–AT1G64580.1|PACid_19651220
Arabidopsis–AT1G64583.1|PACid_19652132
Arabidopsis–AT1G06580.1|PACid_19655415
Malus___MD11G1128800
Kalanchoe___Kalax.0448s0035.1.p
Kalanchoe___Kalax.0063s0108.1.p
Kalanchoe___Kalax.1088s0002.1.p
Kalanchoe___Kalax.0449s0007.1.p
Kalanchoe___Kalax.1167s0006.1.p
Kalanchoe___Kalax.0131s0073.1.p
Kalanchoe___Kalax.0547s0010.1.p
Kalanchoe___Kalax.0295s0012.1.p

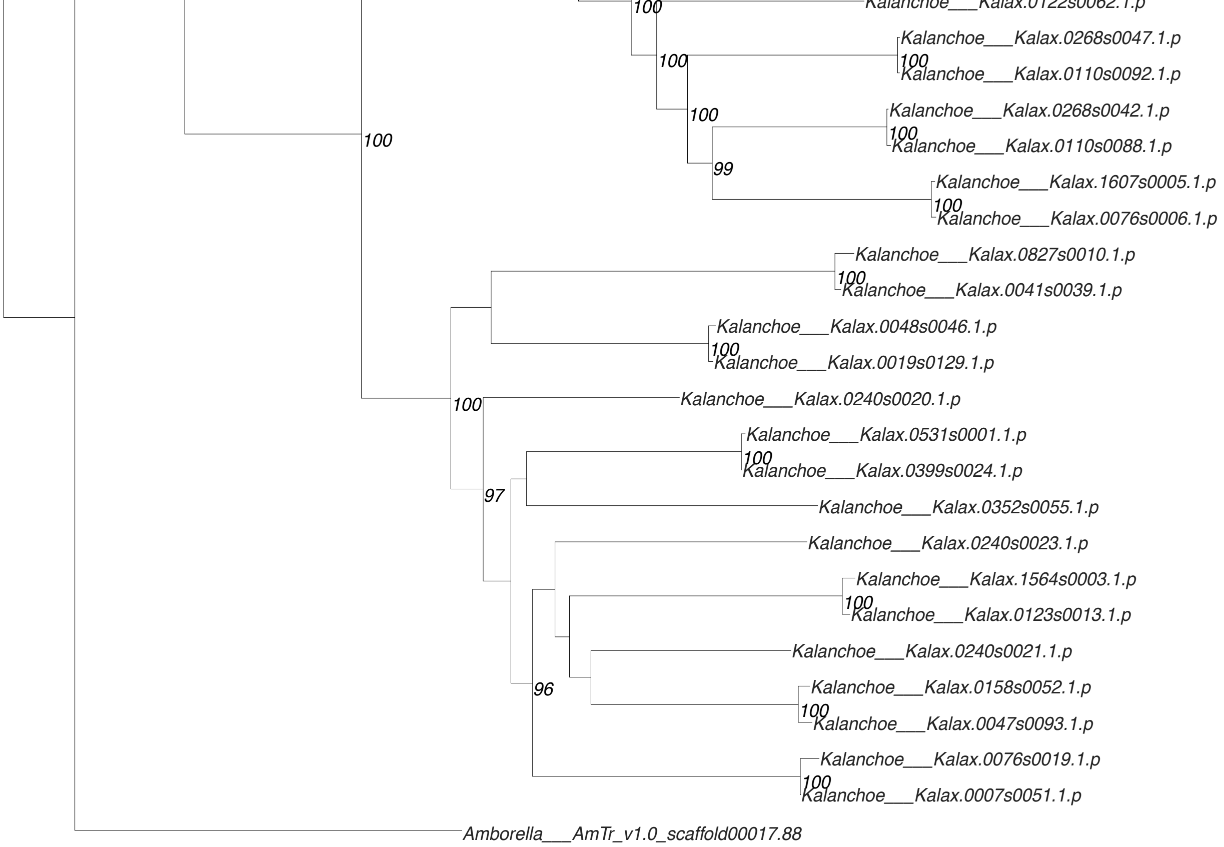

Kalanchoe___Kalax.0122s0062.1.p
100
Kalanchoe___Kalax.0268s0047.1.p
100
Kalanchoe___Kalax.0110s0092.1.p
100
Kalanchoe___Kalax.0268s0042.1.p
100
Kalanchoe___Kalax.0110s0088.1.p
100
99
Kalanchoe___Kalax.1607s0005.1.p
100
Kalanchoe___Kalax.0076s0006.1.p
Kalanchoe___Kalax.0827s0010.1.p
100
Kalanchoe___Kalax.0041s0039.1.p
Kalanchoe___Kalax.0048s0046.1.p
100
Kalanchoe___Kalax.0019s0129.1.p
100
Kalanchoe___Kalax.0240s0020.1.p
Kalanchoe___Kalax.0531s0001.1.p
100
Kalanchoe___Kalax.0399s0024.1.p
97
Kalanchoe___Kalax.0352s0055.1.p
Kalanchoe___Kalax.0240s0023.1.p
Kalanchoe___Kalax.1564s0003.1.p
100
Kalanchoe___Kalax.0123s0013.1.p
Kalanchoe___Kalax.0240s0021.1.p
Kalanchoe___Kalax.0158s0052.1.p
96
100
Kalanchoe___Kalax.0047s0093.1.p
Kalanchoe___Kalax.0076s0019.1.p
100
Kalanchoe___Kalax.0007s0051.1.p
Amborella___AmTr_v1.0_scaffold00017.88

# OG0000470

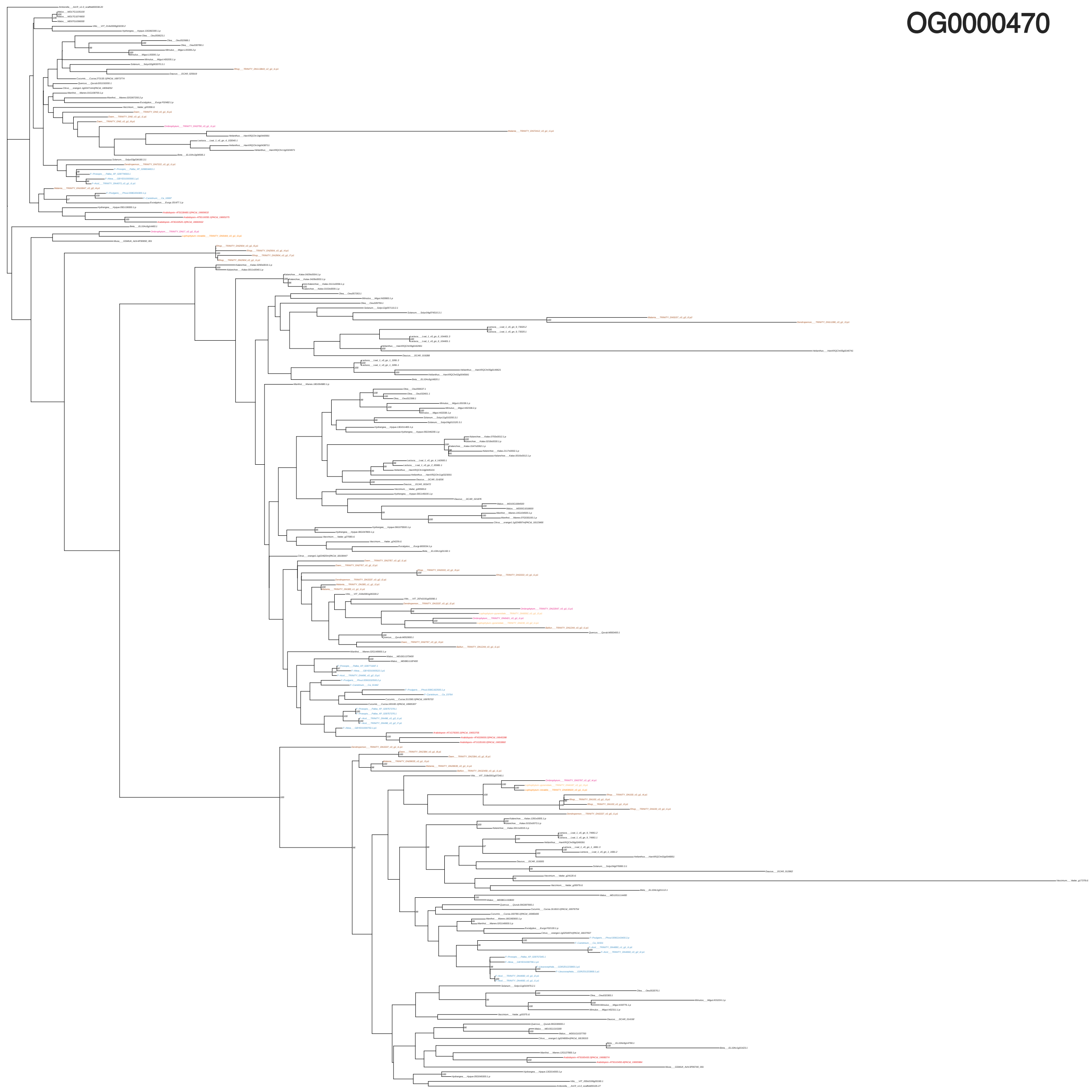

# OG0000564

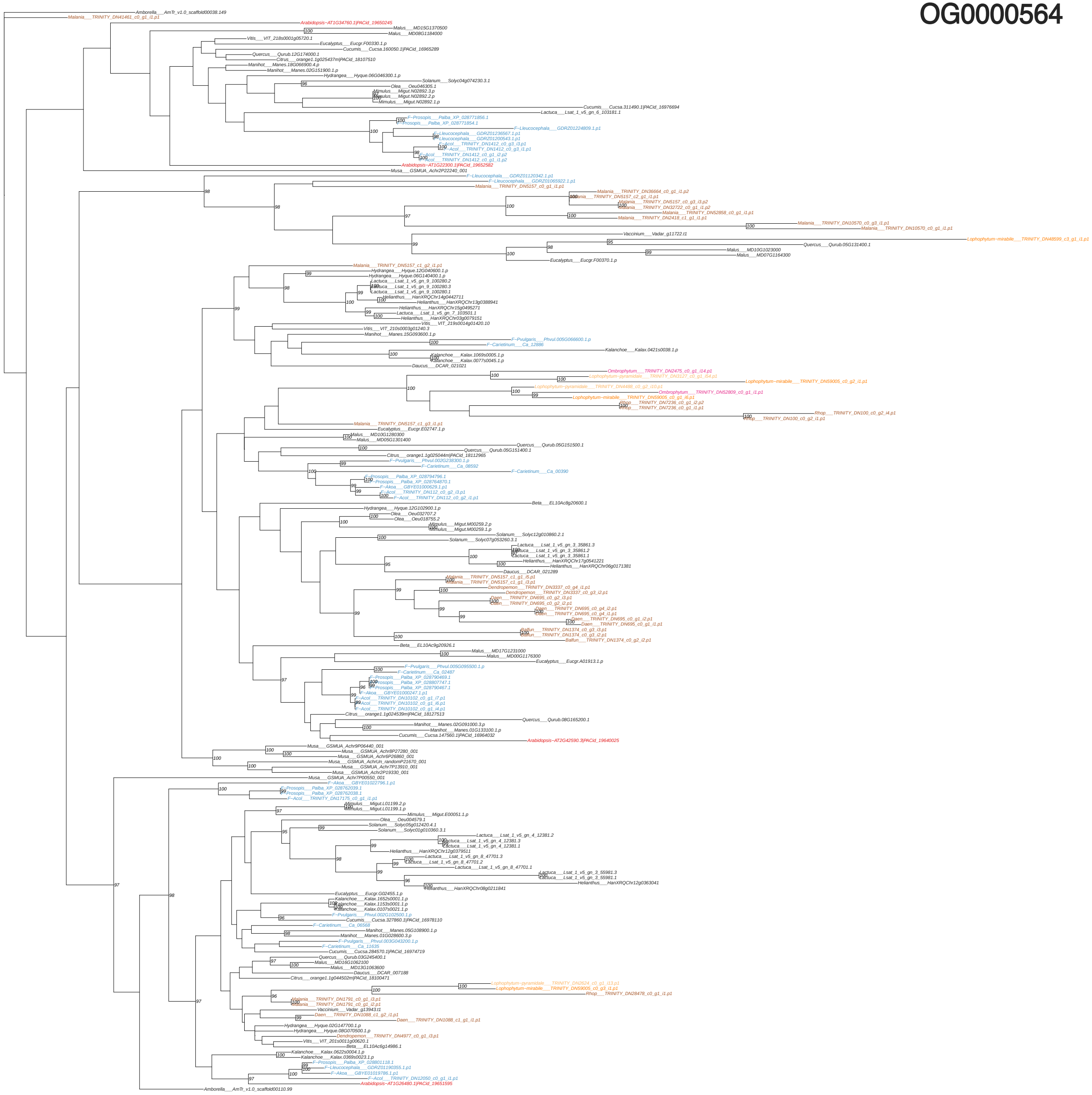

# OG0000636

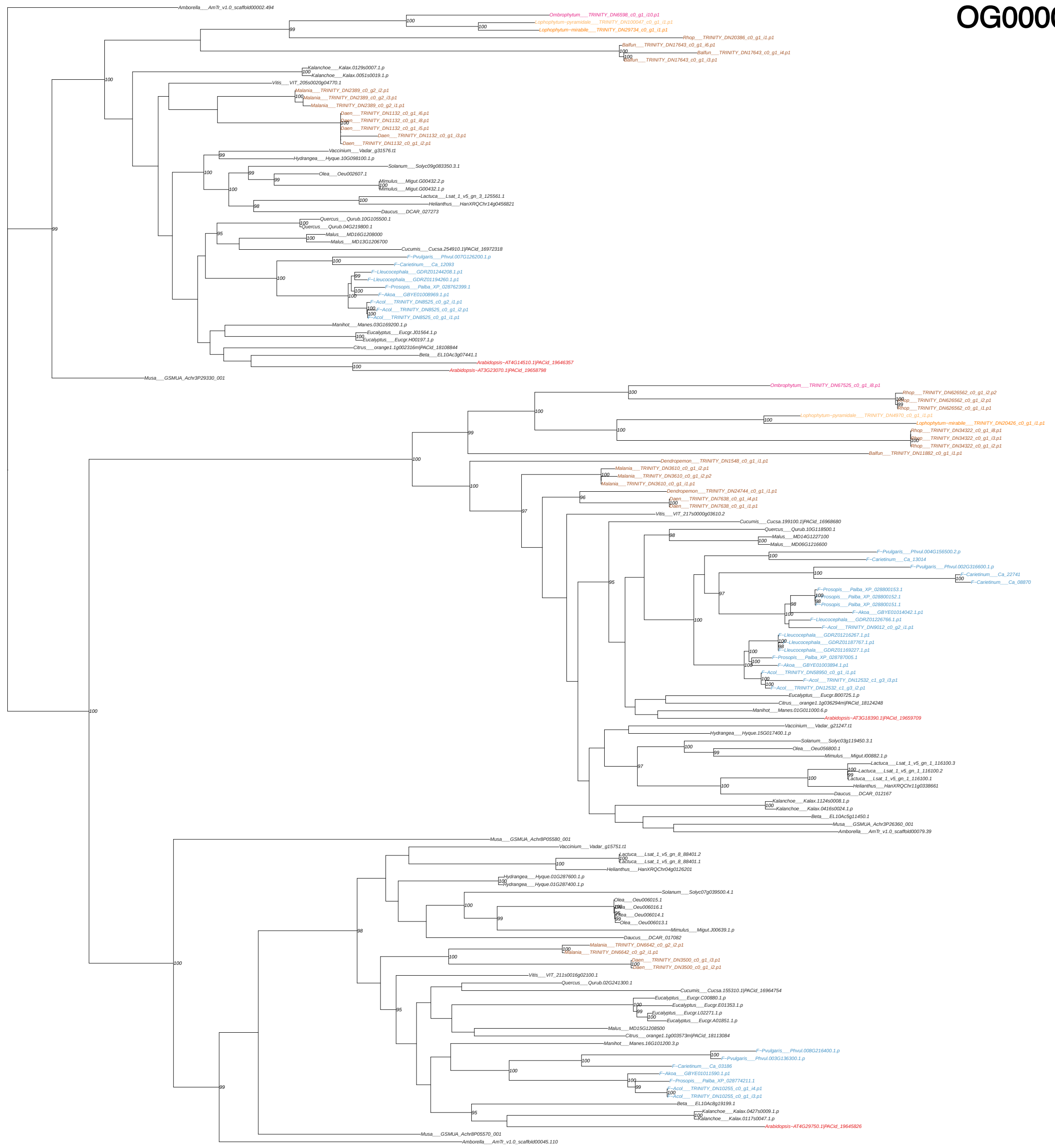

# OG0000730

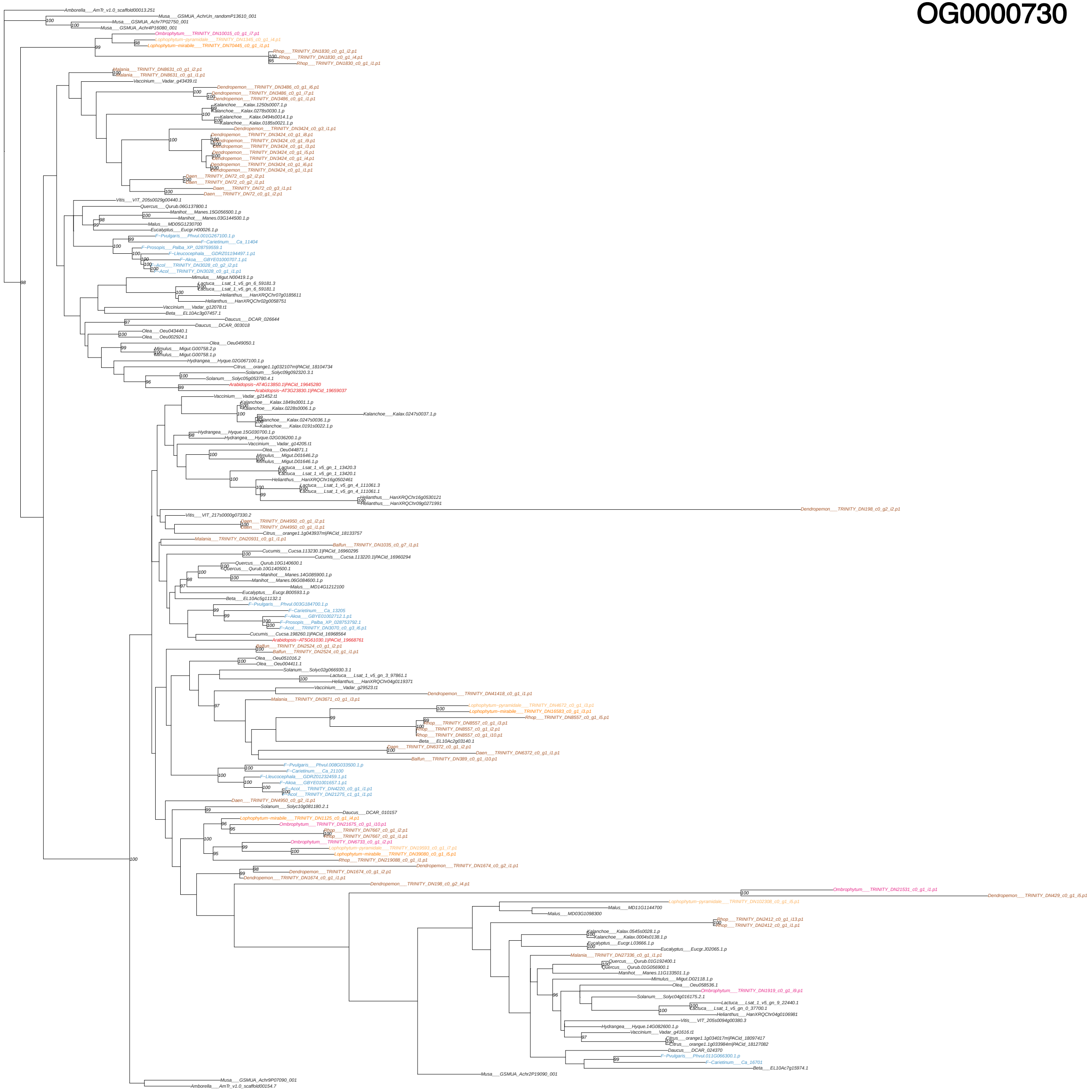

Amborella___AmTr_v1.0_scaffold00013.251
Musa___GSMUA_AchrUn_randomP13610_001
Musa___GSMUA_Achr7P02750_001
Musa___GSMUA_Achr4P16080_001
Ombrophytum___TRINITY_DN10015_c0_g1_i7.p1
Lophophytum~pyramidale___TRINITY_DN1345_c0_g1_i4.p1
Lophophytum~mirabile___TRINITY_DN70445_c0_g1_i1.p1
Rhop___TRINITY_DN1830_c0_g1_i2.p1
Rhop___TRINITY_DN1830_c0_g1_i4.p1
Rhop___TRINITY_DN1830_c0_g1_i1.p1
Malania___TRINITY_DN8631_c0_g1_i2.p1
Malania___TRINITY_DN8631_c0_g1_i1.p1
Vaccinium___Vadar_g43439.t1
Dendropemon___TRINITY_DN3486_c0_g1_i6.p1
Dendropemon___TRINITY_DN3486_c0_g1_i7.p1
Dendropemon___TRINITY_DN3486_c0_g1_i1.p1
Kalanchoe___Kalax.1250s0007.1.p
Kalanchoe___Kalax.0278s0030.1.p
Kalanchoe___Kalax.0494s0014.1.p
Kalanchoe___Kalax.0185s0021.1.p
Dendropemon___TRINITY_DN3424_c0_g3_i1.p1
Dendropemon___TRINITY_DN3424_c0_g1_i8.p1
Dendropemon___TRINITY_DN3424_c0_g1_i9.p1
Dendropemon___TRINITY_DN3424_c0_g1_i3.p1
Dendropemon___TRINITY_DN3424_c0_g1_i5.p1
Dendropemon___TRINITY_DN3424_c0_g1_i4.p1
Dendropemon___TRINITY_DN3424_c0_g1_i6.p1
Dendropemon___TRINITY_DN3424_c0_g1_i1.p1
Daen___TRINITY_DN72_c0_g2_i2.p1
Daen___TRINITY_DN72_c0_g2_i1.p1
Daen___TRINITY_DN72_c0_g3_i1.p1
Daen___TRINITY_DN72_c0_g1_i2.p1
Vitis___VIT_205s0029g00440.1
Quercus___Qurub.06G137800.1
Manihot___Manes.15G056500.1.p
Manihot___Manes.03G144500.1.p
Malus___MD05G1230700
Eucalyptus___Eucgr.H00026.1.p
F~Pvulgaris___Phvul.001G267100.1.p
F~Carietinum___Ca_11404
F~Prosopis___Palba_XP_028759559.1
F~Lleucocephala___GDRZ01194497.1.p1
F~Akoa___GBYE01000707.1.p1
F~Acol___TRINITY_DN3028_c0_g2_i2.p1
F~Acol___TRINITY_DN3028_c0_g1_i1.p1
Mimulus___Migut.N00419.1.p
Lactuca___Lsat_1_v5_gn_6_59181.3
Lactuca___Lsat_1_v5_gn_6_59181.1
Helianthus___HanXRQChr07g0185611
Helianthus___HanXRQChr02g0058751
Vaccinium___Vadar_g12078.t1
Beta___EL10Ac3g07457.1
Daucus___DCAR_026644
Daucus___DCAR_003018
Olea___Oeu043440.1
Olea___Oeu002924.1
Olea___Oeu049050.1
Mimulus___Migut.G00758.2.p
Mimulus___Migut.G00758.1.p
Hydrangea___Hyque.02G067100.1.p
Citrus___orange1.1g032107m|PACid_18104734
Solanum___Solyc09g092320.3.1
Solanum___Solyc05g053780.4.1
Arabidopsis~AT4G13850.1|PACid_19645280
Arabidopsis~AT3G23830.1|PACid_19659037
Vaccinium___Vadar_g21452.t1
Kalanchoe___Kalax.1849s0001.1.p
Kalanchoe___Kalax.0228s0006.1.p
Kalanchoe___Kalax.0247s0037.1.p
Kalanchoe___Kalax.0247s0036.1.p
Kalanchoe___Kalax.0191s0022.1.p
Hydrangea___Hyque.15G030700.1.p
Hydrangea___Hyque.02G036200.1.p
Vaccinium___Vadar_g14205.t1
Olea___Oeu044871.1
Mimulus___Migut.D01646.2.p
Mimulus___Migut.D01646.1.p
Lactuca___Lsat_1_v5_gn_1_13420.3
Lactuca___Lsat_1_v5_gn_1_13420.1
Helianthus___HanXRQChr16g0502461
Lactuca___Lsat_1_v5_gn_4_111061.3
Lactuca___Lsat_1_v5_gn_4_111061.1
Helianthus___HanXRQChr16g0530121
Helianthus___HanXRQChr09g0271991
Dendropemon___TRINITY_DN198_c0_g2_i2.p1
Vitis___VIT_217s0000g07330.2
Daen___TRINITY_DN4950_c0_g1_i2.p1
Daen___TRINITY_DN4950_c0_g1_i1.p1
Citrus___orange1.1g043937m|PACid_18133757
Malania___TRINITY_DN20931_c0_g1_i1.p1
Balfun___TRINITY_DN1035_c0_g7_i1.p1
Cucumis___Cucsa.113230.1|PACid_16960295
Cucumis___Cucsa.113220.1|PACid_16960294
Quercus___Qurub.10G140600.1
Quercus___Qurub.10G140500.1
Manihot___Manes.14G085900.1.p
Manihot___Manes.06G084600.1.p
Malus___MD14G1212100
Eucalyptus___Eucgr.B00593.1.p
Beta___EL10Ac5g11132.1
F~Pvulgaris___Phvul.003G184700.1.p
F~Carietinum___Ca_13205
F~Akoa___GBYE01002712.1.p1
F~Prosopis___Palba_XP_028753792.1
F~Acol___TRINITY_DN3070_c0_g3_i6.p1
Cucumis___Cucsa.198260.1|PACid_16968564
Arabidopsis~AT5G61030.1|PACid_19668761
Balfun___TRINITY_DN2524_c0_g1_i2.p1
Balfun___TRINITY_DN2524_c0_g1_i1.p1
Olea___Oeu051016.2
Olea___Oeu004411.1
Solanum___Solyc02g066930.3.1
Lactuca___Lsat_1_v5_gn_3_97861.1
Helianthus___HanXRQChr04g0119371
Vaccinium___Vadar_g29523.t1
Dendropemon___TRINITY_DN41418_c0_g1_i1.p1
Malania___TRINITY_DN3671_c0_g1_i3.p1
Lophophytum~pyramidale___TRINITY_DN4672_c0_g1_i3.p1
Lophophytum~mirabile___TRINITY_DN16583_c0_g1_i3.p1
Rhop___TRINITY_DN8557_c0_g1_i5.p1
Rhop___TRINITY_DN8557_c0_g1_i3.p1
Rhop___TRINITY_DN8557_c0_g1_i2.p1
Rhop___TRINITY_DN8557_c0_g1_i10.p1
Beta___EL10Ac2g03140.1
Daen___TRINITY_DN6372_c0_g1_i2.p1
Daen___TRINITY_DN6372_c0_g1_i1.p1
Balfun___TRINITY_DN389_c0_g1_i10.p1
F~Pvulgaris___Phvul.008G033500.1.p
F~Carietinum___Ca_21100
F~Lleucocephala___GDRZ01232459.1.p1
F~Akoa___GBYE01001657.1.p1
F~Acol___TRINITY_DN4220_c0_g1_i1.p1
F~Acol___TRINITY_DN21275_c1_g1_i1.p1
Daen___TRINITY_DN4950_c0_g2_i1.p1
Solanum___Solyc10g081180.2.1
Daucus___DCAR_010157
Lophophytum~mirabile___TRINITY_DN1125_c0_g1_i4.p1
Ombrophytum___TRINITY_DN21675_c0_g1_i10.p1
Rhop___TRINITY_DN7667_c0_g1_i2.p1
Rhop___TRINITY_DN7667_c0_g1_i1.p1
Ombrophytum___TRINITY_DN6733_c0_g1_i2.p1
Lophophytum~pyramidale___TRINITY_DN19593_c0_g1_i7.p1
Lophophytum~mirabile___TRINITY_DN39080_c0_g1_i5.p1
Rhop___TRINITY_DN219088_c0_g1_i1.p1
Dendropemon___TRINITY_DN1674_c0_g2_i1.p1
Dendropemon___TRINITY_DN1674_c0_g1_i2.p1
Dendropemon___TRINITY_DN1674_c0_g1_i1.p1
Dendropemon___TRINITY_DN198_c0_g2_i4.p1
Ombrophytum___TRINITY_DN21531_c0_g1_i1.p1
Dendropemon___TRINITY_DN429_c0_g1_i5.p1
Lophophytum~pyramidale___TRINITY_DN102308_c0_g1_i5.p1
Malus___MD11G1144700
Malus___MD03G1098300
Rhop___TRINITY_DN2412_c0_g1_i13.p1
Rhop___TRINITY_DN2412_c0_g1_i1.p1
Kalanchoe___Kalax.0545s0028.1.p
Kalanchoe___Kalax.0004s0138.1.p
Eucalyptus___Eucgr.L03666.1.p
Eucalyptus___Eucgr.J02065.1.p
Malania___TRINITY_DN27336_c0_g1_i1.p1
Quercus___Qurub.01G192400.1
Quercus___Qurub.01G056900.1
Manihot___Manes.11G133501.1.p
Mimulus___Migut.D02118.1.p
Olea___Oeu058536.1
Ombrophytum___TRINITY_DN1919_c0_g1_i9.p1
Solanum___Solyc04g016175.2.1
Lactuca___Lsat_1_v5_gn_9_22440.1
Lactuca___Lsat_1_v5_gn_0_37700.1
Helianthus___HanXRQChr04g0106981
Vitis___VIT_205s0094g00380.3
Hydrangea___Hyque.14G082600.1.p
Vaccinium___Vadar_g41616.t1
Citrus___orange1.1g034017m|PACid_18097417
Citrus___orange1.1g033984m|PACid_18127082
Daucus___DCAR_024370
F~Pvulgaris___Phvul.011G066300.1.p
F~Carietinum___Ca_16701
Beta___EL10Ac7g15974.1
Musa___GSMUA_Achr2P19090_001
Musa___GSMUA_Achr9P07090_001
Amborella___AmTr_v1.0_scaffold00154.7

# OG0000807

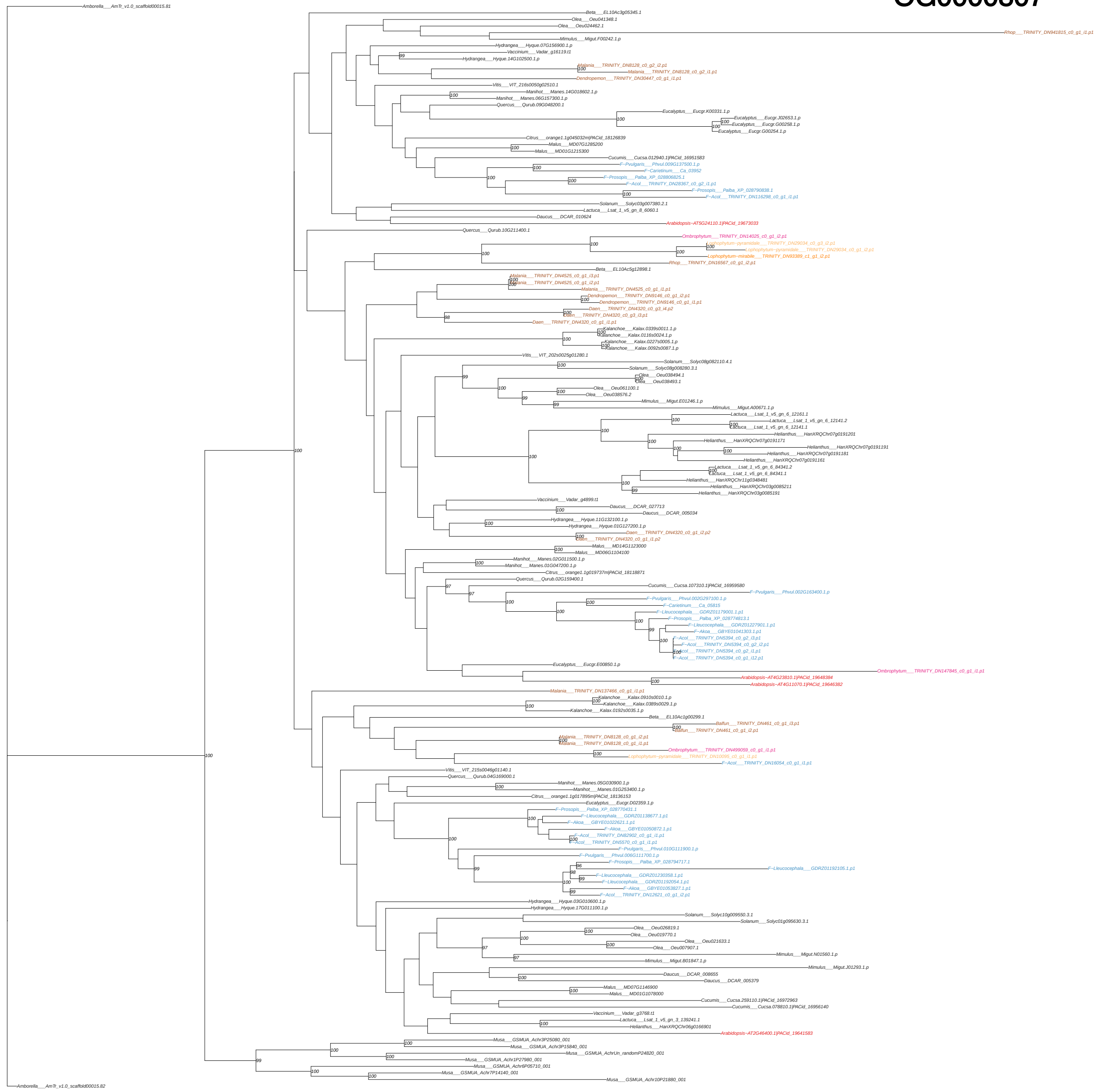

# OG0000848

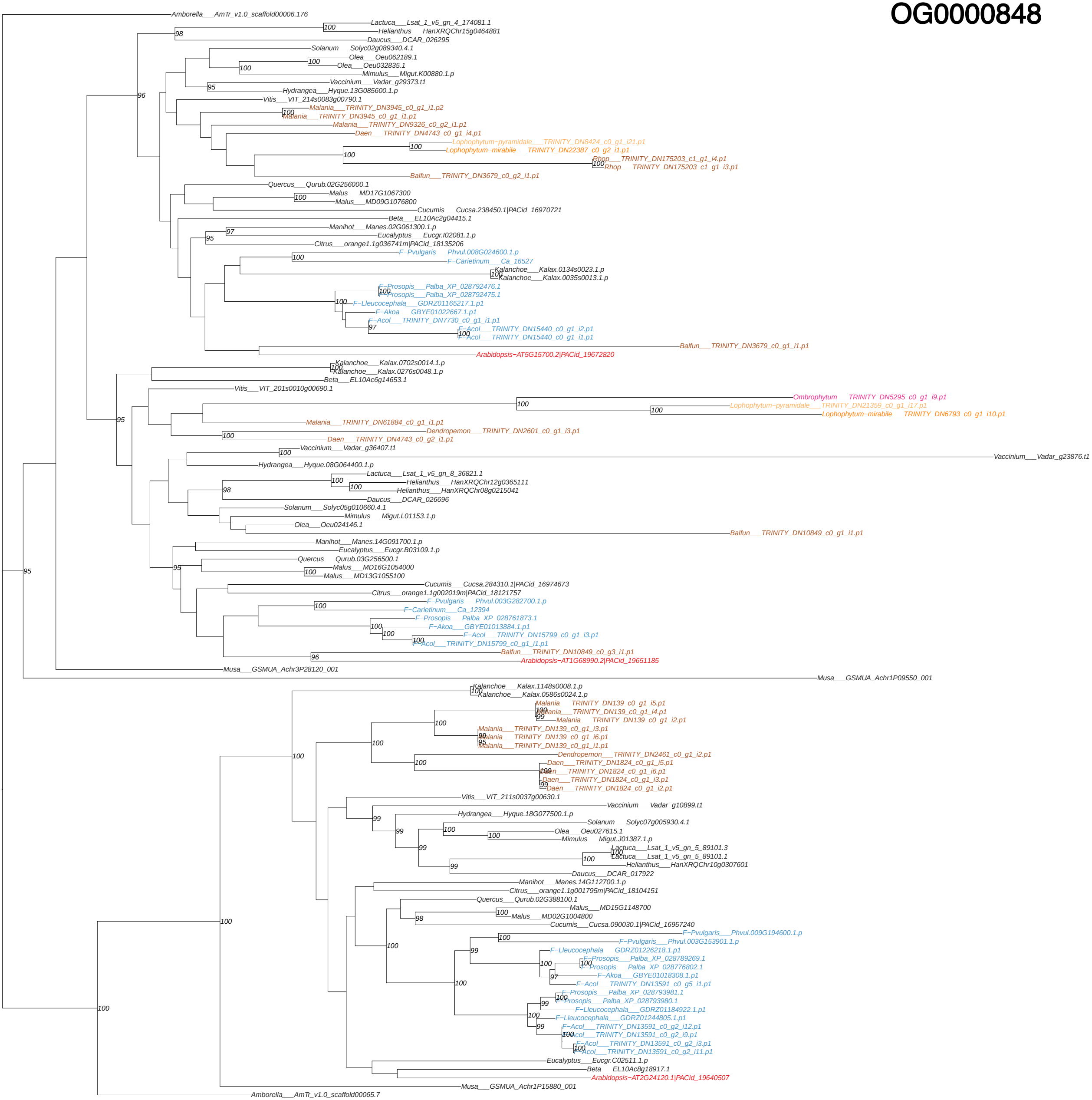

Amborella___AmTr_v1.0_scaffold00006.176
Lactuca___Lsat_1_v5_gn_4_174081.1
Helianthus___HanXRQChr15g0464881
Daucus___DCAR_026295
Solanum___Solyc02g089340.4.1
Olea___Oeu062189.1
Olea___Oeu032835.1
Mimulus___Migut.K00880.1.p
Vaccinium___Vadar_g29373.t1
Hydrangea___Hyque.13G085600.1.p
Vitis___VIT_214s0083g00790.1
Malania___TRINITY_DN3945_c0_g1_i1.p2
Malania___TRINITY_DN3945_c0_g1_i1.p1
Malania___TRINITY_DN9326_c0_g2_i1.p1
Daen___TRINITY_DN4743_c0_g1_i4.p1
Lophophytum−pyramidale___TRINITY_DN8424_c0_g1_i21.p1
Lophophytum−mirabile___TRINITY_DN22387_c0_g2_i1.p1
Rhop___TRINITY_DN175203_c1_g1_i4.p1
Rhop___TRINITY_DN175203_c1_g1_i3.p1
Balfun___TRINITY_DN3679_c0_g2_i1.p1
Quercus___Qurub.02G256000.1
Malus___MD17G1067300
Malus___MD09G1076800
Cucumis___Cucsa.238450.1|PACid_16970721
Beta___EL10Ac2g04415.1
Manihot___Manes.02G061300.1.p
Eucalyptus___Eucgr.I02081.1.p
Citrus___orange1.1g036741m|PACid_18135206
F−Pvulgaris___Phvul.008G024600.1.p
F−Carietinum___Ca_16527
Kalanchoe___Kalax.0134s0023.1.p
Kalanchoe___Kalax.0035s0013.1.p
F−Prosopis___Palba_XP_028792476.1
F−Prosopis___Palba_XP_028792475.1
F−Lleucocephala___GDRZ01165217.1.p1
F−Akoa___GBYE01022667.1.p1
F−Acol___TRINITY_DN7730_c0_g1_i1.p1
F−Acol___TRINITY_DN15440_c0_g1_i2.p1
F−Acol___TRINITY_DN15440_c0_g1_i1.p1
Balfun___TRINITY_DN3679_c0_g1_i1.p1
Arabidopsis−AT5G15700.2|PACid_19672820
Kalanchoe___Kalax.0702s0014.1.p
Kalanchoe___Kalax.0276s0048.1.p
Beta___EL10Ac6g14653.1
Vitis___VIT_201s0010g00690.1
Ombrophytum___TRINITY_DN5295_c0_g1_i9.p1
Lophophytum−pyramidale___TRINITY_DN21359_c0_g1_i17.p1
Lophophytum−mirabile___TRINITY_DN6793_c0_g1_i10.p1
Malania___TRINITY_DN61884_c0_g1_i1.p1
Dendropemon___TRINITY_DN2601_c0_g1_i3.p1
Daen___TRINITY_DN4743_c0_g2_i1.p1
Vaccinium___Vadar_g36407.t1
Vaccinium___Vadar_g23876.t1
Hydrangea___Hyque.08G064400.1.p
Lactuca___Lsat_1_v5_gn_8_36821.1
Helianthus___HanXRQChr12g0365111
Helianthus___HanXRQChr08g0215041
Daucus___DCAR_026696
Solanum___Solyc05g010660.4.1
Mimulus___Migut.L01153.1.p
Olea___Oeu024146.1
Balfun___TRINITY_DN10849_c0_g1_i1.p1
Manihot___Manes.14G091700.1.p
Eucalyptus___Eucgr.B03109.1.p
Quercus___Qurub.03G256500.1
Malus___MD16G1054000
Malus___MD13G1055100
Cucumis___Cucsa.284310.1|PACid_16974673
Citrus___orange1.1g002019m|PACid_18121757
F−Pvulgaris___Phvul.003G282700.1.p
F−Carietinum___Ca_12394
F−Prosopis___Palba_XP_028761873.1
F−Akoa___GBYE01013884.1.p1
F−Acol___TRINITY_DN15799_c0_g1_i3.p1
F−Acol___TRINITY_DN15799_c0_g1_i1.p1
Balfun___TRINITY_DN10849_c0_g3_i1.p1
Arabidopsis−AT1G68990.2|PACid_19651185
Musa___GSMUA_Achr3P28120_001
Musa___GSMUA_Achr1P09550_001
Kalanchoe___Kalax.1148s0008.1.p
Kalanchoe___Kalax.0586s0024.1.p
Malania___TRINITY_DN139_c0_g1_i5.p1
Malania___TRINITY_DN139_c0_g1_i4.p1
Malania___TRINITY_DN139_c0_g1_i2.p1
Malania___TRINITY_DN139_c0_g1_i3.p1
Malania___TRINITY_DN139_c0_g1_i6.p1
Malania___TRINITY_DN139_c0_g1_i1.p1
Dendropemon___TRINITY_DN2461_c0_g1_i2.p1
Daen___TRINITY_DN1824_c0_g1_i5.p1
Daen___TRINITY_DN1824_c0_g1_i6.p1
Daen___TRINITY_DN1824_c0_g1_i3.p1
Daen___TRINITY_DN1824_c0_g1_i2.p1
Vitis___VIT_211s0037g00630.1
Vaccinium___Vadar_g10899.t1
Hydrangea___Hyque.18G077500.1.p
Solanum___Solyc07g005930.4.1
Olea___Oeu027615.1
Mimulus___Migut.J01387.1.p
Lactuca___Lsat_1_v5_gn_5_89101.3
Lactuca___Lsat_1_v5_gn_5_89101.1
Helianthus___HanXRQChr10g0307601
Daucus___DCAR_017922
Manihot___Manes.14G112700.1.p
Citrus___orange1.1g001795m|PACid_18104151
Quercus___Qurub.02G388100.1
Malus___MD15G1148700
Malus___MD02G1004800
Cucumis___Cucsa.090030.1|PACid_16957240
F−Pvulgaris___Phvul.009G194600.1.p
F−Pvulgaris___Phvul.003G153901.1.p
F−Lleucocephala___GDRZ01226218.1.p1
F−Prosopis___Palba_XP_028789269.1
F−Prosopis___Palba_XP_028776802.1
F−Akoa___GBYE01018308.1.p1
F−Acol___TRINITY_DN13591_c0_g5_i1.p1
F−Prosopis___Palba_XP_028793981.1
F−Prosopis___Palba_XP_028793980.1
F−Lleucocephala___GDRZ01184922.1.p1
F−Lleucocephala___GDRZ01244805.1.p1
F−Acol___TRINITY_DN13591_c0_g2_i12.p1
F−Acol___TRINITY_DN13591_c0_g2_i9.p1
F−Acol___TRINITY_DN13591_c0_g2_i3.p1
F−Acol___TRINITY_DN13591_c0_g2_i11.p1
Eucalyptus___Eucgr.C02511.1.p
Beta___EL10Ac8g18917.1
Arabidopsis−AT2G24120.1|PACid_19640507
Musa___GSMUA_Achr1P15880_001
Amborella___AmTr_v1.0_scaffold00065.7

# OG0001036

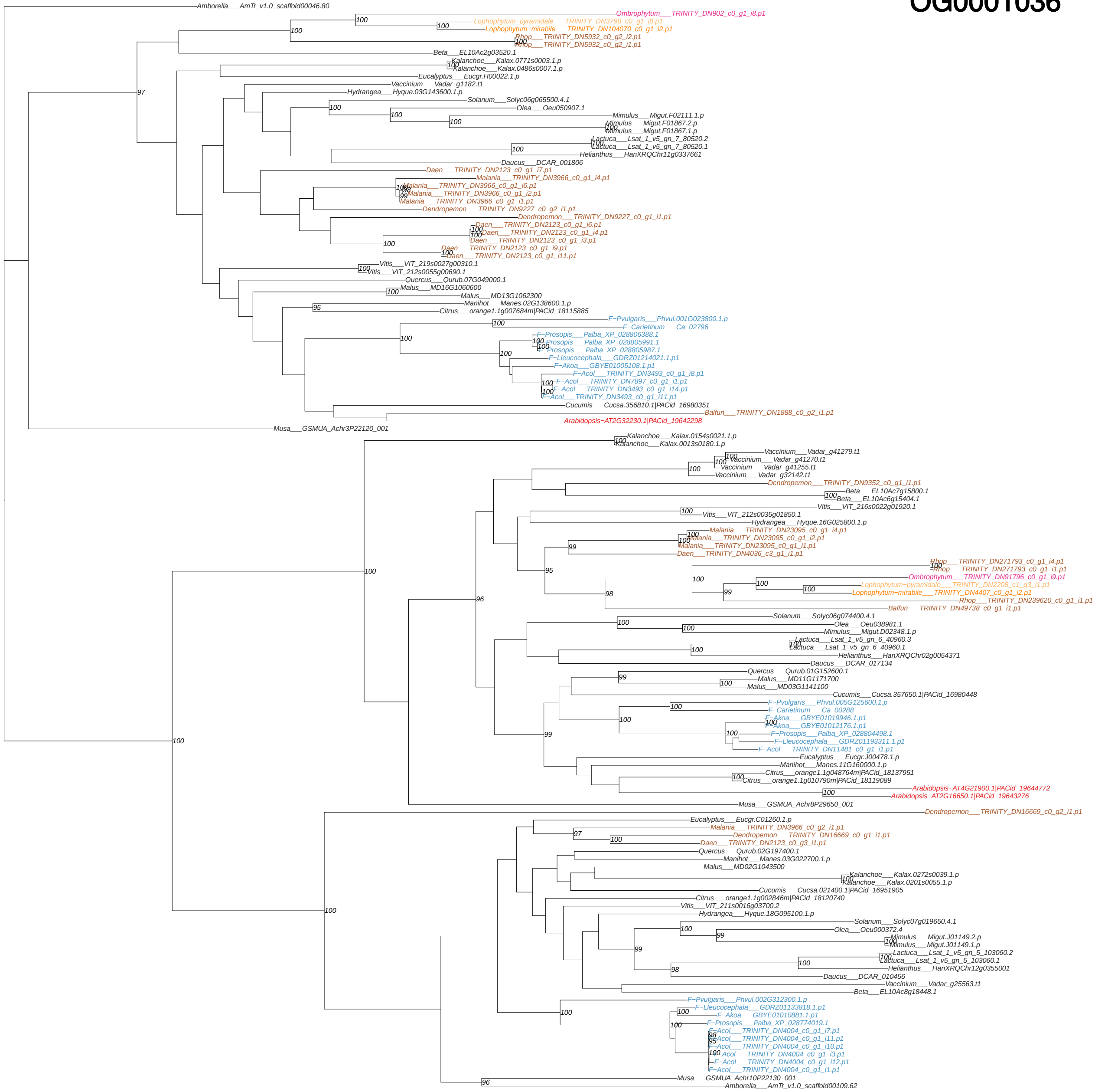

Amborella___AmTr_v1.0_scaffold00046.80
Ombrophytum___TRINITY_DN902_c0_g1_i8.p1
Lophophytum−pyramidale___TRINITY_DN3798_c0_g1_i8.p1
Lophophytum−mirabile___TRINITY_DN104070_c0_g1_i2.p1
Rhop___TRINITY_DN5932_c0_g2_i2.p1
Rhop___TRINITY_DN5932_c0_g2_i1.p1
Beta___EL10Ac2g03520.1
Kalanchoe___Kalax.0771s0003.1.p
Kalanchoe___Kalax.0486s0007.1.p
Eucalyptus___Eucgr.H00022.1.p
Vaccinium___Vadar_g1182.t1
Hydrangea___Hyque.03G143600.1.p
Solanum___Solyc06g065500.4.1
Olea___Oeu050907.1
Mimulus___Migut.F02111.1.p
Mimulus___Migut.F01867.2.p
Mimulus___Migut.F01867.1.p
Lactuca___Lsat_1_v5_gn_7_80520.2
Lactuca___Lsat_1_v5_gn_7_80520.1
Helianthus___HanXRQChr11g0337661
Daucus___DCAR_001806
Daen___TRINITY_DN2123_c0_g1_i7.p1
Malania___TRINITY_DN3966_c0_g1_i4.p1
Malania___TRINITY_DN3966_c0_g1_i6.p1
Malania___TRINITY_DN3966_c0_g1_i2.p1
Malania___TRINITY_DN3966_c0_g1_i1.p1
Dendropemon___TRINITY_DN9227_c0_g2_i1.p1
Dendropemon___TRINITY_DN9227_c0_g1_i1.p1
Daen___TRINITY_DN2123_c0_g1_i6.p1
Daen___TRINITY_DN2123_c0_g1_i4.p1
Daen___TRINITY_DN2123_c0_g1_i3.p1
Daen___TRINITY_DN2123_c0_g1_i9.p1
Daen___TRINITY_DN2123_c0_g1_i11.p1
Vitis___VIT_219s0027g00310.1
Vitis___VIT_212s0055g00690.1
Quercus___Qurub.07G049000.1
Malus___MD16G1060600
Malus___MD13G1062300
Manihot___Manes.02G138600.1.p
Citrus___orange1.1g007684m|PACid_18115885
F−Pvulgaris___Phvul.001G023800.1.p
F−Carietinum___Ca_02796
F−Prosopis___Palba_XP_028806388.1
F−Prosopis___Palba_XP_028805991.1
F−Prosopis___Palba_XP_028805987.1
F−Lleucocephala___GDRZ01214021.1.p1
F−Akoa___GBYE01005108.1.p1
F−Acol___TRINITY_DN3493_c0_g1_i8.p1
F−Acol___TRINITY_DN7897_c0_g1_i1.p1
F−Acol___TRINITY_DN3493_c0_g1_i14.p1
F−Acol___TRINITY_DN3493_c0_g1_i11.p1
Cucumis___Cucsa.356810.1|PACid_16980351
Balfun___TRINITY_DN1888_c0_g2_i1.p1
Arabidopsis−AT2G32230.1|PACid_19642298
Musa___GSMUA_Achr3P22120_001
Kalanchoe___Kalax.0154s0021.1.p
Kalanchoe___Kalax.0013s0180.1.p
Vaccinium___Vadar_g41279.t1
Vaccinium___Vadar_g41270.t1
Vaccinium___Vadar_g41255.t1
Vaccinium___Vadar_g32142.t1
Dendropemon___TRINITY_DN9352_c0_g1_i1.p1
Beta___EL10Ac7g15800.1
Beta___EL10Ac6g15404.1
Vitis___VIT_216s0022g01920.1
Vitis___VIT_212s0035g01850.1
Hydrangea___Hyque.16G025800.1.p
Malania___TRINITY_DN23095_c0_g1_i4.p1
Malania___TRINITY_DN23095_c0_g1_i2.p1
Malania___TRINITY_DN23095_c0_g1_i1.p1
Daen___TRINITY_DN4036_c3_g1_i1.p1
Rhop___TRINITY_DN271793_c0_g1_i4.p1
Rhop___TRINITY_DN271793_c0_g1_i1.p1
Ombrophytum___TRINITY_DN91796_c0_g1_i9.p1
Lophophytum−pyramidale___TRINITY_DN2208_c1_g3_i1.p1
Lophophytum−mirabile___TRINITY_DN4407_c0_g1_i2.p1
Rhop___TRINITY_DN239620_c0_g1_i1.p1
Balfun___TRINITY_DN49738_c0_g1_i1.p1
Solanum___Solyc06g074400.4.1
Olea___Oeu038981.1
Mimulus___Migut.D02348.1.p
Lactuca___Lsat_1_v5_gn_6_40960.3
Lactuca___Lsat_1_v5_gn_6_40960.1
Helianthus___HanXRQChr02g0054371
Daucus___DCAR_017134
Quercus___Qurub.01G152600.1
Malus___MD11G1171700
Malus___MD03G1141100
Cucumis___Cucsa.357650.1|PACid_16980448
F−Pvulgaris___Phvul.005G125600.1.p
F−Carietinum___Ca_00288
F−Akoa___GBYE01019946.1.p1
F−Akoa___GBYE01012176.1.p1
F−Prosopis___Palba_XP_028804498.1
F−Lleucocephala___GDRZ01193311.1.p1
F−Acol___TRINITY_DN11481_c0_g1_i1.p1
Eucalyptus___Eucgr.J00478.1.p
Manihot___Manes.11G160000.1.p
Citrus___orange1.1g048764m|PACid_18137951
Citrus___orange1.1g010790m|PACid_18119089
Arabidopsis−AT4G21900.1|PACid_19644772
Arabidopsis−AT2G16650.1|PACid_19643276
Musa___GSMUA_Achr8P29650_001
Dendropemon___TRINITY_DN16669_c0_g2_i1.p1
Eucalyptus___Eucgr.C01260.1.p
Malania___TRINITY_DN3966_c0_g2_i1.p1
Dendropemon___TRINITY_DN16669_c0_g1_i1.p1
Daen___TRINITY_DN2123_c0_g3_i1.p1
Quercus___Qurub.02G197400.1
Manihot___Manes.03G022700.1.p
Malus___MD02G1043500
Kalanchoe___Kalax.0272s0039.1.p
Kalanchoe___Kalax.0201s0055.1.p
Cucumis___Cucsa.021400.1|PACid_16951905
Citrus___orange1.1g002846m|PACid_18120740
Vitis___VIT_211s0016g03700.2
Hydrangea___Hyque.18G095100.1.p
Solanum___Solyc07g019650.4.1
Olea___Oeu000372.4
Mimulus___Migut.J01149.2.p
Mimulus___Migut.J01149.1.p
Lactuca___Lsat_1_v5_gn_5_103060.2
Lactuca___Lsat_1_v5_gn_5_103060.1
Helianthus___HanXRQChr12g0355001
Daucus___DCAR_010456
Vaccinium___Vadar_g25563.t1
Beta___EL10Ac8g18448.1
F−Pvulgaris___Phvul.002G312300.1.p
F−Lleucocephala___GDRZ01133818.1.p1
F−Akoa___GBYE01010881.1.p1
F−Prosopis___Palba_XP_028774019.1
F−Acol___TRINITY_DN4004_c0_g1_i7.p1
F−Acol___TRINITY_DN4004_c0_g1_i11.p1
F−Acol___TRINITY_DN4004_c0_g1_i10.p1
F−Acol___TRINITY_DN4004_c0_g1_i3.p1
F−Acol___TRINITY_DN4004_c0_g1_i12.p1
F−Acol___TRINITY_DN4004_c0_g1_i1.p1
Musa___GSMUA_Achr10P22130_001
Amborella___AmTr_v1.0_scaffold00109.62

# OG0001078

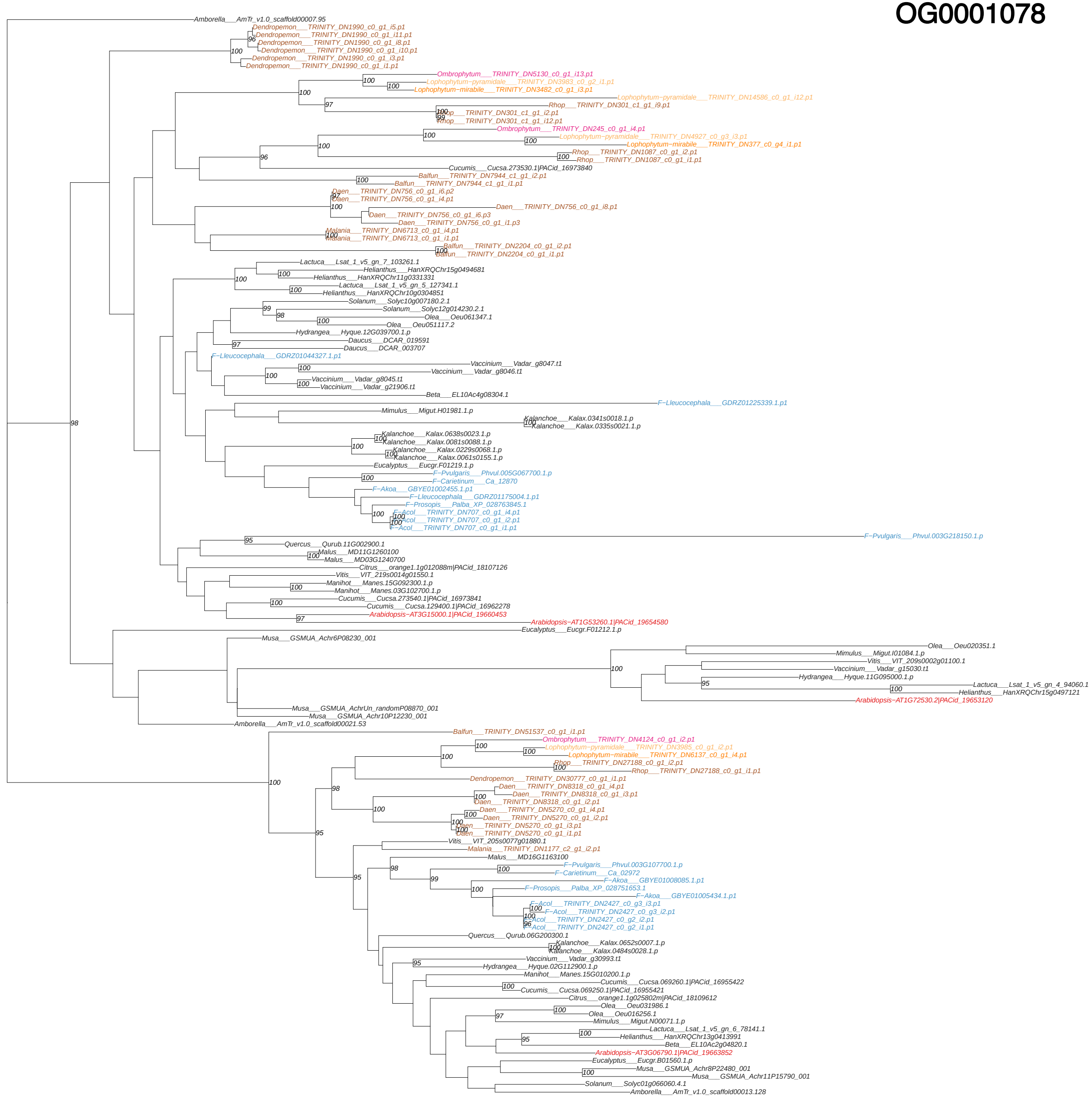

Amborella___AmTr_v1.0_scaffold00007.95
Dendropemon___TRINITY_DN1990_c0_g1_i5.p1
Dendropemon___TRINITY_DN1990_c0_g1_i11.p1
Dendropemon___TRINITY_DN1990_c0_g1_i8.p1
Dendropemon___TRINITY_DN1990_c0_g1_i10.p1
Dendropemon___TRINITY_DN1990_c0_g1_i3.p1
Dendropemon___TRINITY_DN1990_c0_g1_i1.p1
Ombrophytum___TRINITY_DN5130_c0_g1_i13.p1
Lophophytum−pyramidale___TRINITY_DN3983_c0_g2_i1.p1
Lophophytum−mirabile___TRINITY_DN3482_c0_g1_i3.p1
Lophophytum−pyramidale___TRINITY_DN14586_c0_g1_i12.p1
Rhop___TRINITY_DN301_c1_g1_i9.p1
Rhop___TRINITY_DN301_c1_g1_i2.p1
Rhop___TRINITY_DN301_c1_g1_i12.p1
Ombrophytum___TRINITY_DN245_c0_g1_i4.p1
Lophophytum−pyramidale___TRINITY_DN4927_c0_g3_i3.p1
Lophophytum−mirabile___TRINITY_DN377_c0_g4_i1.p1
Rhop___TRINITY_DN1087_c0_g1_i2.p1
Rhop___TRINITY_DN1087_c0_g1_i1.p1
Cucumis___Cucsa.273530.1|PACid_16973840
Balfun___TRINITY_DN7944_c1_g1_i2.p1
Balfun___TRINITY_DN7944_c1_g1_i1.p1
Daen___TRINITY_DN756_c0_g1_i6.p2
Daen___TRINITY_DN756_c0_g1_i4.p1
Daen___TRINITY_DN756_c0_g1_i8.p1
Daen___TRINITY_DN756_c0_g1_i6.p3
Daen___TRINITY_DN756_c0_g1_i1.p3
Malania___TRINITY_DN6713_c0_g1_i4.p1
Malania___TRINITY_DN6713_c0_g1_i1.p1
Balfun___TRINITY_DN2204_c0_g1_i2.p1
Balfun___TRINITY_DN2204_c0_g1_i1.p1
Lactuca___Lsat_1_v5_gn_7_103261.1
Helianthus___HanXRQChr15g0494681
Helianthus___HanXRQChr11g0331331
Lactuca___Lsat_1_v5_gn_5_127341.1
Helianthus___HanXRQChr10g0304851
Solanum___Solyc10g007180.2.1
Solanum___Solyc12g014230.2.1
Olea___Oeu061347.1
Olea___Oeu051117.2
Hydrangea___Hyque.12G039700.1.p
Daucus___DCAR_019591
Daucus___DCAR_003707
F−Lleucocephala___GDRZ01044327.1.p1
Vaccinium___Vadar_g8047.t1
Vaccinium___Vadar_g8046.t1
Vaccinium___Vadar_g8045.t1
Vaccinium___Vadar_g21906.t1
Beta___EL10Ac4g08304.1
F−Lleucocephala___GDRZ01225339.1.p1
Mimulus___Migut.H01981.1.p
Kalanchoe___Kalax.0341s0018.1.p
Kalanchoe___Kalax.0335s0021.1.p
Kalanchoe___Kalax.0638s0023.1.p
Kalanchoe___Kalax.0081s0088.1.p
Kalanchoe___Kalax.0229s0068.1.p
Kalanchoe___Kalax.0061s0155.1.p
Eucalyptus___Eucgr.F01219.1.p
F−Pvulgaris___Phvul.005G067700.1.p
F−Carietinum___Ca_12870
F−Akoa___GBYE01002455.1.p1
F−Lleucocephala___GDRZ01175004.1.p1
F−Prosopis___Palba_XP_028763845.1
F−Acol___TRINITY_DN707_c0_g1_i4.p1
F−Acol___TRINITY_DN707_c0_g1_i2.p1
F−Acol___TRINITY_DN707_c0_g1_i1.p1
F−Pvulgaris___Phvul.003G218150.1.p
Quercus___Qurub.11G002900.1
Malus___MD11G1260100
Malus___MD03G1240700
Citrus___orange1.1g012088m|PACid_18107126
Vitis___VIT_219s0014g01550.1
Manihot___Manes.15G092300.1.p
Manihot___Manes.03G102700.1.p
Cucumis___Cucsa.273540.1|PACid_16973841
Cucumis___Cucsa.129400.1|PACid_16962278
Arabidopsis−AT3G15000.1|PACid_19660453
Arabidopsis−AT1G53260.1|PACid_19654580
Eucalyptus___Eucgr.F01212.1.p
Musa___GSMUA_Achr6P08230_001
Olea___Oeu020351.1
Mimulus___Migut.I01084.1.p
Vitis___VIT_209s0002g01100.1
Vaccinium___Vadar_g15030.t1
Hydrangea___Hyque.11G095000.1.p
Lactuca___Lsat_1_v5_gn_4_94060.1
Helianthus___HanXRQChr15g0497121
Arabidopsis−AT1G72530.2|PACid_19653120
Musa___GSMUA_AchrUn_randomP08870_001
Musa___GSMUA_Achr10P12230_001
Amborella___AmTr_v1.0_scaffold00021.53
Balfun___TRINITY_DN51537_c0_g1_i1.p1
Ombrophytum___TRINITY_DN4124_c0_g1_i2.p1
Lophophytum−pyramidale___TRINITY_DN3985_c0_g1_i2.p1
Lophophytum−mirabile___TRINITY_DN6137_c0_g1_i4.p1
Rhop___TRINITY_DN27188_c0_g1_i2.p1
Rhop___TRINITY_DN27188_c0_g1_i1.p1
Dendropemon___TRINITY_DN30777_c0_g1_i1.p1
Daen___TRINITY_DN8318_c0_g1_i4.p1
Daen___TRINITY_DN8318_c0_g1_i3.p1
Daen___TRINITY_DN8318_c0_g1_i2.p1
Daen___TRINITY_DN5270_c0_g1_i4.p1
Daen___TRINITY_DN5270_c0_g1_i2.p1
Daen___TRINITY_DN5270_c0_g1_i3.p1
Daen___TRINITY_DN5270_c0_g1_i1.p1
Vitis___VIT_205s0077g01880.1
Malania___TRINITY_DN1177_c2_g1_i2.p1
Malus___MD16G1163100
F−Pvulgaris___Phvul.003G107700.1.p
F−Carietinum___Ca_02972
F−Akoa___GBYE01008085.1.p1
F−Prosopis___Palba_XP_028751653.1
F−Akoa___GBYE01005434.1.p1
F−Acol___TRINITY_DN2427_c0_g3_i3.p1
F−Acol___TRINITY_DN2427_c0_g3_i2.p1
F−Acol___TRINITY_DN2427_c0_g2_i2.p1
F−Acol___TRINITY_DN2427_c0_g2_i1.p1
Quercus___Qurub.06G200300.1
Kalanchoe___Kalax.0652s0007.1.p
Kalanchoe___Kalax.0484s0028.1.p
Vaccinium___Vadar_g30993.t1
Hydrangea___Hyque.02G112900.1.p
Manihot___Manes.15G010200.1.p
Cucumis___Cucsa.069260.1|PACid_16955422
Cucumis___Cucsa.069250.1|PACid_16955421
Citrus___orange1.1g025802m|PACid_18109612
Olea___Oeu031986.1
Olea___Oeu016256.1
Mimulus___Migut.N00071.1.p
Lactuca___Lsat_1_v5_gn_6_78141.1
Helianthus___HanXRQChr13g0413991
Beta___EL10Ac2g04820.1
Arabidopsis−AT3G06790.1|PACid_19663852
Eucalyptus___Eucgr.B01560.1.p
Musa___GSMUA_Achr8P22480_001
Musa___GSMUA_Achr11P15790_001
Solanum___Solyc01g066060.4.1
Amborella___AmTr_v1.0_scaffold00013.128

# OG0001238

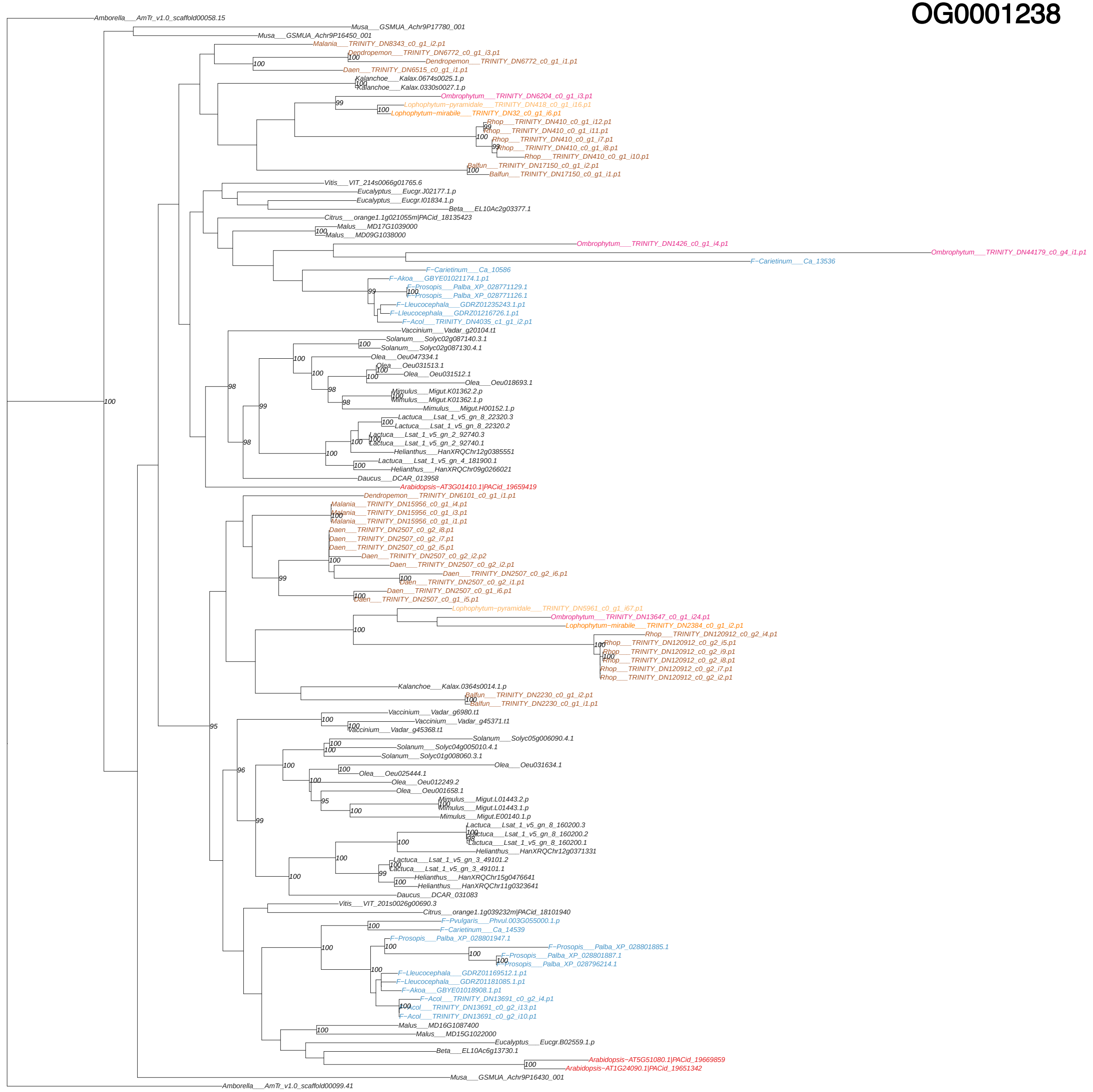
Amborella___AmTr_v1.0_scaffold00058.15
Musa___GSMUA_Achr9P17780_001
Musa___GSMUA_Achr9P16450_001
Malania___TRINITY_DN8343_c0_g1_i2.p1
Dendropemon___TRINITY_DN6772_c0_g1_i3.p1
Dendropemon___TRINITY_DN6772_c0_g1_i1.p1
Daen___TRINITY_DN6515_c0_g1_i1.p1
Kalanchoe___Kalax.0674s0025.1.p
Kalanchoe___Kalax.0330s0027.1.p
Ombrophytum___TRINITY_DN6204_c0_g1_i3.p1
Lophophytum−pyramidale___TRINITY_DN418_c0_g1_i16.p1
Lophophytum−mirabile___TRINITY_DN32_c0_g1_i6.p1
Rhop___TRINITY_DN410_c0_g1_i12.p1
Rhop___TRINITY_DN410_c0_g1_i11.p1
Rhop___TRINITY_DN410_c0_g1_i7.p1
Rhop___TRINITY_DN410_c0_g1_i8.p1
Rhop___TRINITY_DN410_c0_g1_i10.p1
Balfun___TRINITY_DN17150_c0_g1_i2.p1
Balfun___TRINITY_DN17150_c0_g1_i1.p1
Vitis___VIT_214s0066g01765.6
Eucalyptus___Eucgr.J02177.1.p
Eucalyptus___Eucgr.I01834.1.p
Beta___EL10Ac2g03377.1
Citrus___orange1.1g021055m|PACid_18135423
Malus___MD17G1039000
Malus___MD09G1038000
Ombrophytum___TRINITY_DN1426_c0_g1_i4.p1
Ombrophytum___TRINITY_DN44179_c0_g4_i1.p1
F−Carietinum___Ca_13536
F−Carietinum___Ca_10586
F−Akoa___GBYE01021174.1.p1
F−Prosopis___Palba_XP_028771129.1
F−Prosopis___Palba_XP_028771126.1
F−Lleucocephala___GDRZ01235243.1.p1
F−Lleucocephala___GDRZ01216726.1.p1
F−Acol___TRINITY_DN4035_c1_g1_i2.p1
Vaccinium___Vadar_g20104.t1
Solanum___Solyc02g087140.3.1
Solanum___Solyc02g087130.4.1
Olea___Oeu047334.1
Olea___Oeu031513.1
Olea___Oeu031512.1
Olea___Oeu018693.1
Mimulus___Migut.K01362.2.p
Mimulus___Migut.K01362.1.p
Mimulus___Migut.H00152.1.p
Lactuca___Lsat_1_v5_gn_8_22320.3
Lactuca___Lsat_1_v5_gn_8_22320.2
Lactuca___Lsat_1_v5_gn_2_92740.3
Lactuca___Lsat_1_v5_gn_2_92740.1
Helianthus___HanXRQChr12g0385551
Lactuca___Lsat_1_v5_gn_4_181900.1
Helianthus___HanXRQChr09g0266021
Daucus___DCAR_013958
Arabidopsis−AT3G01410.1|PACid_19659419
Dendropemon___TRINITY_DN6101_c0_g1_i1.p1
Malania___TRINITY_DN15956_c0_g1_i4.p1
Malania___TRINITY_DN15956_c0_g1_i3.p1
Malania___TRINITY_DN15956_c0_g1_i1.p1
Daen___TRINITY_DN2507_c0_g2_i8.p1
Daen___TRINITY_DN2507_c0_g2_i7.p1
Daen___TRINITY_DN2507_c0_g2_i5.p1
Daen___TRINITY_DN2507_c0_g2_i2.p2
Daen___TRINITY_DN2507_c0_g2_i2.p1
Daen___TRINITY_DN2507_c0_g2_i6.p1
Daen___TRINITY_DN2507_c0_g2_i1.p1
Daen___TRINITY_DN2507_c0_g1_i6.p1
Daen___TRINITY_DN2507_c0_g1_i5.p1
Lophophytum−pyramidale___TRINITY_DN5961_c0_g1_i67.p1
Ombrophytum___TRINITY_DN13647_c0_g1_i24.p1
Lophophytum−mirabile___TRINITY_DN2384_c0_g1_i2.p1
Rhop___TRINITY_DN120912_c0_g2_i4.p1
Rhop___TRINITY_DN120912_c0_g2_i5.p1
Rhop___TRINITY_DN120912_c0_g2_i9.p1
Rhop___TRINITY_DN120912_c0_g2_i8.p1
Rhop___TRINITY_DN120912_c0_g2_i7.p1
Rhop___TRINITY_DN120912_c0_g2_i2.p1
Kalanchoe___Kalax.0364s0014.1.p
Balfun___TRINITY_DN2230_c0_g1_i2.p1
Balfun___TRINITY_DN2230_c0_g1_i1.p1
Vaccinium___Vadar_g6980.t1
Vaccinium___Vadar_g45371.t1
Vaccinium___Vadar_g45368.t1
Solanum___Solyc05g006090.4.1
Solanum___Solyc04g005010.4.1
Solanum___Solyc01g008060.3.1
Olea___Oeu031634.1
Olea___Oeu025444.1
Olea___Oeu012249.2
Olea___Oeu001658.1
Mimulus___Migut.L01443.2.p
Mimulus___Migut.L01443.1.p
Mimulus___Migut.E00140.1.p
Lactuca___Lsat_1_v5_gn_8_160200.3
Lactuca___Lsat_1_v5_gn_8_160200.2
Lactuca___Lsat_1_v5_gn_8_160200.1
Helianthus___HanXRQChr12g0371331
Lactuca___Lsat_1_v5_gn_3_49101.2
Lactuca___Lsat_1_v5_gn_3_49101.1
Helianthus___HanXRQChr15g0476641
Helianthus___HanXRQChr11g0323641
Daucus___DCAR_031083
Vitis___VIT_201s0026g00690.3
Citrus___orange1.1g039232m|PACid_18101940
F−Pvulgaris___Phvul.003G055000.1.p
F−Carietinum___Ca_14539
F−Prosopis___Palba_XP_028801947.1
F−Prosopis___Palba_XP_028801885.1
F−Prosopis___Palba_XP_028801887.1
F−Prosopis___Palba_XP_028796214.1
F−Lleucocephala___GDRZ01169512.1.p1
F−Lleucocephala___GDRZ01181085.1.p1
F−Akoa___GBYE01018908.1.p1
F−Acol___TRINITY_DN13691_c0_g2_i4.p1
F−Acol___TRINITY_DN13691_c0_g2_i13.p1
F−Acol___TRINITY_DN13691_c0_g2_i10.p1
Malus___MD16G1087400
Malus___MD15G1022000
Eucalyptus___Eucgr.B02559.1.p
Beta___EL10Ac6g13730.1
Arabidopsis−AT5G51080.1|PACid_19669859
Arabidopsis−AT1G24090.1|PACid_19651342
Musa___GSMUA_Achr9P16430_001
Amborella___AmTr_v1.0_scaffold00099.41

# OG0001321

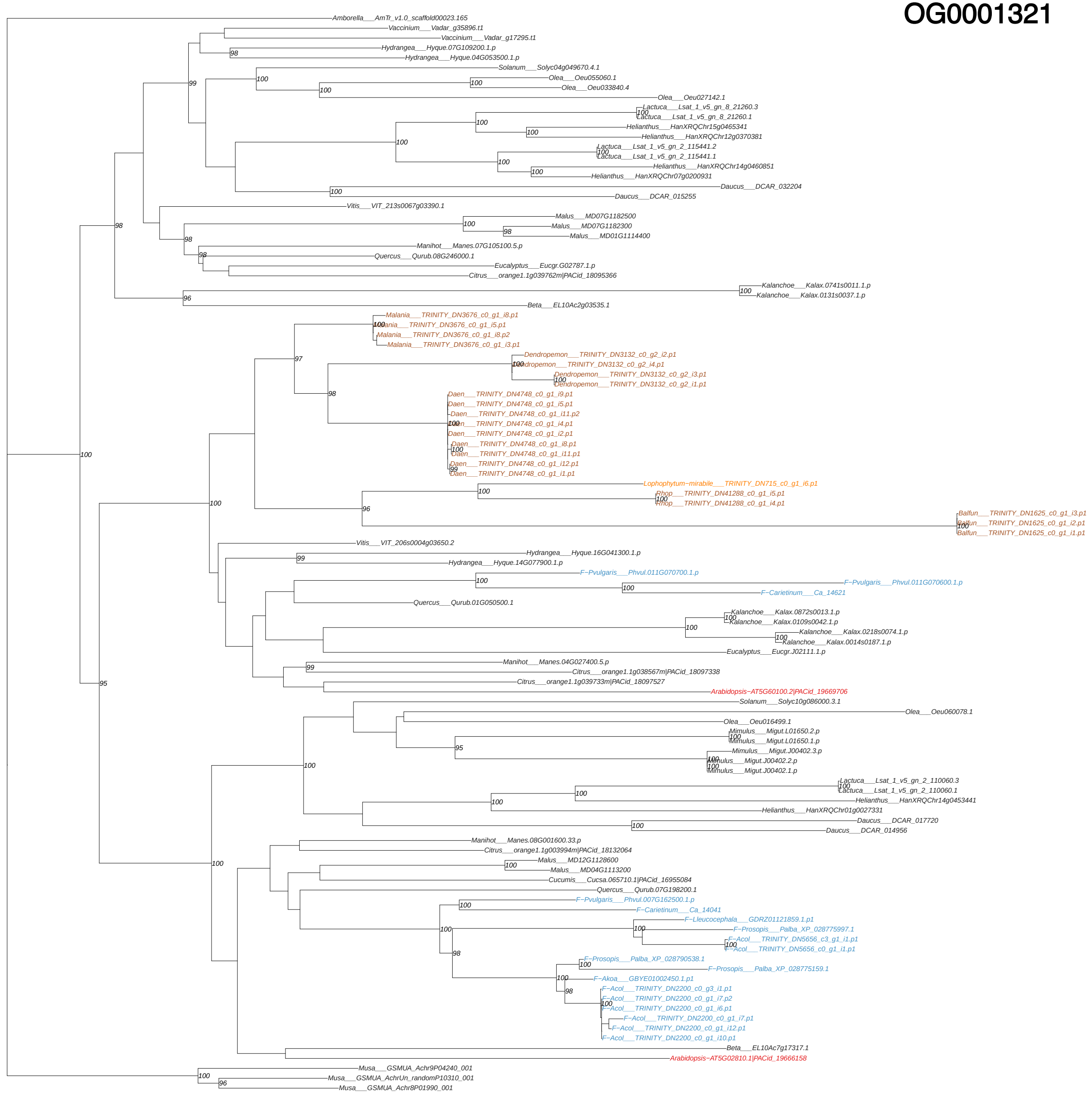

Amborella___AmTr_v1.0_scaffold00023.165
Vaccinium___Vadar_g35896.t1
Vaccinium___Vadar_g17295.t1
Hydrangea___Hyque.07G109200.1.p
Hydrangea___Hyque.04G053500.1.p
Solanum___Solyc04g049670.4.1
Olea___Oeu055060.1
Olea___Oeu033840.4
Olea___Oeu027142.1
Lactuca___Lsat_1_v5_gn_8_21260.3
Lactuca___Lsat_1_v5_gn_8_21260.1
Helianthus___HanXRQChr15g0465341
Helianthus___HanXRQChr12g0370381
Lactuca___Lsat_1_v5_gn_2_115441.2
Lactuca___Lsat_1_v5_gn_2_115441.1
Helianthus___HanXRQChr14g0460851
Helianthus___HanXRQChr07g0200931
Daucus___DCAR_032204
Daucus___DCAR_015255
Vitis___VIT_213s0067g03390.1
Malus___MD07G1182500
Malus___MD07G1182300
Malus___MD01G1114400
Manihot___Manes.07G105100.5.p
Quercus___Qurub.08G246000.1
Eucalyptus___Eucgr.G02787.1.p
Citrus___orange1.1g039762m|PACid_18095366
Kalanchoe___Kalax.0741s0011.1.p
Kalanchoe___Kalax.0131s0037.1.p
Beta___EL10Ac2g03535.1
Malania___TRINITY_DN3676_c0_g1_i8.p1
ania___TRINITY_DN3676_c0_g1_i5.p1
Malania___TRINITY_DN3676_c0_g1_i8.p2
Malania___TRINITY_DN3676_c0_g1_i3.p1
Dendropemon___TRINITY_DN3132_c0_g2_i2.p1
dropemon___TRINITY_DN3132_c0_g2_i4.p1
Dendropemon___TRINITY_DN3132_c0_g2_i3.p1
Dendropemon___TRINITY_DN3132_c0_g2_i1.p1
Daen___TRINITY_DN4748_c0_g1_i9.p1
Daen___TRINITY_DN4748_c0_g1_i5.p1
Daen___TRINITY_DN4748_c0_g1_i11.p2
___TRINITY_DN4748_c0_g1_i4.p1
Daen___TRINITY_DN4748_c0_g1_i2.p1
Daen___TRINITY_DN4748_c0_g1_i8.p1
Daen___TRINITY_DN4748_c0_g1_i11.p1
Daen___TRINITY_DN4748_c0_g1_i12.p1
Daen___TRINITY_DN4748_c0_g1_i1.p1
Lophophytum−mirabile___TRINITY_DN715_c0_g1_i6.p1
Rhop___TRINITY_DN41288_c0_g1_i5.p1
Rhop___TRINITY_DN41288_c0_g1_i4.p1
Balfun___TRINITY_DN1625_c0_g1_i3.p1
___TRINITY_DN1625_c0_g1_i2.p1
Balfun___TRINITY_DN1625_c0_g1_i1.p1
Vitis___VIT_206s0004g03650.2
Hydrangea___Hyque.16G041300.1.p
Hydrangea___Hyque.14G077900.1.p
F−Pvulgaris___Phvul.011G070700.1.p
F−Pvulgaris___Phvul.011G070600.1.p
F−Carietinum___Ca_14621
Quercus___Qurub.01G050500.1
Kalanchoe___Kalax.0872s0013.1.p
Kalanchoe___Kalax.0109s0042.1.p
Kalanchoe___Kalax.0218s0074.1.p
Kalanchoe___Kalax.0014s0187.1.p
Eucalyptus___Eucgr.J02111.1.p
Manihot___Manes.04G027400.5.p
Citrus___orange1.1g038567m|PACid_18097338
Citrus___orange1.1g039733m|PACid_18097527
Arabidopsis−AT5G60100.2|PACid_19669706
Solanum___Solyc10g086000.3.1
Olea___Oeu060078.1
Olea___Oeu016499.1
Mimulus___Migut.L01650.2.p
Mimulus___Migut.L01650.1.p
Mimulus___Migut.J00402.3.p
Mimulus___Migut.J00402.2.p
Mimulus___Migut.J00402.1.p
Lactuca___Lsat_1_v5_gn_2_110060.3
Lactuca___Lsat_1_v5_gn_2_110060.1
Helianthus___HanXRQChr14g0453441
Helianthus___HanXRQChr01g0027331
Daucus___DCAR_017720
Daucus___DCAR_014956
Manihot___Manes.08G001600.33.p
Citrus___orange1.1g003994m|PACid_18132064
Malus___MD12G1128600
Malus___MD04G1113200
Cucumis___Cucsa.065710.1|PACid_16955084
Quercus___Qurub.07G198200.1
F−Pvulgaris___Phvul.007G162500.1.p
F−Carietinum___Ca_14041
F−Lleucocephala___GDRZ01121859.1.p1
F−Prosopis___Palba_XP_028775997.1
F−Acol___TRINITY_DN5656_c3_g1_i1.p1
F−Acol___TRINITY_DN5656_c0_g1_i1.p1
F−Prosopis___Palba_XP_028790538.1
F−Prosopis___Palba_XP_028775159.1
F−Akoa___GBYE01002450.1.p1
F−Acol___TRINITY_DN2200_c0_g3_i1.p1
F−Acol___TRINITY_DN2200_c0_g1_i7.p2
F−Acol___TRINITY_DN2200_c0_g1_i6.p1
F−Acol___TRINITY_DN2200_c0_g1_i7.p1
F−Acol___TRINITY_DN2200_c0_g1_i12.p1
F−Acol___TRINITY_DN2200_c0_g1_i10.p1
Beta___EL10Ac7g17317.1
Arabidopsis−AT5G02810.1|PACid_19666158
Musa___GSMUA_Achr9P04240_001
Musa___GSMUA_AchrUn_randomP10310_001
Musa___GSMUA_Achr8P01990_001

# OG0001336

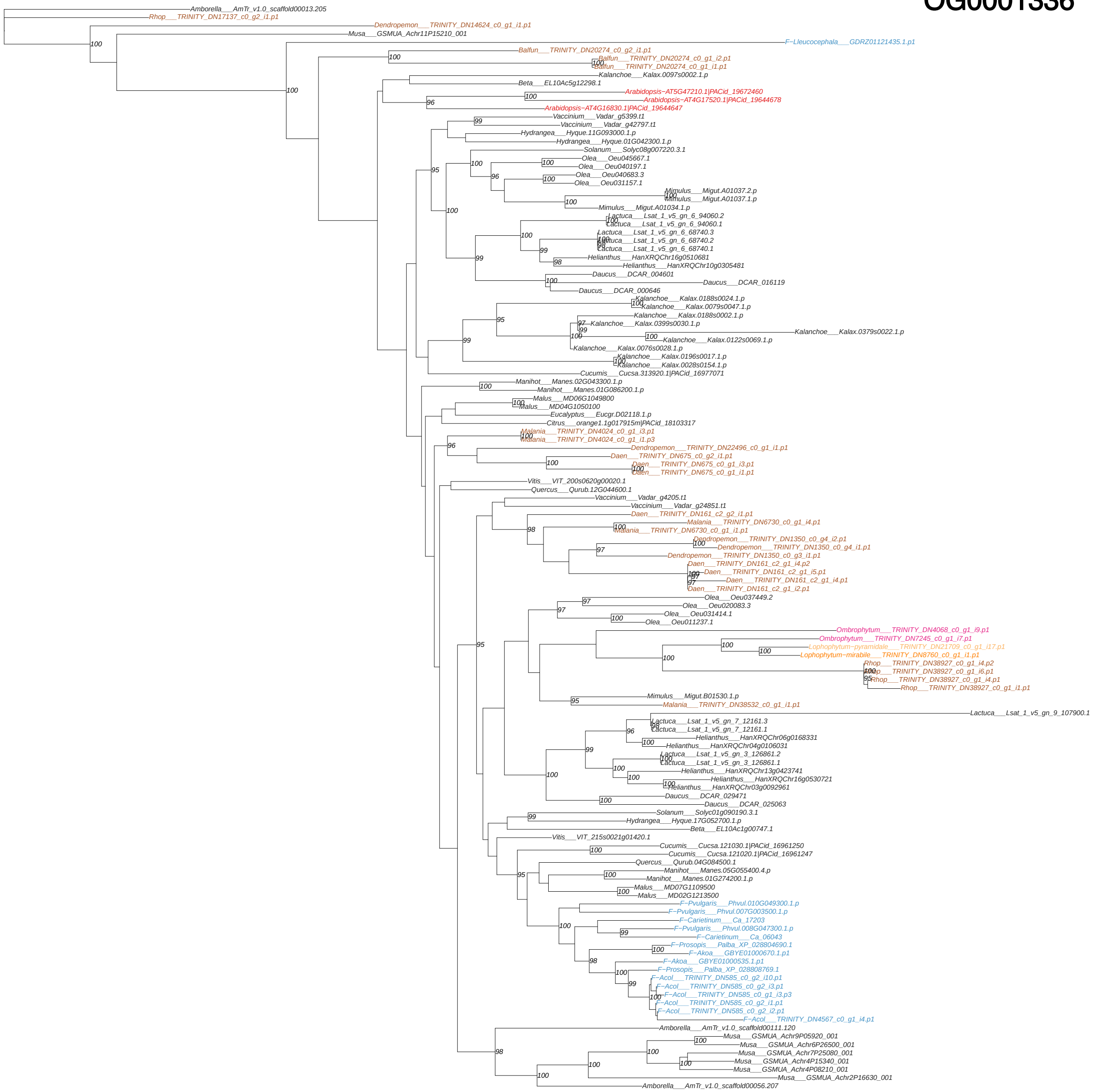

Amborella___AmTr_v1.0_scaffold00013.205
Rhop___TRINITY_DN17137_c0_g2_i1.p1
Dendropemon___TRINITY_DN14624_c0_g1_i1.p1
Musa___GSMUA_Achr11P15210_001
F−Lleucocephala___GDRZ01121435.1.p1
Balfun___TRINITY_DN20274_c0_g2_i1.p1
Balfun___TRINITY_DN20274_c0_g1_i2.p1
Balfun___TRINITY_DN20274_c0_g1_i1.p1
Kalanchoe___Kalax.0097s0002.1.p
Beta___EL10Ac5g12298.1
Arabidopsis−AT5G47210.1|PACid_19672460
Arabidopsis−AT4G17520.1|PACid_19644678
Arabidopsis−AT4G16830.1|PACid_19644647
Vaccinium___Vadar_g5399.t1
Vaccinium___Vadar_g42797.t1
Hydrangea___Hyque.11G093000.1.p
Hydrangea___Hyque.01G042300.1.p
Solanum___Solyc08g007220.3.1
Olea___Oeu045667.1
Olea___Oeu040197.1
Olea___Oeu040683.3
Olea___Oeu031157.1
Mimulus___Migut.A01037.2.p
Mimulus___Migut.A01037.1.p
Mimulus___Migut.A01034.1.p
Lactuca___Lsat_1_v5_gn_6_94060.2
Lactuca___Lsat_1_v5_gn_6_94060.1
Lactuca___Lsat_1_v5_gn_6_68740.3
Lactuca___Lsat_1_v5_gn_6_68740.2
Lactuca___Lsat_1_v5_gn_6_68740.1
Helianthus___HanXRQChr16g0510681
Helianthus___HanXRQChr10g0305481
Daucus___DCAR_004601
Daucus___DCAR_016119
Daucus___DCAR_000646
Kalanchoe___Kalax.0188s0024.1.p
Kalanchoe___Kalax.0079s0047.1.p
Kalanchoe___Kalax.0188s0002.1.p
Kalanchoe___Kalax.0399s0030.1.p
Kalanchoe___Kalax.0379s0022.1.p
Kalanchoe___Kalax.0122s0069.1.p
Kalanchoe___Kalax.0076s0028.1.p
Kalanchoe___Kalax.0196s0017.1.p
Kalanchoe___Kalax.0028s0154.1.p
Cucumis___Cucsa.313920.1|PACid_16977071
Manihot___Manes.02G043300.1.p
Manihot___Manes.01G086200.1.p
Malus___MD06G1049800
Malus___MD04G1050100
Eucalyptus___Eucgr.D02118.1.p
Citrus___orange1.1g017915m|PACid_18103317
Malania___TRINITY_DN4024_c0_g1_i3.p1
Malania___TRINITY_DN4024_c0_g1_i1.p3
Dendropemon___TRINITY_DN22496_c0_g1_i1.p1
Daen___TRINITY_DN675_c0_g2_i1.p1
Daen___TRINITY_DN675_c0_g1_i3.p1
Daen___TRINITY_DN675_c0_g1_i1.p1
Vitis___VIT_200s0620g00020.1
Quercus___Qurub.12G044600.1
Vaccinium___Vadar_g4205.t1
Vaccinium___Vadar_g24851.t1
Daen___TRINITY_DN161_c2_g2_i1.p1
Malania___TRINITY_DN6730_c0_g1_i4.p1
Malania___TRINITY_DN6730_c0_g1_i1.p1
Dendropemon___TRINITY_DN1350_c0_g4_i2.p1
Dendropemon___TRINITY_DN1350_c0_g4_i1.p1
Dendropemon___TRINITY_DN1350_c0_g3_i1.p1
Daen___TRINITY_DN161_c2_g1_i4.p2
Daen___TRINITY_DN161_c2_g1_i5.p1
Daen___TRINITY_DN161_c2_g1_i4.p1
Daen___TRINITY_DN161_c2_g1_i2.p1
Olea___Oeu037449.2
Olea___Oeu020083.3
Olea___Oeu031414.1
Olea___Oeu011237.1
Ombrophytum___TRINITY_DN4068_c0_g1_i9.p1
Ombrophytum___TRINITY_DN7245_c0_g1_i7.p1
Lophophytum−pyramidale___TRINITY_DN21709_c0_g1_i17.p1
Lophophytum−mirabile___TRINITY_DN8760_c0_g1_i1.p1
Rhop___TRINITY_DN38927_c0_g1_i4.p2
Rhop___TRINITY_DN38927_c0_g1_i6.p1
Rhop___TRINITY_DN38927_c0_g1_i4.p1
Rhop___TRINITY_DN38927_c0_g1_i1.p1
Mimulus___Migut.B01530.1.p
Malania___TRINITY_DN38532_c0_g1_i1.p1
Lactuca___Lsat_1_v5_gn_9_107900.1
Lactuca___Lsat_1_v5_gn_7_12161.3
Lactuca___Lsat_1_v5_gn_7_12161.1
Helianthus___HanXRQChr06g0168331
Helianthus___HanXRQChr04g0106031
Lactuca___Lsat_1_v5_gn_3_126861.2
Lactuca___Lsat_1_v5_gn_3_126861.1
Helianthus___HanXRQChr13g0423741
Helianthus___HanXRQChr16g0530721
Helianthus___HanXRQChr03g0092961
Daucus___DCAR_029471
Daucus___DCAR_025063
Solanum___Solyc01g090190.3.1
Hydrangea___Hyque.17G052700.1.p
Beta___EL10Ac1g00747.1
Vitis___VIT_215s0021g01420.1
Cucumis___Cucsa.121030.1|PACid_16961250
Cucumis___Cucsa.121020.1|PACid_16961247
Quercus___Qurub.04G084500.1
Manihot___Manes.05G055400.4.p
Manihot___Manes.01G274200.1.p
Malus___MD07G1109500
Malus___MD02G1213500
F−Pvulgaris___Phvul.010G049300.1.p
F−Pvulgaris___Phvul.007G003500.1.p
F−Carietinum___Ca_17203
F−Pvulgaris___Phvul.008G047300.1.p
F−Carietinum___Ca_06043
F−Prosopis___Palba_XP_028804690.1
F−Akoa___GBYE01000670.1.p1
F−Akoa___GBYE01000535.1.p1
F−Prosopis___Palba_XP_028808769.1
F−Acol___TRINITY_DN585_c0_g2_i10.p1
F−Acol___TRINITY_DN585_c0_g2_i3.p1
F−Acol___TRINITY_DN585_c0_g1_i3.p3
F−Acol___TRINITY_DN585_c0_g2_i1.p1
F−Acol___TRINITY_DN585_c0_g2_i2.p1
F−Acol___TRINITY_DN4567_c0_g1_i4.p1
Amborella___AmTr_v1.0_scaffold00111.120
Musa___GSMUA_Achr9P05920_001
Musa___GSMUA_Achr6P26500_001
Musa___GSMUA_Achr7P25080_001
Musa___GSMUA_Achr4P15340_001
Musa___GSMUA_Achr4P08210_001
Musa___GSMUA_Achr2P16630_001
Amborella___AmTr_v1.0_scaffold00056.207

# OG0001354

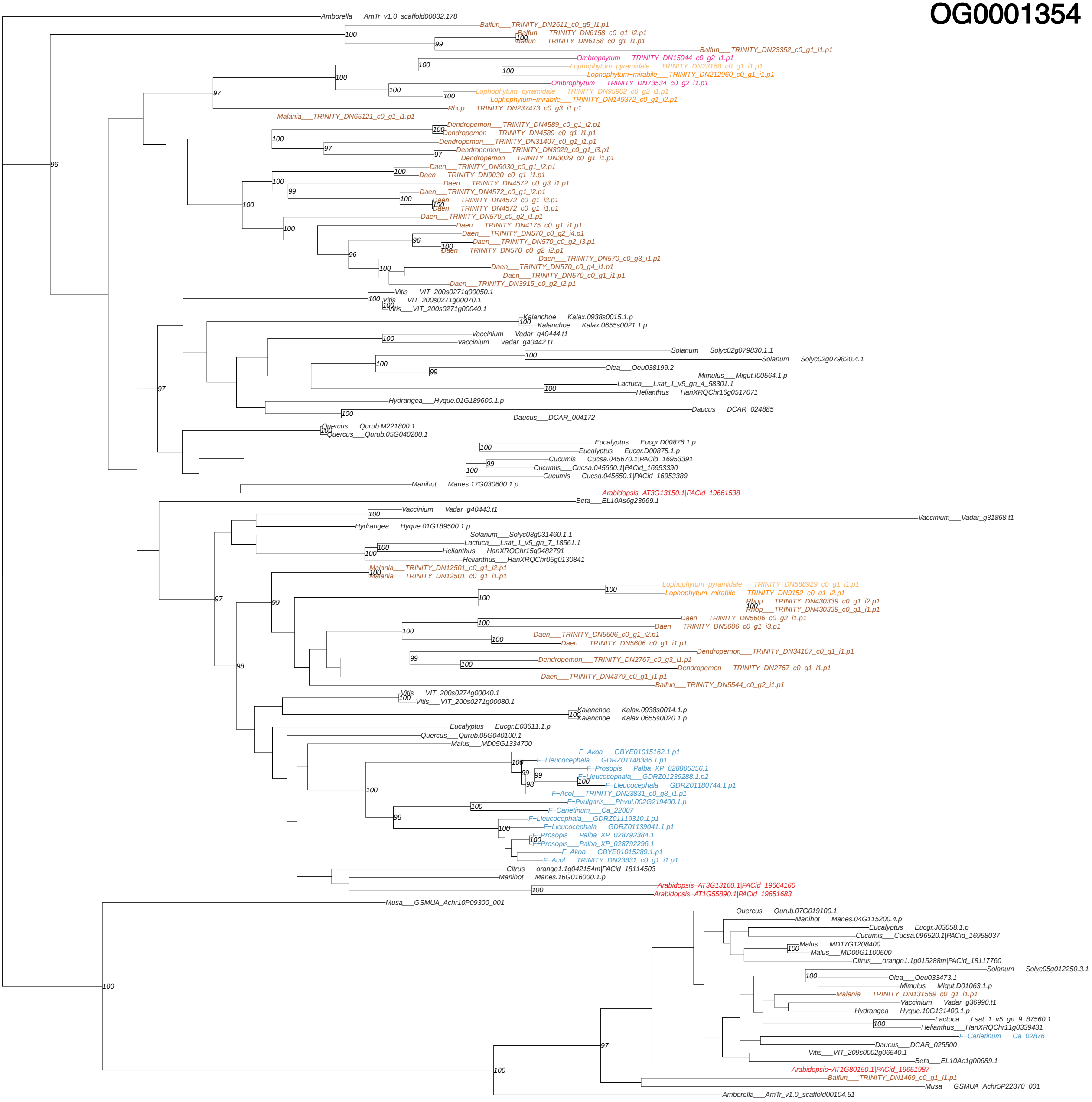

Amborella___AmTr_v1.0_scaffold00032.178
Balfun___TRINITY_DN2611_c0_g5_i1.p1
Balfun___TRINITY_DN6158_c0_g1_i2.p1
Balfun___TRINITY_DN6158_c0_g1_i1.p1
Balfun___TRINITY_DN23352_c0_g1_i1.p1
Ombrophytum___TRINITY_DN15044_c0_g2_i1.p1
Lophophytum−pyramidale___TRINITY_DN23168_c0_g1_i1.p1
Lophophytum−mirabile___TRINITY_DN212960_c0_g1_i1.p1
Ombrophytum___TRINITY_DN73534_c0_g2_i1.p1
Lophophytum−pyramidale___TRINITY_DN95902_c0_g2_i1.p1
Lophophytum−mirabile___TRINITY_DN149372_c0_g1_i2.p1
Rhop___TRINITY_DN237473_c0_g3_i1.p1
Malania___TRINITY_DN65121_c0_g1_i1.p1
Dendropemon___TRINITY_DN4589_c0_g1_i2.p1
Dendropemon___TRINITY_DN4589_c0_g1_i1.p1
Dendropemon___TRINITY_DN31407_c0_g1_i1.p1
Dendropemon___TRINITY_DN3029_c0_g1_i3.p1
Dendropemon___TRINITY_DN3029_c0_g1_i1.p1
Daen___TRINITY_DN9030_c0_g1_i2.p1
Daen___TRINITY_DN9030_c0_g1_i1.p1
Daen___TRINITY_DN4572_c0_g3_i1.p1
Daen___TRINITY_DN4572_c0_g1_i2.p1
Daen___TRINITY_DN4572_c0_g1_i3.p1
Daen___TRINITY_DN4572_c0_g1_i1.p1
Daen___TRINITY_DN570_c0_g2_i1.p1
Daen___TRINITY_DN4175_c0_g1_i1.p1
Daen___TRINITY_DN570_c0_g2_i4.p1
Daen___TRINITY_DN570_c0_g2_i3.p1
Daen___TRINITY_DN570_c0_g2_i2.p1
Daen___TRINITY_DN570_c0_g3_i1.p1
Daen___TRINITY_DN570_c0_g4_i1.p1
Daen___TRINITY_DN570_c0_g1_i1.p1
Daen___TRINITY_DN3915_c0_g2_i2.p1
Vitis___VIT_200s0271g00050.1
Vitis___VIT_200s0271g00070.1
Vitis___VIT_200s0271g00040.1
Kalanchoe___Kalax.0938s0015.1.p
Kalanchoe___Kalax.0655s0021.1.p
Vaccinium___Vadar_g40444.t1
Vaccinium___Vadar_g40442.t1
Solanum___Solyc02g079830.1.1
Solanum___Solyc02g079820.4.1
Olea___Oeu038199.2
Mimulus___Migut.I00564.1.p
Lactuca___Lsat_1_v5_gn_4_58301.1
Helianthus___HanXRQChr16g0517071
Hydrangea___Hyque.01G189600.1.p
Daucus___DCAR_024885
Daucus___DCAR_004172
Quercus___Qurub.M221800.1
Quercus___Qurub.05G040200.1
Eucalyptus___Eucgr.D00876.1.p
Eucalyptus___Eucgr.D00875.1.p
Cucumis___Cucsa.045670.1|PACid_16953391
Cucumis___Cucsa.045660.1|PACid_16953390
Cucumis___Cucsa.045650.1|PACid_16953389
Manihot___Manes.17G030600.1.p
Arabidopsis−AT3G13150.1|PACid_19661538
Beta___EL10As6g23669.1
Vaccinium___Vadar_g40443.t1
Vaccinium___Vadar_g31868.t1
Hydrangea___Hyque.01G189500.1.p
Solanum___Solyc03g031460.1.1
Lactuca___Lsat_1_v5_gn_7_18561.1
Helianthus___HanXRQChr15g0482791
Helianthus___HanXRQChr05g0130841
Malania___TRINITY_DN12501_c0_g1_i2.p1
Malania___TRINITY_DN12501_c0_g1_i1.p1
Lophophytum−pyramidale___TRINITY_DN588929_c0_g1_i1.p1
Lophophytum−mirabile___TRINITY_DN9152_c0_g1_i2.p1
Rhop___TRINITY_DN430339_c0_g1_i2.p1
Rhop___TRINITY_DN430339_c0_g1_i1.p1
Daen___TRINITY_DN5606_c0_g2_i1.p1
Daen___TRINITY_DN5606_c0_g1_i3.p1
Daen___TRINITY_DN5606_c0_g1_i2.p1
Daen___TRINITY_DN5606_c0_g1_i1.p1
Dendropemon___TRINITY_DN34107_c0_g1_i1.p1
Dendropemon___TRINITY_DN2767_c0_g3_i1.p1
Dendropemon___TRINITY_DN2767_c0_g1_i1.p1
Daen___TRINITY_DN4379_c0_g1_i1.p1
Balfun___TRINITY_DN5544_c0_g2_i1.p1
Vitis___VIT_200s0274g00040.1
Vitis___VIT_200s0271g00080.1
Kalanchoe___Kalax.0938s0014.1.p
Kalanchoe___Kalax.0655s0020.1.p
Eucalyptus___Eucgr.E03611.1.p
Quercus___Qurub.05G040100.1
Malus___MD05G1334700
F−Akoa___GBYE01015162.1.p1
F−Lleucocephala___GDRZ01148386.1.p1
F−Prosopis___Palba_XP_028805356.1
F−Lleucocephala___GDRZ01239288.1.p2
F−Lleucocephala___GDRZ01180744.1.p1
F−Acol___TRINITY_DN23831_c0_g3_i1.p1
F−Pvulgaris___Phvul.002G219400.1.p
F−Carietinum___Ca_22007
F−Lleucocephala___GDRZ01119310.1.p1
F−Lleucocephala___GDRZ01139041.1.p1
F−Prosopis___Palba_XP_028792384.1
F−Prosopis___Palba_XP_028792296.1
F−Akoa___GBYE01015289.1.p1
F−Acol___TRINITY_DN23831_c0_g1_i1.p1
Citrus___orange1.1g042154m|PACid_18114503
Manihot___Manes.16G016000.1.p
Arabidopsis−AT3G13160.1|PACid_19664160
Arabidopsis−AT1G55890.1|PACid_19651683
Musa___GSMUA_Achr10P09300_001
Quercus___Qurub.07G019100.1
Manihot___Manes.04G115200.4.p
Eucalyptus___Eucgr.J03058.1.p
Cucumis___Cucsa.096520.1|PACid_16958037
Malus___MD17G1208400
Malus___MD00G1100500
Citrus___orange1.1g015288m|PACid_18117760
Solanum___Solyc05g012250.3.1
Olea___Oeu033473.1
Mimulus___Migut.D01063.1.p
Malania___TRINITY_DN131569_c0_g1_i1.p1
Vaccinium___Vadar_g36990.t1
Hydrangea___Hyque.10G131400.1.p
Lactuca___Lsat_1_v5_gn_9_87560.1
Helianthus___HanXRQChr11g0339431
F−Carietinum___Ca_02876
Daucus___DCAR_025500
Vitis___VIT_209s0002g06540.1
Beta___EL10Ac1g00689.1
Arabidopsis−AT1G80150.1|PACid_19651987
Balfun___TRINITY_DN1469_c0_g1_i1.p1
Musa___GSMUA_Achr5P22370_001
Amborella___AmTr_v1.0_scaffold00104.51

OG0001384

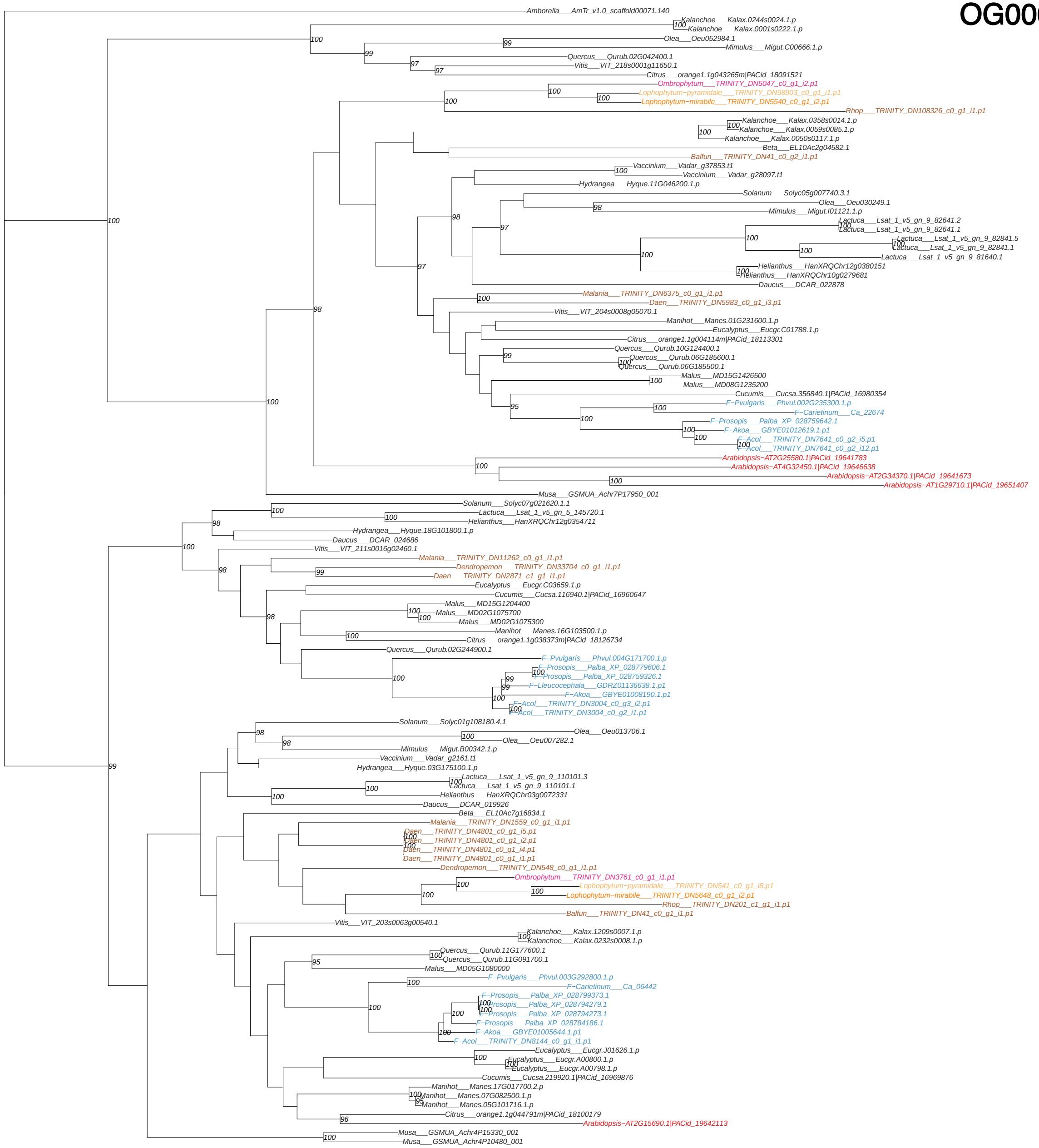
Amborella___AmTr_v1.0_scaffold00071.140
Kalanchoe___Kalax.0244s0024.1.p
Kalanchoe___Kalax.0001s0222.1.p
Olea___Oeu052984.1
Mimulus___Migut.C00666.1.p
Quercus___Qurub.02G042400.1
Vitis___VIT_218s0001g11650.1
Citrus___orange1.1g043265m|PACid_18091521
Ombrophytum___TRINITY_DN5047_c0_g1_i2.p1
Lophophytum−pyramidale___TRINITY_DN98903_c0_g1_i1.p1
Lophophytum−mirabile___TRINITY_DN5540_c0_g1_i2.p1
Rhop___TRINITY_DN108326_c0_g1_i1.p1
Kalanchoe___Kalax.0358s0014.1.p
Kalanchoe___Kalax.0059s0085.1.p
Kalanchoe___Kalax.0050s0117.1.p
Beta___EL10Ac2g04582.1
Balfun___TRINITY_DN41_c0_g2_i1.p1
Vaccinium___Vadar_g37853.t1
Vaccinium___Vadar_g28097.t1
Hydrangea___Hyque.11G046200.1.p
Solanum___Solyc05g007740.3.1
Olea___Oeu030249.1
Mimulus___Migut.I01121.1.p
Lactuca___Lsat_1_v5_gn_9_82641.2
Lactuca___Lsat_1_v5_gn_9_82641.1
Lactuca___Lsat_1_v5_gn_9_82841.5
Lactuca___Lsat_1_v5_gn_9_82841.1
Lactuca___Lsat_1_v5_gn_9_81640.1
Helianthus___HanXRQChr12g0380151
Helianthus___HanXRQChr10g0279681
Daucus___DCAR_022878
Malania___TRINITY_DN6375_c0_g1_i1.p1
Daen___TRINITY_DN5983_c0_g1_i3.p1
Vitis___VIT_204s0008g05070.1
Manihot___Manes.01G231600.1.p
Eucalyptus___Eucgr.C01788.1.p
Citrus___orange1.1g004114m|PACid_18113301
Quercus___Qurub.10G124400.1
Quercus___Qurub.06G185600.1
Quercus___Qurub.06G185500.1
Malus___MD15G1426500
Malus___MD08G1235200
Cucumis___Cucsa.356840.1|PACid_16980354
F−Pvulgaris___Phvul.002G235300.1.p
F−Carietinum___Ca_22674
F−Prosopis___Palba_XP_028759642.1
F−Akoa___GBYE01012619.1.p1
F−Acol___TRINITY_DN7641_c0_g2_i5.p1
F−Acol___TRINITY_DN7641_c0_g2_i12.p1
Arabidopsis−AT2G25580.1|PACid_19641783
Arabidopsis−AT4G32450.1|PACid_19646638
Arabidopsis−AT2G34370.1|PACid_19641673
Arabidopsis−AT1G29710.1|PACid_19651407
Musa___GSMUA_Achr7P17950_001
Solanum___Solyc07g021620.1.1
Lactuca___Lsat_1_v5_gn_5_145720.1
Helianthus___HanXRQChr12g0354711
Hydrangea___Hyque.18G101800.1.p
Daucus___DCAR_024686
Vitis___VIT_211s0016g02460.1
Malania___TRINITY_DN11262_c0_g1_i1.p1
Dendropemon___TRINITY_DN33704_c0_g1_i1.p1
Daen___TRINITY_DN2871_c1_g1_i1.p1
Eucalyptus___Eucgr.C03659.1.p
Cucumis___Cucsa.116940.1|PACid_16960647
Malus___MD15G1204400
Malus___MD02G1075700
Malus___MD02G1075300
Manihot___Manes.16G103500.1.p
Citrus___orange1.1g038373m|PACid_18126734
Quercus___Qurub.02G244900.1
F−Pvulgaris___Phvul.004G171700.1.p
F−Prosopis___Palba_XP_028779606.1
F−Prosopis___Palba_XP_028759326.1
F−Lleucocephala___GDRZ01136638.1.p1
F−Akoa___GBYE01008190.1.p1
F−Acol___TRINITY_DN3004_c0_g3_i2.p1
F−Acol___TRINITY_DN3004_c0_g2_i1.p1
Solanum___Solyc01g108180.4.1
Olea___Oeu013706.1
Olea___Oeu007282.1
Mimulus___Migut.B00342.1.p
Vaccinium___Vadar_g2161.t1
Hydrangea___Hyque.03G175100.1.p
Lactuca___Lsat_1_v5_gn_9_110101.3
Lactuca___Lsat_1_v5_gn_9_110101.1
Helianthus___HanXRQChr03g0072331
Daucus___DCAR_019926
Beta___EL10Ac7g16834.1
Malania___TRINITY_DN1559_c0_g1_i1.p1
Daen___TRINITY_DN4801_c0_g1_i5.p1
Daen___TRINITY_DN4801_c0_g1_i2.p1
Daen___TRINITY_DN4801_c0_g1_i4.p1
Daen___TRINITY_DN4801_c0_g1_i1.p1
Dendropemon___TRINITY_DN548_c0_g1_i1.p1
Ombrophytum___TRINITY_DN3761_c0_g1_i1.p1
Lophophytum−pyramidale___TRINITY_DN541_c0_g1_i8.p1
Lophophytum−mirabile___TRINITY_DN5648_c0_g1_i2.p1
Rhop___TRINITY_DN201_c1_g1_i1.p1
Balfun___TRINITY_DN41_c0_g1_i1.p1
Vitis___VIT_203s0063g00540.1
Kalanchoe___Kalax.1209s0007.1.p
Kalanchoe___Kalax.0232s0008.1.p
Quercus___Qurub.11G177600.1
Quercus___Qurub.11G091700.1
Malus___MD05G1080000
F−Pvulgaris___Phvul.003G292800.1.p
F−Carietinum___Ca_06442
F−Prosopis___Palba_XP_028799373.1
F−Prosopis___Palba_XP_028794279.1
F−Prosopis___Palba_XP_028794273.1
F−Prosopis___Palba_XP_028784186.1
F−Akoa___GBYE01005644.1.p1
F−Acol___TRINITY_DN8144_c0_g1_i1.p1
Eucalyptus___Eucgr.J01626.1.p
Eucalyptus___Eucgr.A00800.1.p
Eucalyptus___Eucgr.A00798.1.p
Cucumis___Cucsa.219920.1|PACid_16969876
Manihot___Manes.17G017700.2.p
Manihot___Manes.07G082500.1.p
Manihot___Manes.05G101716.1.p
Citrus___orange1.1g044791m|PACid_18100179
Arabidopsis−AT2G15690.1|PACid_19642113
Musa___GSMUA_Achr4P15330_001
Musa___GSMUA_Achr4P10480_001

# OG0001977

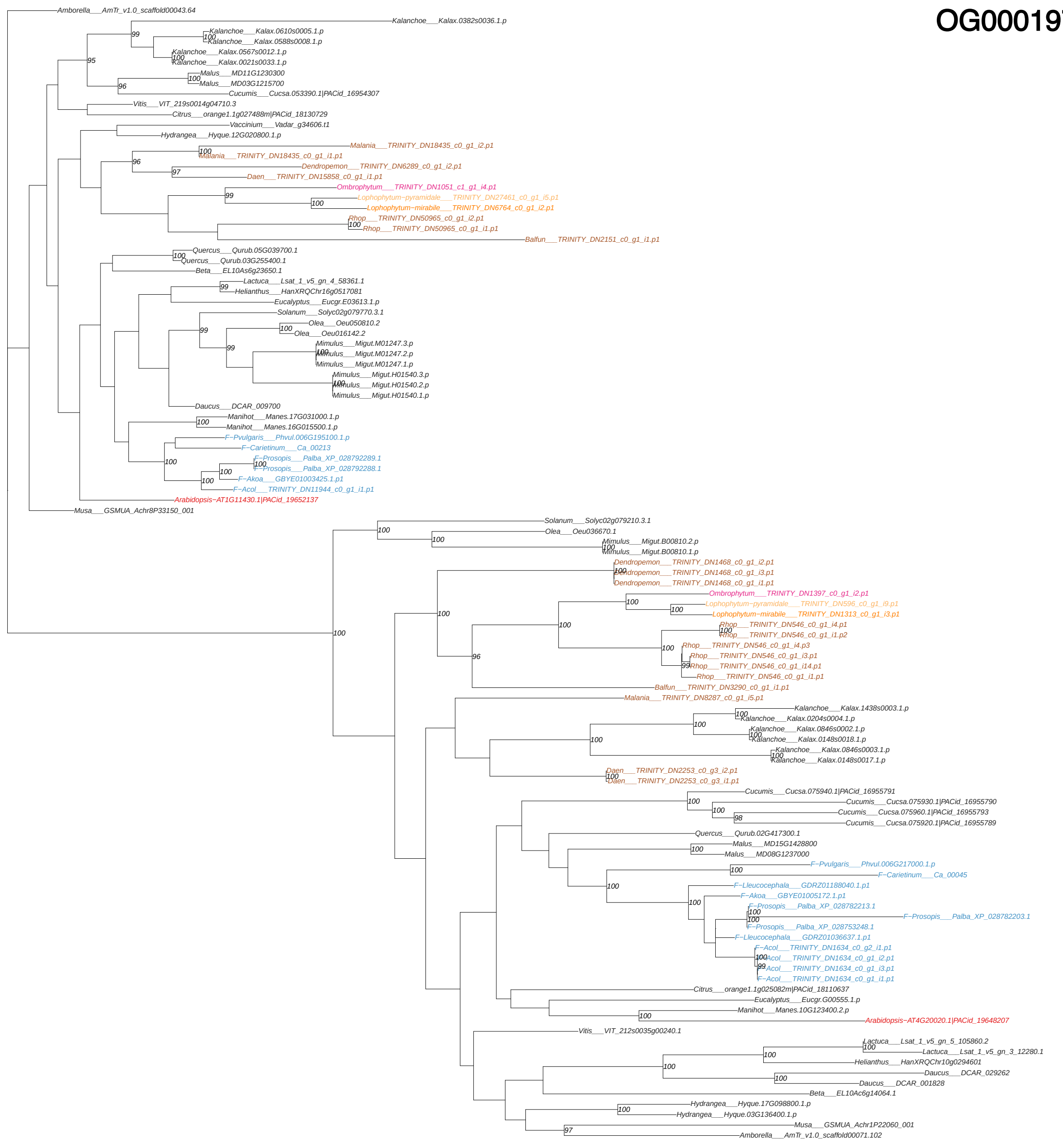
Amborella___AmTr_v1.0_scaffold00043.64
Kalanchoe___Kalax.0382s0036.1.p
99
Kalanchoe___Kalax.0610s0005.1.p
100
Kalanchoe___Kalax.0588s0008.1.p
95
Kalanchoe___Kalax.0567s0012.1.p
100
Kalanchoe___Kalax.0021s0033.1.p
Malus___MD11G1230300
100
96
Malus___MD03G1215700
Cucumis___Cucsa.053390.1|PACid_16954307
Vitis___VIT_219s0014g04710.3
Citrus___orange1.1g027488m|PACid_18130729
Vaccinium___Vadar_g34606.t1
Hydrangea___Hyque.12G020800.1.p
Malania___TRINITY_DN18435_c0_g1_i2.p1
100
Malania___TRINITY_DN18435_c0_g1_i1.p1
96
Dendropemon___TRINITY_DN6289_c0_g1_i2.p1
97
Daen___TRINITY_DN15858_c0_g1_i1.p1
Ombrophytum___TRINITY_DN1051_c1_g1_i4.p1
99
Lophophytum−pyramidale___TRINITY_DN27461_c0_g1_i5.p1
100
Lophophytum−mirabile___TRINITY_DN6764_c0_g1_i2.p1
Rhop___TRINITY_DN50965_c0_g1_i2.p1
100
Rhop___TRINITY_DN50965_c0_g1_i1.p1
Balfun___TRINITY_DN2151_c0_g1_i1.p1
Quercus___Qurub.05G039700.1
100
Quercus___Qurub.03G255400.1
Beta___EL10As6g23650.1
Lactuca___Lsat_1_v5_gn_4_58361.1
99
Helianthus___HanXRQChr16g0517081
Eucalyptus___Eucgr.E03613.1.p
Solanum___Solyc02g079770.3.1
Olea___Oeu050810.2
99
100
Olea___Oeu016142.2
99
Mimulus___Migut.M01247.3.p
100
Mimulus___Migut.M01247.2.p
Mimulus___Migut.M01247.1.p
Mimulus___Migut.H01540.3.p
100
Mimulus___Migut.H01540.2.p
Mimulus___Migut.H01540.1.p
Daucus___DCAR_009700
Manihot___Manes.17G031000.1.p
100
Manihot___Manes.16G015500.1.p
F−Pvulgaris___Phvul.006G195100.1.p
F−Carietinum___Ca_00213
F−Prosopis___Palba_XP_028792289.1
100
100
F−Prosopis___Palba_XP_028792288.1
100
F−Akoa___GBYE01003425.1.p1
100
F−Acol___TRINITY_DN11944_c0_g1_i1.p1
Arabidopsis−AT1G11430.1|PACid_19652137
Musa___GSMUA_Achr8P33150_001
Solanum___Solyc02g079210.3.1
100
Olea___Oeu036670.1
100
Mimulus___Migut.B00810.2.p
100
Mimulus___Migut.B00810.1.p
Dendropemon___TRINITY_DN1468_c0_g1_i2.p1
100
Dendropemon___TRINITY_DN1468_c0_g1_i3.p1
Dendropemon___TRINITY_DN1468_c0_g1_i1.p1
Ombrophytum___TRINITY_DN1397_c0_g1_i2.p1
100
Lophophytum−pyramidale___TRINITY_DN596_c0_g1_i9.p1
100
100
Lophophytum−mirabile___TRINITY_DN1313_c0_g1_i3.p1
100
Rhop___TRINITY_DN546_c0_g1_i4.p1
100
100
Rhop___TRINITY_DN546_c0_g1_i1.p2
Rhop___TRINITY_DN546_c0_g1_i4.p3
96
Rhop___TRINITY_DN546_c0_g1_i3.p1
99
Rhop___TRINITY_DN546_c0_g1_i14.p1
Rhop___TRINITY_DN546_c0_g1_i1.p1
Balfun___TRINITY_DN3290_c0_g1_i1.p1
Malania___TRINITY_DN8287_c0_g1_i5.p1
Kalanchoe___Kalax.1438s0003.1.p
100
Kalanchoe___Kalax.0204s0004.1.p
100
Kalanchoe___Kalax.0846s0002.1.p
100
100
Kalanchoe___Kalax.0148s0018.1.p
Kalanchoe___Kalax.0846s0003.1.p
100
Kalanchoe___Kalax.0148s0017.1.p
Daen___TRINITY_DN2253_c0_g3_i2.p1
100
Daen___TRINITY_DN2253_c0_g3_i1.p1
Cucumis___Cucsa.075940.1|PACid_16955791
100
Cucumis___Cucsa.075930.1|PACid_16955790
100
Cucumis___Cucsa.075960.1|PACid_16955793
98
Cucumis___Cucsa.075920.1|PACid_16955789
Quercus___Qurub.02G417300.1
Malus___MD15G1428800
100
Malus___MD08G1237000
F−Pvulgaris___Phvul.006G217000.1.p
100
F−Carietinum___Ca_00045
F−Lleucocephala___GDRZ01188040.1.p1
100
F−Akoa___GBYE01005172.1.p1
100
F−Prosopis___Palba_XP_028782213.1
100
100
F−Prosopis___Palba_XP_028782203.1
F−Prosopis___Palba_XP_028753248.1
F−Lleucocephala___GDRZ01036637.1.p1
F−Acol___TRINITY_DN1634_c0_g2_i1.p1
100
F−Acol___TRINITY_DN1634_c0_g1_i2.p1
99
F−Acol___TRINITY_DN1634_c0_g1_i3.p1
F−Acol___TRINITY_DN1634_c0_g1_i1.p1
Citrus___orange1.1g025082m|PACid_18110637
Eucalyptus___Eucgr.G00555.1.p
Manihot___Manes.10G123400.2.p
100
Arabidopsis−AT4G20020.1|PACid_19648207
Vitis___VIT_212s0035g00240.1
Lactuca___Lsat_1_v5_gn_5_105860.2
100
100
Lactuca___Lsat_1_v5_gn_3_12280.1
Helianthus___HanXRQChr10g0294601
100
Daucus___DCAR_029262
100
Daucus___DCAR_001828
Beta___EL10Ac6g14064.1
Hydrangea___Hyque.17G098800.1.p
100
Hydrangea___Hyque.03G136400.1.p
Musa___GSMUA_Achr1P22060_001
97
Amborella___AmTr_v1.0_scaffold00071.102

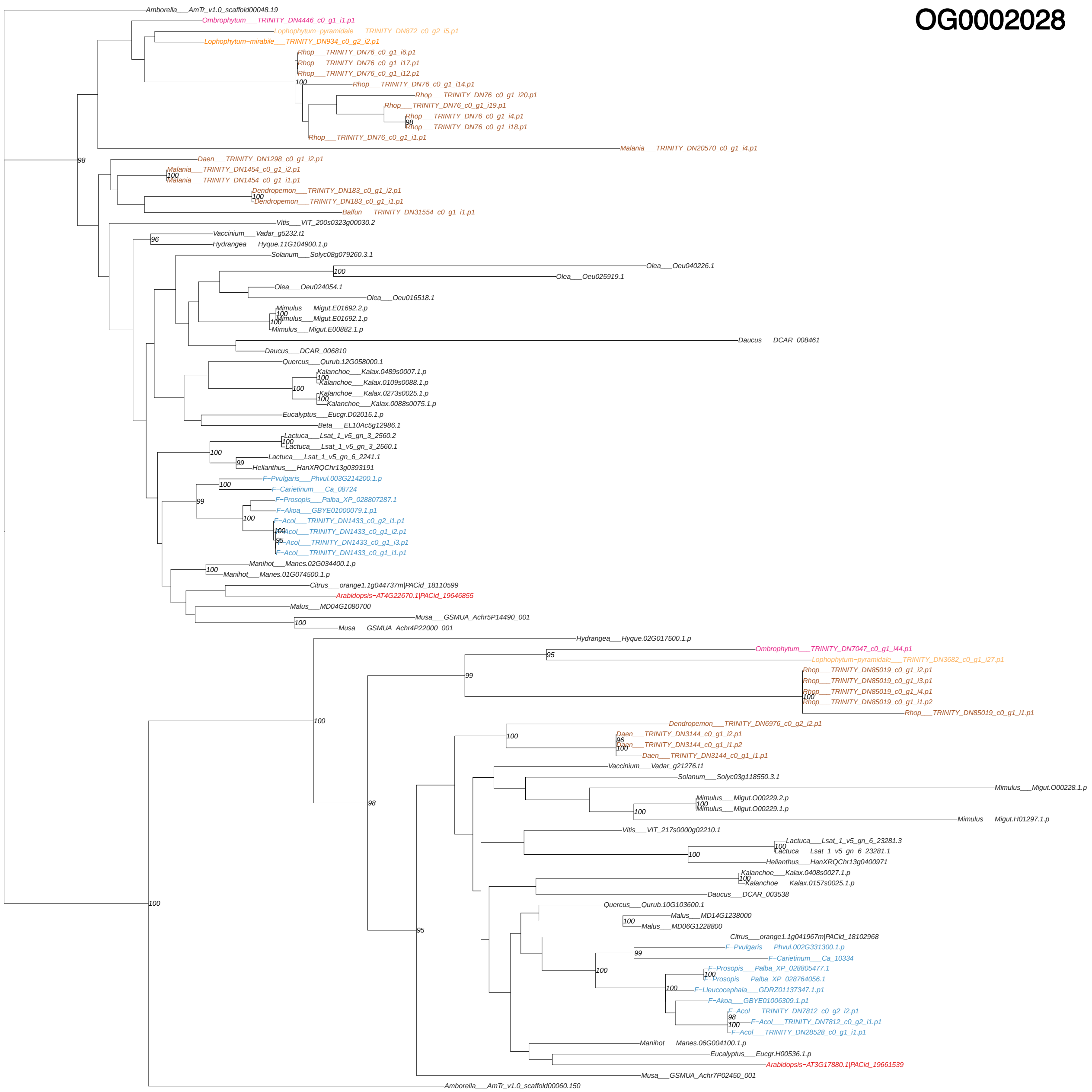

OG0002028
Amborella___AmTr_v1.0_scaffold00048.19
Ombrophytum___TRINITY_DN4446_c0_g1_i1.p1
Lophophytum−pyramidale___TRINITY_DN872_c0_g2_i5.p1
Lophophytum−mirabile___TRINITY_DN934_c0_g2_i2.p1
Rhop___TRINITY_DN76_c0_g1_i6.p1
Rhop___TRINITY_DN76_c0_g1_i17.p1
Rhop___TRINITY_DN76_c0_g1_i12.p1
100
Rhop___TRINITY_DN76_c0_g1_i14.p1
Rhop___TRINITY_DN76_c0_g1_i20.p1
Rhop___TRINITY_DN76_c0_g1_i19.p1
Rhop___TRINITY_DN76_c0_g1_i4.p1
98
Rhop___TRINITY_DN76_c0_g1_i18.p1
Rhop___TRINITY_DN76_c0_g1_i1.p1
Malania___TRINITY_DN20570_c0_g1_i4.p1
Daen___TRINITY_DN1298_c0_g1_i2.p1
98
Malania___TRINITY_DN1454_c0_g1_i2.p1
100
Malania___TRINITY_DN1454_c0_g1_i1.p1
Dendropemon___TRINITY_DN183_c0_g1_i2.p1
100
Dendropemon___TRINITY_DN183_c0_g1_i1.p1
Balfun___TRINITY_DN31554_c0_g1_i1.p1
Vitis___VIT_200s0323g00030.2
Vaccinium___Vadar_g5232.t1
96
Hydrangea___Hyque.11G104900.1.p
Solanum___Solyc08g079260.3.1
Olea___Oeu040226.1
100
Olea___Oeu025919.1
Olea___Oeu024054.1
Olea___Oeu016518.1
Mimulus___Migut.E01692.2.p
100
Mimulus___Migut.E01692.1.p
100
Mimulus___Migut.E00882.1.p
Daucus___DCAR_008461
Daucus___DCAR_006810
Quercus___Qurub.12G058000.1
Kalanchoe___Kalax.0489s0007.1.p
100
Kalanchoe___Kalax.0109s0088.1.p
100
Kalanchoe___Kalax.0273s0025.1.p
100
Kalanchoe___Kalax.0088s0075.1.p
Eucalyptus___Eucgr.D02015.1.p
Beta___EL10Ac5g12986.1
Lactuca___Lsat_1_v5_gn_3_2560.2
100
Lactuca___Lsat_1_v5_gn_3_2560.1
100
Lactuca___Lsat_1_v5_gn_6_2241.1
99
Helianthus___HanXRQChr13g0393191
F−Pvulgaris___Phvul.003G214200.1.p
100
F−Carietinum___Ca_08724
99
F−Prosopis___Palba_XP_028807287.1
F−Akoa___GBYE01000079.1.p1
100
F−Acol___TRINITY_DN1433_c0_g2_i1.p1
100
F−Acol___TRINITY_DN1433_c0_g1_i2.p1
95
F−Acol___TRINITY_DN1433_c0_g1_i3.p1
F−Acol___TRINITY_DN1433_c0_g1_i1.p1
Manihot___Manes.02G034400.1.p
100
Manihot___Manes.01G074500.1.p
Citrus___orange1.1g044737m|PACid_18110599
Arabidopsis−AT4G22670.1|PACid_19646855
Malus___MD04G1080700
Musa___GSMUA_Achr5P14490_001
100
Musa___GSMUA_Achr4P22000_001
Hydrangea___Hyque.02G017500.1.p
Ombrophytum___TRINITY_DN7047_c0_g1_i44.p1
95
Lophophytum−pyramidale___TRINITY_DN3682_c0_g1_i27.p1
99
Rhop___TRINITY_DN85019_c0_g1_i2.p1
Rhop___TRINITY_DN85019_c0_g1_i3.p1
Rhop___TRINITY_DN85019_c0_g1_i4.p1
100
Rhop___TRINITY_DN85019_c0_g1_i1.p2
Rhop___TRINITY_DN85019_c0_g1_i1.p1
100
Dendropemon___TRINITY_DN6976_c0_g2_i2.p1
100
Daen___TRINITY_DN3144_c0_g1_i2.p1
96
Daen___TRINITY_DN3144_c0_g1_i1.p2
100
Daen___TRINITY_DN3144_c0_g1_i1.p1
Vaccinium___Vadar_g21276.t1
Solanum___Solyc03g118550.3.1
Mimulus___Migut.O00228.1.p
98
Mimulus___Migut.O00229.2.p
100
Mimulus___Migut.O00229.1.p
100
Mimulus___Migut.H01297.1.p
Vitis___VIT_217s0000g02210.1
Lactuca___Lsat_1_v5_gn_6_23281.3
100
Lactuca___Lsat_1_v5_gn_6_23281.1
100
Helianthus___HanXRQChr13g0400971
Kalanchoe___Kalax.0408s0027.1.p
100
Kalanchoe___Kalax.0157s0025.1.p
Daucus___DCAR_003538
Quercus___Qurub.10G103600.1
Malus___MD14G1238000
100
Malus___MD06G1228800
Citrus___orange1.1g041967m|PACid_18102968
99
F−Pvulgaris___Phvul.002G331300.1.p
F−Carietinum___Ca_10334
100
F−Prosopis___Palba_XP_028805477.1
100
F−Prosopis___Palba_XP_028764056.1
100
F−Lleucocephala___GDRZ01137347.1.p1
F−Akoa___GBYE01006309.1.p1
F−Acol___TRINITY_DN7812_c0_g2_i2.p1
98
F−Acol___TRINITY_DN7812_c0_g2_i1.p1
100
F−Acol___TRINITY_DN28528_c0_g1_i1.p1
95
Manihot___Manes.06G004100.1.p
Eucalyptus___Eucgr.H00536.1.p
Arabidopsis−AT3G17880.1|PACid_19661539
Musa___GSMUA_Achr7P02450_001
Amborella___AmTr_v1.0_scaffold00060.150

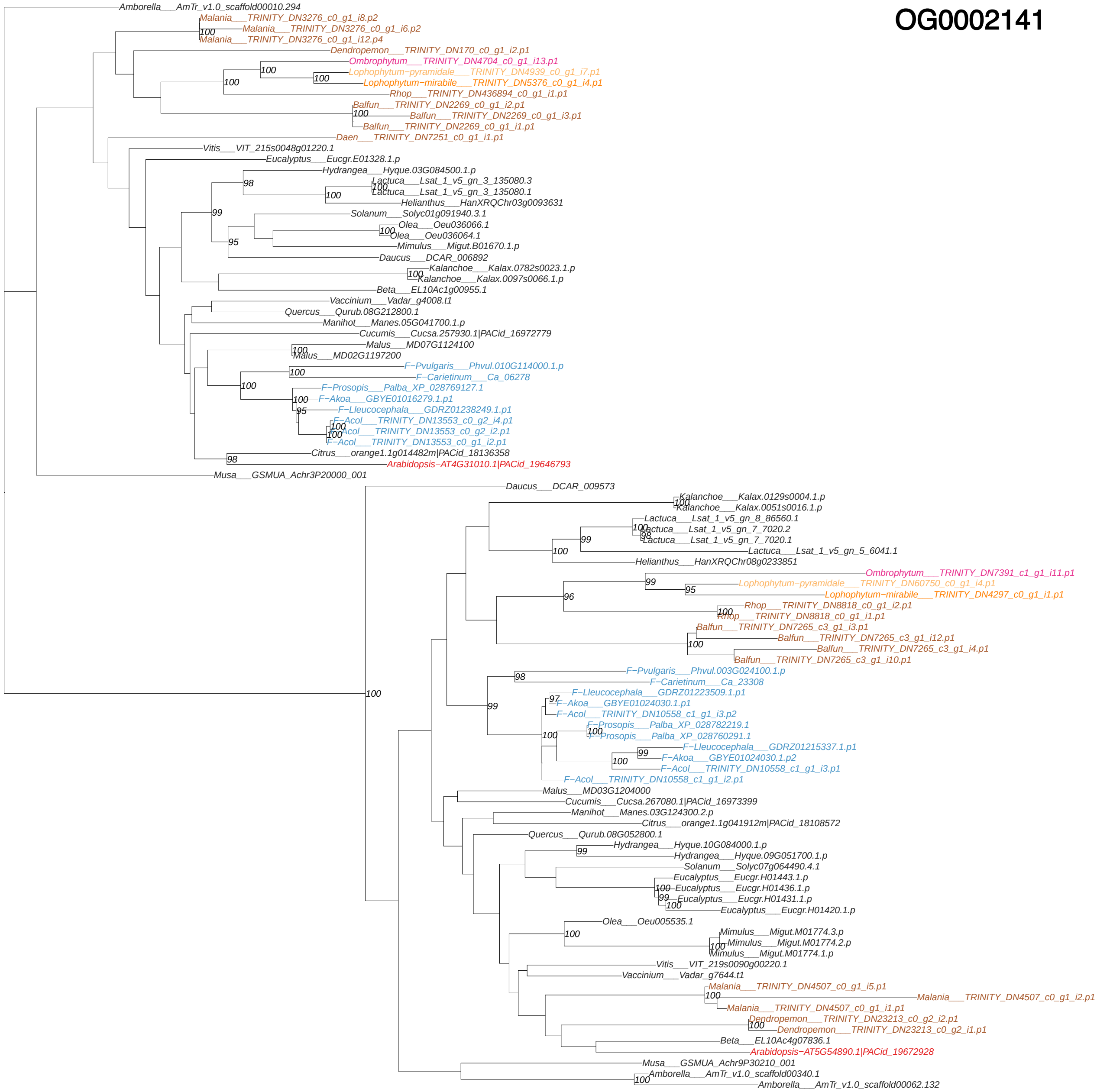
OG0002141
Amborella___AmTr_v1.0_scaffold00010.294
Malania___TRINITY_DN3276_c0_g1_i8.p2
Malania___TRINITY_DN3276_c0_g1_i6.p2
Malania___TRINITY_DN3276_c0_g1_i12.p4
Dendropemon___TRINITY_DN170_c0_g1_i2.p1
Ombrophytum___TRINITY_DN4704_c0_g1_i13.p1
Lophophytum−pyramidale___TRINITY_DN4939_c0_g1_i7.p1
Lophophytum−mirabile___TRINITY_DN5376_c0_g1_i4.p1
Rhop___TRINITY_DN436894_c0_g1_i1.p1
Balfun___TRINITY_DN2269_c0_g1_i2.p1
Balfun___TRINITY_DN2269_c0_g1_i3.p1
Balfun___TRINITY_DN2269_c0_g1_i1.p1
Daen___TRINITY_DN7251_c0_g1_i1.p1
Vitis___VIT_215s0048g01220.1
Eucalyptus___Eucgr.E01328.1.p
Hydrangea___Hyque.03G084500.1.p
Lactuca___Lsat_1_v5_gn_3_135080.3
Lactuca___Lsat_1_v5_gn_3_135080.1
Helianthus___HanXRQChr03g0093631
Solanum___Solyc01g091940.3.1
Olea___Oeu036066.1
Olea___Oeu036064.1
Mimulus___Migut.B01670.1.p
Daucus___DCAR_006892
Kalanchoe___Kalax.0782s0023.1.p
Kalanchoe___Kalax.0097s0066.1.p
Beta___EL10Ac1g00955.1
Vaccinium___Vadar_g4008.t1
Quercus___Qurub.08G212800.1
Manihot___Manes.05G041700.1.p
Cucumis___Cucsa.257930.1|PACid_16972779
Malus___MD07G1124100
Malus___MD02G1197200
F−Pvulgaris___Phvul.010G114000.1.p
F−Carietinum___Ca_06278
F−Prosopis___Palba_XP_028769127.1
F−Akoa___GBYE01016279.1.p1
F−Lleucocephala___GDRZ01238249.1.p1
F−Acol___TRINITY_DN13553_c0_g2_i4.p1
F−Acol___TRINITY_DN13553_c0_g2_i2.p1
F−Acol___TRINITY_DN13553_c0_g1_i2.p1
Citrus___orange1.1g014482m|PACid_18136358
Arabidopsis−AT4G31010.1|PACid_19646793
Musa___GSMUA_Achr3P20000_001
Daucus___DCAR_009573
Kalanchoe___Kalax.0129s0004.1.p
Kalanchoe___Kalax.0051s0016.1.p
Lactuca___Lsat_1_v5_gn_8_86560.1
Lactuca___Lsat_1_v5_gn_7_7020.2
Lactuca___Lsat_1_v5_gn_7_7020.1
Lactuca___Lsat_1_v5_gn_5_6041.1
Helianthus___HanXRQChr08g0233851
Ombrophytum___TRINITY_DN7391_c1_g1_i11.p1
Lophophytum−pyramidale___TRINITY_DN60750_c0_g1_i4.p1
Lophophytum−mirabile___TRINITY_DN4297_c0_g1_i1.p1
Rhop___TRINITY_DN8818_c0_g1_i2.p1
Rhop___TRINITY_DN8818_c0_g1_i1.p1
Balfun___TRINITY_DN7265_c3_g1_i3.p1
Balfun___TRINITY_DN7265_c3_g1_i12.p1
Balfun___TRINITY_DN7265_c3_g1_i4.p1
Balfun___TRINITY_DN7265_c3_g1_i10.p1
F−Pvulgaris___Phvul.003G024100.1.p
F−Carietinum___Ca_23308
F−Lleucocephala___GDRZ01223509.1.p1
F−Akoa___GBYE01024030.1.p1
F−Acol___TRINITY_DN10558_c1_g1_i3.p2
F−Prosopis___Palba_XP_028782219.1
F−Prosopis___Palba_XP_028760291.1
F−Lleucocephala___GDRZ01215337.1.p1
F−Akoa___GBYE01024030.1.p2
F−Acol___TRINITY_DN10558_c1_g1_i3.p1
F−Acol___TRINITY_DN10558_c1_g1_i2.p1
Malus___MD03G1204000
Cucumis___Cucsa.267080.1|PACid_16973399
Manihot___Manes.03G124300.2.p
Citrus___orange1.1g041912m|PACid_18108572
Quercus___Qurub.08G052800.1
Hydrangea___Hyque.10G084000.1.p
Hydrangea___Hyque.09G051700.1.p
Solanum___Solyc07g064490.4.1
Eucalyptus___Eucgr.H01443.1.p
Eucalyptus___Eucgr.H01436.1.p
Eucalyptus___Eucgr.H01431.1.p
Eucalyptus___Eucgr.H01420.1.p
Olea___Oeu005535.1
Mimulus___Migut.M01774.3.p
Mimulus___Migut.M01774.2.p
Mimulus___Migut.M01774.1.p
Vitis___VIT_219s0090g00220.1
Vaccinium___Vadar_g7644.t1
Malania___TRINITY_DN4507_c0_g1_i5.p1
Malania___TRINITY_DN4507_c0_g1_i2.p1
Malania___TRINITY_DN4507_c0_g1_i1.p1
Dendropemon___TRINITY_DN23213_c0_g2_i2.p1
Dendropemon___TRINITY_DN23213_c0_g2_i1.p1
Beta___EL10Ac4g07836.1
Arabidopsis−AT5G54890.1|PACid_19672928
Musa___GSMUA_Achr9P30210_001
Amborella___AmTr_v1.0_scaffold00340.1
Amborella___AmTr_v1.0_scaffold00062.132

# OG0002158

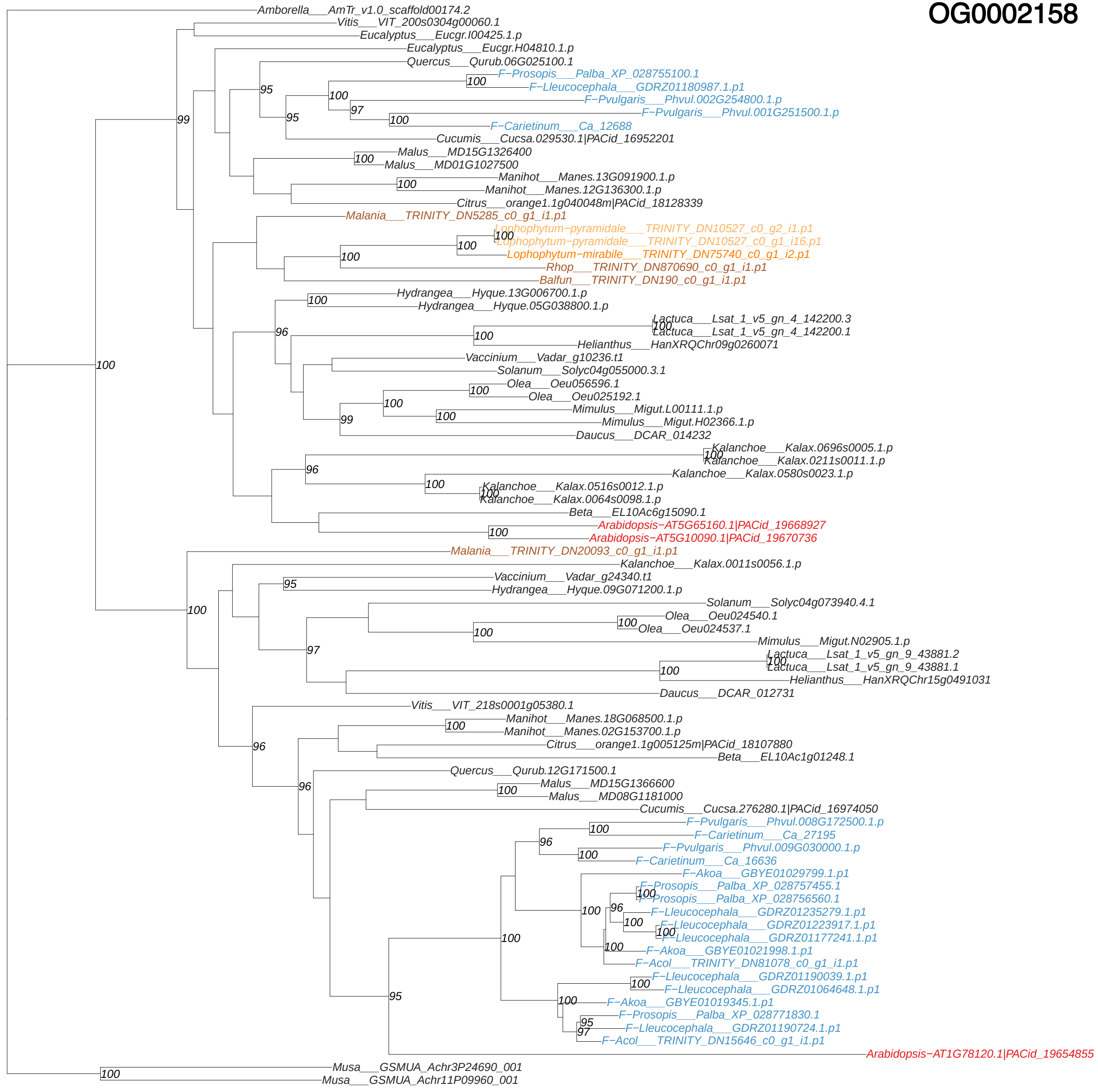

Amborella___AmTr_v1.0_scaffold00174.2
Vitis___VIT_200s0304g00060.1
Eucalyptus___Eucgr.I00425.1.p
Eucalyptus___Eucgr.H04810.1.p
Quercus___Qurub.06G025100.1
F−Prosopis___Palba_XP_028755100.1
F−Lleucocephala___GDRZ01180987.1.p1
F−Pvulgaris___Phvul.002G254800.1.p
F−Pvulgaris___Phvul.001G251500.1.p
F−Carietinum___Ca_12688
Cucumis___Cucsa.029530.1|PACid_16952201
Malus___MD15G1326400
Malus___MD01G1027500
Manihot___Manes.13G091900.1.p
Manihot___Manes.12G136300.1.p
Citrus___orange1.1g040048m|PACid_18128339
Malania___TRINITY_DN5285_c0_g1_i1.p1
Lophophytum−pyramidale___TRINITY_DN10527_c0_g2_i1.p1
Lophophytum−pyramidale___TRINITY_DN10527_c0_g1_i16.p1
Lophophytum−mirabile___TRINITY_DN75740_c0_g1_i2.p1
Rhop___TRINITY_DN870690_c0_g1_i1.p1
Balfun___TRINITY_DN190_c0_g1_i1.p1
Hydrangea___Hyque.13G006700.1.p
Hydrangea___Hyque.05G038800.1.p
Lactuca___Lsat_1_v5_gn_4_142200.3
Lactuca___Lsat_1_v5_gn_4_142200.1
Helianthus___HanXRQChr09g0260071
Vaccinium___Vadar_g10236.t1
Solanum___Solyc04g055000.3.1
Olea___Oeu056596.1
Olea___Oeu025192.1
Mimulus___Migut.L00111.1.p
Mimulus___Migut.H02366.1.p
Daucus___DCAR_014232
Kalanchoe___Kalax.0696s0005.1.p
Kalanchoe___Kalax.0211s0011.1.p
Kalanchoe___Kalax.0580s0023.1.p
Kalanchoe___Kalax.0516s0012.1.p
Kalanchoe___Kalax.0064s0098.1.p
Beta___EL10Ac6g15090.1
Arabidopsis−AT5G65160.1|PACid_19668927
Arabidopsis−AT5G10090.1|PACid_19670736
Malania___TRINITY_DN20093_c0_g1_i1.p1
Kalanchoe___Kalax.0011s0056.1.p
Vaccinium___Vadar_g24340.t1
Hydrangea___Hyque.09G071200.1.p
Solanum___Solyc04g073940.4.1
Olea___Oeu024540.1
Olea___Oeu024537.1
Mimulus___Migut.N02905.1.p
Lactuca___Lsat_1_v5_gn_9_43881.2
Lactuca___Lsat_1_v5_gn_9_43881.1
Helianthus___HanXRQChr15g0491031
Daucus___DCAR_012731
Vitis___VIT_218s0001g05380.1
Manihot___Manes.18G068500.1.p
Manihot___Manes.02G153700.1.p
Citrus___orange1.1g005125m|PACid_18107880
Beta___EL10Ac1g01248.1
Quercus___Qurub.12G171500.1
Malus___MD15G1366600
Malus___MD08G1181000
Cucumis___Cucsa.276280.1|PACid_16974050
F−Pvulgaris___Phvul.008G172500.1.p
F−Carietinum___Ca_27195
F−Pvulgaris___Phvul.009G030000.1.p
F−Carietinum___Ca_16636
F−Akoa___GBYE01029799.1.p1
F−Prosopis___Palba_XP_028757455.1
F−Prosopis___Palba_XP_028756560.1
F−Lleucocephala___GDRZ01235279.1.p1
F−Lleucocephala___GDRZ01223917.1.p1
F−Lleucocephala___GDRZ01177241.1.p1
F−Akoa___GBYE01021998.1.p1
F−Acol___TRINITY_DN81078_c0_g1_i1.p1
F−Lleucocephala___GDRZ01190039.1.p1
F−Lleucocephala___GDRZ01064648.1.p1
F−Akoa___GBYE01019345.1.p1
F−Prosopis___Palba_XP_028771830.1
F−Lleucocephala___GDRZ01190724.1.p1
F−Acol___TRINITY_DN15646_c0_g1_i1.p1
Arabidopsis−AT1G78120.1|PACid_19654855
Musa___GSMUA_Achr3P24690_001
Musa___GSMUA_Achr11P09960_001

# OG0002659

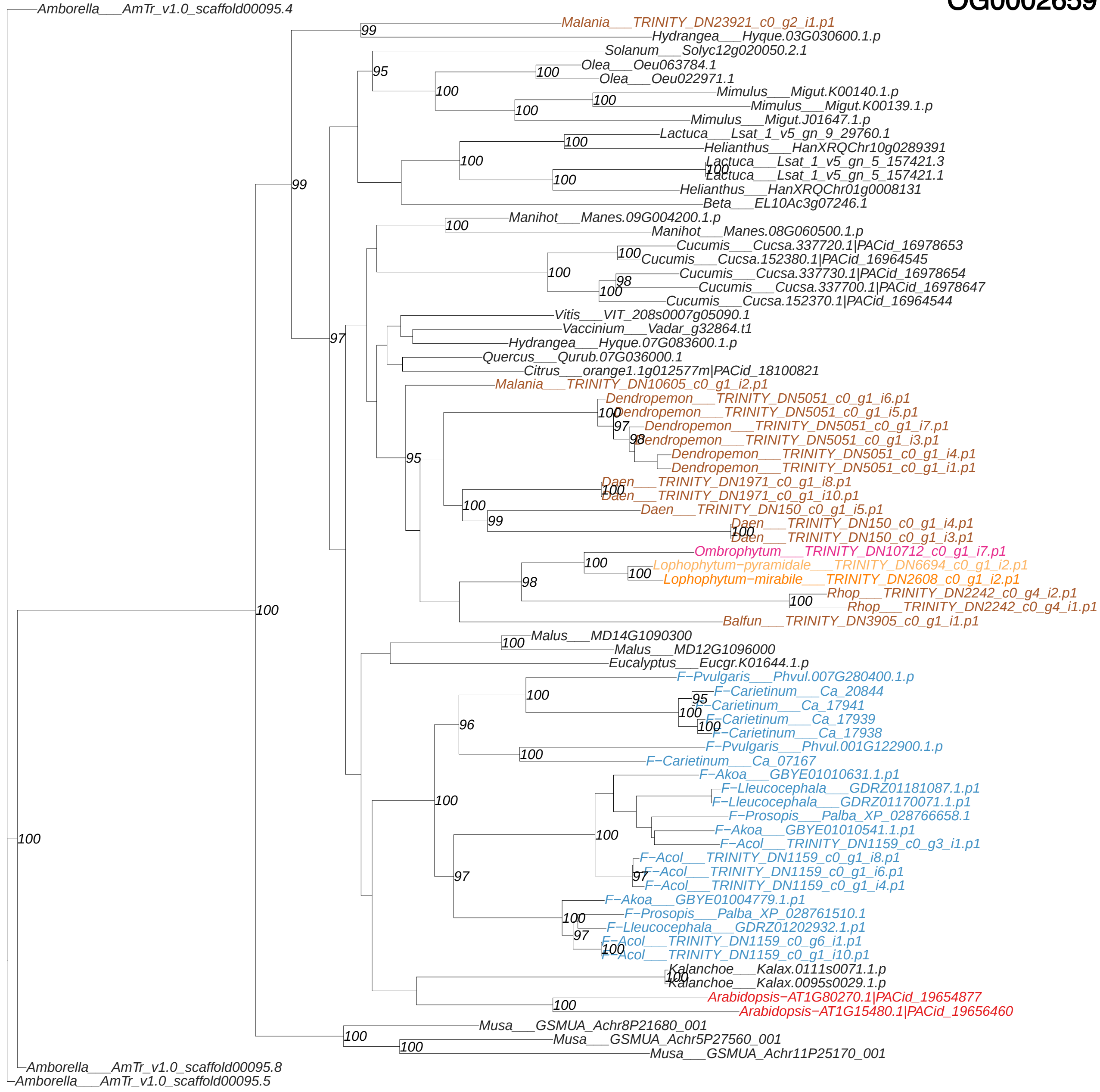

Amborella___AmTr_v1.0_scaffold00095.4
Malania___TRINITY_DN23921_c0_g2_i1.p1
Hydrangea___Hyque.03G030600.1.p
Solanum___Solyc12g020050.2.1
Olea___Oeu063784.1
Olea___Oeu022971.1
Mimulus___Migut.K00140.1.p
Mimulus___Migut.K00139.1.p
Mimulus___Migut.J01647.1.p
Lactuca___Lsat_1_v5_gn_9_29760.1
Helianthus___HanXRQChr10g0289391
Lactuca___Lsat_1_v5_gn_5_157421.3
Lactuca___Lsat_1_v5_gn_5_157421.1
Helianthus___HanXRQChr01g0008131
Beta___EL10Ac3g07246.1
Manihot___Manes.09G004200.1.p
Manihot___Manes.08G060500.1.p
Cucumis___Cucsa.337720.1|PACid_16978653
Cucumis___Cucsa.152380.1|PACid_16964545
Cucumis___Cucsa.337730.1|PACid_16978654
Cucumis___Cucsa.337700.1|PACid_16978647
Cucumis___Cucsa.152370.1|PACid_16964544
Vitis___VIT_208s0007g05090.1
Vaccinium___Vadar_g32864.t1
Hydrangea___Hyque.07G083600.1.p
Quercus___Qurub.07G036000.1
Citrus___orange1.1g012577m|PACid_18100821
Malania___TRINITY_DN10605_c0_g1_i2.p1
Dendropemon___TRINITY_DN5051_c0_g1_i6.p1
Dendropemon___TRINITY_DN5051_c0_g1_i5.p1
Dendropemon___TRINITY_DN5051_c0_g1_i7.p1
Dendropemon___TRINITY_DN5051_c0_g1_i3.p1
Dendropemon___TRINITY_DN5051_c0_g1_i4.p1
Dendropemon___TRINITY_DN5051_c0_g1_i1.p1
Daen___TRINITY_DN1971_c0_g1_i8.p1
Daen___TRINITY_DN1971_c0_g1_i10.p1
Daen___TRINITY_DN150_c0_g1_i5.p1
Daen___TRINITY_DN150_c0_g1_i4.p1
Daen___TRINITY_DN150_c0_g1_i3.p1
Ombrophytum___TRINITY_DN10712_c0_g1_i7.p1
Lophophytum−pyramidale___TRINITY_DN6694_c0_g1_i2.p1
Lophophytum−mirabile___TRINITY_DN2608_c0_g1_i2.p1
Rhop___TRINITY_DN2242_c0_g4_i2.p1
Rhop___TRINITY_DN2242_c0_g4_i1.p1
Balfun___TRINITY_DN3905_c0_g1_i1.p1
Malus___MD14G1090300
Malus___MD12G1096000
Eucalyptus___Eucgr.K01644.1.p
F−Pvulgaris___Phvul.007G280400.1.p
F−Carietinum___Ca_20844
F−Carietinum___Ca_17941
F−Carietinum___Ca_17939
F−Carietinum___Ca_17938
F−Pvulgaris___Phvul.001G122900.1.p
F−Carietinum___Ca_07167
F−Akoa___GBYE01010631.1.p1
F−Lleucocephala___GDRZ01181087.1.p1
F−Lleucocephala___GDRZ01170071.1.p1
F−Prosopis___Palba_XP_028766658.1
F−Akoa___GBYE01010541.1.p1
F−Acol___TRINITY_DN1159_c0_g3_i1.p1
F−Acol___TRINITY_DN1159_c0_g1_i8.p1
F−Acol___TRINITY_DN1159_c0_g1_i6.p1
F−Acol___TRINITY_DN1159_c0_g1_i4.p1
F−Akoa___GBYE01004779.1.p1
F−Prosopis___Palba_XP_028761510.1
F−Lleucocephala___GDRZ01202932.1.p1
F−Acol___TRINITY_DN1159_c0_g6_i1.p1
F−Acol___TRINITY_DN1159_c0_g1_i10.p1
Kalanchoe___Kalax.0111s0071.1.p
Kalanchoe___Kalax.0095s0029.1.p
Arabidopsis−AT1G80270.1|PACid_19654877
Arabidopsis−AT1G15480.1|PACid_19656460
Musa___GSMUA_Achr8P21680_001
Musa___GSMUA_Achr5P27560_001
Musa___GSMUA_Achr11P25170_001
Amborella___AmTr_v1.0_scaffold00095.8
Amborella___AmTr_v1.0_scaffold00095.5

# OG0002993

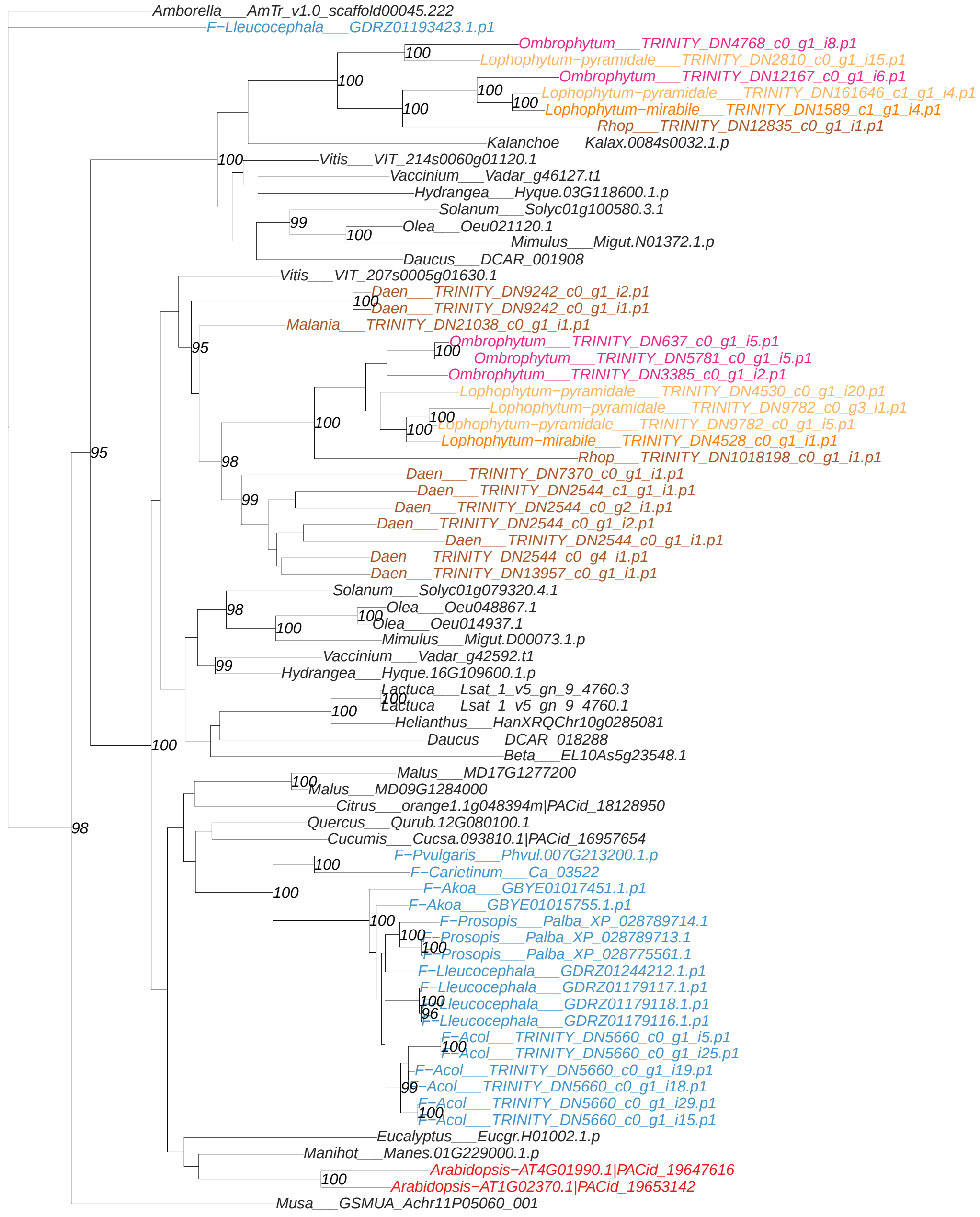
Amborella___AmTr_v1.0_scaffold00045.222
F−Lleucocephala___GDRZ01193423.1.p1
Ombrophytum___TRINITY_DN4768_c0_g1_i8.p1
Lophophytum−pyramidale___TRINITY_DN2810_c0_g1_i15.p1
Ombrophytum___TRINITY_DN12167_c0_g1_i6.p1
Lophophytum−pyramidale___TRINITY_DN161646_c1_g1_i4.p1
Lophophytum−mirabile___TRINITY_DN1589_c1_g1_i4.p1
Rhop___TRINITY_DN12835_c0_g1_i1.p1
Kalanchoe___Kalax.0084s0032.1.p
Vitis___VIT_214s0060g01120.1
Vaccinium___Vadar_g46127.t1
Hydrangea___Hyque.03G118600.1.p
Solanum___Solyc01g100580.3.1
Olea___Oeu021120.1
Mimulus___Migut.N01372.1.p
Daucus___DCAR_001908
Vitis___VIT_207s0005g01630.1
Daen___TRINITY_DN9242_c0_g1_i2.p1
Daen___TRINITY_DN9242_c0_g1_i1.p1
Malania___TRINITY_DN21038_c0_g1_i1.p1
Ombrophytum___TRINITY_DN637_c0_g1_i5.p1
Ombrophytum___TRINITY_DN5781_c0_g1_i5.p1
Ombrophytum___TRINITY_DN3385_c0_g1_i2.p1
Lophophytum−pyramidale___TRINITY_DN4530_c0_g1_i20.p1
Lophophytum−pyramidale___TRINITY_DN9782_c0_g3_i1.p1
Lophophytum−pyramidale___TRINITY_DN9782_c0_g1_i5.p1
Lophophytum−mirabile___TRINITY_DN4528_c0_g1_i1.p1
Rhop___TRINITY_DN1018198_c0_g1_i1.p1
Daen___TRINITY_DN7370_c0_g1_i1.p1
Daen___TRINITY_DN2544_c1_g1_i1.p1
Daen___TRINITY_DN2544_c0_g2_i1.p1
Daen___TRINITY_DN2544_c0_g1_i2.p1
Daen___TRINITY_DN2544_c0_g1_i1.p1
Daen___TRINITY_DN2544_c0_g4_i1.p1
Daen___TRINITY_DN13957_c0_g1_i1.p1
Solanum___Solyc01g079320.4.1
Olea___Oeu048867.1
Olea___Oeu014937.1
Mimulus___Migut.D00073.1.p
Vaccinium___Vadar_g42592.t1
Hydrangea___Hyque.16G109600.1.p
Lactuca___Lsat_1_v5_gn_9_4760.3
Lactuca___Lsat_1_v5_gn_9_4760.1
Helianthus___HanXRQChr10g0285081
Daucus___DCAR_018288
Beta___EL10As5g23548.1
Malus___MD17G1277200
Malus___MD09G1284000
Citrus___orange1.1g048394m|PACid_18128950
Quercus___Qurub.12G080100.1
Cucumis___Cucsa.093810.1|PACid_16957654
F−Pvulgaris___Phvul.007G213200.1.p
F−Carietinum___Ca_03522
F−Akoa___GBYE01017451.1.p1
F−Akoa___GBYE01015755.1.p1
F−Prosopis___Palba_XP_028789714.1
F−Prosopis___Palba_XP_028789713.1
F−Prosopis___Palba_XP_028775561.1
F−Lleucocephala___GDRZ01244212.1.p1
F−Lleucocephala___GDRZ01179117.1.p1
F−Lleucocephala___GDRZ01179118.1.p1
F−Lleucocephala___GDRZ01179116.1.p1
F−Acol___TRINITY_DN5660_c0_g1_i5.p1
F−Acol___TRINITY_DN5660_c0_g1_i25.p1
F−Acol___TRINITY_DN5660_c0_g1_i19.p1
F−Acol___TRINITY_DN5660_c0_g1_i18.p1
F−Acol___TRINITY_DN5660_c0_g1_i29.p1
F−Acol___TRINITY_DN5660_c0_g1_i15.p1
Eucalyptus___Eucgr.H01002.1.p
Manihot___Manes.01G229000.1.p
Arabidopsis−AT4G01990.1|PACid_19647616
Arabidopsis−AT1G02370.1|PACid_19653142
Musa___GSMUA_Achr11P05060_001

OG0003315

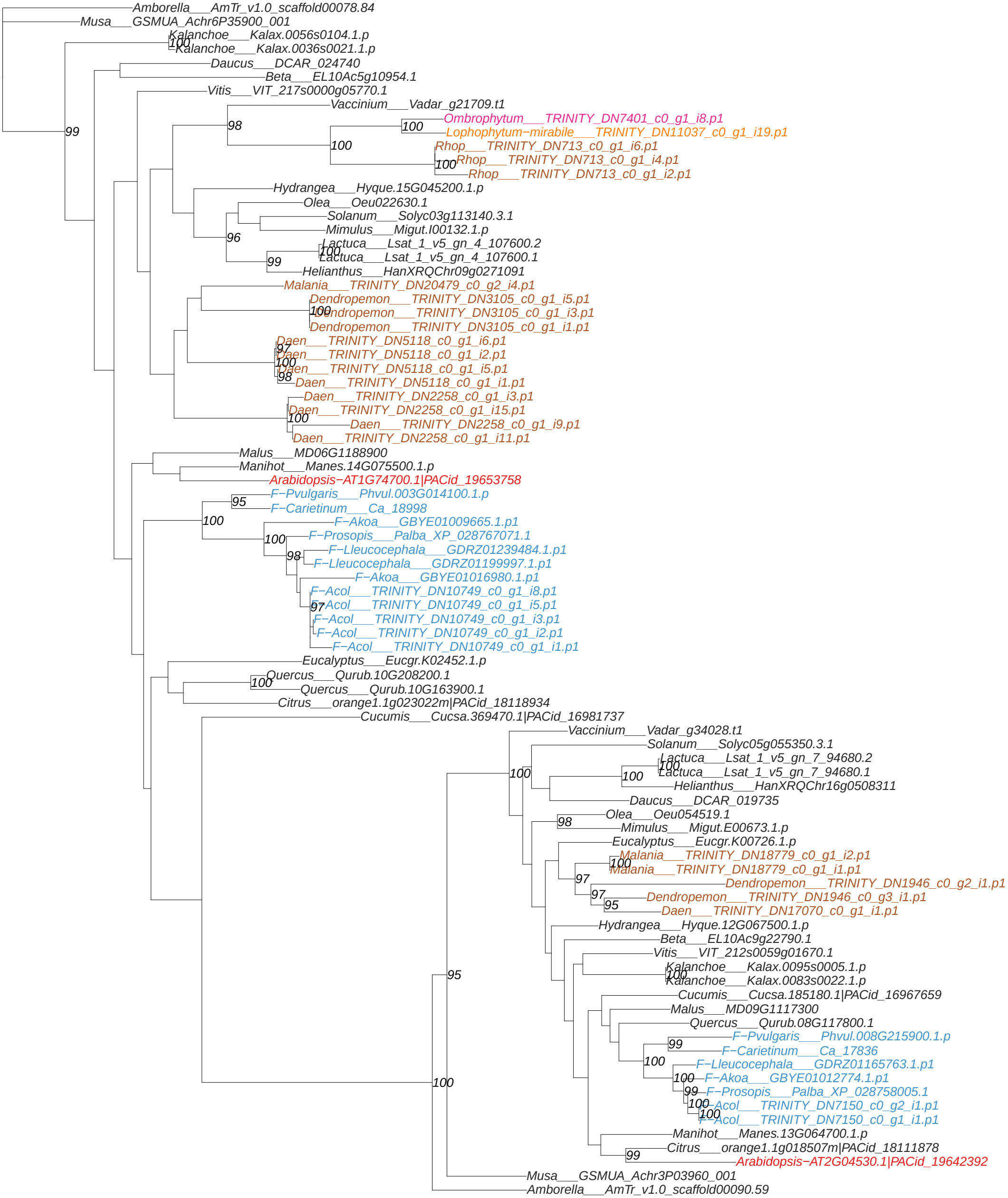

Amborella___AmTr_v1.0_scaffold00078.84
Musa___GSMUA_Achr6P35900_001
Kalanchoe___Kalax.0056s0104.1.p
Kalanchoe___Kalax.0036s0021.1.p
Daucus___DCAR_024740
Beta___EL10Ac5g10954.1
Vitis___VIT_217s0000g05770.1
Vaccinium___Vadar_g21709.t1
Ombrophytum___TRINITY_DN7401_c0_g1_i8.p1
Lophophytum−mirabile___TRINITY_DN11037_c0_g1_i19.p1
Rhop___TRINITY_DN713_c0_g1_i6.p1
Rhop___TRINITY_DN713_c0_g1_i4.p1
Rhop___TRINITY_DN713_c0_g1_i2.p1
Hydrangea___Hyque.15G045200.1.p
Olea___Oeu022630.1
Solanum___Solyc03g113140.3.1
Mimulus___Migut.I00132.1.p
Lactuca___Lsat_1_v5_gn_4_107600.2
Lactuca___Lsat_1_v5_gn_4_107600.1
Helianthus___HanXRQChr09g0271091
Malania___TRINITY_DN20479_c0_g2_i4.p1
Dendropemon___TRINITY_DN3105_c0_g1_i5.p1
Dendropemon___TRINITY_DN3105_c0_g1_i3.p1
Dendropemon___TRINITY_DN3105_c0_g1_i1.p1
Daen___TRINITY_DN5118_c0_g1_i6.p1
Daen___TRINITY_DN5118_c0_g1_i2.p1
Daen___TRINITY_DN5118_c0_g1_i5.p1
Daen___TRINITY_DN5118_c0_g1_i1.p1
Daen___TRINITY_DN2258_c0_g1_i3.p1
Daen___TRINITY_DN2258_c0_g1_i15.p1
Daen___TRINITY_DN2258_c0_g1_i9.p1
Daen___TRINITY_DN2258_c0_g1_i11.p1
Malus___MD06G1188900
Manihot___Manes.14G075500.1.p
Arabidopsis−AT1G74700.1|PACid_19653758
F−Pvulgaris___Phvul.003G014100.1.p
F−Carietinum___Ca_18998
F−Akoa___GBYE01009665.1.p1
F−Prosopis___Palba_XP_028767071.1
F−Lleucocephala___GDRZ01239484.1.p1
F−Lleucocephala___GDRZ01199997.1.p1
F−Akoa___GBYE01016980.1.p1
F−Acol___TRINITY_DN10749_c0_g1_i8.p1
F−Acol___TRINITY_DN10749_c0_g1_i5.p1
F−Acol___TRINITY_DN10749_c0_g1_i3.p1
F−Acol___TRINITY_DN10749_c0_g1_i2.p1
F−Acol___TRINITY_DN10749_c0_g1_i1.p1
Eucalyptus___Eucgr.K02452.1.p
Quercus___Qurub.10G208200.1
Quercus___Qurub.10G163900.1
Citrus___orange1.1g023022m|PACid_18118934
Cucumis___Cucsa.369470.1|PACid_16981737
Vaccinium___Vadar_g34028.t1
Solanum___Solyc05g055350.3.1
Lactuca___Lsat_1_v5_gn_7_94680.2
Lactuca___Lsat_1_v5_gn_7_94680.1
Helianthus___HanXRQChr16g0508311
Daucus___DCAR_019735
Olea___Oeu054519.1
Mimulus___Migut.E00673.1.p
Eucalyptus___Eucgr.K00726.1.p
Malania___TRINITY_DN18779_c0_g1_i2.p1
Malania___TRINITY_DN18779_c0_g1_i1.p1
Dendropemon___TRINITY_DN1946_c0_g2_i1.p1
Dendropemon___TRINITY_DN1946_c0_g3_i1.p1
Daen___TRINITY_DN17070_c0_g1_i1.p1
Hydrangea___Hyque.12G067500.1.p
Beta___EL10Ac9g22790.1
Vitis___VIT_212s0059g01670.1
Kalanchoe___Kalax.0095s0005.1.p
Kalanchoe___Kalax.0083s0022.1.p
Cucumis___Cucsa.185180.1|PACid_16967659
Malus___MD09G1117300
Quercus___Qurub.08G117800.1
F−Pvulgaris___Phvul.008G215900.1.p
F−Carietinum___Ca_17836
F−Lleucocephala___GDRZ01165763.1.p1
F−Akoa___GBYE01012774.1.p1
F−Prosopis___Palba_XP_028758005.1
F−Acol___TRINITY_DN7150_c0_g2_i1.p1
F−Acol___TRINITY_DN7150_c0_g1_i1.p1
Manihot___Manes.13G064700.1.p
Citrus___orange1.1g018507m|PACid_18111878
Arabidopsis−AT2G04530.1|PACid_19642392
Musa___GSMUA_Achr3P03960_001
Amborella___AmTr_v1.0_scaffold00090.59

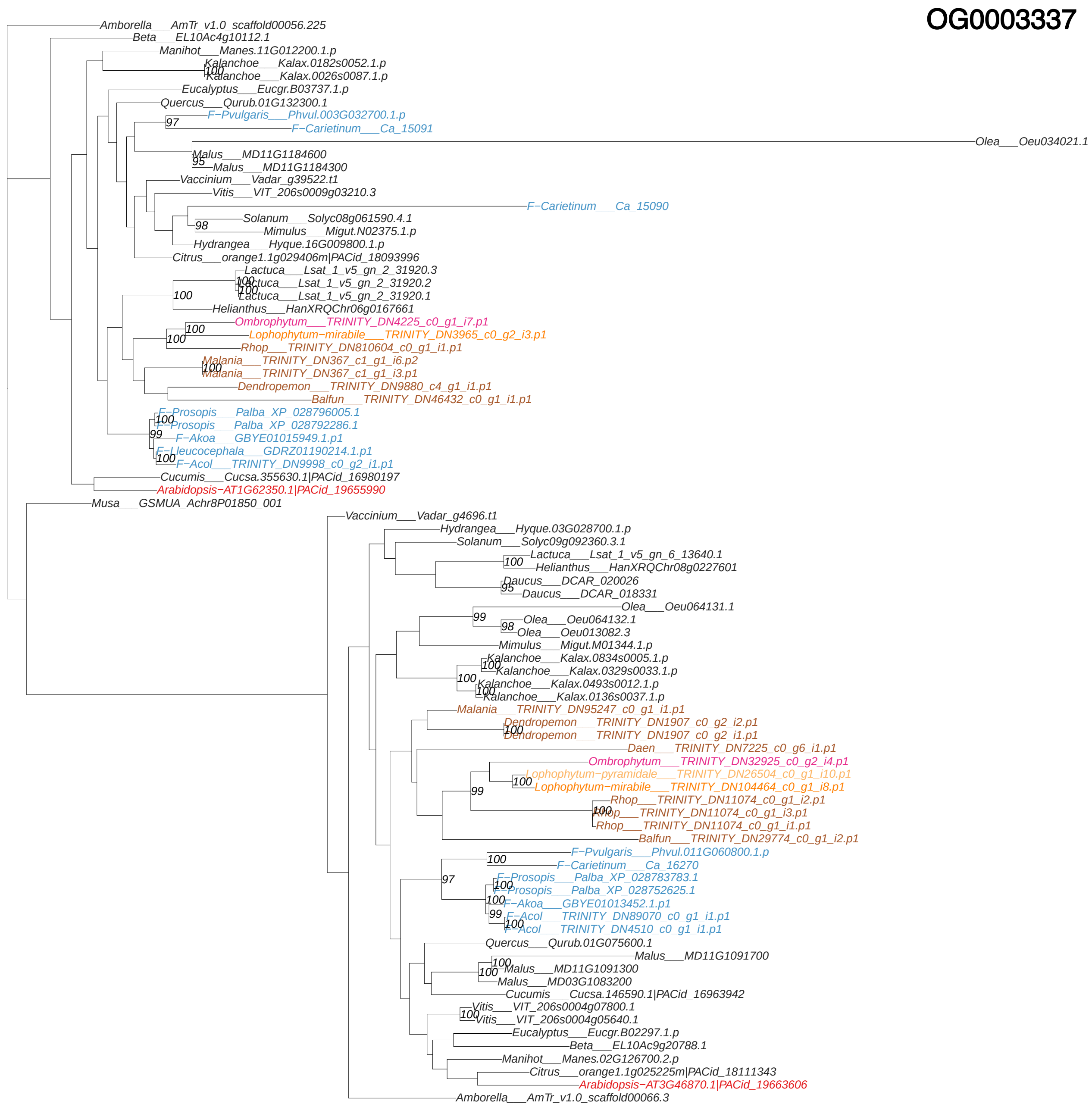
OG0003337
Amborella___AmTr_v1.0_scaffold00056.225
Beta___EL10Ac4g10112.1
Manihot___Manes.11G012200.1.p
Kalanchoe___Kalax.0182s0052.1.p
100
Kalanchoe___Kalax.0026s0087.1.p
Eucalyptus___Eucgr.B03737.1.p
Quercus___Qurub.01G132300.1
F−Pvulgaris___Phvul.003G032700.1.p
97
F−Carietinum___Ca_15091
Olea___Oeu034021.1
Malus___MD11G1184600
95
Malus___MD11G1184300
Vaccinium___Vadar_g39522.t1
Vitis___VIT_206s0009g03210.3
F−Carietinum___Ca_15090
Solanum___Solyc08g061590.4.1
98
Mimulus___Migut.N02375.1.p
Hydrangea___Hyque.16G009800.1.p
Citrus___orange1.1g029406m|PACid_18093996
Lactuca___Lsat_1_v5_gn_2_31920.3
100
Lactuca___Lsat_1_v5_gn_2_31920.2
100
Lactuca___Lsat_1_v5_gn_2_31920.1
100
Helianthus___HanXRQChr06g0167661
Ombrophytum___TRINITY_DN4225_c0_g1_i7.p1
100
Lophophytum−mirabile___TRINITY_DN3965_c0_g2_i3.p1
100
Rhop___TRINITY_DN810604_c0_g1_i1.p1
Malania___TRINITY_DN367_c1_g1_i6.p2
100
Malania___TRINITY_DN367_c1_g1_i3.p1
Dendropemon___TRINITY_DN9880_c4_g1_i1.p1
Balfun___TRINITY_DN46432_c0_g1_i1.p1
F−Prosopis___Palba_XP_028796005.1
100
F−Prosopis___Palba_XP_028792286.1
99
F−Akoa___GBYE01015949.1.p1
F−Lleucocephala___GDRZ01190214.1.p1
100
F−Acol___TRINITY_DN9998_c0_g2_i1.p1
Cucumis___Cucsa.355630.1|PACid_16980197
Arabidopsis−AT1G62350.1|PACid_19655990
Musa___GSMUA_Achr8P01850_001
Vaccinium___Vadar_g4696.t1
Hydrangea___Hyque.03G028700.1.p
Solanum___Solyc09g092360.3.1
Lactuca___Lsat_1_v5_gn_6_13640.1
100
Helianthus___HanXRQChr08g0227601
Daucus___DCAR_020026
95
Daucus___DCAR_018331
Olea___Oeu064131.1
99
Olea___Oeu064132.1
98
Olea___Oeu013082.3
Mimulus___Migut.M01344.1.p
Kalanchoe___Kalax.0834s0005.1.p
100
Kalanchoe___Kalax.0329s0033.1.p
100
Kalanchoe___Kalax.0493s0012.1.p
100
Kalanchoe___Kalax.0136s0037.1.p
Malania___TRINITY_DN95247_c0_g1_i1.p1
Dendropemon___TRINITY_DN1907_c0_g2_i2.p1
100
Dendropemon___TRINITY_DN1907_c0_g2_i1.p1
Daen___TRINITY_DN7225_c0_g6_i1.p1
Ombrophytum___TRINITY_DN32925_c0_g2_i4.p1
Lophophytum−pyramidale___TRINITY_DN26504_c0_g1_i10.p1
100
Lophophytum−mirabile___TRINITY_DN104464_c0_g1_i8.p1
99
Rhop___TRINITY_DN11074_c0_g1_i2.p1
100
Rhop___TRINITY_DN11074_c0_g1_i3.p1
Rhop___TRINITY_DN11074_c0_g1_i1.p1
Balfun___TRINITY_DN29774_c0_g1_i2.p1
F−Pvulgaris___Phvul.011G060800.1.p
100
F−Carietinum___Ca_16270
97
F−Prosopis___Palba_XP_028783783.1
100
F−Prosopis___Palba_XP_028752625.1
100
F−Akoa___GBYE01013452.1.p1
99
F−Acol___TRINITY_DN89070_c0_g1_i1.p1
100
F−Acol___TRINITY_DN4510_c0_g1_i1.p1
Quercus___Qurub.01G075600.1
Malus___MD11G1091700
100
Malus___MD11G1091300
100
Malus___MD03G1083200
Cucumis___Cucsa.146590.1|PACid_16963942
Vitis___VIT_206s0004g07800.1
100
Vitis___VIT_206s0004g05640.1
Eucalyptus___Eucgr.B02297.1.p
Beta___EL10Ac9g20788.1
Manihot___Manes.02G126700.2.p
Citrus___orange1.1g025225m|PACid_18111343
Arabidopsis−AT3G46870.1|PACid_19663606
Amborella___AmTr_v1.0_scaffold00066.3

OG0003791

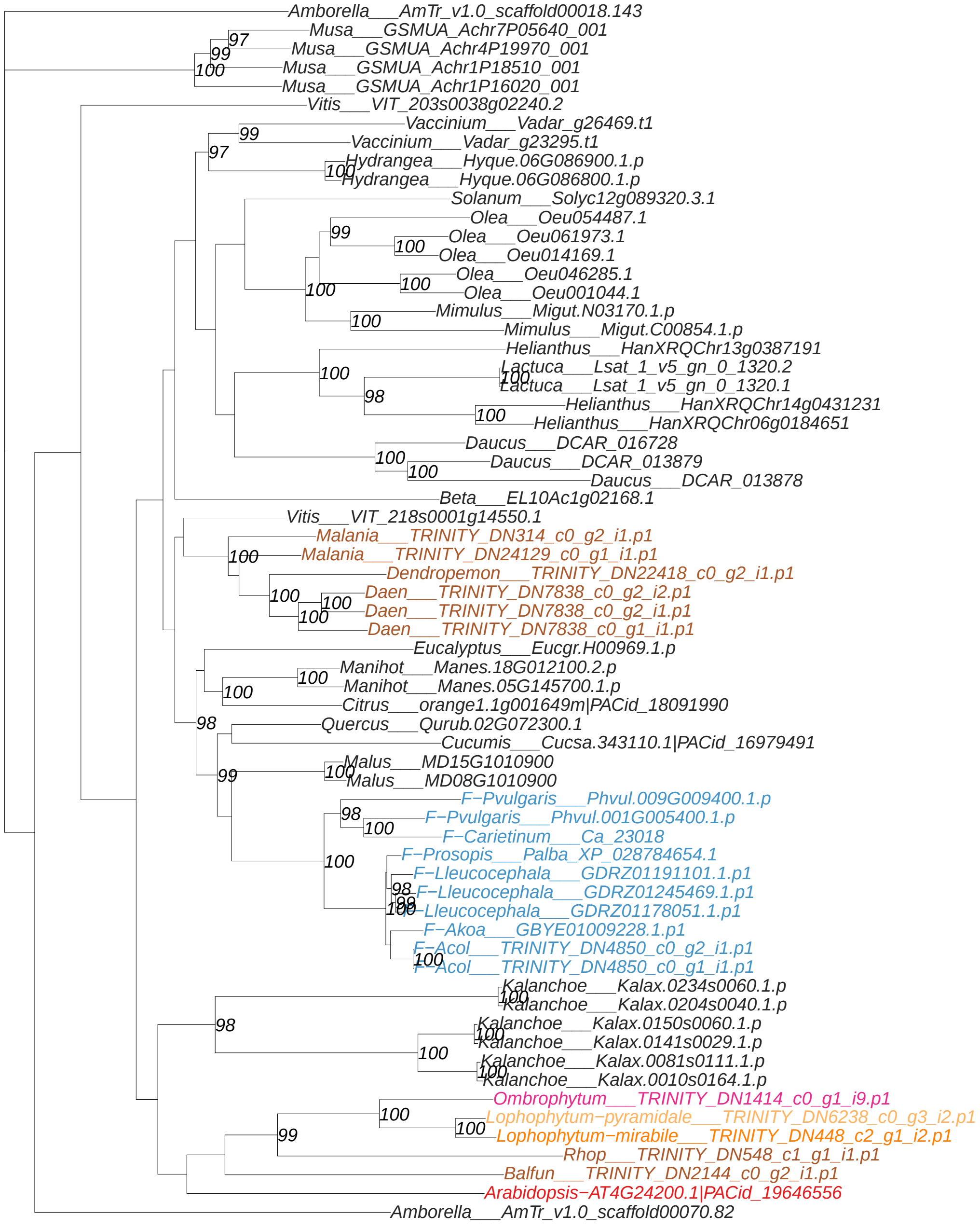
Amborella___AmTr_v1.0_scaffold00018.143
Musa___GSMUA_Achr7P05640_001
Musa___GSMUA_Achr4P19970_001
Musa___GSMUA_Achr1P18510_001
Musa___GSMUA_Achr1P16020_001
Vitis___VIT_203s0038g02240.2
Vaccinium___Vadar_g26469.t1
Vaccinium___Vadar_g23295.t1
Hydrangea___Hyque.06G086900.1.p
Hydrangea___Hyque.06G086800.1.p
Solanum___Solyc12g089320.3.1
Olea___Oeu054487.1
Olea___Oeu061973.1
Olea___Oeu014169.1
Olea___Oeu046285.1
Olea___Oeu001044.1
Mimulus___Migut.N03170.1.p
Mimulus___Migut.C00854.1.p
Helianthus___HanXRQChr13g0387191
Lactuca___Lsat_1_v5_gn_0_1320.2
Lactuca___Lsat_1_v5_gn_0_1320.1
Helianthus___HanXRQChr14g0431231
Helianthus___HanXRQChr06g0184651
Daucus___DCAR_016728
Daucus___DCAR_013879
Daucus___DCAR_013878
Beta___EL10Ac1g02168.1
Vitis___VIT_218s0001g14550.1
Malania___TRINITY_DN314_c0_g2_i1.p1
Malania___TRINITY_DN24129_c0_g1_i1.p1
Dendropemon___TRINITY_DN22418_c0_g2_i1.p1
Daen___TRINITY_DN7838_c0_g2_i2.p1
Daen___TRINITY_DN7838_c0_g2_i1.p1
Daen___TRINITY_DN7838_c0_g1_i1.p1
Eucalyptus___Eucgr.H00969.1.p
Manihot___Manes.18G012100.2.p
Manihot___Manes.05G145700.1.p
Citrus___orange1.1g001649m|PACid_18091990
Quercus___Qurub.02G072300.1
Cucumis___Cucsa.343110.1|PACid_16979491
Malus___MD15G1010900
Malus___MD08G1010900
F−Pvulgaris___Phvul.009G009400.1.p
F−Pvulgaris___Phvul.001G005400.1.p
F−Carietinum___Ca_23018
F−Prosopis___Palba_XP_028784654.1
F−Lleucocephala___GDRZ01191101.1.p1
F−Lleucocephala___GDRZ01245469.1.p1
F−Lleucocephala___GDRZ01178051.1.p1
F−Akoa___GBYE01009228.1.p1
F−Acol___TRINITY_DN4850_c0_g2_i1.p1
F−Acol___TRINITY_DN4850_c0_g1_i1.p1
Kalanchoe___Kalax.0234s0060.1.p
Kalanchoe___Kalax.0204s0040.1.p
Kalanchoe___Kalax.0150s0060.1.p
Kalanchoe___Kalax.0141s0029.1.p
Kalanchoe___Kalax.0081s0111.1.p
Kalanchoe___Kalax.0010s0164.1.p
Ombrophytum___TRINITY_DN1414_c0_g1_i9.p1
Lophophytum−pyramidale___TRINITY_DN6238_c0_g3_i2.p1
Lophophytum−mirabile___TRINITY_DN448_c2_g1_i2.p1
Rhop___TRINITY_DN548_c1_g1_i1.p1
Balfun___TRINITY_DN2144_c0_g2_i1.p1
Arabidopsis−AT4G24200.1|PACid_19646556
Amborella___AmTr_v1.0_scaffold00070.82

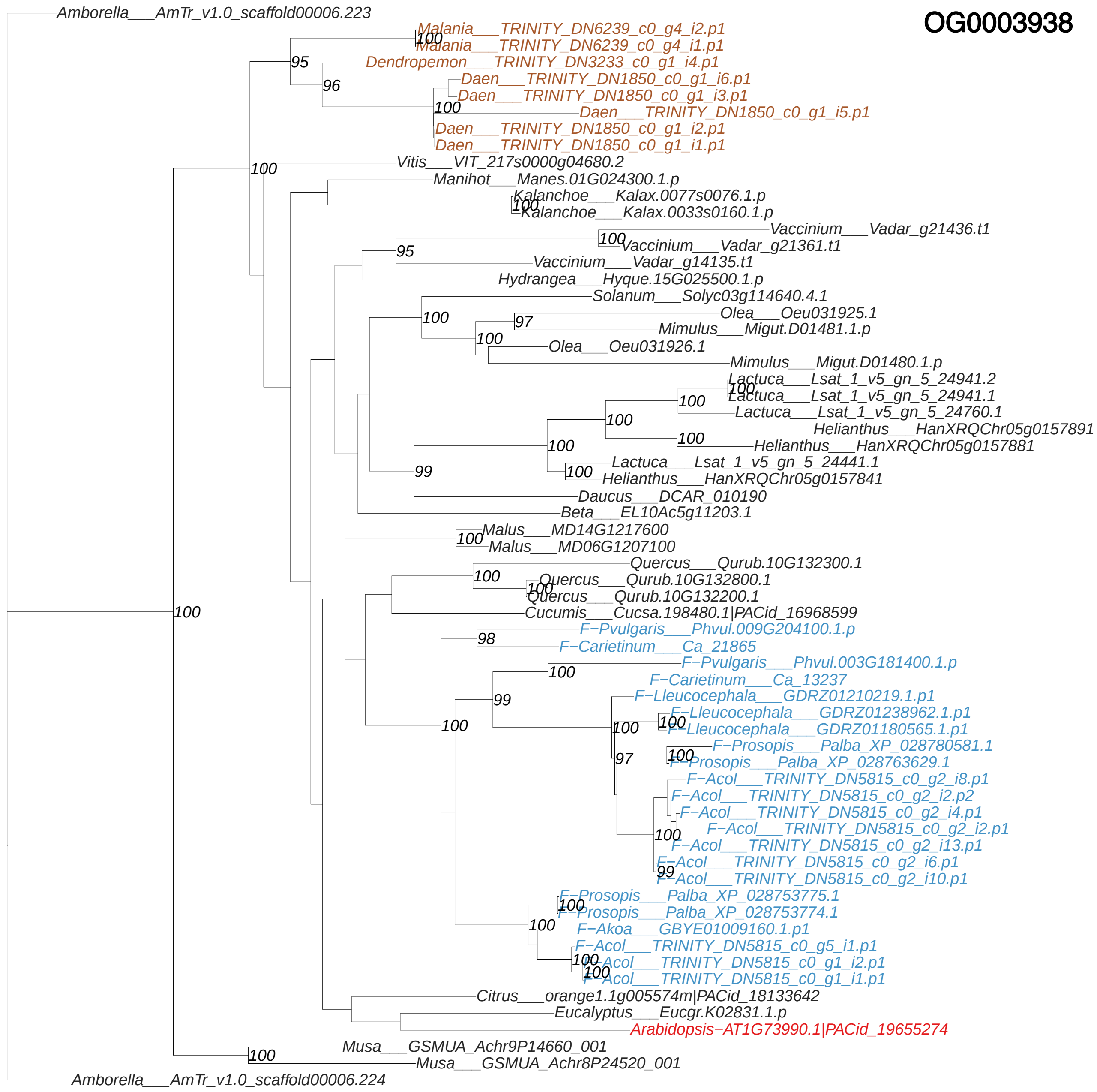
OG0003938
Amborella___AmTr_v1.0_scaffold00006.223
Malania___TRINITY_DN6239_c0_g4_i2.p1
Malania___TRINITY_DN6239_c0_g4_i1.p1
Dendropemon___TRINITY_DN3233_c0_g1_i4.p1
Daen___TRINITY_DN1850_c0_g1_i6.p1
Daen___TRINITY_DN1850_c0_g1_i3.p1
Daen___TRINITY_DN1850_c0_g1_i5.p1
Daen___TRINITY_DN1850_c0_g1_i2.p1
Daen___TRINITY_DN1850_c0_g1_i1.p1
Vitis___VIT_217s0000g04680.2
Manihot___Manes.01G024300.1.p
Kalanchoe___Kalax.0077s0076.1.p
Kalanchoe___Kalax.0033s0160.1.p
Vaccinium___Vadar_g21436.t1
Vaccinium___Vadar_g21361.t1
Vaccinium___Vadar_g14135.t1
Hydrangea___Hyque.15G025500.1.p
Solanum___Solyc03g114640.4.1
Olea___Oeu031925.1
Mimulus___Migut.D01481.1.p
Olea___Oeu031926.1
Mimulus___Migut.D01480.1.p
Lactuca___Lsat_1_v5_gn_5_24941.2
Lactuca___Lsat_1_v5_gn_5_24941.1
Lactuca___Lsat_1_v5_gn_5_24760.1
Helianthus___HanXRQChr05g0157891
Helianthus___HanXRQChr05g0157881
Lactuca___Lsat_1_v5_gn_5_24441.1
Helianthus___HanXRQChr05g0157841
Daucus___DCAR_010190
Beta___EL10Ac5g11203.1
Malus___MD14G1217600
Malus___MD06G1207100
Quercus___Qurub.10G132300.1
Quercus___Qurub.10G132800.1
Quercus___Qurub.10G132200.1
Cucumis___Cucsa.198480.1|PACid_16968599
F−Pvulgaris___Phvul.009G204100.1.p
F−Carietinum___Ca_21865
F−Pvulgaris___Phvul.003G181400.1.p
F−Carietinum___Ca_13237
F−Lleucocephala___GDRZ01210219.1.p1
F−Lleucocephala___GDRZ01238962.1.p1
F−Lleucocephala___GDRZ01180565.1.p1
F−Prosopis___Palba_XP_028780581.1
F−Prosopis___Palba_XP_028763629.1
F−Acol___TRINITY_DN5815_c0_g2_i8.p1
F−Acol___TRINITY_DN5815_c0_g2_i2.p2
F−Acol___TRINITY_DN5815_c0_g2_i4.p1
F−Acol___TRINITY_DN5815_c0_g2_i2.p1
F−Acol___TRINITY_DN5815_c0_g2_i13.p1
F−Acol___TRINITY_DN5815_c0_g2_i6.p1
F−Acol___TRINITY_DN5815_c0_g2_i10.p1
F−Prosopis___Palba_XP_028753775.1
F−Prosopis___Palba_XP_028753774.1
F−Akoa___GBYE01009160.1.p1
F−Acol___TRINITY_DN5815_c0_g5_i1.p1
F−Acol___TRINITY_DN5815_c0_g1_i2.p1
F−Acol___TRINITY_DN5815_c0_g1_i1.p1
Citrus___orange1.1g005574m|PACid_18133642
Eucalyptus___Eucgr.K02831.1.p
Arabidopsis−AT1G73990.1|PACid_19655274
Musa___GSMUA_Achr9P14660_001
Musa___GSMUA_Achr8P24520_001
Amborella___AmTr_v1.0_scaffold00006.224

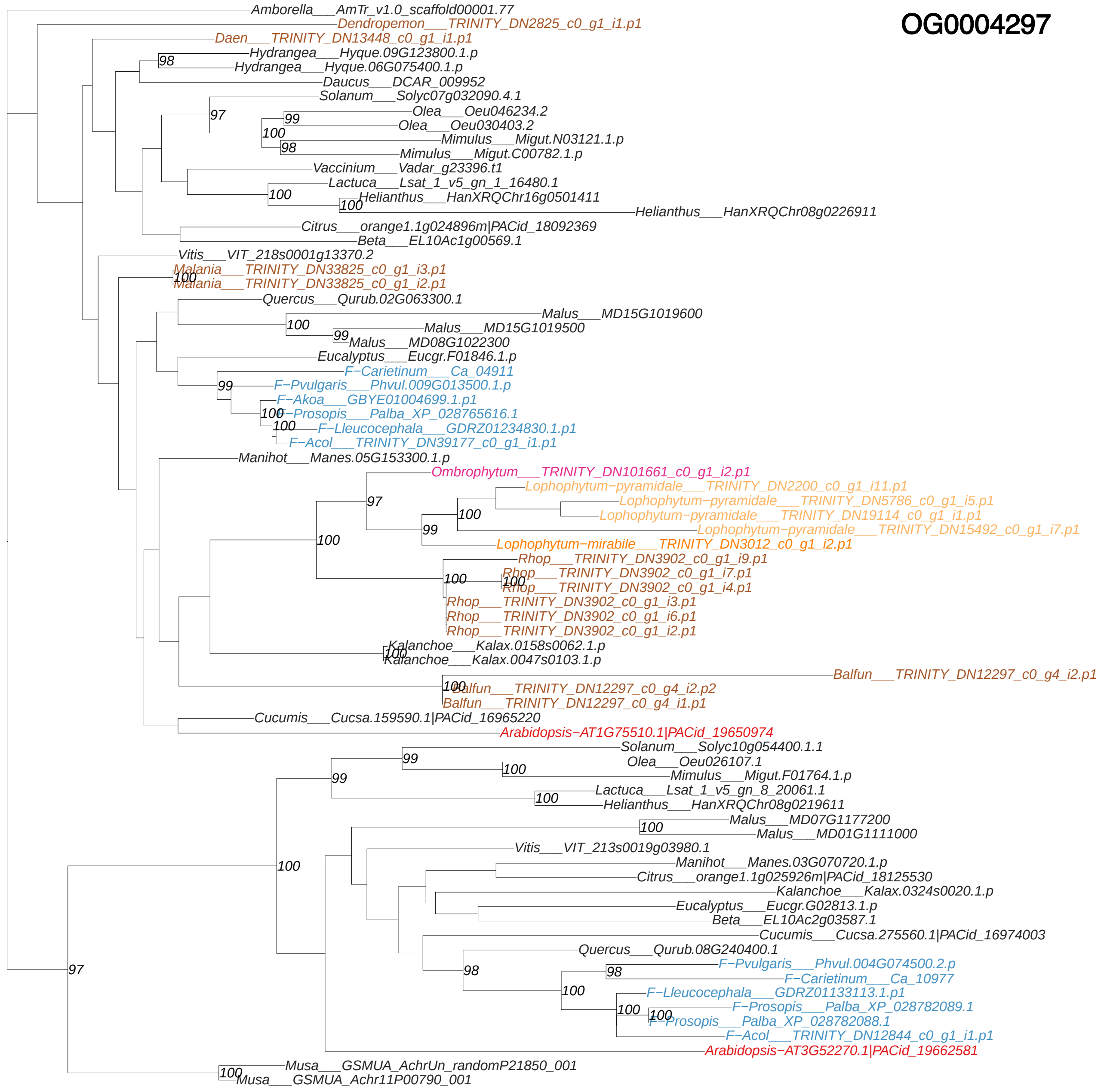
OG0004297
Amborella___AmTr_v1.0_scaffold00001.77
Dendropemon___TRINITY_DN2825_c0_g1_i1.p1
Daen___TRINITY_DN13448_c0_g1_i1.p1
Hydrangea___Hyque.09G123800.1.p
98
Hydrangea___Hyque.06G075400.1.p
Daucus___DCAR_009952
Solanum___Solyc07g032090.4.1
Olea___Oeu046234.2
97
99
Olea___Oeu030403.2
100
Mimulus___Migut.N03121.1.p
98
Mimulus___Migut.C00782.1.p
Vaccinium___Vadar_g23396.t1
Lactuca___Lsat_1_v5_gn_1_16480.1
100
Helianthus___HanXRQChr16g0501411
100
Helianthus___HanXRQChr08g0226911
Citrus___orange1.1g024896m|PACid_18092369
Beta___EL10Ac1g00569.1
Vitis___VIT_218s0001g13370.2
Malania___TRINITY_DN33825_c0_g1_i3.p1
100
Malania___TRINITY_DN33825_c0_g1_i2.p1
Quercus___Qurub.02G063300.1
Malus___MD15G1019600
100
Malus___MD15G1019500
99
Malus___MD08G1022300
Eucalyptus___Eucgr.F01846.1.p
F−Carietinum___Ca_04911
99
F−Pvulgaris___Phvul.009G013500.1.p
F−Akoa___GBYE01004699.1.p1
100
F−Prosopis___Palba_XP_028765616.1
100
F−Lleucocephala___GDRZ01234830.1.p1
F−Acol___TRINITY_DN39177_c0_g1_i1.p1
Manihot___Manes.05G153300.1.p
Ombrophytum___TRINITY_DN101661_c0_g1_i2.p1
Lophophytum−pyramidale___TRINITY_DN2200_c0_g1_i11.p1
97
Lophophytum−pyramidale___TRINITY_DN5786_c0_g1_i5.p1
100
Lophophytum−pyramidale___TRINITY_DN19114_c0_g1_i1.p1
99
Lophophytum−pyramidale___TRINITY_DN15492_c0_g1_i7.p1
100
Lophophytum−mirabile___TRINITY_DN3012_c0_g1_i2.p1
Rhop___TRINITY_DN3902_c0_g1_i9.p1
Rhop___TRINITY_DN3902_c0_g1_i7.p1
100
100
Rhop___TRINITY_DN3902_c0_g1_i4.p1
Rhop___TRINITY_DN3902_c0_g1_i3.p1
Rhop___TRINITY_DN3902_c0_g1_i6.p1
Rhop___TRINITY_DN3902_c0_g1_i2.p1
Kalanchoe___Kalax.0158s0062.1.p
100
Kalanchoe___Kalax.0047s0103.1.p
Balfun___TRINITY_DN12297_c0_g4_i2.p1
100
Balfun___TRINITY_DN12297_c0_g4_i2.p2
Balfun___TRINITY_DN12297_c0_g4_i1.p1
Cucumis___Cucsa.159590.1|PACid_16965220
Arabidopsis−AT1G75510.1|PACid_19650974
Solanum___Solyc10g054400.1.1
99
Olea___Oeu026107.1
100
99
Mimulus___Migut.F01764.1.p
Lactuca___Lsat_1_v5_gn_8_20061.1
100
Helianthus___HanXRQChr08g0219611
Malus___MD07G1177200
100
Malus___MD01G1111000
100
Vitis___VIT_213s0019g03980.1
Manihot___Manes.03G070720.1.p
Citrus___orange1.1g025926m|PACid_18125530
Kalanchoe___Kalax.0324s0020.1.p
Eucalyptus___Eucgr.G02813.1.p
Beta___EL10Ac2g03587.1
Cucumis___Cucsa.275560.1|PACid_16974003
Quercus___Qurub.08G240400.1
97
F−Pvulgaris___Phvul.004G074500.2.p
98
98
F−Carietinum___Ca_10977
100
F−Lleucocephala___GDRZ01133113.1.p1
F−Prosopis___Palba_XP_028782089.1
100
100
F−Prosopis___Palba_XP_028782088.1
F−Acol___TRINITY_DN12844_c0_g1_i1.p1
Arabidopsis−AT3G52270.1|PACid_19662581
Musa___GSMUA_AchrUn_randomP21850_001
100
Musa___GSMUA_Achr11P00790_001

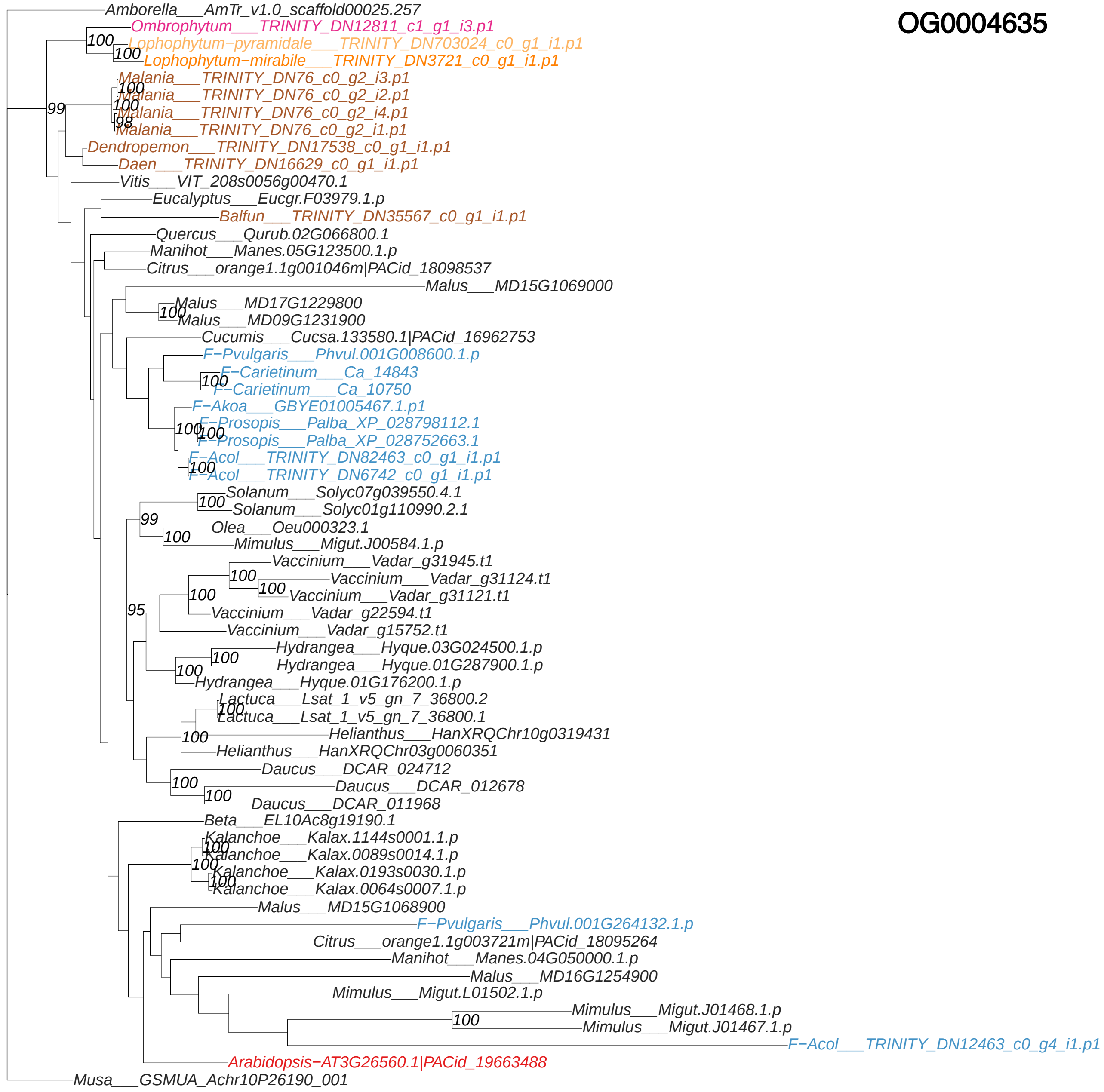

OG0004635
Amborella___AmTr_v1.0_scaffold00025.257
Ombrophytum___TRINITY_DN12811_c1_g1_i3.p1
Lophophytum−pyramidale___TRINITY_DN703024_c0_g1_i1.p1
Lophophytum−mirabile___TRINITY_DN3721_c0_g1_i1.p1
Malania___TRINITY_DN76_c0_g2_i3.p1
Malania___TRINITY_DN76_c0_g2_i2.p1
Malania___TRINITY_DN76_c0_g2_i4.p1
Malania___TRINITY_DN76_c0_g2_i1.p1
Dendropemon___TRINITY_DN17538_c0_g1_i1.p1
Daen___TRINITY_DN16629_c0_g1_i1.p1
Vitis___VIT_208s0056g00470.1
Eucalyptus___Eucgr.F03979.1.p
Balfun___TRINITY_DN35567_c0_g1_i1.p1
Quercus___Qurub.02G066800.1
Manihot___Manes.05G123500.1.p
Citrus___orange1.1g001046m|PACid_18098537
Malus___MD15G1069000
Malus___MD17G1229800
Malus___MD09G1231900
Cucumis___Cucsa.133580.1|PACid_16962753
F−Pvulgaris___Phvul.001G008600.1.p
F−Carietinum___Ca_14843
F−Carietinum___Ca_10750
F−Akoa___GBYE01005467.1.p1
F−Prosopis___Palba_XP_028798112.1
F−Prosopis___Palba_XP_028752663.1
F−Acol___TRINITY_DN82463_c0_g1_i1.p1
F−Acol___TRINITY_DN6742_c0_g1_i1.p1
Solanum___Solyc07g039550.4.1
Solanum___Solyc01g110990.2.1
Olea___Oeu000323.1
Mimulus___Migut.J00584.1.p
Vaccinium___Vadar_g31945.t1
Vaccinium___Vadar_g31124.t1
Vaccinium___Vadar_g31121.t1
Vaccinium___Vadar_g22594.t1
Vaccinium___Vadar_g15752.t1
Hydrangea___Hyque.03G024500.1.p
Hydrangea___Hyque.01G287900.1.p
Hydrangea___Hyque.01G176200.1.p
Lactuca___Lsat_1_v5_gn_7_36800.2
Lactuca___Lsat_1_v5_gn_7_36800.1
Helianthus___HanXRQChr10g0319431
Helianthus___HanXRQChr03g0060351
Daucus___DCAR_024712
Daucus___DCAR_012678
Daucus___DCAR_011968
Beta___EL10Ac8g19190.1
Kalanchoe___Kalax.1144s0001.1.p
Kalanchoe___Kalax.0089s0014.1.p
Kalanchoe___Kalax.0193s0030.1.p
Kalanchoe___Kalax.0064s0007.1.p
Malus___MD15G1068900
F−Pvulgaris___Phvul.001G264132.1.p
Citrus___orange1.1g003721m|PACid_18095264
Manihot___Manes.04G050000.1.p
Malus___MD16G1254900
Mimulus___Migut.L01502.1.p
Mimulus___Migut.J01468.1.p
Mimulus___Migut.J01467.1.p
F−Acol___TRINITY_DN12463_c0_g4_i1.p1
Arabidopsis−AT3G26560.1|PACid_19663488
Musa___GSMUA_Achr10P26190_001

# OG0005072

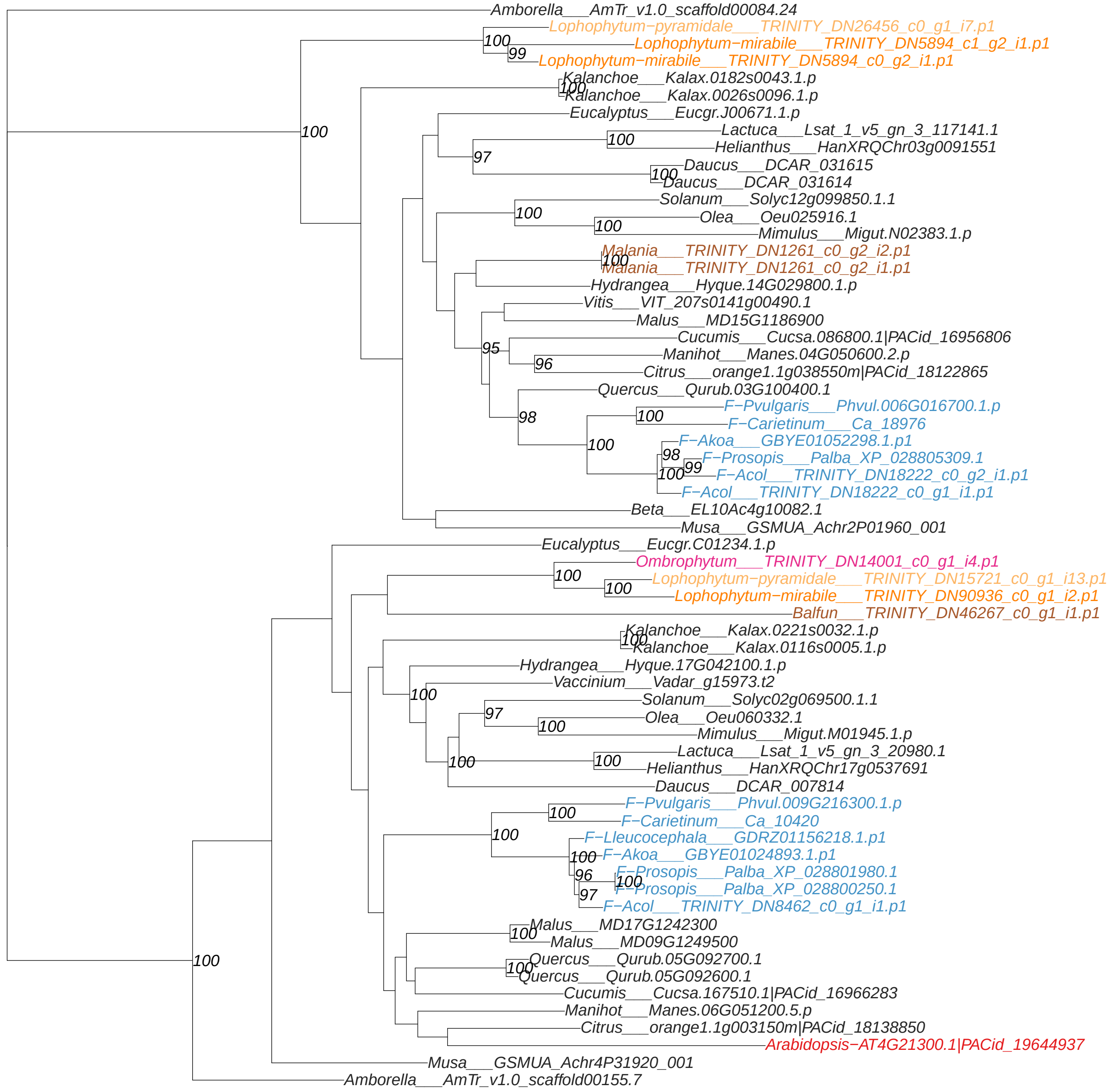

Amborella___AmTr_v1.0_scaffold00084.24
Lophophytum−pyramidale___TRINITY_DN26456_c0_g1_i7.p1
Lophophytum−mirabile___TRINITY_DN5894_c1_g2_i1.p1
Lophophytum−mirabile___TRINITY_DN5894_c0_g2_i1.p1
Kalanchoe___Kalax.0182s0043.1.p
Kalanchoe___Kalax.0026s0096.1.p
Eucalyptus___Eucgr.J00671.1.p
Lactuca___Lsat_1_v5_gn_3_117141.1
Helianthus___HanXRQChr03g0091551
Daucus___DCAR_031615
Daucus___DCAR_031614
Solanum___Solyc12g099850.1.1
Olea___Oeu025916.1
Mimulus___Migut.N02383.1.p
Malania___TRINITY_DN1261_c0_g2_i2.p1
Malania___TRINITY_DN1261_c0_g2_i1.p1
Hydrangea___Hyque.14G029800.1.p
Vitis___VIT_207s0141g00490.1
Malus___MD15G1186900
Cucumis___Cucsa.086800.1|PACid_16956806
Manihot___Manes.04G050600.2.p
Citrus___orange1.1g038550m|PACid_18122865
Quercus___Qurub.03G100400.1
F−Pvulgaris___Phvul.006G016700.1.p
F−Carietinum___Ca_18976
F−Akoa___GBYE01052298.1.p1
F−Prosopis___Palba_XP_028805309.1
F−Acol___TRINITY_DN18222_c0_g2_i1.p1
F−Acol___TRINITY_DN18222_c0_g1_i1.p1
Beta___EL10Ac4g10082.1
Musa___GSMUA_Achr2P01960_001
Eucalyptus___Eucgr.C01234.1.p
Ombrophytum___TRINITY_DN14001_c0_g1_i4.p1
Lophophytum−pyramidale___TRINITY_DN15721_c0_g1_i13.p1
Lophophytum−mirabile___TRINITY_DN90936_c0_g1_i2.p1
Balfun___TRINITY_DN46267_c0_g1_i1.p1
Kalanchoe___Kalax.0221s0032.1.p
Kalanchoe___Kalax.0116s0005.1.p
Hydrangea___Hyque.17G042100.1.p
Vaccinium___Vadar_g15973.t2
Solanum___Solyc02g069500.1.1
Olea___Oeu060332.1
Mimulus___Migut.M01945.1.p
Lactuca___Lsat_1_v5_gn_3_20980.1
Helianthus___HanXRQChr17g0537691
Daucus___DCAR_007814
F−Pvulgaris___Phvul.009G216300.1.p
F−Carietinum___Ca_10420
F−Lleucocephala___GDRZ01156218.1.p1
F−Akoa___GBYE01024893.1.p1
F−Prosopis___Palba_XP_028801980.1
F−Prosopis___Palba_XP_028800250.1
F−Acol___TRINITY_DN8462_c0_g1_i1.p1
Malus___MD17G1242300
Malus___MD09G1249500
Quercus___Qurub.05G092700.1
Quercus___Qurub.05G092600.1
Cucumis___Cucsa.167510.1|PACid_16966283
Manihot___Manes.06G051200.5.p
Citrus___orange1.1g003150m|PACid_18138850
Arabidopsis−AT4G21300.1|PACid_19644937
Musa___GSMUA_Achr4P31920_001
Amborella___AmTr_v1.0_scaffold00155.7

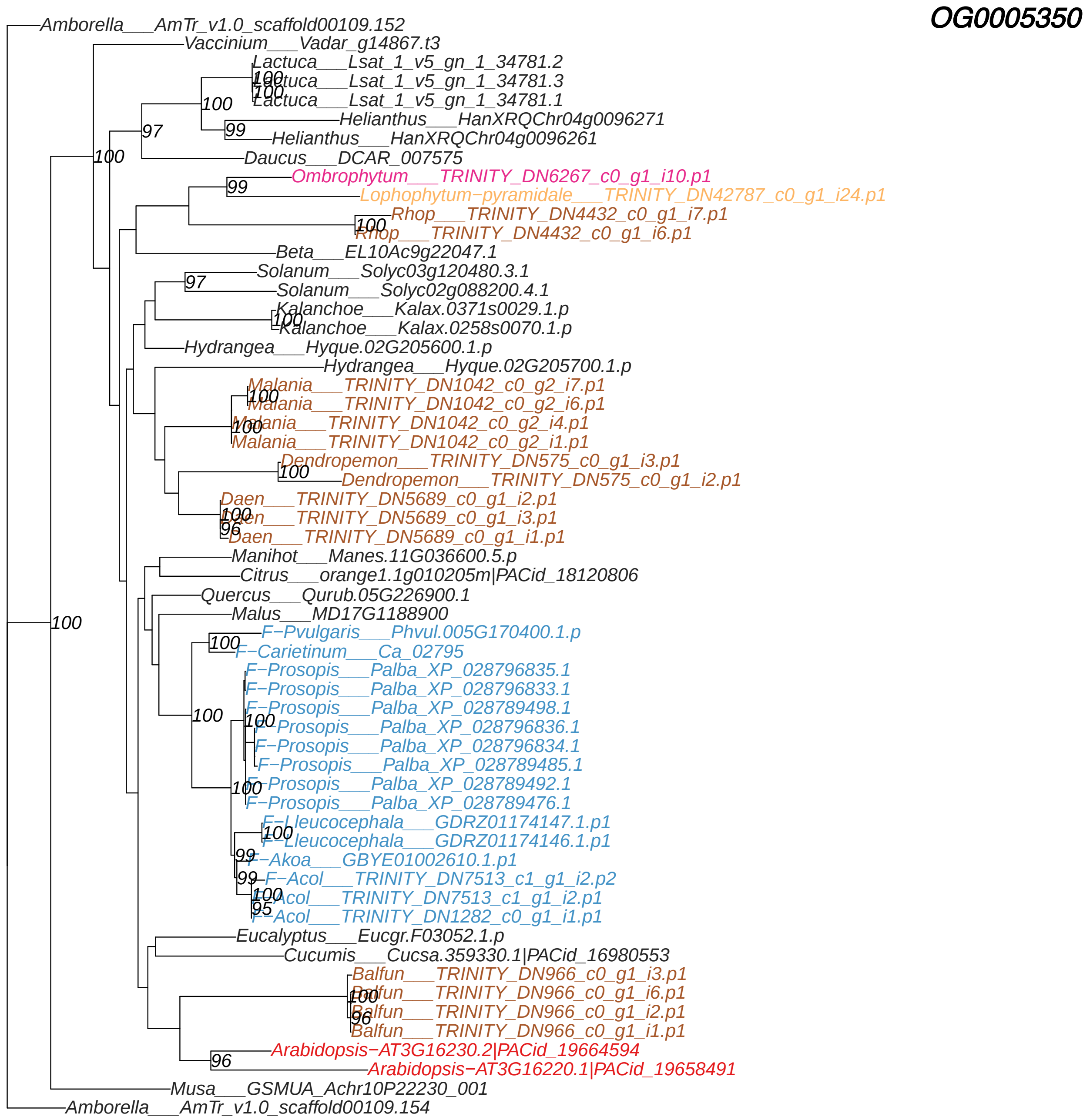
OG0005350
Amborella___AmTr_v1.0_scaffold00109.152
Vaccinium___Vadar_g14867.t3
Lactuca___Lsat_1_v5_gn_1_34781.2
Lactuca___Lsat_1_v5_gn_1_34781.3
Lactuca___Lsat_1_v5_gn_1_34781.1
Helianthus___HanXRQChr04g0096271
Helianthus___HanXRQChr04g0096261
Daucus___DCAR_007575
Ombrophytum___TRINITY_DN6267_c0_g1_i10.p1
Lophophytum−pyramidale___TRINITY_DN42787_c0_g1_i24.p1
Rhop___TRINITY_DN4432_c0_g1_i7.p1
Rhop___TRINITY_DN4432_c0_g1_i6.p1
Beta___EL10Ac9g22047.1
Solanum___Solyc03g120480.3.1
Solanum___Solyc02g088200.4.1
Kalanchoe___Kalax.0371s0029.1.p
Kalanchoe___Kalax.0258s0070.1.p
Hydrangea___Hyque.02G205600.1.p
Hydrangea___Hyque.02G205700.1.p
Malania___TRINITY_DN1042_c0_g2_i7.p1
Malania___TRINITY_DN1042_c0_g2_i6.p1
Malania___TRINITY_DN1042_c0_g2_i4.p1
Malania___TRINITY_DN1042_c0_g2_i1.p1
Dendropemon___TRINITY_DN575_c0_g1_i3.p1
Dendropemon___TRINITY_DN575_c0_g1_i2.p1
Daen___TRINITY_DN5689_c0_g1_i2.p1
Daen___TRINITY_DN5689_c0_g1_i3.p1
Daen___TRINITY_DN5689_c0_g1_i1.p1
Manihot___Manes.11G036600.5.p
Citrus___orange1.1g010205m|PACid_18120806
Quercus___Qurub.05G226900.1
Malus___MD17G1188900
F−Pvulgaris___Phvul.005G170400.1.p
F−Carietinum___Ca_02795
F−Prosopis___Palba_XP_028796835.1
F−Prosopis___Palba_XP_028796833.1
F−Prosopis___Palba_XP_028789498.1
F−Prosopis___Palba_XP_028796836.1
F−Prosopis___Palba_XP_028796834.1
F−Prosopis___Palba_XP_028789485.1
F−Prosopis___Palba_XP_028789492.1
F−Prosopis___Palba_XP_028789476.1
F−Lleucocephala___GDRZ01174147.1.p1
F−Lleucocephala___GDRZ01174146.1.p1
F−Akoa___GBYE01002610.1.p1
F−Acol___TRINITY_DN7513_c1_g1_i2.p2
F−Acol___TRINITY_DN7513_c1_g1_i2.p1
F−Acol___TRINITY_DN1282_c0_g1_i1.p1
Eucalyptus___Eucgr.F03052.1.p
Cucumis___Cucsa.359330.1|PACid_16980553
Balfun___TRINITY_DN966_c0_g1_i3.p1
Balfun___TRINITY_DN966_c0_g1_i6.p1
Balfun___TRINITY_DN966_c0_g1_i2.p1
Balfun___TRINITY_DN966_c0_g1_i1.p1
Arabidopsis−AT3G16230.2|PACid_19664594
Arabidopsis−AT3G16220.1|PACid_19658491
Musa___GSMUA_Achr10P22230_001
Amborella___AmTr_v1.0_scaffold00109.154

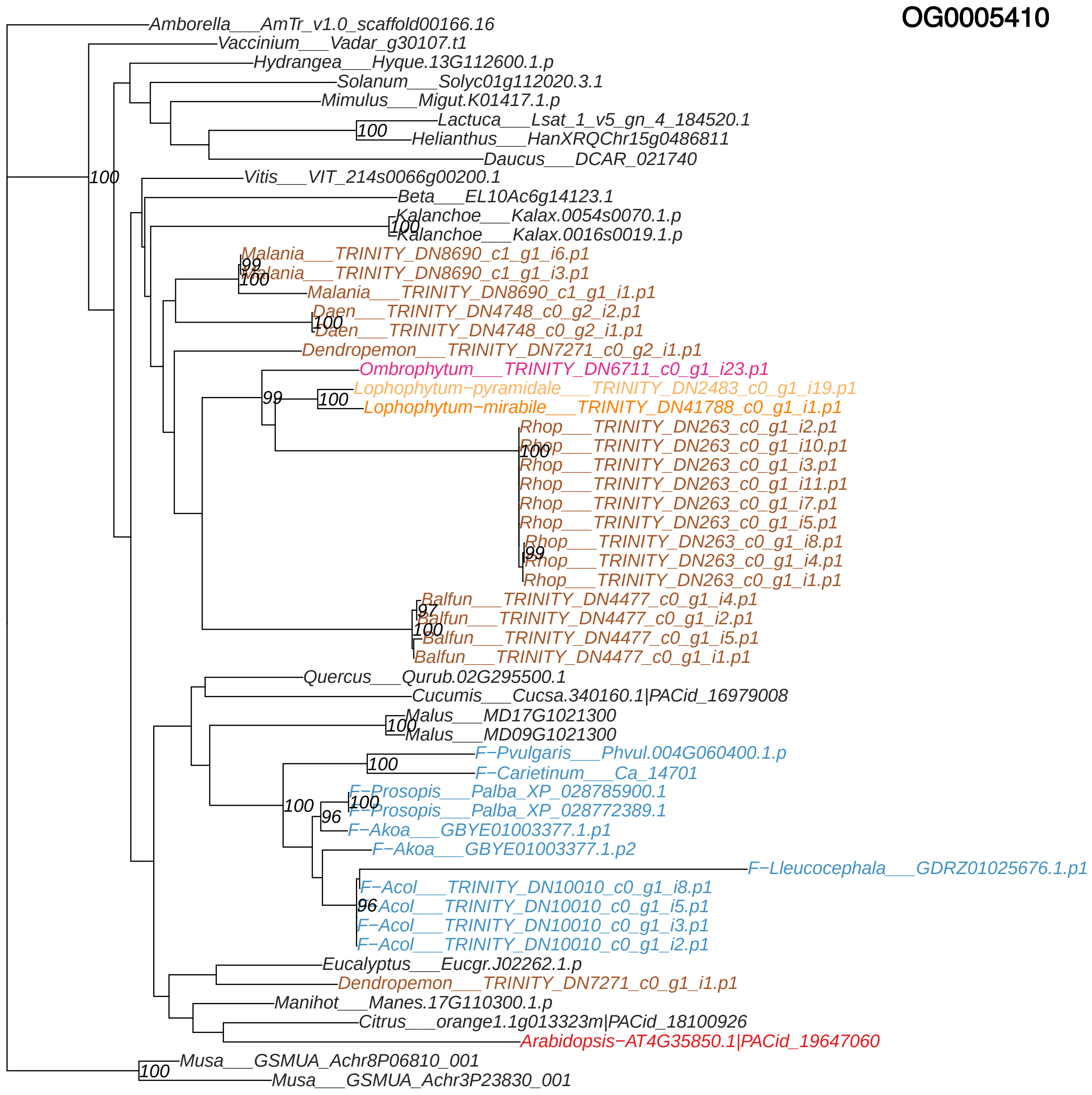
OG0005410
Amborella___AmTr_v1.0_scaffold00166.16
Vaccinium___Vadar_g30107.t1
Hydrangea___Hyque.13G112600.1.p
Solanum___Solyc01g112020.3.1
Mimulus___Migut.K01417.1.p
Lactuca___Lsat_1_v5_gn_4_184520.1
100
Helianthus___HanXRQChr15g0486811
Daucus___DCAR_021740
100
Vitis___VIT_214s0066g00200.1
Beta___EL10Ac6g14123.1
Kalanchoe___Kalax.0054s0070.1.p
100
Kalanchoe___Kalax.0016s0019.1.p
Malania___TRINITY_DN8690_c1_g1_i6.p1
99
Malania___TRINITY_DN8690_c1_g1_i3.p1
100
Malania___TRINITY_DN8690_c1_g1_i1.p1
Daen___TRINITY_DN4748_c0_g2_i2.p1
100
Daen___TRINITY_DN4748_c0_g2_i1.p1
Dendropemon___TRINITY_DN7271_c0_g2_i1.p1
Ombrophytum___TRINITY_DN6711_c0_g1_i23.p1
Lophophytum−pyramidale___TRINITY_DN2483_c0_g1_i19.p1
99
100
Lophophytum−mirabile___TRINITY_DN41788_c0_g1_i1.p1
Rhop___TRINITY_DN263_c0_g1_i2.p1
Rhop___TRINITY_DN263_c0_g1_i10.p1
100
Rhop___TRINITY_DN263_c0_g1_i3.p1
Rhop___TRINITY_DN263_c0_g1_i11.p1
Rhop___TRINITY_DN263_c0_g1_i7.p1
Rhop___TRINITY_DN263_c0_g1_i5.p1
Rhop___TRINITY_DN263_c0_g1_i8.p1
99
Rhop___TRINITY_DN263_c0_g1_i4.p1
Rhop___TRINITY_DN263_c0_g1_i1.p1
Balfun___TRINITY_DN4477_c0_g1_i4.p1
97
Balfun___TRINITY_DN4477_c0_g1_i2.p1
100
Balfun___TRINITY_DN4477_c0_g1_i5.p1
Balfun___TRINITY_DN4477_c0_g1_i1.p1
Quercus___Qurub.02G295500.1
Cucumis___Cucsa.340160.1|PACid_16979008
Malus___MD17G1021300
100
Malus___MD09G1021300
F−Pvulgaris___Phvul.004G060400.1.p
100
F−Carietinum___Ca_14701
F−Prosopis___Palba_XP_028785900.1
100
100
F−Prosopis___Palba_XP_028772389.1
96
F−Akoa___GBYE01003377.1.p1
F−Akoa___GBYE01003377.1.p2
F−Lleucocephala___GDRZ01025676.1.p1
F−Acol___TRINITY_DN10010_c0_g1_i8.p1
96
F−Acol___TRINITY_DN10010_c0_g1_i5.p1
F−Acol___TRINITY_DN10010_c0_g1_i3.p1
F−Acol___TRINITY_DN10010_c0_g1_i2.p1
Eucalyptus___Eucgr.J02262.1.p
Dendropemon___TRINITY_DN7271_c0_g1_i1.p1
Manihot___Manes.17G110300.1.p
Citrus___orange1.1g013323m|PACid_18100926
Arabidopsis−AT4G35850.1|PACid_19647060
Musa___GSMUA_Achr8P06810_001
100
Musa___GSMUA_Achr3P23830_001

# OG0005494

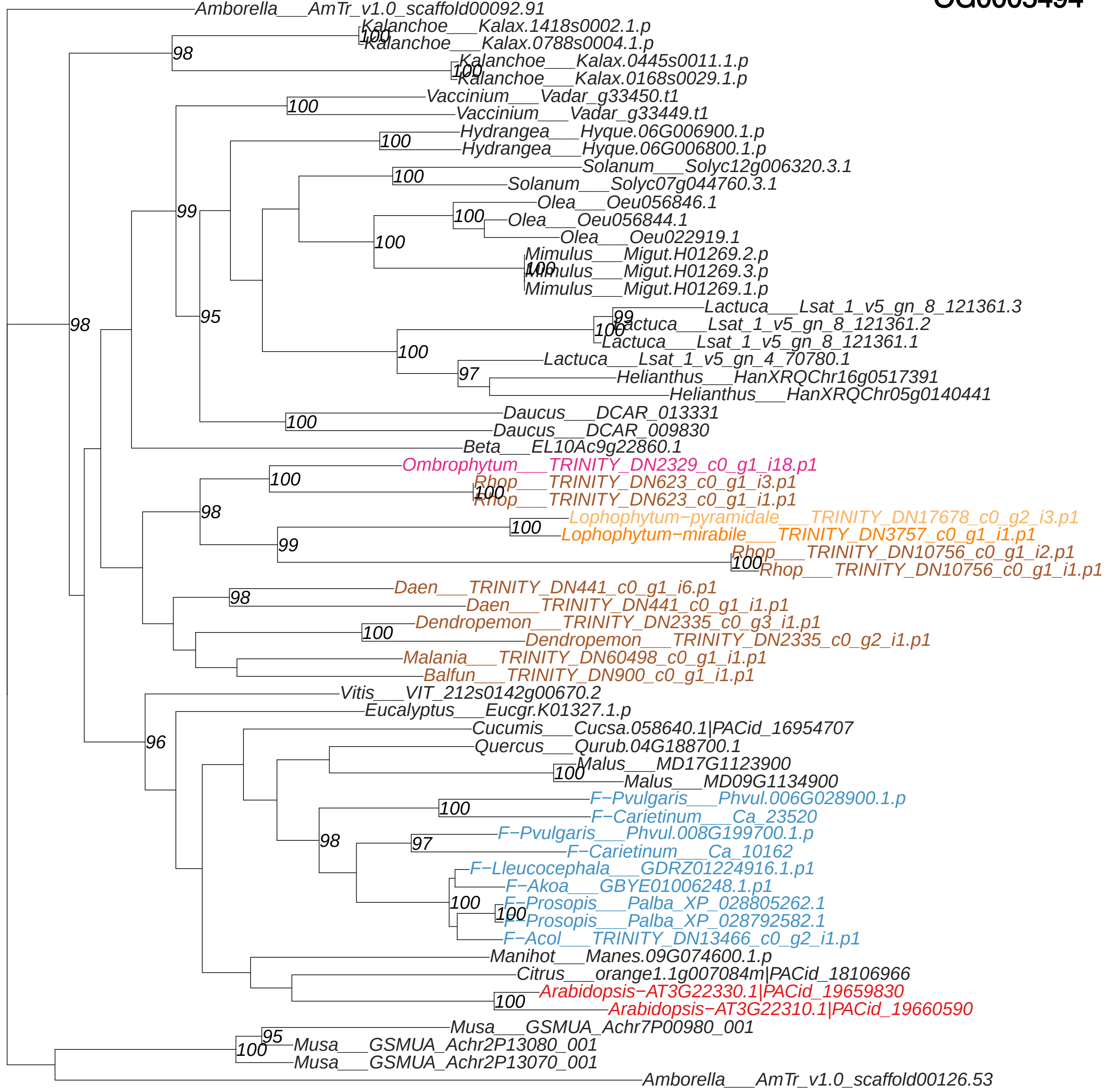
Amborella___AmTr_v1.0_scaffold00092.91
Kalanchoe___Kalax.1418s0002.1.p
Kalanchoe___Kalax.0788s0004.1.p
Kalanchoe___Kalax.0445s0011.1.p
Kalanchoe___Kalax.0168s0029.1.p
Vaccinium___Vadar_g33450.t1
Vaccinium___Vadar_g33449.t1
Hydrangea___Hyque.06G006900.1.p
Hydrangea___Hyque.06G006800.1.p
Solanum___Solyc12g006320.3.1
Solanum___Solyc07g044760.3.1
Olea___Oeu056846.1
Olea___Oeu056844.1
Olea___Oeu022919.1
Mimulus___Migut.H01269.2.p
Mimulus___Migut.H01269.3.p
Mimulus___Migut.H01269.1.p
Lactuca___Lsat_1_v5_gn_8_121361.3
Lactuca___Lsat_1_v5_gn_8_121361.2
Lactuca___Lsat_1_v5_gn_8_121361.1
Lactuca___Lsat_1_v5_gn_4_70780.1
Helianthus___HanXRQChr16g0517391
Helianthus___HanXRQChr05g0140441
Daucus___DCAR_013331
Daucus___DCAR_009830
Beta___EL10Ac9g22860.1
Ombrophytum___TRINITY_DN2329_c0_g1_i18.p1
Rhop___TRINITY_DN623_c0_g1_i3.p1
Rhop___TRINITY_DN623_c0_g1_i1.p1
Lophophytum−pyramidale___TRINITY_DN17678_c0_g2_i3.p1
Lophophytum−mirabile___TRINITY_DN3757_c0_g1_i1.p1
Rhop___TRINITY_DN10756_c0_g1_i2.p1
Rhop___TRINITY_DN10756_c0_g1_i1.p1
Daen___TRINITY_DN441_c0_g1_i6.p1
Daen___TRINITY_DN441_c0_g1_i1.p1
Dendropemon___TRINITY_DN2335_c0_g3_i1.p1
Dendropemon___TRINITY_DN2335_c0_g2_i1.p1
Malania___TRINITY_DN60498_c0_g1_i1.p1
Balfun___TRINITY_DN900_c0_g1_i1.p1
Vitis___VIT_212s0142g00670.2
Eucalyptus___Eucgr.K01327.1.p
Cucumis___Cucsa.058640.1|PACid_16954707
Quercus___Qurub.04G188700.1
Malus___MD17G1123900
Malus___MD09G1134900
F−Pvulgaris___Phvul.006G028900.1.p
F−Carietinum___Ca_23520
F−Pvulgaris___Phvul.008G199700.1.p
F−Carietinum___Ca_10162
F−Lleucocephala___GDRZ01224916.1.p1
F−Akoa___GBYE01006248.1.p1
F−Prosopis___Palba_XP_028805262.1
F−Prosopis___Palba_XP_028792582.1
F−Acol___TRINITY_DN13466_c0_g2_i1.p1
Manihot___Manes.09G074600.1.p
Citrus___orange1.1g007084m|PACid_18106966
Arabidopsis−AT3G22330.1|PACid_19659830
Arabidopsis−AT3G22310.1|PACid_19660590
Musa___GSMUA_Achr7P00980_001
Musa___GSMUA_Achr2P13080_001
Musa___GSMUA_Achr2P13070_001
Amborella___AmTr_v1.0_scaffold00126.53

# OG0005775

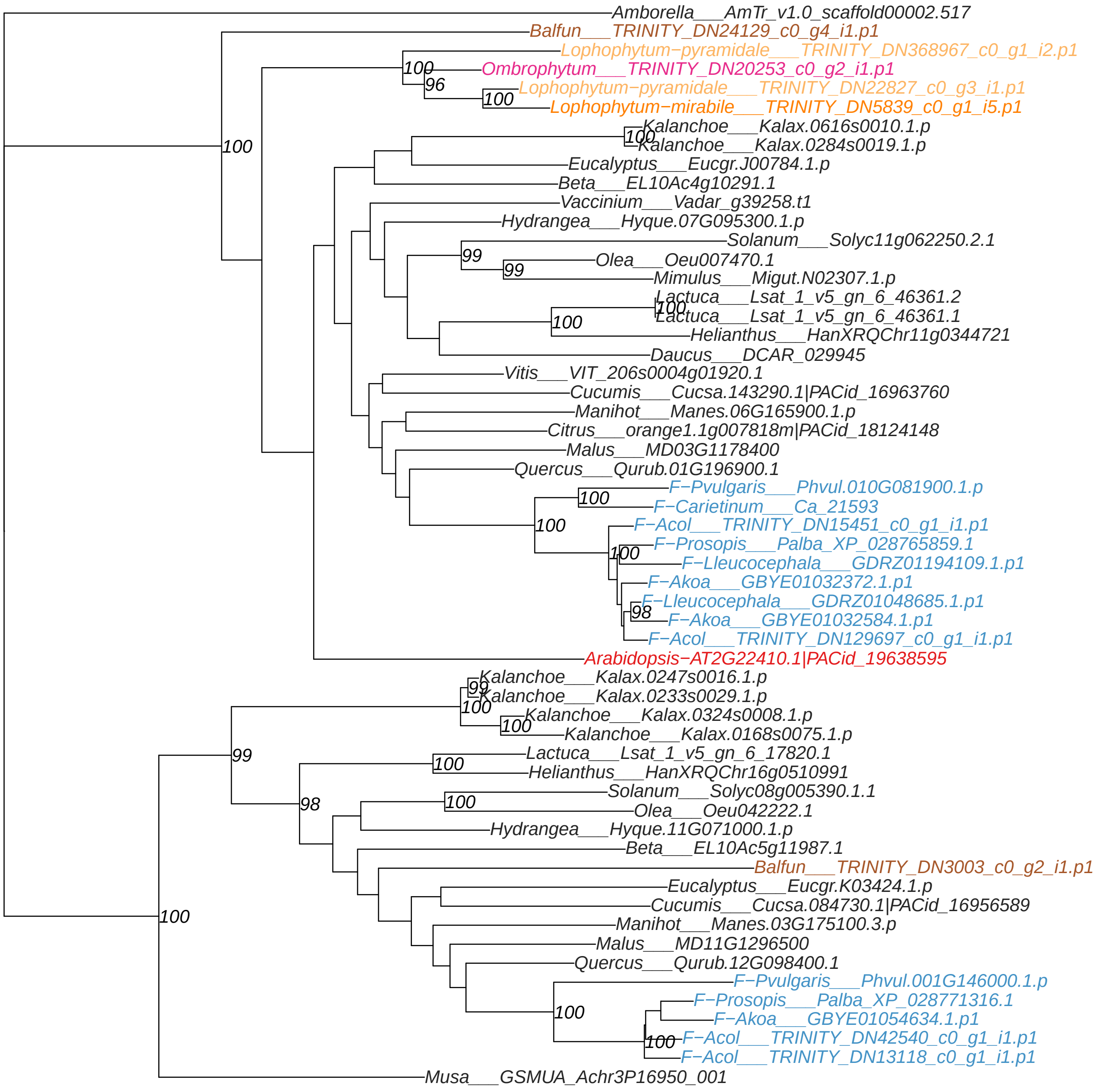

Amborella___AmTr_v1.0_scaffold00002.517
Balfun___TRINITY_DN24129_c0_g4_i1.p1
Lophophytum−pyramidale___TRINITY_DN368967_c0_g1_i2.p1
Ombrophytum___TRINITY_DN20253_c0_g2_i1.p1
Lophophytum−pyramidale___TRINITY_DN22827_c0_g3_i1.p1
Lophophytum−mirabile___TRINITY_DN5839_c0_g1_i5.p1
Kalanchoe___Kalax.0616s0010.1.p
Kalanchoe___Kalax.0284s0019.1.p
Eucalyptus___Eucgr.J00784.1.p
Beta___EL10Ac4g10291.1
Vaccinium___Vadar_g39258.t1
Hydrangea___Hyque.07G095300.1.p
Solanum___Solyc11g062250.2.1
Olea___Oeu007470.1
Mimulus___Migut.N02307.1.p
Lactuca___Lsat_1_v5_gn_6_46361.2
Lactuca___Lsat_1_v5_gn_6_46361.1
Helianthus___HanXRQChr11g0344721
Daucus___DCAR_029945
Vitis___VIT_206s0004g01920.1
Cucumis___Cucsa.143290.1|PACid_16963760
Manihot___Manes.06G165900.1.p
Citrus___orange1.1g007818m|PACid_18124148
Malus___MD03G1178400
Quercus___Qurub.01G196900.1
F−Pvulgaris___Phvul.010G081900.1.p
F−Carietinum___Ca_21593
F−Acol___TRINITY_DN15451_c0_g1_i1.p1
F−Prosopis___Palba_XP_028765859.1
F−Lleucocephala___GDRZ01194109.1.p1
F−Akoa___GBYE01032372.1.p1
F−Lleucocephala___GDRZ01048685.1.p1
F−Akoa___GBYE01032584.1.p1
F−Acol___TRINITY_DN129697_c0_g1_i1.p1
Arabidopsis−AT2G22410.1|PACid_19638595
Kalanchoe___Kalax.0247s0016.1.p
Kalanchoe___Kalax.0233s0029.1.p
Kalanchoe___Kalax.0324s0008.1.p
Kalanchoe___Kalax.0168s0075.1.p
Lactuca___Lsat_1_v5_gn_6_17820.1
Helianthus___HanXRQChr16g0510991
Solanum___Solyc08g005390.1.1
Olea___Oeu042222.1
Hydrangea___Hyque.11G071000.1.p
Beta___EL10Ac5g11987.1
Balfun___TRINITY_DN3003_c0_g2_i1.p1
Eucalyptus___Eucgr.K03424.1.p
Cucumis___Cucsa.084730.1|PACid_16956589
Manihot___Manes.03G175100.3.p
Malus___MD11G1296500
Quercus___Qurub.12G098400.1
F−Pvulgaris___Phvul.001G146000.1.p
F−Prosopis___Palba_XP_028771316.1
F−Akoa___GBYE01054634.1.p1
F−Acol___TRINITY_DN42540_c0_g1_i1.p1
F−Acol___TRINITY_DN13118_c0_g1_i1.p1
Musa___GSMUA_Achr3P16950_001
100
96
100
100
100
99
99
100
100
100
100
100
98
99
100
100
99
100
98
100
100
100
100

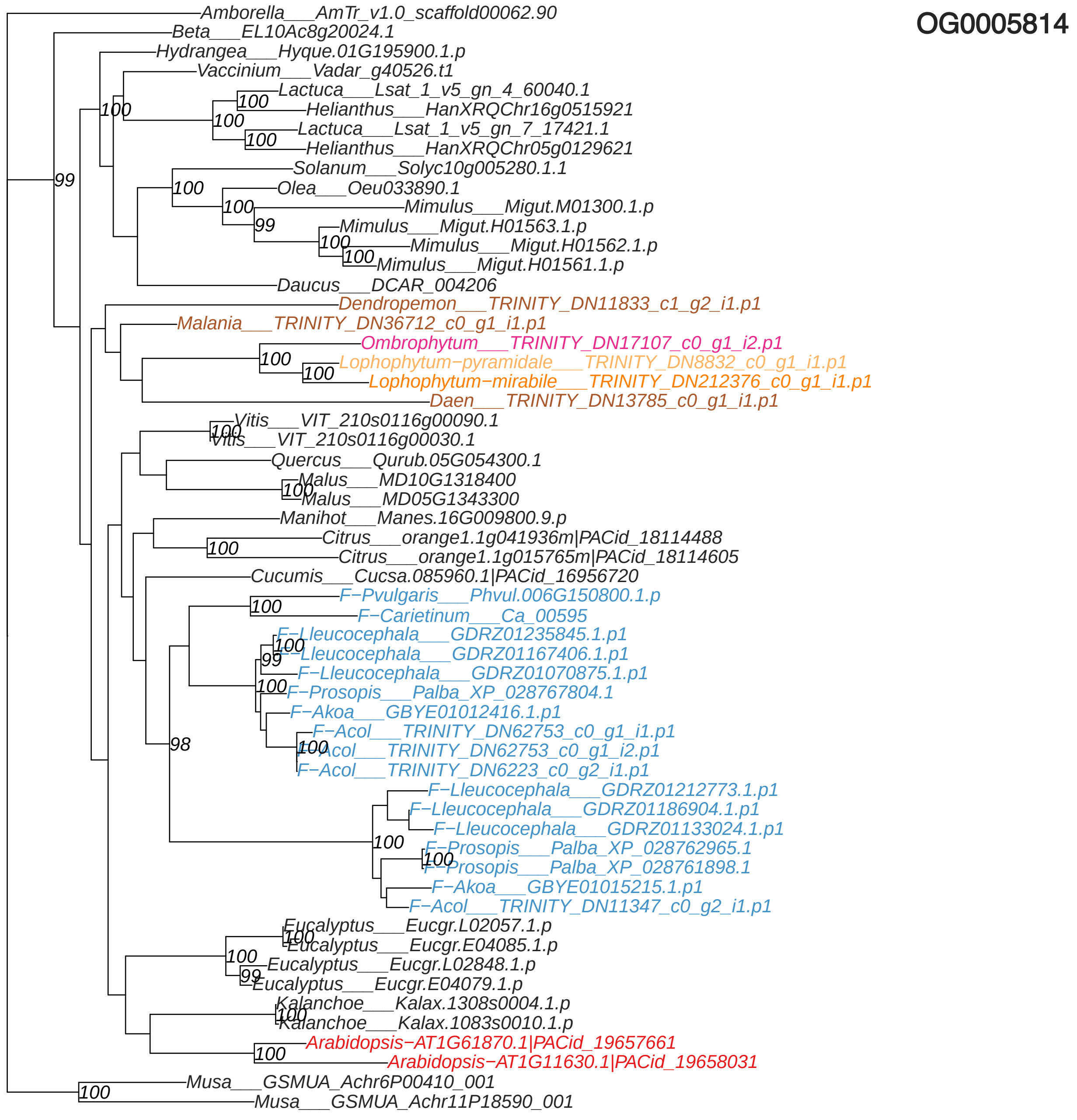
OG0005814
Amborella___AmTr_v1.0_scaffold00062.90
Beta___EL10Ac8g20024.1
Hydrangea___Hyque.01G195900.1.p
Vaccinium___Vadar_g40526.t1
Lactuca___Lsat_1_v5_gn_4_60040.1
Helianthus___HanXRQChr16g0515921
Lactuca___Lsat_1_v5_gn_7_17421.1
Helianthus___HanXRQChr05g0129621
Solanum___Solyc10g005280.1.1
Olea___Oeu033890.1
Mimulus___Migut.M01300.1.p
Mimulus___Migut.H01563.1.p
Mimulus___Migut.H01562.1.p
Mimulus___Migut.H01561.1.p
Daucus___DCAR_004206
Dendropemon___TRINITY_DN11833_c1_g2_i1.p1
Malania___TRINITY_DN36712_c0_g1_i1.p1
Ombrophytum___TRINITY_DN17107_c0_g1_i2.p1
Lophophytum−pyramidale___TRINITY_DN8832_c0_g1_i1.p1
Lophophytum−mirabile___TRINITY_DN212376_c0_g1_i1.p1
Daen___TRINITY_DN13785_c0_g1_i1.p1
Vitis___VIT_210s0116g00090.1
Vitis___VIT_210s0116g00030.1
Quercus___Qurub.05G054300.1
Malus___MD10G1318400
Malus___MD05G1343300
Manihot___Manes.16G009800.9.p
Citrus___orange1.1g041936m|PACid_18114488
Citrus___orange1.1g015765m|PACid_18114605
Cucumis___Cucsa.085960.1|PACid_16956720
F−Pvulgaris___Phvul.006G150800.1.p
F−Carietinum___Ca_00595
F−Lleucocephala___GDRZ01235845.1.p1
F−Lleucocephala___GDRZ01167406.1.p1
F−Lleucocephala___GDRZ01070875.1.p1
F−Prosopis___Palba_XP_028767804.1
F−Akoa___GBYE01012416.1.p1
F−Acol___TRINITY_DN62753_c0_g1_i1.p1
F−Acol___TRINITY_DN62753_c0_g1_i2.p1
F−Acol___TRINITY_DN6223_c0_g2_i1.p1
F−Lleucocephala___GDRZ01212773.1.p1
F−Lleucocephala___GDRZ01186904.1.p1
F−Lleucocephala___GDRZ01133024.1.p1
F−Prosopis___Palba_XP_028762965.1
F−Prosopis___Palba_XP_028761898.1
F−Akoa___GBYE01015215.1.p1
F−Acol___TRINITY_DN11347_c0_g2_i1.p1
Eucalyptus___Eucgr.L02057.1.p
Eucalyptus___Eucgr.E04085.1.p
Eucalyptus___Eucgr.L02848.1.p
Eucalyptus___Eucgr.E04079.1.p
Kalanchoe___Kalax.1308s0004.1.p
Kalanchoe___Kalax.1083s0010.1.p
Arabidopsis−AT1G61870.1|PACid_19657661
Arabidopsis−AT1G11630.1|PACid_19658031
Musa___GSMUA_Achr6P00410_001
Musa___GSMUA_Achr11P18590_001

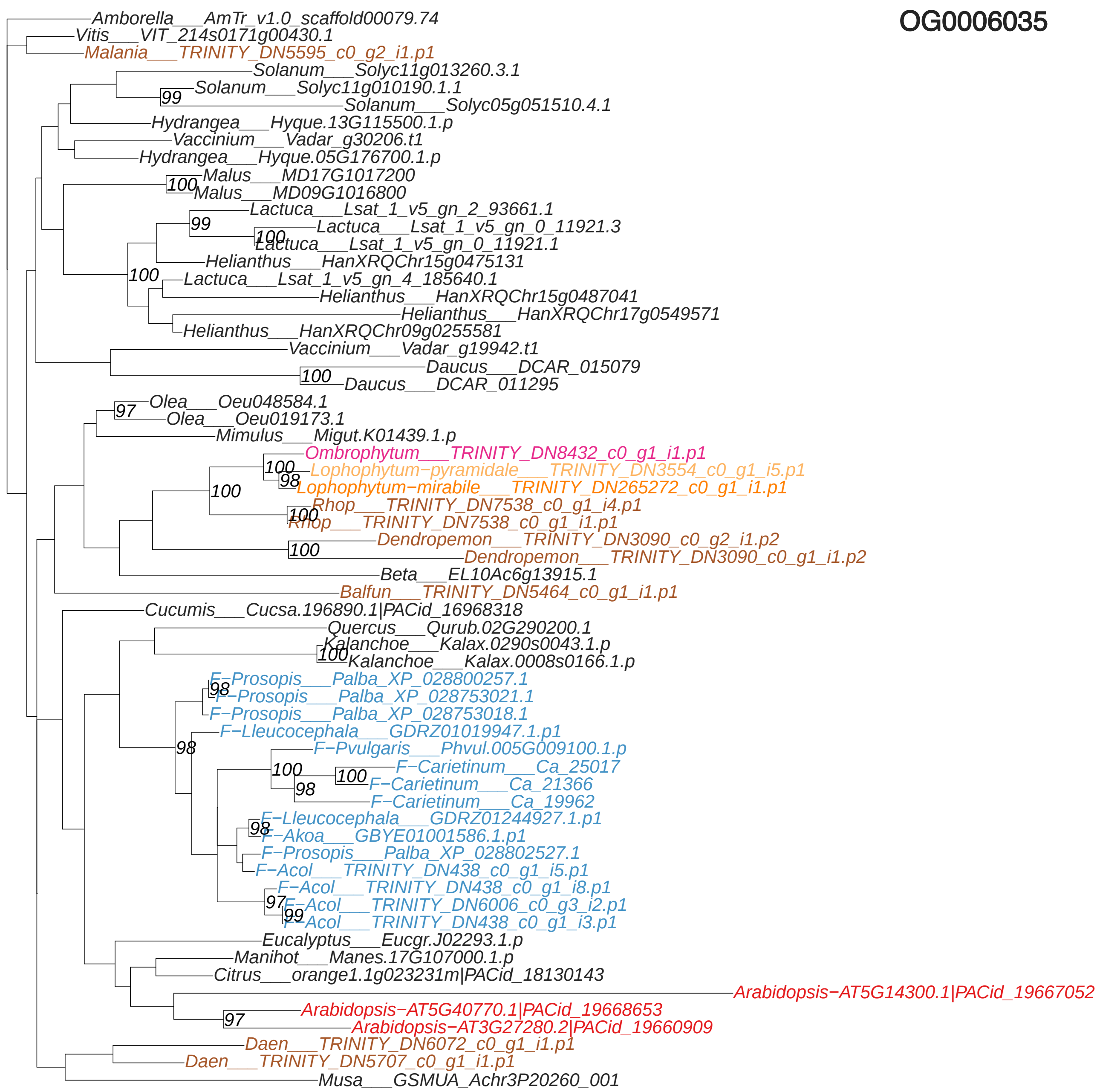
OG0006035
Amborella___AmTr_v1.0_scaffold00079.74
Vitis___VIT_214s0171g00430.1
Malania___TRINITY_DN5595_c0_g2_i1.p1
Solanum___Solyc11g013260.3.1
Solanum___Solyc11g010190.1.1
Solanum___Solyc05g051510.4.1
Hydrangea___Hyque.13G115500.1.p
Vaccinium___Vadar_g30206.t1
Hydrangea___Hyque.05G176700.1.p
Malus___MD17G1017200
Malus___MD09G1016800
Lactuca___Lsat_1_v5_gn_2_93661.1
Lactuca___Lsat_1_v5_gn_0_11921.3
Lactuca___Lsat_1_v5_gn_0_11921.1
Helianthus___HanXRQChr15g0475131
Lactuca___Lsat_1_v5_gn_4_185640.1
Helianthus___HanXRQChr15g0487041
Helianthus___HanXRQChr17g0549571
Helianthus___HanXRQChr09g0255581
Vaccinium___Vadar_g19942.t1
Daucus___DCAR_015079
Daucus___DCAR_011295
Olea___Oeu048584.1
Olea___Oeu019173.1
Mimulus___Migut.K01439.1.p
Ombrophytum___TRINITY_DN8432_c0_g1_i1.p1
Lophophytum−pyramidale___TRINITY_DN3554_c0_g1_i5.p1
Lophophytum−mirabile___TRINITY_DN265272_c0_g1_i1.p1
Rhop___TRINITY_DN7538_c0_g1_i4.p1
Rhop___TRINITY_DN7538_c0_g1_i1.p1
Dendropemon___TRINITY_DN3090_c0_g2_i1.p2
Dendropemon___TRINITY_DN3090_c0_g1_i1.p2
Beta___EL10Ac6g13915.1
Balfun___TRINITY_DN5464_c0_g1_i1.p1
Cucumis___Cucsa.196890.1|PACid_16968318
Quercus___Qurub.02G290200.1
Kalanchoe___Kalax.0290s0043.1.p
Kalanchoe___Kalax.0008s0166.1.p
F−Prosopis___Palba_XP_028800257.1
F−Prosopis___Palba_XP_028753021.1
F−Prosopis___Palba_XP_028753018.1
F−Lleucocephala___GDRZ01019947.1.p1
F−Pvulgaris___Phvul.005G009100.1.p
F−Carietinum___Ca_25017
F−Carietinum___Ca_21366
F−Carietinum___Ca_19962
F−Lleucocephala___GDRZ01244927.1.p1
F−Akoa___GBYE01001586.1.p1
F−Prosopis___Palba_XP_028802527.1
F−Acol___TRINITY_DN438_c0_g1_i5.p1
F−Acol___TRINITY_DN438_c0_g1_i8.p1
F−Acol___TRINITY_DN6006_c0_g3_i2.p1
F−Acol___TRINITY_DN438_c0_g1_i3.p1
Eucalyptus___Eucgr.J02293.1.p
Manihot___Manes.17G107000.1.p
Citrus___orange1.1g023231m|PACid_18130143
Arabidopsis−AT5G14300.1|PACid_19667052
Arabidopsis−AT5G40770.1|PACid_19668653
Arabidopsis−AT3G27280.2|PACid_19660909
Daen___TRINITY_DN6072_c0_g1_i1.p1
Daen___TRINITY_DN5707_c0_g1_i1.p1
Musa___GSMUA_Achr3P20260_001
99
100
99
100
100
100
97
100
98
100
100
100
100
98
98
100
98
100
98
97
99
97

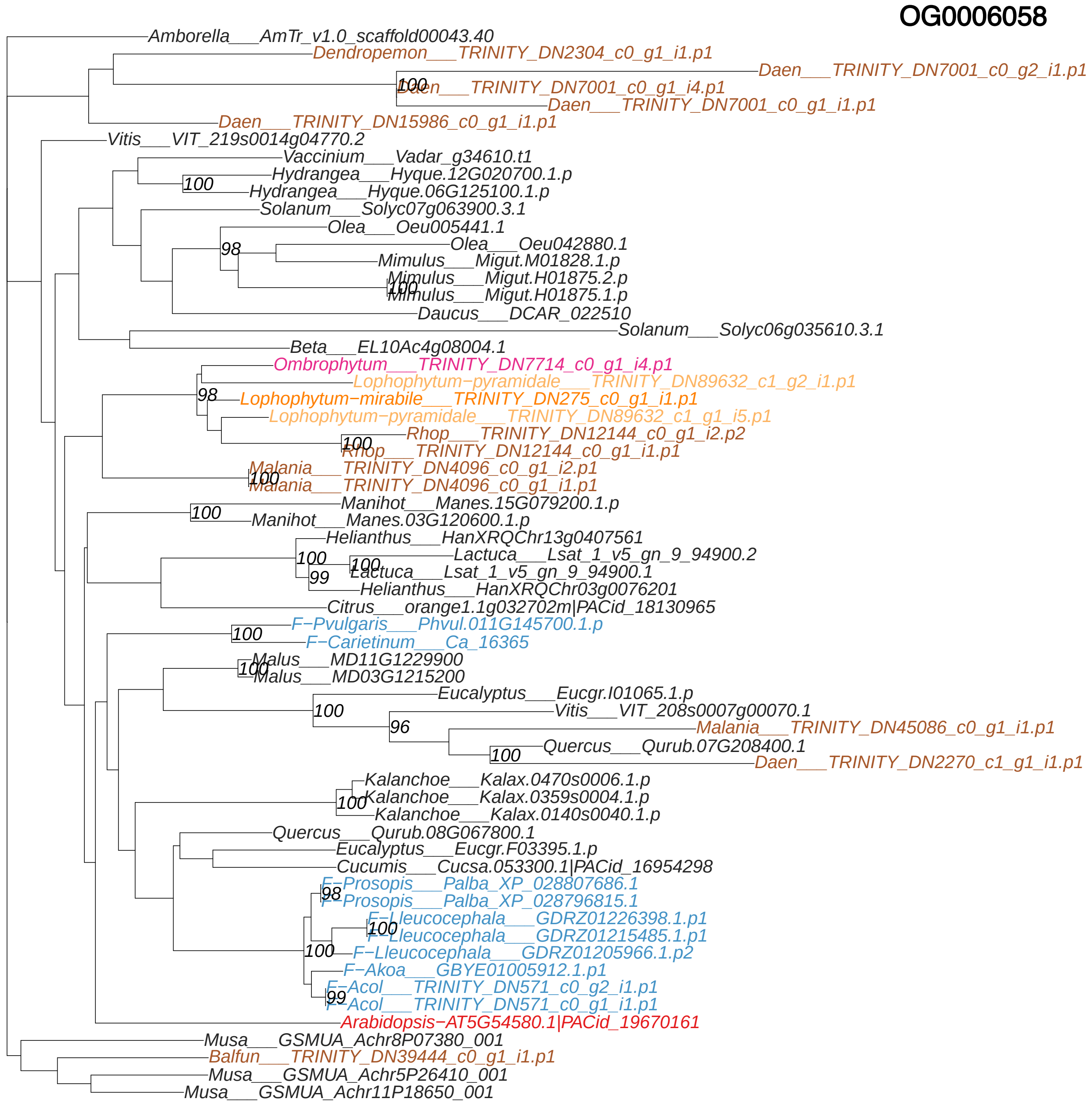
OG0006058
Amborella___AmTr_v1.0_scaffold00043.40
Dendropemon___TRINITY_DN2304_c0_g1_i1.p1
Daen___TRINITY_DN7001_c0_g2_i1.p1
100
Daen___TRINITY_DN7001_c0_g1_i4.p1
Daen___TRINITY_DN7001_c0_g1_i1.p1
Daen___TRINITY_DN15986_c0_g1_i1.p1
Vitis___VIT_219s0014g04770.2
Vaccinium___Vadar_g34610.t1
Hydrangea___Hyque.12G020700.1.p
100
Hydrangea___Hyque.06G125100.1.p
Solanum___Solyc07g063900.3.1
Olea___Oeu005441.1
Olea___Oeu042880.1
98
Mimulus___Migut.M01828.1.p
Mimulus___Migut.H01875.2.p
100
Mimulus___Migut.H01875.1.p
Daucus___DCAR_022510
Solanum___Solyc06g035610.3.1
Beta___EL10Ac4g08004.1
Ombrophytum___TRINITY_DN7714_c0_g1_i4.p1
Lophophytum−pyramidale___TRINITY_DN89632_c1_g2_i1.p1
98
Lophophytum−mirabile___TRINITY_DN275_c0_g1_i1.p1
Lophophytum−pyramidale___TRINITY_DN89632_c1_g1_i5.p1
Rhop___TRINITY_DN12144_c0_g1_i2.p2
100
Rhop___TRINITY_DN12144_c0_g1_i1.p1
Malania___TRINITY_DN4096_c0_g1_i2.p1
100
Malania___TRINITY_DN4096_c0_g1_i1.p1
Manihot___Manes.15G079200.1.p
100
Manihot___Manes.03G120600.1.p
Helianthus___HanXRQChr13g0407561
100
Lactuca___Lsat_1_v5_gn_9_94900.2
100
Lactuca___Lsat_1_v5_gn_9_94900.1
99
Helianthus___HanXRQChr03g0076201
Citrus___orange1.1g032702m|PACid_18130965
F−Pvulgaris___Phvul.011G145700.1.p
100
F−Carietinum___Ca_16365
Malus___MD11G1229900
100
Malus___MD03G1215200
Eucalyptus___Eucgr.I01065.1.p
100
Vitis___VIT_208s0007g00070.1
96
Malania___TRINITY_DN45086_c0_g1_i1.p1
Quercus___Qurub.07G208400.1
100
Daen___TRINITY_DN2270_c1_g1_i1.p1
Kalanchoe___Kalax.0470s0006.1.p
Kalanchoe___Kalax.0359s0004.1.p
100
Kalanchoe___Kalax.0140s0040.1.p
Quercus___Qurub.08G067800.1
Eucalyptus___Eucgr.F03395.1.p
Cucumis___Cucsa.053300.1|PACid_16954298
F−Prosopis___Palba_XP_028807686.1
98
F−Prosopis___Palba_XP_028796815.1
F−Lleucocephala___GDRZ01226398.1.p1
100
F−Lleucocephala___GDRZ01215485.1.p1
100
F−Lleucocephala___GDRZ01205966.1.p2
F−Akoa___GBYE01005912.1.p1
F−Acol___TRINITY_DN571_c0_g2_i1.p1
99
F−Acol___TRINITY_DN571_c0_g1_i1.p1
Arabidopsis−AT5G54580.1|PACid_19670161
Musa___GSMUA_Achr8P07380_001
Balfun___TRINITY_DN39444_c0_g1_i1.p1
Musa___GSMUA_Achr5P26410_001
Musa___GSMUA_Achr11P18650_001

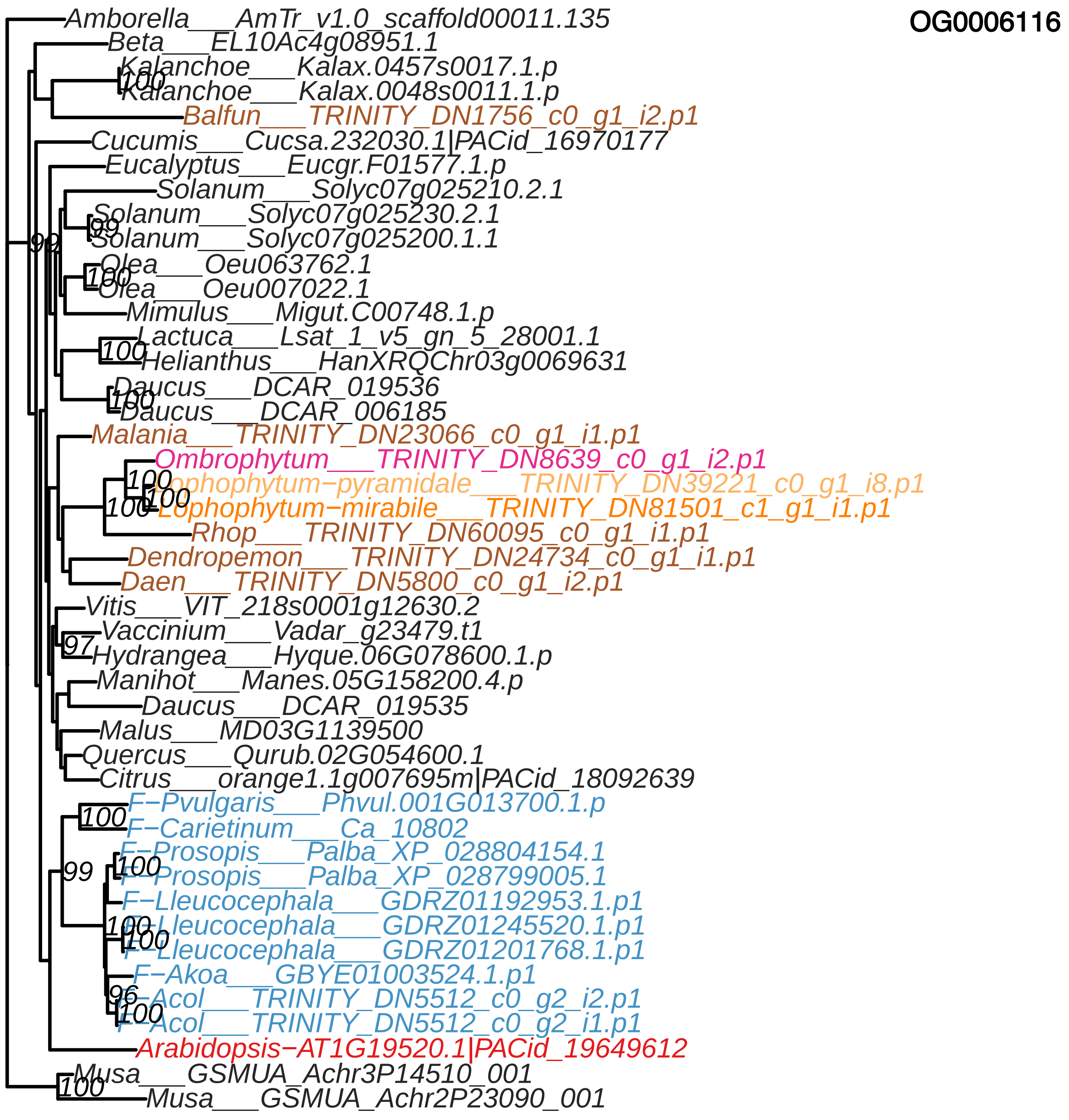
OG0006116
Amborella___AmTr_v1.0_scaffold00011.135
Beta___EL10Ac4g08951.1
Kalanchoe___Kalax.0457s0017.1.p
100
Kalanchoe___Kalax.0048s0011.1.p
Balfun___TRINITY_DN1756_c0_g1_i2.p1
Cucumis___Cucsa.232030.1|PACid_16970177
Eucalyptus___Eucgr.F01577.1.p
Solanum___Solyc07g025210.2.1
Solanum___Solyc07g025230.2.1
99
Solanum___Solyc07g025200.1.1
99
Olea___Oeu063762.1
100
Olea___Oeu007022.1
Mimulus___Migut.C00748.1.p
Lactuca___Lsat_1_v5_gn_5_28001.1
100
Helianthus___HanXRQChr03g0069631
Daucus___DCAR_019536
100
Daucus___DCAR_006185
Malania___TRINITY_DN23066_c0_g1_i1.p1
Ombrophytum___TRINITY_DN8639_c0_g1_i2.p1
100
Lophophytum−pyramidale___TRINITY_DN39221_c0_g1_i8.p1
100
100
Lophophytum−mirabile___TRINITY_DN81501_c1_g1_i1.p1
Rhop___TRINITY_DN60095_c0_g1_i1.p1
Dendropemon___TRINITY_DN24734_c0_g1_i1.p1
Daen___TRINITY_DN5800_c0_g1_i2.p1
Vitis___VIT_218s0001g12630.2
Vaccinium___Vadar_g23479.t1
97
Hydrangea___Hyque.06G078600.1.p
Manihot___Manes.05G158200.4.p
Daucus___DCAR_019535
Malus___MD03G1139500
Quercus___Qurub.02G054600.1
Citrus___orange1.1g007695m|PACid_18092639
F−Pvulgaris___Phvul.001G013700.1.p
100
F−Carietinum___Ca_10802
F−Prosopis___Palba_XP_028804154.1
100
99
F−Prosopis___Palba_XP_028799005.1
F−Lleucocephala___GDRZ01192953.1.p1
100
F−Lleucocephala___GDRZ01245520.1.p1
100
F−Lleucocephala___GDRZ01201768.1.p1
F−Akoa___GBYE01003524.1.p1
96
F−Acol___TRINITY_DN5512_c0_g2_i2.p1
100
F−Acol___TRINITY_DN5512_c0_g2_i1.p1
Arabidopsis−AT1G19520.1|PACid_19649612
Musa___GSMUA_Achr3P14510_001
100
Musa___GSMUA_Achr2P23090_001

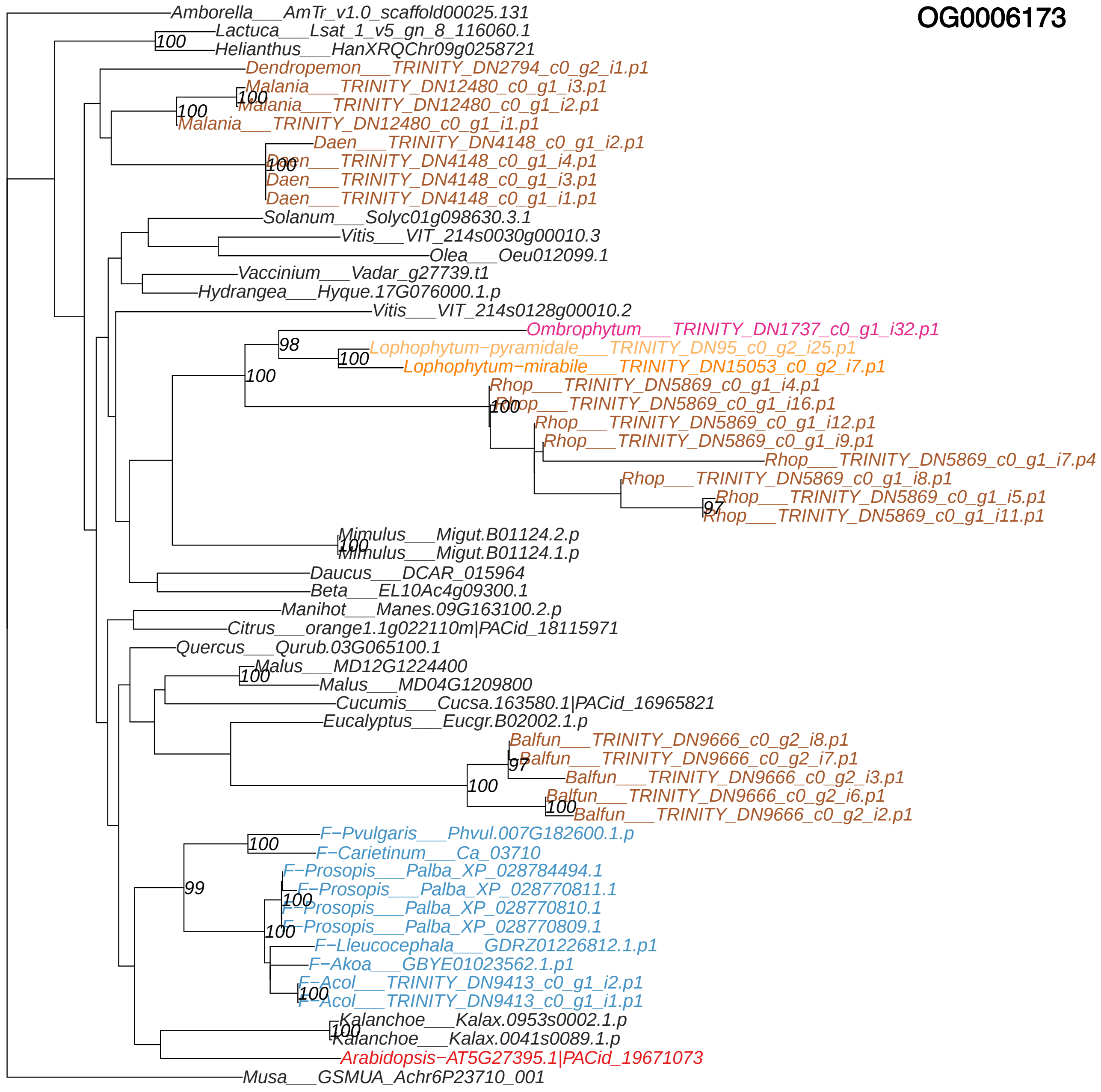

OG0006173
Amborella___AmTr_v1.0_scaffold00025.131
Lactuca___Lsat_1_v5_gn_8_116060.1
Helianthus___HanXRQChr09g0258721
Dendropemon___TRINITY_DN2794_c0_g2_i1.p1
Malania___TRINITY_DN12480_c0_g1_i3.p1
Malania___TRINITY_DN12480_c0_g1_i2.p1
Malania___TRINITY_DN12480_c0_g1_i1.p1
Daen___TRINITY_DN4148_c0_g1_i2.p1
Daen___TRINITY_DN4148_c0_g1_i4.p1
Daen___TRINITY_DN4148_c0_g1_i3.p1
Daen___TRINITY_DN4148_c0_g1_i1.p1
Solanum___Solyc01g098630.3.1
Vitis___VIT_214s0030g00010.3
Olea___Oeu012099.1
Vaccinium___Vadar_g27739.t1
Hydrangea___Hyque.17G076000.1.p
Vitis___VIT_214s0128g00010.2
Ombrophytum___TRINITY_DN1737_c0_g1_i32.p1
Lophophytum−pyramidale___TRINITY_DN95_c0_g2_i25.p1
Lophophytum−mirabile___TRINITY_DN15053_c0_g2_i7.p1
Rhop___TRINITY_DN5869_c0_g1_i4.p1
Rhop___TRINITY_DN5869_c0_g1_i16.p1
Rhop___TRINITY_DN5869_c0_g1_i12.p1
Rhop___TRINITY_DN5869_c0_g1_i9.p1
Rhop___TRINITY_DN5869_c0_g1_i7.p4
Rhop___TRINITY_DN5869_c0_g1_i8.p1
Rhop___TRINITY_DN5869_c0_g1_i5.p1
Rhop___TRINITY_DN5869_c0_g1_i11.p1
Mimulus___Migut.B01124.2.p
Mimulus___Migut.B01124.1.p
Daucus___DCAR_015964
Beta___EL10Ac4g09300.1
Manihot___Manes.09G163100.2.p
Citrus___orange1.1g022110m|PACid_18115971
Quercus___Qurub.03G065100.1
Malus___MD12G1224400
Malus___MD04G1209800
Cucumis___Cucsa.163580.1|PACid_16965821
Eucalyptus___Eucgr.B02002.1.p
Balfun___TRINITY_DN9666_c0_g2_i8.p1
Balfun___TRINITY_DN9666_c0_g2_i7.p1
Balfun___TRINITY_DN9666_c0_g2_i3.p1
Balfun___TRINITY_DN9666_c0_g2_i6.p1
Balfun___TRINITY_DN9666_c0_g2_i2.p1
F−Pvulgaris___Phvul.007G182600.1.p
F−Carietinum___Ca_03710
F−Prosopis___Palba_XP_028784494.1
F−Prosopis___Palba_XP_028770811.1
F−Prosopis___Palba_XP_028770810.1
F−Prosopis___Palba_XP_028770809.1
F−Lleucocephala___GDRZ01226812.1.p1
F−Akoa___GBYE01023562.1.p1
F−Acol___TRINITY_DN9413_c0_g1_i2.p1
F−Acol___TRINITY_DN9413_c0_g1_i1.p1
Kalanchoe___Kalax.0953s0002.1.p
Kalanchoe___Kalax.0041s0089.1.p
Arabidopsis−AT5G27395.1|PACid_19671073
Musa___GSMUA_Achr6P23710_001
100
100
100
100
98
100
100
100
97
100
100
97
100
100
100
99
100
100
100
100

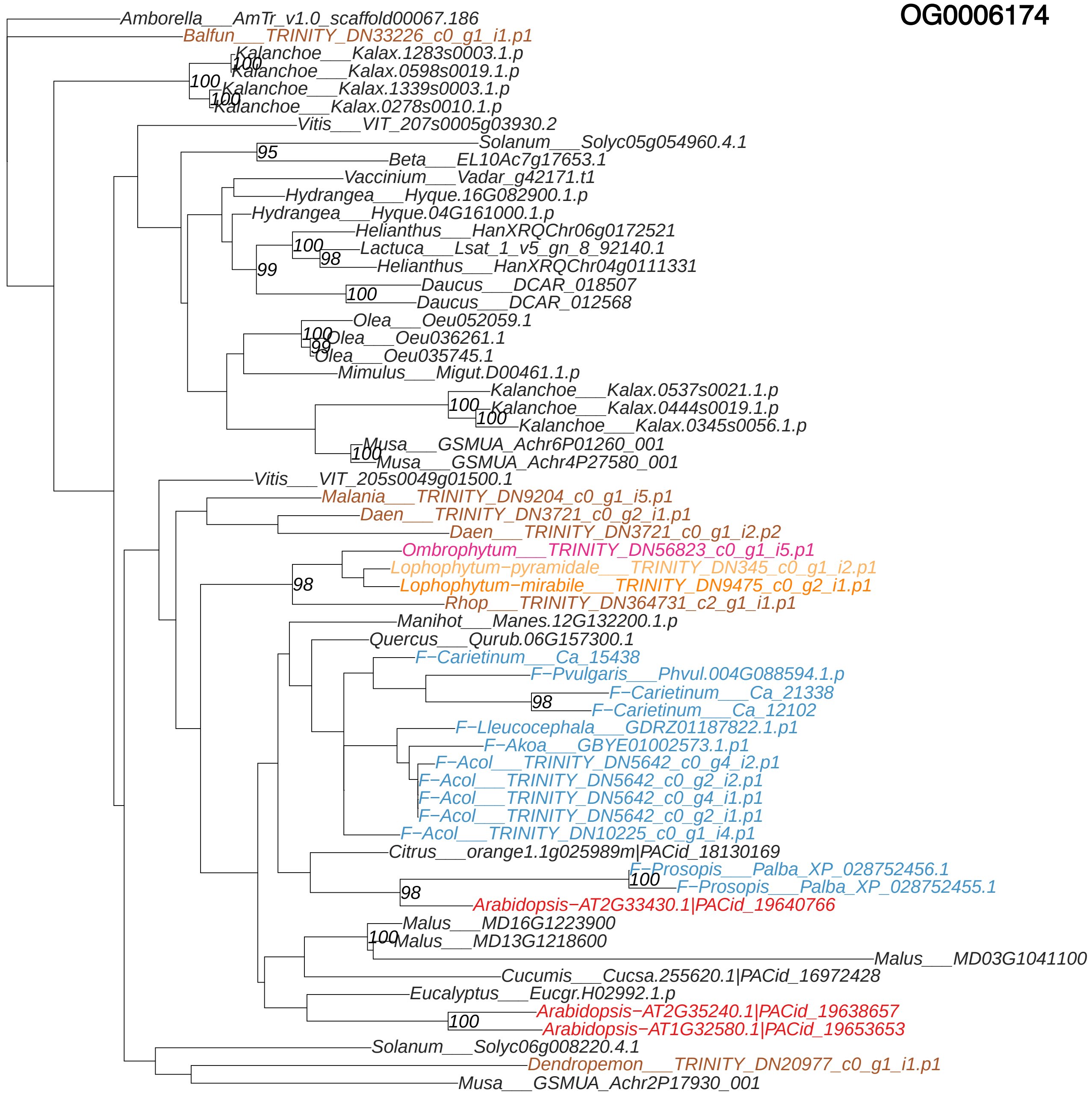
OG0006174
Amborella___AmTr_v1.0_scaffold00067.186
Balfun___TRINITY_DN33226_c0_g1_i1.p1
Kalanchoe___Kalax.1283s0003.1.p
Kalanchoe___Kalax.0598s0019.1.p
Kalanchoe___Kalax.1339s0003.1.p
Kalanchoe___Kalax.0278s0010.1.p
Vitis___VIT_207s0005g03930.2
Solanum___Solyc05g054960.4.1
Beta___EL10Ac7g17653.1
Vaccinium___Vadar_g42171.t1
Hydrangea___Hyque.16G082900.1.p
Hydrangea___Hyque.04G161000.1.p
Helianthus___HanXRQChr06g0172521
Lactuca___Lsat_1_v5_gn_8_92140.1
Helianthus___HanXRQChr04g0111331
Daucus___DCAR_018507
Daucus___DCAR_012568
Olea___Oeu052059.1
Olea___Oeu036261.1
Olea___Oeu035745.1
Mimulus___Migut.D00461.1.p
Kalanchoe___Kalax.0537s0021.1.p
Kalanchoe___Kalax.0444s0019.1.p
Kalanchoe___Kalax.0345s0056.1.p
Musa___GSMUA_Achr6P01260_001
Musa___GSMUA_Achr4P27580_001
Vitis___VIT_205s0049g01500.1
Malania___TRINITY_DN9204_c0_g1_i5.p1
Daen___TRINITY_DN3721_c0_g2_i1.p1
Daen___TRINITY_DN3721_c0_g1_i2.p2
Ombrophytum___TRINITY_DN56823_c0_g1_i5.p1
Lophophytum−pyramidale___TRINITY_DN345_c0_g1_i2.p1
Lophophytum−mirabile___TRINITY_DN9475_c0_g2_i1.p1
Rhop___TRINITY_DN364731_c2_g1_i1.p1
Manihot___Manes.12G132200.1.p
Quercus___Qurub.06G157300.1
F−Carietinum___Ca_15438
F−Pvulgaris___Phvul.004G088594.1.p
F−Carietinum___Ca_21338
F−Carietinum___Ca_12102
F−Lleucocephala___GDRZ01187822.1.p1
F−Akoa___GBYE01002573.1.p1
F−Acol___TRINITY_DN5642_c0_g4_i2.p1
F−Acol___TRINITY_DN5642_c0_g2_i2.p1
F−Acol___TRINITY_DN5642_c0_g4_i1.p1
F−Acol___TRINITY_DN5642_c0_g2_i1.p1
F−Acol___TRINITY_DN10225_c0_g1_i4.p1
Citrus___orange1.1g025989m|PACid_18130169
F−Prosopis___Palba_XP_028752456.1
F−Prosopis___Palba_XP_028752455.1
Arabidopsis−AT2G33430.1|PACid_19640766
Malus___MD16G1223900
Malus___MD13G1218600
Malus___MD03G1041100
Cucumis___Cucsa.255620.1|PACid_16972428
Eucalyptus___Eucgr.H02992.1.p
Arabidopsis−AT2G35240.1|PACid_19638657
Arabidopsis−AT1G32580.1|PACid_19653653
Solanum___Solyc06g008220.4.1
Dendropemon___TRINITY_DN20977_c0_g1_i1.p1
Musa___GSMUA_Achr2P17930_001

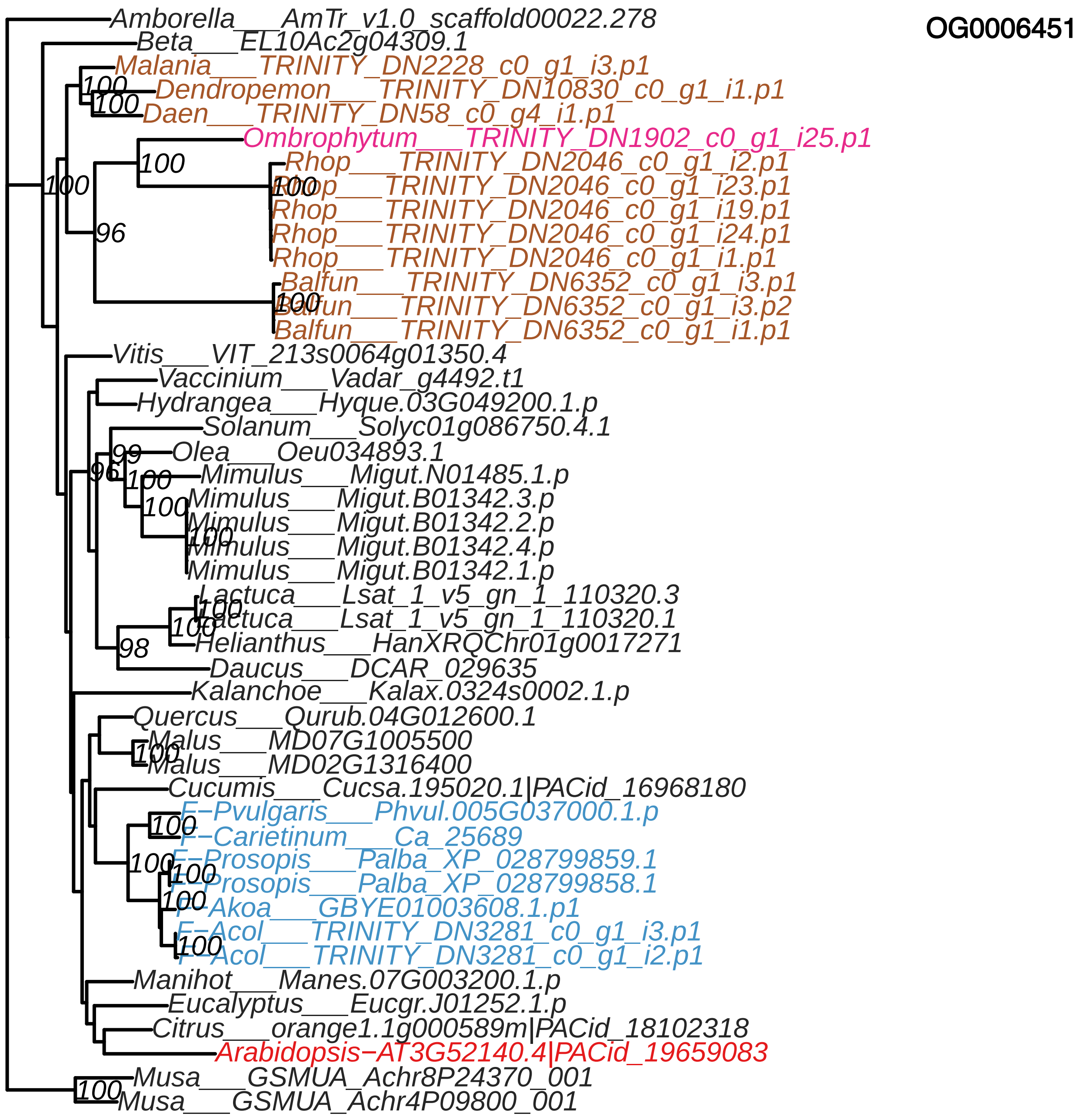
OG0006451
Amborella___AmTr_v1.0_scaffold00022.278
Beta___EL10Ac2g04309.1
Malania___TRINITY_DN2228_c0_g1_i3.p1
Dendropemon___TRINITY_DN10830_c0_g1_i1.p1
Daen___TRINITY_DN58_c0_g4_i1.p1
Ombrophytum___TRINITY_DN1902_c0_g1_i25.p1
Rhop___TRINITY_DN2046_c0_g1_i2.p1
Rhop___TRINITY_DN2046_c0_g1_i23.p1
Rhop___TRINITY_DN2046_c0_g1_i19.p1
Rhop___TRINITY_DN2046_c0_g1_i24.p1
Rhop___TRINITY_DN2046_c0_g1_i1.p1
Balfun___TRINITY_DN6352_c0_g1_i3.p1
Balfun___TRINITY_DN6352_c0_g1_i3.p2
Balfun___TRINITY_DN6352_c0_g1_i1.p1
Vitis___VIT_213s0064g01350.4
Vaccinium___Vadar_g4492.t1
Hydrangea___Hyque.03G049200.1.p
Solanum___Solyc01g086750.4.1
Olea___Oeu034893.1
Mimulus___Migut.N01485.1.p
Mimulus___Migut.B01342.3.p
Mimulus___Migut.B01342.2.p
Mimulus___Migut.B01342.4.p
Mimulus___Migut.B01342.1.p
Lactuca___Lsat_1_v5_gn_1_110320.3
Lactuca___Lsat_1_v5_gn_1_110320.1
Helianthus___HanXRQChr01g0017271
Daucus___DCAR_029635
Kalanchoe___Kalax.0324s0002.1.p
Quercus___Qurub.04G012600.1
Malus___MD07G1005500
Malus___MD02G1316400
Cucumis___Cucsa.195020.1|PACid_16968180
F−Pvulgaris___Phvul.005G037000.1.p
F−Carietinum___Ca_25689
F−Prosopis___Palba_XP_028799859.1
F−Prosopis___Palba_XP_028799858.1
F−Akoa___GBYE01003608.1.p1
F−Acol___TRINITY_DN3281_c0_g1_i3.p1
F−Acol___TRINITY_DN3281_c0_g1_i2.p1
Manihot___Manes.07G003200.1.p
Eucalyptus___Eucgr.J01252.1.p
Citrus___orange1.1g000589m|PACid_18102318
Arabidopsis−AT3G52140.4|PACid_19659083
Musa___GSMUA_Achr8P24370_001
Musa___GSMUA_Achr4P09800_001
100
100
100
100
96
100
100
99
96
100
100
100
100
100
98
100
100
100
100
100
100
100

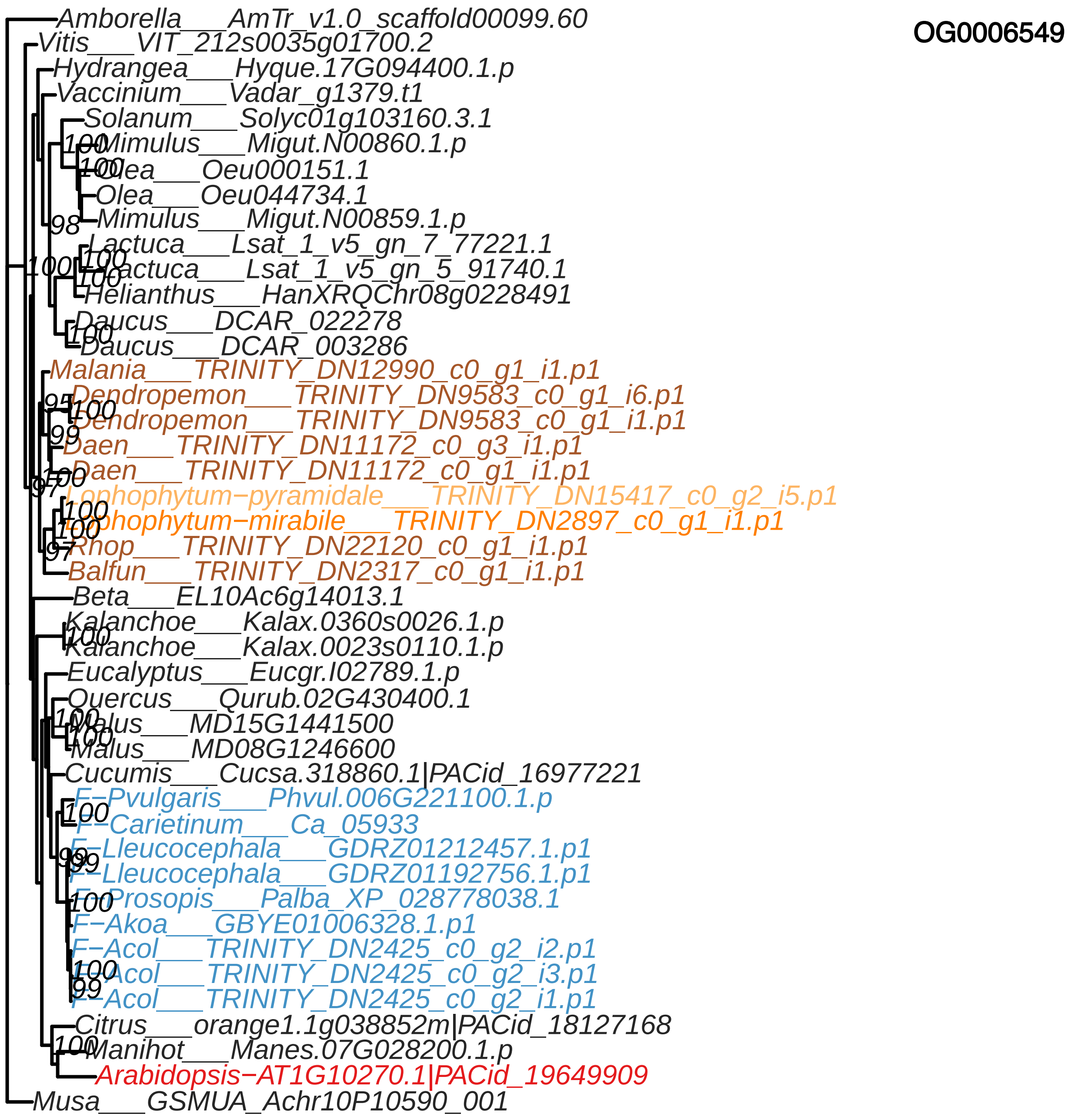
OG0006549
Amborella___AmTr_v1.0_scaffold00099.60
Vitis___VIT_212s0035g01700.2
Hydrangea___Hyque.17G094400.1.p
Vaccinium___Vadar_g1379.t1
Solanum___Solyc01g103160.3.1
Mimulus___Migut.N00860.1.p
Olea___Oeu000151.1
Olea___Oeu044734.1
Mimulus___Migut.N00859.1.p
Lactuca___Lsat_1_v5_gn_7_77221.1
Lactuca___Lsat_1_v5_gn_5_91740.1
Helianthus___HanXRQChr08g0228491
Daucus___DCAR_022278
Daucus___DCAR_003286
Malania___TRINITY_DN12990_c0_g1_i1.p1
Dendropemon___TRINITY_DN9583_c0_g1_i6.p1
Dendropemon___TRINITY_DN9583_c0_g1_i1.p1
Daen___TRINITY_DN11172_c0_g3_i1.p1
Daen___TRINITY_DN11172_c0_g1_i1.p1
Lophophytum−pyramidale___TRINITY_DN15417_c0_g2_i5.p1
Lophophytum−mirabile___TRINITY_DN2897_c0_g1_i1.p1
Rhop___TRINITY_DN22120_c0_g1_i1.p1
Balfun___TRINITY_DN2317_c0_g1_i1.p1
Beta___EL10Ac6g14013.1
Kalanchoe___Kalax.0360s0026.1.p
Kalanchoe___Kalax.0023s0110.1.p
Eucalyptus___Eucgr.I02789.1.p
Quercus___Qurub.02G430400.1
Malus___MD15G1441500
Malus___MD08G1246600
Cucumis___Cucsa.318860.1|PACid_16977221
F−Pvulgaris___Phvul.006G221100.1.p
F−Carietinum___Ca_05933
F−Lleucocephala___GDRZ01212457.1.p1
F−Lleucocephala___GDRZ01192756.1.p1
F−Prosopis___Palba_XP_028778038.1
F−Akoa___GBYE01006328.1.p1
F−Acol___TRINITY_DN2425_c0_g2_i2.p1
F−Acol___TRINITY_DN2425_c0_g2_i3.p1
F−Acol___TRINITY_DN2425_c0_g2_i1.p1
Citrus___orange1.1g038852m|PACid_18127168
Manihot___Manes.07G028200.1.p
Arabidopsis−AT1G10270.1|PACid_19649909
Musa___GSMUA_Achr10P10590_001
100
100
98
100
100
100
100
95
100
99
100
97
100
100
97
100
100
100
100
99
99
100
100
99
100

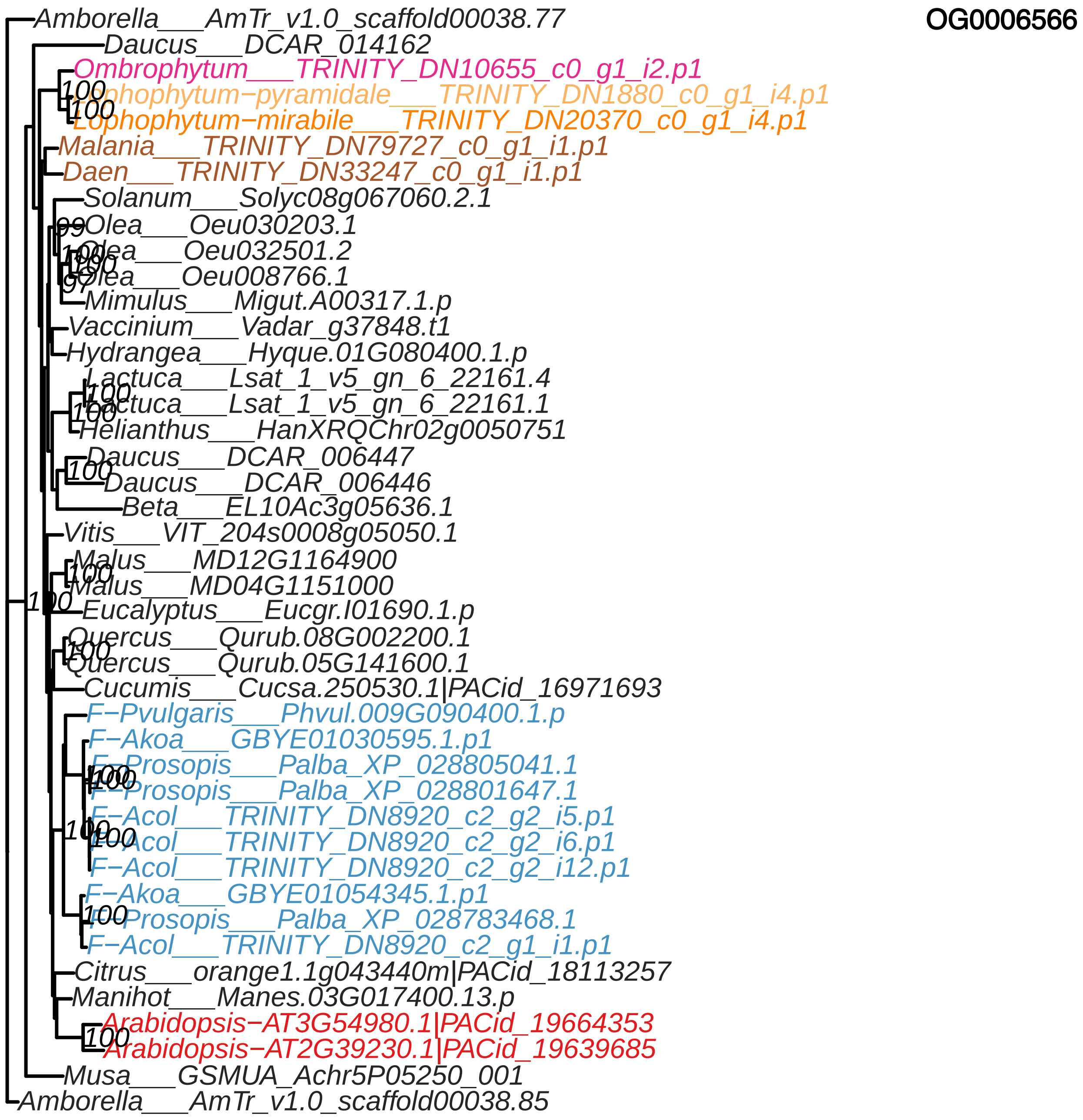
OG0006566
Amborella___AmTr_v1.0_scaffold00038.77
Daucus___DCAR_014162
Ombrophytum___TRINITY_DN10655_c0_g1_i2.p1
Lophophytum−pyramidale___TRINITY_DN1880_c0_g1_i4.p1
Lophophytum−mirabile___TRINITY_DN20370_c0_g1_i4.p1
Malania___TRINITY_DN79727_c0_g1_i1.p1
Daen___TRINITY_DN33247_c0_g1_i1.p1
Solanum___Solyc08g067060.2.1
Olea___Oeu030203.1
Olea___Oeu032501.2
Olea___Oeu008766.1
Mimulus___Migut.A00317.1.p
Vaccinium___Vadar_g37848.t1
Hydrangea___Hyque.01G080400.1.p
Lactuca___Lsat_1_v5_gn_6_22161.4
Lactuca___Lsat_1_v5_gn_6_22161.1
Helianthus___HanXRQChr02g0050751
Daucus___DCAR_006447
Daucus___DCAR_006446
Beta___EL10Ac3g05636.1
Vitis___VIT_204s0008g05050.1
Malus___MD12G1164900
Malus___MD04G1151000
Eucalyptus___Eucgr.I01690.1.p
Quercus___Qurub.08G002200.1
Quercus___Qurub.05G141600.1
Cucumis___Cucsa.250530.1|PACid_16971693
F−Pvulgaris___Phvul.009G090400.1.p
F−Akoa___GBYE01030595.1.p1
F−Prosopis___Palba_XP_028805041.1
F−Prosopis___Palba_XP_028801647.1
F−Acol___TRINITY_DN8920_c2_g2_i5.p1
F−Acol___TRINITY_DN8920_c2_g2_i6.p1
F−Acol___TRINITY_DN8920_c2_g2_i12.p1
F−Akoa___GBYE01054345.1.p1
F−Prosopis___Palba_XP_028783468.1
F−Acol___TRINITY_DN8920_c2_g1_i1.p1
Citrus___orange1.1g043440m|PACid_18113257
Manihot___Manes.03G017400.13.p
Arabidopsis−AT3G54980.1|PACid_19664353
Arabidopsis−AT2G39230.1|PACid_19639685
Musa___GSMUA_Achr5P05250_001
Amborella___AmTr_v1.0_scaffold00038.85
100
100
99
100
100
97
100
100
100
100
100
100
100
100
100
100
100

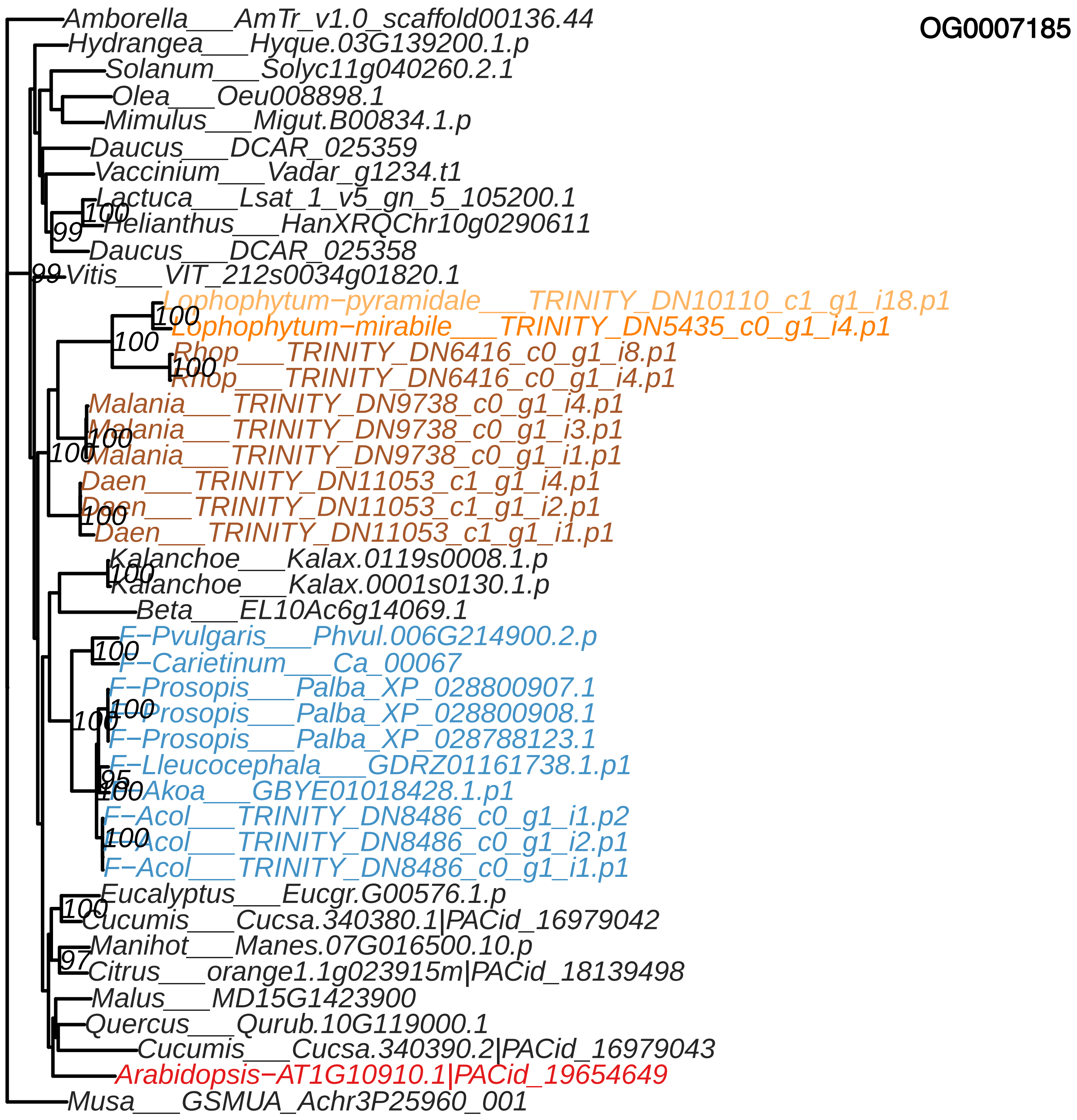

OG0007185
Amborella___AmTr_v1.0_scaffold00136.44
Hydrangea___Hyque.03G139200.1.p
Solanum___Solyc11g040260.2.1
Olea___Oeu008898.1
Mimulus___Migut.B00834.1.p
Daucus___DCAR_025359
Vaccinium___Vadar_g1234.t1
Lactuca___Lsat_1_v5_gn_5_105200.1
Helianthus___HanXRQChr10g0290611
Daucus___DCAR_025358
Vitis___VIT_212s0034g01820.1
Lophophytum−pyramidale___TRINITY_DN10110_c1_g1_i18.p1
Lophophytum−mirabile___TRINITY_DN5435_c0_g1_i4.p1
Rhop___TRINITY_DN6416_c0_g1_i8.p1
Rhop___TRINITY_DN6416_c0_g1_i4.p1
Malania___TRINITY_DN9738_c0_g1_i4.p1
Malania___TRINITY_DN9738_c0_g1_i3.p1
Malania___TRINITY_DN9738_c0_g1_i1.p1
Daen___TRINITY_DN11053_c1_g1_i4.p1
Daen___TRINITY_DN11053_c1_g1_i2.p1
Daen___TRINITY_DN11053_c1_g1_i1.p1
Kalanchoe___Kalax.0119s0008.1.p
Kalanchoe___Kalax.0001s0130.1.p
Beta___EL10Ac6g14069.1
F−Pvulgaris___Phvul.006G214900.2.p
F−Carietinum___Ca_00067
F−Prosopis___Palba_XP_028800907.1
F−Prosopis___Palba_XP_028800908.1
F−Prosopis___Palba_XP_028788123.1
F−Lleucocephala___GDRZ01161738.1.p1
F−Akoa___GBYE01018428.1.p1
F−Acol___TRINITY_DN8486_c0_g1_i1.p2
F−Acol___TRINITY_DN8486_c0_g1_i2.p1
F−Acol___TRINITY_DN8486_c0_g1_i1.p1
Eucalyptus___Eucgr.G00576.1.p
Cucumis___Cucsa.340380.1|PACid_16979042
Manihot___Manes.07G016500.10.p
Citrus___orange1.1g023915m|PACid_18139498
Malus___MD15G1423900
Quercus___Qurub.10G119000.1
Cucumis___Cucsa.340390.2|PACid_16979043
Arabidopsis−AT1G10910.1|PACid_19654649
Musa___GSMUA_Achr3P25960_001
100
99
99
100
100
100
100
100
100
100
100
100
100
95
100
100
100
97

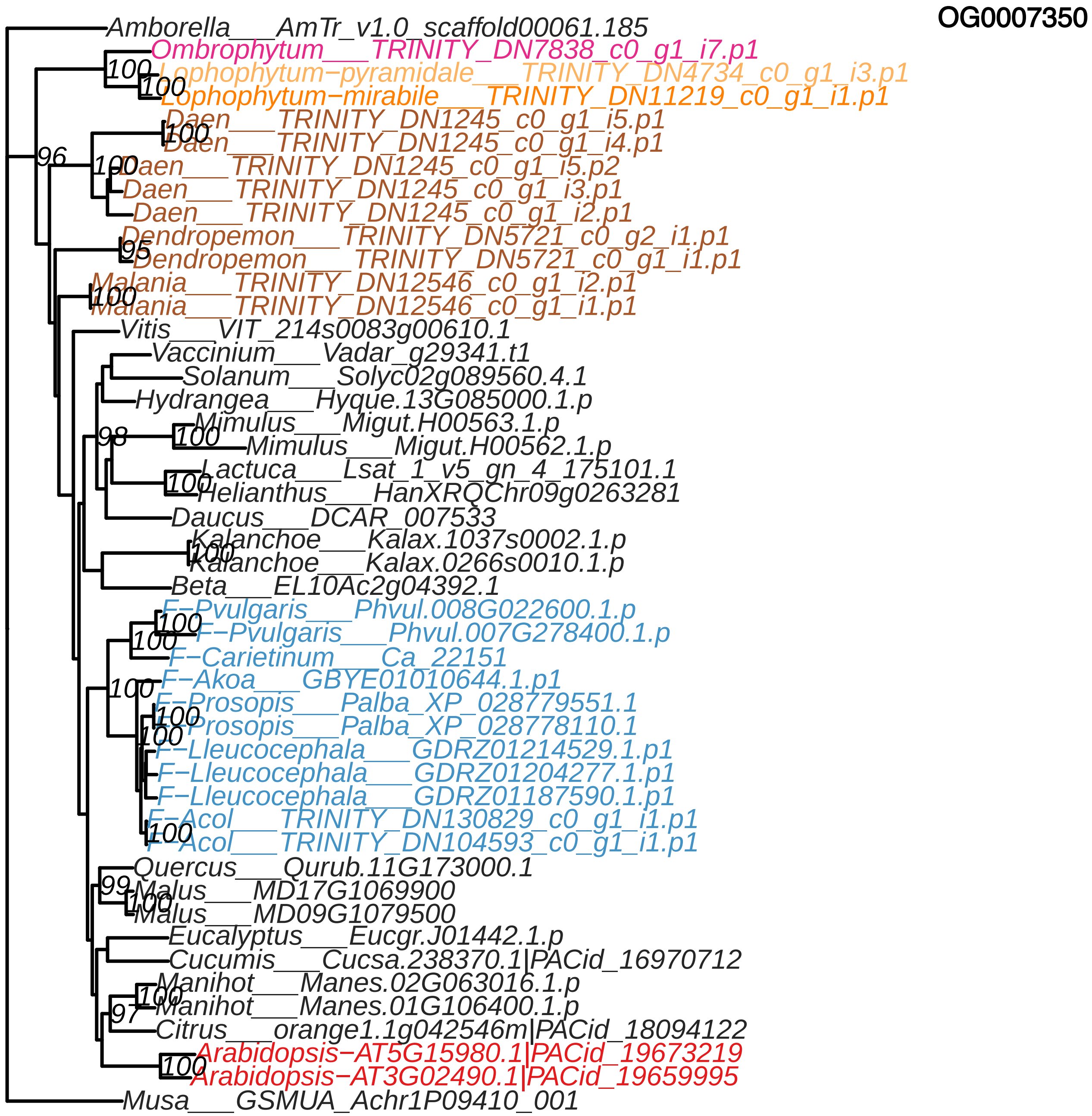
OG0007350
Amborella___AmTr_v1.0_scaffold00061.185
Ombrophytum___TRINITY_DN7838_c0_g1_i7.p1
Lophophytum−pyramidale___TRINITY_DN4734_c0_g1_i3.p1
Lophophytum−mirabile___TRINITY_DN11219_c0_g1_i1.p1
Daen___TRINITY_DN1245_c0_g1_i5.p1
Daen___TRINITY_DN1245_c0_g1_i4.p1
Daen___TRINITY_DN1245_c0_g1_i5.p2
Daen___TRINITY_DN1245_c0_g1_i3.p1
Daen___TRINITY_DN1245_c0_g1_i2.p1
Dendropemon___TRINITY_DN5721_c0_g2_i1.p1
Dendropemon___TRINITY_DN5721_c0_g1_i1.p1
Malania___TRINITY_DN12546_c0_g1_i2.p1
Malania___TRINITY_DN12546_c0_g1_i1.p1
Vitis___VIT_214s0083g00610.1
Vaccinium___Vadar_g29341.t1
Solanum___Solyc02g089560.4.1
Hydrangea___Hyque.13G085000.1.p
Mimulus___Migut.H00563.1.p
Mimulus___Migut.H00562.1.p
Lactuca___Lsat_1_v5_gn_4_175101.1
Helianthus___HanXRQChr09g0263281
Daucus___DCAR_007533
Kalanchoe___Kalax.1037s0002.1.p
Kalanchoe___Kalax.0266s0010.1.p
Beta___EL10Ac2g04392.1
F−Pvulgaris___Phvul.008G022600.1.p
F−Pvulgaris___Phvul.007G278400.1.p
F−Carietinum___Ca_22151
F−Akoa___GBYE01010644.1.p1
F−Prosopis___Palba_XP_028779551.1
F−Prosopis___Palba_XP_028778110.1
F−Lleucocephala___GDRZ01214529.1.p1
F−Lleucocephala___GDRZ01204277.1.p1
F−Lleucocephala___GDRZ01187590.1.p1
F−Acol___TRINITY_DN130829_c0_g1_i1.p1
F−Acol___TRINITY_DN104593_c0_g1_i1.p1
Quercus___Qurub.11G173000.1
Malus___MD17G1069900
Malus___MD09G1079500
Eucalyptus___Eucgr.J01442.1.p
Cucumis___Cucsa.238370.1|PACid_16970712
Manihot___Manes.02G063016.1.p
Manihot___Manes.01G106400.1.p
Citrus___orange1.1g042546m|PACid_18094122
Arabidopsis−AT5G15980.1|PACid_19673219
Arabidopsis−AT3G02490.1|PACid_19659995
Musa___GSMUA_Achr1P09410_001
100
100
100
96
100
95
100
98
100
100
100
100
100
100
100
100
100
99
100
100
97
100

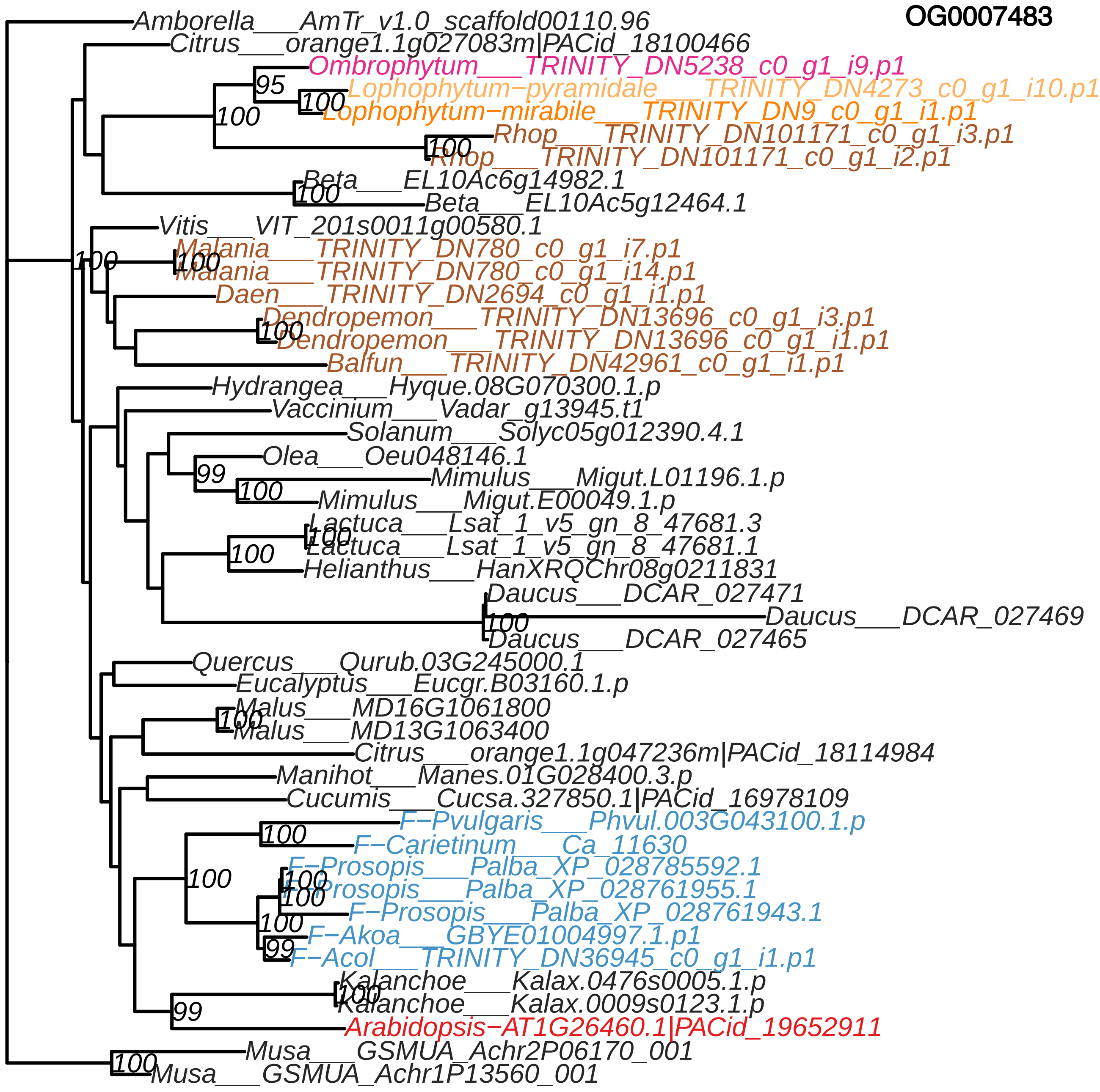
OG0007483
Amborella___AmTr_v1.0_scaffold00110.96
Citrus___orange1.1g027083m|PACid_18100466
Ombrophytum___TRINITY_DN5238_c0_g1_i9.p1
95
Lophophytum−pyramidale___TRINITY_DN4273_c0_g1_i10.p1
100
Lophophytum−mirabile___TRINITY_DN9_c0_g1_i1.p1
100
Rhop___TRINITY_DN101171_c0_g1_i3.p1
100
Rhop___TRINITY_DN101171_c0_g1_i2.p1
Beta___EL10Ac6g14982.1
100
Beta___EL10Ac5g12464.1
Vitis___VIT_201s0011g00580.1
Malania___TRINITY_DN780_c0_g1_i7.p1
100
100
Malania___TRINITY_DN780_c0_g1_i14.p1
Daen___TRINITY_DN2694_c0_g1_i1.p1
Dendropemon___TRINITY_DN13696_c0_g1_i3.p1
100
Dendropemon___TRINITY_DN13696_c0_g1_i1.p1
Balfun___TRINITY_DN42961_c0_g1_i1.p1
Hydrangea___Hyque.08G070300.1.p
Vaccinium___Vadar_g13945.t1
Solanum___Solyc05g012390.4.1
Olea___Oeu048146.1
99
Mimulus___Migut.L01196.1.p
100
Mimulus___Migut.E00049.1.p
Lactuca___Lsat_1_v5_gn_8_47681.3
100
Lactuca___Lsat_1_v5_gn_8_47681.1
100
Helianthus___HanXRQChr08g0211831
Daucus___DCAR_027471
Daucus___DCAR_027469
100
Daucus___DCAR_027465
Quercus___Qurub.03G245000.1
Eucalyptus___Eucgr.B03160.1.p
Malus___MD16G1061800
100
Malus___MD13G1063400
Citrus___orange1.1g047236m|PACid_18114984
Manihot___Manes.01G028400.3.p
Cucumis___Cucsa.327850.1|PACid_16978109
F−Pvulgaris___Phvul.003G043100.1.p
100
F−Carietinum___Ca_11630
F−Prosopis___Palba_XP_028785592.1
100
100
F−Prosopis___Palba_XP_028761955.1
100
F−Prosopis___Palba_XP_028761943.1
100
F−Akoa___GBYE01004997.1.p1
99
F−Acol___TRINITY_DN36945_c0_g1_i1.p1
Kalanchoe___Kalax.0476s0005.1.p
100
Kalanchoe___Kalax.0009s0123.1.p
99
Arabidopsis−AT1G26460.1|PACid_19652911
Musa___GSMUA_Achr2P06170_001
100
Musa___GSMUA_Achr1P13560_001

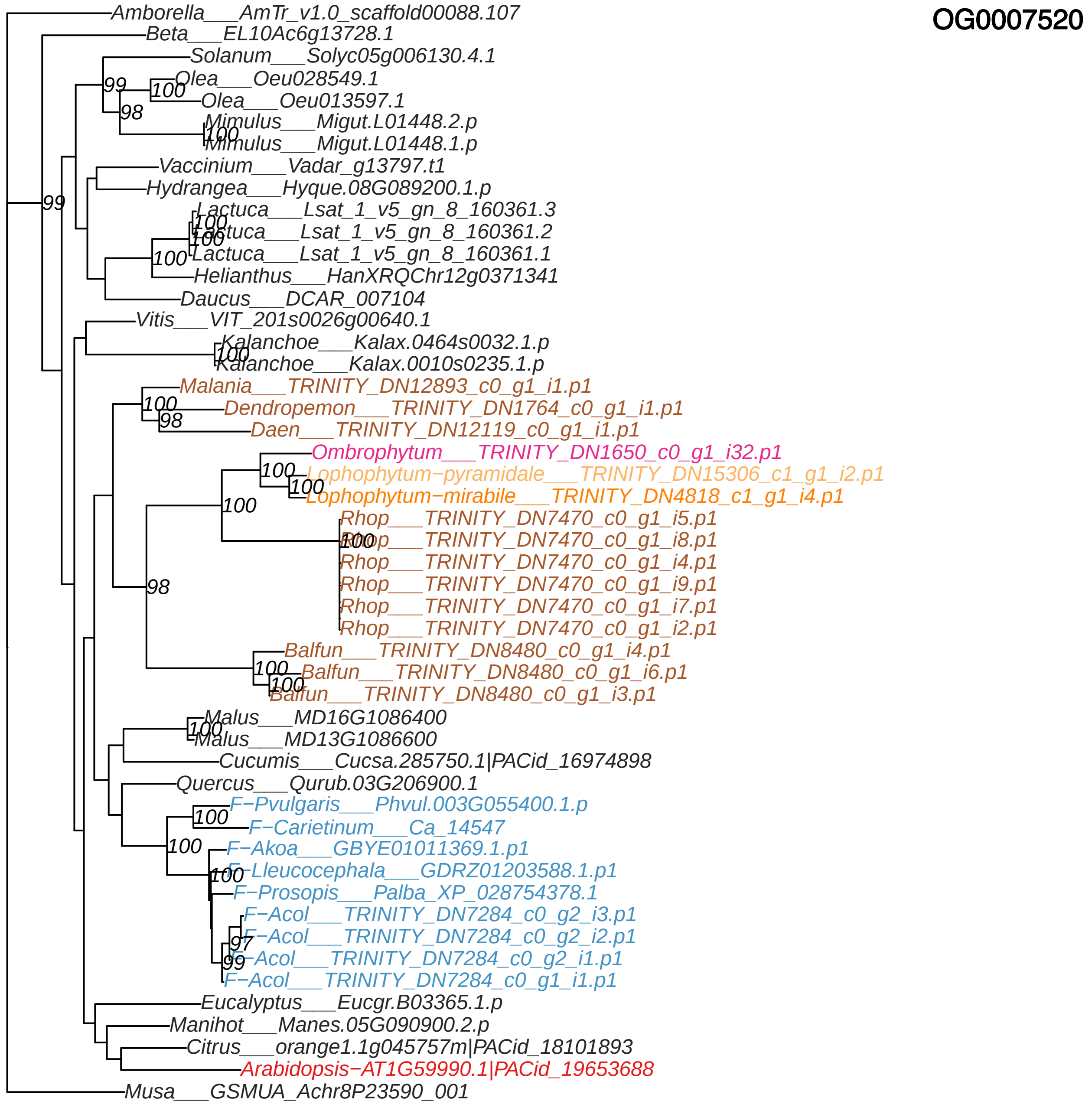
OG0007520
Amborella___AmTr_v1.0_scaffold00088.107
Beta___EL10Ac6g13728.1
Solanum___Solyc05g006130.4.1
Olea___Oeu028549.1
Olea___Oeu013597.1
Mimulus___Migut.L01448.2.p
Mimulus___Migut.L01448.1.p
Vaccinium___Vadar_g13797.t1
Hydrangea___Hyque.08G089200.1.p
Lactuca___Lsat_1_v5_gn_8_160361.3
Lactuca___Lsat_1_v5_gn_8_160361.2
Lactuca___Lsat_1_v5_gn_8_160361.1
Helianthus___HanXRQChr12g0371341
Daucus___DCAR_007104
Vitis___VIT_201s0026g00640.1
Kalanchoe___Kalax.0464s0032.1.p
Kalanchoe___Kalax.0010s0235.1.p
Malania___TRINITY_DN12893_c0_g1_i1.p1
Dendropemon___TRINITY_DN1764_c0_g1_i1.p1
Daen___TRINITY_DN12119_c0_g1_i1.p1
Ombrophytum___TRINITY_DN1650_c0_g1_i32.p1
Lophophytum−pyramidale___TRINITY_DN15306_c1_g1_i2.p1
Lophophytum−mirabile___TRINITY_DN4818_c1_g1_i4.p1
Rhop___TRINITY_DN7470_c0_g1_i5.p1
Rhop___TRINITY_DN7470_c0_g1_i8.p1
Rhop___TRINITY_DN7470_c0_g1_i4.p1
Rhop___TRINITY_DN7470_c0_g1_i9.p1
Rhop___TRINITY_DN7470_c0_g1_i7.p1
Rhop___TRINITY_DN7470_c0_g1_i2.p1
Balfun___TRINITY_DN8480_c0_g1_i4.p1
Balfun___TRINITY_DN8480_c0_g1_i6.p1
Balfun___TRINITY_DN8480_c0_g1_i3.p1
Malus___MD16G1086400
Malus___MD13G1086600
Cucumis___Cucsa.285750.1|PACid_16974898
Quercus___Qurub.03G206900.1
F−Pvulgaris___Phvul.003G055400.1.p
F−Carietinum___Ca_14547
F−Akoa___GBYE01011369.1.p1
F−Lleucocephala___GDRZ01203588.1.p1
F−Prosopis___Palba_XP_028754378.1
F−Acol___TRINITY_DN7284_c0_g2_i3.p1
F−Acol___TRINITY_DN7284_c0_g2_i2.p1
F−Acol___TRINITY_DN7284_c0_g2_i1.p1
F−Acol___TRINITY_DN7284_c0_g1_i1.p1
Eucalyptus___Eucgr.B03365.1.p
Manihot___Manes.05G090900.2.p
Citrus___orange1.1g045757m|PACid_18101893
Arabidopsis−AT1G59990.1|PACid_19653688
Musa___GSMUA_Achr8P23590_001
99
100
98
100
100
100
100
99
100
100
98
100
100
100
100
98
100
100
100
100
100
100
97
99

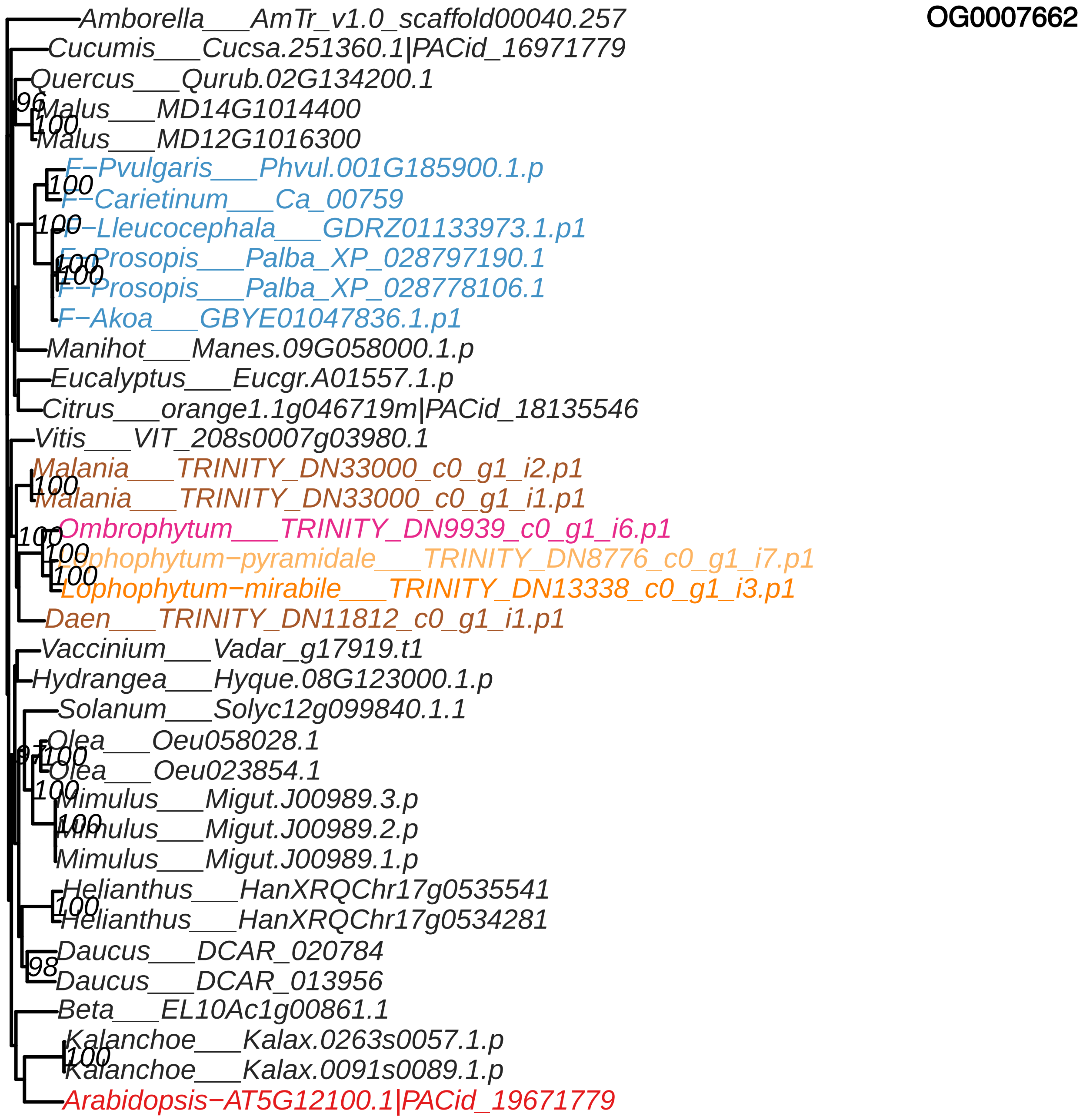
OG0007662
Amborella___AmTr_v1.0_scaffold00040.257
Cucumis___Cucsa.251360.1|PACid_16971779
Quercus___Qurub.02G134200.1
96
Malus___MD14G1014400
100
Malus___MD12G1016300
F−Pvulgaris___Phvul.001G185900.1.p
100
F−Carietinum___Ca_00759
100
F−Lleucocephala___GDRZ01133973.1.p1
100
F−Prosopis___Palba_XP_028797190.1
100
F−Prosopis___Palba_XP_028778106.1
F−Akoa___GBYE01047836.1.p1
Manihot___Manes.09G058000.1.p
Eucalyptus___Eucgr.A01557.1.p
Citrus___orange1.1g046719m|PACid_18135546
Vitis___VIT_208s0007g03980.1
Malania___TRINITY_DN33000_c0_g1_i2.p1
100
Malania___TRINITY_DN33000_c0_g1_i1.p1
100
Ombrophytum___TRINITY_DN9939_c0_g1_i6.p1
100
Lophophytum−pyramidale___TRINITY_DN8776_c0_g1_i7.p1
100
Lophophytum−mirabile___TRINITY_DN13338_c0_g1_i3.p1
Daen___TRINITY_DN11812_c0_g1_i1.p1
Vaccinium___Vadar_g17919.t1
Hydrangea___Hyque.08G123000.1.p
Solanum___Solyc12g099840.1.1
Olea___Oeu058028.1
97
100
Olea___Oeu023854.1
100
Mimulus___Migut.J00989.3.p
100
Mimulus___Migut.J00989.2.p
Mimulus___Migut.J00989.1.p
Helianthus___HanXRQChr17g0535541
100
Helianthus___HanXRQChr17g0534281
Daucus___DCAR_020784
98
Daucus___DCAR_013956
Beta___EL10Ac1g00861.1
Kalanchoe___Kalax.0263s0057.1.p
100
Kalanchoe___Kalax.0091s0089.1.p
Arabidopsis−AT5G12100.1|PACid_19671779

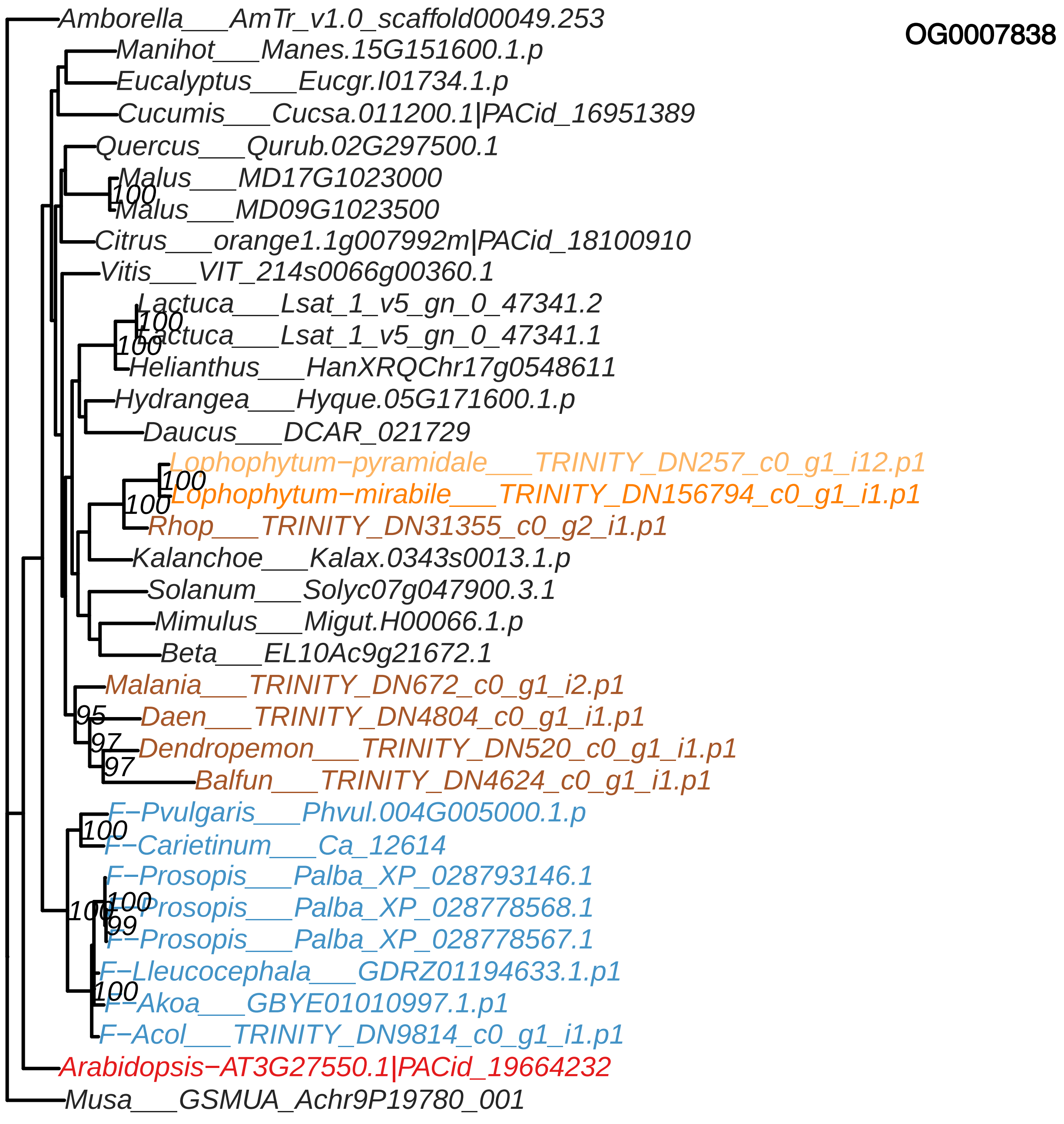

OG0007838
Amborella___AmTr_v1.0_scaffold00049.253
Manihot___Manes.15G151600.1.p
Eucalyptus___Eucgr.I01734.1.p
Cucumis___Cucsa.011200.1|PACid_16951389
Quercus___Qurub.02G297500.1
Malus___MD17G1023000
100
Malus___MD09G1023500
Citrus___orange1.1g007992m|PACid_18100910
Vitis___VIT_214s0066g00360.1
Lactuca___Lsat_1_v5_gn_0_47341.2
100
Lactuca___Lsat_1_v5_gn_0_47341.1
100
Helianthus___HanXRQChr17g0548611
Hydrangea___Hyque.05G171600.1.p
Daucus___DCAR_021729
Lophophytum−pyramidale___TRINITY_DN257_c0_g1_i12.p1
100
Lophophytum−mirabile___TRINITY_DN156794_c0_g1_i1.p1
100
Rhop___TRINITY_DN31355_c0_g2_i1.p1
Kalanchoe___Kalax.0343s0013.1.p
Solanum___Solyc07g047900.3.1
Mimulus___Migut.H00066.1.p
Beta___EL10Ac9g21672.1
Malania___TRINITY_DN672_c0_g1_i2.p1
95
Daen___TRINITY_DN4804_c0_g1_i1.p1
97
Dendropemon___TRINITY_DN520_c0_g1_i1.p1
97
Balfun___TRINITY_DN4624_c0_g1_i1.p1
F−Pvulgaris___Phvul.004G005000.1.p
100
F−Carietinum___Ca_12614
F−Prosopis___Palba_XP_028793146.1
100
F−Prosopis___Palba_XP_028778568.1
100
99
F−Prosopis___Palba_XP_028778567.1
F−Lleucocephala___GDRZ01194633.1.p1
100
F−Akoa___GBYE01010997.1.p1
F−Acol___TRINITY_DN9814_c0_g1_i1.p1
Arabidopsis−AT3G27550.1|PACid_19664232
Musa___GSMUA_Achr9P19780_001

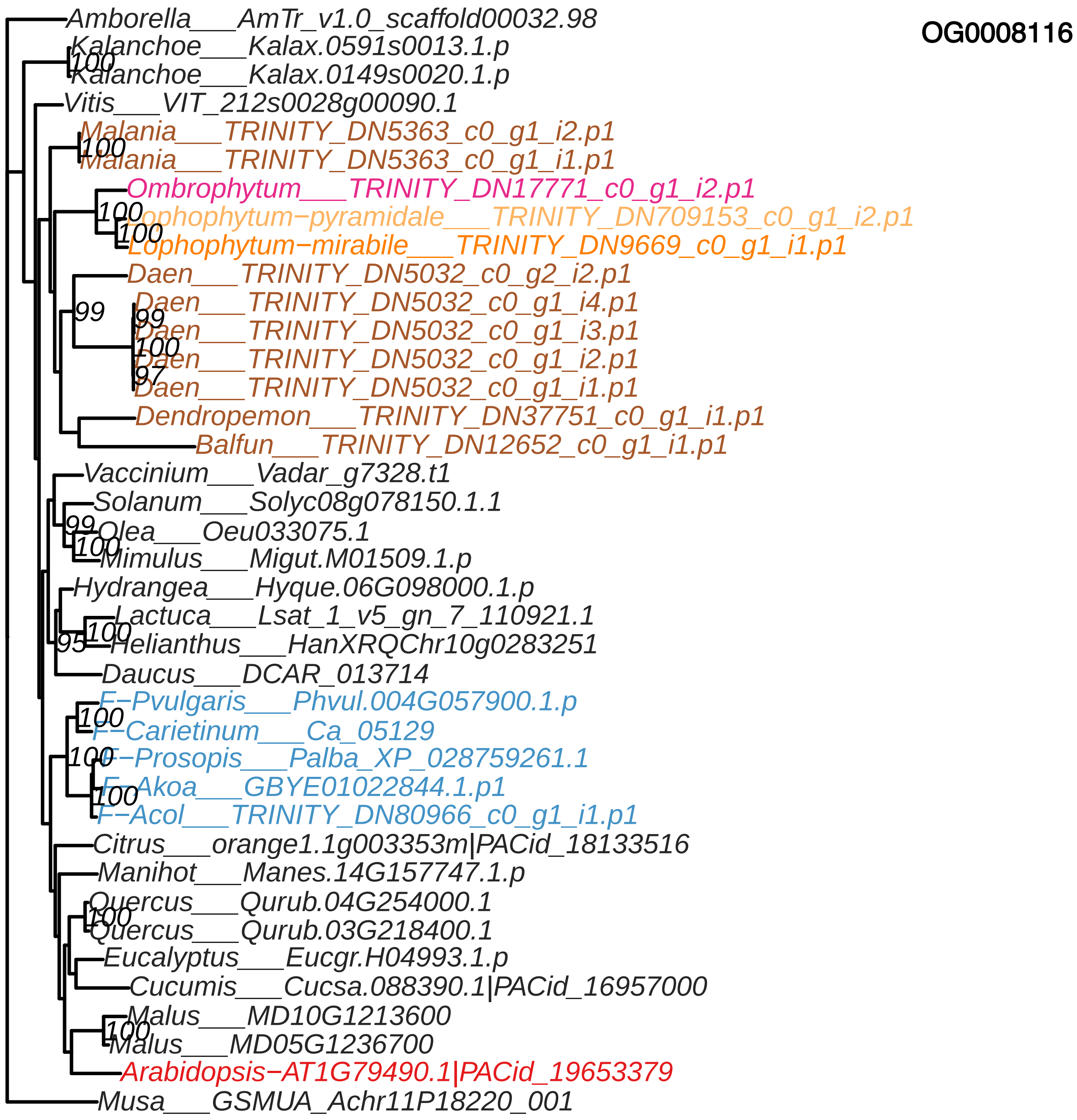
OG0008116
Amborella___AmTr_v1.0_scaffold00032.98
Kalanchoe___Kalax.0591s0013.1.p
100
Kalanchoe___Kalax.0149s0020.1.p
Vitis___VIT_212s0028g00090.1
Malania___TRINITY_DN5363_c0_g1_i2.p1
100
Malania___TRINITY_DN5363_c0_g1_i1.p1
Ombrophytum___TRINITY_DN17771_c0_g1_i2.p1
100
Lophophytum−pyramidale___TRINITY_DN709153_c0_g1_i2.p1
100
Lophophytum−mirabile___TRINITY_DN9669_c0_g1_i1.p1
Daen___TRINITY_DN5032_c0_g2_i2.p1
99
Daen___TRINITY_DN5032_c0_g1_i4.p1
99
Daen___TRINITY_DN5032_c0_g1_i3.p1
100
Daen___TRINITY_DN5032_c0_g1_i2.p1
97
Daen___TRINITY_DN5032_c0_g1_i1.p1
Dendropemon___TRINITY_DN37751_c0_g1_i1.p1
Balfun___TRINITY_DN12652_c0_g1_i1.p1
Vaccinium___Vadar_g7328.t1
Solanum___Solyc08g078150.1.1
99
Olea___Oeu033075.1
100
Mimulus___Migut.M01509.1.p
Hydrangea___Hyque.06G098000.1.p
Lactuca___Lsat_1_v5_gn_7_110921.1
100
95
Helianthus___HanXRQChr10g0283251
Daucus___DCAR_013714
F−Pvulgaris___Phvul.004G057900.1.p
100
F−Carietinum___Ca_05129
100
F−Prosopis___Palba_XP_028759261.1
F−Akoa___GBYE01022844.1.p1
100
F−Acol___TRINITY_DN80966_c0_g1_i1.p1
Citrus___orange1.1g003353m|PACid_18133516
Manihot___Manes.14G157747.1.p
Quercus___Qurub.04G254000.1
100
Quercus___Qurub.03G218400.1
Eucalyptus___Eucgr.H04993.1.p
Cucumis___Cucsa.088390.1|PACid_16957000
Malus___MD10G1213600
100
Malus___MD05G1236700
Arabidopsis−AT1G79490.1|PACid_19653379
Musa___GSMUA_Achr11P18220_001

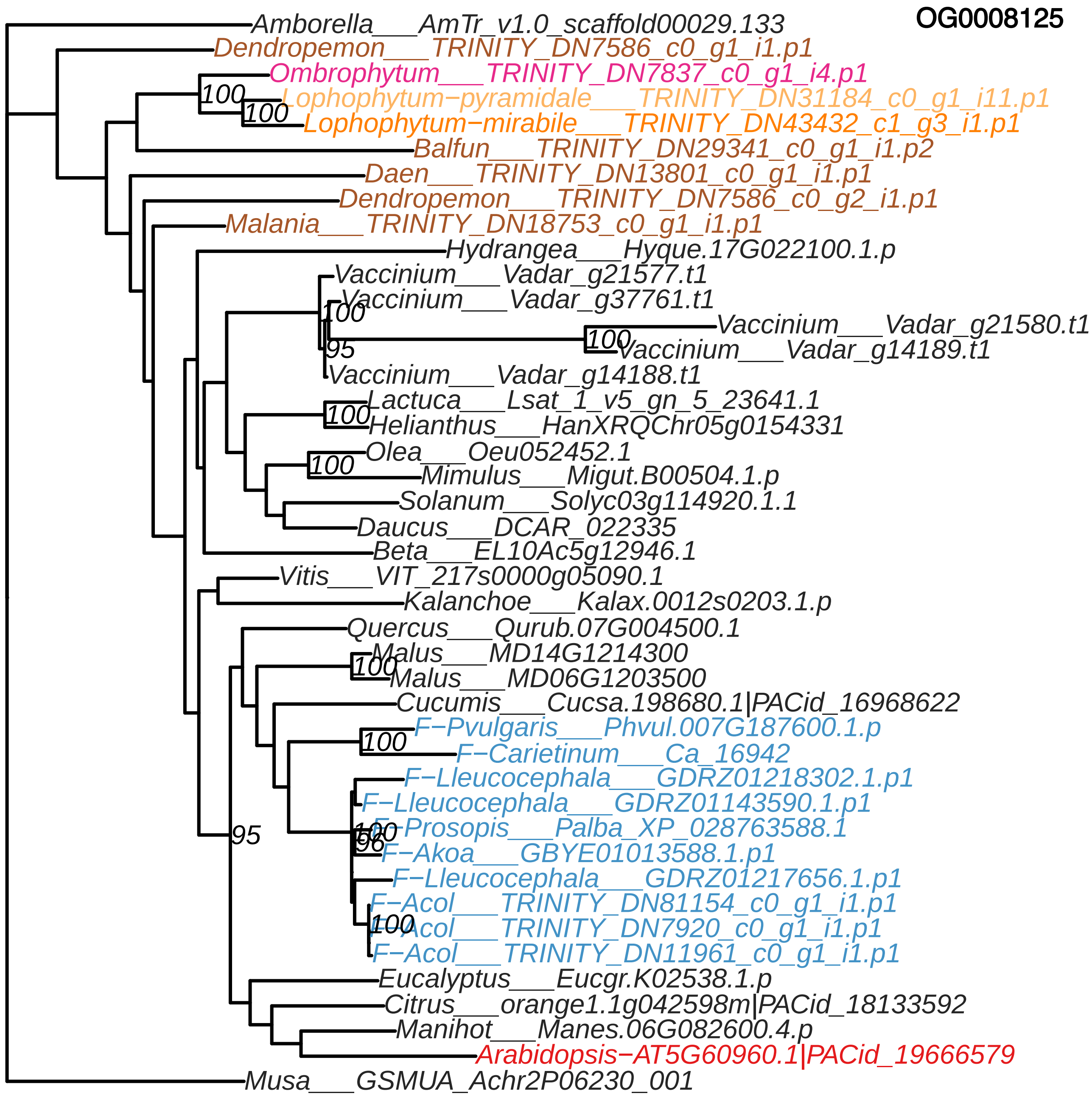
OG0008125
Amborella___AmTr_v1.0_scaffold00029.133
Dendropemon___TRINITY_DN7586_c0_g1_i1.p1
Ombrophytum___TRINITY_DN7837_c0_g1_i4.p1
Lophophytum−pyramidale___TRINITY_DN31184_c0_g1_i11.p1
Lophophytum−mirabile___TRINITY_DN43432_c1_g3_i1.p1
Balfun___TRINITY_DN29341_c0_g1_i1.p2
Daen___TRINITY_DN13801_c0_g1_i1.p1
Dendropemon___TRINITY_DN7586_c0_g2_i1.p1
Malania___TRINITY_DN18753_c0_g1_i1.p1
Hydrangea___Hyque.17G022100.1.p
Vaccinium___Vadar_g21577.t1
Vaccinium___Vadar_g37761.t1
Vaccinium___Vadar_g21580.t1
Vaccinium___Vadar_g14189.t1
Vaccinium___Vadar_g14188.t1
Lactuca___Lsat_1_v5_gn_5_23641.1
Helianthus___HanXRQChr05g0154331
Olea___Oeu052452.1
Mimulus___Migut.B00504.1.p
Solanum___Solyc03g114920.1.1
Daucus___DCAR_022335
Beta___EL10Ac5g12946.1
Vitis___VIT_217s0000g05090.1
Kalanchoe___Kalax.0012s0203.1.p
Quercus___Qurub.07G004500.1
Malus___MD14G1214300
Malus___MD06G1203500
Cucumis___Cucsa.198680.1|PACid_16968622
F−Pvulgaris___Phvul.007G187600.1.p
F−Carietinum___Ca_16942
F−Lleucocephala___GDRZ01218302.1.p1
F−Lleucocephala___GDRZ01143590.1.p1
F−Prosopis___Palba_XP_028763588.1
F−Akoa___GBYE01013588.1.p1
F−Lleucocephala___GDRZ01217656.1.p1
F−Acol___TRINITY_DN81154_c0_g1_i1.p1
F−Acol___TRINITY_DN7920_c0_g1_i1.p1
F−Acol___TRINITY_DN11961_c0_g1_i1.p1
Eucalyptus___Eucgr.K02538.1.p
Citrus___orange1.1g042598m|PACid_18133592
Manihot___Manes.06G082600.4.p
Arabidopsis−AT5G60960.1|PACid_19666579
Musa___GSMUA_Achr2P06230_001
100
100
100
95
100
100
100
100
100
95
100
96
100

OG0008417

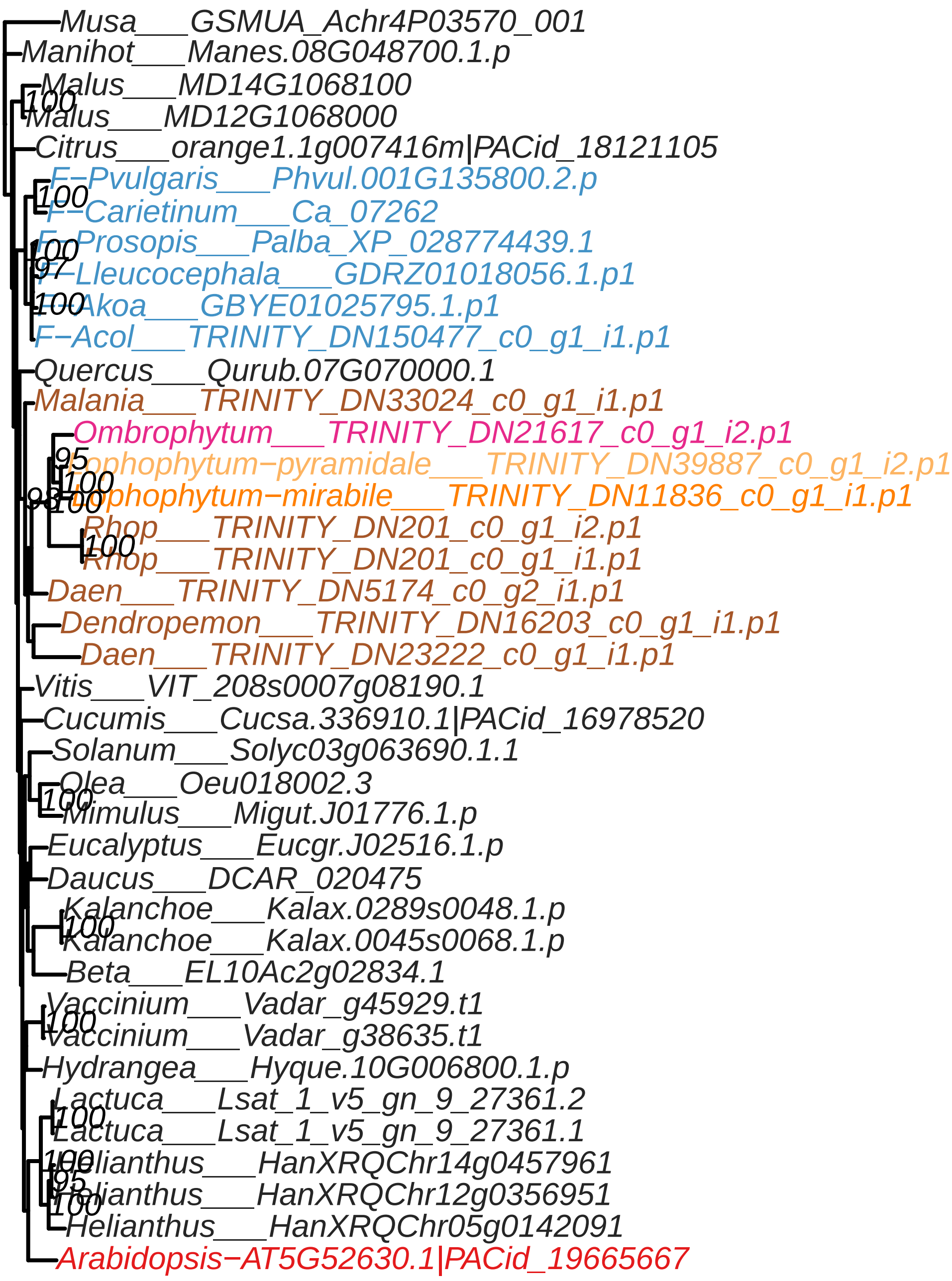

Musa___GSMUA_Achr4P03570_001
Manihot___Manes.08G048700.1.p
Malus___MD14G1068100
100
Malus___MD12G1068000
Citrus___orange1.1g007416m|PACid_18121105
F−Pvulgaris___Phvul.001G135800.2.p
100
F−Carietinum___Ca_07262
F−Prosopis___Palba_XP_028774439.1
100
97
F−Lleucocephala___GDRZ01018056.1.p1
100
F−Akoa___GBYE01025795.1.p1
F−Acol___TRINITY_DN150477_c0_g1_i1.p1
Quercus___Qurub.07G070000.1
Malania___TRINITY_DN33024_c0_g1_i1.p1
Ombrophytum___TRINITY_DN21617_c0_g1_i2.p1
95
ophophytum−pyramidale___TRINITY_DN39887_c0_g1_i2.p1
100
98
100
ophophytum−mirabile___TRINITY_DN11836_c0_g1_i1.p1
Rhop___TRINITY_DN201_c0_g1_i2.p1
100
Rhop___TRINITY_DN201_c0_g1_i1.p1
Daen___TRINITY_DN5174_c0_g2_i1.p1
Dendropemon___TRINITY_DN16203_c0_g1_i1.p1
Daen___TRINITY_DN23222_c0_g1_i1.p1
Vitis___VIT_208s0007g08190.1
Cucumis___Cucsa.336910.1|PACid_16978520
Solanum___Solyc03g063690.1.1
Olea___Oeu018002.3
100
Mimulus___Migut.J01776.1.p
Eucalyptus___Eucgr.J02516.1.p
Daucus___DCAR_020475
Kalanchoe___Kalax.0289s0048.1.p
100
Kalanchoe___Kalax.0045s0068.1.p
Beta___EL10Ac2g02834.1
Vaccinium___Vadar_g45929.t1
100
Vaccinium___Vadar_g38635.t1
Hydrangea___Hyque.10G006800.1.p
Lactuca___Lsat_1_v5_gn_9_27361.2
100
Lactuca___Lsat_1_v5_gn_9_27361.1
100
Helianthus___HanXRQChr14g0457961
95
Helianthus___HanXRQChr12g0356951
100
Helianthus___HanXRQChr05g0142091
Arabidopsis−AT5G52630.1|PACid_19665667

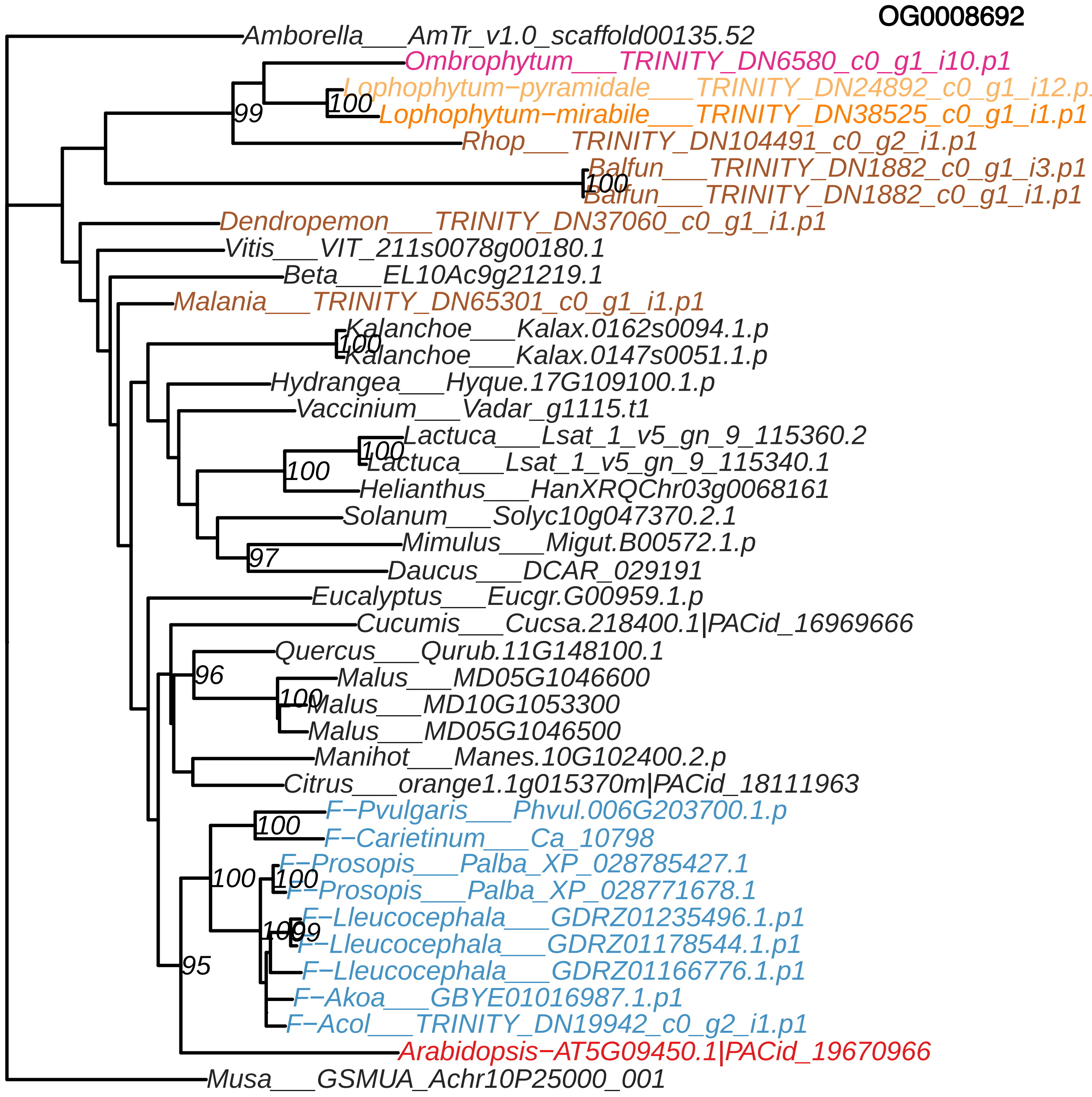

OG0008692
Amborella___AmTr_v1.0_scaffold00135.52
Ombrophytum___TRINITY_DN6580_c0_g1_i10.p1
Lophophytum−pyramidale___TRINITY_DN24892_c0_g1_i12.p
Lophophytum−mirabile___TRINITY_DN38525_c0_g1_i1.p1
Rhop___TRINITY_DN104491_c0_g2_i1.p1
Balfun___TRINITY_DN1882_c0_g1_i3.p1
Balfun___TRINITY_DN1882_c0_g1_i1.p1
Dendropemon___TRINITY_DN37060_c0_g1_i1.p1
Vitis___VIT_211s0078g00180.1
Beta___EL10Ac9g21219.1
Malania___TRINITY_DN65301_c0_g1_i1.p1
Kalanchoe___Kalax.0162s0094.1.p
Kalanchoe___Kalax.0147s0051.1.p
Hydrangea___Hyque.17G109100.1.p
Vaccinium___Vadar_g1115.t1
Lactuca___Lsat_1_v5_gn_9_115360.2
Lactuca___Lsat_1_v5_gn_9_115340.1
Helianthus___HanXRQChr03g0068161
Solanum___Solyc10g047370.2.1
Mimulus___Migut.B00572.1.p
Daucus___DCAR_029191
Eucalyptus___Eucgr.G00959.1.p
Cucumis___Cucsa.218400.1|PACid_16969666
Quercus___Qurub.11G148100.1
Malus___MD05G1046600
Malus___MD10G1053300
Malus___MD05G1046500
Manihot___Manes.10G102400.2.p
Citrus___orange1.1g015370m|PACid_18111963
F−Pvulgaris___Phvul.006G203700.1.p
F−Carietinum___Ca_10798
F−Prosopis___Palba_XP_028785427.1
F−Prosopis___Palba_XP_028771678.1
F−Lleucocephala___GDRZ01235496.1.p1
F−Lleucocephala___GDRZ01178544.1.p1
F−Lleucocephala___GDRZ01166776.1.p1
F−Akoa___GBYE01016987.1.p1
F−Acol___TRINITY_DN19942_c0_g2_i1.p1
Arabidopsis−AT5G09450.1|PACid_19670966
Musa___GSMUA_Achr10P25000_001
99
100
100
100
100
100
97
96
100
100
100
100
100
99
95

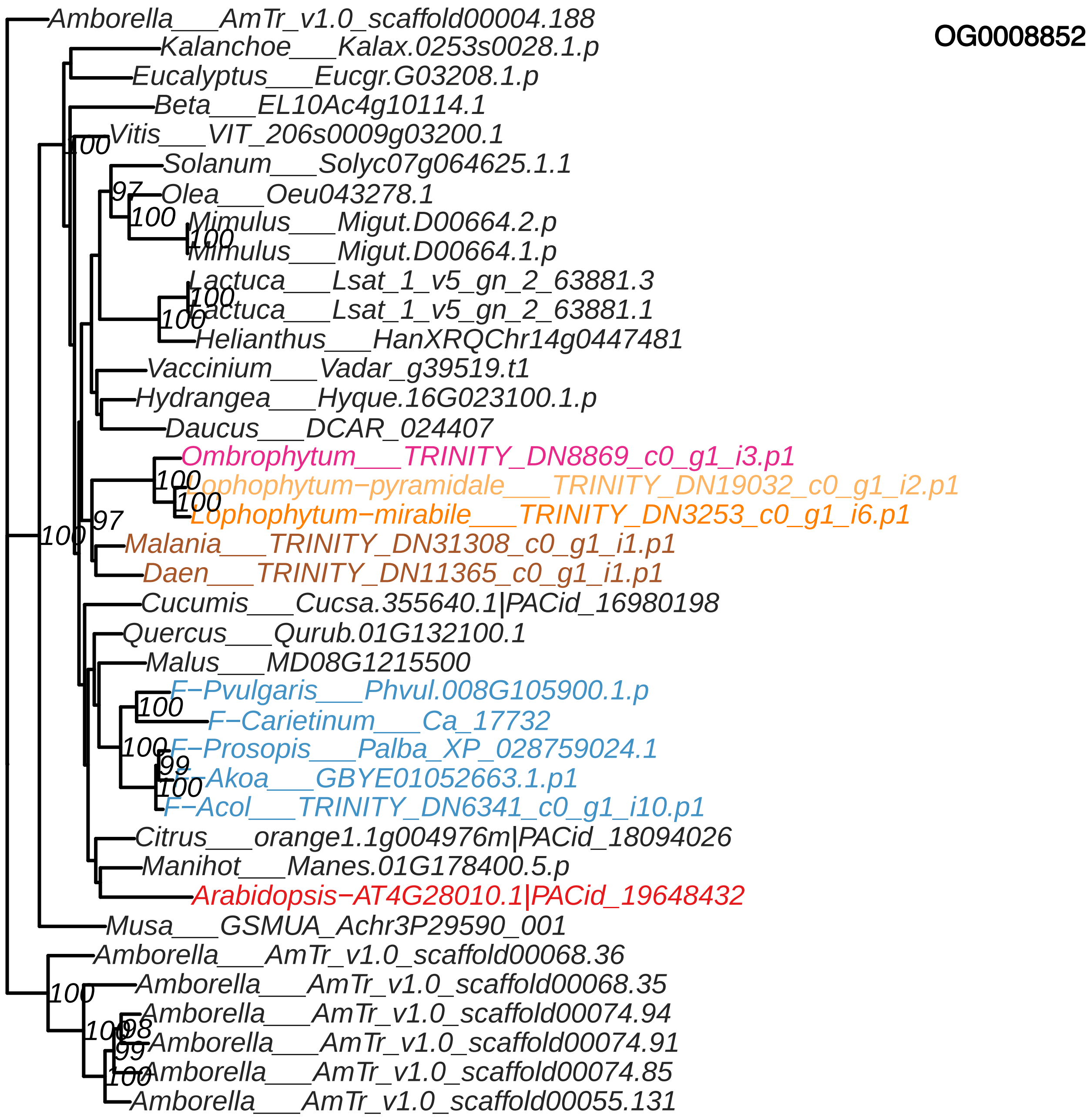
OG0008852
Amborella___AmTr_v1.0_scaffold00004.188
Kalanchoe___Kalax.0253s0028.1.p
Eucalyptus___Eucgr.G03208.1.p
Beta___EL10Ac4g10114.1
Vitis___VIT_206s0009g03200.1
Solanum___Solyc07g064625.1.1
Olea___Oeu043278.1
Mimulus___Migut.D00664.2.p
Mimulus___Migut.D00664.1.p
Lactuca___Lsat_1_v5_gn_2_63881.3
Lactuca___Lsat_1_v5_gn_2_63881.1
Helianthus___HanXRQChr14g0447481
Vaccinium___Vadar_g39519.t1
Hydrangea___Hyque.16G023100.1.p
Daucus___DCAR_024407
Ombrophytum___TRINITY_DN8869_c0_g1_i3.p1
Lophophytum−pyramidale___TRINITY_DN19032_c0_g1_i2.p1
Lophophytum−mirabile___TRINITY_DN3253_c0_g1_i6.p1
Malania___TRINITY_DN31308_c0_g1_i1.p1
Daen___TRINITY_DN11365_c0_g1_i1.p1
Cucumis___Cucsa.355640.1|PACid_16980198
Quercus___Qurub.01G132100.1
Malus___MD08G1215500
F−Pvulgaris___Phvul.008G105900.1.p
F−Carietinum___Ca_17732
F−Prosopis___Palba_XP_028759024.1
F−Akoa___GBYE01052663.1.p1
F−Acol___TRINITY_DN6341_c0_g1_i10.p1
Citrus___orange1.1g004976m|PACid_18094026
Manihot___Manes.01G178400.5.p
Arabidopsis−AT4G28010.1|PACid_19648432
Musa___GSMUA_Achr3P29590_001
Amborella___AmTr_v1.0_scaffold00068.36
Amborella___AmTr_v1.0_scaffold00068.35
Amborella___AmTr_v1.0_scaffold00074.94
Amborella___AmTr_v1.0_scaffold00074.91
Amborella___AmTr_v1.0_scaffold00074.85
Amborella___AmTr_v1.0_scaffold00055.131
100
97
100
100
100
100
100
100
97
100
100
100
99
100
100
100
98
99
100

OG0008911

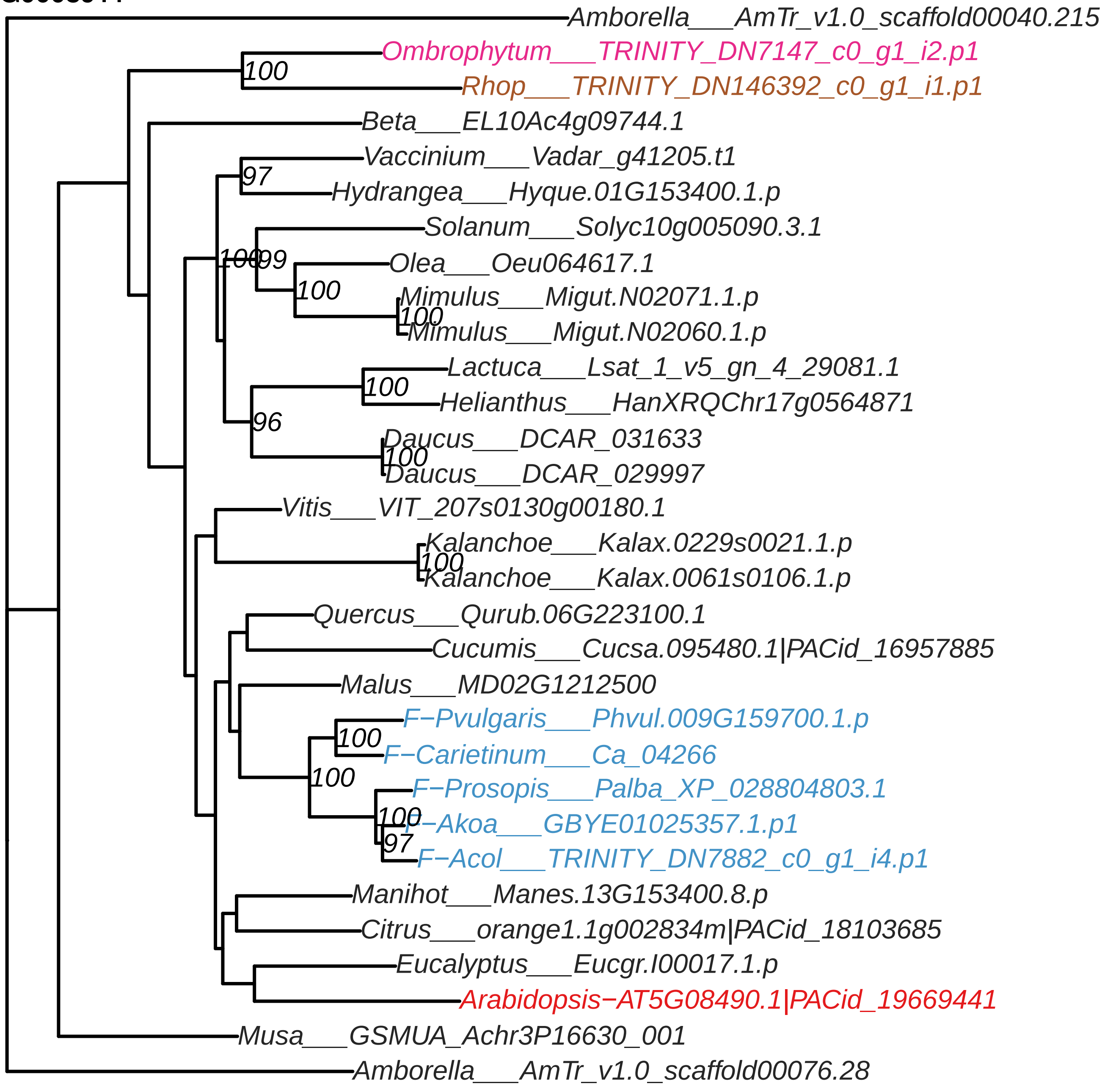
Amborella___AmTr_v1.0_scaffold00040.215
Ombrophytum___TRINITY_DN7147_c0_g1_i2.p1
100
Rhop___TRINITY_DN146392_c0_g1_i1.p1
Beta___EL10Ac4g09744.1
Vaccinium___Vadar_g41205.t1
97
Hydrangea___Hyque.01G153400.1.p
Solanum___Solyc10g005090.3.1
100
99
Olea___Oeu064617.1
100
Mimulus___Migut.N02071.1.p
100
Mimulus___Migut.N02060.1.p
Lactuca___Lsat_1_v5_gn_4_29081.1
100
Helianthus___HanXRQChr17g0564871
96
Daucus___DCAR_031633
100
Daucus___DCAR_029997
Vitis___VIT_207s0130g00180.1
Kalanchoe___Kalax.0229s0021.1.p
100
Kalanchoe___Kalax.0061s0106.1.p
Quercus___Qurub.06G223100.1
Cucumis___Cucsa.095480.1|PACid_16957885
Malus___MD02G1212500
F−Pvulgaris___Phvul.009G159700.1.p
100
F−Carietinum___Ca_04266
100
F−Prosopis___Palba_XP_028804803.1
100
F−Akoa___GBYE01025357.1.p1
97
F−Acol___TRINITY_DN7882_c0_g1_i4.p1
Manihot___Manes.13G153400.8.p
Citrus___orange1.1g002834m|PACid_18103685
Eucalyptus___Eucgr.I00017.1.p
Arabidopsis−AT5G08490.1|PACid_19669441
Musa___GSMUA_Achr3P16630_001
Amborella___AmTr_v1.0_scaffold00076.28

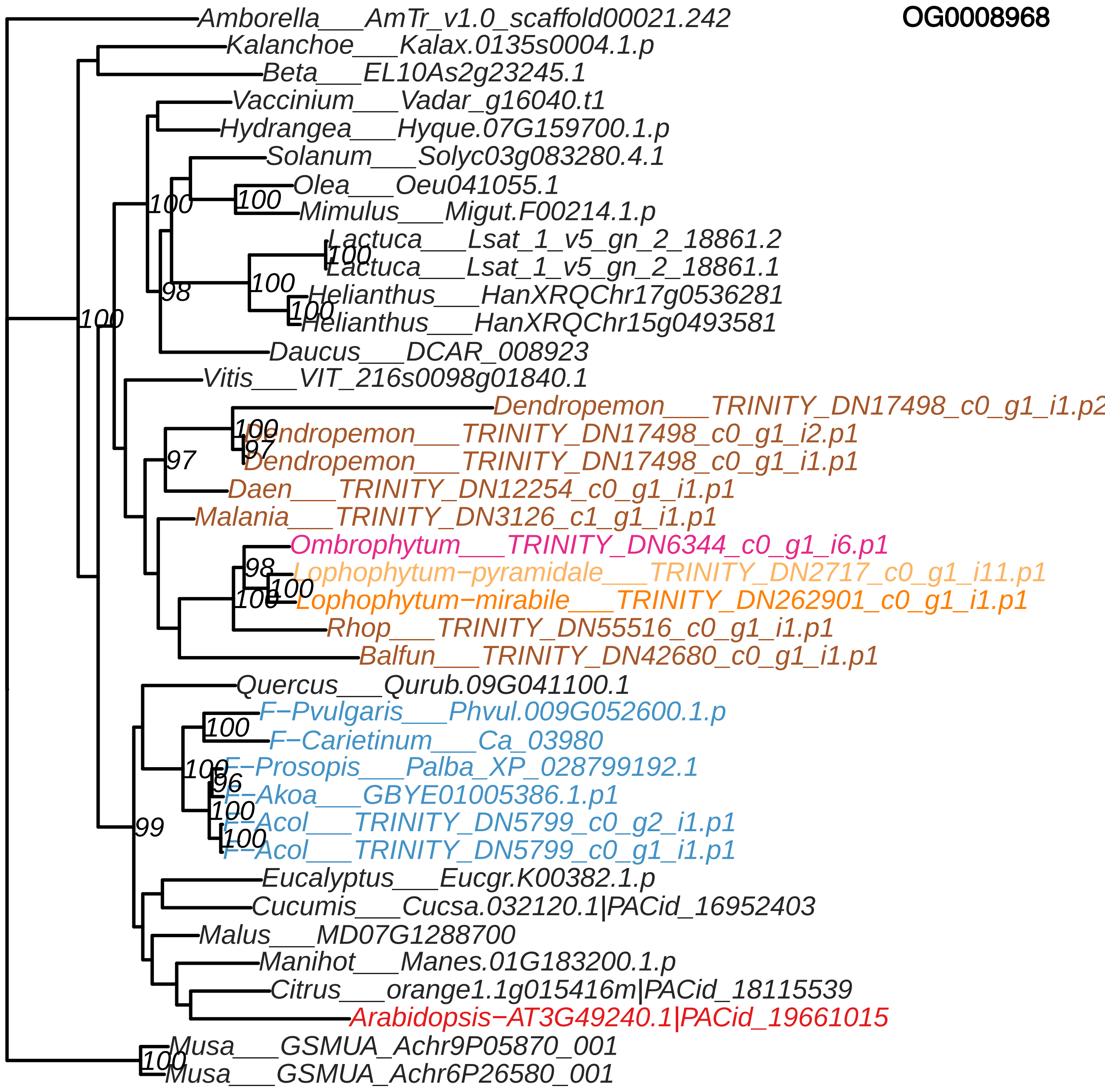
OG0008968
Amborella___AmTr_v1.0_scaffold00021.242
Kalanchoe___Kalax.0135s0004.1.p
Beta___EL10As2g23245.1
Vaccinium___Vadar_g16040.t1
Hydrangea___Hyque.07G159700.1.p
Solanum___Solyc03g083280.4.1
Olea___Oeu041055.1
Mimulus___Migut.F00214.1.p
Lactuca___Lsat_1_v5_gn_2_18861.2
Lactuca___Lsat_1_v5_gn_2_18861.1
Helianthus___HanXRQChr17g0536281
Helianthus___HanXRQChr15g0493581
Daucus___DCAR_008923
Vitis___VIT_216s0098g01840.1
Dendropemon___TRINITY_DN17498_c0_g1_i1.p2
Dendropemon___TRINITY_DN17498_c0_g1_i2.p1
Dendropemon___TRINITY_DN17498_c0_g1_i1.p1
Daen___TRINITY_DN12254_c0_g1_i1.p1
Malania___TRINITY_DN3126_c1_g1_i1.p1
Ombrophytum___TRINITY_DN6344_c0_g1_i6.p1
Lophophytum−pyramidale___TRINITY_DN2717_c0_g1_i11.p1
Lophophytum−mirabile___TRINITY_DN262901_c0_g1_i1.p1
Rhop___TRINITY_DN55516_c0_g1_i1.p1
Balfun___TRINITY_DN42680_c0_g1_i1.p1
Quercus___Qurub.09G041100.1
F−Pvulgaris___Phvul.009G052600.1.p
F−Carietinum___Ca_03980
F−Prosopis___Palba_XP_028799192.1
F−Akoa___GBYE01005386.1.p1
F−Acol___TRINITY_DN5799_c0_g2_i1.p1
F−Acol___TRINITY_DN5799_c0_g1_i1.p1
Eucalyptus___Eucgr.K00382.1.p
Cucumis___Cucsa.032120.1|PACid_16952403
Malus___MD07G1288700
Manihot___Manes.01G183200.1.p
Citrus___orange1.1g015416m|PACid_18115539
Arabidopsis−AT3G49240.1|PACid_19661015
Musa___GSMUA_Achr9P05870_001
Musa___GSMUA_Achr6P26580_001

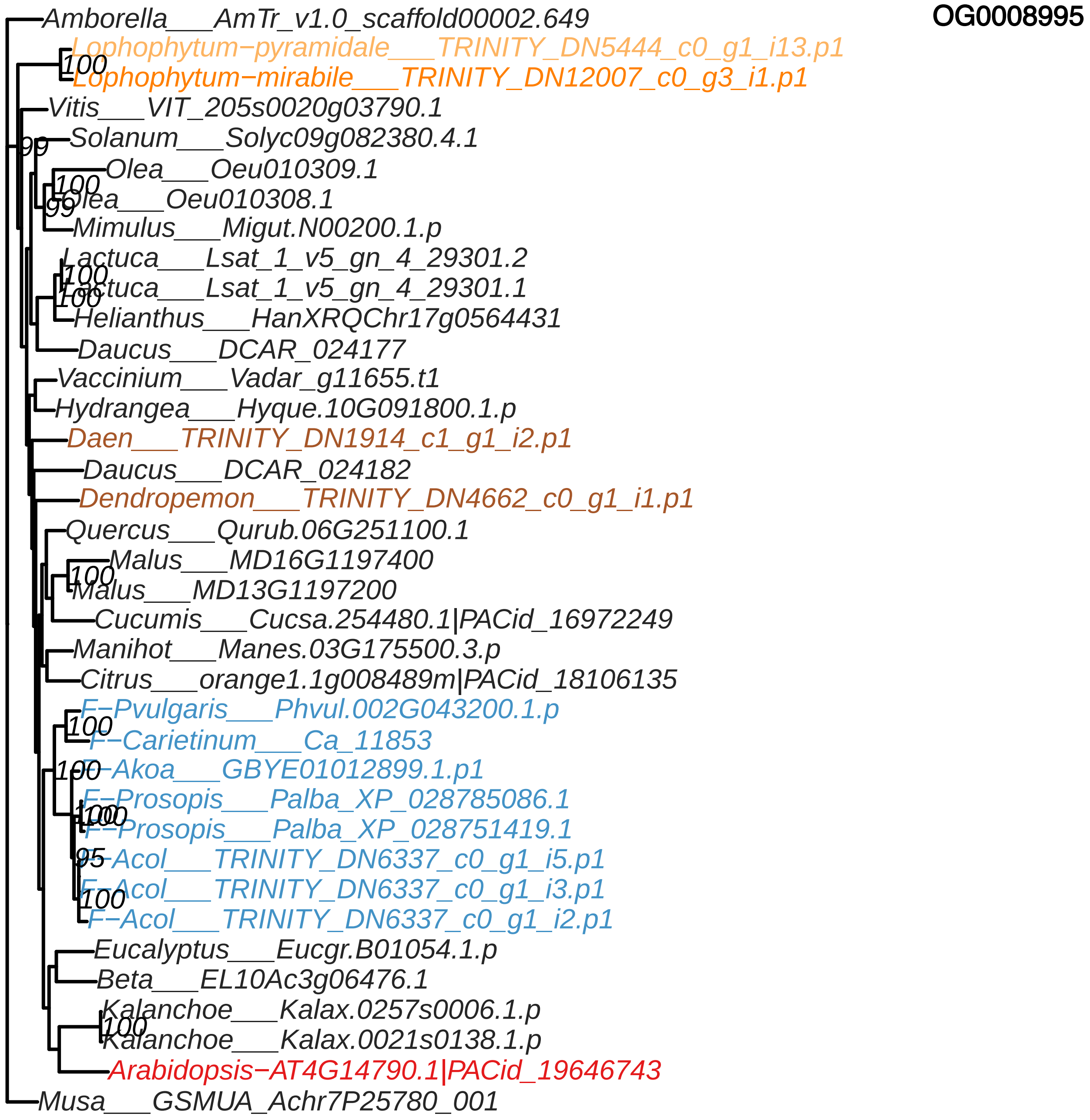
OG0008995
Amborella___AmTr_v1.0_scaffold00002.649
Lophophytum−pyramidale___TRINITY_DN5444_c0_g1_i13.p1
Lophophytum−mirabile___TRINITY_DN12007_c0_g3_i1.p1
Vitis___VIT_205s0020g03790.1
Solanum___Solyc09g082380.4.1
Olea___Oeu010309.1
Olea___Oeu010308.1
Mimulus___Migut.N00200.1.p
Lactuca___Lsat_1_v5_gn_4_29301.2
Lactuca___Lsat_1_v5_gn_4_29301.1
Helianthus___HanXRQChr17g0564431
Daucus___DCAR_024177
Vaccinium___Vadar_g11655.t1
Hydrangea___Hyque.10G091800.1.p
Daen___TRINITY_DN1914_c1_g1_i2.p1
Daucus___DCAR_024182
Dendropemon___TRINITY_DN4662_c0_g1_i1.p1
Quercus___Qurub.06G251100.1
Malus___MD16G1197400
Malus___MD13G1197200
Cucumis___Cucsa.254480.1|PACid_16972249
Manihot___Manes.03G175500.3.p
Citrus___orange1.1g008489m|PACid_18106135
F−Pvulgaris___Phvul.002G043200.1.p
F−Carietinum___Ca_11853
F−Akoa___GBYE01012899.1.p1
F−Prosopis___Palba_XP_028785086.1
F−Prosopis___Palba_XP_028751419.1
F−Acol___TRINITY_DN6337_c0_g1_i5.p1
F−Acol___TRINITY_DN6337_c0_g1_i3.p1
F−Acol___TRINITY_DN6337_c0_g1_i2.p1
Eucalyptus___Eucgr.B01054.1.p
Beta___EL10Ac3g06476.1
Kalanchoe___Kalax.0257s0006.1.p
Kalanchoe___Kalax.0021s0138.1.p
Arabidopsis−AT4G14790.1|PACid_19646743
Musa___GSMUA_Achr7P25780_001
100
99
100
99
100
100
100
100
100
100
95
100
100

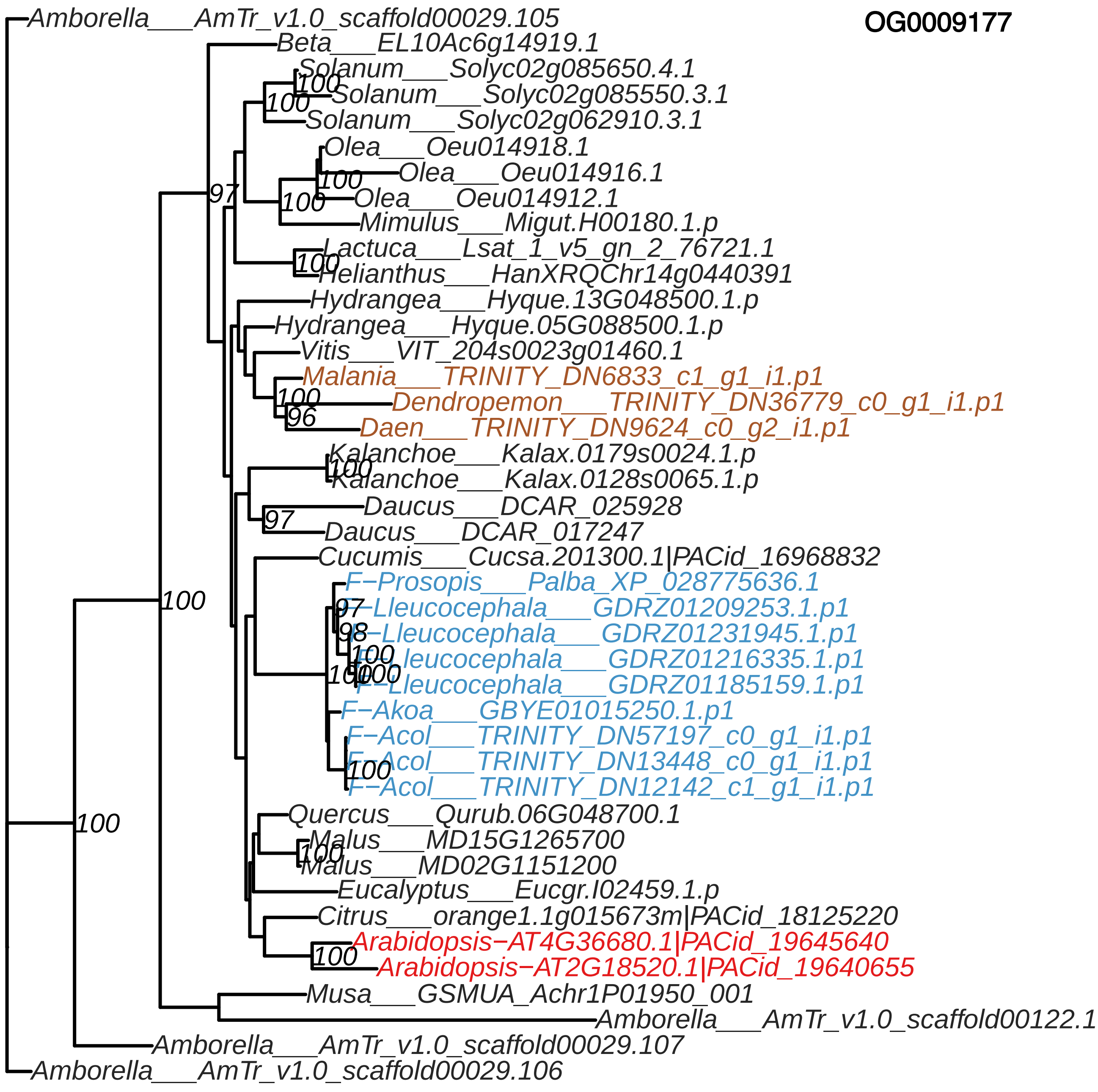
OG0009177
Amborella___AmTr_v1.0_scaffold00029.105
Beta___EL10Ac6g14919.1
Solanum___Solyc02g085650.4.1
Solanum___Solyc02g085550.3.1
Solanum___Solyc02g062910.3.1
Olea___Oeu014918.1
Olea___Oeu014916.1
Olea___Oeu014912.1
Mimulus___Migut.H00180.1.p
Lactuca___Lsat_1_v5_gn_2_76721.1
Helianthus___HanXRQChr14g0440391
Hydrangea___Hyque.13G048500.1.p
Hydrangea___Hyque.05G088500.1.p
Vitis___VIT_204s0023g01460.1
Malania___TRINITY_DN6833_c1_g1_i1.p1
Dendropemon___TRINITY_DN36779_c0_g1_i1.p1
Daen___TRINITY_DN9624_c0_g2_i1.p1
Kalanchoe___Kalax.0179s0024.1.p
Kalanchoe___Kalax.0128s0065.1.p
Daucus___DCAR_025928
Daucus___DCAR_017247
Cucumis___Cucsa.201300.1|PACid_16968832
F−Prosopis___Palba_XP_028775636.1
F−Lleucocephala___GDRZ01209253.1.p1
F−Lleucocephala___GDRZ01231945.1.p1
F−Lleucocephala___GDRZ01216335.1.p1
F−Lleucocephala___GDRZ01185159.1.p1
F−Akoa___GBYE01015250.1.p1
F−Acol___TRINITY_DN57197_c0_g1_i1.p1
F−Acol___TRINITY_DN13448_c0_g1_i1.p1
F−Acol___TRINITY_DN12142_c1_g1_i1.p1
Quercus___Qurub.06G048700.1
Malus___MD15G1265700
Malus___MD02G1151200
Eucalyptus___Eucgr.I02459.1.p
Citrus___orange1.1g015673m|PACid_18125220
Arabidopsis−AT4G36680.1|PACid_19645640
Arabidopsis−AT2G18520.1|PACid_19640655
Musa___GSMUA_Achr1P01950_001
Amborella___AmTr_v1.0_scaffold00122.1
Amborella___AmTr_v1.0_scaffold00029.107
Amborella___AmTr_v1.0_scaffold00029.106
100
100
100
97
100
100
100
96
100
97
100
97
98
100
100
100
100
100
100
100
100

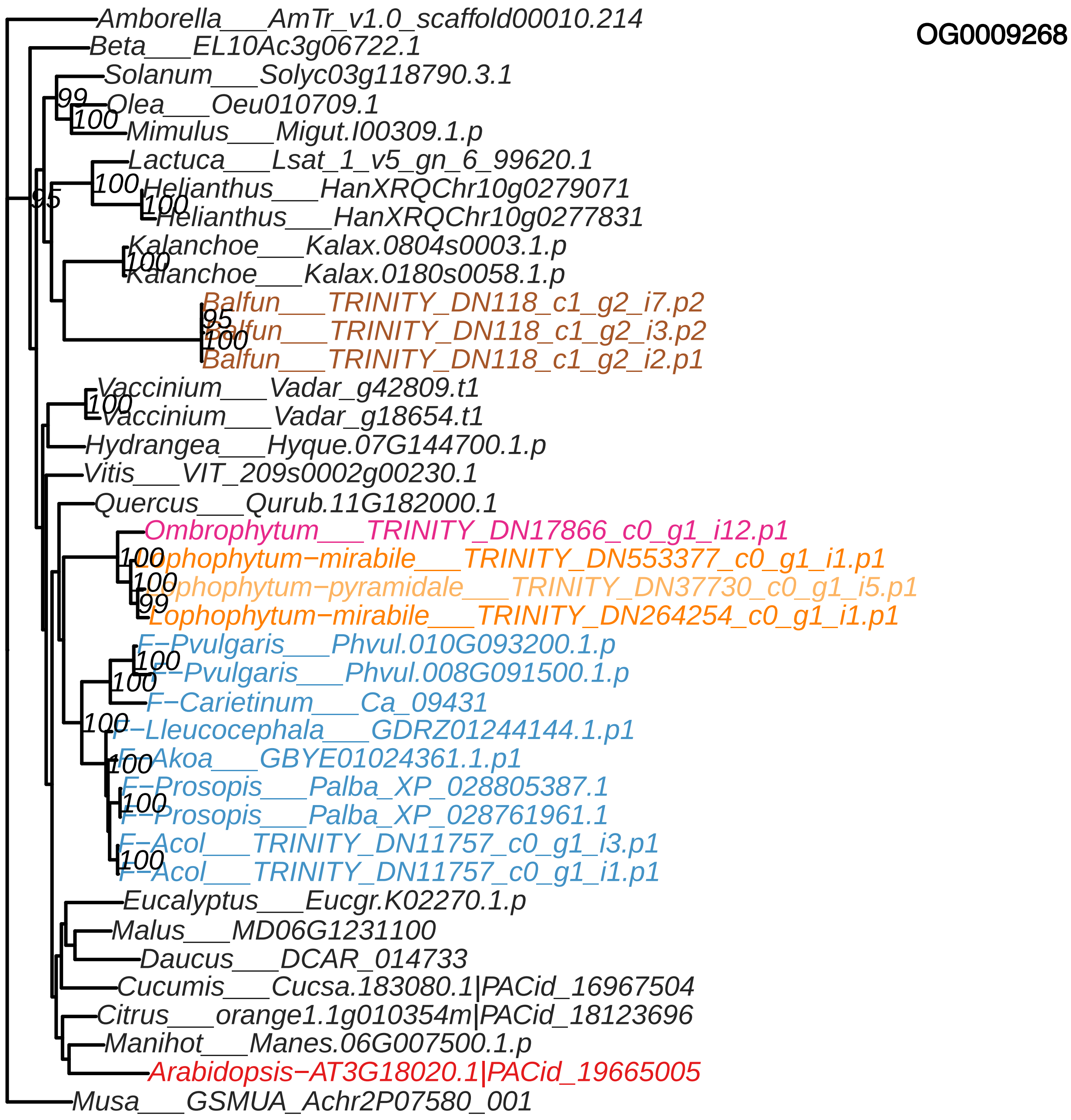
OG0009268
Amborella___AmTr_v1.0_scaffold00010.214
Beta___EL10Ac3g06722.1
Solanum___Solyc03g118790.3.1
99
Olea___Oeu010709.1
100
Mimulus___Migut.I00309.1.p
Lactuca___Lsat_1_v5_gn_6_99620.1
100
Helianthus___HanXRQChr10g0279071
100
Helianthus___HanXRQChr10g0277831
95
Kalanchoe___Kalax.0804s0003.1.p
100
Kalanchoe___Kalax.0180s0058.1.p
Balfun___TRINITY_DN118_c1_g2_i7.p2
95
Balfun___TRINITY_DN118_c1_g2_i3.p2
100
Balfun___TRINITY_DN118_c1_g2_i2.p1
Vaccinium___Vadar_g42809.t1
100
Vaccinium___Vadar_g18654.t1
Hydrangea___Hyque.07G144700.1.p
Vitis___VIT_209s0002g00230.1
Quercus___Qurub.11G182000.1
Ombrophytum___TRINITY_DN17866_c0_g1_i12.p1
100
Lophophytum−mirabile___TRINITY_DN553377_c0_g1_i1.p1
100
Lophophytum−pyramidale___TRINITY_DN37730_c0_g1_i5.p1
99
Lophophytum−mirabile___TRINITY_DN264254_c0_g1_i1.p1
F−Pvulgaris___Phvul.010G093200.1.p
100
F−Pvulgaris___Phvul.008G091500.1.p
100
F−Carietinum___Ca_09431
100
F−Lleucocephala___GDRZ01244144.1.p1
100
F−Akoa___GBYE01024361.1.p1
F−Prosopis___Palba_XP_028805387.1
100
F−Prosopis___Palba_XP_028761961.1
F−Acol___TRINITY_DN11757_c0_g1_i3.p1
100
F−Acol___TRINITY_DN11757_c0_g1_i1.p1
Eucalyptus___Eucgr.K02270.1.p
Malus___MD06G1231100
Daucus___DCAR_014733
Cucumis___Cucsa.183080.1|PACid_16967504
Citrus___orange1.1g010354m|PACid_18123696
Manihot___Manes.06G007500.1.p
Arabidopsis−AT3G18020.1|PACid_19665005
Musa___GSMUA_Achr2P07580_001

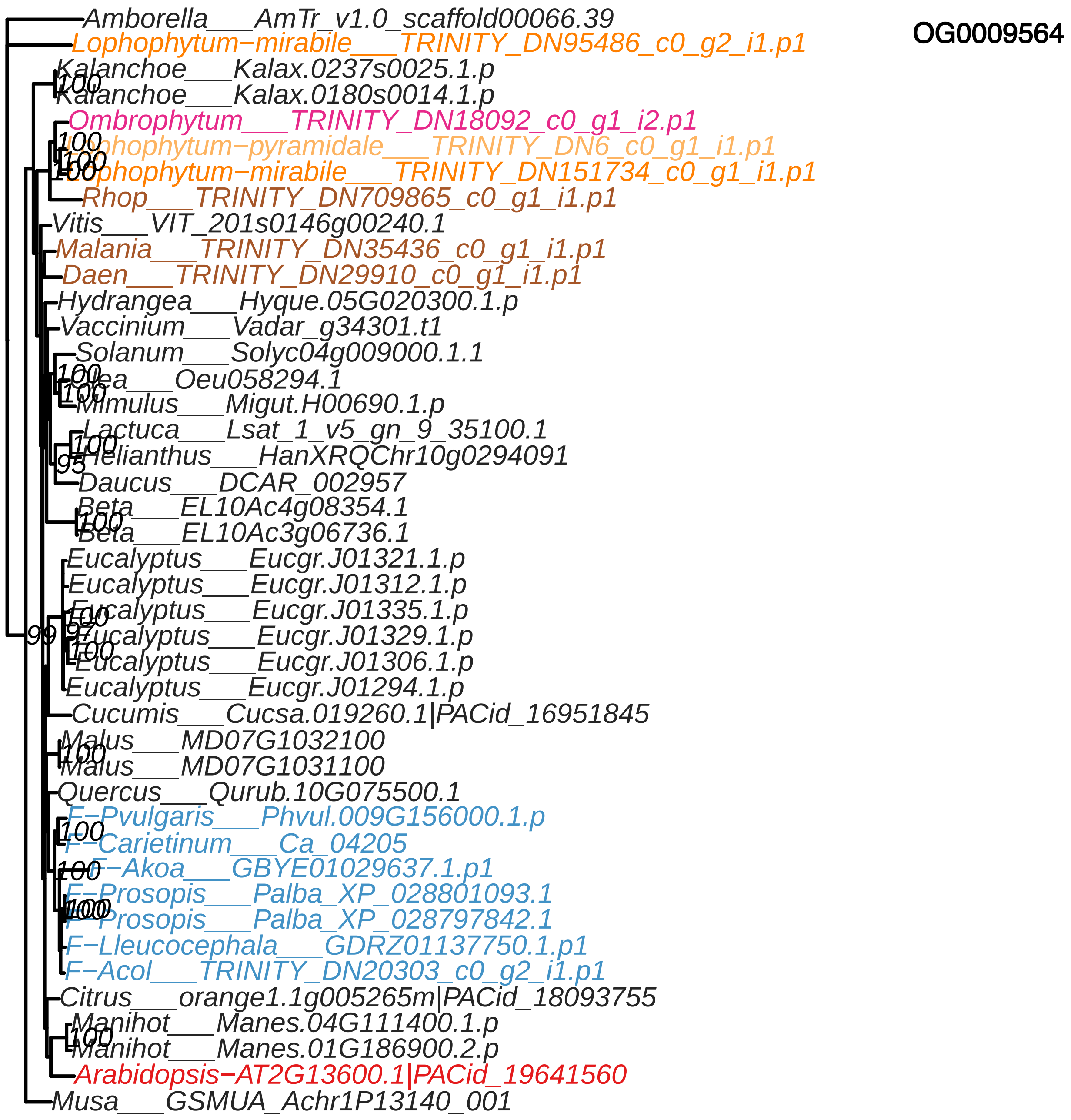
OG0009564
Amborella___AmTr_v1.0_scaffold00066.39
Lophophytum−mirabile___TRINITY_DN95486_c0_g2_i1.p1
Kalanchoe___Kalax.0237s0025.1.p
Kalanchoe___Kalax.0180s0014.1.p
Ombrophytum___TRINITY_DN18092_c0_g1_i2.p1
Lophophytum−pyramidale___TRINITY_DN6_c0_g1_i1.p1
Lophophytum−mirabile___TRINITY_DN151734_c0_g1_i1.p1
Rhop___TRINITY_DN709865_c0_g1_i1.p1
Vitis___VIT_201s0146g00240.1
Malania___TRINITY_DN35436_c0_g1_i1.p1
Daen___TRINITY_DN29910_c0_g1_i1.p1
Hydrangea___Hyque.05G020300.1.p
Vaccinium___Vadar_g34301.t1
Solanum___Solyc04g009000.1.1
Olea___Oeu058294.1
Mimulus___Migut.H00690.1.p
Lactuca___Lsat_1_v5_gn_9_35100.1
Helianthus___HanXRQChr10g0294091
Daucus___DCAR_002957
Beta___EL10Ac4g08354.1
Beta___EL10Ac3g06736.1
Eucalyptus___Eucgr.J01321.1.p
Eucalyptus___Eucgr.J01312.1.p
Eucalyptus___Eucgr.J01335.1.p
Eucalyptus___Eucgr.J01329.1.p
Eucalyptus___Eucgr.J01306.1.p
Eucalyptus___Eucgr.J01294.1.p
Cucumis___Cucsa.019260.1|PACid_16951845
Malus___MD07G1032100
Malus___MD07G1031100
Quercus___Qurub.10G075500.1
F−Pvulgaris___Phvul.009G156000.1.p
F−Carietinum___Ca_04205
F−Akoa___GBYE01029637.1.p1
F−Prosopis___Palba_XP_028801093.1
F−Prosopis___Palba_XP_028797842.1
F−Lleucocephala___GDRZ01137750.1.p1
F−Acol___TRINITY_DN20303_c0_g2_i1.p1
Citrus___orange1.1g005265m|PACid_18093755
Manihot___Manes.04G111400.1.p
Manihot___Manes.01G186900.2.p
Arabidopsis−AT2G13600.1|PACid_19641560
Musa___GSMUA_Achr1P13140_001
100
100
100
100
100
100
100
95
100
99
97
100
100
100
100
100
100
100
100

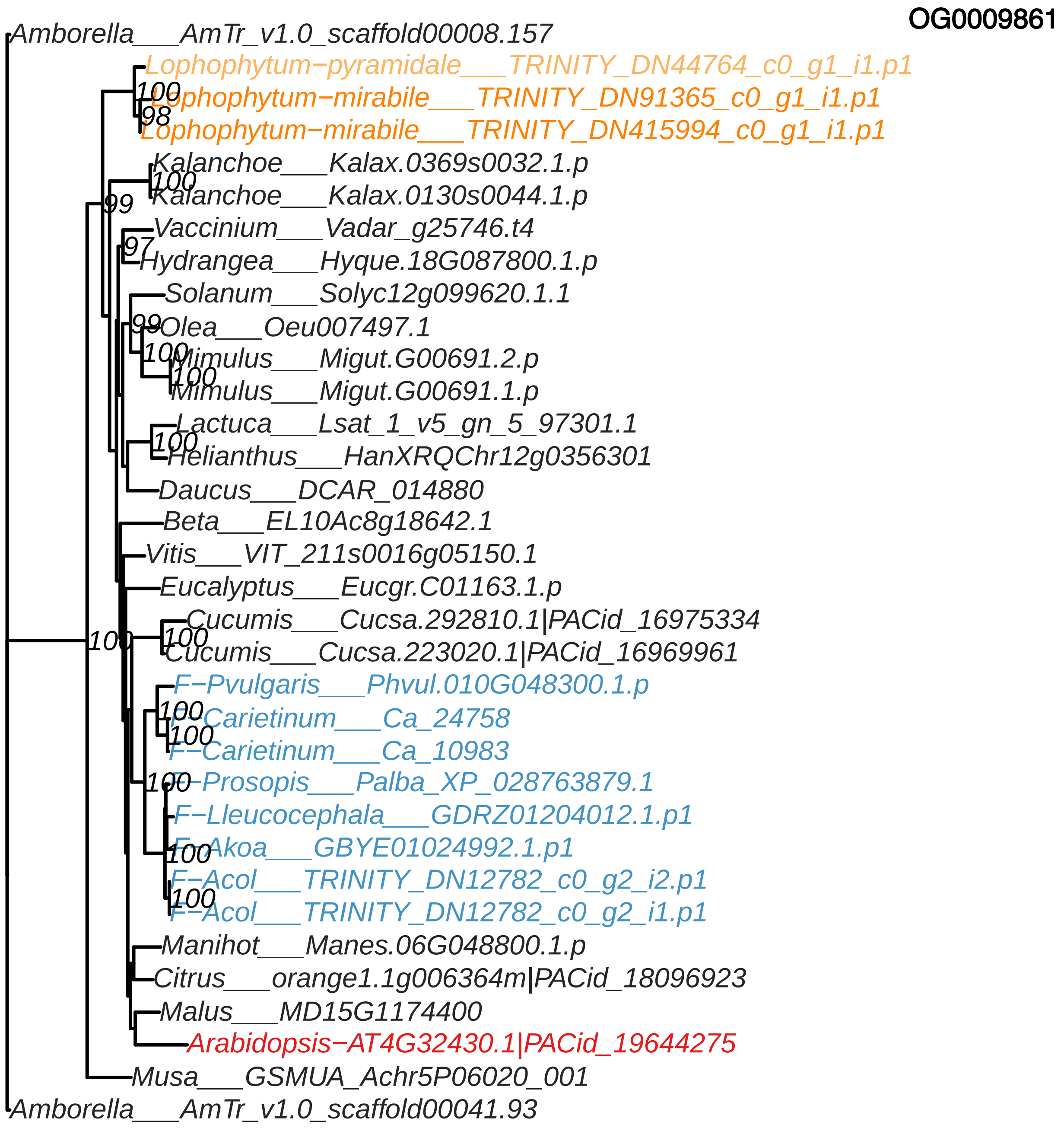
OG0009861
Amborella___AmTr_v1.0_scaffold00008.157
Lophophytum−pyramidale___TRINITY_DN44764_c0_g1_i1.p1
Lophophytum−mirabile___TRINITY_DN91365_c0_g1_i1.p1
Lophophytum−mirabile___TRINITY_DN415994_c0_g1_i1.p1
Kalanchoe___Kalax.0369s0032.1.p
Kalanchoe___Kalax.0130s0044.1.p
Vaccinium___Vadar_g25746.t4
Hydrangea___Hyque.18G087800.1.p
Solanum___Solyc12g099620.1.1
Olea___Oeu007497.1
Mimulus___Migut.G00691.2.p
Mimulus___Migut.G00691.1.p
Lactuca___Lsat_1_v5_gn_5_97301.1
Helianthus___HanXRQChr12g0356301
Daucus___DCAR_014880
Beta___EL10Ac8g18642.1
Vitis___VIT_211s0016g05150.1
Eucalyptus___Eucgr.C01163.1.p
Cucumis___Cucsa.292810.1|PACid_16975334
Cucumis___Cucsa.223020.1|PACid_16969961
F−Pvulgaris___Phvul.010G048300.1.p
F−Carietinum___Ca_24758
F−Carietinum___Ca_10983
F−Prosopis___Palba_XP_028763879.1
F−Lleucocephala___GDRZ01204012.1.p1
F−Akoa___GBYE01024992.1.p1
F−Acol___TRINITY_DN12782_c0_g2_i2.p1
F−Acol___TRINITY_DN12782_c0_g2_i1.p1
Manihot___Manes.06G048800.1.p
Citrus___orange1.1g006364m|PACid_18096923
Malus___MD15G1174400
Arabidopsis−AT4G32430.1|PACid_19644275
Musa___GSMUA_Achr5P06020_001
Amborella___AmTr_v1.0_scaffold00041.93
100
98
100
99
97
99
100
100
100
100
100
100
100
100
100
100

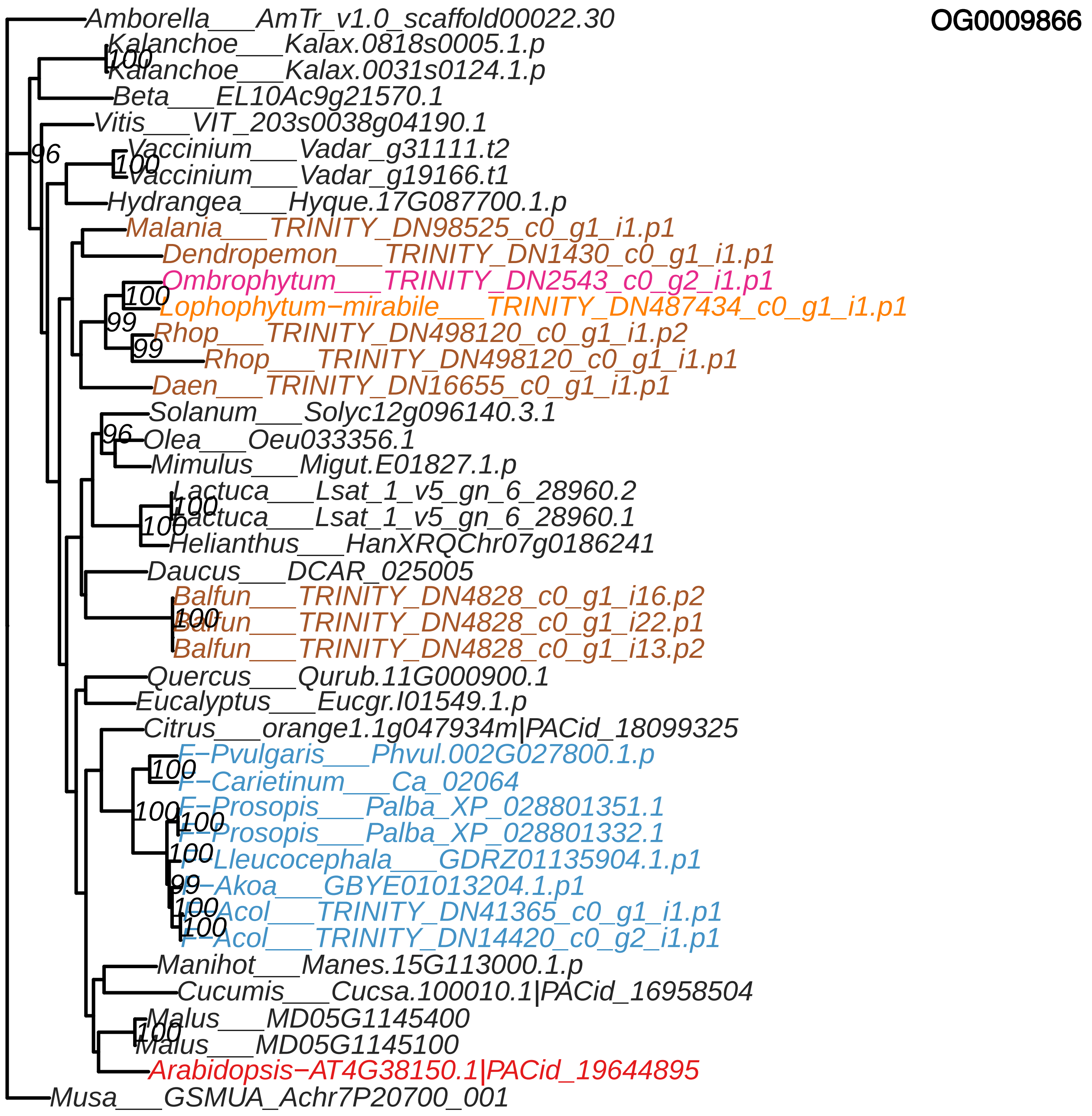
OG0009866
Amborella___AmTr_v1.0_scaffold00022.30
Kalanchoe___Kalax.0818s0005.1.p
Kalanchoe___Kalax.0031s0124.1.p
Beta___EL10Ac9g21570.1
Vitis___VIT_203s0038g04190.1
Vaccinium___Vadar_g31111.t2
Vaccinium___Vadar_g19166.t1
Hydrangea___Hyque.17G087700.1.p
Malania___TRINITY_DN98525_c0_g1_i1.p1
Dendropemon___TRINITY_DN1430_c0_g1_i1.p1
Ombrophytum___TRINITY_DN2543_c0_g2_i1.p1
Lophophytum−mirabile___TRINITY_DN487434_c0_g1_i1.p1
Rhop___TRINITY_DN498120_c0_g1_i1.p2
Rhop___TRINITY_DN498120_c0_g1_i1.p1
Daen___TRINITY_DN16655_c0_g1_i1.p1
Solanum___Solyc12g096140.3.1
Olea___Oeu033356.1
Mimulus___Migut.E01827.1.p
Lactuca___Lsat_1_v5_gn_6_28960.2
Lactuca___Lsat_1_v5_gn_6_28960.1
Helianthus___HanXRQChr07g0186241
Daucus___DCAR_025005
Balfun___TRINITY_DN4828_c0_g1_i16.p2
Balfun___TRINITY_DN4828_c0_g1_i22.p1
Balfun___TRINITY_DN4828_c0_g1_i13.p2
Quercus___Qurub.11G000900.1
Eucalyptus___Eucgr.I01549.1.p
Citrus___orange1.1g047934m|PACid_18099325
F−Pvulgaris___Phvul.002G027800.1.p
F−Carietinum___Ca_02064
F−Prosopis___Palba_XP_028801351.1
F−Prosopis___Palba_XP_028801332.1
F−Lleucocephala___GDRZ01135904.1.p1
F−Akoa___GBYE01013204.1.p1
F−Acol___TRINITY_DN41365_c0_g1_i1.p1
F−Acol___TRINITY_DN14420_c0_g2_i1.p1
Manihot___Manes.15G113000.1.p
Cucumis___Cucsa.100010.1|PACid_16958504
Malus___MD05G1145400
Malus___MD05G1145100
Arabidopsis−AT4G38150.1|PACid_19644895
Musa___GSMUA_Achr7P20700_001

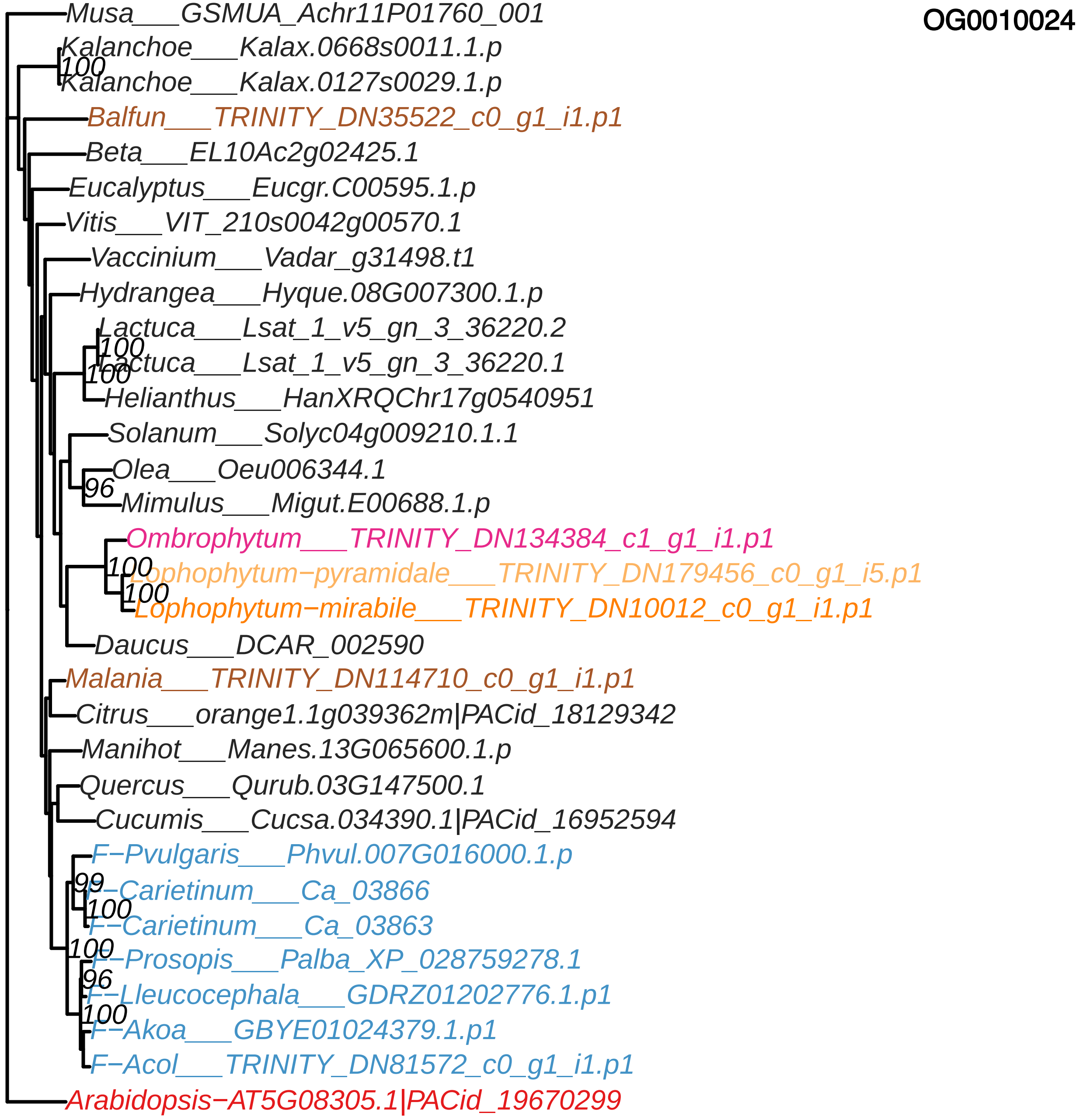
OG0010024
Musa___GSMUA_Achr11P01760_001
Kalanchoe___Kalax.0668s0011.1.p
100
Kalanchoe___Kalax.0127s0029.1.p
Balfun___TRINITY_DN35522_c0_g1_i1.p1
Beta___EL10Ac2g02425.1
Eucalyptus___Eucgr.C00595.1.p
Vitis___VIT_210s0042g00570.1
Vaccinium___Vadar_g31498.t1
Hydrangea___Hyque.08G007300.1.p
Lactuca___Lsat_1_v5_gn_3_36220.2
100
Lactuca___Lsat_1_v5_gn_3_36220.1
100
Helianthus___HanXRQChr17g0540951
Solanum___Solyc04g009210.1.1
Olea___Oeu006344.1
96
Mimulus___Migut.E00688.1.p
Ombrophytum___TRINITY_DN134384_c1_g1_i1.p1
100
Lophophytum−pyramidale___TRINITY_DN179456_c0_g1_i5.p1
100
Lophophytum−mirabile___TRINITY_DN10012_c0_g1_i1.p1
Daucus___DCAR_002590
Malania___TRINITY_DN114710_c0_g1_i1.p1
Citrus___orange1.1g039362m|PACid_18129342
Manihot___Manes.13G065600.1.p
Quercus___Qurub.03G147500.1
Cucumis___Cucsa.034390.1|PACid_16952594
F−Pvulgaris___Phvul.007G016000.1.p
99
F−Carietinum___Ca_03866
100
F−Carietinum___Ca_03863
100
F−Prosopis___Palba_XP_028759278.1
96
F−Lleucocephala___GDRZ01202776.1.p1
100
F−Akoa___GBYE01024379.1.p1
F−Acol___TRINITY_DN81572_c0_g1_i1.p1
Arabidopsis−AT5G08305.1|PACid_19670299

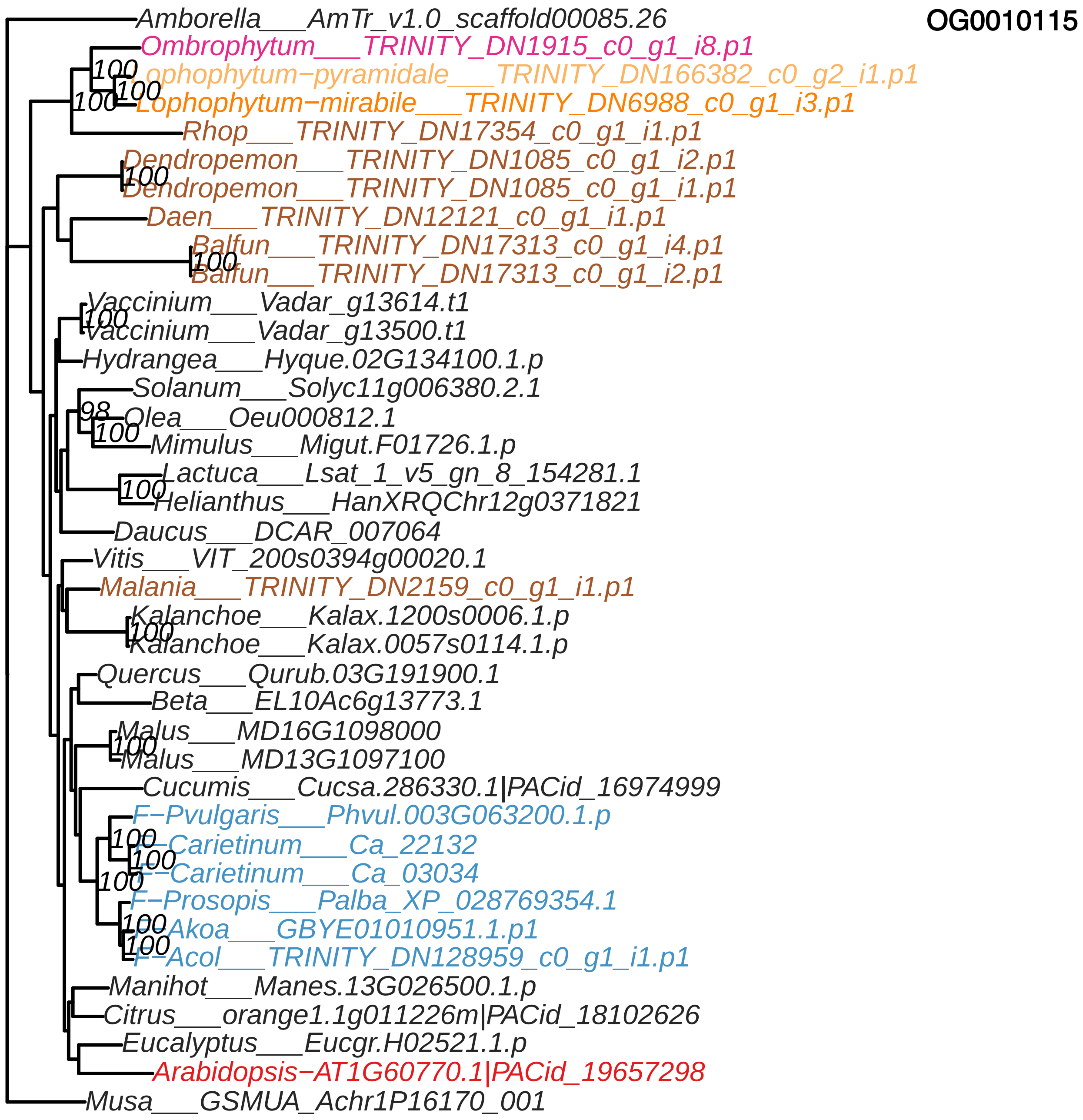
OG0010115
Amborella___AmTr_v1.0_scaffold00085.26
Ombrophytum___TRINITY_DN1915_c0_g1_i8.p1
Lophophytum−pyramidale___TRINITY_DN166382_c0_g2_i1.p1
Lophophytum−mirabile___TRINITY_DN6988_c0_g1_i3.p1
Rhop___TRINITY_DN17354_c0_g1_i1.p1
Dendropemon___TRINITY_DN1085_c0_g1_i2.p1
Dendropemon___TRINITY_DN1085_c0_g1_i1.p1
Daen___TRINITY_DN12121_c0_g1_i1.p1
Balfun___TRINITY_DN17313_c0_g1_i4.p1
Balfun___TRINITY_DN17313_c0_g1_i2.p1
Vaccinium___Vadar_g13614.t1
Vaccinium___Vadar_g13500.t1
Hydrangea___Hyque.02G134100.1.p
Solanum___Solyc11g006380.2.1
Olea___Oeu000812.1
Mimulus___Migut.F01726.1.p
Lactuca___Lsat_1_v5_gn_8_154281.1
Helianthus___HanXRQChr12g0371821
Daucus___DCAR_007064
Vitis___VIT_200s0394g00020.1
Malania___TRINITY_DN2159_c0_g1_i1.p1
Kalanchoe___Kalax.1200s0006.1.p
Kalanchoe___Kalax.0057s0114.1.p
Quercus___Qurub.03G191900.1
Beta___EL10Ac6g13773.1
Malus___MD16G1098000
Malus___MD13G1097100
Cucumis___Cucsa.286330.1|PACid_16974999
F−Pvulgaris___Phvul.003G063200.1.p
F−Carietinum___Ca_22132
F−Carietinum___Ca_03034
F−Prosopis___Palba_XP_028769354.1
F−Akoa___GBYE01010951.1.p1
F−Acol___TRINITY_DN128959_c0_g1_i1.p1
Manihot___Manes.13G026500.1.p
Citrus___orange1.1g011226m|PACid_18102626
Eucalyptus___Eucgr.H02521.1.p
Arabidopsis−AT1G60770.1|PACid_19657298
Musa___GSMUA_Achr1P16170_001

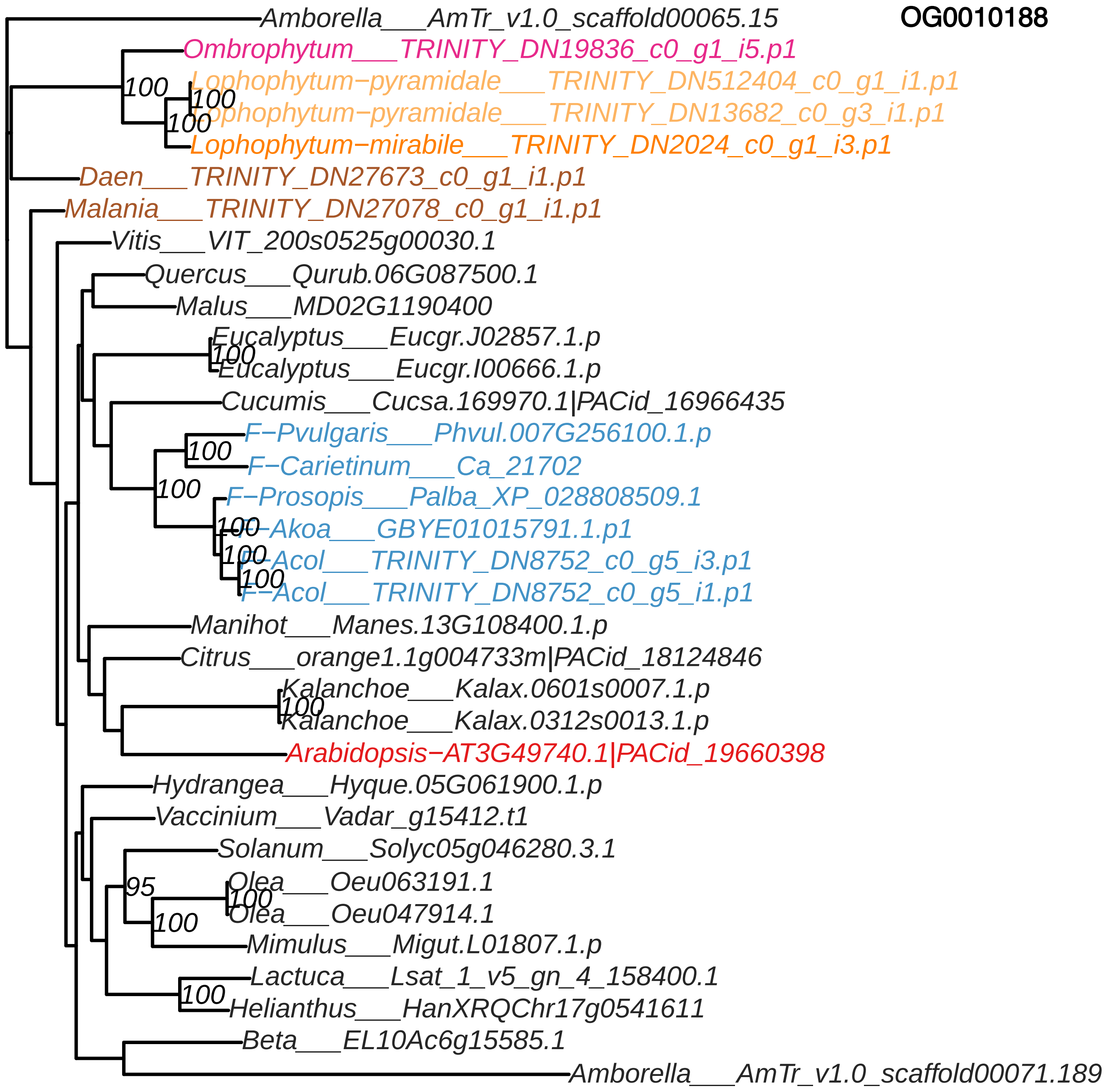
OG0010188
Amborella___AmTr_v1.0_scaffold00065.15
Ombrophytum___TRINITY_DN19836_c0_g1_i5.p1
Lophophytum−pyramidale___TRINITY_DN512404_c0_g1_i1.p1
Lophophytum−pyramidale___TRINITY_DN13682_c0_g3_i1.p1
Lophophytum−mirabile___TRINITY_DN2024_c0_g1_i3.p1
Daen___TRINITY_DN27673_c0_g1_i1.p1
Malania___TRINITY_DN27078_c0_g1_i1.p1
Vitis___VIT_200s0525g00030.1
Quercus___Qurub.06G087500.1
Malus___MD02G1190400
Eucalyptus___Eucgr.J02857.1.p
Eucalyptus___Eucgr.I00666.1.p
Cucumis___Cucsa.169970.1|PACid_16966435
F−Pvulgaris___Phvul.007G256100.1.p
F−Carietinum___Ca_21702
F−Prosopis___Palba_XP_028808509.1
F−Akoa___GBYE01015791.1.p1
F−Acol___TRINITY_DN8752_c0_g5_i3.p1
F−Acol___TRINITY_DN8752_c0_g5_i1.p1
Manihot___Manes.13G108400.1.p
Citrus___orange1.1g004733m|PACid_18124846
Kalanchoe___Kalax.0601s0007.1.p
Kalanchoe___Kalax.0312s0013.1.p
Arabidopsis−AT3G49740.1|PACid_19660398
Hydrangea___Hyque.05G061900.1.p
Vaccinium___Vadar_g15412.t1
Solanum___Solyc05g046280.3.1
Olea___Oeu063191.1
Olea___Oeu047914.1
Mimulus___Migut.L01807.1.p
Lactuca___Lsat_1_v5_gn_4_158400.1
Helianthus___HanXRQChr17g0541611
Beta___EL10Ac6g15585.1
Amborella___AmTr_v1.0_scaffold00071.189
100
100
100
100
100
100
100
100
100
100
95
100
100
100

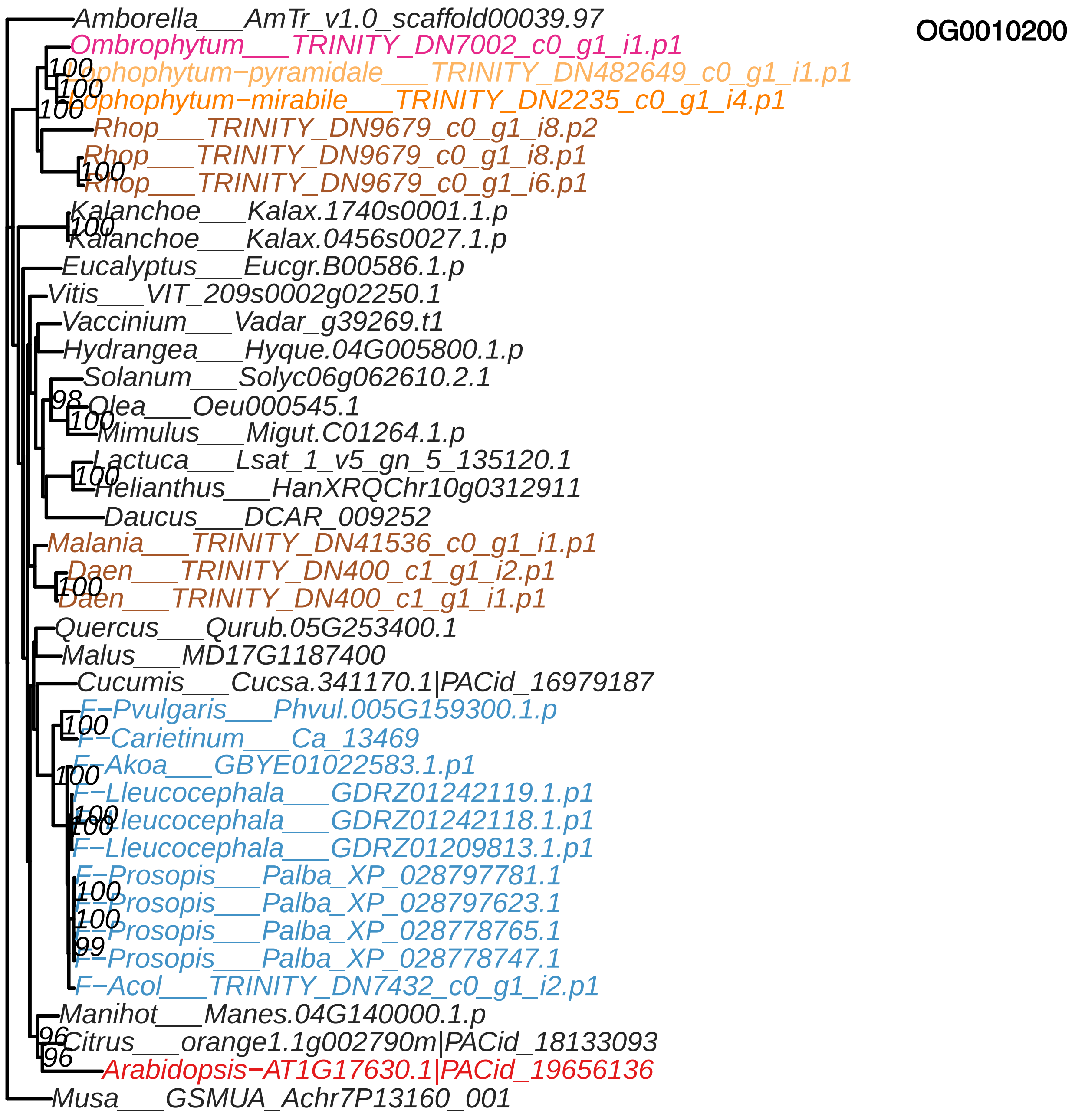
OG0010200
Amborella___AmTr_v1.0_scaffold00039.97
Ombrophytum___TRINITY_DN7002_c0_g1_i1.p1
Lophophytum−pyramidale___TRINITY_DN482649_c0_g1_i1.p1
Lophophytum−mirabile___TRINITY_DN2235_c0_g1_i4.p1
Rhop___TRINITY_DN9679_c0_g1_i8.p2
Rhop___TRINITY_DN9679_c0_g1_i8.p1
Rhop___TRINITY_DN9679_c0_g1_i6.p1
Kalanchoe___Kalax.1740s0001.1.p
Kalanchoe___Kalax.0456s0027.1.p
Eucalyptus___Eucgr.B00586.1.p
Vitis___VIT_209s0002g02250.1
Vaccinium___Vadar_g39269.t1
Hydrangea___Hyque.04G005800.1.p
Solanum___Solyc06g062610.2.1
Olea___Oeu000545.1
Mimulus___Migut.C01264.1.p
Lactuca___Lsat_1_v5_gn_5_135120.1
Helianthus___HanXRQChr10g0312911
Daucus___DCAR_009252
Malania___TRINITY_DN41536_c0_g1_i1.p1
Daen___TRINITY_DN400_c1_g1_i2.p1
Daen___TRINITY_DN400_c1_g1_i1.p1
Quercus___Qurub.05G253400.1
Malus___MD17G1187400
Cucumis___Cucsa.341170.1|PACid_16979187
F−Pvulgaris___Phvul.005G159300.1.p
F−Carietinum___Ca_13469
F−Akoa___GBYE01022583.1.p1
F−Lleucocephala___GDRZ01242119.1.p1
F−Lleucocephala___GDRZ01242118.1.p1
F−Lleucocephala___GDRZ01209813.1.p1
F−Prosopis___Palba_XP_028797781.1
F−Prosopis___Palba_XP_028797623.1
F−Prosopis___Palba_XP_028778765.1
F−Prosopis___Palba_XP_028778747.1
F−Acol___TRINITY_DN7432_c0_g1_i2.p1
Manihot___Manes.04G140000.1.p
Citrus___orange1.1g002790m|PACid_18133093
Arabidopsis−AT1G17630.1|PACid_19656136
Musa___GSMUA_Achr7P13160_001

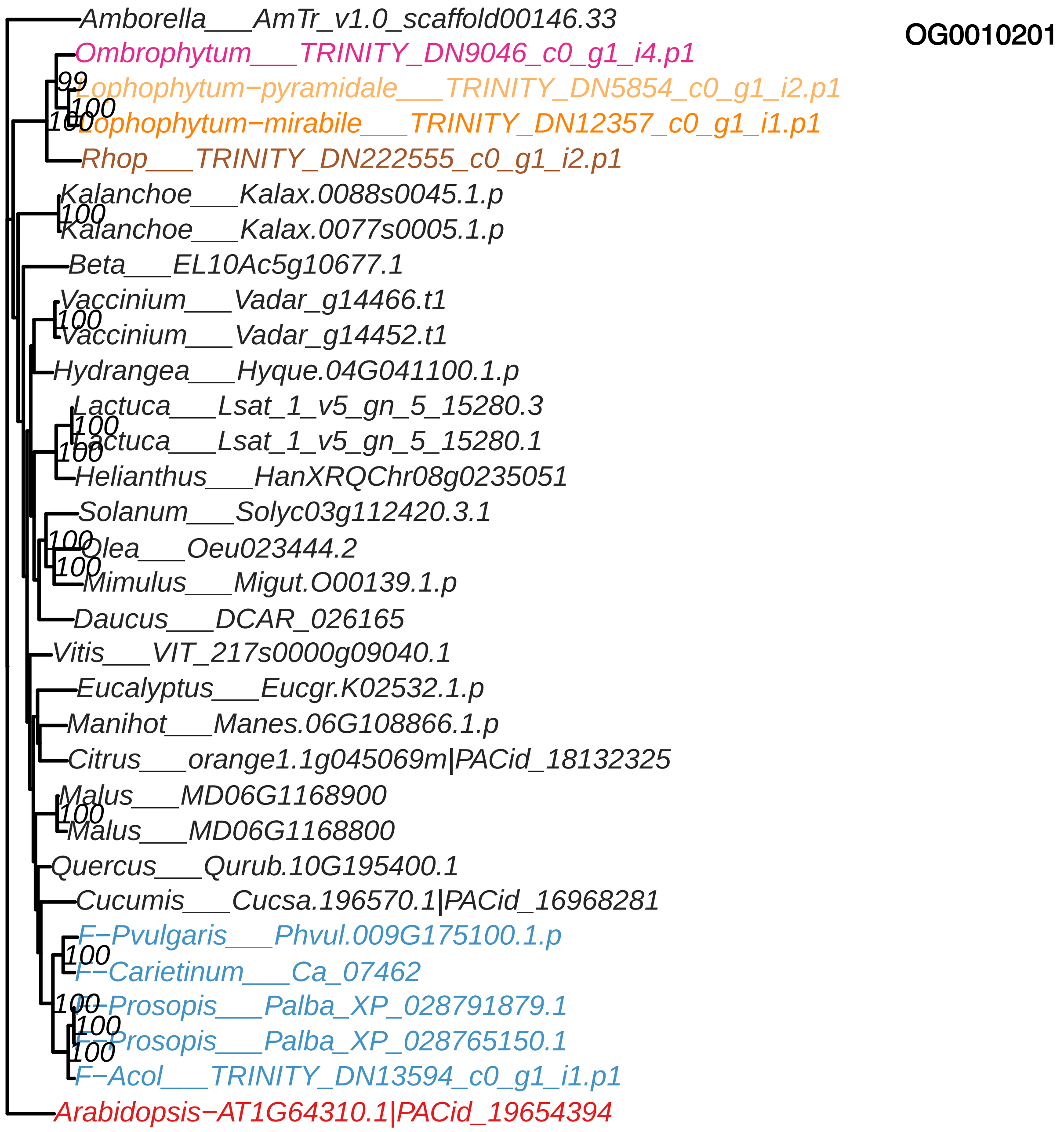
OG0010201
Amborella___AmTr_v1.0_scaffold00146.33
Ombrophytum___TRINITY_DN9046_c0_g1_i4.p1
99
Lophophytum−pyramidale___TRINITY_DN5854_c0_g1_i2.p1
100
100
Lophophytum−mirabile___TRINITY_DN12357_c0_g1_i1.p1
Rhop___TRINITY_DN222555_c0_g1_i2.p1
Kalanchoe___Kalax.0088s0045.1.p
100
Kalanchoe___Kalax.0077s0005.1.p
Beta___EL10Ac5g10677.1
Vaccinium___Vadar_g14466.t1
100
Vaccinium___Vadar_g14452.t1
Hydrangea___Hyque.04G041100.1.p
Lactuca___Lsat_1_v5_gn_5_15280.3
100
Lactuca___Lsat_1_v5_gn_5_15280.1
100
Helianthus___HanXRQChr08g0235051
Solanum___Solyc03g112420.3.1
100
Olea___Oeu023444.2
100
Mimulus___Migut.O00139.1.p
Daucus___DCAR_026165
Vitis___VIT_217s0000g09040.1
Eucalyptus___Eucgr.K02532.1.p
Manihot___Manes.06G108866.1.p
Citrus___orange1.1g045069m|PACid_18132325
Malus___MD06G1168900
100
Malus___MD06G1168800
Quercus___Qurub.10G195400.1
Cucumis___Cucsa.196570.1|PACid_16968281
F−Pvulgaris___Phvul.009G175100.1.p
100
F−Carietinum___Ca_07462
100
F−Prosopis___Palba_XP_028791879.1
100
F−Prosopis___Palba_XP_028765150.1
100
F−Acol___TRINITY_DN13594_c0_g1_i1.p1
Arabidopsis−AT1G64310.1|PACid_19654394

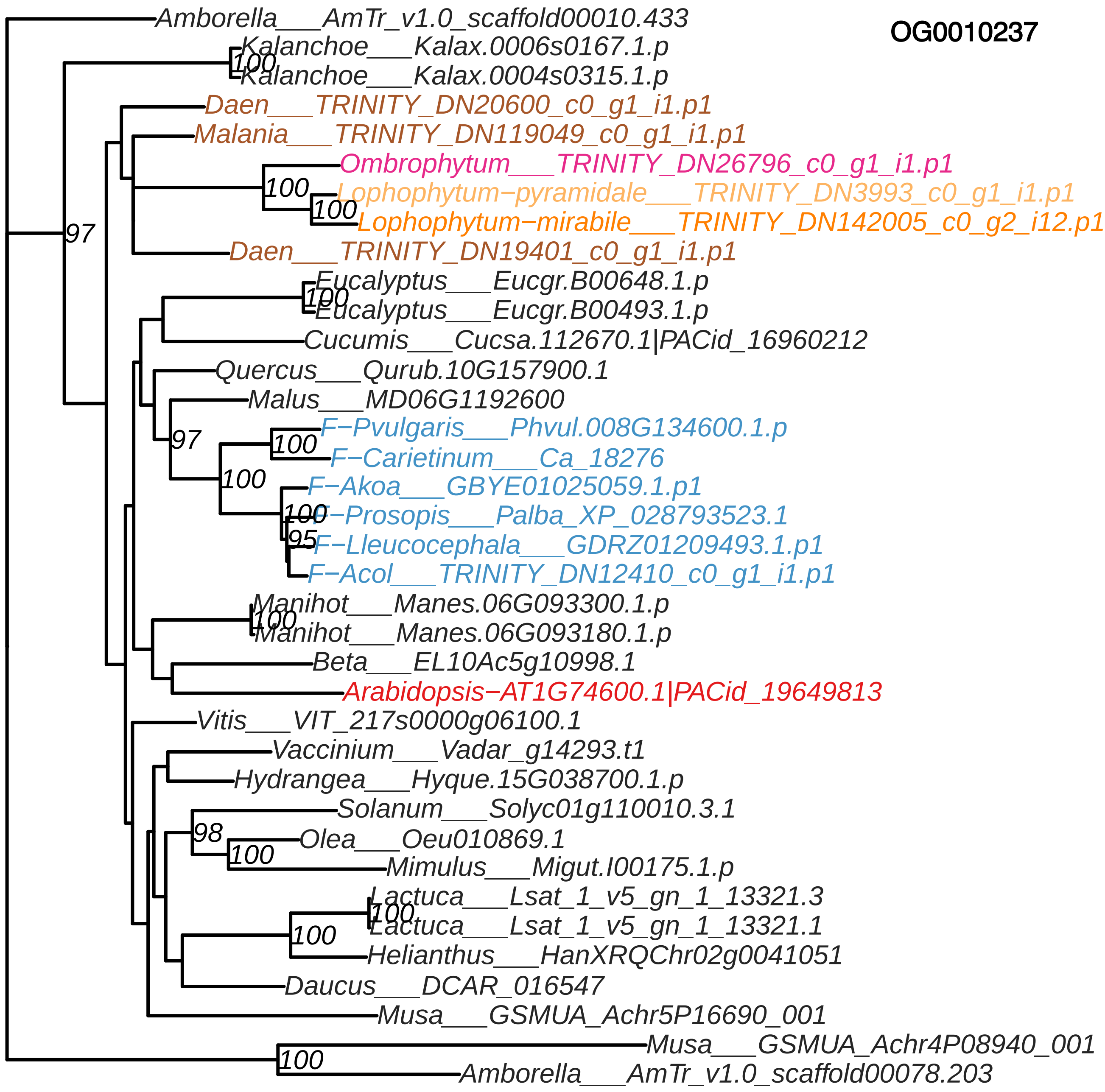

OG0010237
Amborella___AmTr_v1.0_scaffold00010.433
Kalanchoe___Kalax.0006s0167.1.p
100
Kalanchoe___Kalax.0004s0315.1.p
Daen___TRINITY_DN20600_c0_g1_i1.p1
Malania___TRINITY_DN119049_c0_g1_i1.p1
Ombrophytum___TRINITY_DN26796_c0_g1_i1.p1
100
Lophophytum−pyramidale___TRINITY_DN3993_c0_g1_i1.p1
100
Lophophytum−mirabile___TRINITY_DN142005_c0_g2_i12.p1
97
Daen___TRINITY_DN19401_c0_g1_i1.p1
Eucalyptus___Eucgr.B00648.1.p
100
Eucalyptus___Eucgr.B00493.1.p
Cucumis___Cucsa.112670.1|PACid_16960212
Quercus___Qurub.10G157900.1
Malus___MD06G1192600
F−Pvulgaris___Phvul.008G134600.1.p
97
100
F−Carietinum___Ca_18276
F−Akoa___GBYE01025059.1.p1
100
100
F−Prosopis___Palba_XP_028793523.1
95
F−Lleucocephala___GDRZ01209493.1.p1
F−Acol___TRINITY_DN12410_c0_g1_i1.p1
Manihot___Manes.06G093300.1.p
100
Manihot___Manes.06G093180.1.p
Beta___EL10Ac5g10998.1
Arabidopsis−AT1G74600.1|PACid_19649813
Vitis___VIT_217s0000g06100.1
Vaccinium___Vadar_g14293.t1
Hydrangea___Hyque.15G038700.1.p
Solanum___Solyc01g110010.3.1
98
Olea___Oeu010869.1
100
Mimulus___Migut.I00175.1.p
Lactuca___Lsat_1_v5_gn_1_13321.3
100
Lactuca___Lsat_1_v5_gn_1_13321.1
100
Helianthus___HanXRQChr02g0041051
Daucus___DCAR_016547
Musa___GSMUA_Achr5P16690_001
Musa___GSMUA_Achr4P08940_001
100
Amborella___AmTr_v1.0_scaffold00078.203

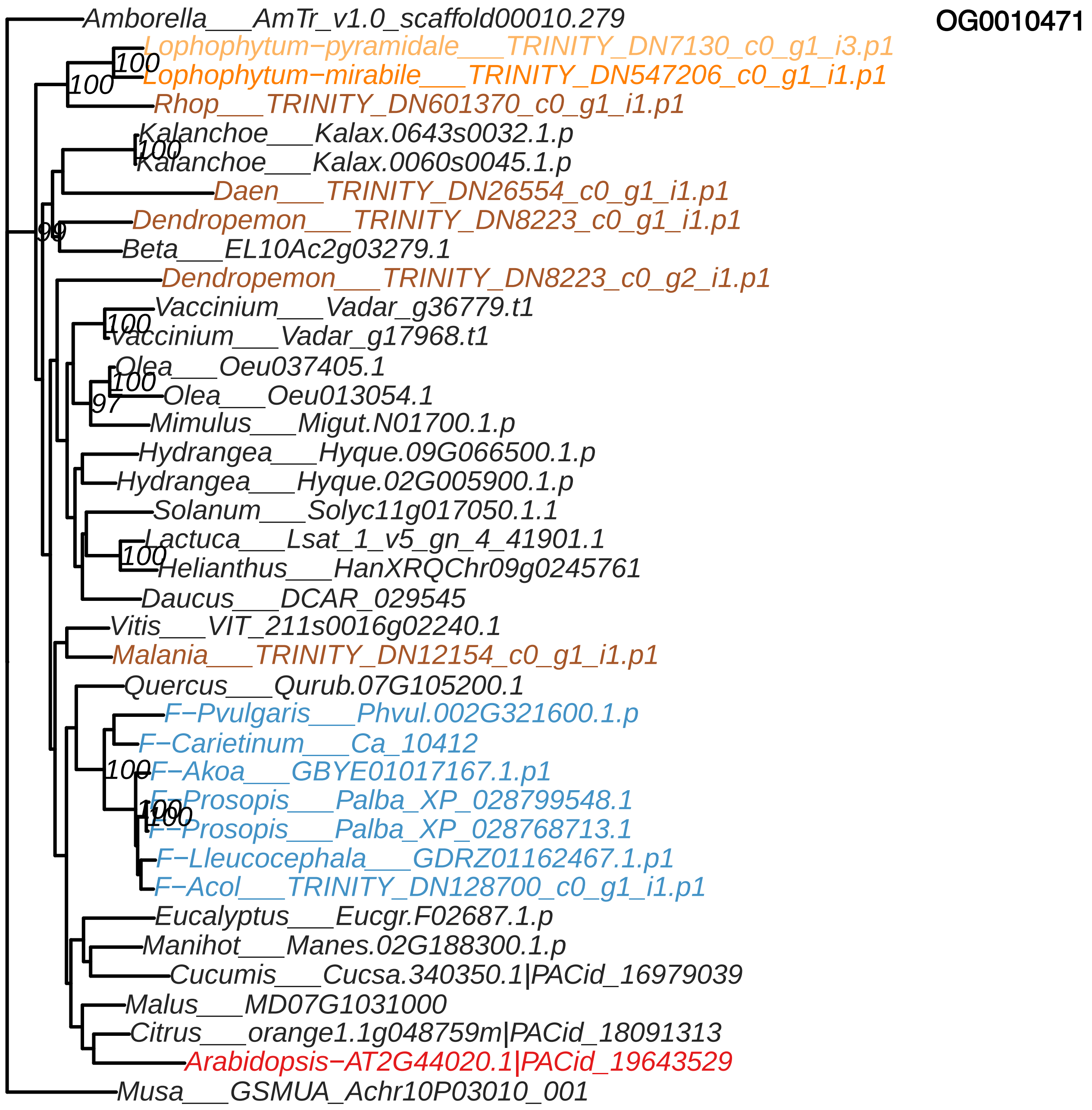
OG0010471
Amborella___AmTr_v1.0_scaffold00010.279
Lophophytum−pyramidale___TRINITY_DN7130_c0_g1_i3.p1
Lophophytum−mirabile___TRINITY_DN547206_c0_g1_i1.p1
Rhop___TRINITY_DN601370_c0_g1_i1.p1
Kalanchoe___Kalax.0643s0032.1.p
Kalanchoe___Kalax.0060s0045.1.p
Daen___TRINITY_DN26554_c0_g1_i1.p1
Dendropemon___TRINITY_DN8223_c0_g1_i1.p1
Beta___EL10Ac2g03279.1
Dendropemon___TRINITY_DN8223_c0_g2_i1.p1
Vaccinium___Vadar_g36779.t1
Vaccinium___Vadar_g17968.t1
Olea___Oeu037405.1
Olea___Oeu013054.1
Mimulus___Migut.N01700.1.p
Hydrangea___Hyque.09G066500.1.p
Hydrangea___Hyque.02G005900.1.p
Solanum___Solyc11g017050.1.1
Lactuca___Lsat_1_v5_gn_4_41901.1
Helianthus___HanXRQChr09g0245761
Daucus___DCAR_029545
Vitis___VIT_211s0016g02240.1
Malania___TRINITY_DN12154_c0_g1_i1.p1
Quercus___Qurub.07G105200.1
F−Pvulgaris___Phvul.002G321600.1.p
F−Carietinum___Ca_10412
F−Akoa___GBYE01017167.1.p1
F−Prosopis___Palba_XP_028799548.1
F−Prosopis___Palba_XP_028768713.1
F−Lleucocephala___GDRZ01162467.1.p1
F−Acol___TRINITY_DN128700_c0_g1_i1.p1
Eucalyptus___Eucgr.F02687.1.p
Manihot___Manes.02G188300.1.p
Cucumis___Cucsa.340350.1|PACid_16979039
Malus___MD07G1031000
Citrus___orange1.1g048759m|PACid_18091313
Arabidopsis−AT2G44020.1|PACid_19643529
Musa___GSMUA_Achr10P03010_001
100
100
100
99
100
100
97
100
100
100
100

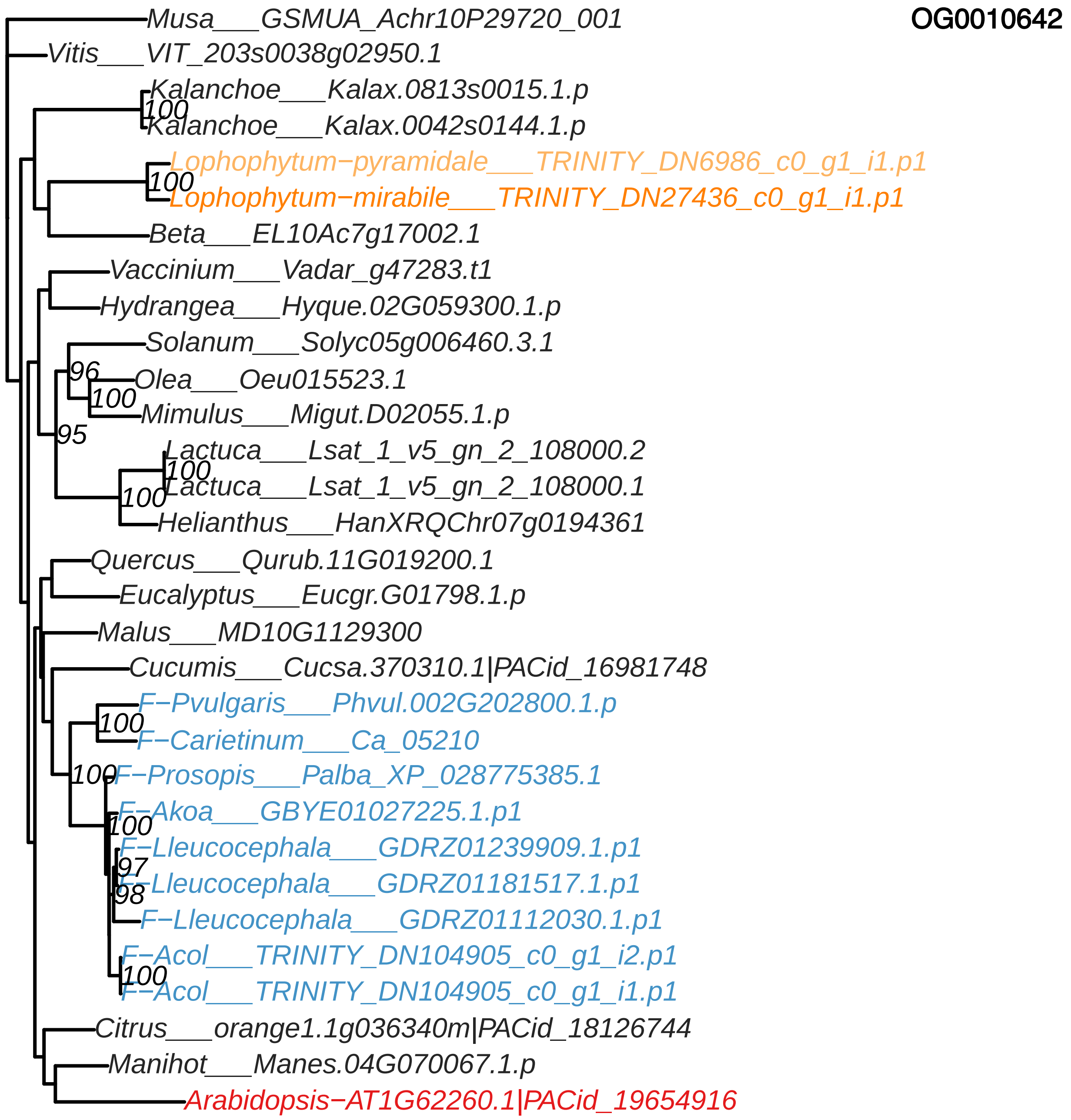
OG0010642
Musa___GSMUA_Achr10P29720_001
Vitis___VIT_203s0038g02950.1
Kalanchoe___Kalax.0813s0015.1.p
100
Kalanchoe___Kalax.0042s0144.1.p
Lophophytum−pyramidale___TRINITY_DN6986_c0_g1_i1.p1
100
Lophophytum−mirabile___TRINITY_DN27436_c0_g1_i1.p1
Beta___EL10Ac7g17002.1
Vaccinium___Vadar_g47283.t1
Hydrangea___Hyque.02G059300.1.p
Solanum___Solyc05g006460.3.1
96
Olea___Oeu015523.1
100
Mimulus___Migut.D02055.1.p
95
Lactuca___Lsat_1_v5_gn_2_108000.2
100
Lactuca___Lsat_1_v5_gn_2_108000.1
100
Helianthus___HanXRQChr07g0194361
Quercus___Qurub.11G019200.1
Eucalyptus___Eucgr.G01798.1.p
Malus___MD10G1129300
Cucumis___Cucsa.370310.1|PACid_16981748
F−Pvulgaris___Phvul.002G202800.1.p
100
F−Carietinum___Ca_05210
100
F−Prosopis___Palba_XP_028775385.1
F−Akoa___GBYE01027225.1.p1
100
F−Lleucocephala___GDRZ01239909.1.p1
97
F−Lleucocephala___GDRZ01181517.1.p1
98
F−Lleucocephala___GDRZ01112030.1.p1
F−Acol___TRINITY_DN104905_c0_g1_i2.p1
100
F−Acol___TRINITY_DN104905_c0_g1_i1.p1
Citrus___orange1.1g036340m|PACid_18126744
Manihot___Manes.04G070067.1.p
Arabidopsis−AT1G62260.1|PACid_19654916

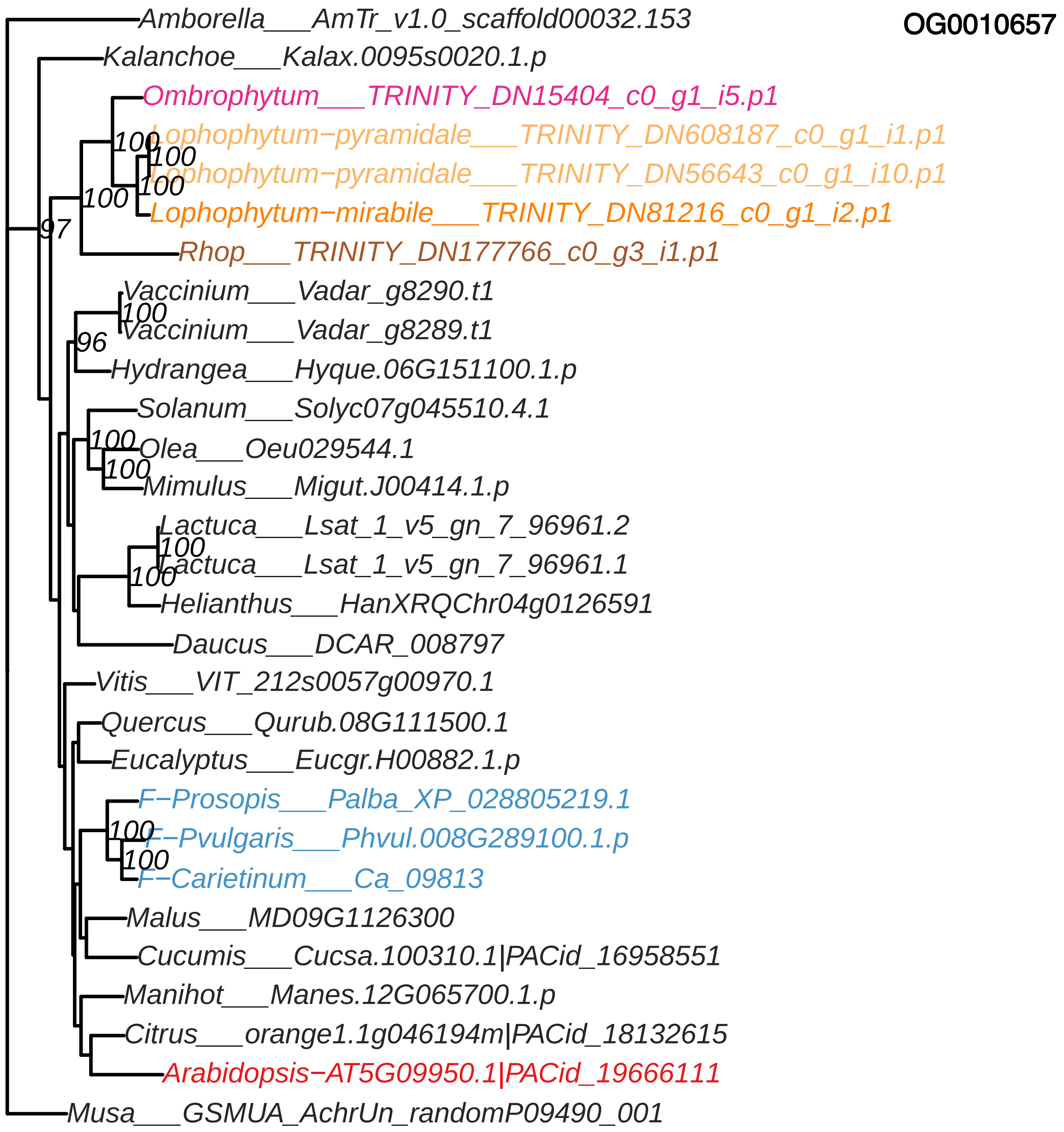
OG0010657
Amborella___AmTr_v1.0_scaffold00032.153
Kalanchoe___Kalax.0095s0020.1.p
Ombrophytum___TRINITY_DN15404_c0_g1_i5.p1
Lophophytum−pyramidale___TRINITY_DN608187_c0_g1_i1.p1
Lophophytum−pyramidale___TRINITY_DN56643_c0_g1_i10.p1
Lophophytum−mirabile___TRINITY_DN81216_c0_g1_i2.p1
Rhop___TRINITY_DN177766_c0_g3_i1.p1
Vaccinium___Vadar_g8290.t1
Vaccinium___Vadar_g8289.t1
Hydrangea___Hyque.06G151100.1.p
Solanum___Solyc07g045510.4.1
Olea___Oeu029544.1
Mimulus___Migut.J00414.1.p
Lactuca___Lsat_1_v5_gn_7_96961.2
Lactuca___Lsat_1_v5_gn_7_96961.1
Helianthus___HanXRQChr04g0126591
Daucus___DCAR_008797
Vitis___VIT_212s0057g00970.1
Quercus___Qurub.08G111500.1
Eucalyptus___Eucgr.H00882.1.p
F−Prosopis___Palba_XP_028805219.1
F−Pvulgaris___Phvul.008G289100.1.p
F−Carietinum___Ca_09813
Malus___MD09G1126300
Cucumis___Cucsa.100310.1|PACid_16958551
Manihot___Manes.12G065700.1.p
Citrus___orange1.1g046194m|PACid_18132615
Arabidopsis−AT5G09950.1|PACid_19666111
Musa___GSMUA_AchrUn_randomP09490_001
100
100
100
100
97
100
96
100
100
100
100
100
100

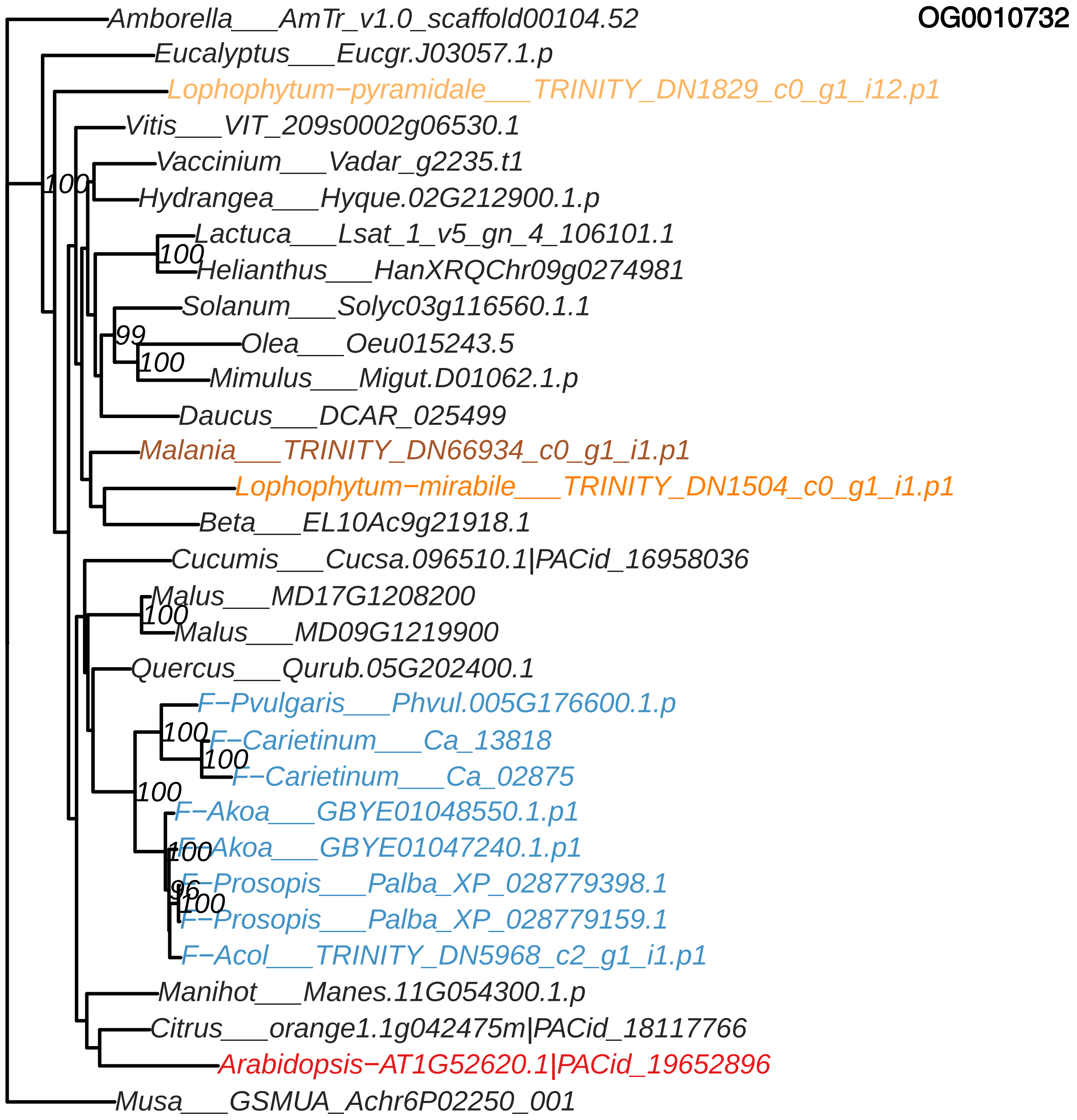
OG0010732
Amborella___AmTr_v1.0_scaffold00104.52
Eucalyptus___Eucgr.J03057.1.p
Lophophytum−pyramidale___TRINITY_DN1829_c0_g1_i12.p1
Vitis___VIT_209s0002g06530.1
Vaccinium___Vadar_g2235.t1
Hydrangea___Hyque.02G212900.1.p
Lactuca___Lsat_1_v5_gn_4_106101.1
Helianthus___HanXRQChr09g0274981
Solanum___Solyc03g116560.1.1
Olea___Oeu015243.5
Mimulus___Migut.D01062.1.p
Daucus___DCAR_025499
Malania___TRINITY_DN66934_c0_g1_i1.p1
Lophophytum−mirabile___TRINITY_DN1504_c0_g1_i1.p1
Beta___EL10Ac9g21918.1
Cucumis___Cucsa.096510.1|PACid_16958036
Malus___MD17G1208200
Malus___MD09G1219900
Quercus___Qurub.05G202400.1
F−Pvulgaris___Phvul.005G176600.1.p
F−Carietinum___Ca_13818
F−Carietinum___Ca_02875
F−Akoa___GBYE01048550.1.p1
F−Akoa___GBYE01047240.1.p1
F−Prosopis___Palba_XP_028779398.1
F−Prosopis___Palba_XP_028779159.1
F−Acol___TRINITY_DN5968_c2_g1_i1.p1
Manihot___Manes.11G054300.1.p
Citrus___orange1.1g042475m|PACid_18117766
Arabidopsis−AT1G52620.1|PACid_19652896
Musa___GSMUA_Achr6P02250_001
100
100
99
100
100
100
100
100
100
96
100

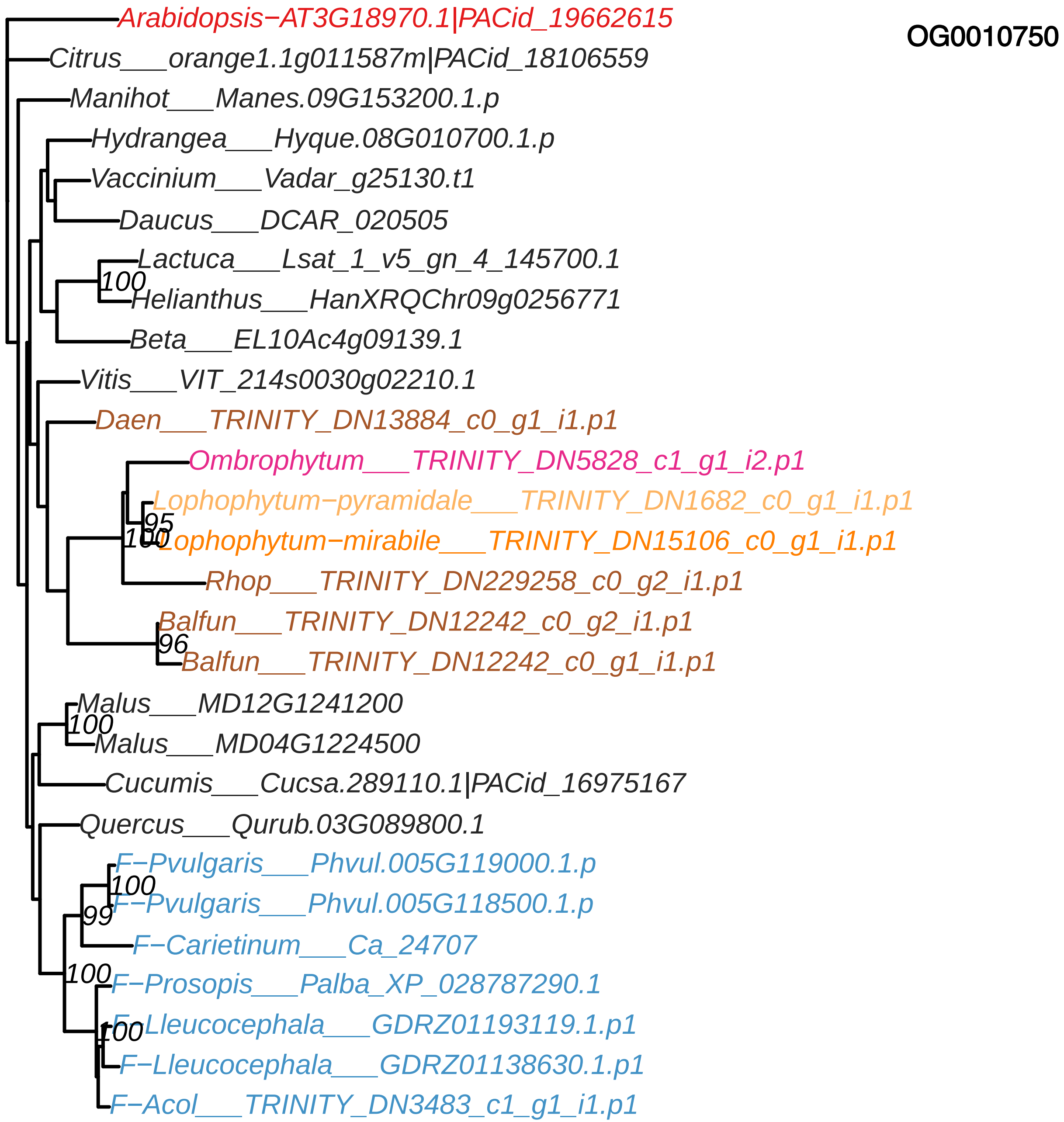
OG0010750
Arabidopsis−AT3G18970.1|PACid_19662615
Citrus___orange1.1g011587m|PACid_18106559
Manihot___Manes.09G153200.1.p
Hydrangea___Hyque.08G010700.1.p
Vaccinium___Vadar_g25130.t1
Daucus___DCAR_020505
Lactuca___Lsat_1_v5_gn_4_145700.1
100
Helianthus___HanXRQChr09g0256771
Beta___EL10Ac4g09139.1
Vitis___VIT_214s0030g02210.1
Daen___TRINITY_DN13884_c0_g1_i1.p1
Ombrophytum___TRINITY_DN5828_c1_g1_i2.p1
Lophophytum−pyramidale___TRINITY_DN1682_c0_g1_i1.p1
95
100
Lophophytum−mirabile___TRINITY_DN15106_c0_g1_i1.p1
Rhop___TRINITY_DN229258_c0_g2_i1.p1
Balfun___TRINITY_DN12242_c0_g2_i1.p1
96
Balfun___TRINITY_DN12242_c0_g1_i1.p1
Malus___MD12G1241200
100
Malus___MD04G1224500
Cucumis___Cucsa.289110.1|PACid_16975167
Quercus___Qurub.03G089800.1
F−Pvulgaris___Phvul.005G119000.1.p
100
F−Pvulgaris___Phvul.005G118500.1.p
99
F−Carietinum___Ca_24707
100
F−Prosopis___Palba_XP_028787290.1
F−Lleucocephala___GDRZ01193119.1.p1
100
F−Lleucocephala___GDRZ01138630.1.p1
F−Acol___TRINITY_DN3483_c1_g1_i1.p1

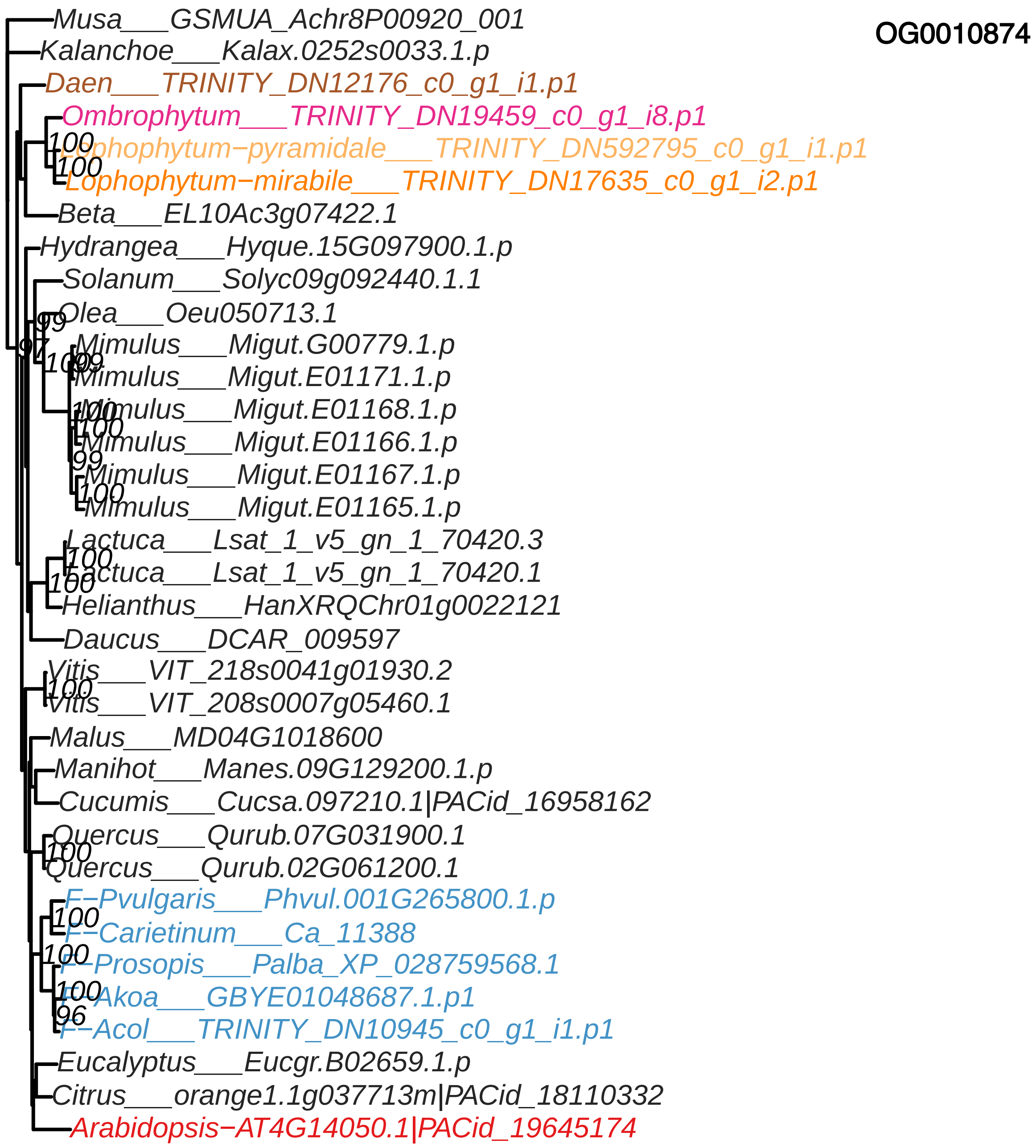
OG0010874
Musa___GSMUA_Achr8P00920_001
Kalanchoe___Kalax.0252s0033.1.p
Daen___TRINITY_DN12176_c0_g1_i1.p1
Ombrophytum___TRINITY_DN19459_c0_g1_i8.p1
Lophophytum−pyramidale___TRINITY_DN592795_c0_g1_i1.p1
Lophophytum−mirabile___TRINITY_DN17635_c0_g1_i2.p1
Beta___EL10Ac3g07422.1
Hydrangea___Hyque.15G097900.1.p
Solanum___Solyc09g092440.1.1
Olea___Oeu050713.1
Mimulus___Migut.G00779.1.p
Mimulus___Migut.E01171.1.p
Mimulus___Migut.E01168.1.p
Mimulus___Migut.E01166.1.p
Mimulus___Migut.E01167.1.p
Mimulus___Migut.E01165.1.p
Lactuca___Lsat_1_v5_gn_1_70420.3
Lactuca___Lsat_1_v5_gn_1_70420.1
Helianthus___HanXRQChr01g0022121
Daucus___DCAR_009597
Vitis___VIT_218s0041g01930.2
Vitis___VIT_208s0007g05460.1
Malus___MD04G1018600
Manihot___Manes.09G129200.1.p
Cucumis___Cucsa.097210.1|PACid_16958162
Quercus___Qurub.07G031900.1
Quercus___Qurub.02G061200.1
F−Pvulgaris___Phvul.001G265800.1.p
F−Carietinum___Ca_11388
F−Prosopis___Palba_XP_028759568.1
F−Akoa___GBYE01048687.1.p1
F−Acol___TRINITY_DN10945_c0_g1_i1.p1
Eucalyptus___Eucgr.B02659.1.p
Citrus___orange1.1g037713m|PACid_18110332
Arabidopsis−AT4G14050.1|PACid_19645174
100
100
99
97
100
99
100
100
99
100
100
100
100
100
100
100
100
96

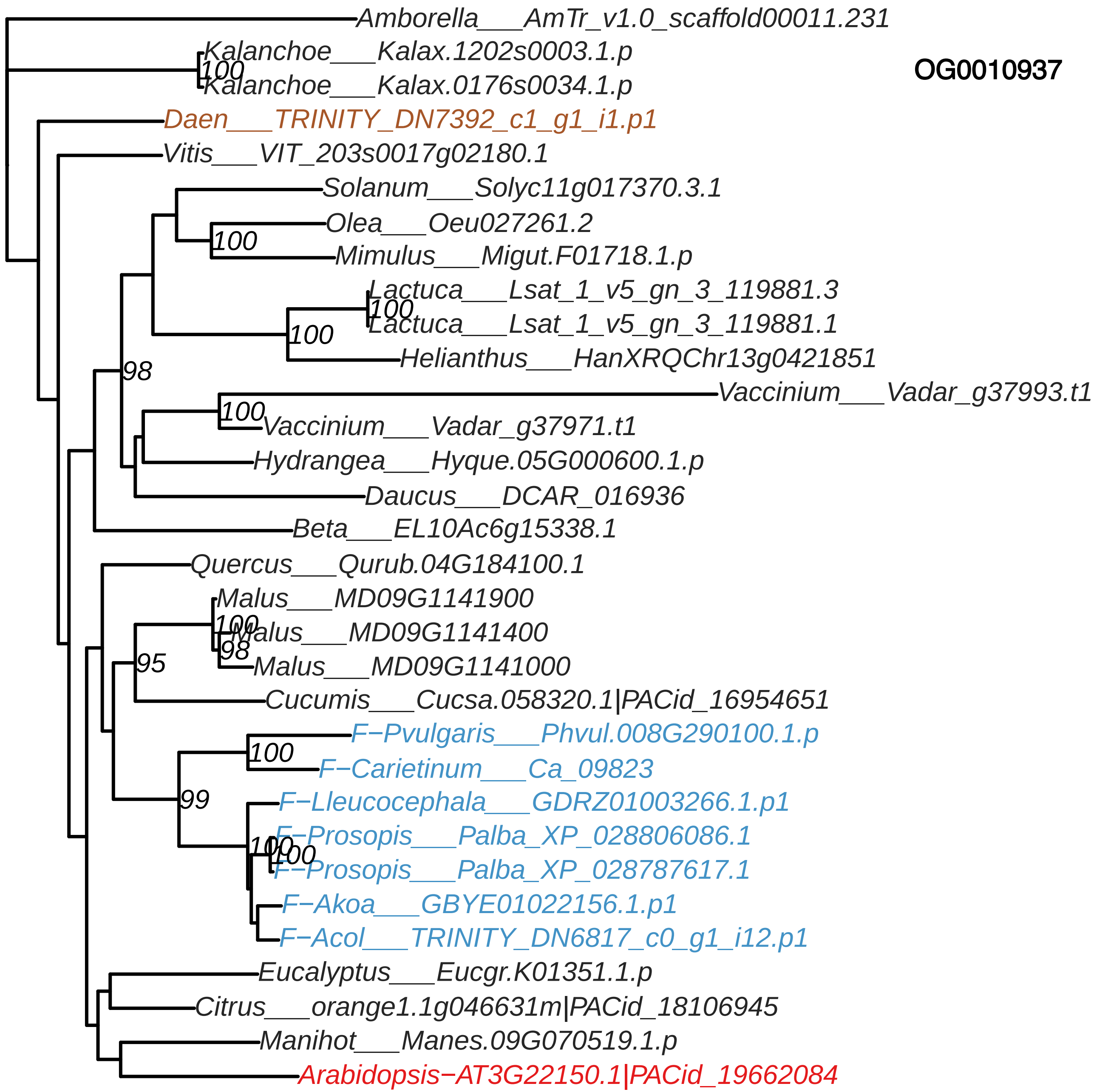
OG0010937
Amborella___AmTr_v1.0_scaffold00011.231
Kalanchoe___Kalax.1202s0003.1.p
100
Kalanchoe___Kalax.0176s0034.1.p
Daen___TRINITY_DN7392_c1_g1_i1.p1
Vitis___VIT_203s0017g02180.1
Solanum___Solyc11g017370.3.1
Olea___Oeu027261.2
100
Mimulus___Migut.F01718.1.p
Lactuca___Lsat_1_v5_gn_3_119881.3
100
Lactuca___Lsat_1_v5_gn_3_119881.1
100
Helianthus___HanXRQChr13g0421851
98
Vaccinium___Vadar_g37993.t1
100
Vaccinium___Vadar_g37971.t1
Hydrangea___Hyque.05G000600.1.p
Daucus___DCAR_016936
Beta___EL10Ac6g15338.1
Quercus___Qurub.04G184100.1
Malus___MD09G1141900
100
Malus___MD09G1141400
98
95
Malus___MD09G1141000
Cucumis___Cucsa.058320.1|PACid_16954651
F−Pvulgaris___Phvul.008G290100.1.p
100
F−Carietinum___Ca_09823
99
F−Lleucocephala___GDRZ01003266.1.p1
F−Prosopis___Palba_XP_028806086.1
100
100
F−Prosopis___Palba_XP_028787617.1
F−Akoa___GBYE01022156.1.p1
F−Acol___TRINITY_DN6817_c0_g1_i12.p1
Eucalyptus___Eucgr.K01351.1.p
Citrus___orange1.1g046631m|PACid_18106945
Manihot___Manes.09G070519.1.p
Arabidopsis−AT3G22150.1|PACid_19662084

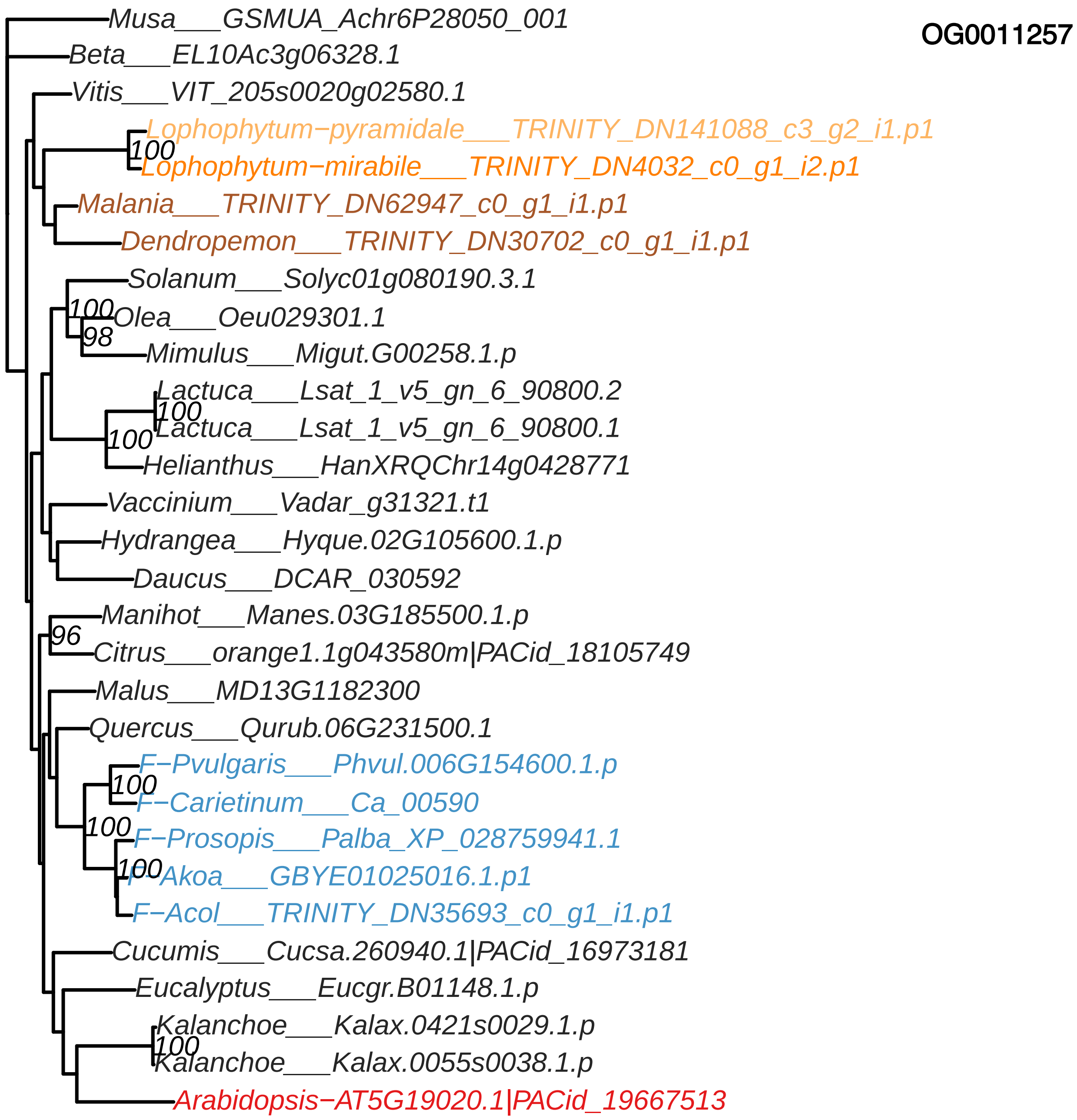
OG0011257
Musa___GSMUA_Achr6P28050_001
Beta___EL10Ac3g06328.1
Vitis___VIT_205s0020g02580.1
Lophophytum−pyramidale___TRINITY_DN141088_c3_g2_i1.p1
100
Lophophytum−mirabile___TRINITY_DN4032_c0_g1_i2.p1
Malania___TRINITY_DN62947_c0_g1_i1.p1
Dendropemon___TRINITY_DN30702_c0_g1_i1.p1
Solanum___Solyc01g080190.3.1
100
Olea___Oeu029301.1
98
Mimulus___Migut.G00258.1.p
Lactuca___Lsat_1_v5_gn_6_90800.2
100
Lactuca___Lsat_1_v5_gn_6_90800.1
100
Helianthus___HanXRQChr14g0428771
Vaccinium___Vadar_g31321.t1
Hydrangea___Hyque.02G105600.1.p
Daucus___DCAR_030592
Manihot___Manes.03G185500.1.p
96
Citrus___orange1.1g043580m|PACid_18105749
Malus___MD13G1182300
Quercus___Qurub.06G231500.1
F−Pvulgaris___Phvul.006G154600.1.p
100
F−Carietinum___Ca_00590
100
F−Prosopis___Palba_XP_028759941.1
100
F−Akoa___GBYE01025016.1.p1
F−Acol___TRINITY_DN35693_c0_g1_i1.p1
Cucumis___Cucsa.260940.1|PACid_16973181
Eucalyptus___Eucgr.B01148.1.p
Kalanchoe___Kalax.0421s0029.1.p
100
Kalanchoe___Kalax.0055s0038.1.p
Arabidopsis−AT5G19020.1|PACid_19667513

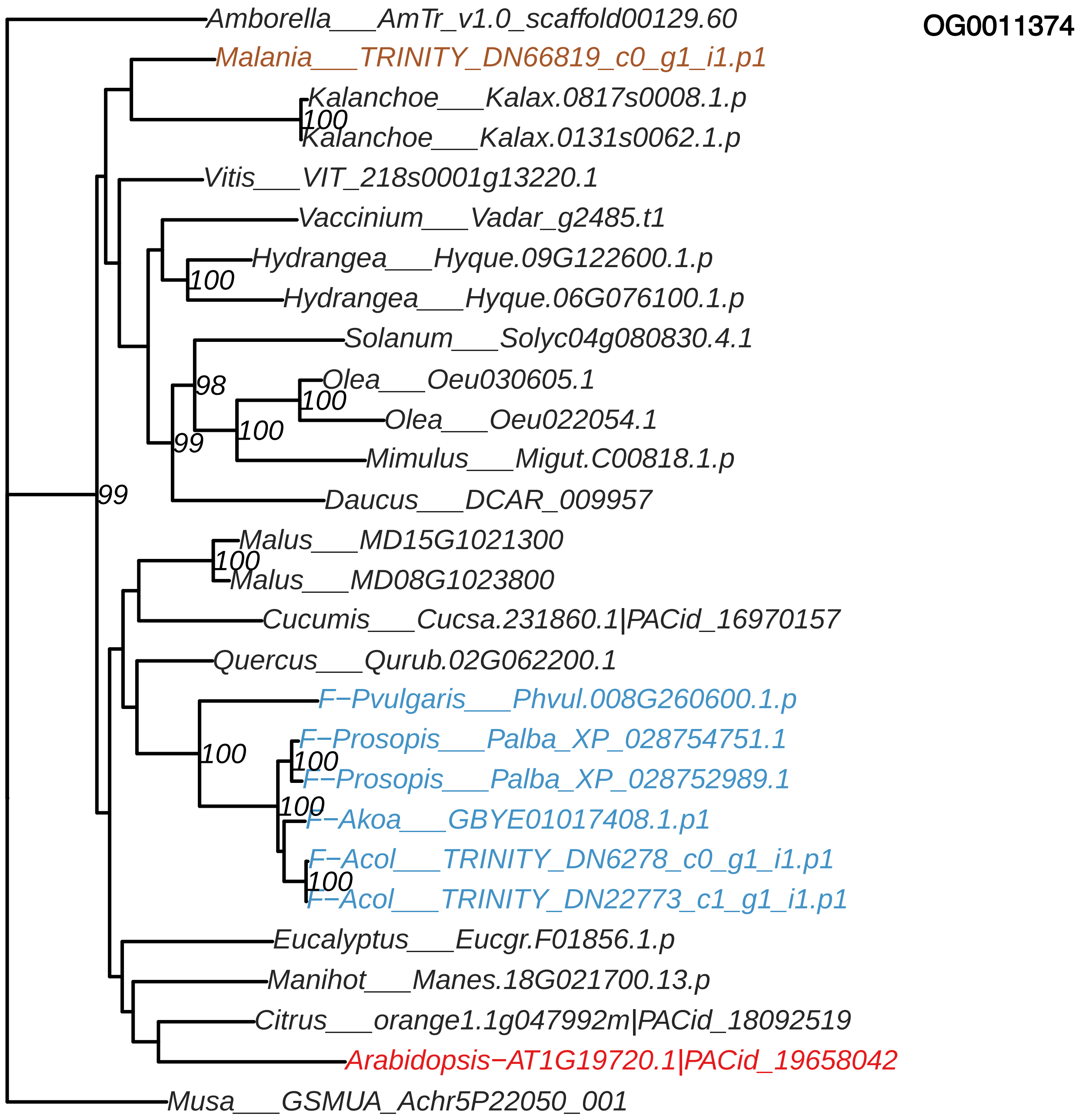

OG0011374
Amborella___AmTr_v1.0_scaffold00129.60
Malania___TRINITY_DN66819_c0_g1_i1.p1
Kalanchoe___Kalax.0817s0008.1.p
100
Kalanchoe___Kalax.0131s0062.1.p
Vitis___VIT_218s0001g13220.1
Vaccinium___Vadar_g2485.t1
Hydrangea___Hyque.09G122600.1.p
100
Hydrangea___Hyque.06G076100.1.p
Solanum___Solyc04g080830.4.1
98
Olea___Oeu030605.1
100
Olea___Oeu022054.1
100
Mimulus___Migut.C00818.1.p
99
Daucus___DCAR_009957
99
Malus___MD15G1021300
100
Malus___MD08G1023800
Cucumis___Cucsa.231860.1|PACid_16970157
Quercus___Qurub.02G062200.1
F−Pvulgaris___Phvul.008G260600.1.p
100
F−Prosopis___Palba_XP_028754751.1
100
F−Prosopis___Palba_XP_028752989.1
100
F−Akoa___GBYE01017408.1.p1
F−Acol___TRINITY_DN6278_c0_g1_i1.p1
100
F−Acol___TRINITY_DN22773_c1_g1_i1.p1
Eucalyptus___Eucgr.F01856.1.p
Manihot___Manes.18G021700.13.p
Citrus___orange1.1g047992m|PACid_18092519
Arabidopsis−AT1G19720.1|PACid_19658042
Musa___GSMUA_Achr5P22050_001

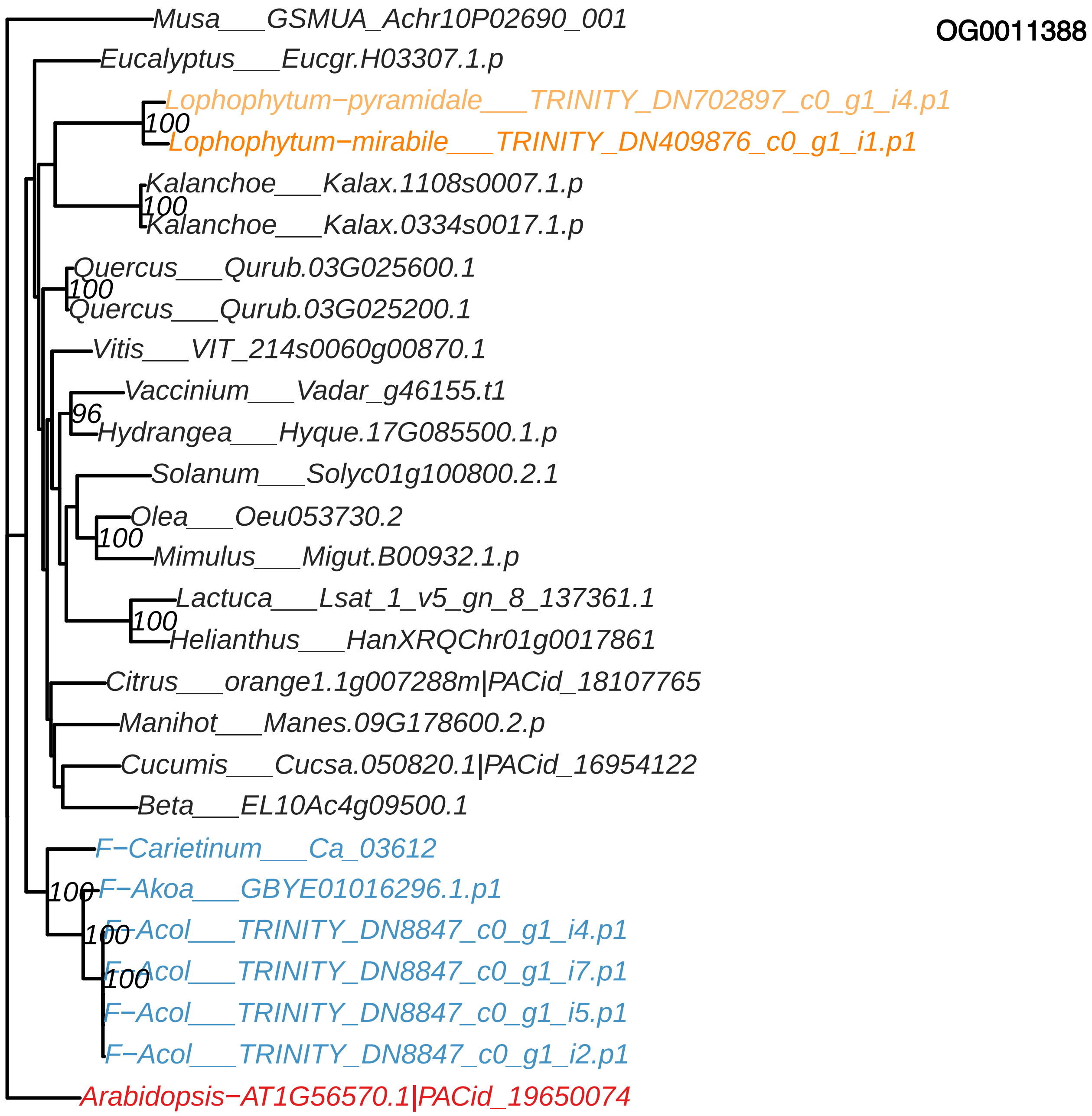
OG0011388
Musa___GSMUA_Achr10P02690_001
Eucalyptus___Eucgr.H03307.1.p
Lophophytum−pyramidale___TRINITY_DN702897_c0_g1_i4.p1
100
Lophophytum−mirabile___TRINITY_DN409876_c0_g1_i1.p1
Kalanchoe___Kalax.1108s0007.1.p
100
Kalanchoe___Kalax.0334s0017.1.p
Quercus___Qurub.03G025600.1
100
Quercus___Qurub.03G025200.1
Vitis___VIT_214s0060g00870.1
Vaccinium___Vadar_g46155.t1
96
Hydrangea___Hyque.17G085500.1.p
Solanum___Solyc01g100800.2.1
Olea___Oeu053730.2
100
Mimulus___Migut.B00932.1.p
Lactuca___Lsat_1_v5_gn_8_137361.1
100
Helianthus___HanXRQChr01g0017861
Citrus___orange1.1g007288m|PACid_18107765
Manihot___Manes.09G178600.2.p
Cucumis___Cucsa.050820.1|PACid_16954122
Beta___EL10Ac4g09500.1
F−Carietinum___Ca_03612
100
F−Akoa___GBYE01016296.1.p1
100
F−Acol___TRINITY_DN8847_c0_g1_i4.p1
100
F−Acol___TRINITY_DN8847_c0_g1_i7.p1
F−Acol___TRINITY_DN8847_c0_g1_i5.p1
F−Acol___TRINITY_DN8847_c0_g1_i2.p1
Arabidopsis−AT1G56570.1|PACid_19650074

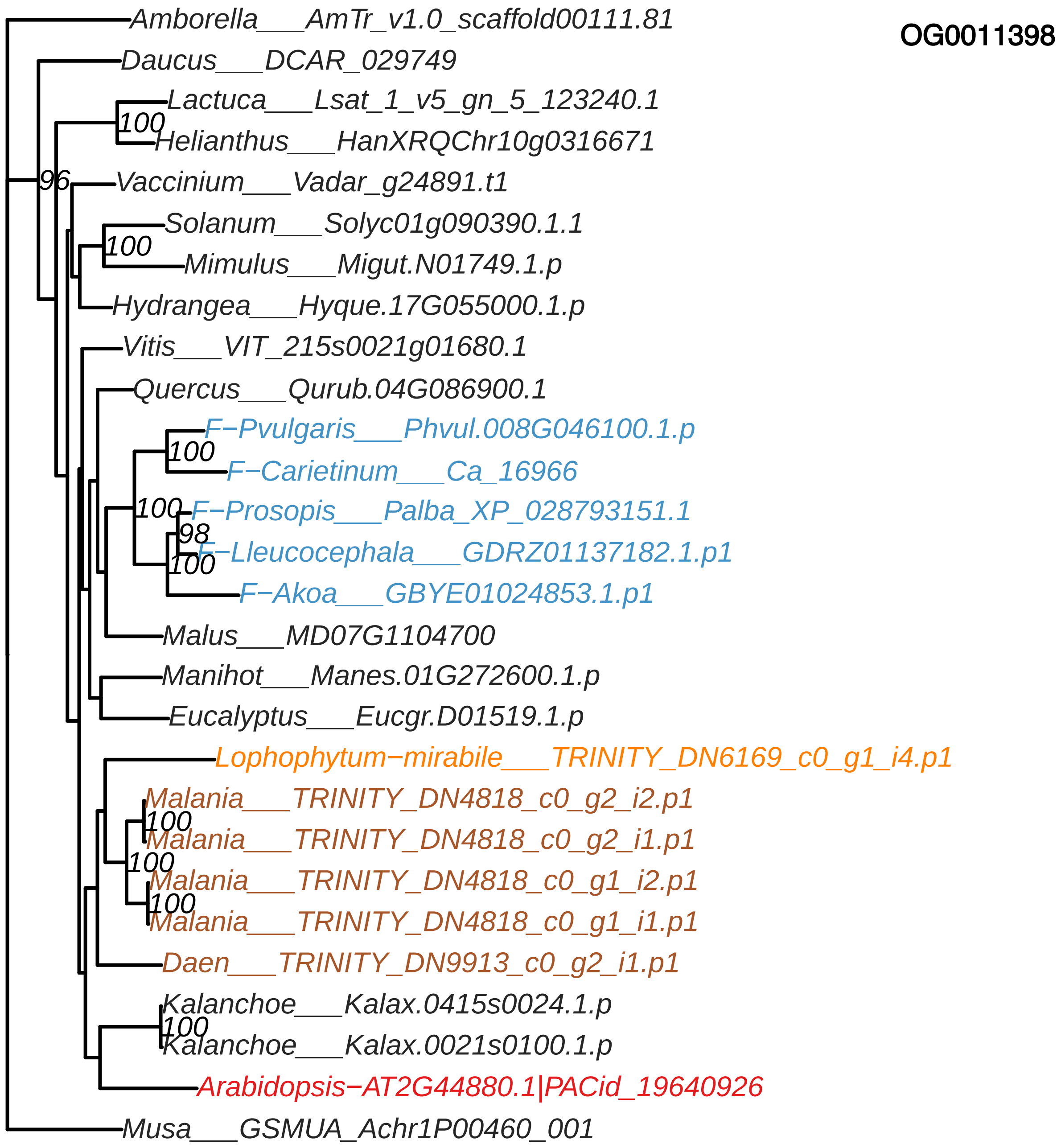
OG0011398
Amborella___AmTr_v1.0_scaffold00111.81
Daucus___DCAR_029749
Lactuca___Lsat_1_v5_gn_5_123240.1
100
Helianthus___HanXRQChr10g0316671
96
Vaccinium___Vadar_g24891.t1
Solanum___Solyc01g090390.1.1
100
Mimulus___Migut.N01749.1.p
Hydrangea___Hyque.17G055000.1.p
Vitis___VIT_215s0021g01680.1
Quercus___Qurub.04G086900.1
F−Pvulgaris___Phvul.008G046100.1.p
100
F−Carietinum___Ca_16966
100
F−Prosopis___Palba_XP_028793151.1
98
F−Lleucocephala___GDRZ01137182.1.p1
100
F−Akoa___GBYE01024853.1.p1
Malus___MD07G1104700
Manihot___Manes.01G272600.1.p
Eucalyptus___Eucgr.D01519.1.p
Lophophytum−mirabile___TRINITY_DN6169_c0_g1_i4.p1
Malania___TRINITY_DN4818_c0_g2_i2.p1
100
Malania___TRINITY_DN4818_c0_g2_i1.p1
100
Malania___TRINITY_DN4818_c0_g1_i2.p1
100
Malania___TRINITY_DN4818_c0_g1_i1.p1
Daen___TRINITY_DN9913_c0_g2_i1.p1
Kalanchoe___Kalax.0415s0024.1.p
100
Kalanchoe___Kalax.0021s0100.1.p
Arabidopsis−AT2G44880.1|PACid_19640926
Musa___GSMUA_Achr1P00460_001

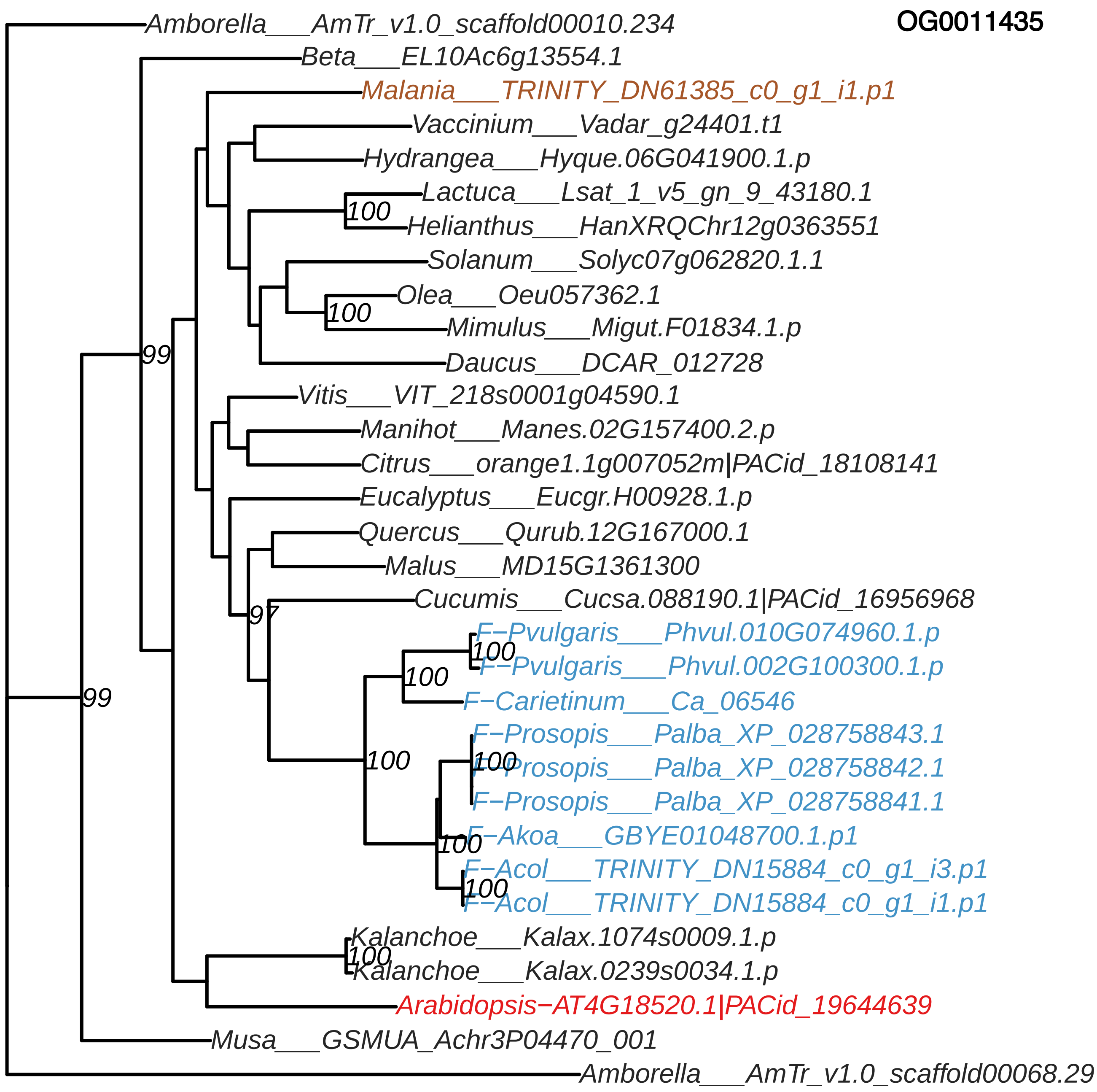
OG0011435
Amborella___AmTr_v1.0_scaffold00010.234
Beta___EL10Ac6g13554.1
Malania___TRINITY_DN61385_c0_g1_i1.p1
Vaccinium___Vadar_g24401.t1
Hydrangea___Hyque.06G041900.1.p
Lactuca___Lsat_1_v5_gn_9_43180.1
100
Helianthus___HanXRQChr12g0363551
Solanum___Solyc07g062820.1.1
Olea___Oeu057362.1
100
Mimulus___Migut.F01834.1.p
99
Daucus___DCAR_012728
Vitis___VIT_218s0001g04590.1
Manihot___Manes.02G157400.2.p
Citrus___orange1.1g007052m|PACid_18108141
Eucalyptus___Eucgr.H00928.1.p
Quercus___Qurub.12G167000.1
Malus___MD15G1361300
Cucumis___Cucsa.088190.1|PACid_16956968
97
F−Pvulgaris___Phvul.010G074960.1.p
100
F−Pvulgaris___Phvul.002G100300.1.p
100
99
F−Carietinum___Ca_06546
F−Prosopis___Palba_XP_028758843.1
100
F−Prosopis___Palba_XP_028758842.1
F−Prosopis___Palba_XP_028758841.1
F−Akoa___GBYE01048700.1.p1
100
F−Acol___TRINITY_DN15884_c0_g1_i3.p1
100
F−Acol___TRINITY_DN15884_c0_g1_i1.p1
Kalanchoe___Kalax.1074s0009.1.p
100
Kalanchoe___Kalax.0239s0034.1.p
Arabidopsis−AT4G18520.1|PACid_19644639
Musa___GSMUA_Achr3P04470_001
Amborella___AmTr_v1.0_scaffold00068.29

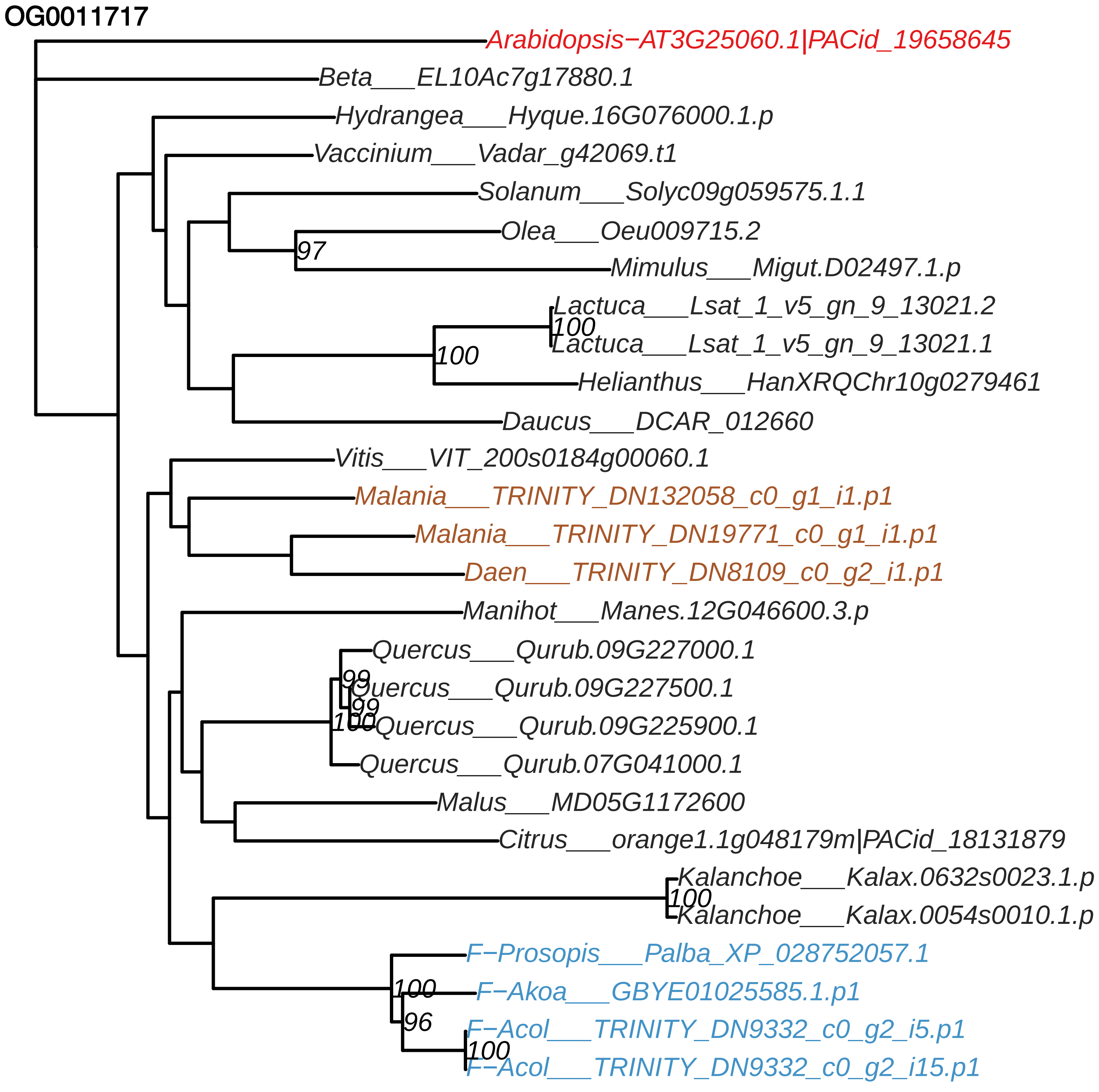
OG0011717
Arabidopsis−AT3G25060.1|PACid_19658645
Beta___EL10Ac7g17880.1
Hydrangea___Hyque.16G076000.1.p
Vaccinium___Vadar_g42069.t1
Solanum___Solyc09g059575.1.1
Olea___Oeu009715.2
97
Mimulus___Migut.D02497.1.p
Lactuca___Lsat_1_v5_gn_9_13021.2
100
Lactuca___Lsat_1_v5_gn_9_13021.1
100
Helianthus___HanXRQChr10g0279461
Daucus___DCAR_012660
Vitis___VIT_200s0184g00060.1
Malania___TRINITY_DN132058_c0_g1_i1.p1
Malania___TRINITY_DN19771_c0_g1_i1.p1
Daen___TRINITY_DN8109_c0_g2_i1.p1
Manihot___Manes.12G046600.3.p
Quercus___Qurub.09G227000.1
99
Quercus___Qurub.09G227500.1
99
100
Quercus___Qurub.09G225900.1
Quercus___Qurub.07G041000.1
Malus___MD05G1172600
Citrus___orange1.1g048179m|PACid_18131879
Kalanchoe___Kalax.0632s0023.1.p
100
Kalanchoe___Kalax.0054s0010.1.p
F−Prosopis___Palba_XP_028752057.1
100
F−Akoa___GBYE01025585.1.p1
96
F−Acol___TRINITY_DN9332_c0_g2_i5.p1
100
F−Acol___TRINITY_DN9332_c0_g2_i15.p1

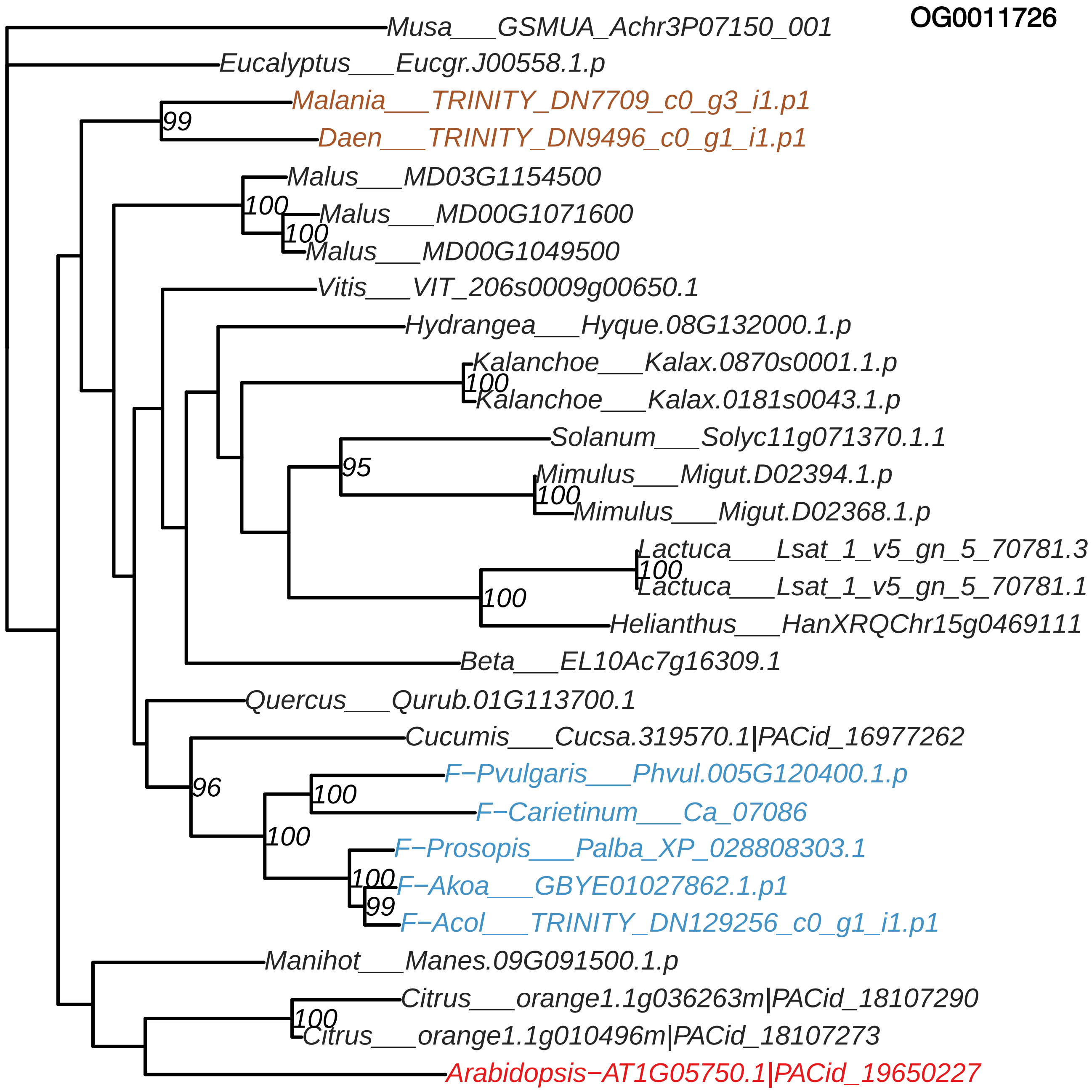

OG0011726
Musa___GSMUA_Achr3P07150_001
Eucalyptus___Eucgr.J00558.1.p
Malania___TRINITY_DN7709_c0_g3_i1.p1
99
Daen___TRINITY_DN9496_c0_g1_i1.p1
Malus___MD03G1154500
100
Malus___MD00G1071600
100
Malus___MD00G1049500
Vitis___VIT_206s0009g00650.1
Hydrangea___Hyque.08G132000.1.p
Kalanchoe___Kalax.0870s0001.1.p
100
Kalanchoe___Kalax.0181s0043.1.p
Solanum___Solyc11g071370.1.1
95
Mimulus___Migut.D02394.1.p
100
Mimulus___Migut.D02368.1.p
Lactuca___Lsat_1_v5_gn_5_70781.3
100
Lactuca___Lsat_1_v5_gn_5_70781.1
100
Helianthus___HanXRQChr15g0469111
Beta___EL10Ac7g16309.1
Quercus___Qurub.01G113700.1
Cucumis___Cucsa.319570.1|PACid_16977262
96
F−Pvulgaris___Phvul.005G120400.1.p
100
F−Carietinum___Ca_07086
100
F−Prosopis___Palba_XP_028808303.1
100
F−Akoa___GBYE01027862.1.p1
99
F−Acol___TRINITY_DN129256_c0_g1_i1.p1
Manihot___Manes.09G091500.1.p
Citrus___orange1.1g036263m|PACid_18107290
100
Citrus___orange1.1g010496m|PACid_18107273
Arabidopsis−AT1G05750.1|PACid_19650227

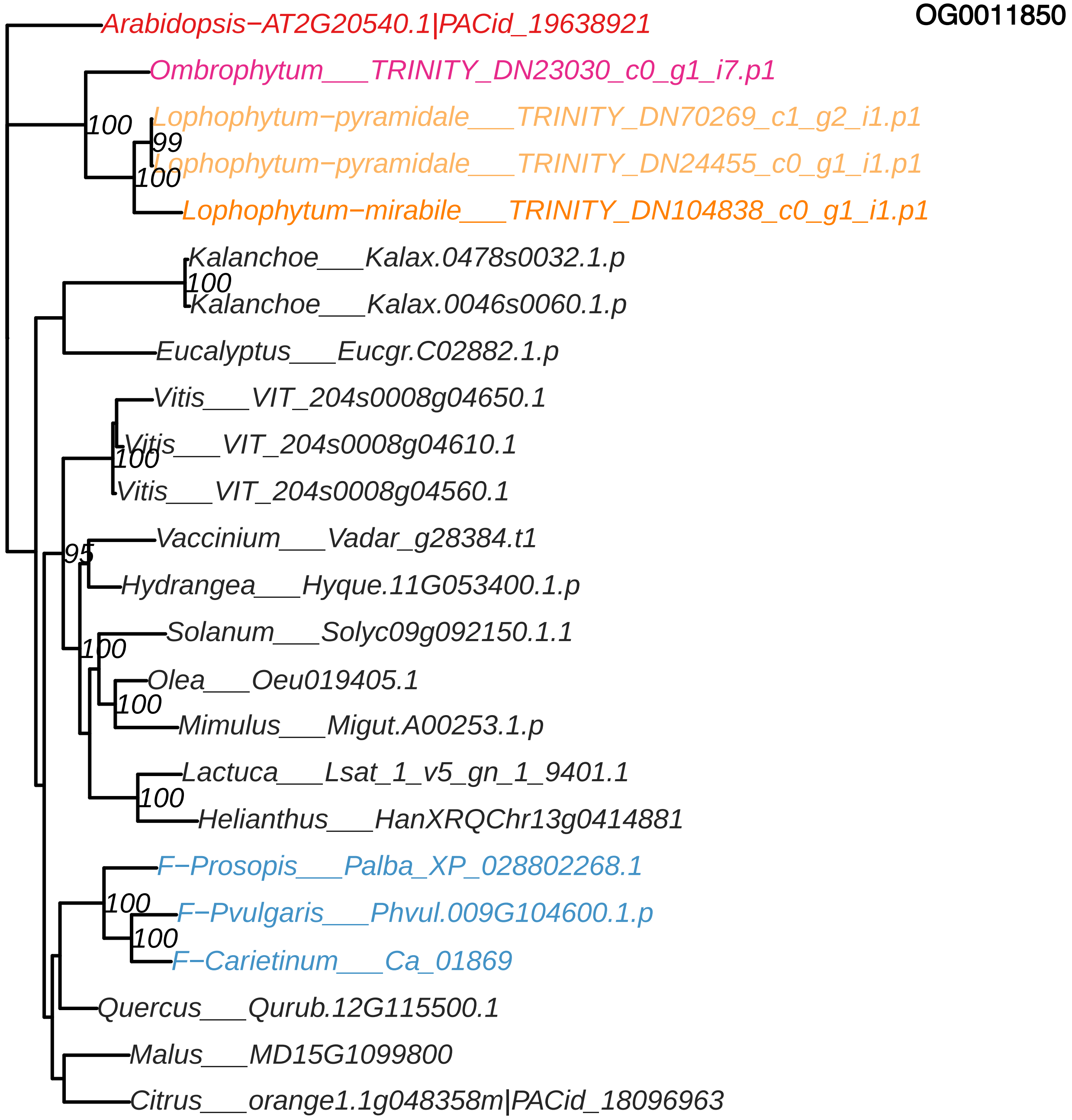
OG0011850
Arabidopsis−AT2G20540.1|PACid_19638921
Ombrophytum___TRINITY_DN23030_c0_g1_i7.p1
Lophophytum−pyramidale___TRINITY_DN70269_c1_g2_i1.p1
Lophophytum−pyramidale___TRINITY_DN24455_c0_g1_i1.p1
Lophophytum−mirabile___TRINITY_DN104838_c0_g1_i1.p1
Kalanchoe___Kalax.0478s0032.1.p
Kalanchoe___Kalax.0046s0060.1.p
Eucalyptus___Eucgr.C02882.1.p
Vitis___VIT_204s0008g04650.1
Vitis___VIT_204s0008g04610.1
Vitis___VIT_204s0008g04560.1
Vaccinium___Vadar_g28384.t1
Hydrangea___Hyque.11G053400.1.p
Solanum___Solyc09g092150.1.1
Olea___Oeu019405.1
Mimulus___Migut.A00253.1.p
Lactuca___Lsat_1_v5_gn_1_9401.1
Helianthus___HanXRQChr13g0414881
F−Prosopis___Palba_XP_028802268.1
F−Pvulgaris___Phvul.009G104600.1.p
F−Carietinum___Ca_01869
Quercus___Qurub.12G115500.1
Malus___MD15G1099800
Citrus___orange1.1g048358m|PACid_18096963
100
99
100
100
100
95
100
100
100
100
100

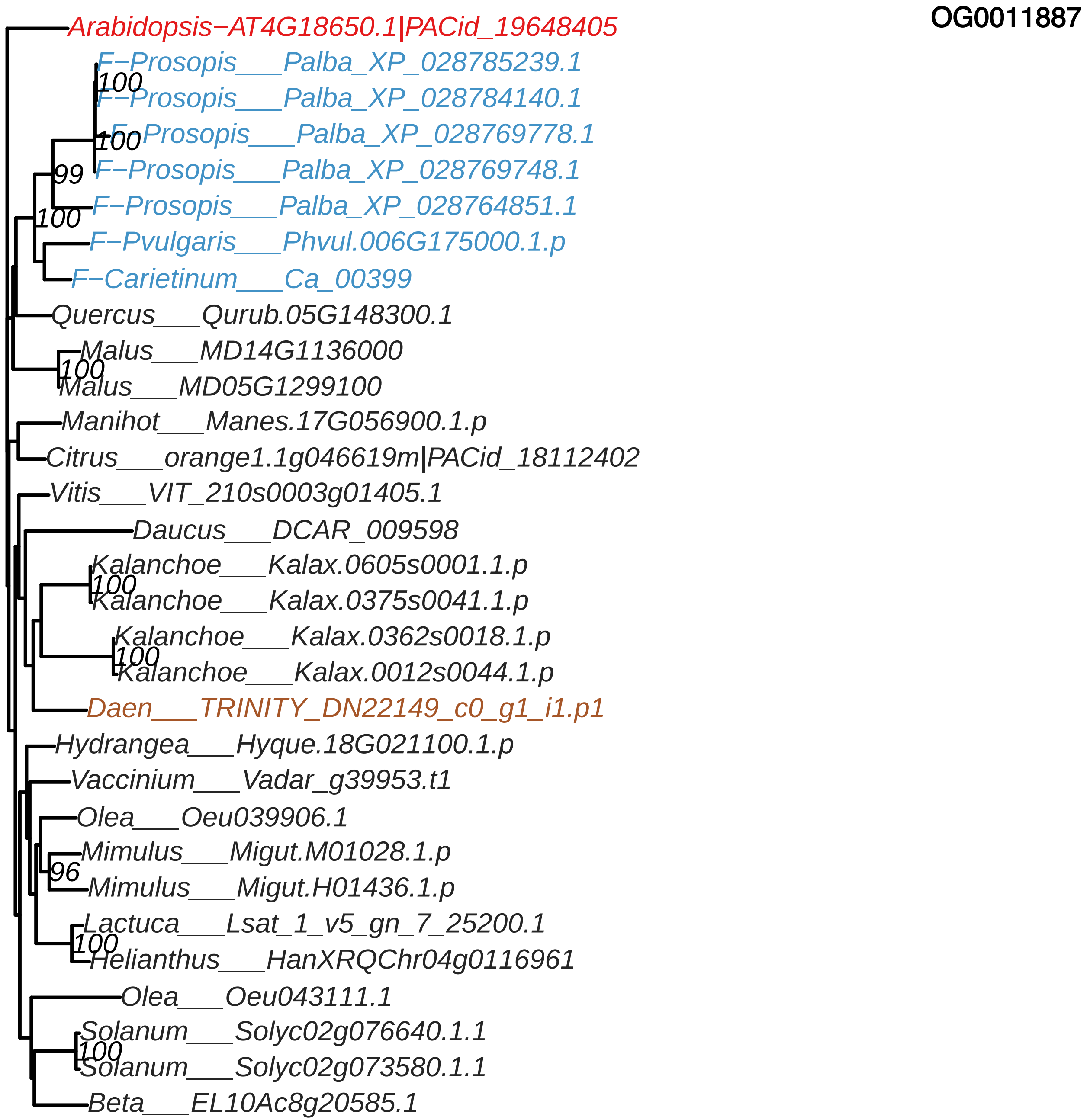
OG0011887
Arabidopsis−AT4G18650.1|PACid_19648405
F−Prosopis___Palba_XP_028785239.1
100
F−Prosopis___Palba_XP_028784140.1
F−Prosopis___Palba_XP_028769778.1
100
F−Prosopis___Palba_XP_028769748.1
99
F−Prosopis___Palba_XP_028764851.1
100
F−Pvulgaris___Phvul.006G175000.1.p
F−Carietinum___Ca_00399
Quercus___Qurub.05G148300.1
Malus___MD14G1136000
100
Malus___MD05G1299100
Manihot___Manes.17G056900.1.p
Citrus___orange1.1g046619m|PACid_18112402
Vitis___VIT_210s0003g01405.1
Daucus___DCAR_009598
Kalanchoe___Kalax.0605s0001.1.p
100
Kalanchoe___Kalax.0375s0041.1.p
Kalanchoe___Kalax.0362s0018.1.p
100
Kalanchoe___Kalax.0012s0044.1.p
Daen___TRINITY_DN22149_c0_g1_i1.p1
Hydrangea___Hyque.18G021100.1.p
Vaccinium___Vadar_g39953.t1
Olea___Oeu039906.1
Mimulus___Migut.M01028.1.p
96
Mimulus___Migut.H01436.1.p
Lactuca___Lsat_1_v5_gn_7_25200.1
100
Helianthus___HanXRQChr04g0116961
Olea___Oeu043111.1
Solanum___Solyc02g076640.1.1
100
Solanum___Solyc02g073580.1.1
Beta___EL10Ac8g20585.1

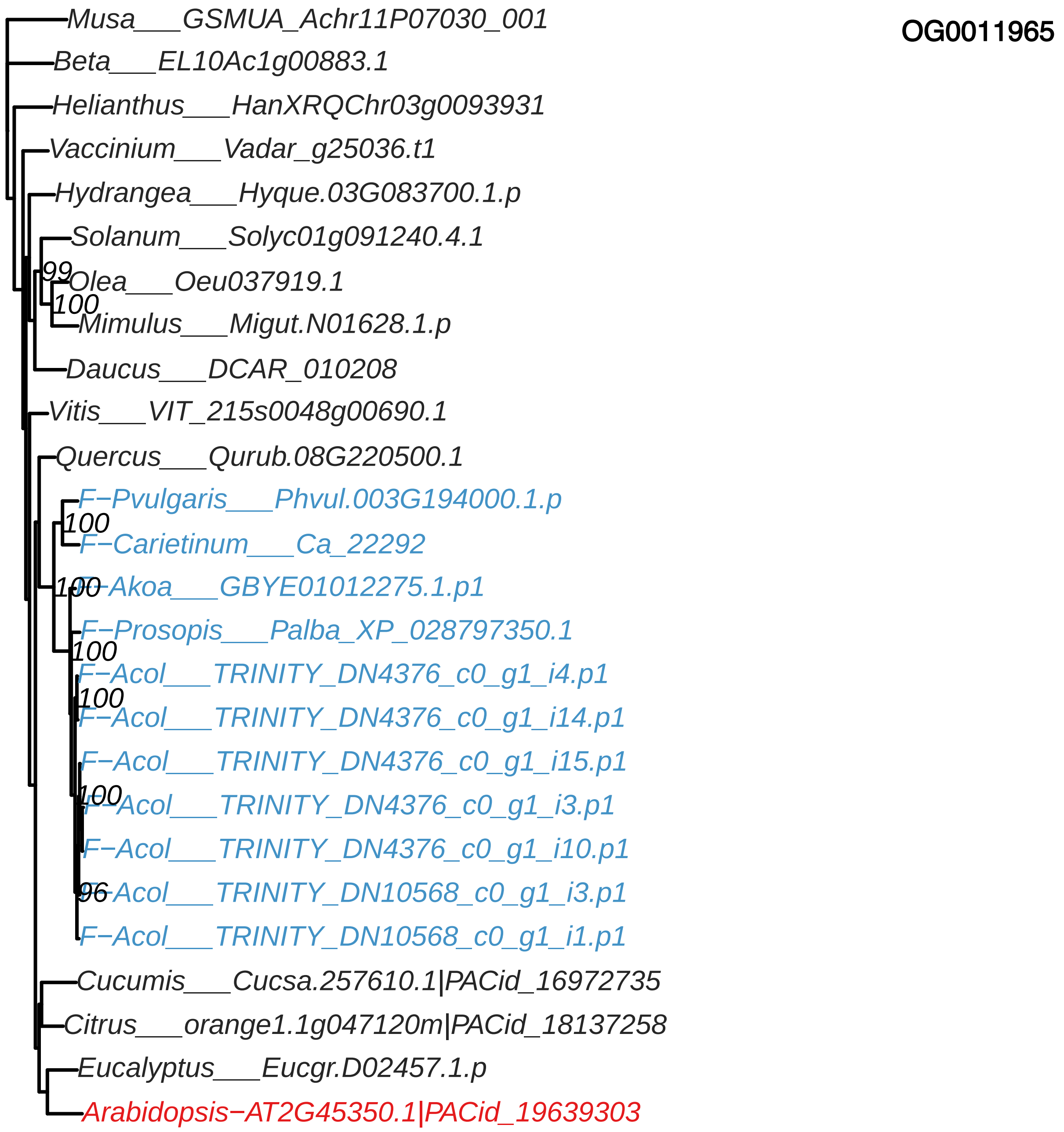

OG0011965
Musa___GSMUA_Achr11P07030_001
Beta___EL10Ac1g00883.1
Helianthus___HanXRQChr03g0093931
Vaccinium___Vadar_g25036.t1
Hydrangea___Hyque.03G083700.1.p
Solanum___Solyc01g091240.4.1
99
Olea___Oeu037919.1
100
Mimulus___Migut.N01628.1.p
Daucus___DCAR_010208
Vitis___VIT_215s0048g00690.1
Quercus___Qurub.08G220500.1
F−Pvulgaris___Phvul.003G194000.1.p
100
F−Carietinum___Ca_22292
100
F−Akoa___GBYE01012275.1.p1
F−Prosopis___Palba_XP_028797350.1
100
F−Acol___TRINITY_DN4376_c0_g1_i4.p1
100
F−Acol___TRINITY_DN4376_c0_g1_i14.p1
F−Acol___TRINITY_DN4376_c0_g1_i15.p1
100
F−Acol___TRINITY_DN4376_c0_g1_i3.p1
F−Acol___TRINITY_DN4376_c0_g1_i10.p1
96
F−Acol___TRINITY_DN10568_c0_g1_i3.p1
F−Acol___TRINITY_DN10568_c0_g1_i1.p1
Cucumis___Cucsa.257610.1|PACid_16972735
Citrus___orange1.1g047120m|PACid_18137258
Eucalyptus___Eucgr.D02457.1.p
Arabidopsis−AT2G45350.1|PACid_19639303

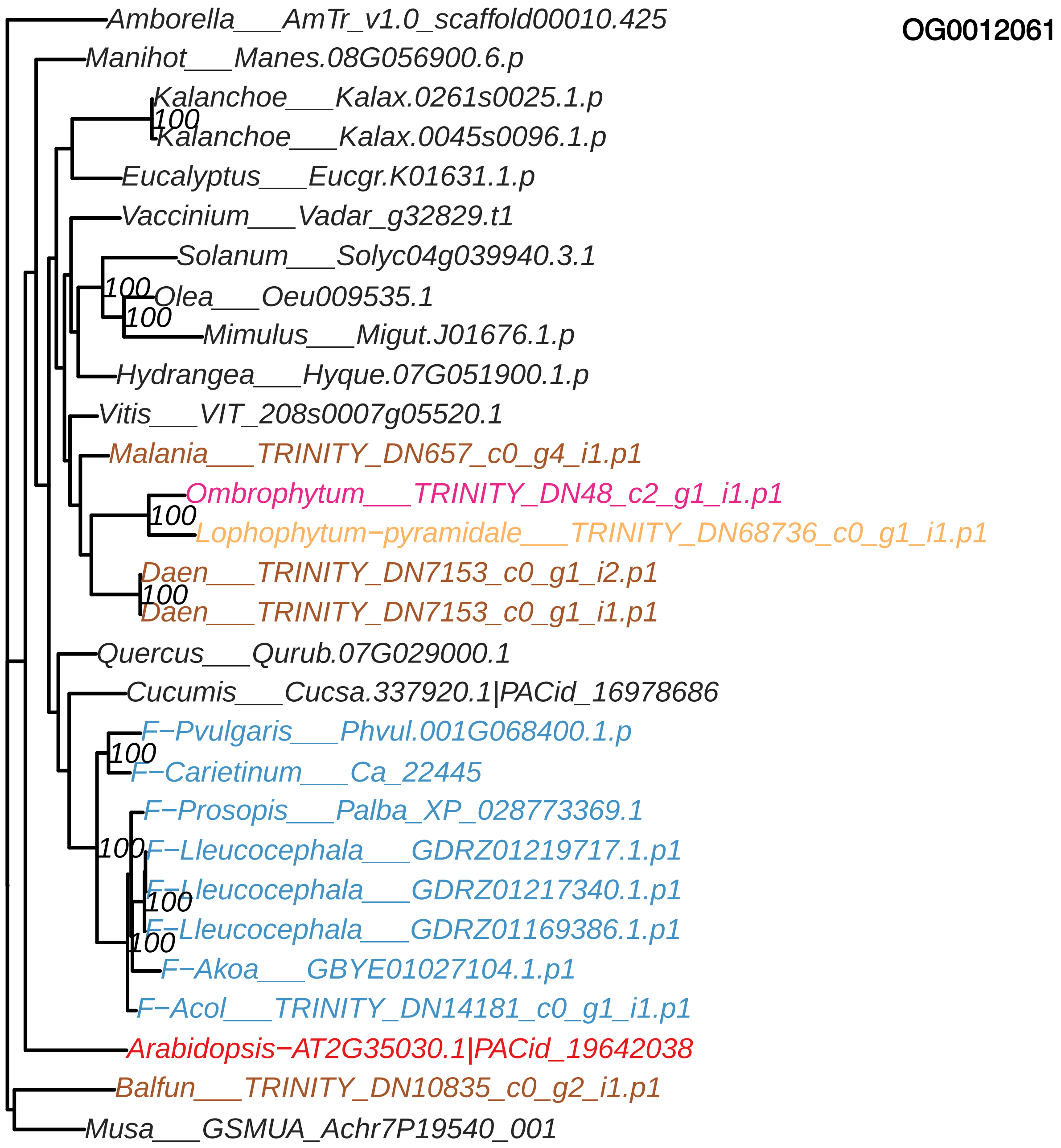

OG0012061
Amborella___AmTr_v1.0_scaffold00010.425
Manihot___Manes.08G056900.6.p
Kalanchoe___Kalax.0261s0025.1.p
100
Kalanchoe___Kalax.0045s0096.1.p
Eucalyptus___Eucgr.K01631.1.p
Vaccinium___Vadar_g32829.t1
Solanum___Solyc04g039940.3.1
100
Olea___Oeu009535.1
100
Mimulus___Migut.J01676.1.p
Hydrangea___Hyque.07G051900.1.p
Vitis___VIT_208s0007g05520.1
Malania___TRINITY_DN657_c0_g4_i1.p1
Ombrophytum___TRINITY_DN48_c2_g1_i1.p1
100
Lophophytum−pyramidale___TRINITY_DN68736_c0_g1_i1.p1
Daen___TRINITY_DN7153_c0_g1_i2.p1
100
Daen___TRINITY_DN7153_c0_g1_i1.p1
Quercus___Qurub.07G029000.1
Cucumis___Cucsa.337920.1|PACid_16978686
F−Pvulgaris___Phvul.001G068400.1.p
100
F−Carietinum___Ca_22445
F−Prosopis___Palba_XP_028773369.1
100
F−Lleucocephala___GDRZ01219717.1.p1
F−Lleucocephala___GDRZ01217340.1.p1
100
F−Lleucocephala___GDRZ01169386.1.p1
100
F−Akoa___GBYE01027104.1.p1
F−Acol___TRINITY_DN14181_c0_g1_i1.p1
Arabidopsis−AT2G35030.1|PACid_19642038
Balfun___TRINITY_DN10835_c0_g2_i1.p1
Musa___GSMUA_Achr7P19540_001

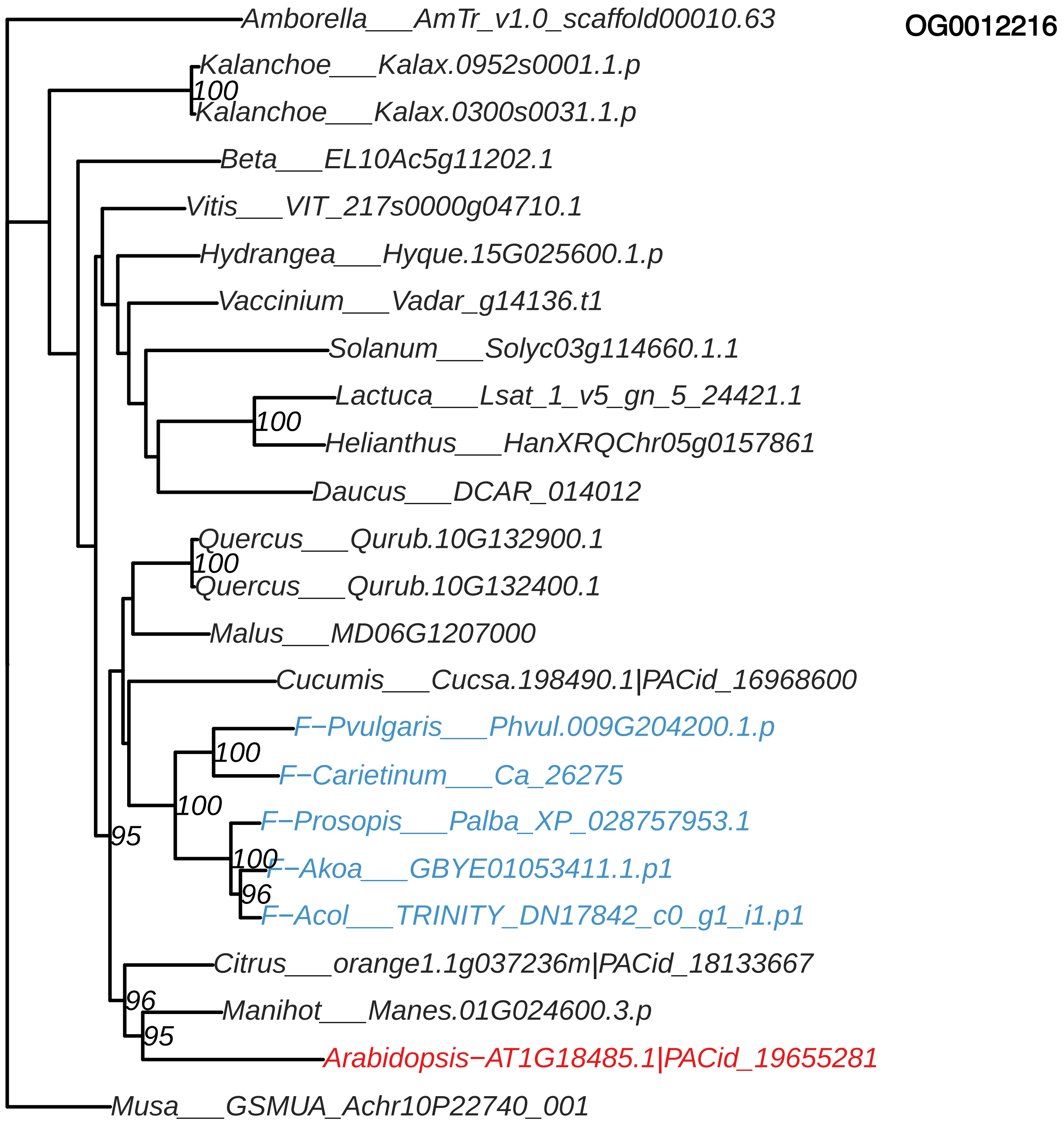
OG0012216
Amborella___AmTr_v1.0_scaffold00010.63
Kalanchoe___Kalax.0952s0001.1.p
100
Kalanchoe___Kalax.0300s0031.1.p
Beta___EL10Ac5g11202.1
Vitis___VIT_217s0000g04710.1
Hydrangea___Hyque.15G025600.1.p
Vaccinium___Vadar_g14136.t1
Solanum___Solyc03g114660.1.1
Lactuca___Lsat_1_v5_gn_5_24421.1
100
Helianthus___HanXRQChr05g0157861
Daucus___DCAR_014012
Quercus___Qurub.10G132900.1
100
Quercus___Qurub.10G132400.1
Malus___MD06G1207000
Cucumis___Cucsa.198490.1|PACid_16968600
F−Pvulgaris___Phvul.009G204200.1.p
100
F−Carietinum___Ca_26275
100
F−Prosopis___Palba_XP_028757953.1
95
100
F−Akoa___GBYE01053411.1.p1
96
F−Acol___TRINITY_DN17842_c0_g1_i1.p1
Citrus___orange1.1g037236m|PACid_18133667
96
Manihot___Manes.01G024600.3.p
95
Arabidopsis−AT1G18485.1|PACid_19655281
Musa___GSMUA_Achr10P22740_001

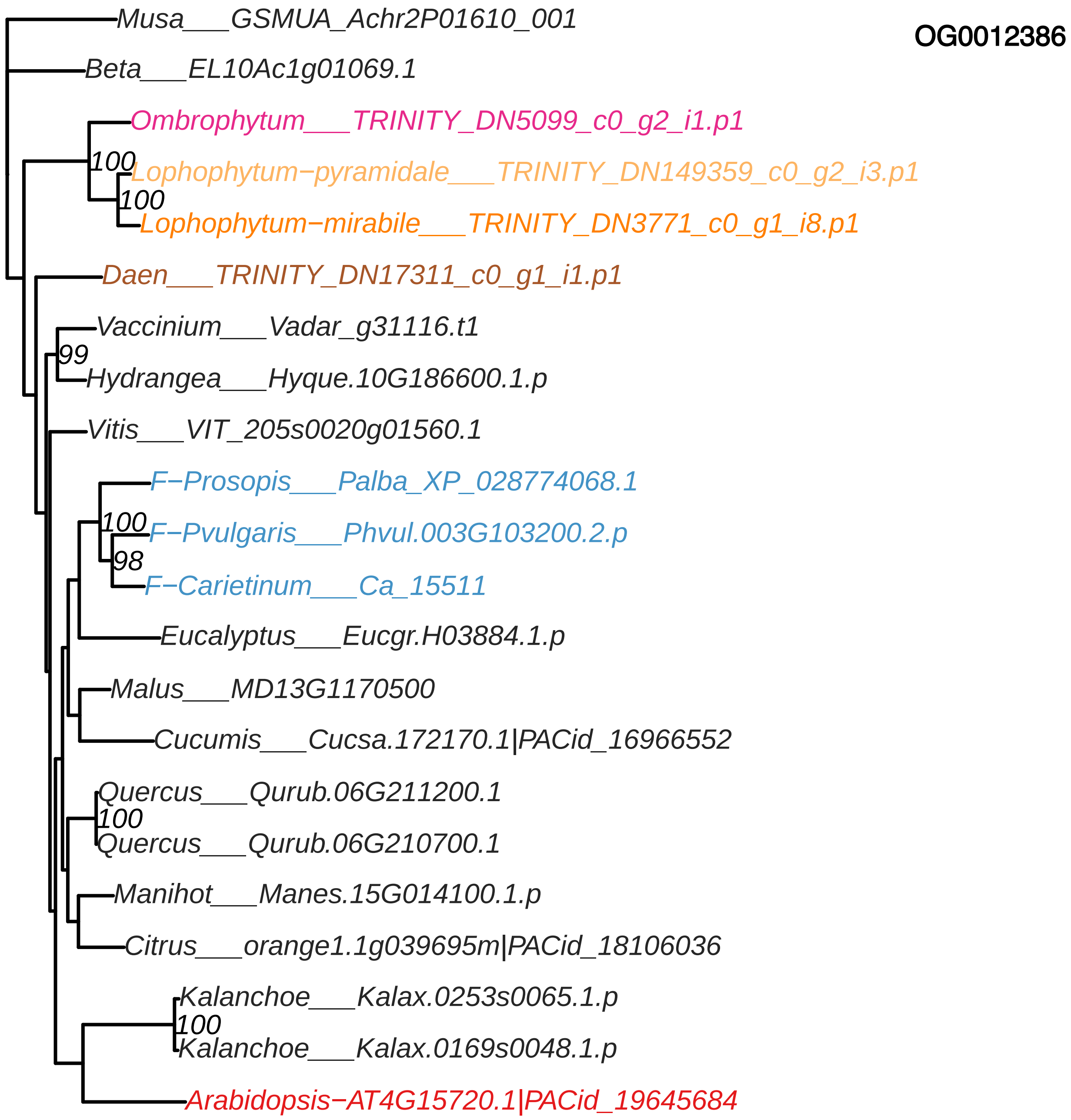
OG0012386
Musa___GSMUA_Achr2P01610_001
Beta___EL10Ac1g01069.1
Ombrophytum___TRINITY_DN5099_c0_g2_i1.p1
100
Lophophytum−pyramidale___TRINITY_DN149359_c0_g2_i3.p1
100
Lophophytum−mirabile___TRINITY_DN3771_c0_g1_i8.p1
Daen___TRINITY_DN17311_c0_g1_i1.p1
Vaccinium___Vadar_g31116.t1
99
Hydrangea___Hyque.10G186600.1.p
Vitis___VIT_205s0020g01560.1
F−Prosopis___Palba_XP_028774068.1
100
F−Pvulgaris___Phvul.003G103200.2.p
98
F−Carietinum___Ca_15511
Eucalyptus___Eucgr.H03884.1.p
Malus___MD13G1170500
Cucumis___Cucsa.172170.1|PACid_16966552
Quercus___Qurub.06G211200.1
100
Quercus___Qurub.06G210700.1
Manihot___Manes.15G014100.1.p
Citrus___orange1.1g039695m|PACid_18106036
Kalanchoe___Kalax.0253s0065.1.p
100
Kalanchoe___Kalax.0169s0048.1.p
Arabidopsis−AT4G15720.1|PACid_19645684

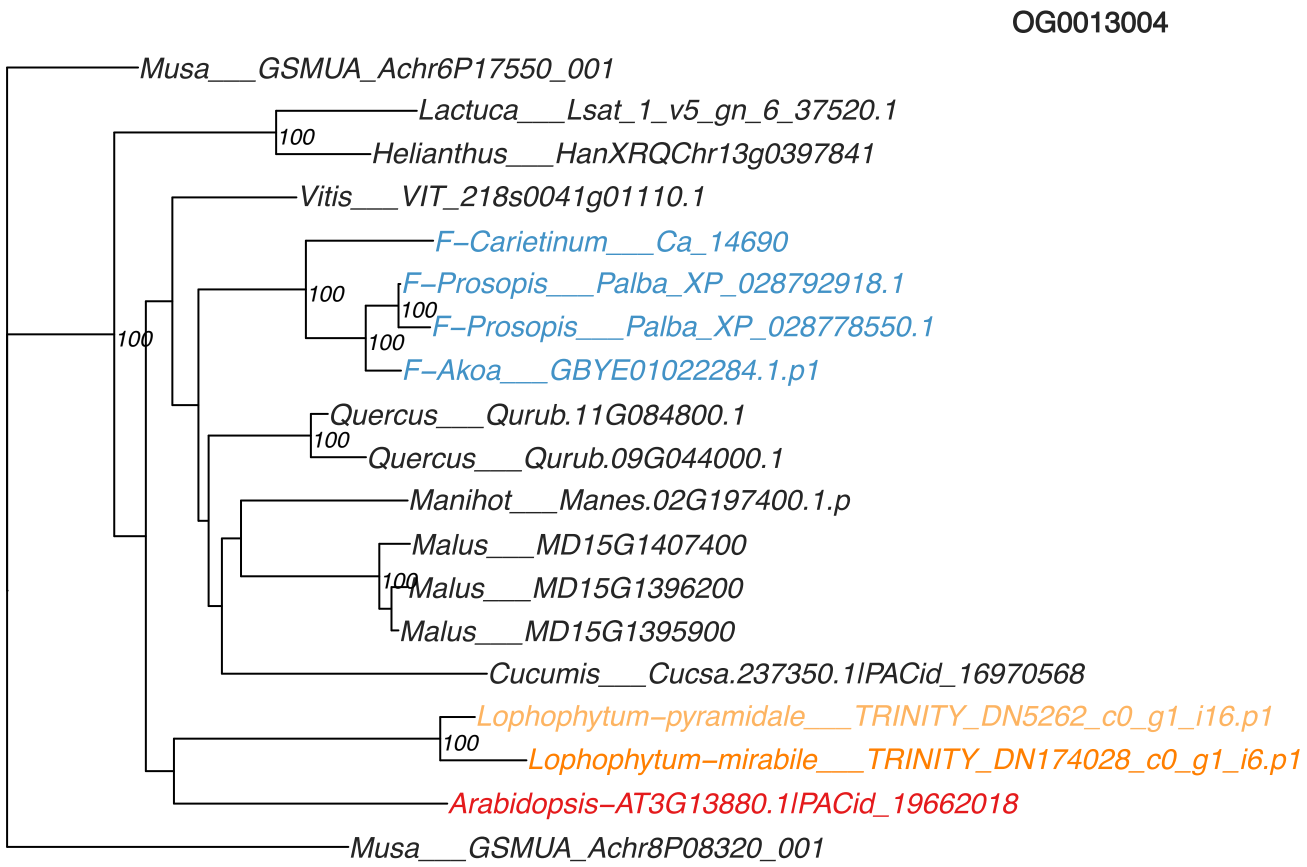
OG0013004
Musa___GSMUA_Achr6P17550_001
Lactuca___Lsat_1_v5_gn_6_37520.1
Helianthus___HanXRQChr13g0397841
Vitis___VIT_218s0041g01110.1
F–Carietinum___Ca_14690
F–Prosopis___Palba_XP_028792918.1
F–Prosopis___Palba_XP_028778550.1
F–Akoa___GBYE01022284.1.p1
Quercus___Qurub.11G084800.1
Quercus___Qurub.09G044000.1
Manihot___Manes.02G197400.1.p
Malus___MD15G1407400
Malus___MD15G1396200
Malus___MD15G1395900
Cucumis___Cucsa.237350.1|PACid_16970568
Lophophytum–pyramidale___TRINITY_DN5262_c0_g1_i16.p1
Lophophytum–mirabile___TRINITY_DN174028_c0_g1_i6.p1
Arabidopsis–AT3G13880.1|PACid_19662018
Musa___GSMUA_Achr8P08320_001
100
100
100
100
100
100
100
100
100

OG0013006

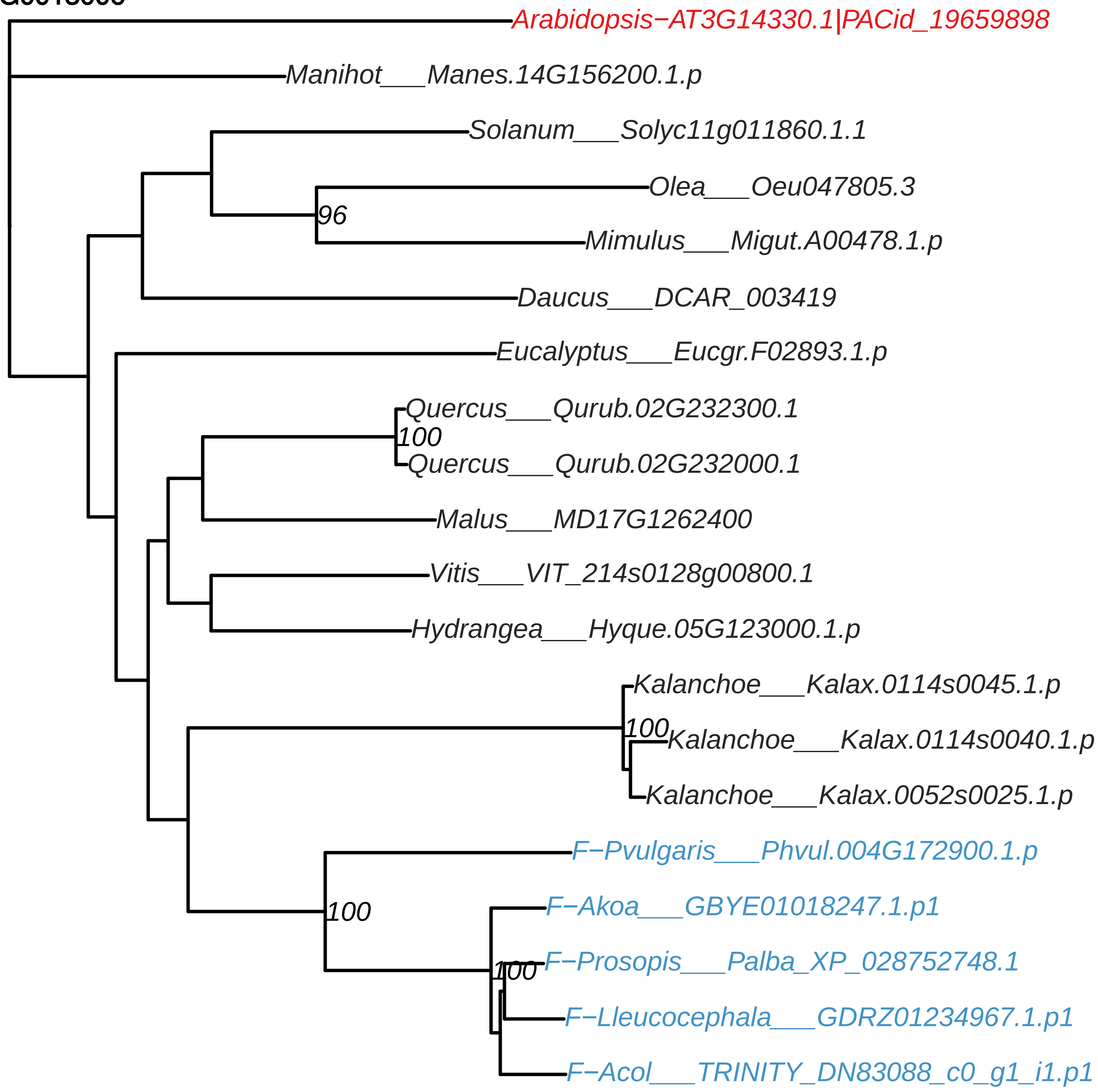

Arabidopsis−AT3G14330.1|PACid_19659898
Manihot___Manes.14G156200.1.p
Solanum___Solyc11g011860.1.1
Olea___Oeu047805.3
96
Mimulus___Migut.A00478.1.p
Daucus___DCAR_003419
Eucalyptus___Eucgr.F02893.1.p
Quercus___Qurub.02G232300.1
100
Quercus___Qurub.02G232000.1
Malus___MD17G1262400
Vitis___VIT_214s0128g00800.1
Hydrangea___Hyque.05G123000.1.p
Kalanchoe___Kalax.0114s0045.1.p
100
Kalanchoe___Kalax.0114s0040.1.p
Kalanchoe___Kalax.0052s0025.1.p
F−Pvulgaris___Phvul.004G172900.1.p
100
F−Akoa___GBYE01018247.1.p1
100
F−Prosopis___Palba_XP_028752748.1
F−Lleucocephala___GDRZ01234967.1.p1
F−Acol___TRINITY_DN83088_c0_g1_i1.p1

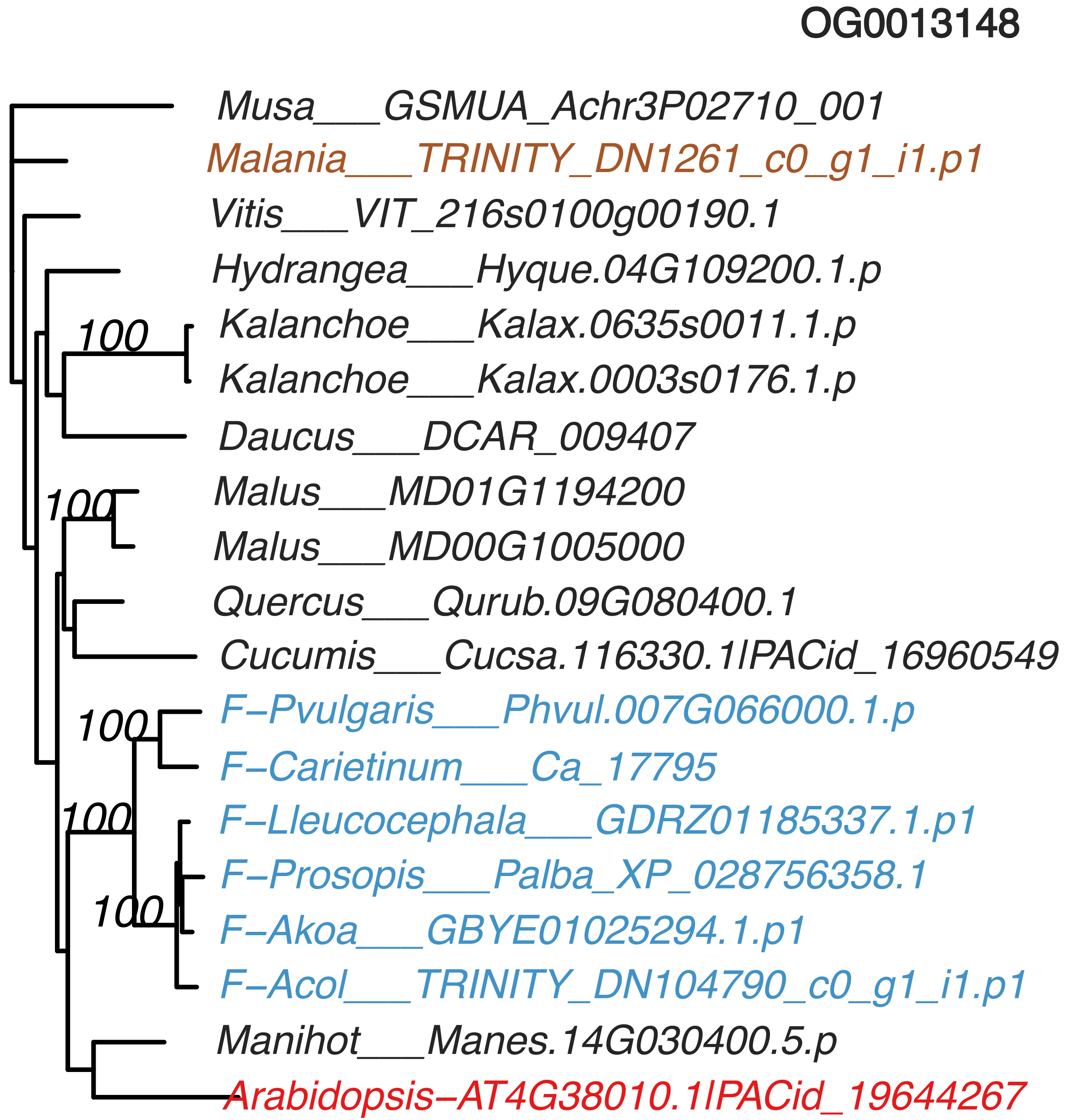

OG0013148
Musa___GSMUA_Achr3P02710_001
Malania___TRINITY_DN1261_c0_g1_i1.p1
Vitis___VIT_216s0100g00190.1
Hydrangea___Hyque.04G109200.1.p
Kalanchoe___Kalax.0635s0011.1.p
Kalanchoe___Kalax.0003s0176.1.p
Daucus___DCAR_009407
Malus___MD01G1194200
Malus___MD00G1005000
Quercus___Qurub.09G080400.1
Cucumis___Cucsa.116330.1|PACid_16960549
F–Pvulgaris___Phvul.007G066000.1.p
F–Carietinum___Ca_17795
F–Lleucocephala___GDRZ01185337.1.p1
F–Prosopis___Palba_XP_028756358.1
F–Akoa___GBYE01025294.1.p1
F–Acol___TRINITY_DN104790_c0_g1_i1.p1
Manihot___Manes.14G030400.5.p
Arabidopsis–AT4G38010.1|PACid_19644267
100
100
100
100
100

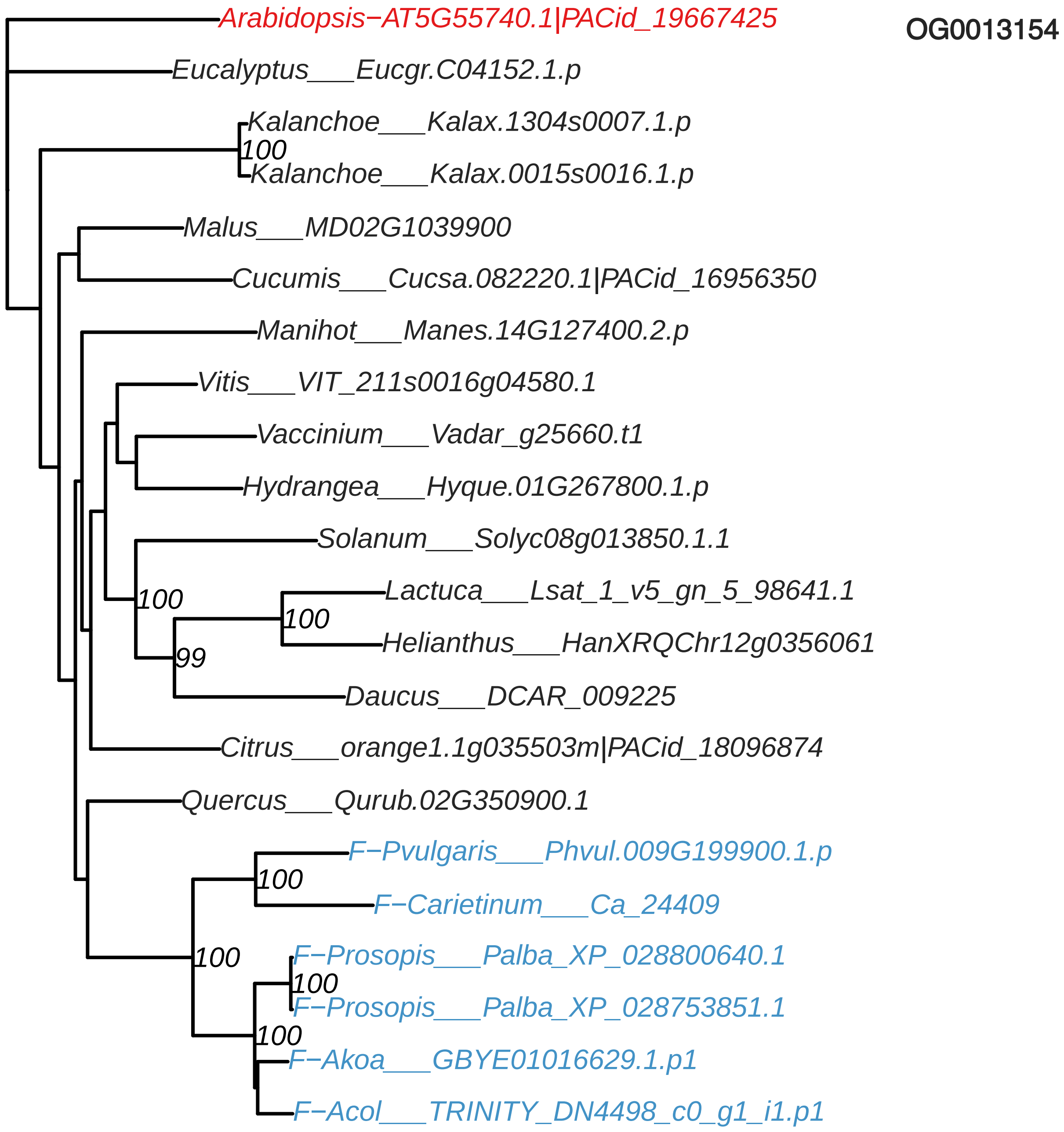
OG0013154
Arabidopsis−AT5G55740.1|PACid_19667425
Eucalyptus___Eucgr.C04152.1.p
Kalanchoe___Kalax.1304s0007.1.p
100
Kalanchoe___Kalax.0015s0016.1.p
Malus___MD02G1039900
Cucumis___Cucsa.082220.1|PACid_16956350
Manihot___Manes.14G127400.2.p
Vitis___VIT_211s0016g04580.1
Vaccinium___Vadar_g25660.t1
Hydrangea___Hyque.01G267800.1.p
Solanum___Solyc08g013850.1.1
100
Lactuca___Lsat_1_v5_gn_5_98641.1
100
Helianthus___HanXRQChr12g0356061
99
Daucus___DCAR_009225
Citrus___orange1.1g035503m|PACid_18096874
Quercus___Qurub.02G350900.1
F−Pvulgaris___Phvul.009G199900.1.p
100
F−Carietinum___Ca_24409
100
F−Prosopis___Palba_XP_028800640.1
100
F−Prosopis___Palba_XP_028753851.1
100
F−Akoa___GBYE01016629.1.p1
F−Acol___TRINITY_DN4498_c0_g1_i1.p1

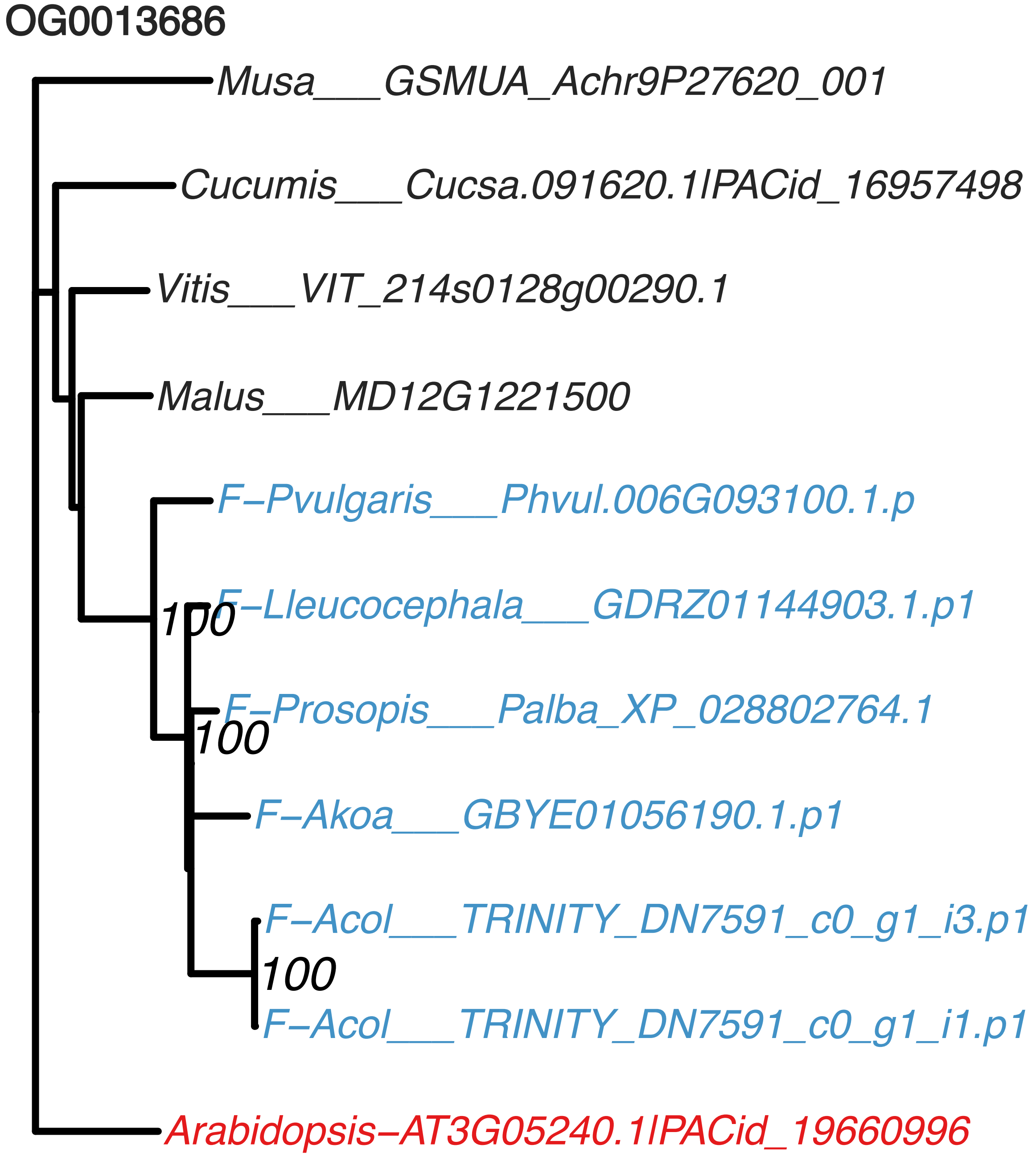

OG0013686
Musa___GSMUA_Achr9P27620_001
Cucumis___Cucsa.091620.1|PACid_16957498
Vitis___VIT_214s0128g00290.1
Malus___MD12G1221500
F–Pvulgaris___Phvul.006G093100.1.p
F–Lleucocephala___GDRZ01144903.1.p1
100
F–Prosopis___Palba_XP_028802764.1
100
F–Akoa___GBYE01056190.1.p1
F–Acol___TRINITY_DN7591_c0_g1_i3.p1
100
F–Acol___TRINITY_DN7591_c0_g1_i1.p1
Arabidopsis–AT3G05240.1|PACid_19660996

# OG0015755

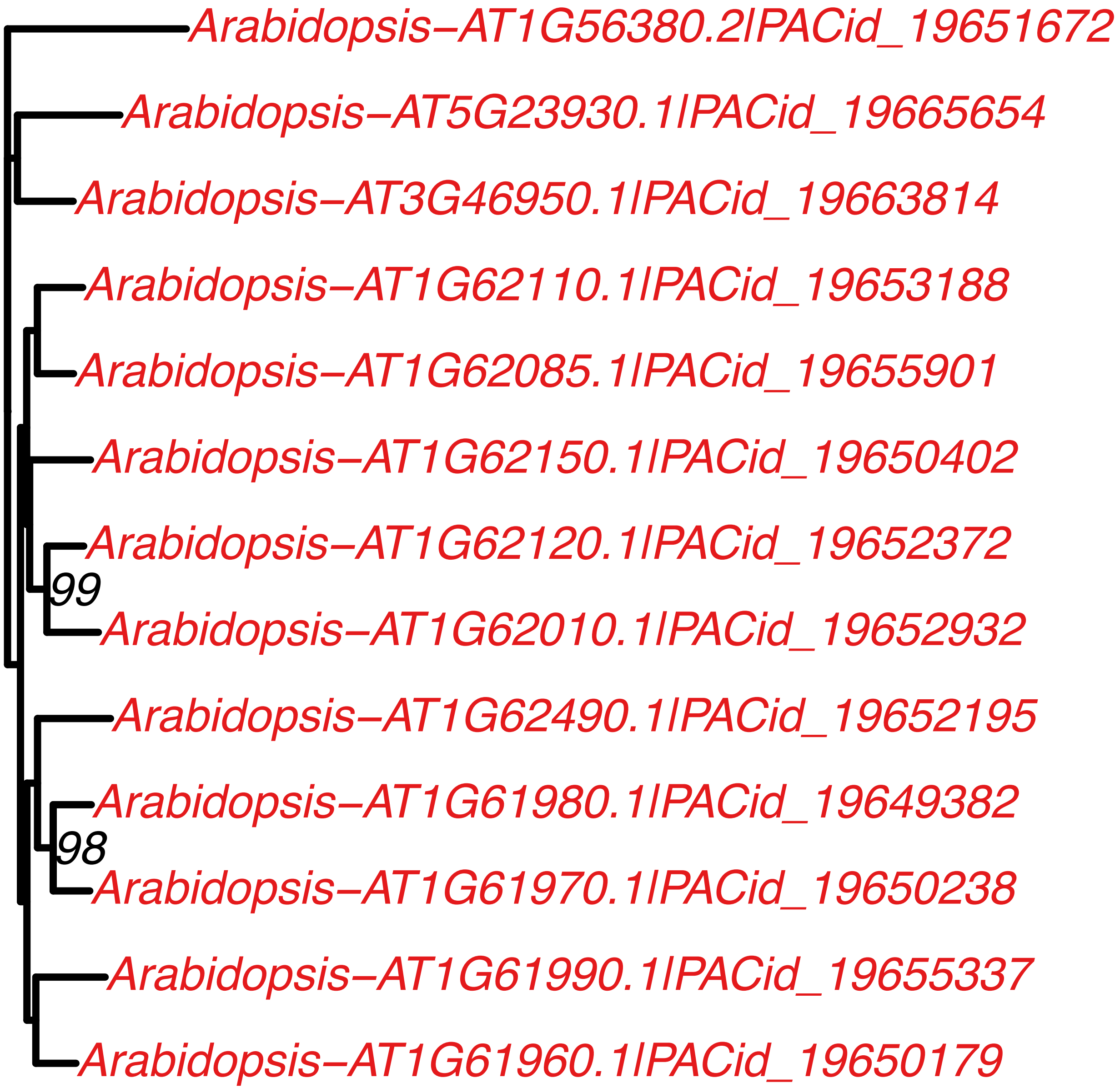

Arabidopsis–AT1G56380.2|PACid_19651672
Arabidopsis–AT5G23930.1|PACid_19665654
Arabidopsis–AT3G46950.1|PACid_19663814
Arabidopsis–AT1G62110.1|PACid_19653188
Arabidopsis–AT1G62085.1|PACid_19655901
Arabidopsis–AT1G62150.1|PACid_19650402
Arabidopsis–AT1G62120.1|PACid_19652372
99
Arabidopsis–AT1G62010.1|PACid_19652932
Arabidopsis–AT1G62490.1|PACid_19652195
Arabidopsis–AT1G61980.1|PACid_19649382
98
Arabidopsis–AT1G61970.1|PACid_19650238
Arabidopsis–AT1G61990.1|PACid_19655337
Arabidopsis–AT1G61960.1|PACid_19650179

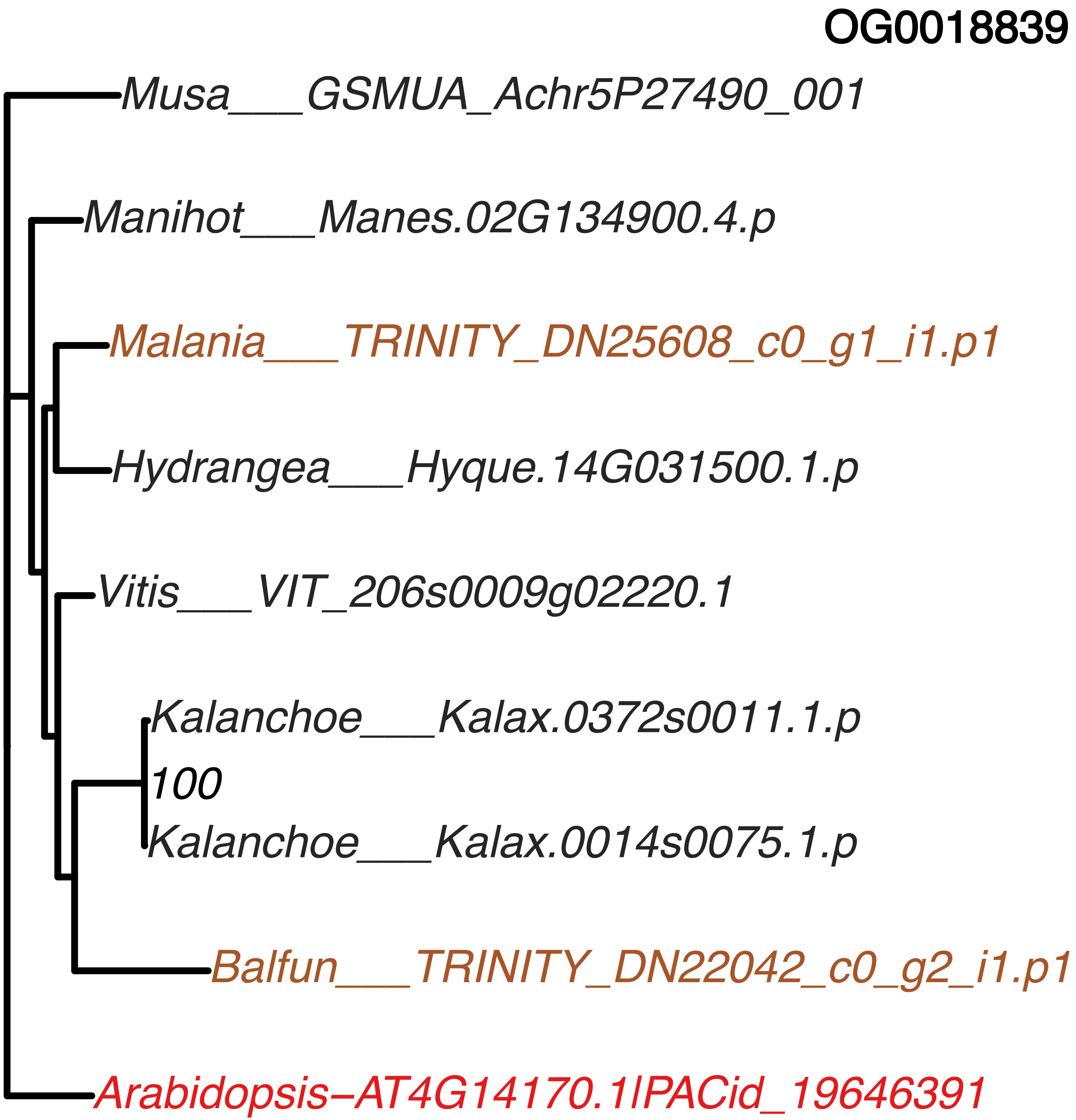
OG0018839
Musa___GSMUA_Achr5P27490_001
Manihot___Manes.02G134900.4.p
Malania___TRINITY_DN25608_c0_g1_i1.p1
Hydrangea___Hyque.14G031500.1.p
Vitis___VIT_206s0009g02220.1
Kalanchoe___Kalax.0372s0011.1.p
100
Kalanchoe___Kalax.0014s0075.1.p
Balfun___TRINITY_DN22042_c0_g2_i1.p1
Arabidopsis–AT4G14170.1|PACid_19646391

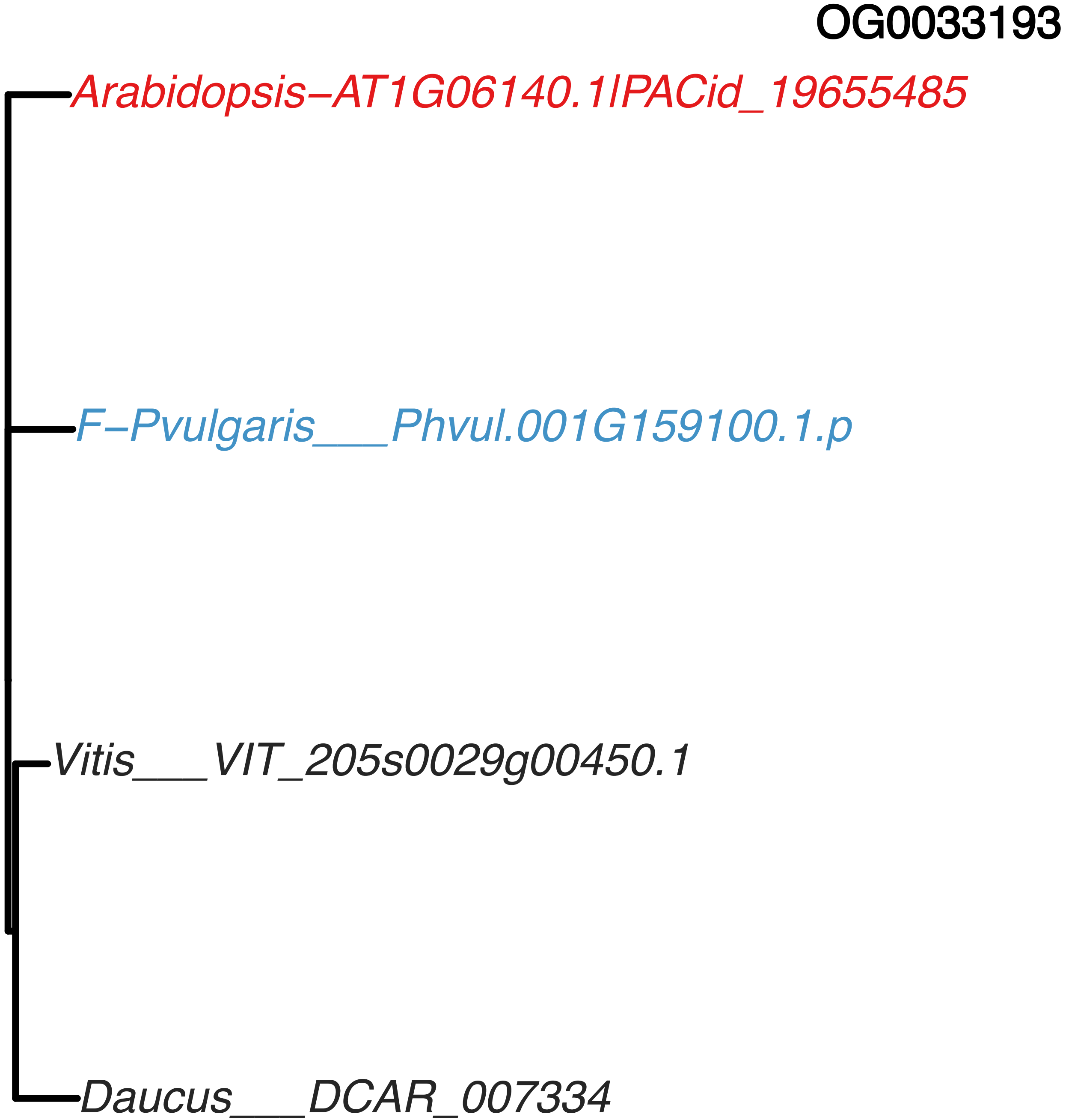
OG0033193
Arabidopsis–AT1G06140.1lPACid_19655485
F–Pvulgaris___Phvul.001G159100.1.p
Vitis___VIT_205s0029g00450.1
Daucus___DCAR_007334

OG0033202

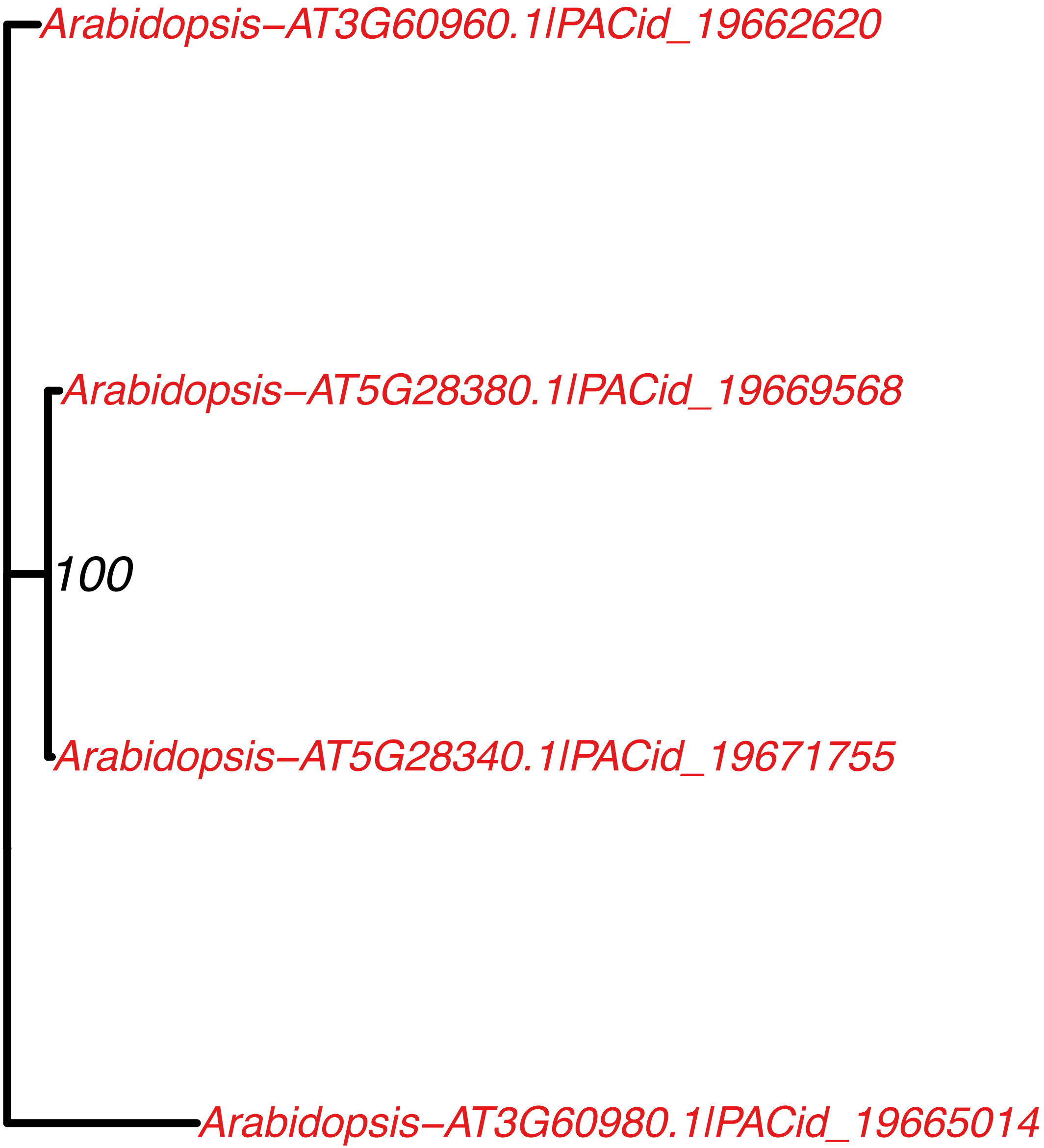

Arabidopsis–AT3G60960.1|PACid_19662620
Arabidopsis–AT5G28380.1|PACid_19669568
100
Arabidopsis–AT5G28340.1|PACid_19671755
Arabidopsis–AT3G60980.1|PACid_19665014

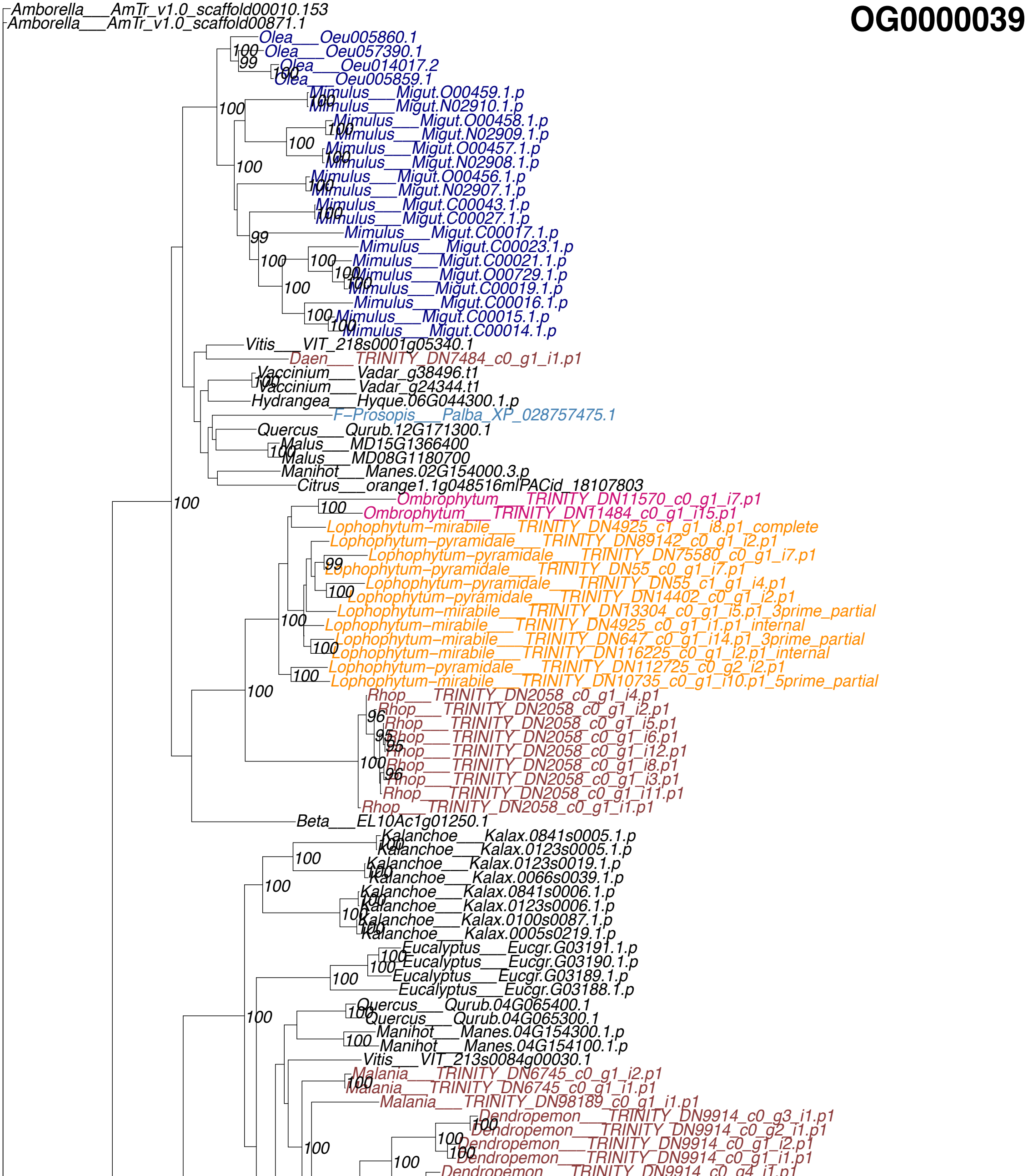
OG0000039
Amborella___AmTr_v1.0_scaffold00010.153
Amborella___AmTr_v1.0_scaffold00871.1
Olea___Oeu005860.1
Olea___Oeu057390.1
Olea___Oeu014017.2
Olea___Oeu005859.1
Mimulus___Migut.O00459.1.p
Mimulus___Migut.N02910.1.p
Mimulus___Migut.O00458.1.p
Mimulus___Migut.N02909.1.p
Mimulus___Migut.O00457.1.p
Mimulus___Migut.N02908.1.p
Mimulus___Migut.O00456.1.p
Mimulus___Migut.N02907.1.p
Mimulus___Migut.C00043.1.p
Mimulus___Migut.C00027.1.p
Mimulus___Migut.C00017.1.p
Mimulus___Migut.C00023.1.p
Mimulus___Migut.C00021.1.p
Mimulus___Migut.O00729.1.p
Mimulus___Migut.C00019.1.p
Mimulus___Migut.C00016.1.p
Mimulus___Migut.C00015.1.p
Mimulus___Migut.C00014.1.p
Vitis___VIT_218s0001g05340.1
Daen___TRINITY_DN7484_c0_g1_i1.p1
Vaccinium___Vadar_g38496.t1
Vaccinium___Vadar_g24344.t1
Hydrangea___Hyque.06G044300.1.p
F−Prosopis___Palba_XP_028757475.1
Quercus___Qurub.12G171300.1
Malus___MD15G1366400
Malus___MD08G1180700
Manihot___Manes.02G154000.3.p
Citrus___orange1.1g048516mlPACid_18107803
Ombrophytum___TRINITY_DN11570_c0_g1_i7.p1
Ombrophytum___TRINITY_DN11484_c0_g1_i15.p1
Lophophytum−mirabile___TRINITY_DN4925_c1_g1_i8.p1_complete
Lophophytum−pyramidale___TRINITY_DN89142_c0_g1_i2.p1
Lophophytum−pyramidale___TRINITY_DN75580_c0_g1_i7.p1
Lophophytum−pyramidale___TRINITY_DN55_c0_g1_i7.p1
Lophophytum−pyramidale___TRINITY_DN55_c1_g1_i4.p1
Lophophytum−pyramidale___TRINITY_DN14402_c0_g1_i2.p1
Lophophytum−mirabile___TRINITY_DN13304_c0_g1_i5.p1_3prime_partial
Lophophytum−mirabile___TRINITY_DN4925_c0_g1_i1.p1_internal
Lophophytum−mirabile___TRINITY_DN647_c0_g1_i14.p1_3prime_partial
Lophophytum−mirabile___TRINITY_DN116225_c0_g1_i2.p1_internal
Lophophytum−pyramidale___TRINITY_DN112725_c0_g2_i2.p1
Lophophytum−mirabile___TRINITY_DN10735_c0_g1_i10.p1_5prime_partial
Rhop___TRINITY_DN2058_c0_g1_i4.p1
Rhop___TRINITY_DN2058_c0_g1_i2.p1
Rhop___TRINITY_DN2058_c0_g1_i5.p1
Rhop___TRINITY_DN2058_c0_g1_i6.p1
Rhop___TRINITY_DN2058_c0_g1_i12.p1
Rhop___TRINITY_DN2058_c0_g1_i8.p1
Rhop___TRINITY_DN2058_c0_g1_i3.p1
Rhop___TRINITY_DN2058_c0_g1_i11.p1
Rhop___TRINITY_DN2058_c0_g1_i1.p1
Beta___EL10Ac1g01250.1
Kalanchoe___Kalax.0841s0005.1.p
Kalanchoe___Kalax.0123s0005.1.p
Kalanchoe___Kalax.0123s0019.1.p
Kalanchoe___Kalax.0066s0039.1.p
Kalanchoe___Kalax.0841s0006.1.p
Kalanchoe___Kalax.0123s0006.1.p
Kalanchoe___Kalax.0100s0087.1.p
Kalanchoe___Kalax.0005s0219.1.p
Eucalyptus___Eucgr.G03191.1.p
Eucalyptus___Eucgr.G03190.1.p
Eucalyptus___Eucgr.G03189.1.p
Eucalyptus___Eucgr.G03188.1.p
Quercus___Qurub.04G065400.1
Quercus___Qurub.04G065300.1
Manihot___Manes.04G154300.1.p
Manihot___Manes.04G154100.1.p
Vitis___VIT_213s0084g00030.1
Malania___TRINITY_DN6745_c0_g1_i2.p1
Malania___TRINITY_DN6745_c0_g1_i1.p1
Malania___TRINITY_DN98189_c0_g1_i1.p1
Dendropemon___TRINITY_DN9914_c0_g3_i1.p1
Dendropemon___TRINITY_DN9914_c0_g2_i1.p1
Dendropemon___TRINITY_DN9914_c0_g1_i2.p1
Dendropemon___TRINITY_DN9914_c0_g1_i1.p1
Dendropemon___TRINITY_DN9914_c0_g4_i1.p1

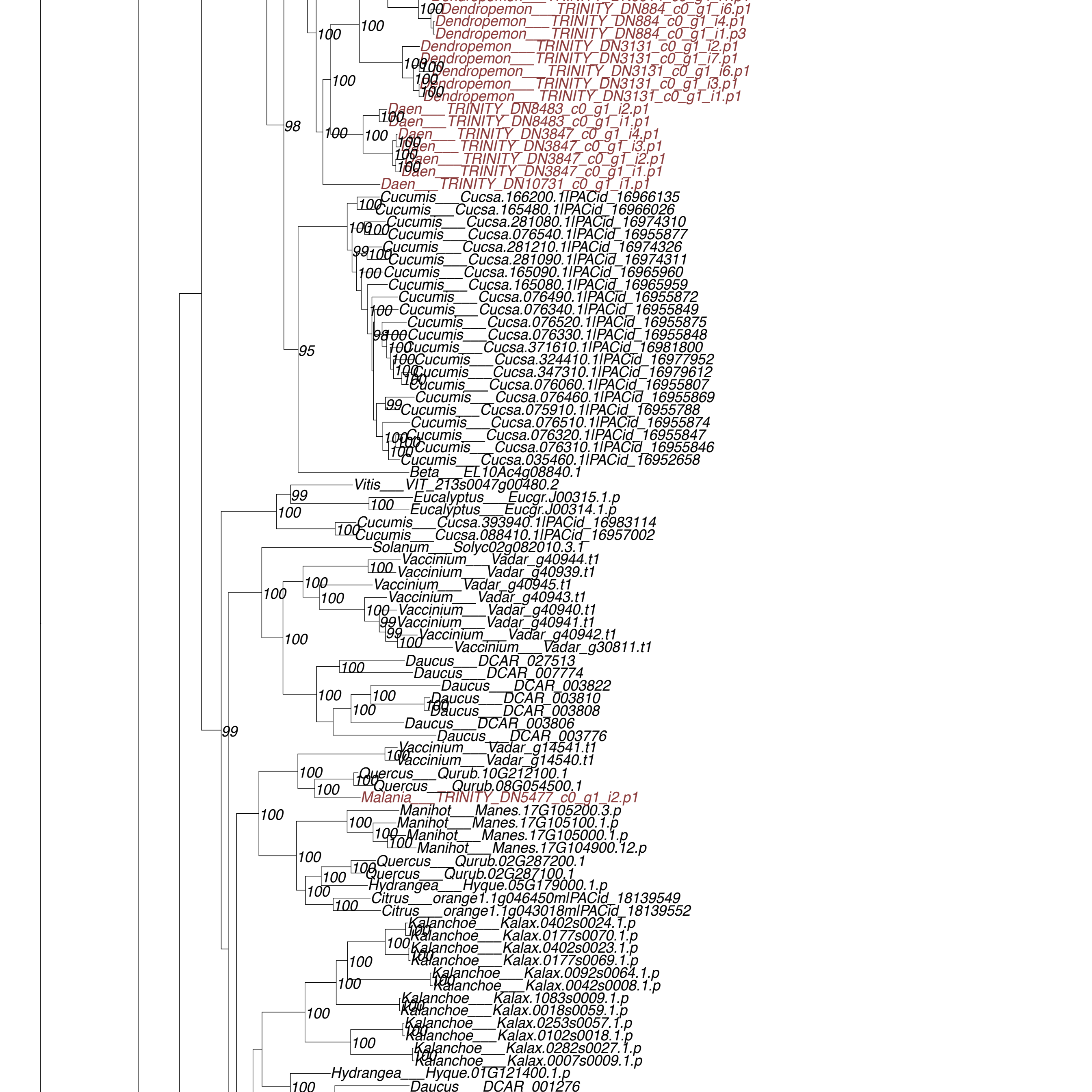

Dendropemon___TRINITY_DN884_c0_g1_i6.p1
Dendropemon___TRINITY_DN884_c0_g1_i4.p1
Dendropemon___TRINITY_DN884_c0_g1_i1.p3
Dendropemon___TRINITY_DN3131_c0_g1_i2.p1
Dendropemon___TRINITY_DN3131_c0_g1_i7.p1
Dendropemon___TRINITY_DN3131_c0_g1_i6.p1
Dendropemon___TRINITY_DN3131_c0_g1_i3.p1
Dendropemon___TRINITY_DN3131_c0_g1_i1.p1
Daen___TRINITY_DN8483_c0_g1_i2.p1
Daen___TRINITY_DN8483_c0_g1_i1.p1
Daen___TRINITY_DN3847_c0_g1_i4.p1
Daen___TRINITY_DN3847_c0_g1_i3.p1
Daen___TRINITY_DN3847_c0_g1_i2.p1
Daen___TRINITY_DN3847_c0_g1_i1.p1
Daen___TRINITY_DN10731_c0_g1_i1.p1
Cucumis___Cucsa.166200.1|PACid_16966135
Cucumis___Cucsa.165480.1|PACid_16966026
Cucumis___Cucsa.281080.1|PACid_16974310
Cucumis___Cucsa.076540.1|PACid_16955877
Cucumis___Cucsa.281210.1|PACid_16974326
Cucumis___Cucsa.281090.1|PACid_16974311
Cucumis___Cucsa.165090.1|PACid_16965960
Cucumis___Cucsa.165080.1|PACid_16965959
Cucumis___Cucsa.076490.1|PACid_16955872
Cucumis___Cucsa.076340.1|PACid_16955849
Cucumis___Cucsa.076520.1|PACid_16955875
Cucumis___Cucsa.076330.1|PACid_16955848
Cucumis___Cucsa.371610.1|PACid_16981800
Cucumis___Cucsa.324410.1|PACid_16977952
Cucumis___Cucsa.347310.1|PACid_16979612
Cucumis___Cucsa.076060.1|PACid_16955807
Cucumis___Cucsa.076460.1|PACid_16955869
Cucumis___Cucsa.075910.1|PACid_16955788
Cucumis___Cucsa.076510.1|PACid_16955874
Cucumis___Cucsa.076320.1|PACid_16955847
Cucumis___Cucsa.076310.1|PACid_16955846
Cucumis___Cucsa.035460.1|PACid_16952658
Beta___EL10Ac4g08840.1
Vitis___VIT_213s0047g00480.2
Eucalyptus___Eucgr.J00315.1.p
Eucalyptus___Eucgr.J00314.1.p
Cucumis___Cucsa.393940.1|PACid_16983114
Cucumis___Cucsa.088410.1|PACid_16957002
Solanum___Solyc02g082010.3.1
Vaccinium___Vadar_g40944.t1
Vaccinium___Vadar_g40939.t1
Vaccinium___Vadar_g40945.t1
Vaccinium___Vadar_g40943.t1
Vaccinium___Vadar_g40940.t1
Vaccinium___Vadar_g40941.t1
Vaccinium___Vadar_g40942.t1
Vaccinium___Vadar_g30811.t1
Daucus___DCAR_027513
Daucus___DCAR_007774
Daucus___DCAR_003822
Daucus___DCAR_003810
Daucus___DCAR_003808
Daucus___DCAR_003806
Daucus___DCAR_003776
Vaccinium___Vadar_g14541.t1
Vaccinium___Vadar_g14540.t1
Quercus___Qurub.10G212100.1
Quercus___Qurub.08G054500.1
Malania___TRINITY_DN5477_c0_g1_i2.p1
Manihot___Manes.17G105200.3.p
Manihot___Manes.17G105100.1.p
Manihot___Manes.17G105000.1.p
Manihot___Manes.17G104900.12.p
Quercus___Qurub.02G287200.1
Quercus___Qurub.02G287100.1
Hydrangea___Hyque.05G179000.1.p
Citrus___orange1.1g046450m|PACid_18139549
Citrus___orange1.1g043018m|PACid_18139552
Kalanchoe___Kalax.0402s0024.1.p
Kalanchoe___Kalax.0177s0070.1.p
Kalanchoe___Kalax.0402s0023.1.p
Kalanchoe___Kalax.0177s0069.1.p
Kalanchoe___Kalax.0092s0064.1.p
Kalanchoe___Kalax.0042s0008.1.p
Kalanchoe___Kalax.1083s0009.1.p
Kalanchoe___Kalax.0018s0059.1.p
Kalanchoe___Kalax.0253s0057.1.p
Kalanchoe___Kalax.0102s0018.1.p
Kalanchoe___Kalax.0282s0027.1.p
Kalanchoe___Kalax.0007s0009.1.p
Hydrangea___Hyque.01G121400.1.p
Daucus___DCAR_001276

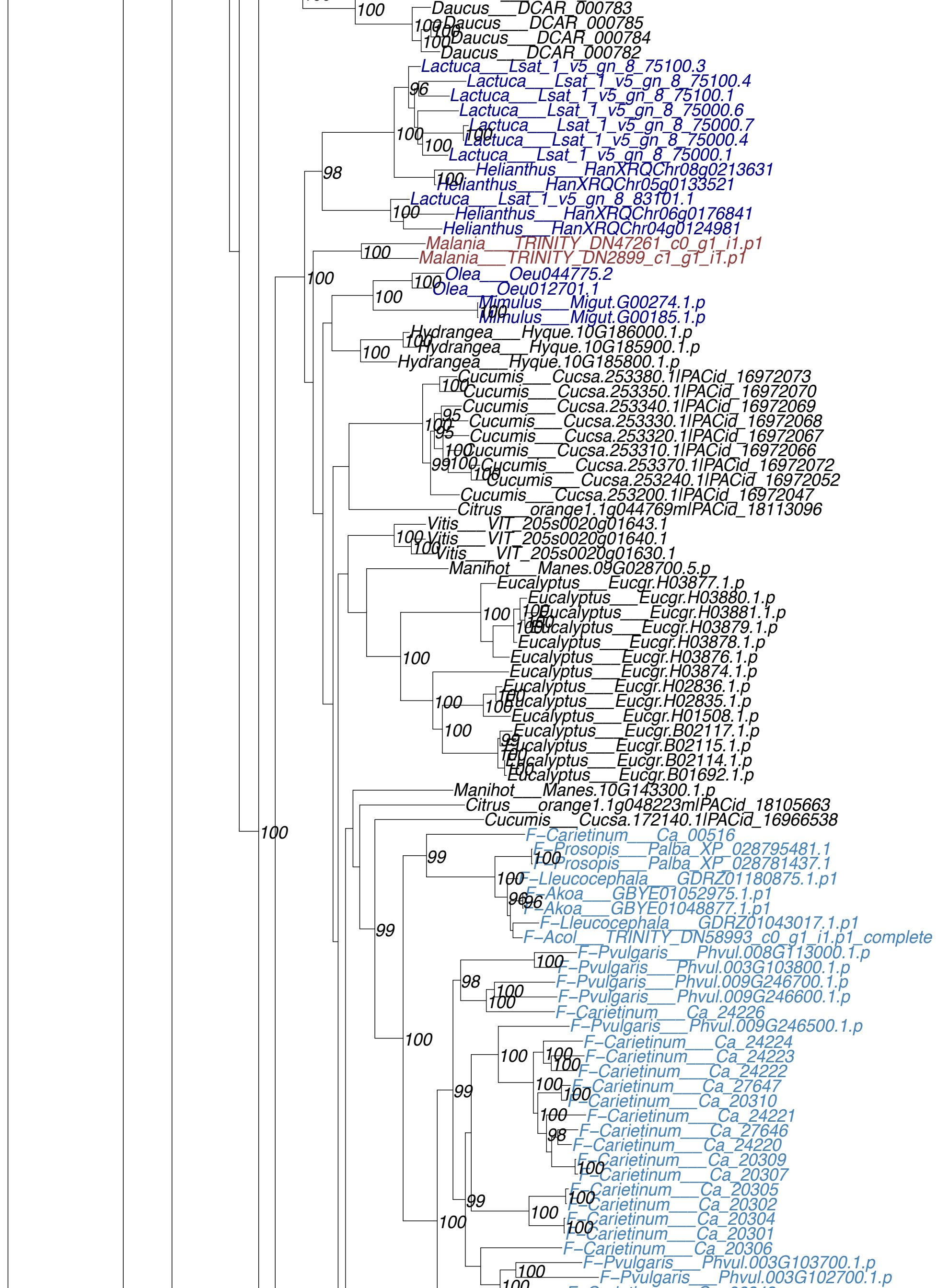
Daucus___DCAR_000783
Daucus___DCAR_000785
Daucus___DCAR_000784
Daucus___DCAR_000782
Lactuca___Lsat_1_v5_gn_8_75100.3
Lactuca___Lsat_1_v5_gn_8_75100.4
Lactuca___Lsat_1_v5_gn_8_75100.1
Lactuca___Lsat_1_v5_gn_8_75000.6
Lactuca___Lsat_1_v5_gn_8_75000.7
Lactuca___Lsat_1_v5_gn_8_75000.4
Lactuca___Lsat_1_v5_gn_8_75000.1
Helianthus___HanXRQChr08g0213631
Helianthus___HanXRQChr05g0133521
Lactuca___Lsat_1_v5_gn_8_83101.1
Helianthus___HanXRQChr06g0176841
Helianthus___HanXRQChr04g0124981
Malania___TRINITY_DN47261_c0_g1_i1.p1
Malania___TRINITY_DN2899_c1_g1_i1.p1
Olea___Oeu044775.2
Olea___Oeu012701.1
Mimulus___Migut.G00274.1.p
Mimulus___Migut.G00185.1.p
Hydrangea___Hyque.10G186000.1.p
Hydrangea___Hyque.10G185900.1.p
Hydrangea___Hyque.10G185800.1.p
Cucumis___Cucsa.253380.1lPACid_16972073
Cucumis___Cucsa.253350.1lPACid_16972070
Cucumis___Cucsa.253340.1lPACid_16972069
Cucumis___Cucsa.253330.1lPACid_16972068
Cucumis___Cucsa.253320.1lPACid_16972067
Cucumis___Cucsa.253310.1lPACid_16972066
Cucumis___Cucsa.253370.1lPACid_16972072
Cucumis___Cucsa.253240.1lPACid_16972052
Cucumis___Cucsa.253200.1lPACid_16972047
Citrus___orange1.1g044769mlPACid_18113096
Vitis___VIT_205s0020g01643.1
Vitis___VIT_205s0020g01640.1
Vitis___VIT_205s0020g01630.1
Manihot___Manes.09G028700.5.p
Eucalyptus___Eucgr.H03877.1.p
Eucalyptus___Eucgr.H03880.1.p
Eucalyptus___Eucgr.H03881.1.p
Eucalyptus___Eucgr.H03879.1.p
Eucalyptus___Eucgr.H03878.1.p
Eucalyptus___Eucgr.H03876.1.p
Eucalyptus___Eucgr.H03874.1.p
Eucalyptus___Eucgr.H02836.1.p
Eucalyptus___Eucgr.H02835.1.p
Eucalyptus___Eucgr.H01508.1.p
Eucalyptus___Eucgr.B02117.1.p
Eucalyptus___Eucgr.B02115.1.p
Eucalyptus___Eucgr.B02114.1.p
Eucalyptus___Eucgr.B01692.1.p
Manihot___Manes.10G143300.1.p
Citrus___orange1.1g048223mlPACid_18105663
Cucumis___Cucsa.172140.1lPACid_16966538
F–Carietinum___Ca_00516
F–Prosopis___Palba_XP_028795481.1
F–Prosopis___Palba_XP_028781437.1
F–Lleucocephala___GDRZ01180875.1.p1
F–Akoa___GBYE01052975.1.p1
F–Akoa___GBYE01048877.1.p1
F–Lleucocephala___GDRZ01043017.1.p1
F–Acol___TRINITY_DN58993_c0_g1_i1.p1_complete
F–Pvulgaris___Phvul.008G113000.1.p
F–Pvulgaris___Phvul.003G103800.1.p
F–Pvulgaris___Phvul.009G246700.1.p
F–Pvulgaris___Phvul.009G246600.1.p
F–Carietinum___Ca_24226
F–Pvulgaris___Phvul.009G246500.1.p
F–Carietinum___Ca_24224
F–Carietinum___Ca_24223
F–Carietinum___Ca_24222
F–Carietinum___Ca_27647
F–Carietinum___Ca_20310
F–Carietinum___Ca_24221
F–Carietinum___Ca_27646
F–Carietinum___Ca_24220
F–Carietinum___Ca_20309
F–Carietinum___Ca_20307
F–Carietinum___Ca_20305
F–Carietinum___Ca_20302
F–Carietinum___Ca_20304
F–Carietinum___Ca_20301
F–Carietinum___Ca_20306
F–Pvulgaris___Phvul.003G103700.1.p
F–Pvulgaris___Phvul.003G102700.1.p

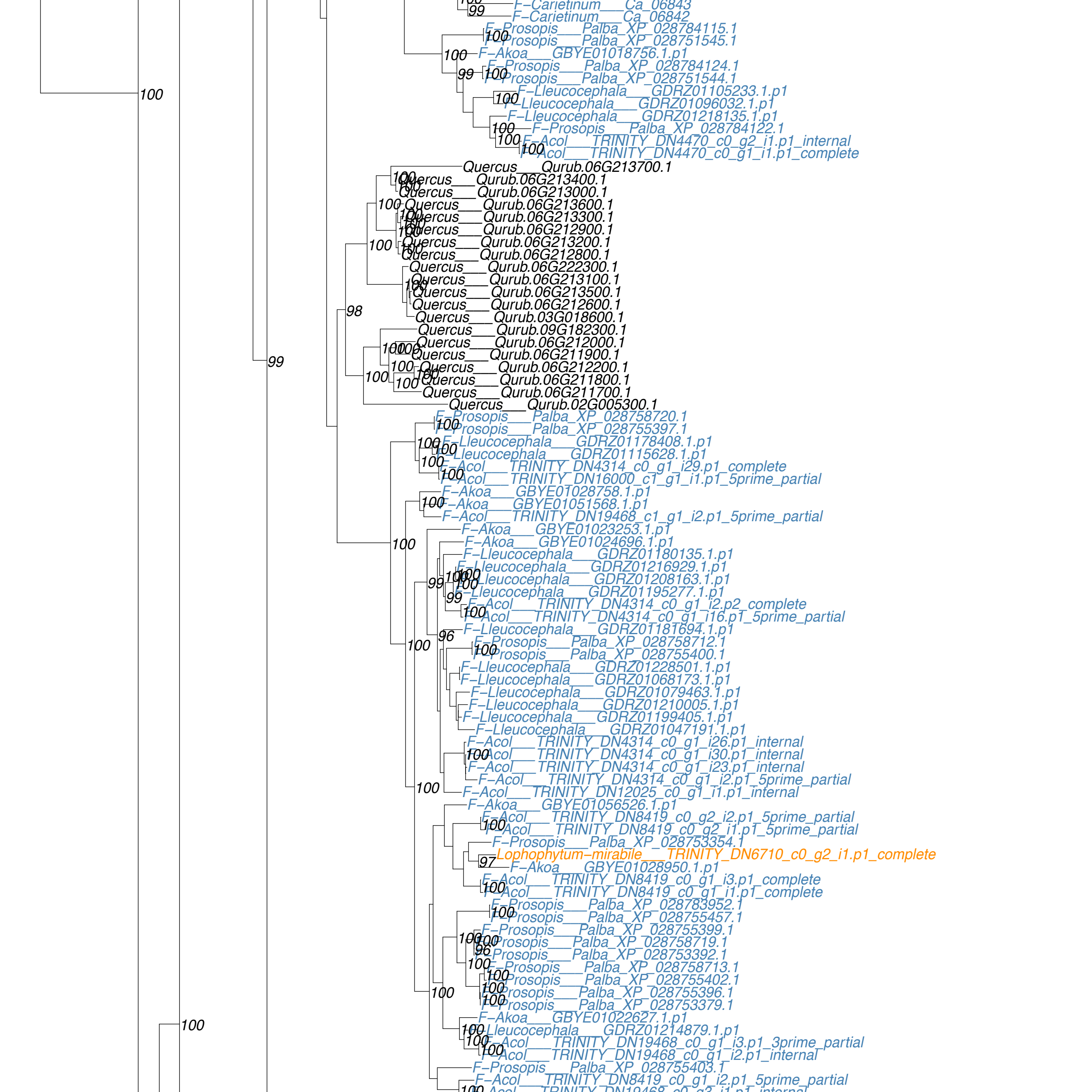

F–Carietinum___Ca_06843
F–Carietinum___Ca_06842
F–Prosopis___Palba_XP_028784115.1
F–Prosopis___Palba_XP_028751545.1
F–Akoa___GBYE01018756.1.p1
F–Prosopis___Palba_XP_028784124.1
F–Prosopis___Palba_XP_028751544.1
F–Lleucocephala___GDRZ01105233.1.p1
F–Lleucocephala___GDRZ01096032.1.p1
F–Lleucocephala___GDRZ01218135.1.p1
F–Prosopis___Palba_XP_028784122.1
F–Acol___TRINITY_DN4470_c0_g2_i1.p1_internal
F–Acol___TRINITY_DN4470_c0_g1_i1.p1_complete
Quercus___Qurub.06G213700.1
Quercus___Qurub.06G213400.1
Quercus___Qurub.06G213000.1
Quercus___Qurub.06G213600.1
Quercus___Qurub.06G213300.1
Quercus___Qurub.06G212900.1
Quercus___Qurub.06G213200.1
Quercus___Qurub.06G212800.1
Quercus___Qurub.06G222300.1
Quercus___Qurub.06G213100.1
Quercus___Qurub.06G213500.1
Quercus___Qurub.06G212600.1
Quercus___Qurub.03G018600.1
Quercus___Qurub.09G182300.1
Quercus___Qurub.06G212000.1
Quercus___Qurub.06G211900.1
Quercus___Qurub.06G212200.1
Quercus___Qurub.06G211800.1
Quercus___Qurub.06G211700.1
Quercus___Qurub.02G005300.1
F–Prosopis___Palba_XP_028758720.1
F–Prosopis___Palba_XP_028755397.1
F–Lleucocephala___GDRZ01178408.1.p1
F–Lleucocephala___GDRZ01115628.1.p1
F–Acol___TRINITY_DN4314_c0_g1_i29.p1_complete
F–Acol___TRINITY_DN16000_c1_g1_i1.p1_5prime_partial
F–Akoa___GBYE01028758.1.p1
F–Akoa___GBYE01051568.1.p1
F–Acol___TRINITY_DN19468_c1_g1_i2.p1_5prime_partial
F–Akoa___GBYE01023253.1.p1
F–Akoa___GBYE01024696.1.p1
F–Lleucocephala___GDRZ01180135.1.p1
F–Lleucocephala___GDRZ01216929.1.p1
F–Lleucocephala___GDRZ01208163.1.p1
F–Lleucocephala___GDRZ01195277.1.p1
F–Acol___TRINITY_DN4314_c0_g1_i2.p2_complete
F–Acol___TRINITY_DN4314_c0_g1_i16.p1_5prime_partial
F–Lleucocephala___GDRZ01181694.1.p1
F–Prosopis___Palba_XP_028758712.1
F–Prosopis___Palba_XP_028755400.1
F–Lleucocephala___GDRZ01228501.1.p1
F–Lleucocephala___GDRZ01068173.1.p1
F–Lleucocephala___GDRZ01079463.1.p1
F–Lleucocephala___GDRZ01210005.1.p1
F–Lleucocephala___GDRZ01199405.1.p1
F–Lleucocephala___GDRZ01047191.1.p1
F–Acol___TRINITY_DN4314_c0_g1_i26.p1_internal
F–Acol___TRINITY_DN4314_c0_g1_i30.p1_internal
F–Acol___TRINITY_DN4314_c0_g1_i23.p1_internal
F–Acol___TRINITY_DN4314_c0_g1_i2.p1_5prime_partial
F–Acol___TRINITY_DN12025_c0_g1_i1.p1_internal
F–Akoa___GBYE01056526.1.p1
F–Acol___TRINITY_DN8419_c0_g2_i2.p1_5prime_partial
F–Acol___TRINITY_DN8419_c0_g2_i1.p1_5prime_partial
F–Prosopis___Palba_XP_028753354.1
Lophophytum–mirabile___TRINITY_DN6710_c0_g2_i1.p1_complete
F–Akoa___GBYE01028950.1.p1
F–Acol___TRINITY_DN8419_c0_g1_i3.p1_complete
F–Acol___TRINITY_DN8419_c0_g1_i1.p1_complete
F–Prosopis___Palba_XP_028783952.1
F–Prosopis___Palba_XP_028755457.1
F–Prosopis___Palba_XP_028755399.1
F–Prosopis___Palba_XP_028758719.1
F–Prosopis___Palba_XP_028753392.1
F–Prosopis___Palba_XP_028758713.1
F–Prosopis___Palba_XP_028755402.1
F–Prosopis___Palba_XP_028755396.1
F–Prosopis___Palba_XP_028753379.1
F–Akoa___GBYE01022627.1.p1
F–Lleucocephala___GDRZ01214879.1.p1
F–Acol___TRINITY_DN19468_c0_g1_i3.p1_3prime_partial
F–Acol___TRINITY_DN19468_c0_g1_i2.p1_internal
F–Prosopis___Palba_XP_028755403.1
F–Acol___TRINITY_DN8419_c0_g1_i2.p1_5prime_partial

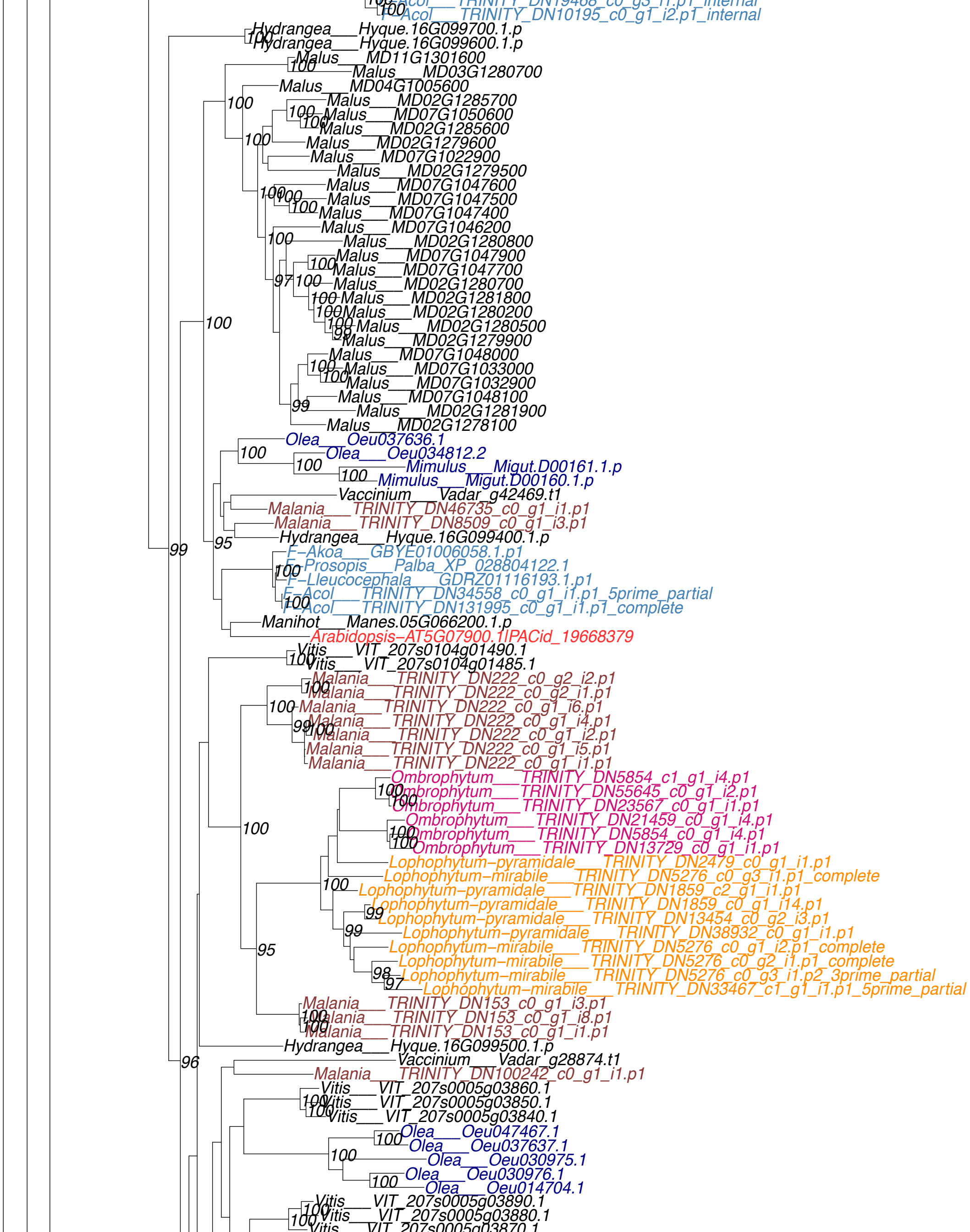

F-Acol___TRINITY_DN10195_c0_g1_i2.p1_internal
Hydrangea___Hyque.16G099700.1.p
Hydrangea___Hyque.16G099600.1.p
Malus___MD11G1301600
Malus___MD03G1280700
Malus___MD04G1005600
Malus___MD02G1285700
Malus___MD07G1050600
Malus___MD02G1285600
Malus___MD02G1279600
Malus___MD07G1022900
Malus___MD02G1279500
Malus___MD07G1047600
Malus___MD07G1047500
Malus___MD07G1047400
Malus___MD07G1046200
Malus___MD02G1280800
Malus___MD07G1047900
Malus___MD07G1047700
Malus___MD02G1280700
Malus___MD02G1281800
Malus___MD02G1280200
Malus___MD02G1280500
Malus___MD02G1279900
Malus___MD07G1048000
Malus___MD07G1033000
Malus___MD07G1032900
Malus___MD07G1048100
Malus___MD02G1281900
Malus___MD02G1278100
Olea___Oeu037636.1
Olea___Oeu034812.2
Mimulus___Migut.D00161.1.p
Mimulus___Migut.D00160.1.p
Vaccinium___Vadar_g42469.t1
Malania___TRINITY_DN46735_c0_g1_i1.p1
Malania___TRINITY_DN8509_c0_g1_i3.p1
Hydrangea___Hyque.16G099400.1.p
F-Akoa___GBYE01006058.1.p1
F-Prosopis___Palba_XP_028804122.1
F-Lleucocephala___GDRZ01116193.1.p1
F-Acol___TRINITY_DN34558_c0_g1_i1.p1_5prime_partial
F-Acol___TRINITY_DN131995_c0_g1_i1.p1_complete
Manihot___Manes.05G066200.1.p
Arabidopsis-AT5G07900.1IPACid_19668379
Vitis___VIT_207s0104g01490.1
Vitis___VIT_207s0104g01485.1
Malania___TRINITY_DN222_c0_g2_i2.p1
Malania___TRINITY_DN222_c0_g2_i1.p1
Malania___TRINITY_DN222_c0_g1_i6.p1
Malania___TRINITY_DN222_c0_g1_i4.p1
Malania___TRINITY_DN222_c0_g1_i2.p1
Malania___TRINITY_DN222_c0_g1_i5.p1
Malania___TRINITY_DN222_c0_g1_i1.p1
Ombrophytum___TRINITY_DN5854_c1_g1_i4.p1
Ombrophytum___TRINITY_DN55645_c0_g1_i2.p1
Ombrophytum___TRINITY_DN23567_c0_g1_i1.p1
Ombrophytum___TRINITY_DN21459_c0_g1_i4.p1
Ombrophytum___TRINITY_DN5854_c0_g1_i4.p1
Ombrophytum___TRINITY_DN13729_c0_g1_i1.p1
Lophophytum-pyramidale___TRINITY_DN2479_c0_g1_i1.p1
Lophophytum-mirabile___TRINITY_DN5276_c0_g3_i1.p1_complete
Lophophytum-pyramidale___TRINITY_DN1859_c2_g1_i1.p1
Lophophytum-pyramidale___TRINITY_DN1859_c0_g1_i14.p1
Lophophytum-pyramidale___TRINITY_DN13454_c0_g2_i3.p1
Lophophytum-pyramidale___TRINITY_DN38932_c0_g1_i1.p1
Lophophytum-mirabile___TRINITY_DN5276_c0_g1_i2.p1_complete
Lophophytum-mirabile___TRINITY_DN5276_c0_g2_i1.p1_complete
Lophophytum-mirabile___TRINITY_DN5276_c0_g3_i1.p2_3prime_partial
Lophophytum-mirabile___TRINITY_DN33467_c1_g1_i1.p1_5prime_partial
Malania___TRINITY_DN153_c0_g1_i3.p1
Malania___TRINITY_DN153_c0_g1_i8.p1
Malania___TRINITY_DN153_c0_g1_i1.p1
Hydrangea___Hyque.16G099500.1.p
Vaccinium___Vadar_g28874.t1
Malania___TRINITY_DN100242_c0_g1_i1.p1
Vitis___VIT_207s0005g03860.1
Vitis___VIT_207s0005g03850.1
Vitis___VIT_207s0005g03840.1
Olea___Oeu047467.1
Olea___Oeu037637.1
Olea___Oeu030975.1
Olea___Oeu030976.1
Olea___Oeu014704.1
Vitis___VIT_207s0005g03890.1
Vitis___VIT_207s0005g03880.1

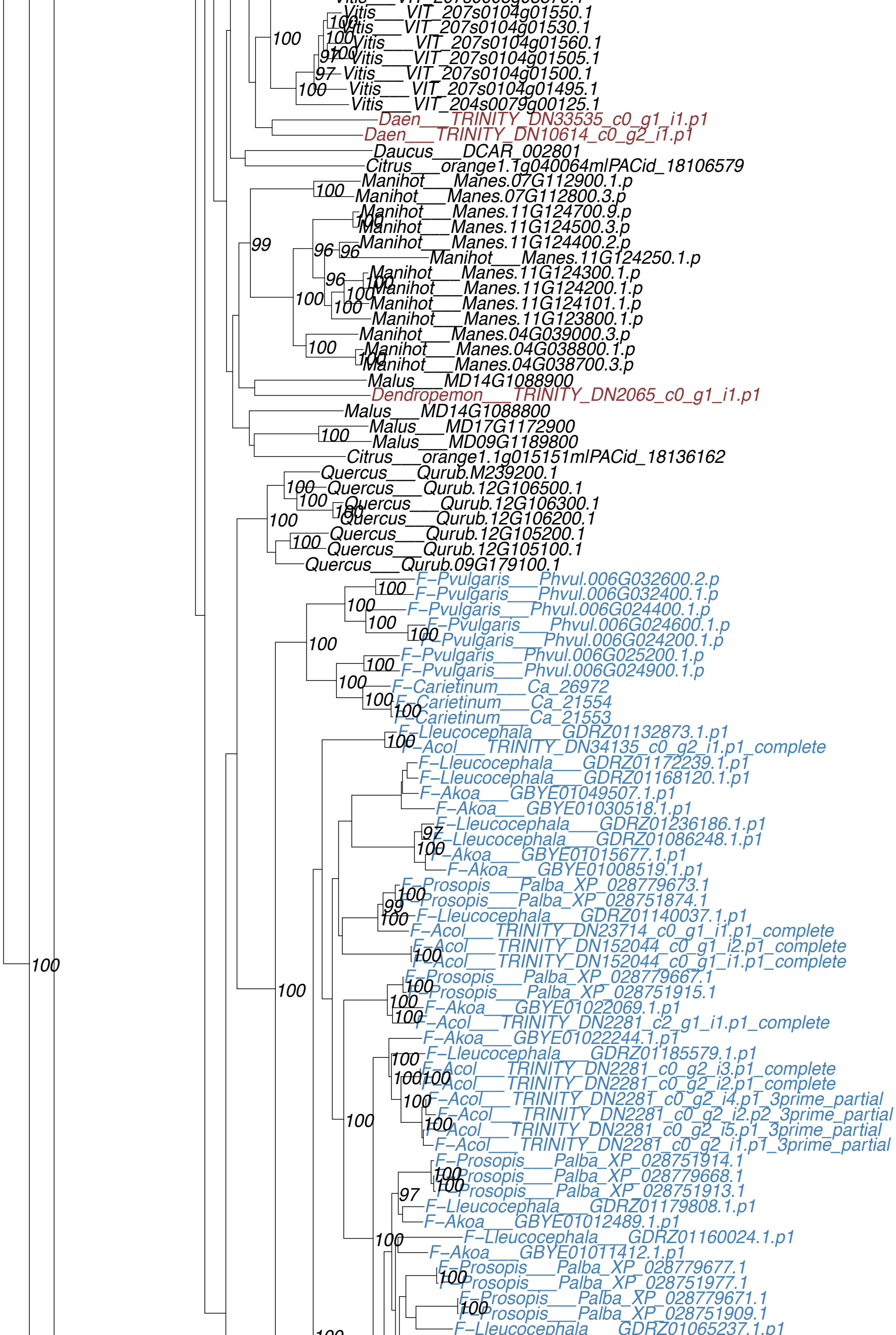
Vitis___VIT_207s0104g01550.1
Vitis___VIT_207s0104g01530.1
Vitis___VIT_207s0104g01560.1
Vitis___VIT_207s0104g01505.1
Vitis___VIT_207s0104g01500.1
Vitis___VIT_207s0104g01495.1
Vitis___VIT_204s0079g00125.1
Daen___TRINITY_DN33535_c0_g1_i1.p1
Daen___TRINITY_DN10614_c0_g2_i1.p1
Daucus___DCAR_002801
Citrus___orange1.1g040064mIPACid_18106579
Manihot___Manes.07G112900.1.p
Manihot___Manes.07G112800.3.p
Manihot___Manes.11G124700.9.p
Manihot___Manes.11G124500.3.p
Manihot___Manes.11G124400.2.p
Manihot___Manes.11G124250.1.p
Manihot___Manes.11G124300.1.p
Manihot___Manes.11G124200.1.p
Manihot___Manes.11G124101.1.p
Manihot___Manes.11G123800.1.p
Manihot___Manes.04G039000.3.p
Manihot___Manes.04G038800.1.p
Manihot___Manes.04G038700.3.p
Malus___MD14G1088900
Dendropemon___TRINITY_DN2065_c0_g1_i1.p1
Malus___MD14G1088800
Malus___MD17G1172900
Malus___MD09G1189800
Citrus___orange1.1g015151mIPACid_18136162
Quercus___Qurub.M239200.1
Quercus___Qurub.12G106500.1
Quercus___Qurub.12G106300.1
Quercus___Qurub.12G106200.1
Quercus___Qurub.12G105200.1
Quercus___Qurub.12G105100.1
Quercus___Qurub.09G179100.1
F–Pvulgaris___Phvul.006G032600.2.p
F–Pvulgaris___Phvul.006G032400.1.p
F–Pvulgaris___Phvul.006G024400.1.p
F–Pvulgaris___Phvul.006G024600.1.p
F–Pvulgaris___Phvul.006G024200.1.p
F–Pvulgaris___Phvul.006G025200.1.p
F–Pvulgaris___Phvul.006G024900.1.p
F–Carietinum___Ca_26972
F–Carietinum___Ca_21554
F–Carietinum___Ca_21553
F–Lleucocephala___GDRZ01132873.1.p1
F–Acol___TRINITY_DN34135_c0_g2_i1.p1_complete
F–Lleucocephala___GDRZ01172239.1.p1
F–Lleucocephala___GDRZ01168120.1.p1
F–Akoa___GBYE01049507.1.p1
F–Akoa___GBYE01030518.1.p1
F–Lleucocephala___GDRZ01236186.1.p1
F–Lleucocephala___GDRZ01086248.1.p1
F–Akoa___GBYE01015677.1.p1
F–Akoa___GBYE01008519.1.p1
F–Prosopis___Palba_XP_028779673.1
F–Prosopis___Palba_XP_028751874.1
F–Lleucocephala___GDRZ01140037.1.p1
F–Acol___TRINITY_DN23714_c0_g1_i1.p1_complete
F–Acol___TRINITY_DN152044_c0_g1_i2.p1_complete
F–Acol___TRINITY_DN152044_c0_g1_i1.p1_complete
F–Prosopis___Palba_XP_028779667.1
F–Prosopis___Palba_XP_028751915.1
F–Akoa___GBYE01022069.1.p1
F–Acol___TRINITY_DN2281_c2_g1_i1.p1_complete
F–Akoa___GBYE01022244.1.p1
F–Lleucocephala___GDRZ01185579.1.p1
F–Acol___TRINITY_DN2281_c0_g2_i3.p1_complete
F–Acol___TRINITY_DN2281_c0_g2_i2.p1_complete
F–Acol___TRINITY_DN2281_c0_g2_i4.p1_3prime_partial
F–Acol___TRINITY_DN2281_c0_g2_i2.p2_3prime_partial
F–Acol___TRINITY_DN2281_c0_g2_i5.p1_3prime_partial
F–Acol___TRINITY_DN2281_c0_g2_i1.p1_3prime_partial
F–Prosopis___Palba_XP_028751914.1
F–Prosopis___Palba_XP_028779668.1
F–Prosopis___Palba_XP_028751913.1
F–Lleucocephala___GDRZ01179808.1.p1
F–Akoa___GBYE01012489.1.p1
F–Lleucocephala___GDRZ01160024.1.p1
F–Akoa___GBYE01011412.1.p1
F–Prosopis___Palba_XP_028779677.1
F–Prosopis___Palba_XP_028751977.1
F–Prosopis___Palba_XP_028779671.1
F–Prosopis___Palba_XP_028751909.1
F–Lleucocephala___GDRZ01065237.1.p1

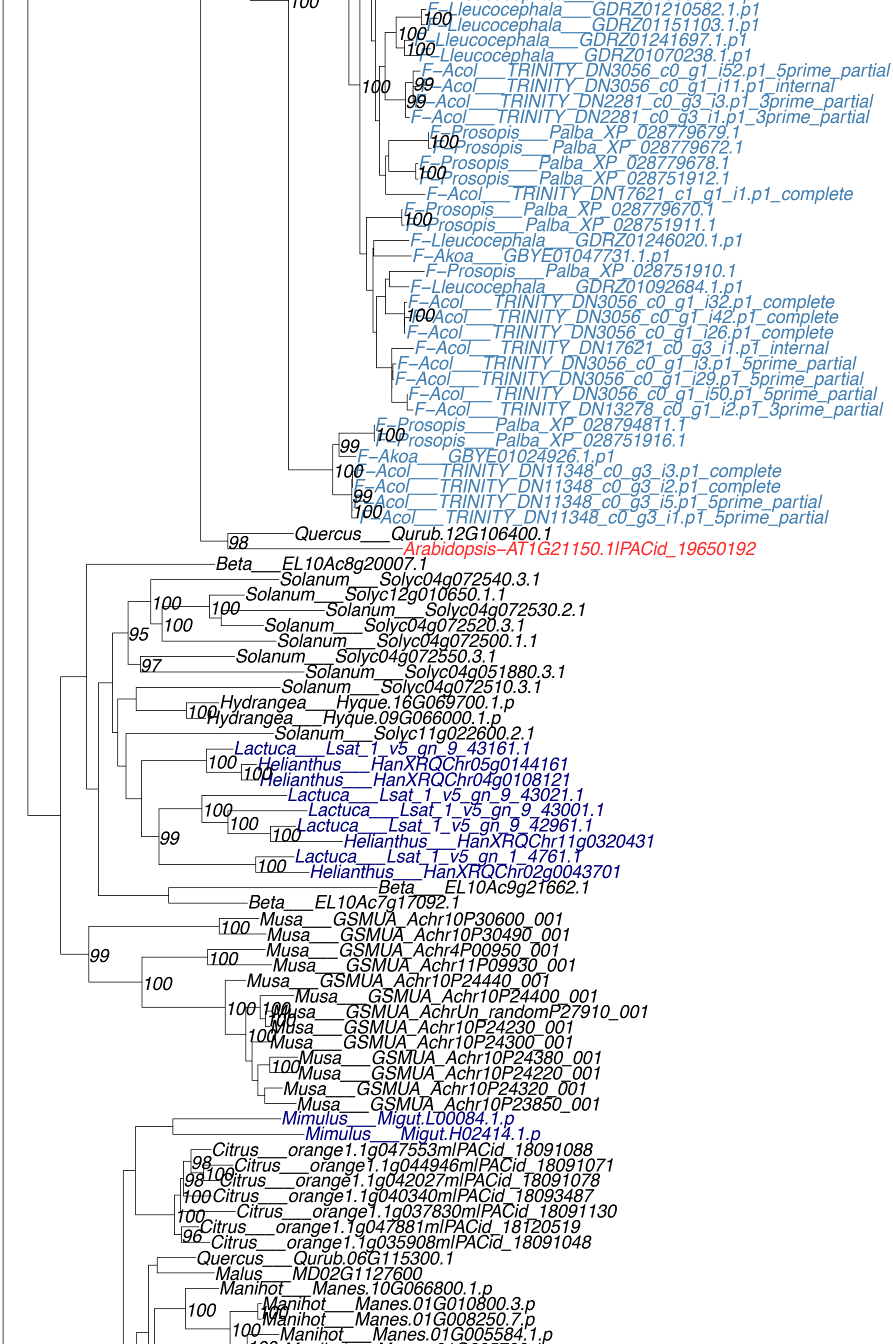
F–Lleucocephala___GDRZ01210582.1.p1
F–Lleucocephala___GDRZ01151103.1.p1
F–Lleucocephala___GDRZ01241697.1.p1
F–Lleucocephala___GDRZ01070238.1.p1
F–Acol___TRINITY_DN3056_c0_g1_i52.p1_5prime_partial
F–Acol___TRINITY_DN3056_c0_g1_i11.p1_internal
F–Acol___TRINITY_DN2281_c0_g3_i3.p1_3prime_partial
F–Acol___TRINITY_DN2281_c0_g3_i1.p1_3prime_partial
F–Prosopis___Palba_XP_028779679.1
F–Prosopis___Palba_XP_028779672.1
F–Prosopis___Palba_XP_028779678.1
F–Prosopis___Palba_XP_028751912.1
F–Acol___TRINITY_DN17621_c1_g1_i1.p1_complete
F–Prosopis___Palba_XP_028779670.1
F–Prosopis___Palba_XP_028751911.1
F–Lleucocephala___GDRZ01246020.1.p1
F–Akoa___GBYE01047731.1.p1
F–Prosopis___Palba_XP_028751910.1
F–Lleucocephala___GDRZ01092684.1.p1
F–Acol___TRINITY_DN3056_c0_g1_i32.p1_complete
F–Acol___TRINITY_DN3056_c0_g1_i42.p1_complete
F–Acol___TRINITY_DN3056_c0_g1_i26.p1_complete
F–Acol___TRINITY_DN17621_c0_g3_i1.p1_internal
F–Acol___TRINITY_DN3056_c0_g1_i3.p1_5prime_partial
F–Acol___TRINITY_DN3056_c0_g1_i29.p1_5prime_partial
F–Acol___TRINITY_DN3056_c0_g1_i50.p1_5prime_partial
F–Acol___TRINITY_DN13278_c0_g1_i2.p1_3prime_partial
F–Prosopis___Palba_XP_028794811.1
F–Prosopis___Palba_XP_028751916.1
F–Akoa___GBYE01024926.1.p1
F–Acol___TRINITY_DN11348_c0_g3_i3.p1_complete
F–Acol___TRINITY_DN11348_c0_g3_i2.p1_complete
F–Acol___TRINITY_DN11348_c0_g3_i5.p1_5prime_partial
F–Acol___TRINITY_DN11348_c0_g3_i1.p1_5prime_partial
Quercus___Qurub.12G106400.1
Arabidopsis–AT1G21150.1lPACid_19650192
Beta___EL10Ac8g20007.1
Solanum___Solyc04g072540.3.1
Solanum___Solyc12g010650.1.1
Solanum___Solyc04g072530.2.1
Solanum___Solyc04g072520.3.1
Solanum___Solyc04g072500.1.1
Solanum___Solyc04g072550.3.1
Solanum___Solyc04g051880.3.1
Solanum___Solyc04g072510.3.1
Hydrangea___Hyque.16G069700.1.p
Hydrangea___Hyque.09G066000.1.p
Solanum___Solyc11g022600.2.1
Lactuca___Lsat_1_v5_gn_9_43161.1
Helianthus___HanXRQChr05g0144161
Helianthus___HanXRQChr04g0108121
Lactuca___Lsat_1_v5_gn_9_43021.1
Lactuca___Lsat_1_v5_gn_9_43001.1
Lactuca___Lsat_1_v5_gn_9_42961.1
Helianthus___HanXRQChr11g0320431
Lactuca___Lsat_1_v5_gn_1_4761.1
Helianthus___HanXRQChr02g0043701
Beta___EL10Ac9g21662.1
Beta___EL10Ac7g17092.1
Musa___GSMUA_Achr10P30600_001
Musa___GSMUA_Achr10P30490_001
Musa___GSMUA_Achr4P00950_001
Musa___GSMUA_Achr11P09930_001
Musa___GSMUA_Achr10P24440_001
Musa___GSMUA_Achr10P24400_001
Musa___GSMUA_AchrUn_randomP27910_001
Musa___GSMUA_Achr10P24230_001
Musa___GSMUA_Achr10P24300_001
Musa___GSMUA_Achr10P24380_001
Musa___GSMUA_Achr10P24220_001
Musa___GSMUA_Achr10P24320_001
Musa___GSMUA_Achr10P23850_001
Mimulus___Migut.L00084.1.p
Mimulus___Migut.H02414.1.p
Citrus___orange1.1g047553mlPACid_18091088
Citrus___orange1.1g044946mlPACid_18091071
Citrus___orange1.1g042027mlPACid_18091078
Citrus___orange1.1g040340mlPACid_18093487
Citrus___orange1.1g037830mlPACid_18091130
Citrus___orange1.1g047881mlPACid_18120519
Citrus___orange1.1g035908mlPACid_18091048
Quercus___Qurub.06G115300.1
Malus___MD02G1127600
Manihot___Manes.10G066800.1.p
Manihot___Manes.01G010800.3.p
Manihot___Manes.01G008250.7.p
Manihot___Manes.01G005584.1.p

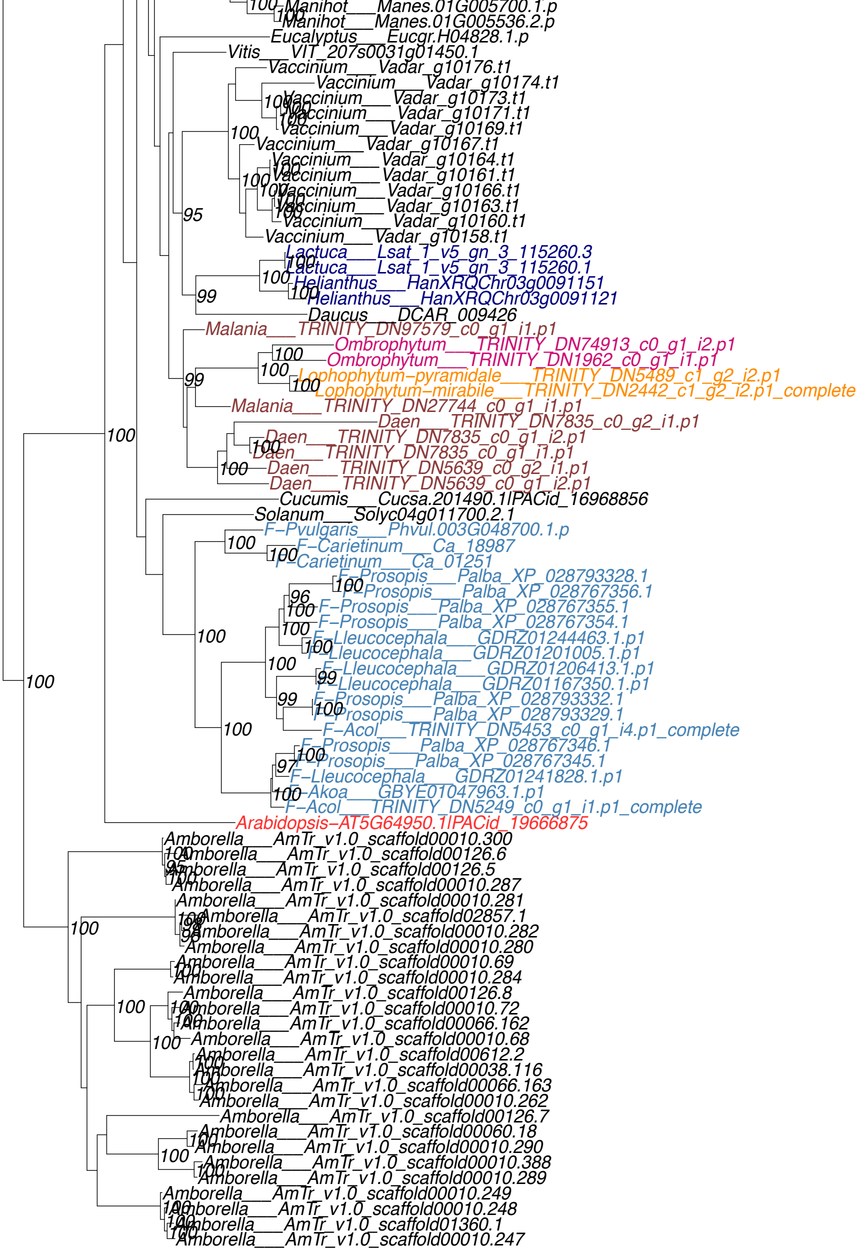

Manihot___Manes.01G005700.1.p
Manihot___Manes.01G005536.2.p
Eucalyptus___Eucgr.H04828.1.p
Vitis___VIT_207s0031g01450.1
Vaccinium___Vadar_g10176.t1
Vaccinium___Vadar_g10174.t1
Vaccinium___Vadar_g10173.t1
Vaccinium___Vadar_g10171.t1
Vaccinium___Vadar_g10169.t1
Vaccinium___Vadar_g10167.t1
Vaccinium___Vadar_g10164.t1
Vaccinium___Vadar_g10161.t1
Vaccinium___Vadar_g10166.t1
Vaccinium___Vadar_g10163.t1
Vaccinium___Vadar_g10160.t1
Vaccinium___Vadar_g10158.t1
Lactuca___Lsat_1_v5_gn_3_115260.3
Lactuca___Lsat_1_v5_gn_3_115260.1
Helianthus___HanXRQChr03g0091151
Helianthus___HanXRQChr03g0091121
Daucus___DCAR_009426
Malania___TRINITY_DN97579_c0_g1_i1.p1
Ombrophytum___TRINITY_DN74913_c0_g1_i2.p1
Ombrophytum___TRINITY_DN1962_c0_g1_i1.p1
Lophophytum–pyramidale___TRINITY_DN5489_c1_g2_i2.p1
Lophophytum–mirabile___TRINITY_DN2442_c1_g2_i2.p1_complete
Malania___TRINITY_DN27744_c0_g1_i1.p1
Daen___TRINITY_DN7835_c0_g2_i1.p1
Daen___TRINITY_DN7835_c0_g1_i2.p1
Daen___TRINITY_DN7835_c0_g1_i1.p1
Daen___TRINITY_DN5639_c0_g2_i1.p1
Daen___TRINITY_DN5639_c0_g1_i2.p1
Cucumis___Cucsa.201490.1lPACid_16968856
Solanum___Solyc04g011700.2.1
F–Pvulgaris___Phvul.003G048700.1.p
F–Carietinum___Ca_18987
F–Carietinum___Ca_01251
F–Prosopis___Palba_XP_028793328.1
F–Prosopis___Palba_XP_028767356.1
F–Prosopis___Palba_XP_028767355.1
F–Prosopis___Palba_XP_028767354.1
F–Lleucocephala___GDRZ01244463.1.p1
F–Lleucocephala___GDRZ01201005.1.p1
F–Lleucocephala___GDRZ01206413.1.p1
F–Lleucocephala___GDRZ01167350.1.p1
F–Prosopis___Palba_XP_028793332.1
F–Prosopis___Palba_XP_028793329.1
F–Acol___TRINITY_DN5453_c0_g1_i4.p1_complete
F–Prosopis___Palba_XP_028767346.1
F–Prosopis___Palba_XP_028767345.1
F–Lleucocephala___GDRZ01241828.1.p1
F–Akoa___GBYE01047963.1.p1
F–Acol___TRINITY_DN5249_c0_g1_i1.p1_complete
Arabidopsis–AT5G64950.1lPACid_19666875
Amborella___AmTr_v1.0_scaffold00010.300
Amborella___AmTr_v1.0_scaffold00126.6
Amborella___AmTr_v1.0_scaffold00126.5
Amborella___AmTr_v1.0_scaffold00010.287
Amborella___AmTr_v1.0_scaffold00010.281
Amborella___AmTr_v1.0_scaffold02857.1
Amborella___AmTr_v1.0_scaffold00010.282
Amborella___AmTr_v1.0_scaffold00010.280
Amborella___AmTr_v1.0_scaffold00010.69
Amborella___AmTr_v1.0_scaffold00010.284
Amborella___AmTr_v1.0_scaffold00126.8
Amborella___AmTr_v1.0_scaffold00010.72
Amborella___AmTr_v1.0_scaffold00066.162
Amborella___AmTr_v1.0_scaffold00010.68
Amborella___AmTr_v1.0_scaffold00612.2
Amborella___AmTr_v1.0_scaffold00038.116
Amborella___AmTr_v1.0_scaffold00066.163
Amborella___AmTr_v1.0_scaffold00010.262
Amborella___AmTr_v1.0_scaffold00126.7
Amborella___AmTr_v1.0_scaffold00060.18
Amborella___AmTr_v1.0_scaffold00010.290
Amborella___AmTr_v1.0_scaffold00010.388
Amborella___AmTr_v1.0_scaffold00010.289
Amborella___AmTr_v1.0_scaffold00010.249
Amborella___AmTr_v1.0_scaffold00010.248
Amborella___AmTr_v1.0_scaffold01360.1
Amborella___AmTr_v1.0_scaffold00010.247

# OG0000645

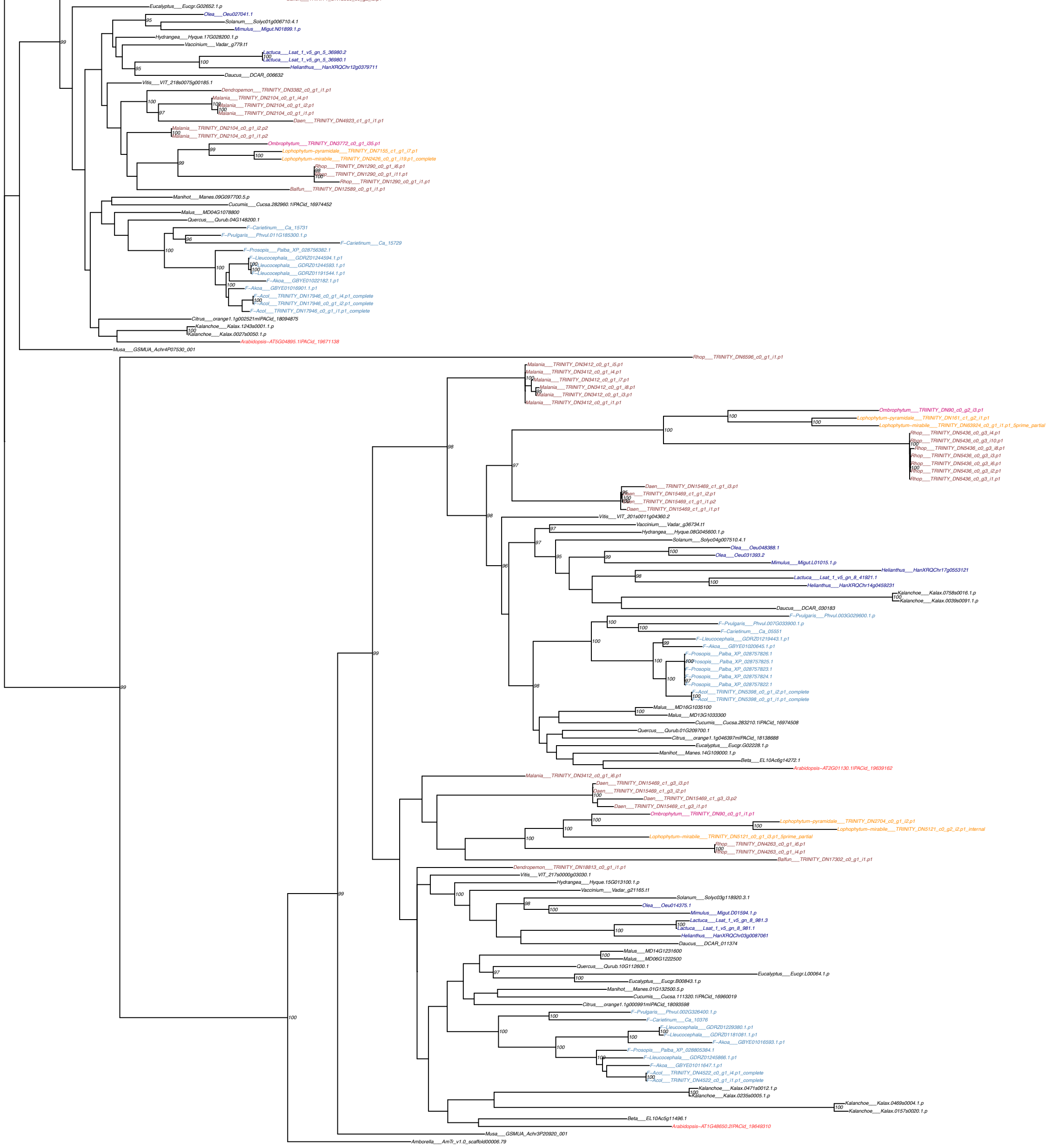

Amborella___AmTr_v1.0_scaffold00004.98
Beta___EL10Ac4g08754.1
Balfun___TRINITY_DN12589_c0_g2_i2.p1
Eucalyptus___Eucgr.G02652.1.p
Olea___Oeu027041.1
Solanum___Solyc01g006710.4.1
Mimulus___Migut.N01899.1.p
Hydrangea___Hyque.17G028200.1.p
Vaccinium___Vadar_g779.t1
Lactuca___Lsat_1_v5_gn_5_36980.2
Lactuca___Lsat_1_v5_gn_5_36980.1
Helianthus___HanXRQChr12g0379711
Daucus___DCAR_006632
Vitis___VIT_218s0075g00185.1
Dendropemon___TRINITY_DN3382_c0_g1_i1.p1
Malania___TRINITY_DN2104_c0_g1_i4.p1
Malania___TRINITY_DN2104_c0_g1_i2.p1
Malania___TRINITY_DN2104_c0_g1_i1.p1
Daen___TRINITY_DN4923_c1_g1_i1.p1
Malania___TRINITY_DN2104_c0_g1_i2.p2
Malania___TRINITY_DN2104_c0_g1_i1.p2
Ombrophytum___TRINITY_DN3772_c0_g1_i35.p1
Lophophytum-pyramidale___TRINITY_DN7155_c1_g1_i7.p1
Lophophytum-mirabile___TRINITY_DN2426_c0_g1_i19.p1_complete
Rhop___TRINITY_DN1290_c0_g1_i6.p1
Rhop___TRINITY_DN1290_c0_g1_i11.p1
Rhop___TRINITY_DN1290_c0_g1_i1.p1
Balfun___TRINITY_DN12589_c0_g1_i1.p1
Manihot___Manes.09G097700.5.p
Cucumis___Cucsa.282960.1|PACid_16974452
Malus___MD04G1078800
Quercus___Qurub.04G148200.1
F-Carietinum___Ca_15731
F-Pvulgaris___Phvul.011G185300.1.p
F-Carietinum___Ca_15729
F-Prosopis___Palba_XP_028756382.1
F-Lleucocephala___GDRZ01244594.1.p1
F-Lleucocephala___GDRZ01244593.1.p1
F-Lleucocephala___GDRZ01191544.1.p1
F-Akoa___GBYE01022182.1.p1
F-Akoa___GBYE01016901.1.p1
F-Acol___TRINITY_DN17946_c0_g1_i4.p1_complete
F-Acol___TRINITY_DN17946_c0_g1_i2.p1_complete
F-Acol___TRINITY_DN17946_c0_g1_i1.p1_complete
Citrus___orange1.1g002521m|PACid_18094875
Kalanchoe___Kalax.1243s0001.1.p
Kalanchoe___Kalax.0027s0050.1.p
Arabidopsis-AT5G04895.1|PACid_19671138
Musa___GSMUA_Achr4P07530_001
Rhop___TRINITY_DN6596_c0_g1_i1.p1
Malania___TRINITY_DN3412_c0_g1_i5.p1
Malania___TRINITY_DN3412_c0_g1_i4.p1
Malania___TRINITY_DN3412_c0_g1_i7.p1
Malania___TRINITY_DN3412_c0_g1_i8.p1
Malania___TRINITY_DN3412_c0_g1_i3.p1
Malania___TRINITY_DN3412_c0_g1_i1.p1
Ombrophytum___TRINITY_DN90_c0_g2_i3.p1
Lophophytum-pyramidale___TRINITY_DN161_c1_g2_i1.p1
Lophophytum-mirabile___TRINITY_DN63924_c0_g1_i1.p1_5prime_partial
Rhop___TRINITY_DN5436_c0_g3_i4.p1
Rhop___TRINITY_DN5436_c0_g3_i10.p1
Rhop___TRINITY_DN5436_c0_g3_i8.p1
Rhop___TRINITY_DN5436_c0_g3_i3.p1
Rhop___TRINITY_DN5436_c0_g3_i6.p1
Rhop___TRINITY_DN5436_c0_g3_i2.p1
Rhop___TRINITY_DN5436_c0_g3_i1.p1
Daen___TRINITY_DN15469_c1_g1_i3.p1
Daen___TRINITY_DN15469_c1_g1_i2.p1
Daen___TRINITY_DN15469_c1_g1_i1.p2
Daen___TRINITY_DN15469_c1_g1_i1.p1
Vitis___VIT_201s0011g04360.2
Vaccinium___Vadar_g36734.t1
Hydrangea___Hyque.08G045600.1.p
Solanum___Solyc04g007510.4.1
Olea___Oeu048388.1
Olea___Oeu031393.2
Mimulus___Migut.L01015.1.p
Helianthus___HanXRQChr17g0553121
Lactuca___Lsat_1_v5_gn_8_41921.1
Helianthus___HanXRQChr14g0459231
Kalanchoe___Kalax.0758s0016.1.p
Kalanchoe___Kalax.0039s0091.1.p
Daucus___DCAR_030183
F-Pvulgaris___Phvul.003G029600.1.p
F-Pvulgaris___Phvul.007G033900.1.p
F-Carietinum___Ca_05551
F-Lleucocephala___GDRZ01219443.1.p1
F-Akoa___GBYE01020645.1.p1
F-Prosopis___Palba_XP_028757826.1
F-Prosopis___Palba_XP_028757825.1
F-Prosopis___Palba_XP_028757823.1
F-Prosopis___Palba_XP_028757824.1
F-Prosopis___Palba_XP_028757822.1
F-Acol___TRINITY_DN5398_c0_g1_i2.p1_complete
F-Acol___TRINITY_DN5398_c0_g1_i1.p1_complete
Malus___MD16G1035100
Malus___MD13G1033300
Cucumis___Cucsa.283210.1|PACid_16974508
Quercus___Qurub.01G209700.1
Citrus___orange1.1g046397m|PACid_18138688
Eucalyptus___Eucgr.G02228.1.p
Manihot___Manes.14G109000.1.p
Beta___EL10Ac6g14272.1
Arabidopsis-AT2G01130.1|PACid_19639162
Malania___TRINITY_DN3412_c0_g1_i6.p1
Daen___TRINITY_DN15469_c1_g3_i3.p1
Daen___TRINITY_DN15469_c1_g3_i2.p1
Daen___TRINITY_DN15469_c1_g3_i3.p2
Daen___TRINITY_DN15469_c1_g3_i1.p1
Ombrophytum___TRINITY_DN90_c0_g1_i1.p1
Lophophytum-pyramidale___TRINITY_DN2704_c0_g1_i2.p1
Lophophytum-mirabile___TRINITY_DN5121_c0_g2_i2.p1_internal
Lophophytum-mirabile___TRINITY_DN5121_c0_g1_i3.p1_5prime_partial
Rhop___TRINITY_DN4263_c0_g1_i6.p1
Rhop___TRINITY_DN4263_c0_g1_i4.p1
Balfun___TRINITY_DN17302_c0_g1_i1.p1
Dendropemon___TRINITY_DN18813_c0_g1_i1.p1
Vitis___VIT_217s0000g03030.1
Hydrangea___Hyque.15G013100.1.p
Vaccinium___Vadar_g21165.t1
Solanum___Solyc03g118920.3.1
Olea___Oeu014375.1
Mimulus___Migut.D01594.1.p
Lactuca___Lsat_1_v5_gn_8_981.3
Lactuca___Lsat_1_v5_gn_8_981.1
Helianthus___HanXRQChr03g0087061
Daucus___DCAR_011374
Malus___MD14G1231600
Malus___MD06G1222500
Quercus___Qurub.10G112600.1
Eucalyptus___Eucgr.L00064.1.p
Eucalyptus___Eucgr.B00843.1.p
Manihot___Manes.01G132500.5.p
Cucumis___Cucsa.111320.1|PACid_16960019
Citrus___orange1.1g000991m|PACid_18093598
F-Pvulgaris___Phvul.002G326400.1.p
F-Carietinum___Ca_10376
F-Lleucocephala___GDRZ01229380.1.p1
F-Lleucocephala___GDRZ01181081.1.p1
F-Akoa___GBYE01016593.1.p1
F-Prosopis___Palba_XP_028805384.1
F-Lleucocephala___GDRZ01245866.1.p1
F-Akoa___GBYE01011647.1.p1
F-Acol___TRINITY_DN4522_c0_g1_i4.p1_complete
F-Acol___TRINITY_DN4522_c0_g1_i1.p1_complete
Kalanchoe___Kalax.0471s0012.1.p
Kalanchoe___Kalax.0235s0005.1.p
Kalanchoe___Kalax.0469s0004.1.p
Kalanchoe___Kalax.0157s0020.1.p
Beta___EL10Ac5g11496.1
Arabidopsis-AT1G48650.2|PACid_19649310
Musa___GSMUA_Achr3P20920_001
Amborella___AmTr_v1.0_scaffold00006.79

## OG0000797

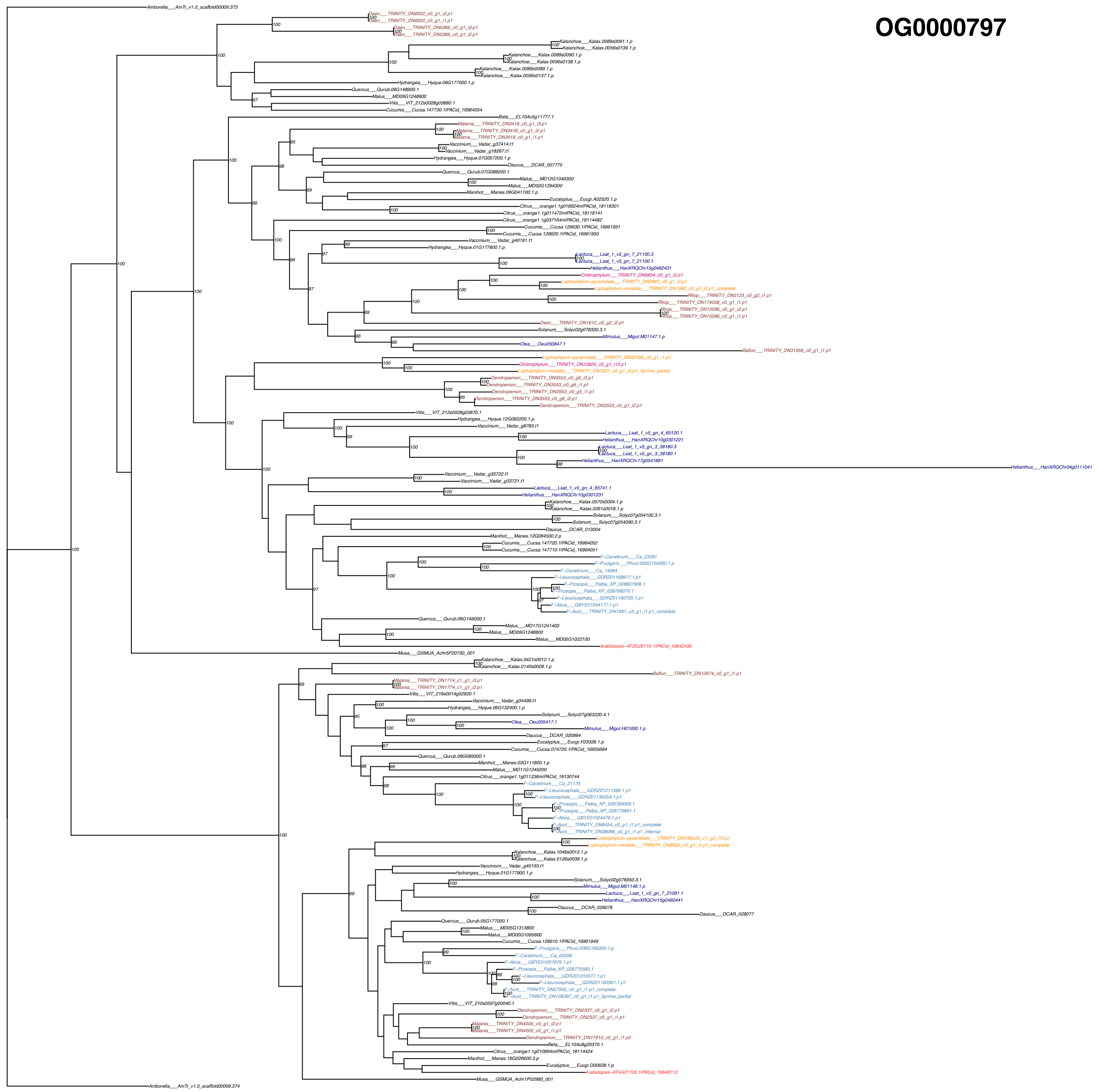

Amborella___AmTr_v1.0_scaffold00009.373
Daen___TRINITY_DN6002_c0_g1_i2.p1
Daen___TRINITY_DN6002_c0_g1_i1.p1
Daen___TRINITY_DN5366_c0_g1_i3.p1
Daen___TRINITY_DN5366_c0_g1_i2.p1
Kalanchoe___Kalax.0089s0091.1.p
Kalanchoe___Kalax.0056s0139.1.p
Kalanchoe___Kalax.0089s0090.1.p
Kalanchoe___Kalax.0056s0138.1.p
Kalanchoe___Kalax.0089s0089.1.p
Kalanchoe___Kalax.0056s0137.1.p
Hydrangea___Hyque.06G177500.1.p
Quercus___Qurub.08G148900.1
Malus___MD09G1248900
Vitis___VIT_212s0028g03880.1
Cucumis___Cucsa.147730.1PACid_16964054
Beta___EL10Ac5g11777.1
Malania___TRINITY_DN3418_c0_g1_i3.p1
Malania___TRINITY_DN3418_c0_g1_i2.p1
Malania___TRINITY_DN3418_c0_g1_i1.p1
Vaccinium___Vadar_g37414.t1
Vaccinium___Vadar_g18267.t1
Hydrangea___Hyque.07G057200.1.p
Daucus___DCAR_007775
Quercus___Qurub.07G088200.1
Malus___MD12G1049300
Malus___MD02G1294300
Manihot___Manes.09G041100.1.p
Eucalyptus___Eucgr.A02520.1.p
Citrus___orange1.1g018924mPACid_18118301
Citrus___orange1.1g011472mPACid_18118141
Citrus___orange1.1g037164mPACid_18114482
Cucumis___Cucsa.126630.1PACid_16961951
Cucumis___Cucsa.126620.1PACid_16961950
Vaccinium___Vadar_g40191.t1
Hydrangea___Hyque.01G177800.1.p
Lactuca___Lsat_1_v5_gn_7_21100.3
Lactuca___Lsat_1_v5_gn_7_21100.1
Helianthus___HanXRQChr15g0482431
Ombrophytum___TRINITY_DN9854_c0_g1_i3.p1
Lophophytum-pyramidale___TRINITY_DN5881_c0_g1_i2.p1
Lophophytum-mirabile___TRINITY_DN1982_c0_g1_i3.p1_complete
Rhop___TRINITY_DN3123_c0_g2_i1.p1
Rhop___TRINITY_DN174058_c0_g1_i1.p1
Rhop___TRINITY_DN15096_c0_g1_i2.p1
Rhop___TRINITY_DN15096_c0_g1_i1.p1
Daen___TRINITY_DN1612_c0_g2_i2.p1
Solanum___Solyc02g078330.3.1
Mimulus___Migut.M01147.1.p
Olea___Oeu050847.1
Balfun___TRINITY_DN31058_c0_g1_i1.p1
Lophophytum-pyramidale___TRINITY_DN32766_c0_g1_i1.p1
Ombrophytum___TRINITY_DN15820_c0_g1_i13.p1
Lophophytum-mirabile___TRINITY_DN7527_c0_g1_i2.p1_5prime_partial
Dendropemon___TRINITY_DN3553_c0_g6_i3.p1
Dendropemon___TRINITY_DN3553_c0_g6_i1.p1
Dendropemon___TRINITY_DN3553_c0_g5_i1.p1
Dendropemon___TRINITY_DN3553_c0_g6_i2.p1
Dendropemon___TRINITY_DN3553_c0_g1_i2.p1
Vitis___VIT_212s0028g03870.1
Hydrangea___Hyque.12G092200.1.p
Vaccinium___Vadar_g8783.t1
Lactuca___Lsat_1_v5_gn_4_65120.1
Helianthus___HanXRQChr10g0301221
Lactuca___Lsat_1_v5_gn_3_38180.3
Lactuca___Lsat_1_v5_gn_3_38180.1
Helianthus___HanXRQChr17g0541881
Helianthus___HanXRQChr04g0111041
Vaccinium___Vadar_g33722.t1
Vaccinium___Vadar_g33721.t1
Lactuca___Lsat_1_v5_gn_4_65741.1
Helianthus___HanXRQChr10g0301231
Kalanchoe___Kalax.0570s0004.1.p
Kalanchoe___Kalax.0261s0018.1.p
Solanum___Solyc07g054100.3.1
Solanum___Solyc07g054090.3.1
Daucus___DCAR_013004
Manihot___Manes.12G084500.2.p
Cucumis___Cucsa.147720.1PACid_16964052
Cucumis___Cucsa.147710.1PACid_16964051
F-Carietinum___Ca_23281
F-Pvulgaris___Phvul.005G104300.1.p
F-Carietinum___Ca_14084
F-Lleucocephala___GDRZ01168977.1.p1
F-Prosopis___Palba_XP_028807908.1
F-Prosopis___Palba_XP_028798070.1
F-Lleucocephala___GDRZ01190725.1.p1
F-Akoa___GBYE01044177.1.p1
F-Acol___TRINITY_DN1391_c3_g1_i1.p1_complete
Quercus___Qurub.08G149000.1
Malus___MD17G1241400
Malus___MD09G1248800
Malus___MD05G1022100
Arabidopsis-AT2G20710.1PACid_19642435
Musa___GSMUA_Achr5P20750_001
Kalanchoe___Kalax.0421s0012.1.p
Kalanchoe___Kalax.0140s0008.1.p
Balfun___TRINITY_DN12674_c0_g1_i1.p1
Malania___TRINITY_DN1774_c1_g1_i3.p1
Malania___TRINITY_DN1774_c1_g1_i2.p1
Vitis___VIT_219s0014g02920.1
Vaccinium___Vadar_g34496.t1
Hydrangea___Hyque.06G132400.1.p
Solanum___Solyc07g063220.4.1
Olea___Oeu005417.1
Mimulus___Migut.H01930.1.p
Daucus___DCAR_020984
Eucalyptus___Eucgr.F03326.1.p
Cucumis___Cucsa.074720.1PACid_16955684
Quercus___Qurub.08G089500.1
Manihot___Manes.03G111800.1.p
Malus___MD11G1245200
Citrus___orange1.1g011236mPACid_18130744
F-Carietinum___Ca_21176
F-Lleucocephala___GDRZ01211380.1.p1
F-Lleucocephala___GDRZ01134354.1.p1
F-Prosopis___Palba_XP_028783068.1
F-Prosopis___Palba_XP_028779891.1
F-Akoa___GBYE01024476.1.p1
F-Acol___TRINITY_DN8454_c0_g1_i1.p1_complete
F-Acol___TRINITY_DN38066_c0_g1_i1.p1_internal
Lophophytum-pyramidale___TRINITY_DN199522_c1_g2_i13.p1
Lophophytum-mirabile___TRINITY_DN8592_c0_g1_i1.p1_complete
Kalanchoe___Kalax.1046s0012.1.p
Kalanchoe___Kalax.0126s0036.1.p
Vaccinium___Vadar_g40193.t1
Hydrangea___Hyque.01G177900.1.p
Solanum___Solyc02g078350.3.1
Mimulus___Migut.M01148.1.p
Lactuca___Lsat_1_v5_gn_7_21081.1
Helianthus___HanXRQChr15g0482441
Daucus___DCAR_028078
Daucus___DCAR_028077
Quercus___Qurub.05G177000.1
Malus___MD05G1313800
Malus___MD00G1095800
Cucumis___Cucsa.126610.1PACid_16961949
F-Pvulgaris___Phvul.006G186200.1.p
F-Carietinum___Ca_00298
F-Akoa___GBYE01007879.1.p1
F-Prosopis___Palba_XP_028775583.1
F-Lleucocephala___GDRZ01210577.1.p1
F-Lleucocephala___GDRZ01182361.1.p1
F-Acol___TRINITY_DN57955_c0_g1_i1.p1_complete
F-Acol___TRINITY_DN128397_c0_g1_i1.p1_5prime_partial
Vitis___VIT_210s0597g00040.1
Dendropemon___TRINITY_DN2337_c0_g1_i2.p1
Dendropemon___TRINITY_DN2337_c0_g1_i1.p1
Malania___TRINITY_DN4505_c0_g1_i2.p1
Malania___TRINITY_DN4505_c0_g1_i1.p1
Dendropemon___TRINITY_DN17910_c0_g1_i1.p2
Beta___EL10Ac8g20370.1
Citrus___orange1.1g010994mPACid_18114424
Manihot___Manes.16G026600.3.p
Eucalyptus___Eucgr.D00628.1.p
Arabidopsis-AT4G21705.1PACid_19646713
Musa___GSMUA_Achr1P02980_001
Amborella___AmTr_v1.0_scaffold00009.374

# OG0003049

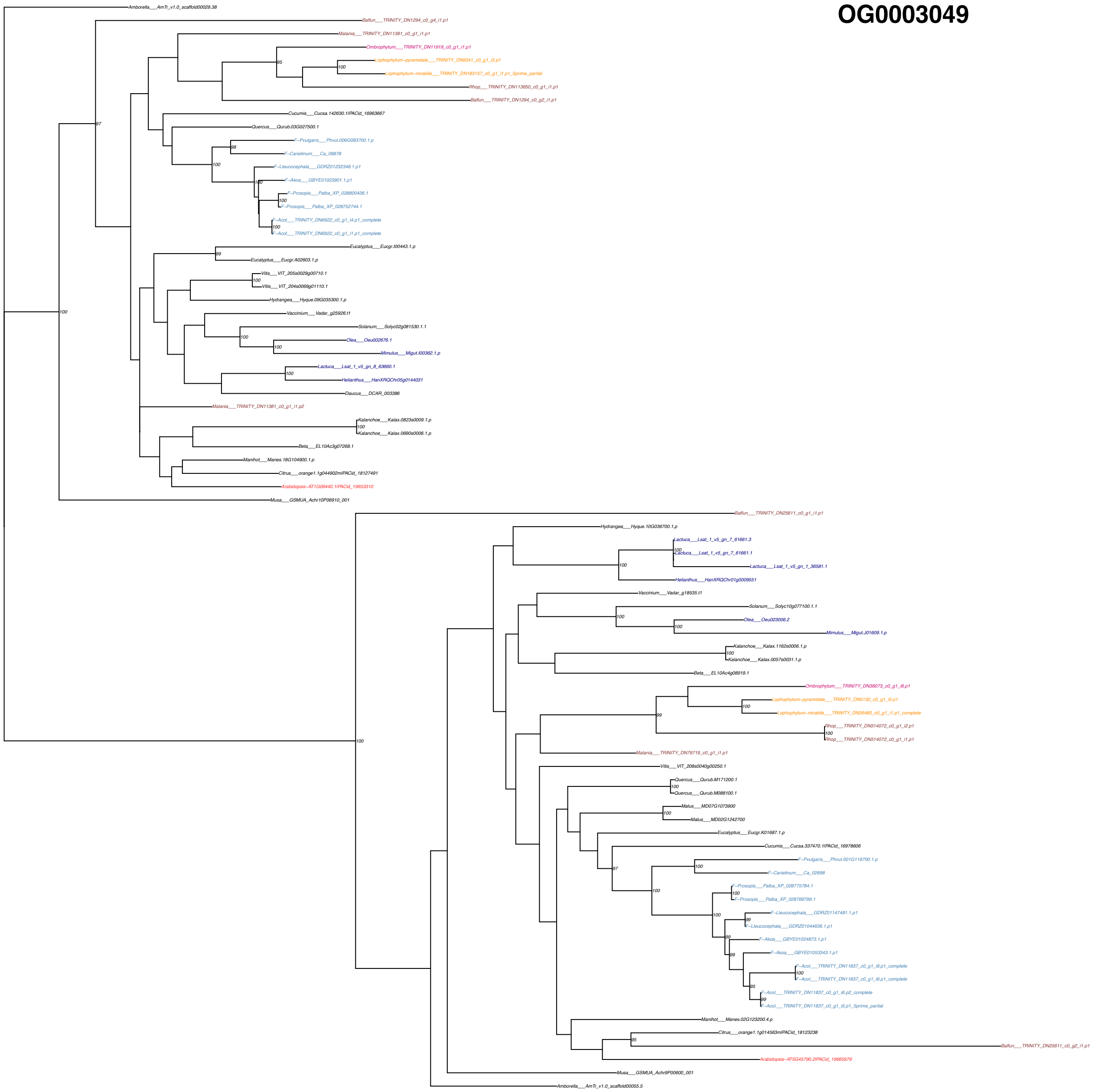

Amborella___AmTr_v1.0_scaffold00029.38
Balfun___TRINITY_DN1294_c0_g4_i1.p1
Malania___TRINITY_DN11381_c0_g1_i1.p1
Ombrophytum___TRINITY_DN11919_c0_g1_i1.p1
Lophophytum-pyramidale___TRINITY_DN9241_c0_g1_i3.p1
Lophophytum-mirabile___TRINITY_DN183157_c0_g1_i1.p1_5prime_partial
Rhop___TRINITY_DN113650_c0_g1_i1.p1
Balfun___TRINITY_DN1294_c0_g2_i1.p1
Cucumis___Cucsa.142630.1IPACid_16963667
Quercus___Qurub.03G027500.1
F-Pvulgaris___Phvul.006G083700.1.p
F-Carietinum___Ca_08878
F-Lleucocephala___GDRZ01232348.1.p1
F-Akoa___GBYE01023901.1.p1
F-Prosopis___Palba_XP_028800436.1
F-Prosopis___Palba_XP_028752744.1
F-Acol___TRINITY_DN6922_c0_g1_i4.p1_complete
F-Acol___TRINITY_DN6922_c0_g1_i1.p1_complete
Eucalyptus___Eucgr.I00443.1.p
Eucalyptus___Eucgr.A02603.1.p
Vitis___VIT_205s0029g00710.1
Vitis___VIT_204s0069g01110.1
Hydrangea___Hyque.09G035300.1.p
Vaccinium___Vadar_g25926.t1
Solanum___Solyc02g081530.1.1
Olea___Oeu002676.1
Mimulus___Migut.I00362.1.p
Lactuca___Lsat_1_v5_gn_8_63660.1
Helianthus___HanXRQChr05g0144031
Daucus___DCAR_003386
Malania___TRINITY_DN11381_c0_g1_i1.p2
Kalanchoe___Kalax.0823s0009.1.p
Kalanchoe___Kalax.0690s0006.1.p
Beta___EL10Ac3g07268.1
Manihot___Manes.18G104900.1.p
Citrus___orange1.1g044902mIPACid_18127491
Arabidopsis-AT1G06440.1IPACid_19653310
Musa___GSMUA_Achr10P06910_001
Balfun___TRINITY_DN25611_c0_g1_i1.p1
Hydrangea___Hyque.10G036700.1.p
Lactuca___Lsat_1_v5_gn_7_61661.3
Lactuca___Lsat_1_v5_gn_7_61661.1
Lactuca___Lsat_1_v5_gn_1_36581.1
Helianthus___HanXRQChr01g0009551
Vaccinium___Vadar_g18535.t1
Solanum___Solyc10g077100.1.1
Olea___Oeu023006.2
Mimulus___Migut.J01609.1.p
Kalanchoe___Kalax.1162s0006.1.p
Kalanchoe___Kalax.0057s0031.1.p
Beta___EL10Ac4g08919.1
Ombrophytum___TRINITY_DN36073_c0_g1_i6.p1
Lophophytum-pyramidale___TRINITY_DN5132_c0_g1_i5.p1
Lophophytum-mirabile___TRINITY_DN26460_c0_g1_i1.p1_complete
Rhop___TRINITY_DN514072_c0_g1_i2.p1
Rhop___TRINITY_DN514072_c0_g1_i1.p1
Malania___TRINITY_DN79719_c0_g1_i1.p1
Vitis___VIT_208s0040g00250.1
Quercus___Qurub.M171200.1
Quercus___Qurub.M088100.1
Malus___MD07G1073900
Malus___MD02G1242700
Eucalyptus___Eucgr.K01687.1.p
Cucumis___Cucsa.337470.1IPACid_16978606
F-Pvulgaris___Phvul.001G116700.1.p
F-Carietinum___Ca_02698
F-Prosopis___Palba_XP_028775784.1
F-Prosopis___Palba_XP_028766799.1
F-Lleucocephala___GDRZ01147481.1.p1
F-Lleucocephala___GDRZ01044636.1.p1
F-Akoa___GBYE01024873.1.p1
F-Akoa___GBYE01053343.1.p1
F-Acol___TRINITY_DN11837_c0_g1_i9.p1_complete
F-Acol___TRINITY_DN11837_c0_g1_i6.p1_complete
F-Acol___TRINITY_DN11837_c0_g1_i6.p2_complete
F-Acol___TRINITY_DN11837_c0_g1_i5.p1_5prime_partial
Manihot___Manes.02G123200.4.p
Citrus___orange1.1g014563mIPACid_18123238
Balfun___TRINITY_DN25611_c0_g2_i1.p1
Arabidopsis-AT5G45790.2IPACid_19665979
Musa___GSMUA_Achr9P00600_001
Amborella___AmTr_v1.0_scaffold00055.5

# OG0003089

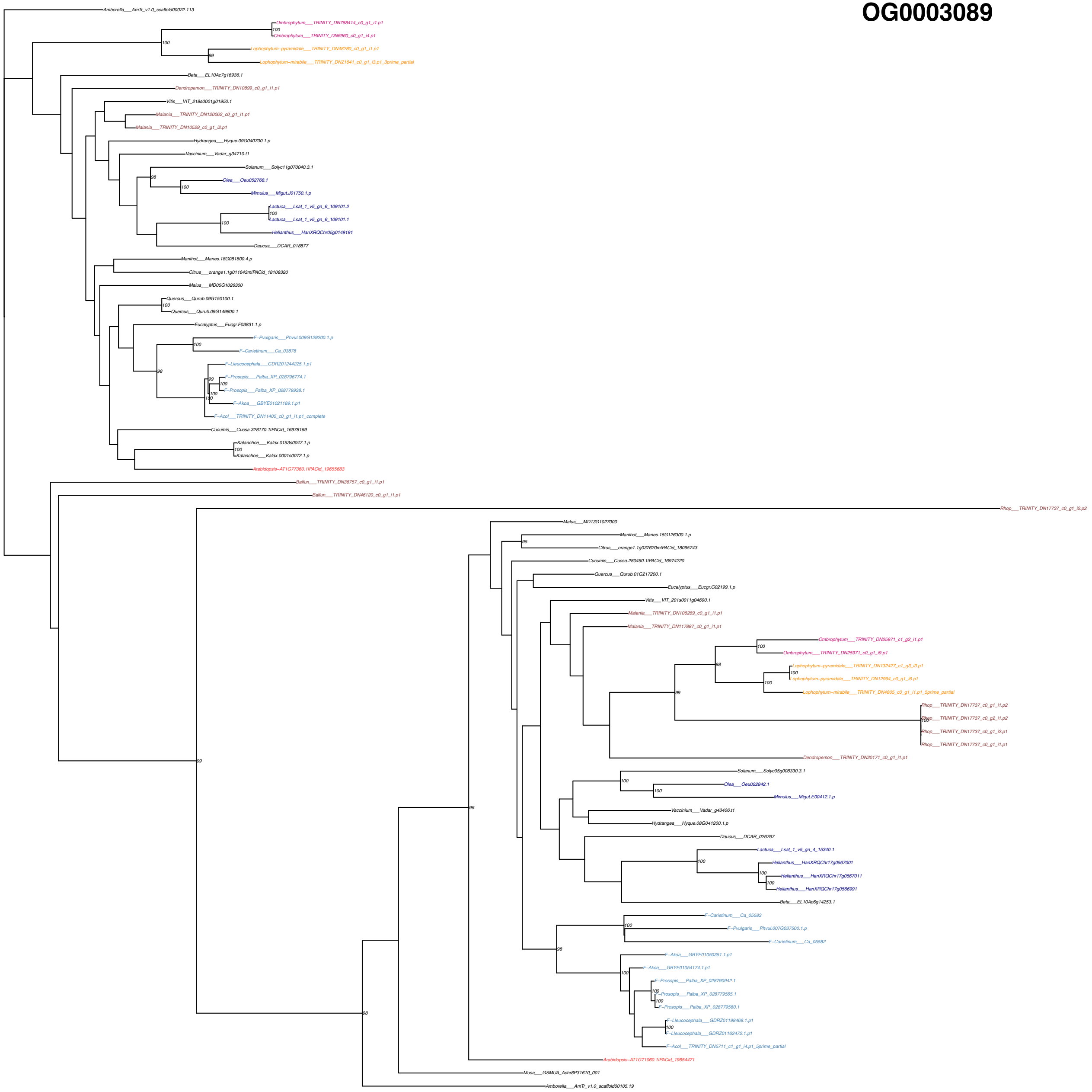

Amborella___AmTr_v1.0_scaffold00022.113
Ombrophytum___TRINITY_DN788414_c0_g1_i1.p1
Ombrophytum___TRINITY_DN6960_c0_g1_i4.p1
Lophophytum-pyramidale___TRINITY_DN48280_c0_g1_i1.p1
Lophophytum-mirabile___TRINITY_DN21641_c0_g1_i3.p1_3prime_partial
Beta___EL10Ac7g16936.1
Dendropemon___TRINITY_DN10899_c0_g1_i1.p1
Vitis___VIT_218s0001g01950.1
Malania___TRINITY_DN120062_c0_g1_i1.p1
Malania___TRINITY_DN10529_c0_g1_i2.p1
Hydrangea___Hyque.09G040700.1.p
Vaccinium___Vadar_g34710.t1
Solanum___Solyc11g070040.3.1
Olea___Oeu052768.1
Mimulus___Migut.J01750.1.p
Lactuca___Lsat_1_v5_gn_6_109101.2
Lactuca___Lsat_1_v5_gn_6_109101.1
Helianthus___HanXRQChr05g0149191
Daucus___DCAR_018877
Manihot___Manes.18G081800.4.p
Citrus___orange1.1g011643mIPACid_18108320
Malus___MD05G1026300
Quercus___Qurub.09G150100.1
Quercus___Qurub.09G149800.1
Eucalyptus___Eucgr.F03831.1.p
F-Pvulgaris___Phvul.009G129200.1.p
F-Carietinum___Ca_03878
F-Lleucocephala___GDRZ01244225.1.p1
F-Prosopis___Palba_XP_028796774.1
F-Prosopis___Palba_XP_028779938.1
F-Akoa___GBYE01021189.1.p1
F-Acol___TRINITY_DN11405_c0_g1_i1.p1_complete
Cucumis___Cucsa.328170.1IPACid_16978169
Kalanchoe___Kalax.0153s0047.1.p
Kalanchoe___Kalax.0001s0072.1.p
Arabidopsis-AT1G77360.1IPACid_19655683
Balfun___TRINITY_DN36757_c0_g1_i1.p1
Balfun___TRINITY_DN46120_c0_g1_i1.p1
Rhop___TRINITY_DN17737_c0_g1_i2.p2
Malus___MD13G1027000
Manihot___Manes.15G126300.1.p
Citrus___orange1.1g037620mIPACid_18095743
Cucumis___Cucsa.280460.1IPACid_16974220
Quercus___Qurub.01G217200.1
Eucalyptus___Eucgr.G02199.1.p
Vitis___VIT_201s0011g04690.1
Malania___TRINITY_DN106269_c0_g1_i1.p1
Malania___TRINITY_DN117887_c0_g1_i1.p1
Ombrophytum___TRINITY_DN25971_c1_g2_i1.p1
Ombrophytum___TRINITY_DN25971_c0_g1_i9.p1
Lophophytum-pyramidale___TRINITY_DN132427_c1_g3_i3.p1
Lophophytum-pyramidale___TRINITY_DN12994_c0_g1_i6.p1
Lophophytum-mirabile___TRINITY_DN4805_c0_g1_i1.p1_5prime_partial
Rhop___TRINITY_DN17737_c0_g1_i1.p2
Rhop___TRINITY_DN17737_c0_g2_i1.p2
Rhop___TRINITY_DN17737_c0_g1_i2.p1
Rhop___TRINITY_DN17737_c0_g1_i1.p1
Dendropemon___TRINITY_DN20171_c0_g1_i1.p1
Solanum___Solyc05g008330.3.1
Olea___Oeu022842.1
Mimulus___Migut.E00412.1.p
Vaccinium___Vadar_g43406.t1
Hydrangea___Hyque.08G041200.1.p
Daucus___DCAR_026767
Lactuca___Lsat_1_v5_gn_4_15340.1
Helianthus___HanXRQChr17g0567001
Helianthus___HanXRQChr17g0567011
Helianthus___HanXRQChr17g0566991
Beta___EL10Ac6g14253.1
F-Carietinum___Ca_05583
F-Pvulgaris___Phvul.007G037500.1.p
F-Carietinum___Ca_05582
F-Akoa___GBYE01050351.1.p1
F-Akoa___GBYE01054174.1.p1
F-Prosopis___Palba_XP_028790942.1
F-Prosopis___Palba_XP_028779565.1
F-Prosopis___Palba_XP_028779560.1
F-Lleucocephala___GDRZ01198468.1.p1
F-Lleucocephala___GDRZ01162472.1.p1
F-Acol___TRINITY_DN5711_c1_g1_i4.p1_5prime_partial
Arabidopsis-AT1G71060.1IPACid_19654471
Musa___GSMUA_Achr8P31610_001
Amborella___AmTr_v1.0_scaffold00105.19

# OG0003131

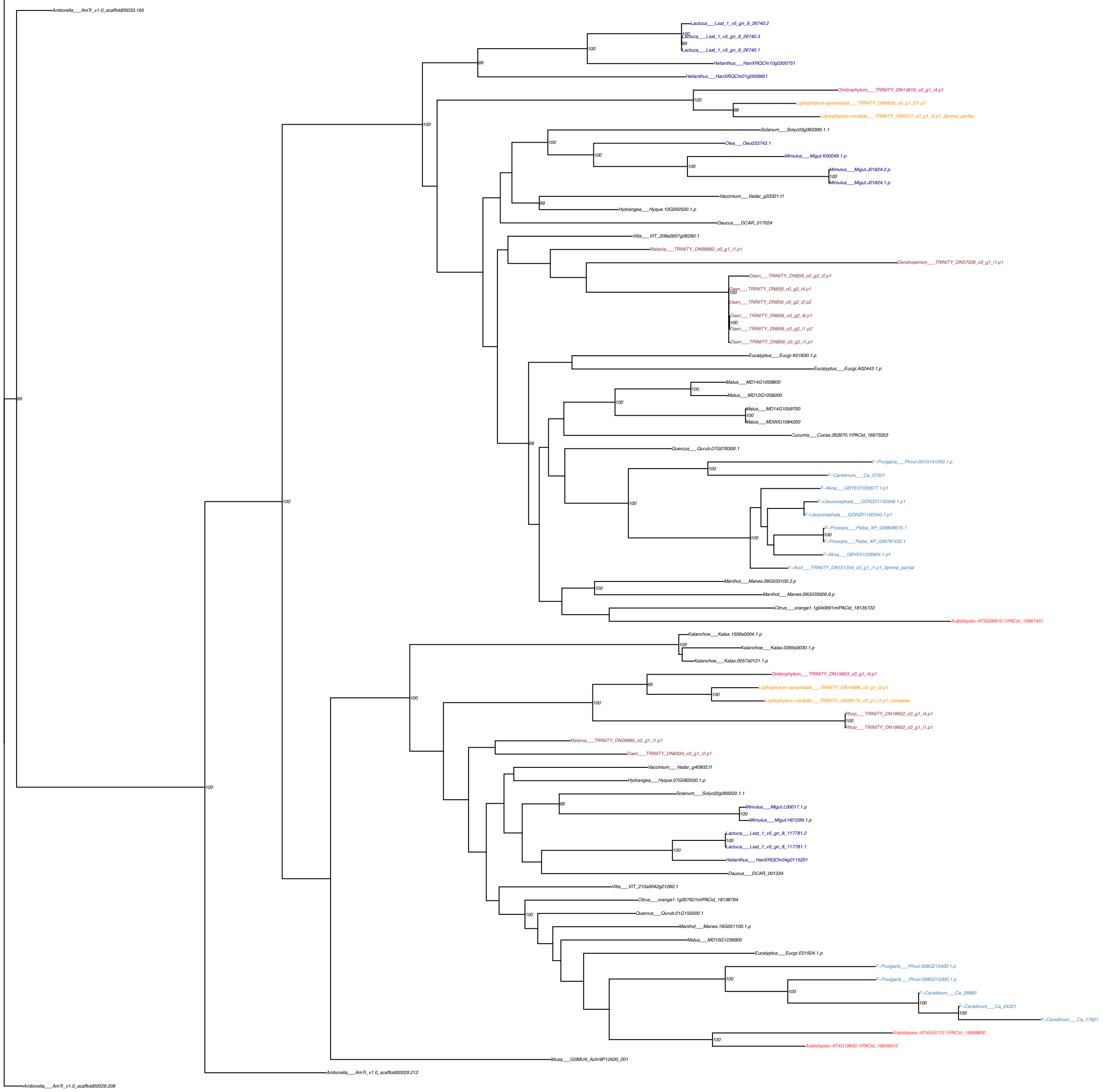

Amborella___AmTr_v1.0_scaffold00027.143
Amborella___AmTr_v1.0_scaffold00033.165
Lactuca___Lsat_1_v5_gn_9_26740.2
Lactuca___Lsat_1_v5_gn_9_26740.3
Lactuca___Lsat_1_v5_gn_9_26740.1
Helianthus___HanXRQChr10g0300751
Helianthus___HanXRQChr01g0009651
Ombrophytum___TRINITY_DN14819_c0_g1_i4.p1
Lophophytum-pyramidale___TRINITY_DN6858_c0_g1_i21.p1
Lophophytum-mirabile___TRINITY_DN5517_c0_g1_i5.p1_3prime_partial
Solanum___Solyc03g063390.1.1
Olea___Oeu033743.1
Mimulus___Migut.K00049.1.p
Mimulus___Migut.J01824.2.p
Mimulus___Migut.J01824.1.p
Vaccinium___Vadar_g33321.t1
Hydrangea___Hyque.10G002500.1.p
Daucus___DCAR_017024
Vitis___VIT_208s0007g06290.1
Malania___TRINITY_DN98892_c0_g1_i1.p1
Dendropemon___TRINITY_DN37026_c0_g1_i1.p1
Daen___TRINITY_DN859_c0_g2_i2.p1
Daen___TRINITY_DN859_c0_g2_i4.p1
Daen___TRINITY_DN859_c0_g2_i2.p2
Daen___TRINITY_DN859_c0_g2_i6.p1
Daen___TRINITY_DN859_c0_g2_i1.p2
Daen___TRINITY_DN859_c0_g2_i1.p1
Eucalyptus___Eucgr.K01830.1.p
Eucalyptus___Eucgr.A02443.1.p
Malus___MD14G1059800
Malus___MD12G1058200
Malus___MD14G1059700
Malus___MD00G1084200
Cucumis___Cucsa.263970.1IPACid_16973253
Quercus___Qurub.07G076300.1
F-Pvulgaris___Phvul.001G141200.1.p
F-Carietinum___Ca_07301
F-Akoa___GBYE01030677.1.p1
F-Lleucocephala___GDRZ01132949.1.p1
F-Lleucocephala___GDRZ01100340.1.p1
F-Prosopis___Palba_XP_028808675.1
F-Prosopis___Palba_XP_028787432.1
F-Akoa___GBYE01028904.1.p1
F-Acol___TRINITY_DN151354_c0_g1_i1.p1_3prime_partial
Manihot___Manes.09G033100.2.p
Manihot___Manes.09G033000.9.p
Citrus___orange1.1g040691mIPACid_18135722
Arabidopsis-AT5G06810.1IPACid_19667451
Kalanchoe___Kalax.1556s0004.1.p
Kalanchoe___Kalax.0395s0030.1.p
Kalanchoe___Kalax.0057s0121.1.p
Ombrophytum___TRINITY_DN15663_c0_g1_i4.p1
Lophophytum-pyramidale___TRINITY_DN14698_c0_g1_i2.p1
Lophophytum-mirabile___TRINITY_DN39175_c0_g1_i1.p1_complete
Rhop___TRINITY_DN18602_c0_g1_i4.p1
Rhop___TRINITY_DN18602_c0_g1_i1.p1
Malania___TRINITY_DN28885_c0_g1_i1.p1
Daen___TRINITY_DN8334_c0_g1_i2.p1
Vaccinium___Vadar_g40902.t1
Hydrangea___Hyque.07G083500.1.p
Solanum___Solyc02g069320.1.1
Mimulus___Migut.L00017.1.p
Mimulus___Migut.H01299.1.p
Lactuca___Lsat_1_v5_gn_8_117781.2
Lactuca___Lsat_1_v5_gn_8_117781.1
Helianthus___HanXRQChr04g0115201
Daucus___DCAR_001334
Vitis___VIT_210s0042g01260.1
Citrus___orange1.1g007621mIPACid_18138794
Quercus___Qurub.01G155000.1
Manihot___Manes.16G051100.1.p
Malus___MD10G1236900
Eucalyptus___Eucgr.E01924.1.p
F-Pvulgaris___Phvul.008G215400.1.p
F-Pvulgaris___Phvul.008G215300.1.p
F-Carietinum___Ca_26683
F-Carietinum___Ca_24321
F-Carietinum___Ca_17831
Arabidopsis-AT5G45113.1IPACid_19668800
Arabidopsis-AT4G19650.1IPACid_19646010
Musa___GSMUA_Achr9P12420_001
Amborella___AmTr_v1.0_scaffold00029.212
Amborella___AmTr_v1.0_scaffold00029.208

OG0003428

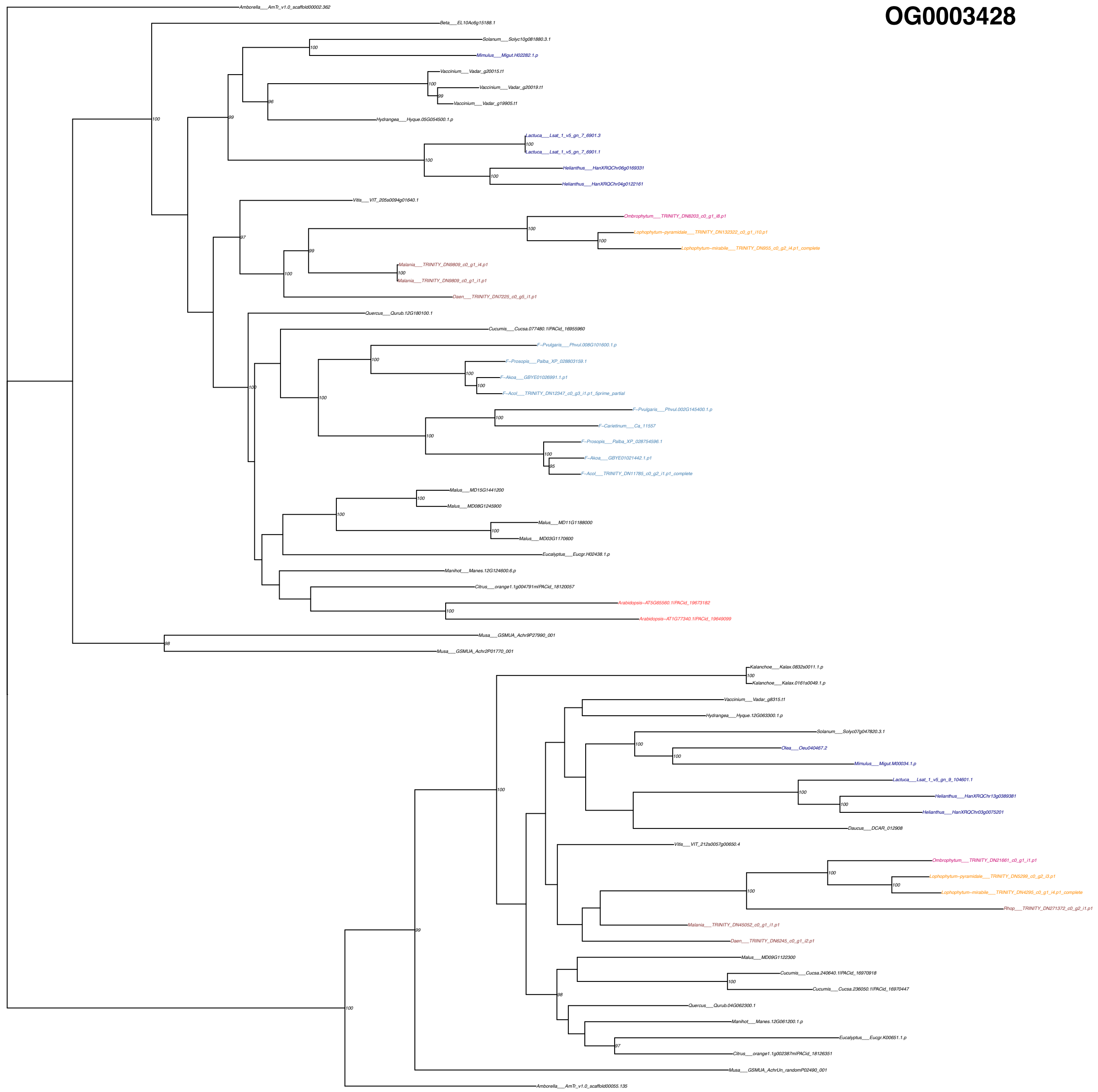

# OG0003768

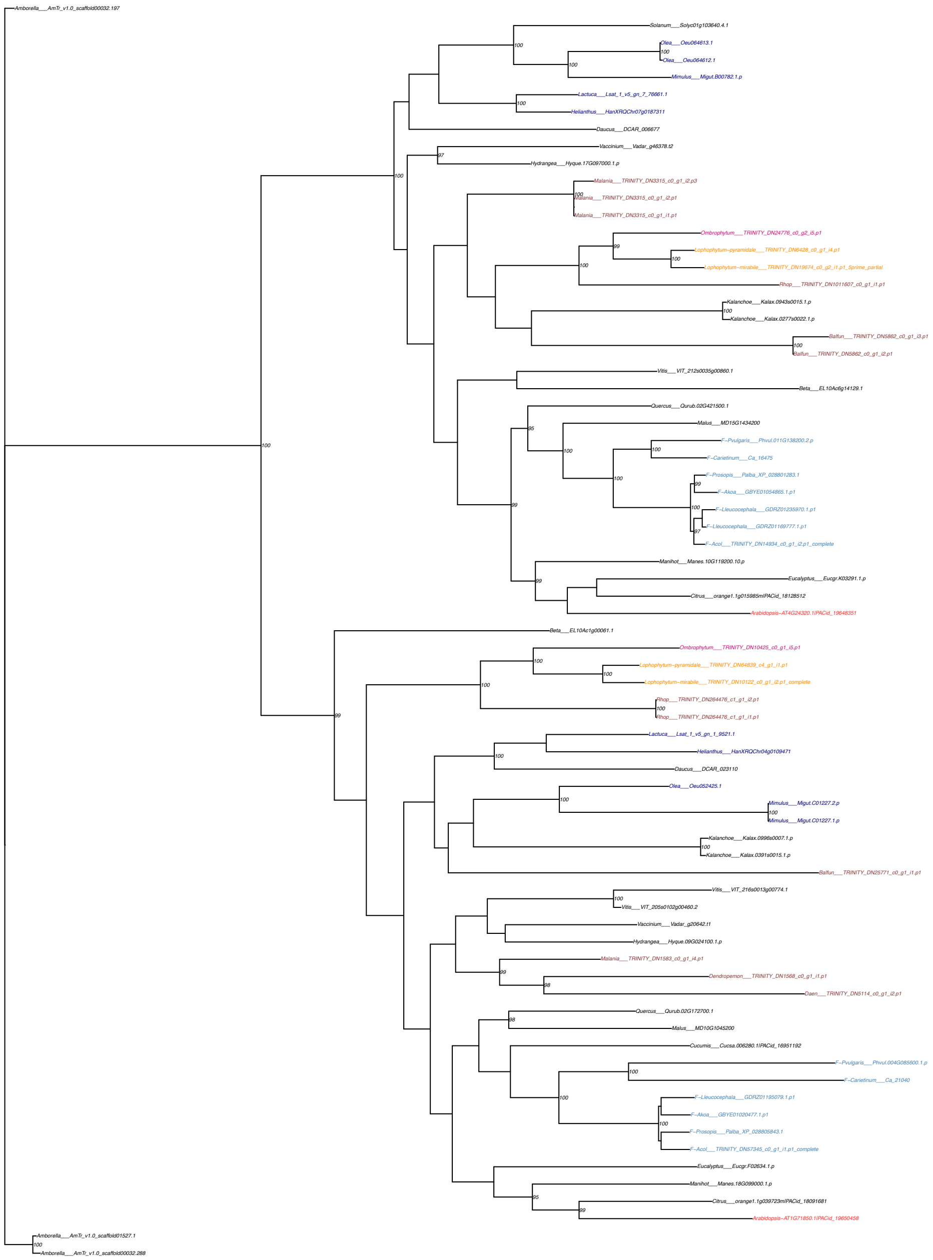

Amborella___AmTr_v1.0_scaffold00032.197
Solanum___Solyc01g103640.4.1
100
Olea___Oeu064613.1
100
Olea___Oeu064612.1
100
Mimulus___Migut.B00782.1.p
Lactuca___Lsat_1_v5_gn_7_76661.1
100
Helianthus___HanXRQChr07g0187311
Daucus___DCAR_006677
Vaccinium___Vadar_g46378.t2
97
Hydrangea___Hyque.17G097000.1.p
100
Malania___TRINITY_DN3315_c0_g1_i2.p3
100
Malania___TRINITY_DN3315_c0_g1_i2.p1
Malania___TRINITY_DN3315_c0_g1_i1.p1
Ombrophytum___TRINITY_DN24776_c0_g2_i5.p1
99
Lophophytum-pyramidale___TRINITY_DN6428_c0_g1_i4.p1
100
Lophophytum-mirabile___TRINITY_DN19674_c0_g2_i1.p1_5prime_partial
100
Rhop___TRINITY_DN1011607_c0_g1_i1.p1
Kalanchoe___Kalax.0943s0015.1.p
100
Kalanchoe___Kalax.0277s0022.1.p
Balfun___TRINITY_DN5862_c0_g1_i3.p1
100
Balfun___TRINITY_DN5862_c0_g1_i2.p1
Vitis___VIT_212s0035g00860.1
Beta___EL10Ac6g14129.1
100
Quercus___Qurub.02G421500.1
95
Malus___MD15G1434200
100
F-Pvulgaris___Phvul.011G138200.2.p
100
F-Carietinum___Ca_16475
100
F-Prosopis___Palba_XP_028801283.1
99
F-Akoa___GBYE01054865.1.p1
100
F-Lleucocephala___GDRZ01235970.1.p1
F-Lleucocephala___GDRZ01169777.1.p1
97
F-Acol___TRINITY_DN14934_c0_g1_i2.p1_complete
99
Manihot___Manes.10G119200.10.p
99
Eucalyptus___Eucgr.K03291.1.p
Citrus___orange1.1g015985mIPACid_18128512
Arabidopsis-AT4G24320.1IPACid_19648351
Beta___EL10Ac1g00061.1
Ombrophytum___TRINITY_DN10425_c0_g1_i5.p1
100
Lophophytum-pyramidale___TRINITY_DN64839_c4_g1_i1.p1
100
Lophophytum-mirabile___TRINITY_DN10122_c0_g1_i2.p1_complete
100
Rhop___TRINITY_DN264476_c1_g1_i2.p1
100
Rhop___TRINITY_DN264476_c1_g1_i1.p1
99
Lactuca___Lsat_1_v5_gn_1_9521.1
100
Helianthus___HanXRQChr04g0109471
Daucus___DCAR_023110
Olea___Oeu052425.1
100
Mimulus___Migut.C01227.2.p
100
Mimulus___Migut.C01227.1.p
Kalanchoe___Kalax.0996s0007.1.p
100
Kalanchoe___Kalax.0391s0015.1.p
Balfun___TRINITY_DN25771_c0_g1_i1.p1
Vitis___VIT_216s0013g00774.1
100
Vitis___VIT_205s0102g00460.2
Vaccinium___Vadar_g20642.t1
Hydrangea___Hyque.09G024100.1.p
Malania___TRINITY_DN1583_c0_g1_i4.p1
99
Dendropemon___TRINITY_DN1568_c0_g1_i1.p1
98
Daen___TRINITY_DN5114_c0_g1_i2.p1
Quercus___Qurub.02G172700.1
98
Malus___MD10G1045200
Cucumis___Cucsa.006280.1IPACid_16951192
F-Pvulgaris___Phvul.004G085600.1.p
100
F-Carietinum___Ca_21040
100
F-Lleucocephala___GDRZ01195079.1.p1
F-Akoa___GBYE01020477.1.p1
100
F-Prosopis___Palba_XP_028805843.1
F-Acol___TRINITY_DN57345_c0_g1_i1.p1_complete
Eucalyptus___Eucgr.F02634.1.p
Manihot___Manes.18G099000.1.p
95
Citrus___orange1.1g039723mIPACid_18091681
99
Arabidopsis-AT1G71850.1IPACid_19650458
Amborella___AmTr_v1.0_scaffold01527.1
100
Amborella___AmTr_v1.0_scaffold00032.288

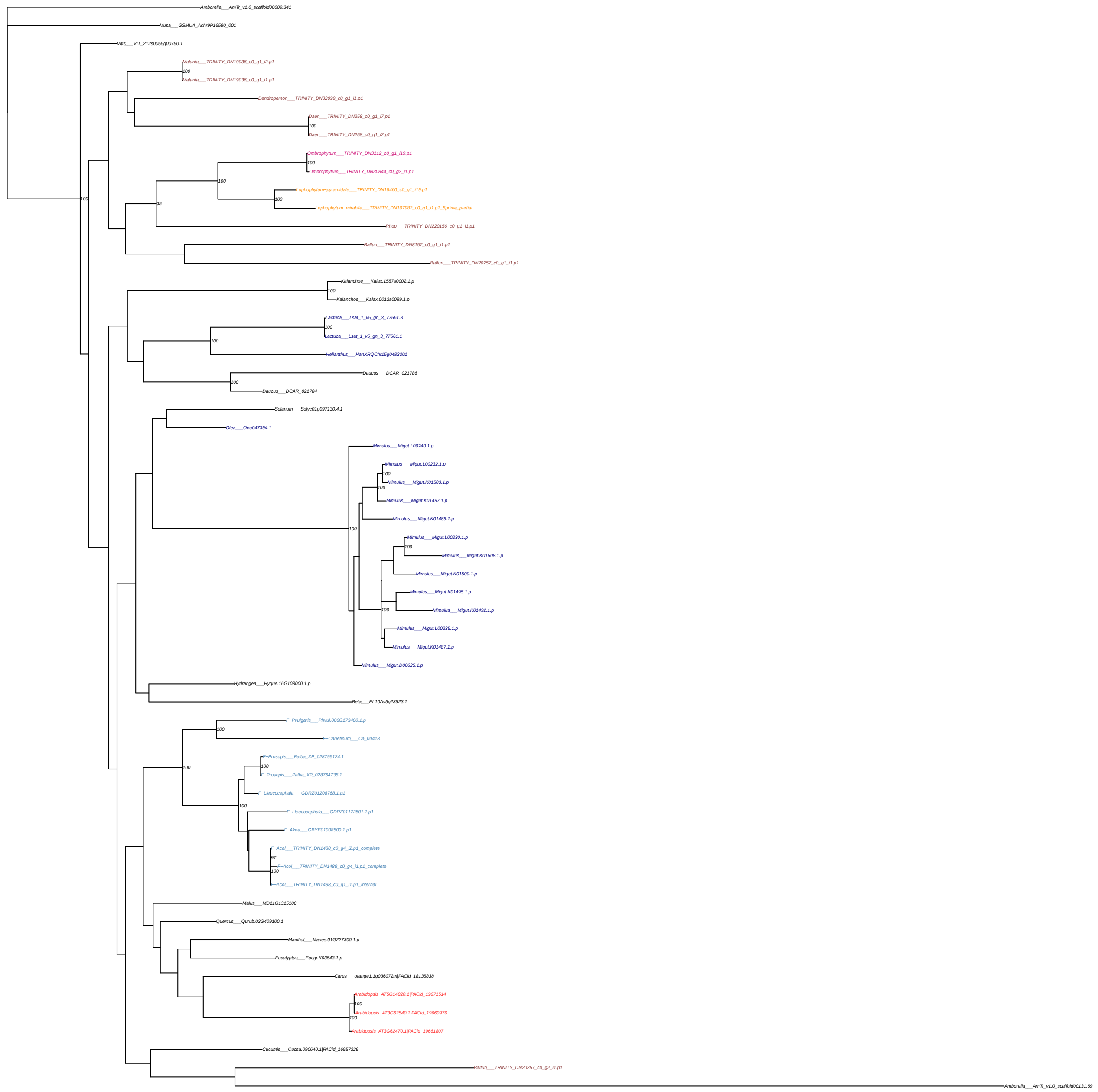
Amborella___AmTr_v1.0_scaffold00009.341
Musa___GSMUA_Achr9P16580_001
Vitis___VIT_212s0055g00750.1
Malania___TRINITY_DN19036_c0_g1_i2.p1
100
Malania___TRINITY_DN19036_c0_g1_i1.p1
Dendropemon___TRINITY_DN32099_c0_g1_i1.p1
Daen___TRINITY_DN258_c0_g1_i7.p1
100
Daen___TRINITY_DN258_c0_g1_i2.p1
Ombrophytum___TRINITY_DN3112_c0_g1_i19.p1
100
Ombrophytum___TRINITY_DN30844_c0_g2_i1.p1
100
Lophophytum~pyramidale___TRINITY_DN18460_c0_g1_i19.p1
100
Lophophytum~mirabile___TRINITY_DN107982_c0_g1_i1.p1_5prime_partial
98
Rhop___TRINITY_DN220156_c0_g1_i1.p1
100
Balfun___TRINITY_DN8157_c0_g1_i1.p1
Balfun___TRINITY_DN20257_c0_g1_i1.p1
Kalanchoe___Kalax.1587s0002.1.p
100
Kalanchoe___Kalax.0012s0089.1.p
Lactuca___Lsat_1_v5_gn_3_77561.3
100
Lactuca___Lsat_1_v5_gn_3_77561.1
100
Helianthus___HanXRQChr15g0482301
Daucus___DCAR_021786
100
Daucus___DCAR_021784
Solanum___Solyc01g097130.4.1
Olea___Oeu047394.1
Mimulus___Migut.L00240.1.p
Mimulus___Migut.L00232.1.p
100
Mimulus___Migut.K01503.1.p
100
Mimulus___Migut.K01497.1.p
Mimulus___Migut.K01489.1.p
100
Mimulus___Migut.L00230.1.p
100
Mimulus___Migut.K01508.1.p
Mimulus___Migut.K01500.1.p
Mimulus___Migut.K01495.1.p
100
Mimulus___Migut.K01492.1.p
Mimulus___Migut.L00235.1.p
Mimulus___Migut.K01487.1.p
Mimulus___Migut.D00625.1.p
Hydrangea___Hyque.16G108000.1.p
Beta___EL10As5g23523.1
F~Pvulgaris___Phvul.006G173400.1.p
100
F~Carietinum___Ca_00418
100
F~Prosopis___Palba_XP_028795124.1
100
F~Prosopis___Palba_XP_028764735.1
F~Lleucocephala___GDRZ01208768.1.p1
100
F~Lleucocephala___GDRZ01172501.1.p1
F~Akoa___GBYE01008500.1.p1
F~Acol___TRINITY_DN1488_c0_g4_i2.p1_complete
97
F~Acol___TRINITY_DN1488_c0_g4_i1.p1_complete
100
F~Acol___TRINITY_DN1488_c0_g1_i1.p1_internal
Malus___MD11G1315100
Quercus___Qurub.02G409100.1
Manihot___Manes.01G227300.1.p
Eucalyptus___Eucgr.K03543.1.p
Citrus___orange1.1g036072m|PACid_18135838
Arabidopsis~AT5G14820.1|PACid_19671514
100
Arabidopsis~AT3G62540.1|PACid_19660976
100
Arabidopsis~AT3G62470.1|PACid_19661807
Cucumis___Cucsa.090640.1|PACid_16957329
Balfun___TRINITY_DN20257_c0_g2_i1.p1
Amborella___AmTr_v1.0_scaffold00131.69

# OG0004376

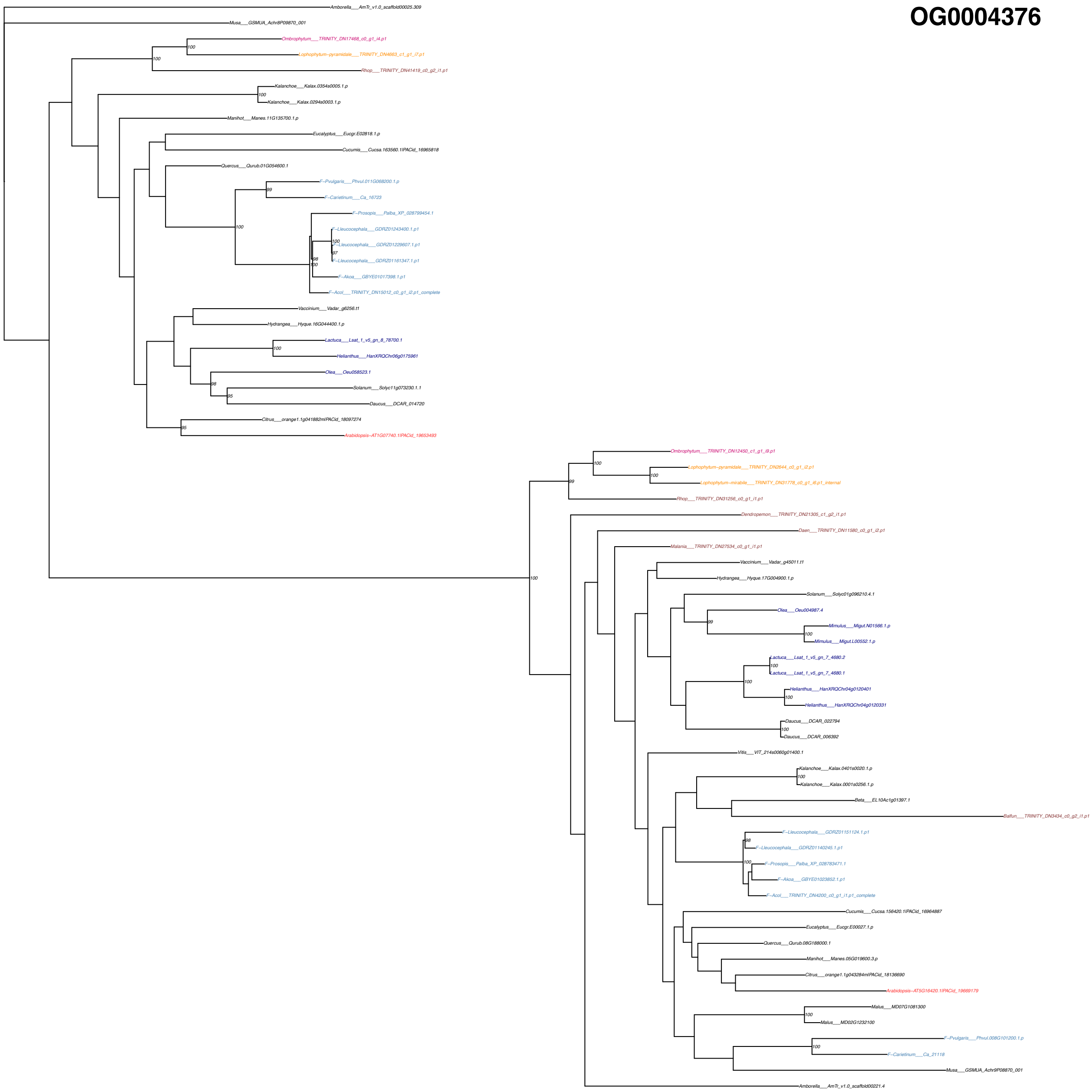
Amborella___AmTr_v1.0_scaffold00025.309
Musa___GSMUA_Achr8P09870_001
Ombrophytum___TRINITY_DN17468_c0_g1_i4.p1
Lophophytum-pyramidale___TRINITY_DN4663_c1_g1_i7.p1
Rhop___TRINITY_DN41419_c0_g2_i1.p1
Kalanchoe___Kalax.0354s0005.1.p
Kalanchoe___Kalax.0294s0003.1.p
Manihot___Manes.11G135700.1.p
Eucalyptus___Eucgr.E02818.1.p
Cucumis___Cucsa.163560.1PACid_16965818
Quercus___Qurub.01G054600.1
F-Pvulgaris___Phvul.011G068200.1.p
F-Carietinum___Ca_16723
F-Prosopis___Palba_XP_028799454.1
F-Lleucocephala___GDRZ01243400.1.p1
F-Lleucocephala___GDRZ01229607.1.p1
F-Lleucocephala___GDRZ01161347.1.p1
F-Akoa___GBYE01017398.1.p1
F-Acol___TRINITY_DN15012_c0_g1_i2.p1_complete
Vaccinium___Vadar_g6256.t1
Hydrangea___Hyque.16G044400.1.p
Lactuca___Lsat_1_v5_gn_8_78700.1
Helianthus___HanXRQChr06g0175961
Olea___Oeu058523.1
Solanum___Solyc11g073230.1.1
Daucus___DCAR_014720
Citrus___orange1.1g041882mPACid_18097274
Arabidopsis-AT1G07740.1PACid_19653493
Ombrophytum___TRINITY_DN12450_c1_g1_i9.p1
Lophophytum-pyramidale___TRINITY_DN2644_c0_g1_i2.p1
Lophophytum-mirabile___TRINITY_DN31778_c0_g1_i6.p1_internal
Rhop___TRINITY_DN31256_c0_g1_i1.p1
Dendropemon___TRINITY_DN21305_c1_g2_i1.p1
Daen___TRINITY_DN11580_c0_g1_i2.p1
Malania___TRINITY_DN27534_c0_g1_i1.p1
Vaccinium___Vadar_g45011.t1
Hydrangea___Hyque.17G004900.1.p
Solanum___Solyc01g096210.4.1
Olea___Oeu004987.4
Mimulus___Migut.N01566.1.p
Mimulus___Migut.L00552.1.p
Lactuca___Lsat_1_v5_gn_7_4680.2
Lactuca___Lsat_1_v5_gn_7_4680.1
Helianthus___HanXRQChr04g0120401
Helianthus___HanXRQChr04g0120331
Daucus___DCAR_022794
Daucus___DCAR_006392
Vitis___VIT_214s0060g01400.1
Kalanchoe___Kalax.0401s0020.1.p
Kalanchoe___Kalax.0001s0256.1.p
Beta___EL10Ac1g01397.1
Balfun___TRINITY_DN3434_c0_g2_i1.p1
F-Lleucocephala___GDRZ01151124.1.p1
F-Lleucocephala___GDRZ01140245.1.p1
F-Prosopis___Palba_XP_028783471.1
F-Akoa___GBYE01023852.1.p1
F-Acol___TRINITY_DN4200_c0_g1_i1.p1_complete
Cucumis___Cucsa.156420.1PACid_16964887
Eucalyptus___Eucgr.E00027.1.p
Quercus___Qurub.08G188000.1
Manihot___Manes.05G019600.3.p
Citrus___orange1.1g043284mPACid_18136690
Arabidopsis-AT5G16420.1PACid_19669179
Malus___MD07G1081300
Malus___MD02G1232100
F-Pvulgaris___Phvul.008G101200.1.p
F-Carietinum___Ca_21118
Musa___GSMUA_Achr9P08870_001
Amborella___AmTr_v1.0_scaffold00221.4

# OG0004578

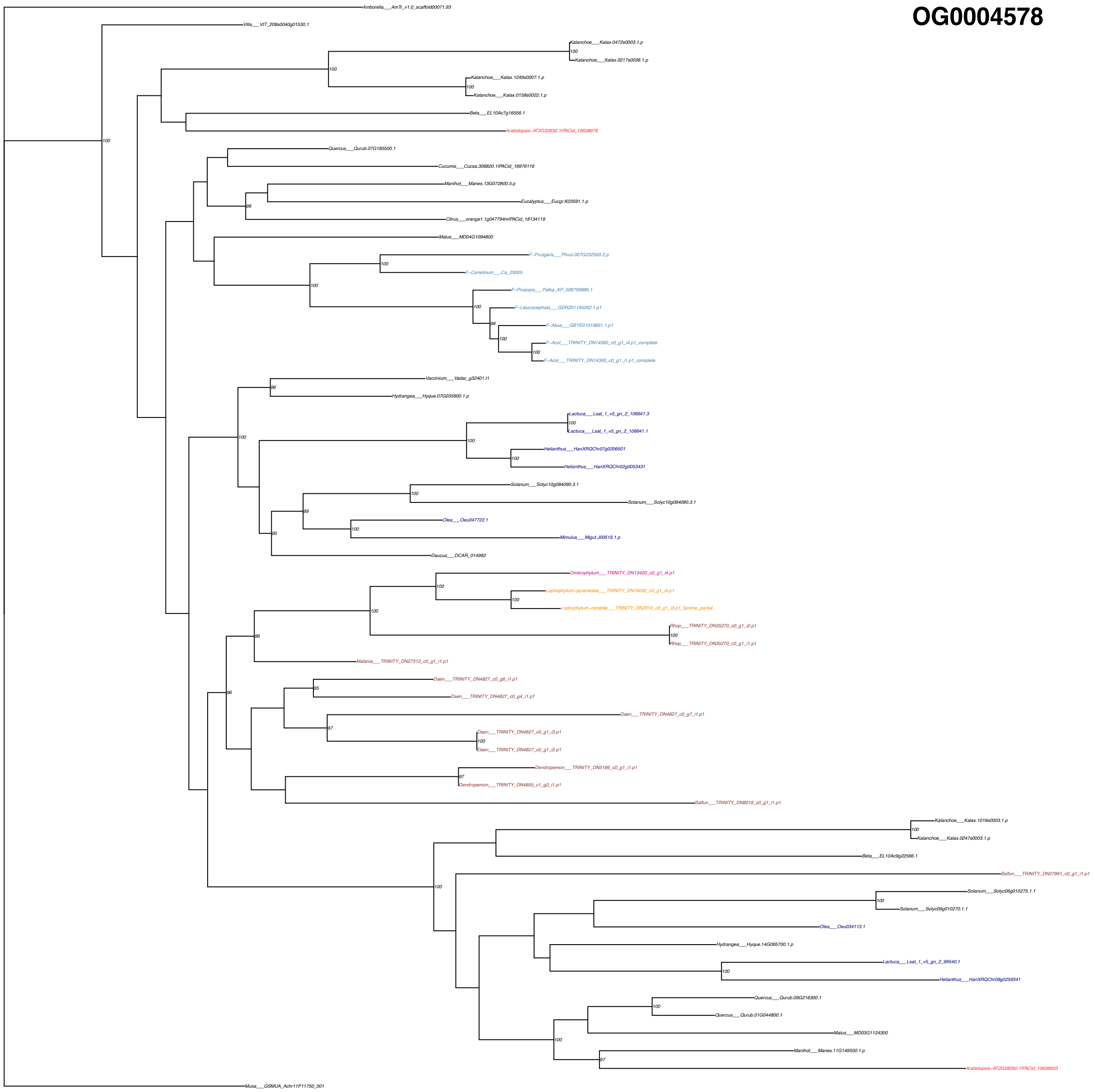

Amborella___AmTr_v1.0_scaffold00071.93
Vitis___VIT_208s0040g01530.1
Kalanchoe___Kalax.0472s0003.1.p
Kalanchoe___Kalax.0217s0038.1.p
Kalanchoe___Kalax.1249s0007.1.p
Kalanchoe___Kalax.0158s0022.1.p
Beta___EL10Ac7g16556.1
Arabidopsis~AT2G32630.1lPACid_19638978
Quercus___Qurub.07G185500.1
Cucumis___Cucsa.306820.1lPACid_16976116
Manihot___Manes.13G072800.5.p
Eucalyptus___Eucgr.K03591.1.p
Citrus___orange1.1g047794mlPACid_18134119
Malus___MD04G1094800
F~Pvulgaris___Phvul.007G232500.2.p
F~Carietinum___Ca_25005
F~Prosopis___Palba_XP_028799986.1
F~Lleucocephala___GDRZ01195262.1.p1
F~Akoa___GBYE01019891.1.p1
F~Acol___TRINITY_DN14360_c0_g1_i4.p1_complete
F~Acol___TRINITY_DN14360_c0_g1_i1.p1_complete
Vaccinium___Vadar_g32401.t1
Hydrangea___Hyque.07G035900.1.p
Lactuca___Lsat_1_v5_gn_2_106641.3
Lactuca___Lsat_1_v5_gn_2_106641.1
Helianthus___HanXRQChr07g0206501
Helianthus___HanXRQChr02g0053431
Solanum___Solyc10g084090.3.1
Solanum___Solyc10g084080.3.1
Olea___Oeu047722.1
Mimulus___Migut.J00519.1.p
Daucus___DCAR_014992
Ombrophytum___TRINITY_DN13420_c0_g1_i4.p1
Lophophytum~pyramidale___TRINITY_DN19035_c0_g1_i4.p1
Lophophytum~mirabile___TRINITY_DN2010_c0_g1_i3.p1_5prime_partial
Rhop___TRINITY_DN35270_c0_g1_i2.p1
Rhop___TRINITY_DN35270_c0_g1_i1.p1
Malania___TRINITY_DN27312_c0_g1_i1.p1
Daen___TRINITY_DN4827_c0_g6_i1.p1
Daen___TRINITY_DN4827_c0_g4_i1.p1
Daen___TRINITY_DN4827_c0_g7_i1.p1
Daen___TRINITY_DN4827_c0_g1_i3.p1
Daen___TRINITY_DN4827_c0_g1_i2.p1
Dendropemon___TRINITY_DN5199_c0_g1_i1.p1
Dendropemon___TRINITY_DN4855_c1_g2_i1.p1
Balfun___TRINITY_DN8019_c0_g1_i1.p1
Kalanchoe___Kalax.1019s0003.1.p
Kalanchoe___Kalax.0247s0003.1.p
Beta___EL10Ac9g22566.1
Balfun___TRINITY_DN27961_c0_g1_i1.p1
Solanum___Solyc06g010275.1.1
Solanum___Solyc06g010270.1.1
Olea___Oeu034113.1
Hydrangea___Hyque.14G065700.1.p
Lactuca___Lsat_1_v5_gn_2_99540.1
Helianthus___HanXRQChr09g0259341
Quercus___Qurub.09G216300.1
Quercus___Qurub.01G044800.1
Malus___MD03G1124300
Manihot___Manes.11G149500.1.p
Arabidopsis~AT2G28050.1lPACid_19638603
Musa___GSMUA_Achr11P11750_001

# OG0005144

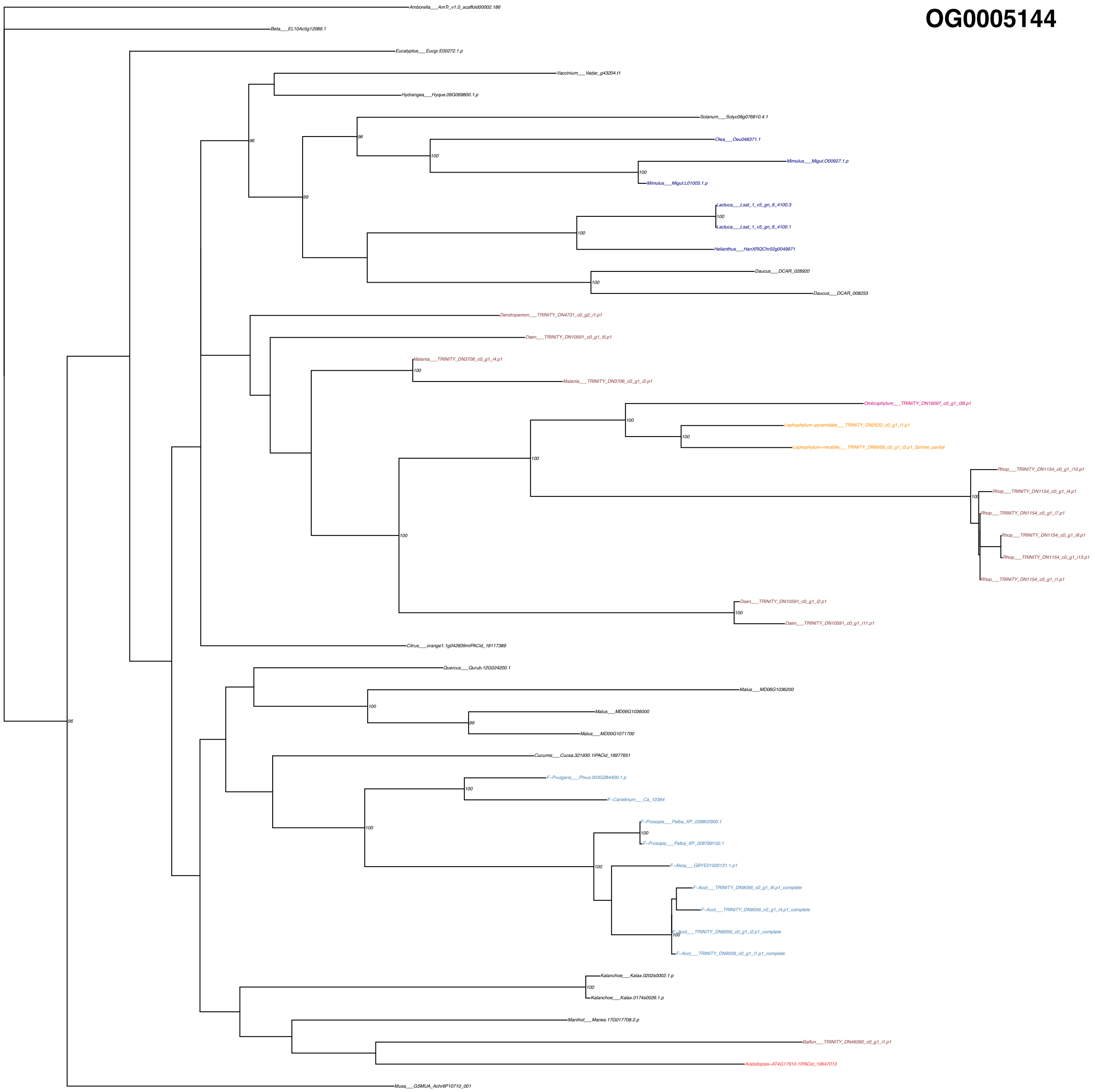

Amborella___AmTr_v1.0_scaffold00002.186
Beta___EL10Ac5g12089.1
Eucalyptus___Eucgr.E00272.1.p
Vaccinium___Vadar_g43204.t1
Hydrangea___Hyque.09G069800.1.p
Solanum___Solyc08g076810.4.1
Olea___Oeu048371.1
Mimulus___Migut.O00927.1.p
Mimulus___Migut.L01003.1.p
Lactuca___Lsat_1_v5_gn_6_4100.3
Lactuca___Lsat_1_v5_gn_6_4100.1
Helianthus___HanXRQChr02g0049971
Daucus___DCAR_028920
Daucus___DCAR_008253
Dendropemon___TRINITY_DN4731_c0_g2_i1.p1
Daen___TRINITY_DN10591_c0_g1_i5.p1
Malania___TRINITY_DN3706_c0_g1_i4.p1
Malania___TRINITY_DN3706_c0_g1_i2.p1
Ombrophytum___TRINITY_DN16097_c0_g1_i38.p1
Lophophytum-pyramidale___TRINITY_DN2532_c0_g1_i1.p1
Lophophytum-mirabile___TRINITY_DN9569_c0_g1_i3.p1_3prime_partial
Rhop___TRINITY_DN1154_c0_g1_i10.p1
Rhop___TRINITY_DN1154_c0_g1_i4.p1
Rhop___TRINITY_DN1154_c0_g1_i7.p1
Rhop___TRINITY_DN1154_c0_g1_i9.p1
Rhop___TRINITY_DN1154_c0_g1_i13.p1
Rhop___TRINITY_DN1154_c0_g1_i1.p1
Daen___TRINITY_DN10591_c0_g1_i2.p1
Daen___TRINITY_DN10591_c0_g1_i11.p1
Citrus___orange1.1g042839mIPACid_18117389
Quercus___Qurub.12G024200.1
Malus___MD06G1036200
Malus___MD06G1036000
Malus___MD00G1071700
Cucumis___Cucsa.321930.1IPACid_16977651
F-Pvulgaris___Phvul.003G284400.1.p
F-Carietinum___Ca_12364
F-Prosopis___Palba_XP_028802900.1
F-Prosopis___Palba_XP_028789150.1
F-Akoa___GBYE01020131.1.p1
F-Acol___TRINITY_DN9056_c0_g1_i6.p1_complete
F-Acol___TRINITY_DN9056_c0_g1_i4.p1_complete
F-Acol___TRINITY_DN9056_c0_g1_i2.p1_complete
F-Acol___TRINITY_DN9056_c0_g1_i1.p1_complete
Kalanchoe___Kalax.0202s0002.1.p
Kalanchoe___Kalax.0174s0026.1.p
Manihot___Manes.17G017708.2.p
Balfun___TRINITY_DN46390_c0_g1_i1.p1
Arabidopsis-AT4G17910.1IPACid_19647013
Musa___GSMUA_Achr6P10710_001
96
96
99
100
100
100
100
100
100
100
100
100
100
100
100
100
99
100
100
100
100
100
100
96

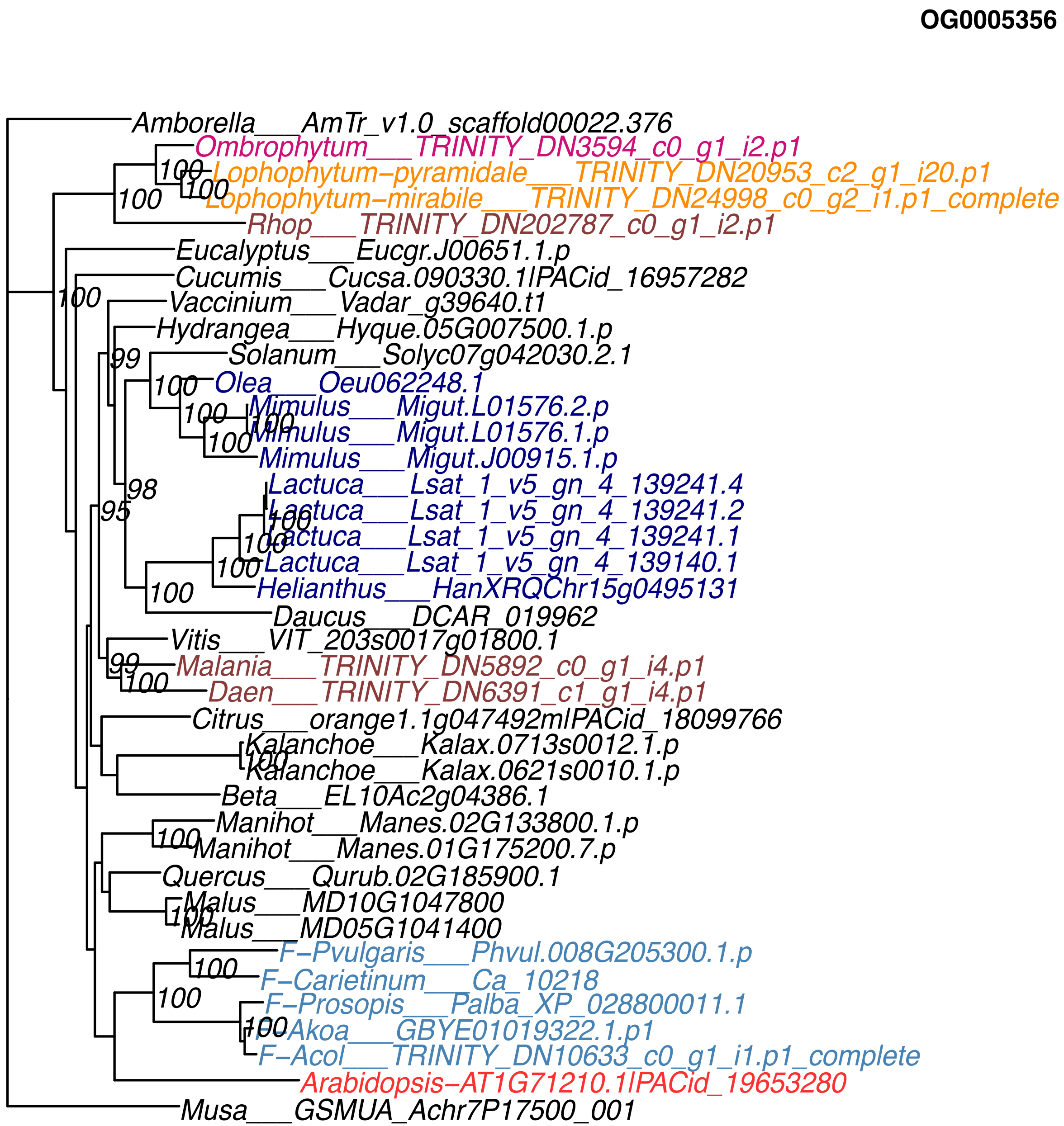
OG0005356
Amborella___AmTr_v1.0_scaffold00022.376
Ombrophytum___TRINITY_DN3594_c0_g1_i2.p1
Lophophytum−pyramidale___TRINITY_DN20953_c2_g1_i20.p1
Lophophytum−mirabile___TRINITY_DN24998_c0_g2_i1.p1_complete
Rhop___TRINITY_DN202787_c0_g1_i2.p1
Eucalyptus___Eucgr.J00651.1.p
Cucumis___Cucsa.090330.1lPACid_16957282
Vaccinium___Vadar_g39640.t1
Hydrangea___Hyque.05G007500.1.p
Solanum___Solyc07g042030.2.1
Olea___Oeu062248.1
Mimulus___Migut.L01576.2.p
Mimulus___Migut.L01576.1.p
Mimulus___Migut.J00915.1.p
Lactuca___Lsat_1_v5_gn_4_139241.4
Lactuca___Lsat_1_v5_gn_4_139241.2
Lactuca___Lsat_1_v5_gn_4_139241.1
Lactuca___Lsat_1_v5_gn_4_139140.1
Helianthus___HanXRQChr15g0495131
Daucus___DCAR_019962
Vitis___VIT_203s0017g01800.1
Malania___TRINITY_DN5892_c0_g1_i4.p1
Daen___TRINITY_DN6391_c1_g1_i4.p1
Citrus___orange1.1g047492mlPACid_18099766
Kalanchoe___Kalax.0713s0012.1.p
Kalanchoe___Kalax.0621s0010.1.p
Beta___EL10Ac2g04386.1
Manihot___Manes.02G133800.1.p
Manihot___Manes.01G175200.7.p
Quercus___Qurub.02G185900.1
Malus___MD10G1047800
Malus___MD05G1041400
F−Pvulgaris___Phvul.008G205300.1.p
F−Carietinum___Ca_10218
F−Prosopis___Palba_XP_028800011.1
F−Akoa___GBYE01019322.1.p1
F−Acol___TRINITY_DN10633_c0_g1_i1.p1_complete
Arabidopsis−AT1G71210.1lPACid_19653280
Musa___GSMUA_Achr7P17500_001

# OG0005452

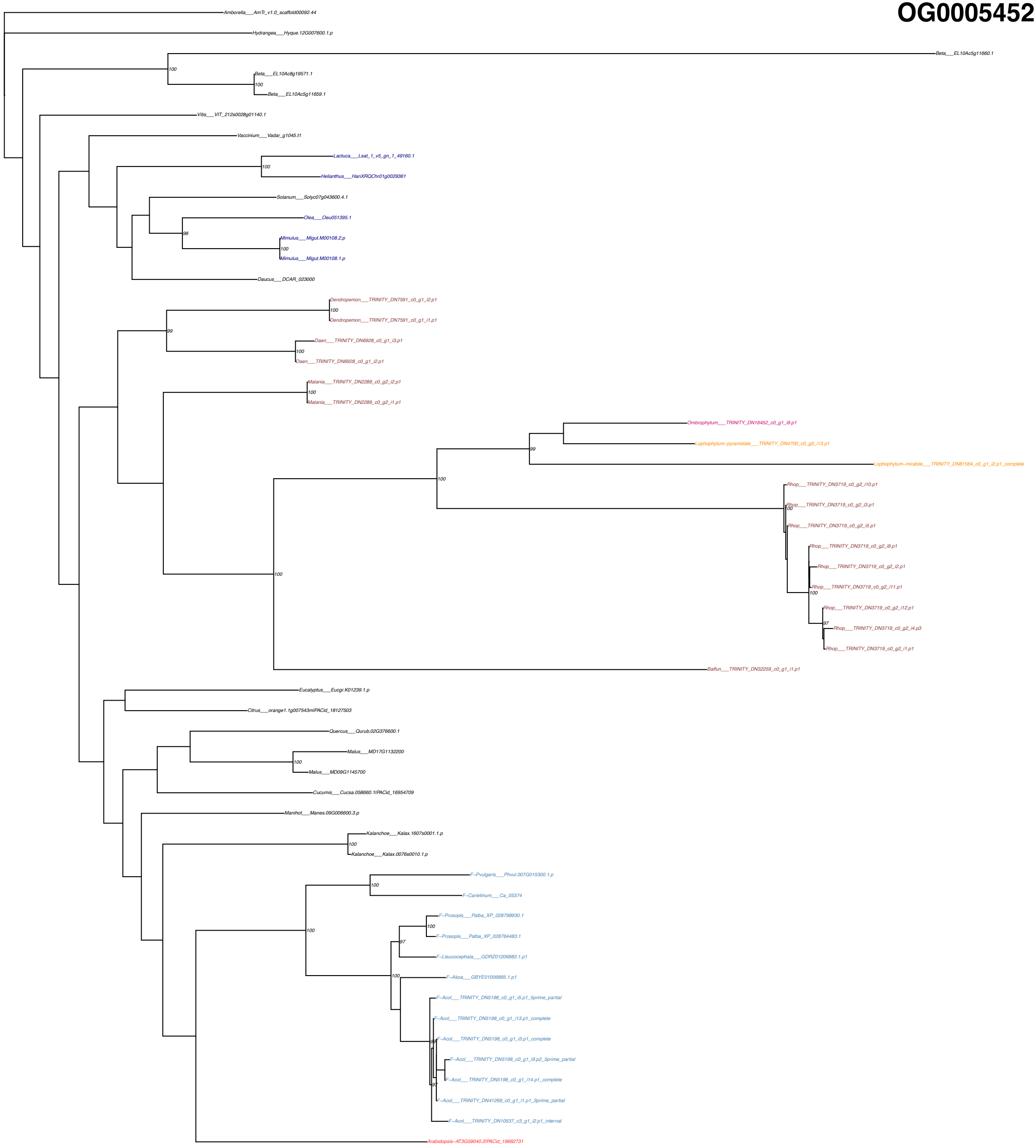

Amborella___AmTr_v1.0_scaffold00092.44
Hydrangea___Hyque.12G007600.1.p
Beta___EL10Ac5g11660.1
100
Beta___EL10Ac8g19571.1
100
Beta___EL10Ac5g11659.1
Vitis___VIT_212s0028g01140.1
Vaccinium___Vadar_g1045.t1
Lactuca___Lsat_1_v5_gn_1_49160.1
100
Helianthus___HanXRQChr01g0029361
Solanum___Solyc07g043600.4.1
Olea___Oeu051395.1
98
Mimulus___Migut.M00108.2.p
100
Mimulus___Migut.M00108.1.p
Daucus___DCAR_023000
Dendropemon___TRINITY_DN7591_c0_g1_i2.p1
100
Dendropemon___TRINITY_DN7591_c0_g1_i1.p1
99
Daen___TRINITY_DN6928_c0_g1_i3.p1
100
Daen___TRINITY_DN6928_c0_g1_i2.p1
Malania___TRINITY_DN2289_c0_g2_i2.p1
100
Malania___TRINITY_DN2289_c0_g2_i1.p1
Ombrophytum___TRINITY_DN16452_c0_g1_i9.p1
Lophophytum-pyramidale___TRINITY_DN4700_c0_g3_i13.p1
99
Lophophytum-mirabile___TRINITY_DN81564_c0_g1_i2.p1_complete
100
Rhop___TRINITY_DN3719_c0_g2_i10.p1
Rhop___TRINITY_DN3719_c0_g2_i3.p1
100
Rhop___TRINITY_DN3719_c0_g2_i5.p1
Rhop___TRINITY_DN3719_c0_g2_i9.p1
Rhop___TRINITY_DN3719_c0_g2_i2.p1
Rhop___TRINITY_DN3719_c0_g2_i11.p1
100
100
Rhop___TRINITY_DN3719_c0_g2_i12.p1
97
Rhop___TRINITY_DN3719_c0_g2_i4.p3
Rhop___TRINITY_DN3719_c0_g2_i1.p1
Balfun___TRINITY_DN32259_c0_g1_i1.p1
Eucalyptus___Eucgr.K01239.1.p
Citrus___orange1.1g007543m|PACid_18127503
Quercus___Qurub.02G376600.1
Malus___MD17G1132200
100
Malus___MD09G1145700
Cucumis___Cucsa.058660.1|PACid_16954709
Manihot___Manes.09G006600.3.p
Kalanchoe___Kalax.1607s0001.1.p
100
Kalanchoe___Kalax.0076s0010.1.p
F-Pvulgaris___Phvul.007G015300.1.p
100
F-Carietinum___Ca_05374
100
F-Prosopis___Palba_XP_028798930.1
100
F-Prosopis___Palba_XP_028764483.1
97
F-Lleucocephala___GDRZ01206883.1.p1
100
F-Akoa___GBYE01006865.1.p1
F-Acol___TRINITY_DN5198_c0_g1_i5.p1_5prime_partial
F-Acol___TRINITY_DN5198_c0_g1_i13.p1_complete
99
F-Acol___TRINITY_DN5198_c0_g1_i3.p1_complete
F-Acol___TRINITY_DN5198_c0_g1_i9.p2_5prime_partial
F-Acol___TRINITY_DN5198_c0_g1_i14.p1_complete
97
F-Acol___TRINITY_DN41269_c0_g1_i1.p1_3prime_partial
F-Acol___TRINITY_DN10537_c3_g1_i2.p1_internal
Arabidopsis-AT3G59040.2|PACid_19662731

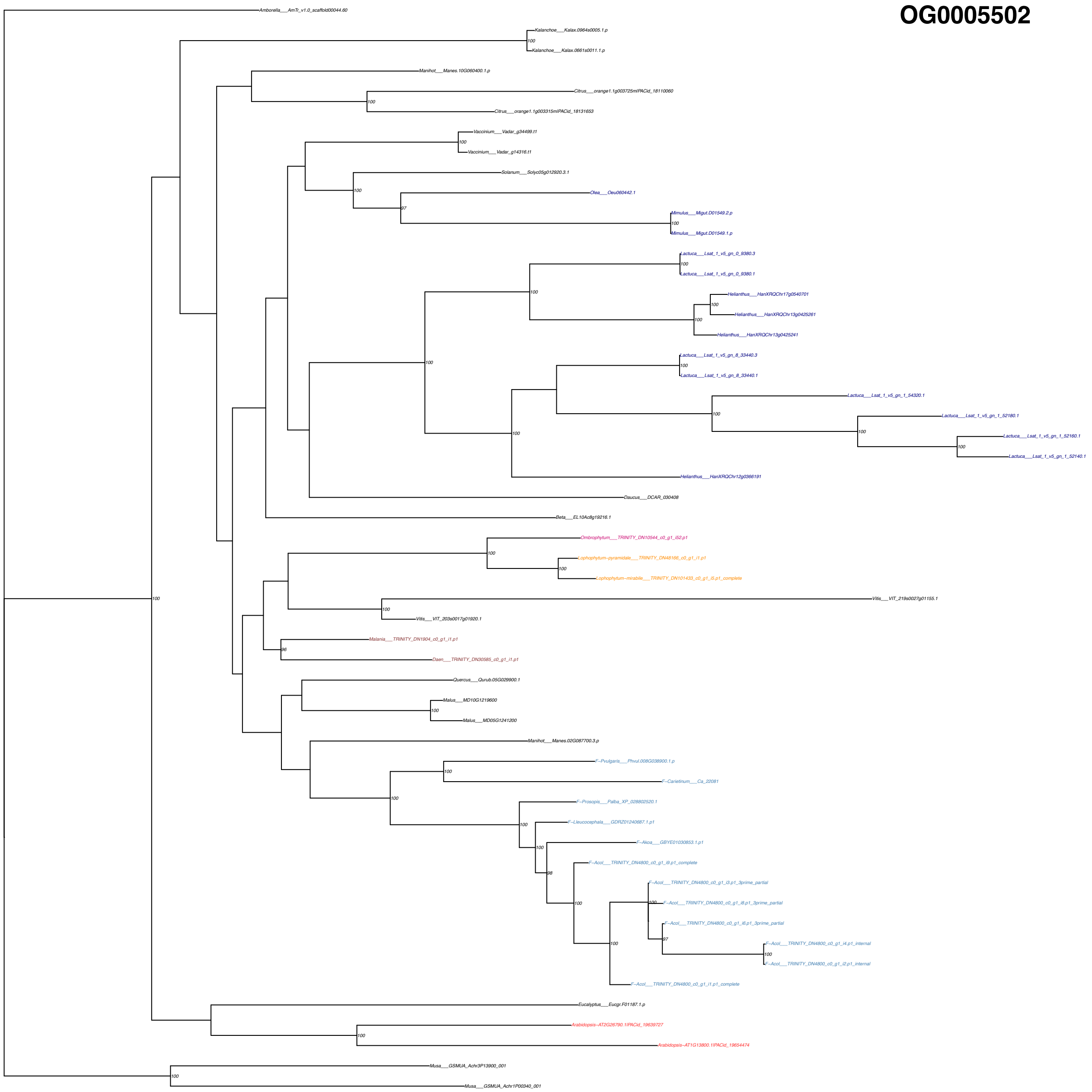
OG0005502
Amborella___AmTr_v1.0_scaffold00044.60
Kalanchoe___Kalax.0964s0005.1.p
Kalanchoe___Kalax.0661s0011.1.p
Manihot___Manes.10G060400.1.p
Citrus___orange1.1g003725mIPACid_18110060
Citrus___orange1.1g003315mIPACid_18131653
Vaccinium___Vadar_g34499.t1
Vaccinium___Vadar_g14316.t1
Solanum___Solyc05g012920.3.1
Olea___Oeu060442.1
Mimulus___Migut.D01549.2.p
Mimulus___Migut.D01549.1.p
Lactuca___Lsat_1_v5_gn_0_9380.3
Lactuca___Lsat_1_v5_gn_0_9380.1
Helianthus___HanXRQChr17g0540701
Helianthus___HanXRQChr13g0425261
Helianthus___HanXRQChr13g0425241
Lactuca___Lsat_1_v5_gn_8_33440.3
Lactuca___Lsat_1_v5_gn_8_33440.1
Lactuca___Lsat_1_v5_gn_1_54320.1
Lactuca___Lsat_1_v5_gn_1_52180.1
Lactuca___Lsat_1_v5_gn_1_52160.1
Lactuca___Lsat_1_v5_gn_1_52140.1
Helianthus___HanXRQChr12g0366191
Daucus___DCAR_030408
Beta___EL10Ac8g19216.1
Ombrophytum___TRINITY_DN10544_c0_g1_i52.p1
Lophophytum-pyramidale___TRINITY_DN48166_c0_g1_i1.p1
Lophophytum-mirabile___TRINITY_DN101433_c0_g1_i5.p1_complete
Vitis___VIT_219s0027g01155.1
Vitis___VIT_203s0017g01920.1
Malania___TRINITY_DN1904_c0_g1_i1.p1
Daen___TRINITY_DN30585_c0_g1_i1.p1
Quercus___Qurub.05G029900.1
Malus___MD10G1219600
Malus___MD05G1241200
Manihot___Manes.02G087700.3.p
F-Pvulgaris___Phvul.008G038900.1.p
F-Carietinum___Ca_22081
F-Prosopis___Palba_XP_028802520.1
F-Lleucocephala___GDRZ01240687.1.p1
F-Akoa___GBYE01030853.1.p1
F-Acol___TRINITY_DN4800_c0_g1_i9.p1_complete
F-Acol___TRINITY_DN4800_c0_g1_i3.p1_3prime_partial
F-Acol___TRINITY_DN4800_c0_g1_i8.p1_3prime_partial
F-Acol___TRINITY_DN4800_c0_g1_i6.p1_3prime_partial
F-Acol___TRINITY_DN4800_c0_g1_i4.p1_internal
F-Acol___TRINITY_DN4800_c0_g1_i2.p1_internal
F-Acol___TRINITY_DN4800_c0_g1_i1.p1_complete
Eucalyptus___Eucgr.F01187.1.p
Arabidopsis-AT2G26790.1IPACid_19639727
Arabidopsis-AT1G13800.1IPACid_19654474
Musa___GSMUA_Achr3P13900_001
Musa___GSMUA_Achr1P00340_001

OG0006083

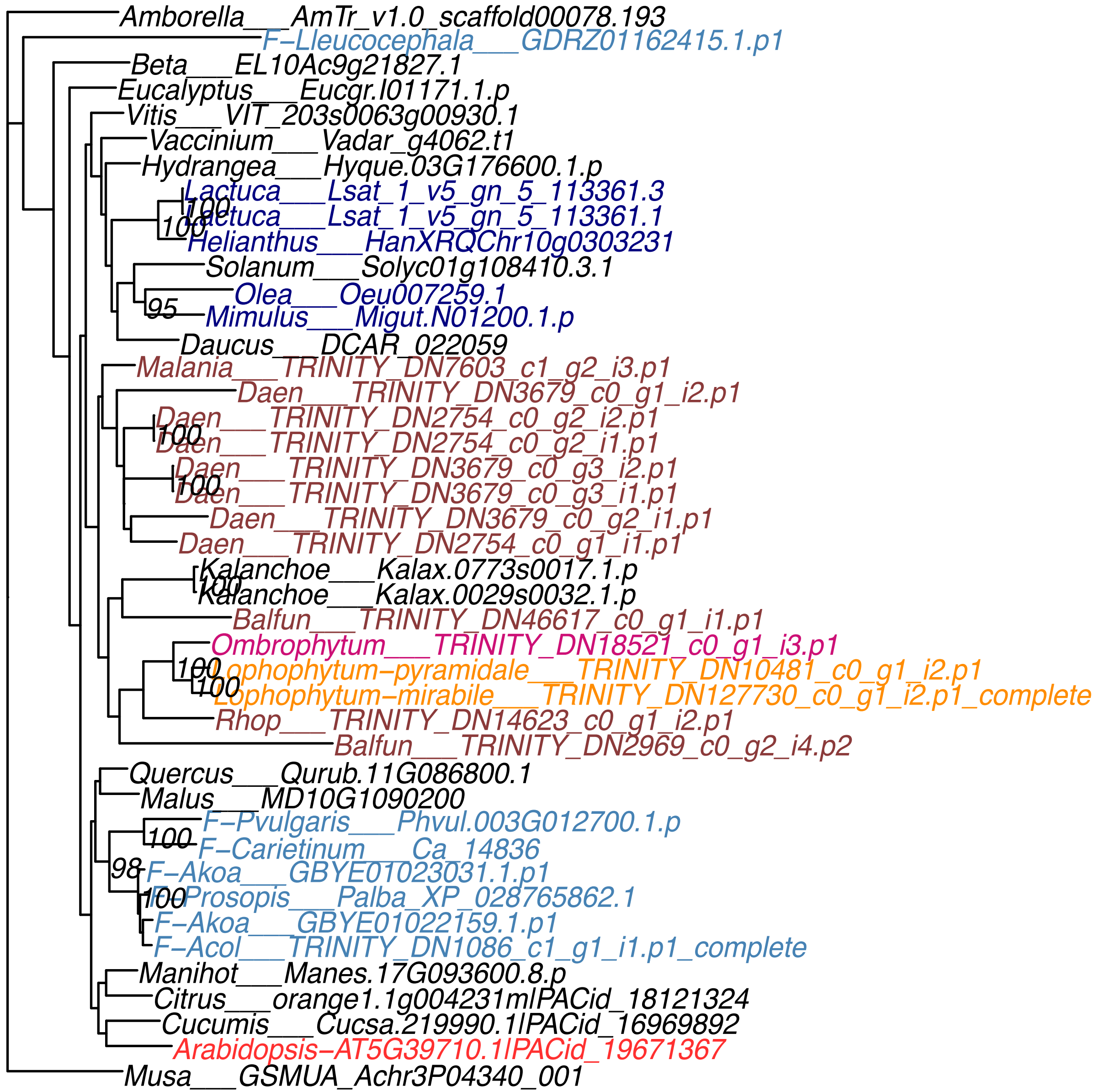
Amborella___AmTr_v1.0_scaffold00078.193
F−Lleucocephala___GDRZ01162415.1.p1
Beta___EL10Ac9g21827.1
Eucalyptus___Eucgr.I01171.1.p
Vitis___VIT_203s0063g00930.1
Vaccinium___Vadar_g4062.t1
Hydrangea___Hyque.03G176600.1.p
Lactuca___Lsat_1_v5_gn_5_113361.3
100
Lactuca___Lsat_1_v5_gn_5_113361.1
100
Helianthus___HanXRQChr10g0303231
Solanum___Solyc01g108410.3.1
Olea___Oeu007259.1
95
Mimulus___Migut.N01200.1.p
Daucus___DCAR_022059
Malania___TRINITY_DN7603_c1_g2_i3.p1
Daen___TRINITY_DN3679_c0_g1_i2.p1
Daen___TRINITY_DN2754_c0_g2_i2.p1
100
Daen___TRINITY_DN2754_c0_g2_i1.p1
Daen___TRINITY_DN3679_c0_g3_i2.p1
100
Daen___TRINITY_DN3679_c0_g3_i1.p1
Daen___TRINITY_DN3679_c0_g2_i1.p1
Daen___TRINITY_DN2754_c0_g1_i1.p1
Kalanchoe___Kalax.0773s0017.1.p
100
Kalanchoe___Kalax.0029s0032.1.p
Balfun___TRINITY_DN46617_c0_g1_i1.p1
Ombrophytum___TRINITY_DN18521_c0_g1_i3.p1
100
Lophophytum−pyramidale___TRINITY_DN10481_c0_g1_i2.p1
100
Lophophytum−mirabile___TRINITY_DN127730_c0_g1_i2.p1_complete
Rhop___TRINITY_DN14623_c0_g1_i2.p1
Balfun___TRINITY_DN2969_c0_g2_i4.p2
Quercus___Qurub.11G086800.1
Malus___MD10G1090200
F−Pvulgaris___Phvul.003G012700.1.p
100
F−Carietinum___Ca_14836
98
F−Akoa___GBYE01023031.1.p1
100
F−Prosopis___Palba_XP_028765862.1
F−Akoa___GBYE01022159.1.p1
F−Acol___TRINITY_DN1086_c1_g1_i1.p1_complete
Manihot___Manes.17G093600.8.p
Citrus___orange1.1g004231m|PACid_18121324
Cucumis___Cucsa.219990.1|PACid_16969892
Arabidopsis−AT5G39710.1|PACid_19671367
Musa___GSMUA_Achr3P04340_001

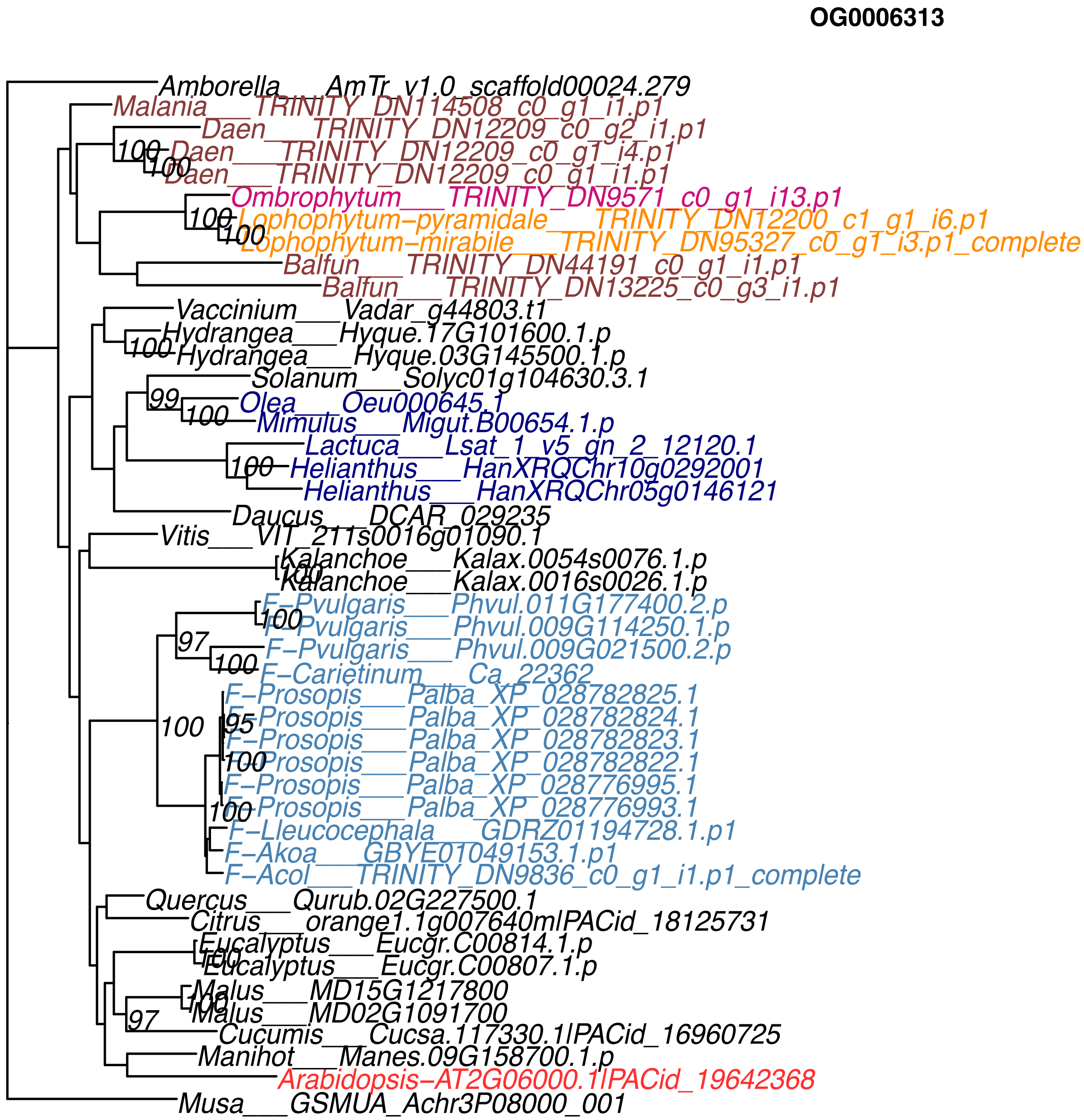
OG0006313
Amborella___AmTr_v1.0_scaffold00024.279
Malania___TRINITY_DN114508_c0_g1_i1.p1
Daen___TRINITY_DN12209_c0_g2_i1.p1
Daen___TRINITY_DN12209_c0_g1_i4.p1
Daen___TRINITY_DN12209_c0_g1_i1.p1
Ombrophytum___TRINITY_DN9571_c0_g1_i13.p1
Lophophytum−pyramidale___TRINITY_DN12200_c1_g1_i6.p1
Lophophytum−mirabile___TRINITY_DN95327_c0_g1_i3.p1_complete
Balfun___TRINITY_DN44191_c0_g1_i1.p1
Balfun___TRINITY_DN13225_c0_g3_i1.p1
Vaccinium___Vadar_g44803.t1
Hydrangea___Hyque.17G101600.1.p
Hydrangea___Hyque.03G145500.1.p
Solanum___Solyc01g104630.3.1
Olea___Oeu000645.1
Mimulus___Migut.B00654.1.p
Lactuca___Lsat_1_v5_gn_2_12120.1
Helianthus___HanXRQChr10g0292001
Helianthus___HanXRQChr05g0146121
Daucus___DCAR_029235
Vitis___VIT_211s0016g01090.1
Kalanchoe___Kalax.0054s0076.1.p
Kalanchoe___Kalax.0016s0026.1.p
F−Pvulgaris___Phvul.011G177400.2.p
F−Pvulgaris___Phvul.009G114250.1.p
F−Pvulgaris___Phvul.009G021500.2.p
F−Carietinum___Ca_22362
F−Prosopis___Palba_XP_028782825.1
F−Prosopis___Palba_XP_028782824.1
F−Prosopis___Palba_XP_028782823.1
F−Prosopis___Palba_XP_028782822.1
F−Prosopis___Palba_XP_028776995.1
F−Prosopis___Palba_XP_028776993.1
F−Lleucocephala___GDRZ01194728.1.p1
F−Akoa___GBYE01049153.1.p1
F−Acol___TRINITY_DN9836_c0_g1_i1.p1_complete
Quercus___Qurub.02G227500.1
Citrus___orange1.1g007640m|PACid_18125731
Eucalyptus___Eucgr.C00814.1.p
Eucalyptus___Eucgr.C00807.1.p
Malus___MD15G1217800
Malus___MD02G1091700
Cucumis___Cucsa.117330.1|PACid_16960725
Manihot___Manes.09G158700.1.p
Arabidopsis−AT2G06000.1|PACid_19642368
Musa___GSMUA_Achr3P08000_001
100
100
100
100
100
99
100
100
100
100
97
100
100
95
100
100
100
100
97

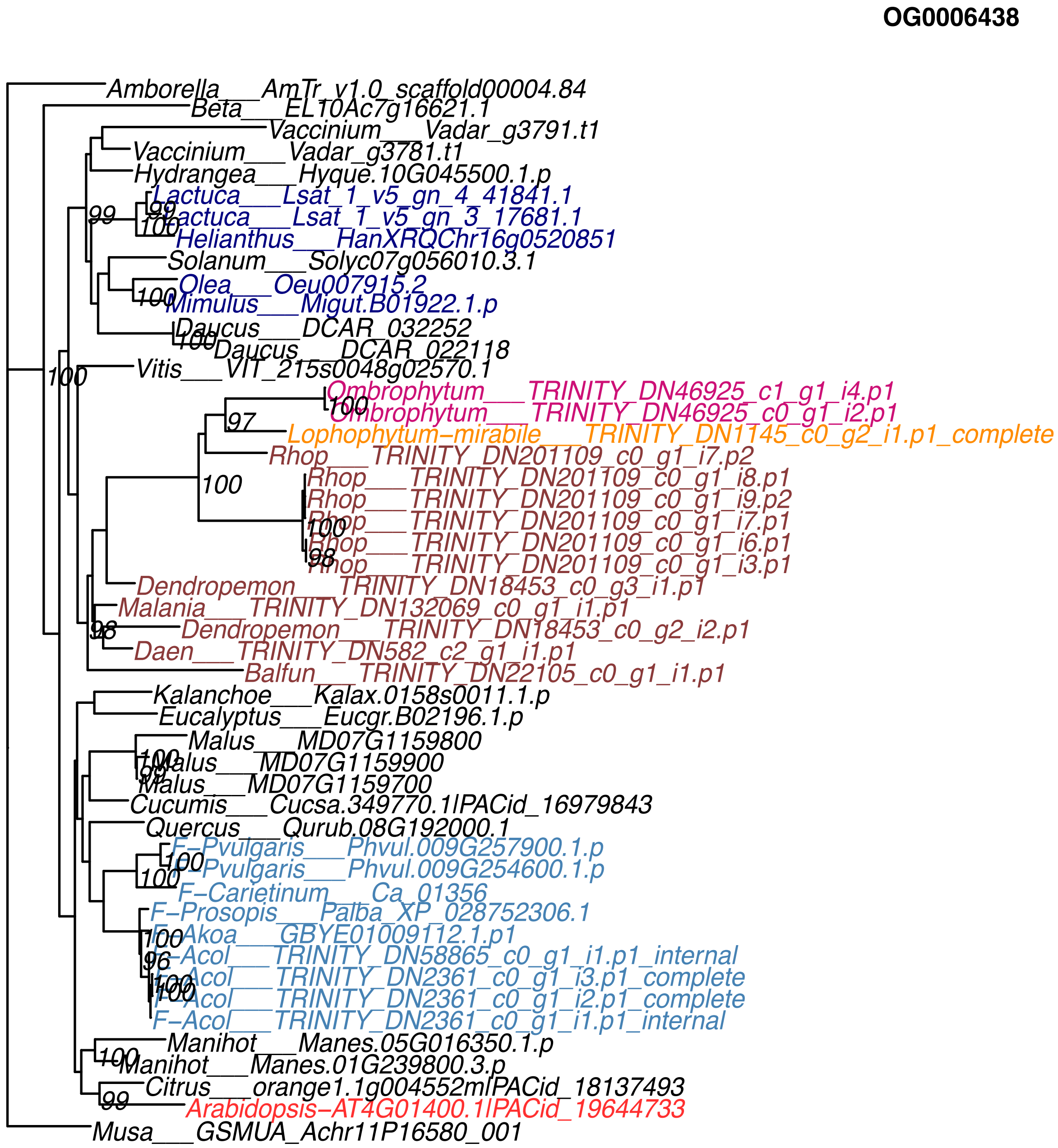
OG0006438
Amborella___AmTr_v1.0_scaffold00004.84
Beta___EL10Ac7g16621.1
Vaccinium___Vadar_g3791.t1
Vaccinium___Vadar_g3781.t1
Hydrangea___Hyque.10G045500.1.p
Lactuca___Lsat_1_v5_gn_4_41841.1
Lactuca___Lsat_1_v5_gn_3_17681.1
Helianthus___HanXRQChr16g0520851
Solanum___Solyc07g056010.3.1
Olea___Oeu007915.2
Mimulus___Migut.B01922.1.p
Daucus___DCAR_032252
Daucus___DCAR_022118
Vitis___VIT_215s0048g02570.1
Ombrophytum___TRINITY_DN46925_c1_g1_i4.p1
Ombrophytum___TRINITY_DN46925_c0_g1_i2.p1
Lophophytum−mirabile___TRINITY_DN1145_c0_g2_i1.p1_complete
Rhop___TRINITY_DN201109_c0_g1_i7.p2
Rhop___TRINITY_DN201109_c0_g1_i8.p1
Rhop___TRINITY_DN201109_c0_g1_i9.p2
Rhop___TRINITY_DN201109_c0_g1_i7.p1
Rhop___TRINITY_DN201109_c0_g1_i6.p1
Rhop___TRINITY_DN201109_c0_g1_i3.p1
Dendropemon___TRINITY_DN18453_c0_g3_i1.p1
Malania___TRINITY_DN132069_c0_g1_i1.p1
Dendropemon___TRINITY_DN18453_c0_g2_i2.p1
Daen___TRINITY_DN582_c2_g1_i1.p1
Balfun___TRINITY_DN22105_c0_g1_i1.p1
Kalanchoe___Kalax.0158s0011.1.p
Eucalyptus___Eucgr.B02196.1.p
Malus___MD07G1159800
Malus___MD07G1159900
Malus___MD07G1159700
Cucumis___Cucsa.349770.1lPACid_16979843
Quercus___Qurub.08G192000.1
F−Pvulgaris___Phvul.009G257900.1.p
F−Pvulgaris___Phvul.009G254600.1.p
F−Carietinum___Ca_01356
F−Prosopis___Palba_XP_028752306.1
F−Akoa___GBYE01009112.1.p1
F−Acol___TRINITY_DN58865_c0_g1_i1.p1_internal
F−Acol___TRINITY_DN2361_c0_g1_i3.p1_complete
F−Acol___TRINITY_DN2361_c0_g1_i2.p1_complete
F−Acol___TRINITY_DN2361_c0_g1_i1.p1_internal
Manihot___Manes.05G016350.1.p
Manihot___Manes.01G239800.3.p
Citrus___orange1.1g004552mlPACid_18137493
Arabidopsis−AT4G01400.1lPACid_19644733
Musa___GSMUA_Achr11P16580_001
99
99
100
100
100
100
100
97
100
100
98
98
100
99
100
100
100
96
100
100
100
99

**OG0006753**

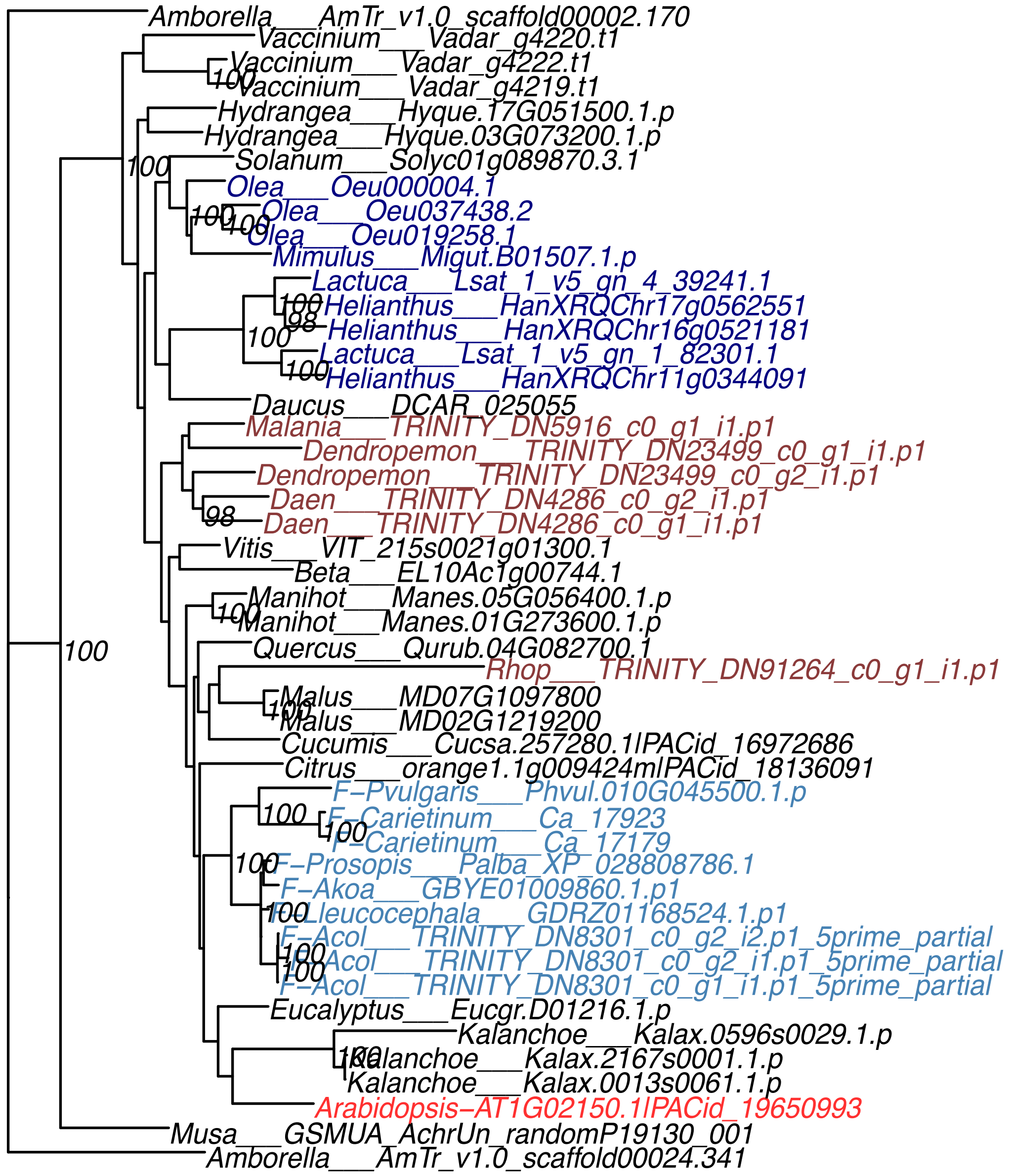
Amborella___AmTr_v1.0_scaffold00002.170
Vaccinium___Vadar_g4220.t1
Vaccinium___Vadar_g4222.t1
100
Vaccinium___Vadar_g4219.t1
Hydrangea___Hyque.17G051500.1.p
Hydrangea___Hyque.03G073200.1.p
100
Solanum___Solyc01g089870.3.1
Olea___Oeu000004.1
100
Olea___Oeu037438.2
100
Olea___Oeu019258.1
Mimulus___Migut.B01507.1.p
Lactuca___Lsat_1_v5_gn_4_39241.1
100
Helianthus___HanXRQChr17g0562551
98
Helianthus___HanXRQChr16g0521181
100
Lactuca___Lsat_1_v5_gn_1_82301.1
100
Helianthus___HanXRQChr11g0344091
Daucus___DCAR_025055
Malania___TRINITY_DN5916_c0_g1_i1.p1
Dendropemon___TRINITY_DN23499_c0_g1_i1.p1
Dendropemon___TRINITY_DN23499_c0_g2_i1.p1
Daen___TRINITY_DN4286_c0_g2_i1.p1
98
Daen___TRINITY_DN4286_c0_g1_i1.p1
Vitis___VIT_215s0021g01300.1
Beta___EL10Ac1g00744.1
Manihot___Manes.05G056400.1.p
100
Manihot___Manes.01G273600.1.p
Quercus___Qurub.04G082700.1
100
Rhop___TRINITY_DN91264_c0_g1_i1.p1
Malus___MD07G1097800
100
Malus___MD02G1219200
Cucumis___Cucsa.257280.1|PACid_16972686
Citrus___orange1.1g009424m|PACid_18136091
F−Pvulgaris___Phvul.010G045500.1.p
100
F−Carietinum___Ca_17923
100
F−Carietinum___Ca_17179
100
F−Prosopis___Palba_XP_028808786.1
F−Akoa___GBYE01009860.1.p1
100
F−Lleucocephala___GDRZ01168524.1.p1
F−Acol___TRINITY_DN8301_c0_g2_i2.p1_5prime_partial
100
F−Acol___TRINITY_DN8301_c0_g2_i1.p1_5prime_partial
100
F−Acol___TRINITY_DN8301_c0_g1_i1.p1_5prime_partial
Eucalyptus___Eucgr.D01216.1.p
Kalanchoe___Kalax.0596s0029.1.p
100
Kalanchoe___Kalax.2167s0001.1.p
Kalanchoe___Kalax.0013s0061.1.p
Arabidopsis−AT1G02150.1|PACid_19650993
Musa___GSMUA_AchrUn_randomP19130_001
Amborella___AmTr_v1.0_scaffold00024.341

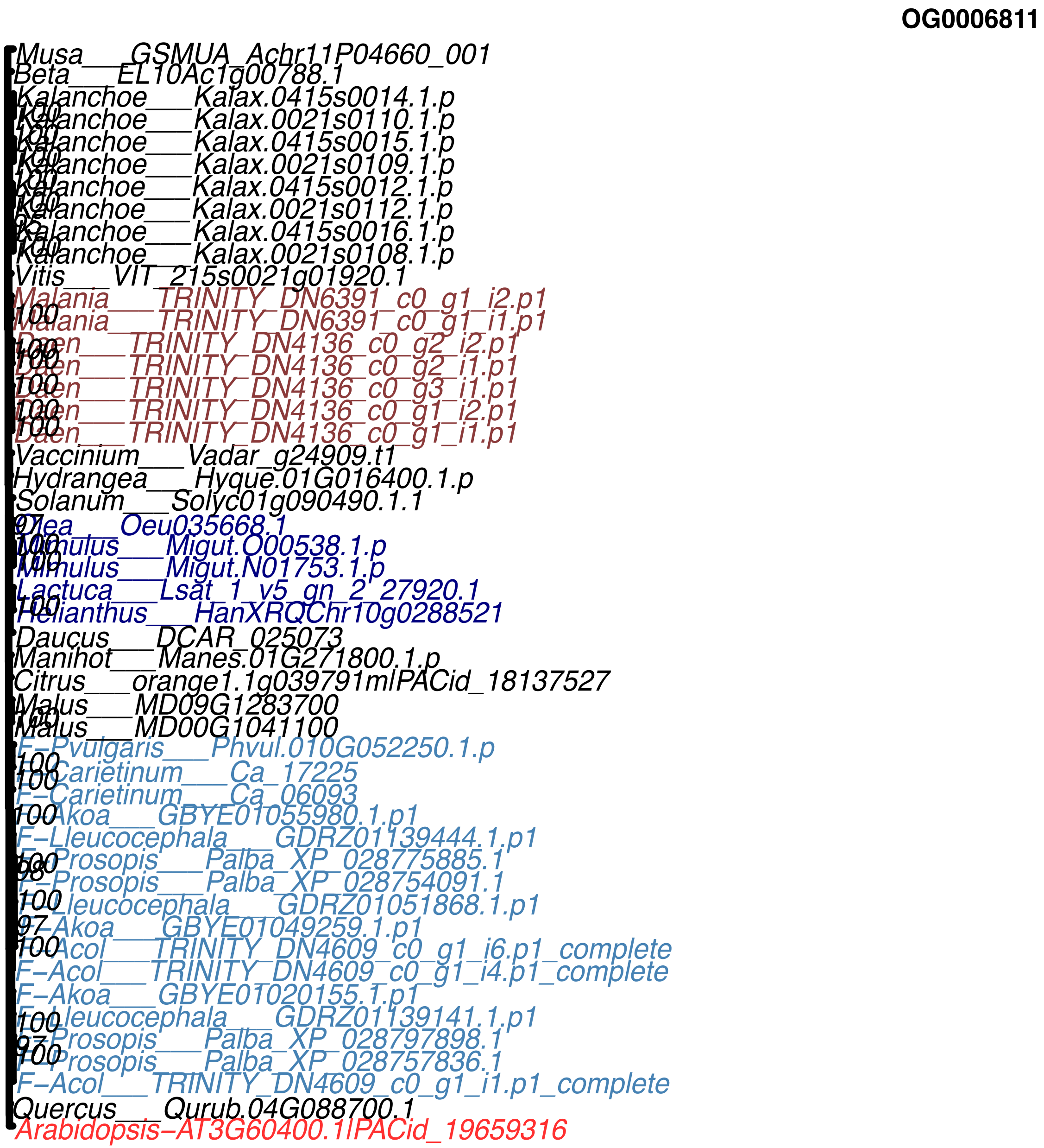
OG0006811
Musa___GSMUA_Achr11P04660_001
Beta___EL10Ac1g00788.1
Kalanchoe___Kalax.0415s0014.1.p
Kalanchoe___Kalax.0021s0110.1.p
Kalanchoe___Kalax.0415s0015.1.p
Kalanchoe___Kalax.0021s0109.1.p
Kalanchoe___Kalax.0415s0012.1.p
Kalanchoe___Kalax.0021s0112.1.p
Kalanchoe___Kalax.0415s0016.1.p
Kalanchoe___Kalax.0021s0108.1.p
Vitis___VIT_215s0021g01920.1
Malania___TRINITY_DN6391_c0_g1_i2.p1
Malania___TRINITY_DN6391_c0_g1_i1.p1
Daen___TRINITY_DN4136_c0_g2_i2.p1
Daen___TRINITY_DN4136_c0_g2_i1.p1
Daen___TRINITY_DN4136_c0_g3_i1.p1
Daen___TRINITY_DN4136_c0_g1_i2.p1
Daen___TRINITY_DN4136_c0_g1_i1.p1
Vaccinium___Vadar_g24909.t1
Hydrangea___Hyque.01G016400.1.p
Solanum___Solyc01g090490.1.1
Olea___Oeu035668.1
Mimulus___Migut.O00538.1.p
Mimulus___Migut.N01753.1.p
Lactuca___Lsat_1_v5_gn_2_27920.1
Helianthus___HanXRQChr10g0288521
Daucus___DCAR_025073
Manihot___Manes.01G271800.1.p
Citrus___orange1.1g039791mlPACid_18137527
Malus___MD09G1283700
Malus___MD00G1041100
F–Pvulgaris___Phvul.010G052250.1.p
F–Carietinum___Ca_17225
F–Carietinum___Ca_06093
F–Akoa___GBYE01055980.1.p1
F–Lleucocephala___GDRZ01139444.1.p1
F–Prosopis___Palba_XP_028775885.1
F–Prosopis___Palba_XP_028754091.1
F–Lleucocephala___GDRZ01051868.1.p1
F–Akoa___GBYE01049259.1.p1
F–Acol___TRINITY_DN4609_c0_g1_i6.p1_complete
F–Acol___TRINITY_DN4609_c0_g1_i4.p1_complete
F–Akoa___GBYE01020155.1.p1
F–Lleucocephala___GDRZ01139141.1.p1
F–Prosopis___Palba_XP_028797898.1
F–Prosopis___Palba_XP_028757836.1
F–Acol___TRINITY_DN4609_c0_g1_i1.p1_complete
Quercus___Qurub.04G088700.1
Arabidopsis–AT3G60400.1lPACid_19659316

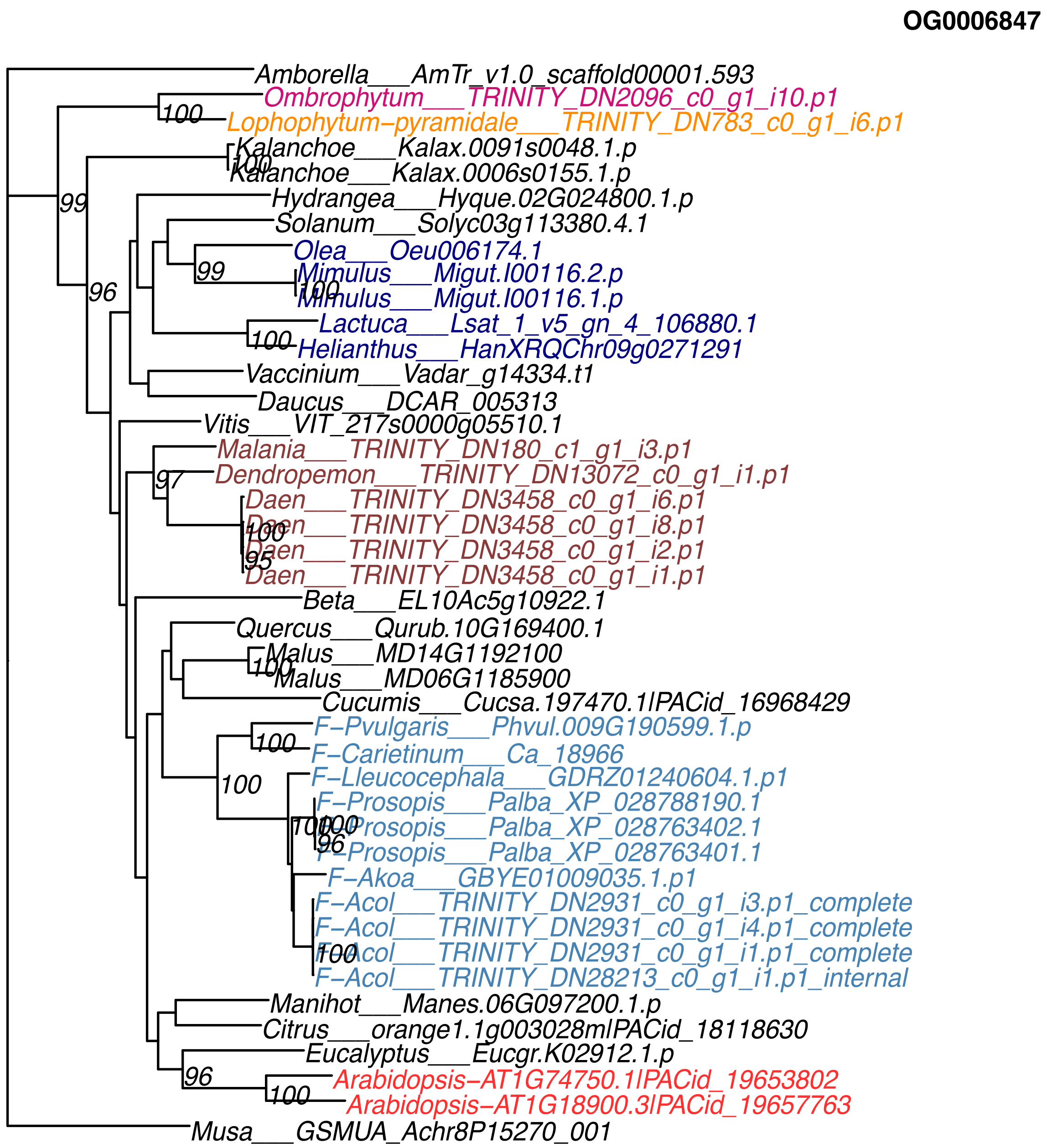
OG0006847
Amborella___AmTr_v1.0_scaffold00001.593
Ombrophytum___TRINITY_DN2096_c0_g1_i10.p1
Lophophytum−pyramidale___TRINITY_DN783_c0_g1_i6.p1
Kalanchoe___Kalax.0091s0048.1.p
Kalanchoe___Kalax.0006s0155.1.p
Hydrangea___Hyque.02G024800.1.p
Solanum___Solyc03g113380.4.1
Olea___Oeu006174.1
Mimulus___Migut.I00116.2.p
Mimulus___Migut.I00116.1.p
Lactuca___Lsat_1_v5_gn_4_106880.1
Helianthus___HanXRQChr09g0271291
Vaccinium___Vadar_g14334.t1
Daucus___DCAR_005313
Vitis___VIT_217s0000g05510.1
Malania___TRINITY_DN180_c1_g1_i3.p1
Dendropemon___TRINITY_DN13072_c0_g1_i1.p1
Daen___TRINITY_DN3458_c0_g1_i6.p1
Daen___TRINITY_DN3458_c0_g1_i8.p1
Daen___TRINITY_DN3458_c0_g1_i2.p1
Daen___TRINITY_DN3458_c0_g1_i1.p1
Beta___EL10Ac5g10922.1
Quercus___Qurub.10G169400.1
Malus___MD14G1192100
Malus___MD06G1185900
Cucumis___Cucsa.197470.1|PACid_16968429
F−Pvulgaris___Phvul.009G190599.1.p
F−Carietinum___Ca_18966
F−Lleucocephala___GDRZ01240604.1.p1
F−Prosopis___Palba_XP_028788190.1
F−Prosopis___Palba_XP_028763402.1
F−Prosopis___Palba_XP_028763401.1
F−Akoa___GBYE01009035.1.p1
F−Acol___TRINITY_DN2931_c0_g1_i3.p1_complete
F−Acol___TRINITY_DN2931_c0_g1_i4.p1_complete
F−Acol___TRINITY_DN2931_c0_g1_i1.p1_complete
F−Acol___TRINITY_DN28213_c0_g1_i1.p1_internal
Manihot___Manes.06G097200.1.p
Citrus___orange1.1g003028m|PACid_18118630
Eucalyptus___Eucgr.K02912.1.p
Arabidopsis−AT1G74750.1|PACid_19653802
Arabidopsis−AT1G18900.3|PACid_19657763
Musa___GSMUA_Achr8P15270_001
100
100
99
99
96
100
100
97
100
95
100
100
100
100
100
96
100
96
100

**OG0007047**

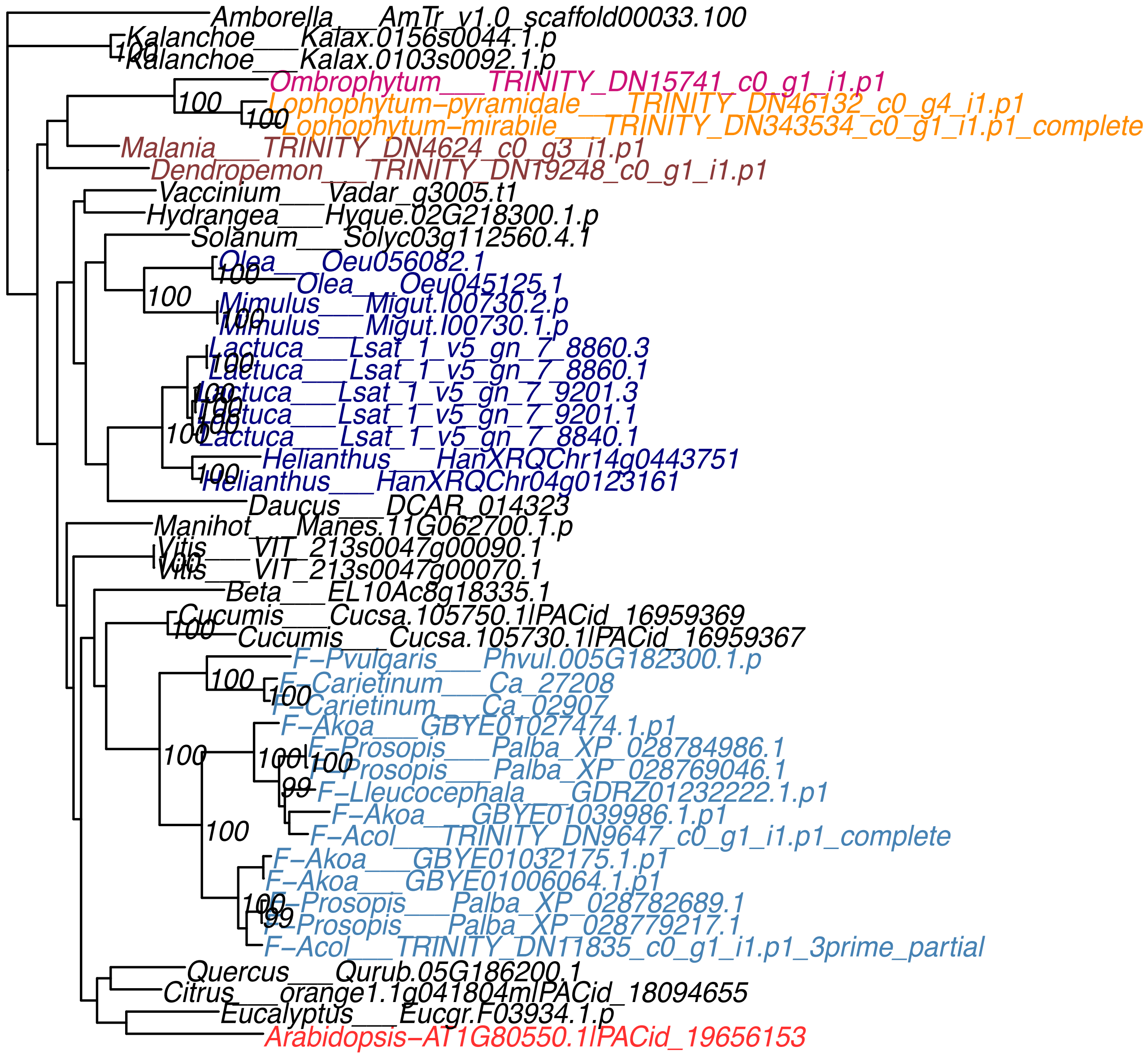
Amborella___AmTr_v1.0_scaffold00033.100
Kalanchoe___Kalax.0156s0044.1.p
Kalanchoe___Kalax.0103s0092.1.p
Ombrophytum___TRINITY_DN15741_c0_g1_i1.p1
Lophophytum–pyramidale___TRINITY_DN46132_c0_g4_i1.p1
Lophophytum–mirabile___TRINITY_DN343534_c0_g1_i1.p1_complete
Malania___TRINITY_DN4624_c0_g3_i1.p1
Dendropemon___TRINITY_DN19248_c0_g1_i1.p1
Vaccinium___Vadar_g3005.t1
Hydrangea___Hyque.02G218300.1.p
Solanum___Solyc03g112560.4.1
Olea___Oeu056082.1
Olea___Oeu045125.1
Mimulus___Migut.I00730.2.p
Mimulus___Migut.I00730.1.p
Lactuca___Lsat_1_v5_gn_7_8860.3
Lactuca___Lsat_1_v5_gn_7_8860.1
Lactuca___Lsat_1_v5_gn_7_9201.3
Lactuca___Lsat_1_v5_gn_7_9201.1
Lactuca___Lsat_1_v5_gn_7_8840.1
Helianthus___HanXRQChr14g0443751
Helianthus___HanXRQChr04g0123161
Daucus___DCAR_014323
Manihot___Manes.11G062700.1.p
Vitis___VIT_213s0047g00090.1
Vitis___VIT_213s0047g00070.1
Beta___EL10Ac8g18335.1
Cucumis___Cucsa.105750.1IPACid_16959369
Cucumis___Cucsa.105730.1IPACid_16959367
F–Pvulgaris___Phvul.005G182300.1.p
F–Carietinum___Ca_27208
F–Carietinum___Ca_02907
F–Akoa___GBYE01027474.1.p1
F–Prosopis___Palba_XP_028784986.1
F–Prosopis___Palba_XP_028769046.1
F–Lleucocephala___GDRZ01232222.1.p1
F–Akoa___GBYE01039986.1.p1
F–Acol___TRINITY_DN9647_c0_g1_i1.p1_complete
F–Akoa___GBYE01032175.1.p1
F–Akoa___GBYE01006064.1.p1
F–Prosopis___Palba_XP_028782689.1
F–Prosopis___Palba_XP_028779217.1
F–Acol___TRINITY_DN11835_c0_g1_i1.p1_3prime_partial
Quercus___Qurub.05G186200.1
Citrus___orange1.1g041804mIPACid_18094655
Eucalyptus___Eucgr.F03934.1.p
Arabidopsis–AT1G80550.1IPACid_19656153

**OG0007057**

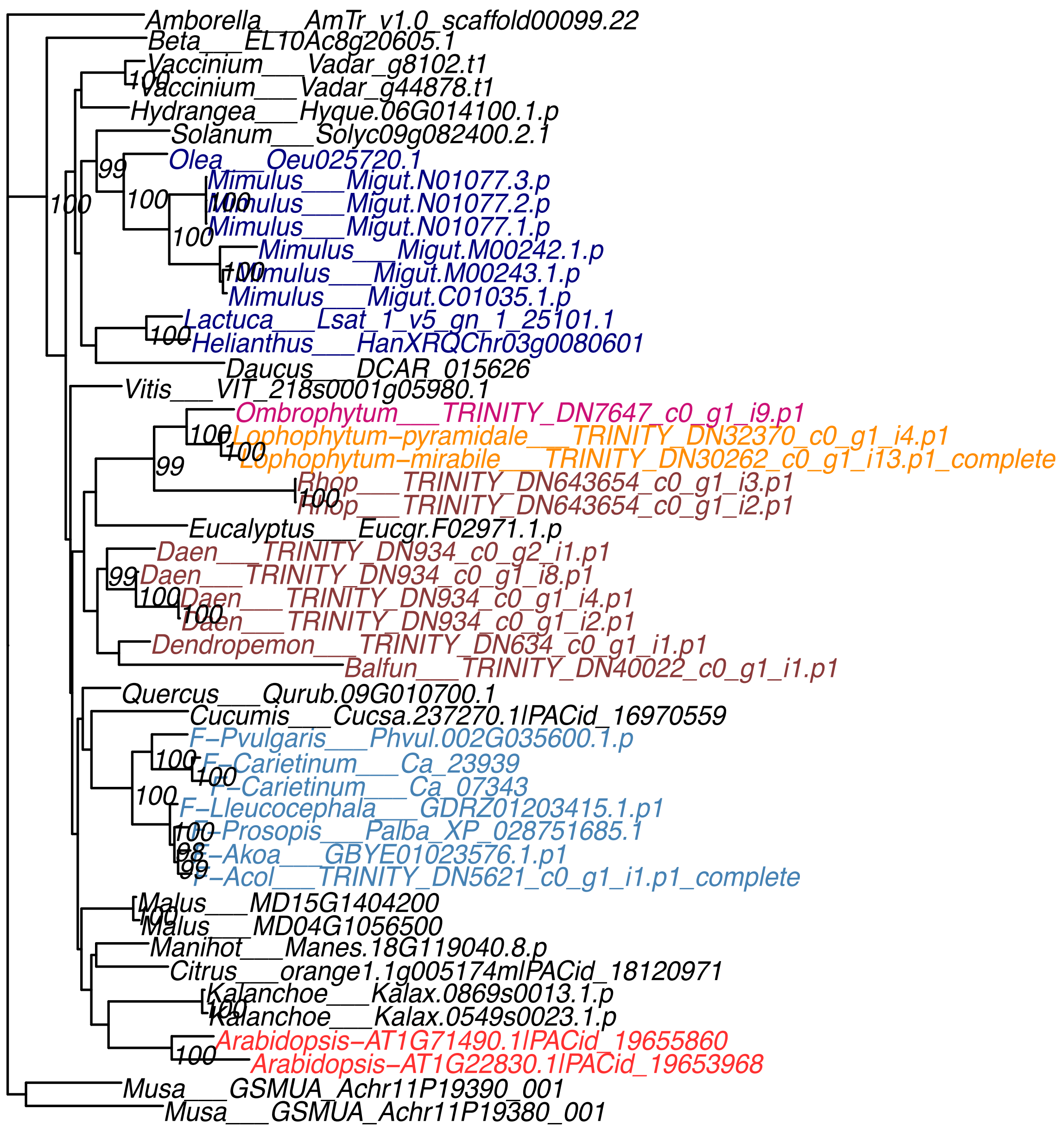
Amborella___AmTr_v1.0_scaffold00099.22
Beta___EL10Ac8g20605.1
Vaccinium___Vadar_g8102.t1
Vaccinium___Vadar_g44878.t1
Hydrangea___Hyque.06G014100.1.p
Solanum___Solyc09g082400.2.1
Olea___Oeu025720.1
Mimulus___Migut.N01077.3.p
Mimulus___Migut.N01077.2.p
Mimulus___Migut.N01077.1.p
Mimulus___Migut.M00242.1.p
Mimulus___Migut.M00243.1.p
Mimulus___Migut.C01035.1.p
Lactuca___Lsat_1_v5_gn_1_25101.1
Helianthus___HanXRQChr03g0080601
Daucus___DCAR_015626
Vitis___VIT_218s0001g05980.1
Ombrophytum___TRINITY_DN7647_c0_g1_i9.p1
Lophophytum−pyramidale___TRINITY_DN32370_c0_g1_i4.p1
Lophophytum−mirabile___TRINITY_DN30262_c0_g1_i13.p1_complete
Rhop___TRINITY_DN643654_c0_g1_i3.p1
Rhop___TRINITY_DN643654_c0_g1_i2.p1
Eucalyptus___Eucgr.F02971.1.p
Daen___TRINITY_DN934_c0_g2_i1.p1
Daen___TRINITY_DN934_c0_g1_i8.p1
Daen___TRINITY_DN934_c0_g1_i4.p1
Daen___TRINITY_DN934_c0_g1_i2.p1
Dendropemon___TRINITY_DN634_c0_g1_i1.p1
Balfun___TRINITY_DN40022_c0_g1_i1.p1
Quercus___Qurub.09G010700.1
Cucumis___Cucsa.237270.1|PACid_16970559
F−Pvulgaris___Phvul.002G035600.1.p
F−Carietinum___Ca_23939
F−Carietinum___Ca_07343
F−Lleucocephala___GDRZ01203415.1.p1
F−Prosopis___Palba_XP_028751685.1
F−Akoa___GBYE01023576.1.p1
F−Acol___TRINITY_DN5621_c0_g1_i1.p1_complete
Malus___MD15G1404200
Malus___MD04G1056500
Manihot___Manes.18G119040.8.p
Citrus___orange1.1g005174m|PACid_18120971
Kalanchoe___Kalax.0869s0013.1.p
Kalanchoe___Kalax.0549s0023.1.p
Arabidopsis−AT1G71490.1|PACid_19655860
Arabidopsis−AT1G22830.1|PACid_19653968
Musa___GSMUA_Achr11P19390_001
Musa___GSMUA_Achr11P19380_001
100
100
99
100
100
100
100
100
100
100
99
99
100
100
100
100
100
100
98
99
100
100
100

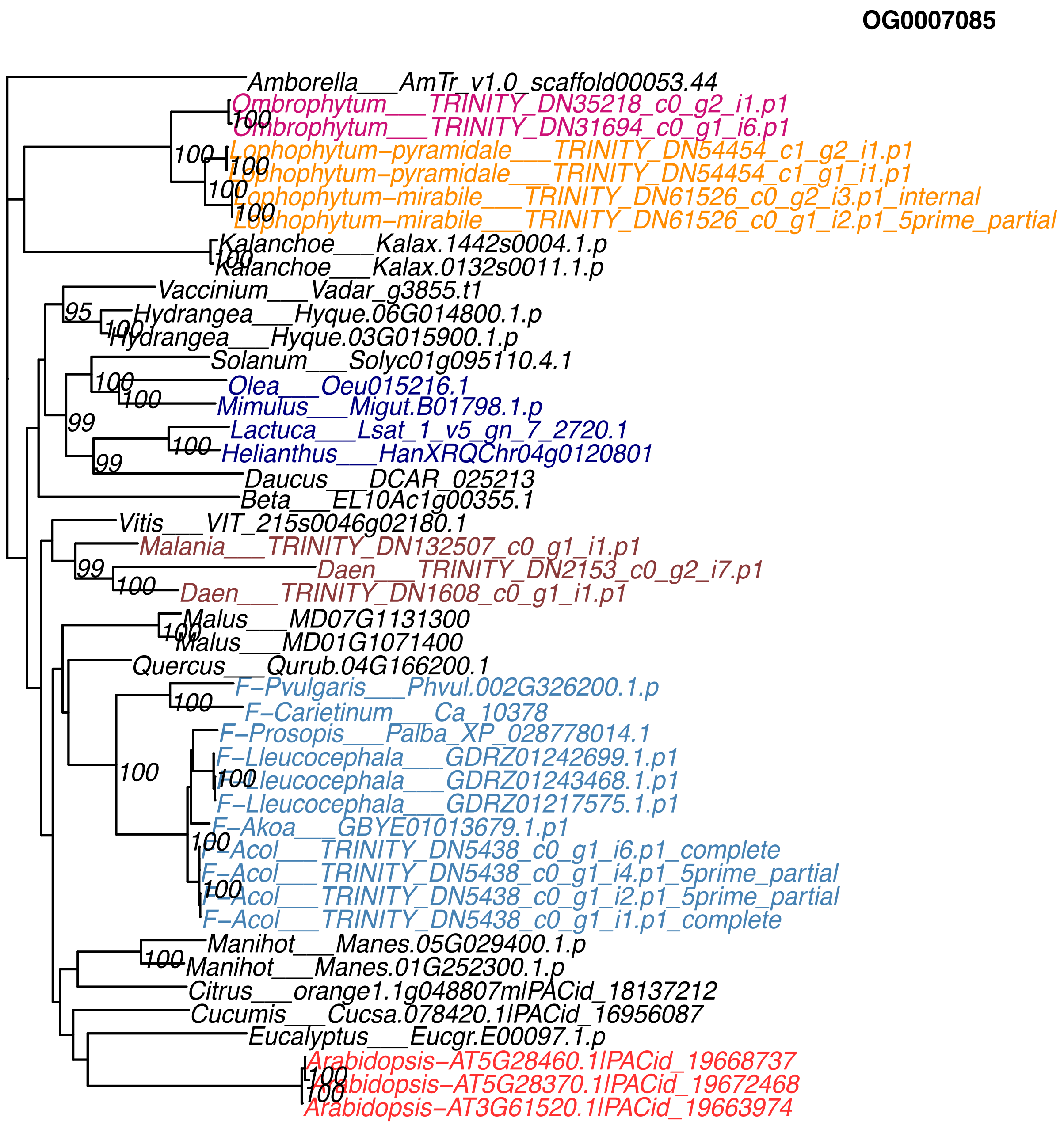
OG0007085
Amborella___AmTr_v1.0_scaffold00053.44
Ombrophytum___TRINITY_DN35218_c0_g2_i1.p1
Ombrophytum___TRINITY_DN31694_c0_g1_i6.p1
Lophophytum-pyramidale___TRINITY_DN54454_c1_g2_i1.p1
Lophophytum-pyramidale___TRINITY_DN54454_c1_g1_i1.p1
Lophophytum-mirabile___TRINITY_DN61526_c0_g2_i3.p1_internal
Lophophytum-mirabile___TRINITY_DN61526_c0_g1_i2.p1_5prime_partial
Kalanchoe___Kalax.1442s0004.1.p
Kalanchoe___Kalax.0132s0011.1.p
Vaccinium___Vadar_g3855.t1
Hydrangea___Hyque.06G014800.1.p
Hydrangea___Hyque.03G015900.1.p
Solanum___Solyc01g095110.4.1
Olea___Oeu015216.1
Mimulus___Migut.B01798.1.p
Lactuca___Lsat_1_v5_gn_7_2720.1
Helianthus___HanXRQChr04g0120801
Daucus___DCAR_025213
Beta___EL10Ac1g00355.1
Vitis___VIT_215s0046g02180.1
Malania___TRINITY_DN132507_c0_g1_i1.p1
Daen___TRINITY_DN2153_c0_g2_i7.p1
Daen___TRINITY_DN1608_c0_g1_i1.p1
Malus___MD07G1131300
Malus___MD01G1071400
Quercus___Qurub.04G166200.1
F-Pvulgaris___Phvul.002G326200.1.p
F-Carietinum___Ca_10378
F-Prosopis___Palba_XP_028778014.1
F-Lleucocephala___GDRZ01242699.1.p1
F-Lleucocephala___GDRZ01243468.1.p1
F-Lleucocephala___GDRZ01217575.1.p1
F-Akoa___GBYE01013679.1.p1
F-Acol___TRINITY_DN5438_c0_g1_i6.p1_complete
F-Acol___TRINITY_DN5438_c0_g1_i4.p1_5prime_partial
F-Acol___TRINITY_DN5438_c0_g1_i2.p1_5prime_partial
F-Acol___TRINITY_DN5438_c0_g1_i1.p1_complete
Manihot___Manes.05G029400.1.p
Manihot___Manes.01G252300.1.p
Citrus___orange1.1g048807m|PACid_18137212
Cucumis___Cucsa.078420.1|PACid_16956087
Eucalyptus___Eucgr.E00097.1.p
Arabidopsis-AT5G28460.1|PACid_19668737
Arabidopsis-AT5G28370.1|PACid_19672468
Arabidopsis-AT3G61520.1|PACid_19663974
100
100
100
100
100
95
100
100
100
99
100
99
99
100
100
100
100
100
100
100
100
100
100

**OG0007116**

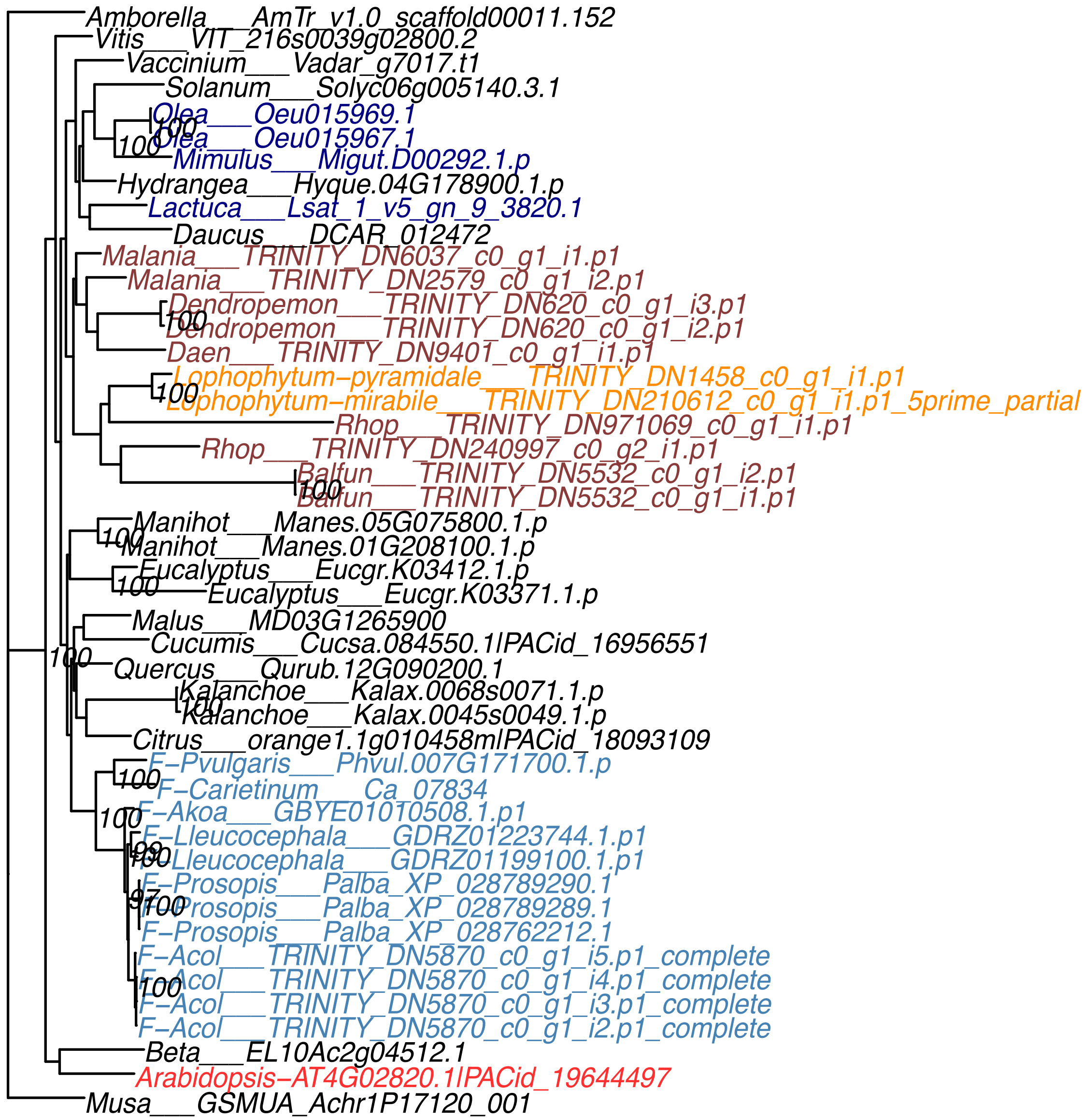

Amborella___AmTr_v1.0_scaffold00011.152
Vitis___VIT_216s0039g02800.2
Vaccinium___Vadar_g7017.t1
Solanum___Solyc06g005140.3.1
Olea___Oeu015969.1
100
Olea___Oeu015967.1
100
Mimulus___Migut.D00292.1.p
Hydrangea___Hyque.04G178900.1.p
Lactuca___Lsat_1_v5_gn_9_3820.1
Daucus___DCAR_012472
Malania___TRINITY_DN6037_c0_g1_i1.p1
Malania___TRINITY_DN2579_c0_g1_i2.p1
Dendropemon___TRINITY_DN620_c0_g1_i3.p1
100
Dendropemon___TRINITY_DN620_c0_g1_i2.p1
Daen___TRINITY_DN9401_c0_g1_i1.p1
Lophophytum−pyramidale___TRINITY_DN1458_c0_g1_i1.p1
100
Lophophytum−mirabile___TRINITY_DN210612_c0_g1_i1.p1_5prime_partial
Rhop___TRINITY_DN971069_c0_g1_i1.p1
Rhop___TRINITY_DN240997_c0_g2_i1.p1
Balfun___TRINITY_DN5532_c0_g1_i2.p1
100
Balfun___TRINITY_DN5532_c0_g1_i1.p1
Manihot___Manes.05G075800.1.p
100
Manihot___Manes.01G208100.1.p
Eucalyptus___Eucgr.K03412.1.p
100
Eucalyptus___Eucgr.K03371.1.p
Malus___MD03G1265900
Cucumis___Cucsa.084550.1|PACid_16956551
100
Quercus___Qurub.12G090200.1
Kalanchoe___Kalax.0068s0071.1.p
100
Kalanchoe___Kalax.0045s0049.1.p
Citrus___orange1.1g010458m|PACid_18093109
F−Pvulgaris___Phvul.007G171700.1.p
100
F−Carietinum___Ca_07834
100
F−Akoa___GBYE01010508.1.p1
F−Lleucocephala___GDRZ01223744.1.p1
99
100
F−Lleucocephala___GDRZ01199100.1.p1
F−Prosopis___Palba_XP_028789290.1
97
100
F−Prosopis___Palba_XP_028789289.1
F−Prosopis___Palba_XP_028762212.1
F−Acol___TRINITY_DN5870_c0_g1_i5.p1_complete
F−Acol___TRINITY_DN5870_c0_g1_i4.p1_complete
100
F−Acol___TRINITY_DN5870_c0_g1_i3.p1_complete
F−Acol___TRINITY_DN5870_c0_g1_i2.p1_complete
Beta___EL10Ac2g04512.1
Arabidopsis−AT4G02820.1|PACid_19644497
Musa___GSMUA_Achr1P17120_001

# OG0007179

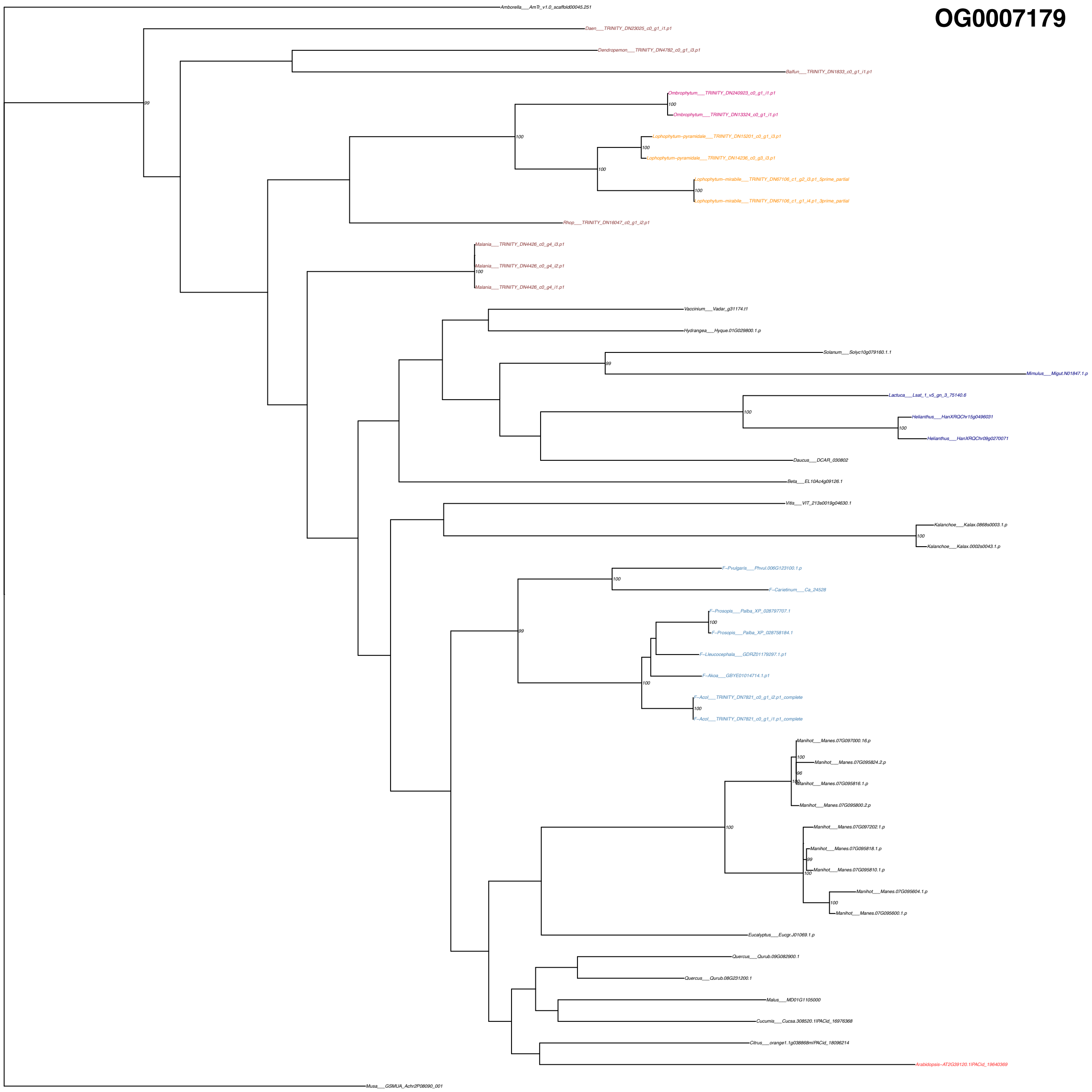

Amborella___AmTr_v1.0_scaffold00045.251
Daen___TRINITY_DN23025_c0_g1_i1.p1
Dendropemon___TRINITY_DN4782_c0_g1_i3.p1
Balfun___TRINITY_DN1833_c0_g1_i1.p1
Ombrophytum___TRINITY_DN240923_c0_g1_i1.p1
Ombrophytum___TRINITY_DN13324_c0_g1_i1.p1
Lophophytum-pyramidale___TRINITY_DN15201_c0_g1_i3.p1
Lophophytum-pyramidale___TRINITY_DN14236_c0_g3_i3.p1
Lophophytum-mirabile___TRINITY_DN67106_c1_g2_i3.p1_5prime_partial
Lophophytum-mirabile___TRINITY_DN67106_c1_g1_i4.p1_3prime_partial
Rhop___TRINITY_DN16047_c0_g1_i2.p1
Malania___TRINITY_DN4426_c0_g4_i3.p1
Malania___TRINITY_DN4426_c0_g4_i2.p1
Malania___TRINITY_DN4426_c0_g4_i1.p1
Vaccinium___Vadar_g31174.t1
Hydrangea___Hyque.01G029800.1.p
Solanum___Solyc10g079160.1.1
Mimulus___Migut.N01847.1.p
Lactuca___Lsat_1_v5_gn_3_75140.6
Helianthus___HanXRQChr15g0496031
Helianthus___HanXRQChr09g0270071
Daucus___DCAR_030802
Beta___EL10Ac4g09126.1
Vitis___VIT_213s0019g04630.1
Kalanchoe___Kalax.0868s0003.1.p
Kalanchoe___Kalax.0002s0043.1.p
F-Pvulgaris___Phvul.006G123100.1.p
F-Carietinum___Ca_24528
F-Prosopis___Palba_XP_028797707.1
F-Prosopis___Palba_XP_028758184.1
F-Lleucocephala___GDRZ01179297.1.p1
F-Akoa___GBYE01014714.1.p1
F-Acol___TRINITY_DN7821_c0_g1_i2.p1_complete
F-Acol___TRINITY_DN7821_c0_g1_i1.p1_complete
Manihot___Manes.07G097000.16.p
Manihot___Manes.07G095824.2.p
Manihot___Manes.07G095816.1.p
Manihot___Manes.07G095800.2.p
Manihot___Manes.07G097202.1.p
Manihot___Manes.07G095818.1.p
Manihot___Manes.07G095810.1.p
Manihot___Manes.07G095604.1.p
Manihot___Manes.07G095600.1.p
Eucalyptus___Eucgr.J01069.1.p
Quercus___Qurub.09G082900.1
Quercus___Qurub.08G231200.1
Malus___MD01G1105000
Cucumis___Cucsa.308520.1lPACid_16976368
Citrus___orange1.1g038868mlPACid_18096214
Arabidopsis-AT2G39120.1lPACid_19640369
Musa___GSMUA_Achr2P08090_001

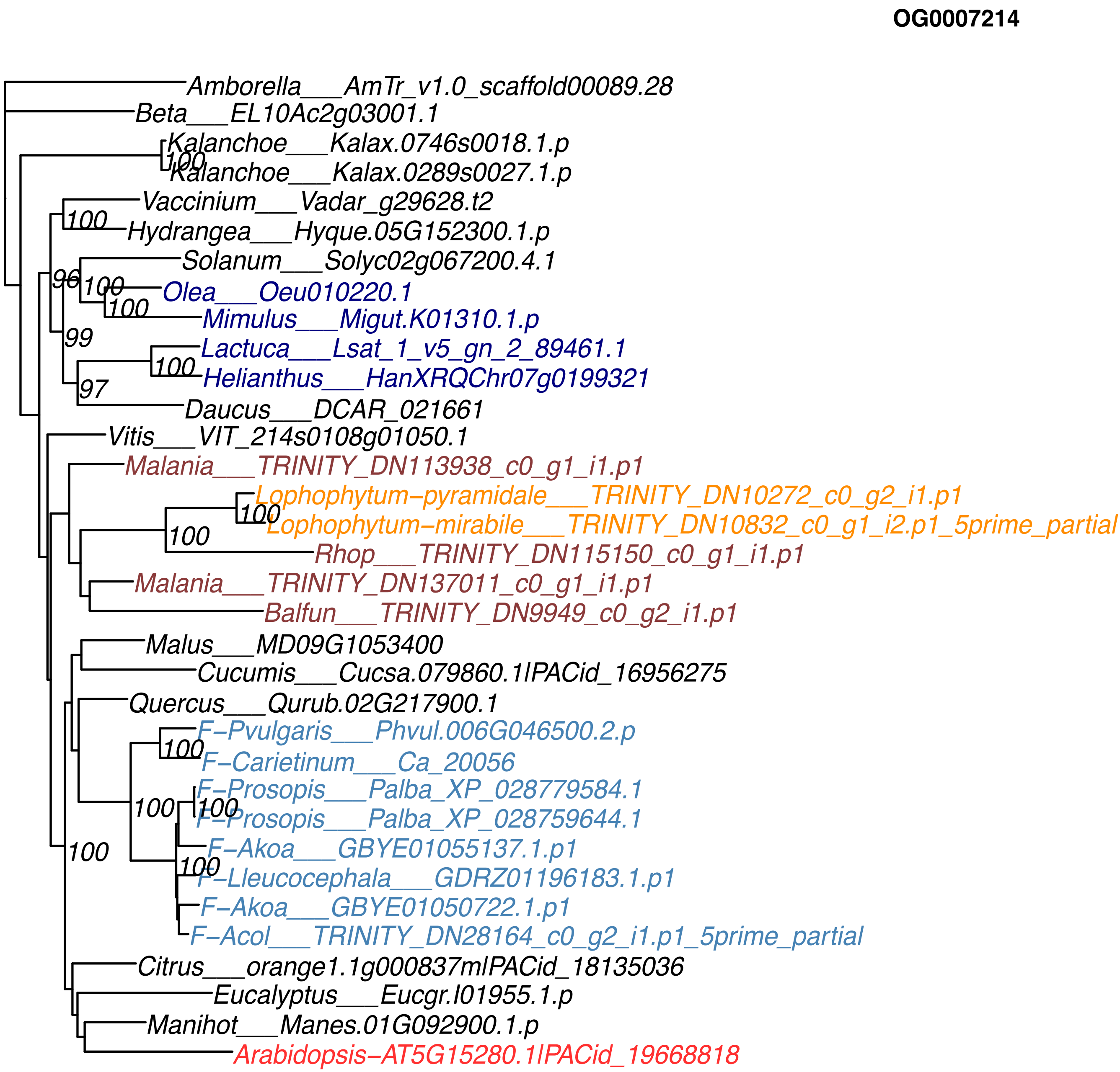
OG0007214
Amborella___AmTr_v1.0_scaffold00089.28
Beta___EL10Ac2g03001.1
Kalanchoe___Kalax.0746s0018.1.p
100
Kalanchoe___Kalax.0289s0027.1.p
Vaccinium___Vadar_g29628.t2
100
Hydrangea___Hyque.05G152300.1.p
Solanum___Solyc02g067200.4.1
96
100
Olea___Oeu010220.1
100
Mimulus___Migut.K01310.1.p
99
Lactuca___Lsat_1_v5_gn_2_89461.1
100
Helianthus___HanXRQChr07g0199321
97
Daucus___DCAR_021661
Vitis___VIT_214s0108g01050.1
Malania___TRINITY_DN113938_c0_g1_i1.p1
Lophophytum−pyramidale___TRINITY_DN10272_c0_g2_i1.p1
100
Lophophytum−mirabile___TRINITY_DN10832_c0_g1_i2.p1_5prime_partial
100
Rhop___TRINITY_DN115150_c0_g1_i1.p1
Malania___TRINITY_DN137011_c0_g1_i1.p1
Balfun___TRINITY_DN9949_c0_g2_i1.p1
Malus___MD09G1053400
Cucumis___Cucsa.079860.1|PACid_16956275
Quercus___Qurub.02G217900.1
F−Pvulgaris___Phvul.006G046500.2.p
100
F−Carietinum___Ca_20056
F−Prosopis___Palba_XP_028779584.1
100
100
F−Prosopis___Palba_XP_028759644.1
F−Akoa___GBYE01055137.1.p1
100
100
F−Lleucocephala___GDRZ01196183.1.p1
F−Akoa___GBYE01050722.1.p1
F−Acol___TRINITY_DN28164_c0_g2_i1.p1_5prime_partial
Citrus___orange1.1g000837m|PACid_18135036
Eucalyptus___Eucgr.I01955.1.p
Manihot___Manes.01G092900.1.p
Arabidopsis−AT5G15280.1|PACid_19668818

**OG0007310**

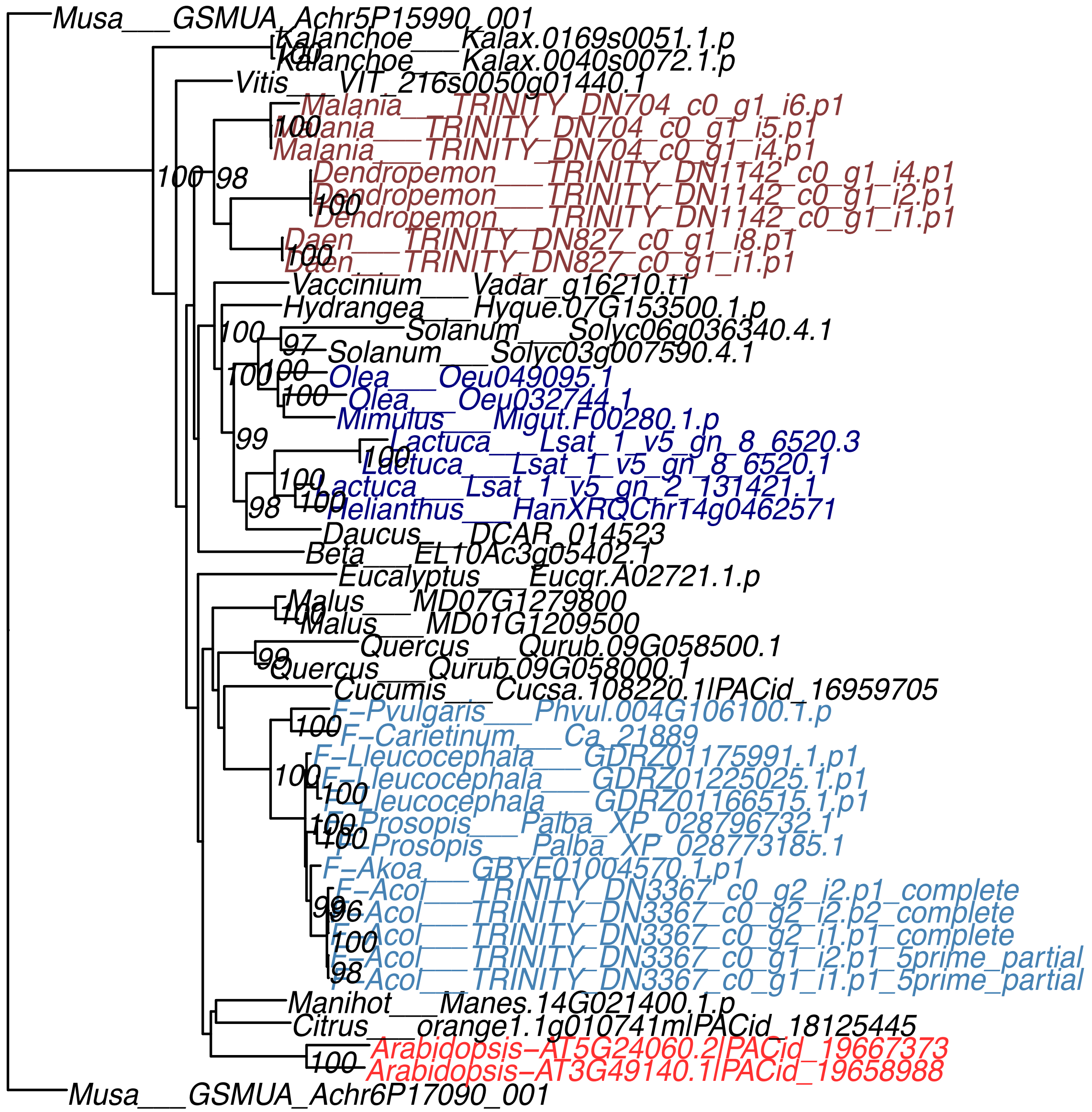

Musa___GSMUA_Achr5P15990_001
Kalanchoe___Kalax.0169s0051.1.p
Kalanchoe___Kalax.0040s0072.1.p
Vitis___VIT_216s0050g01440.1
Malania___TRINITY_DN704_c0_g1_i6.p1
Malania___TRINITY_DN704_c0_g1_i5.p1
Malania___TRINITY_DN704_c0_g1_i4.p1
Dendropemon___TRINITY_DN1142_c0_g1_i4.p1
Dendropemon___TRINITY_DN1142_c0_g1_i2.p1
Dendropemon___TRINITY_DN1142_c0_g1_i1.p1
Daen___TRINITY_DN827_c0_g1_i8.p1
Daen___TRINITY_DN827_c0_g1_i1.p1
Vaccinium___Vadar_g16210.t1
Hydrangea___Hyque.07G153500.1.p
Solanum___Solyc06g036340.4.1
Solanum___Solyc03g007590.4.1
Olea___Oeu049095.1
Olea___Oeu032744.1
Mimulus___Migut.F00280.1.p
Lactuca___Lsat_1_v5_gn_8_6520.3
Lactuca___Lsat_1_v5_gn_8_6520.1
Lactuca___Lsat_1_v5_gn_2_131421.1
Helianthus___HanXRQChr14g0462571
Daucus___DCAR_014523
Beta___EL10Ac3g05402.1
Eucalyptus___Eucgr.A02721.1.p
Malus___MD07G1279800
Malus___MD01G1209500
Quercus___Qurub.09G058500.1
Quercus___Qurub.09G058000.1
Cucumis___Cucsa.108220.1|PACid_16959705
F–Pvulgaris___Phvul.004G106100.1.p
F–Carietinum___Ca_21889
F–Lleucocephala___GDRZ01175991.1.p1
F–Lleucocephala___GDRZ01225025.1.p1
F–Lleucocephala___GDRZ01166515.1.p1
F–Prosopis___Palba_XP_028796732.1
F–Prosopis___Palba_XP_028773185.1
F–Akoa___GBYE01004570.1.p1
F–Acol___TRINITY_DN3367_c0_g2_i2.p1_complete
F–Acol___TRINITY_DN3367_c0_g2_i2.p2_complete
F–Acol___TRINITY_DN3367_c0_g2_i1.p1_complete
F–Acol___TRINITY_DN3367_c0_g1_i2.p1_5prime_partial
F–Acol___TRINITY_DN3367_c0_g1_i1.p1_5prime_partial
Manihot___Manes.14G021400.1.p
Citrus___orange1.1g010741m|PACid_18125445
Arabidopsis–AT5G24060.2|PACid_19667373
Arabidopsis–AT3G49140.1|PACid_19658988
Musa___GSMUA_Achr6P17090_001

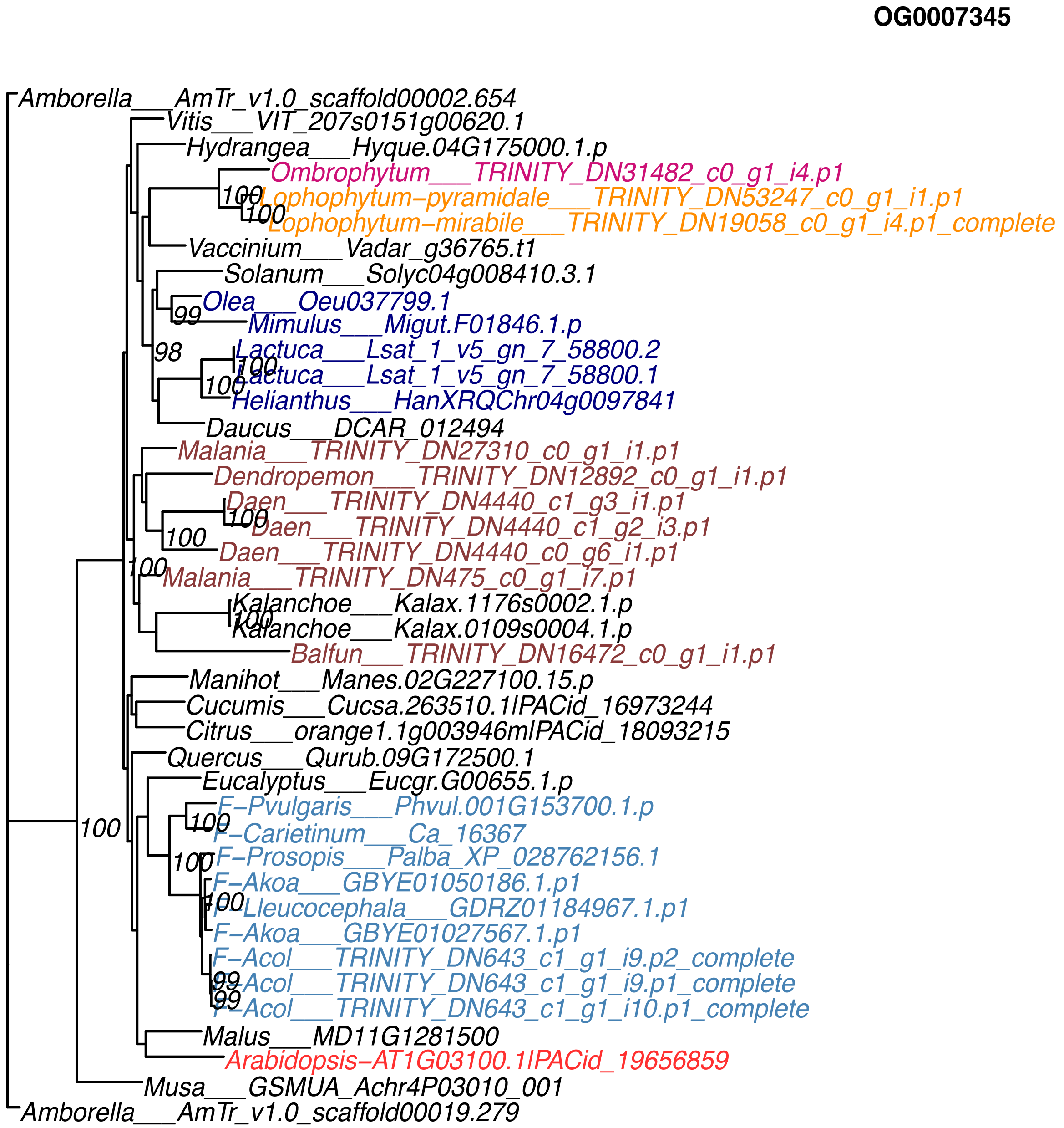

OG0007345
Amborella___AmTr_v1.0_scaffold00002.654
Vitis___VIT_207s0151g00620.1
Hydrangea___Hyque.04G175000.1.p
Ombrophytum___TRINITY_DN31482_c0_g1_i4.p1
Lophophytum−pyramidale___TRINITY_DN53247_c0_g1_i1.p1
Lophophytum−mirabile___TRINITY_DN19058_c0_g1_i4.p1_complete
Vaccinium___Vadar_g36765.t1
Solanum___Solyc04g008410.3.1
Olea___Oeu037799.1
Mimulus___Migut.F01846.1.p
Lactuca___Lsat_1_v5_gn_7_58800.2
Lactuca___Lsat_1_v5_gn_7_58800.1
Helianthus___HanXRQChr04g0097841
Daucus___DCAR_012494
Malania___TRINITY_DN27310_c0_g1_i1.p1
Dendropemon___TRINITY_DN12892_c0_g1_i1.p1
Daen___TRINITY_DN4440_c1_g3_i1.p1
Daen___TRINITY_DN4440_c1_g2_i3.p1
Daen___TRINITY_DN4440_c0_g6_i1.p1
Malania___TRINITY_DN475_c0_g1_i7.p1
Kalanchoe___Kalax.1176s0002.1.p
Kalanchoe___Kalax.0109s0004.1.p
Balfun___TRINITY_DN16472_c0_g1_i1.p1
Manihot___Manes.02G227100.15.p
Cucumis___Cucsa.263510.1|PACid_16973244
Citrus___orange1.1g003946m|PACid_18093215
Quercus___Qurub.09G172500.1
Eucalyptus___Eucgr.G00655.1.p
F−Pvulgaris___Phvul.001G153700.1.p
F−Carietinum___Ca_16367
F−Prosopis___Palba_XP_028762156.1
F−Akoa___GBYE01050186.1.p1
F−Lleucocephala___GDRZ01184967.1.p1
F−Akoa___GBYE01027567.1.p1
F−Acol___TRINITY_DN643_c1_g1_i9.p2_complete
F−Acol___TRINITY_DN643_c1_g1_i9.p1_complete
F−Acol___TRINITY_DN643_c1_g1_i10.p1_complete
Malus___MD11G1281500
Arabidopsis−AT1G03100.1|PACid_19656859
Musa___GSMUA_Achr4P03010_001
Amborella___AmTr_v1.0_scaffold00019.279
100
100
99
98
100
100
100
100
100
100
100
100
100
100
99
99

**OG0007355**

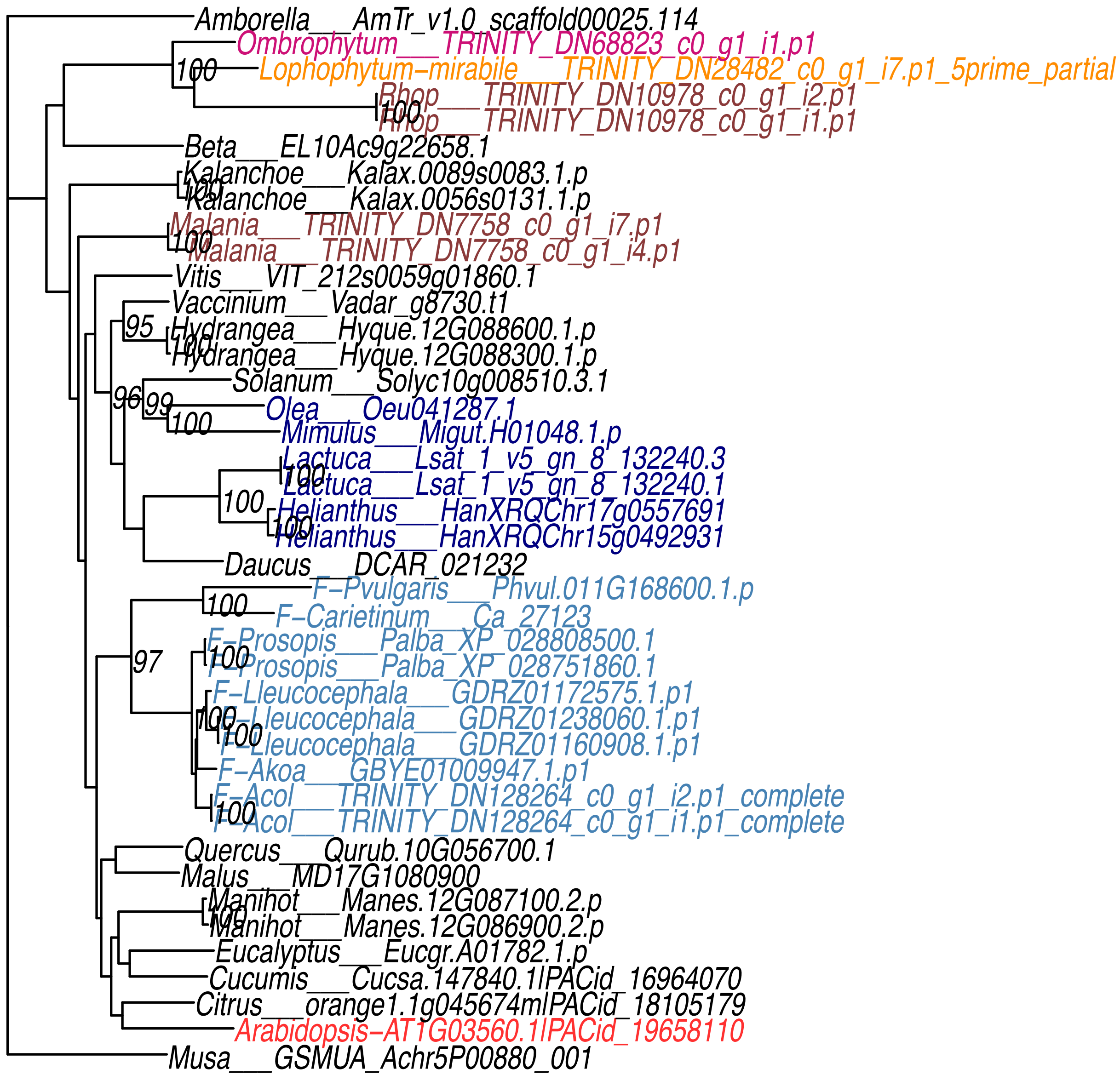

Amborella___AmTr_v1.0_scaffold00025.114
Ombrophytum___TRINITY_DN68823_c0_g1_i1.p1
Lophophytum-mirabile___TRINITY_DN28482_c0_g1_i7.p1_5prime_partial
Rhop___TRINITY_DN10978_c0_g1_i2.p1
Rhop___TRINITY_DN10978_c0_g1_i1.p1
Beta___EL10Ac9g22658.1
Kalanchoe___Kalax.0089s0083.1.p
Kalanchoe___Kalax.0056s0131.1.p
Malania___TRINITY_DN7758_c0_g1_i7.p1
Malania___TRINITY_DN7758_c0_g1_i4.p1
Vitis___VIT_212s0059g01860.1
Vaccinium___Vadar_g8730.t1
Hydrangea___Hyque.12G088600.1.p
Hydrangea___Hyque.12G088300.1.p
Solanum___Solyc10g008510.3.1
Olea___Oeu041287.1
Mimulus___Migut.H01048.1.p
Lactuca___Lsat_1_v5_gn_8_132240.3
Lactuca___Lsat_1_v5_gn_8_132240.1
Helianthus___HanXRQChr17g0557691
Helianthus___HanXRQChr15g0492931
Daucus___DCAR_021232
F−Pvulgaris___Phvul.011G168600.1.p
F−Carietinum___Ca_27123
F−Prosopis___Palba_XP_028808500.1
F−Prosopis___Palba_XP_028751860.1
F−Lleucocephala___GDRZ01172575.1.p1
F−Lleucocephala___GDRZ01238060.1.p1
F−Lleucocephala___GDRZ01160908.1.p1
F−Akoa___GBYE01009947.1.p1
F−Acol___TRINITY_DN128264_c0_g1_i2.p1_complete
F−Acol___TRINITY_DN128264_c0_g1_i1.p1_complete
Quercus___Qurub.10G056700.1
Malus___MD17G1080900
Manihot___Manes.12G087100.2.p
Manihot___Manes.12G086900.2.p
Eucalyptus___Eucgr.A01782.1.p
Cucumis___Cucsa.147840.1|PACid_16964070
Citrus___orange1.1g045674m|PACid_18105179
Arabidopsis−AT1G03560.1|PACid_19658110
Musa___GSMUA_Achr5P00880_001
100
100
100
100
95
100
96
99
100
100
100
100
100
97
100
100
100
100
100

**OG0007387**

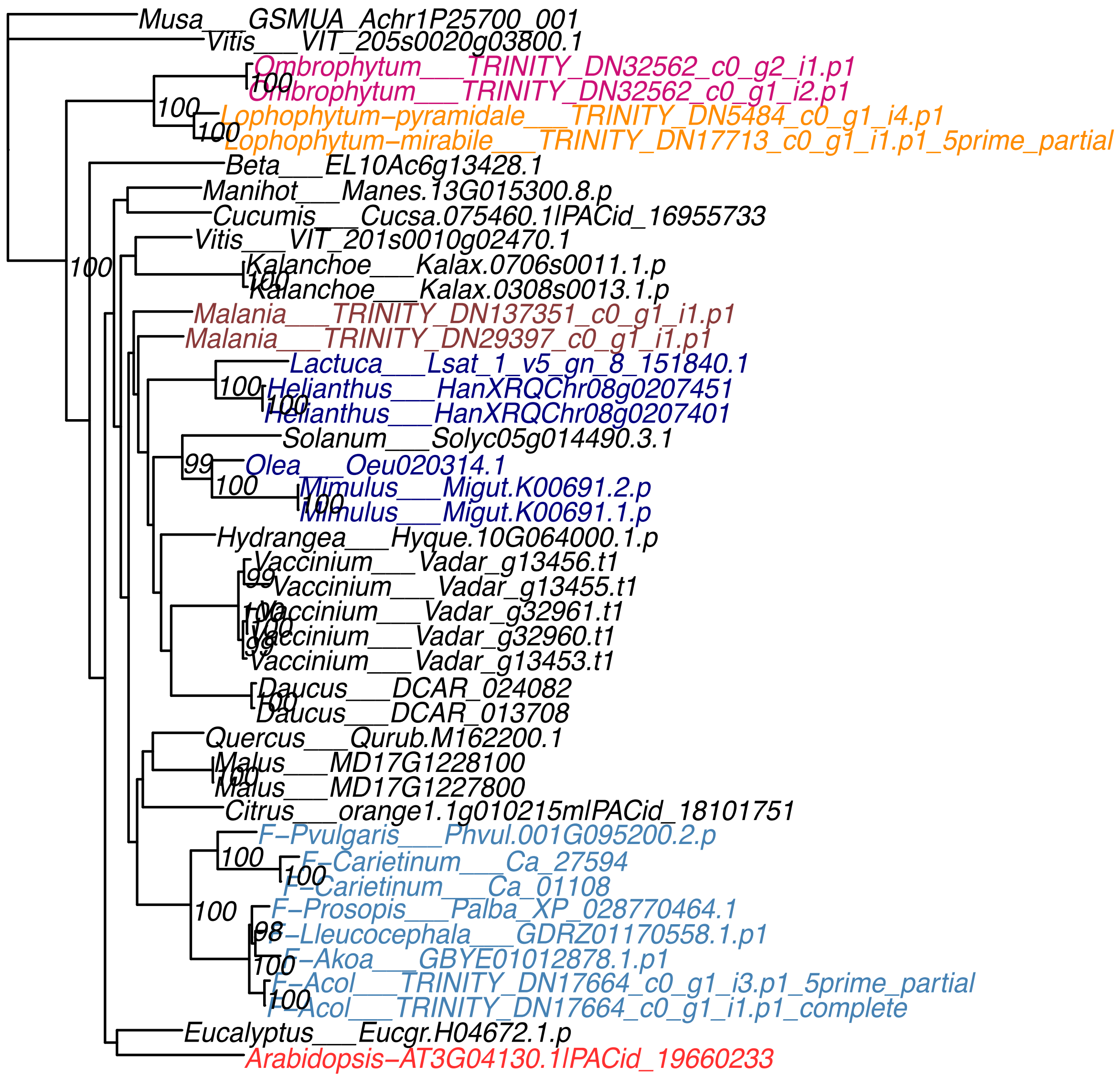

Musa___GSMUA_Achr1P25700_001
Vitis___VIT_205s0020g03800.1
Ombrophytum___TRINITY_DN32562_c0_g2_i1.p1
Ombrophytum___TRINITY_DN32562_c0_g1_i2.p1
Lophophytum–pyramidale___TRINITY_DN5484_c0_g1_i4.p1
Lophophytum–mirabile___TRINITY_DN17713_c0_g1_i1.p1_5prime_partial
Beta___EL10Ac6g13428.1
Manihot___Manes.13G015300.8.p
Cucumis___Cucsa.075460.1|PACid_16955733
Vitis___VIT_201s0010g02470.1
Kalanchoe___Kalax.0706s0011.1.p
Kalanchoe___Kalax.0308s0013.1.p
Malania___TRINITY_DN137351_c0_g1_i1.p1
Malania___TRINITY_DN29397_c0_g1_i1.p1
Lactuca___Lsat_1_v5_gn_8_151840.1
Helianthus___HanXRQChr08g0207451
Helianthus___HanXRQChr08g0207401
Solanum___Solyc05g014490.3.1
Olea___Oeu020314.1
Mimulus___Migut.K00691.2.p
Mimulus___Migut.K00691.1.p
Hydrangea___Hyque.10G064000.1.p
Vaccinium___Vadar_g13456.t1
Vaccinium___Vadar_g13455.t1
Vaccinium___Vadar_g32961.t1
Vaccinium___Vadar_g32960.t1
Vaccinium___Vadar_g13453.t1
Daucus___DCAR_024082
Daucus___DCAR_013708
Quercus___Qurub.M162200.1
Malus___MD17G1228100
Malus___MD17G1227800
Citrus___orange1.1g010215m|PACid_18101751
F–Pvulgaris___Phvul.001G095200.2.p
F–Carietinum___Ca_27594
F–Carietinum___Ca_01108
F–Prosopis___Palba_XP_028770464.1
F–Lleucocephala___GDRZ01170558.1.p1
F–Akoa___GBYE01012878.1.p1
F–Acol___TRINITY_DN17664_c0_g1_i3.p1_5prime_partial
F–Acol___TRINITY_DN17664_c0_g1_i1.p1_complete
Eucalyptus___Eucgr.H04672.1.p
Arabidopsis–AT3G04130.1|PACid_19660233

**OG0007394**

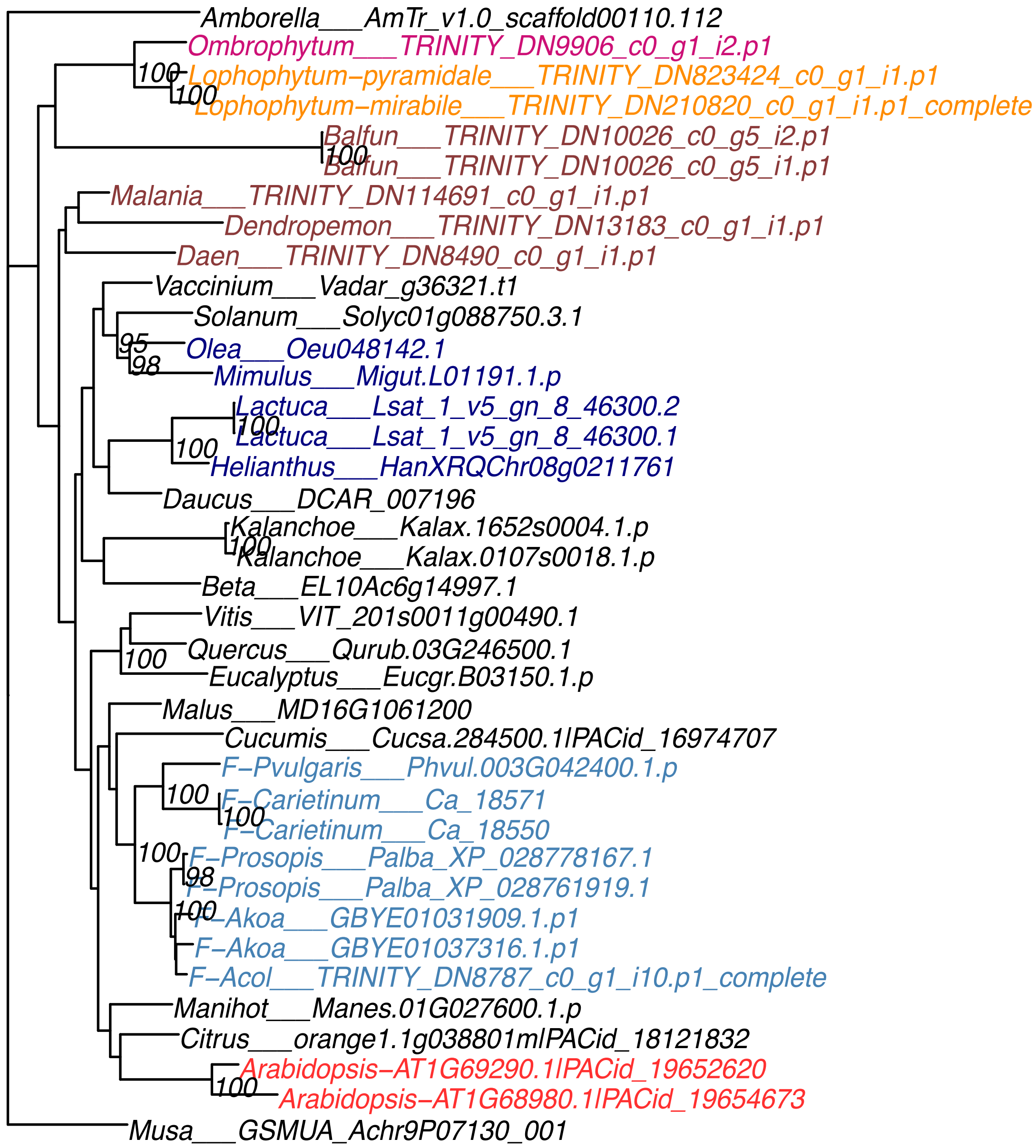

Amborella___AmTr_v1.0_scaffold00110.112
Ombrophytum___TRINITY_DN9906_c0_g1_i2.p1
100
Lophophytum−pyramidale___TRINITY_DN823424_c0_g1_i1.p1
100
Lophophytum−mirabile___TRINITY_DN210820_c0_g1_i1.p1_complete
Balfun___TRINITY_DN10026_c0_g5_i2.p1
100
Balfun___TRINITY_DN10026_c0_g5_i1.p1
Malania___TRINITY_DN114691_c0_g1_i1.p1
Dendropemon___TRINITY_DN13183_c0_g1_i1.p1
Daen___TRINITY_DN8490_c0_g1_i1.p1
Vaccinium___Vadar_g36321.t1
Solanum___Solyc01g088750.3.1
95
Olea___Oeu048142.1
98
Mimulus___Migut.L01191.1.p
Lactuca___Lsat_1_v5_gn_8_46300.2
100
Lactuca___Lsat_1_v5_gn_8_46300.1
100
Helianthus___HanXRQChr08g0211761
Daucus___DCAR_007196
Kalanchoe___Kalax.1652s0004.1.p
100
Kalanchoe___Kalax.0107s0018.1.p
Beta___EL10Ac6g14997.1
Vitis___VIT_201s0011g00490.1
Quercus___Qurub.03G246500.1
100
Eucalyptus___Eucgr.B03150.1.p
Malus___MD16G1061200
Cucumis___Cucsa.284500.1|PACid_16974707
F−Pvulgaris___Phvul.003G042400.1.p
100
F−Carietinum___Ca_18571
100
F−Carietinum___Ca_18550
100
F−Prosopis___Palba_XP_028778167.1
98
F−Prosopis___Palba_XP_028761919.1
100
F−Akoa___GBYE01031909.1.p1
F−Akoa___GBYE01037316.1.p1
F−Acol___TRINITY_DN8787_c0_g1_i10.p1_complete
Manihot___Manes.01G027600.1.p
Citrus___orange1.1g038801m|PACid_18121832
Arabidopsis−AT1G69290.1|PACid_19652620
100
Arabidopsis−AT1G68980.1|PACid_19654673
Musa___GSMUA_Achr9P07130_001

**OG0007494**

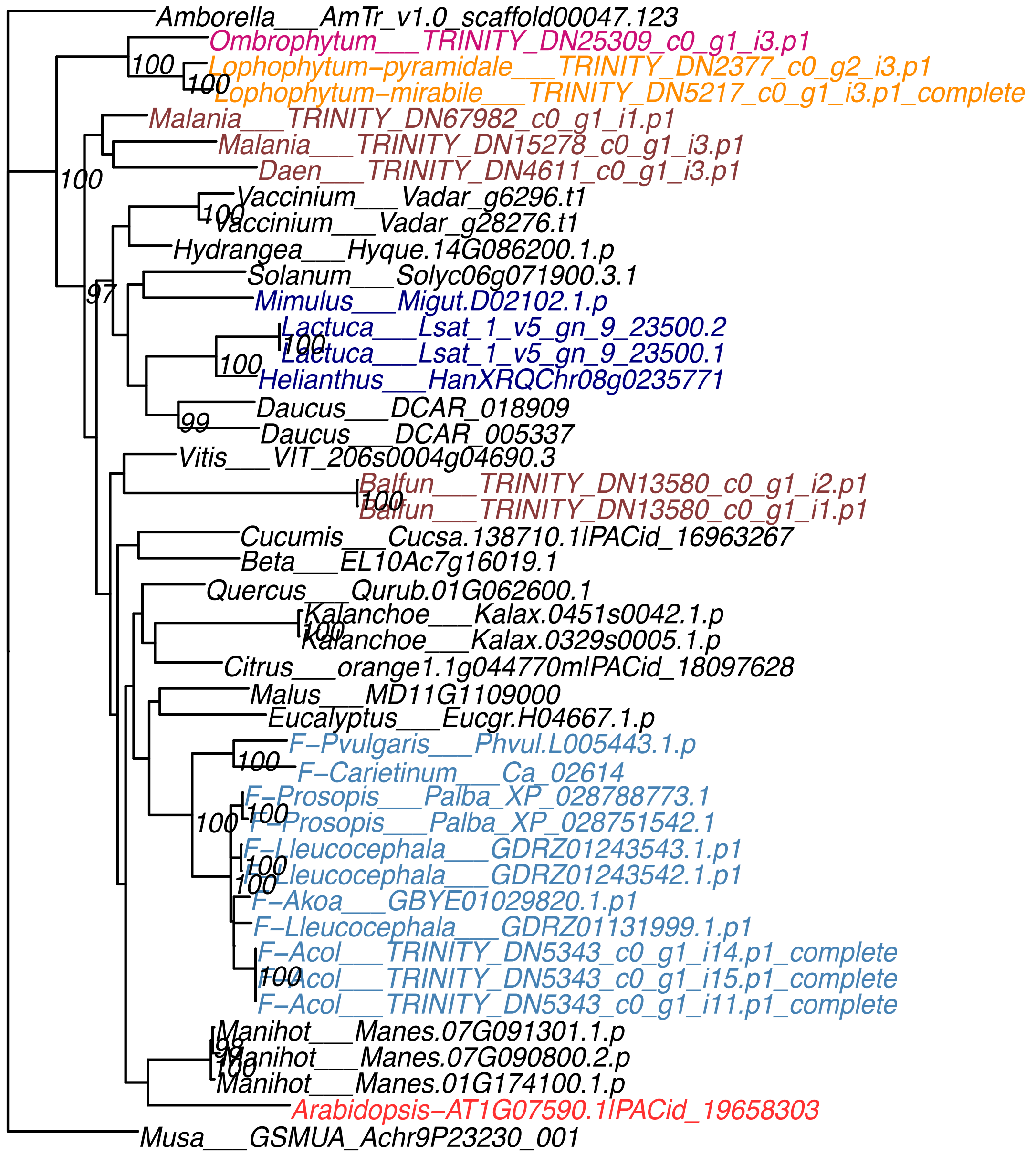

Amborella___AmTr_v1.0_scaffold00047.123
Ombrophytum___TRINITY_DN25309_c0_g1_i3.p1
100
Lophophytum-pyramidale___TRINITY_DN2377_c0_g2_i3.p1
100
Lophophytum-mirabile___TRINITY_DN5217_c0_g1_i3.p1_complete
Malania___TRINITY_DN67982_c0_g1_i1.p1
Malania___TRINITY_DN15278_c0_g1_i3.p1
100
Daen___TRINITY_DN4611_c0_g1_i3.p1
Vaccinium___Vadar_g6296.t1
100
Vaccinium___Vadar_g28276.t1
Hydrangea___Hyque.14G086200.1.p
Solanum___Solyc06g071900.3.1
97
Mimulus___Migut.D02102.1.p
Lactuca___Lsat_1_v5_gn_9_23500.2
100
Lactuca___Lsat_1_v5_gn_9_23500.1
100
Helianthus___HanXRQChr08g0235771
Daucus___DCAR_018909
99
Daucus___DCAR_005337
Vitis___VIT_206s0004g04690.3
Balfun___TRINITY_DN13580_c0_g1_i2.p1
100
Balfun___TRINITY_DN13580_c0_g1_i1.p1
Cucumis___Cucsa.138710.1lPACid_16963267
Beta___EL10Ac7g16019.1
Quercus___Qurub.01G062600.1
Kalanchoe___Kalax.0451s0042.1.p
100
Kalanchoe___Kalax.0329s0005.1.p
Citrus___orange1.1g044770mlPACid_18097628
Malus___MD11G1109000
Eucalyptus___Eucgr.H04667.1.p
F-Pvulgaris___Phvul.L005443.1.p
100
F-Carietinum___Ca_02614
F-Prosopis___Palba_XP_028788773.1
100
100
F-Prosopis___Palba_XP_028751542.1
F-Lleucocephala___GDRZ01243543.1.p1
100
F-Lleucocephala___GDRZ01243542.1.p1
100
F-Akoa___GBYE01029820.1.p1
F-Lleucocephala___GDRZ01131999.1.p1
F-Acol___TRINITY_DN5343_c0_g1_i14.p1_complete
100
F-Acol___TRINITY_DN5343_c0_g1_i15.p1_complete
F-Acol___TRINITY_DN5343_c0_g1_i11.p1_complete
Manihot___Manes.07G091301.1.p
98
Manihot___Manes.07G090800.2.p
100
Manihot___Manes.01G174100.1.p
Arabidopsis-AT1G07590.1lPACid_19658303
Musa___GSMUA_Achr9P23230_001

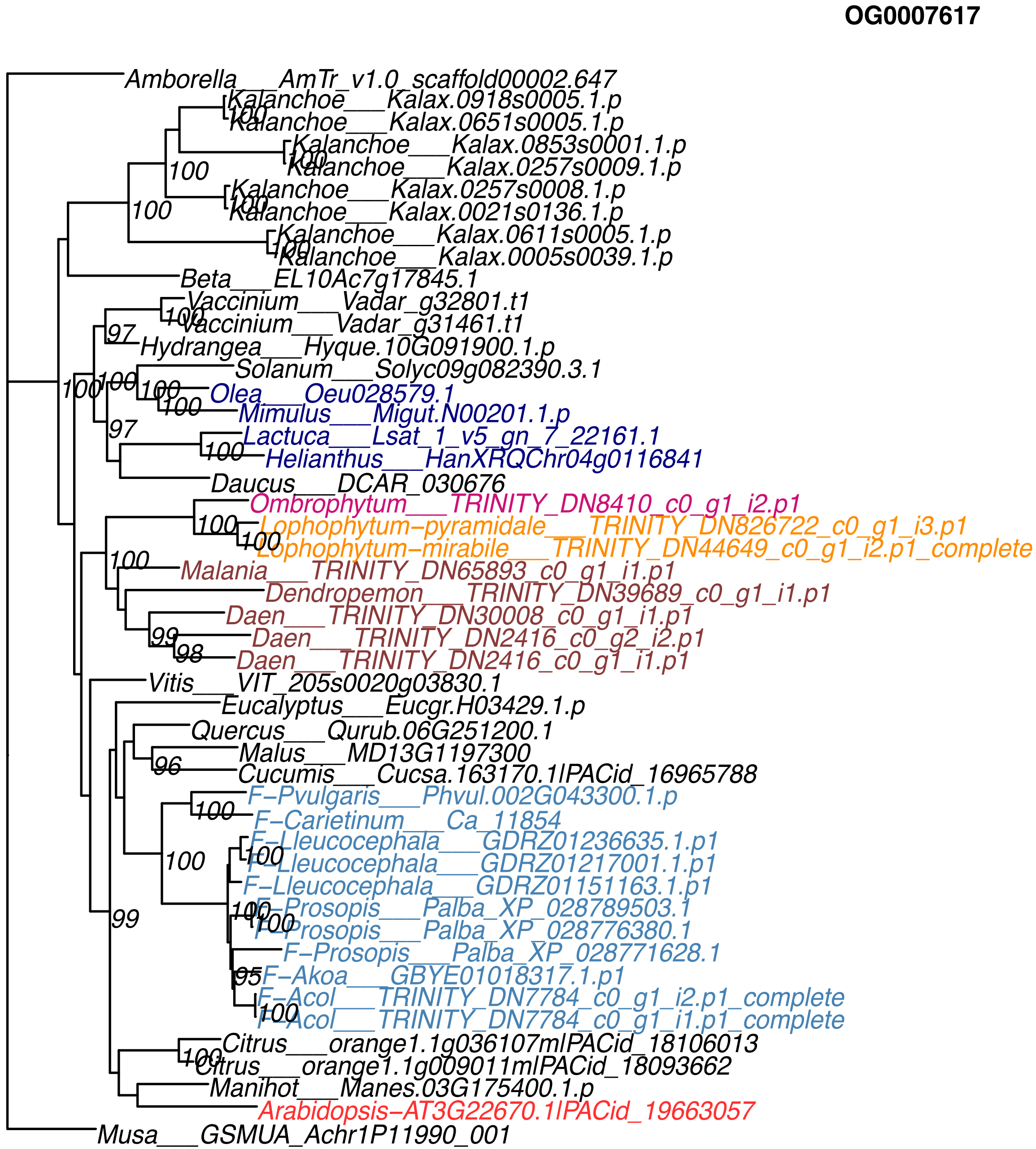
OG0007617
Amborella___AmTr_v1.0_scaffold00002.647
Kalanchoe___Kalax.0918s0005.1.p
Kalanchoe___Kalax.0651s0005.1.p
Kalanchoe___Kalax.0853s0001.1.p
Kalanchoe___Kalax.0257s0009.1.p
Kalanchoe___Kalax.0257s0008.1.p
Kalanchoe___Kalax.0021s0136.1.p
Kalanchoe___Kalax.0611s0005.1.p
Kalanchoe___Kalax.0005s0039.1.p
Beta___EL10Ac7g17845.1
Vaccinium___Vadar_g32801.t1
Vaccinium___Vadar_g31461.t1
Hydrangea___Hyque.10G091900.1.p
Solanum___Solyc09g082390.3.1
Olea___Oeu028579.1
Mimulus___Migut.N00201.1.p
Lactuca___Lsat_1_v5_gn_7_22161.1
Helianthus___HanXRQChr04g0116841
Daucus___DCAR_030676
Ombrophytum___TRINITY_DN8410_c0_g1_i2.p1
Lophophytum−pyramidale___TRINITY_DN826722_c0_g1_i3.p1
Lophophytum−mirabile___TRINITY_DN44649_c0_g1_i2.p1_complete
Malania___TRINITY_DN65893_c0_g1_i1.p1
Dendropemon___TRINITY_DN39689_c0_g1_i1.p1
Daen___TRINITY_DN30008_c0_g1_i1.p1
Daen___TRINITY_DN2416_c0_g2_i2.p1
Daen___TRINITY_DN2416_c0_g1_i1.p1
Vitis___VIT_205s0020g03830.1
Eucalyptus___Eucgr.H03429.1.p
Quercus___Qurub.06G251200.1
Malus___MD13G1197300
Cucumis___Cucsa.163170.1|PACid_16965788
F−Pvulgaris___Phvul.002G043300.1.p
F−Carietinum___Ca_11854
F−Lleucocephala___GDRZ01236635.1.p1
F−Lleucocephala___GDRZ01217001.1.p1
F−Lleucocephala___GDRZ01151163.1.p1
F−Prosopis___Palba_XP_028789503.1
F−Prosopis___Palba_XP_028776380.1
F−Prosopis___Palba_XP_028771628.1
F−Akoa___GBYE01018317.1.p1
F−Acol___TRINITY_DN7784_c0_g1_i2.p1_complete
F−Acol___TRINITY_DN7784_c0_g1_i1.p1_complete
Citrus___orange1.1g036107m|PACid_18106013
Citrus___orange1.1g009011m|PACid_18093662
Manihot___Manes.03G175400.1.p
Arabidopsis−AT3G22670.1|PACid_19663057
Musa___GSMUA_Achr1P11990_001

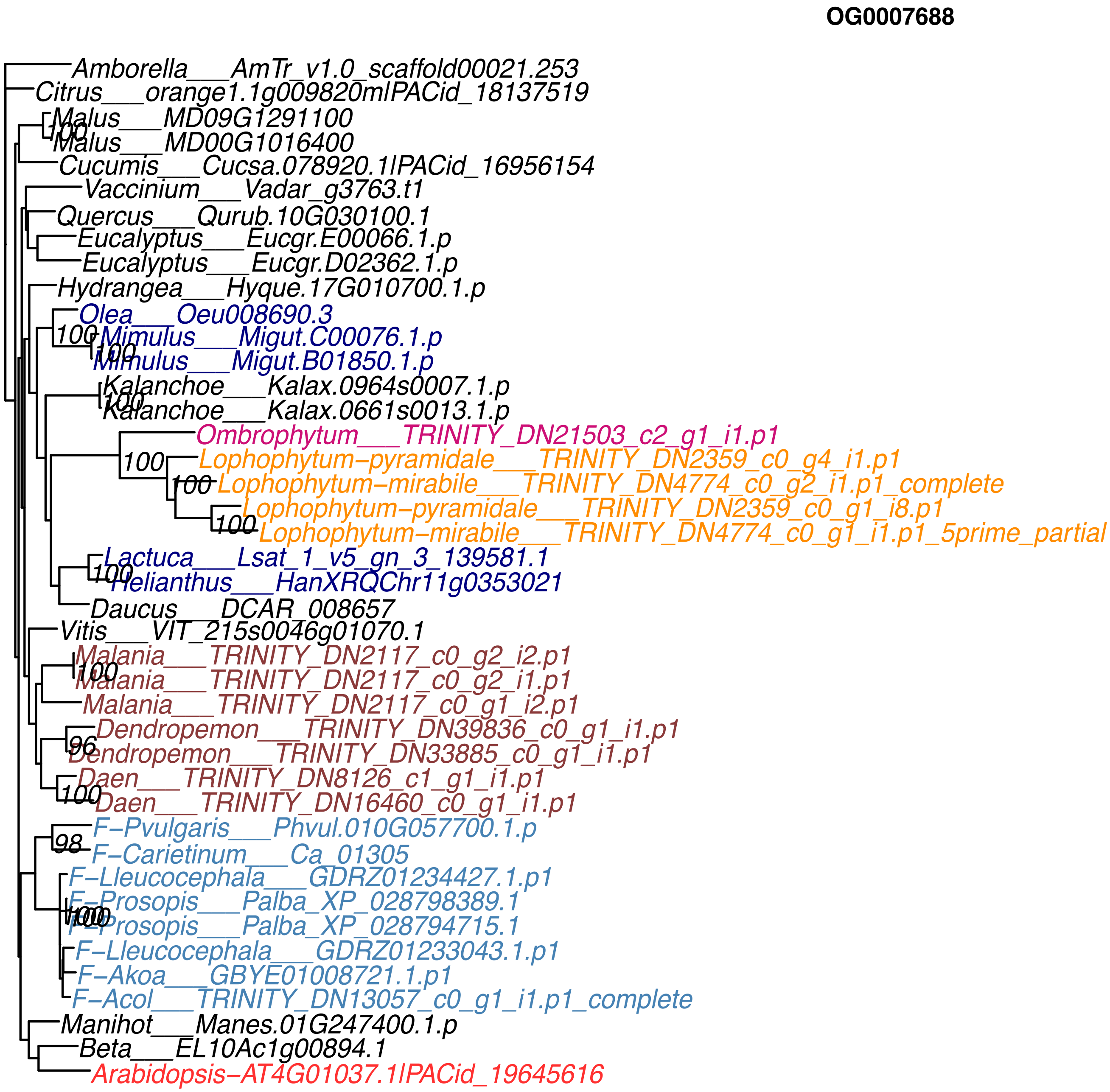

OG0007688
Amborella___AmTr_v1.0_scaffold00021.253
Citrus___orange1.1g009820mIPACid_18137519
Malus___MD09G1291100
100
Malus___MD00G1016400
Cucumis___Cucsa.078920.1IPACid_16956154
Vaccinium___Vadar_g3763.t1
Quercus___Qurub.10G030100.1
Eucalyptus___Eucgr.E00066.1.p
Eucalyptus___Eucgr.D02362.1.p
Hydrangea___Hyque.17G010700.1.p
Olea___Oeu008690.3
100
Mimulus___Migut.C00076.1.p
100
Mimulus___Migut.B01850.1.p
Kalanchoe___Kalax.0964s0007.1.p
100
Kalanchoe___Kalax.0661s0013.1.p
Ombrophytum___TRINITY_DN21503_c2_g1_i1.p1
100
Lophophytum−pyramidale___TRINITY_DN2359_c0_g4_i1.p1
100
Lophophytum−mirabile___TRINITY_DN4774_c0_g2_i1.p1_complete
Lophophytum−pyramidale___TRINITY_DN2359_c0_g1_i8.p1
100
Lophophytum−mirabile___TRINITY_DN4774_c0_g1_i1.p1_5prime_partial
Lactuca___Lsat_1_v5_gn_3_139581.1
100
Helianthus___HanXRQChr11g0353021
Daucus___DCAR_008657
Vitis___VIT_215s0046g01070.1
Malania___TRINITY_DN2117_c0_g2_i2.p1
100
Malania___TRINITY_DN2117_c0_g2_i1.p1
Malania___TRINITY_DN2117_c0_g1_i2.p1
Dendropemon___TRINITY_DN39836_c0_g1_i1.p1
96
Dendropemon___TRINITY_DN33885_c0_g1_i1.p1
Daen___TRINITY_DN8126_c1_g1_i1.p1
100
Daen___TRINITY_DN16460_c0_g1_i1.p1
F−Pvulgaris___Phvul.010G057700.1.p
98
F−Carietinum___Ca_01305
F−Lleucocephala___GDRZ01234427.1.p1
F−Prosopis___Palba_XP_028798389.1
100
100
F−Prosopis___Palba_XP_028794715.1
F−Lleucocephala___GDRZ01233043.1.p1
F−Akoa___GBYE01008721.1.p1
F−Acol___TRINITY_DN13057_c0_g1_i1.p1_complete
Manihot___Manes.01G247400.1.p
Beta___EL10Ac1g00894.1
Arabidopsis−AT4G01037.1IPACid_19645616

## OG0007693

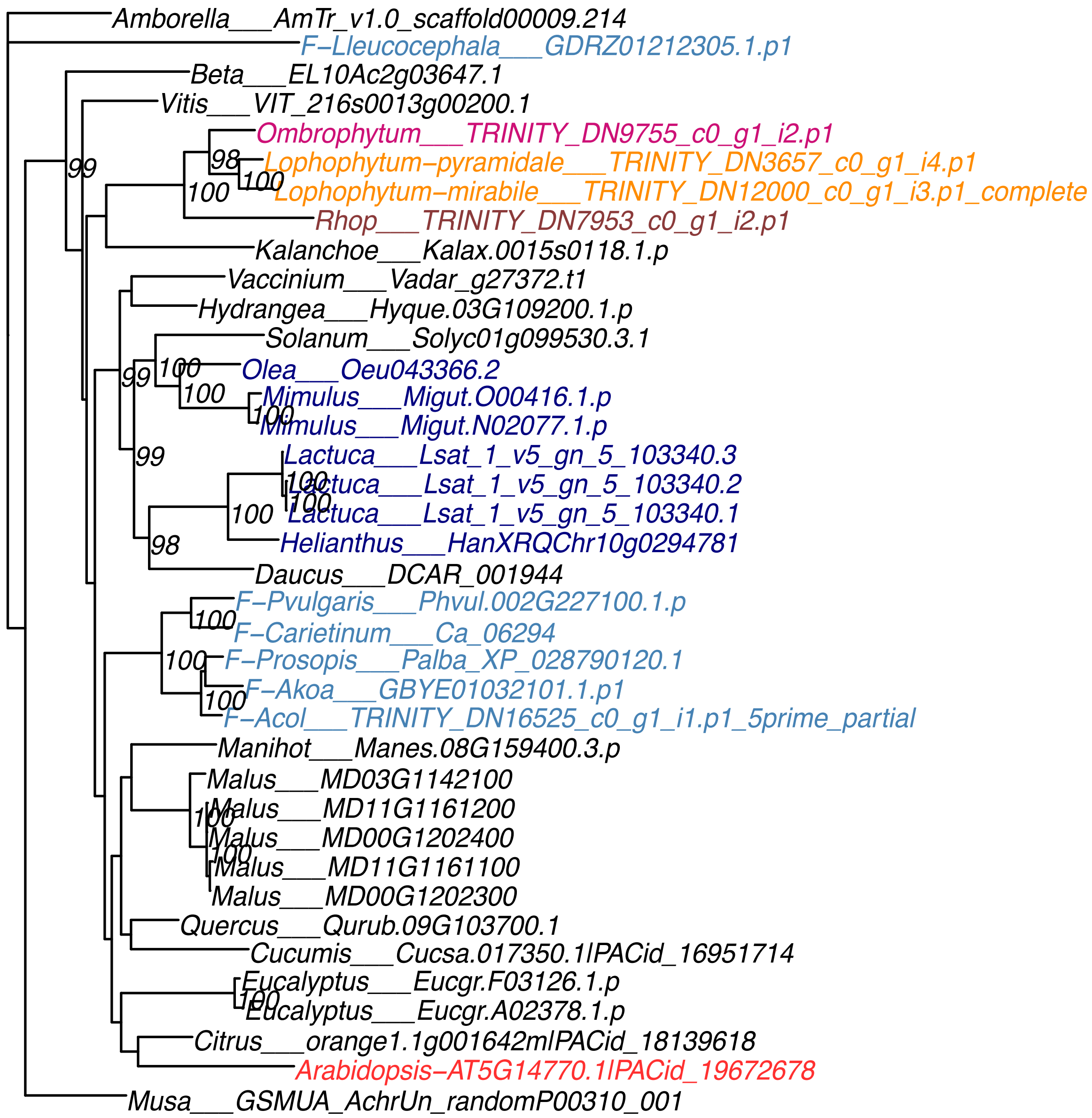
Amborella___AmTr_v1.0_scaffold00009.214
F−Lleucocephala___GDRZ01212305.1.p1
Beta___EL10Ac2g03647.1
Vitis___VIT_216s0013g00200.1
Ombrophytum___TRINITY_DN9755_c0_g1_i2.p1
Lophophytum−pyramidale___TRINITY_DN3657_c0_g1_i4.p1
Lophophytum−mirabile___TRINITY_DN12000_c0_g1_i3.p1_complete
Rhop___TRINITY_DN7953_c0_g1_i2.p1
Kalanchoe___Kalax.0015s0118.1.p
Vaccinium___Vadar_g27372.t1
Hydrangea___Hyque.03G109200.1.p
Solanum___Solyc01g099530.3.1
Olea___Oeu043366.2
Mimulus___Migut.O00416.1.p
Mimulus___Migut.N02077.1.p
Lactuca___Lsat_1_v5_gn_5_103340.3
Lactuca___Lsat_1_v5_gn_5_103340.2
Lactuca___Lsat_1_v5_gn_5_103340.1
Helianthus___HanXRQChr10g0294781
Daucus___DCAR_001944
F−Pvulgaris___Phvul.002G227100.1.p
F−Carietinum___Ca_06294
F−Prosopis___Palba_XP_028790120.1
F−Akoa___GBYE01032101.1.p1
F−Acol___TRINITY_DN16525_c0_g1_i1.p1_5prime_partial
Manihot___Manes.08G159400.3.p
Malus___MD03G1142100
Malus___MD11G1161200
Malus___MD00G1202400
Malus___MD11G1161100
Malus___MD00G1202300
Quercus___Qurub.09G103700.1
Cucumis___Cucsa.017350.1|PACid_16951714
Eucalyptus___Eucgr.F03126.1.p
Eucalyptus___Eucgr.A02378.1.p
Citrus___orange1.1g001642m|PACid_18139618
Arabidopsis−AT5G14770.1|PACid_19672678
Musa___GSMUA_AchrUn_randomP00310_001
99
98
100
100
100
100
99
100
99
100
100
100
98
100
100
100
100
100
100

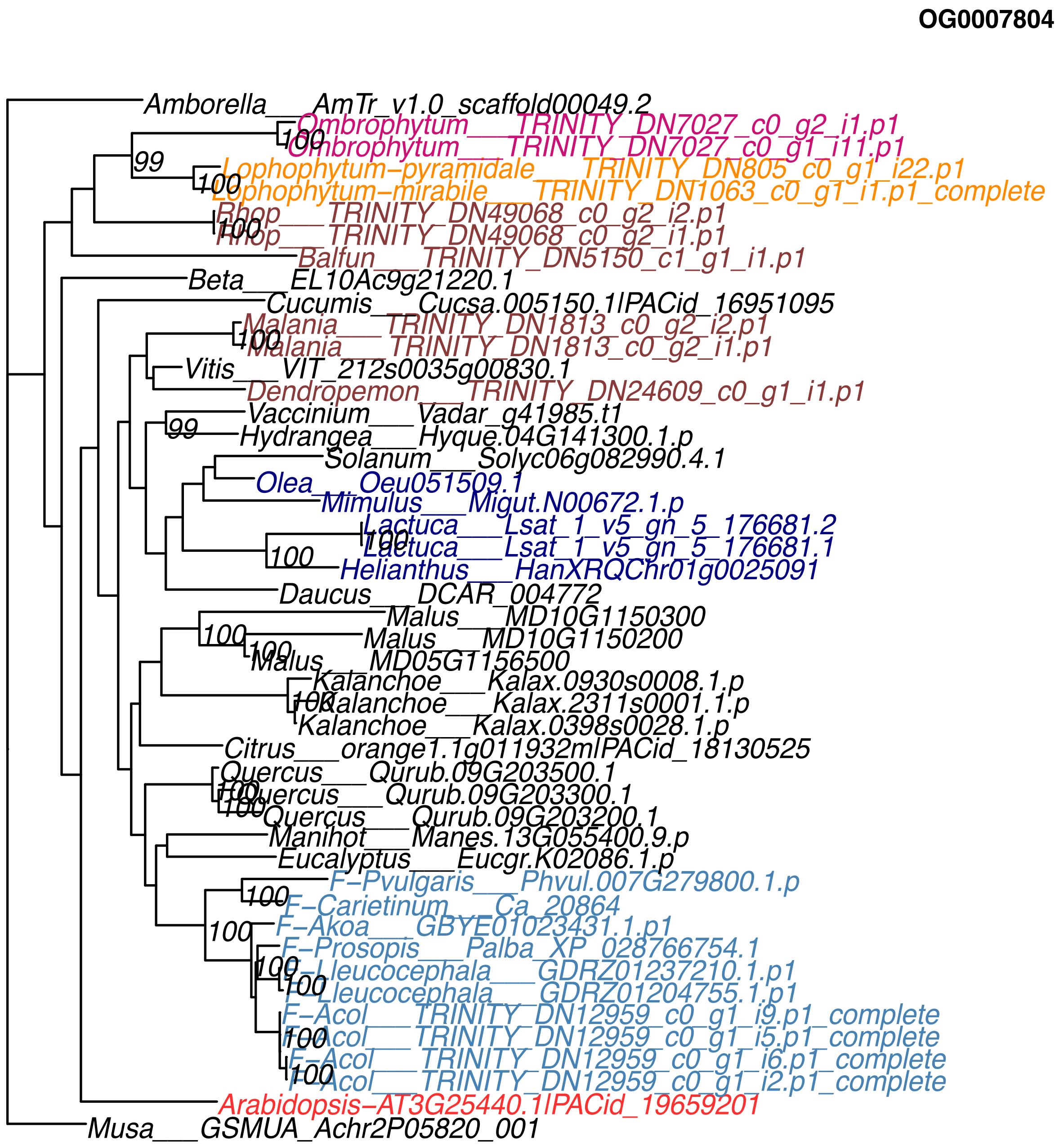
OG0007804
Amborella___AmTr_v1.0_scaffold00049.2
Ombrophytum___TRINITY_DN7027_c0_g2_i1.p1
Ombrophytum___TRINITY_DN7027_c0_g1_i11.p1
Lophophytum−pyramidale___TRINITY_DN805_c0_g1_i22.p1
Lophophytum−mirabile___TRINITY_DN1063_c0_g1_i1.p1_complete
Rhop___TRINITY_DN49068_c0_g2_i2.p1
Rhop___TRINITY_DN49068_c0_g2_i1.p1
Balfun___TRINITY_DN5150_c1_g1_i1.p1
Beta___EL10Ac9g21220.1
Cucumis___Cucsa.005150.1|PACid_16951095
Malania___TRINITY_DN1813_c0_g2_i2.p1
Malania___TRINITY_DN1813_c0_g2_i1.p1
Vitis___VIT_212s0035g00830.1
Dendropemon___TRINITY_DN24609_c0_g1_i1.p1
Vaccinium___Vadar_g41985.t1
Hydrangea___Hyque.04G141300.1.p
Solanum___Solyc06g082990.4.1
Olea___Oeu051509.1
Mimulus___Migut.N00672.1.p
Lactuca___Lsat_1_v5_gn_5_176681.2
Lactuca___Lsat_1_v5_gn_5_176681.1
Helianthus___HanXRQChr01g0025091
Daucus___DCAR_004772
Malus___MD10G1150300
Malus___MD10G1150200
Malus___MD05G1156500
Kalanchoe___Kalax.0930s0008.1.p
Kalanchoe___Kalax.2311s0001.1.p
Kalanchoe___Kalax.0398s0028.1.p
Citrus___orange1.1g011932m|PACid_18130525
Quercus___Qurub.09G203500.1
Quercus___Qurub.09G203300.1
Quercus___Qurub.09G203200.1
Manihot___Manes.13G055400.9.p
Eucalyptus___Eucgr.K02086.1.p
F−Pvulgaris___Phvul.007G279800.1.p
F−Carietinum___Ca_20864
F−Akoa___GBYE01023431.1.p1
F−Prosopis___Palba_XP_028766754.1
F−Lleucocephala___GDRZ01237210.1.p1
F−Lleucocephala___GDRZ01204755.1.p1
F−Acol___TRINITY_DN12959_c0_g1_i9.p1_complete
F−Acol___TRINITY_DN12959_c0_g1_i5.p1_complete
F−Acol___TRINITY_DN12959_c0_g1_i6.p1_complete
F−Acol___TRINITY_DN12959_c0_g1_i2.p1_complete
Arabidopsis−AT3G25440.1|PACid_19659201
Musa___GSMUA_Achr2P05820_001
100
99
100
100
100
99
100
100
100
100
100
100
100
100
100
100
100
100
100

OG0007844

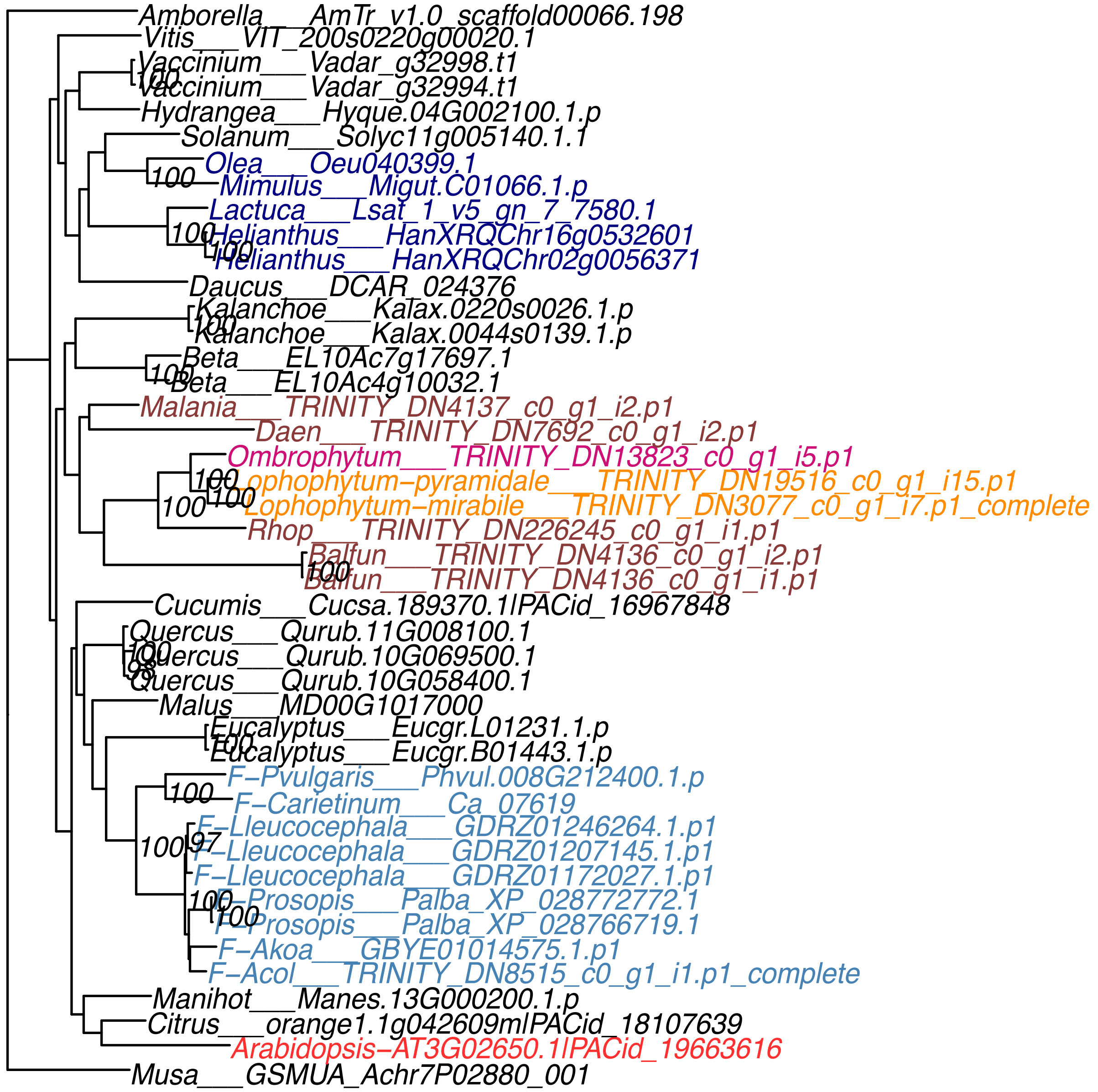

Amborella___AmTr_v1.0_scaffold00066.198
Vitis___VIT_200s0220g00020.1
Vaccinium___Vadar_g32998.t1
100
Vaccinium___Vadar_g32994.t1
Hydrangea___Hyque.04G002100.1.p
Solanum___Solyc11g005140.1.1
Olea___Oeu040399.1
100
Mimulus___Migut.C01066.1.p
Lactuca___Lsat_1_v5_gn_7_7580.1
100
Helianthus___HanXRQChr16g0532601
100
Helianthus___HanXRQChr02g0056371
Daucus___DCAR_024376
Kalanchoe___Kalax.0220s0026.1.p
100
Kalanchoe___Kalax.0044s0139.1.p
Beta___EL10Ac7g17697.1
100
Beta___EL10Ac4g10032.1
Malania___TRINITY_DN4137_c0_g1_i2.p1
Daen___TRINITY_DN7692_c0_g1_i2.p1
Ombrophytum___TRINITY_DN13823_c0_g1_i5.p1
100
Lophophytum-pyramidale___TRINITY_DN19516_c0_g1_i15.p1
100
Lophophytum-mirabile___TRINITY_DN3077_c0_g1_i7.p1_complete
100
Rhop___TRINITY_DN226245_c0_g1_i1.p1
Balfun___TRINITY_DN4136_c0_g1_i2.p1
100
Balfun___TRINITY_DN4136_c0_g1_i1.p1
Cucumis___Cucsa.189370.1IPACid_16967848
Quercus___Qurub.11G008100.1
100
Quercus___Qurub.10G069500.1
98
Quercus___Qurub.10G058400.1
Malus___MD00G1017000
Eucalyptus___Eucgr.L01231.1.p
100
Eucalyptus___Eucgr.B01443.1.p
F-Pvulgaris___Phvul.008G212400.1.p
100
F-Carietinum___Ca_07619
F-Lleucocephala___GDRZ01246264.1.p1
100
97
F-Lleucocephala___GDRZ01207145.1.p1
F-Lleucocephala___GDRZ01172027.1.p1
100
F-Prosopis___Palba_XP_028772772.1
100
F-Prosopis___Palba_XP_028766719.1
F-Akoa___GBYE01014575.1.p1
F-Acol___TRINITY_DN8515_c0_g1_i1.p1_complete
Manihot___Manes.13G000200.1.p
Citrus___orange1.1g042609mIPACid_18107639
Arabidopsis-AT3G02650.1IPACid_19663616
Musa___GSMUA_Achr7P02880_001

OG0007917

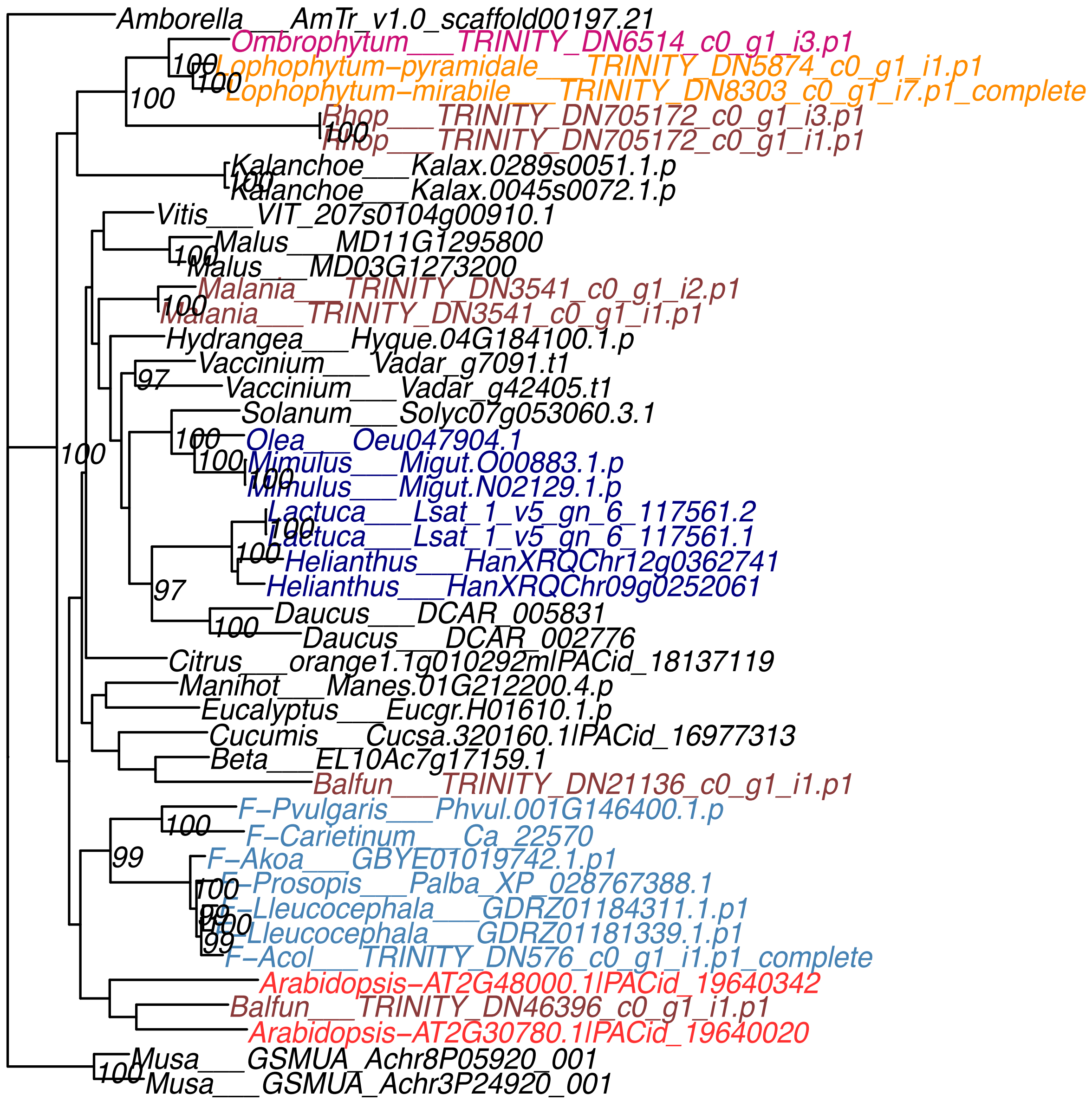

Amborella___AmTr_v1.0_scaffold00197.21
Ombrophytum___TRINITY_DN6514_c0_g1_i3.p1
Lophophytum-pyramidale___TRINITY_DN5874_c0_g1_i1.p1
Lophophytum-mirabile___TRINITY_DN8303_c0_g1_i7.p1_complete
Rhop___TRINITY_DN705172_c0_g1_i3.p1
Rhop___TRINITY_DN705172_c0_g1_i1.p1
Kalanchoe___Kalax.0289s0051.1.p
Kalanchoe___Kalax.0045s0072.1.p
Vitis___VIT_207s0104g00910.1
Malus___MD11G1295800
Malus___MD03G1273200
Malania___TRINITY_DN3541_c0_g1_i2.p1
Malania___TRINITY_DN3541_c0_g1_i1.p1
Hydrangea___Hyque.04G184100.1.p
Vaccinium___Vadar_g7091.t1
Vaccinium___Vadar_g42405.t1
Solanum___Solyc07g053060.3.1
Olea___Oeu047904.1
Mimulus___Migut.O00883.1.p
Mimulus___Migut.N02129.1.p
Lactuca___Lsat_1_v5_gn_6_117561.2
Lactuca___Lsat_1_v5_gn_6_117561.1
Helianthus___HanXRQChr12g0362741
Helianthus___HanXRQChr09g0252061
Daucus___DCAR_005831
Daucus___DCAR_002776
Citrus___orange1.1g010292mIPACid_18137119
Manihot___Manes.01G212200.4.p
Eucalyptus___Eucgr.H01610.1.p
Cucumis___Cucsa.320160.1IPACid_16977313
Beta___EL10Ac7g17159.1
Balfun___TRINITY_DN21136_c0_g1_i1.p1
F-Pvulgaris___Phvul.001G146400.1.p
F-Carietinum___Ca_22570
F-Akoa___GBYE01019742.1.p1
F-Prosopis___Palba_XP_028767388.1
F-Lleucocephala___GDRZ01184311.1.p1
F-Lleucocephala___GDRZ01181339.1.p1
F-Acol___TRINITY_DN576_c0_g1_i1.p1_complete
Arabidopsis-AT2G48000.1IPACid_19640342
Balfun___TRINITY_DN46396_c0_g1_i1.p1
Arabidopsis-AT2G30780.1IPACid_19640020
Musa___GSMUA_Achr8P05920_001
Musa___GSMUA_Achr3P24920_001
100
100
100
100
100
100
100
100
97
100
100
100
100
100
97
100
100
100
99
100
99
100
99
100

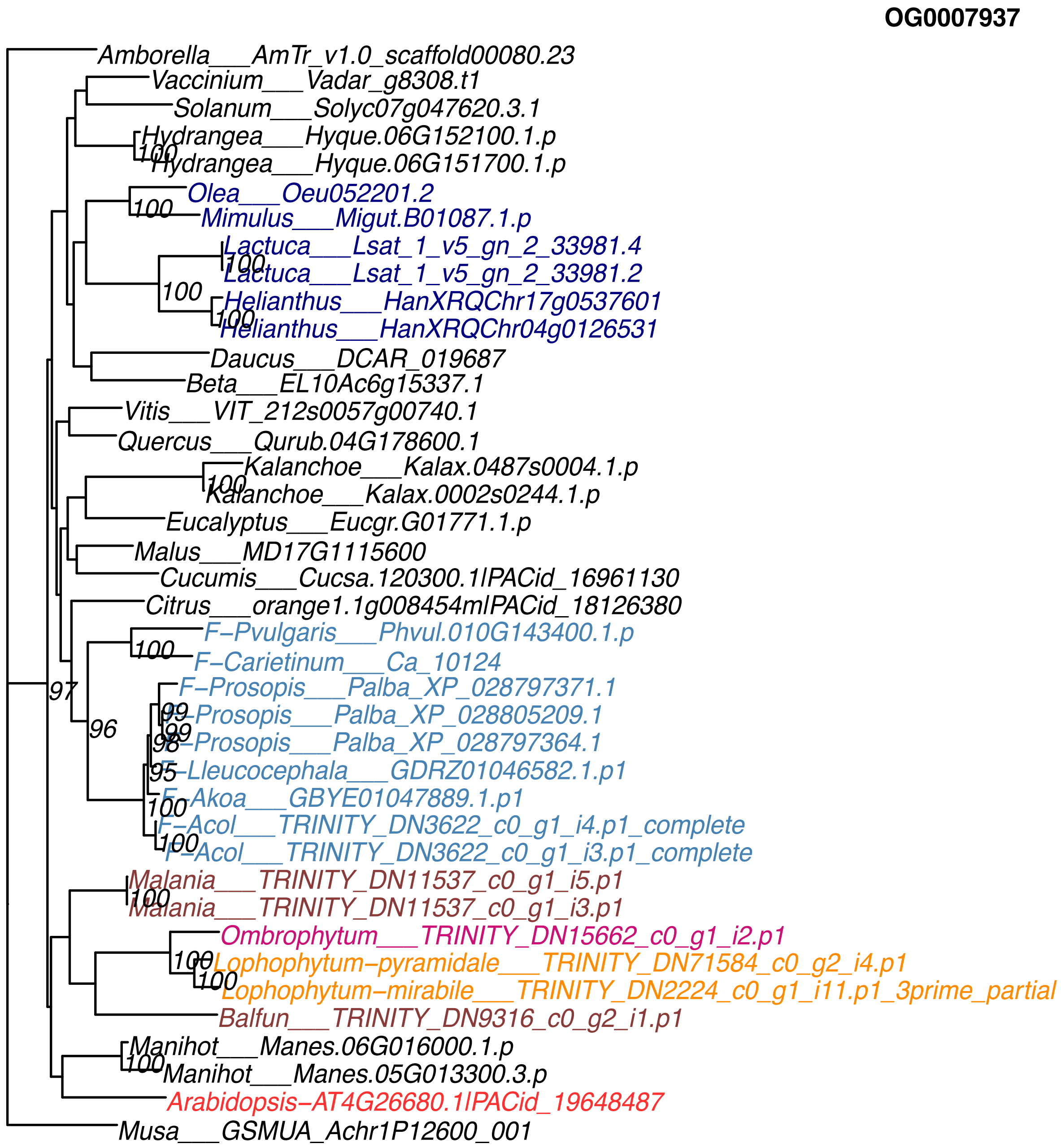

OG0007937
Amborella___AmTr_v1.0_scaffold00080.23
Vaccinium___Vadar_g8308.t1
Solanum___Solyc07g047620.3.1
Hydrangea___Hyque.06G152100.1.p
100
Hydrangea___Hyque.06G151700.1.p
Olea___Oeu052201.2
100
Mimulus___Migut.B01087.1.p
Lactuca___Lsat_1_v5_gn_2_33981.4
100
Lactuca___Lsat_1_v5_gn_2_33981.2
100
Helianthus___HanXRQChr17g0537601
100
Helianthus___HanXRQChr04g0126531
Daucus___DCAR_019687
Beta___EL10Ac6g15337.1
Vitis___VIT_212s0057g00740.1
Quercus___Qurub.04G178600.1
Kalanchoe___Kalax.0487s0004.1.p
100
Kalanchoe___Kalax.0002s0244.1.p
Eucalyptus___Eucgr.G01771.1.p
Malus___MD17G1115600
Cucumis___Cucsa.120300.1|PACid_16961130
Citrus___orange1.1g008454m|PACid_18126380
F−Pvulgaris___Phvul.010G143400.1.p
100
F−Carietinum___Ca_10124
97
F−Prosopis___Palba_XP_028797371.1
99
F−Prosopis___Palba_XP_028805209.1
99
96
98
F−Prosopis___Palba_XP_028797364.1
95
F−Lleucocephala___GDRZ01046582.1.p1
F−Akoa___GBYE01047889.1.p1
100
F−Acol___TRINITY_DN3622_c0_g1_i4.p1_complete
100
F−Acol___TRINITY_DN3622_c0_g1_i3.p1_complete
Malania___TRINITY_DN11537_c0_g1_i5.p1
100
Malania___TRINITY_DN11537_c0_g1_i3.p1
Ombrophytum___TRINITY_DN15662_c0_g1_i2.p1
100
Lophophytum−pyramidale___TRINITY_DN71584_c0_g2_i4.p1
100
Lophophytum−mirabile___TRINITY_DN2224_c0_g1_i11.p1_3prime_partial
Balfun___TRINITY_DN9316_c0_g2_i1.p1
Manihot___Manes.06G016000.1.p
100
Manihot___Manes.05G013300.3.p
Arabidopsis−AT4G26680.1|PACid_19648487
Musa___GSMUA_Achr1P12600_001

# OG0007948

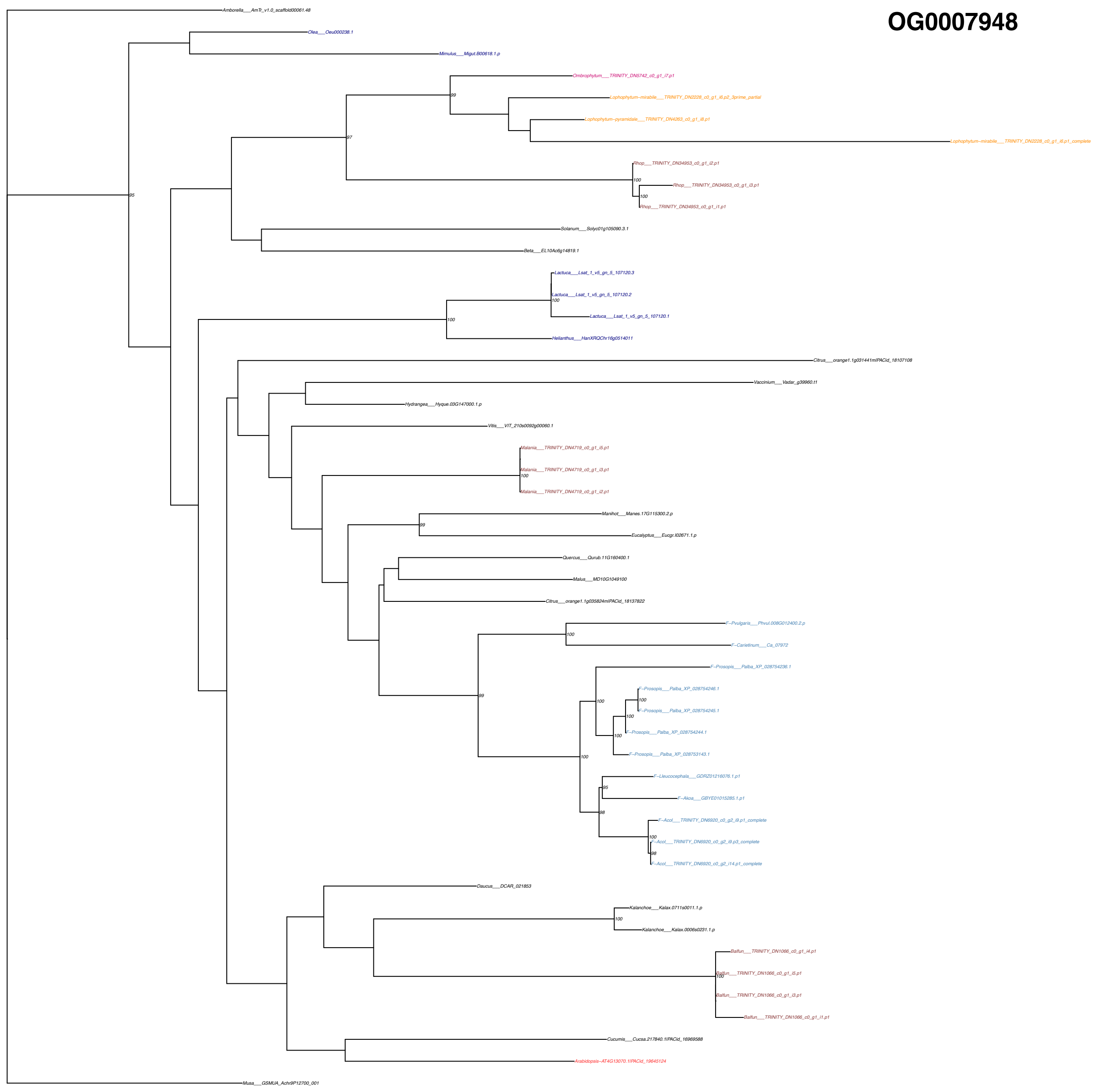

Amborella___AmTr_v1.0_scaffold00061.48
Olea___Oeu000238.1
Mimulus___Migut.B00618.1.p
95
Ombrophytum___TRINITY_DN5742_c0_g1_i7.p1
99
Lophophytum-mirabile___TRINITY_DN2228_c0_g1_i6.p2_3prime_partial
Lophophytum-pyramidale___TRINITY_DN4263_c0_g1_i8.p1
Lophophytum-mirabile___TRINITY_DN2228_c0_g1_i6.p1_complete
97
Rhop___TRINITY_DN34953_c0_g1_i2.p1
100
Rhop___TRINITY_DN34953_c0_g1_i3.p1
100
Rhop___TRINITY_DN34953_c0_g1_i1.p1
Solanum___Solyc01g105090.3.1
Beta___EL10Ac6g14819.1
Lactuca___Lsat_1_v5_gn_5_107120.3
Lactuca___Lsat_1_v5_gn_5_107120.2
100
Lactuca___Lsat_1_v5_gn_5_107120.1
100
Helianthus___HanXRQChr16g0514011
Citrus___orange1.1g031441mIPACid_18107108
Vaccinium___Vadar_g39960.t1
Hydrangea___Hyque.03G147000.1.p
Vitis___VIT_210s0092g00060.1
Malania___TRINITY_DN4719_c0_g1_i5.p1
Malania___TRINITY_DN4719_c0_g1_i3.p1
100
Malania___TRINITY_DN4719_c0_g1_i2.p1
Manihot___Manes.17G115300.2.p
99
Eucalyptus___Eucgr.I02671.1.p
Quercus___Qurub.11G160400.1
Malus___MD10G1049100
Citrus___orange1.1g035824mIPACid_18137822
F-Pvulgaris___Phvul.008G012400.2.p
100
F-Carietinum___Ca_07972
99
F-Prosopis___Palba_XP_028754236.1
100
F-Prosopis___Palba_XP_028754246.1
100
F-Prosopis___Palba_XP_028754245.1
100
F-Prosopis___Palba_XP_028754244.1
100
F-Prosopis___Palba_XP_028753143.1
100
F-Lleucocephala___GDRZ01216076.1.p1
95
F-Akoa___GBYE01015285.1.p1
98
F-Acol___TRINITY_DN6920_c0_g2_i9.p1_complete
100
F-Acol___TRINITY_DN6920_c0_g2_i9.p3_complete
98
F-Acol___TRINITY_DN6920_c0_g2_i14.p1_complete
Daucus___DCAR_021853
Kalanchoe___Kalax.0711s0011.1.p
100
Kalanchoe___Kalax.0006s0231.1.p
Balfun___TRINITY_DN1066_c0_g1_i4.p1
Balfun___TRINITY_DN1066_c0_g1_i5.p1
100
Balfun___TRINITY_DN1066_c0_g1_i3.p1
Balfun___TRINITY_DN1066_c0_g1_i1.p1
Cucumis___Cucsa.217840.1IPACid_16969588
Arabidopsis-AT4G13070.1IPACid_19645124
Musa___GSMUA_Achr9P12700_001

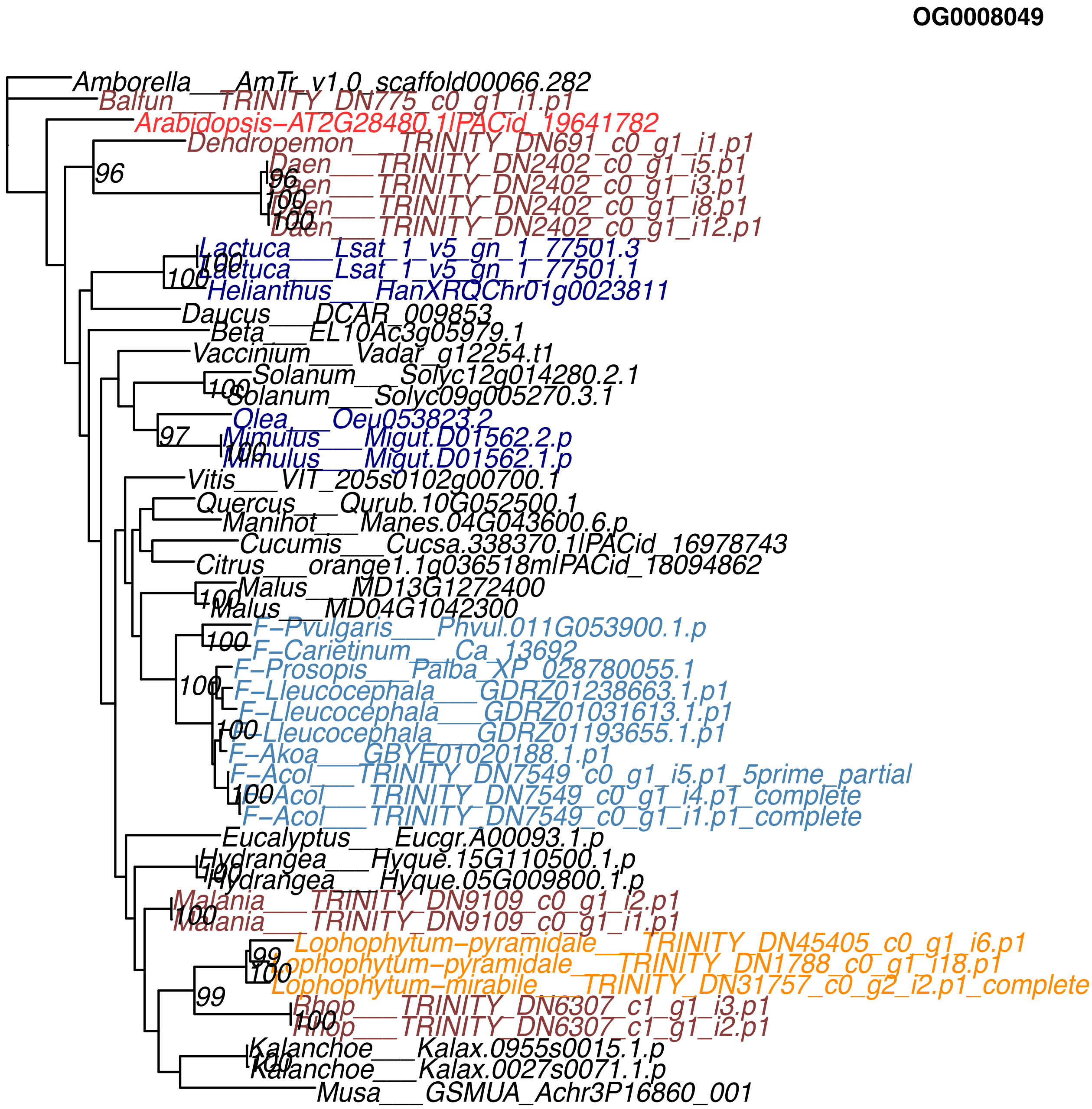
OG0008049
Amborella___AmTr_v1.0_scaffold00066.282
Balfun___TRINITY_DN775_c0_g1_i1.p1
Arabidopsis–AT2G28480.1|PACid_19641782
Dendropemon___TRINITY_DN691_c0_g1_i1.p1
Daen___TRINITY_DN2402_c0_g1_i5.p1
Daen___TRINITY_DN2402_c0_g1_i3.p1
Daen___TRINITY_DN2402_c0_g1_i8.p1
Daen___TRINITY_DN2402_c0_g1_i12.p1
Lactuca___Lsat_1_v5_gn_1_77501.3
Lactuca___Lsat_1_v5_gn_1_77501.1
Helianthus___HanXRQChr01g0023811
Daucus___DCAR_009853
Beta___EL10Ac3g05979.1
Vaccinium___Vadar_g12254.t1
Solanum___Solyc12g014280.2.1
Solanum___Solyc09g005270.3.1
Olea___Oeu053823.2
Mimulus___Migut.D01562.2.p
Mimulus___Migut.D01562.1.p
Vitis___VIT_205s0102g00700.1
Quercus___Qurub.10G052500.1
Manihot___Manes.04G043600.6.p
Cucumis___Cucsa.338370.1|PACid_16978743
Citrus___orange1.1g036518m|PACid_18094862
Malus___MD13G1272400
Malus___MD04G1042300
F–Pvulgaris___Phvul.011G053900.1.p
F–Carietinum___Ca_13692
F–Prosopis___Palba_XP_028780055.1
F–Lleucocephala___GDRZ01238663.1.p1
F–Lleucocephala___GDRZ01031613.1.p1
F–Lleucocephala___GDRZ01193655.1.p1
F–Akoa___GBYE01020188.1.p1
F–Acol___TRINITY_DN7549_c0_g1_i5.p1_5prime_partial
F–Acol___TRINITY_DN7549_c0_g1_i4.p1_complete
F–Acol___TRINITY_DN7549_c0_g1_i1.p1_complete
Eucalyptus___Eucgr.A00093.1.p
Hydrangea___Hyque.15G110500.1.p
Hydrangea___Hyque.05G009800.1.p
Malania___TRINITY_DN9109_c0_g1_i2.p1
Malania___TRINITY_DN9109_c0_g1_i1.p1
Lophophytum–pyramidale___TRINITY_DN45405_c0_g1_i6.p1
Lophophytum–pyramidale___TRINITY_DN1788_c0_g1_i18.p1
Lophophytum–mirabile___TRINITY_DN31757_c0_g2_i2.p1_complete
Rhop___TRINITY_DN6307_c1_g1_i3.p1
Rhop___TRINITY_DN6307_c1_g1_i2.p1
Kalanchoe___Kalax.0955s0015.1.p
Kalanchoe___Kalax.0027s0071.1.p
Musa___GSMUA_Achr3P16860_001

**OG0008078**

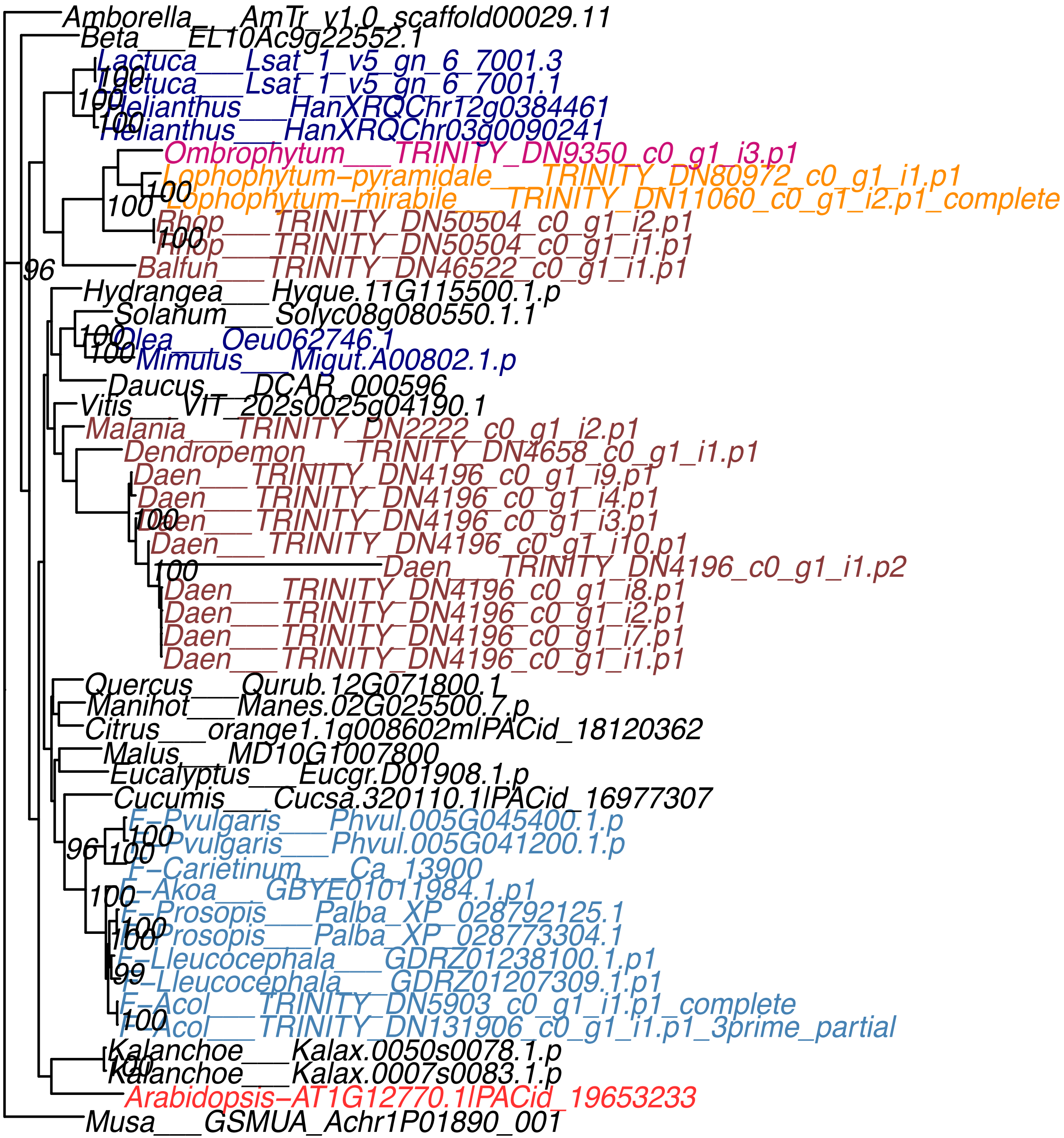
Amborella___AmTr_v1.0_scaffold00029.11
Beta___EL10Ac9g22552.1
Lactuca___Lsat_1_v5_gn_6_7001.3
Lactuca___Lsat_1_v5_gn_6_7001.1
Helianthus___HanXRQChr12g0384461
Helianthus___HanXRQChr03g0090241
Ombrophytum___TRINITY_DN9350_c0_g1_i3.p1
Lophophytum–pyramidale___TRINITY_DN80972_c0_g1_i1.p1
Lophophytum–mirabile___TRINITY_DN11060_c0_g1_i2.p1_complete
Rhop___TRINITY_DN50504_c0_g1_i2.p1
Rhop___TRINITY_DN50504_c0_g1_i1.p1
Balfun___TRINITY_DN46522_c0_g1_i1.p1
Hydrangea___Hyque.11G115500.1.p
Solanum___Solyc08g080550.1.1
Olea___Oeu062746.1
Mimulus___Migut.A00802.1.p
Daucus___DCAR_000596
Vitis___VIT_202s0025g04190.1
Malania___TRINITY_DN2222_c0_g1_i2.p1
Dendropemon___TRINITY_DN4658_c0_g1_i1.p1
Daen___TRINITY_DN4196_c0_g1_i9.p1
Daen___TRINITY_DN4196_c0_g1_i4.p1
Daen___TRINITY_DN4196_c0_g1_i3.p1
Daen___TRINITY_DN4196_c0_g1_i10.p1
Daen___TRINITY_DN4196_c0_g1_i1.p2
Daen___TRINITY_DN4196_c0_g1_i8.p1
Daen___TRINITY_DN4196_c0_g1_i2.p1
Daen___TRINITY_DN4196_c0_g1_i7.p1
Daen___TRINITY_DN4196_c0_g1_i1.p1
Quercus___Qurub.12G071800.1
Manihot___Manes.02G025500.7.p
Citrus___orange1.1g008602m|PACid_18120362
Malus___MD10G1007800
Eucalyptus___Eucgr.D01908.1.p
Cucumis___Cucsa.320110.1|PACid_16977307
F–Pvulgaris___Phvul.005G045400.1.p
F–Pvulgaris___Phvul.005G041200.1.p
F–Carietinum___Ca_13900
F–Akoa___GBYE01011984.1.p1
F–Prosopis___Palba_XP_028792125.1
F–Prosopis___Palba_XP_028773304.1
F–Lleucocephala___GDRZ01238100.1.p1
F–Lleucocephala___GDRZ01207309.1.p1
F–Acol___TRINITY_DN5903_c0_g1_i1.p1_complete
F–Acol___TRINITY_DN131906_c0_g1_i1.p1_3prime_partial
Kalanchoe___Kalax.0050s0078.1.p
Kalanchoe___Kalax.0007s0083.1.p
Arabidopsis–AT1G12770.1|PACid_19653233
Musa___GSMUA_Achr1P01890_001
100
100
100
100
100
100
96
100
100
100
100
96
100
100
100
100
99
100
100

**OG0008144**

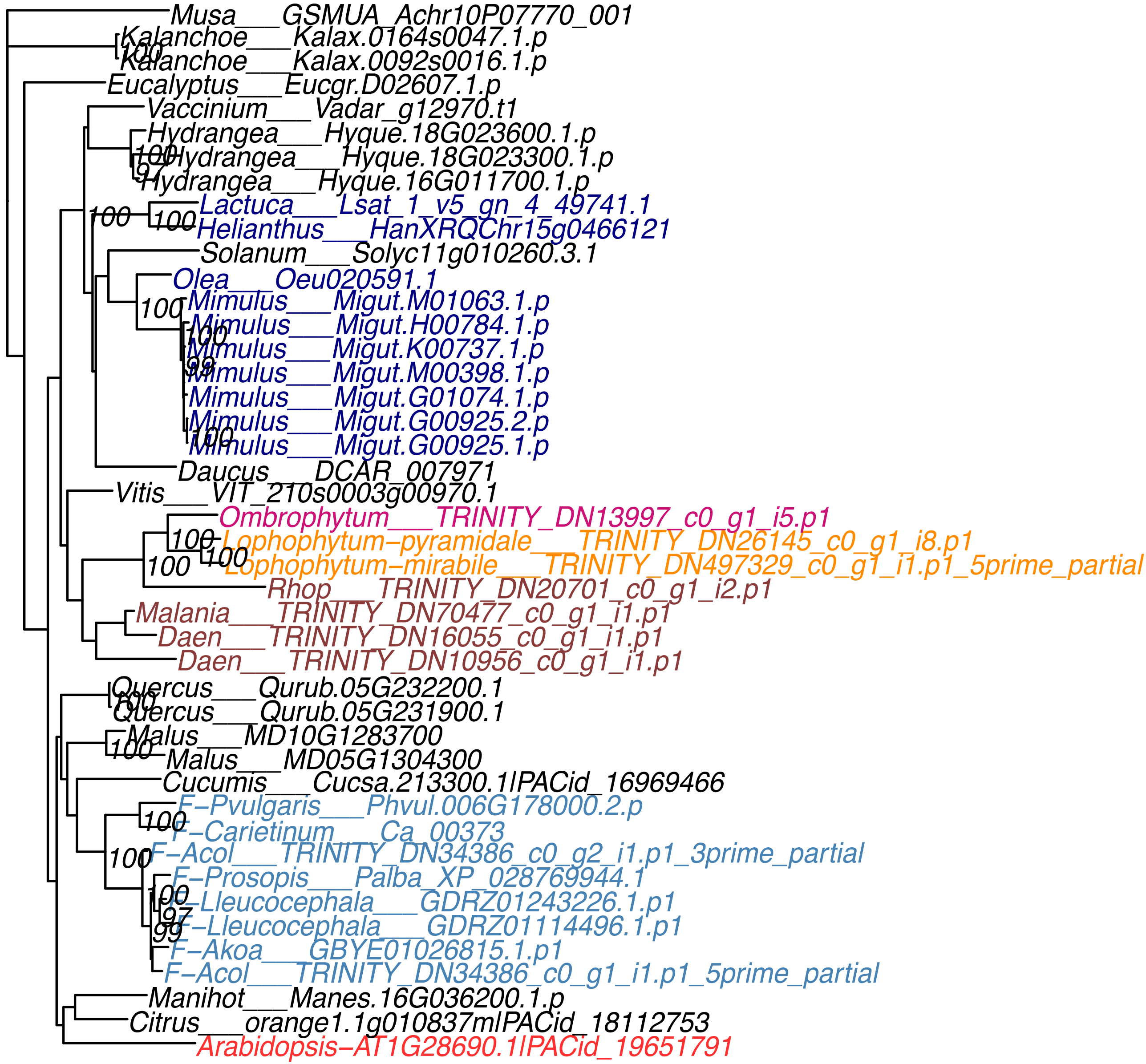

Musa___GSMUA_Achr10P07770_001
Kalanchoe___Kalax.0164s0047.1.p
100
Kalanchoe___Kalax.0092s0016.1.p
Eucalyptus___Eucgr.D02607.1.p
Vaccinium___Vadar_g12970.t1
Hydrangea___Hyque.18G023600.1.p
100
Hydrangea___Hyque.18G023300.1.p
97
Hydrangea___Hyque.16G011700.1.p
100
Lactuca___Lsat_1_v5_gn_4_49741.1
100
Helianthus___HanXRQChr15g0466121
Solanum___Solyc11g010260.3.1
Olea___Oeu020591.1
100
Mimulus___Migut.M01063.1.p
Mimulus___Migut.H00784.1.p
100
Mimulus___Migut.K00737.1.p
99
Mimulus___Migut.M00398.1.p
Mimulus___Migut.G01074.1.p
Mimulus___Migut.G00925.2.p
100
Mimulus___Migut.G00925.1.p
Daucus___DCAR_007971
Vitis___VIT_210s0003g00970.1
Ombrophytum___TRINITY_DN13997_c0_g1_i5.p1
100
Lophophytum-pyramidale___TRINITY_DN26145_c0_g1_i8.p1
100
100
Lophophytum-mirabile___TRINITY_DN497329_c0_g1_i1.p1_5prime_partial
Rhop___TRINITY_DN20701_c0_g1_i2.p1
Malania___TRINITY_DN70477_c0_g1_i1.p1
Daen___TRINITY_DN16055_c0_g1_i1.p1
Daen___TRINITY_DN10956_c0_g1_i1.p1
Quercus___Qurub.05G232200.1
100
Quercus___Qurub.05G231900.1
Malus___MD10G1283700
100
Malus___MD05G1304300
Cucumis___Cucsa.213300.1|PACid_16969466
F-Pvulgaris___Phvul.006G178000.2.p
100
F-Carietinum___Ca_00373
100
F-Acol___TRINITY_DN34386_c0_g2_i1.p1_3prime_partial
F-Prosopis___Palba_XP_028769944.1
100
F-Lleucocephala___GDRZ01243226.1.p1
97
F-Lleucocephala___GDRZ01114496.1.p1
99
F-Akoa___GBYE01026815.1.p1
F-Acol___TRINITY_DN34386_c0_g1_i1.p1_5prime_partial
Manihot___Manes.16G036200.1.p
Citrus___orange1.1g010837m|PACid_18112753
Arabidopsis-AT1G28690.1|PACid_19651791

**OG0008160**

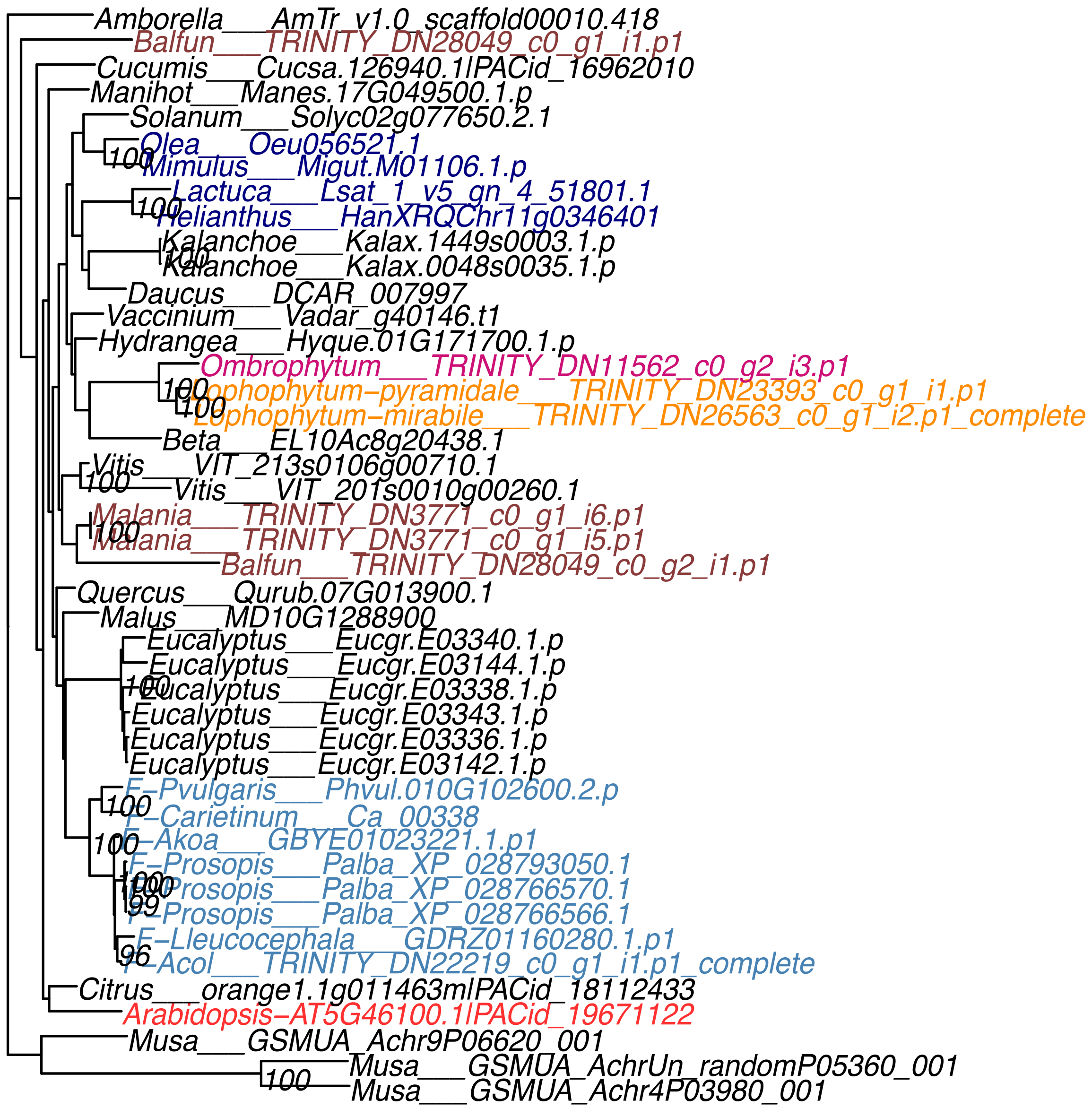
Amborella___AmTr_v1.0_scaffold00010.418
Balfun___TRINITY_DN28049_c0_g1_i1.p1
Cucumis___Cucsa.126940.1|PACid_16962010
Manihot___Manes.17G049500.1.p
Solanum___Solyc02g077650.2.1
Olea___Oeu056521.1
100
Mimulus___Migut.M01106.1.p
Lactuca___Lsat_1_v5_gn_4_51801.1
100
Helianthus___HanXRQChr11g0346401
Kalanchoe___Kalax.1449s0003.1.p
100
Kalanchoe___Kalax.0048s0035.1.p
Daucus___DCAR_007997
Vaccinium___Vadar_g40146.t1
Hydrangea___Hyque.01G171700.1.p
Ombrophytum___TRINITY_DN11562_c0_g2_i3.p1
100
Lophophytum−pyramidale___TRINITY_DN23393_c0_g1_i1.p1
100
Lophophytum−mirabile___TRINITY_DN26563_c0_g1_i2.p1_complete
Beta___EL10Ac8g20438.1
Vitis___VIT_213s0106g00710.1
100
Vitis___VIT_201s0010g00260.1
Malania___TRINITY_DN3771_c0_g1_i6.p1
100
Malania___TRINITY_DN3771_c0_g1_i5.p1
Balfun___TRINITY_DN28049_c0_g2_i1.p1
Quercus___Qurub.07G013900.1
Malus___MD10G1288900
Eucalyptus___Eucgr.E03340.1.p
Eucalyptus___Eucgr.E03144.1.p
100
Eucalyptus___Eucgr.E03338.1.p
Eucalyptus___Eucgr.E03343.1.p
Eucalyptus___Eucgr.E03336.1.p
Eucalyptus___Eucgr.E03142.1.p
F−Pvulgaris___Phvul.010G102600.2.p
100
F−Carietinum___Ca_00338
100
F−Akoa___GBYE01023221.1.p1
F−Prosopis___Palba_XP_028793050.1
100
100
F−Prosopis___Palba_XP_028766570.1
99
F−Prosopis___Palba_XP_028766566.1
F−Lleucocephala___GDRZ01160280.1.p1
96
F−Acol___TRINITY_DN22219_c0_g1_i1.p1_complete
Citrus___orange1.1g011463m|PACid_18112433
Arabidopsis−AT5G46100.1|PACid_19671122
Musa___GSMUA_Achr9P06620_001
Musa___GSMUA_AchrUn_randomP05360_001
100
Musa___GSMUA_Achr4P03980_001

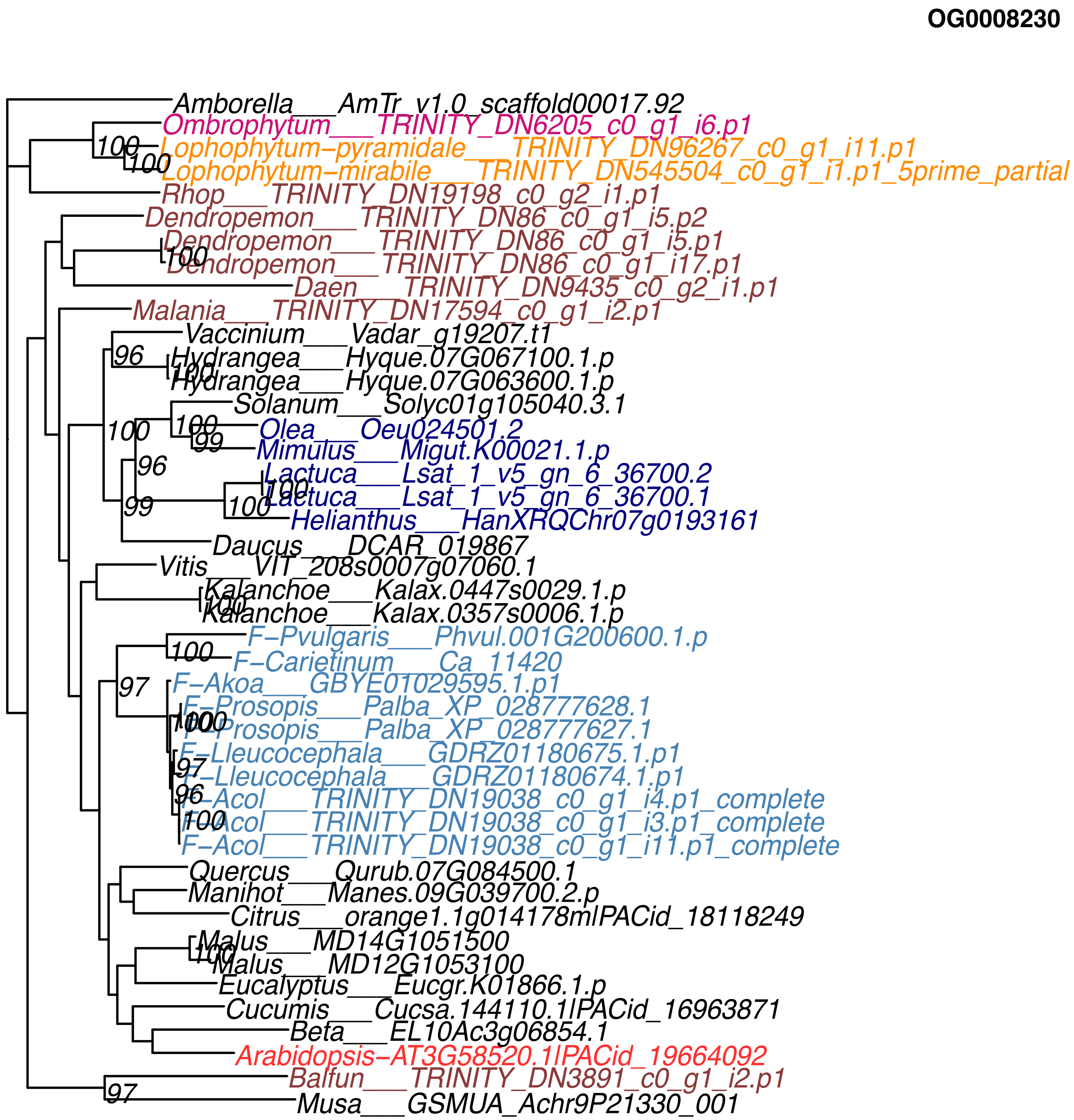
OG0008230
Amborella___AmTr_v1.0_scaffold00017.92
Ombrophytum___TRINITY_DN6205_c0_g1_i6.p1
Lophophytum-pyramidale___TRINITY_DN96267_c0_g1_i11.p1
Lophophytum-mirabile___TRINITY_DN545504_c0_g1_i1.p1_5prime_partial
Rhop___TRINITY_DN19198_c0_g2_i1.p1
Dendropemon___TRINITY_DN86_c0_g1_i5.p2
Dendropemon___TRINITY_DN86_c0_g1_i5.p1
Dendropemon___TRINITY_DN86_c0_g1_i17.p1
Daen___TRINITY_DN9435_c0_g2_i1.p1
Malania___TRINITY_DN17594_c0_g1_i2.p1
Vaccinium___Vadar_g19207.t1
Hydrangea___Hyque.07G067100.1.p
Hydrangea___Hyque.07G063600.1.p
Solanum___Solyc01g105040.3.1
Olea___Oeu024501.2
Mimulus___Migut.K00021.1.p
Lactuca___Lsat_1_v5_gn_6_36700.2
Lactuca___Lsat_1_v5_gn_6_36700.1
Helianthus___HanXRQChr07g0193161
Daucus___DCAR_019867
Vitis___VIT_208s0007g07060.1
Kalanchoe___Kalax.0447s0029.1.p
Kalanchoe___Kalax.0357s0006.1.p
F-Pvulgaris___Phvul.001G200600.1.p
F-Carietinum___Ca_11420
F-Akoa___GBYE01029595.1.p1
F-Prosopis___Palba_XP_028777628.1
F-Prosopis___Palba_XP_028777627.1
F-Lleucocephala___GDRZ01180675.1.p1
F-Lleucocephala___GDRZ01180674.1.p1
F-Acol___TRINITY_DN19038_c0_g1_i4.p1_complete
F-Acol___TRINITY_DN19038_c0_g1_i3.p1_complete
F-Acol___TRINITY_DN19038_c0_g1_i11.p1_complete
Quercus___Qurub.07G084500.1
Manihot___Manes.09G039700.2.p
Citrus___orange1.1g014178mIPACid_18118249
Malus___MD14G1051500
Malus___MD12G1053100
Eucalyptus___Eucgr.K01866.1.p
Cucumis___Cucsa.144110.1IPACid_16963871
Beta___EL10Ac3g06854.1
Arabidopsis-AT3G58520.1IPACid_19664092
Balfun___TRINITY_DN3891_c0_g1_i2.p1
Musa___GSMUA_Achr9P21330_001
100
100
100
96
100
100
100
99
96
100
100
99
100
100
97
100
97
96
100
100
97

**OG0008319**

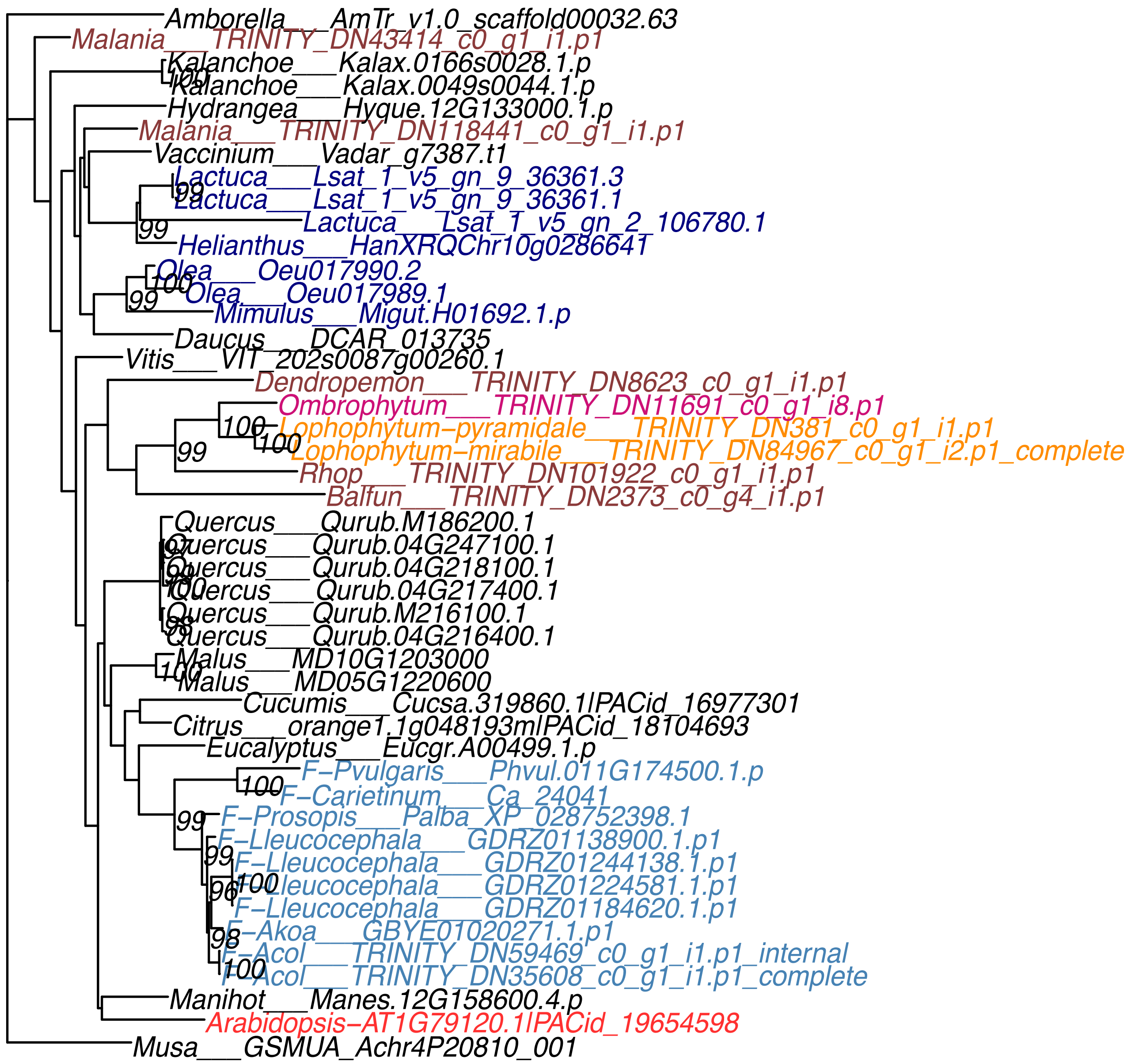

Amborella___AmTr_v1.0_scaffold00032.63
Malania___TRINITY_DN43414_c0_g1_i1.p1
Kalanchoe___Kalax.0166s0028.1.p
100
Kalanchoe___Kalax.0049s0044.1.p
Hydrangea___Hyque.12G133000.1.p
Malania___TRINITY_DN118441_c0_g1_i1.p1
Vaccinium___Vadar_g7387.t1
Lactuca___Lsat_1_v5_gn_9_36361.3
99
Lactuca___Lsat_1_v5_gn_9_36361.1
Lactuca___Lsat_1_v5_gn_2_106780.1
99
Helianthus___HanXRQChr10g0286641
Olea___Oeu017990.2
100
Olea___Oeu017989.1
99
Mimulus___Migut.H01692.1.p
Daucus___DCAR_013735
Vitis___VIT_202s0087g00260.1
Dendropemon___TRINITY_DN8623_c0_g1_i1.p1
Ombrophytum___TRINITY_DN11691_c0_g1_i8.p1
100
Lophophytum−pyramidale___TRINITY_DN381_c0_g1_i1.p1
100
Lophophytum−mirabile___TRINITY_DN84967_c0_g1_i2.p1_complete
99
Rhop___TRINITY_DN101922_c0_g1_i1.p1
Balfun___TRINITY_DN2373_c0_g4_i1.p1
Quercus___Qurub.M186200.1
97
Quercus___Qurub.04G247100.1
99
Quercus___Qurub.04G218100.1
100
Quercus___Qurub.04G217400.1
Quercus___Qurub.M216100.1
98
Quercus___Qurub.04G216400.1
Malus___MD10G1203000
100
Malus___MD05G1220600
Cucumis___Cucsa.319860.1|PACid_16977301
Citrus___orange1.1g048193m|PACid_18104693
Eucalyptus___Eucgr.A00499.1.p
F−Pvulgaris___Phvul.011G174500.1.p
100
F−Carietinum___Ca_24041
99
F−Prosopis___Palba_XP_028752398.1
F−Lleucocephala___GDRZ01138900.1.p1
99
F−Lleucocephala___GDRZ01244138.1.p1
100
96
F−Lleucocephala___GDRZ01224581.1.p1
F−Lleucocephala___GDRZ01184620.1.p1
F−Akoa___GBYE01020271.1.p1
98
F−Acol___TRINITY_DN59469_c0_g1_i1.p1_internal
100
F−Acol___TRINITY_DN35608_c0_g1_i1.p1_complete
Manihot___Manes.12G158600.4.p
Arabidopsis−AT1G79120.1|PACid_19654598
Musa___GSMUA_Achr4P20810_001

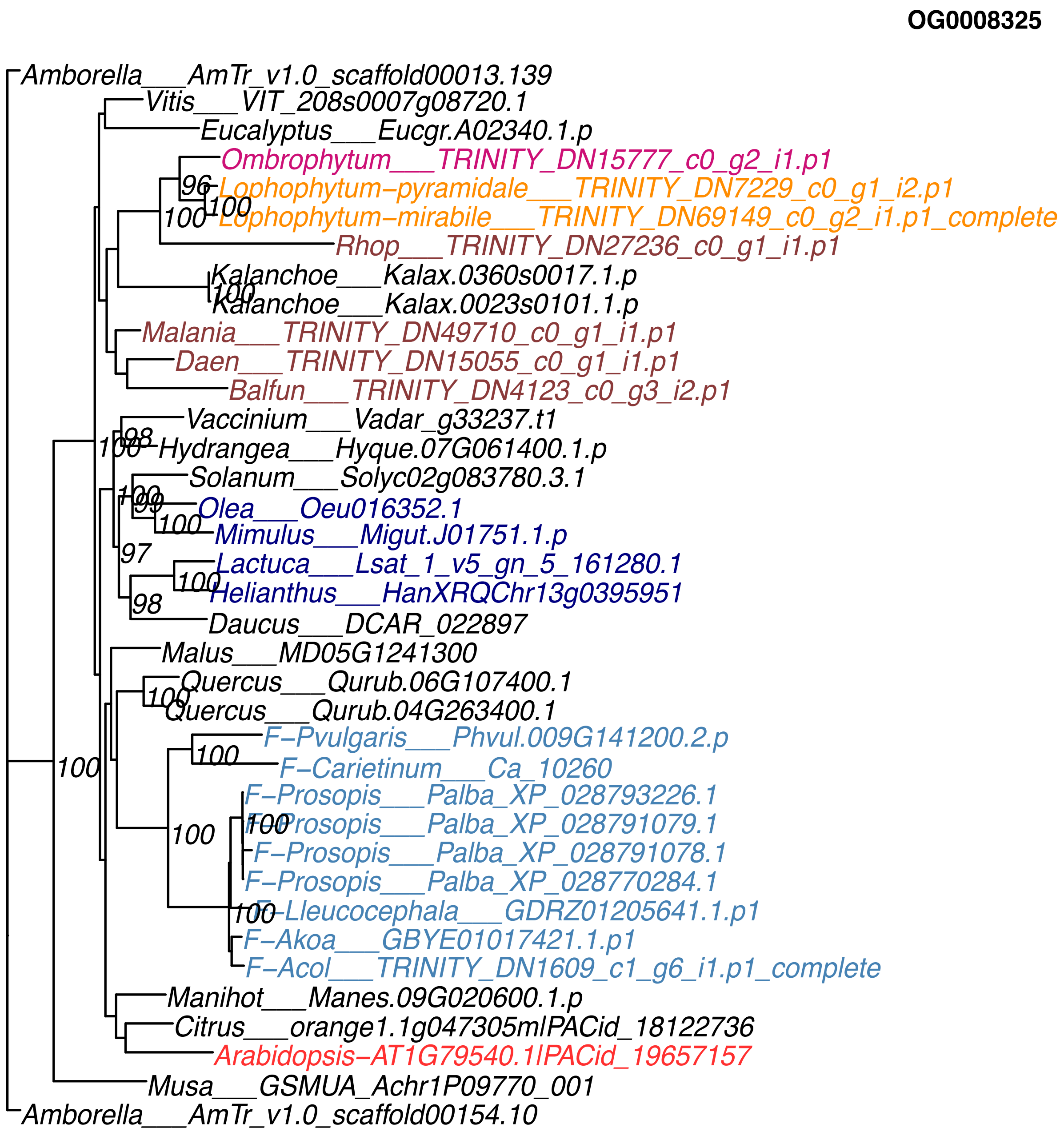
OG0008325
Amborella___AmTr_v1.0_scaffold00013.139
Vitis___VIT_208s0007g08720.1
Eucalyptus___Eucgr.A02340.1.p
Ombrophytum___TRINITY_DN15777_c0_g2_i1.p1
Lophophytum–pyramidale___TRINITY_DN7229_c0_g1_i2.p1
Lophophytum–mirabile___TRINITY_DN69149_c0_g2_i1.p1_complete
Rhop___TRINITY_DN27236_c0_g1_i1.p1
Kalanchoe___Kalax.0360s0017.1.p
Kalanchoe___Kalax.0023s0101.1.p
Malania___TRINITY_DN49710_c0_g1_i1.p1
Daen___TRINITY_DN15055_c0_g1_i1.p1
Balfun___TRINITY_DN4123_c0_g3_i2.p1
Vaccinium___Vadar_g33237.t1
Hydrangea___Hyque.07G061400.1.p
Solanum___Solyc02g083780.3.1
Olea___Oeu016352.1
Mimulus___Migut.J01751.1.p
Lactuca___Lsat_1_v5_gn_5_161280.1
Helianthus___HanXRQChr13g0395951
Daucus___DCAR_022897
Malus___MD05G1241300
Quercus___Qurub.06G107400.1
Quercus___Qurub.04G263400.1
F–Pvulgaris___Phvul.009G141200.2.p
F–Carietinum___Ca_10260
F–Prosopis___Palba_XP_028793226.1
F–Prosopis___Palba_XP_028791079.1
F–Prosopis___Palba_XP_028791078.1
F–Prosopis___Palba_XP_028770284.1
F–Lleucocephala___GDRZ01205641.1.p1
F–Akoa___GBYE01017421.1.p1
F–Acol___TRINITY_DN1609_c1_g6_i1.p1_complete
Manihot___Manes.09G020600.1.p
Citrus___orange1.1g047305m|PACid_18122736
Arabidopsis–AT1G79540.1|PACid_19657157
Musa___GSMUA_Achr1P09770_001
Amborella___AmTr_v1.0_scaffold00154.10
96
100
100
100
98
100
100
99
100
97
100
98
100
100
100
100
100
100

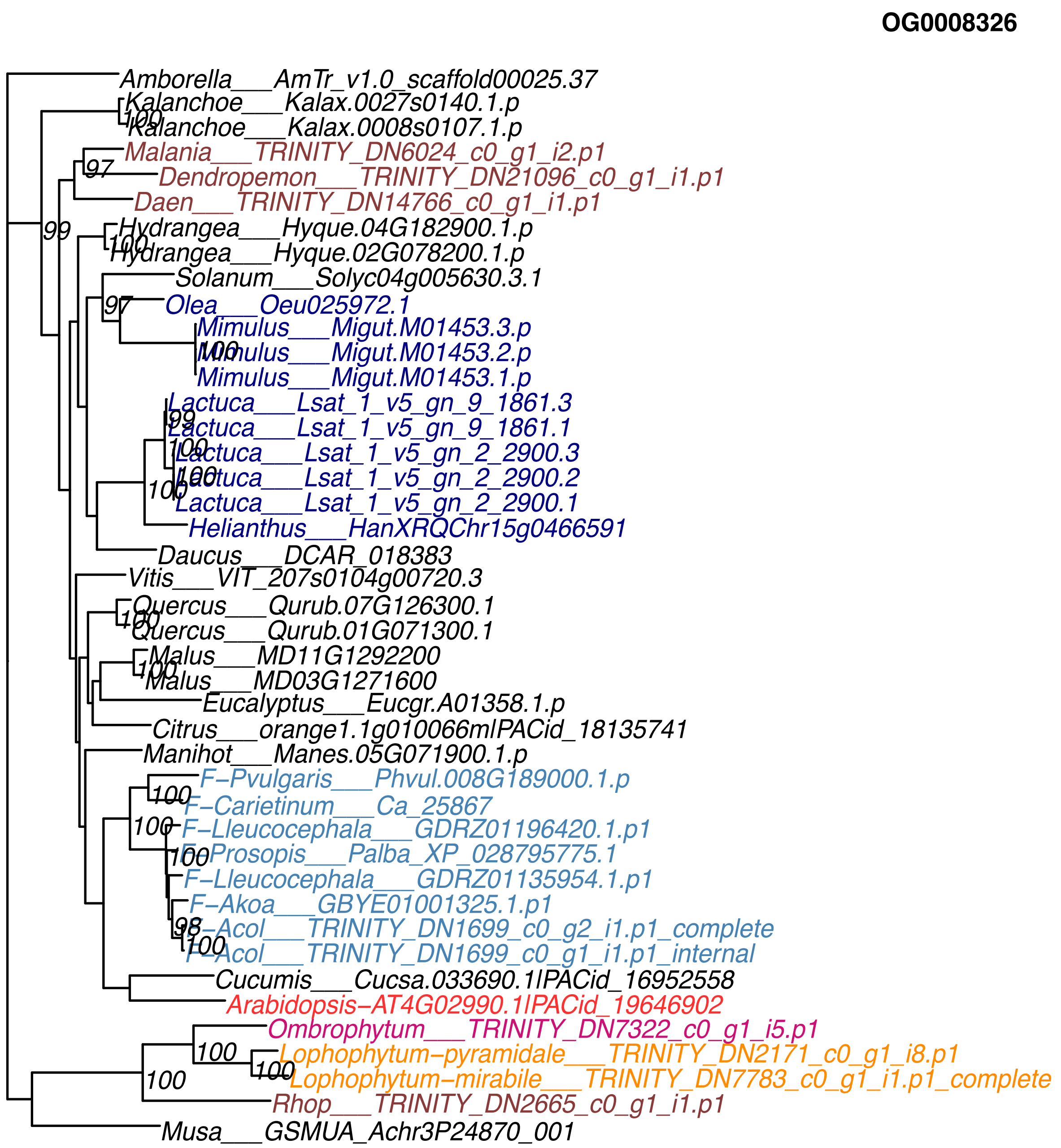
OG0008326
Amborella___AmTr_v1.0_scaffold00025.37
Kalanchoe___Kalax.0027s0140.1.p
100
Kalanchoe___Kalax.0008s0107.1.p
Malania___TRINITY_DN6024_c0_g1_i2.p1
97
Dendropemon___TRINITY_DN21096_c0_g1_i1.p1
Daen___TRINITY_DN14766_c0_g1_i1.p1
99
Hydrangea___Hyque.04G182900.1.p
100
Hydrangea___Hyque.02G078200.1.p
Solanum___Solyc04g005630.3.1
97
Olea___Oeu025972.1
Mimulus___Migut.M01453.3.p
100
Mimulus___Migut.M01453.2.p
Mimulus___Migut.M01453.1.p
Lactuca___Lsat_1_v5_gn_9_1861.3
99
Lactuca___Lsat_1_v5_gn_9_1861.1
100
Lactuca___Lsat_1_v5_gn_2_2900.3
100
Lactuca___Lsat_1_v5_gn_2_2900.2
100
Lactuca___Lsat_1_v5_gn_2_2900.1
Helianthus___HanXRQChr15g0466591
Daucus___DCAR_018383
Vitis___VIT_207s0104g00720.3
Quercus___Qurub.07G126300.1
100
Quercus___Qurub.01G071300.1
Malus___MD11G1292200
100
Malus___MD03G1271600
Eucalyptus___Eucgr.A01358.1.p
Citrus___orange1.1g010066mlPACid_18135741
Manihot___Manes.05G071900.1.p
F−Pvulgaris___Phvul.008G189000.1.p
100
F−Carietinum___Ca_25867
100
F−Lleucocephala___GDRZ01196420.1.p1
F−Prosopis___Palba_XP_028795775.1
100
F−Lleucocephala___GDRZ01135954.1.p1
F−Akoa___GBYE01001325.1.p1
98
F−Acol___TRINITY_DN1699_c0_g2_i1.p1_complete
100
F−Acol___TRINITY_DN1699_c0_g1_i1.p1_internal
Cucumis___Cucsa.033690.1lPACid_16952558
Arabidopsis−AT4G02990.1lPACid_19646902
Ombrophytum___TRINITY_DN7322_c0_g1_i5.p1
100
Lophophytum−pyramidale___TRINITY_DN2171_c0_g1_i8.p1
100
Lophophytum−mirabile___TRINITY_DN7783_c0_g1_i1.p1_complete
100
Rhop___TRINITY_DN2665_c0_g1_i1.p1
Musa___GSMUA_Achr3P24870_001

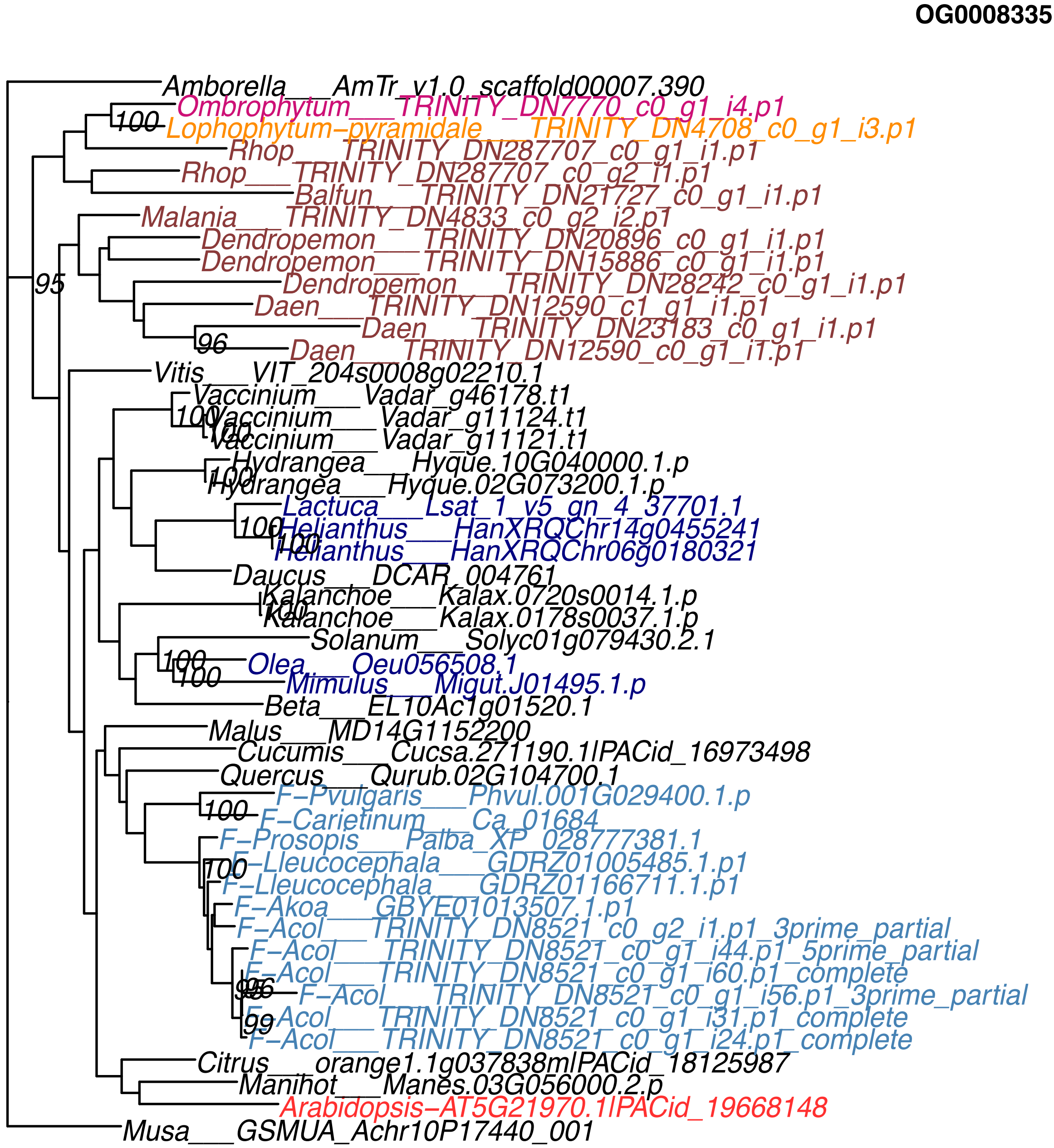

OG0008335
Amborella___AmTr_v1.0_scaffold00007.390
Ombrophytum___TRINITY_DN7770_c0_g1_i4.p1
Lophophytum-pyramidale___TRINITY_DN4708_c0_g1_i3.p1
Rhop___TRINITY_DN287707_c0_g1_i1.p1
Rhop___TRINITY_DN287707_c0_g2_i1.p1
Balfun___TRINITY_DN21727_c0_g1_i1.p1
Malania___TRINITY_DN4833_c0_g2_i2.p1
Dendropemon___TRINITY_DN20896_c0_g1_i1.p1
Dendropemon___TRINITY_DN15886_c0_g1_i1.p1
Dendropemon___TRINITY_DN28242_c0_g1_i1.p1
Daen___TRINITY_DN12590_c1_g1_i1.p1
Daen___TRINITY_DN23183_c0_g1_i1.p1
Daen___TRINITY_DN12590_c0_g1_i1.p1
Vitis___VIT_204s0008g02210.1
Vaccinium___Vadar_g46178.t1
Vaccinium___Vadar_g11124.t1
Vaccinium___Vadar_g11121.t1
Hydrangea___Hyque.10G040000.1.p
Hydrangea___Hyque.02G073200.1.p
Lactuca___Lsat_1_v5_gn_4_37701.1
Helianthus___HanXRQChr14g0455241
Helianthus___HanXRQChr06g0180321
Daucus___DCAR_004761
Kalanchoe___Kalax.0720s0014.1.p
Kalanchoe___Kalax.0178s0037.1.p
Solanum___Solyc01g079430.2.1
Olea___Oeu056508.1
Mimulus___Migut.J01495.1.p
Beta___EL10Ac1g01520.1
Malus___MD14G1152200
Cucumis___Cucsa.271190.1|PACid_16973498
Quercus___Qurub.02G104700.1
F-Pvulgaris___Phvul.001G029400.1.p
F-Carietinum___Ca_01684
F-Prosopis___Palba_XP_028777381.1
F-Lleucocephala___GDRZ01005485.1.p1
F-Lleucocephala___GDRZ01166711.1.p1
F-Akoa___GBYE01013507.1.p1
F-Acol___TRINITY_DN8521_c0_g2_i1.p1_3prime_partial
F-Acol___TRINITY_DN8521_c0_g1_i44.p1_5prime_partial
F-Acol___TRINITY_DN8521_c0_g1_i60.p1_complete
F-Acol___TRINITY_DN8521_c0_g1_i56.p1_3prime_partial
F-Acol___TRINITY_DN8521_c0_g1_i31.p1_complete
F-Acol___TRINITY_DN8521_c0_g1_i24.p1_complete
Citrus___orange1.1g037838m|PACid_18125987
Manihot___Manes.03G056000.2.p
Arabidopsis-AT5G21970.1|PACid_19668148
Musa___GSMUA_Achr10P17440_001

**OG0008357**

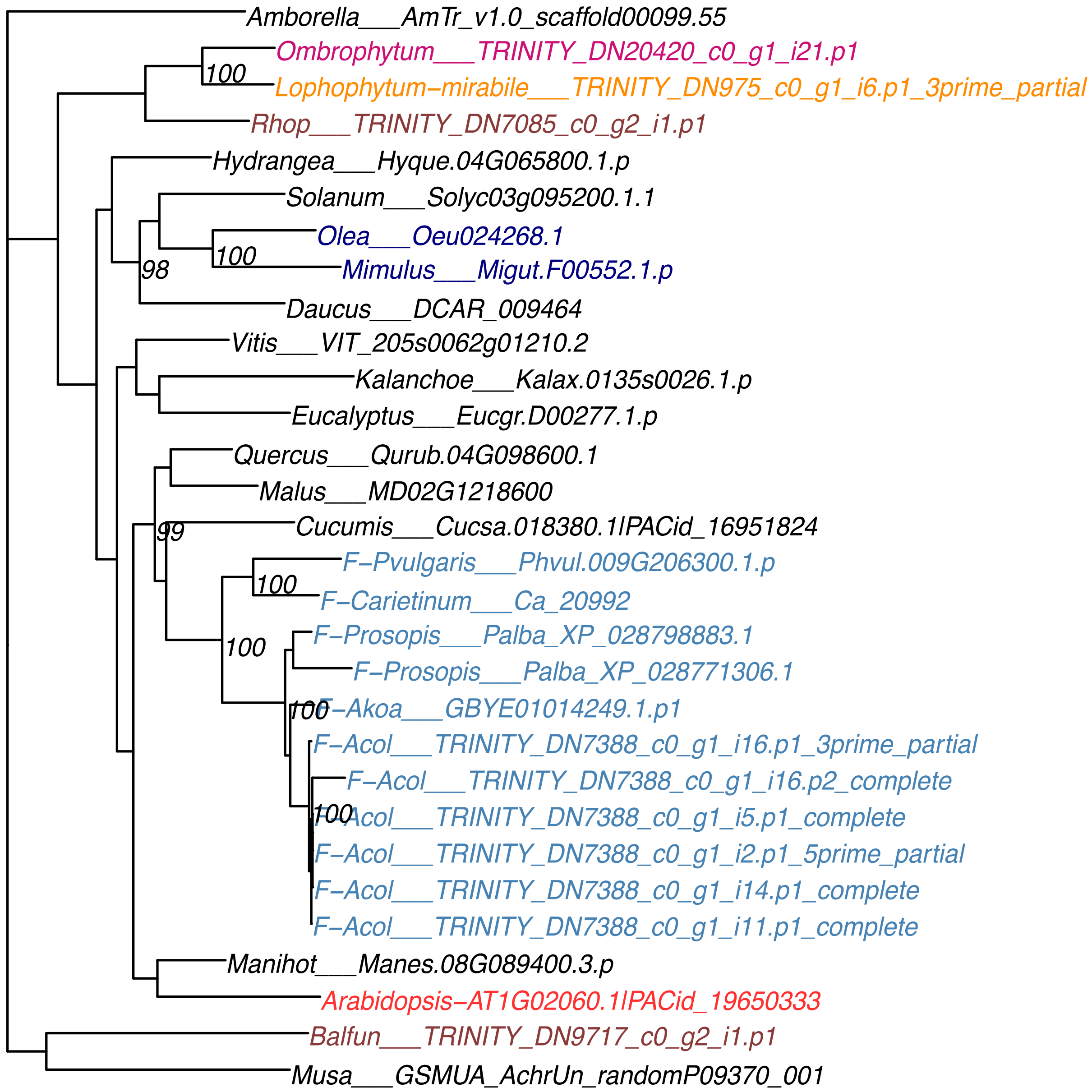
Amborella___AmTr_v1.0_scaffold00099.55
Ombrophytum___TRINITY_DN20420_c0_g1_i21.p1
100
Lophophytum−mirabile___TRINITY_DN975_c0_g1_i6.p1_3prime_partial
Rhop___TRINITY_DN7085_c0_g2_i1.p1
Hydrangea___Hyque.04G065800.1.p
Solanum___Solyc03g095200.1.1
Olea___Oeu024268.1
100
Mimulus___Migut.F00552.1.p
98
Daucus___DCAR_009464
Vitis___VIT_205s0062g01210.2
Kalanchoe___Kalax.0135s0026.1.p
Eucalyptus___Eucgr.D00277.1.p
Quercus___Qurub.04G098600.1
Malus___MD02G1218600
Cucumis___Cucsa.018380.1|PACid_16951824
99
F−Pvulgaris___Phvul.009G206300.1.p
100
F−Carietinum___Ca_20992
100
F−Prosopis___Palba_XP_028798883.1
F−Prosopis___Palba_XP_028771306.1
100
F−Akoa___GBYE01014249.1.p1
F−Acol___TRINITY_DN7388_c0_g1_i16.p1_3prime_partial
F−Acol___TRINITY_DN7388_c0_g1_i16.p2_complete
100
F−Acol___TRINITY_DN7388_c0_g1_i5.p1_complete
F−Acol___TRINITY_DN7388_c0_g1_i2.p1_5prime_partial
F−Acol___TRINITY_DN7388_c0_g1_i14.p1_complete
F−Acol___TRINITY_DN7388_c0_g1_i11.p1_complete
Manihot___Manes.08G089400.3.p
Arabidopsis−AT1G02060.1|PACid_19650333
Balfun___TRINITY_DN9717_c0_g2_i1.p1
Musa___GSMUA_AchrUn_randomP09370_001

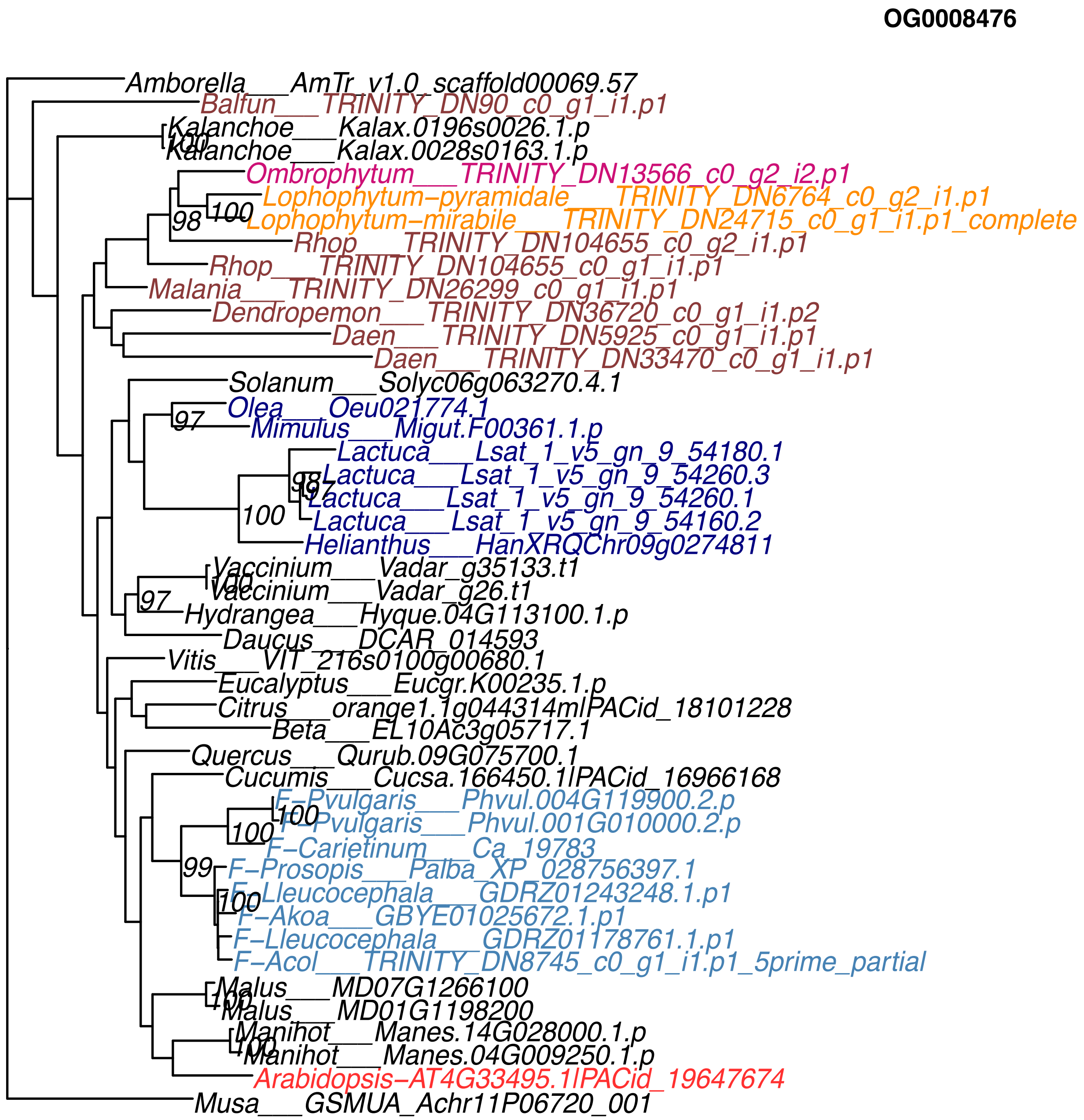
OG0008476
Amborella___AmTr_v1.0_scaffold00069.57
Balfun___TRINITY_DN90_c0_g1_i1.p1
Kalanchoe___Kalax.0196s0026.1.p
100
Kalanchoe___Kalax.0028s0163.1.p
Ombrophytum___TRINITY_DN13566_c0_g2_i2.p1
Lophophytum−pyramidale___TRINITY_DN6764_c0_g2_i1.p1
98
100
Lophophytum−mirabile___TRINITY_DN24715_c0_g1_i1.p1_complete
Rhop___TRINITY_DN104655_c0_g2_i1.p1
Rhop___TRINITY_DN104655_c0_g1_i1.p1
Malania___TRINITY_DN26299_c0_g1_i1.p1
Dendropemon___TRINITY_DN36720_c0_g1_i1.p2
Daen___TRINITY_DN5925_c0_g1_i1.p1
Daen___TRINITY_DN33470_c0_g1_i1.p1
Solanum___Solyc06g063270.4.1
Olea___Oeu021774.1
97
Mimulus___Migut.F00361.1.p
Lactuca___Lsat_1_v5_gn_9_54180.1
Lactuca___Lsat_1_v5_gn_9_54260.3
98
97
Lactuca___Lsat_1_v5_gn_9_54260.1
100
Lactuca___Lsat_1_v5_gn_9_54160.2
Helianthus___HanXRQChr09g0274811
Vaccinium___Vadar_g35133.t1
100
Vaccinium___Vadar_g26.t1
97
Hydrangea___Hyque.04G113100.1.p
Daucus___DCAR_014593
Vitis___VIT_216s0100g00680.1
Eucalyptus___Eucgr.K00235.1.p
Citrus___orange1.1g044314mIPACid_18101228
Beta___EL10Ac3g05717.1
Quercus___Qurub.09G075700.1
Cucumis___Cucsa.166450.1IPACid_16966168
F−Pvulgaris___Phvul.004G119900.2.p
100
F−Pvulgaris___Phvul.001G010000.2.p
100
F−Carietinum___Ca_19783
99
F−Prosopis___Palba_XP_028756397.1
F−Lleucocephala___GDRZ01243248.1.p1
100
F−Akoa___GBYE01025672.1.p1
F−Lleucocephala___GDRZ01178761.1.p1
F−Acol___TRINITY_DN8745_c0_g1_i1.p1_5prime_partial
Malus___MD07G1266100
100
Malus___MD01G1198200
Manihot___Manes.14G028000.1.p
100
Manihot___Manes.04G009250.1.p
Arabidopsis−AT4G33495.1IPACid_19647674
Musa___GSMUA_Achr11P06720_001

**OG0008583**

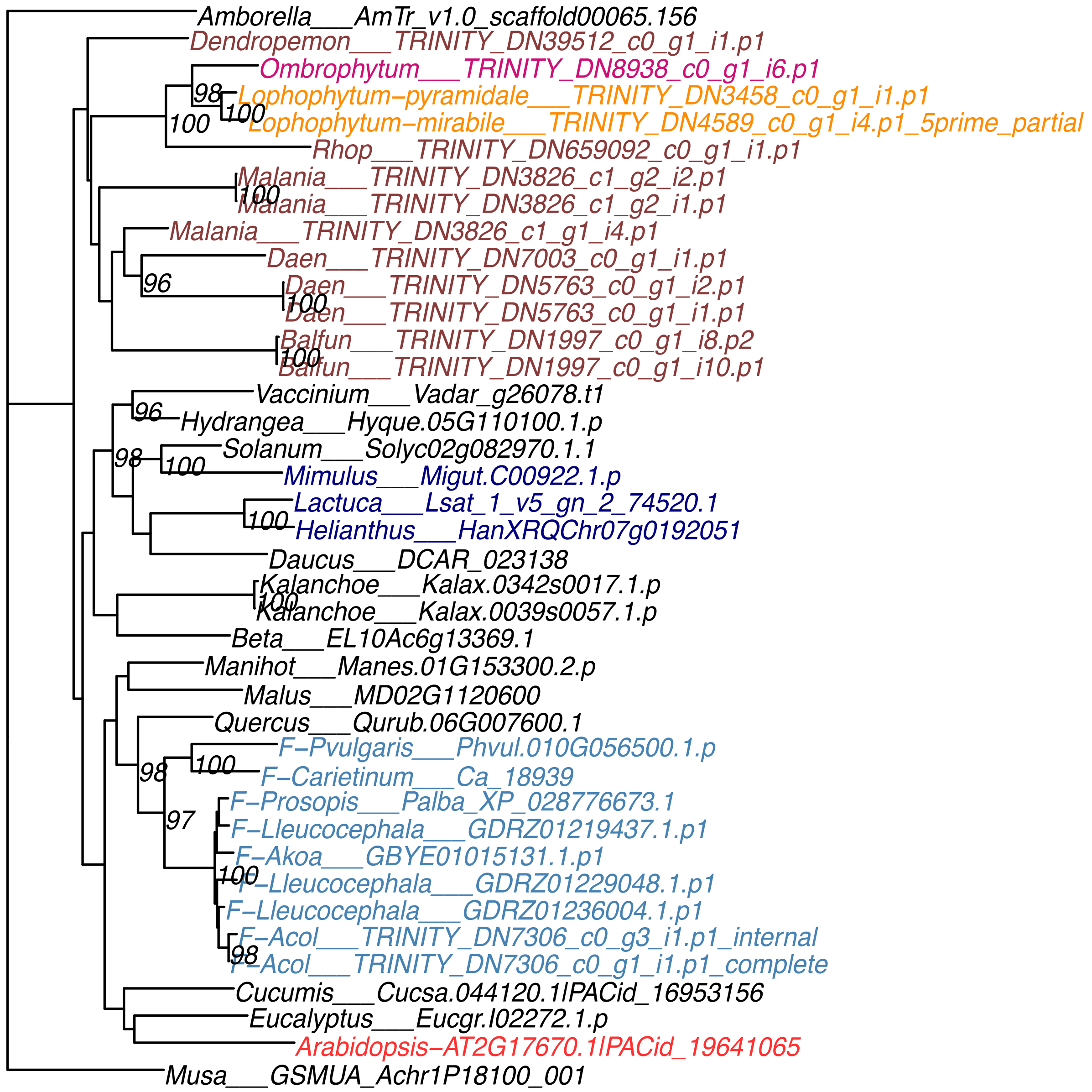
Amborella___AmTr_v1.0_scaffold00065.156
Dendropemon___TRINITY_DN39512_c0_g1_i1.p1
Ombrophytum___TRINITY_DN8938_c0_g1_i6.p1
Lophophytum−pyramidale___TRINITY_DN3458_c0_g1_i1.p1
Lophophytum−mirabile___TRINITY_DN4589_c0_g1_i4.p1_5prime_partial
Rhop___TRINITY_DN659092_c0_g1_i1.p1
Malania___TRINITY_DN3826_c1_g2_i2.p1
Malania___TRINITY_DN3826_c1_g2_i1.p1
Malania___TRINITY_DN3826_c1_g1_i4.p1
Daen___TRINITY_DN7003_c0_g1_i1.p1
Daen___TRINITY_DN5763_c0_g1_i2.p1
Daen___TRINITY_DN5763_c0_g1_i1.p1
Balfun___TRINITY_DN1997_c0_g1_i8.p2
Balfun___TRINITY_DN1997_c0_g1_i10.p1
Vaccinium___Vadar_g26078.t1
Hydrangea___Hyque.05G110100.1.p
Solanum___Solyc02g082970.1.1
Mimulus___Migut.C00922.1.p
Lactuca___Lsat_1_v5_gn_2_74520.1
Helianthus___HanXRQChr07g0192051
Daucus___DCAR_023138
Kalanchoe___Kalax.0342s0017.1.p
Kalanchoe___Kalax.0039s0057.1.p
Beta___EL10Ac6g13369.1
Manihot___Manes.01G153300.2.p
Malus___MD02G1120600
Quercus___Qurub.06G007600.1
F−Pvulgaris___Phvul.010G056500.1.p
F−Carietinum___Ca_18939
F−Prosopis___Palba_XP_028776673.1
F−Lleucocephala___GDRZ01219437.1.p1
F−Akoa___GBYE01015131.1.p1
F−Lleucocephala___GDRZ01229048.1.p1
F−Lleucocephala___GDRZ01236004.1.p1
F−Acol___TRINITY_DN7306_c0_g3_i1.p1_internal
F−Acol___TRINITY_DN7306_c0_g1_i1.p1_complete
Cucumis___Cucsa.044120.1lPACid_16953156
Eucalyptus___Eucgr.I02272.1.p
Arabidopsis−AT2G17670.1lPACid_19641065
Musa___GSMUA_Achr1P18100_001
98
100
100
100
96
100
100
96
98
100
100
100
100
98
97
100
98

**OG0008743**

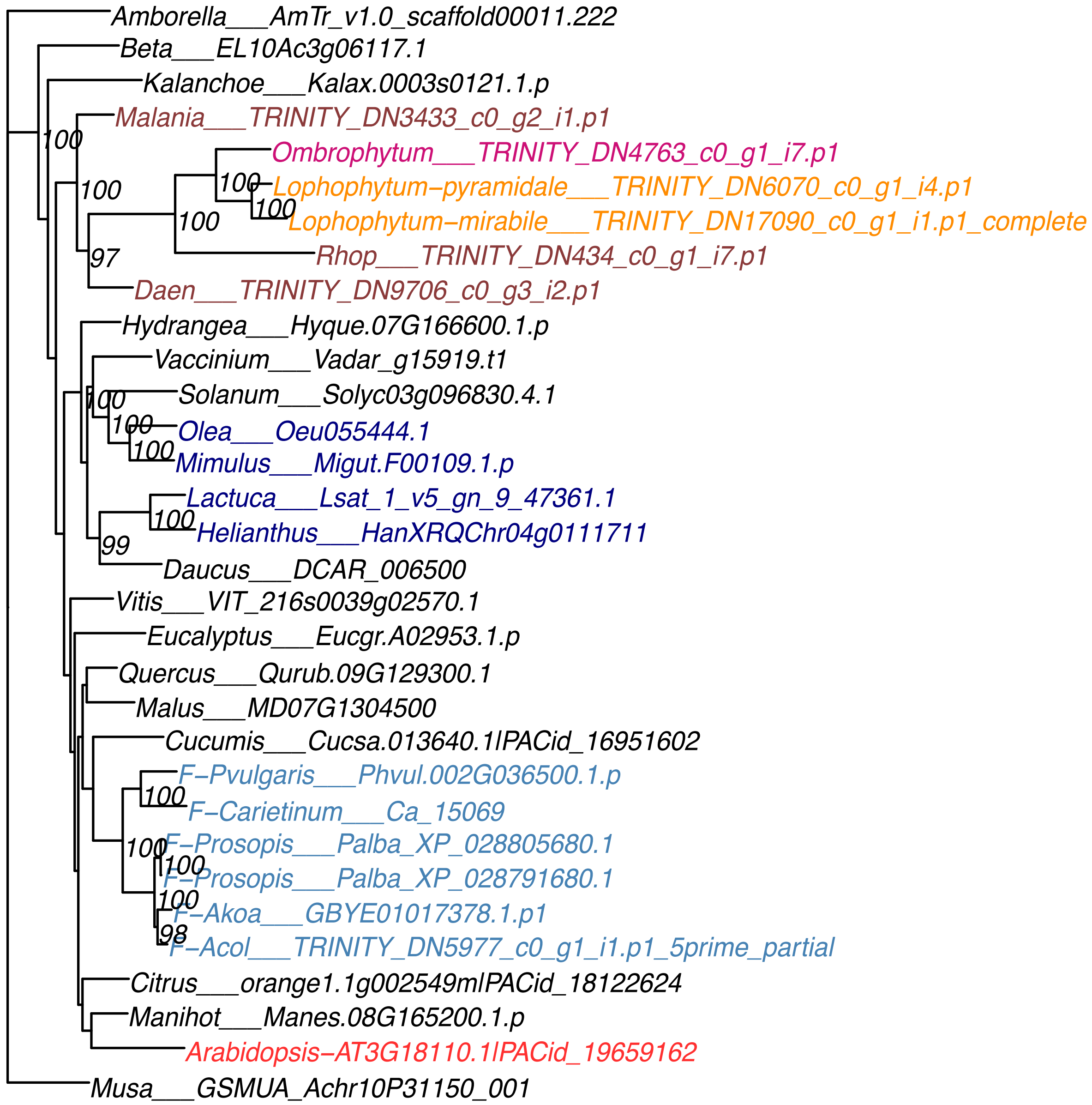
Amborella___AmTr_v1.0_scaffold00011.222
Beta___EL10Ac3g06117.1
Kalanchoe___Kalax.0003s0121.1.p
Malania___TRINITY_DN3433_c0_g2_i1.p1
Ombrophytum___TRINITY_DN4763_c0_g1_i7.p1
Lophophytum−pyramidale___TRINITY_DN6070_c0_g1_i4.p1
Lophophytum−mirabile___TRINITY_DN17090_c0_g1_i1.p1_complete
Rhop___TRINITY_DN434_c0_g1_i7.p1
Daen___TRINITY_DN9706_c0_g3_i2.p1
Hydrangea___Hyque.07G166600.1.p
Vaccinium___Vadar_g15919.t1
Solanum___Solyc03g096830.4.1
Olea___Oeu055444.1
Mimulus___Migut.F00109.1.p
Lactuca___Lsat_1_v5_gn_9_47361.1
Helianthus___HanXRQChr04g0111711
Daucus___DCAR_006500
Vitis___VIT_216s0039g02570.1
Eucalyptus___Eucgr.A02953.1.p
Quercus___Qurub.09G129300.1
Malus___MD07G1304500
Cucumis___Cucsa.013640.1|PACid_16951602
F−Pvulgaris___Phvul.002G036500.1.p
F−Carietinum___Ca_15069
F−Prosopis___Palba_XP_028805680.1
F−Prosopis___Palba_XP_028791680.1
F−Akoa___GBYE01017378.1.p1
F−Acol___TRINITY_DN5977_c0_g1_i1.p1_5prime_partial
Citrus___orange1.1g002549m|PACid_18122624
Manihot___Manes.08G165200.1.p
Arabidopsis−AT3G18110.1|PACid_19659162
Musa___GSMUA_Achr10P31150_001
100
100
100
100
100
97
100
100
100
100
99
100
100
100
100
98

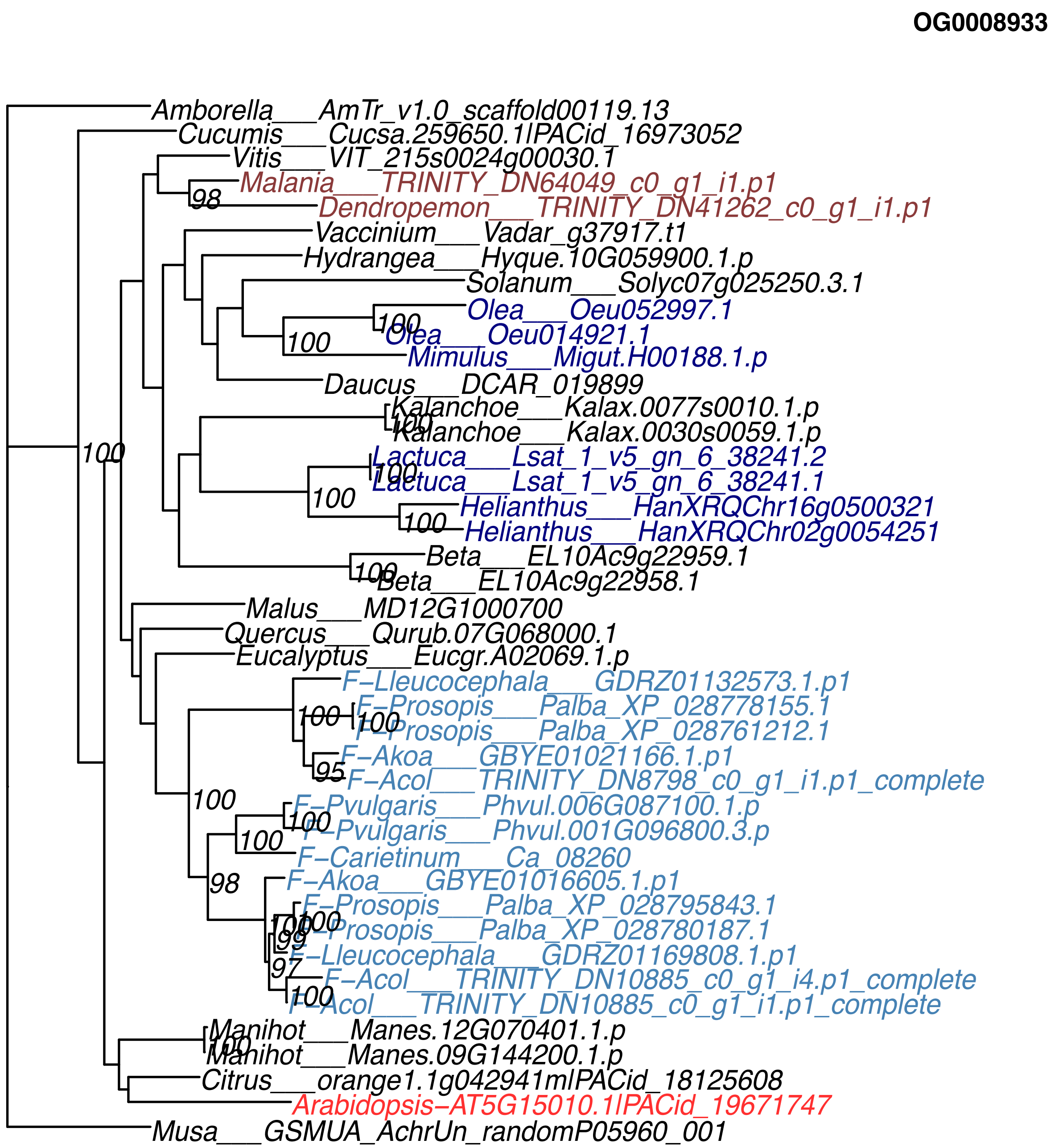
OG0008933
Amborella___AmTr_v1.0_scaffold00119.13
Cucumis___Cucsa.259650.1|PACid_16973052
Vitis___VIT_215s0024g00030.1
Malania___TRINITY_DN64049_c0_g1_i1.p1
98
Dendropemon___TRINITY_DN41262_c0_g1_i1.p1
Vaccinium___Vadar_g37917.t1
Hydrangea___Hyque.10G059900.1.p
Solanum___Solyc07g025250.3.1
Olea___Oeu052997.1
100
Olea___Oeu014921.1
100
Mimulus___Migut.H00188.1.p
Daucus___DCAR_019899
Kalanchoe___Kalax.0077s0010.1.p
100
Kalanchoe___Kalax.0030s0059.1.p
100
Lactuca___Lsat_1_v5_gn_6_38241.2
100
Lactuca___Lsat_1_v5_gn_6_38241.1
100
Helianthus___HanXRQChr16g0500321
100
Helianthus___HanXRQChr02g0054251
Beta___EL10Ac9g22959.1
100
Beta___EL10Ac9g22958.1
Malus___MD12G1000700
Quercus___Qurub.07G068000.1
Eucalyptus___Eucgr.A02069.1.p
F−Lleucocephala___GDRZ01132573.1.p1
100
F−Prosopis___Palba_XP_028778155.1
100
F−Prosopis___Palba_XP_028761212.1
F−Akoa___GBYE01021166.1.p1
95
F−Acol___TRINITY_DN8798_c0_g1_i1.p1_complete
100
F−Pvulgaris___Phvul.006G087100.1.p
100
F−Pvulgaris___Phvul.001G096800.3.p
100
F−Carietinum___Ca_08260
98
F−Akoa___GBYE01016605.1.p1
F−Prosopis___Palba_XP_028795843.1
100
100
F−Prosopis___Palba_XP_028780187.1
99
F−Lleucocephala___GDRZ01169808.1.p1
97
F−Acol___TRINITY_DN10885_c0_g1_i4.p1_complete
100
F−Acol___TRINITY_DN10885_c0_g1_i1.p1_complete
Manihot___Manes.12G070401.1.p
100
Manihot___Manes.09G144200.1.p
Citrus___orange1.1g042941m|PACid_18125608
Arabidopsis−AT5G15010.1|PACid_19671747
Musa___GSMUA_AchrUn_randomP05960_001

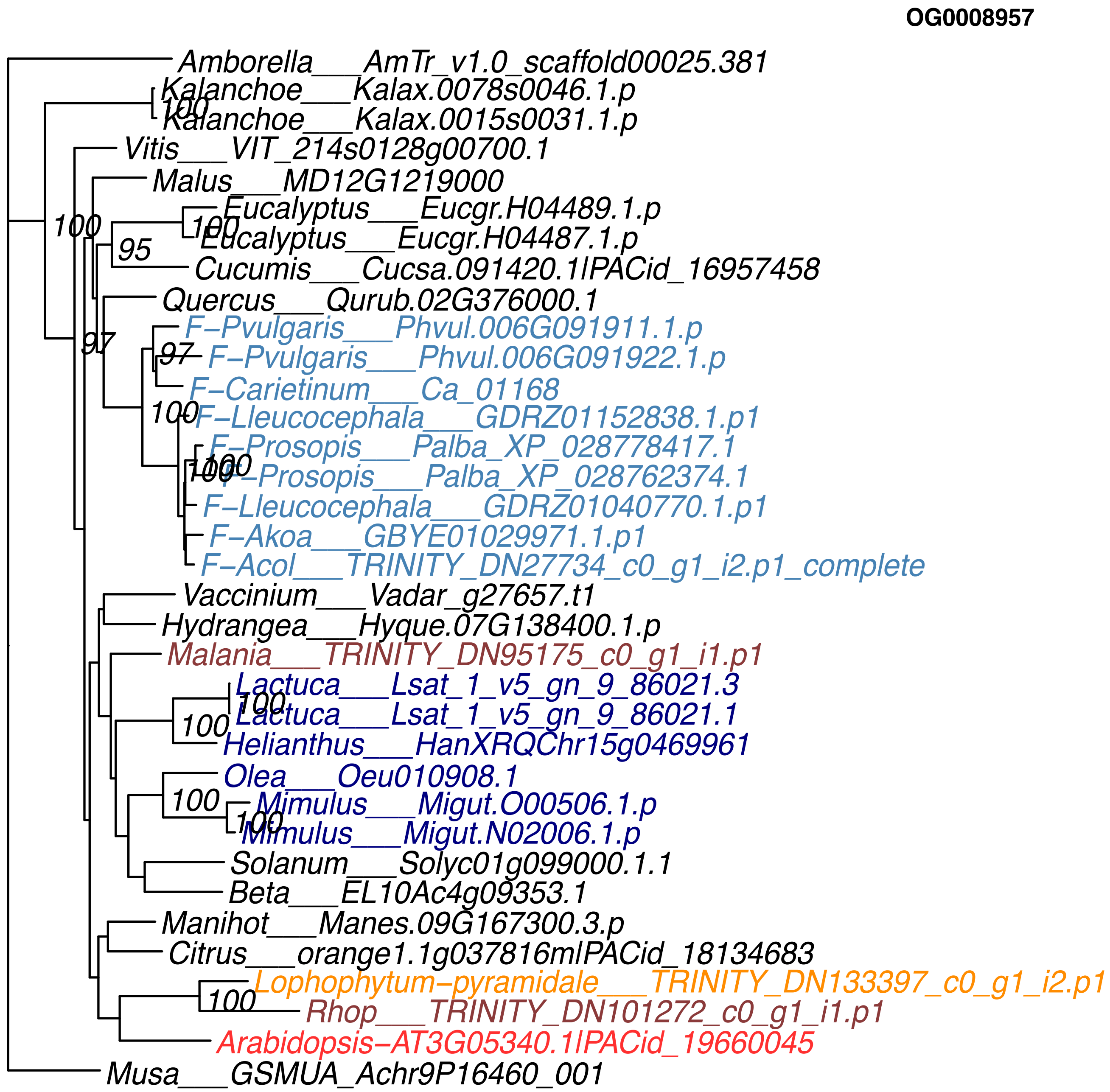

OG0008957
Amborella___AmTr_v1.0_scaffold00025.381
Kalanchoe___Kalax.0078s0046.1.p
100
Kalanchoe___Kalax.0015s0031.1.p
Vitis___VIT_214s0128g00700.1
Malus___MD12G1219000
Eucalyptus___Eucgr.H04489.1.p
100
Eucalyptus___Eucgr.H04487.1.p
100
95
Cucumis___Cucsa.091420.1|PACid_16957458
Quercus___Qurub.02G376000.1
F−Pvulgaris___Phvul.006G091911.1.p
97
97
F−Pvulgaris___Phvul.006G091922.1.p
F−Carietinum___Ca_01168
100
F−Lleucocephala___GDRZ01152838.1.p1
F−Prosopis___Palba_XP_028778417.1
100
100
F−Prosopis___Palba_XP_028762374.1
F−Lleucocephala___GDRZ01040770.1.p1
F−Akoa___GBYE01029971.1.p1
F−Acol___TRINITY_DN27734_c0_g1_i2.p1_complete
Vaccinium___Vadar_g27657.t1
Hydrangea___Hyque.07G138400.1.p
Malania___TRINITY_DN95175_c0_g1_i1.p1
Lactuca___Lsat_1_v5_gn_9_86021.3
100
Lactuca___Lsat_1_v5_gn_9_86021.1
100
Helianthus___HanXRQChr15g0469961
Olea___Oeu010908.1
100
Mimulus___Migut.O00506.1.p
100
Mimulus___Migut.N02006.1.p
Solanum___Solyc01g099000.1.1
Beta___EL10Ac4g09353.1
Manihot___Manes.09G167300.3.p
Citrus___orange1.1g037816m|PACid_18134683
Lophophytum−pyramidale___TRINITY_DN133397_c0_g1_i2.p1
100
Rhop___TRINITY_DN101272_c0_g1_i1.p1
Arabidopsis−AT3G05340.1|PACid_19660045
Musa___GSMUA_Achr9P16460_001

**OG0008987**

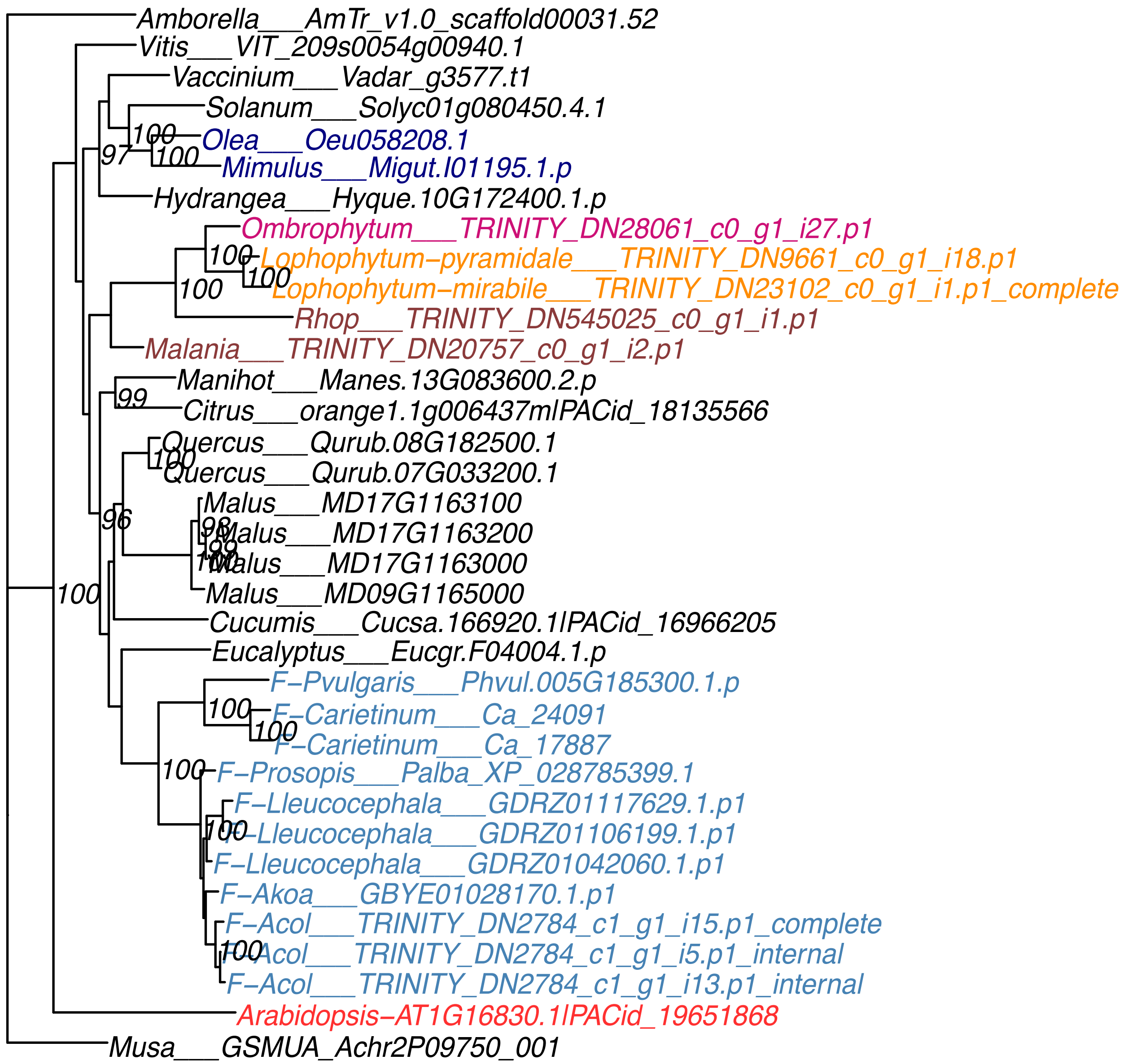

Amborella___AmTr_v1.0_scaffold00031.52
Vitis___VIT_209s0054g00940.1
Vaccinium___Vadar_g3577.t1
Solanum___Solyc01g080450.4.1
100
Olea___Oeu058208.1
97
100
Mimulus___Migut.I01195.1.p
Hydrangea___Hyque.10G172400.1.p
Ombrophytum___TRINITY_DN28061_c0_g1_i27.p1
100
Lophophytum−pyramidale___TRINITY_DN9661_c0_g1_i18.p1
100
100
Lophophytum−mirabile___TRINITY_DN23102_c0_g1_i1.p1_complete
Rhop___TRINITY_DN545025_c0_g1_i1.p1
Malania___TRINITY_DN20757_c0_g1_i2.p1
Manihot___Manes.13G083600.2.p
99
Citrus___orange1.1g006437mIPACid_18135566
Quercus___Qurub.08G182500.1
100
Quercus___Qurub.07G033200.1
Malus___MD17G1163100
96
98
Malus___MD17G1163200
99
100
Malus___MD17G1163000
Malus___MD09G1165000
100
Cucumis___Cucsa.166920.1IPACid_16966205
Eucalyptus___Eucgr.F04004.1.p
F−Pvulgaris___Phvul.005G185300.1.p
100
F−Carietinum___Ca_24091
100
F−Carietinum___Ca_17887
100
F−Prosopis___Palba_XP_028785399.1
F−Lleucocephala___GDRZ01117629.1.p1
100
F−Lleucocephala___GDRZ01106199.1.p1
F−Lleucocephala___GDRZ01042060.1.p1
F−Akoa___GBYE01028170.1.p1
F−Acol___TRINITY_DN2784_c1_g1_i15.p1_complete
100
F−Acol___TRINITY_DN2784_c1_g1_i5.p1_internal
F−Acol___TRINITY_DN2784_c1_g1_i13.p1_internal
Arabidopsis−AT1G16830.1IPACid_19651868
Musa___GSMUA_Achr2P09750_001

**OG0009083**

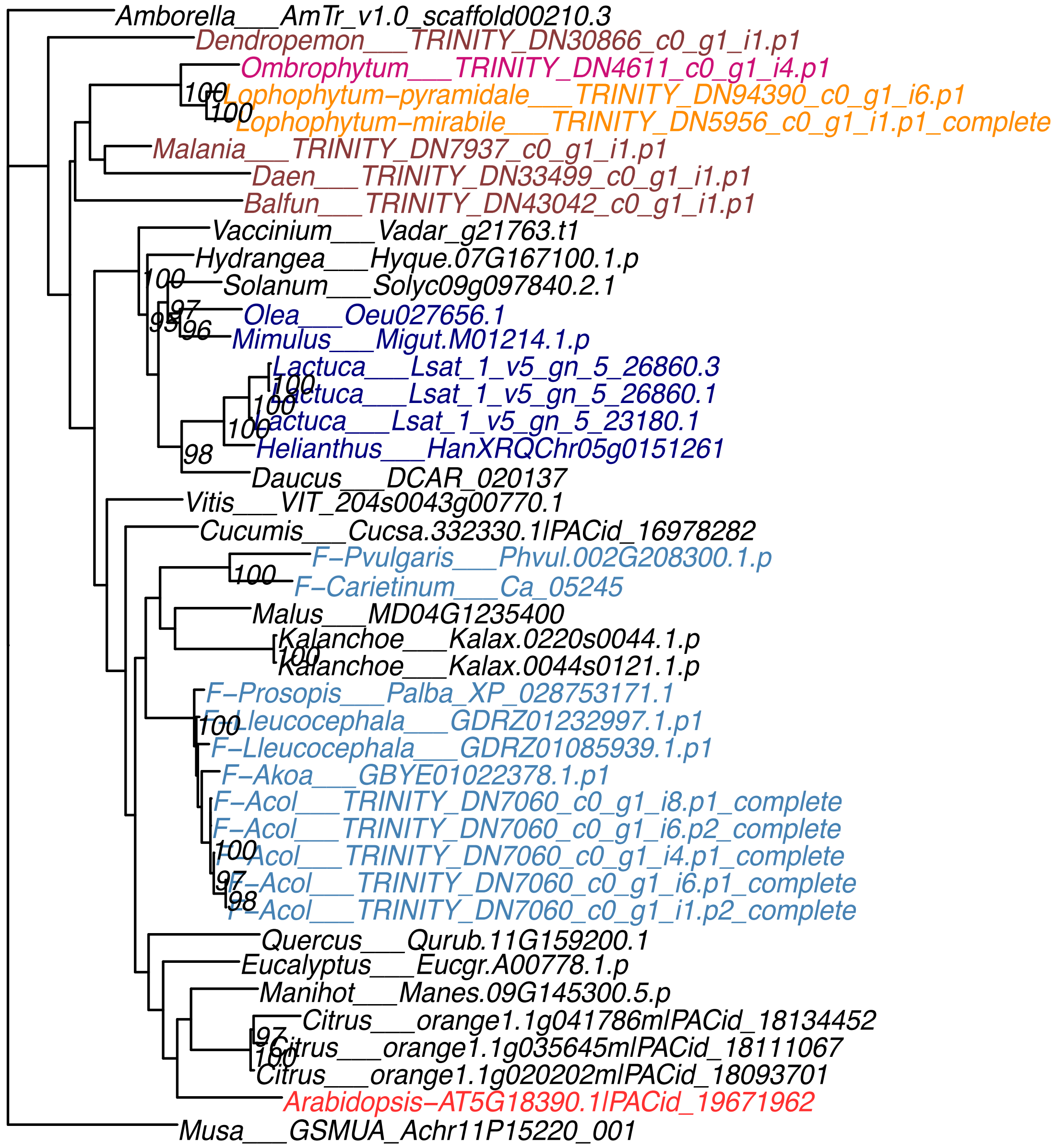
Amborella___AmTr_v1.0_scaffold00210.3
Dendropemon___TRINITY_DN30866_c0_g1_i1.p1
Ombrophytum___TRINITY_DN4611_c0_g1_i4.p1
Lophophytum−pyramidale___TRINITY_DN94390_c0_g1_i6.p1
Lophophytum−mirabile___TRINITY_DN5956_c0_g1_i1.p1_complete
Malania___TRINITY_DN7937_c0_g1_i1.p1
Daen___TRINITY_DN33499_c0_g1_i1.p1
Balfun___TRINITY_DN43042_c0_g1_i1.p1
Vaccinium___Vadar_g21763.t1
Hydrangea___Hyque.07G167100.1.p
Solanum___Solyc09g097840.2.1
Olea___Oeu027656.1
Mimulus___Migut.M01214.1.p
Lactuca___Lsat_1_v5_gn_5_26860.3
Lactuca___Lsat_1_v5_gn_5_26860.1
Lactuca___Lsat_1_v5_gn_5_23180.1
Helianthus___HanXRQChr05g0151261
Daucus___DCAR_020137
Vitis___VIT_204s0043g00770.1
Cucumis___Cucsa.332330.1lPACid_16978282
F−Pvulgaris___Phvul.002G208300.1.p
F−Carietinum___Ca_05245
Malus___MD04G1235400
Kalanchoe___Kalax.0220s0044.1.p
Kalanchoe___Kalax.0044s0121.1.p
F−Prosopis___Palba_XP_028753171.1
F−Lleucocephala___GDRZ01232997.1.p1
F−Lleucocephala___GDRZ01085939.1.p1
F−Akoa___GBYE01022378.1.p1
F−Acol___TRINITY_DN7060_c0_g1_i8.p1_complete
F−Acol___TRINITY_DN7060_c0_g1_i6.p2_complete
F−Acol___TRINITY_DN7060_c0_g1_i4.p1_complete
F−Acol___TRINITY_DN7060_c0_g1_i6.p1_complete
F−Acol___TRINITY_DN7060_c0_g1_i1.p2_complete
Quercus___Qurub.11G159200.1
Eucalyptus___Eucgr.A00778.1.p
Manihot___Manes.09G145300.5.p
Citrus___orange1.1g041786mlPACid_18134452
Citrus___orange1.1g035645mlPACid_18111067
Citrus___orange1.1g020202mlPACid_18093701
Arabidopsis−AT5G18390.1lPACid_19671962
Musa___GSMUA_Achr11P15220_001
100
100
100
95
97
96
100
100
100
98
100
100
100
100
97
98
97
100

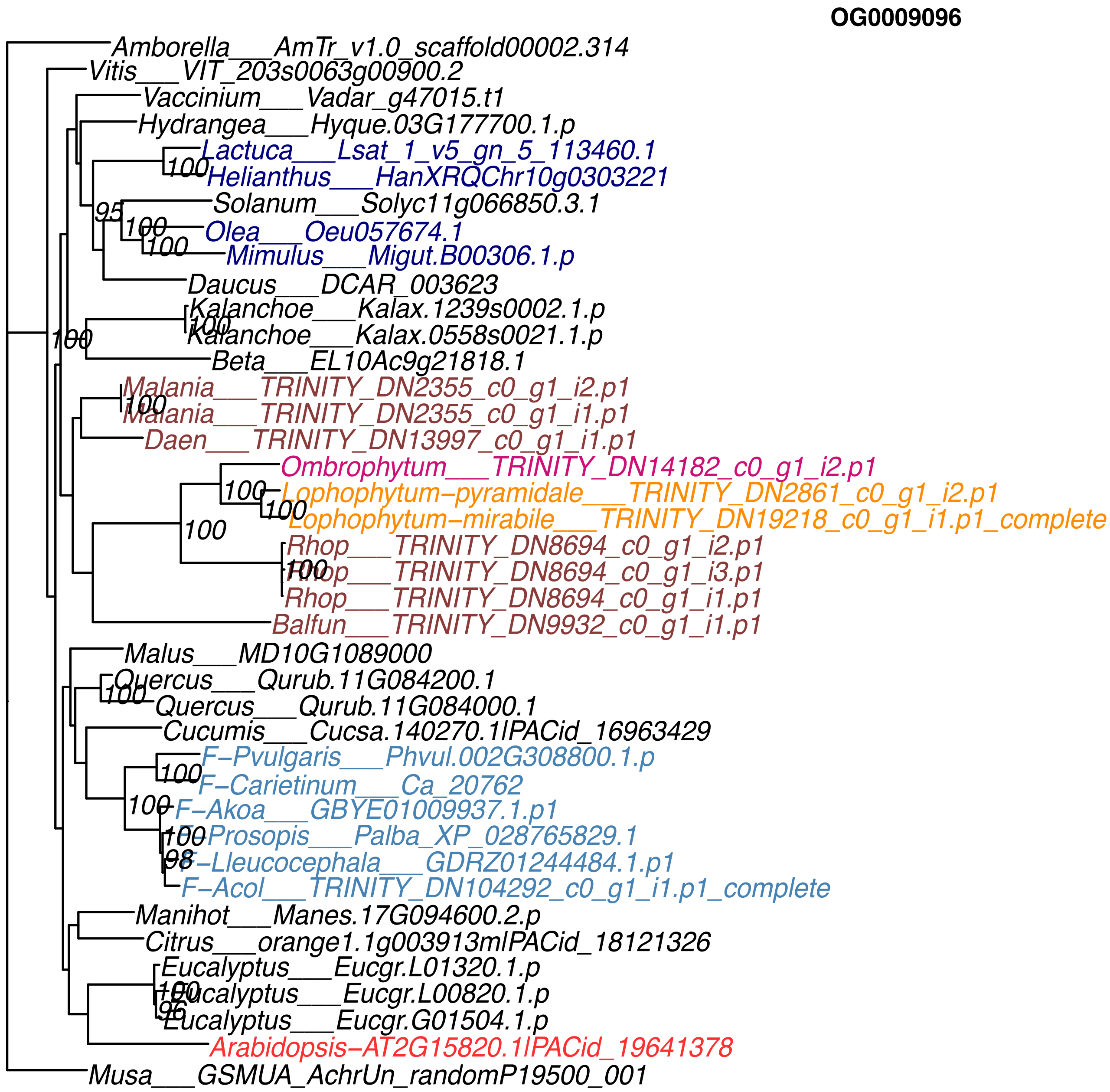

OG0009096
Amborella___AmTr_v1.0_scaffold00002.314
Vitis___VIT_203s0063g00900.2
Vaccinium___Vadar_g47015.t1
Hydrangea___Hyque.03G177700.1.p
Lactuca___Lsat_1_v5_gn_5_113460.1
Helianthus___HanXRQChr10g0303221
Solanum___Solyc11g066850.3.1
Olea___Oeu057674.1
Mimulus___Migut.B00306.1.p
Daucus___DCAR_003623
Kalanchoe___Kalax.1239s0002.1.p
Kalanchoe___Kalax.0558s0021.1.p
Beta___EL10Ac9g21818.1
Malania___TRINITY_DN2355_c0_g1_i2.p1
Malania___TRINITY_DN2355_c0_g1_i1.p1
Daen___TRINITY_DN13997_c0_g1_i1.p1
Ombrophytum___TRINITY_DN14182_c0_g1_i2.p1
Lophophytum−pyramidale___TRINITY_DN2861_c0_g1_i2.p1
Lophophytum−mirabile___TRINITY_DN19218_c0_g1_i1.p1_complete
Rhop___TRINITY_DN8694_c0_g1_i2.p1
Rhop___TRINITY_DN8694_c0_g1_i3.p1
Rhop___TRINITY_DN8694_c0_g1_i1.p1
Balfun___TRINITY_DN9932_c0_g1_i1.p1
Malus___MD10G1089000
Quercus___Qurub.11G084200.1
Quercus___Qurub.11G084000.1
Cucumis___Cucsa.140270.1lPACid_16963429
F−Pvulgaris___Phvul.002G308800.1.p
F−Carietinum___Ca_20762
F−Akoa___GBYE01009937.1.p1
F−Prosopis___Palba_XP_028765829.1
F−Lleucocephala___GDRZ01244484.1.p1
F−Acol___TRINITY_DN104292_c0_g1_i1.p1_complete
Manihot___Manes.17G094600.2.p
Citrus___orange1.1g003913mlPACid_18121326
Eucalyptus___Eucgr.L01320.1.p
Eucalyptus___Eucgr.L00820.1.p
Eucalyptus___Eucgr.G01504.1.p
Arabidopsis−AT2G15820.1lPACid_19641378
Musa___GSMUA_AchrUn_randomP19500_001
100
95
100
100
100
100
100
100
100
100
100
100
100
100
100
98
100
96

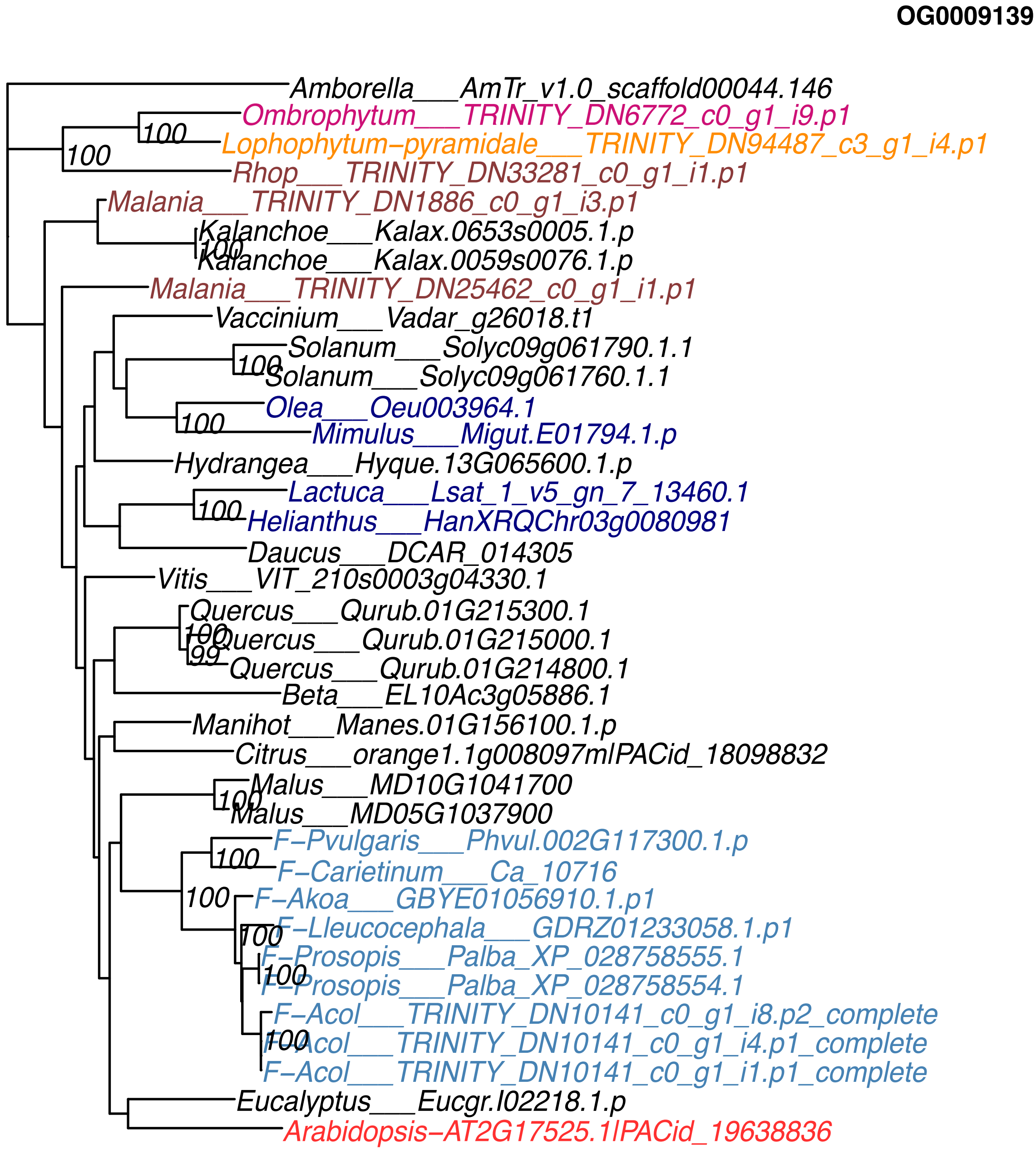
OG0009139
Amborella___AmTr_v1.0_scaffold00044.146
Ombrophytum___TRINITY_DN6772_c0_g1_i9.p1
Lophophytum−pyramidale___TRINITY_DN94487_c3_g1_i4.p1
Rhop___TRINITY_DN33281_c0_g1_i1.p1
Malania___TRINITY_DN1886_c0_g1_i3.p1
Kalanchoe___Kalax.0653s0005.1.p
Kalanchoe___Kalax.0059s0076.1.p
Malania___TRINITY_DN25462_c0_g1_i1.p1
Vaccinium___Vadar_g26018.t1
Solanum___Solyc09g061790.1.1
Solanum___Solyc09g061760.1.1
Olea___Oeu003964.1
Mimulus___Migut.E01794.1.p
Hydrangea___Hyque.13G065600.1.p
Lactuca___Lsat_1_v5_gn_7_13460.1
Helianthus___HanXRQChr03g0080981
Daucus___DCAR_014305
Vitis___VIT_210s0003g04330.1
Quercus___Qurub.01G215300.1
Quercus___Qurub.01G215000.1
Quercus___Qurub.01G214800.1
Beta___EL10Ac3g05886.1
Manihot___Manes.01G156100.1.p
Citrus___orange1.1g008097mIPACid_18098832
Malus___MD10G1041700
Malus___MD05G1037900
F−Pvulgaris___Phvul.002G117300.1.p
F−Carietinum___Ca_10716
F−Akoa___GBYE01056910.1.p1
F−Lleucocephala___GDRZ01233058.1.p1
F−Prosopis___Palba_XP_028758555.1
F−Prosopis___Palba_XP_028758554.1
F−Acol___TRINITY_DN10141_c0_g1_i8.p2_complete
F−Acol___TRINITY_DN10141_c0_g1_i4.p1_complete
F−Acol___TRINITY_DN10141_c0_g1_i1.p1_complete
Eucalyptus___Eucgr.I02218.1.p
Arabidopsis−AT2G17525.1IPACid_19638836
100
100
100
100
100
100
100
99
100
100
100
100
100
100

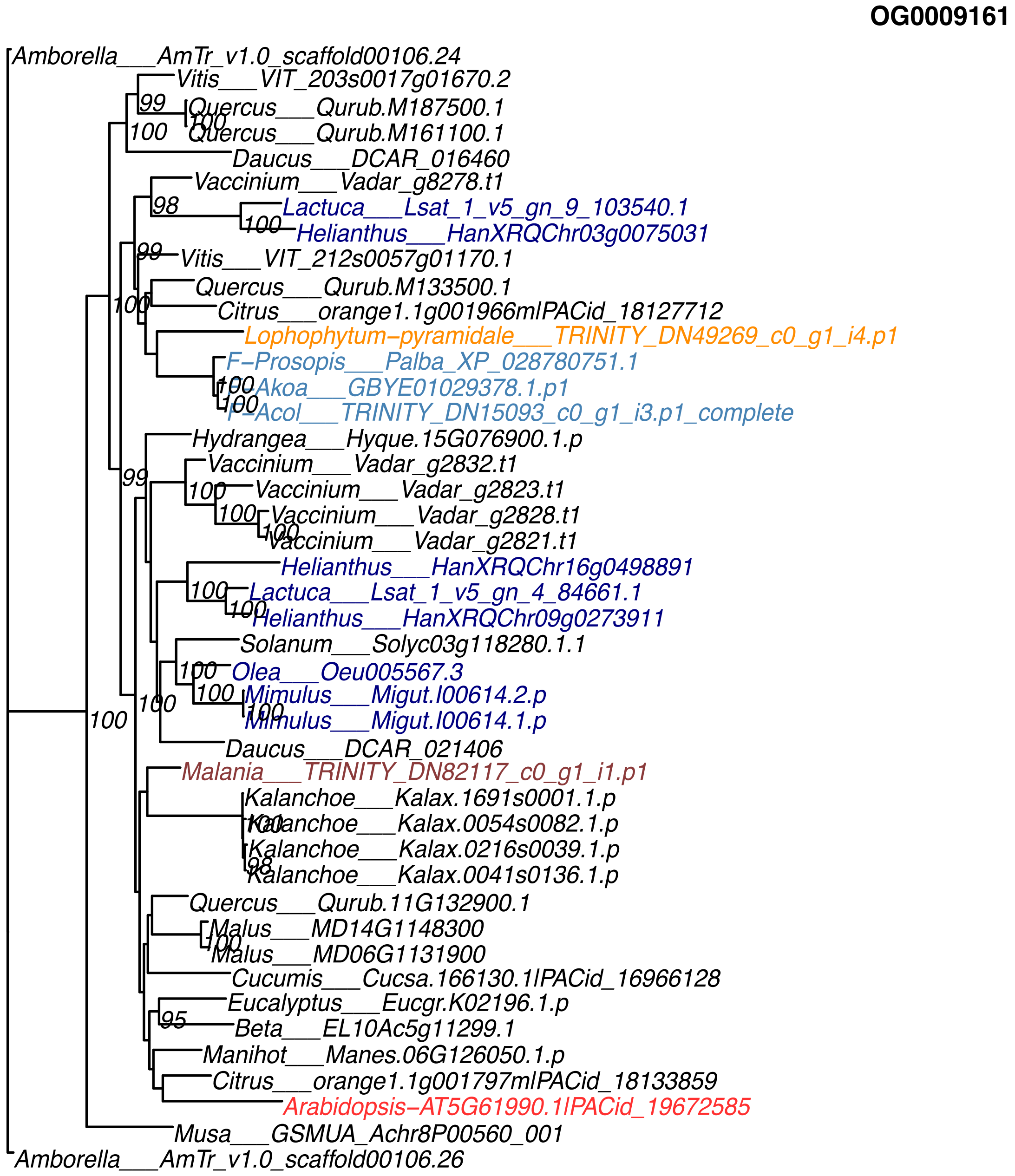

OG0009161
Amborella___AmTr_v1.0_scaffold00106.24
Vitis___VIT_203s0017g01670.2
99
Quercus___Qurub.M187500.1
100
100
Quercus___Qurub.M161100.1
Daucus___DCAR_016460
Vaccinium___Vadar_g8278.t1
98
Lactuca___Lsat_1_v5_gn_9_103540.1
100
Helianthus___HanXRQChr03g0075031
99
Vitis___VIT_212s0057g01170.1
Quercus___Qurub.M133500.1
100
Citrus___orange1.1g001966mIPACid_18127712
Lophophytum-pyramidale___TRINITY_DN49269_c0_g1_i4.p1
F-Prosopis___Palba_XP_028780751.1
100
F-Akoa___GBYE01029378.1.p1
100
F-Acol___TRINITY_DN15093_c0_g1_i3.p1_complete
Hydrangea___Hyque.15G076900.1.p
99
Vaccinium___Vadar_g2832.t1
100
Vaccinium___Vadar_g2823.t1
100
Vaccinium___Vadar_g2828.t1
100
Vaccinium___Vadar_g2821.t1
Helianthus___HanXRQChr16g0498891
100
Lactuca___Lsat_1_v5_gn_4_84661.1
100
Helianthus___HanXRQChr09g0273911
Solanum___Solyc03g118280.1.1
100
Olea___Oeu005567.3
100
Mimulus___Migut.I00614.2.p
100
100
Mimulus___Migut.I00614.1.p
100
Daucus___DCAR_021406
Malania___TRINITY_DN82117_c0_g1_i1.p1
Kalanchoe___Kalax.1691s0001.1.p
100
Kalanchoe___Kalax.0054s0082.1.p
Kalanchoe___Kalax.0216s0039.1.p
98
Kalanchoe___Kalax.0041s0136.1.p
Quercus___Qurub.11G132900.1
Malus___MD14G1148300
100
Malus___MD06G1131900
Cucumis___Cucsa.166130.1IPACid_16966128
Eucalyptus___Eucgr.K02196.1.p
95
Beta___EL10Ac5g11299.1
Manihot___Manes.06G126050.1.p
Citrus___orange1.1g001797mIPACid_18133859
Arabidopsis-AT5G61990.1IPACid_19672585
Musa___GSMUA_Achr8P00560_001
Amborella___AmTr_v1.0_scaffold00106.26

**OG0009186**

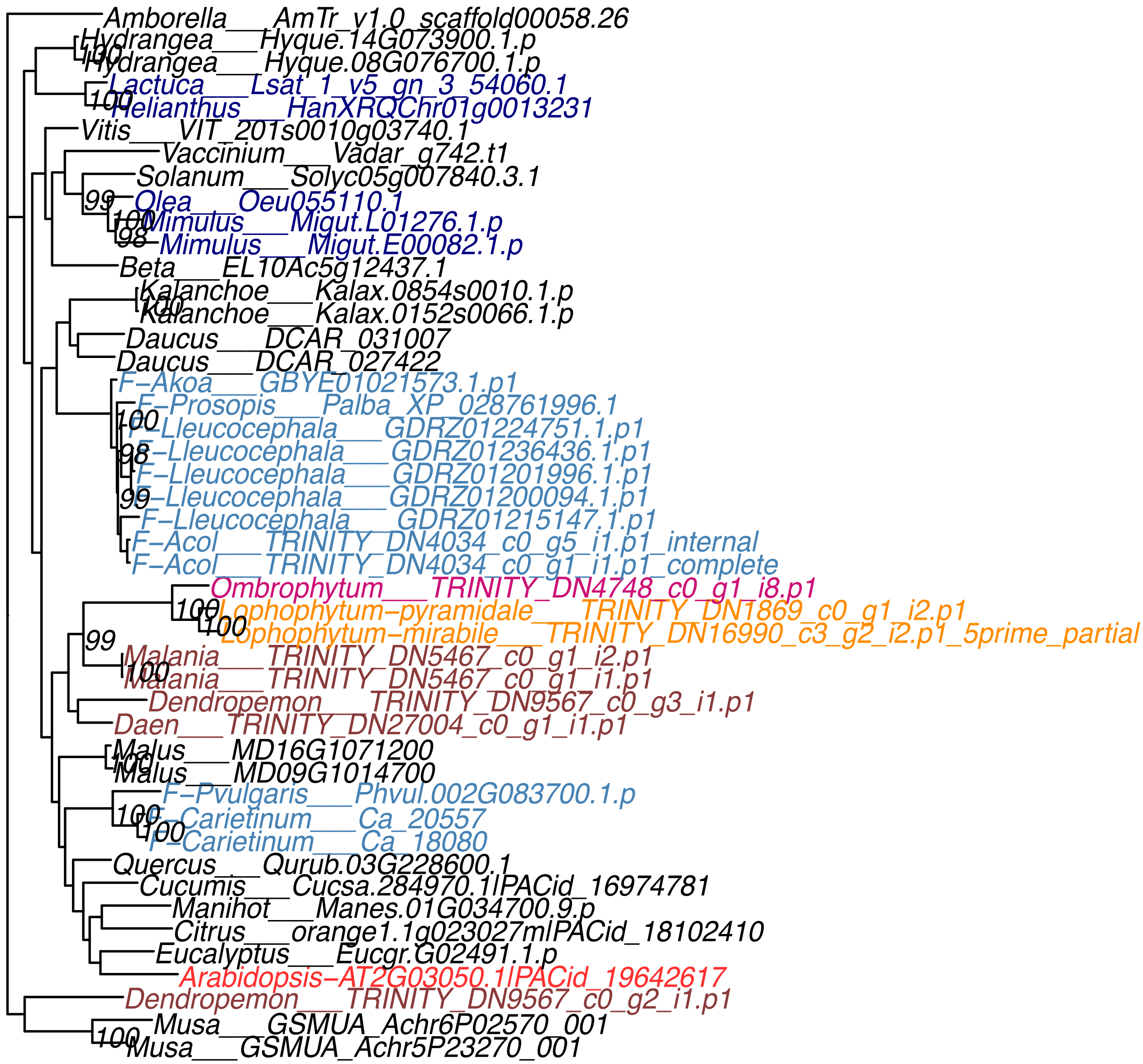
Amborella___AmTr_v1.0_scaffold00058.26
Hydrangea___Hyque.14G073900.1.p
100
Hydrangea___Hyque.08G076700.1.p
Lactuca___Lsat_1_v5_gn_3_54060.1
100
Helianthus___HanXRQChr01g0013231
Vitis___VIT_201s0010g03740.1
Vaccinium___Vadar_g742.t1
Solanum___Solyc05g007840.3.1
99
Olea___Oeu055110.1
100
Mimulus___Migut.L01276.1.p
98
Mimulus___Migut.E00082.1.p
Beta___EL10Ac5g12437.1
Kalanchoe___Kalax.0854s0010.1.p
100
Kalanchoe___Kalax.0152s0066.1.p
Daucus___DCAR_031007
Daucus___DCAR_027422
F–Akoa___GBYE01021573.1.p1
F–Prosopis___Palba_XP_028761996.1
100
F–Lleucocephala___GDRZ01224751.1.p1
98
F–Lleucocephala___GDRZ01236436.1.p1
F–Lleucocephala___GDRZ01201996.1.p1
99
F–Lleucocephala___GDRZ01200094.1.p1
F–Lleucocephala___GDRZ01215147.1.p1
F–Acol___TRINITY_DN4034_c0_g5_i1.p1_internal
F–Acol___TRINITY_DN4034_c0_g1_i1.p1_complete
Ombrophytum___TRINITY_DN4748_c0_g1_i8.p1
100
Lophophytum–pyramidale___TRINITY_DN1869_c0_g1_i2.p1
100
Lophophytum–mirabile___TRINITY_DN16990_c3_g2_i2.p1_5prime_partial
99
Malania___TRINITY_DN5467_c0_g1_i2.p1
100
Malania___TRINITY_DN5467_c0_g1_i1.p1
Dendropemon___TRINITY_DN9567_c0_g3_i1.p1
Daen___TRINITY_DN27004_c0_g1_i1.p1
Malus___MD16G1071200
100
Malus___MD09G1014700
F–Pvulgaris___Phvul.002G083700.1.p
100
F–Carietinum___Ca_20557
100
F–Carietinum___Ca_18080
Quercus___Qurub.03G228600.1
Cucumis___Cucsa.284970.1|PACid_16974781
Manihot___Manes.01G034700.9.p
Citrus___orange1.1g023027m|PACid_18102410
Eucalyptus___Eucgr.G02491.1.p
Arabidopsis–AT2G03050.1|PACid_19642617
Dendropemon___TRINITY_DN9567_c0_g2_i1.p1
Musa___GSMUA_Achr6P02570_001
100
Musa___GSMUA_Achr5P23270_001

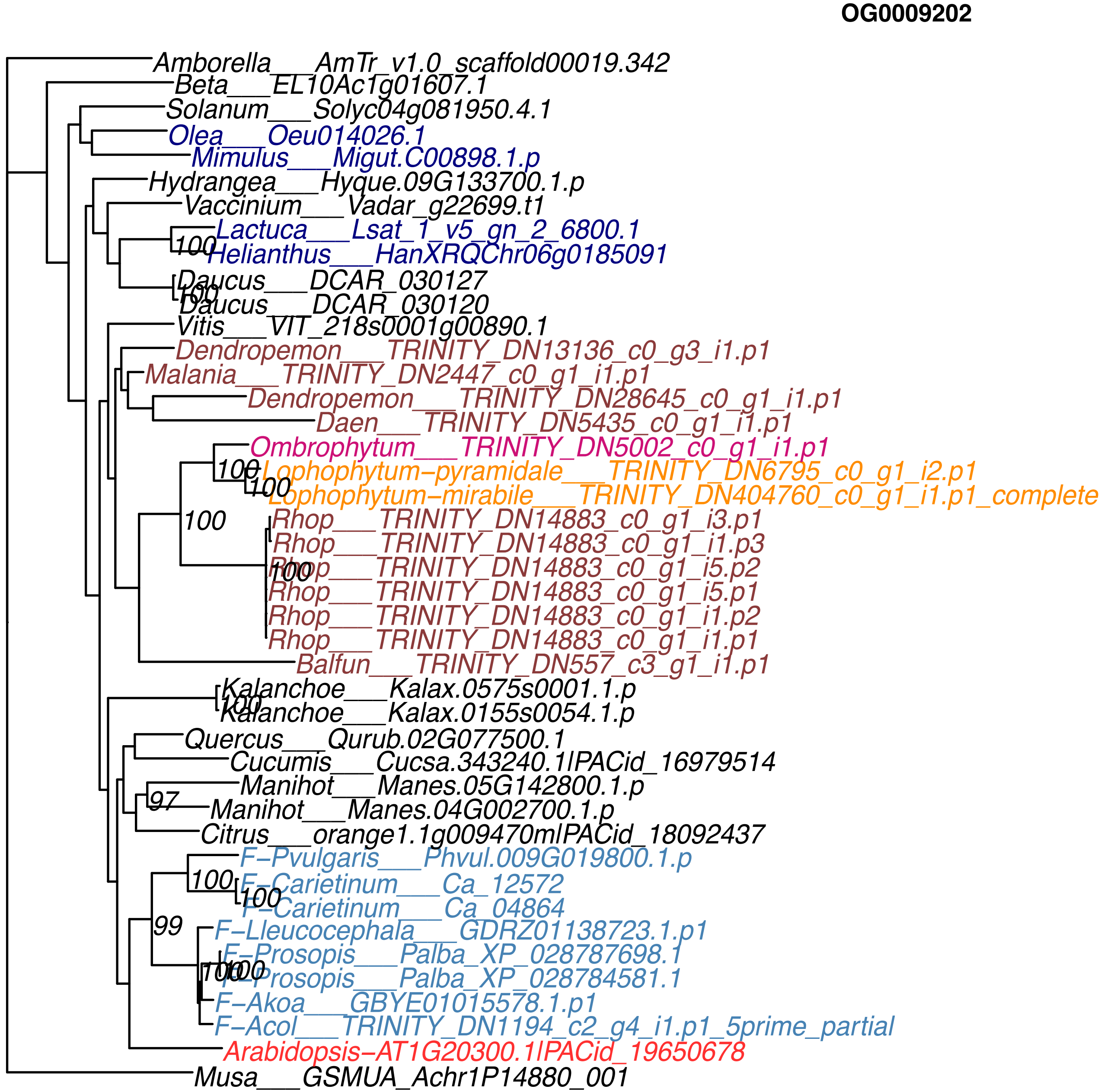

OG0009202
Amborella___AmTr_v1.0_scaffold00019.342
Beta___EL10Ac1g01607.1
Solanum___Solyc04g081950.4.1
Olea___Oeu014026.1
Mimulus___Migut.C00898.1.p
Hydrangea___Hyque.09G133700.1.p
Vaccinium___Vadar_g22699.t1
Lactuca___Lsat_1_v5_gn_2_6800.1
Helianthus___HanXRQChr06g0185091
Daucus___DCAR_030127
Daucus___DCAR_030120
Vitis___VIT_218s0001g00890.1
Dendropemon___TRINITY_DN13136_c0_g3_i1.p1
Malania___TRINITY_DN2447_c0_g1_i1.p1
Dendropemon___TRINITY_DN28645_c0_g1_i1.p1
Daen___TRINITY_DN5435_c0_g1_i1.p1
Ombrophytum___TRINITY_DN5002_c0_g1_i1.p1
Lophophytum−pyramidale___TRINITY_DN6795_c0_g1_i2.p1
Lophophytum−mirabile___TRINITY_DN404760_c0_g1_i1.p1_complete
Rhop___TRINITY_DN14883_c0_g1_i3.p1
Rhop___TRINITY_DN14883_c0_g1_i1.p3
Rhop___TRINITY_DN14883_c0_g1_i5.p2
Rhop___TRINITY_DN14883_c0_g1_i5.p1
Rhop___TRINITY_DN14883_c0_g1_i1.p2
Rhop___TRINITY_DN14883_c0_g1_i1.p1
Balfun___TRINITY_DN557_c3_g1_i1.p1
Kalanchoe___Kalax.0575s0001.1.p
Kalanchoe___Kalax.0155s0054.1.p
Quercus___Qurub.02G077500.1
Cucumis___Cucsa.343240.1lPACid_16979514
Manihot___Manes.05G142800.1.p
Manihot___Manes.04G002700.1.p
Citrus___orange1.1g009470mlPACid_18092437
F−Pvulgaris___Phvul.009G019800.1.p
F−Carietinum___Ca_12572
F−Carietinum___Ca_04864
F−Lleucocephala___GDRZ01138723.1.p1
F−Prosopis___Palba_XP_028787698.1
F−Prosopis___Palba_XP_028784581.1
F−Akoa___GBYE01015578.1.p1
F−Acol___TRINITY_DN1194_c2_g4_i1.p1_5prime_partial
Arabidopsis−AT1G20300.1lPACid_19650678
Musa___GSMUA_Achr1P14880_001
100
100
100
100
100
100
100
97
100
100
99
100
100

**OG0009208**

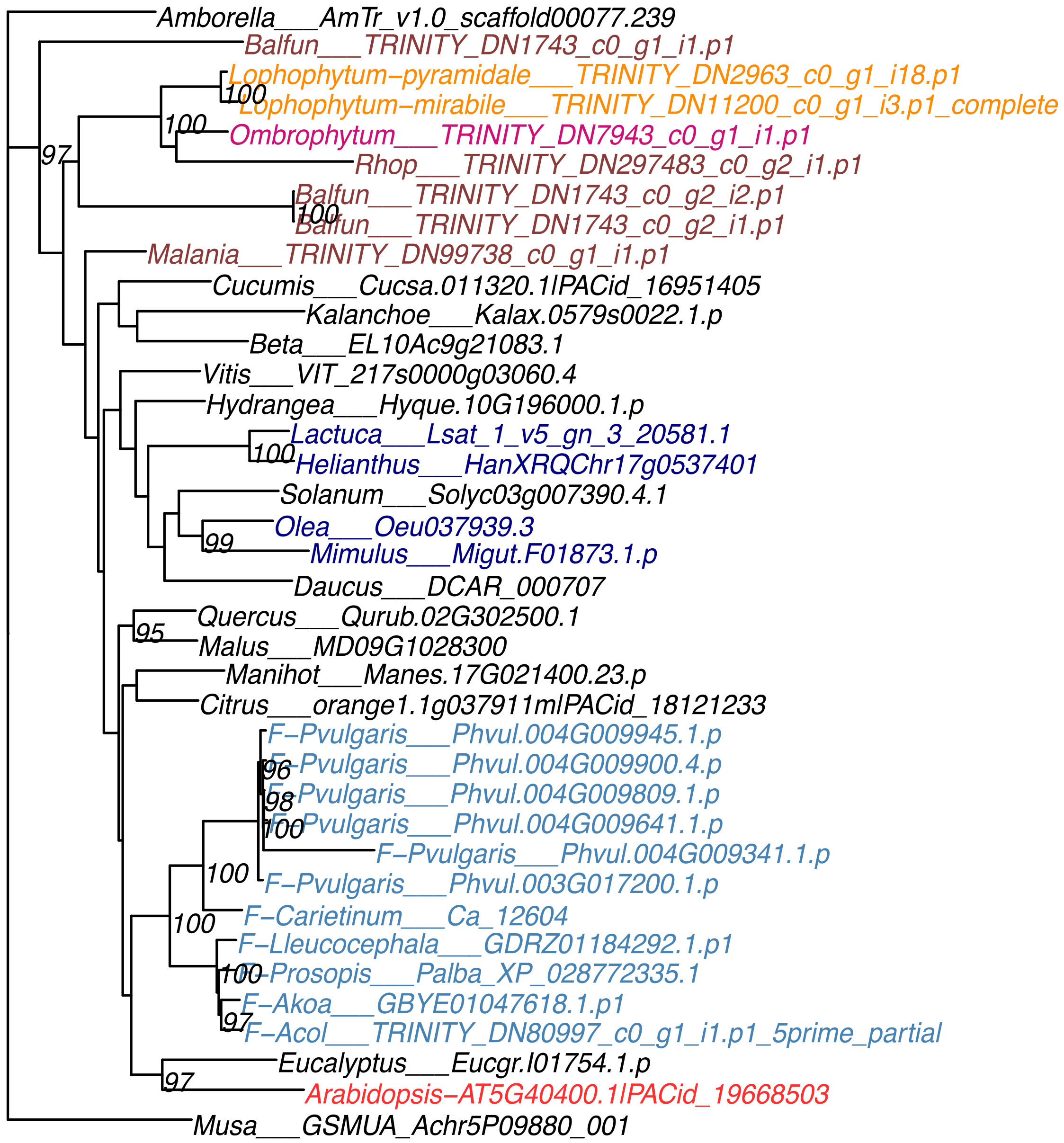
Amborella___AmTr_v1.0_scaffold00077.239
Balfun___TRINITY_DN1743_c0_g1_i1.p1
Lophophytum−pyramidale___TRINITY_DN2963_c0_g1_i18.p1
100
Lophophytum−mirabile___TRINITY_DN11200_c0_g1_i3.p1_complete
100
Ombrophytum___TRINITY_DN7943_c0_g1_i1.p1
97
Rhop___TRINITY_DN297483_c0_g2_i1.p1
Balfun___TRINITY_DN1743_c0_g2_i2.p1
100
Balfun___TRINITY_DN1743_c0_g2_i1.p1
Malania___TRINITY_DN99738_c0_g1_i1.p1
Cucumis___Cucsa.011320.1|PACid_16951405
Kalanchoe___Kalax.0579s0022.1.p
Beta___EL10Ac9g21083.1
Vitis___VIT_217s0000g03060.4
Hydrangea___Hyque.10G196000.1.p
Lactuca___Lsat_1_v5_gn_3_20581.1
100
Helianthus___HanXRQChr17g0537401
Solanum___Solyc03g007390.4.1
Olea___Oeu037939.3
99
Mimulus___Migut.F01873.1.p
Daucus___DCAR_000707
Quercus___Qurub.02G302500.1
95
Malus___MD09G1028300
Manihot___Manes.17G021400.23.p
Citrus___orange1.1g037911m|PACid_18121233
F−Pvulgaris___Phvul.004G009945.1.p
F−Pvulgaris___Phvul.004G009900.4.p
96
F−Pvulgaris___Phvul.004G009809.1.p
98
F−Pvulgaris___Phvul.004G009641.1.p
100
F−Pvulgaris___Phvul.004G009341.1.p
100
F−Pvulgaris___Phvul.003G017200.1.p
F−Carietinum___Ca_12604
100
F−Lleucocephala___GDRZ01184292.1.p1
100
F−Prosopis___Palba_XP_028772335.1
F−Akoa___GBYE01047618.1.p1
97
F−Acol___TRINITY_DN80997_c0_g1_i1.p1_5prime_partial
Eucalyptus___Eucgr.I01754.1.p
97
Arabidopsis−AT5G40400.1|PACid_19668503
Musa___GSMUA_Achr5P09880_001

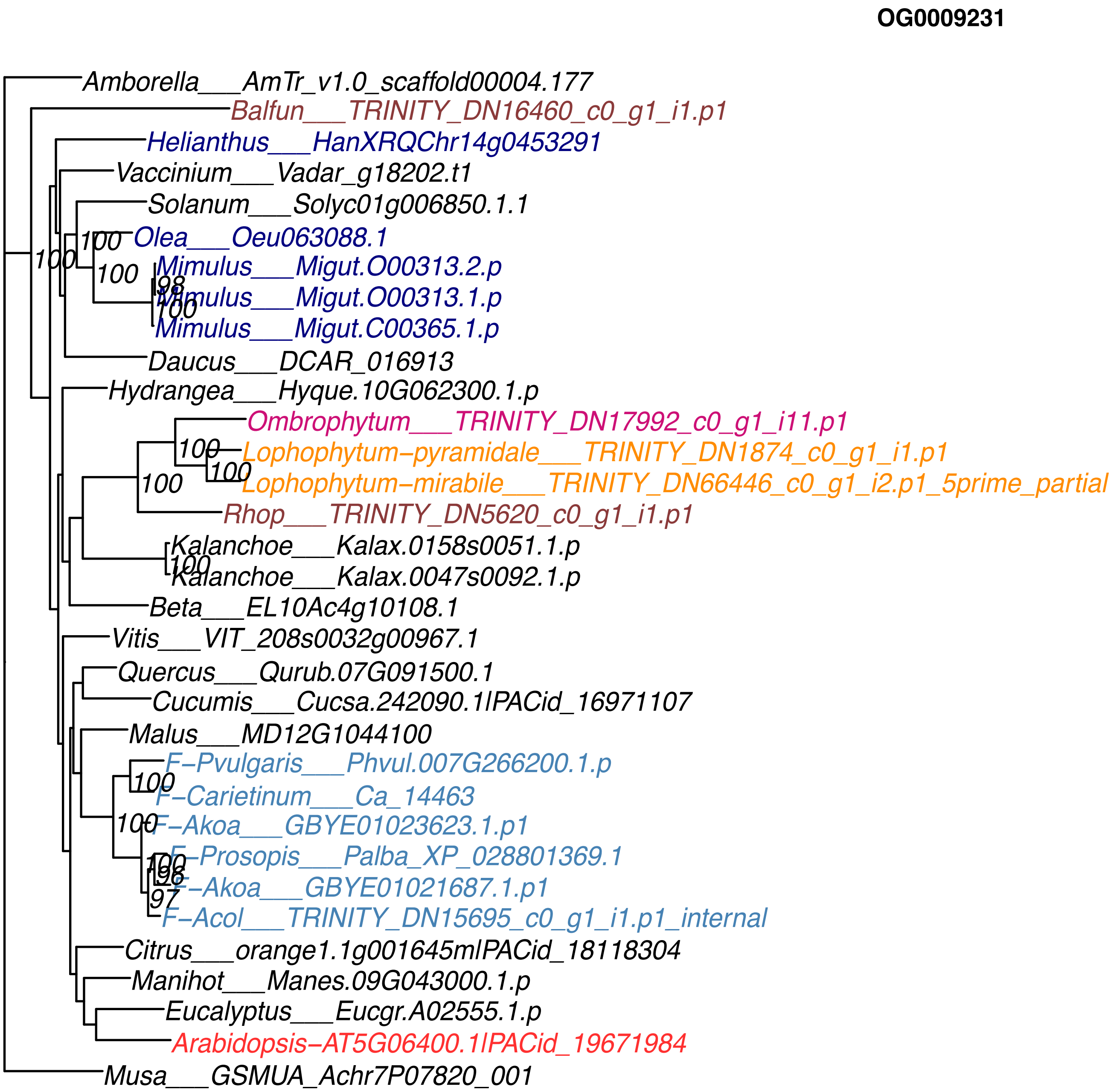
OG0009231
Amborella___AmTr_v1.0_scaffold00004.177
Balfun___TRINITY_DN16460_c0_g1_i1.p1
Helianthus___HanXRQChr14g0453291
Vaccinium___Vadar_g18202.t1
Solanum___Solyc01g006850.1.1
Olea___Oeu063088.1
Mimulus___Migut.O00313.2.p
Mimulus___Migut.O00313.1.p
Mimulus___Migut.C00365.1.p
Daucus___DCAR_016913
Hydrangea___Hyque.10G062300.1.p
Ombrophytum___TRINITY_DN17992_c0_g1_i11.p1
Lophophytum−pyramidale___TRINITY_DN1874_c0_g1_i1.p1
Lophophytum−mirabile___TRINITY_DN66446_c0_g1_i2.p1_5prime_partial
Rhop___TRINITY_DN5620_c0_g1_i1.p1
Kalanchoe___Kalax.0158s0051.1.p
Kalanchoe___Kalax.0047s0092.1.p
Beta___EL10Ac4g10108.1
Vitis___VIT_208s0032g00967.1
Quercus___Qurub.07G091500.1
Cucumis___Cucsa.242090.1lPACid_16971107
Malus___MD12G1044100
F−Pvulgaris___Phvul.007G266200.1.p
F−Carietinum___Ca_14463
F−Akoa___GBYE01023623.1.p1
F−Prosopis___Palba_XP_028801369.1
F−Akoa___GBYE01021687.1.p1
F−Acol___TRINITY_DN15695_c0_g1_i1.p1_internal
Citrus___orange1.1g001645mlPACid_18118304
Manihot___Manes.09G043000.1.p
Eucalyptus___Eucgr.A02555.1.p
Arabidopsis−AT5G06400.1lPACid_19671984
Musa___GSMUA_Achr7P07820_001

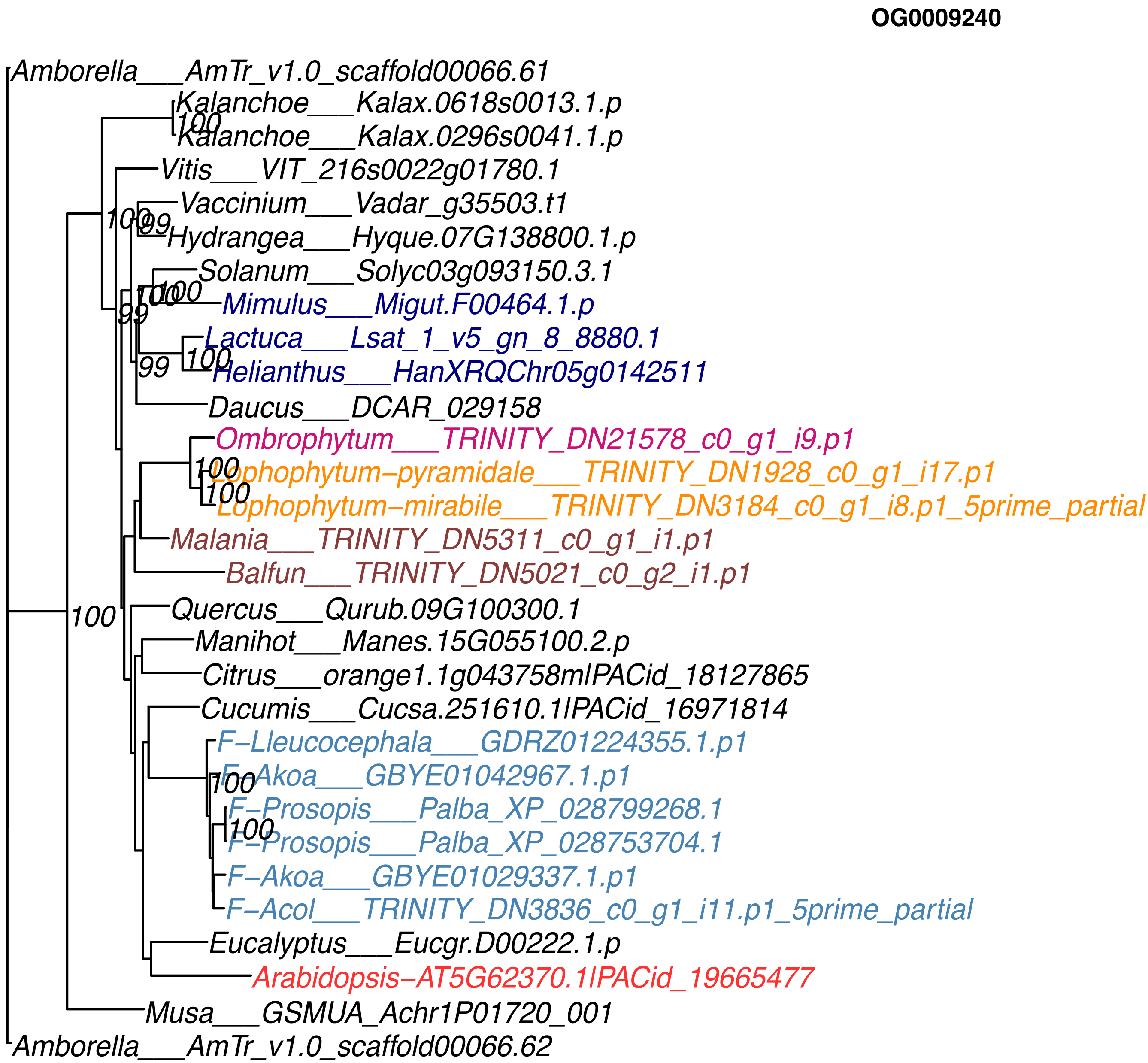

OG0009240
Amborella___AmTr_v1.0_scaffold00066.61
Kalanchoe___Kalax.0618s0013.1.p
100
Kalanchoe___Kalax.0296s0041.1.p
Vitis___VIT_216s0022g01780.1
Vaccinium___Vadar_g35503.t1
100
99
Hydrangea___Hyque.07G138800.1.p
Solanum___Solyc03g093150.3.1
100
100
Mimulus___Migut.F00464.1.p
99
Lactuca___Lsat_1_v5_gn_8_8880.1
100
99
Helianthus___HanXRQChr05g0142511
Daucus___DCAR_029158
Ombrophytum___TRINITY_DN21578_c0_g1_i9.p1
100
Lophophytum-pyramidale___TRINITY_DN1928_c0_g1_i17.p1
100
Lophophytum-mirabile___TRINITY_DN3184_c0_g1_i8.p1_5prime_partial
Malania___TRINITY_DN5311_c0_g1_i1.p1
Balfun___TRINITY_DN5021_c0_g2_i1.p1
Quercus___Qurub.09G100300.1
100
Manihot___Manes.15G055100.2.p
Citrus___orange1.1g043758m|PACid_18127865
Cucumis___Cucsa.251610.1|PACid_16971814
F-Lleucocephala___GDRZ01224355.1.p1
F-Akoa___GBYE01042967.1.p1
100
F-Prosopis___Palba_XP_028799268.1
100
F-Prosopis___Palba_XP_028753704.1
F-Akoa___GBYE01029337.1.p1
F-Acol___TRINITY_DN3836_c0_g1_i11.p1_5prime_partial
Eucalyptus___Eucgr.D00222.1.p
Arabidopsis-AT5G62370.1|PACid_19665477
Musa___GSMUA_Achr1P01720_001
Amborella___AmTr_v1.0_scaffold00066.62

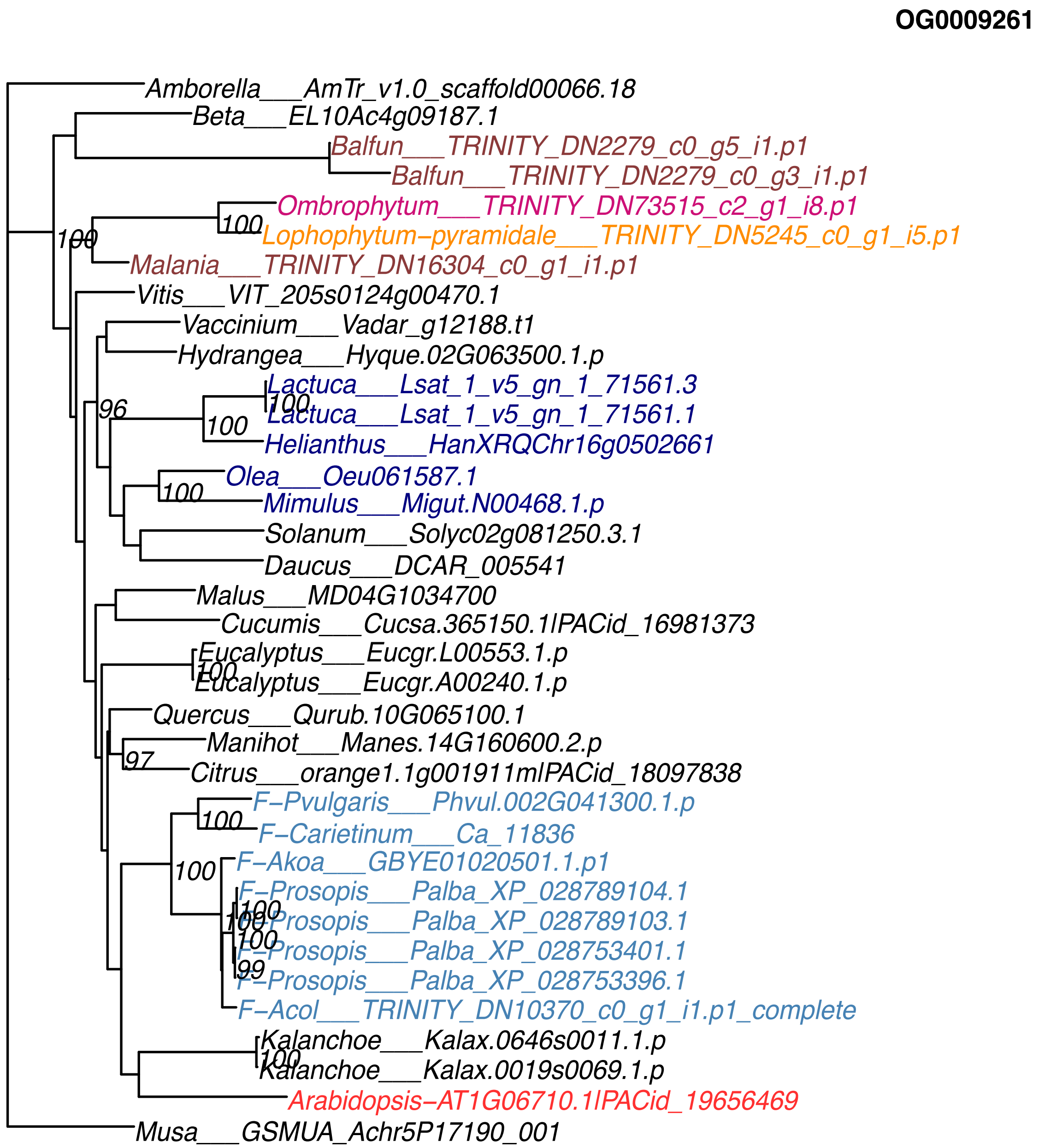

OG0009261
Amborella___AmTr_v1.0_scaffold00066.18
Beta___EL10Ac4g09187.1
Balfun___TRINITY_DN2279_c0_g5_i1.p1
Balfun___TRINITY_DN2279_c0_g3_i1.p1
Ombrophytum___TRINITY_DN73515_c2_g1_i8.p1
Lophophytum−pyramidale___TRINITY_DN5245_c0_g1_i5.p1
Malania___TRINITY_DN16304_c0_g1_i1.p1
Vitis___VIT_205s0124g00470.1
Vaccinium___Vadar_g12188.t1
Hydrangea___Hyque.02G063500.1.p
Lactuca___Lsat_1_v5_gn_1_71561.3
Lactuca___Lsat_1_v5_gn_1_71561.1
Helianthus___HanXRQChr16g0502661
Olea___Oeu061587.1
Mimulus___Migut.N00468.1.p
Solanum___Solyc02g081250.3.1
Daucus___DCAR_005541
Malus___MD04G1034700
Cucumis___Cucsa.365150.1|PACid_16981373
Eucalyptus___Eucgr.L00553.1.p
Eucalyptus___Eucgr.A00240.1.p
Quercus___Qurub.10G065100.1
Manihot___Manes.14G160600.2.p
Citrus___orange1.1g001911m|PACid_18097838
F−Pvulgaris___Phvul.002G041300.1.p
F−Carietinum___Ca_11836
F−Akoa___GBYE01020501.1.p1
F−Prosopis___Palba_XP_028789104.1
F−Prosopis___Palba_XP_028789103.1
F−Prosopis___Palba_XP_028753401.1
F−Prosopis___Palba_XP_028753396.1
F−Acol___TRINITY_DN10370_c0_g1_i1.p1_complete
Kalanchoe___Kalax.0646s0011.1.p
Kalanchoe___Kalax.0019s0069.1.p
Arabidopsis−AT1G06710.1|PACid_19656469
Musa___GSMUA_Achr5P17190_001
100
100
96
100
100
100
100
97
100
100
100
100
99
100

**OG0009267**

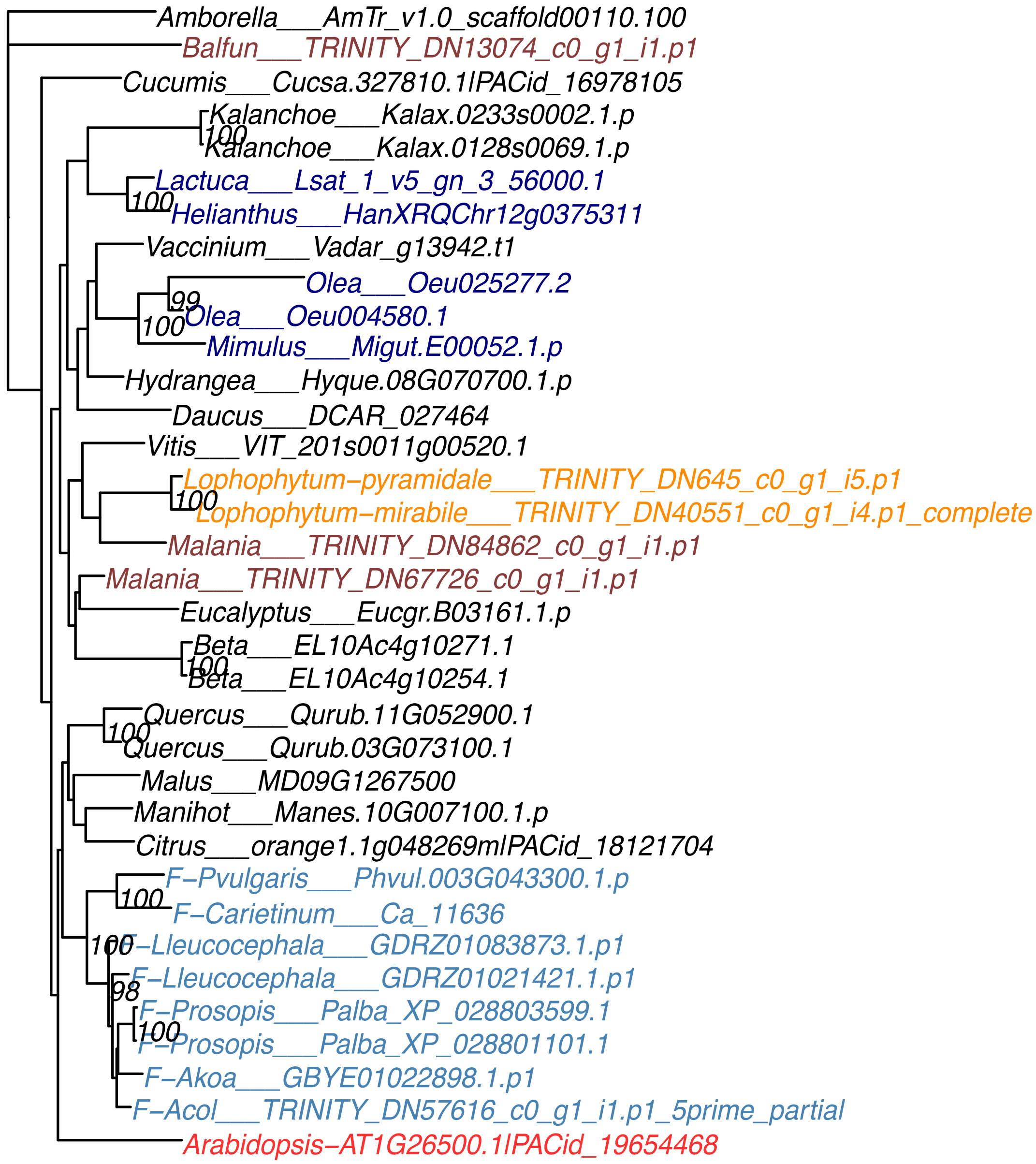

Amborella___AmTr_v1.0_scaffold00110.100
Balfun___TRINITY_DN13074_c0_g1_i1.p1
Cucumis___Cucsa.327810.1|PACid_16978105
Kalanchoe___Kalax.0233s0002.1.p
100
Kalanchoe___Kalax.0128s0069.1.p
Lactuca___Lsat_1_v5_gn_3_56000.1
100
Helianthus___HanXRQChr12g0375311
Vaccinium___Vadar_g13942.t1
Olea___Oeu025277.2
99
Olea___Oeu004580.1
100
Mimulus___Migut.E00052.1.p
Hydrangea___Hyque.08G070700.1.p
Daucus___DCAR_027464
Vitis___VIT_201s0011g00520.1
Lophophytum−pyramidale___TRINITY_DN645_c0_g1_i5.p1
100
Lophophytum−mirabile___TRINITY_DN40551_c0_g1_i4.p1_complete
Malania___TRINITY_DN84862_c0_g1_i1.p1
Malania___TRINITY_DN67726_c0_g1_i1.p1
Eucalyptus___Eucgr.B03161.1.p
Beta___EL10Ac4g10271.1
100
Beta___EL10Ac4g10254.1
Quercus___Qurub.11G052900.1
100
Quercus___Qurub.03G073100.1
Malus___MD09G1267500
Manihot___Manes.10G007100.1.p
Citrus___orange1.1g048269m|PACid_18121704
F−Pvulgaris___Phvul.003G043300.1.p
100
F−Carietinum___Ca_11636
100
F−Lleucocephala___GDRZ01083873.1.p1
F−Lleucocephala___GDRZ01021421.1.p1
98
F−Prosopis___Palba_XP_028803599.1
100
F−Prosopis___Palba_XP_028801101.1
F−Akoa___GBYE01022898.1.p1
F−Acol___TRINITY_DN57616_c0_g1_i1.p1_5prime_partial
Arabidopsis−AT1G26500.1|PACid_19654468

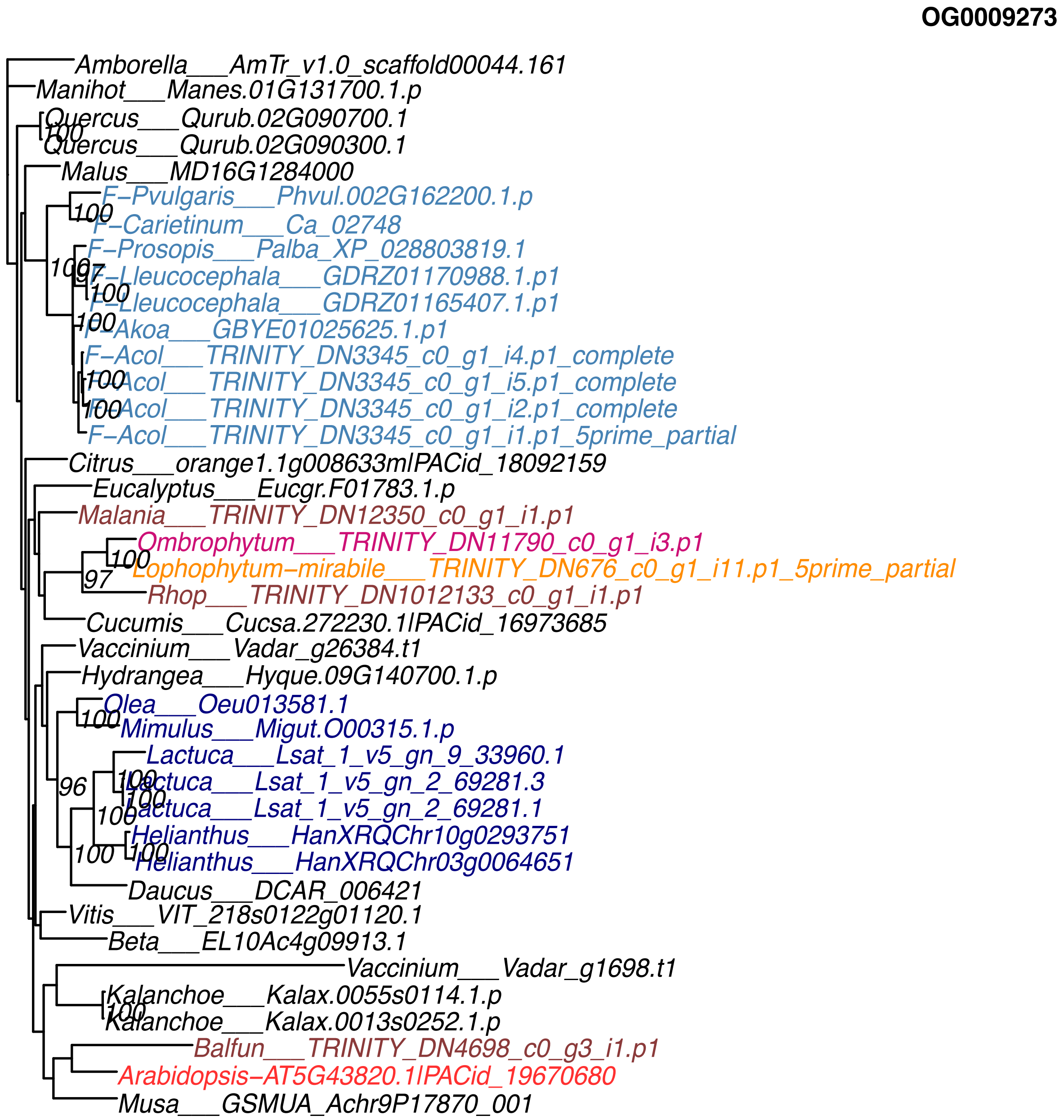
OG0009273
Amborella___AmTr_v1.0_scaffold00044.161
Manihot___Manes.01G131700.1.p
Quercus___Qurub.02G090700.1
100
Quercus___Qurub.02G090300.1
Malus___MD16G1284000
F−Pvulgaris___Phvul.002G162200.1.p
100
F−Carietinum___Ca_02748
F−Prosopis___Palba_XP_028803819.1
100
97
F−Lleucocephala___GDRZ01170988.1.p1
100
F−Lleucocephala___GDRZ01165407.1.p1
100
F−Akoa___GBYE01025625.1.p1
F−Acol___TRINITY_DN3345_c0_g1_i4.p1_complete
100
F−Acol___TRINITY_DN3345_c0_g1_i5.p1_complete
100
F−Acol___TRINITY_DN3345_c0_g1_i2.p1_complete
F−Acol___TRINITY_DN3345_c0_g1_i1.p1_5prime_partial
Citrus___orange1.1g008633m|PACid_18092159
Eucalyptus___Eucgr.F01783.1.p
Malania___TRINITY_DN12350_c0_g1_i1.p1
Ombrophytum___TRINITY_DN11790_c0_g1_i3.p1
100
97
Lophophytum−mirabile___TRINITY_DN676_c0_g1_i11.p1_5prime_partial
Rhop___TRINITY_DN1012133_c0_g1_i1.p1
Cucumis___Cucsa.272230.1|PACid_16973685
Vaccinium___Vadar_g26384.t1
Hydrangea___Hyque.09G140700.1.p
Olea___Oeu013581.1
100
Mimulus___Migut.O00315.1.p
Lactuca___Lsat_1_v5_gn_9_33960.1
100
96
Lactuca___Lsat_1_v5_gn_2_69281.3
100
100
Lactuca___Lsat_1_v5_gn_2_69281.1
Helianthus___HanXRQChr10g0293751
100
100
Helianthus___HanXRQChr03g0064651
Daucus___DCAR_006421
Vitis___VIT_218s0122g01120.1
Beta___EL10Ac4g09913.1
Vaccinium___Vadar_g1698.t1
Kalanchoe___Kalax.0055s0114.1.p
100
Kalanchoe___Kalax.0013s0252.1.p
Balfun___TRINITY_DN4698_c0_g3_i1.p1
Arabidopsis−AT5G43820.1|PACid_19670680
Musa___GSMUA_Achr9P17870_001

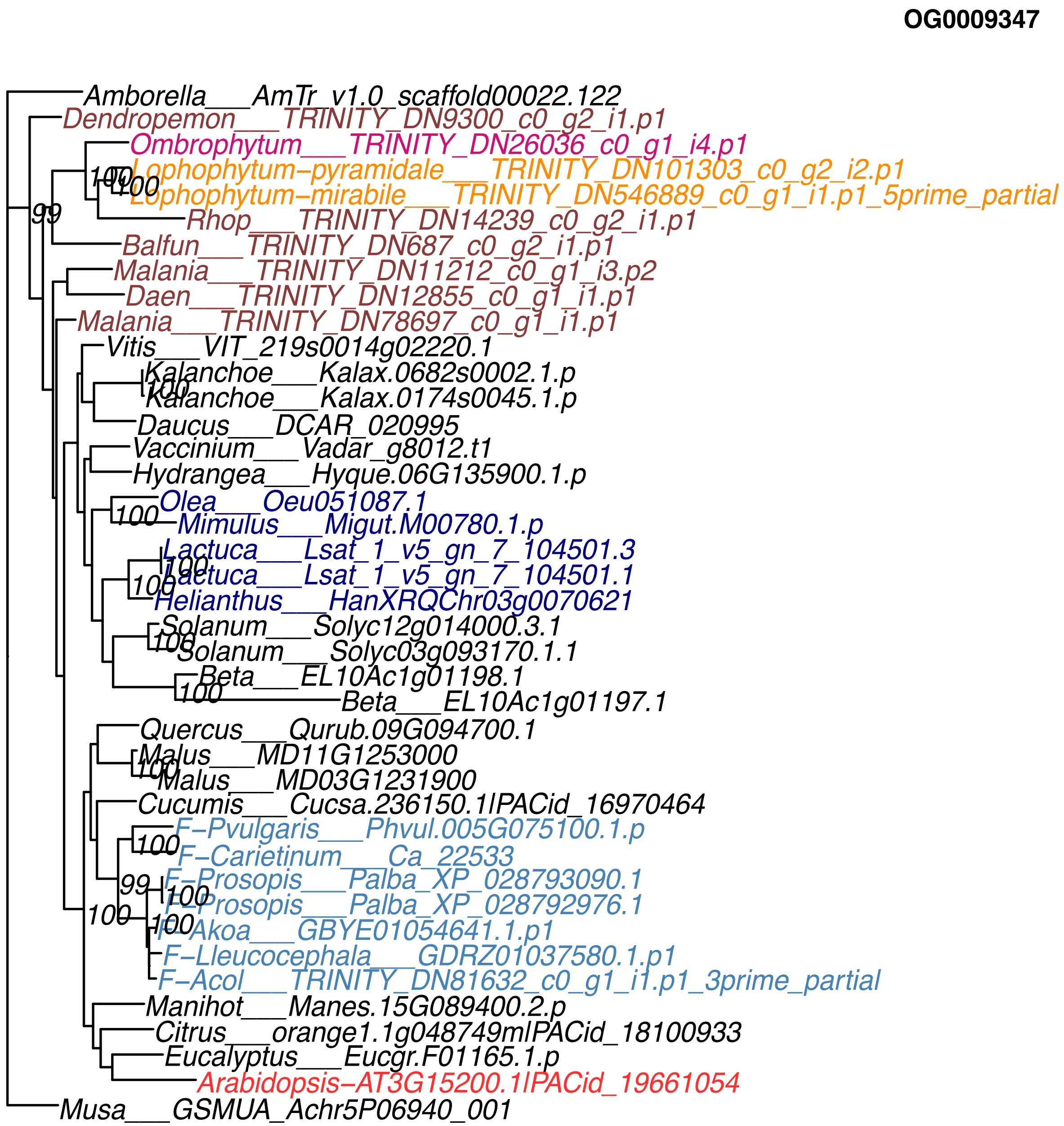
OG0009347
Amborella___AmTr_v1.0_scaffold00022.122
Dendropemon___TRINITY_DN9300_c0_g2_i1.p1
Ombrophytum___TRINITY_DN26036_c0_g1_i4.p1
Lophophytum-pyramidale___TRINITY_DN101303_c0_g2_i2.p1
Lophophytum-mirabile___TRINITY_DN546889_c0_g1_i1.p1_5prime_partial
Rhop___TRINITY_DN14239_c0_g2_i1.p1
Balfun___TRINITY_DN687_c0_g2_i1.p1
Malania___TRINITY_DN11212_c0_g1_i3.p2
Daen___TRINITY_DN12855_c0_g1_i1.p1
Malania___TRINITY_DN78697_c0_g1_i1.p1
Vitis___VIT_219s0014g02220.1
Kalanchoe___Kalax.0682s0002.1.p
Kalanchoe___Kalax.0174s0045.1.p
Daucus___DCAR_020995
Vaccinium___Vadar_g8012.t1
Hydrangea___Hyque.06G135900.1.p
Olea___Oeu051087.1
Mimulus___Migut.M00780.1.p
Lactuca___Lsat_1_v5_gn_7_104501.3
Lactuca___Lsat_1_v5_gn_7_104501.1
Helianthus___HanXRQChr03g0070621
Solanum___Solyc12g014000.3.1
Solanum___Solyc03g093170.1.1
Beta___EL10Ac1g01198.1
Beta___EL10Ac1g01197.1
Quercus___Qurub.09G094700.1
Malus___MD11G1253000
Malus___MD03G1231900
Cucumis___Cucsa.236150.1|PACid_16970464
F-Pvulgaris___Phvul.005G075100.1.p
F-Carietinum___Ca_22533
F-Prosopis___Palba_XP_028793090.1
F-Prosopis___Palba_XP_028792976.1
F-Akoa___GBYE01054641.1.p1
F-Lleucocephala___GDRZ01037580.1.p1
F-Acol___TRINITY_DN81632_c0_g1_i1.p1_3prime_partial
Manihot___Manes.15G089400.2.p
Citrus___orange1.1g048749m|PACid_18100933
Eucalyptus___Eucgr.F01165.1.p
Arabidopsis-AT3G15200.1|PACid_19661054
Musa___GSMUA_Achr5P06940_001
100
100
99
100
100
100
100
100
100
100
100
99
100
100
100

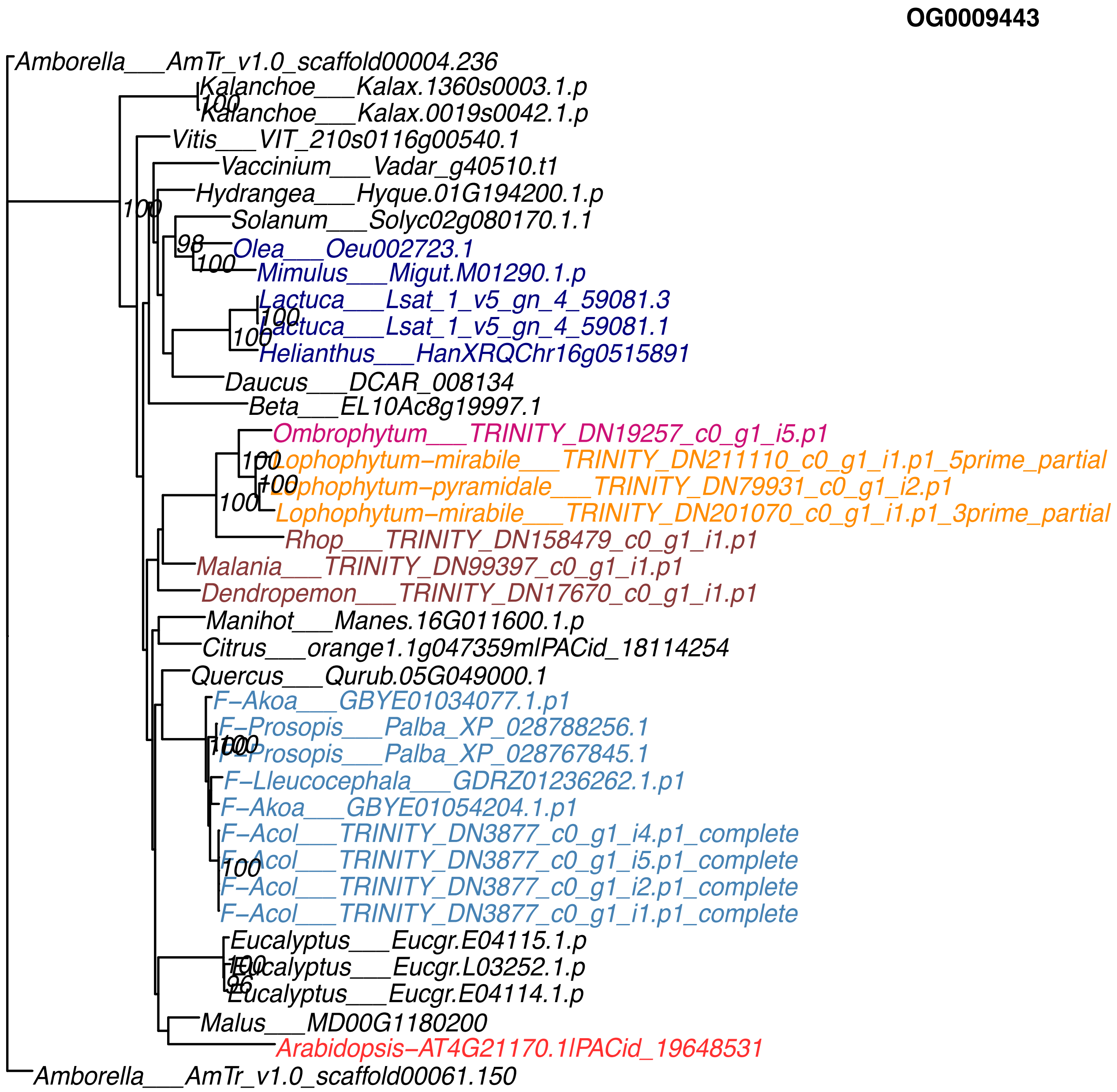
OG0009443
Amborella___AmTr_v1.0_scaffold00004.236
Kalanchoe___Kalax.1360s0003.1.p
100
Kalanchoe___Kalax.0019s0042.1.p
Vitis___VIT_210s0116g00540.1
Vaccinium___Vadar_g40510.t1
Hydrangea___Hyque.01G194200.1.p
100
Solanum___Solyc02g080170.1.1
98
Olea___Oeu002723.1
100
Mimulus___Migut.M01290.1.p
Lactuca___Lsat_1_v5_gn_4_59081.3
100
Lactuca___Lsat_1_v5_gn_4_59081.1
100
Helianthus___HanXRQChr16g0515891
Daucus___DCAR_008134
Beta___EL10Ac8g19997.1
Ombrophytum___TRINITY_DN19257_c0_g1_i5.p1
100
Lophophytum−mirabile___TRINITY_DN211110_c0_g1_i1.p1_5prime_partial
100
Lophophytum−pyramidale___TRINITY_DN79931_c0_g1_i2.p1
100
Lophophytum−mirabile___TRINITY_DN201070_c0_g1_i1.p1_3prime_partial
Rhop___TRINITY_DN158479_c0_g1_i1.p1
Malania___TRINITY_DN99397_c0_g1_i1.p1
Dendropemon___TRINITY_DN17670_c0_g1_i1.p1
Manihot___Manes.16G011600.1.p
Citrus___orange1.1g047359m|PACid_18114254
Quercus___Qurub.05G049000.1
F−Akoa___GBYE01034077.1.p1
F−Prosopis___Palba_XP_028788256.1
100
100
F−Prosopis___Palba_XP_028767845.1
F−Lleucocephala___GDRZ01236262.1.p1
F−Akoa___GBYE01054204.1.p1
F−Acol___TRINITY_DN3877_c0_g1_i4.p1_complete
F−Acol___TRINITY_DN3877_c0_g1_i5.p1_complete
100
F−Acol___TRINITY_DN3877_c0_g1_i2.p1_complete
F−Acol___TRINITY_DN3877_c0_g1_i1.p1_complete
Eucalyptus___Eucgr.E04115.1.p
100
Eucalyptus___Eucgr.L03252.1.p
96
Eucalyptus___Eucgr.E04114.1.p
Malus___MD00G1180200
Arabidopsis−AT4G21170.1|PACid_19648531
Amborella___AmTr_v1.0_scaffold00061.150

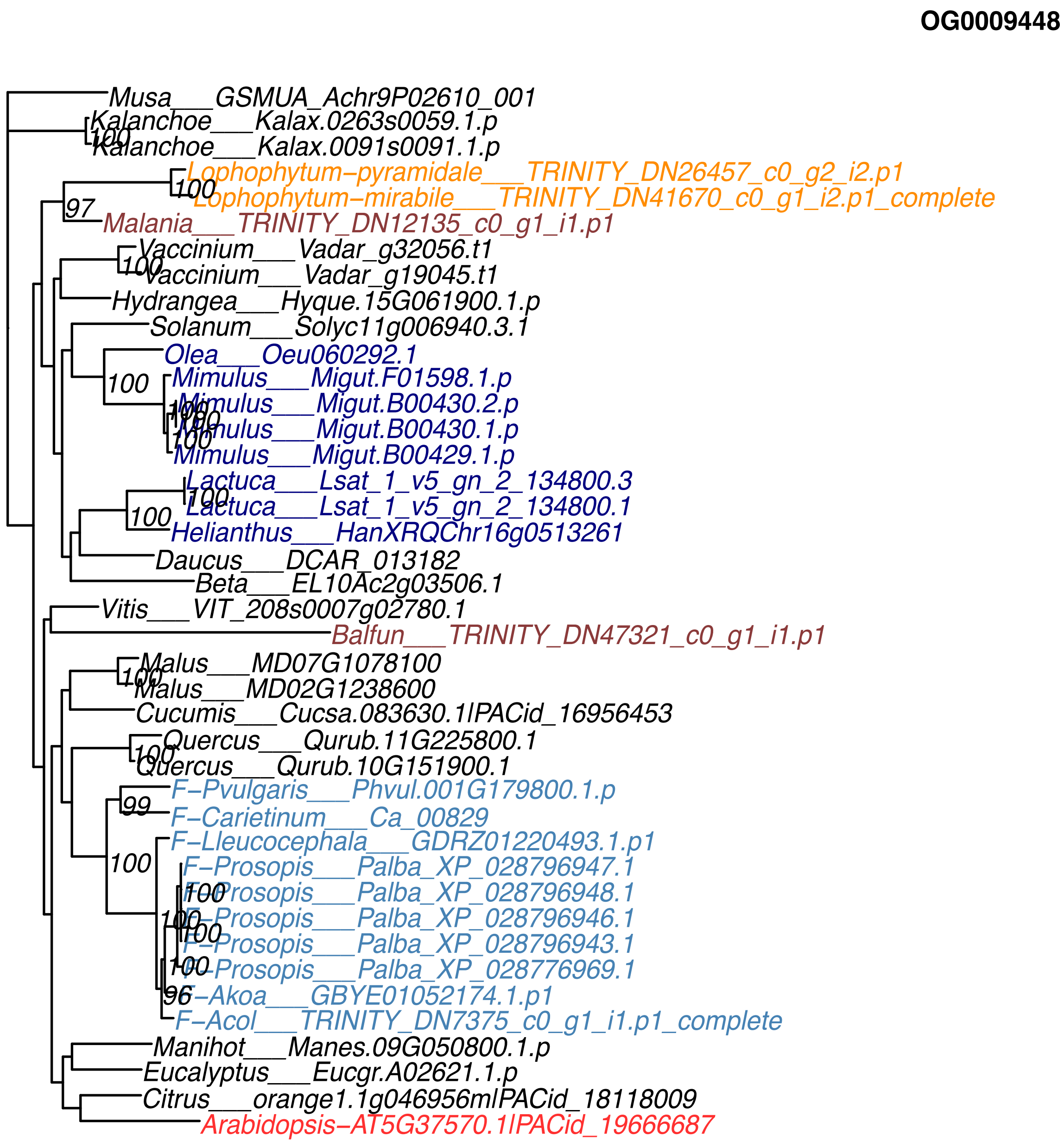

OG0009448
Musa___GSMUA_Achr9P02610_001
Kalanchoe___Kalax.0263s0059.1.p
Kalanchoe___Kalax.0091s0091.1.p
Lophophytum−pyramidale___TRINITY_DN26457_c0_g2_i2.p1
Lophophytum−mirabile___TRINITY_DN41670_c0_g1_i2.p1_complete
Malania___TRINITY_DN12135_c0_g1_i1.p1
Vaccinium___Vadar_g32056.t1
Vaccinium___Vadar_g19045.t1
Hydrangea___Hyque.15G061900.1.p
Solanum___Solyc11g006940.3.1
Olea___Oeu060292.1
Mimulus___Migut.F01598.1.p
Mimulus___Migut.B00430.2.p
Mimulus___Migut.B00430.1.p
Mimulus___Migut.B00429.1.p
Lactuca___Lsat_1_v5_gn_2_134800.3
Lactuca___Lsat_1_v5_gn_2_134800.1
Helianthus___HanXRQChr16g0513261
Daucus___DCAR_013182
Beta___EL10Ac2g03506.1
Vitis___VIT_208s0007g02780.1
Balfun___TRINITY_DN47321_c0_g1_i1.p1
Malus___MD07G1078100
Malus___MD02G1238600
Cucumis___Cucsa.083630.1|PACid_16956453
Quercus___Qurub.11G225800.1
Quercus___Qurub.10G151900.1
F−Pvulgaris___Phvul.001G179800.1.p
F−Carietinum___Ca_00829
F−Lleucocephala___GDRZ01220493.1.p1
F−Prosopis___Palba_XP_028796947.1
F−Prosopis___Palba_XP_028796948.1
F−Prosopis___Palba_XP_028796946.1
F−Prosopis___Palba_XP_028796943.1
F−Prosopis___Palba_XP_028776969.1
F−Akoa___GBYE01052174.1.p1
F−Acol___TRINITY_DN7375_c0_g1_i1.p1_complete
Manihot___Manes.09G050800.1.p
Eucalyptus___Eucgr.A02621.1.p
Citrus___orange1.1g046956m|PACid_18118009
Arabidopsis−AT5G37570.1|PACid_19666687

**OG0009468**

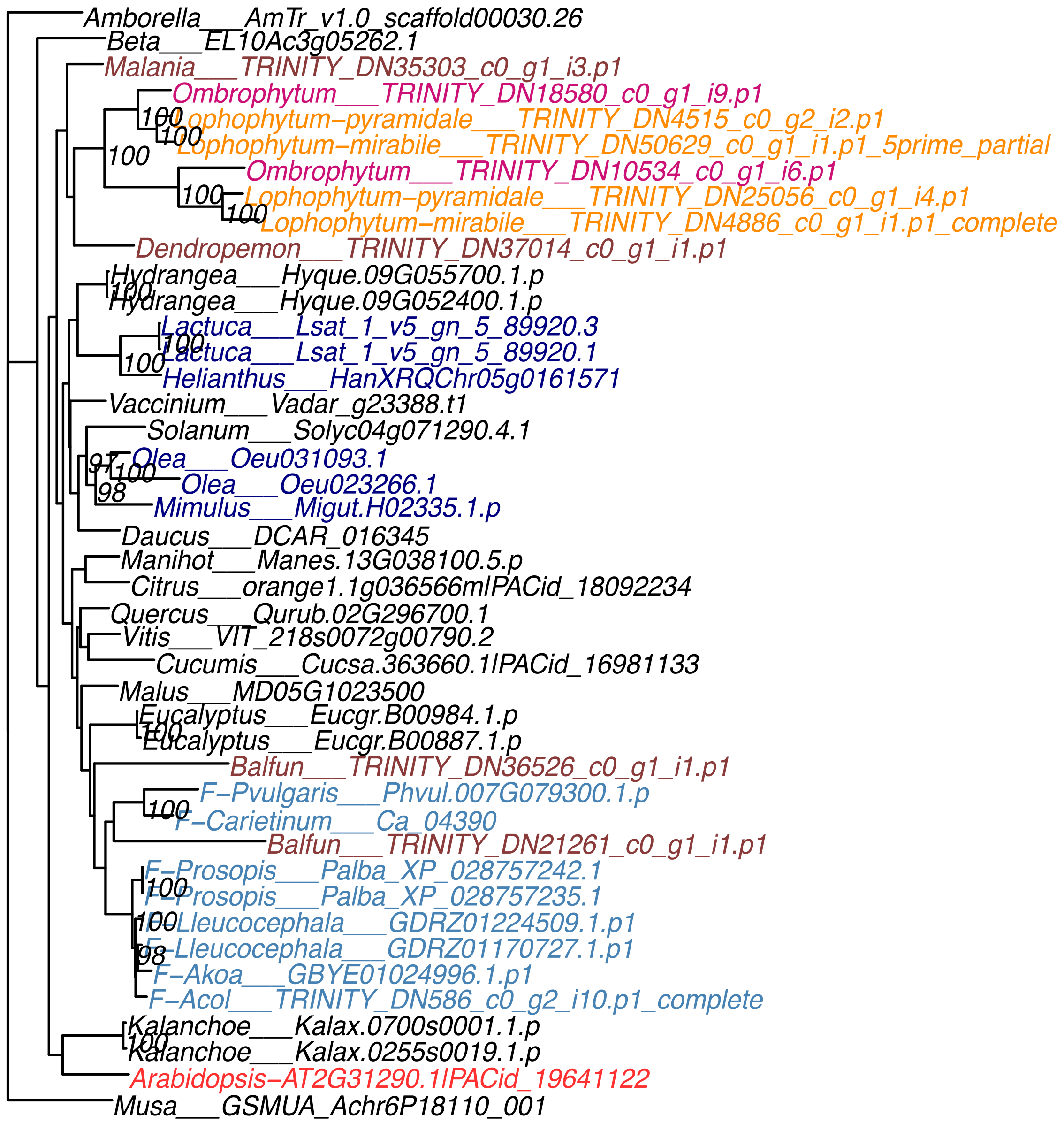
Amborella___AmTr_v1.0_scaffold00030.26
Beta___EL10Ac3g05262.1
Malania___TRINITY_DN35303_c0_g1_i3.p1
Ombrophytum___TRINITY_DN18580_c0_g1_i9.p1
Lophophytum−pyramidale___TRINITY_DN4515_c0_g2_i2.p1
Lophophytum−mirabile___TRINITY_DN50629_c0_g1_i1.p1_5prime_partial
Ombrophytum___TRINITY_DN10534_c0_g1_i6.p1
Lophophytum−pyramidale___TRINITY_DN25056_c0_g1_i4.p1
Lophophytum−mirabile___TRINITY_DN4886_c0_g1_i1.p1_complete
Dendropemon___TRINITY_DN37014_c0_g1_i1.p1
Hydrangea___Hyque.09G055700.1.p
Hydrangea___Hyque.09G052400.1.p
Lactuca___Lsat_1_v5_gn_5_89920.3
Lactuca___Lsat_1_v5_gn_5_89920.1
Helianthus___HanXRQChr05g0161571
Vaccinium___Vadar_g23388.t1
Solanum___Solyc04g071290.4.1
Olea___Oeu031093.1
Olea___Oeu023266.1
Mimulus___Migut.H02335.1.p
Daucus___DCAR_016345
Manihot___Manes.13G038100.5.p
Citrus___orange1.1g036566m|PACid_18092234
Quercus___Qurub.02G296700.1
Vitis___VIT_218s0072g00790.2
Cucumis___Cucsa.363660.1|PACid_16981133
Malus___MD05G1023500
Eucalyptus___Eucgr.B00984.1.p
Eucalyptus___Eucgr.B00887.1.p
Balfun___TRINITY_DN36526_c0_g1_i1.p1
F−Pvulgaris___Phvul.007G079300.1.p
F−Carietinum___Ca_04390
Balfun___TRINITY_DN21261_c0_g1_i1.p1
F−Prosopis___Palba_XP_028757242.1
F−Prosopis___Palba_XP_028757235.1
F−Lleucocephala___GDRZ01224509.1.p1
F−Lleucocephala___GDRZ01170727.1.p1
F−Akoa___GBYE01024996.1.p1
F−Acol___TRINITY_DN586_c0_g2_i10.p1_complete
Kalanchoe___Kalax.0700s0001.1.p
Kalanchoe___Kalax.0255s0019.1.p
Arabidopsis−AT2G31290.1|PACid_19641122
Musa___GSMUA_Achr6P18110_001
100
100
100
100
100
100
100
100
97
100
98
100
100
100
100
98
100

OG0009485

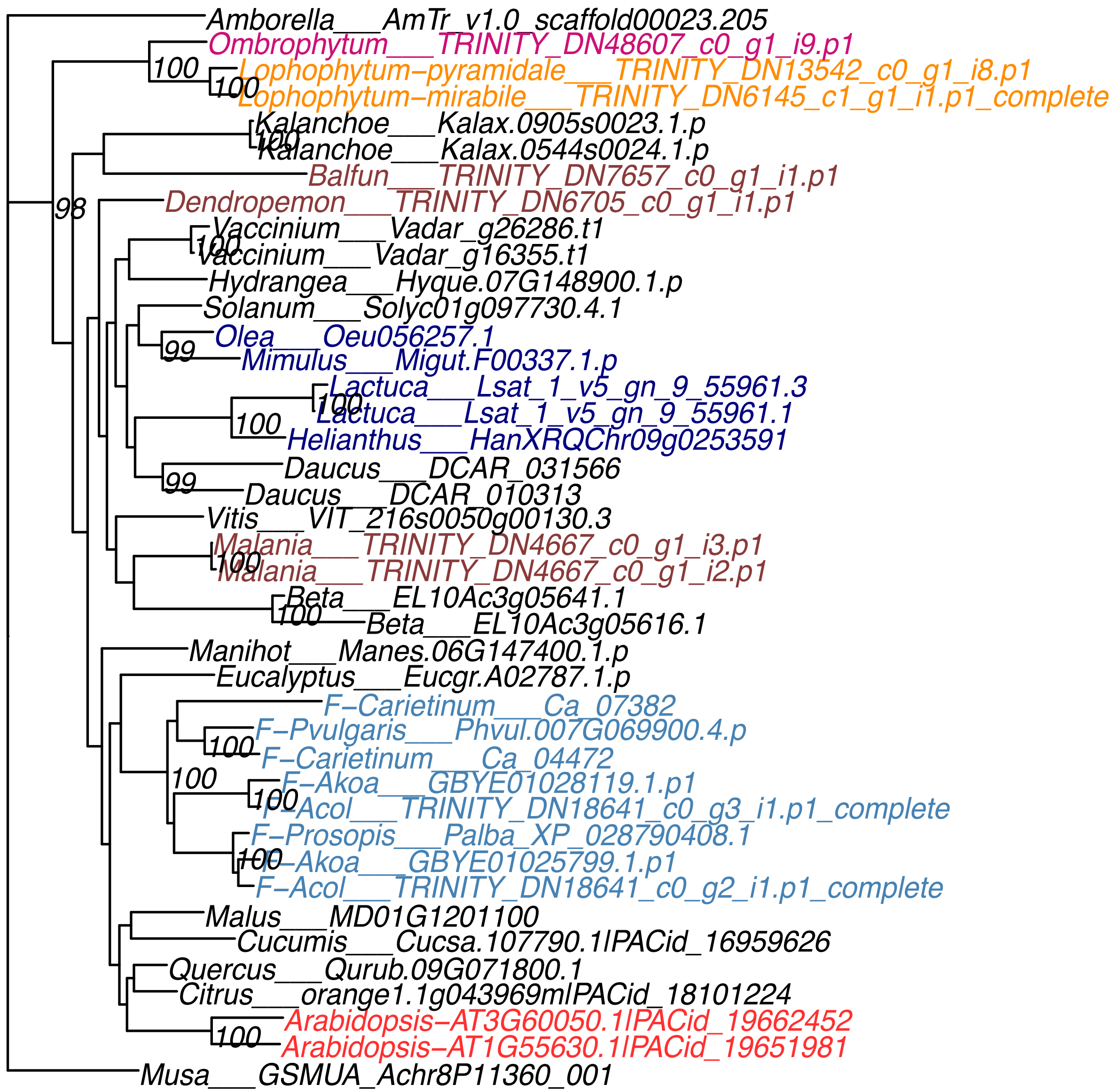

Amborella___AmTr_v1.0_scaffold00023.205
Ombrophytum___TRINITY_DN48607_c0_g1_i9.p1
100
Lophophytum−pyramidale___TRINITY_DN13542_c0_g1_i8.p1
100
Lophophytum−mirabile___TRINITY_DN6145_c1_g1_i1.p1_complete
Kalanchoe___Kalax.0905s0023.1.p
100
Kalanchoe___Kalax.0544s0024.1.p
Balfun___TRINITY_DN7657_c0_g1_i1.p1
98
Dendropemon___TRINITY_DN6705_c0_g1_i1.p1
Vaccinium___Vadar_g26286.t1
100
Vaccinium___Vadar_g16355.t1
Hydrangea___Hyque.07G148900.1.p
Solanum___Solyc01g097730.4.1
Olea___Oeu056257.1
99
Mimulus___Migut.F00337.1.p
Lactuca___Lsat_1_v5_gn_9_55961.3
100
Lactuca___Lsat_1_v5_gn_9_55961.1
100
Helianthus___HanXRQChr09g0253591
Daucus___DCAR_031566
99
Daucus___DCAR_010313
Vitis___VIT_216s0050g00130.3
Malania___TRINITY_DN4667_c0_g1_i3.p1
100
Malania___TRINITY_DN4667_c0_g1_i2.p1
Beta___EL10Ac3g05641.1
100
Beta___EL10Ac3g05616.1
Manihot___Manes.06G147400.1.p
Eucalyptus___Eucgr.A02787.1.p
F−Carietinum___Ca_07382
F−Pvulgaris___Phvul.007G069900.4.p
100
F−Carietinum___Ca_04472
100
F−Akoa___GBYE01028119.1.p1
100
F−Acol___TRINITY_DN18641_c0_g3_i1.p1_complete
F−Prosopis___Palba_XP_028790408.1
100
F−Akoa___GBYE01025799.1.p1
F−Acol___TRINITY_DN18641_c0_g2_i1.p1_complete
Malus___MD01G1201100
Cucumis___Cucsa.107790.1|PACid_16959626
Quercus___Qurub.09G071800.1
Citrus___orange1.1g043969m|PACid_18101224
Arabidopsis−AT3G60050.1|PACid_19662452
100
Arabidopsis−AT1G55630.1|PACid_19651981
Musa___GSMUA_Achr8P11360_001

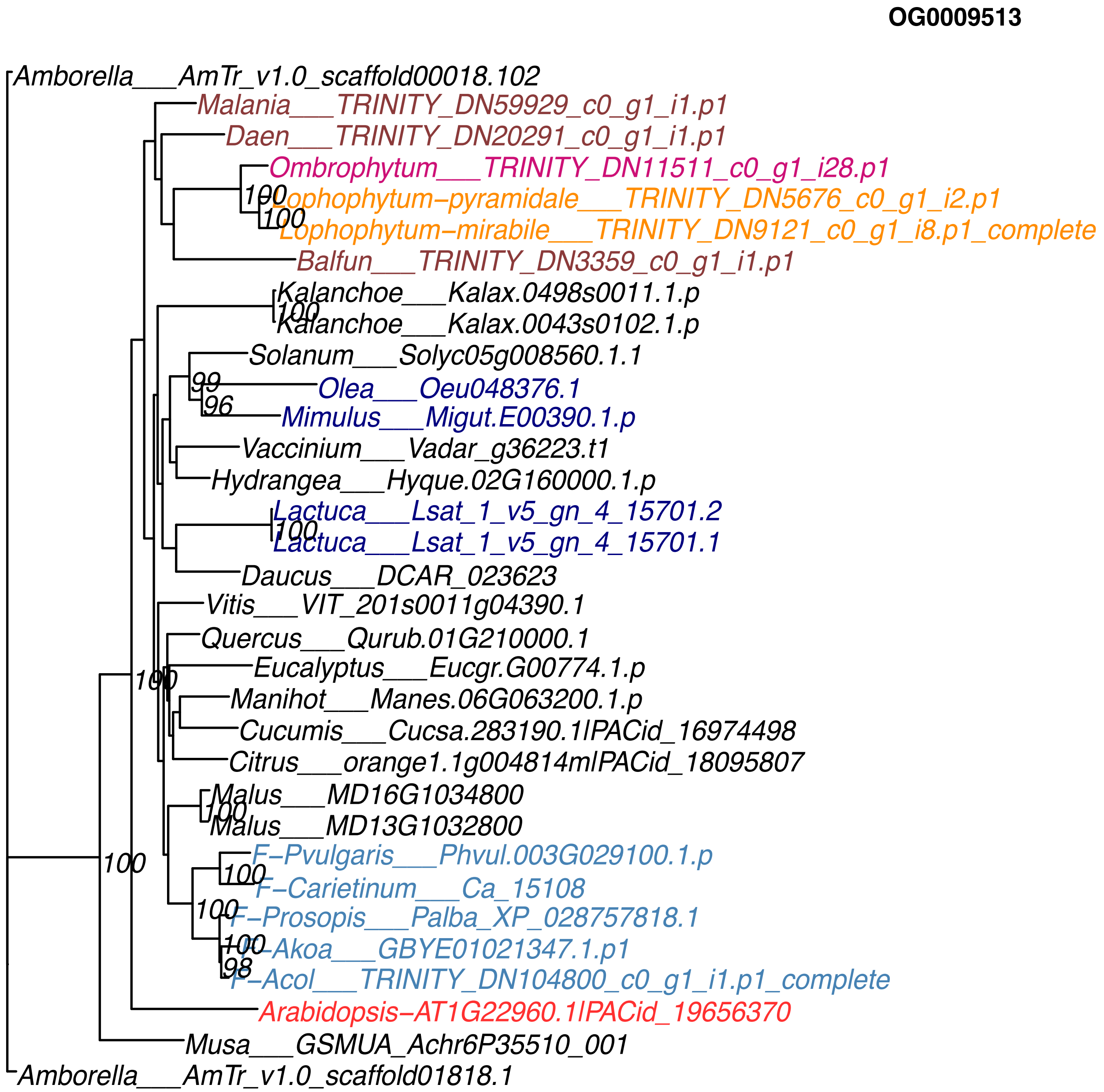

OG0009513
Amborella___AmTr_v1.0_scaffold00018.102
Malania___TRINITY_DN59929_c0_g1_i1.p1
Daen___TRINITY_DN20291_c0_g1_i1.p1
Ombrophytum___TRINITY_DN11511_c0_g1_i28.p1
Lophophytum−pyramidale___TRINITY_DN5676_c0_g1_i2.p1
Lophophytum−mirabile___TRINITY_DN9121_c0_g1_i8.p1_complete
Balfun___TRINITY_DN3359_c0_g1_i1.p1
Kalanchoe___Kalax.0498s0011.1.p
Kalanchoe___Kalax.0043s0102.1.p
Solanum___Solyc05g008560.1.1
Olea___Oeu048376.1
Mimulus___Migut.E00390.1.p
Vaccinium___Vadar_g36223.t1
Hydrangea___Hyque.02G160000.1.p
Lactuca___Lsat_1_v5_gn_4_15701.2
Lactuca___Lsat_1_v5_gn_4_15701.1
Daucus___DCAR_023623
Vitis___VIT_201s0011g04390.1
Quercus___Qurub.01G210000.1
Eucalyptus___Eucgr.G00774.1.p
Manihot___Manes.06G063200.1.p
Cucumis___Cucsa.283190.1|PACid_16974498
Citrus___orange1.1g004814m|PACid_18095807
Malus___MD16G1034800
Malus___MD13G1032800
F−Pvulgaris___Phvul.003G029100.1.p
F−Carietinum___Ca_15108
F−Prosopis___Palba_XP_028757818.1
F−Akoa___GBYE01021347.1.p1
F−Acol___TRINITY_DN104800_c0_g1_i1.p1_complete
Arabidopsis−AT1G22960.1|PACid_19656370
Musa___GSMUA_Achr6P35510_001
Amborella___AmTr_v1.0_scaffold01818.1
100
100
100
99
96
100
100
100
100
100
100
100
98

# OG0009520

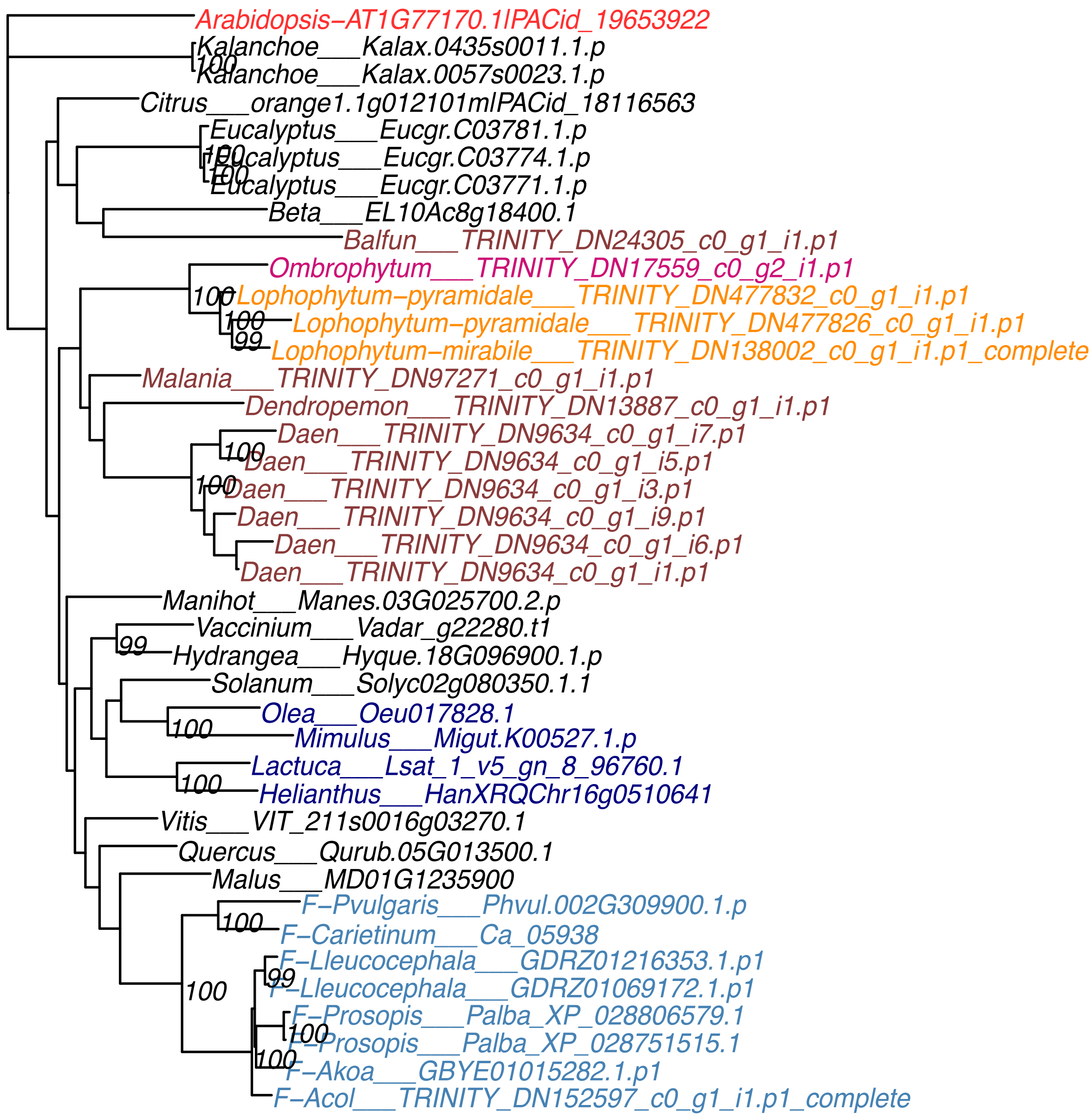

Arabidopsis−AT1G77170.1IPACid_19653922
Kalanchoe___Kalax.0435s0011.1.p
100
Kalanchoe___Kalax.0057s0023.1.p
Citrus___orange1.1g012101mIPACid_18116563
Eucalyptus___Eucgr.C03781.1.p
100
Eucalyptus___Eucgr.C03774.1.p
100
Eucalyptus___Eucgr.C03771.1.p
Beta___EL10Ac8g18400.1
Balfun___TRINITY_DN24305_c0_g1_i1.p1
Ombrophytum___TRINITY_DN17559_c0_g2_i1.p1
100
Lophophytum−pyramidale___TRINITY_DN477832_c0_g1_i1.p1
100
Lophophytum−pyramidale___TRINITY_DN477826_c0_g1_i1.p1
99
Lophophytum−mirabile___TRINITY_DN138002_c0_g1_i1.p1_complete
Malania___TRINITY_DN97271_c0_g1_i1.p1
Dendropemon___TRINITY_DN13887_c0_g1_i1.p1
Daen___TRINITY_DN9634_c0_g1_i7.p1
100
Daen___TRINITY_DN9634_c0_g1_i5.p1
100
Daen___TRINITY_DN9634_c0_g1_i3.p1
Daen___TRINITY_DN9634_c0_g1_i9.p1
Daen___TRINITY_DN9634_c0_g1_i6.p1
Daen___TRINITY_DN9634_c0_g1_i1.p1
Manihot___Manes.03G025700.2.p
Vaccinium___Vadar_g22280.t1
99
Hydrangea___Hyque.18G096900.1.p
Solanum___Solyc02g080350.1.1
Olea___Oeu017828.1
100
Mimulus___Migut.K00527.1.p
Lactuca___Lsat_1_v5_gn_8_96760.1
100
Helianthus___HanXRQChr16g0510641
Vitis___VIT_211s0016g03270.1
Quercus___Qurub.05G013500.1
Malus___MD01G1235900
F−Pvulgaris___Phvul.002G309900.1.p
100
F−Carietinum___Ca_05938
F−Lleucocephala___GDRZ01216353.1.p1
99
F−Lleucocephala___GDRZ01069172.1.p1
100
F−Prosopis___Palba_XP_028806579.1
100
F−Prosopis___Palba_XP_028751515.1
100
F−Akoa___GBYE01015282.1.p1
F−Acol___TRINITY_DN152597_c0_g1_i1.p1_complete

**OG0009596**

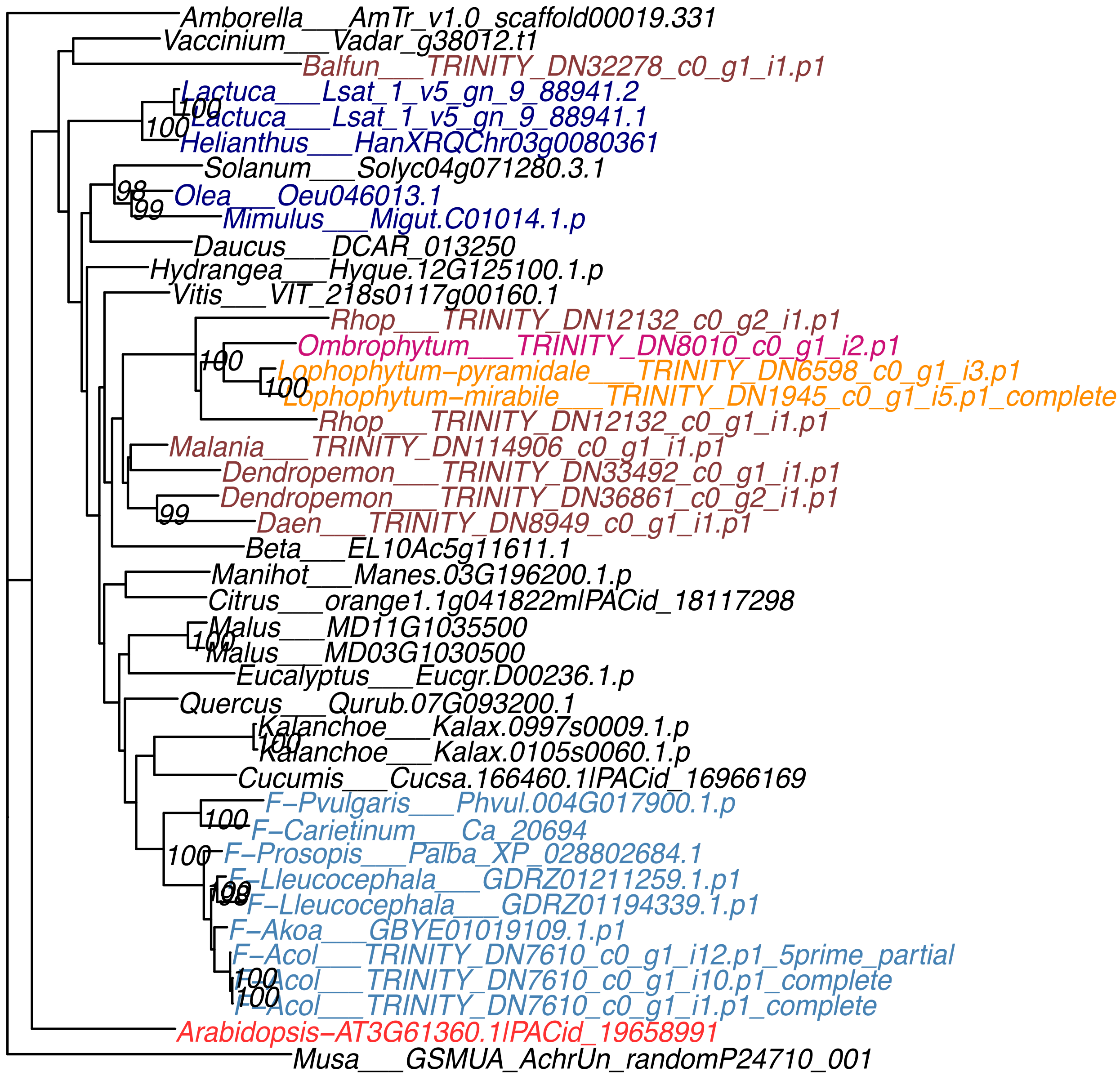
Amborella___AmTr_v1.0_scaffold00019.331
Vaccinium___Vadar_g38012.t1
Balfun___TRINITY_DN32278_c0_g1_i1.p1
Lactuca___Lsat_1_v5_gn_9_88941.2
100
Lactuca___Lsat_1_v5_gn_9_88941.1
100
Helianthus___HanXRQChr03g0080361
Solanum___Solyc04g071280.3.1
98
Olea___Oeu046013.1
99
Mimulus___Migut.C01014.1.p
Daucus___DCAR_013250
Hydrangea___Hyque.12G125100.1.p
Vitis___VIT_218s0117g00160.1
Rhop___TRINITY_DN12132_c0_g2_i1.p1
Ombrophytum___TRINITY_DN8010_c0_g1_i2.p1
100
Lophophytum−pyramidale___TRINITY_DN6598_c0_g1_i3.p1
100
Lophophytum−mirabile___TRINITY_DN1945_c0_g1_i5.p1_complete
Rhop___TRINITY_DN12132_c0_g1_i1.p1
Malania___TRINITY_DN114906_c0_g1_i1.p1
Dendropemon___TRINITY_DN33492_c0_g1_i1.p1
Dendropemon___TRINITY_DN36861_c0_g2_i1.p1
99
Daen___TRINITY_DN8949_c0_g1_i1.p1
Beta___EL10Ac5g11611.1
Manihot___Manes.03G196200.1.p
Citrus___orange1.1g041822mIPACid_18117298
Malus___MD11G1035500
100
Malus___MD03G1030500
Eucalyptus___Eucgr.D00236.1.p
Quercus___Qurub.07G093200.1
Kalanchoe___Kalax.0997s0009.1.p
100
Kalanchoe___Kalax.0105s0060.1.p
Cucumis___Cucsa.166460.1IPACid_16966169
F−Pvulgaris___Phvul.004G017900.1.p
100
F−Carietinum___Ca_20694
100
F−Prosopis___Palba_XP_028802684.1
F−Lleucocephala___GDRZ01211259.1.p1
F−Lleucocephala___GDRZ01194339.1.p1
F−Akoa___GBYE01019109.1.p1
F−Acol___TRINITY_DN7610_c0_g1_i12.p1_5prime_partial
100
F−Acol___TRINITY_DN7610_c0_g1_i10.p1_complete
100
F−Acol___TRINITY_DN7610_c0_g1_i1.p1_complete
Arabidopsis−AT3G61360.1IPACid_19658991
Musa___GSMUA_AchrUn_randomP24710_001

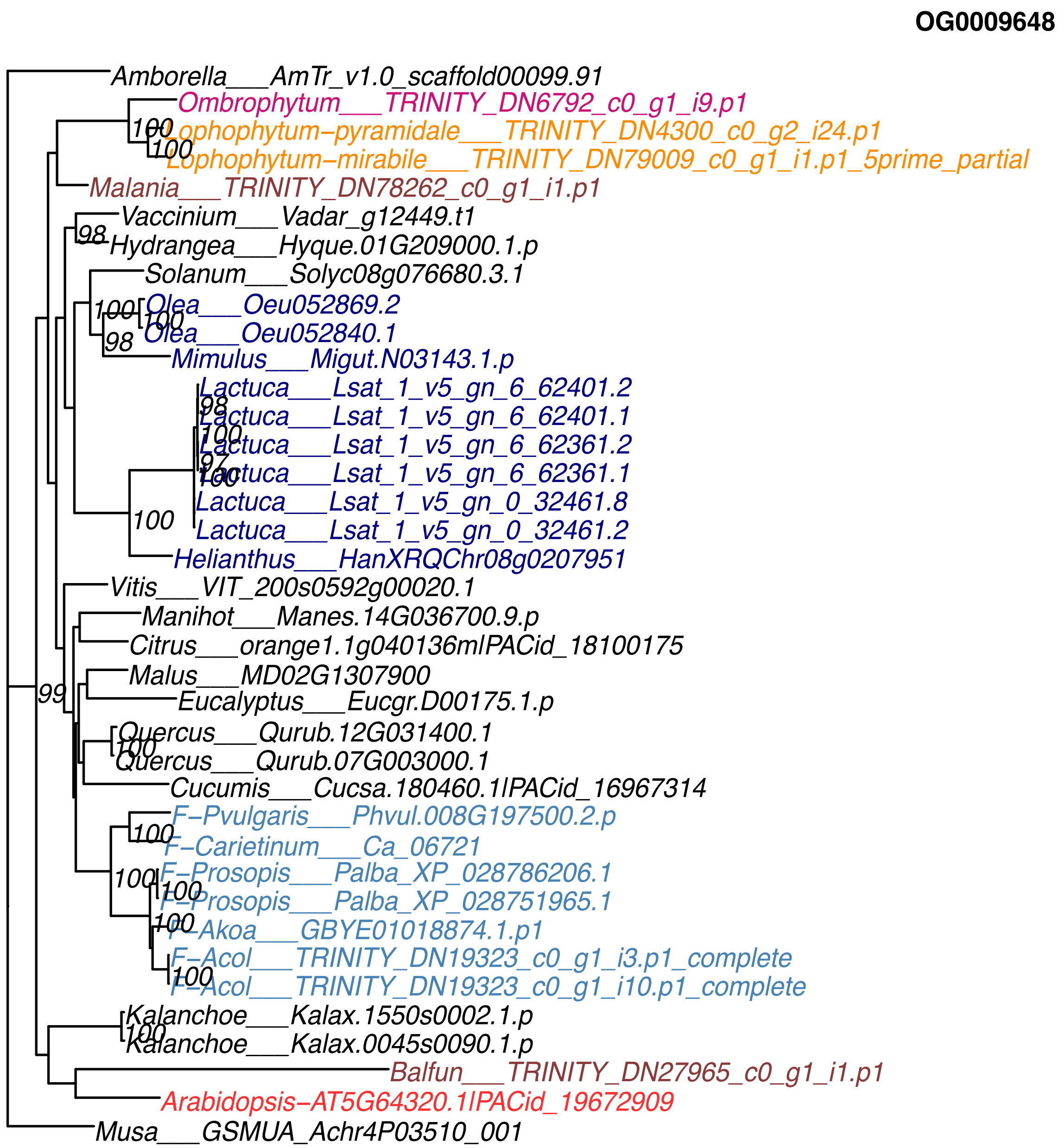

OG0009648
Amborella___AmTr_v1.0_scaffold00099.91
Ombrophytum___TRINITY_DN6792_c0_g1_i9.p1
Lophophytum−pyramidale___TRINITY_DN4300_c0_g2_i24.p1
Lophophytum−mirabile___TRINITY_DN79009_c0_g1_i1.p1_5prime_partial
Malania___TRINITY_DN78262_c0_g1_i1.p1
Vaccinium___Vadar_g12449.t1
Hydrangea___Hyque.01G209000.1.p
Solanum___Solyc08g076680.3.1
Olea___Oeu052869.2
Olea___Oeu052840.1
Mimulus___Migut.N03143.1.p
Lactuca___Lsat_1_v5_gn_6_62401.2
Lactuca___Lsat_1_v5_gn_6_62401.1
Lactuca___Lsat_1_v5_gn_6_62361.2
Lactuca___Lsat_1_v5_gn_6_62361.1
Lactuca___Lsat_1_v5_gn_0_32461.8
Lactuca___Lsat_1_v5_gn_0_32461.2
Helianthus___HanXRQChr08g0207951
Vitis___VIT_200s0592g00020.1
Manihot___Manes.14G036700.9.p
Citrus___orange1.1g040136mlPACid_18100175
Malus___MD02G1307900
Eucalyptus___Eucgr.D00175.1.p
Quercus___Qurub.12G031400.1
Quercus___Qurub.07G003000.1
Cucumis___Cucsa.180460.1lPACid_16967314
F−Pvulgaris___Phvul.008G197500.2.p
F−Carietinum___Ca_06721
F−Prosopis___Palba_XP_028786206.1
F−Prosopis___Palba_XP_028751965.1
F−Akoa___GBYE01018874.1.p1
F−Acol___TRINITY_DN19323_c0_g1_i3.p1_complete
F−Acol___TRINITY_DN19323_c0_g1_i10.p1_complete
Kalanchoe___Kalax.1550s0002.1.p
Kalanchoe___Kalax.0045s0090.1.p
Balfun___TRINITY_DN27965_c0_g1_i1.p1
Arabidopsis−AT5G64320.1lPACid_19672909
Musa___GSMUA_Achr4P03510_001
100
100
98
100
100
98
98
100
97
100
100
99
100
100
100
100
100
100
100

**OG0009670**

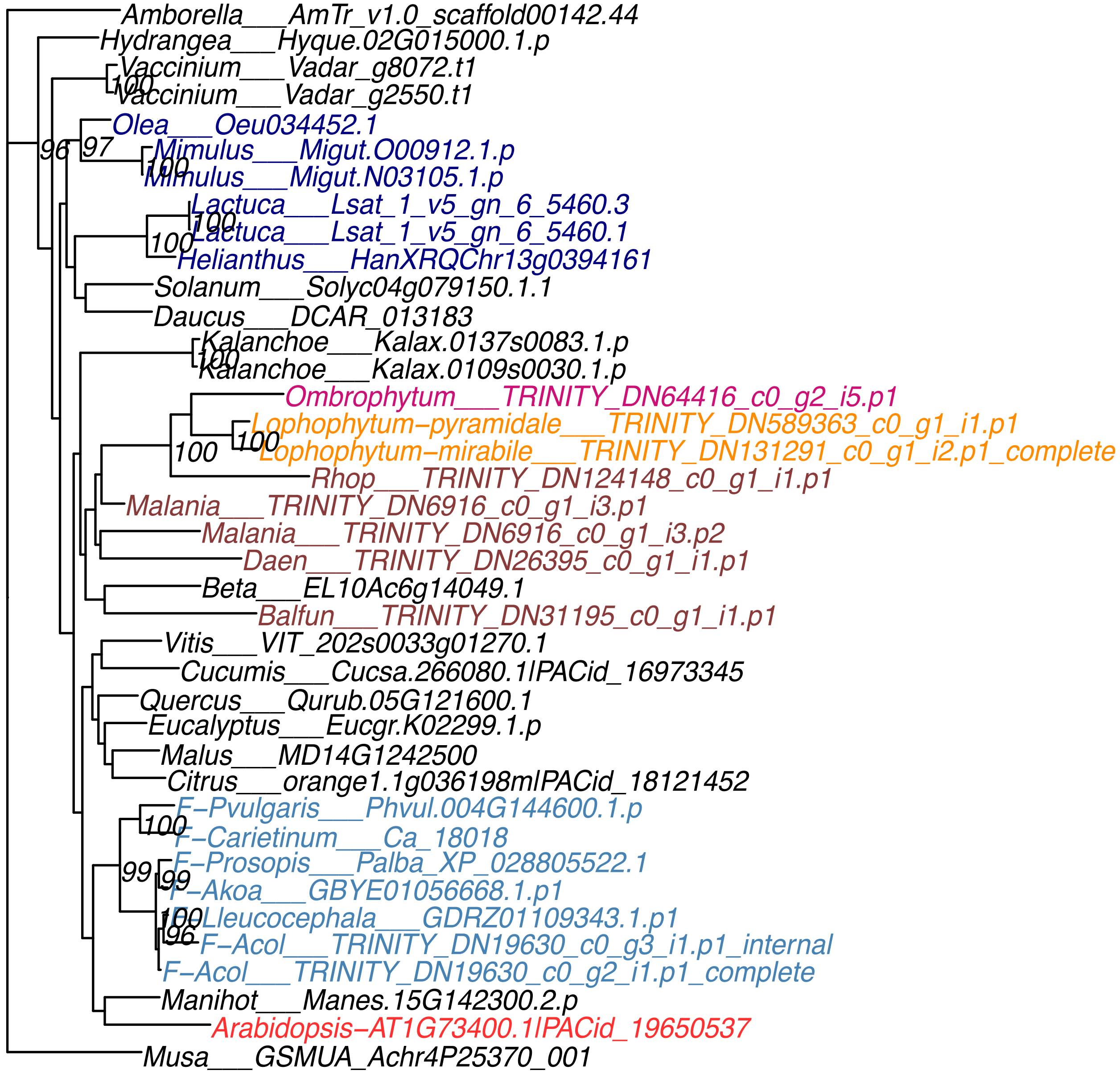
Amborella___AmTr_v1.0_scaffold00142.44
Hydrangea___Hyque.02G015000.1.p
Vaccinium___Vadar_g8072.t1
100
Vaccinium___Vadar_g2550.t1
Olea___Oeu034452.1
96
97
Mimulus___Migut.O00912.1.p
100
Mimulus___Migut.N03105.1.p
Lactuca___Lsat_1_v5_gn_6_5460.3
100
Lactuca___Lsat_1_v5_gn_6_5460.1
100
Helianthus___HanXRQChr13g0394161
Solanum___Solyc04g079150.1.1
Daucus___DCAR_013183
Kalanchoe___Kalax.0137s0083.1.p
100
Kalanchoe___Kalax.0109s0030.1.p
Ombrophytum___TRINITY_DN64416_c0_g2_i5.p1
Lophophytum−pyramidale___TRINITY_DN589363_c0_g1_i1.p1
100
100
Lophophytum−mirabile___TRINITY_DN131291_c0_g1_i2.p1_complete
Rhop___TRINITY_DN124148_c0_g1_i1.p1
Malania___TRINITY_DN6916_c0_g1_i3.p1
Malania___TRINITY_DN6916_c0_g1_i3.p2
Daen___TRINITY_DN26395_c0_g1_i1.p1
Beta___EL10Ac6g14049.1
Balfun___TRINITY_DN31195_c0_g1_i1.p1
Vitis___VIT_202s0033g01270.1
Cucumis___Cucsa.266080.1|PACid_16973345
Quercus___Qurub.05G121600.1
Eucalyptus___Eucgr.K02299.1.p
Malus___MD14G1242500
Citrus___orange1.1g036198m|PACid_18121452
F−Pvulgaris___Phvul.004G144600.1.p
100
F−Carietinum___Ca_18018
F−Prosopis___Palba_XP_028805522.1
99
99
F−Akoa___GBYE01056668.1.p1
100
F−Lleucocephala___GDRZ01109343.1.p1
96
F−Acol___TRINITY_DN19630_c0_g3_i1.p1_internal
F−Acol___TRINITY_DN19630_c0_g2_i1.p1_complete
Manihot___Manes.15G142300.2.p
Arabidopsis−AT1G73400.1|PACid_19650537
Musa___GSMUA_Achr4P25370_001

**OG0009682**

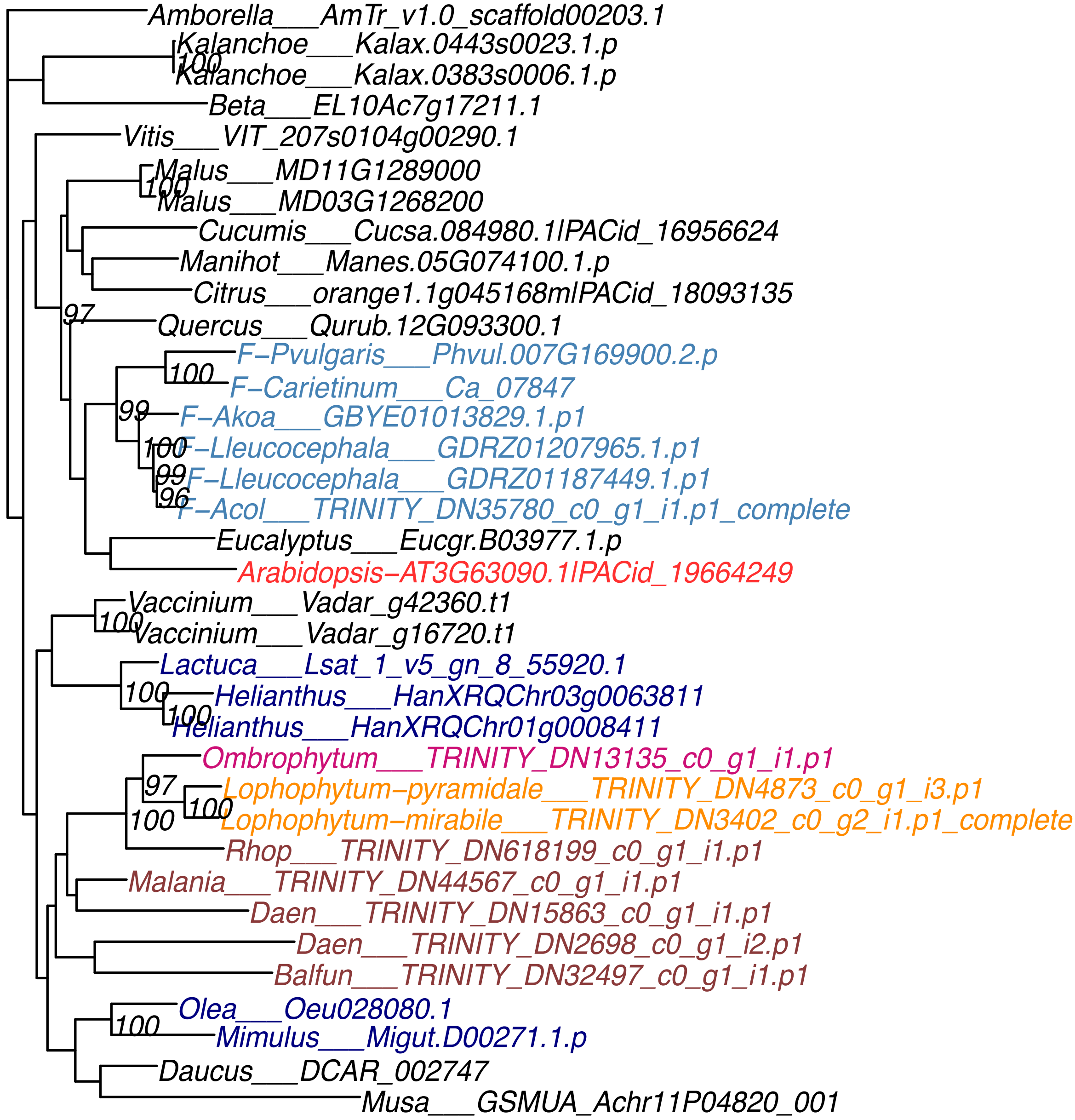

Amborella___AmTr_v1.0_scaffold00203.1
Kalanchoe___Kalax.0443s0023.1.p
100
Kalanchoe___Kalax.0383s0006.1.p
Beta___EL10Ac7g17211.1
Vitis___VIT_207s0104g00290.1
Malus___MD11G1289000
100
Malus___MD03G1268200
Cucumis___Cucsa.084980.1lPACid_16956624
Manihot___Manes.05G074100.1.p
Citrus___orange1.1g045168mlPACid_18093135
97
Quercus___Qurub.12G093300.1
F−Pvulgaris___Phvul.007G169900.2.p
100
F−Carietinum___Ca_07847
99
F−Akoa___GBYE01013829.1.p1
100
F−Lleucocephala___GDRZ01207965.1.p1
99
F−Lleucocephala___GDRZ01187449.1.p1
96
F−Acol___TRINITY_DN35780_c0_g1_i1.p1_complete
Eucalyptus___Eucgr.B03977.1.p
Arabidopsis−AT3G63090.1lPACid_19664249
Vaccinium___Vadar_g42360.t1
100
Vaccinium___Vadar_g16720.t1
Lactuca___Lsat_1_v5_gn_8_55920.1
100
Helianthus___HanXRQChr03g0063811
100
Helianthus___HanXRQChr01g0008411
Ombrophytum___TRINITY_DN13135_c0_g1_i1.p1
97
Lophophytum−pyramidale___TRINITY_DN4873_c0_g1_i3.p1
100
100
Lophophytum−mirabile___TRINITY_DN3402_c0_g2_i1.p1_complete
Rhop___TRINITY_DN618199_c0_g1_i1.p1
Malania___TRINITY_DN44567_c0_g1_i1.p1
Daen___TRINITY_DN15863_c0_g1_i1.p1
Daen___TRINITY_DN2698_c0_g1_i2.p1
Balfun___TRINITY_DN32497_c0_g1_i1.p1
Olea___Oeu028080.1
100
Mimulus___Migut.D00271.1.p
Daucus___DCAR_002747
Musa___GSMUA_Achr11P04820_001

OG0009690

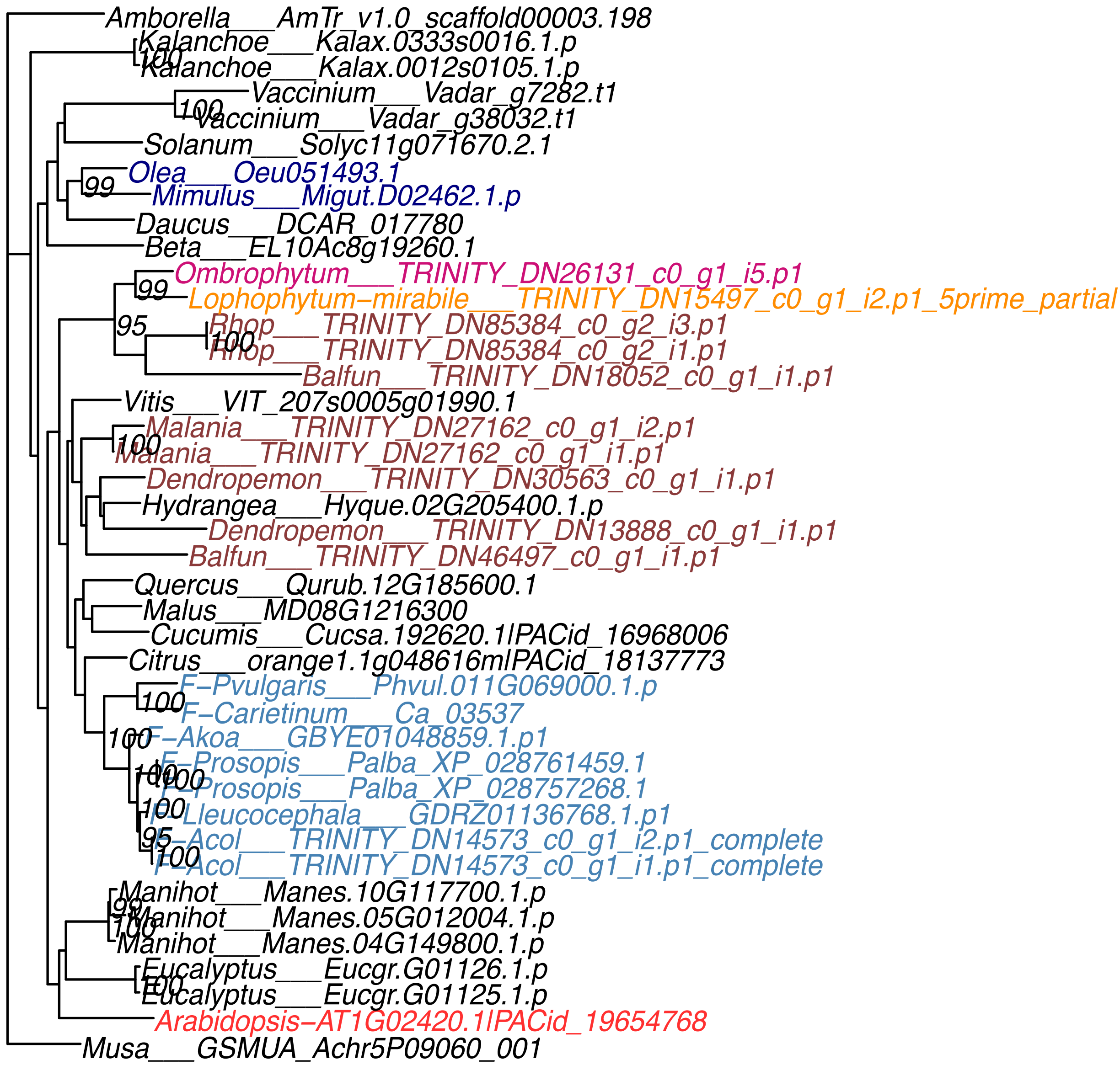

Amborella___AmTr_v1.0_scaffold00003.198
Kalanchoe___Kalax.0333s0016.1.p
100
Kalanchoe___Kalax.0012s0105.1.p
Vaccinium___Vadar_g7282.t1
100
Vaccinium___Vadar_g38032.t1
Solanum___Solyc11g071670.2.1
Olea___Oeu051493.1
99
Mimulus___Migut.D02462.1.p
Daucus___DCAR_017780
Beta___EL10Ac8g19260.1
Ombrophytum___TRINITY_DN26131_c0_g1_i5.p1
99
Lophophytum−mirabile___TRINITY_DN15497_c0_g1_i2.p1_5prime_partial
95
Rhop___TRINITY_DN85384_c0_g2_i3.p1
100
Rhop___TRINITY_DN85384_c0_g2_i1.p1
Balfun___TRINITY_DN18052_c0_g1_i1.p1
Vitis___VIT_207s0005g01990.1
Malania___TRINITY_DN27162_c0_g1_i2.p1
100
Malania___TRINITY_DN27162_c0_g1_i1.p1
Dendropemon___TRINITY_DN30563_c0_g1_i1.p1
Hydrangea___Hyque.02G205400.1.p
Dendropemon___TRINITY_DN13888_c0_g1_i1.p1
Balfun___TRINITY_DN46497_c0_g1_i1.p1
Quercus___Qurub.12G185600.1
Malus___MD08G1216300
Cucumis___Cucsa.192620.1|PACid_16968006
Citrus___orange1.1g048616m|PACid_18137773
F−Pvulgaris___Phvul.011G069000.1.p
100
F−Carietinum___Ca_03537
100
F−Akoa___GBYE01048859.1.p1
F−Prosopis___Palba_XP_028761459.1
100
100
F−Prosopis___Palba_XP_028757268.1
100
F−Lleucocephala___GDRZ01136768.1.p1
95
F−Acol___TRINITY_DN14573_c0_g1_i2.p1_complete
100
F−Acol___TRINITY_DN14573_c0_g1_i1.p1_complete
Manihot___Manes.10G117700.1.p
99
Manihot___Manes.05G012004.1.p
100
Manihot___Manes.04G149800.1.p
Eucalyptus___Eucgr.G01126.1.p
100
Eucalyptus___Eucgr.G01125.1.p
Arabidopsis−AT1G02420.1|PACid_19654768
Musa___GSMUA_Achr5P09060_001

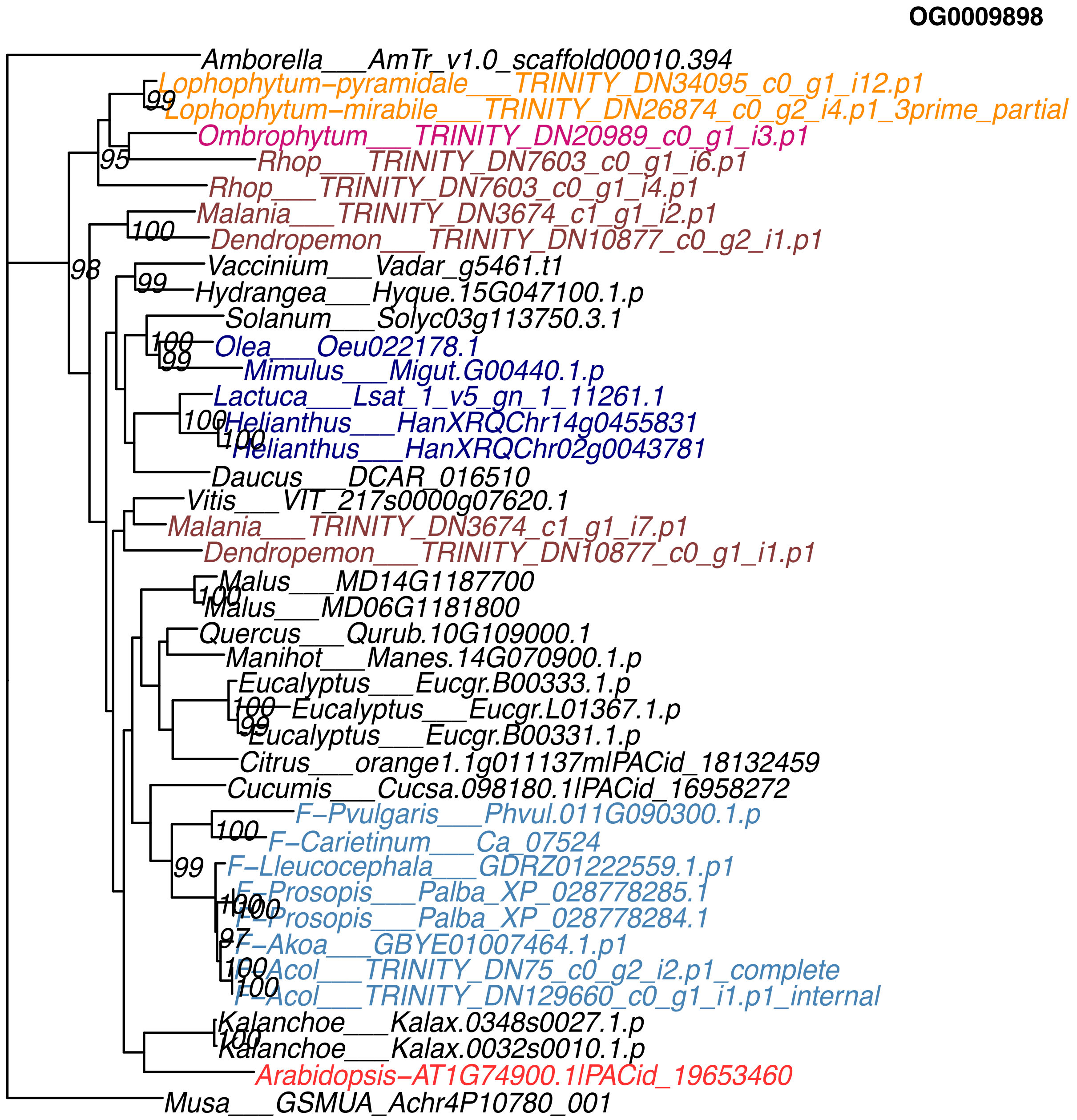

OG0009898
Amborella___AmTr_v1.0_scaffold00010.394
Lophophytum−pyramidale___TRINITY_DN34095_c0_g1_i12.p1
99
Lophophytum−mirabile___TRINITY_DN26874_c0_g2_i4.p1_3prime_partial
Ombrophytum___TRINITY_DN20989_c0_g1_i3.p1
95
Rhop___TRINITY_DN7603_c0_g1_i6.p1
Rhop___TRINITY_DN7603_c0_g1_i4.p1
Malania___TRINITY_DN3674_c1_g1_i2.p1
100
Dendropemon___TRINITY_DN10877_c0_g2_i1.p1
98
Vaccinium___Vadar_g5461.t1
99
Hydrangea___Hyque.15G047100.1.p
Solanum___Solyc03g113750.3.1
100
Olea___Oeu022178.1
99
Mimulus___Migut.G00440.1.p
Lactuca___Lsat_1_v5_gn_1_11261.1
100
Helianthus___HanXRQChr14g0455831
100
Helianthus___HanXRQChr02g0043781
Daucus___DCAR_016510
Vitis___VIT_217s0000g07620.1
Malania___TRINITY_DN3674_c1_g1_i7.p1
Dendropemon___TRINITY_DN10877_c0_g1_i1.p1
Malus___MD14G1187700
100
Malus___MD06G1181800
Quercus___Qurub.10G109000.1
Manihot___Manes.14G070900.1.p
Eucalyptus___Eucgr.B00333.1.p
100
Eucalyptus___Eucgr.L01367.1.p
99
Eucalyptus___Eucgr.B00331.1.p
Citrus___orange1.1g011137mlPACid_18132459
Cucumis___Cucsa.098180.1lPACid_16958272
F−Pvulgaris___Phvul.011G090300.1.p
100
F−Carietinum___Ca_07524
99
F−Lleucocephala___GDRZ01222559.1.p1
F−Prosopis___Palba_XP_028778285.1
100
100
F−Prosopis___Palba_XP_028778284.1
97
F−Akoa___GBYE01007464.1.p1
100
F−Acol___TRINITY_DN75_c0_g2_i2.p1_complete
100
F−Acol___TRINITY_DN129660_c0_g1_i1.p1_internal
Kalanchoe___Kalax.0348s0027.1.p
100
Kalanchoe___Kalax.0032s0010.1.p
Arabidopsis−AT1G74900.1lPACid_19653460
Musa___GSMUA_Achr4P10780_001

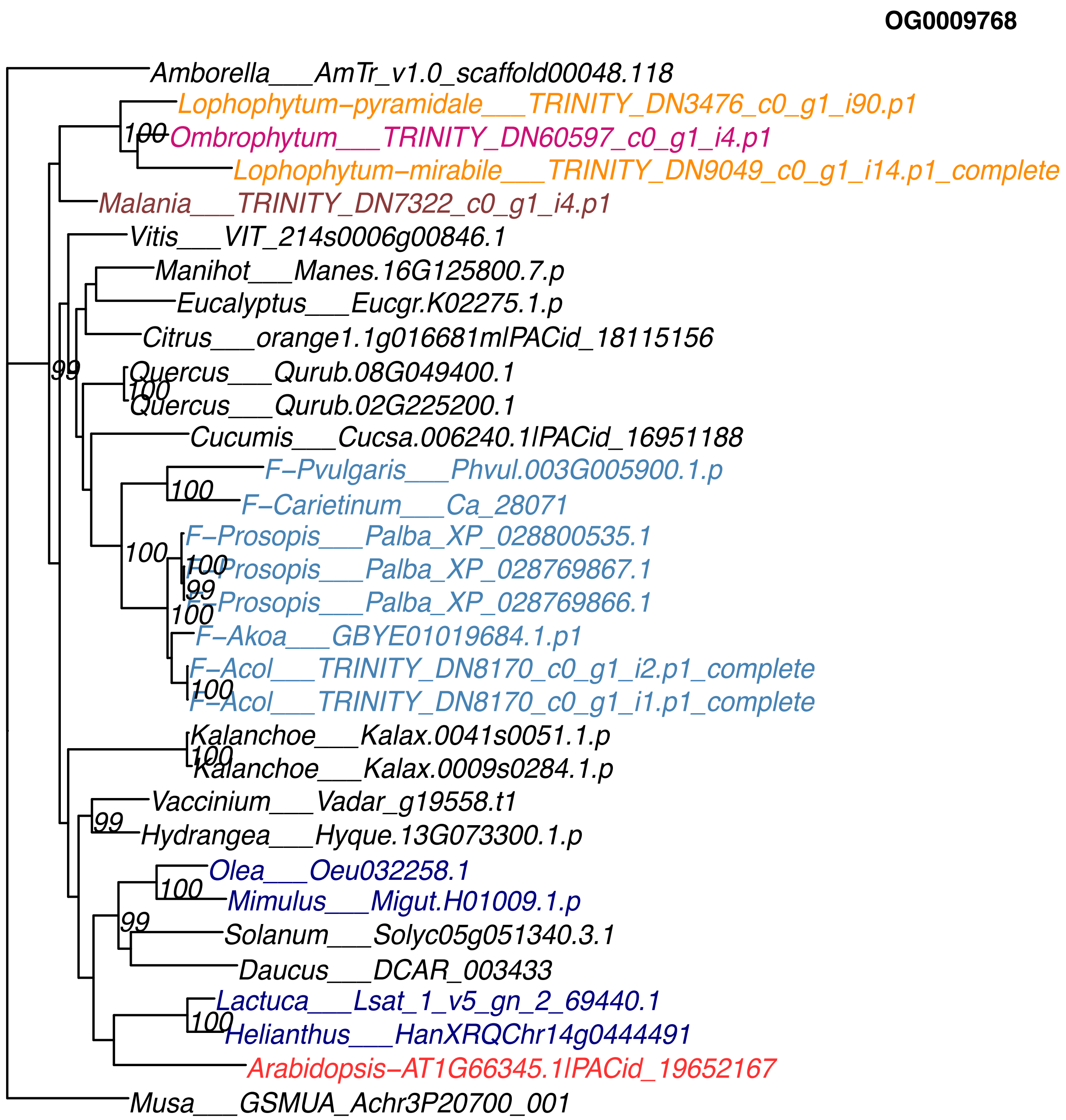
OG0009768
Amborella___AmTr_v1.0_scaffold00048.118
Lophophytum−pyramidale___TRINITY_DN3476_c0_g1_i90.p1
100
Ombrophytum___TRINITY_DN60597_c0_g1_i4.p1
Lophophytum−mirabile___TRINITY_DN9049_c0_g1_i14.p1_complete
Malania___TRINITY_DN7322_c0_g1_i4.p1
Vitis___VIT_214s0006g00846.1
Manihot___Manes.16G125800.7.p
Eucalyptus___Eucgr.K02275.1.p
Citrus___orange1.1g016681mlPACid_18115156
99
Quercus___Qurub.08G049400.1
100
Quercus___Qurub.02G225200.1
Cucumis___Cucsa.006240.1lPACid_16951188
F−Pvulgaris___Phvul.003G005900.1.p
100
F−Carietinum___Ca_28071
F−Prosopis___Palba_XP_028800535.1
100
100
F−Prosopis___Palba_XP_028769867.1
99
F−Prosopis___Palba_XP_028769866.1
100
F−Akoa___GBYE01019684.1.p1
F−Acol___TRINITY_DN8170_c0_g1_i2.p1_complete
100
F−Acol___TRINITY_DN8170_c0_g1_i1.p1_complete
Kalanchoe___Kalax.0041s0051.1.p
100
Kalanchoe___Kalax.0009s0284.1.p
Vaccinium___Vadar_g19558.t1
99
Hydrangea___Hyque.13G073300.1.p
Olea___Oeu032258.1
100
Mimulus___Migut.H01009.1.p
99
Solanum___Solyc05g051340.3.1
Daucus___DCAR_003433
Lactuca___Lsat_1_v5_gn_2_69440.1
100
Helianthus___HanXRQChr14g0444491
Arabidopsis−AT1G66345.1lPACid_19652167
Musa___GSMUA_Achr3P20700_001

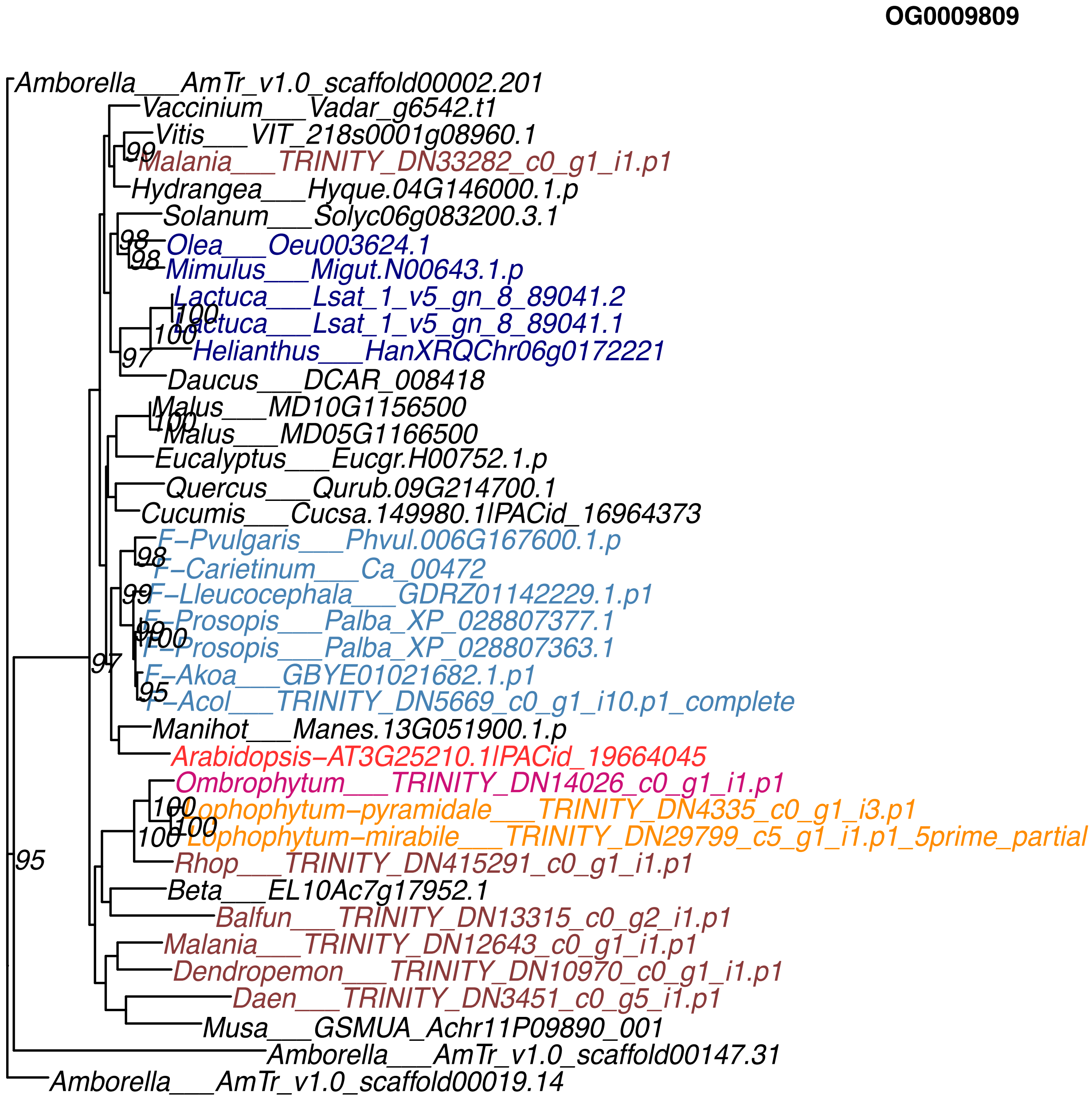
OG0009809
Amborella___AmTr_v1.0_scaffold00002.201
Vaccinium___Vadar_g6542.t1
Vitis___VIT_218s0001g08960.1
99
Malania___TRINITY_DN33282_c0_g1_i1.p1
Hydrangea___Hyque.04G146000.1.p
Solanum___Solyc06g083200.3.1
98
Olea___Oeu003624.1
98
Mimulus___Migut.N00643.1.p
Lactuca___Lsat_1_v5_gn_8_89041.2
100
Lactuca___Lsat_1_v5_gn_8_89041.1
100
Helianthus___HanXRQChr06g0172221
97
Daucus___DCAR_008418
Malus___MD10G1156500
100
Malus___MD05G1166500
Eucalyptus___Eucgr.H00752.1.p
Quercus___Qurub.09G214700.1
Cucumis___Cucsa.149980.1|PACid_16964373
F−Pvulgaris___Phvul.006G167600.1.p
98
F−Carietinum___Ca_00472
99
F−Lleucocephala___GDRZ01142229.1.p1
F−Prosopis___Palba_XP_028807377.1
99
100
F−Prosopis___Palba_XP_028807363.1
97
F−Akoa___GBYE01021682.1.p1
95
F−Acol___TRINITY_DN5669_c0_g1_i10.p1_complete
Manihot___Manes.13G051900.1.p
Arabidopsis−AT3G25210.1|PACid_19664045
Ombrophytum___TRINITY_DN14026_c0_g1_i1.p1
100
Lophophytum−pyramidale___TRINITY_DN4335_c0_g1_i3.p1
100
100
Lophophytum−mirabile___TRINITY_DN29799_c5_g1_i1.p1_5prime_partial
95
Rhop___TRINITY_DN415291_c0_g1_i1.p1
Beta___EL10Ac7g17952.1
Balfun___TRINITY_DN13315_c0_g2_i1.p1
Malania___TRINITY_DN12643_c0_g1_i1.p1
Dendropemon___TRINITY_DN10970_c0_g1_i1.p1
Daen___TRINITY_DN3451_c0_g5_i1.p1
Musa___GSMUA_Achr11P09890_001
Amborella___AmTr_v1.0_scaffold00147.31
Amborella___AmTr_v1.0_scaffold00019.14

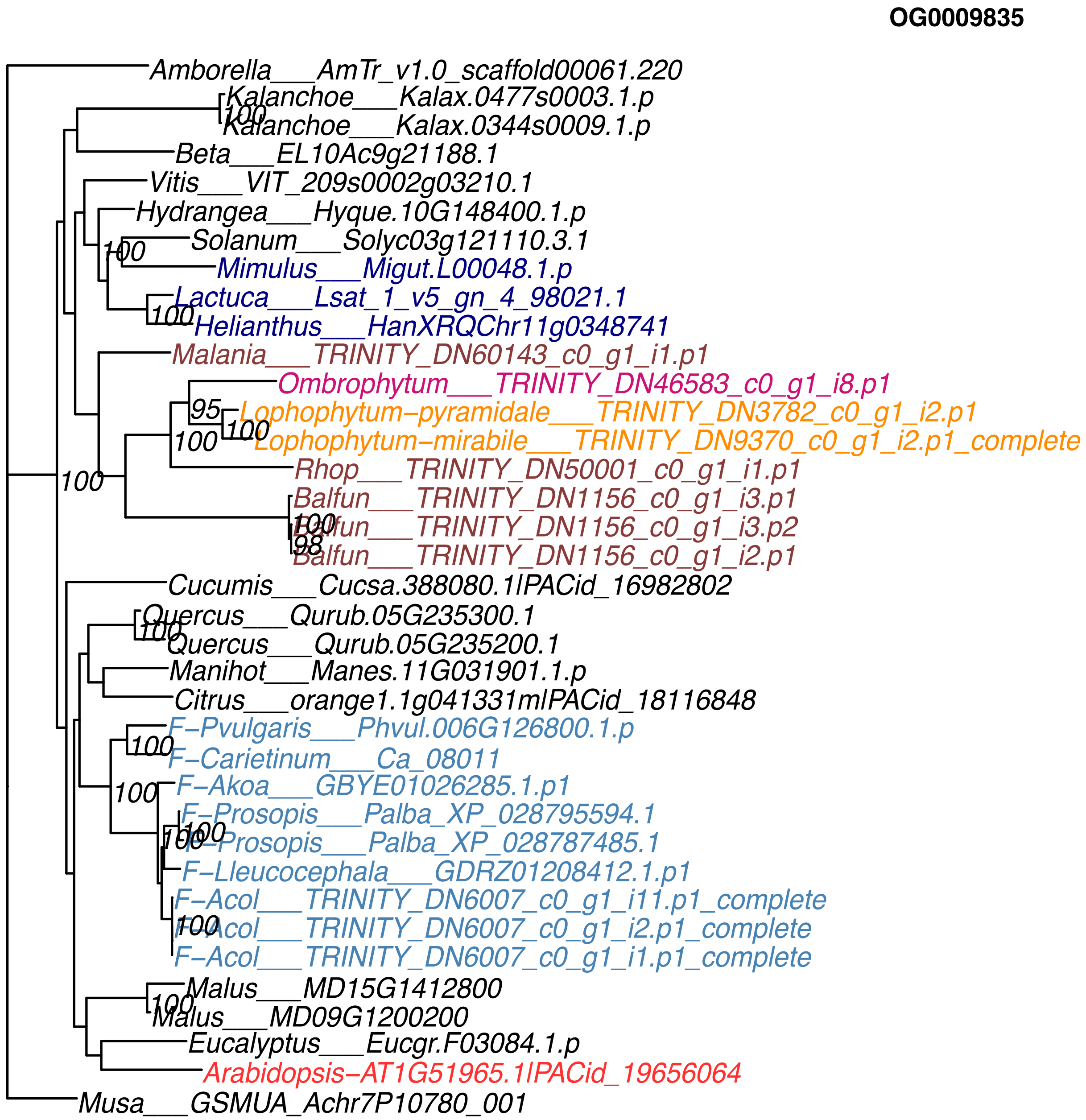
OG0009835
Amborella___AmTr_v1.0_scaffold00061.220
Kalanchoe___Kalax.0477s0003.1.p
Kalanchoe___Kalax.0344s0009.1.p
Beta___EL10Ac9g21188.1
Vitis___VIT_209s0002g03210.1
Hydrangea___Hyque.10G148400.1.p
Solanum___Solyc03g121110.3.1
Mimulus___Migut.L00048.1.p
Lactuca___Lsat_1_v5_gn_4_98021.1
Helianthus___HanXRQChr11g0348741
Malania___TRINITY_DN60143_c0_g1_i1.p1
Ombrophytum___TRINITY_DN46583_c0_g1_i8.p1
Lophophytum−pyramidale___TRINITY_DN3782_c0_g1_i2.p1
Lophophytum−mirabile___TRINITY_DN9370_c0_g1_i2.p1_complete
Rhop___TRINITY_DN50001_c0_g1_i1.p1
Balfun___TRINITY_DN1156_c0_g1_i3.p1
Balfun___TRINITY_DN1156_c0_g1_i3.p2
Balfun___TRINITY_DN1156_c0_g1_i2.p1
Cucumis___Cucsa.388080.1lPACid_16982802
Quercus___Qurub.05G235300.1
Quercus___Qurub.05G235200.1
Manihot___Manes.11G031901.1.p
Citrus___orange1.1g041331mlPACid_18116848
F−Pvulgaris___Phvul.006G126800.1.p
F−Carietinum___Ca_08011
F−Akoa___GBYE01026285.1.p1
F−Prosopis___Palba_XP_028795594.1
F−Prosopis___Palba_XP_028787485.1
F−Lleucocephala___GDRZ01208412.1.p1
F−Acol___TRINITY_DN6007_c0_g1_i11.p1_complete
F−Acol___TRINITY_DN6007_c0_g1_i2.p1_complete
F−Acol___TRINITY_DN6007_c0_g1_i1.p1_complete
Malus___MD15G1412800
Malus___MD09G1200200
Eucalyptus___Eucgr.F03084.1.p
Arabidopsis−AT1G51965.1lPACid_19656064
Musa___GSMUA_Achr7P10780_001

OG0009838

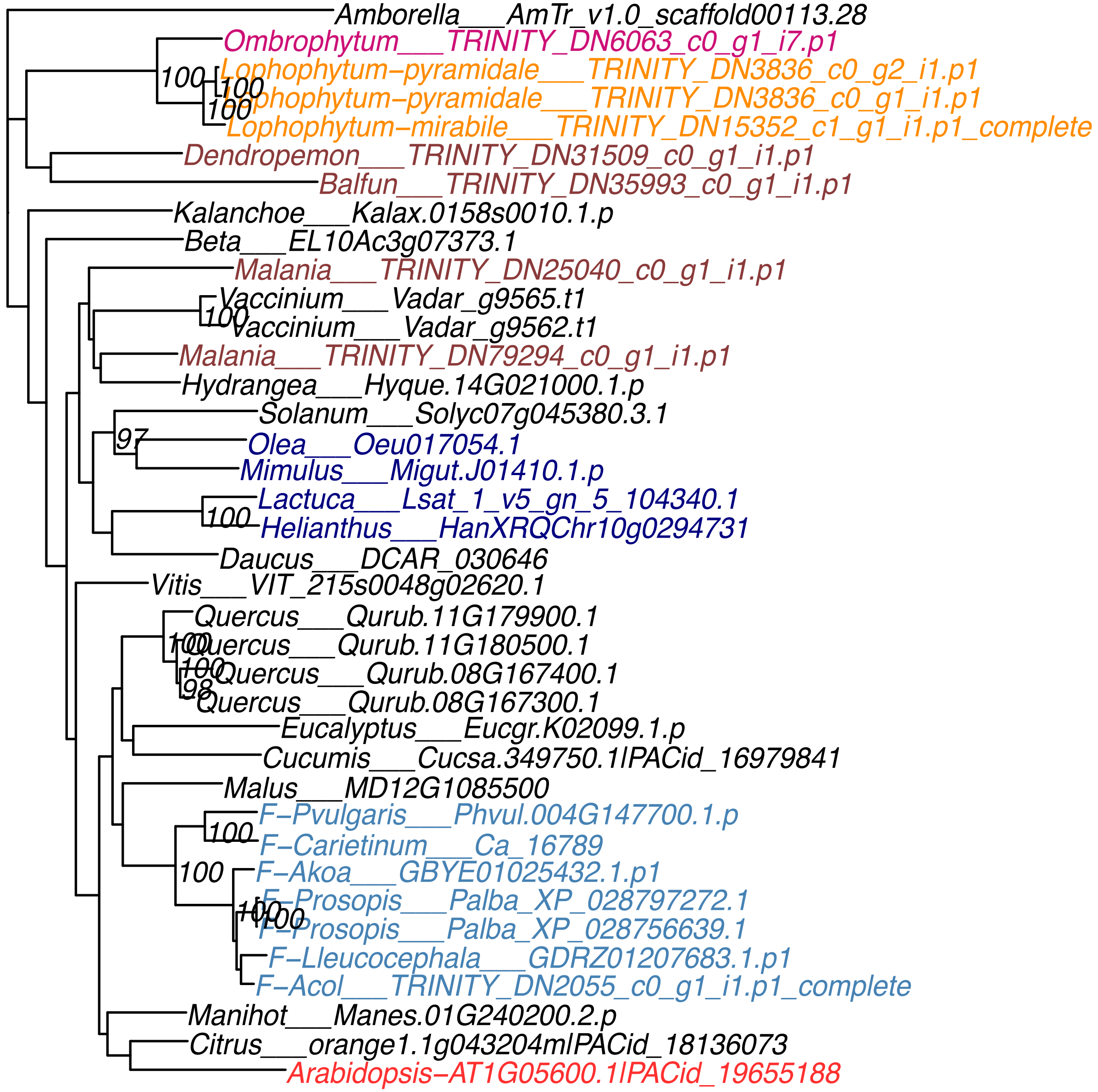
Amborella___AmTr_v1.0_scaffold00113.28
Ombrophytum___TRINITY_DN6063_c0_g1_i7.p1
Lophophytum−pyramidale___TRINITY_DN3836_c0_g2_i1.p1
Lophophytum−pyramidale___TRINITY_DN3836_c0_g1_i1.p1
Lophophytum−mirabile___TRINITY_DN15352_c1_g1_i1.p1_complete
Dendropemon___TRINITY_DN31509_c0_g1_i1.p1
Balfun___TRINITY_DN35993_c0_g1_i1.p1
Kalanchoe___Kalax.0158s0010.1.p
Beta___EL10Ac3g07373.1
Malania___TRINITY_DN25040_c0_g1_i1.p1
Vaccinium___Vadar_g9565.t1
Vaccinium___Vadar_g9562.t1
Malania___TRINITY_DN79294_c0_g1_i1.p1
Hydrangea___Hyque.14G021000.1.p
Solanum___Solyc07g045380.3.1
Olea___Oeu017054.1
Mimulus___Migut.J01410.1.p
Lactuca___Lsat_1_v5_gn_5_104340.1
Helianthus___HanXRQChr10g0294731
Daucus___DCAR_030646
Vitis___VIT_215s0048g02620.1
Quercus___Qurub.11G179900.1
Quercus___Qurub.11G180500.1
Quercus___Qurub.08G167400.1
Quercus___Qurub.08G167300.1
Eucalyptus___Eucgr.K02099.1.p
Cucumis___Cucsa.349750.1lPACid_16979841
Malus___MD12G1085500
F−Pvulgaris___Phvul.004G147700.1.p
F−Carietinum___Ca_16789
F−Akoa___GBYE01025432.1.p1
F−Prosopis___Palba_XP_028797272.1
F−Prosopis___Palba_XP_028756639.1
F−Lleucocephala___GDRZ01207683.1.p1
F−Acol___TRINITY_DN2055_c0_g1_i1.p1_complete
Manihot___Manes.01G240200.2.p
Citrus___orange1.1g043204mlPACid_18136073
Arabidopsis−AT1G05600.1lPACid_19655188
100
100
100
100
97
100
100
100
98
100
100
100
100

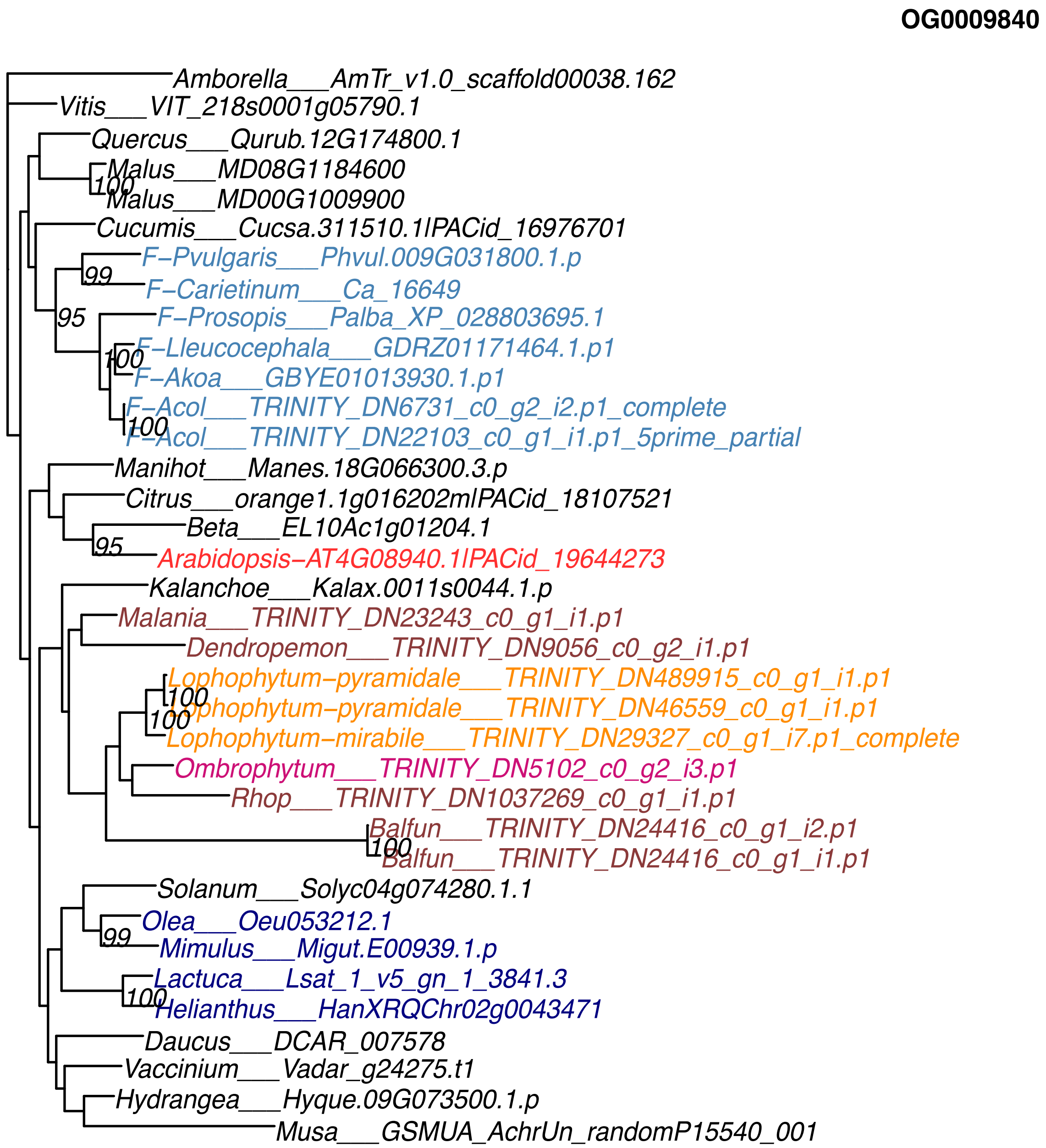
OG0009840
Amborella___AmTr_v1.0_scaffold00038.162
Vitis___VIT_218s0001g05790.1
Quercus___Qurub.12G174800.1
Malus___MD08G1184600
100
Malus___MD00G1009900
Cucumis___Cucsa.311510.1|PACid_16976701
F−Pvulgaris___Phvul.009G031800.1.p
99
F−Carietinum___Ca_16649
95
F−Prosopis___Palba_XP_028803695.1
F−Lleucocephala___GDRZ01171464.1.p1
100
F−Akoa___GBYE01013930.1.p1
F−Acol___TRINITY_DN6731_c0_g2_i2.p1_complete
100
F−Acol___TRINITY_DN22103_c0_g1_i1.p1_5prime_partial
Manihot___Manes.18G066300.3.p
Citrus___orange1.1g016202m|PACid_18107521
Beta___EL10Ac1g01204.1
95
Arabidopsis−AT4G08940.1|PACid_19644273
Kalanchoe___Kalax.0011s0044.1.p
Malania___TRINITY_DN23243_c0_g1_i1.p1
Dendropemon___TRINITY_DN9056_c0_g2_i1.p1
Lophophytum−pyramidale___TRINITY_DN489915_c0_g1_i1.p1
100
Lophophytum−pyramidale___TRINITY_DN46559_c0_g1_i1.p1
100
Lophophytum−mirabile___TRINITY_DN29327_c0_g1_i7.p1_complete
Ombrophytum___TRINITY_DN5102_c0_g2_i3.p1
Rhop___TRINITY_DN1037269_c0_g1_i1.p1
Balfun___TRINITY_DN24416_c0_g1_i2.p1
100
Balfun___TRINITY_DN24416_c0_g1_i1.p1
Solanum___Solyc04g074280.1.1
Olea___Oeu053212.1
99
Mimulus___Migut.E00939.1.p
Lactuca___Lsat_1_v5_gn_1_3841.3
100
Helianthus___HanXRQChr02g0043471
Daucus___DCAR_007578
Vaccinium___Vadar_g24275.t1
Hydrangea___Hyque.09G073500.1.p
Musa___GSMUA_AchrUn_randomP15540_001

**OG0009997**

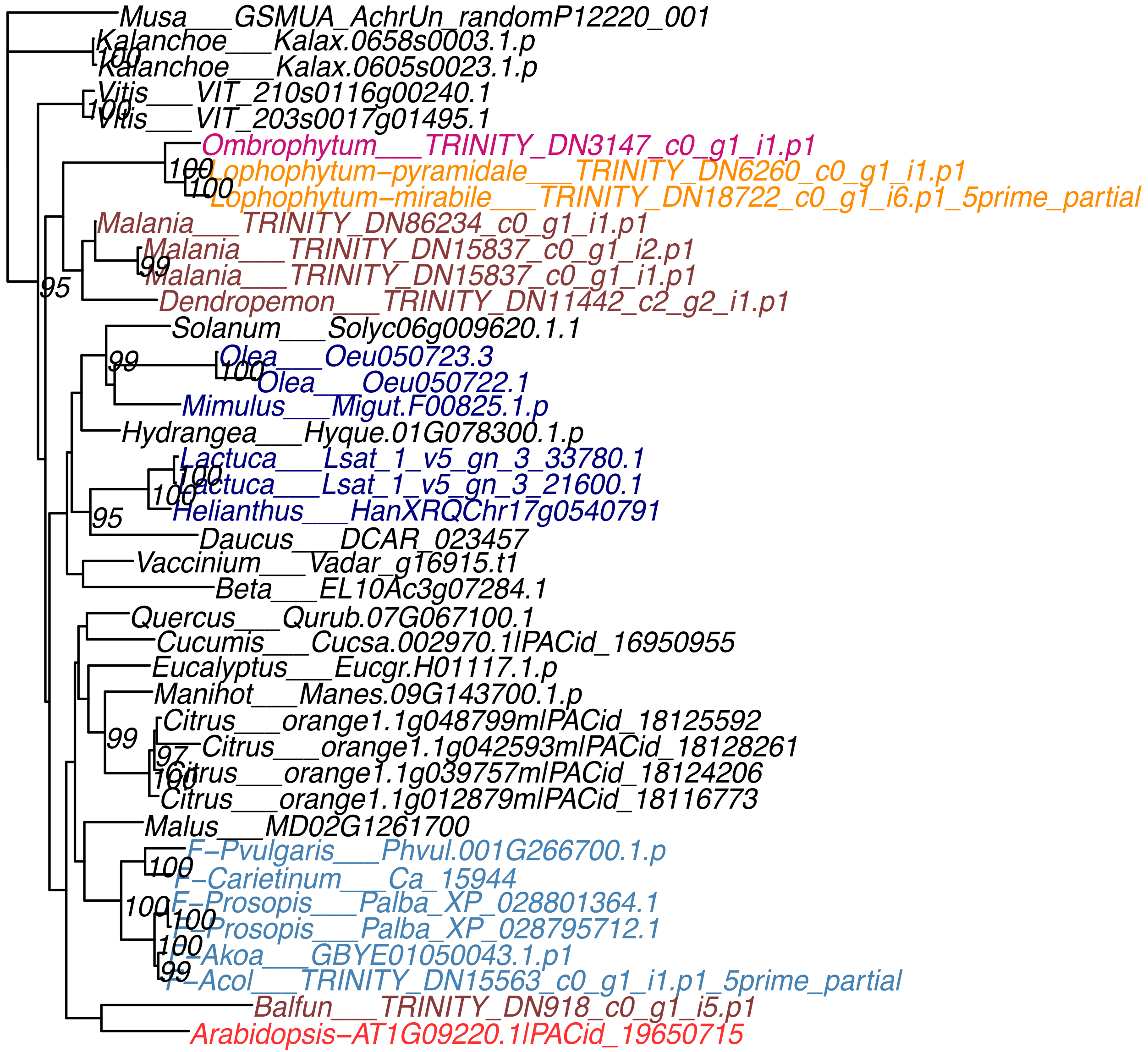
Musa___GSMUA_AchrUn_randomP12220_001
Kalanchoe___Kalax.0658s0003.1.p
100
Kalanchoe___Kalax.0605s0023.1.p
Vitis___VIT_210s0116g00240.1
100
Vitis___VIT_203s0017g01495.1
Ombrophytum___TRINITY_DN3147_c0_g1_i1.p1
100
Lophophytum−pyramidale___TRINITY_DN6260_c0_g1_i1.p1
100
Lophophytum−mirabile___TRINITY_DN18722_c0_g1_i6.p1_5prime_partial
Malania___TRINITY_DN86234_c0_g1_i1.p1
Malania___TRINITY_DN15837_c0_g1_i2.p1
99
Malania___TRINITY_DN15837_c0_g1_i1.p1
95
Dendropemon___TRINITY_DN11442_c2_g2_i1.p1
Solanum___Solyc06g009620.1.1
99
Olea___Oeu050723.3
100
Olea___Oeu050722.1
Mimulus___Migut.F00825.1.p
Hydrangea___Hyque.01G078300.1.p
Lactuca___Lsat_1_v5_gn_3_33780.1
100
Lactuca___Lsat_1_v5_gn_3_21600.1
100
Helianthus___HanXRQChr17g0540791
95
Daucus___DCAR_023457
Vaccinium___Vadar_g16915.t1
Beta___EL10Ac3g07284.1
Quercus___Qurub.07G067100.1
Cucumis___Cucsa.002970.1|PACid_16950955
Eucalyptus___Eucgr.H01117.1.p
Manihot___Manes.09G143700.1.p
Citrus___orange1.1g048799m|PACid_18125592
99
Citrus___orange1.1g042593m|PACid_18128261
97
Citrus___orange1.1g039757m|PACid_18124206
100
Citrus___orange1.1g012879m|PACid_18116773
Malus___MD02G1261700
F−Pvulgaris___Phvul.001G266700.1.p
100
F−Carietinum___Ca_15944
100
F−Prosopis___Palba_XP_028801364.1
100
F−Prosopis___Palba_XP_028795712.1
100
F−Akoa___GBYE01050043.1.p1
99
F−Acol___TRINITY_DN15563_c0_g1_i1.p1_5prime_partial
Balfun___TRINITY_DN918_c0_g1_i5.p1
Arabidopsis−AT1G09220.1|PACid_19650715

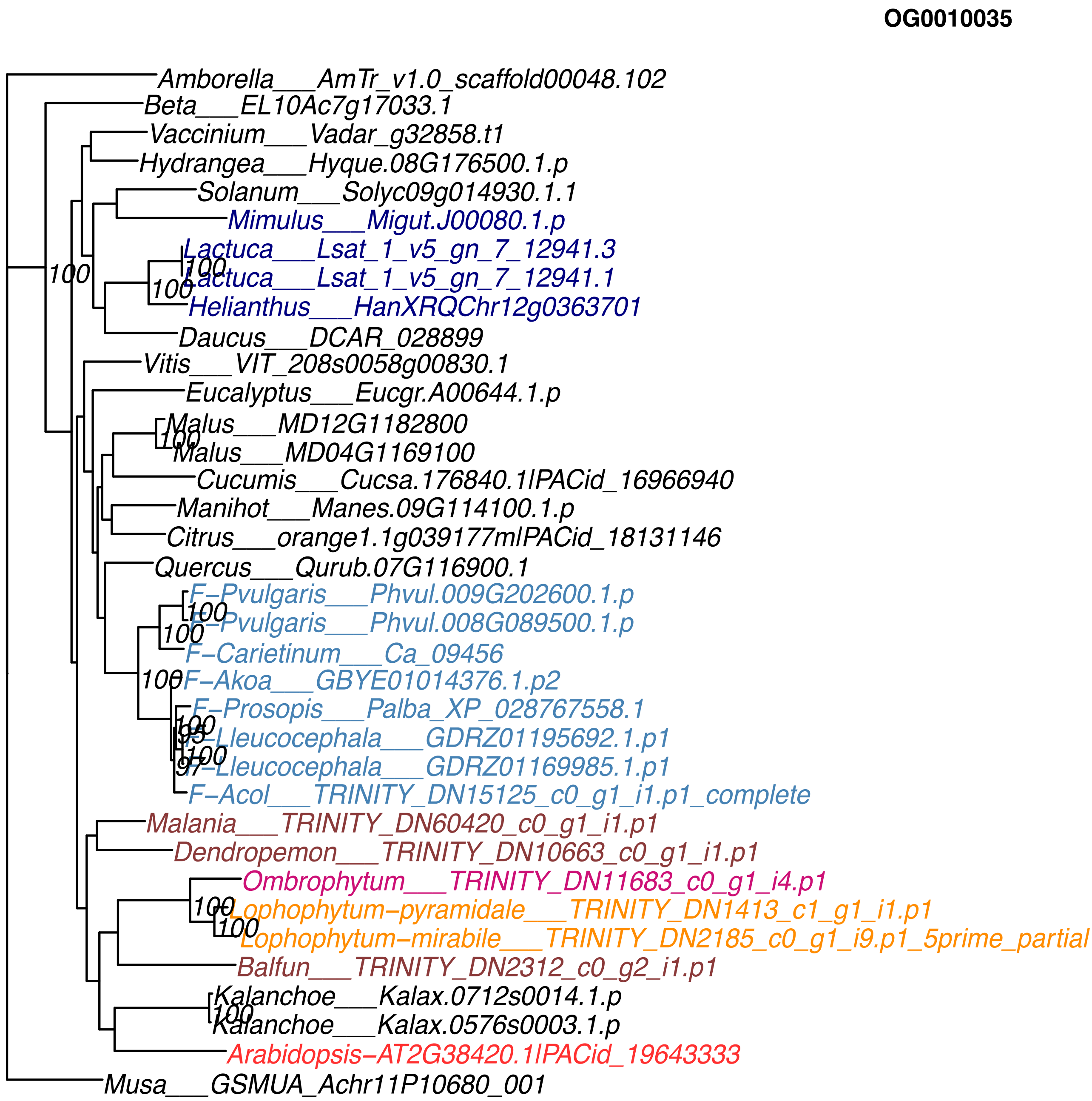
OG0010035
Amborella___AmTr_v1.0_scaffold00048.102
Beta___EL10Ac7g17033.1
Vaccinium___Vadar_g32858.t1
Hydrangea___Hyque.08G176500.1.p
Solanum___Solyc09g014930.1.1
Mimulus___Migut.J00080.1.p
Lactuca___Lsat_1_v5_gn_7_12941.3
Lactuca___Lsat_1_v5_gn_7_12941.1
Helianthus___HanXRQChr12g0363701
Daucus___DCAR_028899
Vitis___VIT_208s0058g00830.1
Eucalyptus___Eucgr.A00644.1.p
Malus___MD12G1182800
Malus___MD04G1169100
Cucumis___Cucsa.176840.1|PACid_16966940
Manihot___Manes.09G114100.1.p
Citrus___orange1.1g039177m|PACid_18131146
Quercus___Qurub.07G116900.1
F–Pvulgaris___Phvul.009G202600.1.p
F–Pvulgaris___Phvul.008G089500.1.p
F–Carietinum___Ca_09456
F–Akoa___GBYE01014376.1.p2
F–Prosopis___Palba_XP_028767558.1
F–Lleucocephala___GDRZ01195692.1.p1
F–Lleucocephala___GDRZ01169985.1.p1
F–Acol___TRINITY_DN15125_c0_g1_i1.p1_complete
Malania___TRINITY_DN60420_c0_g1_i1.p1
Dendropemon___TRINITY_DN10663_c0_g1_i1.p1
Ombrophytum___TRINITY_DN11683_c0_g1_i4.p1
Lophophytum–pyramidale___TRINITY_DN1413_c1_g1_i1.p1
Lophophytum–mirabile___TRINITY_DN2185_c0_g1_i9.p1_5prime_partial
Balfun___TRINITY_DN2312_c0_g2_i1.p1
Kalanchoe___Kalax.0712s0014.1.p
Kalanchoe___Kalax.0576s0003.1.p
Arabidopsis–AT2G38420.1|PACid_19643333
Musa___GSMUA_Achr11P10680_001
100
100
100
100
100
100
100
100
95
100
97
100
100
100

**OG0010043**

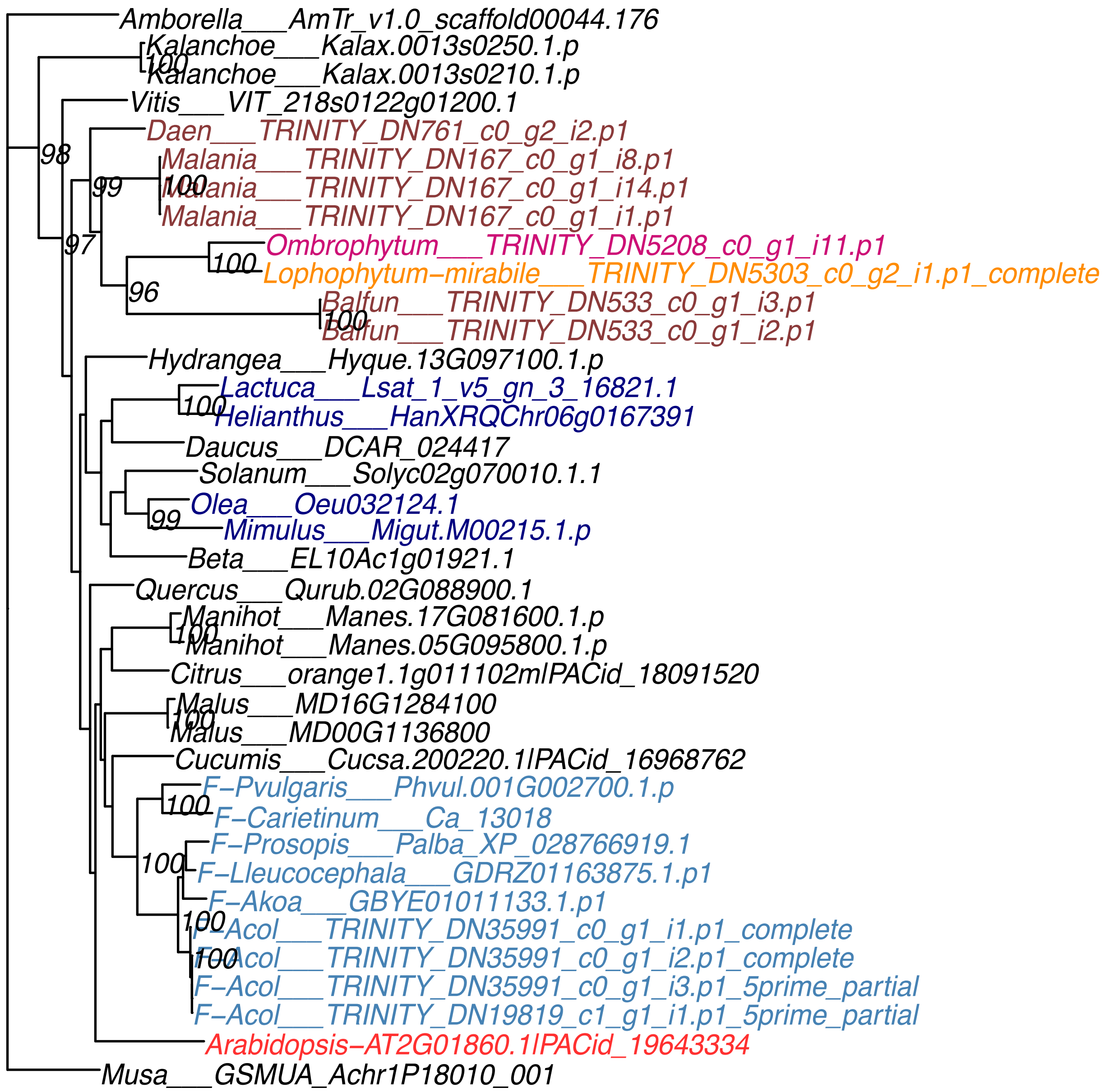

Amborella___AmTr_v1.0_scaffold00044.176
Kalanchoe___Kalax.0013s0250.1.p
100
Kalanchoe___Kalax.0013s0210.1.p
Vitis___VIT_218s0122g01200.1
98
Daen___TRINITY_DN761_c0_g2_i2.p1
Malania___TRINITY_DN167_c0_g1_i8.p1
100
Malania___TRINITY_DN167_c0_g1_i14.p1
99
Malania___TRINITY_DN167_c0_g1_i1.p1
97
Ombrophytum___TRINITY_DN5208_c0_g1_i11.p1
100
Lophophytum−mirabile___TRINITY_DN5303_c0_g2_i1.p1_complete
96
Balfun___TRINITY_DN533_c0_g1_i3.p1
100
Balfun___TRINITY_DN533_c0_g1_i2.p1
Hydrangea___Hyque.13G097100.1.p
Lactuca___Lsat_1_v5_gn_3_16821.1
100
Helianthus___HanXRQChr06g0167391
Daucus___DCAR_024417
Solanum___Solyc02g070010.1.1
Olea___Oeu032124.1
99
Mimulus___Migut.M00215.1.p
Beta___EL10Ac1g01921.1
Quercus___Qurub.02G088900.1
Manihot___Manes.17G081600.1.p
100
Manihot___Manes.05G095800.1.p
Citrus___orange1.1g011102mlPACid_18091520
Malus___MD16G1284100
100
Malus___MD00G1136800
Cucumis___Cucsa.200220.1lPACid_16968762
F−Pvulgaris___Phvul.001G002700.1.p
100
F−Carietinum___Ca_13018
F−Prosopis___Palba_XP_028766919.1
100
F−Lleucocephala___GDRZ01163875.1.p1
F−Akoa___GBYE01011133.1.p1
100
F−Acol___TRINITY_DN35991_c0_g1_i1.p1_complete
100
F−Acol___TRINITY_DN35991_c0_g1_i2.p1_complete
F−Acol___TRINITY_DN35991_c0_g1_i3.p1_5prime_partial
F−Acol___TRINITY_DN19819_c1_g1_i1.p1_5prime_partial
Arabidopsis−AT2G01860.1lPACid_19643334
Musa___GSMUA_Achr1P18010_001

**OG0010044**

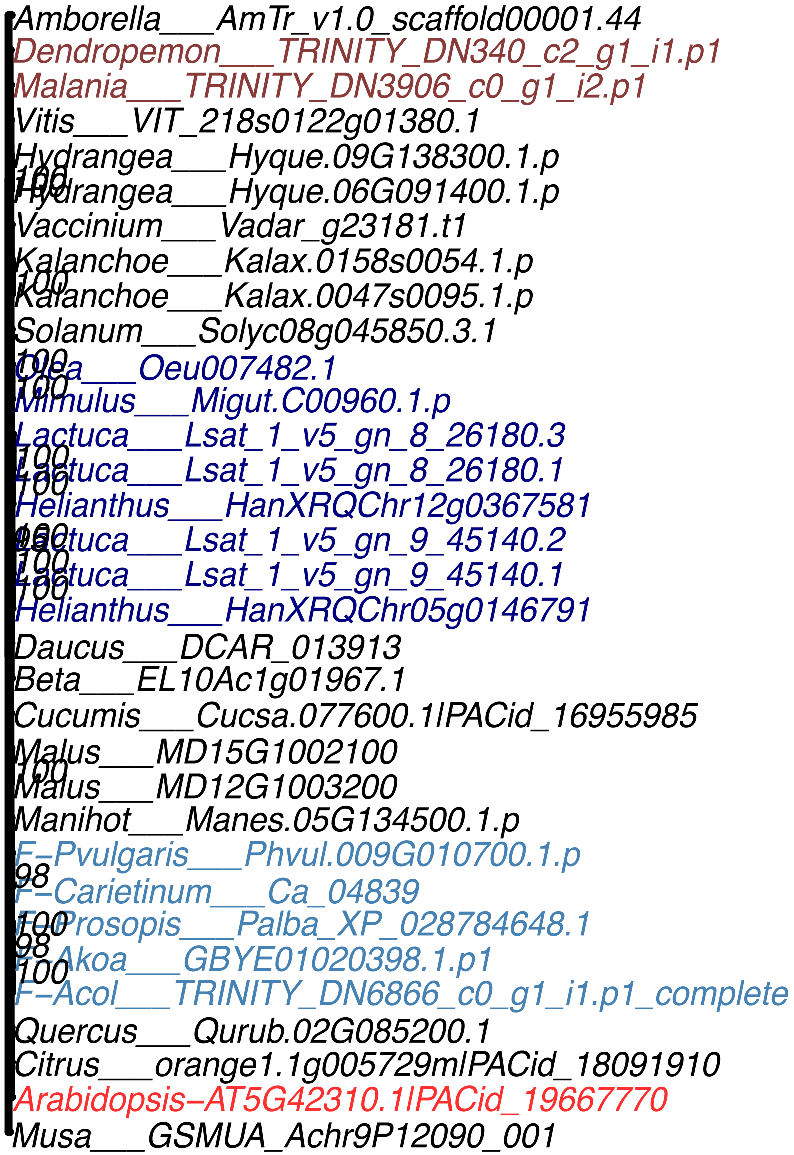

Amborella___AmTr_v1.0_scaffold00001.44
Dendropemon___TRINITY_DN340_c2_g1_i1.p1
Malania___TRINITY_DN3906_c0_g1_i2.p1
Vitis___VIT_218s0122g01380.1
Hydrangea___Hyque.09G138300.1.p
100
Hydrangea___Hyque.06G091400.1.p
Vaccinium___Vadar_g23181.t1
Kalanchoe___Kalax.0158s0054.1.p
100
Kalanchoe___Kalax.0047s0095.1.p
Solanum___Solyc08g045850.3.1
100
Olea___Oeu007482.1
100
Mimulus___Migut.C00960.1.p
Lactuca___Lsat_1_v5_gn_8_26180.3
100
Lactuca___Lsat_1_v5_gn_8_26180.1
100
Helianthus___HanXRQChr12g0367581
Lactuca___Lsat_1_v5_gn_9_45140.2
100
Lactuca___Lsat_1_v5_gn_9_45140.1
100
Helianthus___HanXRQChr05g0146791
Daucus___DCAR_013913
Beta___EL10Ac1g01967.1
Cucumis___Cucsa.077600.1|PACid_16955985
Malus___MD15G1002100
100
Malus___MD12G1003200
Manihot___Manes.05G134500.1.p
F–Pvulgaris___Phvul.009G010700.1.p
98
F–Carietinum___Ca_04839
100
F–Prosopis___Palba_XP_028784648.1
98
F–Akoa___GBYE01020398.1.p1
100
F–Acol___TRINITY_DN6866_c0_g1_i1.p1_complete
Quercus___Qurub.02G085200.1
Citrus___orange1.1g005729m|PACid_18091910
Arabidopsis–AT5G42310.1|PACid_19667770
Musa___GSMUA_Achr9P12090_001

OG0010261

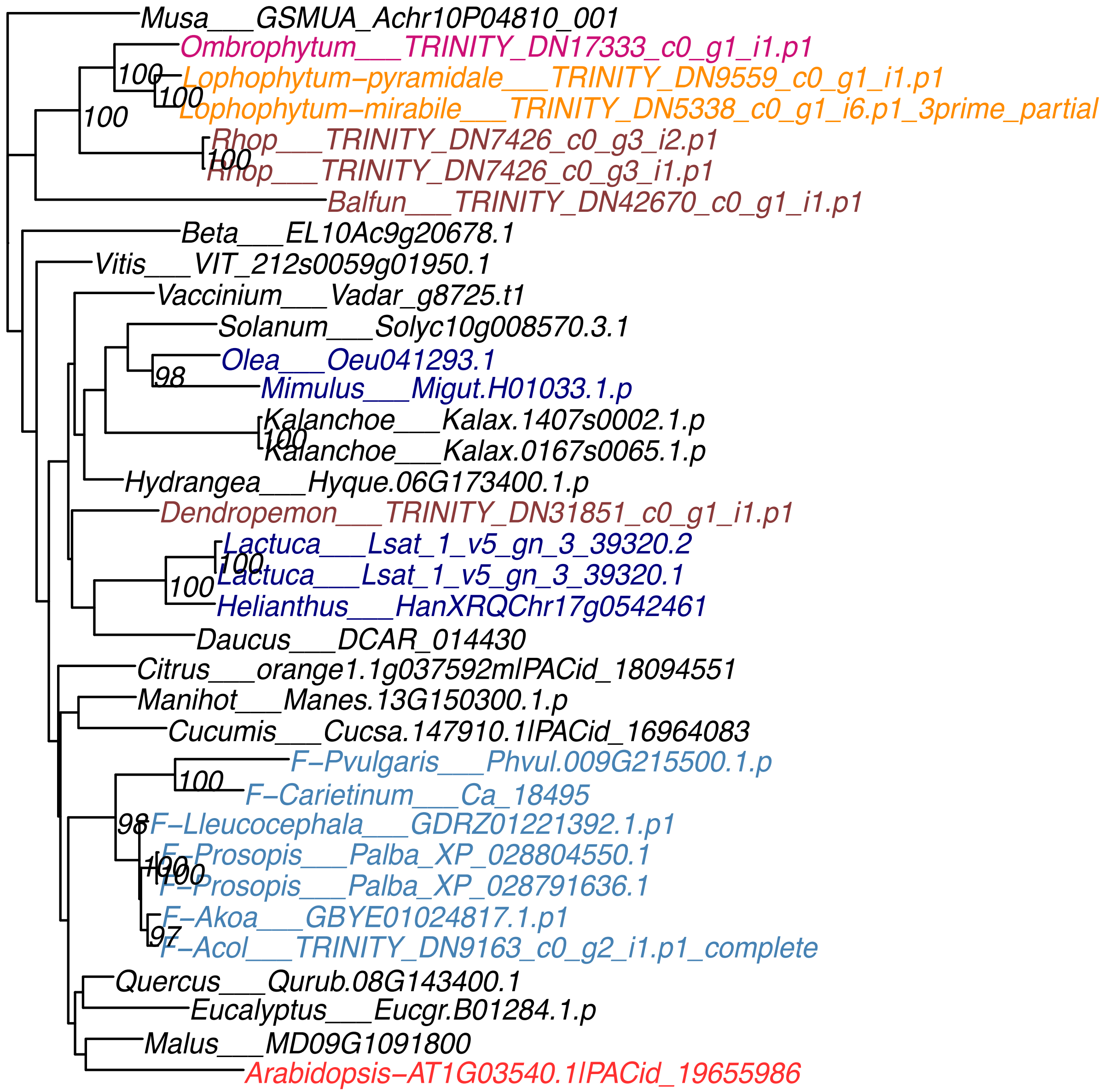
Musa___GSMUA_Achr10P04810_001
Ombrophytum___TRINITY_DN17333_c0_g1_i1.p1
Lophophytum−pyramidale___TRINITY_DN9559_c0_g1_i1.p1
Lophophytum−mirabile___TRINITY_DN5338_c0_g1_i6.p1_3prime_partial
Rhop___TRINITY_DN7426_c0_g3_i2.p1
Rhop___TRINITY_DN7426_c0_g3_i1.p1
Balfun___TRINITY_DN42670_c0_g1_i1.p1
Beta___EL10Ac9g20678.1
Vitis___VIT_212s0059g01950.1
Vaccinium___Vadar_g8725.t1
Solanum___Solyc10g008570.3.1
Olea___Oeu041293.1
Mimulus___Migut.H01033.1.p
Kalanchoe___Kalax.1407s0002.1.p
Kalanchoe___Kalax.0167s0065.1.p
Hydrangea___Hyque.06G173400.1.p
Dendropemon___TRINITY_DN31851_c0_g1_i1.p1
Lactuca___Lsat_1_v5_gn_3_39320.2
Lactuca___Lsat_1_v5_gn_3_39320.1
Helianthus___HanXRQChr17g0542461
Daucus___DCAR_014430
Citrus___orange1.1g037592mIPACid_18094551
Manihot___Manes.13G150300.1.p
Cucumis___Cucsa.147910.1IPACid_16964083
F−Pvulgaris___Phvul.009G215500.1.p
F−Carietinum___Ca_18495
F−Lleucocephala___GDRZ01221392.1.p1
F−Prosopis___Palba_XP_028804550.1
F−Prosopis___Palba_XP_028791636.1
F−Akoa___GBYE01024817.1.p1
F−Acol___TRINITY_DN9163_c0_g2_i1.p1_complete
Quercus___Qurub.08G143400.1
Eucalyptus___Eucgr.B01284.1.p
Malus___MD09G1091800
Arabidopsis−AT1G03540.1IPACid_19655986
100
100
100
100
98
100
100
100
100
98
100
100
97

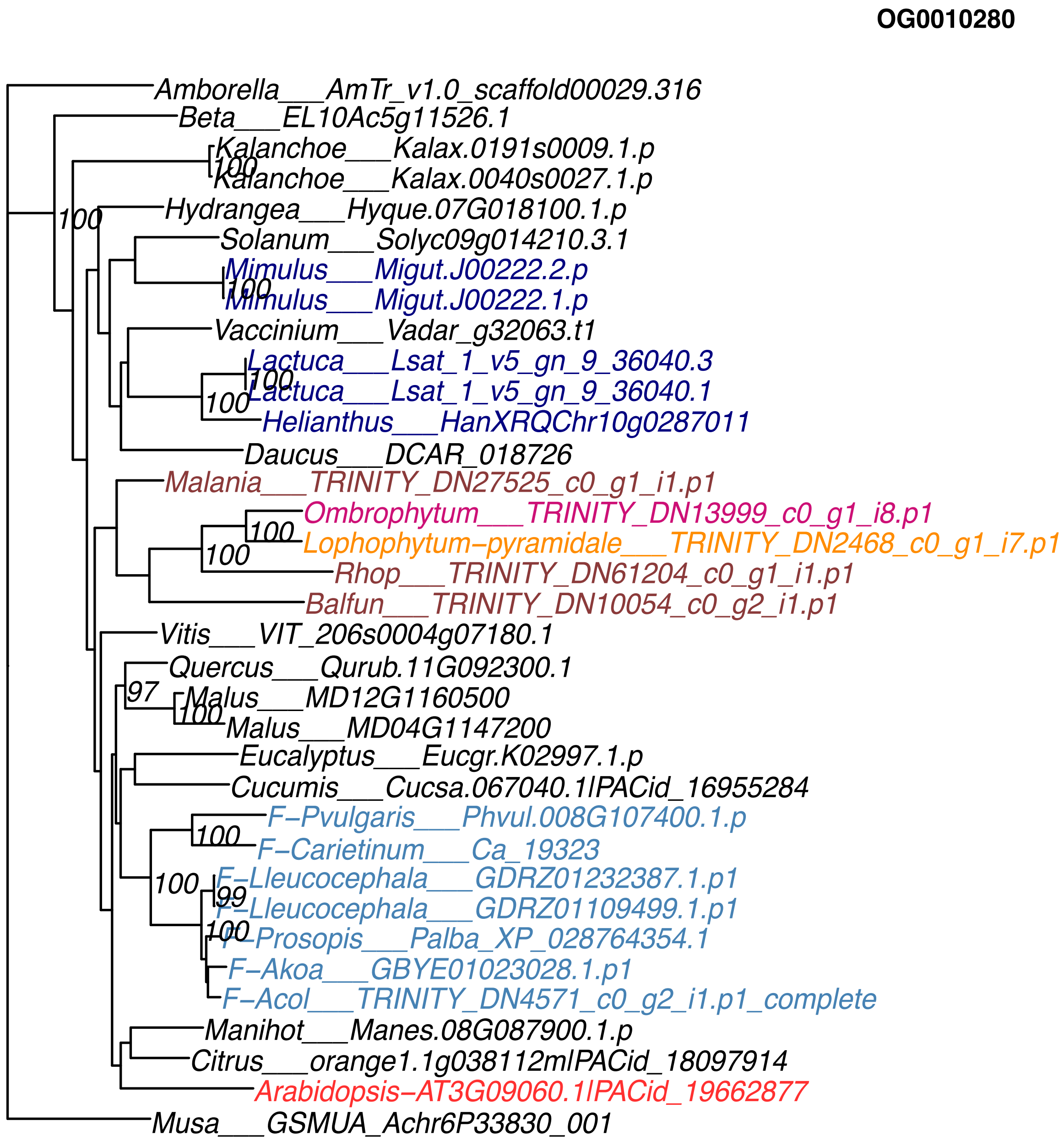
OG0010280
Amborella___AmTr_v1.0_scaffold00029.316
Beta___EL10Ac5g11526.1
Kalanchoe___Kalax.0191s0009.1.p
100
Kalanchoe___Kalax.0040s0027.1.p
100
Hydrangea___Hyque.07G018100.1.p
Solanum___Solyc09g014210.3.1
Mimulus___Migut.J00222.2.p
100
Mimulus___Migut.J00222.1.p
Vaccinium___Vadar_g32063.t1
Lactuca___Lsat_1_v5_gn_9_36040.3
100
Lactuca___Lsat_1_v5_gn_9_36040.1
100
Helianthus___HanXRQChr10g0287011
Daucus___DCAR_018726
Malania___TRINITY_DN27525_c0_g1_i1.p1
Ombrophytum___TRINITY_DN13999_c0_g1_i8.p1
100
Lophophytum–pyramidale___TRINITY_DN2468_c0_g1_i7.p1
100
Rhop___TRINITY_DN61204_c0_g1_i1.p1
Balfun___TRINITY_DN10054_c0_g2_i1.p1
Vitis___VIT_206s0004g07180.1
Quercus___Qurub.11G092300.1
97
Malus___MD12G1160500
100
Malus___MD04G1147200
Eucalyptus___Eucgr.K02997.1.p
Cucumis___Cucsa.067040.1|PACid_16955284
F–Pvulgaris___Phvul.008G107400.1.p
100
F–Carietinum___Ca_19323
100
F–Lleucocephala___GDRZ01232387.1.p1
99
F–Lleucocephala___GDRZ01109499.1.p1
100
F–Prosopis___Palba_XP_028764354.1
F–Akoa___GBYE01023028.1.p1
F–Acol___TRINITY_DN4571_c0_g2_i1.p1_complete
Manihot___Manes.08G087900.1.p
Citrus___orange1.1g038112m|PACid_18097914
Arabidopsis–AT3G09060.1|PACid_19662877
Musa___GSMUA_Achr6P33830_001

**OG0010287**

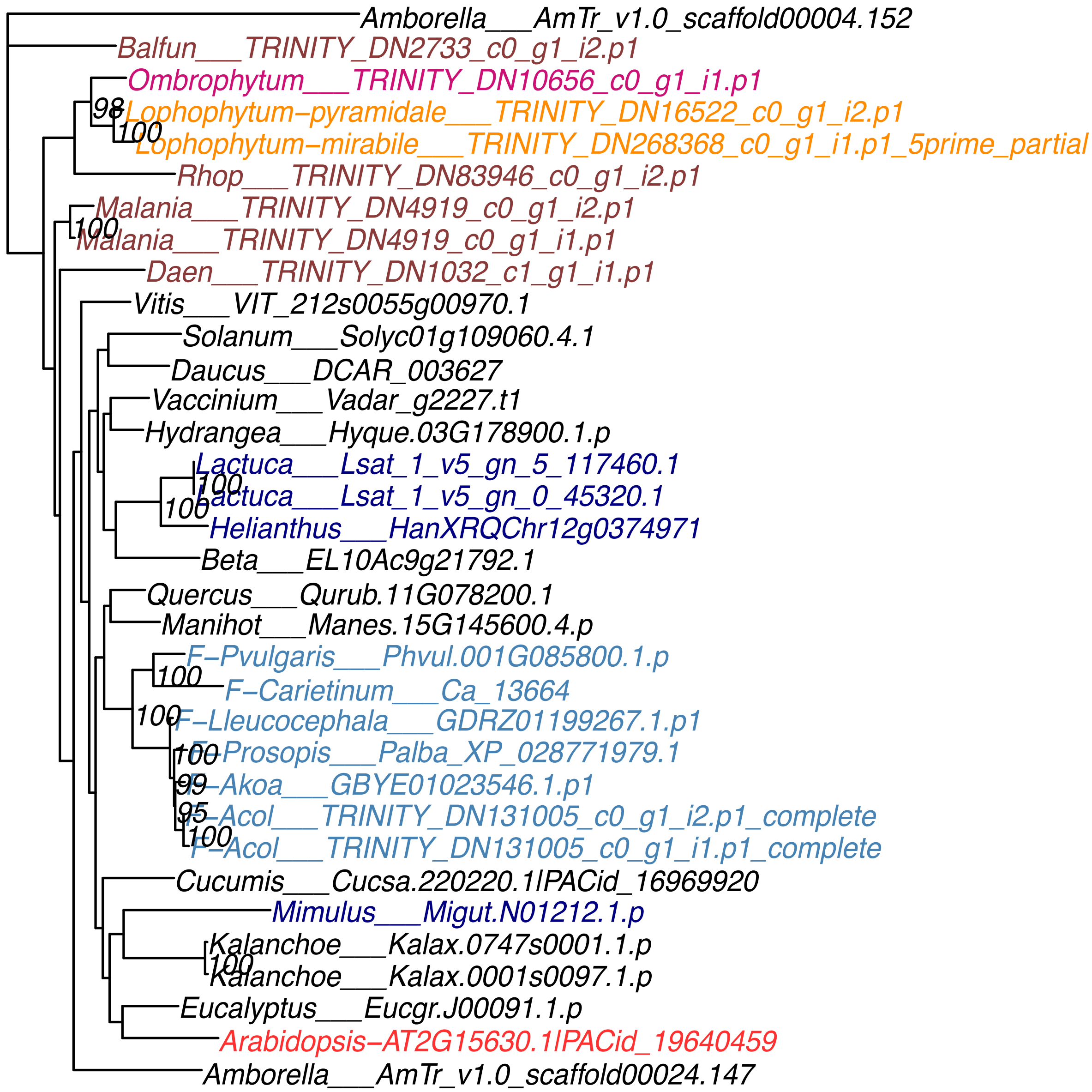
Amborella___AmTr_v1.0_scaffold00004.152
Balfun___TRINITY_DN2733_c0_g1_i2.p1
Ombrophytum___TRINITY_DN10656_c0_g1_i1.p1
98
Lophophytum−pyramidale___TRINITY_DN16522_c0_g1_i2.p1
100
Lophophytum−mirabile___TRINITY_DN268368_c0_g1_i1.p1_5prime_partial
Rhop___TRINITY_DN83946_c0_g1_i2.p1
Malania___TRINITY_DN4919_c0_g1_i2.p1
100
Malania___TRINITY_DN4919_c0_g1_i1.p1
Daen___TRINITY_DN1032_c1_g1_i1.p1
Vitis___VIT_212s0055g00970.1
Solanum___Solyc01g109060.4.1
Daucus___DCAR_003627
Vaccinium___Vadar_g2227.t1
Hydrangea___Hyque.03G178900.1.p
Lactuca___Lsat_1_v5_gn_5_117460.1
100
Lactuca___Lsat_1_v5_gn_0_45320.1
100
Helianthus___HanXRQChr12g0374971
Beta___EL10Ac9g21792.1
Quercus___Qurub.11G078200.1
Manihot___Manes.15G145600.4.p
F−Pvulgaris___Phvul.001G085800.1.p
100
F−Carietinum___Ca_13664
100
F−Lleucocephala___GDRZ01199267.1.p1
100
F−Prosopis___Palba_XP_028771979.1
99
F−Akoa___GBYE01023546.1.p1
95
F−Acol___TRINITY_DN131005_c0_g1_i2.p1_complete
100
F−Acol___TRINITY_DN131005_c0_g1_i1.p1_complete
Cucumis___Cucsa.220220.1lPACid_16969920
Mimulus___Migut.N01212.1.p
Kalanchoe___Kalax.0747s0001.1.p
100
Kalanchoe___Kalax.0001s0097.1.p
Eucalyptus___Eucgr.J00091.1.p
Arabidopsis−AT2G15630.1lPACid_19640459
Amborella___AmTr_v1.0_scaffold00024.147

**OG0010291**

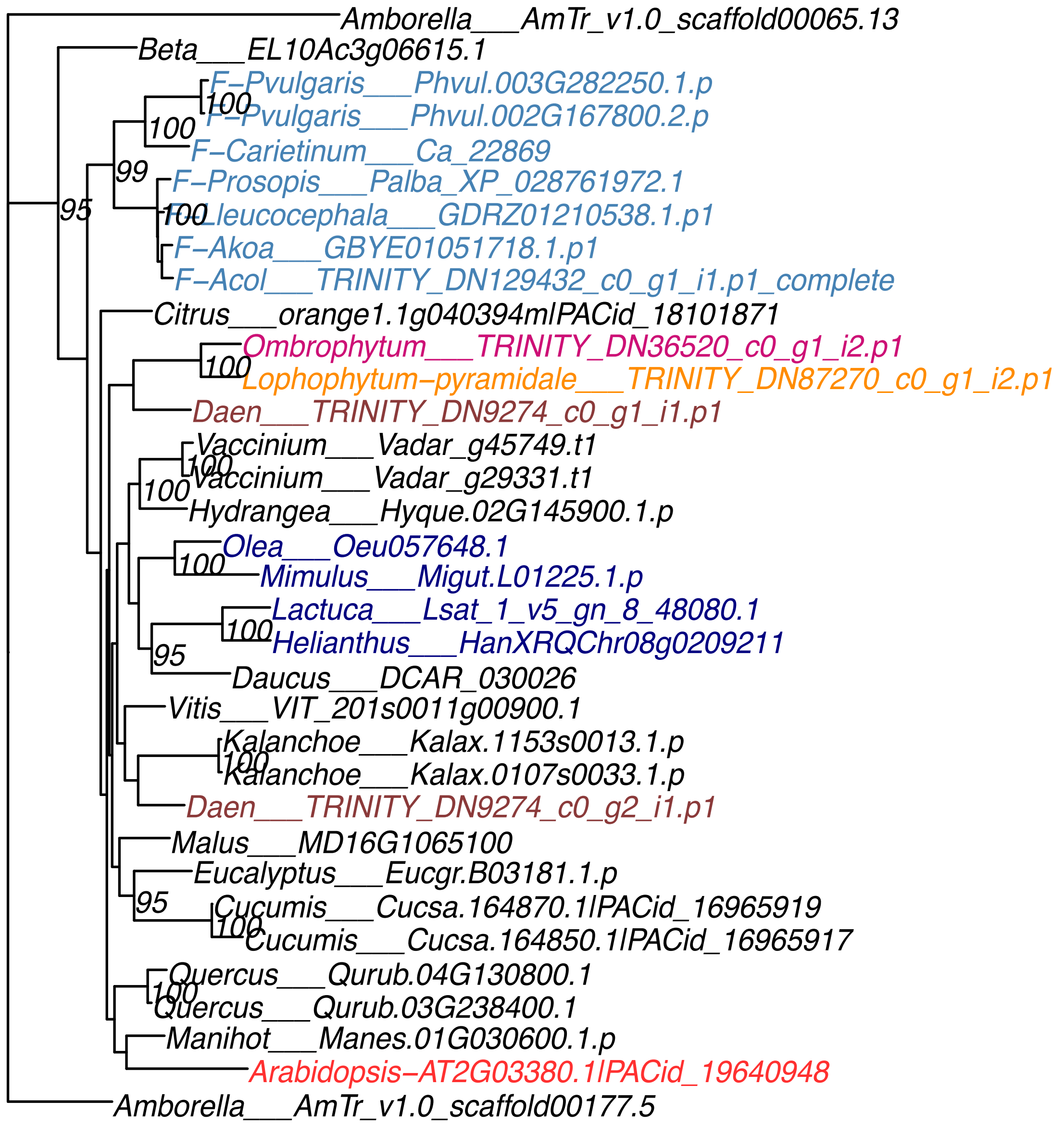

Amborella___AmTr_v1.0_scaffold00065.13
Beta___EL10Ac3g06615.1
F−Pvulgaris___Phvul.003G282250.1.p
100
F−Pvulgaris___Phvul.002G167800.2.p
100
F−Carietinum___Ca_22869
99
F−Prosopis___Palba_XP_028761972.1
95
100
F−Lleucocephala___GDRZ01210538.1.p1
F−Akoa___GBYE01051718.1.p1
F−Acol___TRINITY_DN129432_c0_g1_i1.p1_complete
Citrus___orange1.1g040394m|PACid_18101871
Ombrophytum___TRINITY_DN36520_c0_g1_i2.p1
100
Lophophytum−pyramidale___TRINITY_DN87270_c0_g1_i2.p1
Daen___TRINITY_DN9274_c0_g1_i1.p1
Vaccinium___Vadar_g45749.t1
100
Vaccinium___Vadar_g29331.t1
100
Hydrangea___Hyque.02G145900.1.p
Olea___Oeu057648.1
100
Mimulus___Migut.L01225.1.p
Lactuca___Lsat_1_v5_gn_8_48080.1
100
Helianthus___HanXRQChr08g0209211
95
Daucus___DCAR_030026
Vitis___VIT_201s0011g00900.1
Kalanchoe___Kalax.1153s0013.1.p
100
Kalanchoe___Kalax.0107s0033.1.p
Daen___TRINITY_DN9274_c0_g2_i1.p1
Malus___MD16G1065100
Eucalyptus___Eucgr.B03181.1.p
95
Cucumis___Cucsa.164870.1|PACid_16965919
100
Cucumis___Cucsa.164850.1|PACid_16965917
Quercus___Qurub.04G130800.1
100
Quercus___Qurub.03G238400.1
Manihot___Manes.01G030600.1.p
Arabidopsis−AT2G03380.1|PACid_19640948
Amborella___AmTr_v1.0_scaffold00177.5

**OG0010334**

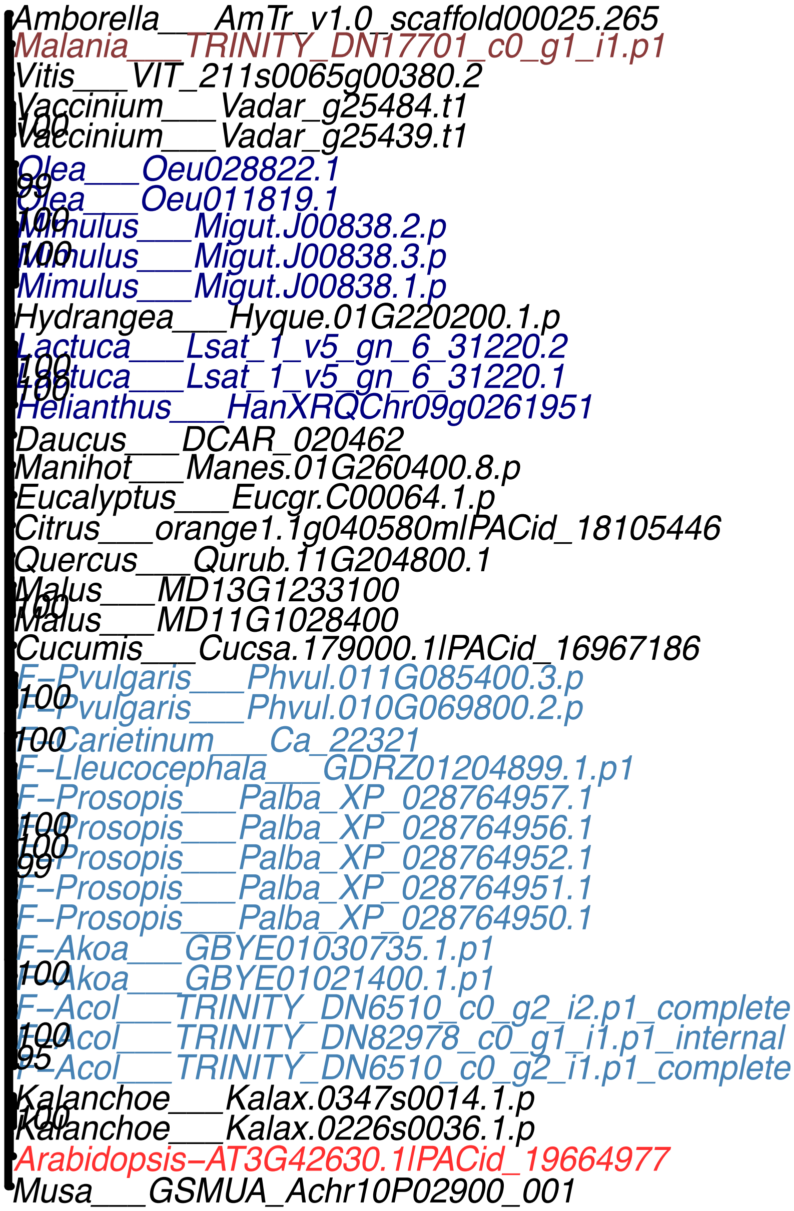
Amborella___AmTr_v1.0_scaffold00025.265
Malania___TRINITY_DN17701_c0_g1_i1.p1
Vitis___VIT_211s0065g00380.2
Vaccinium___Vadar_g25484.t1
Vaccinium___Vadar_g25439.t1
Olea___Oeu028822.1
Olea___Oeu011819.1
Mimulus___Migut.J00838.2.p
Mimulus___Migut.J00838.3.p
Mimulus___Migut.J00838.1.p
Hydrangea___Hyque.01G220200.1.p
Lactuca___Lsat_1_v5_gn_6_31220.2
Lactuca___Lsat_1_v5_gn_6_31220.1
Helianthus___HanXRQChr09g0261951
Daucus___DCAR_020462
Manihot___Manes.01G260400.8.p
Eucalyptus___Eucgr.C00064.1.p
Citrus___orange1.1g040580m|PACid_18105446
Quercus___Qurub.11G204800.1
Malus___MD13G1233100
Malus___MD11G1028400
Cucumis___Cucsa.179000.1|PACid_16967186
F–Pvulgaris___Phvul.011G085400.3.p
F–Pvulgaris___Phvul.010G069800.2.p
F–Carietinum___Ca_22321
F–Lleucocephala___GDRZ01204899.1.p1
F–Prosopis___Palba_XP_028764957.1
F–Prosopis___Palba_XP_028764956.1
F–Prosopis___Palba_XP_028764952.1
F–Prosopis___Palba_XP_028764951.1
F–Prosopis___Palba_XP_028764950.1
F–Akoa___GBYE01030735.1.p1
F–Akoa___GBYE01021400.1.p1
F–Acol___TRINITY_DN6510_c0_g2_i2.p1_complete
F–Acol___TRINITY_DN82978_c0_g1_i1.p1_internal
F–Acol___TRINITY_DN6510_c0_g2_i1.p1_complete
Kalanchoe___Kalax.0347s0014.1.p
Kalanchoe___Kalax.0226s0036.1.p
Arabidopsis–AT3G42630.1|PACid_19664977
Musa___GSMUA_Achr10P02900_001

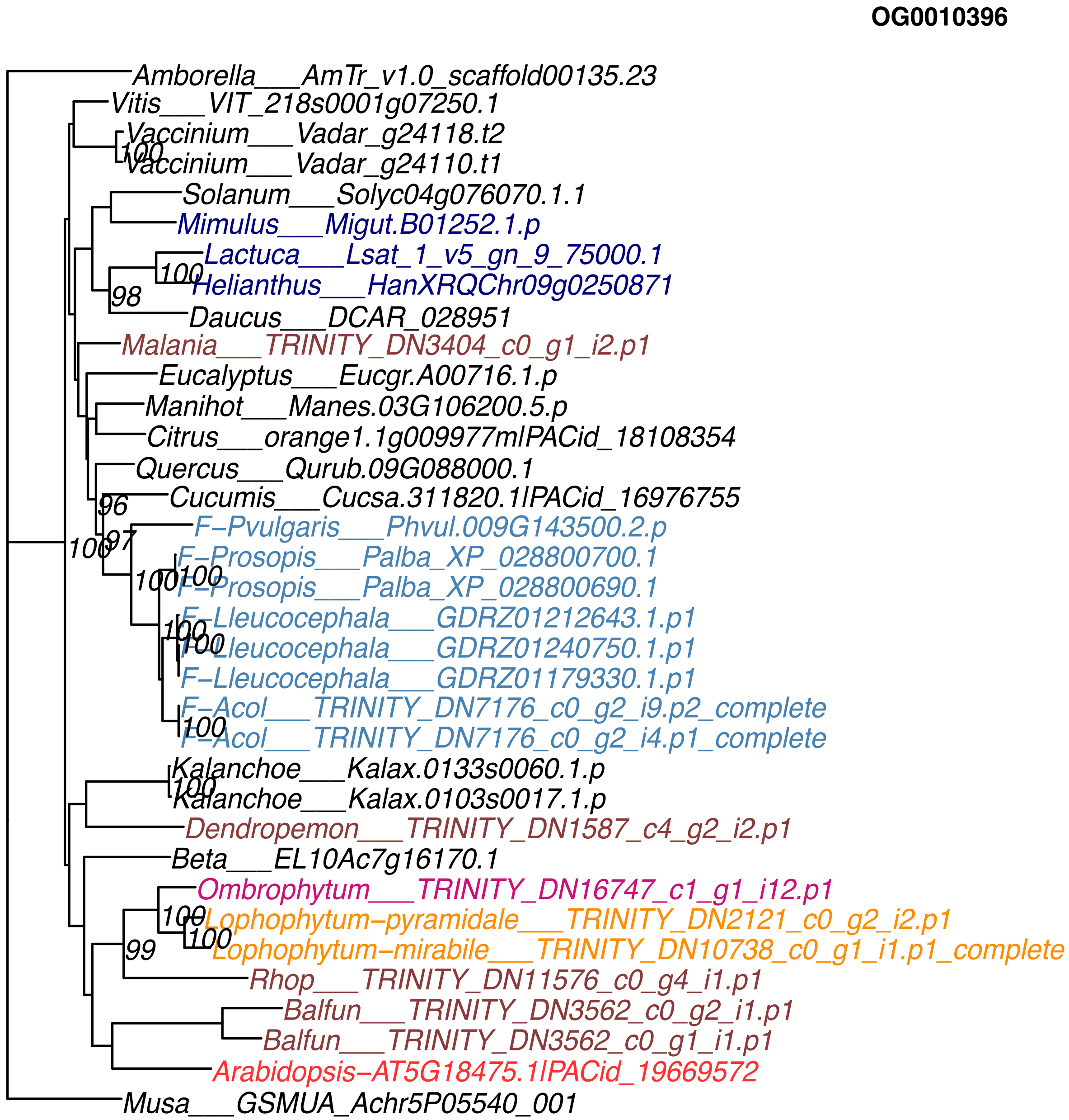
OG0010396
Amborella___AmTr_v1.0_scaffold00135.23
Vitis___VIT_218s0001g07250.1
Vaccinium___Vadar_g24118.t2
100
Vaccinium___Vadar_g24110.t1
Solanum___Solyc04g076070.1.1
Mimulus___Migut.B01252.1.p
Lactuca___Lsat_1_v5_gn_9_75000.1
100
Helianthus___HanXRQChr09g0250871
98
Daucus___DCAR_028951
Malania___TRINITY_DN3404_c0_g1_i2.p1
Eucalyptus___Eucgr.A00716.1.p
Manihot___Manes.03G106200.5.p
Citrus___orange1.1g009977mIPACid_18108354
Quercus___Qurub.09G088000.1
Cucumis___Cucsa.311820.1IPACid_16976755
96
F−Pvulgaris___Phvul.009G143500.2.p
100
97
F−Prosopis___Palba_XP_028800700.1
100
100
F−Prosopis___Palba_XP_028800690.1
F−Lleucocephala___GDRZ01212643.1.p1
100
100
F−Lleucocephala___GDRZ01240750.1.p1
F−Lleucocephala___GDRZ01179330.1.p1
F−Acol___TRINITY_DN7176_c0_g2_i9.p2_complete
100
F−Acol___TRINITY_DN7176_c0_g2_i4.p1_complete
Kalanchoe___Kalax.0133s0060.1.p
100
Kalanchoe___Kalax.0103s0017.1.p
Dendropemon___TRINITY_DN1587_c4_g2_i2.p1
Beta___EL10Ac7g16170.1
Ombrophytum___TRINITY_DN16747_c1_g1_i12.p1
100
Lophophytum−pyramidale___TRINITY_DN2121_c0_g2_i2.p1
100
99
Lophophytum−mirabile___TRINITY_DN10738_c0_g1_i1.p1_complete
Rhop___TRINITY_DN11576_c0_g4_i1.p1
Balfun___TRINITY_DN3562_c0_g2_i1.p1
Balfun___TRINITY_DN3562_c0_g1_i1.p1
Arabidopsis−AT5G18475.1IPACid_19669572
Musa___GSMUA_Achr5P05540_001

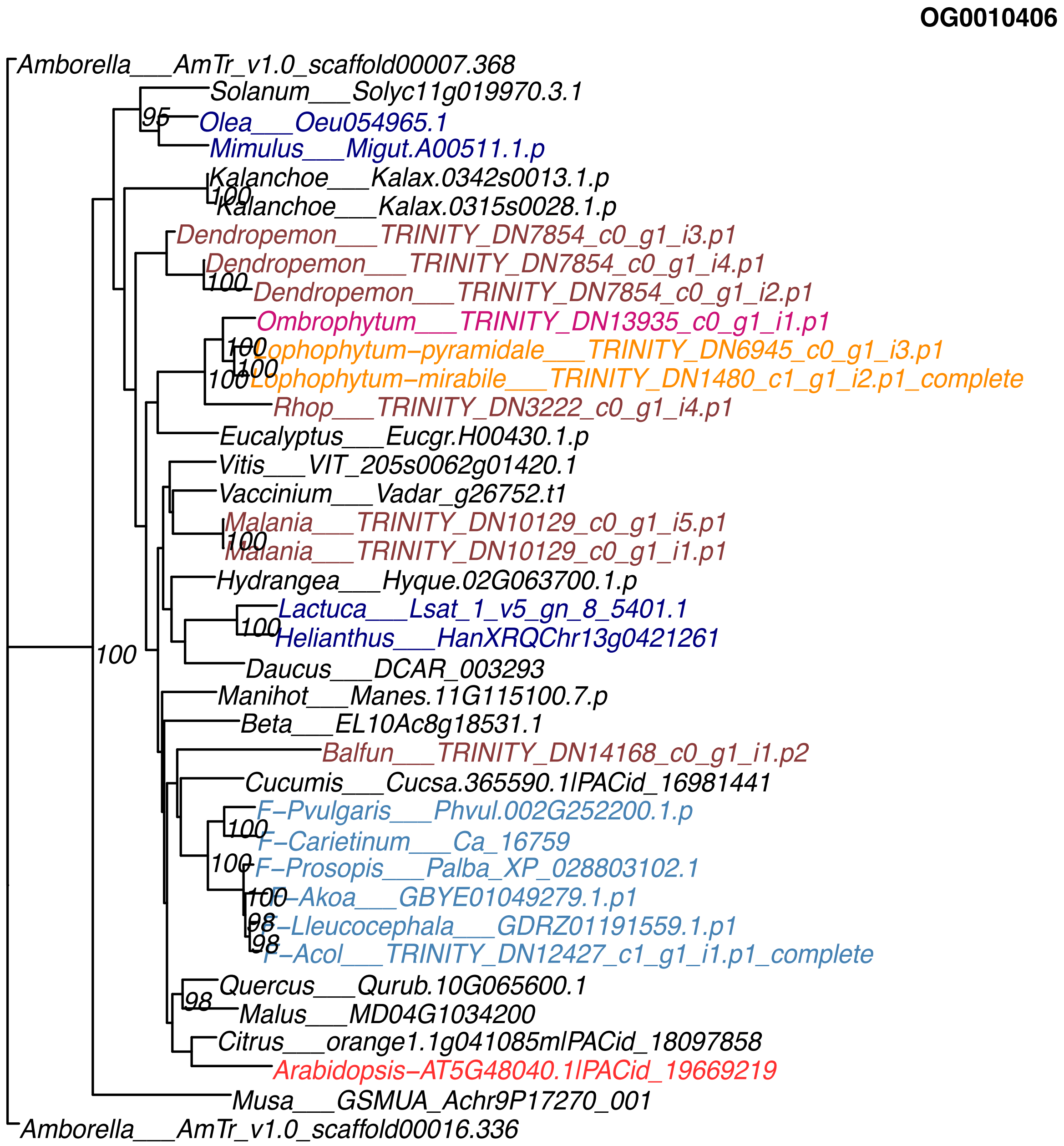
OG0010406
Amborella___AmTr_v1.0_scaffold00007.368
Solanum___Solyc11g019970.3.1
Olea___Oeu054965.1
Mimulus___Migut.A00511.1.p
Kalanchoe___Kalax.0342s0013.1.p
Kalanchoe___Kalax.0315s0028.1.p
Dendropemon___TRINITY_DN7854_c0_g1_i3.p1
Dendropemon___TRINITY_DN7854_c0_g1_i4.p1
Dendropemon___TRINITY_DN7854_c0_g1_i2.p1
Ombrophytum___TRINITY_DN13935_c0_g1_i1.p1
Lophophytum−pyramidale___TRINITY_DN6945_c0_g1_i3.p1
Lophophytum−mirabile___TRINITY_DN1480_c1_g1_i2.p1_complete
Rhop___TRINITY_DN3222_c0_g1_i4.p1
Eucalyptus___Eucgr.H00430.1.p
Vitis___VIT_205s0062g01420.1
Vaccinium___Vadar_g26752.t1
Malania___TRINITY_DN10129_c0_g1_i5.p1
Malania___TRINITY_DN10129_c0_g1_i1.p1
Hydrangea___Hyque.02G063700.1.p
Lactuca___Lsat_1_v5_gn_8_5401.1
Helianthus___HanXRQChr13g0421261
Daucus___DCAR_003293
Manihot___Manes.11G115100.7.p
Beta___EL10Ac8g18531.1
Balfun___TRINITY_DN14168_c0_g1_i1.p2
Cucumis___Cucsa.365590.1|PACid_16981441
F−Pvulgaris___Phvul.002G252200.1.p
F−Carietinum___Ca_16759
F−Prosopis___Palba_XP_028803102.1
F−Akoa___GBYE01049279.1.p1
F−Lleucocephala___GDRZ01191559.1.p1
F−Acol___TRINITY_DN12427_c1_g1_i1.p1_complete
Quercus___Qurub.10G065600.1
Malus___MD04G1034200
Citrus___orange1.1g041085m|PACid_18097858
Arabidopsis−AT5G48040.1|PACid_19669219
Musa___GSMUA_Achr9P17270_001
Amborella___AmTr_v1.0_scaffold00016.336
95
100
100
100
100
100
100
100
100
100
100
100
98
98
98

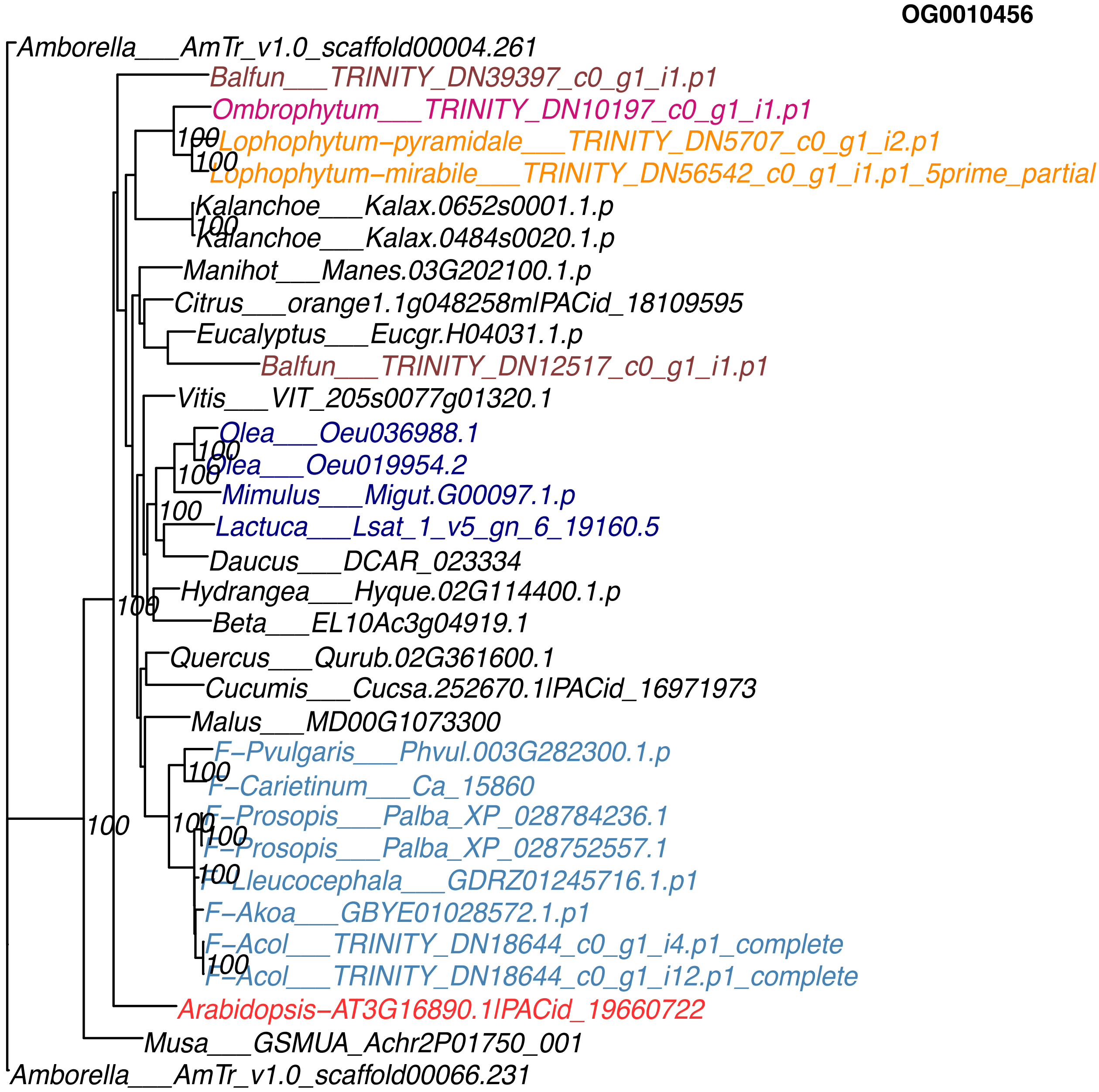

OG0010456
Amborella___AmTr_v1.0_scaffold00004.261
Balfun___TRINITY_DN39397_c0_g1_i1.p1
Ombrophytum___TRINITY_DN10197_c0_g1_i1.p1
100
Lophophytum−pyramidale___TRINITY_DN5707_c0_g1_i2.p1
100
Lophophytum−mirabile___TRINITY_DN56542_c0_g1_i1.p1_5prime_partial
Kalanchoe___Kalax.0652s0001.1.p
100
Kalanchoe___Kalax.0484s0020.1.p
Manihot___Manes.03G202100.1.p
Citrus___orange1.1g048258m|PACid_18109595
Eucalyptus___Eucgr.H04031.1.p
Balfun___TRINITY_DN12517_c0_g1_i1.p1
Vitis___VIT_205s0077g01320.1
Olea___Oeu036988.1
100
Olea___Oeu019954.2
100
Mimulus___Migut.G00097.1.p
100
Lactuca___Lsat_1_v5_gn_6_19160.5
Daucus___DCAR_023334
Hydrangea___Hyque.02G114400.1.p
100
Beta___EL10Ac3g04919.1
Quercus___Qurub.02G361600.1
Cucumis___Cucsa.252670.1|PACid_16971973
Malus___MD00G1073300
F−Pvulgaris___Phvul.003G282300.1.p
100
F−Carietinum___Ca_15860
F−Prosopis___Palba_XP_028784236.1
100
100
F−Prosopis___Palba_XP_028752557.1
100
F−Lleucocephala___GDRZ01245716.1.p1
F−Akoa___GBYE01028572.1.p1
F−Acol___TRINITY_DN18644_c0_g1_i4.p1_complete
100
F−Acol___TRINITY_DN18644_c0_g1_i12.p1_complete
Arabidopsis−AT3G16890.1|PACid_19660722
100
Musa___GSMUA_Achr2P01750_001
Amborella___AmTr_v1.0_scaffold00066.231

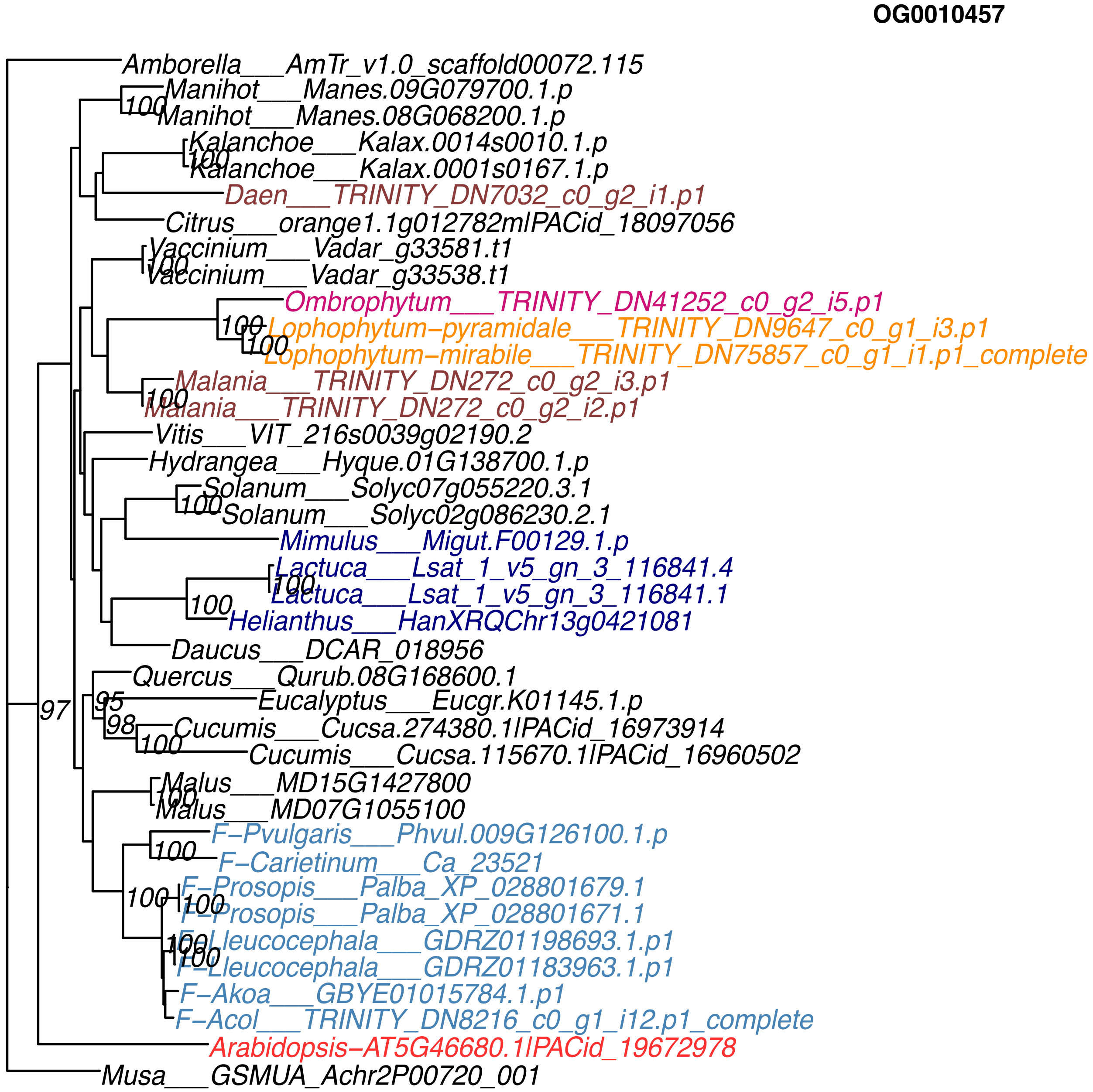

OG0010457
Amborella___AmTr_v1.0_scaffold00072.115
Manihot___Manes.09G079700.1.p
100
Manihot___Manes.08G068200.1.p
Kalanchoe___Kalax.0014s0010.1.p
100
Kalanchoe___Kalax.0001s0167.1.p
Daen___TRINITY_DN7032_c0_g2_i1.p1
Citrus___orange1.1g012782mIPACid_18097056
Vaccinium___Vadar_g33581.t1
100
Vaccinium___Vadar_g33538.t1
Ombrophytum___TRINITY_DN41252_c0_g2_i5.p1
100
Lophophytum–pyramidale___TRINITY_DN9647_c0_g1_i3.p1
100
Lophophytum–mirabile___TRINITY_DN75857_c0_g1_i1.p1_complete
Malania___TRINITY_DN272_c0_g2_i3.p1
100
Malania___TRINITY_DN272_c0_g2_i2.p1
Vitis___VIT_216s0039g02190.2
Hydrangea___Hyque.01G138700.1.p
Solanum___Solyc07g055220.3.1
100
Solanum___Solyc02g086230.2.1
Mimulus___Migut.F00129.1.p
Lactuca___Lsat_1_v5_gn_3_116841.4
100
Lactuca___Lsat_1_v5_gn_3_116841.1
100
Helianthus___HanXRQChr13g0421081
Daucus___DCAR_018956
Quercus___Qurub.08G168600.1
95
Eucalyptus___Eucgr.K01145.1.p
98
Cucumis___Cucsa.274380.1IPACid_16973914
100
Cucumis___Cucsa.115670.1IPACid_16960502
97
Malus___MD15G1427800
100
Malus___MD07G1055100
F–Pvulgaris___Phvul.009G126100.1.p
100
F–Carietinum___Ca_23521
100
F–Prosopis___Palba_XP_028801679.1
100
F–Prosopis___Palba_XP_028801671.1
100
F–Lleucocephala___GDRZ01198693.1.p1
100
F–Lleucocephala___GDRZ01183963.1.p1
F–Akoa___GBYE01015784.1.p1
F–Acol___TRINITY_DN8216_c0_g1_i12.p1_complete
Arabidopsis–AT5G46680.1IPACid_19672978
Musa___GSMUA_Achr2P00720_001

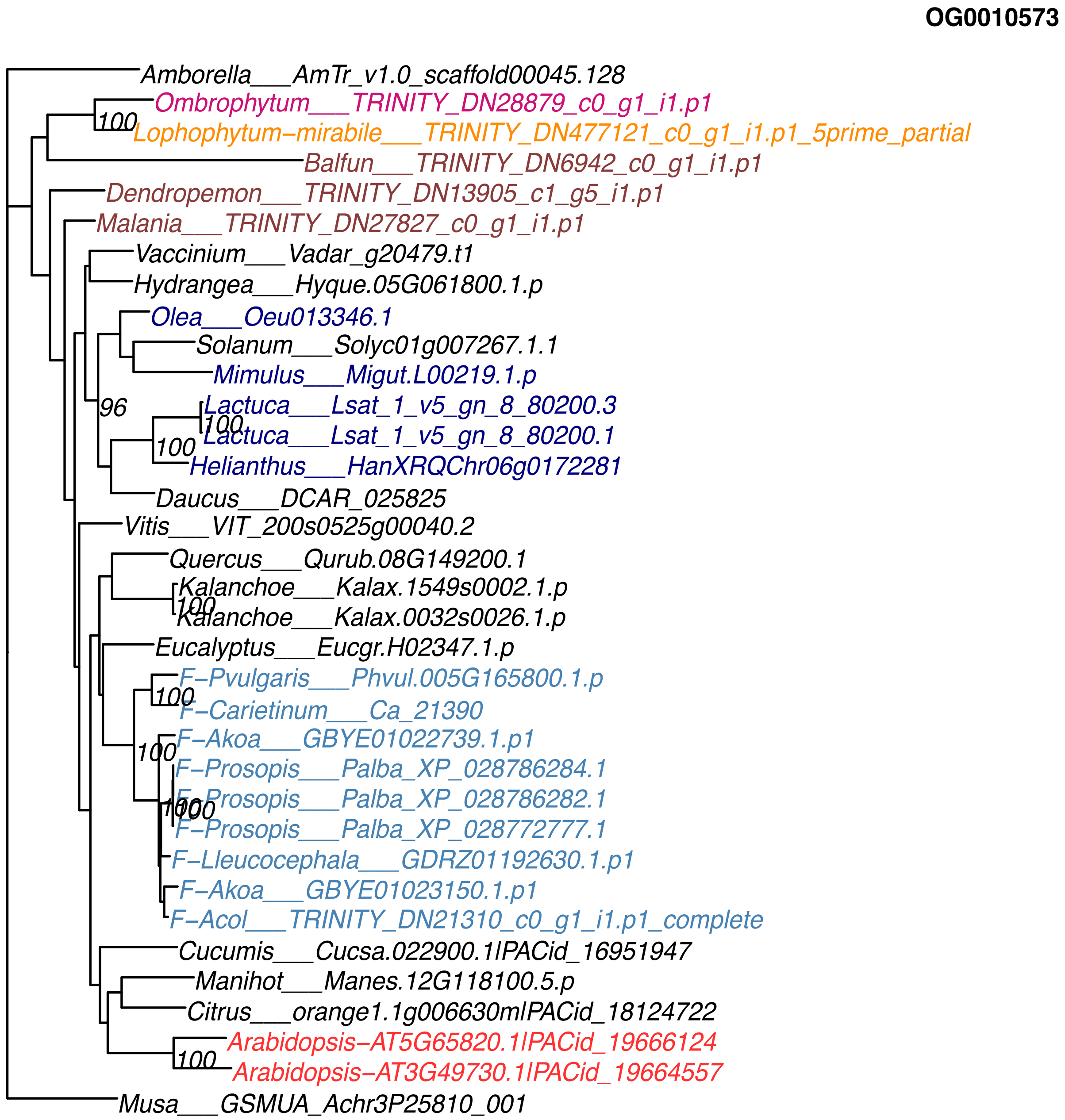
OG0010573
Amborella___AmTr_v1.0_scaffold00045.128
Ombrophytum___TRINITY_DN28879_c0_g1_i1.p1
100
Lophophytum−mirabile___TRINITY_DN477121_c0_g1_i1.p1_5prime_partial
Balfun___TRINITY_DN6942_c0_g1_i1.p1
Dendropemon___TRINITY_DN13905_c1_g5_i1.p1
Malania___TRINITY_DN27827_c0_g1_i1.p1
Vaccinium___Vadar_g20479.t1
Hydrangea___Hyque.05G061800.1.p
Olea___Oeu013346.1
Solanum___Solyc01g007267.1.1
Mimulus___Migut.L00219.1.p
96
Lactuca___Lsat_1_v5_gn_8_80200.3
100
Lactuca___Lsat_1_v5_gn_8_80200.1
100
Helianthus___HanXRQChr06g0172281
Daucus___DCAR_025825
Vitis___VIT_200s0525g00040.2
Quercus___Qurub.08G149200.1
Kalanchoe___Kalax.1549s0002.1.p
100
Kalanchoe___Kalax.0032s0026.1.p
Eucalyptus___Eucgr.H02347.1.p
F−Pvulgaris___Phvul.005G165800.1.p
100
F−Carietinum___Ca_21390
F−Akoa___GBYE01022739.1.p1
100
F−Prosopis___Palba_XP_028786284.1
F−Prosopis___Palba_XP_028786282.1
100
F−Prosopis___Palba_XP_028772777.1
F−Lleucocephala___GDRZ01192630.1.p1
F−Akoa___GBYE01023150.1.p1
F−Acol___TRINITY_DN21310_c0_g1_i1.p1_complete
Cucumis___Cucsa.022900.1|PACid_16951947
Manihot___Manes.12G118100.5.p
Citrus___orange1.1g006630m|PACid_18124722
Arabidopsis−AT5G65820.1|PACid_19666124
100
Arabidopsis−AT3G49730.1|PACid_19664557
Musa___GSMUA_Achr3P25810_001

**OG0010616**

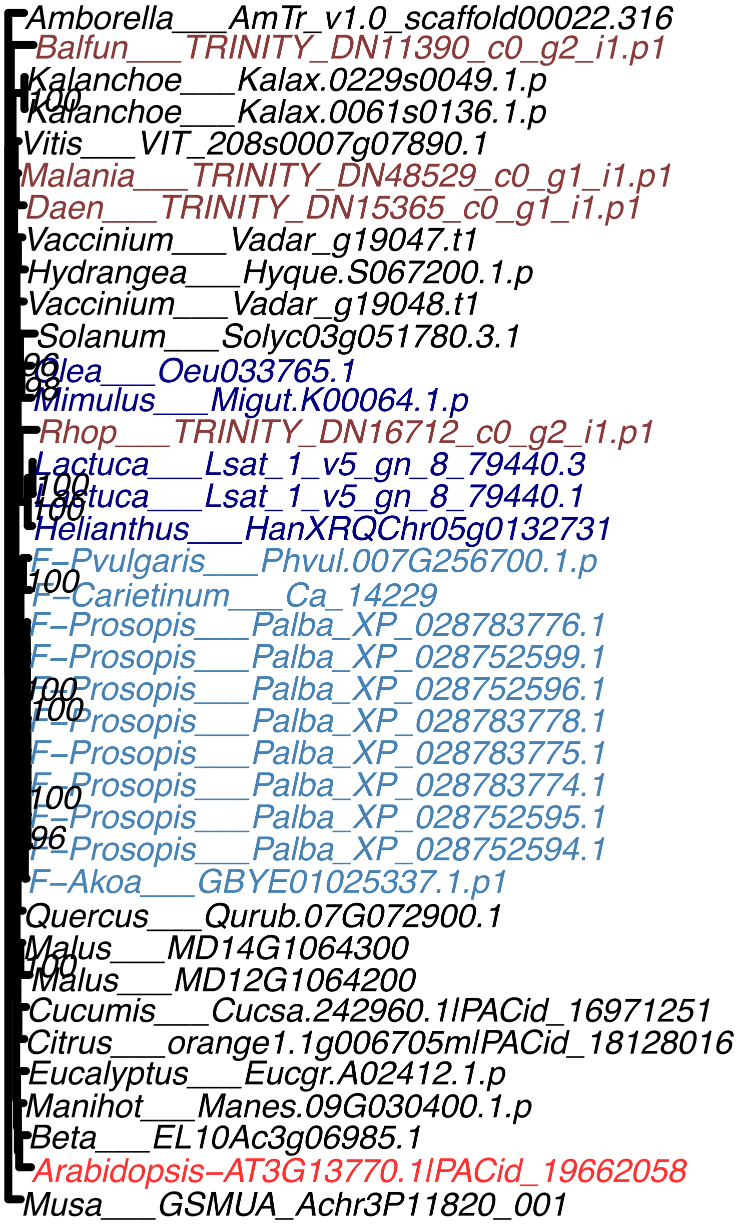
Amborella___AmTr_v1.0_scaffold00022.316
Balfun___TRINITY_DN11390_c0_g2_i1.p1
Kalanchoe___Kalax.0229s0049.1.p
100
Kalanchoe___Kalax.0061s0136.1.p
Vitis___VIT_208s0007g07890.1
Malania___TRINITY_DN48529_c0_g1_i1.p1
Daen___TRINITY_DN15365_c0_g1_i1.p1
Vaccinium___Vadar_g19047.t1
Hydrangea___Hyque.S067200.1.p
Vaccinium___Vadar_g19048.t1
Solanum___Solyc03g051780.3.1
96
Olea___Oeu033765.1
98
Mimulus___Migut.K00064.1.p
Rhop___TRINITY_DN16712_c0_g2_i1.p1
Lactuca___Lsat_1_v5_gn_8_79440.3
100
Lactuca___Lsat_1_v5_gn_8_79440.1
100
Helianthus___HanXRQChr05g0132731
F–Pvulgaris___Phvul.007G256700.1.p
100
F–Carietinum___Ca_14229
F–Prosopis___Palba_XP_028783776.1
F–Prosopis___Palba_XP_028752599.1
100
F–Prosopis___Palba_XP_028752596.1
100
F–Prosopis___Palba_XP_028783778.1
F–Prosopis___Palba_XP_028783775.1
F–Prosopis___Palba_XP_028783774.1
100
F–Prosopis___Palba_XP_028752595.1
96
F–Prosopis___Palba_XP_028752594.1
F–Akoa___GBYE01025337.1.p1
Quercus___Qurub.07G072900.1
Malus___MD14G1064300
100
Malus___MD12G1064200
Cucumis___Cucsa.242960.1lPACid_16971251
Citrus___orange1.1g006705mlPACid_18128016
Eucalyptus___Eucgr.A02412.1.p
Manihot___Manes.09G030400.1.p
Beta___EL10Ac3g06985.1
Arabidopsis–AT3G13770.1lPACid_19662058
Musa___GSMUA_Achr3P11820_001

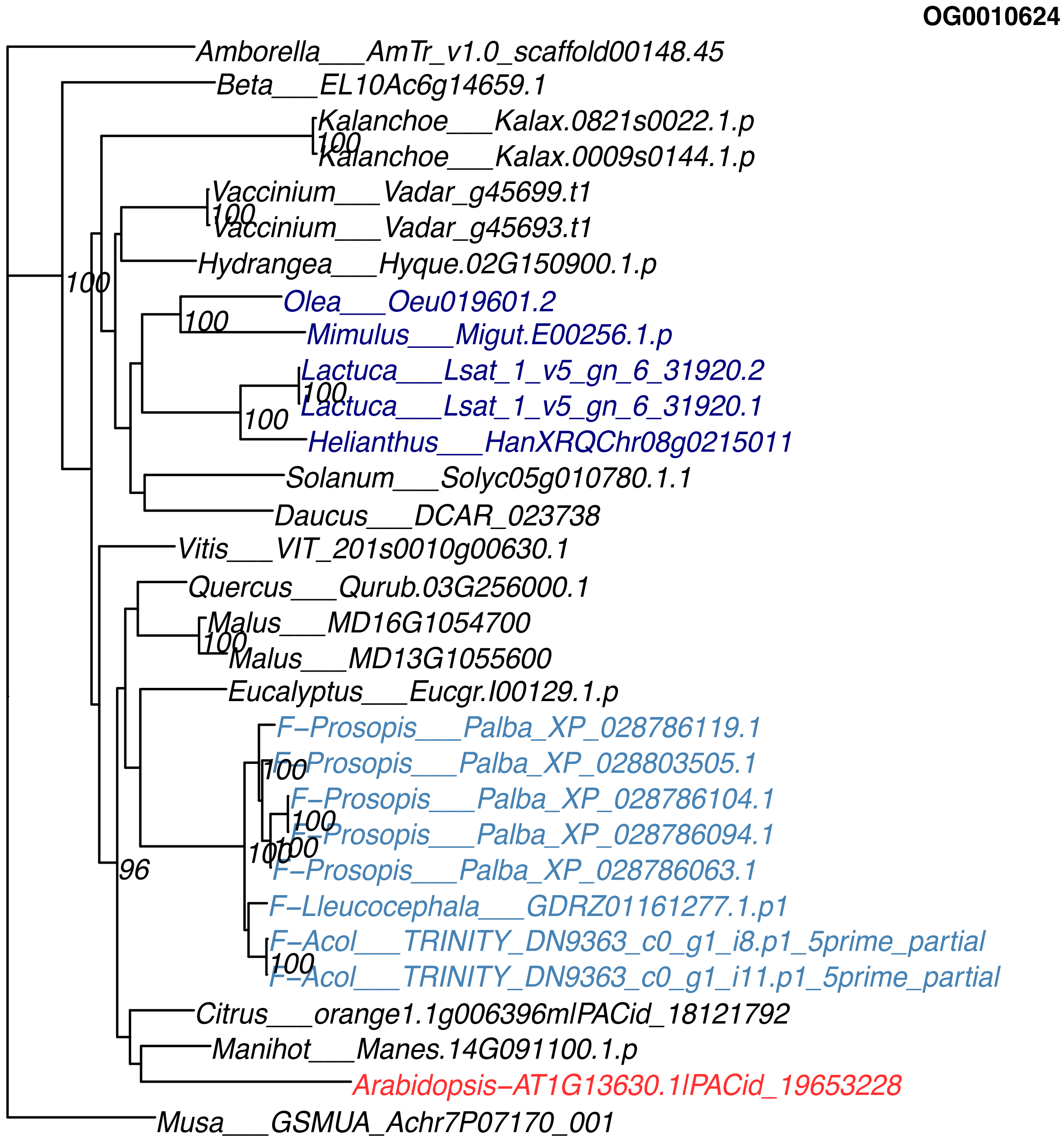
OG0010624
Amborella___AmTr_v1.0_scaffold00148.45
Beta___EL10Ac6g14659.1
Kalanchoe___Kalax.0821s0022.1.p
100
Kalanchoe___Kalax.0009s0144.1.p
Vaccinium___Vadar_g45699.t1
100
Vaccinium___Vadar_g45693.t1
Hydrangea___Hyque.02G150900.1.p
100
Olea___Oeu019601.2
100
Mimulus___Migut.E00256.1.p
Lactuca___Lsat_1_v5_gn_6_31920.2
100
Lactuca___Lsat_1_v5_gn_6_31920.1
100
Helianthus___HanXRQChr08g0215011
Solanum___Solyc05g010780.1.1
Daucus___DCAR_023738
Vitis___VIT_201s0010g00630.1
Quercus___Qurub.03G256000.1
Malus___MD16G1054700
100
Malus___MD13G1055600
Eucalyptus___Eucgr.I00129.1.p
F−Prosopis___Palba_XP_028786119.1
F−Prosopis___Palba_XP_028803505.1
100
F−Prosopis___Palba_XP_028786104.1
100
F−Prosopis___Palba_XP_028786094.1
100
100
F−Prosopis___Palba_XP_028786063.1
96
F−Lleucocephala___GDRZ01161277.1.p1
F−Acol___TRINITY_DN9363_c0_g1_i8.p1_5prime_partial
100
F−Acol___TRINITY_DN9363_c0_g1_i11.p1_5prime_partial
Citrus___orange1.1g006396mIPACid_18121792
Manihot___Manes.14G091100.1.p
Arabidopsis−AT1G13630.1IPACid_19653228
Musa___GSMUA_Achr7P07170_001

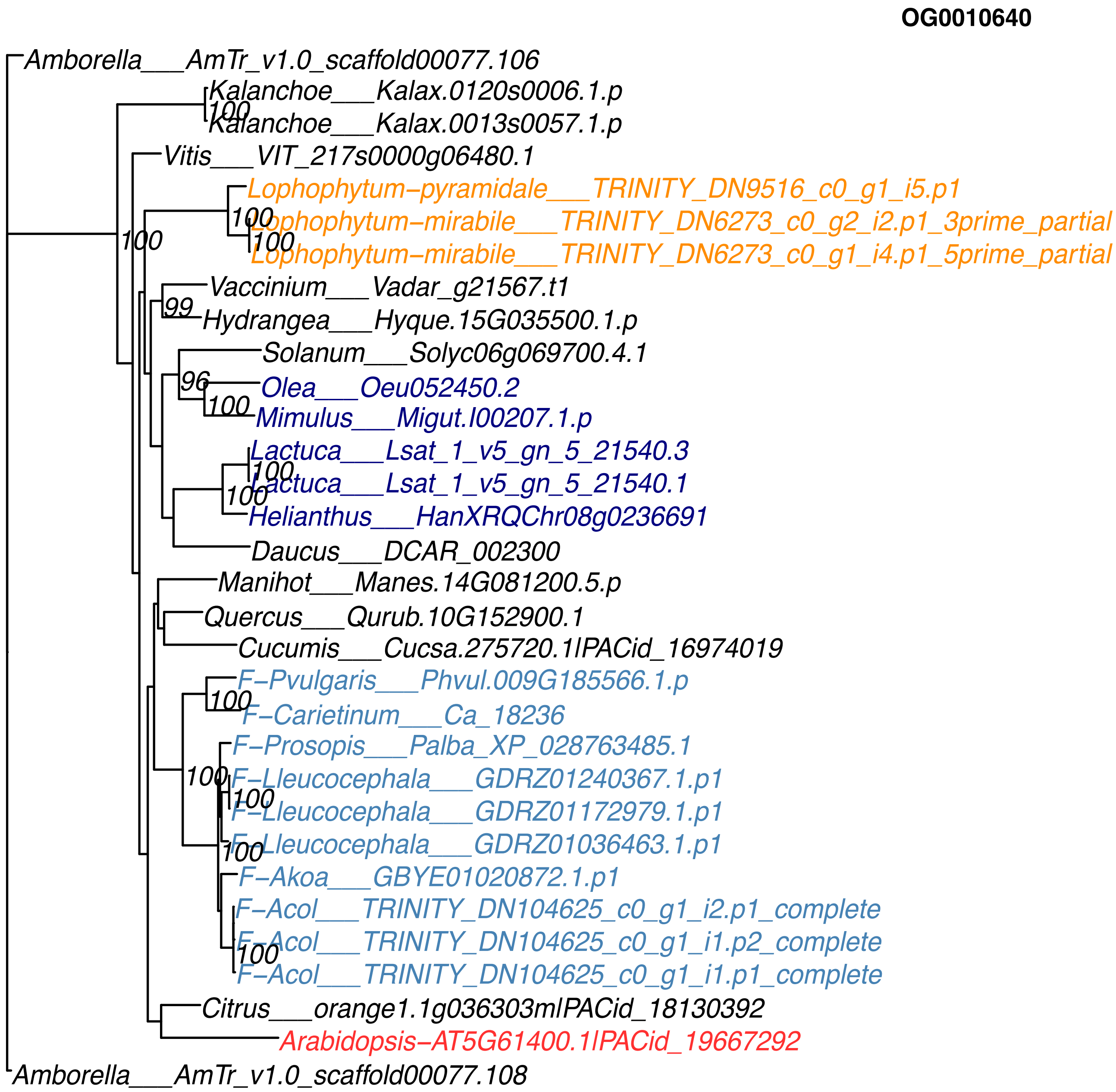

OG0010640
Amborella___AmTr_v1.0_scaffold00077.106
Kalanchoe___Kalax.0120s0006.1.p
100
Kalanchoe___Kalax.0013s0057.1.p
Vitis___VIT_217s0000g06480.1
Lophophytum−pyramidale___TRINITY_DN9516_c0_g1_i5.p1
100
Lophophytum−mirabile___TRINITY_DN6273_c0_g2_i2.p1_3prime_partial
100
100
Lophophytum−mirabile___TRINITY_DN6273_c0_g1_i4.p1_5prime_partial
Vaccinium___Vadar_g21567.t1
99
Hydrangea___Hyque.15G035500.1.p
Solanum___Solyc06g069700.4.1
96
Olea___Oeu052450.2
100
Mimulus___Migut.I00207.1.p
Lactuca___Lsat_1_v5_gn_5_21540.3
100
Lactuca___Lsat_1_v5_gn_5_21540.1
100
Helianthus___HanXRQChr08g0236691
Daucus___DCAR_002300
Manihot___Manes.14G081200.5.p
Quercus___Qurub.10G152900.1
Cucumis___Cucsa.275720.1lPACid_16974019
F−Pvulgaris___Phvul.009G185566.1.p
100
F−Carietinum___Ca_18236
F−Prosopis___Palba_XP_028763485.1
100
F−Lleucocephala___GDRZ01240367.1.p1
100
F−Lleucocephala___GDRZ01172979.1.p1
F−Lleucocephala___GDRZ01036463.1.p1
100
F−Akoa___GBYE01020872.1.p1
F−Acol___TRINITY_DN104625_c0_g1_i2.p1_complete
F−Acol___TRINITY_DN104625_c0_g1_i1.p2_complete
100
F−Acol___TRINITY_DN104625_c0_g1_i1.p1_complete
Citrus___orange1.1g036303mlPACid_18130392
Arabidopsis−AT5G61400.1lPACid_19667292
Amborella___AmTr_v1.0_scaffold00077.108

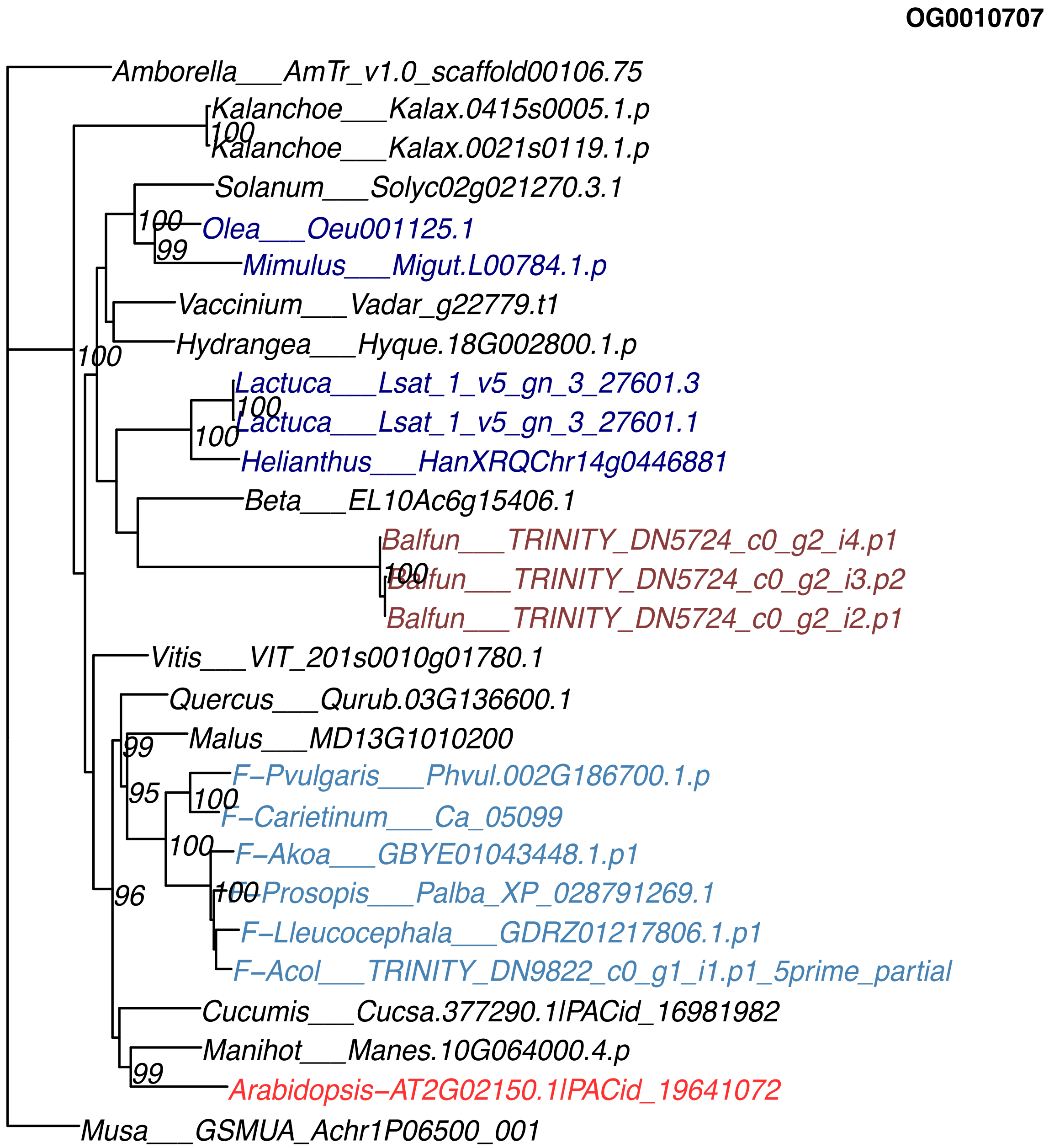
OG0010707
Amborella___AmTr_v1.0_scaffold00106.75
Kalanchoe___Kalax.0415s0005.1.p
100
Kalanchoe___Kalax.0021s0119.1.p
Solanum___Solyc02g021270.3.1
100
Olea___Oeu001125.1
99
Mimulus___Migut.L00784.1.p
Vaccinium___Vadar_g22779.t1
Hydrangea___Hyque.18G002800.1.p
100
Lactuca___Lsat_1_v5_gn_3_27601.3
100
Lactuca___Lsat_1_v5_gn_3_27601.1
100
Helianthus___HanXRQChr14g0446881
Beta___EL10Ac6g15406.1
Balfun___TRINITY_DN5724_c0_g2_i4.p1
100
Balfun___TRINITY_DN5724_c0_g2_i3.p2
Balfun___TRINITY_DN5724_c0_g2_i2.p1
Vitis___VIT_201s0010g01780.1
Quercus___Qurub.03G136600.1
Malus___MD13G1010200
99
F−Pvulgaris___Phvul.002G186700.1.p
95
100
F−Carietinum___Ca_05099
100
F−Akoa___GBYE01043448.1.p1
100
F−Prosopis___Palba_XP_028791269.1
96
F−Lleucocephala___GDRZ01217806.1.p1
F−Acol___TRINITY_DN9822_c0_g1_i1.p1_5prime_partial
Cucumis___Cucsa.377290.1|PACid_16981982
Manihot___Manes.10G064000.4.p
99
Arabidopsis−AT2G02150.1|PACid_19641072
Musa___GSMUA_Achr1P06500_001

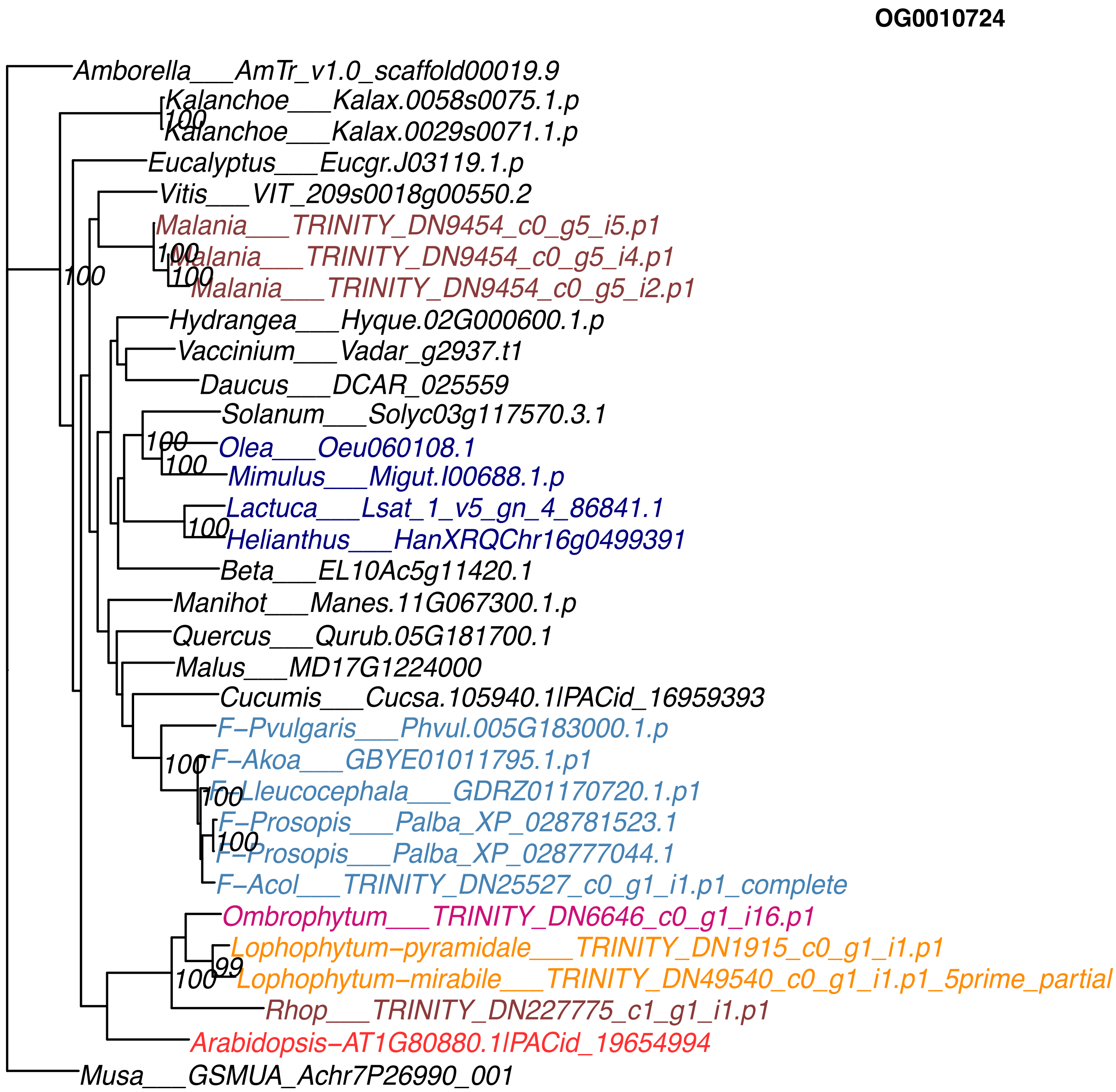
OG0010724
Amborella___AmTr_v1.0_scaffold00019.9
Kalanchoe___Kalax.0058s0075.1.p
100
Kalanchoe___Kalax.0029s0071.1.p
Eucalyptus___Eucgr.J03119.1.p
Vitis___VIT_209s0018g00550.2
Malania___TRINITY_DN9454_c0_g5_i5.p1
100
Malania___TRINITY_DN9454_c0_g5_i4.p1
100
100
Malania___TRINITY_DN9454_c0_g5_i2.p1
Hydrangea___Hyque.02G000600.1.p
Vaccinium___Vadar_g2937.t1
Daucus___DCAR_025559
Solanum___Solyc03g117570.3.1
100
Olea___Oeu060108.1
100
Mimulus___Migut.I00688.1.p
Lactuca___Lsat_1_v5_gn_4_86841.1
100
Helianthus___HanXRQChr16g0499391
Beta___EL10Ac5g11420.1
Manihot___Manes.11G067300.1.p
Quercus___Qurub.05G181700.1
Malus___MD17G1224000
Cucumis___Cucsa.105940.1IPACid_16959393
F−Pvulgaris___Phvul.005G183000.1.p
100
F−Akoa___GBYE01011795.1.p1
F−Lleucocephala___GDRZ01170720.1.p1
100
F−Prosopis___Palba_XP_028781523.1
100
F−Prosopis___Palba_XP_028777044.1
F−Acol___TRINITY_DN25527_c0_g1_i1.p1_complete
Ombrophytum___TRINITY_DN6646_c0_g1_i16.p1
Lophophytum−pyramidale___TRINITY_DN1915_c0_g1_i1.p1
99
100
Lophophytum−mirabile___TRINITY_DN49540_c0_g1_i1.p1_5prime_partial
Rhop___TRINITY_DN227775_c1_g1_i1.p1
Arabidopsis−AT1G80880.1IPACid_19654994
Musa___GSMUA_Achr7P26990_001

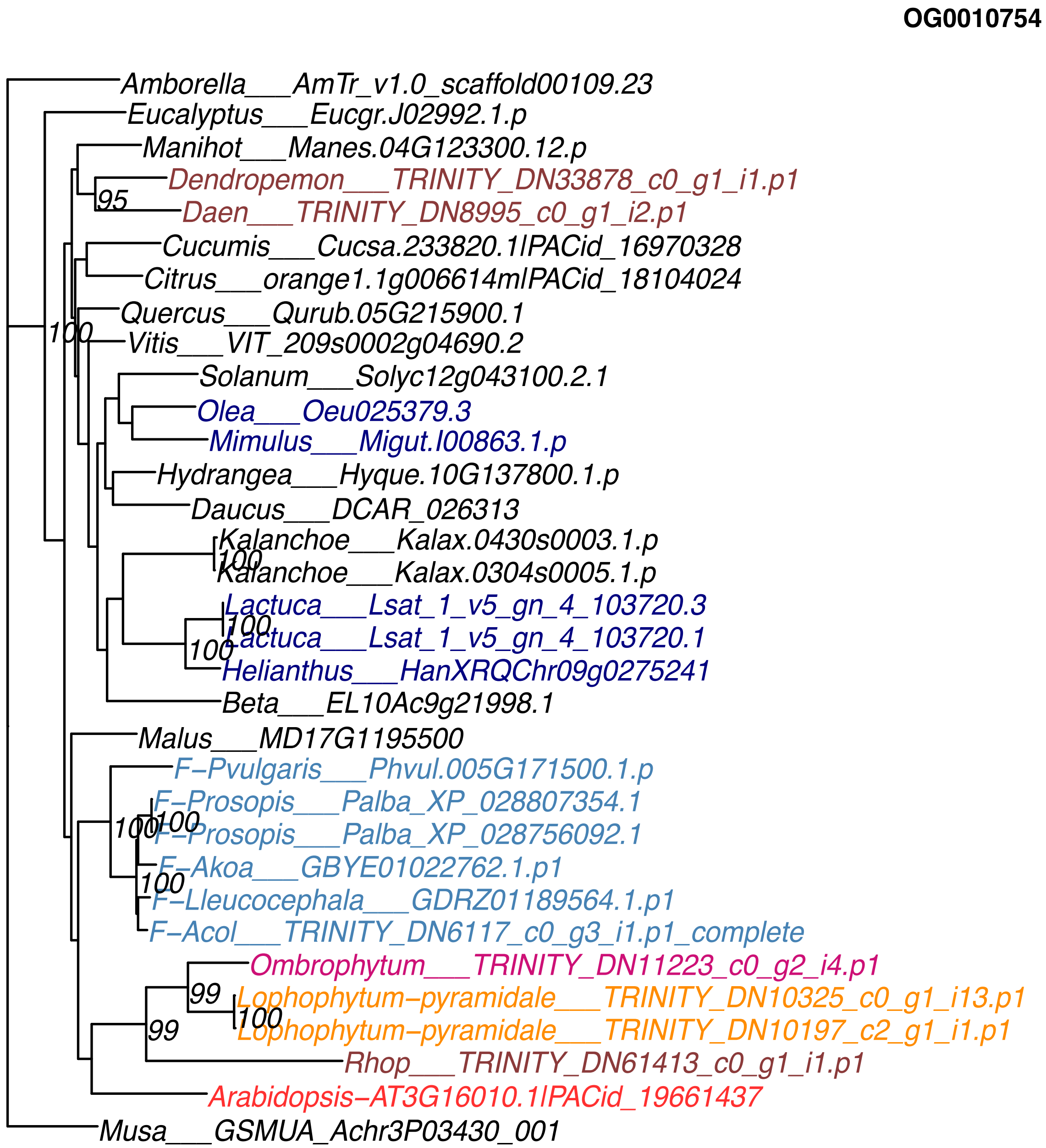

OG0010754
Amborella___AmTr_v1.0_scaffold00109.23
Eucalyptus___Eucgr.J02992.1.p
Manihot___Manes.04G123300.12.p
Dendropemon___TRINITY_DN33878_c0_g1_i1.p1
95
Daen___TRINITY_DN8995_c0_g1_i2.p1
Cucumis___Cucsa.233820.1lPACid_16970328
Citrus___orange1.1g006614mlPACid_18104024
Quercus___Qurub.05G215900.1
100
Vitis___VIT_209s0002g04690.2
Solanum___Solyc12g043100.2.1
Olea___Oeu025379.3
Mimulus___Migut.I00863.1.p
Hydrangea___Hyque.10G137800.1.p
Daucus___DCAR_026313
Kalanchoe___Kalax.0430s0003.1.p
100
Kalanchoe___Kalax.0304s0005.1.p
Lactuca___Lsat_1_v5_gn_4_103720.3
100
Lactuca___Lsat_1_v5_gn_4_103720.1
100
Helianthus___HanXRQChr09g0275241
Beta___EL10Ac9g21998.1
Malus___MD17G1195500
F−Pvulgaris___Phvul.005G171500.1.p
F−Prosopis___Palba_XP_028807354.1
100
100
F−Prosopis___Palba_XP_028756092.1
F−Akoa___GBYE01022762.1.p1
100
F−Lleucocephala___GDRZ01189564.1.p1
F−Acol___TRINITY_DN6117_c0_g3_i1.p1_complete
Ombrophytum___TRINITY_DN11223_c0_g2_i4.p1
99
Lophophytum−pyramidale___TRINITY_DN10325_c0_g1_i13.p1
100
99
Lophophytum−pyramidale___TRINITY_DN10197_c2_g1_i1.p1
Rhop___TRINITY_DN61413_c0_g1_i1.p1
Arabidopsis−AT3G16010.1lPACid_19661437
Musa___GSMUA_Achr3P03430_001

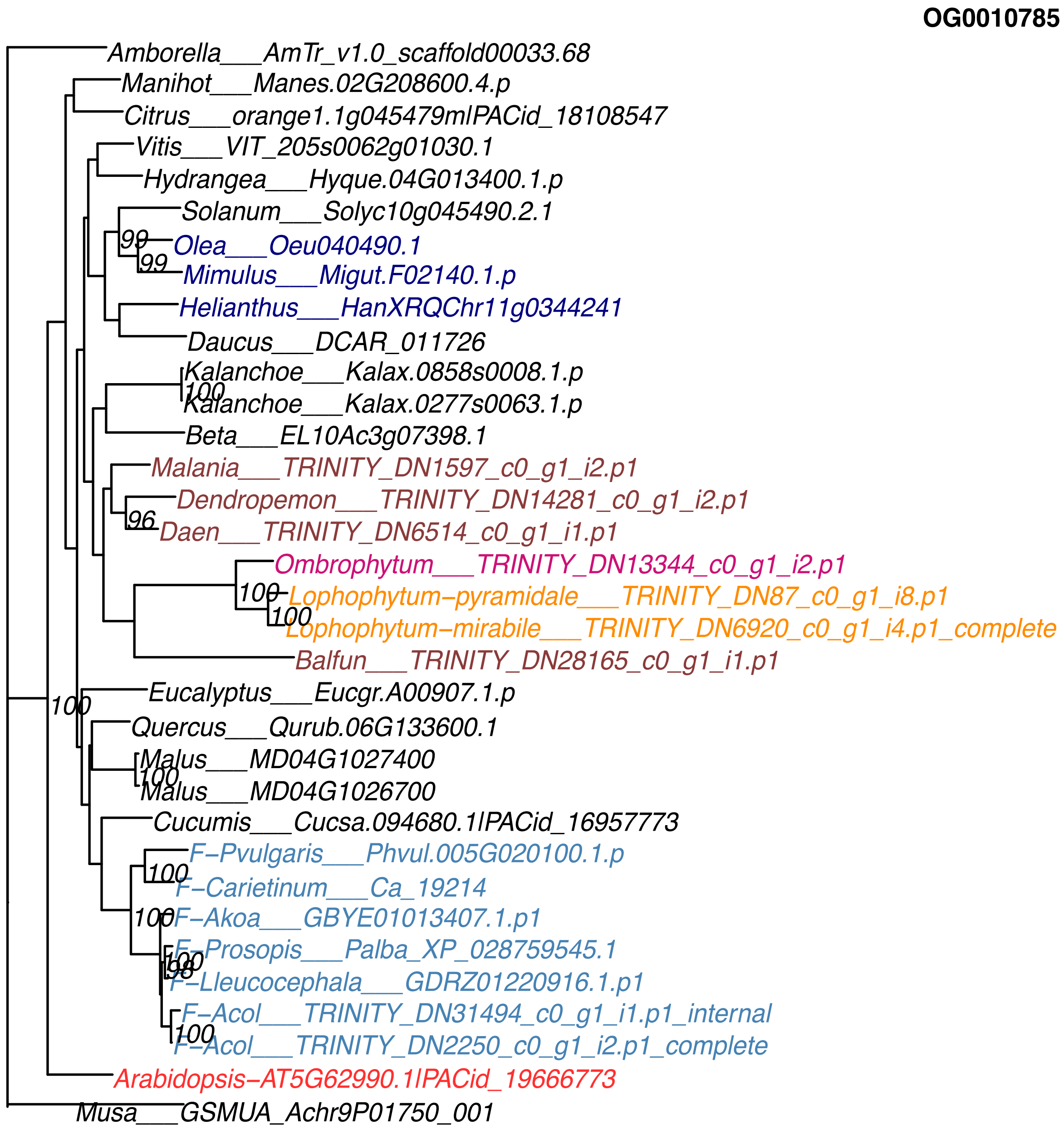
OG0010785
Amborella___AmTr_v1.0_scaffold00033.68
Manihot___Manes.02G208600.4.p
Citrus___orange1.1g045479m|PACid_18108547
Vitis___VIT_205s0062g01030.1
Hydrangea___Hyque.04G013400.1.p
Solanum___Solyc10g045490.2.1
99
Olea___Oeu040490.1
99
Mimulus___Migut.F02140.1.p
Helianthus___HanXRQChr11g0344241
Daucus___DCAR_011726
Kalanchoe___Kalax.0858s0008.1.p
100
Kalanchoe___Kalax.0277s0063.1.p
Beta___EL10Ac3g07398.1
Malania___TRINITY_DN1597_c0_g1_i2.p1
Dendropemon___TRINITY_DN14281_c0_g1_i2.p1
96
Daen___TRINITY_DN6514_c0_g1_i1.p1
Ombrophytum___TRINITY_DN13344_c0_g1_i2.p1
100
Lophophytum−pyramidale___TRINITY_DN87_c0_g1_i8.p1
100
Lophophytum−mirabile___TRINITY_DN6920_c0_g1_i4.p1_complete
Balfun___TRINITY_DN28165_c0_g1_i1.p1
Eucalyptus___Eucgr.A00907.1.p
100
Quercus___Qurub.06G133600.1
Malus___MD04G1027400
100
Malus___MD04G1026700
Cucumis___Cucsa.094680.1|PACid_16957773
F−Pvulgaris___Phvul.005G020100.1.p
100
F−Carietinum___Ca_19214
100
F−Akoa___GBYE01013407.1.p1
F−Prosopis___Palba_XP_028759545.1
100
98
F−Lleucocephala___GDRZ01220916.1.p1
F−Acol___TRINITY_DN31494_c0_g1_i1.p1_internal
100
F−Acol___TRINITY_DN2250_c0_g1_i2.p1_complete
Arabidopsis−AT5G62990.1|PACid_19666773
Musa___GSMUA_Achr9P01750_001

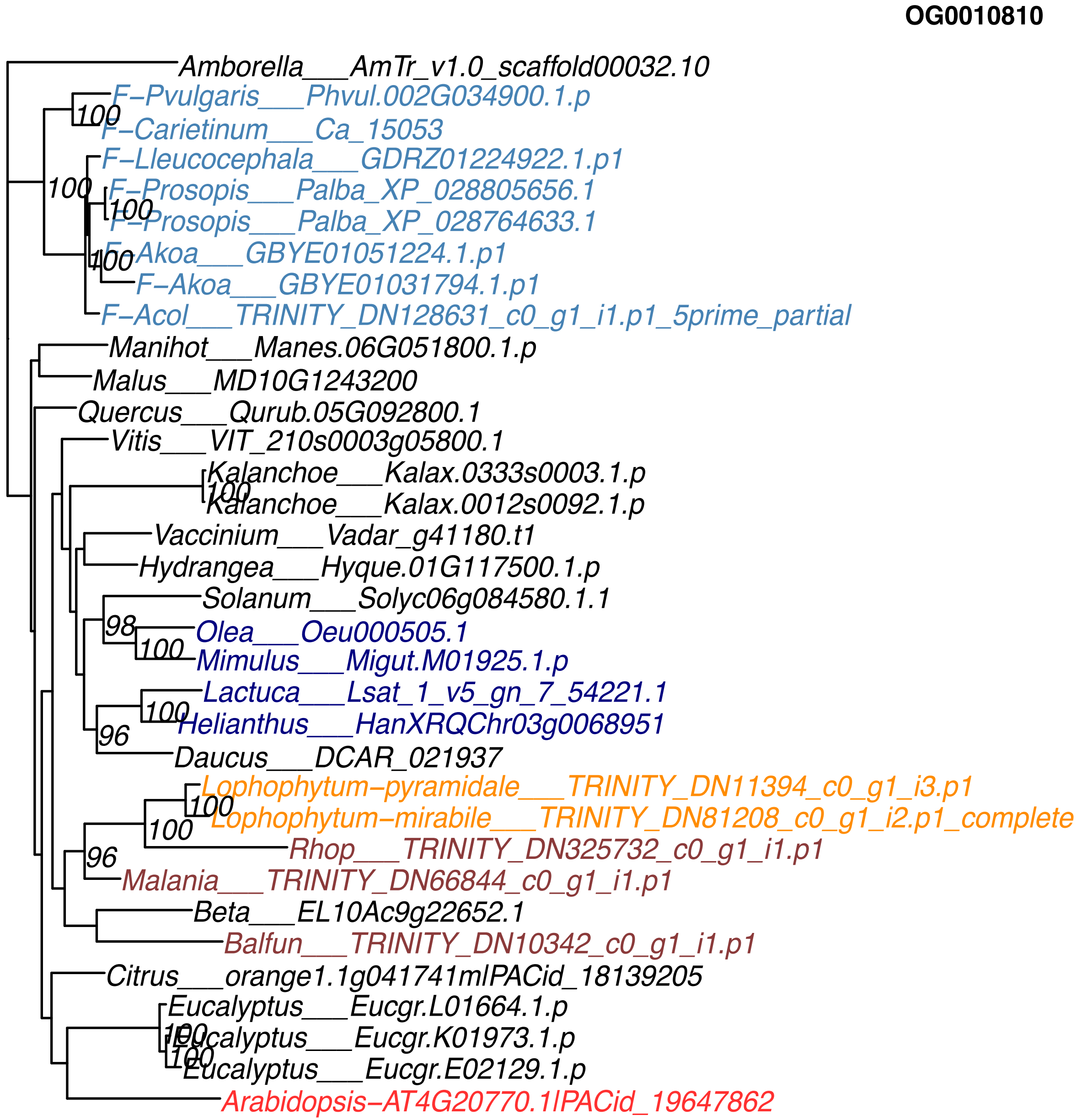
OG0010810
Amborella___AmTr_v1.0_scaffold00032.10
F−Pvulgaris___Phvul.002G034900.1.p
100
F−Carietinum___Ca_15053
F−Lleucocephala___GDRZ01224922.1.p1
100
F−Prosopis___Palba_XP_028805656.1
100
F−Prosopis___Palba_XP_028764633.1
F−Akoa___GBYE01051224.1.p1
100
F−Akoa___GBYE01031794.1.p1
F−Acol___TRINITY_DN128631_c0_g1_i1.p1_5prime_partial
Manihot___Manes.06G051800.1.p
Malus___MD10G1243200
Quercus___Qurub.05G092800.1
Vitis___VIT_210s0003g05800.1
Kalanchoe___Kalax.0333s0003.1.p
100
Kalanchoe___Kalax.0012s0092.1.p
Vaccinium___Vadar_g41180.t1
Hydrangea___Hyque.01G117500.1.p
Solanum___Solyc06g084580.1.1
98
Olea___Oeu000505.1
100
Mimulus___Migut.M01925.1.p
Lactuca___Lsat_1_v5_gn_7_54221.1
100
Helianthus___HanXRQChr03g0068951
96
Daucus___DCAR_021937
Lophophytum−pyramidale___TRINITY_DN11394_c0_g1_i3.p1
100
Lophophytum−mirabile___TRINITY_DN81208_c0_g1_i2.p1_complete
100
Rhop___TRINITY_DN325732_c0_g1_i1.p1
96
Malania___TRINITY_DN66844_c0_g1_i1.p1
Beta___EL10Ac9g22652.1
Balfun___TRINITY_DN10342_c0_g1_i1.p1
Citrus___orange1.1g041741m|PACid_18139205
Eucalyptus___Eucgr.L01664.1.p
100
Eucalyptus___Eucgr.K01973.1.p
100
Eucalyptus___Eucgr.E02129.1.p
Arabidopsis−AT4G20770.1|PACid_19647862

**OG0010896**

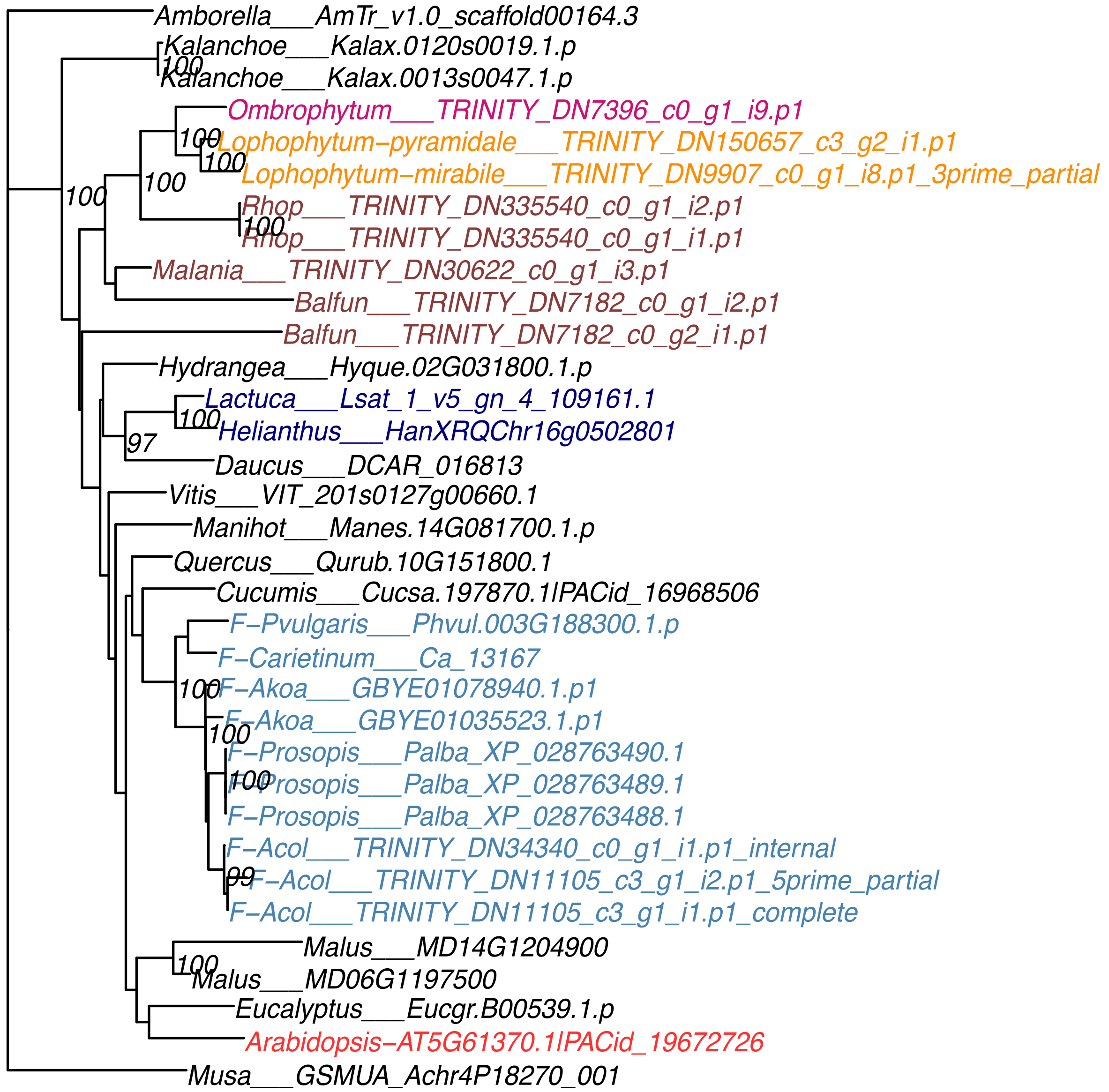
Amborella___AmTr_v1.0_scaffold00164.3
Kalanchoe___Kalax.0120s0019.1.p
100
Kalanchoe___Kalax.0013s0047.1.p
Ombrophytum___TRINITY_DN7396_c0_g1_i9.p1
100
Lophophytum−pyramidale___TRINITY_DN150657_c3_g2_i1.p1
100
Lophophytum−mirabile___TRINITY_DN9907_c0_g1_i8.p1_3prime_partial
100
100
Rhop___TRINITY_DN335540_c0_g1_i2.p1
100
Rhop___TRINITY_DN335540_c0_g1_i1.p1
Malania___TRINITY_DN30622_c0_g1_i3.p1
Balfun___TRINITY_DN7182_c0_g1_i2.p1
Balfun___TRINITY_DN7182_c0_g2_i1.p1
Hydrangea___Hyque.02G031800.1.p
Lactuca___Lsat_1_v5_gn_4_109161.1
100
Helianthus___HanXRQChr16g0502801
97
Daucus___DCAR_016813
Vitis___VIT_201s0127g00660.1
Manihot___Manes.14G081700.1.p
Quercus___Qurub.10G151800.1
Cucumis___Cucsa.197870.1|PACid_16968506
F−Pvulgaris___Phvul.003G188300.1.p
F−Carietinum___Ca_13167
100
F−Akoa___GBYE01078940.1.p1
F−Akoa___GBYE01035523.1.p1
100
F−Prosopis___Palba_XP_028763490.1
100
F−Prosopis___Palba_XP_028763489.1
F−Prosopis___Palba_XP_028763488.1
F−Acol___TRINITY_DN34340_c0_g1_i1.p1_internal
99
F−Acol___TRINITY_DN11105_c3_g1_i2.p1_5prime_partial
F−Acol___TRINITY_DN11105_c3_g1_i1.p1_complete
Malus___MD14G1204900
100
Malus___MD06G1197500
Eucalyptus___Eucgr.B00539.1.p
Arabidopsis−AT5G61370.1|PACid_19672726
Musa___GSMUA_Achr4P18270_001

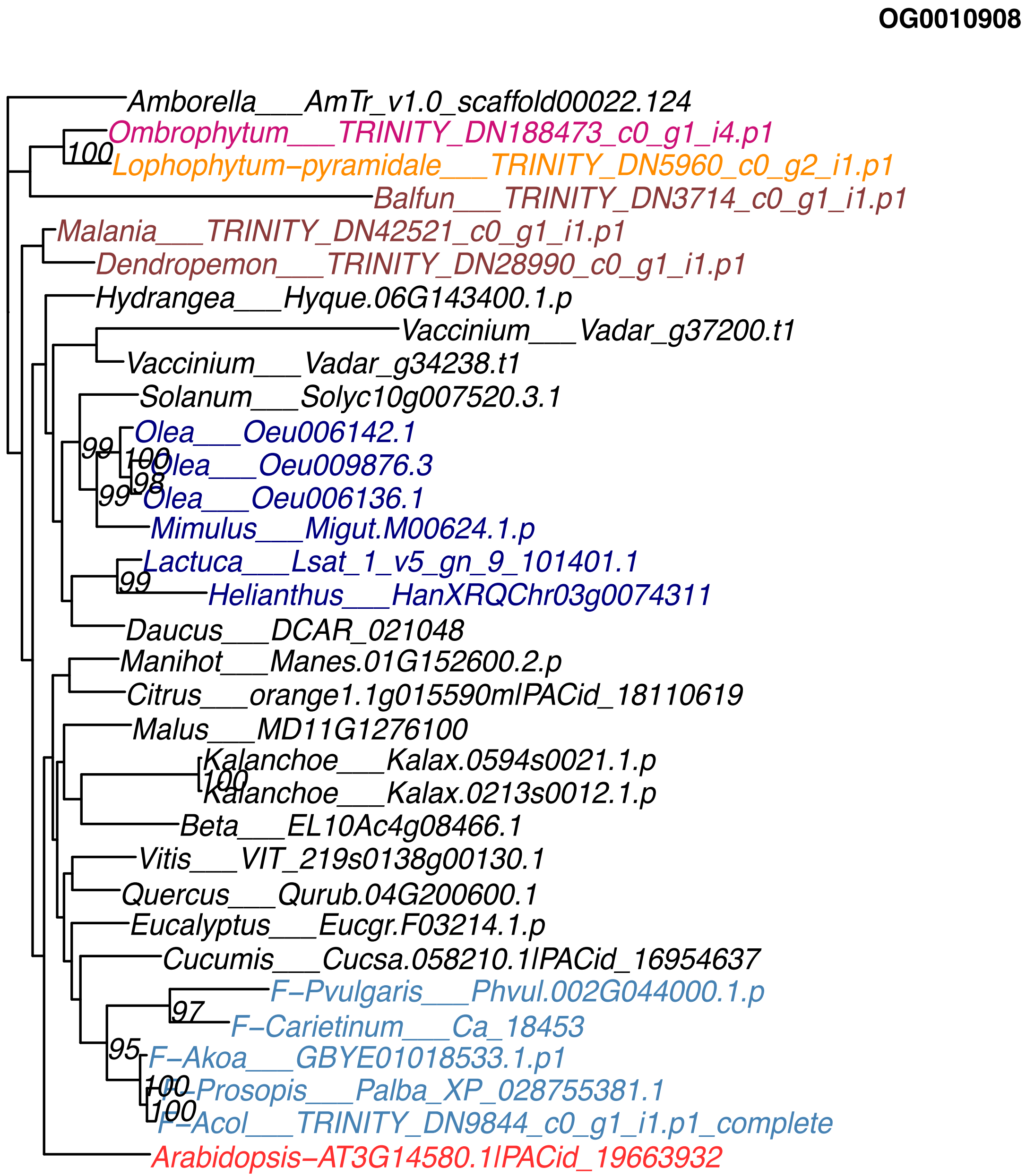
OG0010908
Amborella___AmTr_v1.0_scaffold00022.124
Ombrophytum___TRINITY_DN188473_c0_g1_i4.p1
100
Lophophytum−pyramidale___TRINITY_DN5960_c0_g2_i1.p1
Balfun___TRINITY_DN3714_c0_g1_i1.p1
Malania___TRINITY_DN42521_c0_g1_i1.p1
Dendropemon___TRINITY_DN28990_c0_g1_i1.p1
Hydrangea___Hyque.06G143400.1.p
Vaccinium___Vadar_g37200.t1
Vaccinium___Vadar_g34238.t1
Solanum___Solyc10g007520.3.1
Olea___Oeu006142.1
99
100
Olea___Oeu009876.3
98
99
Olea___Oeu006136.1
Mimulus___Migut.M00624.1.p
Lactuca___Lsat_1_v5_gn_9_101401.1
99
Helianthus___HanXRQChr03g0074311
Daucus___DCAR_021048
Manihot___Manes.01G152600.2.p
Citrus___orange1.1g015590mIPACid_18110619
Malus___MD11G1276100
Kalanchoe___Kalax.0594s0021.1.p
100
Kalanchoe___Kalax.0213s0012.1.p
Beta___EL10Ac4g08466.1
Vitis___VIT_219s0138g00130.1
Quercus___Qurub.04G200600.1
Eucalyptus___Eucgr.F03214.1.p
Cucumis___Cucsa.058210.1IPACid_16954637
F−Pvulgaris___Phvul.002G044000.1.p
97
F−Carietinum___Ca_18453
95
F−Akoa___GBYE01018533.1.p1
100
F−Prosopis___Palba_XP_028755381.1
100
F−Acol___TRINITY_DN9844_c0_g1_i1.p1_complete
Arabidopsis−AT3G14580.1IPACid_19663932

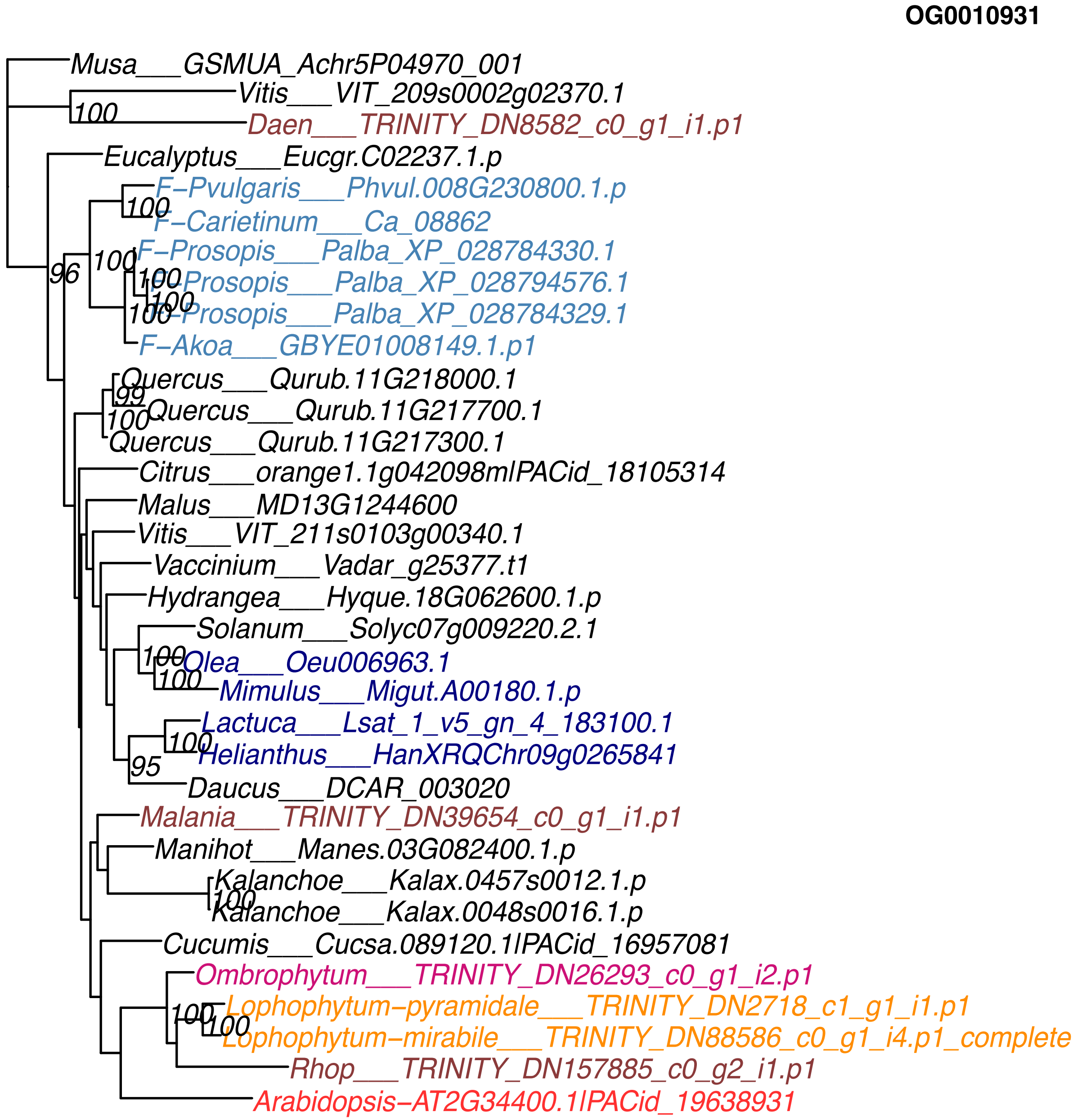
OG0010931
Musa___GSMUA_Achr5P04970_001
Vitis___VIT_209s0002g02370.1
100
Daen___TRINITY_DN8582_c0_g1_i1.p1
Eucalyptus___Eucgr.C02237.1.p
F−Pvulgaris___Phvul.008G230800.1.p
100
F−Carietinum___Ca_08862
96
100
F−Prosopis___Palba_XP_028784330.1
100
F−Prosopis___Palba_XP_028794576.1
100
100
F−Prosopis___Palba_XP_028784329.1
F−Akoa___GBYE01008149.1.p1
Quercus___Qurub.11G218000.1
99
100
Quercus___Qurub.11G217700.1
Quercus___Qurub.11G217300.1
Citrus___orange1.1g042098m|PACid_18105314
Malus___MD13G1244600
Vitis___VIT_211s0103g00340.1
Vaccinium___Vadar_g25377.t1
Hydrangea___Hyque.18G062600.1.p
Solanum___Solyc07g009220.2.1
100
Olea___Oeu006963.1
100
Mimulus___Migut.A00180.1.p
Lactuca___Lsat_1_v5_gn_4_183100.1
100
Helianthus___HanXRQChr09g0265841
95
Daucus___DCAR_003020
Malania___TRINITY_DN39654_c0_g1_i1.p1
Manihot___Manes.03G082400.1.p
Kalanchoe___Kalax.0457s0012.1.p
100
Kalanchoe___Kalax.0048s0016.1.p
Cucumis___Cucsa.089120.1|PACid_16957081
Ombrophytum___TRINITY_DN26293_c0_g1_i2.p1
100
Lophophytum−pyramidale___TRINITY_DN2718_c1_g1_i1.p1
100
Lophophytum−mirabile___TRINITY_DN88586_c0_g1_i4.p1_complete
Rhop___TRINITY_DN157885_c0_g2_i1.p1
Arabidopsis−AT2G34400.1|PACid_19638931

**OG0010959**

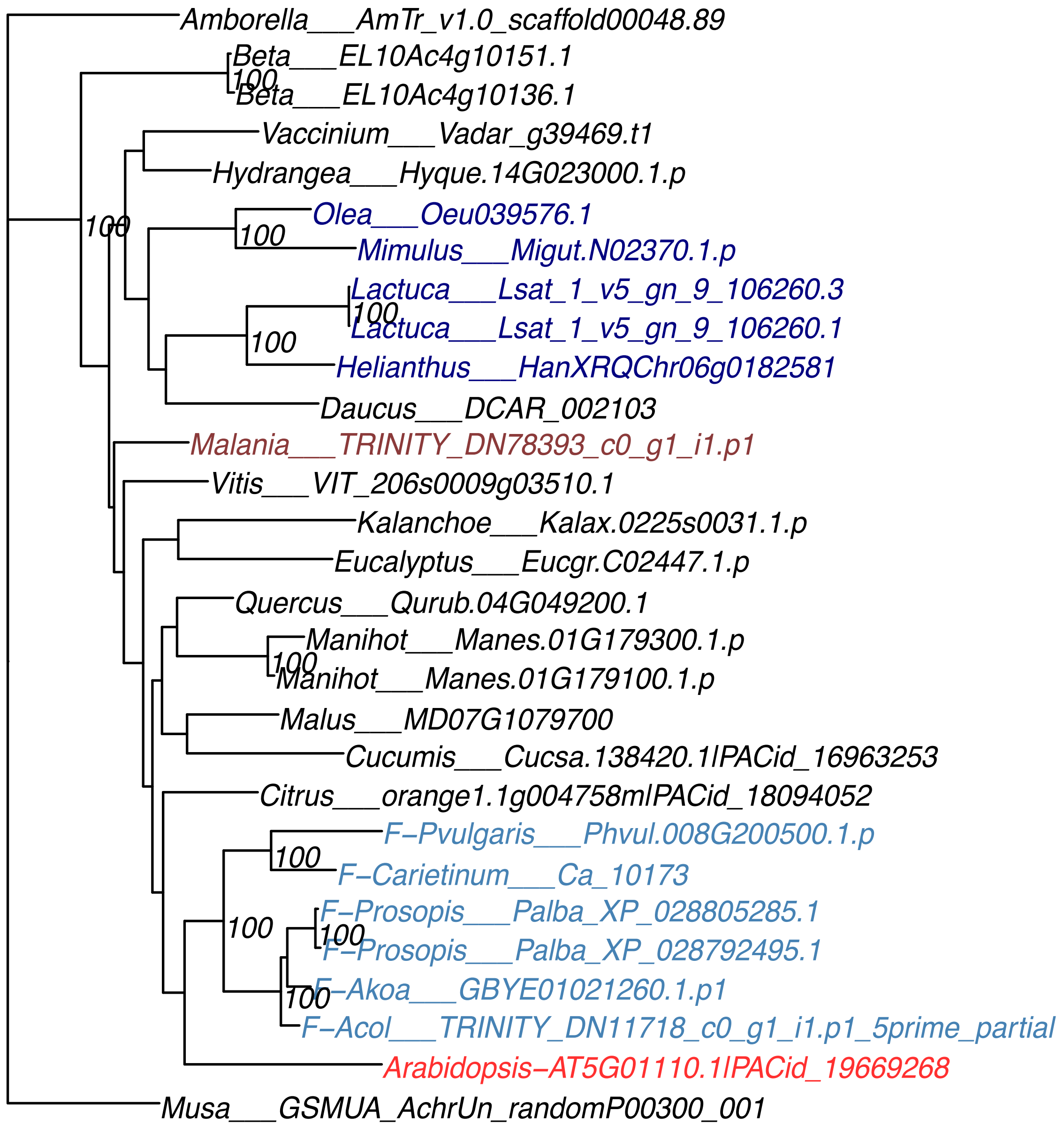
Amborella___AmTr_v1.0_scaffold00048.89
Beta___EL10Ac4g10151.1
100
Beta___EL10Ac4g10136.1
Vaccinium___Vadar_g39469.t1
Hydrangea___Hyque.14G023000.1.p
100
Olea___Oeu039576.1
100
Mimulus___Migut.N02370.1.p
Lactuca___Lsat_1_v5_gn_9_106260.3
100
Lactuca___Lsat_1_v5_gn_9_106260.1
100
Helianthus___HanXRQChr06g0182581
Daucus___DCAR_002103
Malania___TRINITY_DN78393_c0_g1_i1.p1
Vitis___VIT_206s0009g03510.1
Kalanchoe___Kalax.0225s0031.1.p
Eucalyptus___Eucgr.C02447.1.p
Quercus___Qurub.04G049200.1
Manihot___Manes.01G179300.1.p
100
Manihot___Manes.01G179100.1.p
Malus___MD07G1079700
Cucumis___Cucsa.138420.1lPACid_16963253
Citrus___orange1.1g004758mlPACid_18094052
F−Pvulgaris___Phvul.008G200500.1.p
100
F−Carietinum___Ca_10173
F−Prosopis___Palba_XP_028805285.1
100
100
F−Prosopis___Palba_XP_028792495.1
F−Akoa___GBYE01021260.1.p1
100
F−Acol___TRINITY_DN11718_c0_g1_i1.p1_5prime_partial
Arabidopsis−AT5G01110.1lPACid_19669268
Musa___GSMUA_AchrUn_randomP00300_001

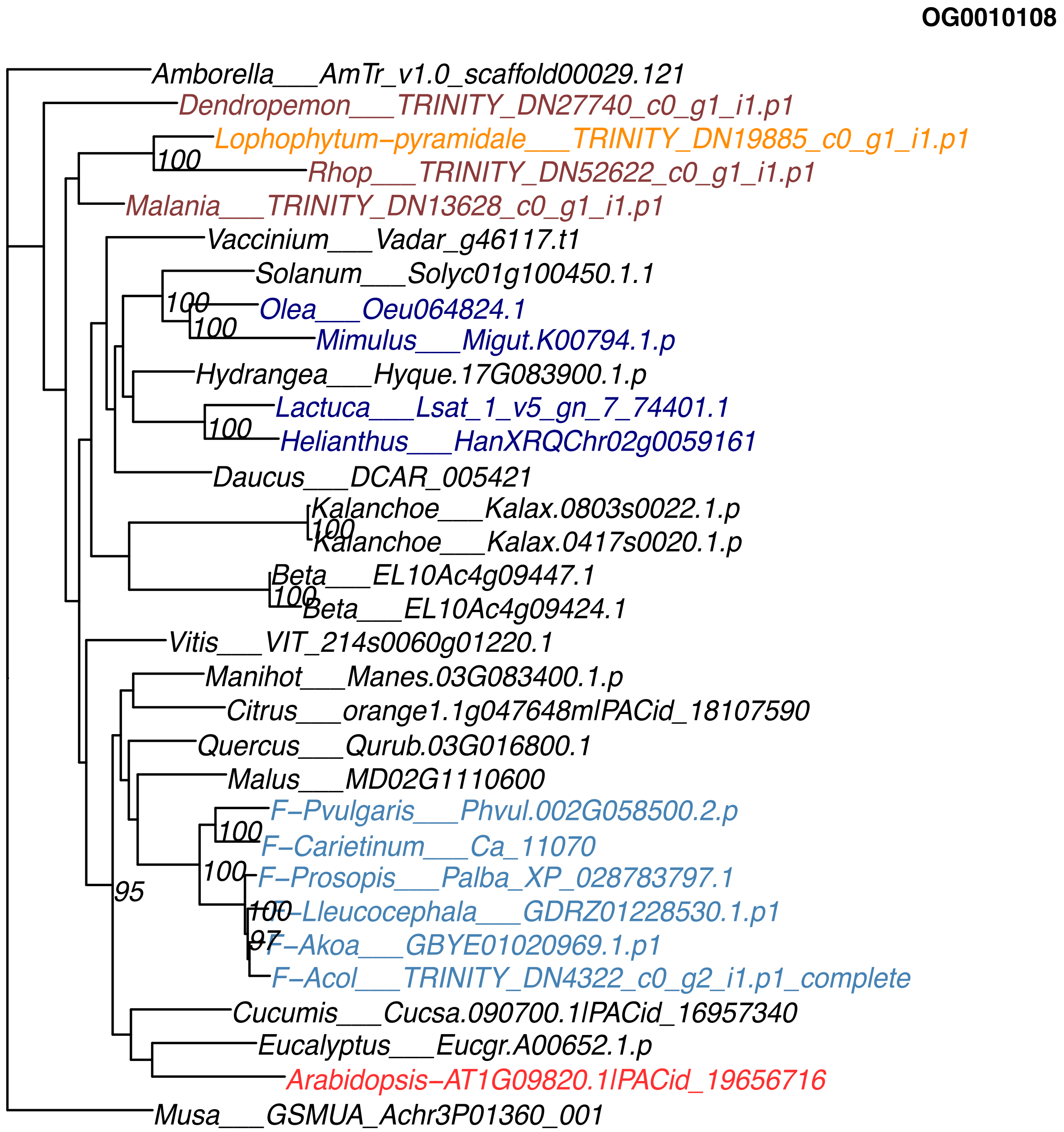
OG0010108
Amborella___AmTr_v1.0_scaffold00029.121
Dendropemon___TRINITY_DN27740_c0_g1_i1.p1
Lophophytum−pyramidale___TRINITY_DN19885_c0_g1_i1.p1
100
Rhop___TRINITY_DN52622_c0_g1_i1.p1
Malania___TRINITY_DN13628_c0_g1_i1.p1
Vaccinium___Vadar_g46117.t1
Solanum___Solyc01g100450.1.1
100
Olea___Oeu064824.1
100
Mimulus___Migut.K00794.1.p
Hydrangea___Hyque.17G083900.1.p
Lactuca___Lsat_1_v5_gn_7_74401.1
100
Helianthus___HanXRQChr02g0059161
Daucus___DCAR_005421
Kalanchoe___Kalax.0803s0022.1.p
100
Kalanchoe___Kalax.0417s0020.1.p
Beta___EL10Ac4g09447.1
100
Beta___EL10Ac4g09424.1
Vitis___VIT_214s0060g01220.1
Manihot___Manes.03G083400.1.p
Citrus___orange1.1g047648mIPACid_18107590
Quercus___Qurub.03G016800.1
Malus___MD02G1110600
F−Pvulgaris___Phvul.002G058500.2.p
100
F−Carietinum___Ca_11070
100
F−Prosopis___Palba_XP_028783797.1
95
100
F−Lleucocephala___GDRZ01228530.1.p1
97
F−Akoa___GBYE01020969.1.p1
F−Acol___TRINITY_DN4322_c0_g2_i1.p1_complete
Cucumis___Cucsa.090700.1IPACid_16957340
Eucalyptus___Eucgr.A00652.1.p
Arabidopsis−AT1G09820.1IPACid_19656716
Musa___GSMUA_Achr3P01360_001

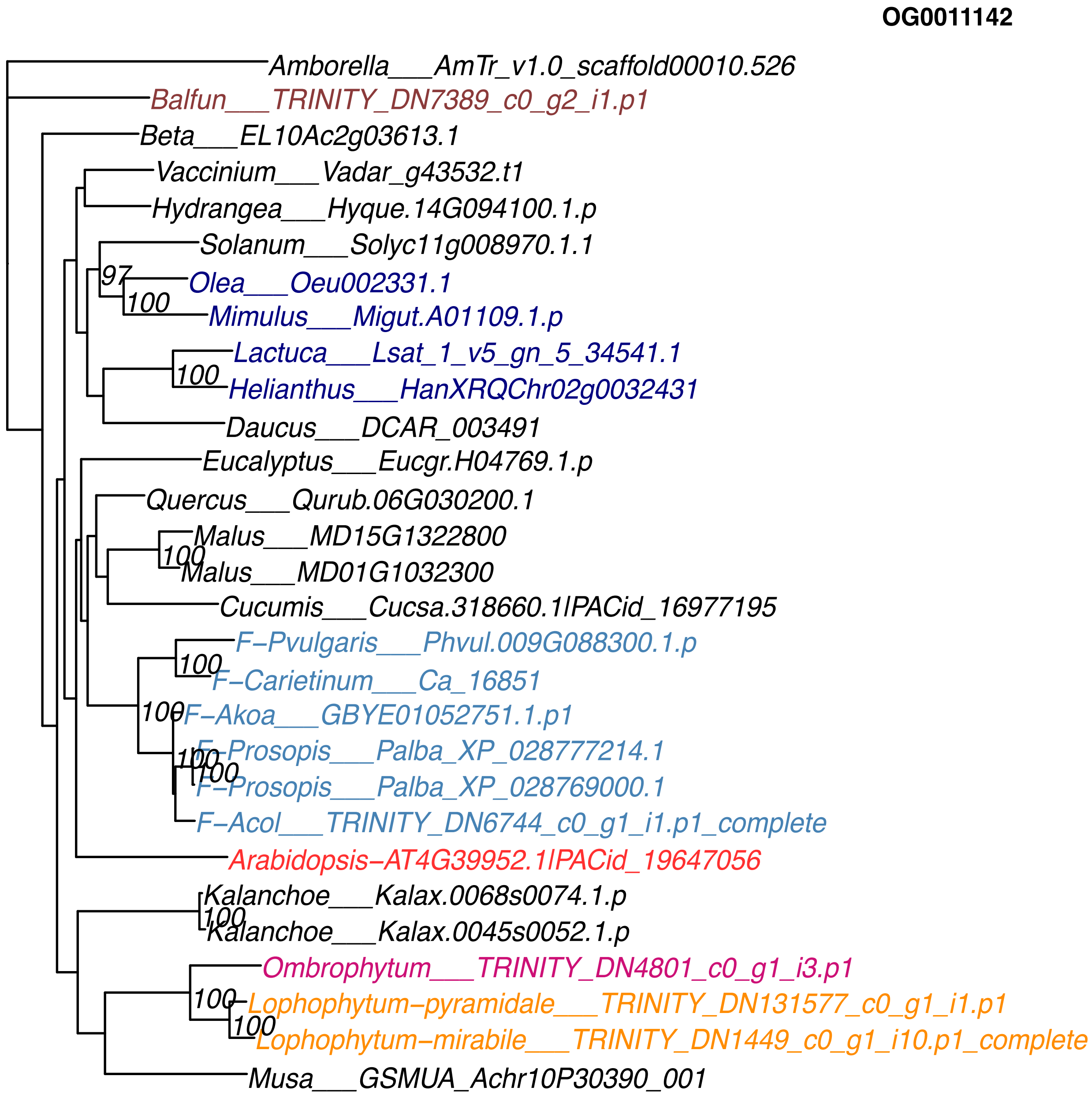
OG0011142
Amborella___AmTr_v1.0_scaffold00010.526
Balfun___TRINITY_DN7389_c0_g2_i1.p1
Beta___EL10Ac2g03613.1
Vaccinium___Vadar_g43532.t1
Hydrangea___Hyque.14G094100.1.p
Solanum___Solyc11g008970.1.1
97
Olea___Oeu002331.1
100
Mimulus___Migut.A01109.1.p
Lactuca___Lsat_1_v5_gn_5_34541.1
100
Helianthus___HanXRQChr02g0032431
Daucus___DCAR_003491
Eucalyptus___Eucgr.H04769.1.p
Quercus___Qurub.06G030200.1
Malus___MD15G1322800
100
Malus___MD01G1032300
Cucumis___Cucsa.318660.1|PACid_16977195
F−Pvulgaris___Phvul.009G088300.1.p
100
F−Carietinum___Ca_16851
100
F−Akoa___GBYE01052751.1.p1
F−Prosopis___Palba_XP_028777214.1
100
100
F−Prosopis___Palba_XP_028769000.1
F−Acol___TRINITY_DN6744_c0_g1_i1.p1_complete
Arabidopsis−AT4G39952.1|PACid_19647056
Kalanchoe___Kalax.0068s0074.1.p
100
Kalanchoe___Kalax.0045s0052.1.p
Ombrophytum___TRINITY_DN4801_c0_g1_i3.p1
100
Lophophytum−pyramidale___TRINITY_DN131577_c0_g1_i1.p1
100
Lophophytum−mirabile___TRINITY_DN1449_c0_g1_i10.p1_complete
Musa___GSMUA_Achr10P30390_001

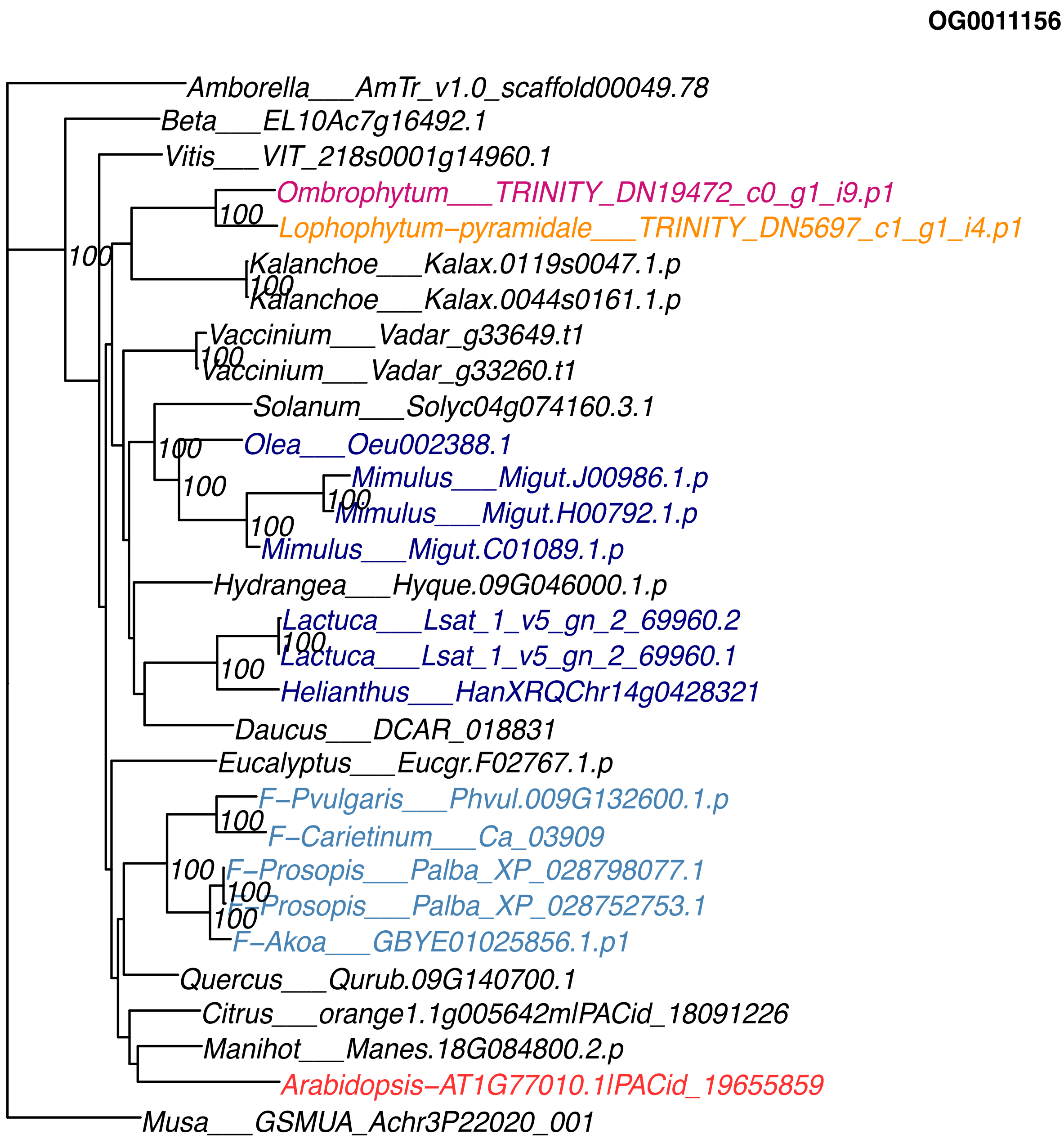
OG0011156
Amborella___AmTr_v1.0_scaffold00049.78
Beta___EL10Ac7g16492.1
Vitis___VIT_218s0001g14960.1
Ombrophytum___TRINITY_DN19472_c0_g1_i9.p1
Lophophytum−pyramidale___TRINITY_DN5697_c1_g1_i4.p1
Kalanchoe___Kalax.0119s0047.1.p
Kalanchoe___Kalax.0044s0161.1.p
Vaccinium___Vadar_g33649.t1
Vaccinium___Vadar_g33260.t1
Solanum___Solyc04g074160.3.1
Olea___Oeu002388.1
Mimulus___Migut.J00986.1.p
Mimulus___Migut.H00792.1.p
Mimulus___Migut.C01089.1.p
Hydrangea___Hyque.09G046000.1.p
Lactuca___Lsat_1_v5_gn_2_69960.2
Lactuca___Lsat_1_v5_gn_2_69960.1
Helianthus___HanXRQChr14g0428321
Daucus___DCAR_018831
Eucalyptus___Eucgr.F02767.1.p
F−Pvulgaris___Phvul.009G132600.1.p
F−Carietinum___Ca_03909
F−Prosopis___Palba_XP_028798077.1
F−Prosopis___Palba_XP_028752753.1
F−Akoa___GBYE01025856.1.p1
Quercus___Qurub.09G140700.1
Citrus___orange1.1g005642mlPACid_18091226
Manihot___Manes.18G084800.2.p
Arabidopsis−AT1G77010.1lPACid_19655859
Musa___GSMUA_Achr3P22020_001
100
100
100
100
100
100
100
100
100
100
100
100
100
100

**OG0011157**

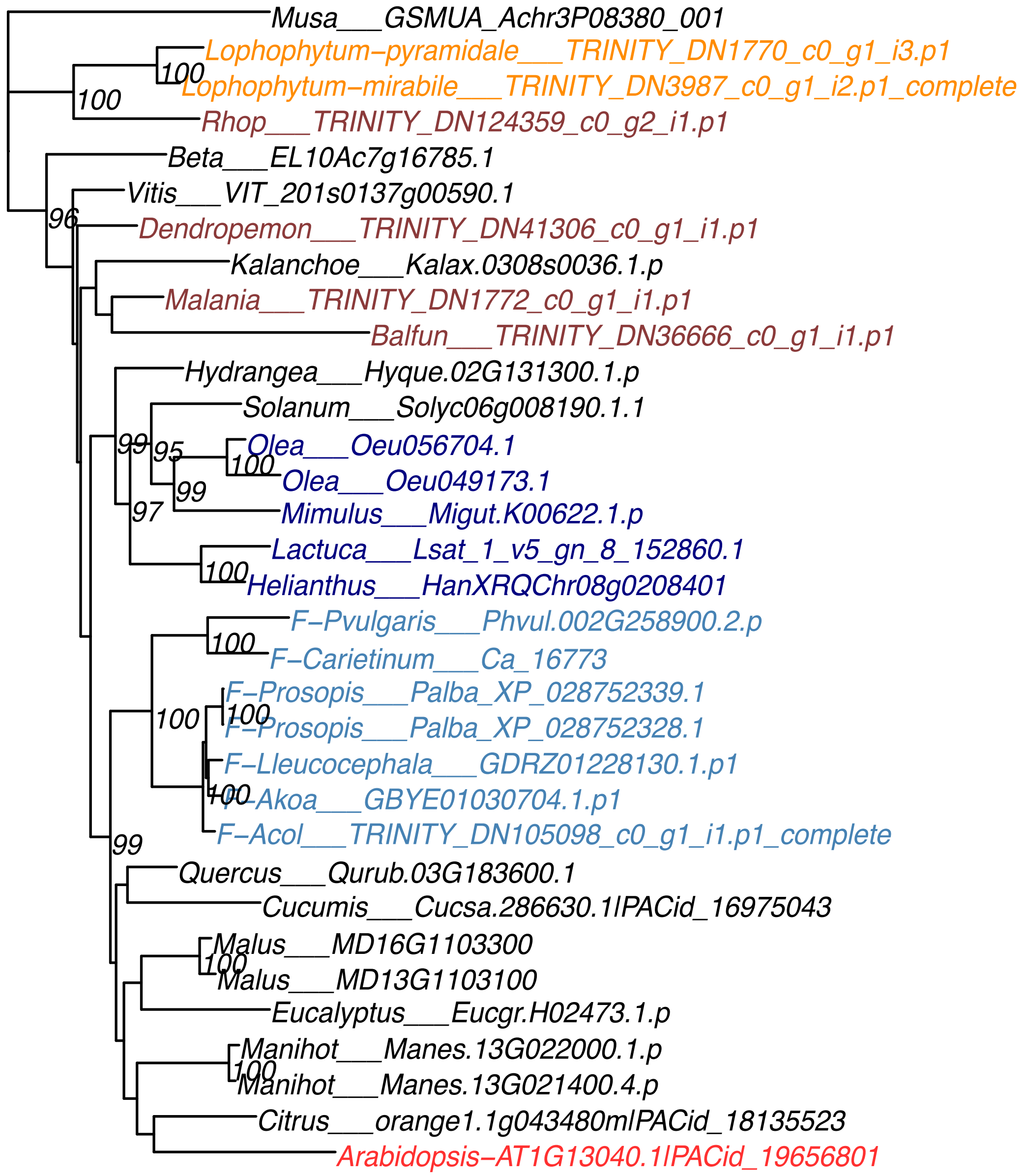

Musa___GSMUA_Achr3P08380_001
Lophophytum−pyramidale___TRINITY_DN1770_c0_g1_i3.p1
Lophophytum−mirabile___TRINITY_DN3987_c0_g1_i2.p1_complete
Rhop___TRINITY_DN124359_c0_g2_i1.p1
Beta___EL10Ac7g16785.1
Vitis___VIT_201s0137g00590.1
Dendropemon___TRINITY_DN41306_c0_g1_i1.p1
Kalanchoe___Kalax.0308s0036.1.p
Malania___TRINITY_DN1772_c0_g1_i1.p1
Balfun___TRINITY_DN36666_c0_g1_i1.p1
Hydrangea___Hyque.02G131300.1.p
Solanum___Solyc06g008190.1.1
Olea___Oeu056704.1
Olea___Oeu049173.1
Mimulus___Migut.K00622.1.p
Lactuca___Lsat_1_v5_gn_8_152860.1
Helianthus___HanXRQChr08g0208401
F−Pvulgaris___Phvul.002G258900.2.p
F−Carietinum___Ca_16773
F−Prosopis___Palba_XP_028752339.1
F−Prosopis___Palba_XP_028752328.1
F−Lleucocephala___GDRZ01228130.1.p1
F−Akoa___GBYE01030704.1.p1
F−Acol___TRINITY_DN105098_c0_g1_i1.p1_complete
Quercus___Qurub.03G183600.1
Cucumis___Cucsa.286630.1|PACid_16975043
Malus___MD16G1103300
Malus___MD13G1103100
Eucalyptus___Eucgr.H02473.1.p
Manihot___Manes.13G022000.1.p
Manihot___Manes.13G021400.4.p
Citrus___orange1.1g043480m|PACid_18135523
Arabidopsis−AT1G13040.1|PACid_19656801
100
100
96
99
95
100
99
97
100
100
100
100
100
99
100
100

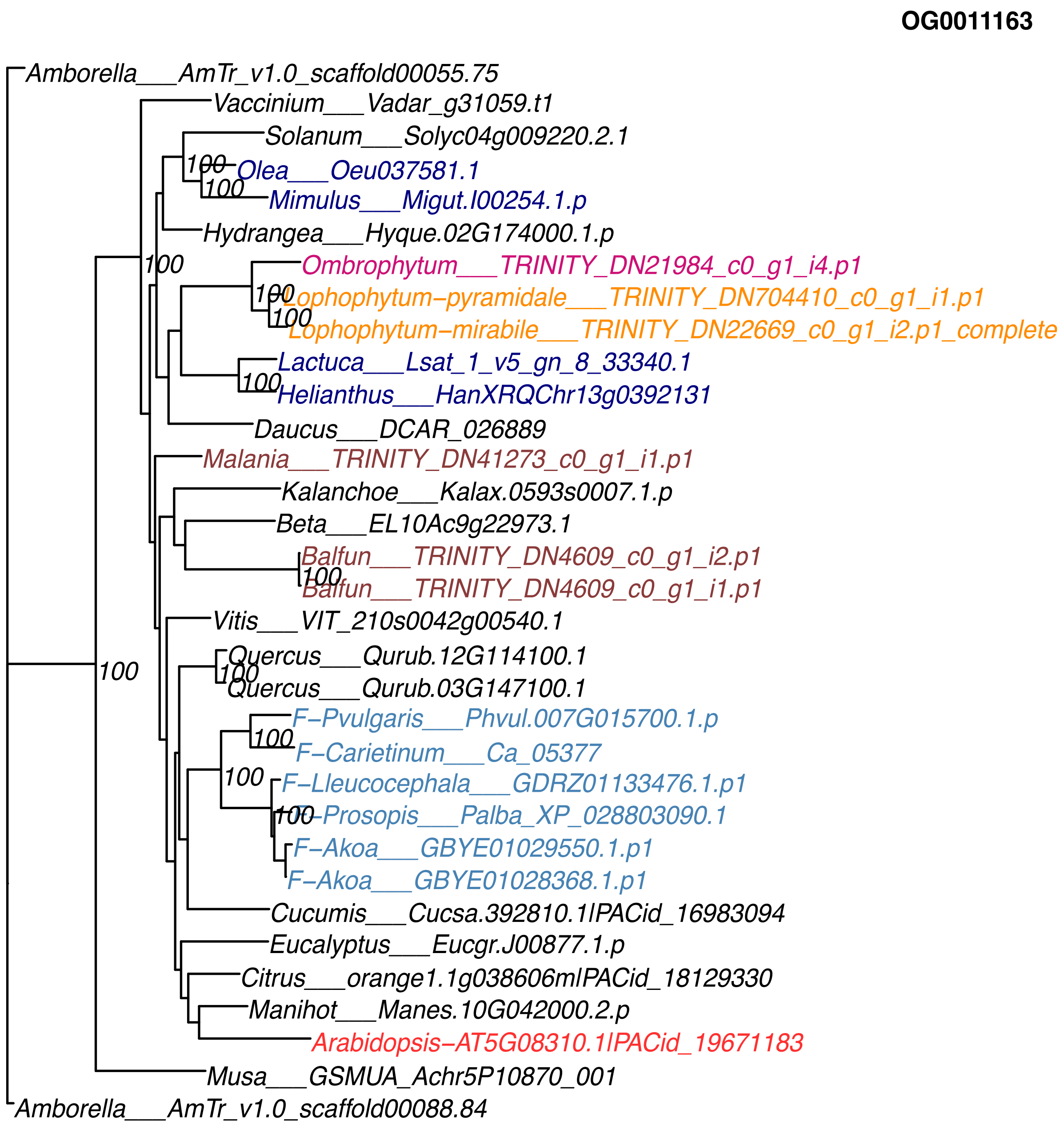

OG0011163
Amborella___AmTr_v1.0_scaffold00055.75
Vaccinium___Vadar_g31059.t1
Solanum___Solyc04g009220.2.1
Olea___Oeu037581.1
Mimulus___Migut.I00254.1.p
Hydrangea___Hyque.02G174000.1.p
Ombrophytum___TRINITY_DN21984_c0_g1_i4.p1
Lophophytum−pyramidale___TRINITY_DN704410_c0_g1_i1.p1
Lophophytum−mirabile___TRINITY_DN22669_c0_g1_i2.p1_complete
Lactuca___Lsat_1_v5_gn_8_33340.1
Helianthus___HanXRQChr13g0392131
Daucus___DCAR_026889
Malania___TRINITY_DN41273_c0_g1_i1.p1
Kalanchoe___Kalax.0593s0007.1.p
Beta___EL10Ac9g22973.1
Balfun___TRINITY_DN4609_c0_g1_i2.p1
Balfun___TRINITY_DN4609_c0_g1_i1.p1
Vitis___VIT_210s0042g00540.1
Quercus___Qurub.12G114100.1
Quercus___Qurub.03G147100.1
F−Pvulgaris___Phvul.007G015700.1.p
F−Carietinum___Ca_05377
F−Lleucocephala___GDRZ01133476.1.p1
F−Prosopis___Palba_XP_028803090.1
F−Akoa___GBYE01029550.1.p1
F−Akoa___GBYE01028368.1.p1
Cucumis___Cucsa.392810.1|PACid_16983094
Eucalyptus___Eucgr.J00877.1.p
Citrus___orange1.1g038606m|PACid_18129330
Manihot___Manes.10G042000.2.p
Arabidopsis−AT5G08310.1|PACid_19671183
Musa___GSMUA_Achr5P10870_001
Amborella___AmTr_v1.0_scaffold00088.84
100
100
100
100
100
100
100
100
100
100
100
100
100

**OG0011248**

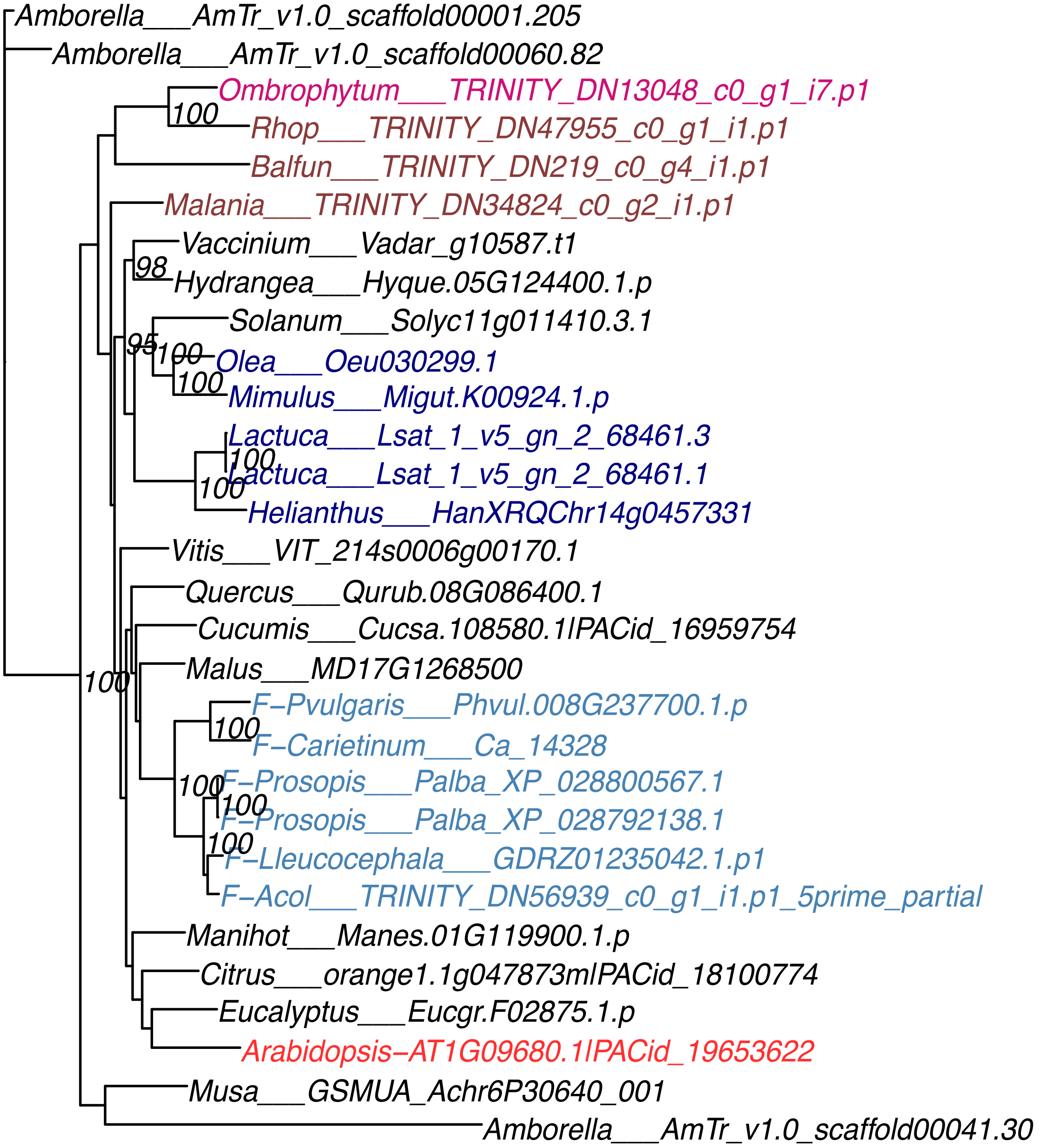
Amborella___AmTr_v1.0_scaffold00001.205
Amborella___AmTr_v1.0_scaffold00060.82
Ombrophytum___TRINITY_DN13048_c0_g1_i7.p1
100
Rhop___TRINITY_DN47955_c0_g1_i1.p1
Balfun___TRINITY_DN219_c0_g4_i1.p1
Malania___TRINITY_DN34824_c0_g2_i1.p1
Vaccinium___Vadar_g10587.t1
98
Hydrangea___Hyque.05G124400.1.p
Solanum___Solyc11g011410.3.1
95
100
Olea___Oeu030299.1
100
Mimulus___Migut.K00924.1.p
Lactuca___Lsat_1_v5_gn_2_68461.3
100
Lactuca___Lsat_1_v5_gn_2_68461.1
100
Helianthus___HanXRQChr14g0457331
Vitis___VIT_214s0006g00170.1
Quercus___Qurub.08G086400.1
Cucumis___Cucsa.108580.1|PACid_16959754
Malus___MD17G1268500
100
F−Pvulgaris___Phvul.008G237700.1.p
100
F−Carietinum___Ca_14328
100
F−Prosopis___Palba_XP_028800567.1
100
F−Prosopis___Palba_XP_028792138.1
100
F−Lleucocephala___GDRZ01235042.1.p1
F−Acol___TRINITY_DN56939_c0_g1_i1.p1_5prime_partial
Manihot___Manes.01G119900.1.p
Citrus___orange1.1g047873m|PACid_18100774
Eucalyptus___Eucgr.F02875.1.p
Arabidopsis−AT1G09680.1|PACid_19653622
Musa___GSMUA_Achr6P30640_001
Amborella___AmTr_v1.0_scaffold00041.30

**OG0011357**

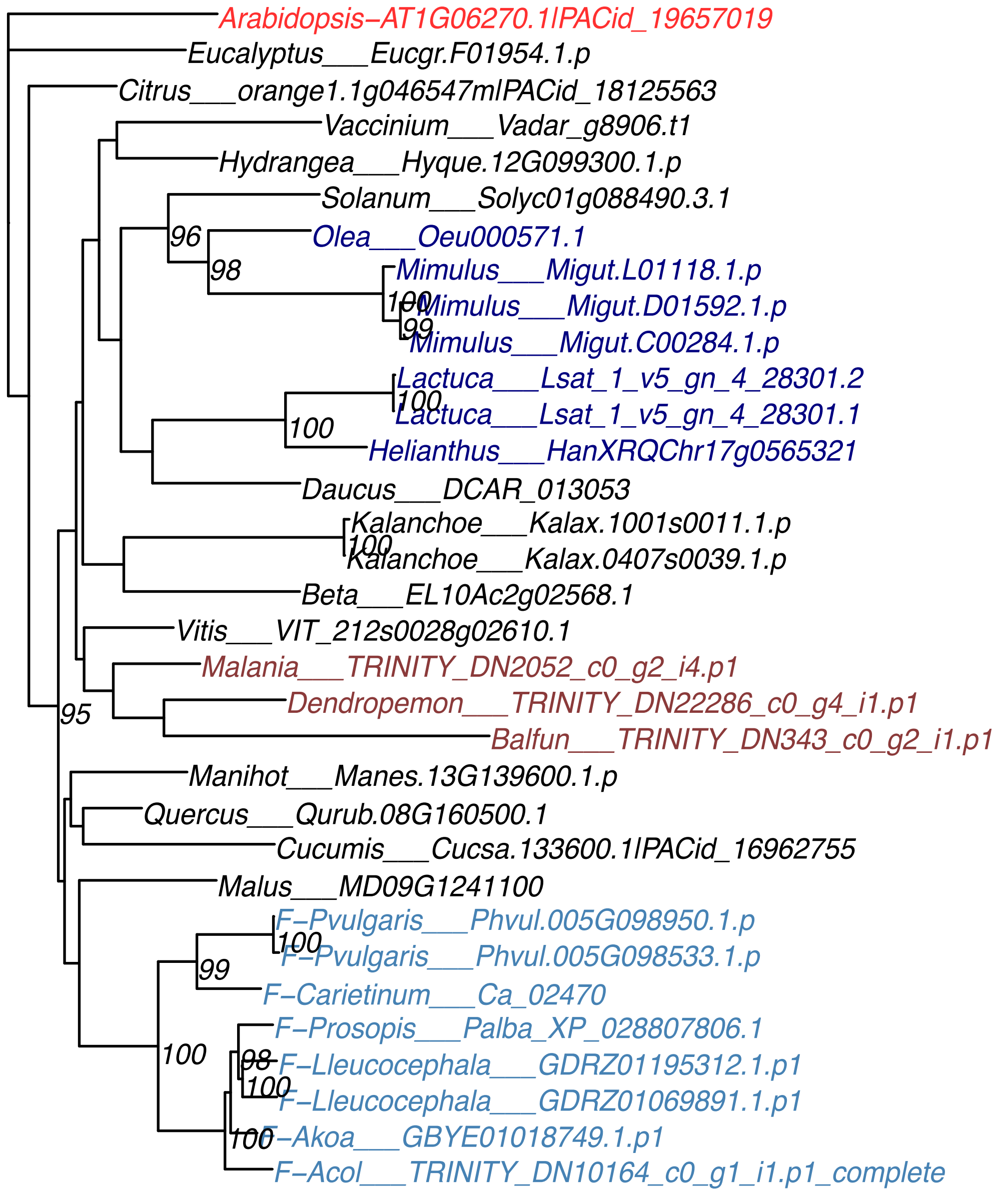

Arabidopsis−AT1G06270.1|PACid_19657019
Eucalyptus___Eucgr.F01954.1.p
Citrus___orange1.1g046547m|PACid_18125563
Vaccinium___Vadar_g8906.t1
Hydrangea___Hyque.12G099300.1.p
Solanum___Solyc01g088490.3.1
96
Olea___Oeu000571.1
98
Mimulus___Migut.L01118.1.p
100
Mimulus___Migut.D01592.1.p
99
Mimulus___Migut.C00284.1.p
Lactuca___Lsat_1_v5_gn_4_28301.2
100
Lactuca___Lsat_1_v5_gn_4_28301.1
100
Helianthus___HanXRQChr17g0565321
Daucus___DCAR_013053
Kalanchoe___Kalax.1001s0011.1.p
100
Kalanchoe___Kalax.0407s0039.1.p
Beta___EL10Ac2g02568.1
Vitis___VIT_212s0028g02610.1
Malania___TRINITY_DN2052_c0_g2_i4.p1
Dendropemon___TRINITY_DN22286_c0_g4_i1.p1
95
Balfun___TRINITY_DN343_c0_g2_i1.p1
Manihot___Manes.13G139600.1.p
Quercus___Qurub.08G160500.1
Cucumis___Cucsa.133600.1|PACid_16962755
Malus___MD09G1241100
F−Pvulgaris___Phvul.005G098950.1.p
100
F−Pvulgaris___Phvul.005G098533.1.p
99
F−Carietinum___Ca_02470
F−Prosopis___Palba_XP_028807806.1
100
98
F−Lleucocephala___GDRZ01195312.1.p1
100
F−Lleucocephala___GDRZ01069891.1.p1
100
F−Akoa___GBYE01018749.1.p1
F−Acol___TRINITY_DN10164_c0_g1_i1.p1_complete

**OG0011375**

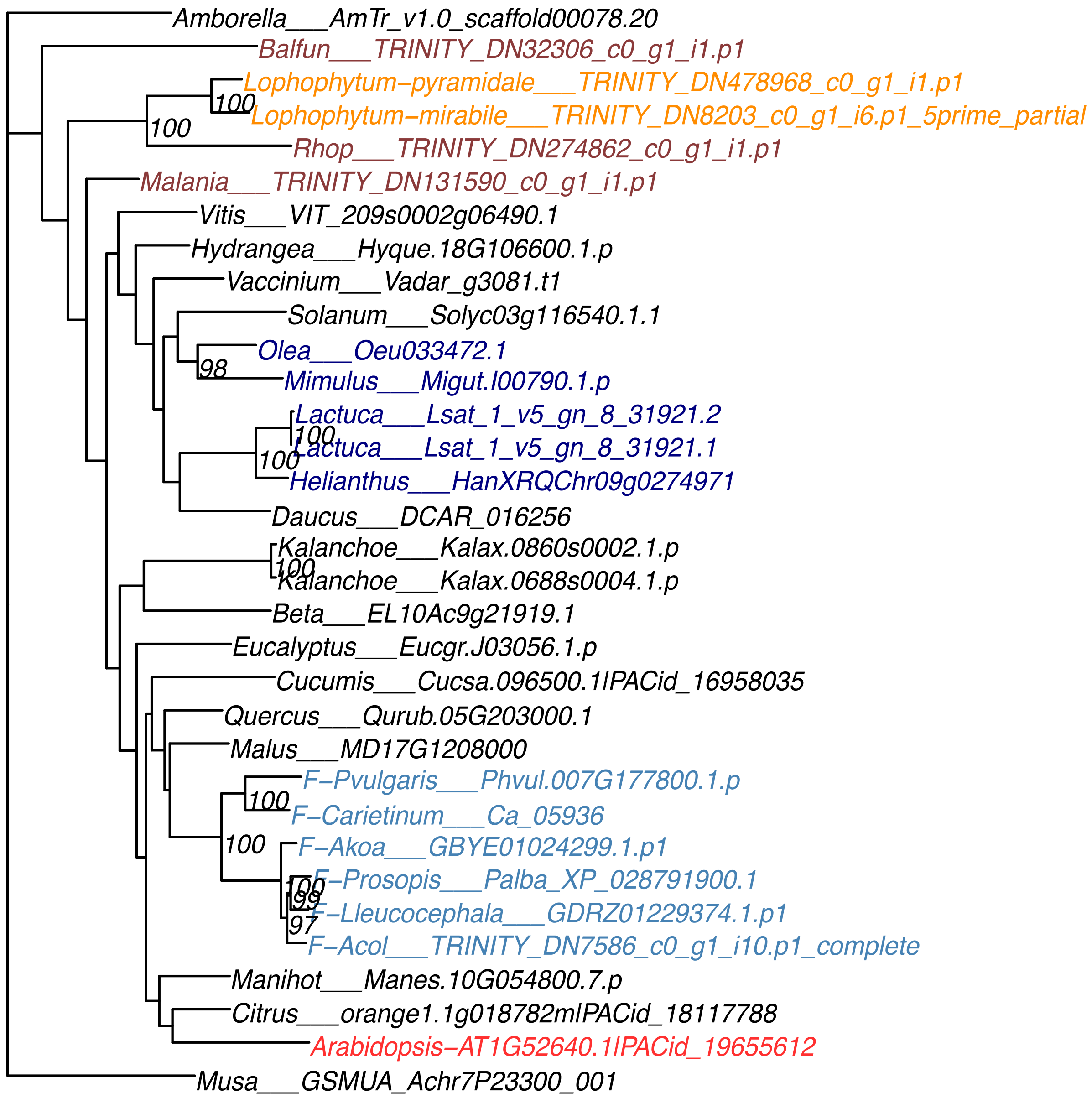

Amborella___AmTr_v1.0_scaffold00078.20
Balfun___TRINITY_DN32306_c0_g1_i1.p1
Lophophytum−pyramidale___TRINITY_DN478968_c0_g1_i1.p1
100
Lophophytum−mirabile___TRINITY_DN8203_c0_g1_i6.p1_5prime_partial
100
Rhop___TRINITY_DN274862_c0_g1_i1.p1
Malania___TRINITY_DN131590_c0_g1_i1.p1
Vitis___VIT_209s0002g06490.1
Hydrangea___Hyque.18G106600.1.p
Vaccinium___Vadar_g3081.t1
Solanum___Solyc03g116540.1.1
Olea___Oeu033472.1
98
Mimulus___Migut.I00790.1.p
Lactuca___Lsat_1_v5_gn_8_31921.2
100
Lactuca___Lsat_1_v5_gn_8_31921.1
100
Helianthus___HanXRQChr09g0274971
Daucus___DCAR_016256
Kalanchoe___Kalax.0860s0002.1.p
100
Kalanchoe___Kalax.0688s0004.1.p
Beta___EL10Ac9g21919.1
Eucalyptus___Eucgr.J03056.1.p
Cucumis___Cucsa.096500.1|PACid_16958035
Quercus___Qurub.05G203000.1
Malus___MD17G1208000
F−Pvulgaris___Phvul.007G177800.1.p
100
F−Carietinum___Ca_05936
100
F−Akoa___GBYE01024299.1.p1
F−Prosopis___Palba_XP_028791900.1
100
99
F−Lleucocephala___GDRZ01229374.1.p1
97
F−Acol___TRINITY_DN7586_c0_g1_i10.p1_complete
Manihot___Manes.10G054800.7.p
Citrus___orange1.1g018782m|PACid_18117788
Arabidopsis−AT1G52640.1|PACid_19655612
Musa___GSMUA_Achr7P23300_001

OG0011432

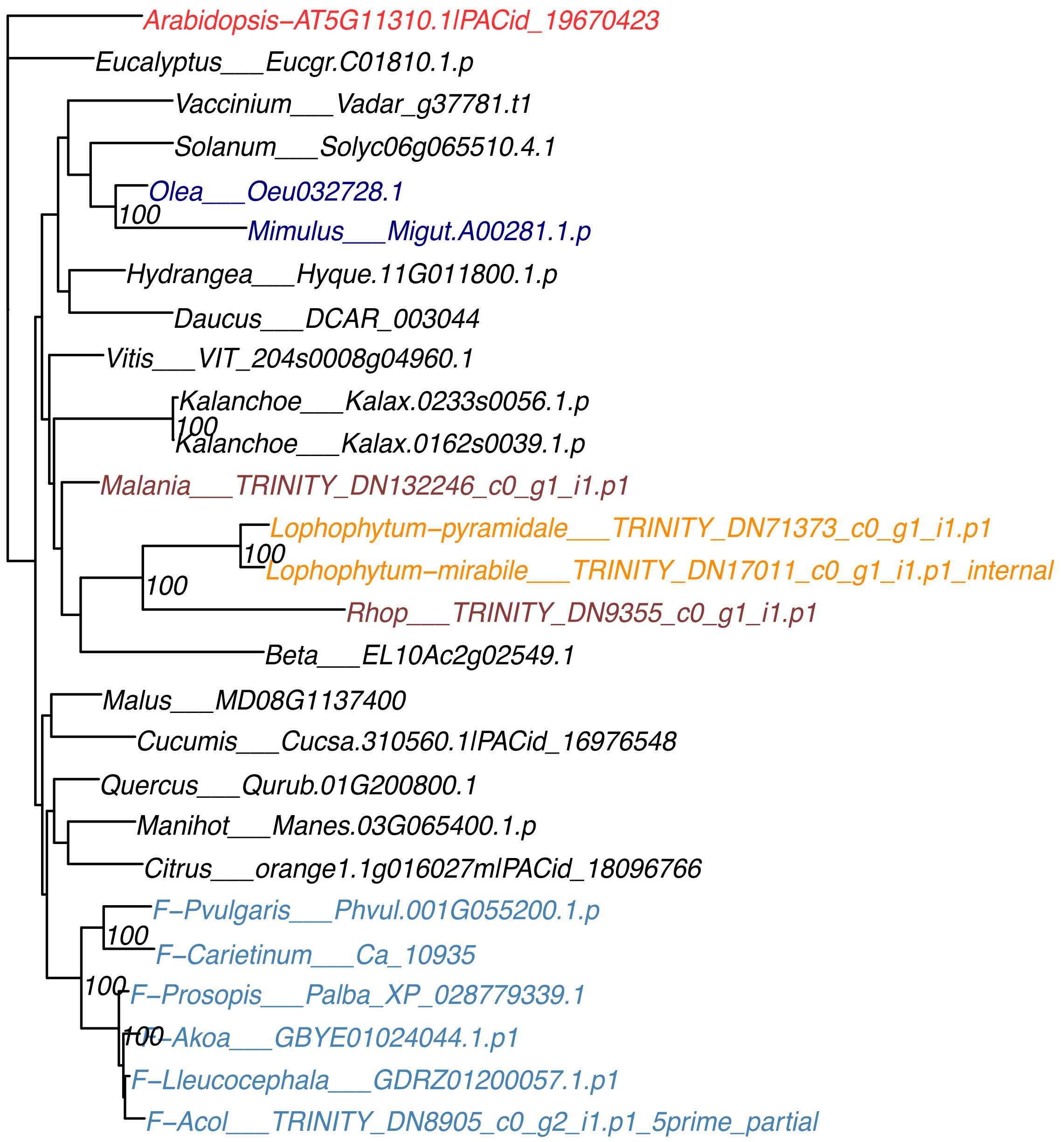
Arabidopsis−AT5G11310.1|PACid_19670423
Eucalyptus___Eucgr.C01810.1.p
Vaccinium___Vadar_g37781.t1
Solanum___Solyc06g065510.4.1
Olea___Oeu032728.1
100
Mimulus___Migut.A00281.1.p
Hydrangea___Hyque.11G011800.1.p
Daucus___DCAR_003044
Vitis___VIT_204s0008g04960.1
Kalanchoe___Kalax.0233s0056.1.p
100
Kalanchoe___Kalax.0162s0039.1.p
Malania___TRINITY_DN132246_c0_g1_i1.p1
Lophophytum−pyramidale___TRINITY_DN71373_c0_g1_i1.p1
100
Lophophytum−mirabile___TRINITY_DN17011_c0_g1_i1.p1_internal
100
Rhop___TRINITY_DN9355_c0_g1_i1.p1
Beta___EL10Ac2g02549.1
Malus___MD08G1137400
Cucumis___Cucsa.310560.1|PACid_16976548
Quercus___Qurub.01G200800.1
Manihot___Manes.03G065400.1.p
Citrus___orange1.1g016027m|PACid_18096766
F−Pvulgaris___Phvul.001G055200.1.p
100
F−Carietinum___Ca_10935
100
F−Prosopis___Palba_XP_028779339.1
100
F−Akoa___GBYE01024044.1.p1
F−Lleucocephala___GDRZ01200057.1.p1
F−Acol___TRINITY_DN8905_c0_g2_i1.p1_5prime_partial

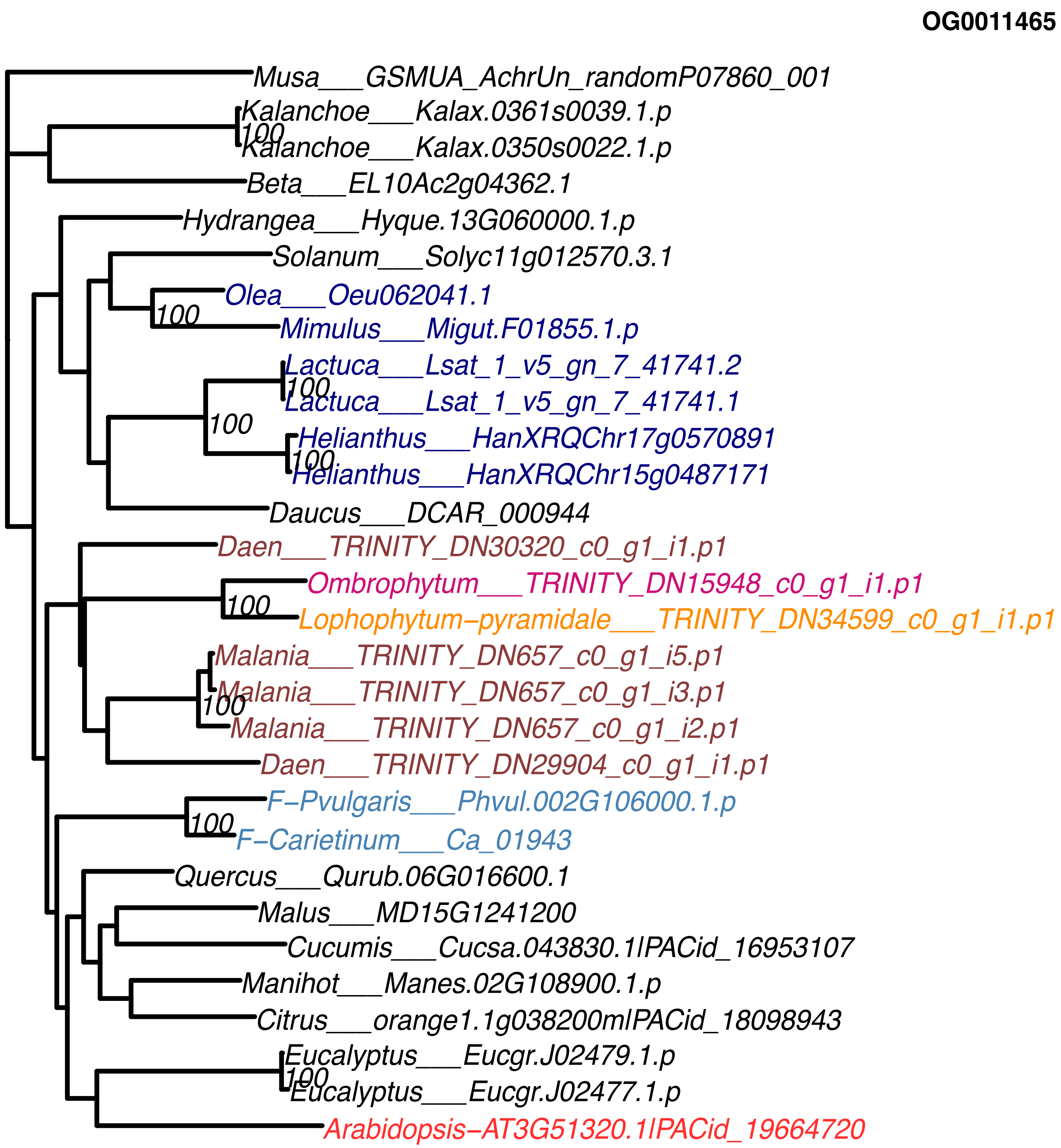
OG0011465
Musa___GSMUA_AchrUn_randomP07860_001
Kalanchoe___Kalax.0361s0039.1.p
100
Kalanchoe___Kalax.0350s0022.1.p
Beta___EL10Ac2g04362.1
Hydrangea___Hyque.13G060000.1.p
Solanum___Solyc11g012570.3.1
Olea___Oeu062041.1
100
Mimulus___Migut.F01855.1.p
Lactuca___Lsat_1_v5_gn_7_41741.2
100
Lactuca___Lsat_1_v5_gn_7_41741.1
100
Helianthus___HanXRQChr17g0570891
100
Helianthus___HanXRQChr15g0487171
Daucus___DCAR_000944
Daen___TRINITY_DN30320_c0_g1_i1.p1
Ombrophytum___TRINITY_DN15948_c0_g1_i1.p1
100
Lophophytum−pyramidale___TRINITY_DN34599_c0_g1_i1.p1
Malania___TRINITY_DN657_c0_g1_i5.p1
Malania___TRINITY_DN657_c0_g1_i3.p1
100
Malania___TRINITY_DN657_c0_g1_i2.p1
Daen___TRINITY_DN29904_c0_g1_i1.p1
F−Pvulgaris___Phvul.002G106000.1.p
100
F−Carietinum___Ca_01943
Quercus___Qurub.06G016600.1
Malus___MD15G1241200
Cucumis___Cucsa.043830.1|PACid_16953107
Manihot___Manes.02G108900.1.p
Citrus___orange1.1g038200m|PACid_18098943
Eucalyptus___Eucgr.J02479.1.p
100
Eucalyptus___Eucgr.J02477.1.p
Arabidopsis−AT3G51320.1|PACid_19664720

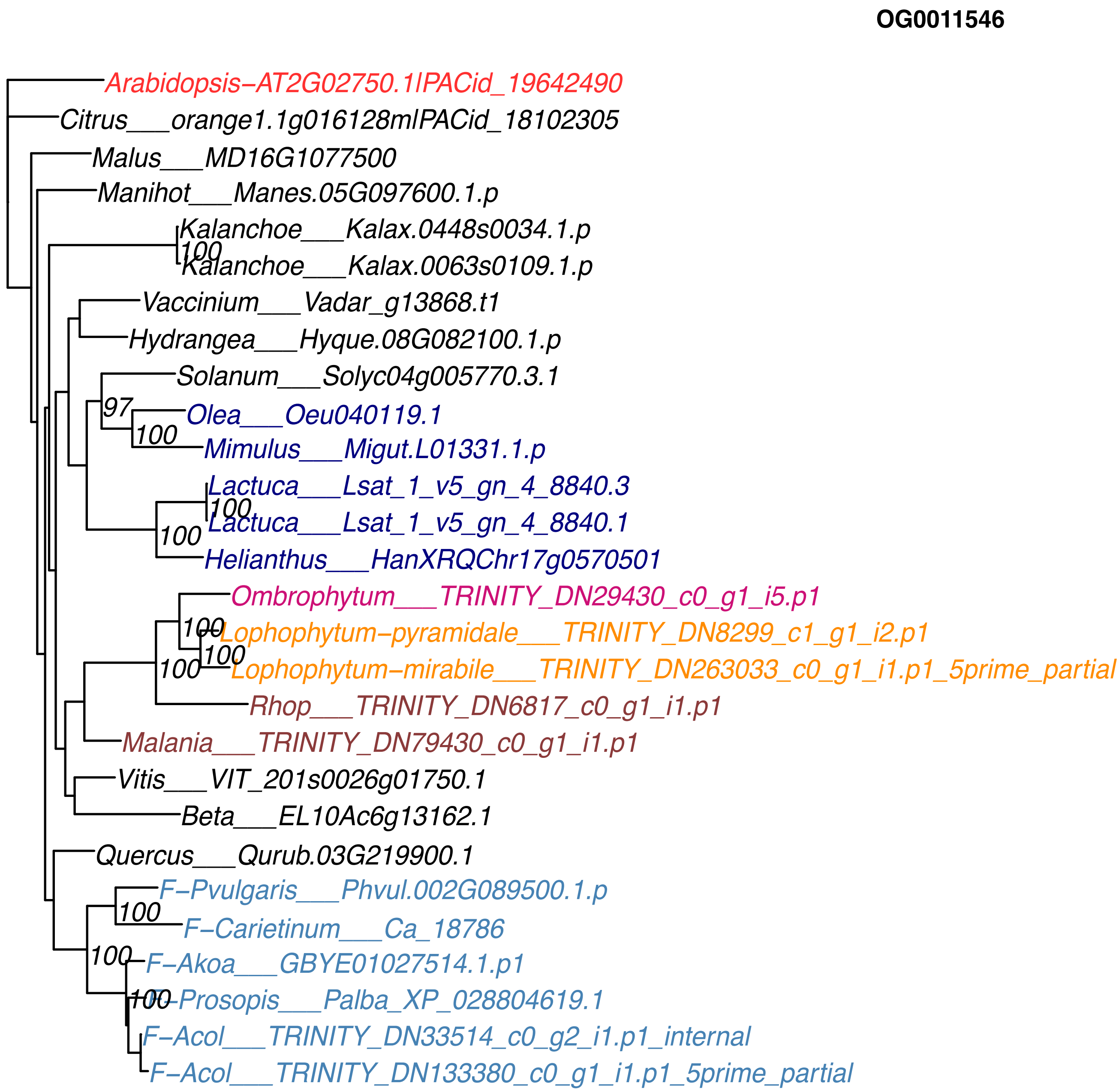
OG0011546
Arabidopsis−AT2G02750.1lPACid_19642490
Citrus___orange1.1g016128mlPACid_18102305
Malus___MD16G1077500
Manihot___Manes.05G097600.1.p
Kalanchoe___Kalax.0448s0034.1.p
100
Kalanchoe___Kalax.0063s0109.1.p
Vaccinium___Vadar_g13868.t1
Hydrangea___Hyque.08G082100.1.p
Solanum___Solyc04g005770.3.1
97
Olea___Oeu040119.1
100
Mimulus___Migut.L01331.1.p
Lactuca___Lsat_1_v5_gn_4_8840.3
100
Lactuca___Lsat_1_v5_gn_4_8840.1
100
Helianthus___HanXRQChr17g0570501
Ombrophytum___TRINITY_DN29430_c0_g1_i5.p1
100
Lophophytum−pyramidale___TRINITY_DN8299_c1_g1_i2.p1
100
100
Lophophytum−mirabile___TRINITY_DN263033_c0_g1_i1.p1_5prime_partial
Rhop___TRINITY_DN6817_c0_g1_i1.p1
Malania___TRINITY_DN79430_c0_g1_i1.p1
Vitis___VIT_201s0026g01750.1
Beta___EL10Ac6g13162.1
Quercus___Qurub.03G219900.1
F−Pvulgaris___Phvul.002G089500.1.p
100
F−Carietinum___Ca_18786
100
F−Akoa___GBYE01027514.1.p1
100
F−Prosopis___Palba_XP_028804619.1
F−Acol___TRINITY_DN33514_c0_g2_i1.p1_internal
F−Acol___TRINITY_DN133380_c0_g1_i1.p1_5prime_partial

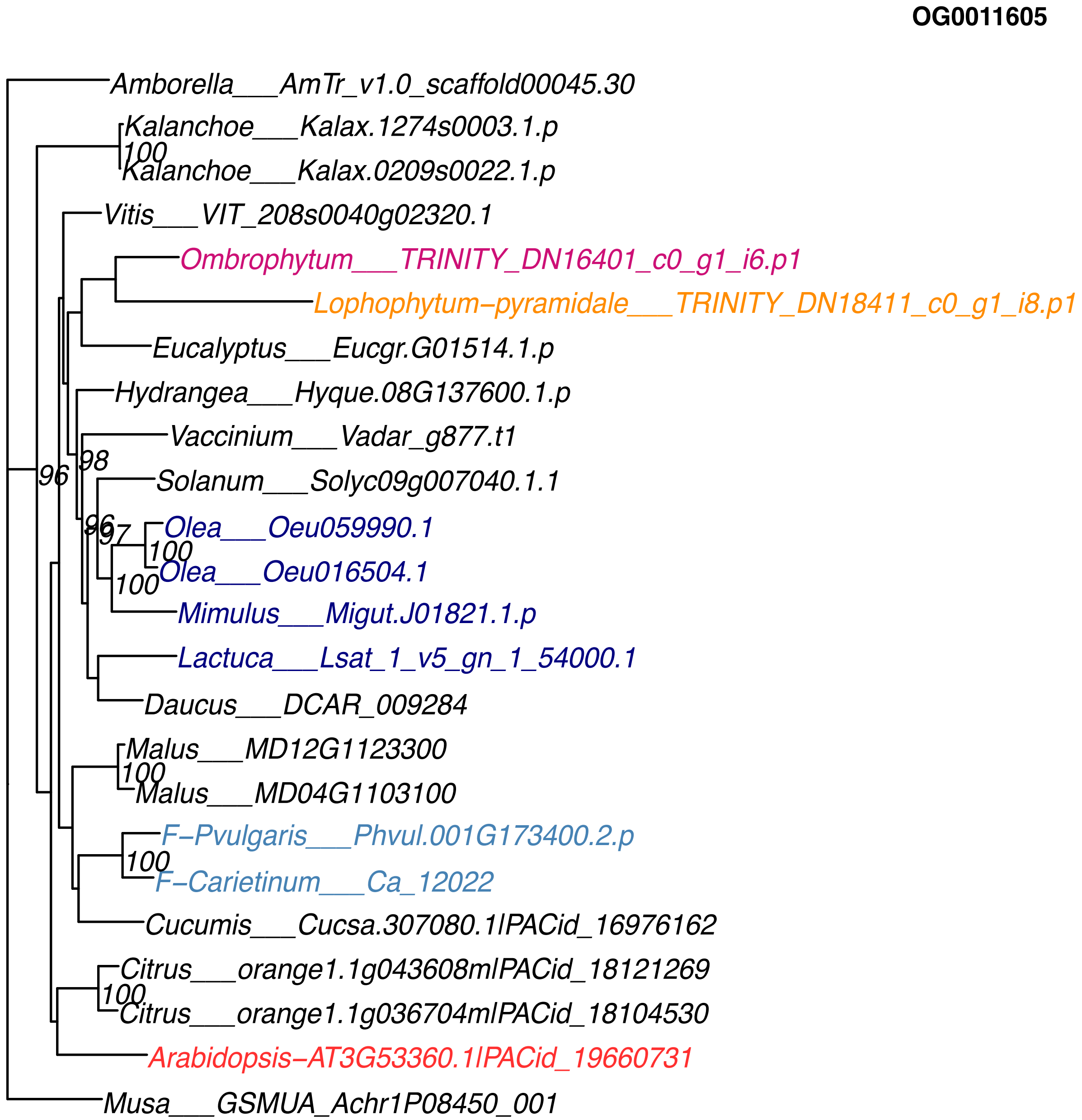
OG0011605
Amborella___AmTr_v1.0_scaffold00045.30
Kalanchoe___Kalax.1274s0003.1.p
100
Kalanchoe___Kalax.0209s0022.1.p
Vitis___VIT_208s0040g02320.1
Ombrophytum___TRINITY_DN16401_c0_g1_i6.p1
Lophophytum−pyramidale___TRINITY_DN18411_c0_g1_i8.p1
Eucalyptus___Eucgr.G01514.1.p
Hydrangea___Hyque.08G137600.1.p
Vaccinium___Vadar_g877.t1
98
96
Solanum___Solyc09g007040.1.1
96
97
Olea___Oeu059990.1
100
Olea___Oeu016504.1
100
Mimulus___Migut.J01821.1.p
Lactuca___Lsat_1_v5_gn_1_54000.1
Daucus___DCAR_009284
Malus___MD12G1123300
100
Malus___MD04G1103100
F−Pvulgaris___Phvul.001G173400.2.p
100
F−Carietinum___Ca_12022
Cucumis___Cucsa.307080.1|PACid_16976162
Citrus___orange1.1g043608m|PACid_18121269
100
Citrus___orange1.1g036704m|PACid_18104530
Arabidopsis−AT3G53360.1|PACid_19660731
Musa___GSMUA_Achr1P08450_001

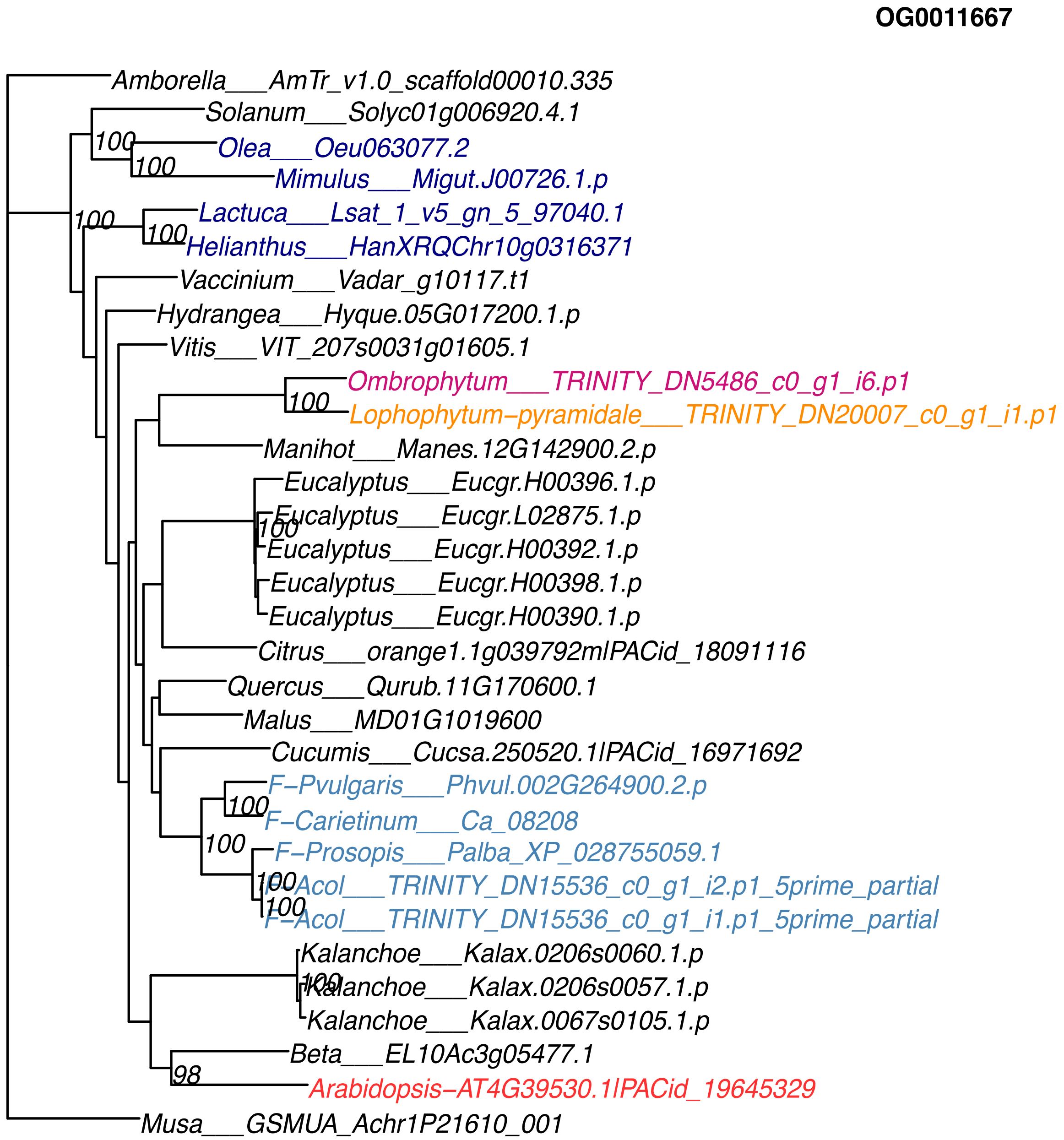
OG0011667
Amborella___AmTr_v1.0_scaffold00010.335
Solanum___Solyc01g006920.4.1
100
Olea___Oeu063077.2
100
Mimulus___Migut.J00726.1.p
Lactuca___Lsat_1_v5_gn_5_97040.1
100
100
Helianthus___HanXRQChr10g0316371
Vaccinium___Vadar_g10117.t1
Hydrangea___Hyque.05G017200.1.p
Vitis___VIT_207s0031g01605.1
Ombrophytum___TRINITY_DN5486_c0_g1_i6.p1
100
Lophophytum−pyramidale___TRINITY_DN20007_c0_g1_i1.p1
Manihot___Manes.12G142900.2.p
Eucalyptus___Eucgr.H00396.1.p
Eucalyptus___Eucgr.L02875.1.p
100
Eucalyptus___Eucgr.H00392.1.p
Eucalyptus___Eucgr.H00398.1.p
Eucalyptus___Eucgr.H00390.1.p
Citrus___orange1.1g039792mIPACid_18091116
Quercus___Qurub.11G170600.1
Malus___MD01G1019600
Cucumis___Cucsa.250520.1IPACid_16971692
F−Pvulgaris___Phvul.002G264900.2.p
100
F−Carietinum___Ca_08208
100
F−Prosopis___Palba_XP_028755059.1
100
F−Acol___TRINITY_DN15536_c0_g1_i2.p1_5prime_partial
100
F−Acol___TRINITY_DN15536_c0_g1_i1.p1_5prime_partial
Kalanchoe___Kalax.0206s0060.1.p
100
Kalanchoe___Kalax.0206s0057.1.p
Kalanchoe___Kalax.0067s0105.1.p
Beta___EL10Ac3g05477.1
98
Arabidopsis−AT4G39530.1IPACid_19645329
Musa___GSMUA_Achr1P21610_001

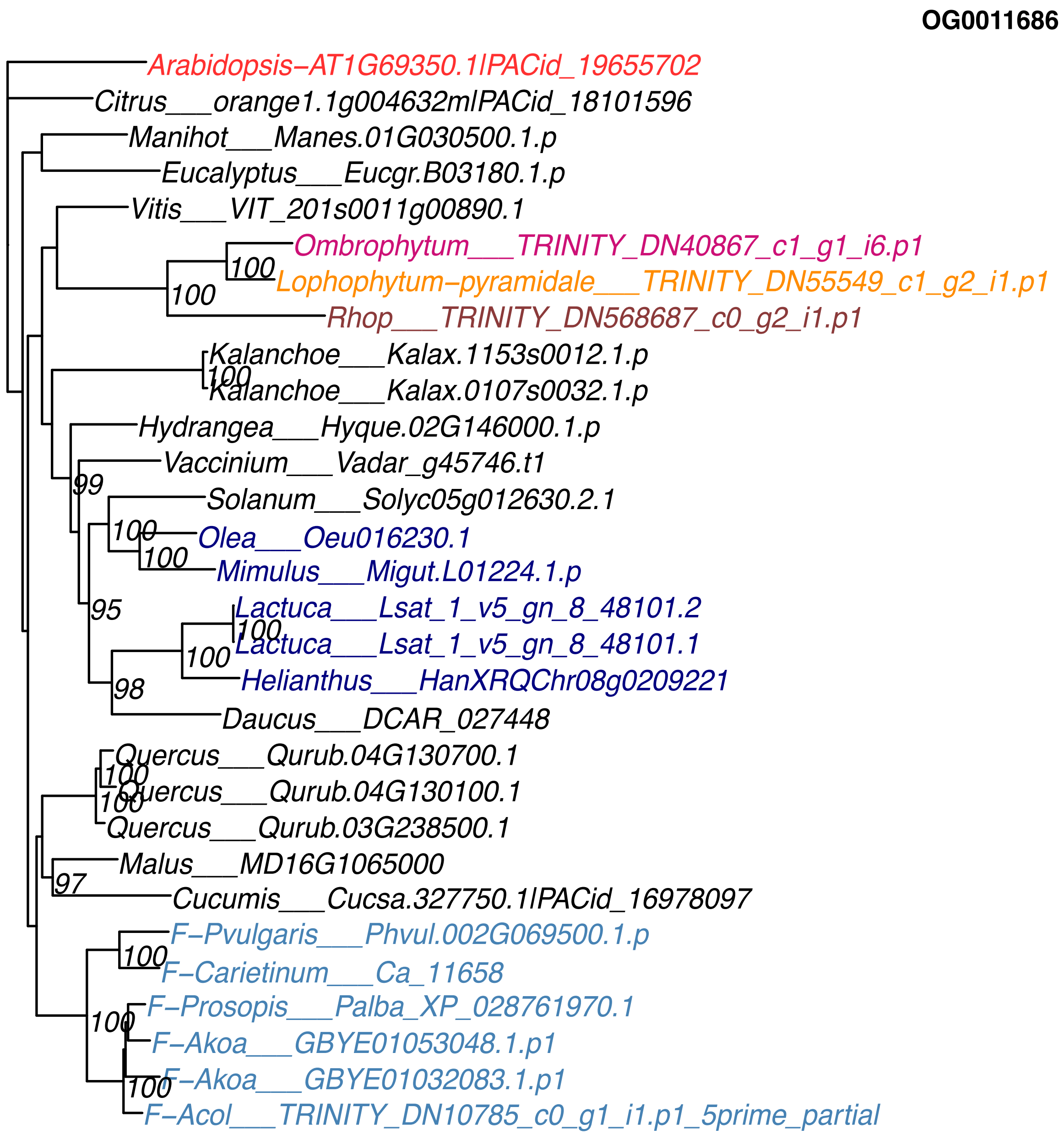
OG0011686
Arabidopsis–AT1G69350.1|PACid_19655702
Citrus___orange1.1g004632m|PACid_18101596
Manihot___Manes.01G030500.1.p
Eucalyptus___Eucgr.B03180.1.p
Vitis___VIT_201s0011g00890.1
Ombrophytum___TRINITY_DN40867_c1_g1_i6.p1
Lophophytum–pyramidale___TRINITY_DN55549_c1_g2_i1.p1
Rhop___TRINITY_DN568687_c0_g2_i1.p1
Kalanchoe___Kalax.1153s0012.1.p
Kalanchoe___Kalax.0107s0032.1.p
Hydrangea___Hyque.02G146000.1.p
Vaccinium___Vadar_g45746.t1
Solanum___Solyc05g012630.2.1
Olea___Oeu016230.1
Mimulus___Migut.L01224.1.p
Lactuca___Lsat_1_v5_gn_8_48101.2
Lactuca___Lsat_1_v5_gn_8_48101.1
Helianthus___HanXRQChr08g0209221
Daucus___DCAR_027448
Quercus___Qurub.04G130700.1
Quercus___Qurub.04G130100.1
Quercus___Qurub.03G238500.1
Malus___MD16G1065000
Cucumis___Cucsa.327750.1|PACid_16978097
F–Pvulgaris___Phvul.002G069500.1.p
F–Carietinum___Ca_11658
F–Prosopis___Palba_XP_028761970.1
F–Akoa___GBYE01053048.1.p1
F–Akoa___GBYE01032083.1.p1
F–Acol___TRINITY_DN10785_c0_g1_i1.p1_5prime_partial
100
100
100
99
100
100
95
100
100
98
100
100
97
100
100
100

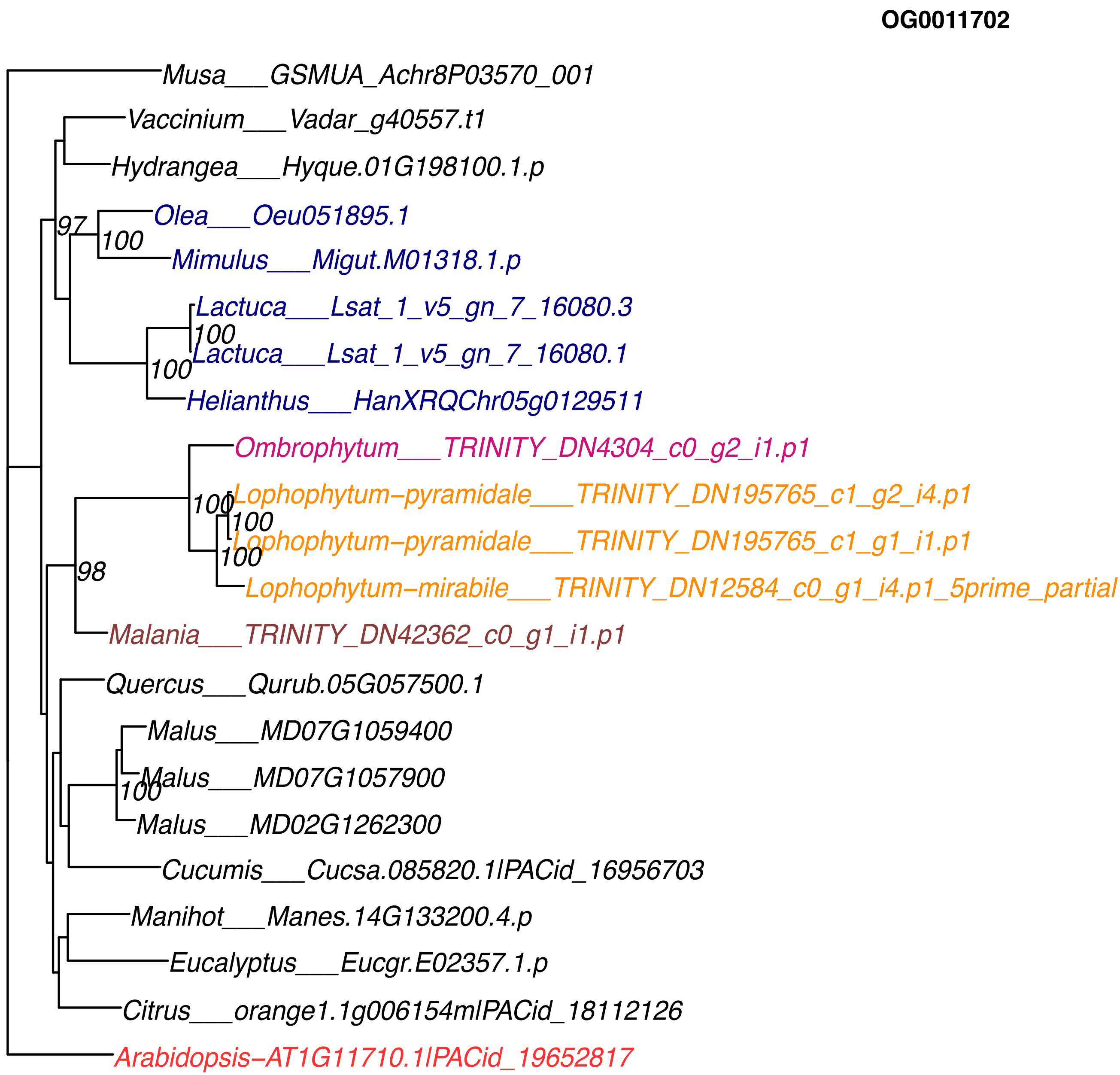

OG0011702
Musa___GSMUA_Achr8P03570_001
Vaccinium___Vadar_g40557.t1
Hydrangea___Hyque.01G198100.1.p
Olea___Oeu051895.1
Mimulus___Migut.M01318.1.p
Lactuca___Lsat_1_v5_gn_7_16080.3
Lactuca___Lsat_1_v5_gn_7_16080.1
Helianthus___HanXRQChr05g0129511
Ombrophytum___TRINITY_DN4304_c0_g2_i1.p1
Lophophytum−pyramidale___TRINITY_DN195765_c1_g2_i4.p1
Lophophytum−pyramidale___TRINITY_DN195765_c1_g1_i1.p1
Lophophytum−mirabile___TRINITY_DN12584_c0_g1_i4.p1_5prime_partial
Malania___TRINITY_DN42362_c0_g1_i1.p1
Quercus___Qurub.05G057500.1
Malus___MD07G1059400
Malus___MD07G1057900
Malus___MD02G1262300
Cucumis___Cucsa.085820.1|PACid_16956703
Manihot___Manes.14G133200.4.p
Eucalyptus___Eucgr.E02357.1.p
Citrus___orange1.1g006154m|PACid_18112126
Arabidopsis−AT1G11710.1|PACid_19652817
97
100
100
100
100
100
100
98
100

**OG0011826**

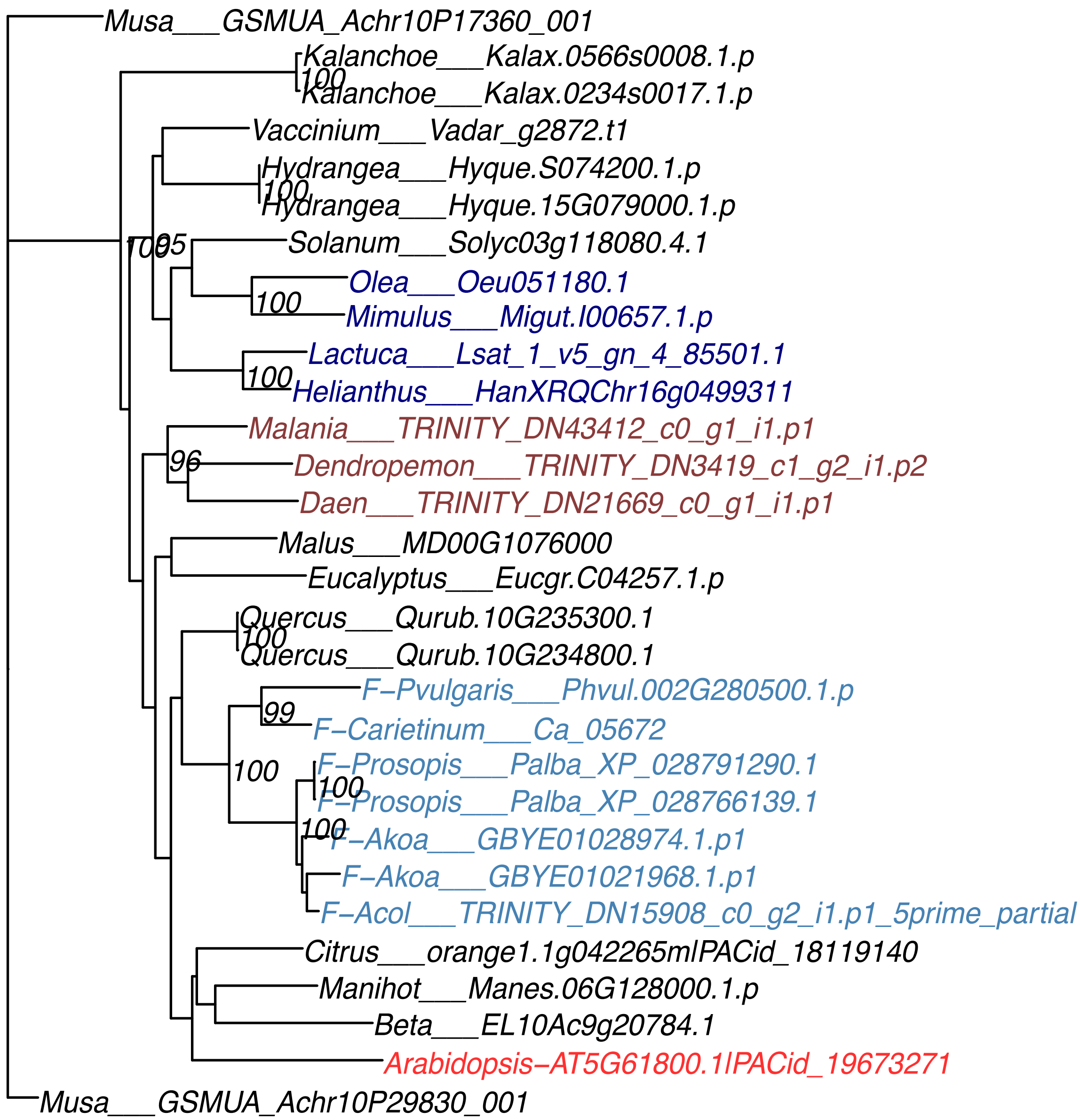

Musa___GSMUA_Achr10P17360_001
Kalanchoe___Kalax.0566s0008.1.p
100
Kalanchoe___Kalax.0234s0017.1.p
Vaccinium___Vadar_g2872.t1
Hydrangea___Hyque.S074200.1.p
100
Hydrangea___Hyque.15G079000.1.p
100
95
Solanum___Solyc03g118080.4.1
Olea___Oeu051180.1
100
Mimulus___Migut.I00657.1.p
Lactuca___Lsat_1_v5_gn_4_85501.1
100
Helianthus___HanXRQChr16g0499311
Malania___TRINITY_DN43412_c0_g1_i1.p1
96
Dendropemon___TRINITY_DN3419_c1_g2_i1.p2
Daen___TRINITY_DN21669_c0_g1_i1.p1
Malus___MD00G1076000
Eucalyptus___Eucgr.C04257.1.p
Quercus___Qurub.10G235300.1
100
Quercus___Qurub.10G234800.1
F−Pvulgaris___Phvul.002G280500.1.p
99
F−Carietinum___Ca_05672
100
F−Prosopis___Palba_XP_028791290.1
100
F−Prosopis___Palba_XP_028766139.1
100
F−Akoa___GBYE01028974.1.p1
F−Akoa___GBYE01021968.1.p1
F−Acol___TRINITY_DN15908_c0_g2_i1.p1_5prime_partial
Citrus___orange1.1g042265mIPACid_18119140
Manihot___Manes.06G128000.1.p
Beta___EL10Ac9g20784.1
Arabidopsis−AT5G61800.1IPACid_19673271
Musa___GSMUA_Achr10P29830_001

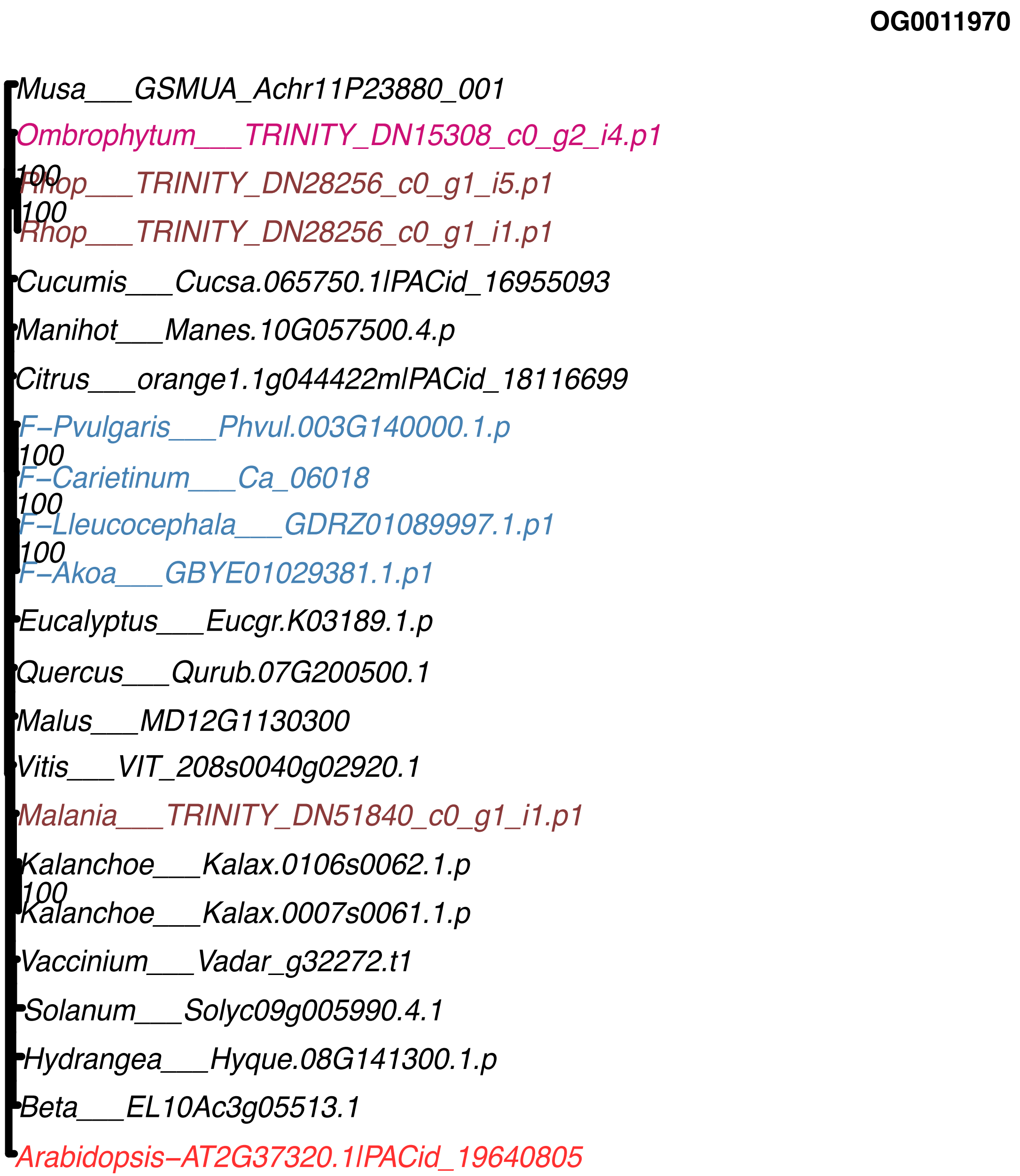
OG0011970
Musa___GSMUA_Achr11P23880_001
Ombrophytum___TRINITY_DN15308_c0_g2_i4.p1
100
Rhop___TRINITY_DN28256_c0_g1_i5.p1
100
Rhop___TRINITY_DN28256_c0_g1_i1.p1
Cucumis___Cucsa.065750.1|PACid_16955093
Manihot___Manes.10G057500.4.p
Citrus___orange1.1g044422m|PACid_18116699
F–Pvulgaris___Phvul.003G140000.1.p
100
F–Carietinum___Ca_06018
100
F–Lleucocephala___GDRZ01089997.1.p1
100
F–Akoa___GBYE01029381.1.p1
Eucalyptus___Eucgr.K03189.1.p
Quercus___Qurub.07G200500.1
Malus___MD12G1130300
Vitis___VIT_208s0040g02920.1
Malania___TRINITY_DN51840_c0_g1_i1.p1
Kalanchoe___Kalax.0106s0062.1.p
100
Kalanchoe___Kalax.0007s0061.1.p
Vaccinium___Vadar_g32272.t1
Solanum___Solyc09g005990.4.1
Hydrangea___Hyque.08G141300.1.p
Beta___EL10Ac3g05513.1
Arabidopsis–AT2G37320.1|PACid_19640805

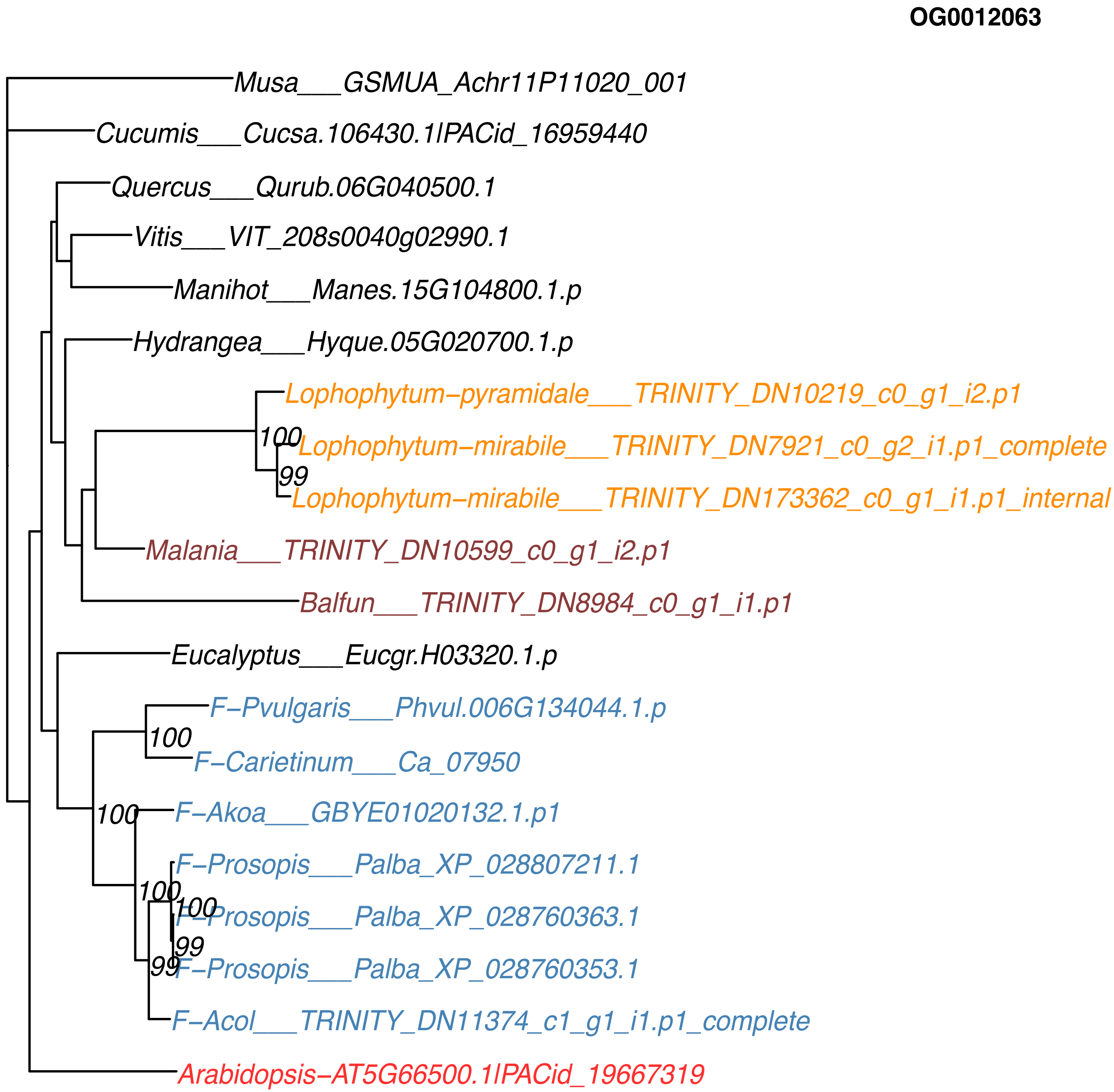
OG0012063
Musa___GSMUA_Achr11P11020_001
Cucumis___Cucsa.106430.1lPACid_16959440
Quercus___Qurub.06G040500.1
Vitis___VIT_208s0040g02990.1
Manihot___Manes.15G104800.1.p
Hydrangea___Hyque.05G020700.1.p
Lophophytum−pyramidale___TRINITY_DN10219_c0_g1_i2.p1
100
Lophophytum−mirabile___TRINITY_DN7921_c0_g2_i1.p1_complete
99
Lophophytum−mirabile___TRINITY_DN173362_c0_g1_i1.p1_internal
Malania___TRINITY_DN10599_c0_g1_i2.p1
Balfun___TRINITY_DN8984_c0_g1_i1.p1
Eucalyptus___Eucgr.H03320.1.p
F−Pvulgaris___Phvul.006G134044.1.p
100
F−Carietinum___Ca_07950
100
F−Akoa___GBYE01020132.1.p1
F−Prosopis___Palba_XP_028807211.1
100
100
F−Prosopis___Palba_XP_028760363.1
99
99
F−Prosopis___Palba_XP_028760353.1
F−Acol___TRINITY_DN11374_c1_g1_i1.p1_complete
Arabidopsis−AT5G66500.1lPACid_19667319

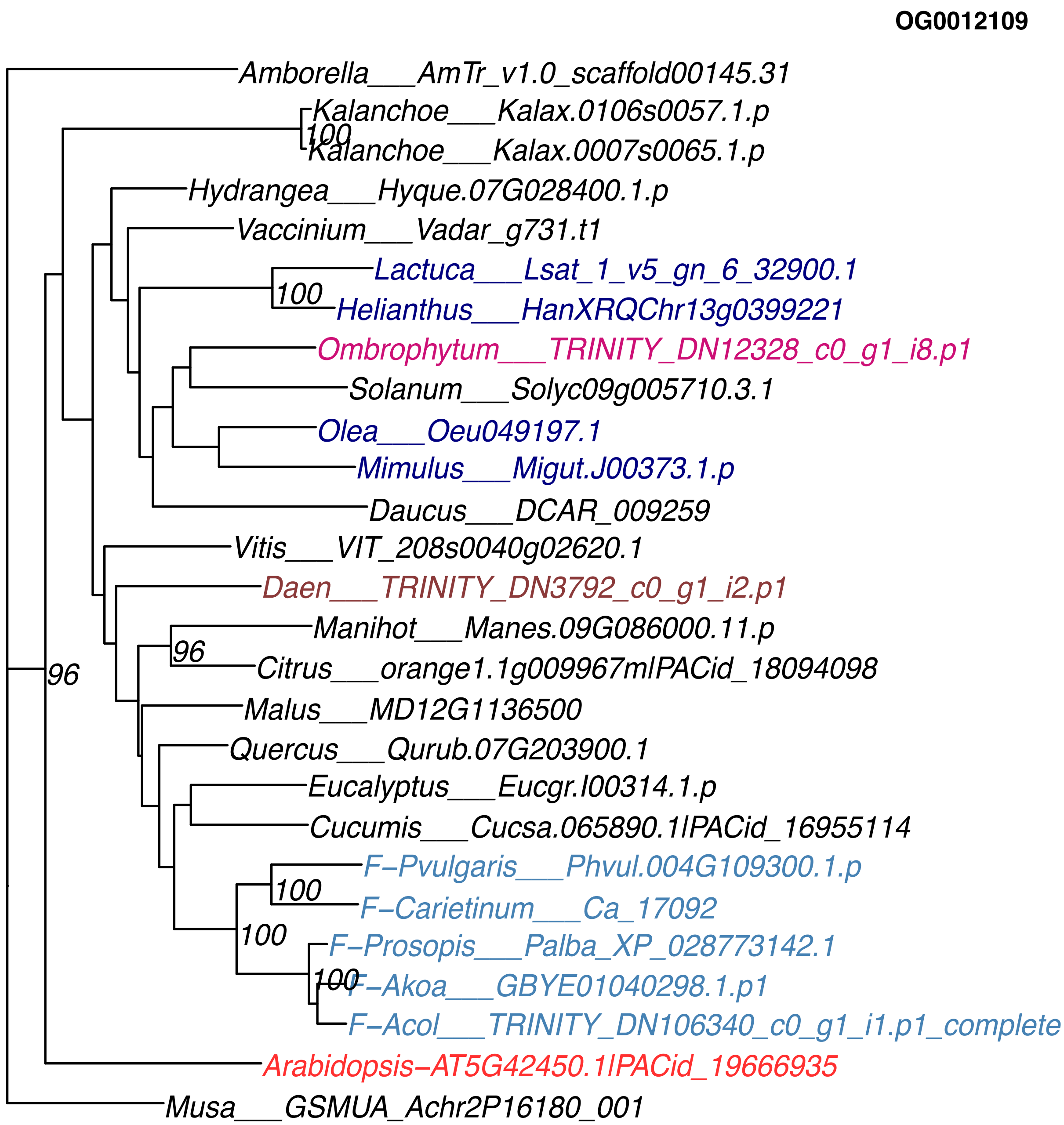

OG0012109
Amborella___AmTr_v1.0_scaffold00145.31
Kalanchoe___Kalax.0106s0057.1.p
100
Kalanchoe___Kalax.0007s0065.1.p
Hydrangea___Hyque.07G028400.1.p
Vaccinium___Vadar_g731.t1
Lactuca___Lsat_1_v5_gn_6_32900.1
100
Helianthus___HanXRQChr13g0399221
Ombrophytum___TRINITY_DN12328_c0_g1_i8.p1
Solanum___Solyc09g005710.3.1
Olea___Oeu049197.1
Mimulus___Migut.J00373.1.p
Daucus___DCAR_009259
Vitis___VIT_208s0040g02620.1
Daen___TRINITY_DN3792_c0_g1_i2.p1
Manihot___Manes.09G086000.11.p
96
Citrus___orange1.1g009967m|PACid_18094098
96
Malus___MD12G1136500
Quercus___Qurub.07G203900.1
Eucalyptus___Eucgr.I00314.1.p
Cucumis___Cucsa.065890.1|PACid_16955114
F−Pvulgaris___Phvul.004G109300.1.p
100
F−Carietinum___Ca_17092
100
F−Prosopis___Palba_XP_028773142.1
100
F−Akoa___GBYE01040298.1.p1
F−Acol___TRINITY_DN106340_c0_g1_i1.p1_complete
Arabidopsis−AT5G42450.1|PACid_19666935
Musa___GSMUA_Achr2P16180_001

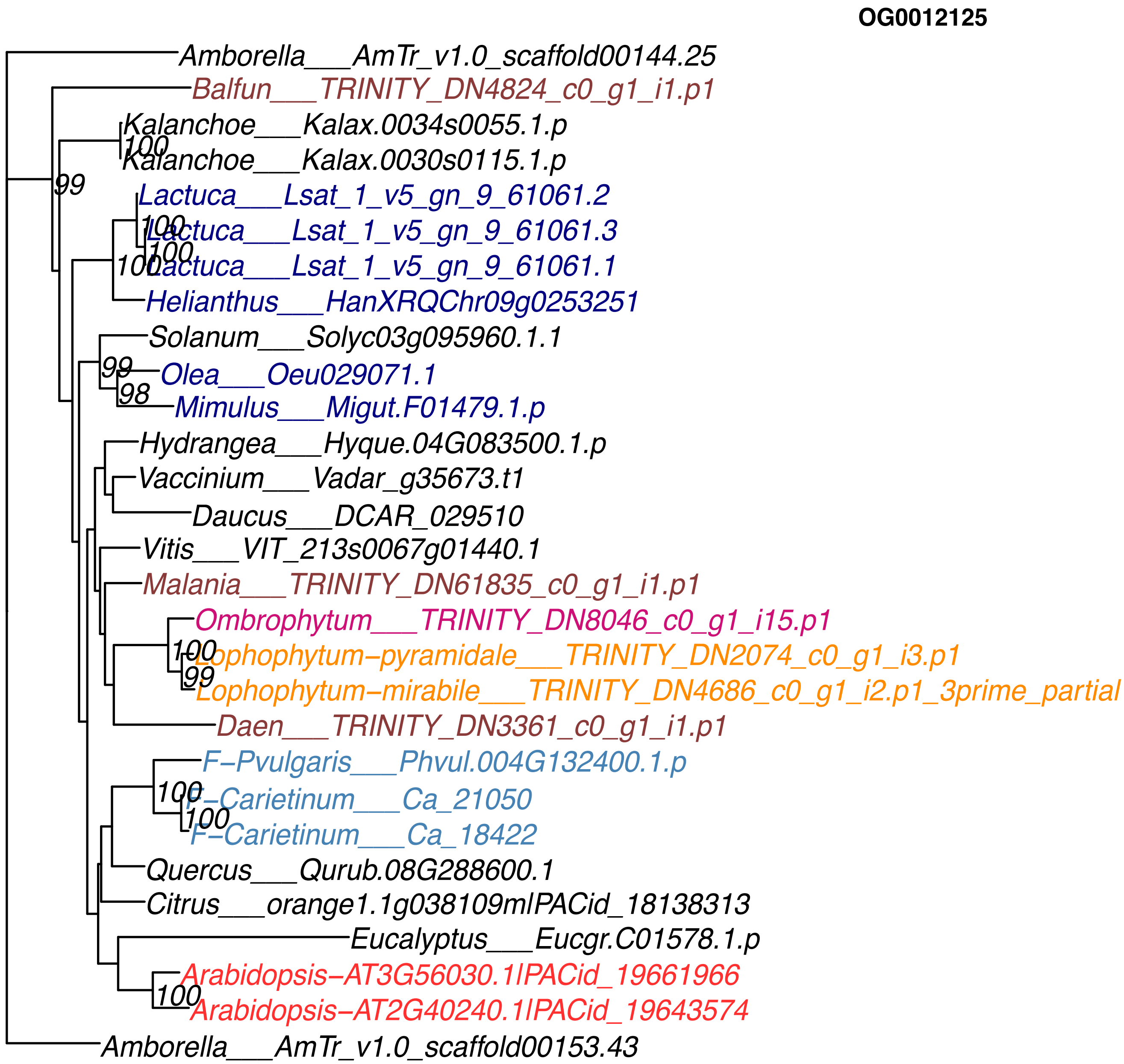
OG0012125
Amborella___AmTr_v1.0_scaffold00144.25
Balfun___TRINITY_DN4824_c0_g1_i1.p1
Kalanchoe___Kalax.0034s0055.1.p
100
Kalanchoe___Kalax.0030s0115.1.p
99
Lactuca___Lsat_1_v5_gn_9_61061.2
100
Lactuca___Lsat_1_v5_gn_9_61061.3
100
100
Lactuca___Lsat_1_v5_gn_9_61061.1
Helianthus___HanXRQChr09g0253251
Solanum___Solyc03g095960.1.1
99
Olea___Oeu029071.1
98
Mimulus___Migut.F01479.1.p
Hydrangea___Hyque.04G083500.1.p
Vaccinium___Vadar_g35673.t1
Daucus___DCAR_029510
Vitis___VIT_213s0067g01440.1
Malania___TRINITY_DN61835_c0_g1_i1.p1
Ombrophytum___TRINITY_DN8046_c0_g1_i15.p1
100
Lophophytum−pyramidale___TRINITY_DN2074_c0_g1_i3.p1
99
Lophophytum−mirabile___TRINITY_DN4686_c0_g1_i2.p1_3prime_partial
Daen___TRINITY_DN3361_c0_g1_i1.p1
F−Pvulgaris___Phvul.004G132400.1.p
100
F−Carietinum___Ca_21050
100
F−Carietinum___Ca_18422
Quercus___Qurub.08G288600.1
Citrus___orange1.1g038109m|PACid_18138313
Eucalyptus___Eucgr.C01578.1.p
Arabidopsis−AT3G56030.1|PACid_19661966
100
Arabidopsis−AT2G40240.1|PACid_19643574
Amborella___AmTr_v1.0_scaffold00153.43

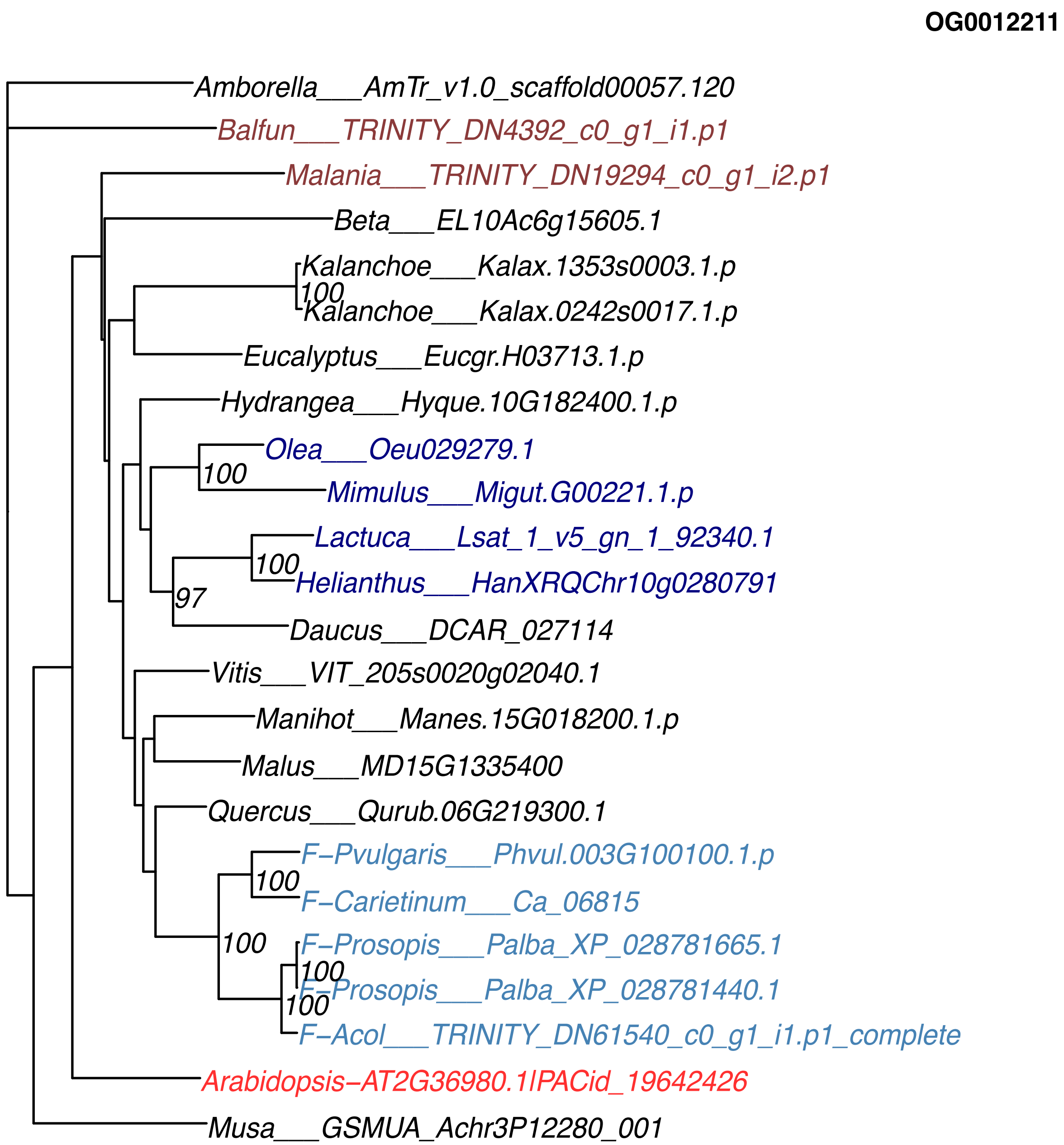
OG0012211
Amborella___AmTr_v1.0_scaffold00057.120
Balfun___TRINITY_DN4392_c0_g1_i1.p1
Malania___TRINITY_DN19294_c0_g1_i2.p1
Beta___EL10Ac6g15605.1
Kalanchoe___Kalax.1353s0003.1.p
100
Kalanchoe___Kalax.0242s0017.1.p
Eucalyptus___Eucgr.H03713.1.p
Hydrangea___Hyque.10G182400.1.p
Olea___Oeu029279.1
100
Mimulus___Migut.G00221.1.p
Lactuca___Lsat_1_v5_gn_1_92340.1
100
Helianthus___HanXRQChr10g0280791
97
Daucus___DCAR_027114
Vitis___VIT_205s0020g02040.1
Manihot___Manes.15G018200.1.p
Malus___MD15G1335400
Quercus___Qurub.06G219300.1
F−Pvulgaris___Phvul.003G100100.1.p
100
F−Carietinum___Ca_06815
100
F−Prosopis___Palba_XP_028781665.1
100
F−Prosopis___Palba_XP_028781440.1
100
F−Acol___TRINITY_DN61540_c0_g1_i1.p1_complete
Arabidopsis−AT2G36980.1|PACid_19642426
Musa___GSMUA_Achr3P12280_001

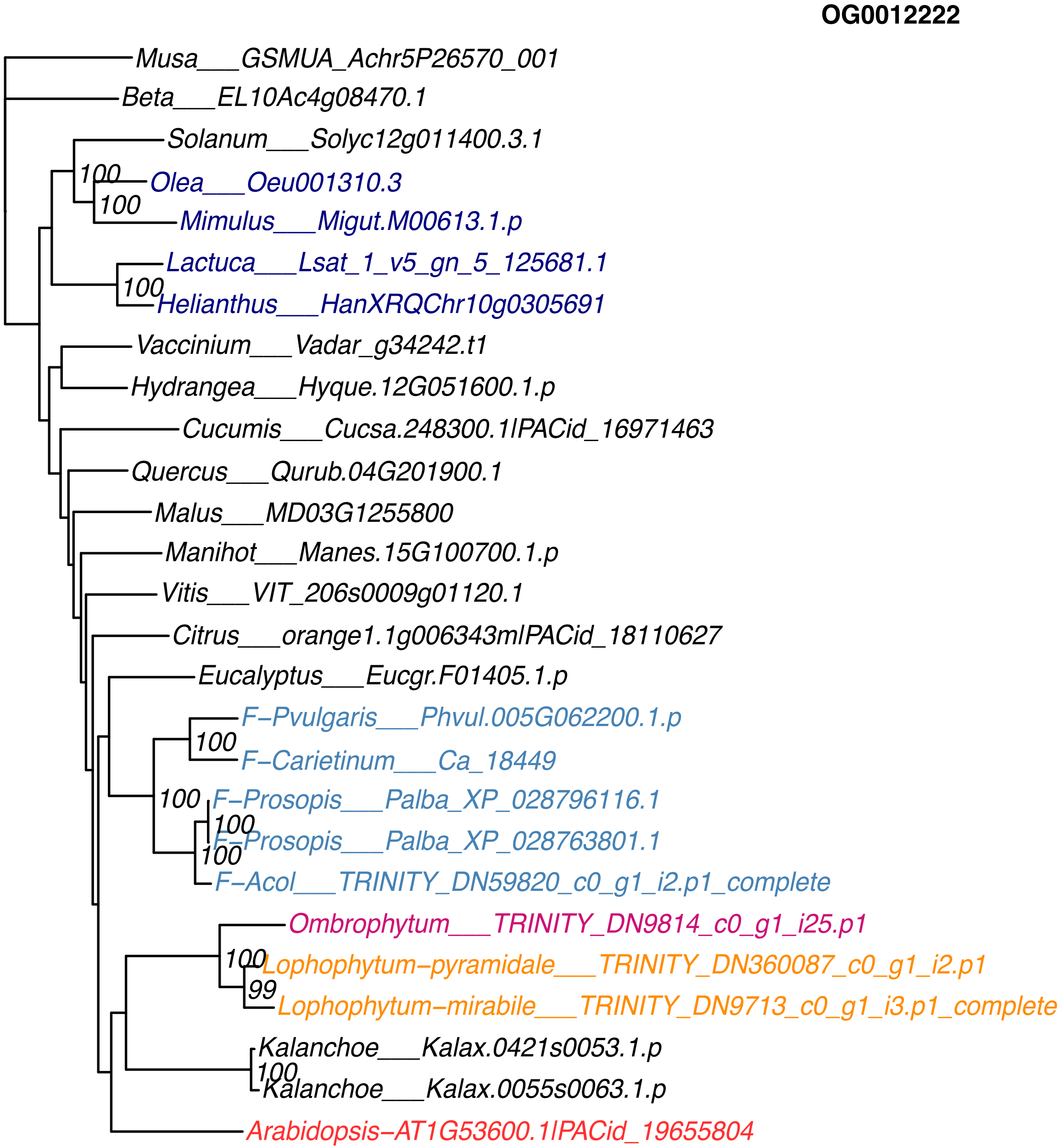
OG0012222
Musa___GSMUA_Achr5P26570_001
Beta___EL10Ac4g08470.1
Solanum___Solyc12g011400.3.1
100
Olea___Oeu001310.3
100
Mimulus___Migut.M00613.1.p
Lactuca___Lsat_1_v5_gn_5_125681.1
100
Helianthus___HanXRQChr10g0305691
Vaccinium___Vadar_g34242.t1
Hydrangea___Hyque.12G051600.1.p
Cucumis___Cucsa.248300.1|PACid_16971463
Quercus___Qurub.04G201900.1
Malus___MD03G1255800
Manihot___Manes.15G100700.1.p
Vitis___VIT_206s0009g01120.1
Citrus___orange1.1g006343m|PACid_18110627
Eucalyptus___Eucgr.F01405.1.p
F–Pvulgaris___Phvul.005G062200.1.p
100
F–Carietinum___Ca_18449
100
F–Prosopis___Palba_XP_028796116.1
100
F–Prosopis___Palba_XP_028763801.1
100
F–Acol___TRINITY_DN59820_c0_g1_i2.p1_complete
Ombrophytum___TRINITY_DN9814_c0_g1_i25.p1
100
Lophophytum–pyramidale___TRINITY_DN360087_c0_g1_i2.p1
99
Lophophytum–mirabile___TRINITY_DN9713_c0_g1_i3.p1_complete
Kalanchoe___Kalax.0421s0053.1.p
100
Kalanchoe___Kalax.0055s0063.1.p
Arabidopsis–AT1G53600.1|PACid_19655804

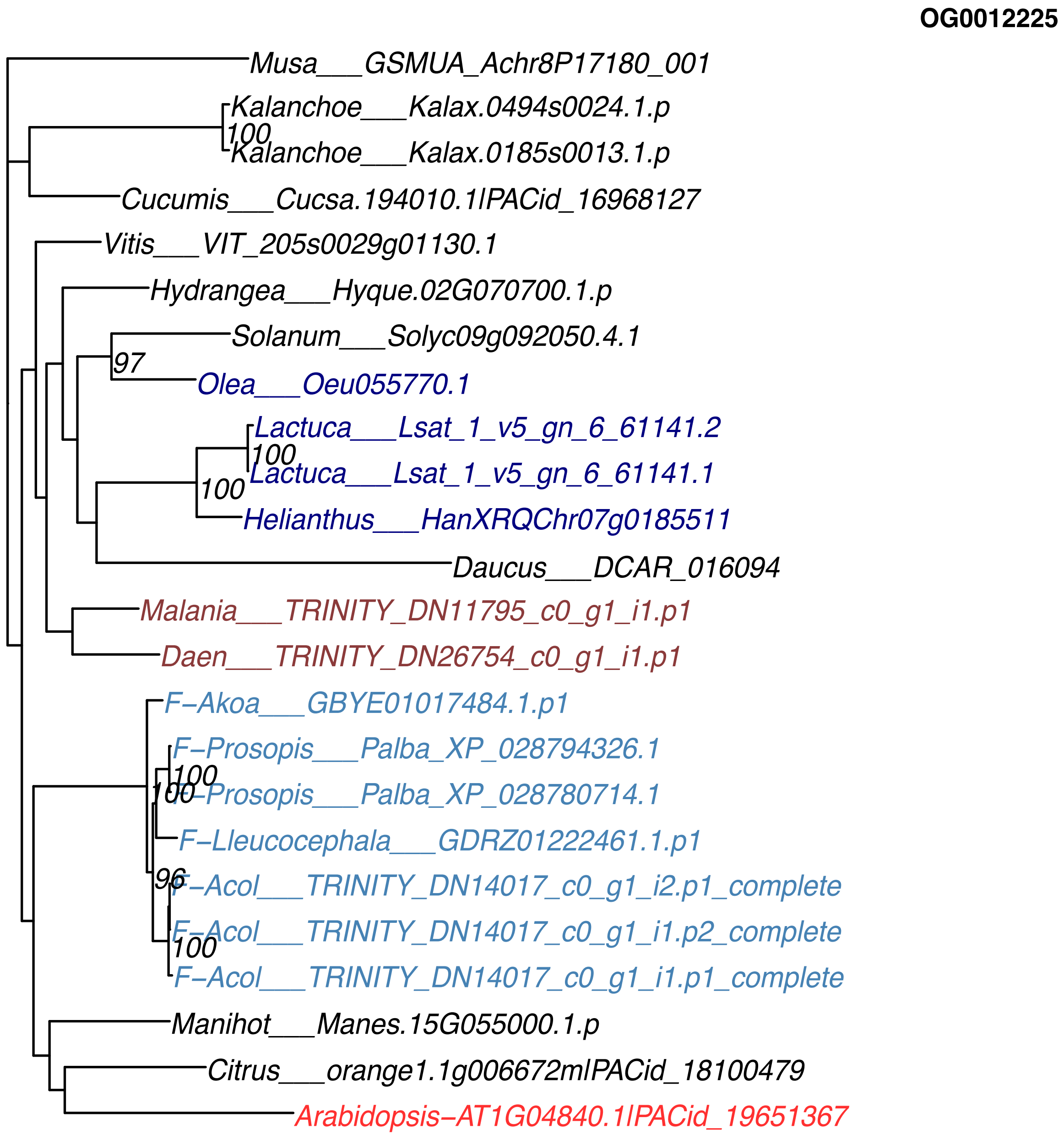
OG0012225
Musa___GSMUA_Achr8P17180_001
Kalanchoe___Kalax.0494s0024.1.p
100
Kalanchoe___Kalax.0185s0013.1.p
Cucumis___Cucsa.194010.1lPACid_16968127
Vitis___VIT_205s0029g01130.1
Hydrangea___Hyque.02G070700.1.p
Solanum___Solyc09g092050.4.1
97
Olea___Oeu055770.1
Lactuca___Lsat_1_v5_gn_6_61141.2
100
Lactuca___Lsat_1_v5_gn_6_61141.1
100
Helianthus___HanXRQChr07g0185511
Daucus___DCAR_016094
Malania___TRINITY_DN11795_c0_g1_i1.p1
Daen___TRINITY_DN26754_c0_g1_i1.p1
F−Akoa___GBYE01017484.1.p1
F−Prosopis___Palba_XP_028794326.1
100
100
F−Prosopis___Palba_XP_028780714.1
F−Lleucocephala___GDRZ01222461.1.p1
96
F−Acol___TRINITY_DN14017_c0_g1_i2.p1_complete
F−Acol___TRINITY_DN14017_c0_g1_i1.p2_complete
100
F−Acol___TRINITY_DN14017_c0_g1_i1.p1_complete
Manihot___Manes.15G055000.1.p
Citrus___orange1.1g006672mlPACid_18100479
Arabidopsis−AT1G04840.1lPACid_19651367

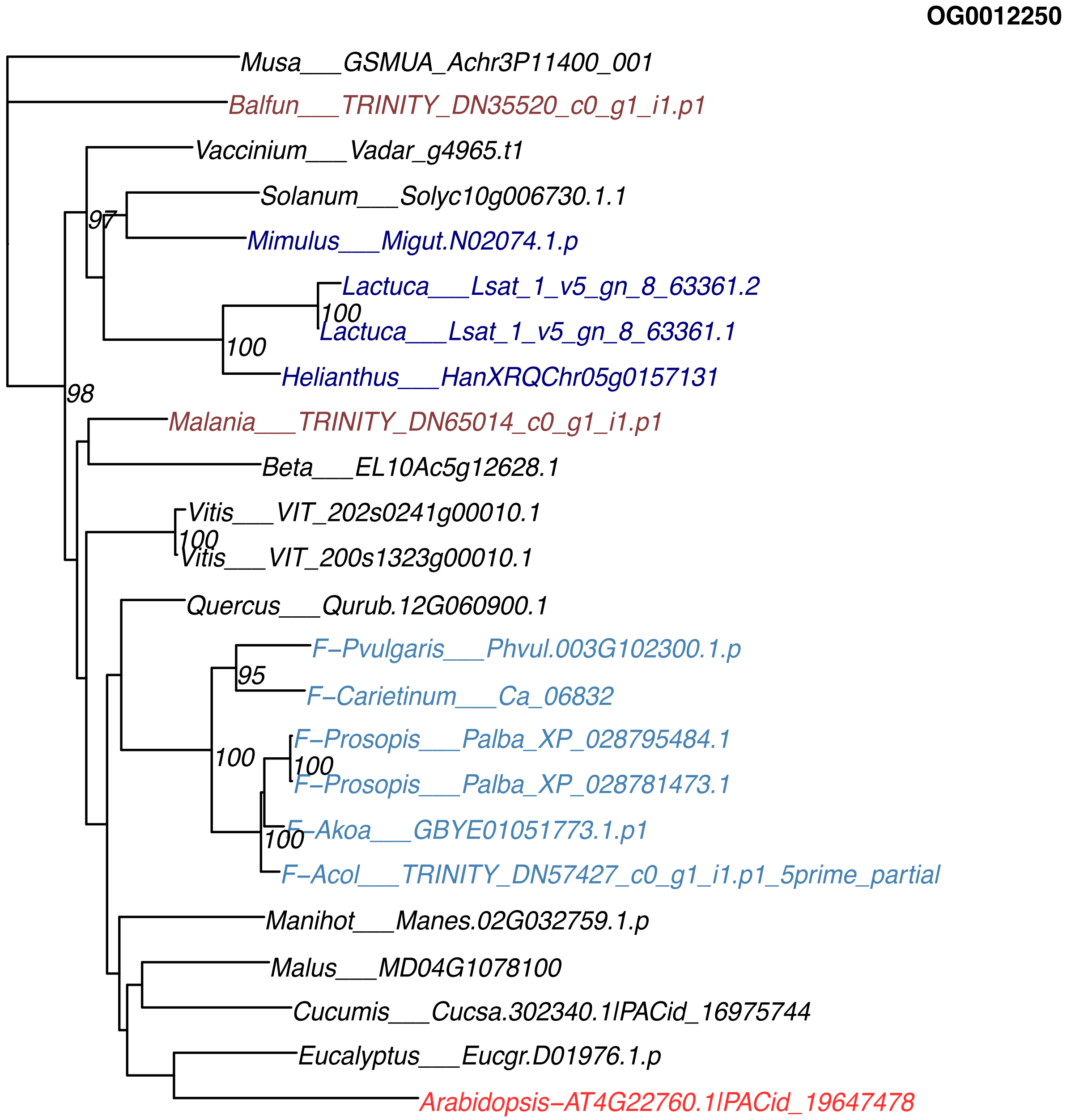
OG0012250
Musa___GSMUA_Achr3P11400_001
Balfun___TRINITY_DN35520_c0_g1_i1.p1
Vaccinium___Vadar_g4965.t1
Solanum___Solyc10g006730.1.1
Mimulus___Migut.N02074.1.p
Lactuca___Lsat_1_v5_gn_8_63361.2
Lactuca___Lsat_1_v5_gn_8_63361.1
Helianthus___HanXRQChr05g0157131
Malania___TRINITY_DN65014_c0_g1_i1.p1
Beta___EL10Ac5g12628.1
Vitis___VIT_202s0241g00010.1
Vitis___VIT_200s1323g00010.1
Quercus___Qurub.12G060900.1
F−Pvulgaris___Phvul.003G102300.1.p
F−Carietinum___Ca_06832
F−Prosopis___Palba_XP_028795484.1
F−Prosopis___Palba_XP_028781473.1
F−Akoa___GBYE01051773.1.p1
F−Acol___TRINITY_DN57427_c0_g1_i1.p1_5prime_partial
Manihot___Manes.02G032759.1.p
Malus___MD04G1078100
Cucumis___Cucsa.302340.1|PACid_16975744
Eucalyptus___Eucgr.D01976.1.p
Arabidopsis−AT4G22760.1|PACid_19647478
97
100
100
98
100
95
100
100
100

OG0012264

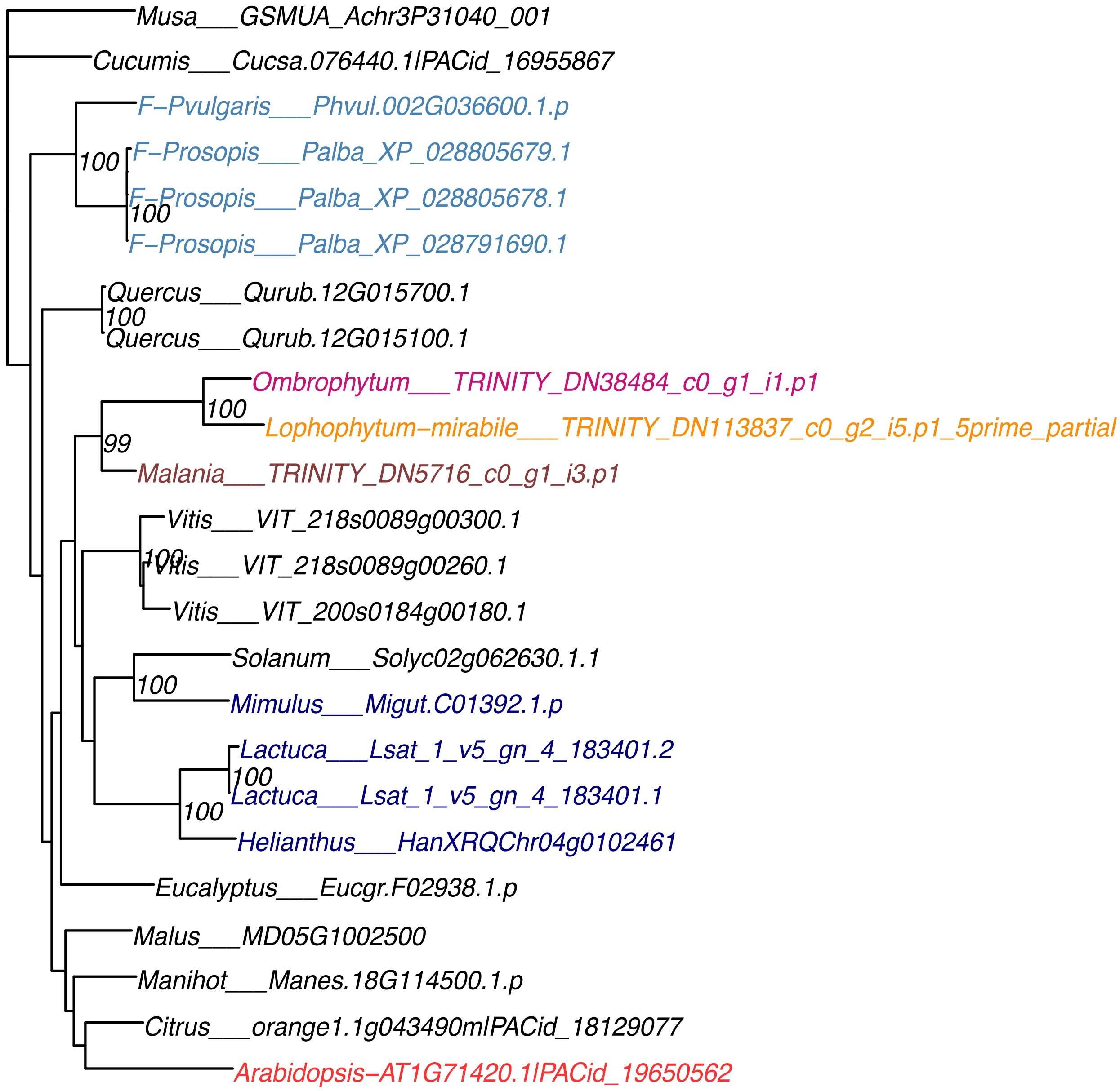

Musa___GSMUA_Achr3P31040_001
Cucumis___Cucsa.076440.1|PACid_16955867
F−Pvulgaris___Phvul.002G036600.1.p
100
F−Prosopis___Palba_XP_028805679.1
F−Prosopis___Palba_XP_028805678.1
100
F−Prosopis___Palba_XP_028791690.1
Quercus___Qurub.12G015700.1
100
Quercus___Qurub.12G015100.1
Ombrophytum___TRINITY_DN38484_c0_g1_i1.p1
100
Lophophytum−mirabile___TRINITY_DN113837_c0_g2_i5.p1_5prime_partial
99
Malania___TRINITY_DN5716_c0_g1_i3.p1
Vitis___VIT_218s0089g00300.1
100
Vitis___VIT_218s0089g00260.1
Vitis___VIT_200s0184g00180.1
Solanum___Solyc02g062630.1.1
100
Mimulus___Migut.C01392.1.p
Lactuca___Lsat_1_v5_gn_4_183401.2
100
Lactuca___Lsat_1_v5_gn_4_183401.1
100
Helianthus___HanXRQChr04g0102461
Eucalyptus___Eucgr.F02938.1.p
Malus___MD05G1002500
Manihot___Manes.18G114500.1.p
Citrus___orange1.1g043490m|PACid_18129077
Arabidopsis−AT1G71420.1|PACid_19650562

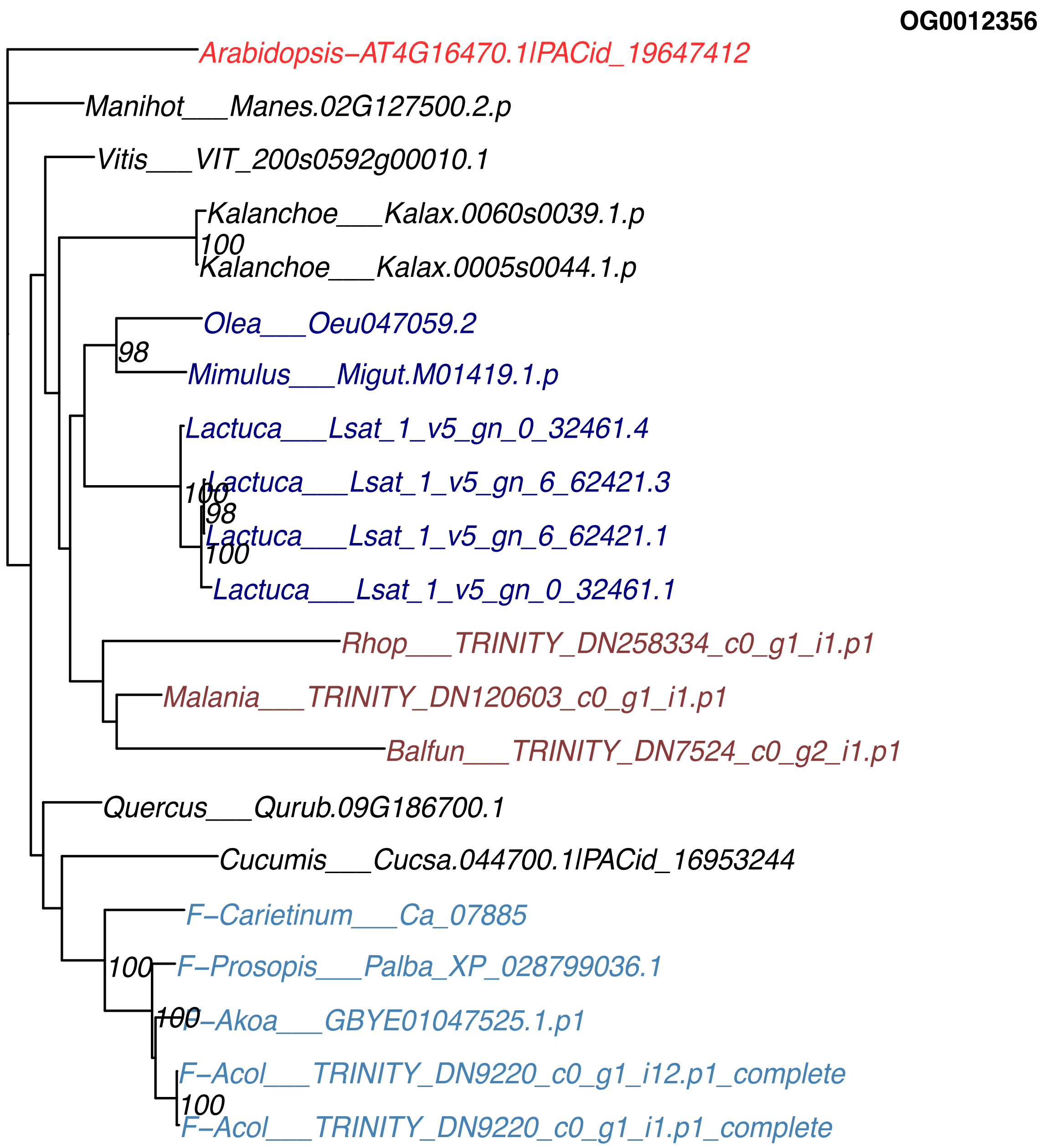
OG0012356
Arabidopsis−AT4G16470.1|PACid_19647412
Manihot___Manes.02G127500.2.p
Vitis___VIT_200s0592g00010.1
Kalanchoe___Kalax.0060s0039.1.p
100
Kalanchoe___Kalax.0005s0044.1.p
Olea___Oeu047059.2
98
Mimulus___Migut.M01419.1.p
Lactuca___Lsat_1_v5_gn_0_32461.4
Lactuca___Lsat_1_v5_gn_6_62421.3
100
98
Lactuca___Lsat_1_v5_gn_6_62421.1
100
Lactuca___Lsat_1_v5_gn_0_32461.1
Rhop___TRINITY_DN258334_c0_g1_i1.p1
Malania___TRINITY_DN120603_c0_g1_i1.p1
Balfun___TRINITY_DN7524_c0_g2_i1.p1
Quercus___Qurub.09G186700.1
Cucumis___Cucsa.044700.1|PACid_16953244
F−Carietinum___Ca_07885
100
F−Prosopis___Palba_XP_028799036.1
100
F−Akoa___GBYE01047525.1.p1
F−Acol___TRINITY_DN9220_c0_g1_i12.p1_complete
100
F−Acol___TRINITY_DN9220_c0_g1_i1.p1_complete

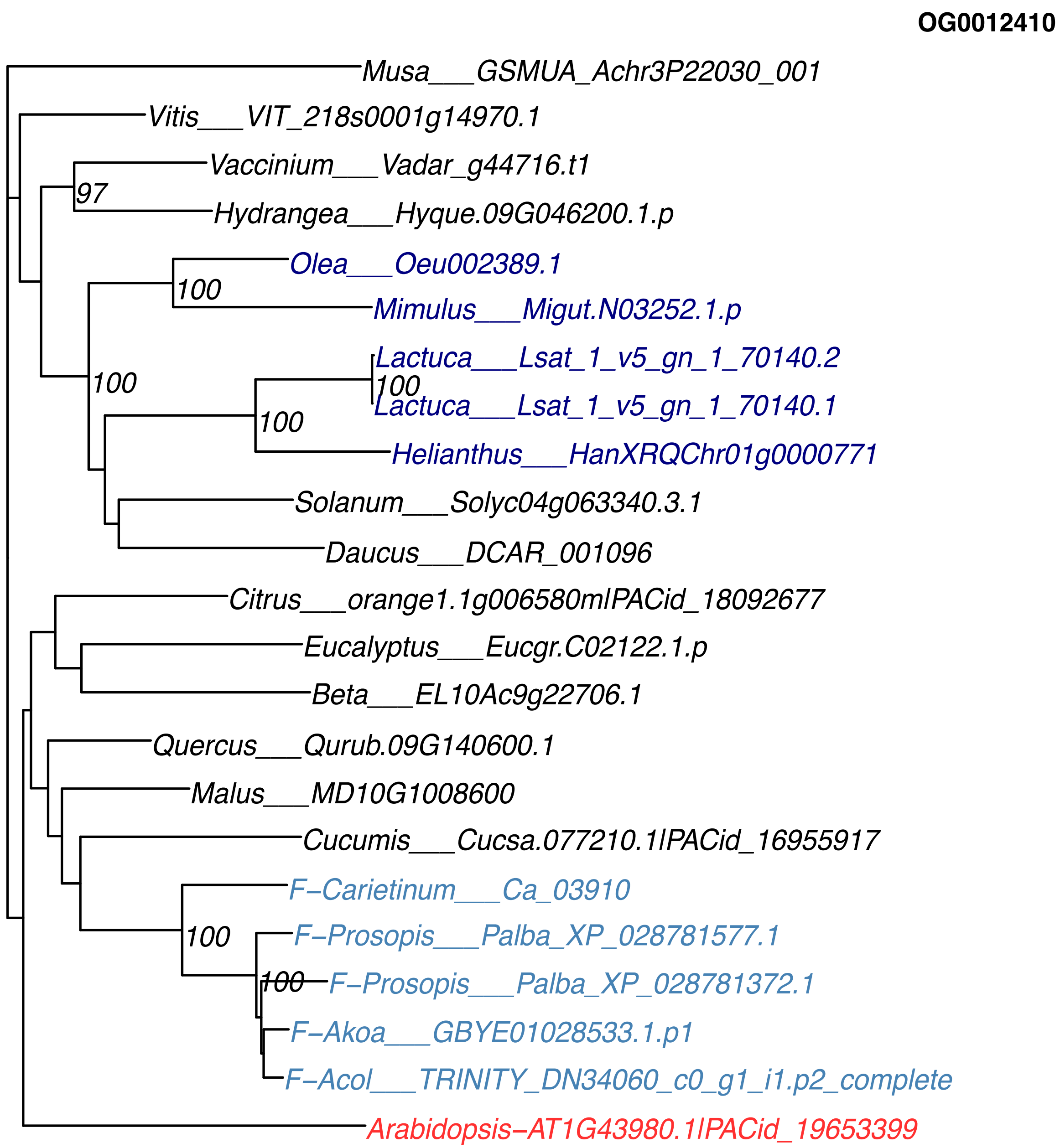
OG0012410
Musa___GSMUA_Achr3P22030_001
Vitis___VIT_218s0001g14970.1
Vaccinium___Vadar_g44716.t1
97
Hydrangea___Hyque.09G046200.1.p
Olea___Oeu002389.1
100
Mimulus___Migut.N03252.1.p
Lactuca___Lsat_1_v5_gn_1_70140.2
100
100
Lactuca___Lsat_1_v5_gn_1_70140.1
100
Helianthus___HanXRQChr01g0000771
Solanum___Solyc04g063340.3.1
Daucus___DCAR_001096
Citrus___orange1.1g006580m|PACid_18092677
Eucalyptus___Eucgr.C02122.1.p
Beta___EL10Ac9g22706.1
Quercus___Qurub.09G140600.1
Malus___MD10G1008600
Cucumis___Cucsa.077210.1|PACid_16955917
F−Carietinum___Ca_03910
100
F−Prosopis___Palba_XP_028781577.1
100
F−Prosopis___Palba_XP_028781372.1
F−Akoa___GBYE01028533.1.p1
F−Acol___TRINITY_DN34060_c0_g1_i1.p2_complete
Arabidopsis−AT1G43980.1|PACid_19653399

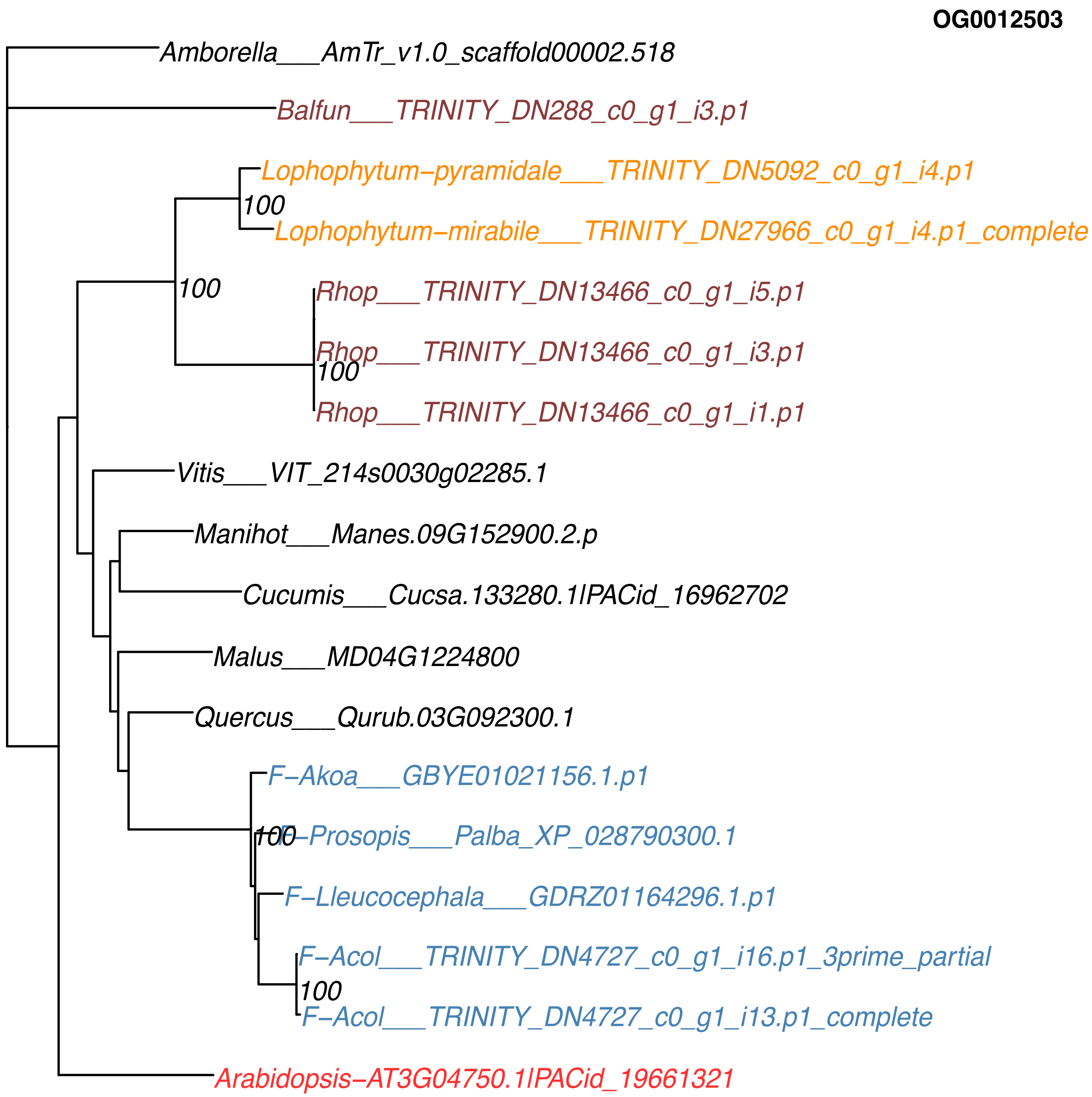
OG0012503
Amborella___AmTr_v1.0_scaffold00002.518
Balfun___TRINITY_DN288_c0_g1_i3.p1
Lophophytum−pyramidale___TRINITY_DN5092_c0_g1_i4.p1
100
Lophophytum−mirabile___TRINITY_DN27966_c0_g1_i4.p1_complete
100
Rhop___TRINITY_DN13466_c0_g1_i5.p1
Rhop___TRINITY_DN13466_c0_g1_i3.p1
100
Rhop___TRINITY_DN13466_c0_g1_i1.p1
Vitis___VIT_214s0030g02285.1
Manihot___Manes.09G152900.2.p
Cucumis___Cucsa.133280.1|PACid_16962702
Malus___MD04G1224800
Quercus___Qurub.03G092300.1
F−Akoa___GBYE01021156.1.p1
100
F−Prosopis___Palba_XP_028790300.1
F−Lleucocephala___GDRZ01164296.1.p1
F−Acol___TRINITY_DN4727_c0_g1_i16.p1_3prime_partial
100
F−Acol___TRINITY_DN4727_c0_g1_i13.p1_complete
Arabidopsis−AT3G04750.1|PACid_19661321

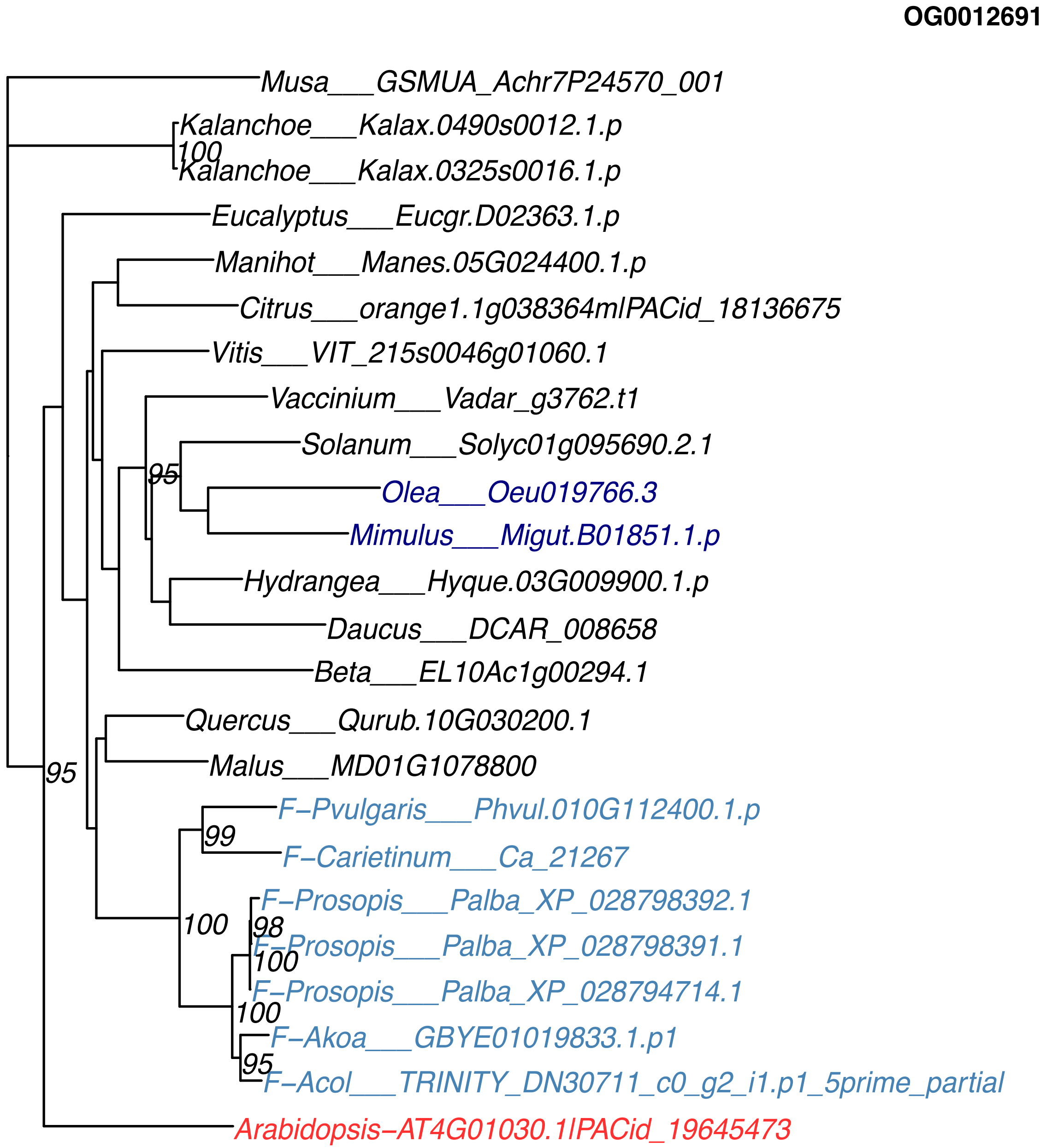

OG0012691
Musa___GSMUA_Achr7P24570_001
Kalanchoe___Kalax.0490s0012.1.p
100
Kalanchoe___Kalax.0325s0016.1.p
Eucalyptus___Eucgr.D02363.1.p
Manihot___Manes.05G024400.1.p
Citrus___orange1.1g038364m|PACid_18136675
Vitis___VIT_215s0046g01060.1
Vaccinium___Vadar_g3762.t1
Solanum___Solyc01g095690.2.1
95
Olea___Oeu019766.3
Mimulus___Migut.B01851.1.p
Hydrangea___Hyque.03G009900.1.p
Daucus___DCAR_008658
Beta___EL10Ac1g00294.1
Quercus___Qurub.10G030200.1
Malus___MD01G1078800
95
F–Pvulgaris___Phvul.010G112400.1.p
99
F–Carietinum___Ca_21267
F–Prosopis___Palba_XP_028798392.1
100
98
F–Prosopis___Palba_XP_028798391.1
100
F–Prosopis___Palba_XP_028794714.1
100
F–Akoa___GBYE01019833.1.p1
95
F–Acol___TRINITY_DN30711_c0_g2_i1.p1_5prime_partial
Arabidopsis–AT4G01030.1|PACid_19645473

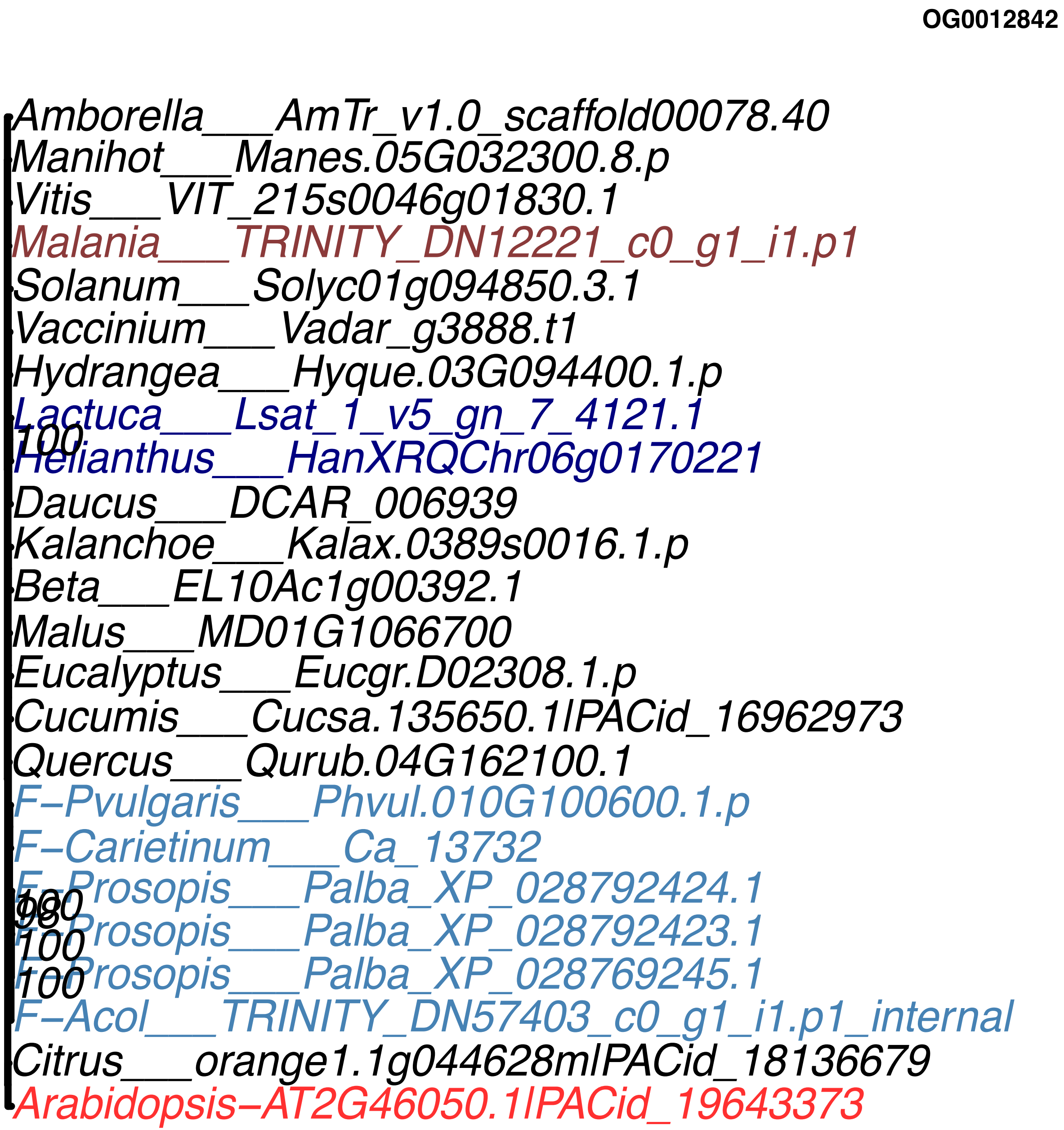
OG0012842
Amborella___AmTr_v1.0_scaffold00078.40
Manihot___Manes.05G032300.8.p
Vitis___VIT_215s0046g01830.1
Malania___TRINITY_DN12221_c0_g1_i1.p1
Solanum___Solyc01g094850.3.1
Vaccinium___Vadar_g3888.t1
Hydrangea___Hyque.03G094400.1.p
Lactuca___Lsat_1_v5_gn_7_4121.1
100
Helianthus___HanXRQChr06g0170221
Daucus___DCAR_006939
Kalanchoe___Kalax.0389s0016.1.p
Beta___EL10Ac1g00392.1
Malus___MD01G1066700
Eucalyptus___Eucgr.D02308.1.p
Cucumis___Cucsa.135650.1|PACid_16962973
Quercus___Qurub.04G162100.1
F−Pvulgaris___Phvul.010G100600.1.p
F−Carietinum___Ca_13732
F−Prosopis___Palba_XP_028792424.1
100
98
F−Prosopis___Palba_XP_028792423.1
100
F−Prosopis___Palba_XP_028769245.1
100
F−Acol___TRINITY_DN57403_c0_g1_i1.p1_internal
Citrus___orange1.1g044628m|PACid_18136679
Arabidopsis−AT2G46050.1|PACid_19643373

**OG0012870**

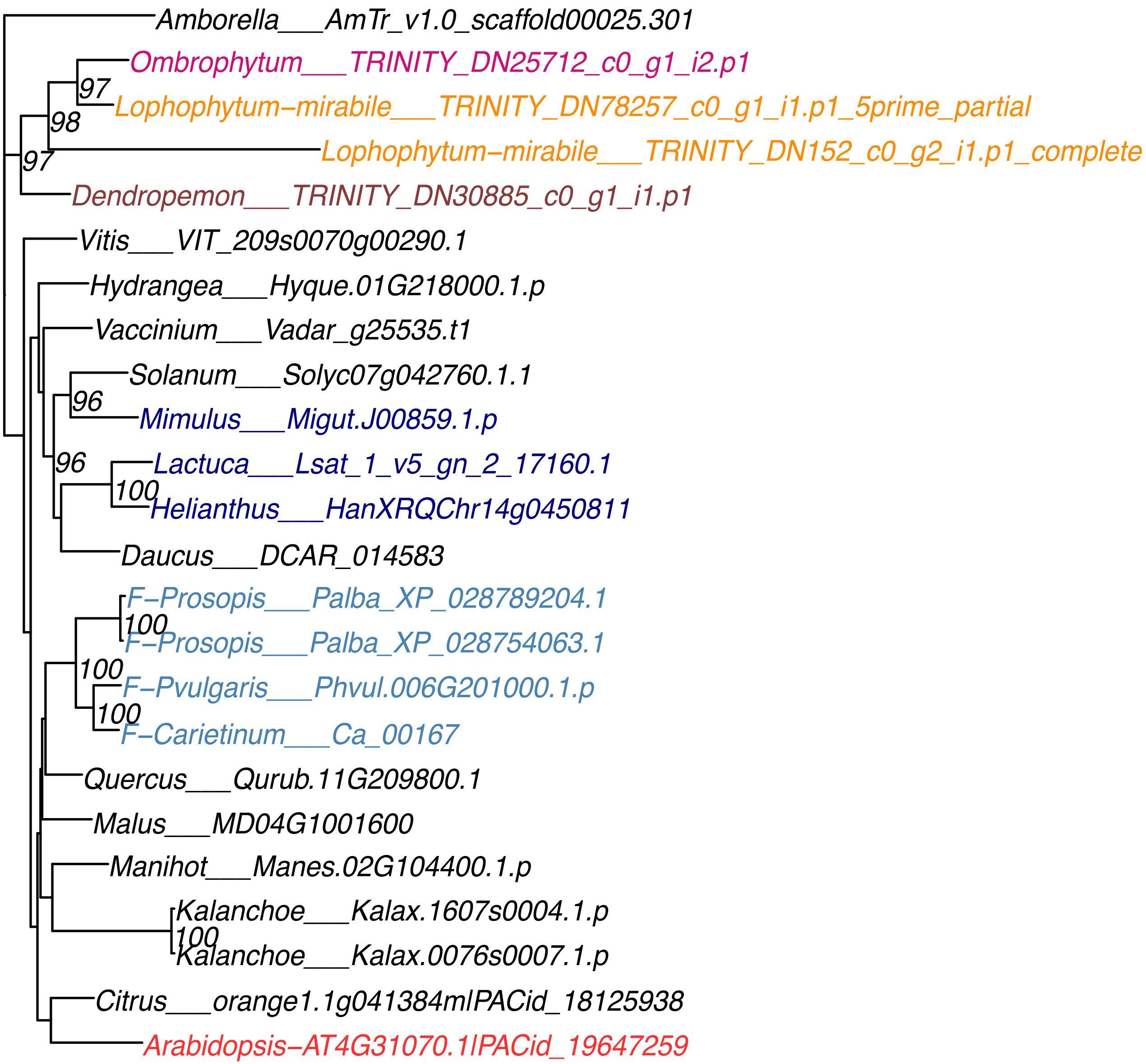

Amborella___AmTr_v1.0_scaffold00025.301
Ombrophytum___TRINITY_DN25712_c0_g1_i2.p1
97
Lophophytum−mirabile___TRINITY_DN78257_c0_g1_i1.p1_5prime_partial
98
Lophophytum−mirabile___TRINITY_DN152_c0_g2_i1.p1_complete
97
Dendropemon___TRINITY_DN30885_c0_g1_i1.p1
Vitis___VIT_209s0070g00290.1
Hydrangea___Hyque.01G218000.1.p
Vaccinium___Vadar_g25535.t1
Solanum___Solyc07g042760.1.1
96
Mimulus___Migut.J00859.1.p
96
Lactuca___Lsat_1_v5_gn_2_17160.1
100
Helianthus___HanXRQChr14g0450811
Daucus___DCAR_014583
F−Prosopis___Palba_XP_028789204.1
100
F−Prosopis___Palba_XP_028754063.1
100
F−Pvulgaris___Phvul.006G201000.1.p
100
F−Carietinum___Ca_00167
Quercus___Qurub.11G209800.1
Malus___MD04G1001600
Manihot___Manes.02G104400.1.p
Kalanchoe___Kalax.1607s0004.1.p
100
Kalanchoe___Kalax.0076s0007.1.p
Citrus___orange1.1g041384mlPACid_18125938
Arabidopsis−AT4G31070.1lPACid_19647259

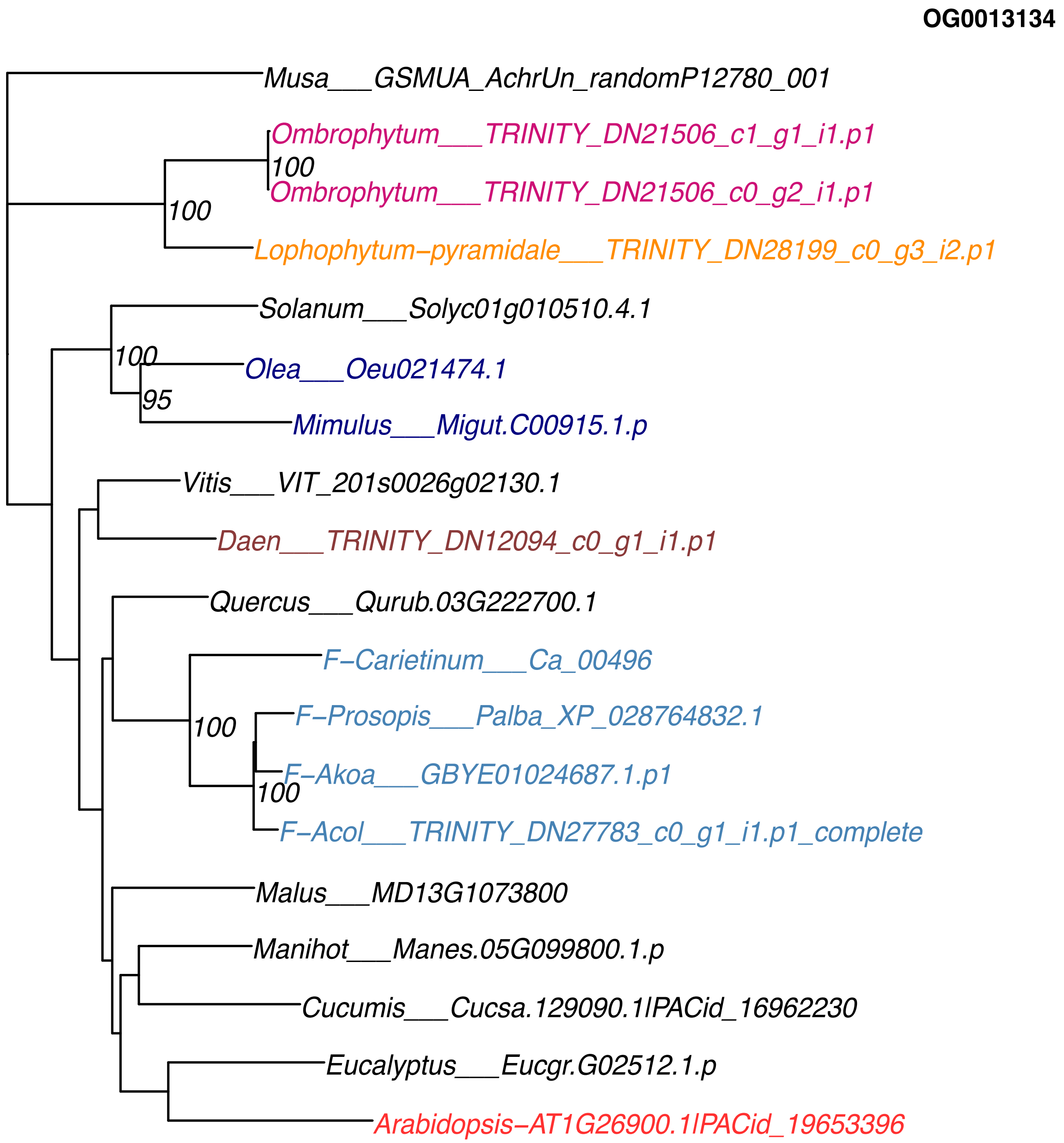

OG0013134
Musa___GSMUA_AchrUn_randomP12780_001
Ombrophytum___TRINITY_DN21506_c1_g1_i1.p1
100
Ombrophytum___TRINITY_DN21506_c0_g2_i1.p1
100
Lophophytum−pyramidale___TRINITY_DN28199_c0_g3_i2.p1
Solanum___Solyc01g010510.4.1
100
Olea___Oeu021474.1
95
Mimulus___Migut.C00915.1.p
Vitis___VIT_201s0026g02130.1
Daen___TRINITY_DN12094_c0_g1_i1.p1
Quercus___Qurub.03G222700.1
F−Carietinum___Ca_00496
F−Prosopis___Palba_XP_028764832.1
100
F−Akoa___GBYE01024687.1.p1
100
F−Acol___TRINITY_DN27783_c0_g1_i1.p1_complete
Malus___MD13G1073800
Manihot___Manes.05G099800.1.p
Cucumis___Cucsa.129090.1|PACid_16962230
Eucalyptus___Eucgr.G02512.1.p
Arabidopsis−AT1G26900.1|PACid_19653396

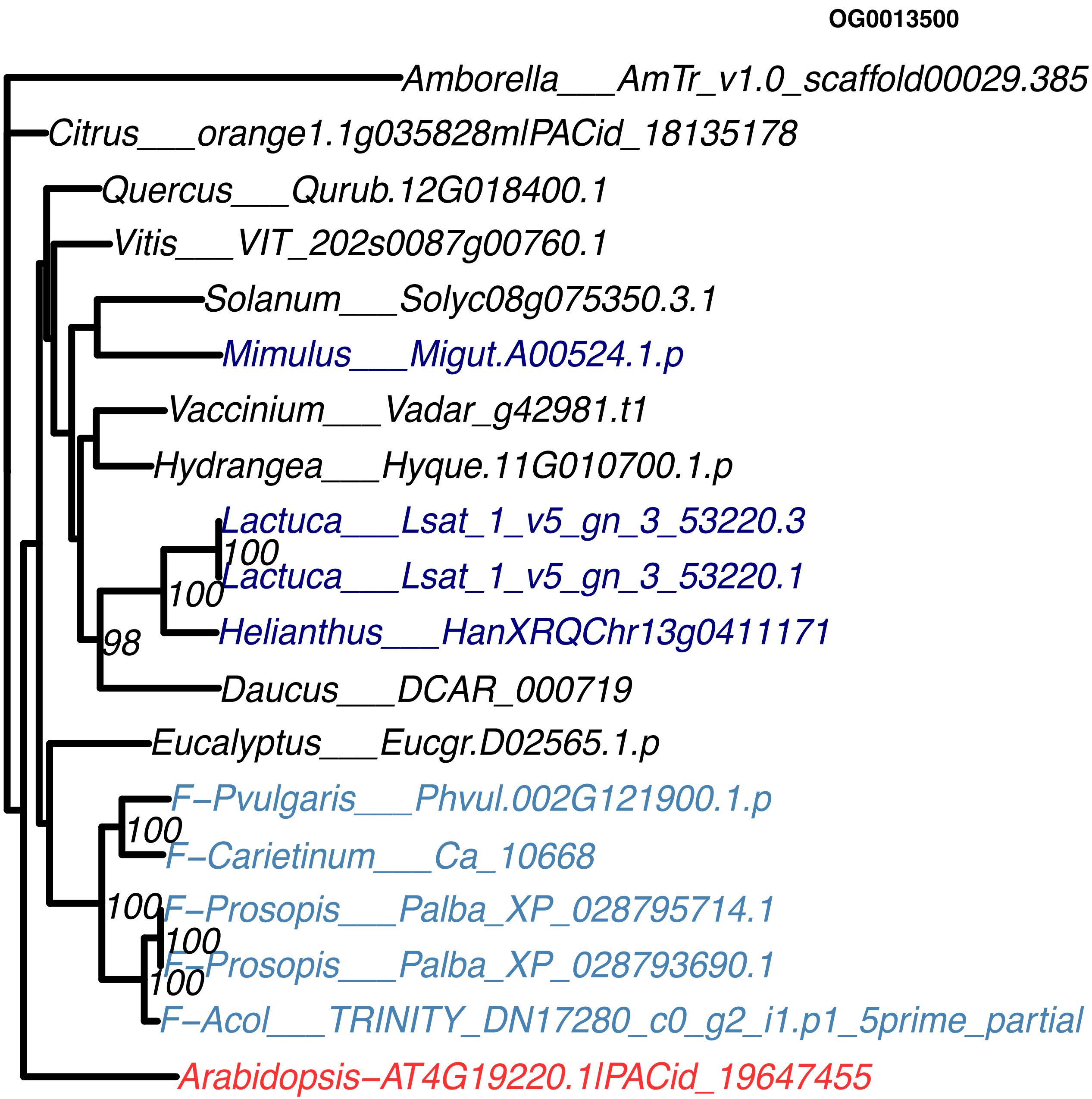

OG0013500
Amborella___AmTr_v1.0_scaffold00029.385
Citrus___orange1.1g035828mlPACid_18135178
Quercus___Qurub.12G018400.1
Vitis___VIT_202s0087g00760.1
Solanum___Solyc08g075350.3.1
Mimulus___Migut.A00524.1.p
Vaccinium___Vadar_g42981.t1
Hydrangea___Hyque.11G010700.1.p
Lactuca___Lsat_1_v5_gn_3_53220.3
100
Lactuca___Lsat_1_v5_gn_3_53220.1
100
Helianthus___HanXRQChr13g0411171
98
Daucus___DCAR_000719
Eucalyptus___Eucgr.D02565.1.p
F−Pvulgaris___Phvul.002G121900.1.p
100
F−Carietinum___Ca_10668
100
F−Prosopis___Palba_XP_028795714.1
100
F−Prosopis___Palba_XP_028793690.1
100
F−Acol___TRINITY_DN17280_c0_g2_i1.p1_5prime_partial
Arabidopsis−AT4G19220.1lPACid_19647455

**OG0013731**

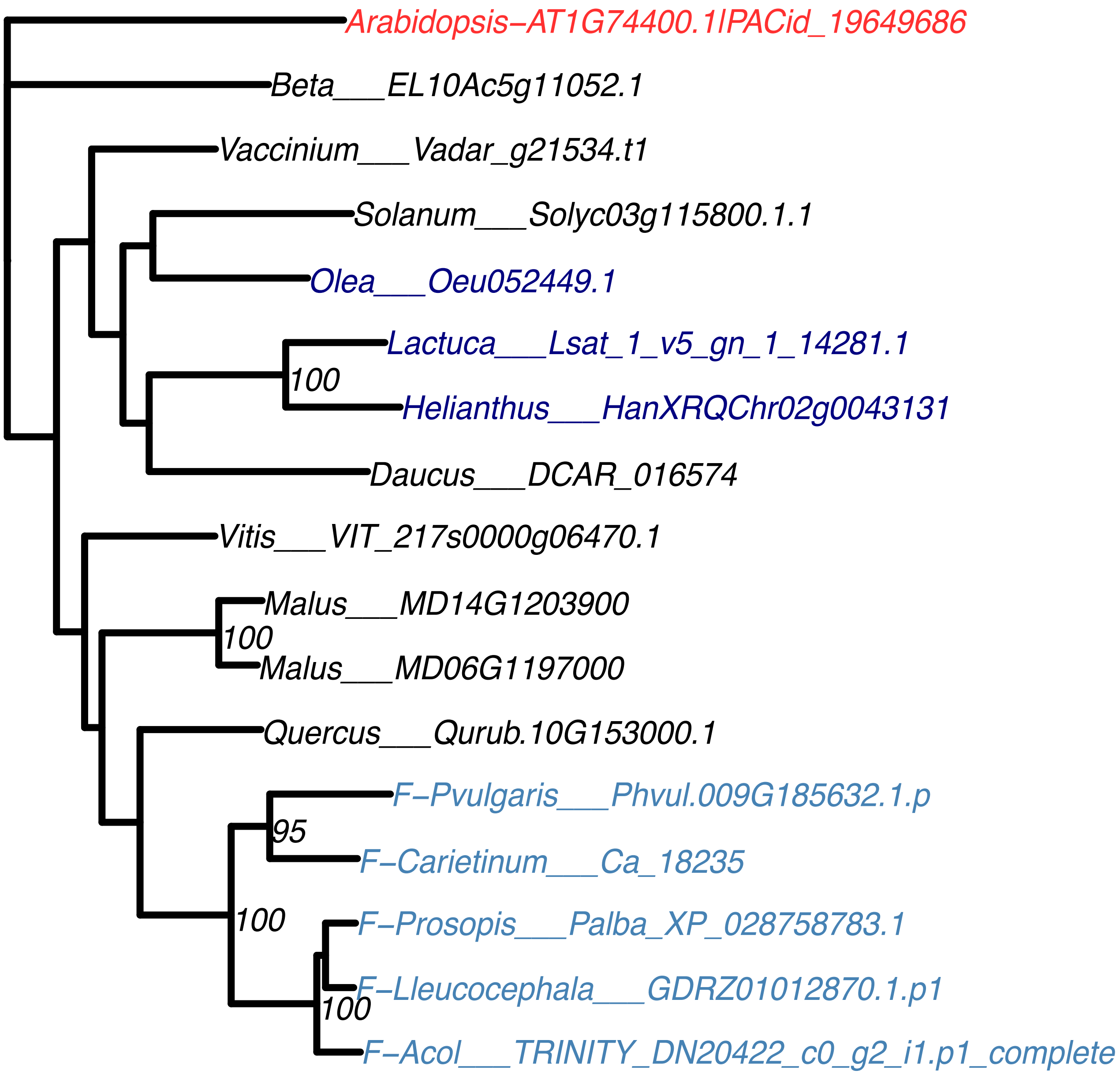
Arabidopsis–AT1G74400.1|PACid_19649686
Beta___EL10Ac5g11052.1
Vaccinium___Vadar_g21534.t1
Solanum___Solyc03g115800.1.1
Olea___Oeu052449.1
Lactuca___Lsat_1_v5_gn_1_14281.1
100
Helianthus___HanXRQChr02g0043131
Daucus___DCAR_016574
Vitis___VIT_217s0000g06470.1
Malus___MD14G1203900
100
Malus___MD06G1197000
Quercus___Qurub.10G153000.1
F–Pvulgaris___Phvul.009G185632.1.p
95
F–Carietinum___Ca_18235
100
F–Prosopis___Palba_XP_028758783.1
F–Lleucocephala___GDRZ01012870.1.p1
100
F–Acol___TRINITY_DN20422_c0_g2_i1.p1_complete

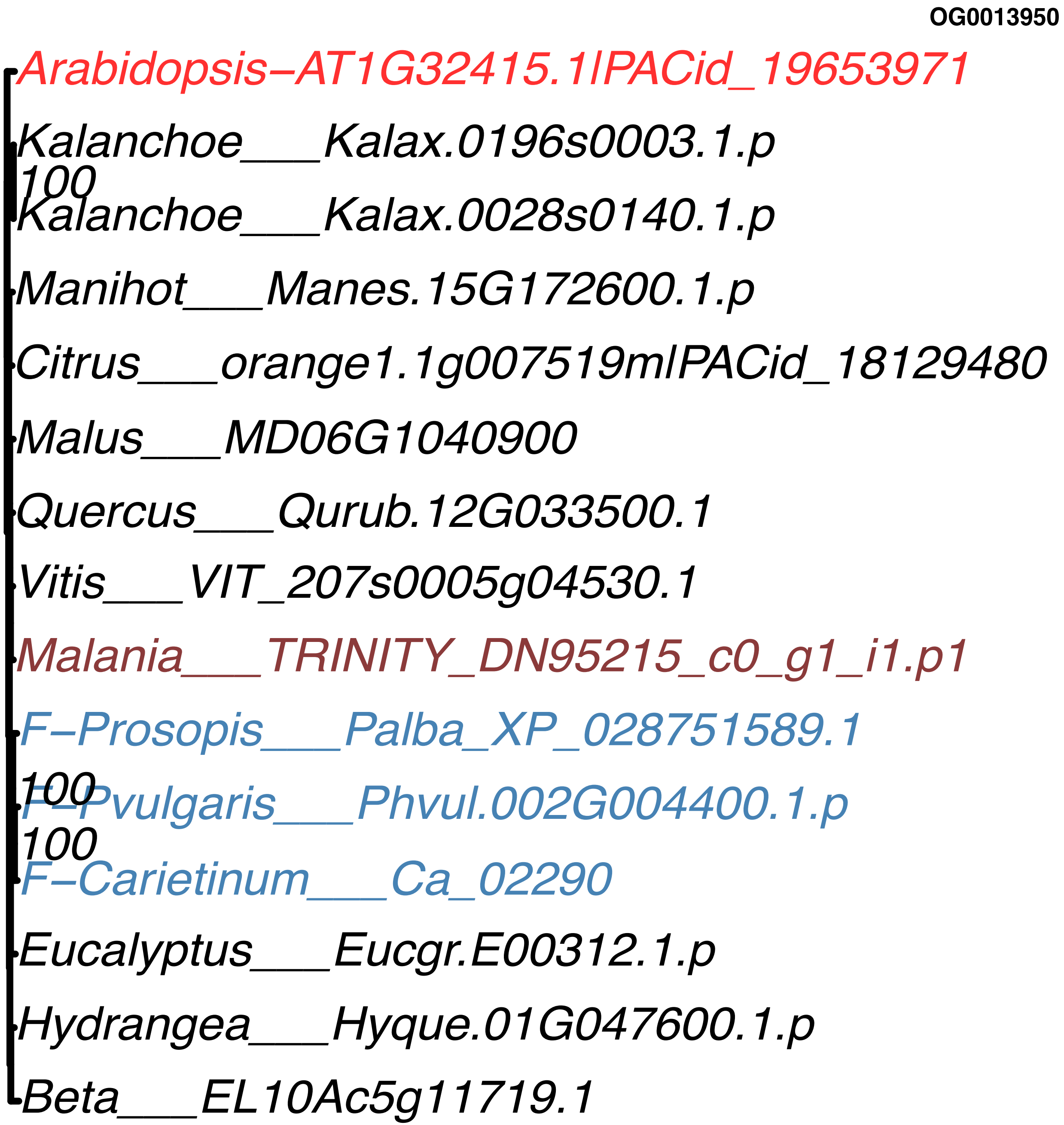
OG0013950
Arabidopsis–AT1G32415.1|PACid_19653971
Kalanchoe___Kalax.0196s0003.1.p
100
Kalanchoe___Kalax.0028s0140.1.p
Manihot___Manes.15G172600.1.p
Citrus___orange1.1g007519m|PACid_18129480
Malus___MD06G1040900
Quercus___Qurub.12G033500.1
Vitis___VIT_207s0005g04530.1
Malania___TRINITY_DN95215_c0_g1_i1.p1
F–Prosopis___Palba_XP_028751589.1
100
F–Pvulgaris___Phvul.002G004400.1.p
100
F–Carietinum___Ca_02290
Eucalyptus___Eucgr.E00312.1.p
Hydrangea___Hyque.01G047600.1.p
Beta___EL10Ac5g11719.1

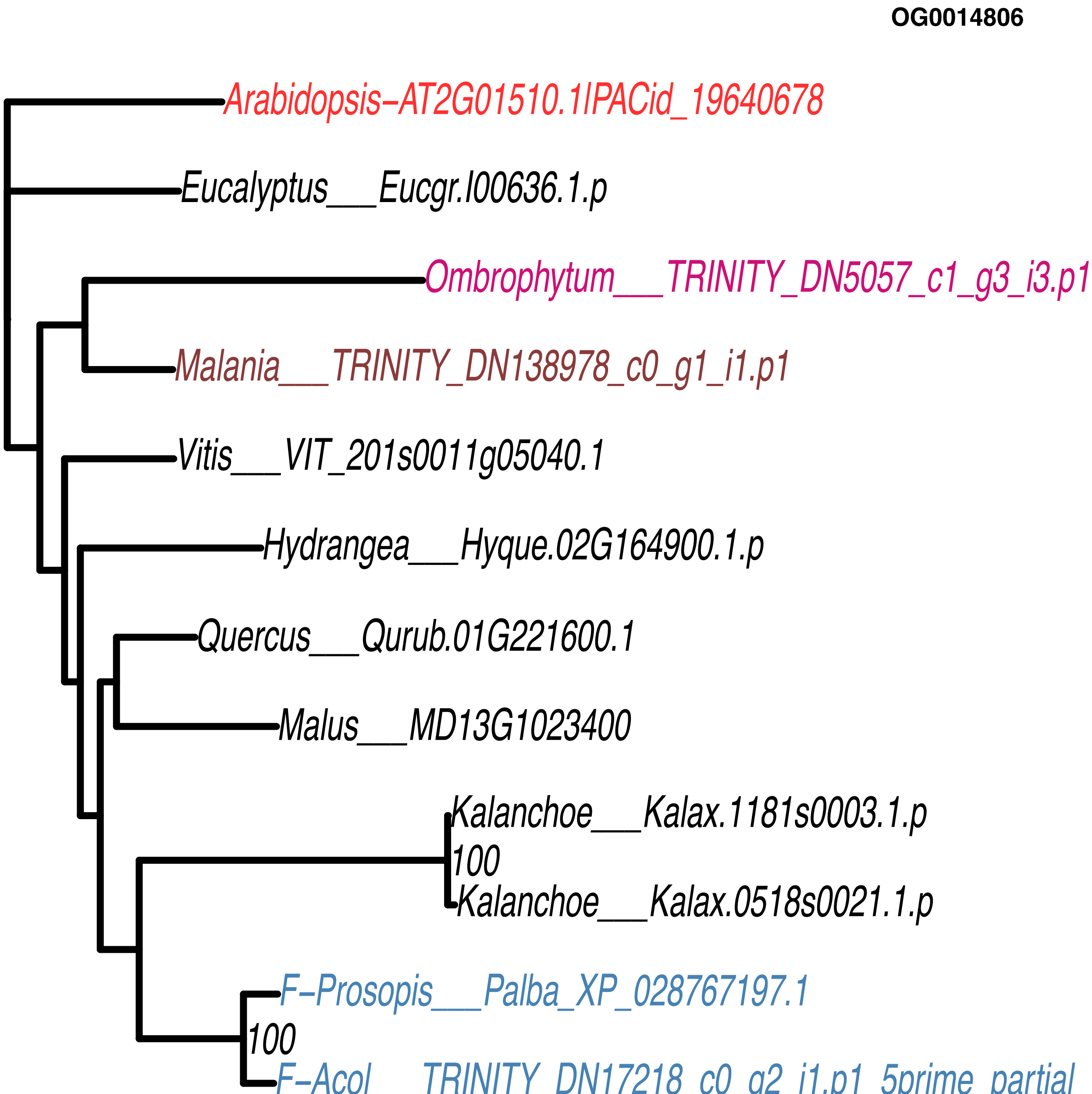
OG0014806
Arabidopsis–AT2G01510.1lPACid_19640678
Eucalyptus___Eucgr.I00636.1.p
Ombrophytum___TRINITY_DN5057_c1_g3_i3.p1
Malania___TRINITY_DN138978_c0_g1_i1.p1
Vitis___VIT_201s0011g05040.1
Hydrangea___Hyque.02G164900.1.p
Quercus___Qurub.01G221600.1
Malus___MD13G1023400
Kalanchoe___Kalax.1181s0003.1.p
100
Kalanchoe___Kalax.0518s0021.1.p
F–Prosopis___Palba_XP_028767197.1
100
F–Acol___TRINITY_DN17218_c0_g2_i1.p1_5prime_partial

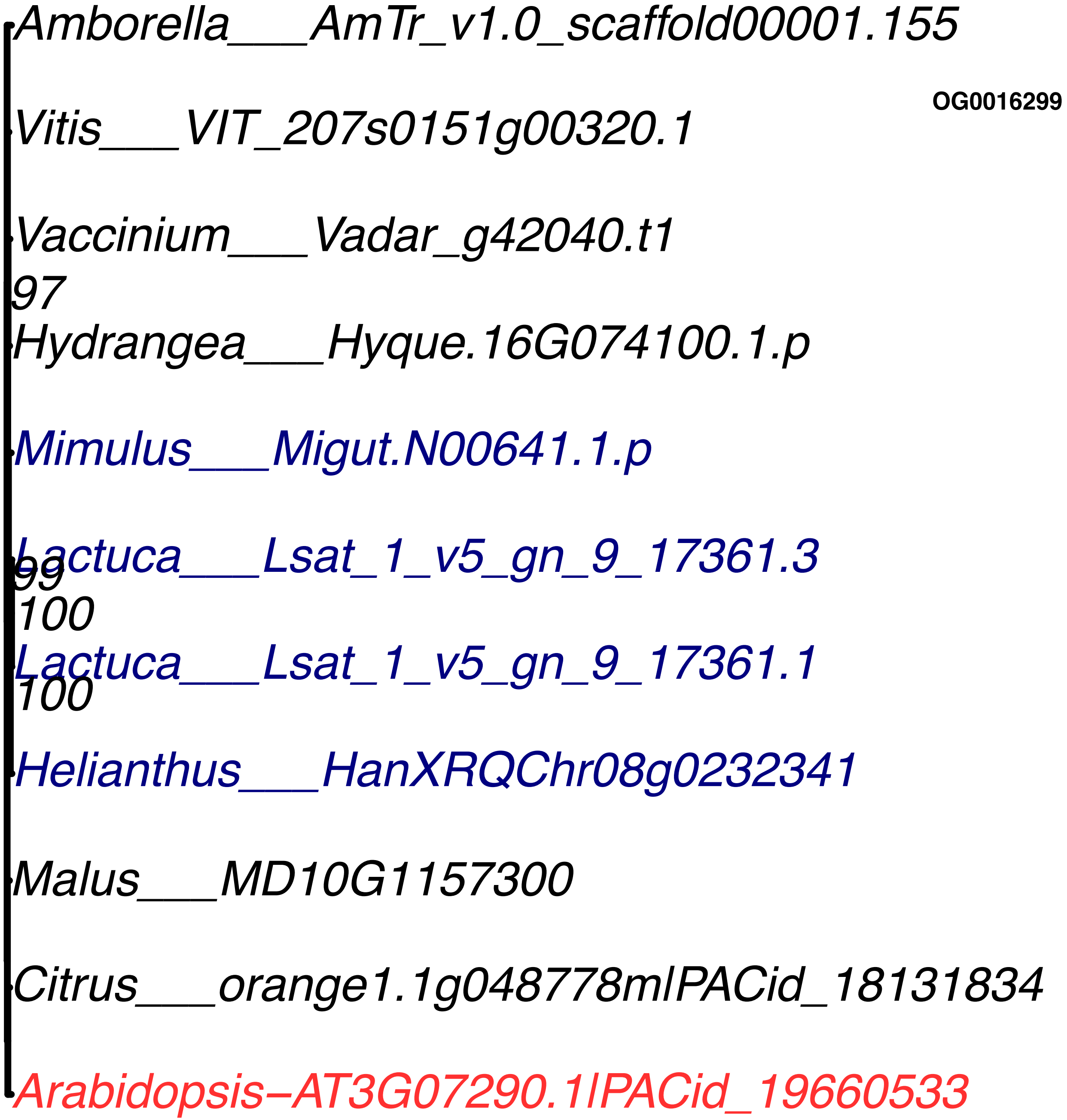
OG0016299
Amborella___AmTr_v1.0_scaffold00001.155
Vitis___VIT_207s0151g00320.1
Vaccinium___Vadar_g42040.t1
97
Hydrangea___Hyque.16G074100.1.p
Mimulus___Migut.N00641.1.p
Lactuca___Lsat_1_v5_gn_9_17361.3
99
100
Lactuca___Lsat_1_v5_gn_9_17361.1
100
Helianthus___HanXRQChr08g0232341
Malus___MD10G1157300
Citrus___orange1.1g048778m|PACid_18131834
Arabidopsis–AT3G07290.1|PACid_19660533

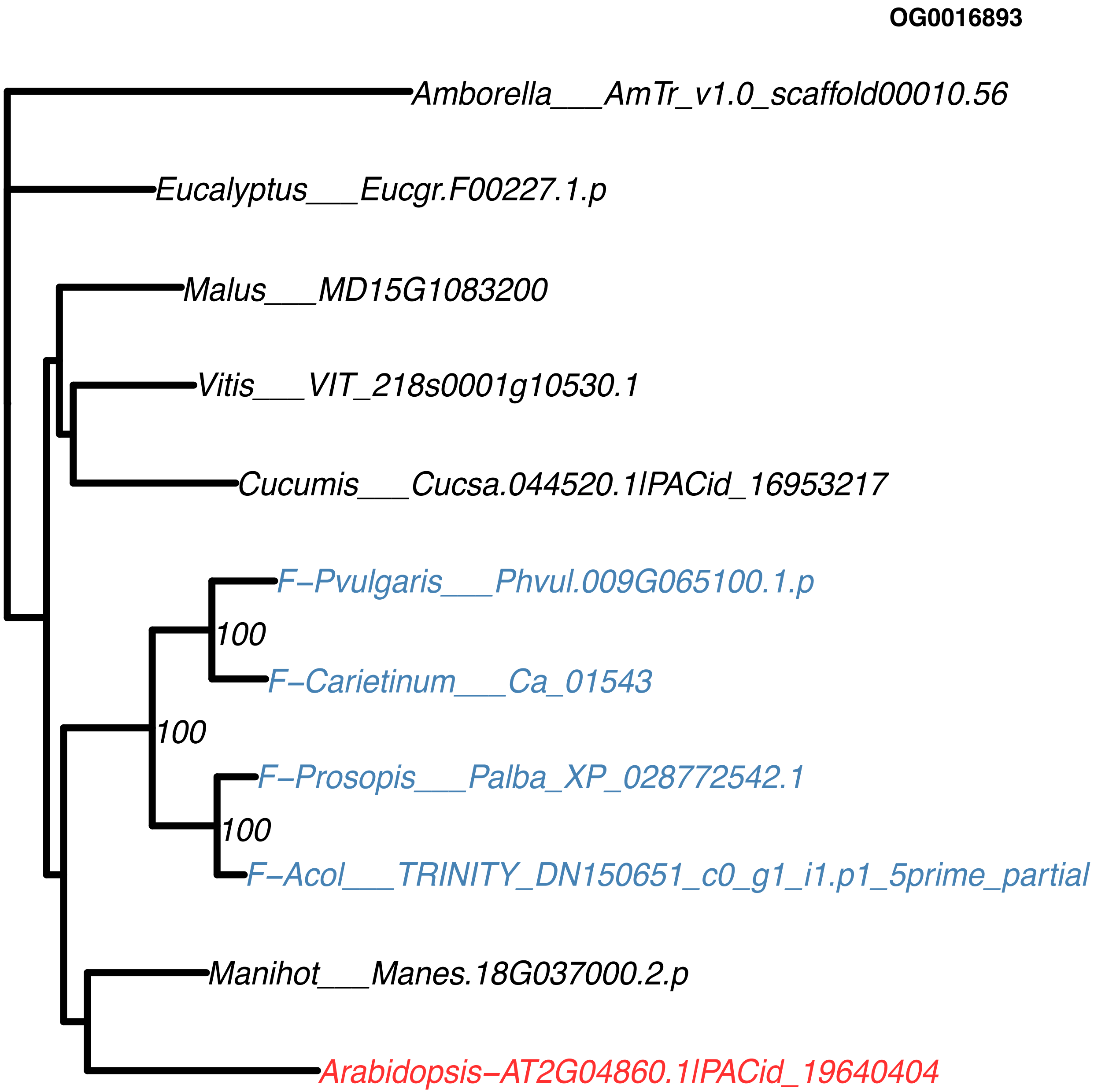
OG0016893
Amborella___AmTr_v1.0_scaffold00010.56
Eucalyptus___Eucgr.F00227.1.p
Malus___MD15G1083200
Vitis___VIT_218s0001g10530.1
Cucumis___Cucsa.044520.1|PACid_16953217
F−Pvulgaris___Phvul.009G065100.1.p
100
F−Carietinum___Ca_01543
100
F−Prosopis___Palba_XP_028772542.1
100
F−Acol___TRINITY_DN150651_c0_g1_i1.p1_5prime_partial
Manihot___Manes.18G037000.2.p
Arabidopsis−AT2G04860.1|PACid_19640404

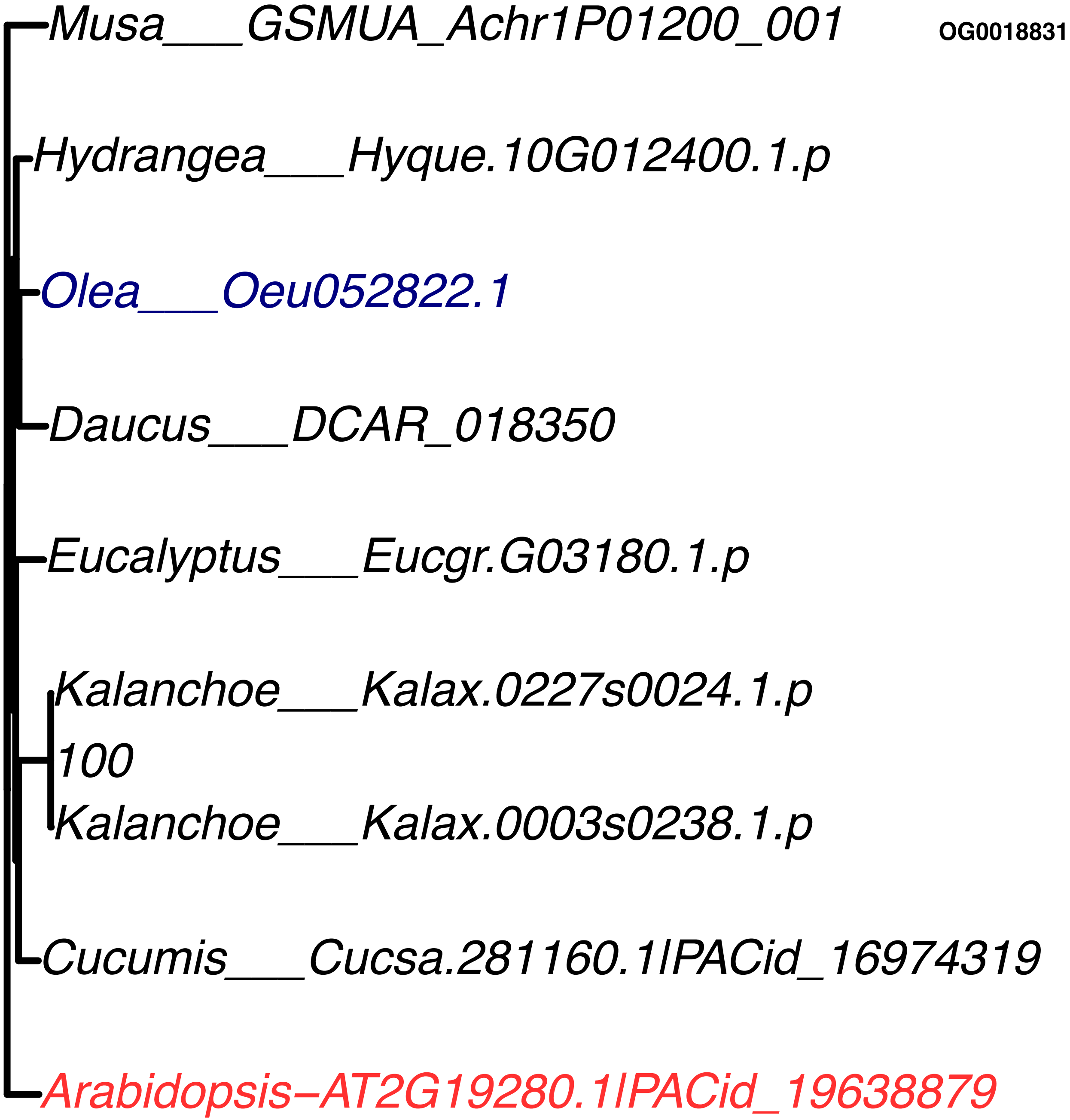
OG0018831
Musa___GSMUA_Achr1P01200_001
Hydrangea___Hyque.10G012400.1.p
Olea___Oeu052822.1
Daucus___DCAR_018350
Eucalyptus___Eucgr.G03180.1.p
Kalanchoe___Kalax.0227s0024.1.p
100
Kalanchoe___Kalax.0003s0238.1.p
Cucumis___Cucsa.281160.1|PACid_16974319
Arabidopsis–AT2G19280.1|PACid_19638879

OG0018833

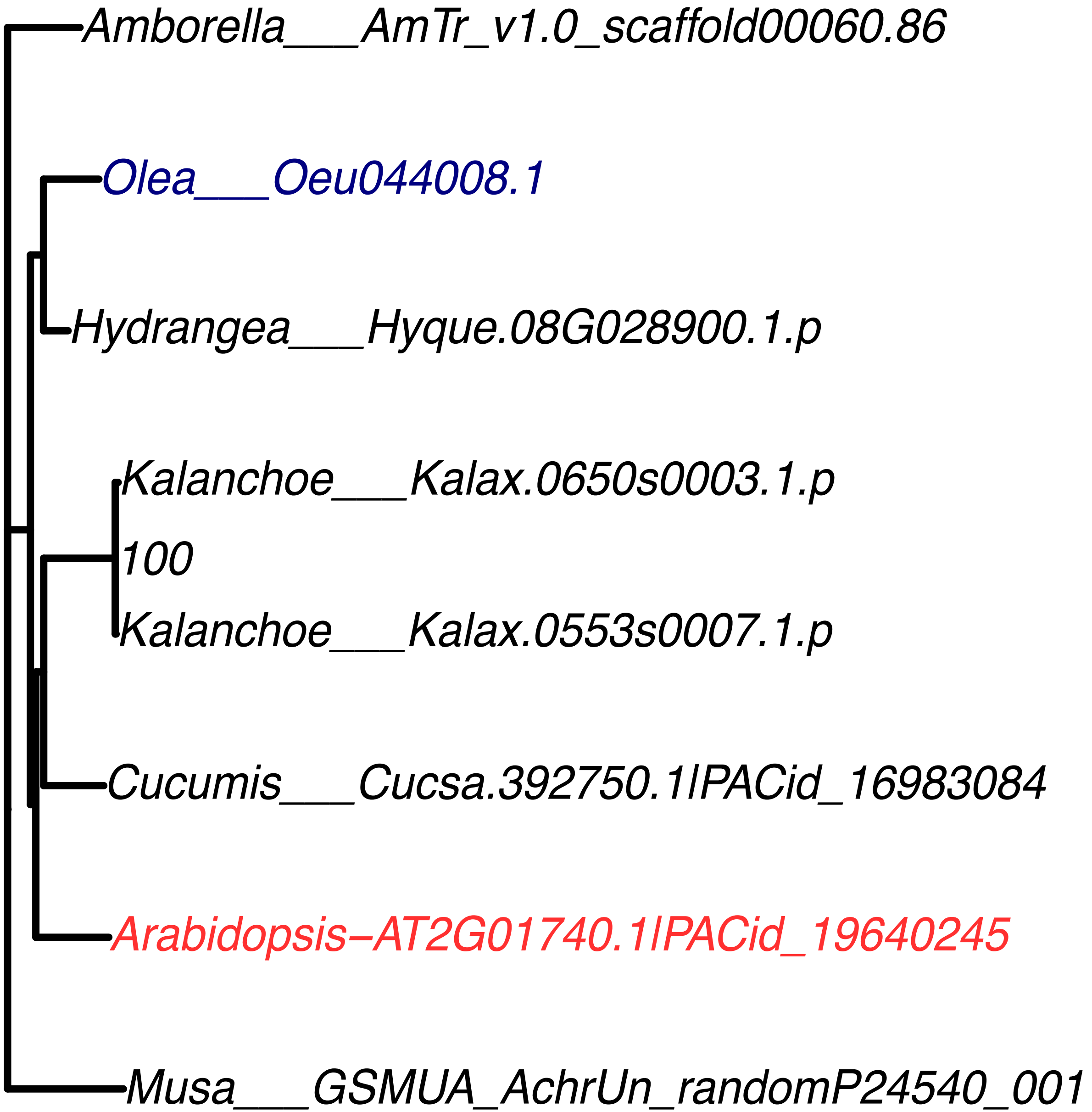
Amborella___AmTr_v1.0_scaffold00060.86
Olea___Oeu044008.1
Hydrangea___Hyque.08G028900.1.p
Kalanchoe___Kalax.0650s0003.1.p
100
Kalanchoe___Kalax.0553s0007.1.p
Cucumis___Cucsa.392750.1|PACid_16983084
Arabidopsis–AT2G01740.1|PACid_19640245
Musa___GSMUA_AchrUn_randomP24540_001

OG0027900

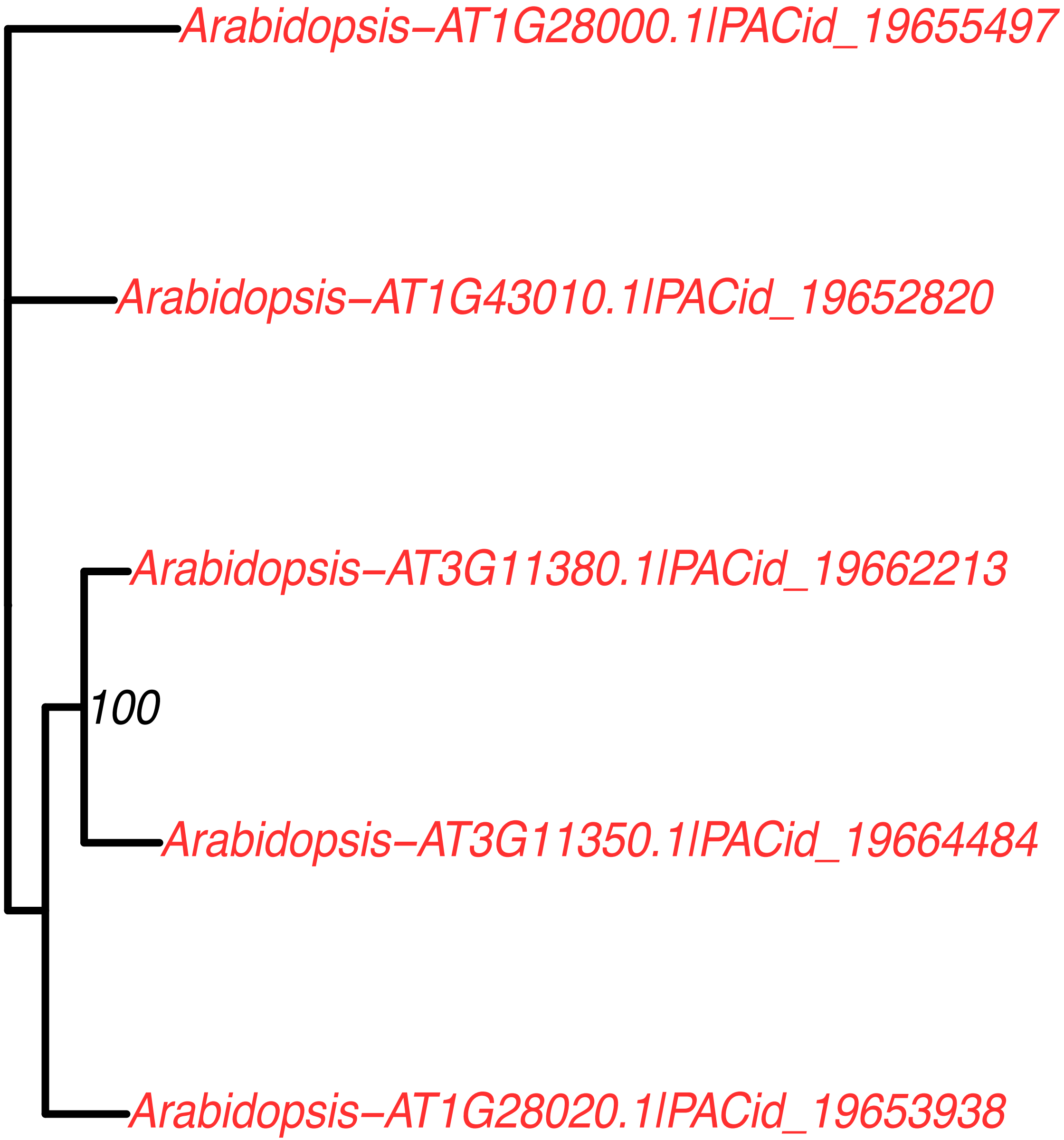
Arabidopsis–AT1G28000.1|PACid_19655497
Arabidopsis–AT1G43010.1|PACid_19652820
Arabidopsis–AT3G11380.1|PACid_19662213
100
Arabidopsis–AT3G11350.1|PACid_19664484
Arabidopsis–AT1G28020.1|PACid_19653938
